# SILAC phosphoproteomics reveals unique signaling circuits in CAR-T cells and the inhibition of B cell-activating phosphorylation in target cells

**DOI:** 10.1101/2021.09.10.459784

**Authors:** Alijah A. Griffith, Kenneth P. Callahan, Nathan Gordo King, Qian Xiao, Xiaolei Su, Arthur R. Salomon

## Abstract

Chimeric antigen receptor (CAR) is a single-pass transmembrane receptor designed to specifically target and eliminate cancers. While CARs prove highly efficacious against B cell malignancies, the intracellular signaling events which promote CAR T cell activity remain elusive. To gain further insight into both CAR T cell signaling and the potential signaling response of cells targeted by CAR, we analyzed phosphopeptides captured by two separate phopshoenrichment strategies from third generation CD19-CAR T cells cocultured with SILAC labeled Raji B cells by liquid chromatography-tandem mass spectrometry (LC-MS/MS). Here, we report that CD19-CAR T cells upregulated several key phosphorylation events also observed in canonical T cell receptor (TCR) signaling while Raji B cells exhibited a significant decrease in B cell receptor-signaling related phosphorylation events in response to coculture. Our data suggest that CD19-CAR stimulation activates a mixture of unique CD19-CAR-specific signaling pathways and canonical TCR signaling while global phosphorylation in Raji B cells is reduced after association with the CD19-CAR T cells.

## Introduction

The ability to modify patient T cells to express chimeric antigen receptors (CARs) that target specific tumor-associated antigens has revolutionized cancer immunotherapy. CARs are synthetic single-pass transmembrane receptors comprised of an extracellular single-chain antibody fragment (scFv), enabling antigen specificity, along with multiple intracellular signaling domains which act cooperatively to initiate T cell receptor (TCR)-like signaling upon CAR association with its target antigen.^1–10^ CD19-targeted CARs have proven highly efficacious in patients with B cell malignancies, yielding complete responses in both refractory and relapsed diffuse large B cell lymphoma, chronic lymphocytic leukemia, and B cell acute lymphoblastic leukemia.^11^ Nevertheless, the efficacy of CAR T cell therapy is overshadowed by potential side effects including cytokine release syndrome (CRS) and immune effector cell-associated neurotoxicity (ICANs).^4,10,12,13^ While factors predisposing patients to certain clinical outcomes are defined and there is a fundamental understanding of the endogenous signaling molecules incorporated into CAR constructs, it is without question that there is limited insight regarding the mechanism underlying CAR-based signal transduction and how CAR design may influence this process. It is necessary to comprehensively study CAR-based signaling to improve future CAR efficacy while minimizing CAR toxicity. ^14^

Endogenous TCR signaling is initiated by the association of the TCR with antigenic peptides presented by major histocompatibility complexes (pMHC) expressed on the surface of antigen-presenting cells.^15^ pMHC-TCR ligation promotes immunological synapse formation, during which costimulatory signaling receptors are recruited and the Src family protein tyrosine kinase Lck is activated. Active Lck phosphorylates immunoreceptor tyrosine-based activation motifs (ITAMs) present within the cytoplasmic tails of CD3 subunits (TCR*ζ*, CD3*ϵ*/*δ*/*γ*) complexed with the TCR.^15^ Upon phosphorylation, ITAMs serve as docking sites for Zap70, a critical Lck substrate that is central to the signal propagation necessary to initiate an effective immune response.^13,15–17^ Through the presence of CAR, CAR-T cells can bypass both the MHC-based antigen recognition and antigen processing requirements inherent to endogenous T cell activation, allowing for streamlined recognition and targeting of intact antigenic molecules present on malignant cells.^18–21^ Downstream effects of TCR stimulation, including T cell proliferation, cytotoxicity, and cytokine release are all observed after CAR activation, indicating that CAR signal transduction likely involves similar molecular machinery as canonical TCR signaling. Nevertheless, many questions remain as few studies have comprehensively characterized CAR-based signal transduction. ^22–24^

Phosphoproteomics enables the simultaneous monitoring of cooperative phosphorylation-based signaling networks in response to cellular perturbations and is a well-suited approach to elucidate CAR T cell signaling transduction. ^25–27^ The first proteomic study of CAR was reported by Salter *et al*. and investigated kinase-mediated signaling of two clinically relevant CD19-targeting CARs stimulated by *α*-CD19 antibody. After pTyr-1000 (phosphotyrosine-specific enrichment) and immobilized metal affinity chromatography (IMAC; pan-specific phosphorylation enrichment), a total of 571 phosphotyrosine (pTyr), 4,647 phosphothreonine (pThr) and 21,586 phosphoserine (pSer) sites were identified from three independent experiments. The phosphorylation events identified by Salter *et al*. exhibited minor exceptions from those commonly observed by antibody-stimulated TCR phosphoproteomic studies, namely a lack of CD3*ϵ*/*δ* phosphorylation post-stimulation.^17,28^ While notable as the first proteomic investigation of CAR signaling, antibody stimulation disregards the potential influence of the interaction between other surface molecules expressed by both the T and target cell and, therefore, may not fully recapitulate physiological activation of the CAR. A later study by Ramello *et al*. addressed this consideration by stimulating prostate stem cell antigen (PSCA)-targeting CAR T cells with PSCA-expressing human pancreatic ade-nocarcinoma cells (HPACs) in coculture, leveraging stable isotopic labeling of amino acids in cell culture (SILAC) to determine the cell of origin for each captured peptide. Much like Salter et al., pTyr-1000 enrichment was performed but suffered from poor sequencing depth, reporting only 40 pTyr and 751 pSer/pThr sites. In this study, only 20 pTyr and 20 pSer/pThr peptides changed significantly after CAR stimulation. Though many TCR-related pTyr phosphopeptides were observed, the phosphopeptides originating from the target cells (HPACs) were not reported by this study.^29^

Ideally, quantification of as many phosphorylation sites as possible in both CAR T and B cells during their interaction would provide the most detailed and biologically relevant view possible of phosphorylation-mediated signaling networks associated with CAR T cell therapy. We recently compared the commonly used pTyr-1000 enrichment to Src Superbinder (sSH2), a highly efficient pTyr enrichment scheme that uses a modified form of the pTyr-specific Src-homology 2 (SH2) domain as an affinity reagent.^30,31^ Our comparison demonstrated that sSH2 enrichment exhibited twice as many unique pTyr sites along with an 82% increase in pTyr peptide abundance than results obtained by pTyr-1000.^31^ The superior performance of sSH2 indicates its potential usefulness in situations wherein detection of high numbers of unique pTyr sites are desired, such as in the characterization of CAR T cell signaling. In this study, we simultaneously characterized changes in wide-scale pTyr-specific and total phosphorylation in both CAR T cells and target cells upon CAR activation by coculturing CD19-CAR Jurkat T cells (CD19-CAR T cells) with SILAC-labeled CD19-expressing Raji B cells (Raji B cells) and combining the principles of SILAC, pTyr specific sSH2 enrichment, and pan-specific titanium dioxide phosphoenrichment (TiO_2_). Here, we report the kinase-mediated signaling responses during coculture in both CD19-CAR T cells and Raji B cells utilizing these optimized phosphoproteome enrichment strategies.

## Materials and Methods

### Cell lines, Generation of CD19-expressing Raji B cells and Jurkat CD19-CAR T cells

Both Jurkat T cell (clone E6.1) and Raji B cell lines (UCSF Cell Culture Facility) were grown in RPMI 1640 with 10% (v/v) fetal bovine sera (FBS), 2 mM 2 mM L-glutamine, 100 U/mL penicillin, 100 *μ*g/mL streptomycin, and 10 mM HEPES, pH 7.4. Raji B cells were stained for CD19 with PE anti-human CD19 antibody (Biolegend, #392506, clone 4G7) and the top 3% highest CD19-expressing cells (Raji B cells) were collected by fluorescence activated cell sorting. The number of CD19 molecules expressed on the cell surface was estimated by flow cytometry in combination with the BD Quantibrite kit (BD Biosciences, #340495) according to the manufacturer’s instructions. CD19-CAR T cells were generated using a lentiviral vector of third generation CD19 FM63 scFV CAR tagged with superfold GFP (pHR-CD19 ScFv-myc-CD8 stalk-CD8 TM-CD28-41BB-CD3*ζ*-sfGFP). Lentiviral particles were produced by co-transfection of HEK293T cells with pHR plasmids and the second-generation lentiviral packaging plasmids, pMD2.G (Addgene, plasmid #12259) and psPAX2 (Addgene, plasmid #12260) using Genejuice transfection reagent (EMD Millipore, #70967-3). Cell culture media containing viral particles were harvested 48 hours post-transfection, centrifuged, and filtered through 0.45*μ*m pore size filters before infection of Jurkat T cells in RPMI media for 72 hours. Jurkat T cells expressing CD19-CAR-GFP were sorted by flow cytometry.

### Cell Culture & SILAC labeling of Raji B cells

CD19-CAR T cells and Raji B cells were initially cultured in RPMI 1640 supplemented with 10% (v/v) FBS, 2mM L-glutamine, 100*μ*/mL penicillin, and 100 *μ*g/mL streptomycin and maintained under standard conditions (5% CO_2_, 37 °C). After 12 days in culture, Raji B cells were collected and washed twice with SILAC RPMI 1640 (Thermo Fisher Scientific, Waltham, MA) before reconstitution in SILAC RPMI 1640 medium (Thermo Fisher Scientific, Waltham, MA) containing 0.38 mM ^13^C_6_, ^15^N_4_ Arginine, 0.22 mM ^13^C_6_, ^15^N_2_ Lysine (Cambridge Isotope Laboratories, Andover, MA), 10% dialyzed FBS (Gibco Life Technologies, Guilford, CT), 2 mM L-glutamine, 100 U/mL Penicillin, and 100 *μ*g/mL Streptomycin (Cytiva, Malborough, MA) and maintained for a total of 8 doublings under standard conditions (5% CO_2_, 37 °C). During this time, CD19-CAR T cells were expanded and maintained in RPMI 1640 medium supplemented with 10% (v/v) FBS, 2 mM L-glutamine, 100 U/mL Penicillin, and 100 *μ*g/mL Streptomycin.

### Co-incubation, lysis, alkylation, reduction, and digestion for Phosphoproteomics

CD19-CAR T cells and Raji B cells were collected and re-suspended individually at a concentration of 2.6 *×* 10^8^ cell/mL in pre-warmed RPMI 1640 without phenol red (Sigma Aldrich, St. Louis, MO), followed by a rest period of 5 minutes at 37 °C. To initiate coculture, CD19-CAR T cells were mixed with Raji B cells in a 1 : 1 ratio, spun at 500g for 30 seconds at room temperature (RT)^32,33^ and cocultured for either 2- or 5-minutes. Cocultured cells were lysed through an 5 : 1 dilution using a concentrated urea lysis buffer (final concentrations of 8M Urea, 20 mM HEPES, 1 mM sodium orthovanadate, 2.5 mM sodium pyrophosphate, 1 mM *β*-glycerophosphate, pH 8). The resulting lysates were chilled at 4°C for 15 minutes before sonication at 75% amplitude for 30 seconds, twice. Samples were centrifuged at 21,000g for 5 minutes at 20 °C. A negative control was generated by performing the previous steps for CD19-CAR T cells and Raji B cells independently and combining the resulting lysates from both cell lines after sonication. Lysate protein concentrations for all samples were determined by Pierce BCA Assay (Thermo Fisher Scientific, Franklin, Massachusetts) as per the manufacturer’s instructions. Total protein across all samples was normalized to 32 mg per replicate with each coculture time-point representing a total of 5 biological replicates. Samples were treated with 10 mM DTT for 30 minutes at 37 °C and 10 mM Iodoacetamide for 30 minutes at RT in the dark. The resulting protein samples were diluted to a final concentration of 2M Urea using 20 mM HEPES (pH 8) and digested overnight at 37°C using sequencing grade modified Trypsin (Promega, Madison, WI) in a 1 : 50 (w/w) trypsin-to-protein ratio.

Following tryptic digestion, peptides were acidified to 1% (v/v) trifluoroacetic acid (TFA), centrifuged at 1,800g for 5 minutes at RT and desalted using Sep-Pak C18 Plus cartridges (Waters, Milford, MA). Each individual sample was split into three respective portions and desalted in series with 12 mL 0.1% TFA. Desalted peptides were eluted from the C18 SepPak column using 7 mL 0.1% TFA in 40% acetonitrile (ACN) and the eluted peptides from the three original portions were combined. One-tenth of total protein content as determined by BCA (e.g., 3.2 mg) was used as starting material for global phosphopeptide enrichment by titanium dioxide (TiO_2_) and dried by speed-vac prior to enrichment. The remaining nine-tenths of each sample were diluted 1 : 1 in 0.1% TFA to reduce the final ACN concentration to 20% (v/v), stored at −80 °C overnight, and lyophilized for 48 hours prior to phosphotyrosine enrichment by Src SH2 superbinder.

### Global Phosphoenrichment by Titanium Dioxide

Global enrichment of phosphorylated peptides was performed using Titansphere Phos-Tio Kit (GL Sciences, Japan) as described.^34^ Briefly, 3.2 mg of lyophilized, trypsinized peptides were reconstituted in 100 *μ*L TiO_2_ buffer (GL Sciences, Japan) and centrifuged at 12,000g for 5 minutes. Titanium dioxide phosphotips (GL Sciences, Tokyo, Japan) were washed once using 20 *μ*L of Buffer A (0.4% v/v TFA in ACN) followed by 20 *μ*L Buffer B (6.25% v/v Lactic acid, 3% v/v TFA in ACN). Tips were centrifuged at 3,000g for 2 minutes at RT between each solvent. Fifty microliters of the reconstituted sample were loaded onto the TiO_2_ phosphotips with 100 *μ*L Buffer B, mixed briefly by pipetting, and spun at 1,000g for 10 minutes at RT. This process was then repeated for the remaining 50 *μ*L of reconstituted sample. After the entire sample was loaded onto the TiO_2_ phosphotips, the tips were first washed once with 20*μ*L Buffer B followed by four washes with 20 *μ*L Buffer A. Tips were centrifuged at 3,000g for 2 minutes at RT following each wash. Phosphopeptides were eluted from the TiO_2_ tips by first adding 50 *μ*L of 5% ammonium hydroxide followed by 50 *μ*L 5% pyrrolidine solution and centrifuging the tips at 1,000g for 5 minutes at RT after the addition of each solvent. Eluted phosphopeptides were dried overnight by speed-vac before LC-MS/MS analysis.

### sSH2 Superbinder Phosphotyrosine Enrichment

Phosphotyrosine enrichment was performed using the superbinder SH2 (sSH2) domain as described previously.^30,31^ Briefly, purified superbinder SH2 conjugated to CNBr-activated Sepharose beads (GE Healthcare, Wauwatosa, Wisconsin) was used to capture pTyr containing peptides in our samples as described. ^31^ Approximately 28 mg of desalted, trypsinized peptides was re-suspended in 3 mL IAP buffer (10 mM Sodium phosphate monobasic monohydrate, 50 mM sodium chloride, 500 mM MOPs, pH 7.2) before a 2 hour incubation with superbinder beads containing 1 mg of superbinder at 4°C on a 4 rpm-rotator. Beads were washed three times with 1 mL ice-cold IAP buffer and 1 mL ice-cold unbuffered HPLC-grade water before elution. Captured peptides were eluted from the beads by mixing sSH2-conjugated beads with 100 *μ*L 0.15% (v/v) TFA for 10 minutes on a 1,150 rpm mixer at RT, twice. Eluted peptides were desalted using C18 tips (Thermo Scientific, Waltham, MA) according to the manufacturer’s guidelines before speed-vac and subsequent LC-MS/MS analysis.

### Western Blotting

Western Blot samples were prepared by lysing 2 *×* 10^6^ cells in 2*×* Laemmli sample loading buffer (4% SDS, 125 mM Tris-HCL pH 6.8, 20% glycerol, 5% *β*-mercaptoethanol, 0.01% bromophenol blue) to a final concentration 2 *×* 10^7^ cell/mL, followed by boiling for 10 min at 90 °C, and spinning at 13,000 rpm for 5 minutes. Proteins were separated by 4-20% TEO-Tricine gradient gel (Abcam, Cambridge, MA) and transferred to a PVDF Immobilon membrane (EMD Millipore, Billerica, MA), which was blocked using Odyssey blocking buffer (Li-Cor, Lincoln, NE) prior to incubation with primary antibody cocktails. In cases where total protein stain was performed, REVERT Total Protein Stain (LiCor, Lincoln, NE) was used according to manufacturer’s instructions prior to blocking. *α*-pERK1/2 (T202/Y204) antibody (Cell Signaling cat. 9101, Danvers, MA), *α*-Erk1/2 (Cell Signaling cat. 9107, Danvers, MA), *α*-Lck (Cell Signaling Technologies, cat. 2657, Danvers, MA), *α*-4G10 (Millipore Sigma cat. 05-321, Billerica, MA), *α*-GAPDH (Sigma Aldrich cat. G9545, St. Louis, MO) were used as primary antibodies. Donkey-*α*-mouse IgG (LiCor, Lincoln, NE) and goat-*α*-rabbit IgG (LiCor, Lincoln, NE) secondary antibodies were used for the Odyssey CLx Imaging System (Li-Cor, Lincoln, NE).

### Liquid Chromatography-Mass Spectrometry (LC-MS/MS)

Enriched tryptic peptides were separated on a reversed-phase, in-line analytical column (150 mm by 75 *μ*m) packed in house with XSelect CSH C18 2.5 um resin. Peptides were eluted for 90 min using a gradient of beginning in a Buffer A (0.1% formic acid, 0.5% HPLC-grade acetonitrile, 99.4% HPLC-grade water) to Buffer B (0.1% formic acid, 99.9% HPLC-grade acetonitrile) at a starting Buffer A:B ratio of 98 : 2 progressing to a final ratio of 65 : 35. Eluted peptides were ionized by electrospray (2.2 kV, positive spray) and analyzed by an Orbitrap Exploris 480 mass spectrometer (Thermo) in DDA mode with a cycle time of 2.5 s. MS1 were acquired for precursor ions with a charge state between +2 and +6 and within a range of 375 m/z to 1500 m/z using a Fourier Transform mass spectrometer (FTMS) in profile mode at a mass resolution of 120,000, a maximum injection time of 50 ms, a dynamic exclusion time of 20 s and a normalized target automatic gain control (AGC) of 300%. After higher energy collisional dissociation, MS2 spectra for selected fragment ions were acquired in a 0.7 m/z window with a normalized collision energy of 30%, normalized target AGC of 50% and a resolution of 15,000 using the FTMS in centroid mode.

### Database Searching

Raw files generated by the Orbitrap Exploris 480 were processed through the high-throughput autonomous proteomic pipeline (HTAPP) and PeptideDepot, a custom phosphoproteomics software created in house.^35,36^ Briefly, MS2 spectra for each experiment were compared with a database containing 98,300 non-redundant forward sequence proteins from the UniProt complete proteome data set (Homo sapiens, downloaded on 2019-08-30) and an equal number of reverse sequence decoys generated by Mascot (version 2.4.1).^37^ The following parameters were used for the MASCOT search: trypsin enzyme cleavage (tolerating up to 2 missed cleavages), 7 ppm mass tolerance for precursor ions, 20 mmu mass tolerance for fragment ions, variable modifications for phosphorylation (serine, threonine, tyrosine: +79.9963 Da), oxidation (methionine: +15.9949 Da), and a static modification of carbamidomethylation (cysteine: +57.0215 Da). Mascot results were filtered by MOWSE score to a false discovery rate (*FDR*) of 1%.^26,27^ The ambiguity score (Ascore) algorithm^38^ was applied to determine the confidence in phosphorylation site assignment for each phosphopeptide spectrum.

### Relative Phosphopeptide Abundance

Relative phosphopeptide abundance was determined by integration of the selected ion chromatogram (SIC) peaks for each phosphopeptide. Retention times were aligned for replicate analyses as previously described,^27^ while peak areas were calculated using an in-house software package coupled with the R Bioconductor package XCMS (version 1.42.0).^39^ Heat maps depicting the phosphopeptide abundance were produced in PeptideDepot, wherein each heat map box corresponds to a specific time point. For a given phosphopeptide, the geometric mean of all replicates across all time points was represented by the color black, while above and below average abundances were indicated by either a yellow or blue color, respectively. A given peptide for which a SIC was not identified any of the replicates for a specific time point was represented by a blank heat map square.^40^ A *FDR* less than 0.05 indicates statistical significance, and was indicated by a white dot present within in the center of a heat map square. To estimate the *FDR*, *p*-values were first determined for *log*_2_ transformed intensities using a two-tailed Student’s T-test. For each phosphopeptide, Student’s T-tests compared the means of all time points to the time point with the lowest mean abundance and the resulting list of *p*-values was used to estimate the *FDR* by applying the algorithm of Benjamini & Hochberg using the R package QVALUE (version 1.43.0), as previously described. ^26,41,42^ Replicate reproducibility was evaluated by computing the pairwise Pearson correlation coefficient for all time points between replicates.

### Post-translational Modification Signature Enrichment Analysis (PTM-SEA)

The relationship between the observed phosphorylation events and PhosphoSitePlus annotations was determined by PTM-SEA, a modified version of ssGSEA.^43^ Briefly, phosphorylated peptides with an Ascore > 13 (localization probability > 0.73) were centered around the phosphorylation site(s) with seven flanking amino acids on each side. ^38,43^ For peptides with multiple confidently assigned phosphorylation sites, flanking sequences were included for each site and the original intensity values were duplicated for each flanking sequence. For each flanking sequence, Student’s T-Tests were performed on log_2_ transformed peak areas between all time points for which three or more replicate peak areas were measured, and the resulting list of *p*-values were adjusted to *q*-values using the method of Benjamini & Hochberg. ^41,42,44^ As input for PTM-SEA, replicates were condensed by transforming the *q*-values as follows:

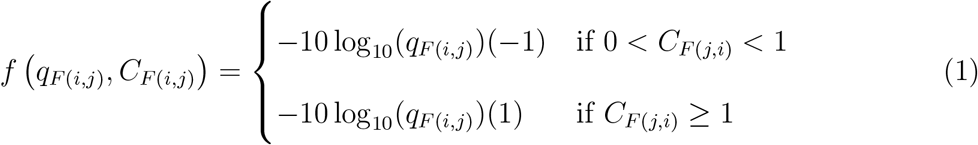

where *j* > *i*, *q_F_* _(*i,j*)_ was the *FDR* corrected significance between time points *i* and *j* for flanking sequence *F*, and *C_F_* _(*j,i*)_ = *μ_j_/μ_i_* qas the ratio of the mean intensities between time point *j* and time point *i* for flanking sequence *F*, as described previously.^43^ The flanking sequences and accompanying transformed *q*-values were written in GCT 1.3 format and used as input for PTM-SEA. The settings for PTM-SEA of transformed *q*-values were as follows: the values were not rank normalized and the weight parameter was set to 1, incorporating replicate variance into the enrichment score calculation. ^28,43^ Normalized Enrichment Scores (*NES*) were calculated for each signature set *S* in a group *G* by rank ordering transformed *q*-values for one condition, determining group membership for modifications annotated as upregulated (*ES*_u_) and downregulated (*ES*_d_), calculating *ES*_S_ = *ES*_u_ − *ES*_d_, and normalizing *ES*_S_ to its null distribution.^43^ Statistical significance for each *S* was determined by a permutation test and corrected for false discoveries using the method of Benjamini & Hochberg. ^42–44^

### Code and Data Availability

Our in-house HTAPP^35^ and analysis software PeptideDepot ^36^ were used for initial analysis of mass spectrometry data, generation of abundance heat maps and determination of KEGG/GO categorical membership. The script “automate ptm sea.py” was used to perform all steps of PTM-SEA. Data management and statistical tests were performed using Python (version 3.8.10) with the following non-base modules: matplotlib (version 3.3.2), NumPy (version 1.19.2), SciPy (version 1.6.1) and Pandas (version 1.2.3). PTM-SEA was run using R (version 4.1.0) with the script “ssGSEA2.0.R” and the human flanking PTM-sigDB (version 1.9.0).^43^ All scripts and data are available in Supplementary Folder 1 and on Github (https://github.com/drsalomon/griffith_callahan_2021_CART_code).

## Results and Discussion

### CAR Expression, SILAC labeling, Sample Collection, and Enrichment Depth

We used third-generation CD19-CAR T cells and CD19-expressing Raji B cells to characterize the kinase-mediated signaling events induced in both cell populations upon CAR-CD19 ligation. A previously generated third-generation CD19-CAR construct^10^ (Figure 1) was introduced in Jurkat T cells by lentiviral transduction. CD19-CAR expression was confirmed post-transduction using flow cytometry (Supplementary Figure 1). CD19-CAR T cells were activated by coculture^32,33^ with SILAC-labeled Raji B cells to better simulate physiological conditions of CAR stimulation. SILAC-labeling of the Raji B cells occurred for a total of 8 doublings prior to coculture so that phosphopeptide origin could be determined after LC-MS/MS. 99.7% of Raji B cell peptides exhibited ^13^C_6_, ^15^N_4_ Arginine, ^13^C_6_, ^15^N_2_ Lysine labeling with 0.7% arginine-to-proline conversion within this labeling period (Supplementary Tables 1, 2, 3). Three coculture time-points (0-, 2- and 5-minutes) were assessed and represented by five biological replicates each. Coculture peptides were enriched for either pTyr containing peptides by Src SH2 superbinder enrichment (sSH2) or total phosphorylation by titanium dioxide enrichment (TiO_2_) prior to LC-MS/MS (Figure 2).

**Figure 1:**
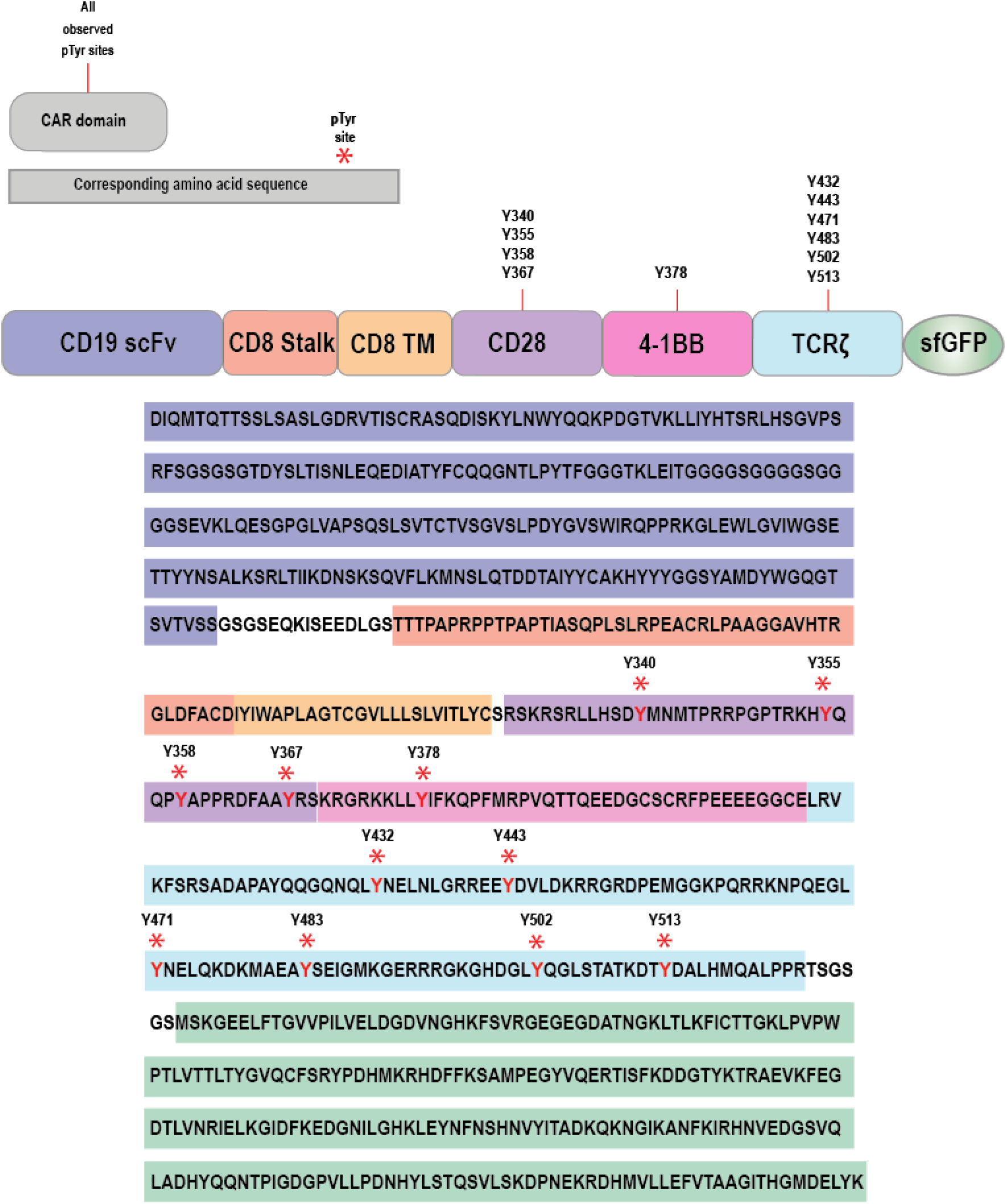
Amino acid sequence for the CD19-targeting CAR used in our study. The colors used to highlight portions of the CAR sequence correspond to the domains used to construct the CAR. Red tyrosine (Y) amino acids with a star (*) character above them indicate tyrosine phosphorylation sites frequently observed in the native proteins after TCR stimulation.

**Figure 2:**
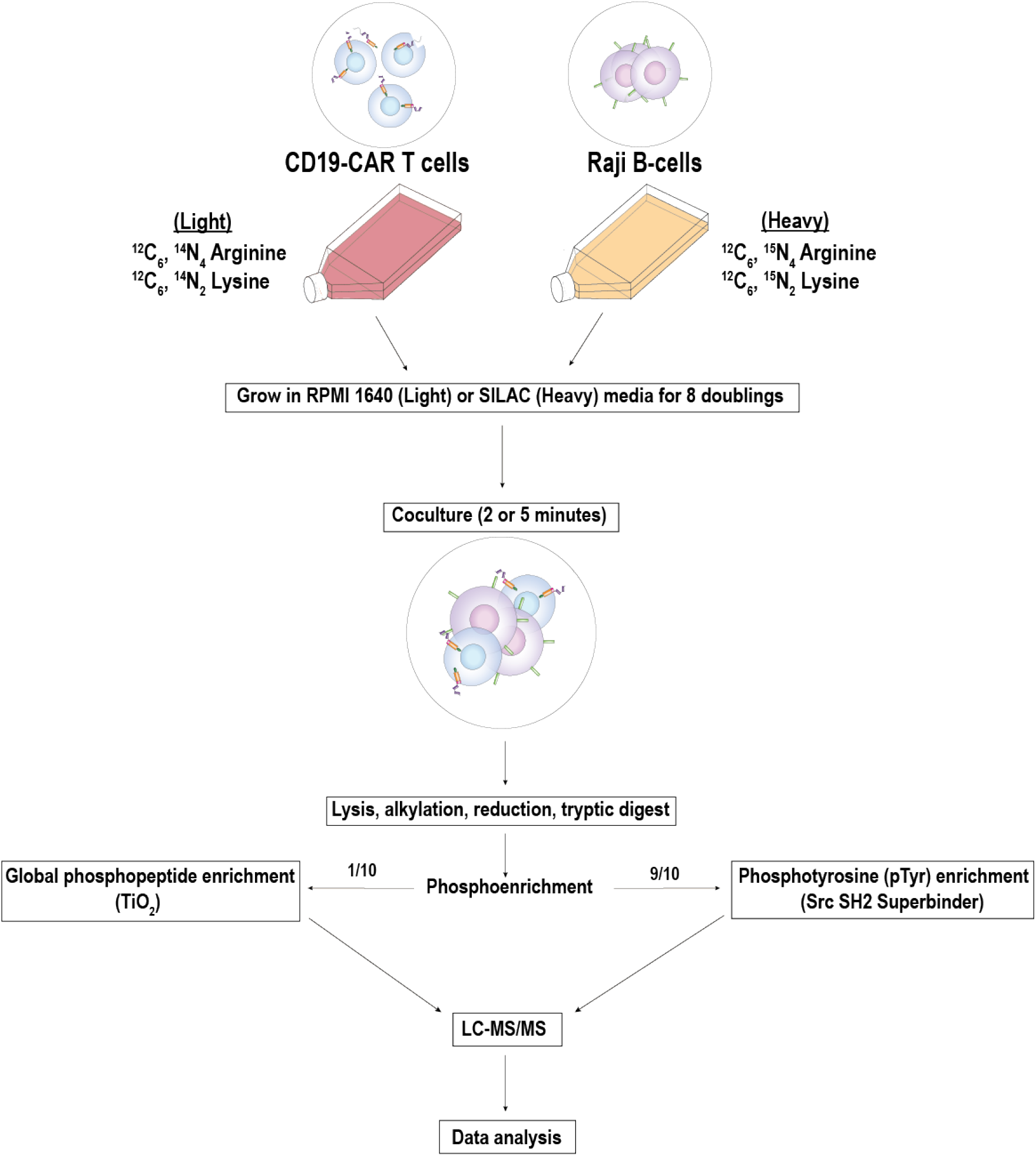
Experimental Methods Schematic. Raji B cells were “heavy” labeled with ^13^C_6_, ^15^N_4_ Arginine, and ^13^C_6_, ^15^N_2_ Lysine for 8 doublings, while CD19-CAR T cells were “light” labeled with ^12^C_6_, ^14^N_4_ Arginine, and ^12^C_6_, ^14^N_2_ Lysine. The CD19-CAR T cells and Raji B cells were cocultured together for 2- or 5-minutes, or mixed after lysis (0-minutes; negative control). One-tenth of the total peptide sample was subjected to TiO_2_ phosphopeptide enrichment while the remaining nine-tenths was used for superbinder sSH2 enrichment before LC-MS/MS analysis.

Peptide spectrum matches, phosphorylation site assignments, and the relative abundance of each phosphopeptide were determined as described in the Materials and Methods. Data reproducibility was evaluated by calculating the pairwise correlation coefficient between replicate peak areas across all of the time points and replicates (Supplementary Figures 2, 3). A complete list of all unique phosphopeptides observed in this study is available (Supplementary Tables 4, 5 and and Supplementary Figures4, 5) and the mass spectrometry proteomic data was deposited to the ProteomeXchange consortium (http://proteomecentral.proteomexchange.org) *via* the PRIDE partner repository with the data set identifier PXD028109.

A total of 590 unique pTyr sites were identified from 386 proteins at an *FDR* < 0.01 after sSH2 enrichment. A slight majority of the unique pTyr sites were identified in the Raji B cells (375 pTyr sites) compared to CD19 CAR T cells (319 pTyr sites). A total of 8,731 unique sites (353 unique pTyr [4%], 7,006 unique pSer [80%] and 1,372 unique pThr [16%] sites) were identified from 3,494 proteins at an *FDR* < 0.01 after TiO_2_ enrichment. More of the phosphopeptides observed after TiO_2_ enrichment were identified in the CD19-CAR T cells (6,465 phosphorylation sites) compared to Raji B cells (4,708 phosphorylation sites). Average correlation coefficients (*ρ*) obtained from the sSH2 data revealed high replicate reproducibility for both CD19-CAR T cells (0.8839*±*0.0303) and Raji B cells (0.8770*±*0.0284). Compared to the sSH2 data, the TiO_2_ data set exhibited lower replicate reproducibility for CD19-CAR T cells (0.6458 *±* 0.0829) and Raji B cells (0.6896 *±* 0.0653) but was sufficient to identify 2,324 significantly changed phosphorylation site-containing PSMs.

### Coculture Promotes Phosphotyrosine Signaling Cascades in CD19-CAR T cells

Anti-phosphotyrosine Western blot of coculture lysates revealed increased global tyrosine phosphorylation upon coculture, indicating signal transduction had occurred in response to coculture (Figure 3B). Post-translational modification enrichment analysis (PTM-SEA) of the sSH2 data set attributed the increase of global pTyr in the coculture cell lysates to CD19-CAR T cells, as Raji B cells curiously exhibited decreasing pTyr perturbation signatures post-coculture (Supplementary Figure 6B, 7B). Inclusion of TCR*ζ* within the CD19-CAR (Figure 1) may activate canonical TCR signaling pathways such as MAPK signaling, resulting in phosphorylation of Erk1/2.^28^ PTM-SEA of TiO_2_ data revealed both Erk1 and Erk2 kinase substrate signatures significantly increased in CD19-CAR T cells post-coculture (Figure 4A). Western blot further supported Erk1/2 phosphorylation in response to CAR activation, along with the increased abundance observed for phosphorylated kinase-activating sites,^45^ Erk1^T202Y204^ and Erk2^T185Y187^ (Figure 3A,C). Similarly, PTM-SEA of both data sets indicated that pathway phosphorylation signatures significantly increased in CD19-CAR T cells after CAR activation (Figure 4C, Supplementary Figure 6C). Together, our data suggested that CD19-CAR activation promoted a global increase in tyrosine phosphorylation and that CAR may activate pathways involved in canonical TCR signaling upon association with its target antigen.

**Figure 3:**
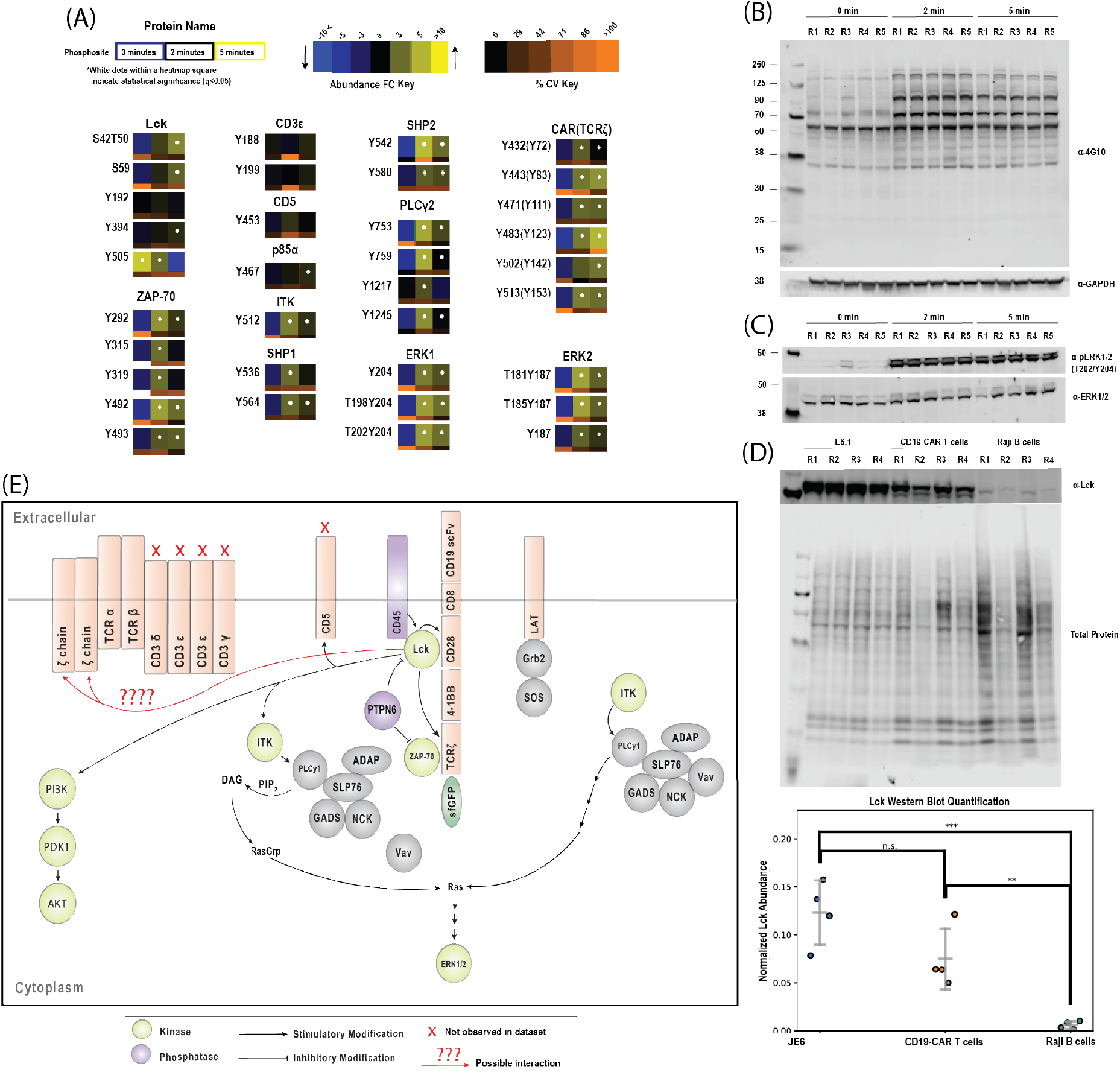
Stimulation of the CD19-CAR promoted activation of tyrosine kinase phosphorylation cascades associated with canonical TCR signaling. (A) Phosphopeptide abundance changes for Lck phosphorylation sites, Lck kinase substrate sites, and other proteins related to initiation and propagation of canonical TCR signaling. Refer to the abundance FC key for interpretation of the presented heatmaps. (B) Global tyrosine phosphorylation in cocultured CD19-CAR T cells and Raji B cells as determined by *α*-pTyr (4G10) Western blot. (C) Erk1^T202Y204^ and Erk2^T185Y187^ phosphorylation in CD19-CAR T cells and Raji B cells, as probed by *α*-pErk Western blot. (D) Lck protein abundance in Jurkat cells (E6.1), CD19 CAR-expressing Jurkat cells, and Raji B cells, as determined by quantification of Western blot using *α*-Lck antibodies and normalized to REVERT total protein stain. (E) Pathway map showing Lck-related phosphorylation events.

**Figure 4:**
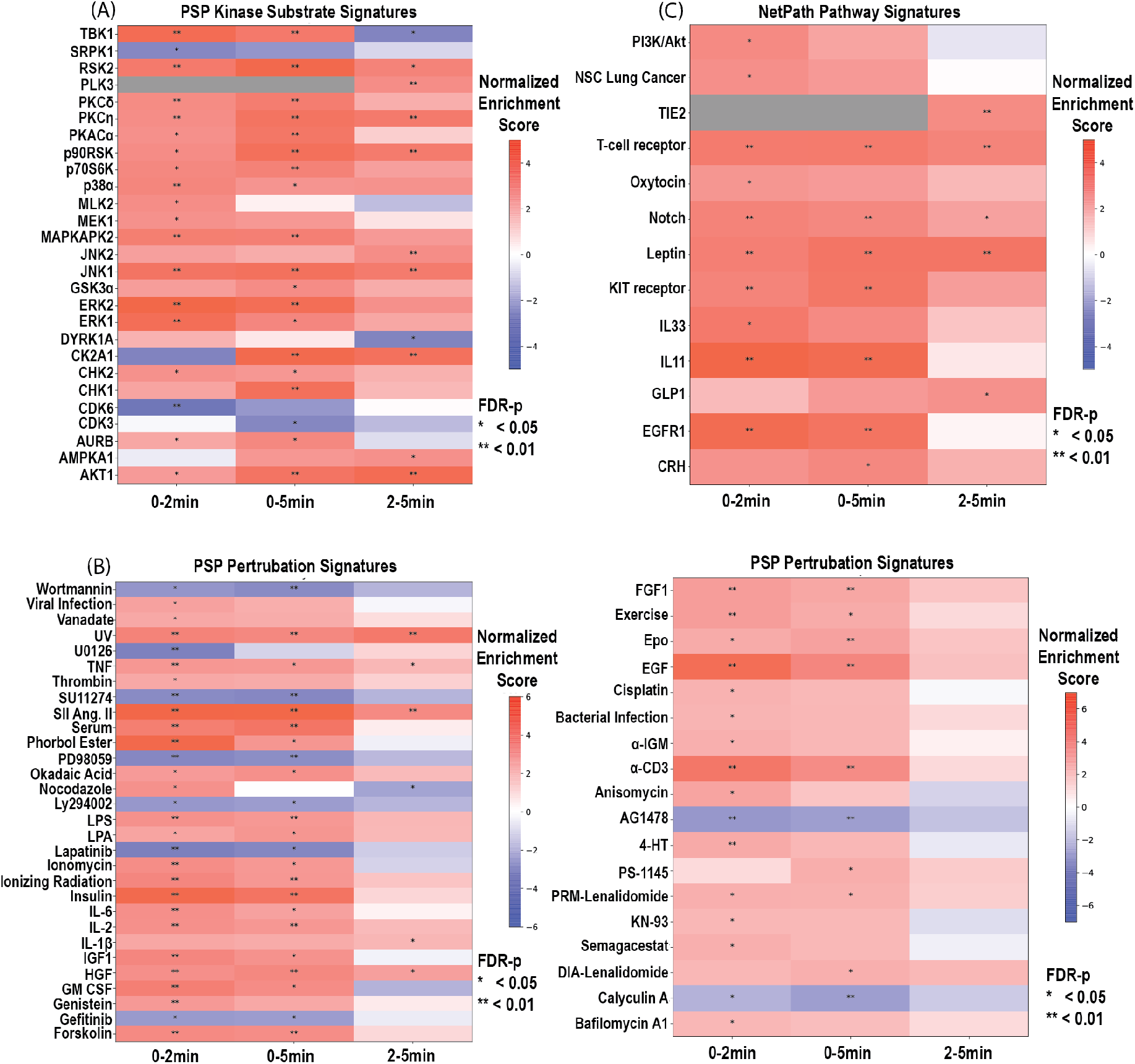
PTM-SEA of phosphopeptides originating from the CD19-CAR T cells that were observed in the TiO_2_ enrichment data. Signature sets in the PTM signature database (PTMsigDB, version 1.9.0) were grouped by (A) PhosphoSitePlus annotated kinase substrate modifications, (B) PhosphoSitePlus annotated modifications that change in response to perturbagens, and (C) NetPath annotated modifications associated with a particular signaling pathway.^43^ Red colors indicate positive correlations and blue colors indicate negative correlations between signature-associated phosphopeptide abundance changes observed between time points. On each heatmap block, one star (*) indicates an *q* < 0.05 and two stars (**) indicates an *q* < 0.01.

### Robust Activation of Lck in CAR T cells from the CD19-CAR

Canonical TCR signaling is initiated by TCR-pMHC ligation, promoting CD45-mediated dephosphorylation of Lck^Y505^ followed by subsequent autophosphorylation of Lck^Y394^. These phosphorylation events cooperatively activate Lck and initiate phosphotyrosine signaling cascades responsible for an effective immune response. ^15^ If CD19-CAR stimulated proximal TCR signaling by activation of Lck, an increased abundance of the activating Lck^Y394^ site accompanied by a decreased abundance of the inhibitory Lck^Y505^ site would be expected shortly after CAR activation. sSH2 enrichment data indicated that Lck was activated by the CD19-CAR as Lck^Y394^ abundance significantly increased by 2.1-fold 5-minutes after CD19-CAR T cell stimulation. Additionally, Lck^Y505^ abundance significantly decreased 2- and 5-minutes after coculture (Figure 3A). Due to the lack of Lys or Arg residues, and therefore SILAC labeling, in the C-terminal tryptic fragment of Lck containing Lck^Y505^, the abundance observed for this PSM could possibly represent a combined signal between both cell populations. We attributed the observed abundance of this PSM to CD19-CAR T cells as Western blot revealed that Raji B cells expressed negligible levels of Lck protein and that the difference in Lck expression between the parental Jurkat E6 and CD19-CAR T cells was not significant (Figure 3D, Supplementary Figure 9, Supplementary Table 6). Overall, a 38-fold decrease in Lck^Y505^ was observed 5-minutes after stimulation, which was in contrast to a 31-fold increase that was previously observed in a separate phosphoproteomic study of TCR signaling using soluble antibody stimulation. ^46^ Furthermore, TiO_2_ global phosphoenrichment data revealed a significant increase in Lck^S59^ phosphopeptide abundance. Lck^S59^ is significant as it may regulate the Erk1/2 positive feedback loop in TCR signaling. ^25,47,48^ Together, our data revealed that Lck activation is differentially regulated in CD19-CAR T cells compared to normal TCR signaling.

When activated, Lck characteristically phosphorylates a number of proteins necessary for signal propagation, including CD3*ϵ*/*δ*/*γ*, TCR*ζ* and Zap70. ^15^ It is worth noting that CD19-CAR T cells express TCR*ζ* both within the endogenous TCR and the CD19-CAR, ^49^ therefore any of the observed TCR*ζ* phosphosites potentially originated from either receptor. The abundance of many Lck substrates significantly increased within the sSH2 data set at both 2- and 5-minutes after CD19-CAR stimulation in CAR T cells (Itk^Y512^, Zap70^Y292^, Zap70^Y315^, Zap70^Y319^, Zap70^Y492^, Zap70^Y493^, TCR*ζ*^Y72^, TCR*ζ*^Y83^, TCR*ζ*^Y111^, TCR*ζ*^Y123^, TCR*ζ*^Y153^, PTPN6^Y564^, PLC*γ*2^Y753^, PLC*γ*2^Y759^ and PKC*δ*^Y313^).^50–57^ These patterns were consistent with phosphoproteomic studies of anti-TCR signaling. ^25–27,46^ The abundances of nearly all identified Lck substrates exhibited a subtle decrease between 2- and 5-minutes coculture (Figure 3A), an observation that was supported by PTM-SEA of the sSH2 pTyr enrichment data (Supplementary Figure 6A). Interestingly, the abundances of some Lck substrate sites failed to significantly change (Figure 3A), most notably CD5^Y453^, a pTyr site on the CD5 costimulatory receptor,^58,59^ and the ITAM sites CD3*ϵ*^Y188^, CD3*ϵ*^Y199^, CD3*δ*^Y160^ and CD3*δ*^Y171^.^54,60^ These sites were previously observed to significantly increase by at least 15 fold with anti-TCR antibody stimulation. ^46^ Overall, our data were consistent with existing literature detailing CAR signaling^17^ but also suggested divergence between early CAR and TCR signaling which will require additional study to be fully characterized.

### The CD28 costimulatory Domain Enhances Canonical TCR Signaling

CD28 is often included in addition to TCR*ζ* in CAR constructs as a costimulatory domain to promote the direct recruitment of Lck, Itk and phosphoinositide-3-kinase (PI3K) to the CAR and subsequently enhance signaling of the TCR*ζ* domain.^22,23^ A number of tyrosine residues present on the intracellular tail of CD28 are phosphorylated after *α*-CD3 or *α*-CD28 treatment of wild-type T cells. Although these sites can be phosphorylated by either Itk and Lck *in vitro*, more recent data from Itk deficient mice suggest these sites are Lck substrate sites. ^61–63^ CD28^Y191^, CD28^Y206^, CD28^Y209^ and CD28^Y218^ were all significantly increased in the sSH2 data set (Figure 5A). The abundance of PLC*γ*1^Y783^, a site phosphorylated by Itk, also significantly increased within the same data set. Activation of PLC*γ*1 by Itk phosphorylation converts phosphoinositol-4,5-bisphosphate (PIP_2_) into diacylglycerol (DAG) and inositol-1,4,5-trisphosphate (IP_3_) promoting MAPK/Erk, PKC/NF-*κ*B, and calcium signaling. ^26^ PTM-SEA of our TiO_2_ data set revealed that Jnk1/2, PKC*δ* and p38*α*/*δ* kinase substrate signatures all significantly increased after CD19-CAR stimulation in the CD19-CAR T cells (Figure 4A), which is notable as Jnk1/2, PKC*δ*, and p38*α*/*δ* are all targets of activated PLC*γ*1. Phosphorylation of the CD28 ITAMs also promotes the activation of the PI3K/Akt pathway,^22,23,63,64^ therefore, we expected to observe increased abundance of phosphopeptides that are associated with PI3K/Akt signaling in CD19-CAR T cells after CD19-CAR stimulation. sSH2 pTyr enrichment revealed that only four PI3K-related phosphopeptides exhibited significant abundance changes after CD19-CAR stimulation in the CD19-CAR T cells, namely p85*α*^Y467^, p85*α*^Y607^, p85*δ*^Y460^, p85*δ*^Y464^ and p85*δ*^Y605^ (Figure 3A). The physiological relevance of these sites has not yet been determined in any signaling context although p85*α*^Y607^ and p85*δ*^Y605^ are both predicted Lck substrates on PhosphoNET (www.phosphonet.ca).^65^ TiO_2_ enrichment provided evidence that the PI3K/Akt signaling pathway was activated in CD19-CAR T cells, as we observed a significant increase in PI3K signaling pathway signatures by 2-minutes and a significant increase in Akt1 kinase substrate signatures after CD19-CAR stimulation by PTM-SEA in this cell population (Figure 4A,C). Notably, the Akt1 kinase substrate site PLC*γ*1^S1248^ (Figure 5A) significantly increased, suggesting long term downregulation of PLC*γ*1 kinase activity by the PI3K/Akt pathway. ^66^ When combined with the observation of Erk1/2 activation (Figures 3A,C, 4A) and the current knowledge of CD28 costimulatory action, ^22,23,63,64^ our data supported a model of signal transduction in CD19-CAR T cells wherein the CD28 costimulatory domain of the CAR promoted Itk and PLC*γ*1 activation and downstream activation of the PI3K/Akt pathway.

**Figure 5:**
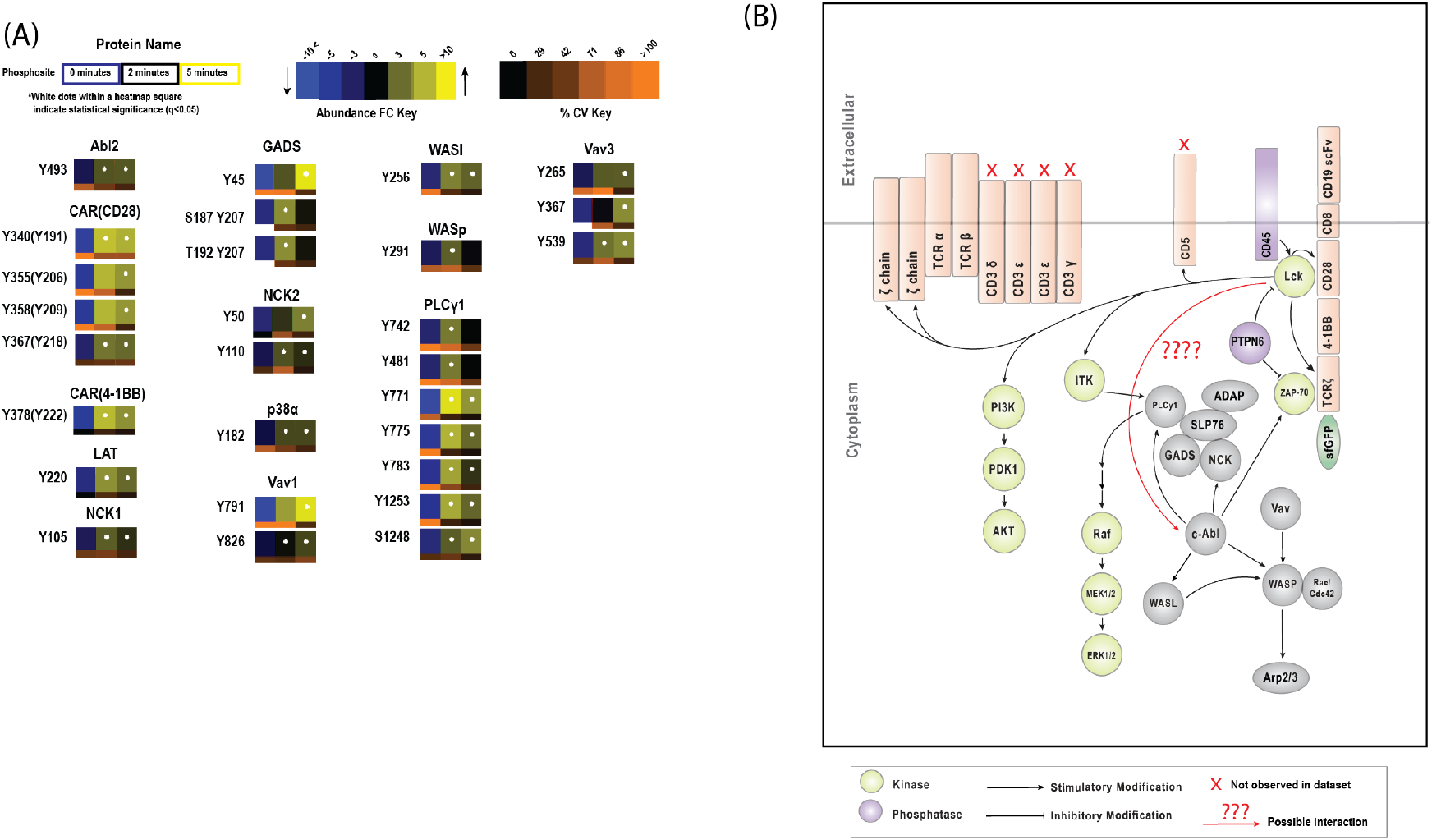
Stimulation of the CD19-CAR in CAR T cells promotes phosphorylation events associated with the CD28 costimulatory domain, the LAT-SLP76 complex and the nonreceptor tyrosine kinases Abl1/2. (A) Phosphopeptide abundance changes for CD28-related and LAT-SLP76-related phosphorylation sites and Abl2 kinase substrate sites. For heat map block color information, refer to the Abundance FC Key or the Materials and Methods. (B) Truncated pathway map illustrating SLP76 and Abl1/2-related phosphorylation events.

### Signaling from the LAT-SLP76 Complex

Following TCR-pMHC binding, Lck-activated Zap70 characteristically phosphorylates a number of proteins that are required for signal propagation from the TCR, including the linker for activation of T cells (LAT) and the SH2 domain containing leukocyte protein of 76kD (SLP76).^67–69^ Phosphorylation of LAT by Zap70 triggers the recruitment of SLP76 and formation of the SLP76 complex, which includes PLC*γ*1, Itk, GADS, ADAP, and Vav1.^70^ Our superbinder sSH2 pTyr enrichment data revealed that the Zap70 substrate site LAT^Y220^ (LAT^Y191^ in LAT isoform 2), a phosphorylation site that improves PLC*γ*1 and Grb2/SOS binding to the LAT,^71,72^ increased significantly after CD19-CAR stimulation in CD19-CAR T cells. Another Zap70 substrate site, ADAP^Y571^, also increased significantly within the CD19-CAR T cells the same data set. The appearance of ADAP^Y571^ was particularly interesting as it promotes both ADAP/STAT3 interaction and subsequent STAT3 phosphorylation,^73^ the latter of which was also observed in the form of STAT3^T716S727^ by TiO_2_ enrichment (Figure 5A). Crucially, sSH2 enrichment indicated increased phosphorylation of GADS^Y45^ (Figure 5A), an Itk substrate site that is phosphorylated exclusively within the LAT-SLP76 complex. ^74^ The primary function of SLP76 complex is to activate critical signaling pathways downstream of the TCR, including p38/Jnk, MAPK/Erk1/2 and subsequent pathways regulating the actin cytoskeleton. ^15,75^ The guanine exchange factor, Vav1, is a component of the SLP76 complex known to activate Rho/Rac GTPases after TCR stimulation.^15,76^ sSH2 pTyr enrichment revealed that Vav1^Y791^, a pTyr site known to increase in response to *α*-CD3 and *α*-CD28 treatment, significantly increased upon CAR activation. However, the physiological relevance of this site is not known.^76^ We additionally observed a significant positive enrichment of Jnk1 kinase substrate signatures in our TiO_2_ data, which supports a model of active Vav1 on the SLP76 complex (Figure 4A). Collectively, these data suggested that the LAT-SLP76 complex formed after CD19-CAR stimulation of the CAR T cells and contributed to signal transduction.

### CD19-CAR T cell Stimulation and Abl Kinase Activation

Abl family kinases (Abl1/2) are non-receptor tyrosine kinases necessary for canonical TCR signaling and for their regulation of the actin cytoskeleton through the SLP76 complex. ^77–79^ Activation of Abl family kinases can occur either by phosphorylation by Src family kinases (at tyrosines Abl1^Y412^, Abl2^Y439^) or by autophosphorylation of a specific residues on each Abl family kinase. ^77,80^ sSH2 enrichment revealed several phosphorylation events specific to Abl1/2 (Figure 5A). In particular, we observed a significant increase in the abundance of Abl1 kinase substrate sites, namely PLC*γ*1^Y771^, Zap70^Y319^, WASp^Y291^, and Nck1^Y105^.^78,81–83^ PTM-SEA of the sSH2 enrichment data supported Abl1 activation by CD19-CAR in CAR T cells as Abl1 kinase substrate signatures showed significant positive enrichment between 0- and 2-minutes after CD19-CAR stimulation (Supplementary Figure 6A). In addition to the activation of Abl1, Abl2^Y439^ phosphopeptide abundance and the Abl2 kinase substrate site WASl^Y256^ significantly increased (Figure 5A). Phosphorylation of WASp by Abl1 promotes WASp-Arb2/3 complex formation and subsequent regulation of actin polymerization. ^75,82^ Further, phosphorylation of the WASp homolog WASl by Abl2 was shown to regulate actin cytoskeletal dynamics with Pak2 after Pak2^S141^ autophosphorylation, ^84–86^ a phosphorylation site which significantly increased in our TiO_2_ data (Figure 5A). Interestingly, we observed a significant increase in the abundance of TNFRSF9^Y222^, a PhosphoNET^65^ predicted Abl1/2 kinase substrate site in the 4-1BB costimulatory domain of our CD19-CAR (Y378 of our CAR; Figure 5A). The importance of phosphorylation in 4-1BB signaling remains unclear, ^87^ and TNFRSF9^Y222^ was not observed by previous phosphoproteomic studies to our knowledge. Collectively, our data supported the hypothesis that Abl1/2 becomes activated by CD19-CAR stimulation, participates in actin cytoskeletal rearrangement, and may have roles in the phosphorylation of the 4-1BB domain within the CD19-CAR.

### Downregulation of Phosphorylation in Raji B cells

SILAC “Heavy“labeling of Raji B cells (Figure 2) allowed for phosphorylation changes to be associated with either CD19-CAR T cells or Raji B cells. Changes in Raji B cell pTyr peptide abundance were not expected because the CD19-CAR targets CD19, a B cell costimulatory receptor with no known extracellular ligand. ^88^ sSH2 enrichment revealed that pTyr peptide abundance broadly decreased in Raji B cells in response to coculture with CD19-CAR T cells (Figure 6). The abundance of several pTyr peptides corresponding to BCR signaling-related proteins (Lyn^Y306^, Lyn^Y316^, Btk^Y223^, Pag1^Y163^, Pag1^Y181^, Pag1^Y317^, Rab7^Y183^, Hgal^S69Y86^, Ship1^Y914^, Ship2^Y886^, Dok3^Y398^, Scimp^Y131^) and actin cytoskeleton-related proteins (Axna6^Y30^, Ezrin^Y116^, Fli^Y737^, Cofilin1^Y68^, Lcp1^Y28^, Moesin^Y116^) decreased significantly after coculture (Figure 6. Only a subset of these sites were previously character-ized, namely Btk^Y223^, an autophosphorylation site, and the Lyn substrate site, HgalS69Y86. Observed cryptic phosphopeptides containing Lyn^Y306^ and Lyn^Y316^ also significantly decreased, though Lyn^Y316^ was predicted to be a Src family kinase substrate by PhosphoNET. ^65^ The abundance of Syk^Y323^, a kinase activating phosphorylation site,^89^ also decreased though this decrease was not considered statistically significant (*q* = 0.629). PTM-SEA of Raji B cell peptides confirmed a general trend of decreased phosphopeptide abundance after coculture with CD19-CAR T cells (Supplementary Figure 7, 10). Overall, our analyses of phosphopeptide abundance revealed a novel decrease in Raji B cell phosphopeptide abundance after coculture of CD19-targeting CAR T cells and CD19-expressing Raji B cells.

**Figure 6:**
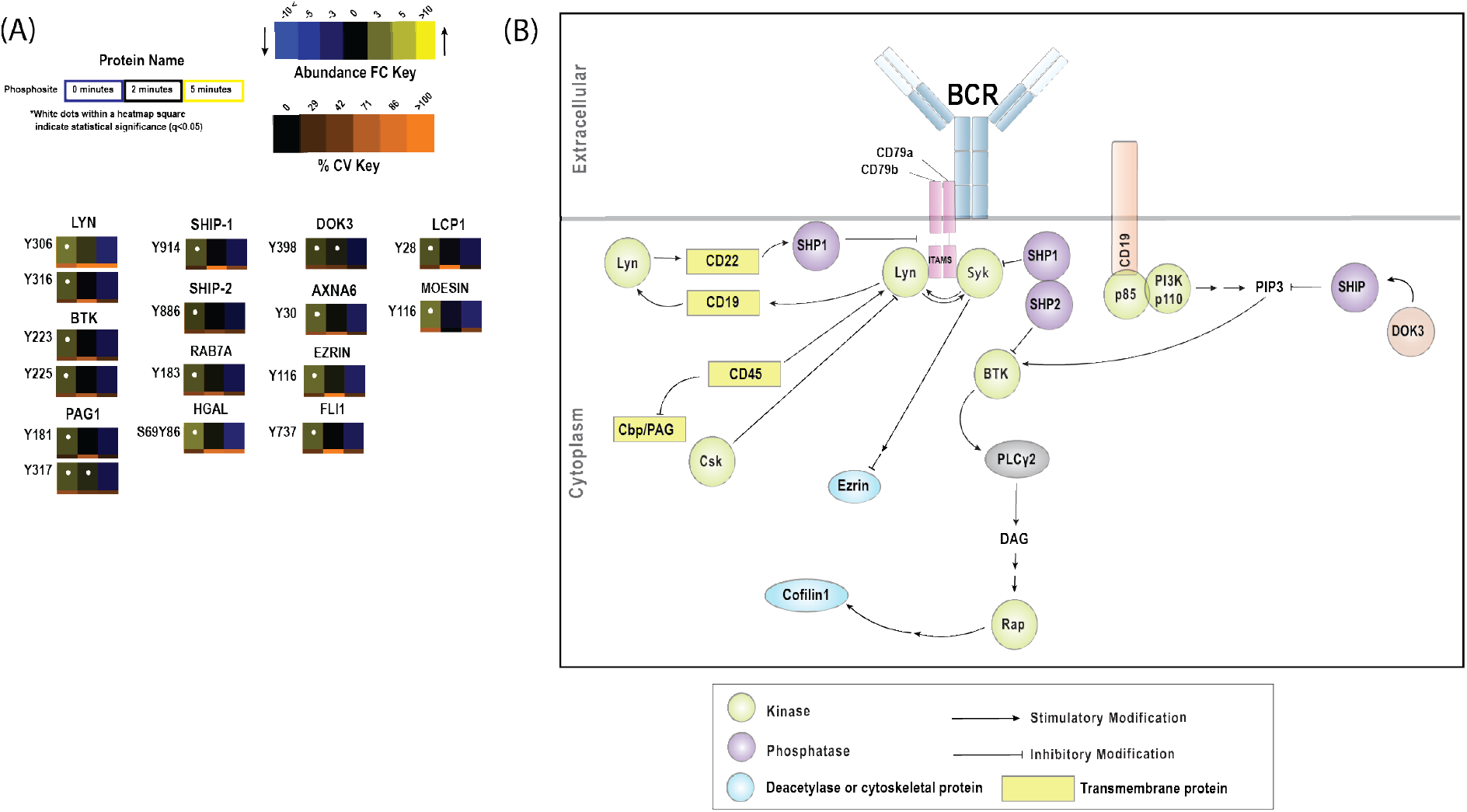
Interactions between the CD19-CAR and CD19 on Raji B cells reduced phosphopeptide abundance from proteins involved in the B cell receptor pathway in Raji B cells. Abundance heat maps for significantly changing proteins related to the BCR signaling pathway. For heat map block color information, refer to the Abundance FC Key or the Materials and Methods. (B) A pathway map depicting proteins involved in the BCR pathway that were observed in our sSH2 pTyr enrichment data.

**Figure 7:**
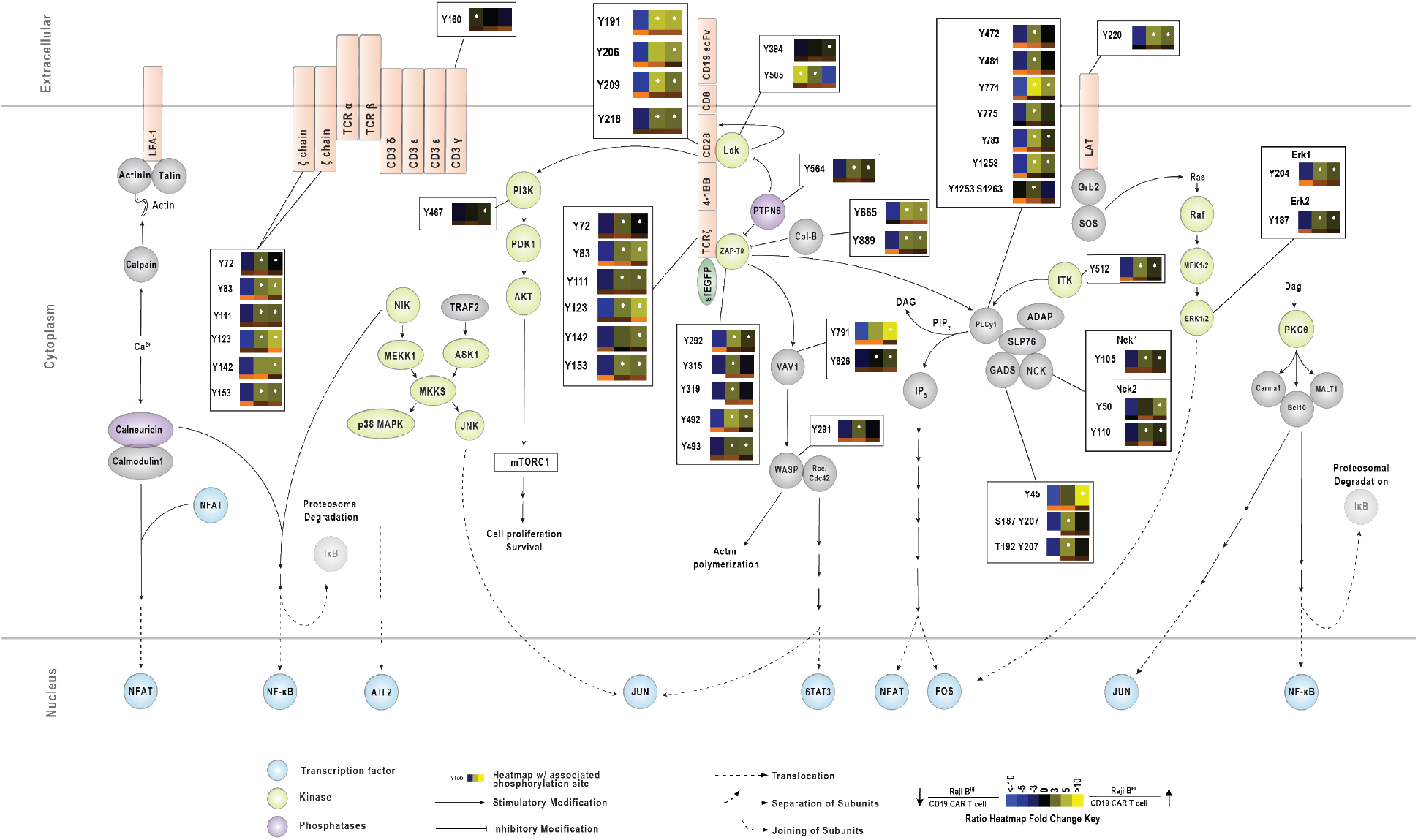
A full representation of signaling from the CD19-CAR. Heat maps representing phosphopeptide abundance changes for a subset of phosphopeptides we observed in the CD19-CAR T cells that are critical or unique to signal transduction from our CD19-CAR.

## Conclusion

Examining phosphotyrosine signaling after physiologically relevant CAR stimulation is necessary to ensure the safety and efficacy of CARs. Through coculture of a third generation CD19-CAR T cells with Raji B cells, we showed that an evaluation of signal transduction in both the CAR-expressing cells and the CAR-targeting cells is possible by using SILAC-based phosphoproteomics coupled with phosphopeptide enrichment. Our data suggest that, in CD19-CAR T cells, CAR stimulation by Raji B cells promotes strong signal transduction through canonical TCR- and CAR-specific pathways. In Raji B cells, we observed a decrease in global phosphorylation in response to CAR interaction, which may be critical for clinical efficacy of CD19-targeting CARs. With an in depth understanding of signal transduction in CAR-expressing T cells and ligand-expressing target cells, researchers will be able to make informed design choices to reduce off-target effects and increase efficacy of CARs in patients.

## Supporting information

Supplementary Table 1 SILAC labeling test

Supplementary Table 2 sSH2 peptide all data

Supplementary Table 3 TiO2 peptide all data

Supplementary Table 4 sSH2 unique peptides

Supplementary Table 5 TiO2 unique peptides

Supplementary Table 6 Lck Western Quantification

## Acknowledgements

The authors wish to thank Dr. Ricky Edmondson and Dr. Samuel G. Mackintosh from University of Arkansas for Medical Sciences (UAMS) for collecting the proteomic data. Arthur R. Salomon was supported by the NIH grants R01AI083636, P20GM121293, and R24GM137786. Xiaolei Su was supported by the Charles H. Hood Foundation Child Health Research Awards, the Rally Foundation and Bear Necessities Foundation a Collaborative Pediatric Cancer Research Awards Program, the Rally Young Scholar Program, the Frederick A. DeLuca Foundation Award, the Yale SPORE in skin cancer DRP Award (CA121974), and the NIGMS MIRA (R35) program (GM138299).

## TOC Graphic

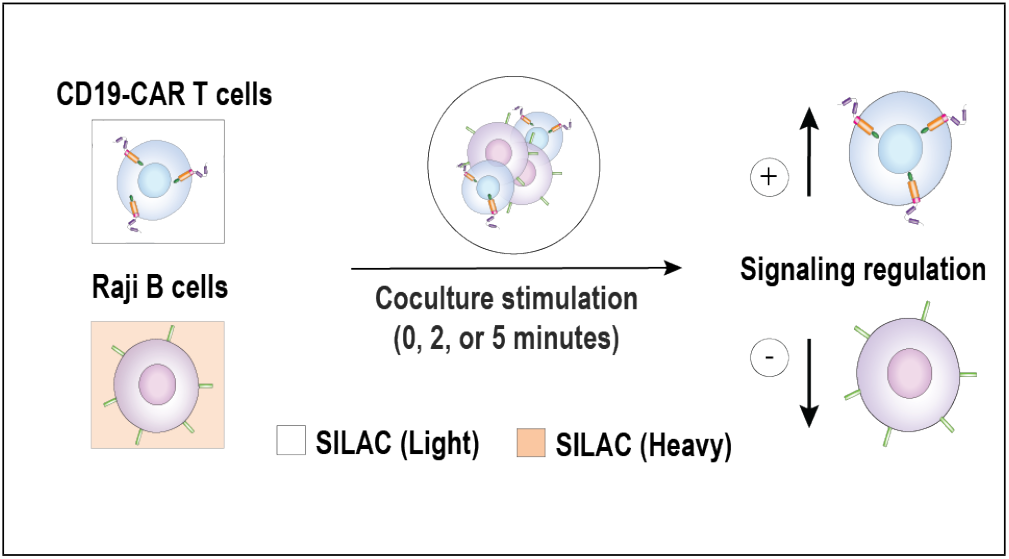

