## Supplementary Figures for "SILAC phosphoproteomics reveals unique signaling circuits in CAR-T cells and the inhibition of B cell-activating phosphorylation in target cells"

Supplementary Table 1: Complete list of peptides identified after heavy isotope labeling of Raji B cells and their labeling status. Included are the protein name, assigned peptide sequence, MOWSE score, mass error, SILAC labeling status, heavy proline inclusion, isolated mass, scan number, charge state, UNIPROT accesssion number, UNIPROT gene name, NCBI gi and HPRD accession number for each peptide identified. In total, we observed 5,038 heavy, 11 light and 5 mixed isotope labeled peptides after 8 doublings of Raji B cells in SILAC media.

Supplementary Table 2: Complete list of all peptides identified from pTyr enrichment after coculture of CD19-CAR T cells and Raji B cells. All LC-MS/MS data corresponding to each replicate and each condition are listed as separated worksheets. The assigned names are listed, as well as the position of phosphorylation in the assigned peptide sequence. The star character (*) represents a phosphorylated amino acid, the pound character (#) represents methionine oxidation, and the period characters (.) represent the ends of the identified peptide. For each peptide, the MOWSE score, mass error, SILAC labeling status, heavy proline inclusion, isolated mass, scan number, charge state, UNIPROT accesssion number, UNIPROT gene name, NCBI gi and HPRD accession number are reported.

Supplementary Table 3: Complete list of all peptides identified from $TiO_2$ phosphopeptide enrichment after coculture of CD19-CAR T cells and Raji B cells. All LC-MS/MS data corresponding to each replicate and each condition are listed as separated worksheets. The assigned names are listed, as well as the position of phosphorylation in the assigned peptide sequence. The star character (*) represents a phosphorylated amino acid, the pound character (#) represents methionine oxidation, and the period characters (.) represent the ends of the identified peptide. For each peptide, the MOWSE score, mass error, SILAC labeling status, heavy proline inclusion, isolated mass, scan number, charge state, UNIPROT accession number, UNIPROT gene name, NCBI gi and HPRD accession number are reported.

Supplementary Table 4: List of unique peptides identified from pTyr enrichment after coculture of CD19-CAR T cells and Raji B cells. For each peptide, the SIC peak areas for each replicate measurment and time point, $q$-values, accession numbers and KEGG/GO annotations are provided. Phosphopeptide assignments were filtered by a MOWSE score greater than 20, forward hits only, and assigned an Ascore.

Supplementary Table 5: List of unique peptides identified from $TiO_2$ phosphopeptide enrichment after coculture of CD19-CAR T cells and Raji B cells. For each peptide, the SIC peak areas for each replicate measurment and time point, $q$-values, accession numbers and KEGG/GO annotations are provided. Phosphopeptide assignments were filtered by a MOWSE score greater than 20, forward hits only, and assigned an Ascore.

Supplementary Table 6: Raw signal intensity measurements from the $\alpha$-Lck and Total Protein Western blots and corresponding statistics. Signal intensities were measured using the LiCor Image Studio system and exported to Microsoft Excel. Statistical calculations were performed in Python 3 using the script generate_lck_abundance_graph.py (See Supplementary Folder 1) and exported to Microsoft Excel. The two Excel files were combined to create this table.

Supplementary Folder 1: All Python 3 code used to run PTM-SEA, generate PTM-SEA heatmaps, generate KEGG/GO membership bar charts and perform statistics on Lck Western blots. Included in this folder are all scripts (and helper scripts) used for PTM-SEA, KEGG/GO and Lck Western blot graph generation, the required input files for each script, and the raw output from each script. For a more detailed description of use for these scripts, refer to the docstring of each script or our GitHub page (https://github.com/drsalomon/griffith_callahan_2021_CART_code).

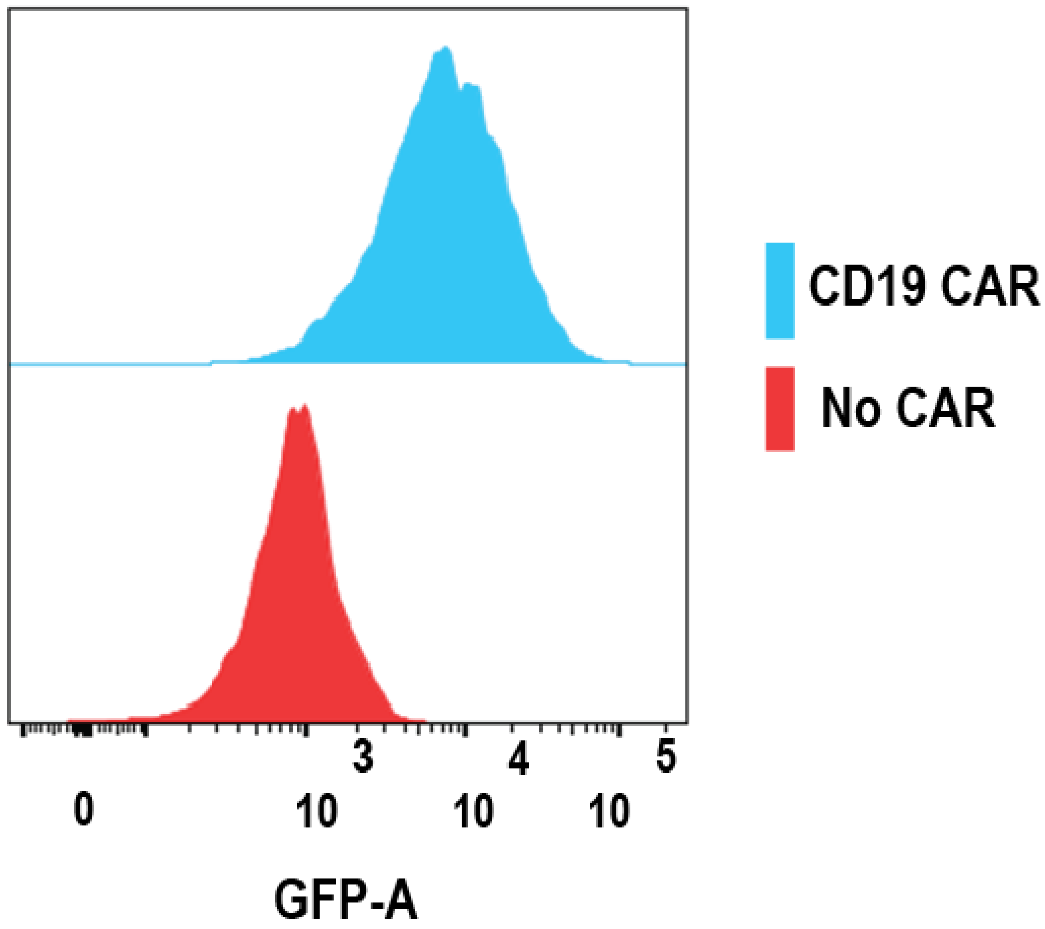


Supplementary Figure 1: Fluorescence activated cell sorting (FACS) determines the population of Jurkat T-cells expressing the CD19 (CD28/4-1BB/CD3ζ/sGFP) CAR construct.

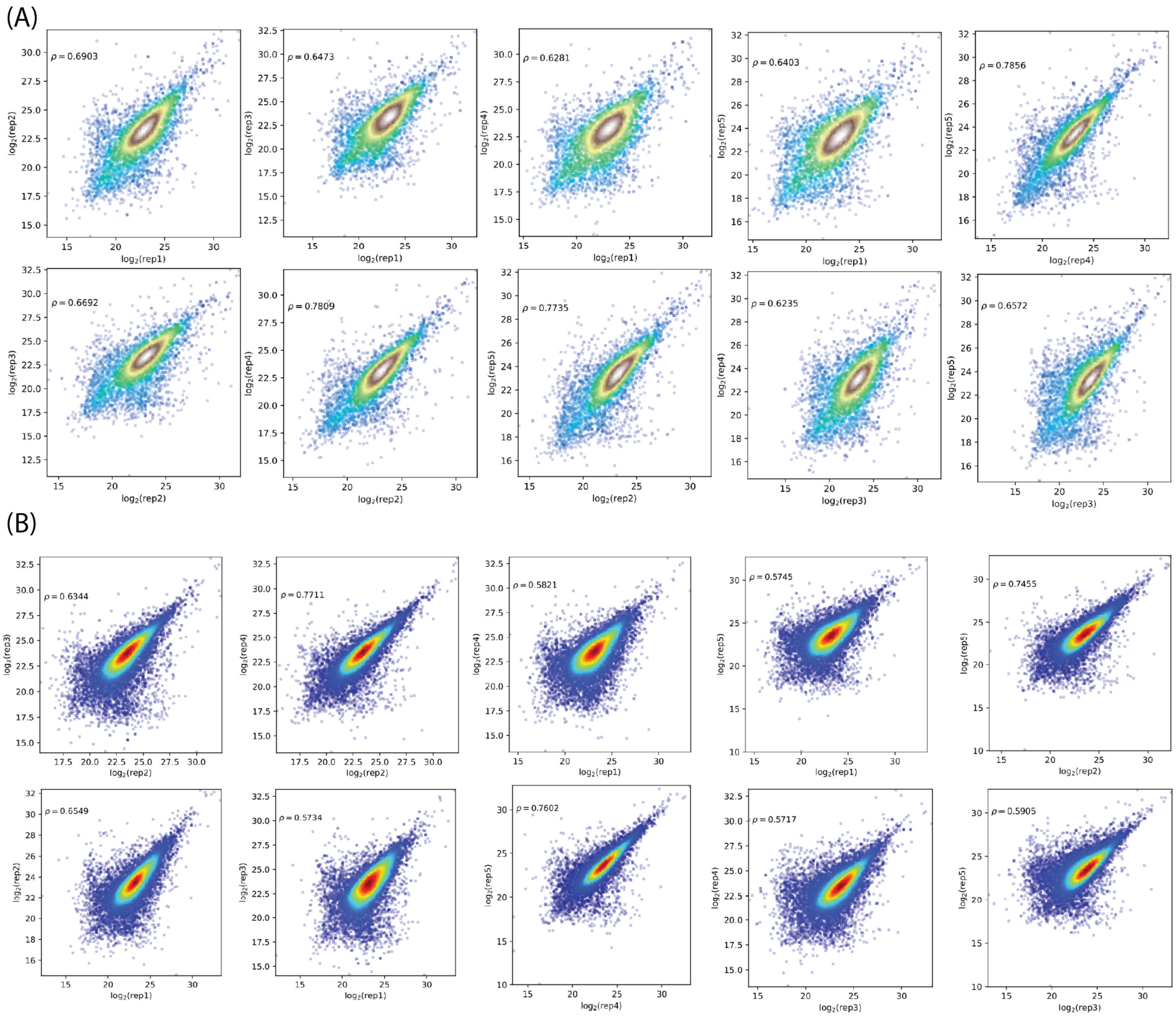


Supplementary Figure 2: Pairwise comparisons between replicates of $\log_2$ transformed phosphopeptide abundances with the same assigned sequence and charge state obtained by $TiO_2$ phosphoenrichment. (A) Phosphopeptide abundances measured from $CD19^{HI}$ Raji B-cells and (B) Phosphopeptide abundances measured from CD19 CAR T-cells. For each comparison, we used the Pearson's correlation coefficient (denoted $\rho$) as an estimate of the linear correlation between replicates and thus a measure of reproducibility between replicates.

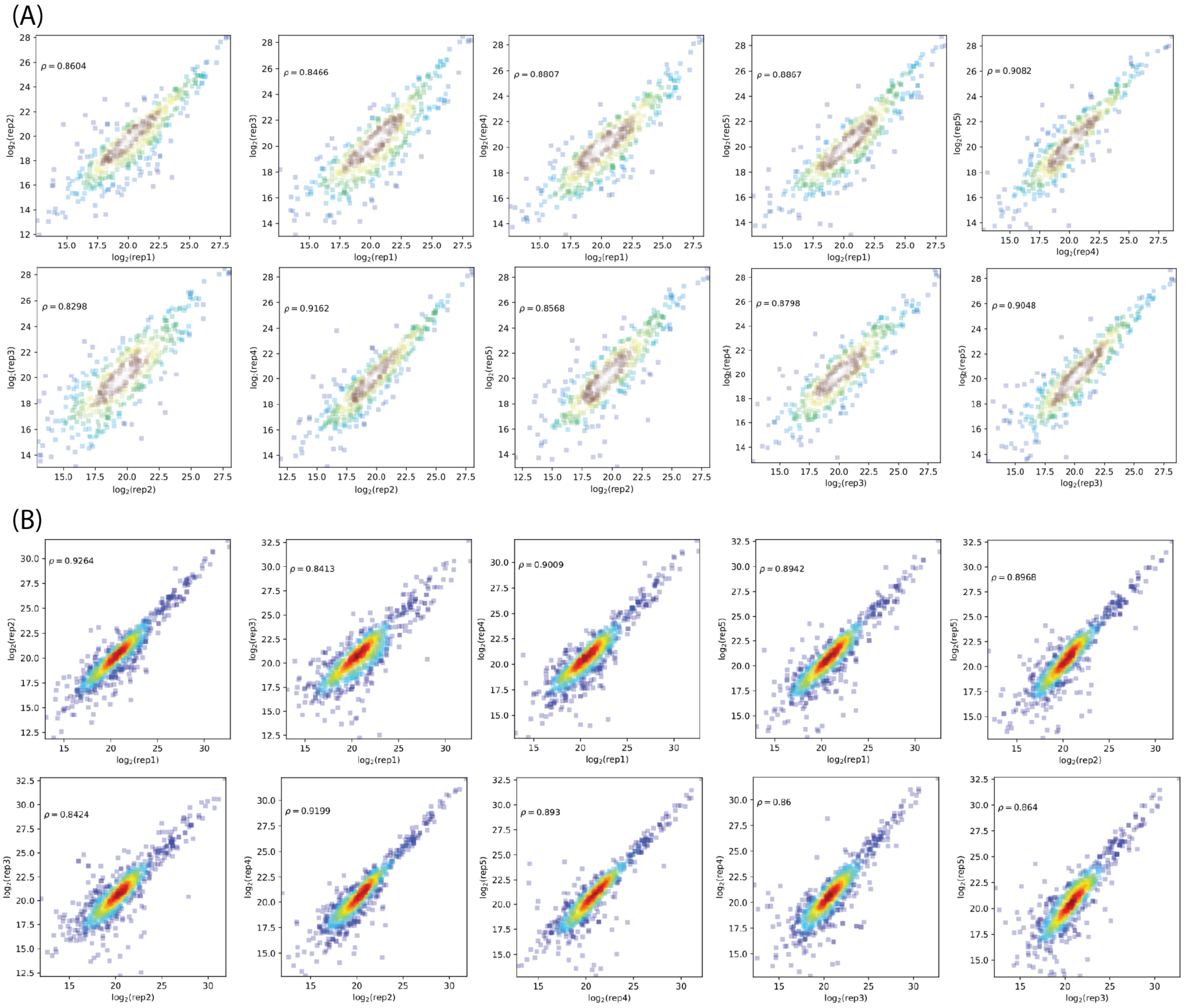

Supplementary Figure 3: Pairwise comparisons between replicates of $\log_2$ transformed phosphopeptide abundances with the same assigned sequence and charge state obtained by sSH2 phosphotyrosine enrichment. (A) Phosphopeptide abundances measured from CD19$^{HI}$ Raji B-cells and (B) Phosphopeptide abundances measured from CD19 CAR T-cells. For each comparison, we used the Pearson's correlation coefficient (denoted $\rho$) as an estimate of the linear correlation between replicates and thus a measure of reproducibility between replicates.

Supplementary Figure 4: A list of all abundance heat maps for unique phosphopeptides identified by pTyr enrichment after coculture of CD19-CAR T cells and Raji B cells. Included are the protein name, gene name, phosphosite(s), abundance heat maps with significance indicators (if applicable), maximum Ascore, MOWSE score and assigned peptide sequence.

Supplementary Figure 5: A list of all abundance heat maps for unique phosphopeptides identified by $TiO_2$ phosphopeptide enrichment after coculture of CD19-CAR T cells and Raji B cells. Included are the protein name, gene name, phosphosite(s), abundance heat maps with significance indicators (if applicable), maximum Ascore, MOWSE score and assigned peptide sequence.

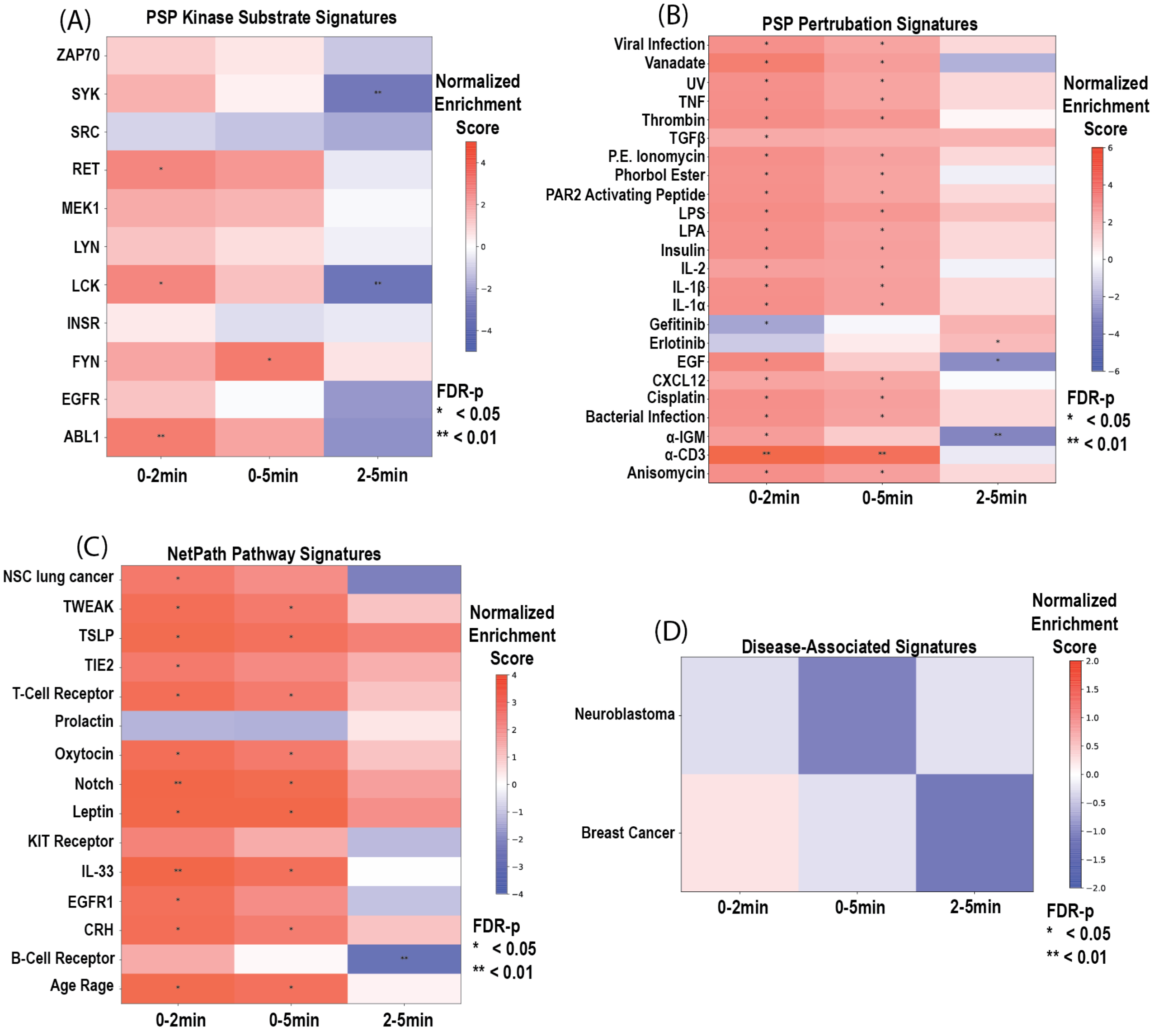


Supplementary Figure 6: PTM-SEA on CD19 CAR T-cell originating phosphopeptides observed in the sSH2 enrichment data. Signature sets ($S$) in the PTM signature database (PTMsigDB, version 1.9.0) are grouped by (A) PhosphoSitePlus annotated kinase substrate modifications, (B) PhosphoSitePlus annotated modifications that change in response to perturbagens, (C) NetPath annoated modifications associated with a particular signaling pathway, and (D) modifications associated with the progression of a disease. Red colors indicate positive correlations and blue colors indicate negative correlations between signature-associated phosphopeptide abundance changes observed between time points. On each heatmap block, one star (*) indicates an $q < 0.05$ and two stars (**) indicates an $q < 0.01$.

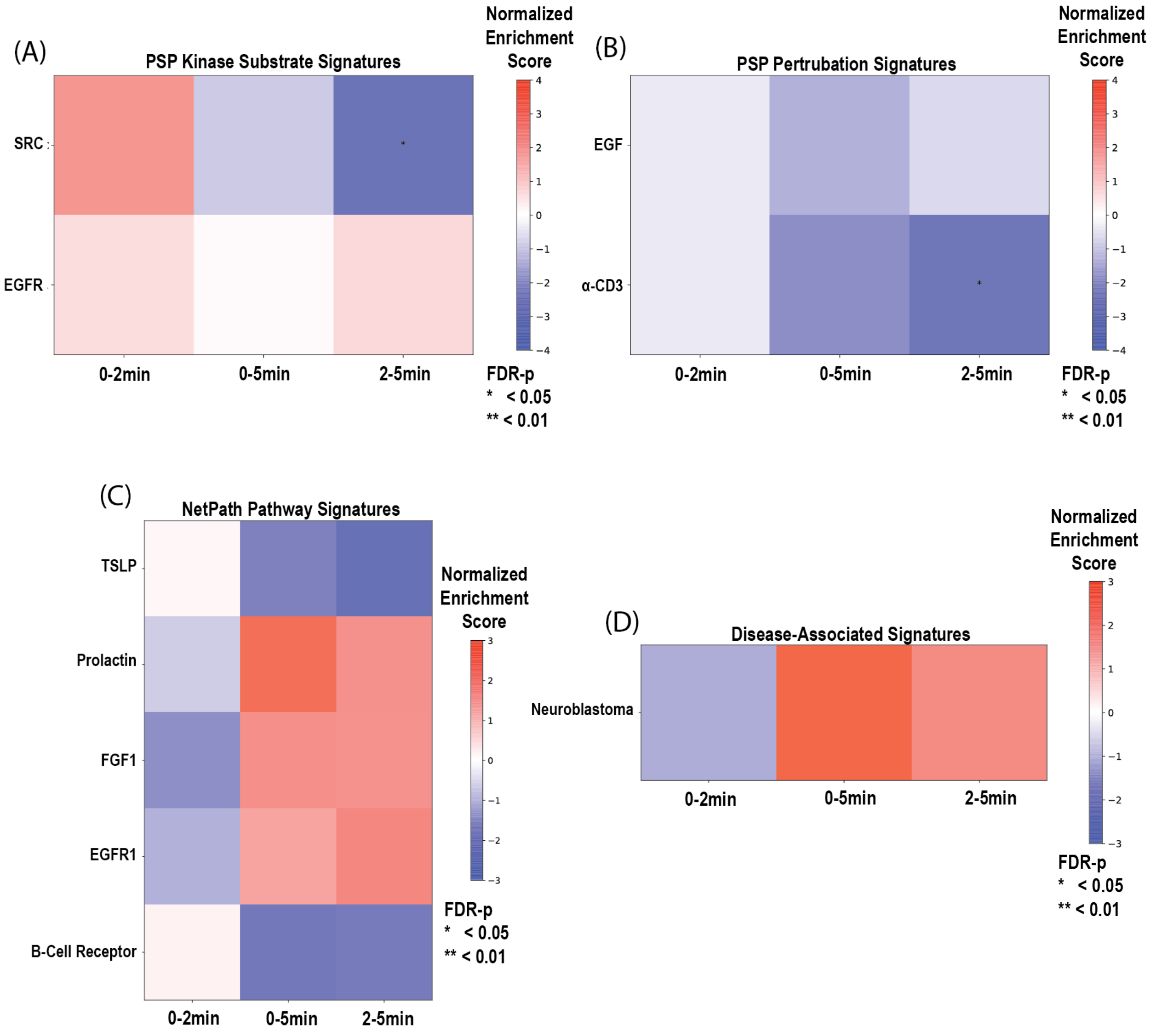


Supplementary Figure 7: PTM-SEA on CD19$^{\mathrm{HI}}$ Raji B-cell originating phosphopeptides observed in the sSH2 enrichment data. Signature sets ($S$) in the PTM signature database (PTMsigDB, version 1.9.0) are grouped by (A) PhosphoSitePlus annotated kinase substrate modifications, (B) PhosphoSitePlus annotated modifications that change in response to perturbagens, (C) NetPath annoated modifications associated with a particular signaling pathway, and (D) modifications associated with the progression of a disease. Red colors indicate positive correlations and blue colors indicate negative correlations between signature-associated phosphopeptide abundance changes observed between time points. On each heatmap block, one star (*) indicates an $q < 0.05$ and two stars (**) indicates an $q < 0.01$.

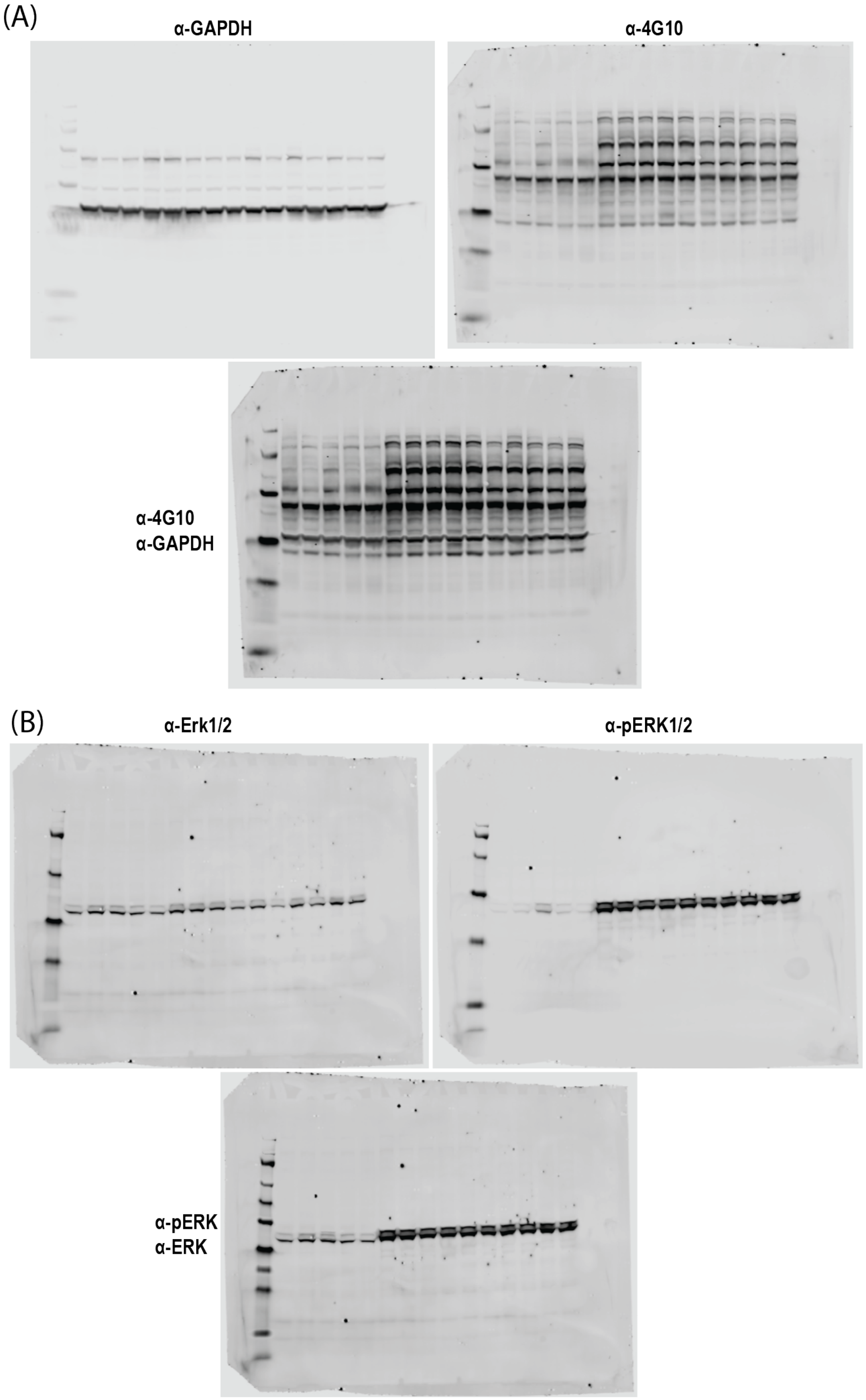


Supplementary Figure 8: Full, uncropped Western blots for (A) $\alpha$-pTyr (4G10) and (B) $\alpha$-pErk. Each full blot includes the loading control [$\alpha$-GAPDH and $\alpha$-Erk for (A) and (B), respectively], the target, and the merged blots.

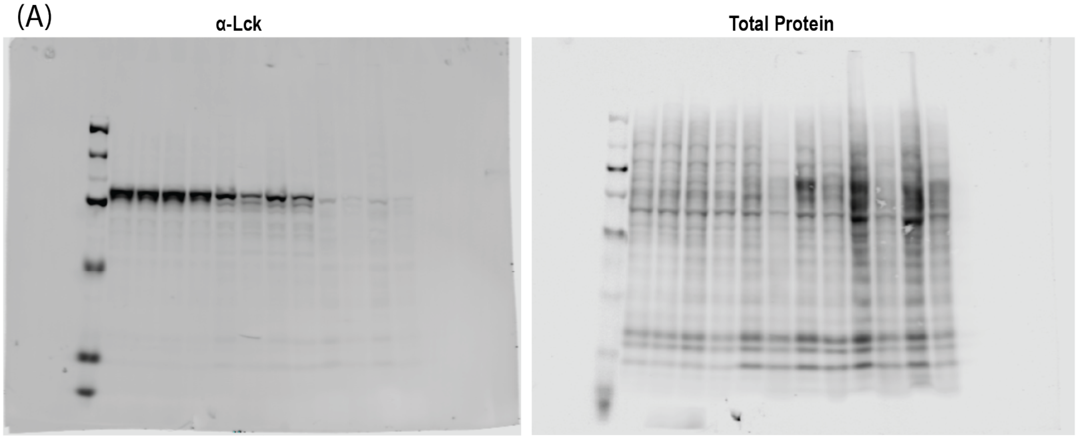


Supplementary Figure 9: (A) Full, uncropped Western blots showing the abundance of Lck protein and total protein abundance in JE6 (Lanes 2-5), CD19-CAR T cells (Lanes 6-9) and Raji B cells (Lanes 10-13). (B) Quantification of Lck abundance for the Western blots in (A). The $y$-axis represents Lck protein abundance normalized to the total protein loaded into the lane (Lck/Total Protein). Black bars with significance indicators above them show the results of a two-tailed Student's T-test performed between the two connected groups. For the raw data and statistical output, refer to Supplementary Table 6.

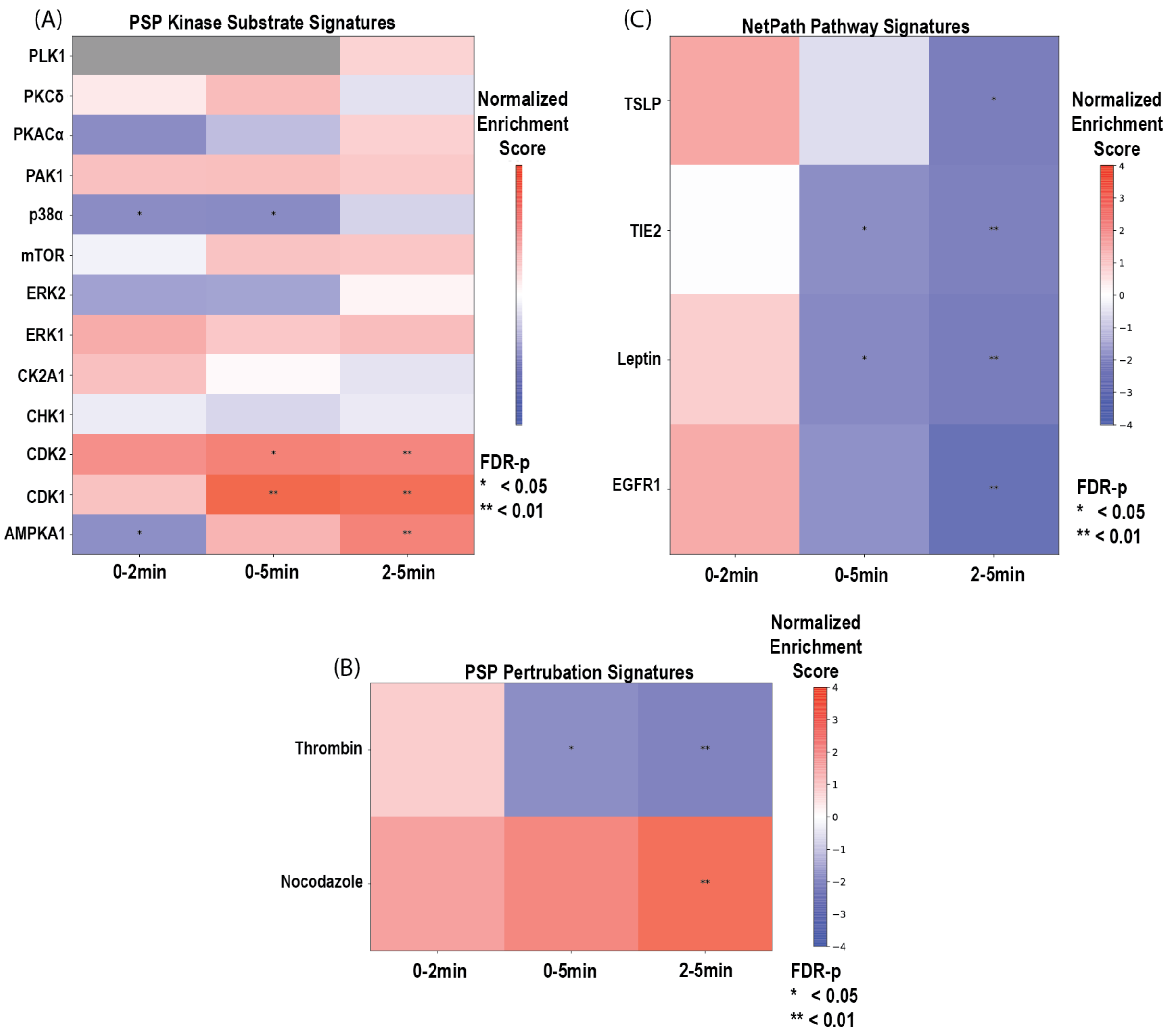


Supplementary Figure 10: PTM-SEA on phosphopeptides originating from CD19$^{HI}$ Raji B-cell observed in the $TiO_2$ enrichment data. Signature sets ($S$) in the PTM signature database (PTMsigDB, version 1.9.0) are grouped by (A) PhosphoSitePlus annotated kinase substrate modifications, (B) PhosphoSitePlus annotated modifications that change in response to perturbagens and (C) NetPath annoated modifications associated with a particular signaling pathway. Red colors indicate positive correlations and blue colors indicate negative correlations between signature-associated phosphopeptide abundance changes observed between time points. On each heat map block, one star (*) indicates an $q < 0.05$ and two stars (**) indicates an $q < 0.01$.
