## Supplementary Figure 4 for "SILAC phosphoproteomics reveals unique signaling circuits in CAR-T cells and the inhibition of B cell-activating phosphorylation in target cells"

Peak Area: <-10, -5, -3, 0, 3, 5, >10

%CV: 0, 29, 43, 71, 86, >100

**White dots**- Significant change in peptide abundance at 5 %FDR compared to the timepoint with the minimum peak area for a given PSM

| Protein Name | Gene | Phosphosites | Ascor | MOWSE | Sequence |
|---|---|---|---|---|---|
| A6RP | TWF2 | Y309 | 175.4 | 111.2 | K.KIEIGDGAELTAEFLY*DEVHPK.Q |
| ABL interactor 1 | ABI1 | Y213 | 22.0 | 50.0 | K.TLEPVKPPTVPNDY*MTSPAR.L |
| ABL interactor 2 | ABI2 | Y213 | 31.4 | 49.1 | R.TLEPVRPPVVPNDY*VPSPTR.N |
| Actin alpha, cardiac muscle | ACTB | Y93 | 42.4 | 27.6 | K.IWHHTFY*NELR.V |
| Actin alpha, cardiac muscle | ACTB | Y242 | -0.2 | 66.0 | R.SY*ELPDGQVITIGNER.F |
| Actin binding LIM protein 1 | ABLIM1 | Y373 | 11.5 | 19.8 | R.TSSESIYSRPGSSIPGSPGHTIY*AK.V |
| Actin binding LIM protein 1 | ABLIM1 | Y406 | 12.3 | 15.9 | K.AIYDIERPDLITY*EPFYTSGYDDK.Q |
| Actin gamma 1 | ACTB | Y294 | 31.5 | 102.9 | K.DLY*ANTVLSGGTTM#YPGIADR.M |
| Actin gamma 1 | ACTB | Y198 | 30.2 | 43.3 | R.GY*SFTTTAER.E |
| Actin gamma 1 | ACTB | Y218 | 48.5 | 47.0 | K.LCY*VALDFEQEM#ATAASSSSLEK.S |
| Actin related protein 10 | ACTR10 | Y4 | 105.1 | 119.3 | M.PLY*EGLGSGGEK.T |
| Actin related protein 3 | ACTR3 | Y109 | 56.5 | 27.6 | R.AEPEDHY*FLLTEPPLNTPENR.E |
| Acyl CoA synthetase long chain family member 4 | ACSL4 | Y582 | 26.0 | 63.4 | K.LQAGEY*VSLGK.V |
| ADAM9 | ADAM9 | Y769 | 100.0 | 26.5 | R.EVPIY*ANR.F |
| Adenosine deaminase | ADA | Y29 | 13.0 | 46.5 | K.VELHVHLDGSIKPETILY*YGR.R |
| Aldolase 3 | ALDOC | Y358 | 35.4 | 56.8 | K.YEGSGEDGGAAAQSLY*IANHAY.- |
| ALS2CR7 | CDK14 | Y63 | 16.9 | 34.4 | K.LGEGSY*ATVYK.G |
| Angiotensin II receptor associated protein | AGTRAP | Y133 | 10.9 | 39.7 | R.SAY*QTIDSAEAPADPFAVPEGR.S |
| Ankyrin repeat domain 13 | ANKRD13 | Y485 | 136.4 | 77.7 | R.NVHLQDEDY*EIMQFAIQQSLLESSR.S |
| Annexin II | ANXA2 | Y24 | 12.2 | 80.8 | K.LSLEGDHSTPPSAY*GSVK.A |
| Annexin VI | ANXA6 | Y30 | 38.4 | 64.7 | R.GSIHDFPGFDPNQDAEALY*TAMK.G |
| Annexin VI | ANXA6 | Y302 | 22.5 | 20.1 | K.SLY*SM#IK.N |
| ARP2/3 protein complex subunit p21 | ARPC3 | Y47 | 13.9 | 62.3 | K.DTDIVDEAIY*YFK.A |
| Ataxin 2 domain protein | ATXN2L | Y349 | 100.0 | 24.1 | K.Y*IPLPQR.V |
| ATP binding cassette 50 | ABCF1 | Y378 | 100.0 | 31.1 | K.VY*EELR.A |
| ATP citrate lyase | ACLY | Y682 | 74.7 | 68.6 | R.TTDGVY*EGVAIGGDRYPGSTFMDHVLR.Y |
| ATP citrate lyase | ACLY | Y131 | 100.0 | 55.0 | R.EGDY*VLFHHEGGVDVGDVDAK.A |
| ATP1A1 | ATP1A1 | Y260 | 16.9 | 30.9 | R.GIVVY*TGDR.T |
| BCR | BCR | Y177 | 100.0 | 50.9 | K.GHGQPGADAEKPFY*VNVEFHHER.G |
| BCR downstream signaling 1 | STAP1 | Y168 | 15.9 | 46.1 | K.EKPTEDY*VDVLNPM#PACFYTVSR.K |
| BH3 interacting domain death agonist | BID | Y54 | 40.7 | 44.3 | R.ELDALGHELPVLAPQWEGY*DELQTDGNR.S |
| BLK | BLK | Y501 | 100.0 | 23.6 | R.QY*ELQP.- |
| BLK | BLK | Y187 | 22.0 | 79.0 | R.CLDEGGY*YISPR.I |
| BLK | BLK | Y389 | 32.1 | 91.9 | R.IIDSEY*TAQEGAK.F |
| BLK | BLK | Y188 | 16.9 | 88.8 | R.CLDEGGYY*ISPR.I |
| Bruton`s tyrosine kinase | BTK | Y551 | 16.9 | 86.8 | R.YVLDDEY*TSSVGSK.F |
| Bruton`s tyrosine kinase | BTK | Y361 | 26.9 | 62.8 | K.HLFSTIPELINY*HQHNSAGLISR.L |
| Bruton`s tyrosine kinase | BTK | Y223 | 16.0 | 54.9 | K.VVALY*DYMPM#NANDLQLR.K |
| Bruton`s tyrosine kinase | BTK | Y225 | 7.3 | 54.7 | K.VVALYDY*MPM#NANDLQLR.K |
| Bruton`s tyrosine kinase | BTK | Y344 | 6.2 | 39.2 | R.HYVVCSTPQSQY*YLAEK.H |
| Bruton`s tyrosine kinase | BTK | Y334 | 55.5 | 48.1 | R.HY*VVCSTPQSQYYLAEK.H |
| C20ORF77 protein | RPRD1B | Y165 | 5.2 | 61.8 | R.TFQQIQEEEDDDYPGSY*SPQDPSAGPLLTEELIK.A |
| C4orf14 protein | NOA1 | Y77 | 95.3 | 31.0 | R.FLFPEY*ILDPEPQPTR.E |
| Calmodulin 1 | CALM2 | Y100 | 58.0 | 85.0 | R.VFDKDGNGY*ISAAELR.H |
| Calmodulin 1 | CALM3 | Y139 | 50.6 | 43.3 | R.EADIDGDGQVNY*EEFVQMMTAK.- |
| Calumenin | CALU | Y47 | 3.0 | 26.2 | K.VHNDAQSFDY*DHDAFLGAEEAK.T |
| Cask-interacting protein 2 | CASKIN2 | Y253 | 27.4 | 123.7 | R.NTY*NQTALDIVNQFTTSQASR.E |
| CBL | CBL | Y674 | 24.1 | 52.1 | K.IKPSSSANAIY*SLAARPLPVPK.L |
| CBL | CBL | S667Y674 | 14.0 | 34.7 | K.IKPS*SSANAIY*SLAARPLPVPK.L |
| CblB | CBLB | Y665S672 | 32.0 | 37.8 | K.VFSNGHLGSEEY*DVPPRLS*PPPPVTTLLPSIK.C |
| CblB | CBLB | T885Y889 | 7.9 | 26.9 | K.TNRT*SQDY*DQLPSCSDGSQAPARPPKPRPR.R |
| CblB | CBLB | Y889 | 30.5 | 33.0 | R.TSQDY*DQLPSCSDGSQAPARPPKPRPR.R |
| CblB | CBLB | Y665 | 9.2 | 16.8 | K.VFSNGHLGSEEY*DVPPR.L |
| CD19 | CD19 | Y508 | 100.0 | 42.2 | R.GILY*AAPQLR.S |
| CD19 | CD19 | Y348 | 28.4 | 41.5 | K.VTPPPGSGPQNQY*GNVLSLPTPTSGLGR.A |
| CD2 associated protein | CD2AP | Y361 | 17.8 | 24.4 | K.Y*FSLKPEEK.D |
| CD28 | | Y191T195 | 21.1 | 27.7 | R.LLHSDY*MNM#T*PR.R |
| CD28 | | Y191 | 19.3 | 40.1 | R.LLHSDY*MNMTPR.R |
| CD28 | | Y209 | 37.1 | 21.1 | R.KHYQPY*APPR.D |
| CD28 | | S189Y191 | 67.1 | 31.8 | R.LLHS*DY*M#NM#TPR.R |
| CD28 | | Y218 | 100.0 | 35.7 | R.DFAAY*R.S |
| CD28 | | Y206 | 27.3 | 19.6 | R.KHY*QPYAPPR.D |

Peak Area: <-10, -5, -3, 0, 3, 5, >10

%CV: 0, 29, 42, 71, 86, >100

**White dots-** Significant change in peptide abundance at **5** %FDR compared to the timepoint with the minimum peak area for a given PSM

CarT: 0m, 2m, 5m — RajiB: 0m, 2m, 5m

| Protein Name | Gene | Phosphosites | Ascor | MOWSE | Sequence |
|---|---|---|---|---|---|
| **CD3 gamma** | CD3G | Y160 | **80.6** | **50.1** | K.QTLLPNDQLY*QPLKDR.E |
| **CD3 gamma** | CD3G | Y171 | **13.9** | **13.5** | R.EDDQY*SHLQGNQLR.R |
| **CD31** | PECAM1 | Y713 | **27.6** | **61.0** | K.DTETVY*SEVR.K |
| **CD37** | CD37 | Y274 | **100.0** | **36.4** | R.NLDHVY*NR.L |
| **CD3E** | CD3E | Y199 | **39.9** | **49.3** | R.DLY*SGLNQR.R |
| **CD3E** | CD3E | Y188 | **100.0** | **41.2** | K.ERPPPVPNPDY*EPIRK.G |
| **CD5** | CD5 | S439Y453 | **24.1** | **24.2** | R.S*HAENPTASHVDNEY*SQPPR.N |
| **CD5** | CD5 | Y453 | **25.2** | **18.3** | R.SHAENPTASHVDNEY*SQPPR.N |
| **CD7** | CD7 | Y222 | **38.2** | **38.3** | R.DKNSAACVVY*EDMSHSR.C |
| **CD84** | CD84 | Y262 | **39.8** | **57.7** | K.TIY*TYIMASR.N |
| **CD84** | CD84 | Y299 | **32.3** | **94.6** | K.EEPVNTVY*SEVQFADK.M |
| **CD84** | CD84 | Y324 | **12.1** | **63.9** | K.ASTQDSKPPGTSSY*EIVI.- |
| **CD84** | CD84 | Y279 | **112.4** | **34.0** | R.IY*DEILQSK.V |
| **CDC2** | CDK1 | Y15 | **16.9** | **76.3** | K.IGEGTY*GVVYK.A |
| **CDC2** | CDK1 | T14Y15 | **69.8** | **74.1** | K.IGEGT*Y*GVVYK.A |
| **CDC2** | CDK1 | Y19 | **33.2** | **47.9** | K.IGEGTYGVVY*K.A |
| **CDC37** | CDC37 | Y298 | **58.0** | **38.4** | R.LGPGGLDPVEVY*ESLPEELQK.C |
| **CDV3 homolog** | CDV3 | Y190 | **39.3** | **83.9** | R.KTPQGPPEIY*SDTQFPSLQSTAK.H |
| **CDV3 homolog** | CDV3 | Y244 | **100.0** | **102.8** | K.LQLDNQY*AVLENQK.S |
| **CDV3 homolog** | CDV3 | Y95 | **16.9** | **21.1** | K.EVDY*SGLR.V |
| **Centaurin delta 2** | ARAP1 | Y497 | **16.4** | **56.5** | K.HY*SVVLPTVSHSGFLYK.T |
| **Centaurin delta 2** | ARAP1 | Y231 | **11.5** | **20.9** | R.LFPEFDDSDY*DEVPEEGPGAPAR.V |
| **Chaperonin containing T complex polypeptide 1 subunit 4** | CCT4 | Y269 | **20.7** | **30.7** | K.TDM#DNQIVVSDY*AQM#DR.V |
| **Chaperonin containing T complex polypeptide 1, subunit 7** | CCT7 | Y263 | **40.0** | **88.2** | R.VHTVEDY*QAIVDAEWNILYDK.L |
| **Chemokine orphan receptor 1** | ACKR3 | Y354 | **32.1** | **97.9** | R.VSETEY*SALEQSTK.- |
| **CHERP** | CHERP | Y894 | | **31.0** | K.GVGVALDDPY*ENYRR.N |
| **Chromosome 11 open reading frame 59** | LAMTOR1 | Y40 | **26.0** | **38.0** | K.ALNGAEPNY*HSLPSAR.T |
| **Chromosome 12 open reading frame 44** | ATG101 | Y164 | **100.0** | **26.1** | R.HEY*LPK.M |
| **Chromosome 9 open reading frame 86** | RABL6 | Y683 | **100.0** | **33.4** | R.HPGGGDY*EEL.- |
| **Chromosome condensation protein G** | NCAPG | Y929 | **10.9** | **17.7** | K.EVY*M#TPLR.G |
| **Cingulin** | CGN | Y105 | **101.1** | **24.5** | K.GANDQGASGALSSDLELPENPY*SQVK.G |
| **Clathrin, heavy polypeptide** | CLTC | Y634 | **93.7** | **54.8** | R.ALEHFTDLY*DIKR.A |
| **Clathrin, heavy polypeptide** | CLTC | Y430 | **52.2** | **24.9** | K.Y*ESLELCRPVLQQGR.K |
| **Clathrin, heavy polypeptide** | CLTC | Y1477 | **37.7** | **85.3** | K.SVNESLNNLFITEEDY*QALR.T |
| **Clathrin, heavy polypeptide** | CLTC | Y1096 | **100.0** | **45.4** | R.AY*EFAER.C |
| **Clathrin, heavy polypeptide** | CLTC | Y1487 | **18.4** | **56.7** | R.TSIDAY*DNFDNISLAQR.L |
| **CLNS1A** | CLNS1A | Y214 | **29.6** | **86.9** | R.TEDSIRDY*EDGM#EVDTTPTVAGQFEDADVDH.- |
| **Cofilin 1** | CFL1 | Y89 | **16.9** | **51.3** | R.YALYDATY*ETK.E |
| **Cofilin 1** | CFL1 | Y68 | **22.0** | **71.0** | K.EILVGDVGQTVDDPY*ATFVK.M |
| **Cofilin 1** | CFL1 | Y140 | **100.0** | **75.8** | K.HELQANCY*EEVKDR.C |
| **Complement receptor 2** | CR2 | Y1029 | **29.3** | **15.4** | R.EVYSVDPY*NPAS.- |
| **Coronin 1C** | CORO1C | Y301 | **33.4** | **45.3** | R.YFEITDESPY*VHYLNTFSSK.E |
| **Cortactin** | CTTN | Y421 | **55.8** | **61.0** | R.LPSSPVY*EDAASFK.A |
| **CRK** | CRK | Y190 | | **49.2** | K. Y*[illegible] |
| **CRKL** | CRKL | Y132 | **15.1** | **43.1** | R.TLY*DFPGNDAEDLPFKK.G |
| **Cyclin dependent kinase 5** | CDK5 | Y15 | **16.9** | **67.6** | K.IGEGTY*GTVFK.A |
| **Cyclin M3** | CNNM3 | Y301 | **19.2** | **15.0** | R.GGGDPY*SDLSK.G |
| **Cysteine string protein** | DNAJC5 | Y149 | **26.0** | **52.8** | K.APEGEETEFY*VSPEDLEAQLQSDER.E |
| **Cytoplasmic FMR1 interacting protein 1** | CYFIP2 | Y108 | **100.0** | **13.6** | R.VEIY*EK.T |
| **Cytoskeleton associated protein 1** | TBCB | Y98 | **82.3** | **76.1** | R.LGEY*EDVSR.V |
| **D4, zinc and double PHD fingers family 2** | DPF2 | Y172 | **55.5** | **54.6** | R.ILEPDDFLDDLDDEDY*EEDTPK.R |
| **DAPP1** | DAPP1 | Y139 | **9.1** | **38.7** | K.VEEPSIY*ESVR.V |
| **DDX20** | DDX20 | Y756 | **62.9** | **38.4** | R.LQTEAQEDDWY*DCHR.E |
| **DDX3** | DDX3X | Y69 | **16.9** | **65.4** | K.DKDAY*SSFGSR.S |
| **DDX3** | DDX3X | Y104 | **45.2** | **58.8** | R.SDY*DGIGSR.G |
| **DDX3** | DDX3X | Y283 | **100.0** | **56.0** | R.ELAVQIY*EEAR.K |
| **DDX3** | DDX3X | Y462 | **39.0** | **51.5** | K.KGADSLEDFLY*HEGYACTSIHGDR.S |
| **DDX3** | DDX3X | Y525 | **62.7** | **32.2** | K.HVINFDLPSDIEEY*VHR.I |
| **DDX3** | DDX3X | Y243 | **13.9** | **58.2** | K.TAAFLLPILSQIY*SDGPGEALR.A |
| **DDX3Y** | DDX3Y | Y103 | **45.2** | **57.4** | R.SDY*DGIGNR.E |
| **Decapping enzyme hDcp1b** | DCP1B | Y191 | **32.5** | **50.0** | K.KITSSSAIY*DNPNLIKPIPVKPSENQQQR.I |
| **Dedicator of cytokinesis 2** | DOCK2 | Y212 | **28.2** | **33.0** | K.DQPDYAM#Y*SR.I |

Peak Area: <-10, -5, -3, 0, 3, 5, >10

%CV: 0, 29, 42, 71, 86, >100

**White dots-** Significant change in peptide abundance at **5** %FDR compared to the timepoint with the minimum peak area for a given PSM

CarT (0m, 2m, 5m) RajiB (0m, 2m, 5m)

| Protein Name | Gene | Phosphosites | Ascor | MOWSE | Sequence |
|---|---|---|---|---|---|
| Diaphanous related formin 1 | DIAPH1 | Y374 | 10.9 | 93.1 | R.VQLNVFDEQGEEDSY*DLKGR.L |
| Diazepam binding inhibitor | DBI | Y46 | 31.5 | 67.8 | K.TKPSDEEM#LFIY*GHYK.Q |
| Diazepam binding inhibitor | DBI | Y49 | 9.1 | 34.6 | K.TKPSDEEM#LFIYGHY*K.Q |
| Dicer1, Dcr-1 homolog | DICER1 | Y1438 | 49.9 | 29.3 | K.EEADY*EDDFLEYDQEHIR.F |
| DNA methyltransferase 1 | DNMT1 | Y399 | 13.9 | 58.7 | K.LSIFDANESGFESY*EALPQHK.L |
| DNA topoisomerase I | TOP1 | Y723 | 100.0 | 14.6 | K.LNY*LDPR.I |
| Docking protein 1 | DOK1 | Y315 | 22.5 | 67.8 | R.IAPCPSQDSLY*SDPLDSTSAQAGEGVQR.K |
| Docking protein 1 | DOK1 | Y409 | 40.7 | 70.5 | R.VKEEGYELPYNPATDDY*AVPPPR.S |
| Docking protein 1 | DOK1 | Y362 | 109.7 | 40.6 | K.EDPIY*DEPEGLAPVPPQGLYDLPR.E |
| Docking protein 1 | DOK1 | Y296 | 38.9 | 49.4 | K.LPSPPGPQELLDSPPALY*AEPLDSLR.I |
| Docking protein 1 | DOK1 | Y449 | 90.6 | 72.8 | K.SHNSALY*SQVQK.S |
| Docking protein 2, 56kDa | DOK2 | Y299 | 100.0 | 98.1 | R.GQEGEY*AVPFDAVAR.S |
| Docking protein 3 | DOK3 | Y398 | 44.9 | 53.9 | R.ASGPPGNEHLY*ENLCVLEASPTLHGGEPEPHEGPGSR.S |
| Docking protein 3 | DOK3 | S389Y398 | | 15.1 | K.RAS*GPPGNEHLY*ENLCVLEASPTLHGGEPEPHEGPGSR.S |
| Docking protein 3 | DOK3 | Y432 | 12.0 | 57.7 | R.SPTTSPIY*HNGQDLSWPGPANDSTLEAQYR.R |
| Dual-specificitytyrosine phosphorylation-regulated kinase 2 | DYRK4 | Y382 | 22.5 | 55.7 | R.VYTY*IQSR.F |
| Dynamitin | DCTN2 | Y91 | 39.4 | 96.3 | R.TGYESGEY*EM#LGEGLGVK.E |
| Dynamitin | DCTN2 | Y318 | 14.7 | 16.5 | K.VHQLY*ETIQR.W |
| DYRK1B | DYRK1A | Y271 | 33.4 | 49.5 | R.IY*QYIQSR.F |
| DYRK1B | DYRK1A | Y273 | 39.3 | 44.8 | R.IYQY*IQSR.F |
| DYRK3 | DYRK3 | Y369 | 22.5 | 32.5 | K.LYTY*IQSR.F |
| EBNA2 coactivator p100 | SND1 | Y113 | 5.8 | 17.0 | R.EYGM#IY*LGK.D |
| EBNA2 coactivator p100 | SND1 | Y109 | 18.6 | 26.4 | R.EY*GM#IYLGK.D |
| EBNA2 coactivator p100 | SND1 | Y908 | 12.1 | 45.5 | R.ADDADEFGY*SR.- |
| EH domain containing 1 | EHD1 | Y453 | 12.1 | 27.0 | K.DKPTYDEIFY*TLSPVNGK.I |
| Elongation factor 1 alpha 1 | EEF1A1 | Y86 | 10.6 | 50.5 | K.YY*VTIIDAPGHR.D |
| Elongation factor 1 alpha 1 | EEF1A1 | Y141 | 23.9 | 65.7 | R.EHALLAY*TLGVK.Q |
| Elongation factor 1 alpha 1 | EEF1A1 | Y85 | 8.2 | 44.9 | K.Y*YVTIIDAPGHR.D |
| Elongation protein 3 | ELP3 | Y202 | 50.5 | 57.2 | R.NLHDALSGHTSNNIY*EAVK.Y |
| Emerin | EMD | Y161 | 55.6 | 55.4 | R.DSAY*QSITHYRPVSASR.S |
| Engulfment and cell motility 2 | ELMO2 | Y717 | 19.8 | 58.5 | K.EPSSYDFVY*HYG.- |
| Enhancer of filamentation 1 | NEDD9 | Y92 | 100.0 | 22.9 | K.LY*QVPNPQAAPR.D |
| Enhancer of filamentation 1 | NEDD9 | Y164 | 10.9 | 15.3 | R.TGHGY*VYEYPSR.Y |
| Enhancer of filamentation 1 | NEDD9 | Y345 | 100.0 | 30.7 | R.DGVY*DVPLHNPPDAK.G |
| Enhancer of filamentation 1 | NEDD9 | Y166 | 27.8 | 23.4 | R.TGHGYVY*EYPSR.Y |
| Enhancer of filamentation 1 | NEDD9 | Y317 | 6.3 | 50.1 | R.RHQSLSPNHPPPQLGQSVGSQNDAY*DVPR.G |
| Enhancer of filamentation 1 | NEDD9 | Y261 | 40.4 | 15.1 | R.QAGRPDLRPEGVY*DIPPTCTKPAGK.D |
| Enolase 1 | ENO1 | Y44 | 40.0 | 46.9 | R.AAVPSGASTGIY*EALELRDNDK.T |
| Enolase 2 | ENO3 | Y44 | 93.0 | 104.8 | R.AAVPSGASTGIY*EALELR.D |
| EPS15R | EPS15L1 | Y564 | 63.3 | 60.0 | R.SLEQY*DQVLDGAHGASLTDLANLSEGVSLAER.G |
| Erbin | ERBIN | Y884 | 40.7 | 63.2 | K.IY*DILSDNGPQQPSTTVK.I |
| Erbin | ERBIN | Y1104 | 32.3 | 55.6 | R.AQIPEGDY*LSYR.E |
| ERK1 | MAPK3 | Y204 | 25.5 | 111.1 | R.IADPEHDHTGFLTEY*VATR.W |
| ERK1 | MAPK3 | T198Y204 | 16.5 | 52.8 | R.IADPEHDHT*GFLTEY*VATR.W |
| ERK1 | MAPK3 | T202Y204 | 40.7 | 95.4 | R.IADPEHDHTGFLT*EY*VATR.W |
| ERK2 | MAPK1 | Y187 | 31.5 | 104.6 | R.VADPDHDHTGFLTEY*VATR.W |
| ERK2 | MAPK1 | T185Y187 | 16.5 | 75.7 | R.VADPDHDHTGFLT*EY*VATR.W |
| ERK2 | MAPK1 | T181Y187 | 11.7 | 47.0 | R.VADPDHDHT*GFLTEY*VATR.W |
| Erythrocyte membrane protein band 4.1 like 5 | EPB41L5 | Y499 | 8.0 | 34.1 | R.LPGLGEPEVEY*ETLKDTSEK.L |
| Eukaryotic translation elongation factor 1 beta 2 | EEF1B2 | Y79 | | 48.7 | K.Y*GPADVEDTTGSGATDSKDDDDIDLFGSDDEEESEEAKR.L |
| Eukaryotic translation elongation factor 2 | EEF2 | Y760 | 77.9 | 21.8 | R.LMEPIYLVEIQCPEQVVGGIY*GVLNR.K |
| Eukaryotic translation initiation factor 2A | EIF2A | Y250 | 45.3 | 88.5 | K.TGASY*YGEQTLHYIATNGESAVVQLPK.N |
| Expressed in T cells and eosinophils in atopic dermatitis | FAF2 | Y79 | 29.0 | 15.4 | R.IY*SYVVSRPQPR.G |
| Ezrin | EZR | Y354 | 100.0 | 33.9 | R.LQDY*EEK.T |
| Ezrin | EZR | Y116 | 16.5 | 41.1 | K.EGILSDEIY*CPPETAVLLGSYAVQAK.F |
| Fatty acid synthase | FASN | Y1047 | 28.4 | 15.3 | K.HGLY*LPTR.V |
| Fc receptor like protein 2 | FCRL2 | Y502 | 16.9 | 22.0 | K.DSQVIY*SSVK.S |
| Fc receptor like protein 3 | FCRL3 | Y722 | 100.0 | 89.9 | R.AHEEDDEENY*ENVPR.V |
| Fc receptor like protein 3 | FCRL3 | Y692 | 15.2 | 35.0 | K.ENSANCPMMHQEHEELTVLY*SELK.K |
| FGFR1 oncogene partner | FGFR1OP | Y337 | 48.0 | 79.0 | K.IGSLGLGTGEDDDY*VDDFNSTSHR.S |
| Fgr | FGR | Y208 | 33.8 | 82.4 | K.LDMGGY*YITTR.V |
| Fgr | FGR | Y34 | 88.1 | 63.1 | R.SYGAADHY*GPDPTK.A |

Peak Area: <-10, -5, -3, 0, 3, 5, >10

%CV: 0, 29, 42, 71, 86, >100

**White dots-** Significant change in peptide abundance at **5** %FDR compared to the timepoint with the minimum peak area for a given PSM

CarT: 0m, 2m, 5m — RajiB: 0m, 2m, 5m

| Protein Name | Gene | Phosphosites | Ascor | MOWSE | Sequence |
|---|---|---|---|---|---|
| Fgr | FGR | Y145 | 22.9 | 22.1 | K.TGCIPSNYVAPVDSIQAEEWY*FGK.I |
| Filamin B | FLNB | Y2502 | 32.1 | 35.5 | R.SSTETCY*SAIPK.A |
| FK506 binding protein 4 | FKBP4 | Y220 | 46.5 | 35.0 | K.GEHSIVY*LKPSYAFGSVGK.E |
| Flightless 1 | FLII | Y737 | 120.1 | 63.5 | K.VGLGLGY*LELPQINYK.L |
| Flightless 1 | FLII | Y946 | 60.4 | 17.7 | R.YWVPVEY*EEEEK.K |
| Fodrin beta | SPTBN1 | Y777 | 3.8 | 33.4 | K.IVSSSDVGHDEY*STQSLVK.K |
| Fumarase | FH | Y491 | 14.3 | 64.3 | K.ETAIELGY*LTAEQFDEWVKPK.D |
| FYB | FYB1 | Y801Y803 | 100.0 | 32.6 | K.Y*GY*VLR.S |
| FYB | FYB1 | Y571 | 32.3 | 79.2 | K.TTAVEIDY*DSLK.L |
| FYB | FYB1 | Y801 | 44.7 | 41.1 | K.Y*GYVLR.S |
| FYB | FYB1 | Y559 | 10.9 | 23.3 | R.GSY*GYIK.T |
| FYB | FYB1 | Y747 | 100.0 | 28.4 | K.FKY*DGEIR.V |
| FYB | FYB1 | Y803 | 16.9 | 38.3 | K.YGY*VLR.S |
| Fyn | YES1 | Y185 | 39.9 | 51.3 | K.GAY*SLSIR.D |
| Fyn | YES1 | Y420 | 16.9 | 76.4 | R.LIEDNEY*TAR.Q |
| FYVE, RhoGEF and PH domain containing 6 | FGD6 | Y760 | 31.1 | 45.1 | R.HYEEIPEY*ENLPFIMAIR.K |
| G protein signalling modulator 3 | GPSM3 | Y108 | 11.0 | 24.0 | R.EQLY*STILSHQCQR.M |
| GAB2 | GAB2 | Y476 | 9.4 | 20.2 | R.AGDNSQSVY*IPM#SPGAHHFDSLGYPSTTLPVHR.G |
| GART | GART | Y348 | 42.3 | 17.1 | K.GYPGDY*TK.G |
| GDP dissociation inhibitor 2 | GDI2 | Y203 | 20.5 | 41.9 | R.TDDYLDQPCY*ETINR.I |
| GGA2 | GGA2 | Y269 | 100.0 | 60.1 | R.RPGQAPPDQEALQVVY*ER.C |
| GIT1 | GIT1 | Y545 | 23.4 | 47.0 | R.LQPFHSTELEDDAIY*SVHVPAGLYR.I |
| Glucocorticoid receptor DNA binding factor 1 | ARHGAP3 | Y1105 | 16.9 | 59.9 | R.NEEENIY*SVPHDSTQGK.I |
| Glucocorticoid receptor DNA binding factor 1 | ARHGAP3 | Y1087 | 7.9 | 40.9 | K.SVSSSPWLPQDGFDPSDY*AEPM#DAVVKPR.N |
| Glucose-6-phosphate dehydrogenase | G6PD | Y533 | 46.9 | 64.4 | R.VGFQY*EGTYK.W |
| Glucose-6-phosphate dehydrogenase | G6PD | Y431 | | 43.7 | R.VQPNEAVY*TK.M |
| Glutamate dehydrogenase 1 | GLUD1 | Y464 | 30.5 | 63.8 | R.DSNY*HLLMSVQESLER.K |
| Glutamate oxaloacetate transaminase, mitochondrial | GOT2 | Y96 | 100.0 | 44.2 | K.NLDKEY*LPIGGLAEFCK.A |
| Glutathione S transferase 3 | GSTP1 | Y8 | 76.7 | 59.4 | M.PPYTVVY*FPVR.G |
| Glycogen debranching enzyme | AGL | Y584 | 28.1 | 56.5 | R.EAM#SAY*NSHEEGR.L |
| Glycogen phosphorylase, brain type | PYGB | Y197 | 106.0 | 22.4 | K.ARPEY*M#LPVHFYGR.V |
| Glycogen synthase kinase 3 beta | GSK3B | Y216 | 16.9 | 51.4 | R.GEPNVSY*ICSR.Y |
| Glycogen synthase kinase 3 beta | GSK3B | Y216S219 | 29.0 | 31.4 | R.GEPNVSY*ICS*R.Y |
| Glycogen synthase kinase 3 beta | GSK3B | S215Y216 | 22.1 | 39.2 | R.GEPNVS*Y*ICSR.Y |
| Glyoxylate reductase | GRHPR | Y255 | 78.2 | 79.5 | R.GDVVNQDDLY*QALASGK.I |
| Golgi autoantigen, golgin subfamily A, 4 | GOLGA4 | Y2148 | 13.9 | 21.3 | K.NVY*ATTVGTPYK.G |
| Golgi phosphoprotein 4 | GOLIM4 | Y483 | 65.0 | 77.0 | R.QQAHY*DAMDNDIVQGAEDQGIQGEEGAYER.D |
| Golgi phosphoprotein 4 | GOLIM4 | Y673 | 100.0 | 30.7 | R.EEHY*EEEEEEEDGAAVAEK.S |
| Golgin 160 | GOLGA3 | Y329 | 10.9 | 27.1 | R.GTY*GILSK.T |
| Golgin 160 | GOLGA3 | Y210 | 16.9 | 19.3 | K.EY*SFLR.T |
| GRB2 associated binding protein 3 | GAB3 | Y560 | 100.0 | 59.8 | R.VDY*VQVDEQK.T |
| Grb4 | NCK2 | Y110 | 100.0 | 65.1 | R.IY*DLNIPAFVK.F |
| Grb4 | NCK2 | S90Y110 | 25.5 | 35.7 | R.DAS*PTPSTDAEYPANGSGADRIY*DLNIPAFVK.F |
| Grb4 | NCK2 | Y50 | 14.7 | 35.7 | R.TGY*VPSNYVER.K |
| GRID | GRAP2 | Y207 | 45.3 | 47.5 | K.LSDHPPTLPLQQHQHQPQPPQY*APAPQQLQQPPQQR.Y |
| GRID | GRAP2 | Y45 | 42.3 | 72.6 | K.AELGSQEGY*VPK.N |
| GRID | GRAP2 | T192Y207 | 0.7 | 27.9 | R.KLSDHPPT*LPLQQHQHQPQPPQY*APAPQQLQQPPQQR.Y |
| GRID | GRAP2 | S187Y207 | 11.0 | 36.6 | R.KLS*DHPPTLPLQQHQHQPQPPQY*APAPQQLQQPPQQR.Y |
| GRID | GRAP2 | Y222 | 100.0 | 28.1 | R.Y*LQHHHFHQER.R |
| Growth arrest specific 7 | GAS7 | Y333 | 0.5 | 17.4 | R.Y*ASVEKARK.A |
| GRP1 associated scaffold protein | GRASP | Y237 | 8.4 | 30.1 | R.LVHGLVVKDPSIY*DTLESVR.S |
| H2B Histone family, member E | HIST1H2B | Y41 | 9.1 | 24.3 | R.KESYSVY*VYK.V |
| H4 Histone family, member A | HIST1H4A | Y52 | 32.3 | 50.5 | R.ISGLIY*EETR.G |
| HBS1L | HBS1L | Y58 | 25.4 | 47.4 | R.DKPSVEPVEEYDY*EDLK.E |
| HBS1L | HBS1L | Y56 | 21.2 | 23.6 | R.DKPSVEPVEEY*DYEDLKESSNSVSNHQLSGFDQAR.L |
| Hck | HCK | Y411 | 16.9 | 82.6 | R.VIEDNEY*TAR.E |
| Heat shock 10 KD protein | HSPE1 | Y76 | 18.4 | 53.4 | K.VLLPEY*GGTK.V |
| Heat shock 60 KD protein 1 (chaperonin) | HSPD1 | Y227 | 39.3 | 45.8 | R.GYISPY*FINTSK.G |
| Heat shock 70 KD protein 1A | HSPA1L | Y41 | 16.9 | 90.3 | R.TTPSY*VAFTDTER.L |
| Heat shock 70 kDa protein 8 | HSPA8 | Y15 | 27.7 | 116.7 | K.GPAVGIDLGTTY*SCVGVFQHGK.V |
| Heat shock 70kDa protein 4 | HSPA4 | Y336 | 143.9 | 66.0 | K.EDIY*AVEIVGGATR.I |
| Heat shock 70kDa protein 4 | HSPA4 | Y626 | 32.1 | 61.3 | K.NAVEEYVY*EM#R.D |

Peak Area: <-10, -5, -3, 0, 3, 5, >10

%CV: 0, 29, 43, 71, 86, >100

**White dots**- Significant change in peptide abundance at 5 %FDR compared to the timepoint with the minimum peak area for a given PSM

CarT: 0m 2m 5m — RajiB: 0m 2m 5m

| Protein Name | Gene | Phosphosites | Ascor | MOWSE | Sequence |
|---|---|---|---|---|---|
| **Heat shock 70kDa protein 4** | HSPA4 | Y624 | 32.1 | 56.2 | K.NAVEEY*VYEMR.D |
| **Heat shock 70kDa protein 4** | HSPA4 | Y660 | 46.6 | 18.9 | K.LEDTENWLY*EDGEDQPK.Q |
| **Heat shock protein 105 kDa** | HSPH1 | Y641 | 32.1 | 56.5 | K.NAVEEY*VYEFR.D |
| **Heat shock protein 105 kDa** | HSPH1 | Y677 | 62.1 | 54.6 | R.LLTETEDWLY*EEGEDQAK.Q |
| **Helicase ATP binding 1** | ASCC3 | Y459 | 38.3 | 16.6 | R.IPYSEPM#PLSFEEKPVY*IQDLDEIGQLAFK.G |
| **Hematopoietic cell specific LYN substrate 1** | HCLS1 | Y378 | 35.3 | 23.5 | R.TLEGLQVEEEPVYEAEPEPEPEPEPEPENDY*EDVEEM#DR.H |
| **Hematopoietic cell specific LYN substrate 1** | HCLS1 | Y360 | 25.8 | 34.7 | R.TLEGLQVEEEPVY*EAEPEPEPEPEPEPENDYEDVEEM#DR.H |
| **Hematopoietic cell specific LYN substrate 1** | HCLS1 | Y103 | 67.1 | 58.2 | K.SAVGHEY*VAEVEK.H |
| **Heterogeneous nuclear ribonucleoprotein C** | HNRNPC | Y137 | 30.2 | 19.0 | R.MY*SYPAR.V |
| **Heterogeneous nuclear ribonucleoprotein H1** | HNRNPH1 | Y306 | 7.8 | 35.8 | K.ATENDIY*NFFSPLNPVR.V |
| **Heterogeneous nuclear ribonucleoprotein H3** | HNRNPH3 | Y296 | 22.4 | 85.5 | R.DGMDNQGGY*GSVGR.M |
| **Heterogeneous nuclear ribonucleoprotein M** | HNRNPM | Y681 | 100.0 | 23.1 | K.DKFNECGHVLY*ADIK.M |
| **Heterogeneous nuclear ribonucleoprotein U** | HNRNPU | Y466 | 6.6 | 49.1 | K.EKPY*FPIPEEYTFIQNVPLEDR.V |
| **Heterogeneous nuclear ribonucleoprotein U** | HNRNPU | Y473 | 13.1 | 34.0 | K.EKPYFPIPEEY*TFIQNVPLEDR.V |
| **Hexokinase domain containing 1** | HKDC1 | Y497 | 100.0 | 24.7 | R.AELEY*GLKKK.S |
| **HGAL** | GCSAM | Y107 | 40.2 | 49.7 | R.VLCTRPSGNSAEEYY*ENVPCK.A |
| **HGAL** | GCSAM | Y148 | 66.2 | 53.4 | R.SPEDEY*ELLM#PHR.I |
| **HGAL** | GCSAM | Y128 | 14.0 | 34.7 | R.ESLGGTETEY*SLLHMPSTDPR.H |
| **HGAL** | GCSAM | Y86 | 15.2 | 67.0 | R.MSSTPIQDNVDQTYSEELCY*TLINHR.V |
| **HGAL** | GCSAM | S99Y107 | 12.8 | 19.5 | R.VLCTRPS*GNSAEEYY*ENVPCK.A |
| **HGAL** | GCSAM | Y106 | 66.5 | 44.4 | R.VLCTRPSGNSAEEY*YENVPCK.A |
| **HGAL** | GCSAM | S102Y107 | 12.7 | 35.5 | R.VLCTRPSGNS*AEEYY*ENVPCK.A |
| **HGAL** | GCSAM | S69Y86 | 13.7 | 31.2 | R.M#SS*TPIQDNVDQTYSEELCY*TLINHR.V |
| **HIP55** | DBNL | Y162 | 32.2 | 52.6 | R.FQDVGPQAPVGSVY*QK.T |
| **HIPK1** | HIPK2 | Y352 | 39.9 | 67.8 | K.AVCSTY*LQSR.Y |
| **HIV 1 REV binding protein** | AGFG1 | Y327 | 20.9 | 61.2 | K.AGLQTADKY*AALANLDNIFSAGQGGDQGSGFGTTGK.A |
| **HLA DR associated protein I** | ANP32A | Y148 | 33.2 | 36.8 | K.LLPQLTYLDGY*DR.E |
| **HLA-DMB** | HLA-DMB | Y248 | 8.0 | 37.6 | R.AGHSSY*TPLPGSNYSEGWHIS.- |
| **Homeodomain interacting protein kinase 3** | HIPK3 | Y359 | 16.9 | 68.0 | K.TVCSTY*LQSR.Y |
| **HP1-BP74** | HP1BP3 | Y379 | 19.3 | 41.4 | K.YVLENHPGTNSNY*QMHLLK.K |
| **HSP90A** | HSP90AA1 | Y492 | 48.6 | 70.9 | K.HIY*YITGETK.D |
| **HSP90A** | HSP90AB1 | Y197 | 32.1 | 67.9 | K.EDQTEY*LEER.W |
| **HSP90A** | HSP90AB1 | Y284 | 100.0 | 17.5 | K.Y*IDQEELNK.T |
| **HSP90B** | HSP90AB1 | Y484 | 33.8 | 53.1 | K.SIY*YITGESK.E |
| **Hypothetical protein FLJ10769** | NAXD | Y85 | 35.4 | 66.2 | R.IGVVGGCQEY*TGAPYFAAISALK.V |
| **Hypothetical protein FLJ38482** | TMEM192 | Y209 | 28.2 | 64.6 | K.IY*AYPSNITSETGFR.T |
| **ICT1** | MRPL58 | Y49 | 19.3 | 26.2 | K.SIYSLDKLY*PESQGSDTAWR.V |
| **Ig alpha** | CD79A | Y210 | 15.2 | 71.3 | R.GLQGTY*QDVGSLNIGDVQLEKP.- |
| **Ig beta** | CD79B | Y207 | 12.1 | 54.3 | K.AGM#EEDHTYEGLDIDQTATY*EDIVTLR.T |
| **IGF-I receptor** | INSR | Y1165 | 26.0 | 34.4 | R.DIYETDY*YR.K |
| **IK factor** | IK | Y114 | 40.7 | 60.9 | R.DGVNKDY*EETELISTTANYR.A |
| **Interferon gamma receptor 1** | IFNGR1 | Y304 | 29.1 | 70.0 | K.Y*VSLITSYQPFSLEK.E |
| **Intersectin 2** | ITSN2 | Y967 | 100.0 | 66.5 | R.EEPEALY*AAVNK.K |
| **Intersectin 2** | ITSN2 | Y552 | 100.0 | 49.4 | K.LIY*LVPEK.Q |
| **Intestinal cell (MAK-like) kinase** | ICK | T157Y159 | 12.9 | 26.5 | R.SKPPYT*DY*VSTR.W |
| **Intestinal cell (MAK-like) kinase** | ICK | Y156T157 | 7.6 | 24.4 | R.SKPPY*T*DYVSTR.W |
| **Intestinal cell (MAK-like) kinase** | ICK | Y156Y159 | -0.2 | 14.9 | R.SKPPY*TDY*VSTR.W |
| **IQ motif and Sec7 domain 1** | IQSEC1 | Y343 | 100.0 | 27.9 | R.AGGAAPDY*WALAHKEDK.A |
| **ITK** | ITK | Y512 | 40.2 | 56.3 | R.FVLDDQY*TSSTGTKFPVK.W |
| **Janus kinase 2** | JAK2 | Y570 | 22.3 | 16.1 | R.EVGDY*GQLHETEVLLK.V |
| **Janus kinase 3** | JAK3 | Y785 | 5.7 | 54.9 | R.DLNSLISSDY*ELLSDPTPGALAPR.D |
| **JAW1** | LRMP | Y26 | 30.2 | 46.3 | R.EY*SSLPLPR.H |
| **Junction adhesion molecule 1** | F11R | Y280 | 16.9 | 15.3 | K.VIY*SQPSAR.S |
| **Karyopherin beta 1** | KPNB1 | Y752 | -0.9 | 18.8 | K.SDY*DMVDYLNELR.E |
| **Karyopherin beta 3** | IPO5 | Y838 | 135.3 | 94.1 | K.RQDEDY*DEQVEESLQDEDDNDVYILTK.V |
| **KCC1** | SLC12A4 | Y17 | 85.5 | 40.1 | R.RGDY*DNLEGLSWVDYGER.A |
| **KIAA0317** | AREL1 | Y47 | 35.1 | 15.0 | R.TIY*DYVR.G |
| **KIAA1949** | PPP1R18 | Y230 | 6.8 | 26.6 | R.LSPGESAY*QK.L |
| **Lactate dehydrogenase A** | LDHA | Y239 | 39.3 | 56.0 | K.QVVESAY*EVIK.L |
| **Lactate dehydrogenase A** | LDHA | Y10 | 100.0 | 37.0 | K.DQLIY*NLLK.E |
| **Lactate dehydrogenase B** | | Y240 | 26.0 | 48.8 | K.MVVESAY*EVIK.L |
| **Laminin receptor 1** | RPSA | Y139 | 19.1 | 87.3 | R.ADHQPLTEASY*VNLPTIALCNTDSPLR.Y |

Peak Area: <-10, -5, -3, 0, 3, 5, >10

%CV: 0, 29, 42, 71, 86, >100

**White dots-** Significant change in peptide abundance at **5** %FDR compared to the timepoint with the minimum peak area for a given PSM

| Protein Name | Gene | Phosphosites | CarT (0m, 2m, 5m) | RajiB (0m, 2m, 5m) | Ascor | MOWSE | Sequence |
|---|---|---|---|---|---|---|---|
| LAT like membrane associated protein | LAX1 | Y294 | | | 142.7 | 57.9 | R.DY*ENVPAADPSGSQQQAEK.D |
| LAT like membrane associated protein | LAX1 | Y93 | | | 100.0 | 42.8 | K.NIY*DILPWR.Q |
| LAT like membrane associated protein | LAX1 | Y373 | | | 31.5 | 88.3 | K.HREEM#SNEDSSDY*ENVLTAK.L |
| LBA | LRBA | Y1110 | | | 164.7 | 74.8 | K.SIVEEEEDDDY*VELK.V |
| Lck | LCK | Y192 | | | 44.7 | 70.4 | R.NLDNGGFY*ISPR.I |
| Lck | LCK | Y505 | | | 19.7 | 29.8 | R.SVLEDFFTATEGQY*QPQP.- |
| Lck | LCK | Y414 | | | 26.0 | 58.7 | K.WTAPEAINY*GTFTIK.S |
| Lck | LCK | Y470 | | | 100.0 | 19.4 | R.M#VRPDNCPEELY*QLMR.L |
| Lck interacting transmembrane adaptor 1 | LIME1 | Y235 | | | 25.4 | 39.1 | K.GQGAILALAGDLAY*QTLPLR.A |
| Leucyl cystinyl aminopeptidase | LNPEP | Y70 | | | 28.2 | 49.6 | R.GLGEHEM#EEDEEDY*ESSAK.L |
| Leupaxin | LPXN | Y22 | | | 20.8 | 53.4 | R.STLQDSDEY*SNPAPLPLDQHSR.K |
| Lim and SH3 protein 1 | LASP1 | Y171 | | | 7.4 | 31.2 | R.RPLEQQQPHHIPTSAPVY*QQPQQQPVAQSYGGYK.E |
| LIM domain only 7 | LMO7 | Y185 | | | 50.1 | 41.7 | K.AQSNPY*YNGPHLNLK.A |
| LIM domain only 7 | LMO7 | Y186 | | | 9.5 | 33.4 | K.AQSNPYY*NGPHLNLK.A |
| Linker for activation of T cells | LAT | Y220 | | | 120.8 | 94.5 | R.EY*VNVSQELHPGAAK.T |
| Linker for activation of T cells | LAT | Y220S224 | | | 100.0 | 51.7 | R.EY*VNVS*QELHPGAAK.T |
| Liver-specific bHLH-Zip transcription factor | LSR | Y551 | | | 37.5 | 14.0 | R.SRDPHY*DDFR.S |
| Lupus brain antigen 1 | TRANK1 | Y779 | | | 42.4 | 15.7 | R.LTEEVY*K.K |
| Lupus brain antigen 1 | TRANK1 | Y2350 | | | 100.0 | 55.5 | K.EGIVQEDDY*ENEVEDFGELRPR.R |
| Lymphocyte antigen 9 | LY9 | Y626 | | | 32.1 | 73.2 | K.EESSATIY*CSIR.K |
| Lymphocyte cytosolic protein 1 | LCP1 | Y299 | | | | 77.7 | K.AY*YHLLEQVAPK.G |
| Lymphocyte cytosolic protein 1 | PLS3 | Y124 | | | 100.0 | 51.1 | K.Y*AFVNWINK.A |
| Lymphocyte cytosolic protein 1 | LCP1 | Y28 | | | 67.6 | 160.6 | K.VDTDGNGY*ISFNELNDLFK.A |
| Lymphocyte cytosolic protein 1 | LCP1 | Y598 | | | 100.0 | 47.3 | R.VY*ALPEDLVEVNPK.M |
| Lymphocyte cytosolic protein 1 | LCP1 | Y300 | | | -0.1 | 72.8 | K.AYY*HLLEQVAPK.G |
| Lymphocyte cytosolic protein 1 | LCP1 | Y417 | | | 8.0 | 29.5 | R.VNHLY*SDLSDALVIFQLYEK.I |
| Lymphocyte specific protein | LSP1 | Y125 | | | 62.5 | 50.9 | R.SPEGEQEDRPGLHAY*EK.E |
| Lymphocyte specific protein | LSP1 | S111Y125 | | | 100.0 | 37.5 | R.S*PEGEQEDRPGLHAY*EK.E |
| Lymphocyte specific protein | LSP1 | Y234 | | | 33.8 | 122.1 | K.IDQWLEQY*TQAIETAGR.T |
| Lyn | LYN | Y194 | | | 39.9 | 79.3 | R.SLDNGGYY*ISPR.I |
| Lyn | LYN | Y473 | | | 100.0 | 55.0 | R.VENCPDELY*DIM#K.M |
| Lyn | LYN | Y316 | | | 37.7 | 59.0 | R.EEPIY*IITEYMAK.G |
| Lyn | LYN | Y508 | | | 67.3 | 39.5 | K.EKAEERPTFDYLQSVLDDFYTATEGQY*QQQP.- |
| Lyn | LYN | Y501 | | | 30.1 | 35.9 | K.EKAEERPTFDYLQSVLDDFY*TATEGQYQQQP.- |
| Lyn | LYN | Y306 | | | 87.6 | 41.8 | R.LY*AVVTR.E |
| LysM, peptidoglycan-binding, domain containing 2 | LYSMD2 | Y208 | | | 22.4 | 27.9 | R.DEESPY*ATSLYHS.- |
| MAGOH | MAGOH | Y40 | | | 7.3 | 41.5 | R.YANNSNY*KNDVM#IR.K |
| Malate dehydrogenase mitochondrial | MDH2 | Y56 | | | 40.5 | 82.7 | R.LTLY*DIAHTPGVAADLSHIETK.A |
| MAP4K4 | MINK1 | Y36 | | | 10.3 | 88.7 | R.DPAGIFELVEVVGNGTY*GQVYK.G |
| MAPK11 | MAPK11 | Y190 | | | 10.9 | 70.6 | R.QADEEMTGY*VATR.W |
| MAPK12 | MAPK12 | Y185 | | | 32.1 | 104.6 | R.QADSEMTGY*VVTR.W |
| MAPK14 | MAPK14 | Y182 | | | 41.9 | 66.2 | R.HTDDEMTGY*VATR.W |
| MAPK14 | MAPK14 | T180Y182 | | | 51.8 | 101.0 | R.HTDDEMT*GY*VATR.W |
| MAPK14 | MAPK14 | T175Y182 | | | 28.4 | 19.1 | R.HT*DDEM#TGY*VATR.W |
| Matrin 3 | MATR3 | Y250 | | | 32.1 | 45.4 | K.FDSEY*ER.M |
| Matrin 3 | MATR3 | Y219 | | | 100.0 | 26.2 | R.MDY*EDDRLR.D |
| MBC2 | ESYT1 | Y822 | | | 26.4 | 71.1 | K.HLSPY*ATLTVGDSSHK.T |
| MCM2 minichromosome maintenance deficient 2, mitotin | MCM2 | Y137 | | | 1.6 | 30.1 | R.GLLY*DSDEEDEERPAR.K |
| MCM3 | MCM3 | Y708 | | | 14.6 | 29.4 | K.DGDSYDPY*DFSDTEEEM#PQVHTPK.T |
| MCM3 | MCM3 | Y708T722 | | | 20.8 | 26.2 | K.DGDSYDPY*DFSDTEEEM#PQVHT*PK.T |
| Melanoregulin | MREG | Y37 | | | 25.4 | 40.0 | R.ALPEKEPLVSDNNPY*SSFGATLVR.D |
| Microtubule associated protein 4 | MAP4 | Y47 | | | 10.4 | 55.1 | K.TDY*IPLLDVDEK.T |
| Moesin | MSN | Y116 | | | 11.9 | 44.8 | K.EGILNDDIY*CPPETAVLLASYAVQSK.Y |
| Myelin protein zero like 1 | MPZL1 | Y263 | | | 46.9 | 41.4 | K.SESVVY*ADIR.K |
| Myosin heavy chain 9, nonmuscle | MYH9 | Y1408 | | | 100.0 | 37.6 | K.VAAY*DKLEK.T |
| Myosin heavy chain 9, nonmuscle | MYH9 | Y151 | | | 12.9 | 28.3 | R.HEM#PPHIY*AITDTAYR.S |
| Myosin heavy chain 9, nonmuscle | MYH9 | Y11 | | | 20.1 | 54.8 | K.YLY*VDKNFINNPLAQADWAAK.K |
| Myosin IG | MYO1G | Y72 | | | 46.5 | 36.7 | R.ELY*ERPPHLYAVANAAYK.A |
| Myotubularin related protein 10 | MTMR10 | Y708 | | | 76.6 | 80.1 | R.SGPLEACY*GELGQSR.M |
| NCK1 | NCK1 | Y105 | | | 211.4 | 68.7 | R.LY*DLNMPAYVK.F |
| NCK1 | NCK1 | S85Y105 | | | 12.9 | 36.3 | K.RKPS*VPDSASPADDSFVDPGERLY*DLNMPAYVK.F |
| Nectin 1 | NECTIN1 | Y468 | | | 5.6 | 62.3 | K.YDEDAKRPY*FTVDEAEAR.Q |

**Peak Area**: <-10, -5, -3, 0, 3, 5, >10

**%CV**: 0, 29, 42, 71, 86, >100

**White dots-** Significant change in peptide abundance at **5** %FDR compared to the timepoint with the minimum peak area for a given PSM

| Protein Name | Gene | Phosphosites | CarT 0m | CarT 2m | CarT 5m | RajiB 0m | RajiB 2m | RajiB 5m | Ascor | MOWSE | Sequence |
|---|---|---|---|---|---|---|---|---|---|---|---|
| **NF kappa B inhibitor epsilon** | NFKBIE | Y155 | | | | | | | **33.2** | **77.3** | R.KGPDEAEESQY*DSGIESLR.S |
| **NFKB3** | RELA | Y306 | | | | | | | **-0.2** | **14.0** | R.TY*ETFK.S |
| **NipSnap1** | NIPSNAP1 | Y261 | | | | | | | **32.1** | **44.9** | R.GWDENVY*YTVPLVR.H |
| **NSFL1C** | NSFL1C | Y167 | | | | | | | **22.4** | **67.4** | R.LGAAPEEESAY*VAGEK.R |
| **NTBA** | SLAMF6 | Y308 | | | | | | | **26.0** | **51.3** | R.ENDTITIY*STINHSK.E |
| **NTBA** | SLAMF6 | Y284 | | | | | | | **28.1** | **86.9** | R.NLEYVSVSPTNNTVY*ASVTHSNR.E |
| **NTBA** | SLAMF6 | Y273 | | | | | | | **55.8** | **74.6** | R.NLEY*VSVSPTNNTVYASVTHSNR.E |
| **NTBA** | SLAMF6 | Y273Y284 | | | | | | | **37.0** | **51.4** | R.NLEY*VSVSPTNNTVY*ASVTHSNR.E |
| **Nuclear factor kappa B, subunit 2** | NFKB2 | Y39 | | | | | | | **65.1** | **43.2** | K.EPAPETADGPY*LVIVEQPK.Q |
| **Nuclear ubiquitous casein kinase and cyclin dependent kinase substrate** | NUCKS1 | Y13 | | | | | | | **12.1** | **76.7** | K.VVDY*SQFQESDDADEDYGR.D |
| **Nucleoside diphosphate kinase A** | NME1- | Y52 | | | | | | | **100.0** | **38.3** | K.EHY*VDLK.D |
| **Nucleoside diphosphate kinase B** | NME1- | Y151 | | | | | | | **71.6** | **37.2** | K.SCAHDWVY*E.- |
| **OCIA domain containing 1** | OCIAD1 | Y199 | | | | | | | **13.9** | **30.7** | K.NITY*EELR.N |
| **Odin** | ANKS1A | Y455 | | | | | | | **44.7** | **30.9** | R.EEDEHPY*ELLLTAETK.K |
| **PAG** | PAG1 | Y359 | | | | | | | **60.4** | **113.3** | R.SPSSCNDLY*ATVK.D |
| **PAG** | PAG1 | Y227 | | | | | | | **32.1** | **61.5** | K.AEFAEY*ASVDR.N |
| **PAG** | PAG1 | Y417 | | | | | | | **39.3** | **110.0** | K.ENDY*ESISDLQQGR.D |
| **PAG** | PAG1 | Y163 | | | | | | | **197.9** | **113.3** | R.SVDGDQGLGM#EGPY*EVLK.D |
| **PAG** | PAG1 | S353Y359 | | | | | | | **24.6** | **82.8** | R.SPS*SCNDLY*ATVK.D |
| **PAG** | PAG1 | Y341 | | | | | | | **13.9** | **62.7** | K.SGQSLTVPESTY*TSIQGDPQR.S |
| **PAG** | PAG1 | Y387 | | | | | | | **56.5** | **48.8** | K.TPNSTLPPAGRPSEEPEPDY*EAIQTLNR.E |
| **PAG** | PAG1 | Y181 | | | | | | | **39.4** | **76.5** | K.DSSSQENMVEDCLY*ETVK.E |
| **PAG** | PAG1 | Y317 | | | | | | | **49.2** | **102.8** | R.EEDPTLTEEEISAMY*SSVNKPGQLVNK.S |
| **PAG** | PAG1 | S354Y359 | | | | | | | **20.5** | **83.2** | R.SPSS*CNDLY*ATVK.D |
| **PAG** | PAG1 | S170Y181 | | | | | | | **19.1** | **66.6** | K.DSS*SQENM#VEDCLY*ETVK.E |
| **PAG** | PAG1 | S301Y317 | | | | | | | **9.3** | **36.0** | K.S*REEDPTLTEEEISAM#Y*SSVNKPGQLVNK.S |
| **PAG** | PAG1 | S169Y181 | | | | | | | **76.7** | **63.1** | K.DS*SSQENM#VEDCLY*ETVK.E |
| **Paxillin** | PXN | Y118 | | | | | | | **6.8** | **17.5** | R.VGEEEHVY*SFPNK.Q |
| **PCAF** | KAT2B | Y729 | | | | | | | | **36.0** | R.DPDQLY*STLK.S |
| **PCTAIRE protein kinase 1** | CDK16 | Y176 | | | | | | | **16.9** | **20.2** | K.LGEGTY*ATVYK.G |
| **PDZ and LIM domain 1** | PDLIM1 | Y151 | | | | | | | **25.4** | **58.1** | R.VITNQYNNPAGLY*SSENISNFNNALESK.T |
| **PDZ and LIM domain 1** | PDLIM1 | Y321 | | | | | | | **17.6** | **41.2** | R.VTPPEGY*EVVTVFPK.- |
| **Pentatricopeptide repeat domain 3** | PTCD3 | Y144 | | | | | | | **100.0** | **23.0** | K.DIAEPHIPCLMPEY*FEPQIK.D |
| **Pericentriolar material 1** | PCM1 | Y1176 | | | | | | | **28.9** | **87.1** | K.TEY*M#AFPKPFESSSSIGAEKPR.N |
| **PEX5** | PEX5 | Y304 | | | | | | | **36.8** | **59.0** | R.DAEAHPWLSDY*DDLTSATYDK.G |
| **Phosphatidylinositol 3 kinase regulatory subunit, alpha** | PIK3R1 | Y607 | | | | | | | **15.2** | **69.9** | K.LNEWLGNENTEDQY*SLVEDDEDLPHHDEK.T |
| **Phosphatidylinositol 3 kinase regulatory subunit, alpha** | PIK3R1 | Y580 | | | | | | | **97.5** | **26.9** | R.DQY*LM#WLTQK.G |
| **Phosphatidylinositol 3 kinase regulatory subunit, alpha** | PIK3R1 | Y452 | | | | | | | **51.2** | **71.5** | K.LHEY*NTQFQEK.S |
| **Phosphatidylinositol 3 kinase regulatory subunit, alpha** | PIK3R1 | Y467 | | | | | | | **90.6** | **50.5** | R.LY*EEYTR.T |
| **Phosphatidylinositol 3 kinase, catalytic subunit, delta** | PIK3CD | Y548 | | | | | | | **50.5** | **51.7** | R.GSGELY*EHEK.D |
| **Phosphatidylinositol 4-kinase alpha** | PI4KA | Y1096 | | | | | | | **56.6** | **36.0** | R.YAGEVY*GM#IR.F |
| **Phosphoglycerate mutase 1** | PGAM2 | Y92 | | | | | | | **41.9** | **33.2** | R.HY*GGLTGLNK.A |
| **Phosphoglycerate mutase 1** | PGAM1 | Y133 | | | | | | | **12.1** | **19.8** | R.SYDVPPPPM#EPDHPFY*SNISK.D |
| **Phosphoglycerate mutase 1** | PGAM1 | Y26 | | | | | | | **7.4** | **17.8** | R.FSGWY*DADLSPAGHEEAK.R |
| **Phosphoinositide 3 kinase adaptor protein 1** | PIK3AP1 | Y570 | | | | | | | **49.2** | **42.7** | K.SQERPGNFY*VSSESIR.K |
| **Phosphoinositide 3 kinase adaptor protein 1** | PIK3AP1 | Y694 | | | | | | | **13.9** | **65.3** | K.VEFGVY*ESGPR.K |
| **Phosphoinositide 3 kinase adaptor protein 1** | PIK3AP1 | Y594 | | | | | | | **42.4** | **32.5** | R.DRPQSSIY*DPFAGM#K.T |
| **Phosphoinositide 3 kinase adaptor protein 1** | PIK3AP1 | Y513 | | | | | | | | **26.9** | R. [illegible] |
| **Phosphoinositide 3 kinase adaptor protein 1** | PIK3AP1 | Y419 | | | | | | | **20.9** | **34.4** | K.EELMHGEEADAVY*ESMAHLSTDLLM#K.C |
| **Phospholipase C like 2** | PLCL2 | Y779 | | | | | | | **13.9** | **15.9** | R.EY*ASLR.T |
| **Phospholipase C, gamma 1** | PLCG1 | Y771 | | | | | | | **71.3** | **92.5** | K.IGTAEPDY*GALYEGR.N |
| **Phospholipase C, gamma 1** | PLCG1 | Y775 | | | | | | | **77.7** | **82.7** | K.IGTAEPDYGALY*EGR.N |
| **Phospholipase C, gamma 1** | PLCG1 | Y1253 | | | | | | | **100.0** | **51.2** | R.Y*QQPFEDFR.I |
| **Phospholipase C, gamma 1** | PLCG1 | Y481 | | | | | | | **15.3** | **60.2** | K.LAEGSAYEEVPTSM#M#Y*SENDISNSIK.N |
| **Phospholipase C, gamma 1** | PLCG1 | Y783 | | | | | | | **195.0** | **82.6** | R.NPGFY*VEANPMPTFK.C |
| **Phospholipase C, gamma 1** | PLCG1 | Y775Y783 | | | | | | | **59.8** | **39.8** | K.IGTAEPDYGALY*EGRNPGFY*VEANPM#PTFK.C |
| **Phospholipase C, gamma 1** | PLCG1 | Y472 | | | | | | | **18.1** | **72.4** | K.LAEGSAY*EEVPTSM#M#YSENDISNSIK.N |
| **Phospholipase C, gamma 1** | PLCG1 | Y771Y775 | | | | | | | **54.0** | **43.3** | K.IGTAEPDY*GALY*EGR.N |
| **Phospholipase C, gamma 1** | PLCG1 | Y1253S1263 | | | | | | | **39.0** | **13.8** | R.Y*QQPFEDFRIS*QEHLADHFDSR.E |
| **Phospholipase C, gamma 1** | PLCG1 | T766Y783 | | | | | | | **17.2** | **16.7** | K.IGT*AEPDYGALYEGRNPGFY*VEANPM#PTFK.C |
| **Phospholysine phosphohistidine inorganic pyrophosphate phosphatase** | LHPP | Y158 | | | | | | | **12.2** | **15.1** | R.Y*YKETSGLM#LDVGPYM#K.A |
| **PI 3 kinase, regulatory subunit beta** | PIK3R2 | Y464 | | | | | | | **34.8** | **37.6** | R.EYDQLY*EEYTR.T |

Peak Area: <-10, -5, -3, 0, 3, 5, >10

%CV: 0, 29, 42, 71, 86, >100

**White dots-** Significant change in peptide abundance at 5 %FDR compared to the timepoint with the minimum peak area for a given PSM

CarT (0m, 2m, 5m) RajiB (0m, 2m, 5m)

| Protein Name | Gene | Phosphosites | Ascor | MOWSE | Sequence |
|---|---|---|---|---|---|
| PI 3 kinase, regulatory subunit beta | PIK3R2 | Y605 | 6.1 | 37.3 | K.NETEDQY*ALM#EDEDDLPHHEER.T |
| PI 3 kinase, regulatory subunit beta | PIK3R2 | Y460 | 10.8 | 37.4 | R.EY*DQLYEEYTR.T |
| PLC, gamma 2 | PLCG2 | Y680 | 16.9 | 88.0 | R.EGSDSY*AITFR.A |
| PLC, gamma 2 | PLCG2 | Y753 | 41.9 | 63.1 | R.DINSLY*DVSR.M |
| PLC, gamma 2 | PLCG2 | Y1245 | 106.8 | 73.8 | K.EFSVNENQLQLY*QEK.C |
| PLC, gamma 2 | PLCG2 | Y1217 | 46.4 | 68.8 | R.QEELNNQLFLY*DTHQNLR.N |
| PLC, gamma 2 | PLCG2 | Y759 | 20.9 | 38.0 | R.M#Y*VDPSEINPSMPQR.T |
| Pleckstrin homology domain-containing family G member 1 | PLEKHG1 | Y1280 | 75.6 | 57.5 | K.ETDGDEDDY*VEIK.S |
| Poly(rC) binding protein 2 | PCBP2 | Y201 | 61.3 | 50.1 | K.GVTIPYRPKPSSSPVIFAGGQAYTIQGQY*AIPQPDLTK.L |
| Polyadenylate binding protein 1 | PABPC1 | Y364 | 70.0 | 67.5 | R.IVATKPLY*VALAQR.K |
| Polypyrimidine tract binding protein 1 | PTBP1 | Y127 | 49.2 | 45.3 | R.GQPIY*IQFSNHK.E |
| Prolyl endopeptidase | PREP | Y71 | 32.1 | 61.2 | R.M#TELY*DYPK.Y |
| Proteasome 26S subunit, non-ATPase, 14 | PSMD14 | Y32 | 28.2 | 72.9 | R.LGGGM#PGLGQGPPTDAPAVDTAEQVY*ISSLALLK.M |
| Proteasome subunit alpha type 2 | PSMA2 | Y101 | 33.4 | 56.3 | K.LAQQYYLVY*QEPIPTAQLVQR.V |
| Proteasome subunit alpha type 2 | PSMA2 | Y57 | 44.6 | 23.8 | K.SILY*DER.S |
| Protein kinase C delta | PRKCD | Y313 | 120.0 | 53.6 | R.RSDSASSEPVGIY*QGFEK.K |
| Protein kinase C delta | PRKCD | S304Y313 | 27.6 | 48.3 | R.SDS*ASSEPVGIY*QGFEK.K |
| Protein kinase C delta | PRKCD | Y374 | 100.0 | 14.1 | R.GEY*FAIK.A |
| Protein kinase C delta | PRKCD | S306Y313 | 32.2 | 18.6 | R.RSDSAS*SEPVGIY*QGFEK.K |
| Protein kinase C delta | PRKCD | Y64 | 40.7 | 26.8 | K.STFDAHIY*EGR.V |
| Protein THEMIS2 | THEMIS2 | Y632 | 100.0 | 53.6 | R.QDLDDDEHDY*EEILEQFQK.T |
| Protein tyrosine kinase TXK | TXK | Y420 | 32.1 | 68.2 | R.YVLDDEY*VSSFGAK.F |
| Protein tyrosine phosphatase receptor type C | PTPRC | Y681 | 112.6 | 17.0 | R.Y*VDILPYDYNR.V |
| Protein tyrosine phosphatase, non-receptor type 11 | PTPN11 | Y580 | 100.0 | 91.4 | R.VY*ENVGLMQQQK.S |
| Protein tyrosine phosphatase, non-receptor type 11 | PTPN11 | Y62 | 22.0 | 100.5 | K.IQNTGDY*YDLYGGEK.F |
| Protein tyrosine phosphatase, non-receptor type 11 | PTPN11 | Y542 | 16.9 | 46.7 | R.KGHEY*TNIK.Y |
| Protein tyrosine phosphatase, non-receptor type 6 | PTPN6 | Y536 | 13.9 | 56.3 | K.GQESEY*GNITYPPAM#K.N |
| Protein tyrosine phosphatase, non-receptor type 6 | PTPN6 | Y564 | 111.4 | 53.4 | K.HKEDVY*ENLHTK.N |
| Protein tyrosine phosphatase, non-receptor type 6 | PTPN6 | Y301 | 26.0 | 66.9 | R.DSNIPGSDY*INANYIK.N |
| Protein tyrosine phosphatase, receptor type, alpha | PTPRA | Y798 | 26.7 | 70.1 | K.VVQEYIDAFSDY*ANFK.- |
| PRP4 pre-mRNA processing factor 4 homolog B | PRPF4B | Y849 | 52.3 | 82.5 | K.LCDFGSASHVADNDITPY*LVSR.F |
| PRP4 pre-mRNA processing factor 4 homolog B | PRPF4B | S839Y849 | 26.4 | 58.7 | K.LCDFGSAS*HVADNDITPY*LVSR.F |
| PRP4 pre-mRNA processing factor 4 homolog B | PRPF4B | S837Y849 | 31.0 | 69.8 | K.LCDFGS*ASHVADNDITPY*LVSR.F |
| PTK2B protein tyrosine kinase 2 beta | PTK2B | Y580 | 16.9 | 58.6 | R.YIEDEDYY*K.A |
| PTK2B protein tyrosine kinase 2 beta | PTK2B | Y579Y580 | 40.0 | 64.4 | R.YIEDEDY*Y*KASVTR.L |
| PTK2B protein tyrosine kinase 2 beta | PTK2B | Y849 | 77.7 | 57.3 | K.EVGY*LEFTGPPQKPPR.L |
| PTK2B protein tyrosine kinase 2 beta | PTK2B | Y402 | 89.5 | 99.9 | R.SHLSESCSIESDIY*AEIPDETLR.R |
| PTK2B protein tyrosine kinase 2 beta | PTK2B | Y579 | 25.4 | 23.5 | R.YIEDEDY*YKASVTR.L |
| PTK2B protein tyrosine kinase 2 beta | PTK2B | Y819 | 100.0 | 45.8 | K.QM#VEDY*QWLR.Q |
| PTPRF interacting protein binding protein 1 | PPFIBP1 | Y336 | 32.8 | 22.5 | K.GKDGEY*EELLNSSSISSLLDAQGFSDLEK.S |
| Putative translation initiation factor | EIF1 | Y30 | 36.4 | 35.5 | K.GDDLLPAGTEDY*IHIR.I |
| Pyruvate dehydrogenase complex, E1-alpha polypeptide 1 | PDHA1 | Y366 | 24.1 | 23.1 | K.EIEDAAQFATADPEPPLEELGY*HIYSSDPPFEVR.G |
| Pyruvate dehydrogenase complex, E1-alpha polypeptide 1 | PDHA1 | Y369 | 6.9 | 19.3 | R.KEIEDAAQFATADPEPPLEELGYHIY*SSDPPFEVR.G |
| Pyruvate dehydrogenase E1 alpha subunit, testis specific form | PDHA1 | S293Y299 | 3.1 | 34.0 | R.YHGHSM#S*DPGVSY*R.T |
| Pyruvate dehydrogenase E1 alpha subunit, testis specific form | PDHA1 | S291Y299 | 6.4 | 21.3 | R.YHGHS*MSDPGVSY*R.T |
| Pyruvate dehydrogenase E1 alpha subunit, testis specific form | PDHA1 | Y287S293 | 0.7 | 20.8 | R.Y*HGHSMS*DPGVSYR.T |
| Pyruvate kinase 3 | PKM | Y105 | 59.4 | 70.4 | R.TATESFASDPILY*RPVAVALDTK.G |
| Pyruvate kinase 3 | PKM | Y148 | 96.5 | 49.3 | K.ITLDNAY*MEK.C |
| Pyruvate kinase 3 | PKM | Y390 | 100.0 | 49.1 | R.EAEAAIY*HLQLFEELRR.L |
| Pyruvate kinase 3 | PKM | Y175 | 193.2 | 81.7 | K.IY*VDDGLISLQVK.Q |
| RAB GTPase activating protein 1 like | RABGAP1 | Y455 | 80.2 | 70.7 | K.GHTNAGDAIY*EVVSLQR.E |
| RAB10 | RAB10 | Y6 | 12.1 | 32.5 | K.KTY*DLLFK.L |
| RAB7 | RAB7A | Y183 | 20.6 | 28.2 | K.QETEVELY*NEFPEPIKLDK.N |
| RAC GTPase activating protein 1 | RACGAP1 | Y241 | 7.3 | 22.0 | K.TTVTVPNDGGPIEAVSTIETVPY*WTR.S |
| Raf kinase inhibitor protein | PEBP1 | Y181 | 96.6 | 45.3 | K.LY*EQLSGK.- |
| Raft linking protein | RFTN1 | Y20 | 30.2 | 42.2 | K.RPGNIY*STLK.R |
| Raft linking protein | RFTN1 | Y122 | 21.4 | 57.3 | K.TDLHNEGY*ILELDCCSSLDHPTDQK.L |
| Rap guanine nucleotide exchange factor GEF 6 | RAPGEF6 | Y29 | 19.3 | 23.1 | R.TPEDLNTIY*SYLHGMEILSNLR.E |
| Ras related protein Rab35 | RAB35 | Y5 | 100.0 | 14.6 | R.DY*DHLFK.L |
| RasGAP | RASA1 | Y460 | 44.7 | 29.6 | K.EIY*NTIR.R |
| Replication factor C1 | RFC1 | Y282S283 | 6.0 | 21.9 | R.SY*S*PR.S |
| Retinoblastoma 1 | RB1 | Y790S794 | 23.2 | 34.1 | R.SPY*KFPS*SPLR.I |

Peak Area: <-10, -5, -3, 0, 3, 5, >10

%CV: 0, 29, 42, 71, 86, >100

**White dots-** Significant change in peptide abundance at 5 %FDR compared to the timepoint with the minimum peak area for a given PSM

| Protein Name | Gene | Phosphosites | Ascor | MOWSE | Sequence |
|---|---|---|---|---|---|
| Rho GTPase-activating protein 27 | ARHGAP2 | Y28 | 100.0 | 49.5 | R.ALPAQVDDPPEPVY*ANIER.Q |
| Rho guanine nucleotide exchange factor 2 | ARHGEF2 | Y893 | 7.8 | 46.3 | R.SLPAGDALY*LSFNPPQPSR.G |
| Ribosomal protein L10a | RPL10A | Y11 | 28.9 | 28.2 | R.DTLY*EAVR.E |
| Ribosomal protein L3 | RPL3 | Y307 | 32.1 | 73.1 | K.NNASTDY*DLSDK.S |
| Ribosomal protein S10 | RPS10- | Y12 | 100.0 | 50.4 | R.IAIY*ELLFK.E |
| Ribosomal protein S2 | RPS2 | Y133 | 100.0 | 58.5 | K.AFVAIGDY*NGHVGLGVK.C |
| Ribosomal protein S3a | RPS3A | Y256 | 66.6 | 27.3 | R.ADGY*EPPVQESV.- |
| Ribosomal protein S8 | RPS8 | Y117 | 15.7 | 20.4 | R.QWYESHY*ALPLGR.K |
| Ribosomal protein, large, P0 | RPLP0 | Y24 | 100.0 | 49.1 | K.IIQLLDDY*PK.C |
| RNA polymerase II subunit 2 | POLR2B | Y845 | 100.0 | 67.6 | R.HAIY*DKLDDDGLIAPGVR.V |
| ROCK2 | ROCK2 | Y722 | 67.6 | 60.7 | K.IY*ESIEEAK.S |
| SEC16 homolog A | SEC16A | Y991 | 32.1 | 88.1 | K.ANHSSHQEDTY*GALDFTLSR.T |
| Secretory carrier membrane protein 3 | SCAMP3 | Y35 | 60.4 | 64.6 | R.QY*ATLDVYNPFETR.E |
| septin 1 | SEPT1 | Y5 | 100.0 | 49.3 | K.EY*VGFAALPNQLHR.K |
| Septin 9 | SEPT9 | Y278 | 56.9 | 48.7 | K.APVDFGY*VGIDSILEQM#R.R |
| Serine/threonine protein kinase 9 | CDKL5 | Y171 | 32.1 | 108.1 | R.NLSEGNNANYTEY*VATR.W |
| SERPINE1 mRNA binding protein 1 | SERBP1 | Y207 | 13.9 | 19.4 | R.SSFSHY*SGLK.H |
| Seryl tRNA synthetase 2 | SARS2 | Y52 | 10.0 | 46.7 | R.EGY*SALPQLDIER.F |
| SET protein | SET | Y133 | 89.3 | 53.2 | R.IDFYFDENPY*FENK.V |
| SH3 protein expressed in lymphocytes | SASH3 | Y189 | 42.4 | 59.2 | R.VHTDFTPSPY*DHDSLK.L |
| SHC (Src homology 2 domain containing) transforming protein 1 | SHC1 | Y427 | 21.2 | 85.9 | R.ELFDDPSY*VNVQNLDK.A |
| SHC (Src homology 2 domain containing) transforming protein 1 | SHC1 | Y349 | 89.6 | 25.2 | R.M#AGFDGSAWDEEEEEPPDHQY*YNDFPGK.E |
| SHIP1 | INPP5D | Y864 | 100.0 | 36.1 | K.LY*DFVK.T |
| SHIP1 | INPP5D | Y914 | 68.0 | 60.9 | R.APPCSGSSITEIINPNY*M#GVGPFGPPM#PLHVK.Q |
| SHIP1 | INPP5D | Y1021 | 24.2 | 42.7 | K.NAGDTLPQEDLPLTKPEM#FENPLY*GSLSSFPKPAPR.K |
| SHIP1 | INPP5D | Y795 | 39.1 | 62.4 | K.LKPIISDPEY*LLDQHILISIK.S |
| SHIP1 | INPP5D | Y372 | 56.6 | 21.6 | K.EY*VFADSK.K |
| SHIP1 | INPP5D | Y1021S1023 | 10.3 | 20.0 | K.NAGDTLPQEDLPLTKPEM#FENPLY*GS*LSSFPKPAPR.K |
| SHIP2 | INPPL1 | Y1135 | 76.7 | 82.2 | K.TLSEVDY*APAGPAR.S |
| SHIP2 | INPPL1 | Y886 | 146.2 | 80.4 | R.LY*EWISIDKDEAGAK.S |
| SHIP2 | INPPL1 | Y986 | 62.7 | 55.5 | K.NSFNNPAY*YVLEGVPHQLLPPEPPSPAR.A |
| Signal recognition particle 14 kDa | SRP14 | Y27 | 26.0 | 63.2 | R.TSGSVY*ITLK.K |
| Signaling lymphocytic activation molecule | SLAMF1 | Y281 | 55.2 | 46.6 | K.SLTIY*AQVQKPGPLQK.K |
| Single stranded DNA binding protein 1 | SSBP1 | Y73 | 48.0 | 50.2 | R.SGDSEVY*QLGDVSQK.T |
| SIT | SIT1 | Y148 | 161.6 | 79.5 | K.Y*SEVVLDSEPK.S |
| SIT | SIT1 | Y90 | 33.5 | 60.2 | R.SGESVEEVPLY*GNLHYLQTGR.L |
| SIT | SIT1 | Y127 | 13.9 | 38.9 | R.AAEEVM#CY*TSLQLRPPQGR.I |
| SIT | SIT1 | Y95 | 24.3 | 67.7 | R.SGESVEEVPLYGNLHY*LQTGR.L |
| SIT | SIT1 | Y188 | 13.3 | 21.8 | R.ASFPDQAY*ANSQPAAS.- |
| SKAP55 | SKAP1 | Y232 | 22.8 | 62.4 | K.EETY*DDIDGFDSPSCGSQCRPTILPGSVGIK.E |
| SKAP55 | SKAP1 | Y271 | 13.8 | 42.0 | K.EPTEEKEEEDIY*EVLPDEEHDLEEDESGTR.R |
| Small nuclear ribonucleoprotein 70 kD | SNRNP70 | Y126 | 100.0 | 33.9 | R.EFEVY*GPIKR.I |
| SMARCA6 | HELLS | S60Y65 | 25.4 | 13.9 | R.ES*TEIRY*R.R |
| Solute carrier family 38, member 2 | SLC38A2 | Y41 | 10.8 | 58.3 | K.SHY*ADVDPENQNFLLESNLGK.K |
| Splicing factor 3A, subunit 3 | SF3A3 | Y479 | 25.3 | 63.1 | R.WQPDTEEEY*EDSSGNVVNKK.T |
| Splicing factor 3B subunit 4 | SF3B4 | Y16 | 32.1 | 32.7 | R.NQDATVY*VGGLDEK.V |
| Splicing factor, proline and glutamine rich | SFPQ | Y488 | 25.4 | 18.5 | R.FAQHGTFEY*EYSQR.W |
| Sprouty homolog 1 | SPRY1 | Y53 | 16.9 | 73.9 | R.GSNEY*TEGPSVVK.R |
| SRp30c | SRSF9 | S211Y214 | 16.8 | 32.5 | R.GS*PHY*FSPFRPY.- |
| SRp30c | SRSF9 | Y214S216 | 5.2 | 25.7 | R.GSPHY*FS*PFRPY.- |
| SRp30c | SRSF9 | Y214 | 3.3 | 42.1 | R.GSPHY*FSPFRPY.- |
| STE20 like kinase MST1 | STK4 | Y433 | 100.0 | 41.9 | K.IPQDGDY*EFLK.S |
| Structure specific recognition protein 1 | SSRP1 | Y441 | 3.1 | 40.7 | K.EGM#NPSYDEY*ADSDEDQHDAYLER.M |
| SUGT1 | SUGT1 | Y317 | 26.0 | 29.6 | R.LFQQIY*SDGSDEVKR.A |
| Switch associated protein 70 | SWAP70 | Y517 | 100.0 | 18.8 | R.KQALEQY*EEVKK.K |
| SYK | SYK | Y352 | 42.4 | 58.6 | R.EALPM#DTEVYESPY*ADPEEIRPK.E |
| SYK | SYK | Y323 | 40.6 | 63.6 | R.QESTVSFNPY*EPELAPWAADK.G |
| SYK | SYK | Y348 | 33.2 | 62.6 | R.EALPMDTEVY*ESPYADPEEIRPK.E |
| SYK | SYK | Y74 | 16.9 | 76.7 | R.ELNGTY*AIAGGR.T |
| SYK | SYK | Y28 | 34.8 | 41.4 | R.EEAEDY*LVQGGMSDGLYLLR.Q |
| Synapse associated protein 1 | SYAP1 | Y327 | 51.4 | 61.3 | K.ELQQELQEY*EVVTESEKR.D |
| T cell antigen receptor, zeta | | Y142 | 68.3 | 88.3 | R.GKGHDGLY*QGLSTATK.D |

**Peak Area**: <-10, -5, -3, 0, 3, 5, >10

**%CV**: 0, 29, 42, 71, 86, >100

**White dots-** Significant change in peptide abundance at **5** %FDR compared to the timepoint with the minimum peak area for a given PSM

| Protein Name | Gene | Phosphosites | CarT 0m | CarT 2m | CarT 5m | RajiB 0m | RajiB 2m | RajiB 5m | Ascor | MOWSE | Sequence |
|---|---|---|---|---|---|---|---|---|---|---|---|
| T cell antigen receptor, zeta | | Y83 | | | | | | | 100.0 | 44.3 | R.REEY*DVLDK.R |
| T cell antigen receptor, zeta | | Y153 | | | | | | | 33.7 | 38.6 | K.DTY*DALHMQALPPR.T |
| T cell antigen receptor, zeta | | Y72 | | | | | | | 54.3 | 25.1 | R.SADAPAYQQGQNQLY*NELNLGRR.E |
| T cell antigen receptor, zeta | | Y64Y72 | | | | | | | 61.4 | 56.7 | R.SADAPAY*QQGQNQLY*NELNLGR.R |
| T cell antigen receptor, zeta | | Y111 | | | | | | | 100.0 | 31.6 | R.RKNPQEGLY*NELQK.D |
| T cell antigen receptor, zeta | | Y123 | | | | | | | | 14.2 | K.DKM#AEAY*SEIGM#K.G |
| T cell receptor T3 delta chain | CD3D | Y149 | | | | | | | 100.0 | 37.4 | R.NDQVY*QPLR.D |
| T cell receptor T3 delta chain | CD3D | Y160 | | | | | | | 10.3 | 18.5 | R.DDAQY*SHLGGNWAR.N |
| T Complex 1 | TCP1 | Y545 | | | | | | | -0.1 | 14.2 | K.DDKHGSY*EDAVHSGALND.- |
| T Complex 1 | TCP1 | Y181 | | | | | | | 16.9 | 23.5 | K.Y*TDIR.G |
| Tara like protein | TRIOBP | Y612 | | | | | | | 41.5 | 35.0 | K.YQDVY*VELSHIK.T |
| TATSF1 | HTATSF1 | Y650 | | | | | | | | 27.6 | K.VFDDESDEKEDEEY*ADEK.G |
| TBC1 domain family member 15 (Fragment) | TBC1D15 | Y116 | | | | | | | 74.9 | 55.6 | K.SLSQSFENLLDEPAY*GLIQK.I |
| Tec tyrosine kinase | TEC | Y519 | | | | | | | 16.9 | 65.7 | R.YVLDDQY*TSSSGAK.F |
| Tensin 3 | TNS3 | S332Y354 | | | | | | | 57.7 | 66.6 | R.WDS*YENLSADGEVLHTQGPVDGSLY*AK.V |
| Tensin 3 | TNS3 | Y333Y354 | | | | | | | 10.2 | 29.1 | R.WDSY*ENLSADGEVLHTQGPVDGSLY*AK.V |
| Thioredoxin reductase 1 | TXNRD1 | Y183 | | | | | | | 56.3 | 57.8 | K.VVYENAY*GQFIGPHR.I |
| Tight junction protein 2 | TJP2 | Y1118 | | | | | | | 100.0 | 19.8 | K.HPDIY*AVPIK.T |
| TRA2A | TRA2A | Y87T88 | | | | | | | -0.1 | 21.4 | R.SY*T*PEYR.R |
| Transgelin 2 | TAGLN3 | Y192 | | | | | | | 13.9 | 45.9 | R.GASQAGMTGY*GMPR.Q |
| Transgelin 2 | TAGLN2 | Y8 | | | | | | | 15.8 | 29.3 | R.GPAY*GLSR.E |
| Transketolase | TKT | Y275 | | | | | | | 12.3 | 66.4 | K.NMAEQIIQEIY*SQIQSK.K |
| Tripartite motif containing protein 9 | TRIM9 | Y60 | | | | | | | 32.1 | 73.3 | R.AAGSGVSDY*DYLDLDK.M |
| Tripeptidyl peptidase II | TPP2 | Y645 | | | | | | | 11.7 | 36.8 | K.VNESSHY*DLAFTDVHFKPGQIR.R |
| Tropomyosin 1 alpha chain | TPM2 | Y162 | | | | | | | 100.0 | 36.8 | K.Y*EEVAR.K |
| Tropomyosin 3 | TPM3 | Y162 | | | | | | | 100.0 | 17.2 | K.HIAEEADRKY*EEVAR.K |
| Tubulin, beta | TUBB | Y50 | | | | | | | 13.9 | 23.3 | R.ISVY*YNEATGGK.Y |
| TUFM | TUFM | Y249 | | | | | | | 16.9 | 70.9 | K.LLDAVDTY*IPVPAR.D |
| TUFM | TUFM | Y269 | | | | | | | 15.2 | 69.4 | R.DLEKPFLLPVEAVY*SVPGR.G |
| Tumor necrosis factor receptor superfamily, member 9 | | Y222 | | | | | | | 100.0 | 26.5 | K.LLY*IFK.Q |
| TYK2 | TYK2 | Y292 | | | | | | | 100.0 | 40.7 | R.LLAQAEGEPCY*IR.D |
| TYK2 | TYK2 | Y433 | | | | | | | 10.9 | 43.4 | R.LTADSSHY*LCHEVAPPR.L |
| Tyrosine kinase, non-receptor, 2 | TNK2 | Y284 | | | | | | | 100.0 | 41.8 | R.ALPQNDDHY*VMQEHR.K |
| Tyrosine kinase, non-receptor, 2 | TNK2 | Y827 | | | | | | | 29.1 | 56.1 | K.Y*ATPQVIQAPGPR.A |
| Tyrosine kinase, non-receptor, 2 | TNK2 | Y859 | | | | | | | 10.3 | 14.3 | K.VSSTHY*YLLPERPSYLER.Y |
| Tyrosine protein kinase ABL2 | ABL1 | Y439 | | | | | | | 10.9 | 41.9 | R.LM#TGDTY*TAHAGAK.F |
| Tyrosine protein kinase ABL2 | ABL2 | Y231 | | | | | | | 53.8 | 58.6 | K.VY*VTAESR.F |
| Tyrosine-protein kinase PRAG1 | PRAG1 | Y413 | | | | | | | 17.3 | 19.5 | R.EATQPEPIY*AESTKR.K |
| Ubiquitin activating enzyme 1 | UBA1 | Y55 | | | | | | | 86.5 | 75.1 | K.NGSEADIDEGLY*SR.Q |
| UNC112 related protein 2 | FERMT3 | Y11 | | | | | | | 46.9 | 72.9 | K.TASGDY*IDSSWELR.V |
| UNC119 | UNC119 | Y194 | | | | | | | 6.3 | 32.7 | K.NTCEHIY*DFPPLSEELISEMIR.H |
| UNQ5783 | SCIMP | Y131 | | | | | | | 22.1 | 31.2 | K.TVSIPSYIEPEDDY*DDVEIPANTEK.A |
| Uridine 5`-monophosphate synthase | UMPS | Y37 | | | | | | | 46.9 | 56.6 | K.SGLSSPIY*IDLR.G |
| Vacuolar protein sorting 35 | VPS35 | Y791 | | | | | | | 48.5 | 27.6 | R.ESPESEGPIY*EGLIL.- |
| Vasodilator stimulated phosphoprotein | VASP | Y39 | | | | | | | 14.8 | 14.5 | R.VQIY*HNPTANSFR.V |
| VAV1 | VAV1 | Y267 | | | | | | | 91.2 | 92.5 | K.EALGTPGAANLY*QVFIK.Y |
| VAV1 | VAV1 | Y844 | | | | | | | 30.2 | 20.0 | R.VGWFPANYVEEDYSEY*C.- |
| VAV1 | VAV1 | Y791 | | | | | | | 100.0 | 25.4 | K.ARY*DFCAR.D |
| VAV1 | VAV1 | Y826 | | | | | | | 100.0 | 20.5 | R.GEIY*GR.V |
| VAV2 | VAV2 | Y142 | | | | | | | 37.1 | 64.0 | K.GIRPFPSEETTENDDDVY*R.S |
| VAV3 | VAV3 | Y539 | | | | | | | 28.2 | 49.8 | R.GTFY*QGYLCFK.C |
| VAV3 | VAV3 | Y265 | | | | | | | 93.0 | 100.9 | K.NDQNLY*QVFINYK.E |
| VAV3 | VAV3 | Y367 | | | | | | | 100.0 | 26.5 | K.DLAQY*VNEVK.R |
| Vigilin | HDLBP | Y437 | | | | | | | 100.0 | 16.8 | R.MDY*VEINIDHK.F |
| Vimentin | VIM | Y61 | | | | | | | 43.2 | 63.1 | R.SLYASSPGGVY*ATR.S |
| Vimentin | VIM | Y53 | | | | | | | 13.4 | 26.9 | R.SLY*ASSPGGVYATR.S |
| WD repeat protein 1 | WDR1 | Y238 | | | | | | | 53.7 | 109.7 | K.AHDGGIY*AISWSPDSTHLLSASGDK.T |
| Williams Beuren syndrome chromosome region 1 | EIF4H | Y12 | | | | | | | 33.8 | 33.7 | R.AY*SSFGGGR.G |
| Wiskott-aldrich syndrome protein | WAS | Y291 | | | | | | | 100.0 | 94.4 | K.LIY*DFIEDQGGLEAVR.Q |
| Wiskott-Aldrich syndrome-like | WASL | Y256 | | | | | | | 100.0 | 39.1 | K.VIY*DFIEK.T |
| ZAP70 | ZAP70 | Y492 | | | | | | | 16.9 | 97.4 | K.ALGADDSY*YTAR.S |
| ZAP70 | ZAP70 | Y292 | | | | | | | 32.3 | 55.2 | R.IDTLNSDGY*TPEPAR.I |

**Peak Area**: <-10, -5, -3, 0, 3, 5, >10

**%CV**: 0, 29, 42, 71, 86, >100

**White dots-** Significant change in peptide abundance at **5** %FDR compared to the timepoint with the minimum peak area for a given PSM

| Protein Name | Gene | Phosphosites | CarT 0m | CarT 2m | CarT 5m | RajiB 0m | RajiB 2m | RajiB 5m | Ascor | MOWSE | Sequence |
|---|---|---|---|---|---|---|---|---|---|---|---|
| **ZAP70** | ZAP70 | Y397 | | | | | | | **100.0** | **54.3** | R.EAQIMHQLDNPY*IVR.L |
| **ZAP70** | ZAP70 | Y492Y493 | | | | | | | **28.4** | **81.2** | K.ALGADDSY*Y*TAR.S |
| **ZAP70** | ZAP70 | Y493 | | | | | | | **16.9** | **79.0** | K.ALGADDSYY*TAR.S |
| **ZAP70** | ZAP70 | S491Y493 | | | | | | | **12.1** | **27.4** | K.ALGADDS*YY*TAR.S |
| **ZAP70** | ZAP70 | T286Y292 | | | | | | | **9.7** | **45.8** | R.RIDT*LNSDGY*TPEPAR.I |
| **ZAP70** | ZAP70 | Y319 | | | | | | | **7.3** | **30.7** | R.ITSPDKPRPMPMDTSVYESPY*SDPEELKDK.K |
| **ZAP70** | ZAP70 | Y315 | | | | | | | **9.4** | **19.5** | R.ITSPDKPRPMPMDTSVY*ESPYSDPEELKDK.K |
| **Zinc finger protein 147** | TRIM25 | Y278 | | | | | | | **39.3** | **57.9** | K.FDTIY*QILLK.K |
| **Zinc finger protein 289, ID1 regulated** | ARFGAP2 | Y445 | | | | | | | **100.0** | **21.0** | R.EVDAEY*EAR.S |
| **Zinc finger protein 598** | ZNF598 | Y306 | | | | | | | **100.0** | **71.1** | R.RNEGVVGGEDY*EEVDR.Y |
| **Zinc finger protein A20** | TNFAIP3 | Y443 | | | | | | | **20.8** | **43.0** | R.GEAY*EPLAWNPEESTGGPHSAPPTAPSPFLFSETTAM#K.C |
| **Zinc finger protein, subfamily 1A, member 3** | IKZF3 | Y96 | | | | | | | **46.9** | **20.1** | R.EYNEY*ENIK.L |
| **ZAP70** | ZAP70 | Y319 | | | | | | | **7.3** | **30.7** | R.ITSPDKPRPMPMDTSVYESPY*SDPEELKDK.K |
