## Supplementary Figure 5 for "SILAC phosphoproteomics reveals unique signaling circuits in CAR-T cells and the inhibition of B cell-activating phosphorylation in target cells"

Peak Area: <-10, -5, -3, 0, 3, 5, >10

%CV: 0, 29, 42, 71, 86, >100

**White dots-** Significant change in peptide abundance at **5** %FDR compared to the timepoint with the minimum peak area for a given PSM

| Protein Name | Gene | Phosphosites | CarT 0m | CarT 2m | CarT 5m | RajiB 0m | RajiB 2m | RajiB 5m | Ascor | MOWSE | Sequence |
|---|---|---|---|---|---|---|---|---|---|---|---|
| **14-3-3 epsilon** | YWHAE | S210 | | | | | | | **13.4** | **57.0** | K.AAFDDAIAELDTLS*EESYK.D |
| **14-3-3 epsilon** | YWHAE | T208 | | | | | | | **9.1** | **53.4** | K.AAFDDAIAELDT*LSEESYK.D |
| **14-3-3 Eta** | YWHAH | S46 | | | | | | | **37.7** | **36.8** | R.NLLS*VAYK.N |
| **14-3-3 theta** | YWHAQ | S230 | | | | | | | **16.9** | **27.8** | R.DNLTLWTS*DSAGEECDAAEGAEN.- |
| **14-3-3 theta** | YWHAQ | S232 | | | | | | | **15.3** | **38.5** | R.DNLTLWTSDS*AGEECDAAEGAEN.- |
| **14-3-3 zeta** | YWHAZ | S207 | | | | | | | **24.1** | **93.9** | K.TAFDEAIAELDTLS*EESYK.D |
| **14-3-3 zeta** | YWHAZ | T205 | | | | | | | **10.2** | **43.5** | K.TAFDEAIAELDT*LSEESYK.D |
| **3 Phosphoinositide dependent protein kinase 1** | PDPK1 | S241 | | | | | | | **57.2** | **129.2** | R.ANS*FVGTAQYVSPELLTEK.S |
| **3 Phosphoinositide dependent protein kinase 1** | PDPK1 | T245 | | | | | | | **9.2** | **38.2** | R.ANSFVGT*AQYVSPELLTEK.S |
| **5 azacytidine induced 1** | CEP131 | S89 | | | | | | | **10.1** | **59.8** | R.SGS*PRPTEPTDFLMLFEGSPSGK.K |
| **5 azacytidine induced 1** | CEP131 | S87S105 | | | | | | | **11.3** | **35.9** | R.S*GSPRPTEPTDFLMLFEGS*PSGK.K |
| | CEP131 | S89S105 | | | | | | | **3.2** | **32.4** | R.SGS*PRPTEPTDFLMLFEGS*PSGKK.R |
| **5 azacytidine induced 1** | CEP131 | T96S105 | | | | | | | **9.8** | **15.7** | R.SGSPRPTEPT*DFLMLFEGS*PSGKK.R |
| **5 azacytidine induced 1** | CEP131 | T93 | | | | | | | **9.2** | **37.1** | R.SGSPRPT*EPTDFLMLFEGSPSGK.K |
| **5 azacytidine induced 1** | CEP131 | T96S107 | | | | | | | **9.6** | **21.5** | R.SGSPRPTEPT*DFLMLFEGSPS*GKK.R |
| **5\`-3\` exoribonuclease 2** | XRN2 | S448 | | | | | | | **46.9** | **26.4** | R.NS*PGSQVASNPR.Q |
| **5\`-3\` exoribonuclease 2** | XRN2 | S499S501 | | | | | | | **100.0** | **105.2** | R.KAEDS*DS*EPEPEDNVR.L |
| **5\`-3\` exoribonuclease 2** | XRN2 | T478 | | | | | | | **15.2** | **59.4** | R.MQNNSSPSISPNT*SFTSDGSPSPLGGIK.R |
| **5\`-3\` exoribonuclease 2** | XRN2 | S473 | | | | | | | **8.4** | **133.5** | R.MQNNSSPS*ISPNTSFTSDGSPSPLGGIK.R |
| **5\`-3\` exoribonuclease 2** | XRN2 | S482 | | | | | | | **-0.2** | **19.5** | R.MQNNSSPSISPNTSFTS*DGSPSPLGGIK.R |
| **5\`-3\` exoribonuclease 2** | XRN2 | S471 | | | | | | | **2.3** | **86.7** | R.MQNNSS*PSISPNTSFTSDGSPSPLGGIK.R |
| **5\`-3\` exoribonuclease 2** | XRN2 | S475 | | | | | | | **6.0** | **63.6** | R.MQNNSSPSIS*PNTSFTSDGSPSPLGGIK.R |
| **5\`-3\` exoribonuclease 2** | XRN2 | S471S473 | | | | | | | **12.3** | **79.6** | R.MQNNSS*PS*ISPNTSFTSDGSPSPLGGIK.R |
| **5\`-3\` exoribonuclease 2** | XRN2 | S475T478 | | | | | | | **3.2** | **79.5** | R.MQNNSSPSIS*PNT*SFTSDGSPSPLGGIK.R |
| **5\`-3\` exoribonuclease 2** | XRN2 | S473S475 | | | | | | | | **11.4** | R.MQNNSSPS*IS*PNTSFTSDGSPSPLGGIK.R |
| **5\`-3\` exoribonuclease 2** | XRN2 | S471S475 | | | | | | | **2.4** | **90.4** | R.MQNNSS*PSIS*PNTSFTSDGSPSPLGGIK.R |
| **5\`-3\` exoribonuclease 2** | XRN2 | S470S471 | | | | | | | **11.3** | **38.1** | R.MQNNS*S*PSISPNTSFTSDGSPSPLGGIK.R |
| **5\`-3\` exoribonuclease 2** | XRN2 | T478S479 | | | | | | | **13.6** | **11.8** | R.MQNNSSPSISPNT*S*FTSDGSPSPLGGIK.R |
| **5\`-3\` exoribonuclease 2** | XRN2 | S470 | | | | | | | **7.4** | **25.7** | R.MQNNS*SPSISPNTSFTSDGSPSPLGGIK.R |
| **5\`-3\` exoribonuclease 2** | XRN2 | S471T478 | | | | | | | **5.4** | **73.1** | R.MQNNSS*PSISPNT*SFTSDGSPSPLGGIK.R |
| **5\`, 3\`-Nucleotidase, cytosolic** | NT5C | S182 | | | | | | | **51.2** | **58.2** | R.LLS*WSDNWR.E |
| **6-phosphofructo-2 kinase/fructose-2,6-biphosphatase 2** | PFKFB2 | S466 | | | | | | | **51.2** | **50.9** | R.RNS*FTPLSSSNTIR.R |
| **6-phosphofructo-2 kinase/fructose-2,6-biphosphatase 2** | PFKFB2 | Y482 | | | | | | | **9.0** | **14.5** | R.NY*SVGSRPLKPLSPLR.A |
| **6-phosphofructo-2 kinase/fructose-2,6-biphosphatase 2** | PFKFB2 | S483 | | | | | | | **24.1** | **33.3** | R.NYS*VGSRPLKPLSPLR.A |
| **6-phosphofructo-2 kinase/fructose-2,6-biphosphatase 2** | PFKFB2 | S486S493 | | | | | | | **37.9** | **18.0** | R.NYSVGS*RPLKPLS*PLR.A |
| **6-phosphofructo-2 kinase/fructose-2,6-biphosphatase 2** | PFKFB2 | S483S493 | | | | | | | **13.0** | **21.2** | R.NYS*VGSRPLKPLS*PLR.A |
| **6-phosphofructo-2 kinase/fructose-2,6-biphosphatase 2** | PFKFB2 | S486 | | | | | | | **16.1** | **33.2** | R.NYSVGS*RPLKPLSPLR.A |
| **6-phosphofructo-2 kinase/fructose-2,6-biphosphatase 2** | PFKFB2 | Y482S493 | | | | | | | **8.3** | **14.2** | R.NY*SVGSRPLKPLS*PLR.A |
| **60S ribosomal protein L22-like 1** | RPL22L1 | S117 | | | | | | | **12.1** | **40.6** | R.YFQISQDEDES*ESED.- |
| **7-dehydrocholesterol reductase** | DHCR7 | S14 | | | | | | | **46.8** | **13.8** | K.S*LDGVTNDR.T |
| **A kinase anchor protein 11** | AKAP11 | S1337S1340 | | | | | | | **10.6** | **23.5** | K.YPS*CES*VTDEYAGHLIQILK.Q |
| **A kinase anchor protein 11** | AKAP11 | Y1335S1340 | | | | | | | **23.9** | **58.3** | K.Y*PSCES*VTDEYAGHLIQILK.Q |
| **A kinase anchor protein 11** | AKAP11 | S1242 | | | | | | | **-0.5** | **22.1** | R.SVS*PTFLNPSDENLK.T |
| **A kinase anchor protein 2** | AKAP2 | S383 | | | | | | | **17.8** | **82.0** | R.DALGDSLQVPVS*PSSTTSSR.C |
| **A kinase anchor protein 2** | AKAP2 | S951 | | | | | | | **33.6** | **63.1** | R.TLS*MIEEEIR.A |
| **A kinase anchor protein 2** | AKAP2 | S979 | | | | | | | **39.3** | **31.6** | R.QVLQSTQS*PR.T |
| **A kinase anchor protein 2** | AKAP2 | T949 | | | | | | | **10.6** | **60.7** | R.T*LSMIEEEIR.A |
| **A kinase anchor protein 2** | AKAP2 | S385 | | | | | | | **7.8** | **47.0** | R.DALGDSLQVPVSPS*STTSSR.C |
| **A kinase anchor protein 2** | AKAP2 | T1012 | | | | | | | **1.6** | **12.1** | K.VKPPPSPTT*EGPSLQPDLAPEEAAGTQRPK.N |
| **A kinase anchor protein 2** | AKAP2 | S1016 | | | | | | | **18.4** | **13.2** | K.VKPPPSPTTEGPS*LQPDLAPEEAAGTQRPK.N |
| **A kinase anchor protein 2** | AKAP2 | S862 | | | | | | | **13.4** | **12.9** | R.DGAEQQGPEATVEEAEAAAFGSEKPQSMFEPPQVSS*PVQEK.R |
| **A kinase anchor protein 2** | AKAP2 | T837 | | | | | | | | **16.3** | R.DGAEQQGPEAT*VEEAEAAAFGSEKPQSMFEPPQVSSPVQEK.R |
| **A kinase anchoring protein 13** | AKAP13 | S2732 | | | | | | | **19.8** | **56.2** | K.SGSLDSELSVS*PK.R |
| **A kinase anchoring protein 13** | AKAP13 | S1880 | | | | | | | **104.4** | **85.5** | R.S*AVLLVDETATTPIFANR.R |
| **A kinase anchoring protein 13** | AKAP13 | S1931S1936 | | | | | | | **14.4** | **22.5** | K.FLS*HSTDS*LNK.I |
| **A kinase anchoring protein 13** | AKAP13 | S1559 | | | | | | | **9.9** | **16.7** | R.SLS*PFRR.H |
| **A kinase anchoring protein 13** | AKAP13 | S1910 | | | | | | | **5.1** | **28.6** | K.SVS*IQNITGVGNDENMSNTWK.F |
| **A kinase anchoring protein 13** | AKAP13 | S1931S1933 | | | | | | | **14.7** | **17.6** | K.FLS*HS*TDSLNK.I |
| **A kinase PRKA anchor protein 8 like** | AKAP8L | S297S300S302 | | | | | | | **1.7** | **68.6** | R.TDCS*DNS*DS*DNDEGTEGEATEGLEGTEAVEK.G |
| **A kinase PRKA anchor protein 8 like** | AKAP8L | S297S302T308 | | | | | | | **2.4** | **40.3** | R.TDCS*DNSDS*DNDEGT*EGEATEGLEGTEAVEK.G |
| **A_Raf** | ARAF | S299 | | | | | | | **9.8** | **27.1** | R.DS*GYYWEVPPSEVQLLK.R |
| **A_Raf** | ARAF | T213 | | | | | | | **-0.3** | **41.3** | R.ST*STPNVHMVSTTAPMDSNLIQLTGQSFSTDAAGSR.G |

Peak Area: <-10, -5, -3, 0, 3, 5, >10

%CV: 0, 29, 42, 71, 86, >100

**White dots-** Significant change in peptide abundance at **5** %FDR compared to the timepoint with the minimum peak area for a given PSM

| Protein Name | Gene | Phosphosites | CarT 0m | CarT 2m | CarT 5m | RajiB 0m | RajiB 2m | RajiB 5m | Ascor | MOWSE | Sequence |
|---|---|---|---|---|---|---|---|---|---|---|---|
| **A_Raf** | ARAF | S212 | | | | | | | 11.5 | 39.9 | R.S*TSTPNVHMVSTTAPMDSNLIQLTGQSFSTDAAGSR.G |
| **ABL interactor 1** | ABI1 | S231 | | | | | | | 11.4 | 24.0 | R.TAS*LNQRPR.T |
| **Absent in melanoma 1** | CRYBG1 | T20 | | | | | | | -0.1 | 44.5 | K.ST*DSPGADAELPESAAR.D |
| **Acetyl-CoA carboxylase alpha** | ACACA | S117 | | | | | | | 22.5 | 62.3 | R.SSMS*GLHLVK.Q |
| **Acetyl-CoA carboxylase alpha** | ACACA | S62 | | | | | | | 10.9 | 61.9 | R.FIIGSVS*EDNSEDEISNLVK.L |
| **Acetyl-CoA carboxylase alpha** | ACACA | S60S66 | | | | | | | 46.7 | 105.3 | R.FIIGS*VSEDNS*EDEISNLVK.L |
| **Acetyl-CoA carboxylase alpha** | ACACA | S62S66 | | | | | | | 41.9 | 115.7 | R.FIIGSVS*EDNS*EDEISNLVK.L |
| **Acetyl-CoA carboxylase alpha** | ACACA | S60S62 | | | | | | | 20.5 | 14.2 | R.FIIGS*VS*EDNSEDEISNLVK.L |
| **Acetyl-CoA carboxylase alpha** | ACACA | T1249S1253 | | | | | | | 6.5 | 11.6 | R.MSFSSNLNHYGMT*HVAS*VSDVLLDNSFTPPCQR.M |
| **Acetyl-CoA carboxylase alpha** | ACACA | S87S90 | | | | | | | 3.1 | 38.3 | K.EGSLS*PAS*VGSDTLSDLGISSLQDGLALHIR.S |
| **Acetyl-CoA carboxylase alpha** | ACACA | S85S87S90 | | | | | | | 1.2 | 43.1 | K.EGS*LS*PAS*VGSDTLSDLGISSLQDGLALHIR.S |
| **Acetyl-CoA carboxylase alpha** | ACACA | S85S87S93 | | | | | | | 5.4 | 32.8 | K.EGS*LS*PASVGS*DTLSDLGISSLQDGLALHIR.S |
| **Acetyl-CoA carboxylase alpha** | ACACA | S1296 | | | | | | | 7.4 | 11.5 | R.IFDEVMGCFSDS*PPQSPTFPEAGHTSLYDEDKVPR.D |
| **Acetyl-CoA carboxylase alpha** | ACACA | S1296T1302 | | | | | | | 22.2 | 70.5 | R.IFDEVMGCFSDS*PPQSPT*FPEAGHTSLYDEDKVPR.D |
| **Acetyl-CoA carboxylase alpha** | ACACA | S1296S1300 | | | | | | | 26.0 | 52.8 | R.IFDEVMGCFSDS*PPQS*PTFPEAGHTSLYDEDKVPR.D |
| **Acetyl-CoA carboxylase alpha** | ACACA | S1294S1296 | | | | | | | | 46.6 | R.IFDEVMGCFS*DS*PPQSPTFPEAGHTSLYDEDKVPR.D |
| **Acetyl-CoA carboxylase alpha** | ACACA | S85S87 | | | | | | | 7.3 | 43.4 | K.EGS*LS*PASVGSDTLSDLGISSLQDGLALHIR.S |
| **Acetyl-CoA carboxylase alpha** | ACACA | S85S90 | | | | | | | 12.2 | 63.9 | K.EGS*LSPAS*VGSDTLSDLGISSLQDGLALHIR.S |
| **Acetyl-CoA carboxylase alpha** | ACACA | S85S87S90T95 | | | | | | | 5.1 | 57.3 | K.EGS*LS*PAS*VGSDT*LSDLGISSLQDGLALHIR.S |
| **Acetyl-CoA carboxylase alpha** | ACACA | S87S90S93S97 | | | | | | | 5.0 | 44.5 | K.EGSLS*PAS*VGS*DTLS*DLGISSLQDGLALHIR.S |
| **Acetyl-CoA carboxylase alpha** | ACACA | S66 | | | | | | | 9.1 | 74.6 | R.FIIGSVSEDNS*EDEISNLVK.L |
| **Acetyl-CoA carboxylase alpha** | ACACA | S60 | | | | | | | 5.5 | 69.1 | R.FIIGS*VSEDNSEDEISNLVK.L |
| **Acetyl-CoA carboxylase alpha** | ACACA | S1253S1255 | | | | | | | 2.2 | 24.5 | R.MSFSSNLNHYGMTHVAS*VS*DVLLDNSFTPPCQR.M |
| **Acetyl-CoA carboxylase alpha** | ACACA | S1238T1249 | | | | | | | 2.5 | 24.2 | R.MS*FSSNLNHYGMT*HVASVSDVLLDNSFTPPCQR.M |
| **Acetyl-CoA carboxylase alpha** | ACACA | Y1246S1255 | | | | | | | 10.0 | 58.8 | R.MSFSSNLNHY*GMTHVASVS*DVLLDNSFTPPCQR.M |
| **Acetyl-CoA carboxylase alpha** | ACACA | Y1246T1249 | | | | | | | 3.3 | 12.8 | R.MSFSSNLNHY*GMT*HVASVSDVLLDNSFTPPCQR.M |
| **Acetyl-CoA carboxylase alpha** | ACACA | S1238S1240 | | | | | | | 2.2 | 23.7 | R.MS*FS*SNLNHYGMTHVASVSDVLLDNSFTPPCQR.M |
| **Acetyl-CoA carboxylase alpha** | ACACA | S1300T1302 | | | | | | | 27.8 | 17.3 | R.IFDEVMGCFSDSPPQS*PT*FPEAGHTSLYDEDKVPR.D |
| **Acetyl-CoA carboxylase alpha** | ACACA | S1294T1302 | | | | | | | 11.9 | 38.0 | R.IFDEVMGCFS*DSPPQSPT*FPEAGHTSLYDEDK.V |
| | ACACA | S1294S1300 | | | | | | | 6.4 | 20.2 | R.IFDEVMGCFS*DSPPQS*PTFPEAGHTSLYDEDK.V |
| **Acetyl-CoA carboxylase alpha** | ACACA | S85T95 | | | | | | | 6.1 | 21.3 | K.EGS*LSPASVGSDT*LSDLGISSLQDGLALHIR.S |
| **Acetyl-CoA carboxylase alpha** | ACACA | S87S90S93 | | | | | | | 5.8 | 36.6 | K.EGSLS*PAS*VGS*DTLSDLGISSLQDGLALHIR.S |
| **Acetyl-CoA carboxylase alpha** | ACACA | S85S87S93T95 | | | | | | | 17.0 | 71.4 | K.EGS*LS*PASVGS*DT*LSDLGISSLQDGLALHIR.S |
| **Acetylserotonin methyltransferase-like** | ASMTL | S239 | | | | | | | 66.0 | 120.0 | K.HDSIPAADTFEDLS*DVEGGGSEPTQR.D |
| **Acetylserotonin methyltransferase-like** | ASMTL | T234 | | | | | | | 16.5 | 13.0 | K.HDSIPAADT*FEDLSDVEGGGSEPTQR.D |
| **Acheron** | FAM27E3 | S247S248 | | | | | | | 100.0 | 28.2 | R.RIS*S*R.G |
| **TRAF3-interacting JNK-activating modulator** | TRAF3IP3 | S110 | | | | | | | 100.0 | 30.6 | R.RIS*SPR.E |
| **ACINUS** | ACIN1 | T563 | | | | | | | 18.4 | 41.9 | R.RASHT*LLPSHR.L |
| **ACINUS** | ACIN1 | S1004 | | | | | | | 71.2 | 36.0 | R.TAQVPS*PPR.G |
| **ACINUS** | ACIN1 | S490 | | | | | | | 121.6 | 70.3 | K.AES*PAEKVPEESVLPLVQK.S |
| **ACINUS** | ACIN1 | T408 | | | | | | | 7.9 | 58.3 | K.ASLVALPEQT*ASEEETPPPLLTK.E |
| **ACINUS** | ACIN1 | T408T414 | | | | | | | 12.4 | 27.7 | K.ASLVALPEQT*ASEEET*PPPLLTK.E |
| **ACINUS** | ACIN1 | S386S388 | | | | | | | 32.4 | 24.1 | K.SKS*PS*PPR.L |
| **ACINUS** | ACIN1 | S729 | | | | | | | 100.0 | 29.5 | R.LQPERGS*PK.K |
| **ACINUS** | ACIN1 | S605 | | | | | | | 35.1 | 25.1 | R.SHS*PLR.S |
| **ACINUS** | ACIN1 | S898 | | | | | | | 61.4 | 57.3 | R.IS*EDETER.N |
| **ACINUS** | ACIN1 | S710 | | | | | | | 46.5 | 54.6 | R.RLS*QPESAEK.H |
| **ACINUS** | ACIN1 | S655S657 | | | | | | | 60.8 | 48.6 | K.S*LS*PGVSR.D |
| **ACINUS** | ACIN1 | S478S482 | | | | | | | 26.4 | 48.8 | R.ELLVSQHTVQLVGGLS*PLSS*PSDTK.A |
| **ACINUS** | ACIN1 | S478S481 | | | | | | | 24.0 | 60.3 | R.ELLVSQHTVQLVGGLS*PLS*SPSDTK.A |
| **ACINUS** | ACIN1 | S216 | | | | | | | 16.9 | 63.3 | K.SSSISEEKGDS*DDEKPR.K |
| **ACINUS** | ACIN1 | S895 | | | | | | | 44.6 | 24.2 | K.SLIPDIKPLAGQEAVVDLHADDS*RISEDETER.N |
| **ACINUS** | ACIN1 | S467T470S478 | | | | | | | | 47.2 | R.ELLVS*QHT*VQLVGGLS*PLSSPSDTKAESPAEKVPEESVLPLVQK.S |
| **ACINUS** | ACIN1 | S1329S1331T1 | | | | | | | 100.0 | 20.2 | R.S*RS*T*PVRDR.G |
| **ACINUS** | ACIN1 | S166S169 | | | | | | | 14.0 | 20.5 | R.EAAELEEAS*AES*EDEM#IHPEGVASLLPPDFQSSLERPELELSR.H |
| **ACINUS** | ACIN1 | S1331T1332 | | | | | | | 10.4 | 23.9 | R.SRS*T*PVRDR.G |
| **ACINUS** | ACIN1 | S603 | | | | | | | 24.6 | 27.8 | R.S*HSPLR.S |
| **ACINUS** | ACIN1 | S561 | | | | | | | 25.8 | 32.8 | R.RAS*HTLLPSHR.L |
| **ACINUS** | ACIN1 | S240S243 | | | | | | | 61.8 | 39.1 | K.LS*EGS*QPAEEEEDQETPSR.N |
| **ACINUS** | ACIN1 | S410 | | | | | | | 2.2 | 67.7 | K.ASLVALPEQTAS*EEETPPPLLTK.E |
| **ACINUS** | ACIN1 | S410T414 | | | | | | | 3.8 | 22.7 | K.ASLVALPEQTAS*EEET*PPPLLTK.E |
| **ACINUS** | ACIN1 | S1284 | | | | | | | 100.0 | 12.9 | K.RREHS*R.E |

Peak Area: <-10, -5, -3, 0, 3, 5, >10

%CV: 0, 29, 42, 71, 86, >100

**White dots-** Significant change in peptide abundance at **5** %FDR compared to the timepoint with the minimum peak area for a given PSM

CarT (0m, 2m, 5m) — RajiB (0m, 2m, 5m)

| Protein Name | Gene | Phosphosites | Ascor | MOWSE | Sequence |
|---|---|---|---|---|---|
| **ACINUS** | ACIN1 | S365 | **9.1** | **24.9** | K.TTS*PLEEEER.E |
| **ACINUS** | ACIN1 | S657S661 | **8.4** | **12.4** | K.SLS*PGVS*R.D |
| **ACINUS** | ACIN1 | S478 | **24.4** | **17.0** | R.ELLVSQHTVQLVGGLS*PLSSPSDTK.A |
| **ACINUS** | ACIN1 | S210 | **11.9** | **42.3** | K.SSSIS*EEKGDSDDEKPR.K |
| **Actin alpha, cardiac muscle** | ACTB | Y242 | **-0.3** | **93.2** | R.SY*ELPDGQVITIGNER.F |
| **Actin binding LIM protein 1** | ABLIM1 | S452 | **13.8** | **76.7** | R.STS*QGSINSPVYSR.H |
| **Actin binding LIM protein 1** | ABLIM1 | S655 | **25.4** | **24.1** | K.TAS*LPGYGR.N |
| **Actin binding LIM protein 1** | ABLIM1 | S431 | **25.3** | **80.5** | R.TLS*PTPSAEGYQDVR.D |
| **Actin binding LIM protein 1** | ABLIM1 | S185 | **98.4** | **40.1** | R.S*PQHFHRPDQGINIYR.K |
| **Actin binding LIM protein 1** | ABLIM1 | S352 | **9.8** | **53.6** | R.TS*SESIYSRPGSSIPGSPGHTIYAK.V |
| **Actin binding LIM protein 1** | ABLIM1 | S352S367 | **16.1** | **35.1** | R.TS*SESIYSRPGSSIPGS*PGHTIYAK.V |
| **Actin binding LIM protein 1** | ABLIM1 | S367 | **38.4** | **24.9** | R.TSSESIYSRPGSSIPGS*PGHTIYAK.V |
| **Actin binding LIM protein 1** | ABLIM1 | S353 | **1.3** | **32.2** | R.TSS*ESIYSRPGSSIPGSPGHTIYAK.V |
| **Actin binding LIM protein 1** | ABLIM1 | S353S367 | **13.8** | **21.9** | R.TSS*ESIYSRPGSSIPGS*PGHTIYAK.V |
| **Actin filament associated protein** | AFAP1 | S342S343 | **6.5** | **24.0** | K.KKPSTDEQTS*S*AEEDVPTCGYLNVLSNSR.W |
| **Actin gamma 1** | ACTB | S232 | **28.2** | **12.3** | K.LCYVALDFEQEMATAAS*SSSLEK.S |
| **Actin-bundling protein with BAIAP2 homology** | MTSS1L | S639T643 | | **13.1** | K.RLS*LPNT*AWGSPSPEAAGYPGAGAEDEQQQLAANRHSLVEK.L |
| **Actin-bundling protein with BAIAP2 homology** | MTSS1L | S639S673 | **10.1** | **12.0** | K.RLS*LPNTAWGSPSPEAAGYPGAGAEDEQQQLAANRHS*LVEK.L |
| **Actin-like 6A** | ACTL6A | S233 | **100.0** | **34.2** | R.EGS*PANWK.R |
| **Activating transcription factor 1** | PKP4 | S63 | **13.9** | **19.2** | R.RPS*YR.K |
| **Activating transcription factor 2** | ATF2 | T69T71 | **16.4** | **37.6** | R.NDSVIVADQT*PT*PTR.F |
| **Active BCR related Gene** | ABR | T30S33 | **15.3** | **49.0** | K.VLEDEDVFLLEECELGT*PTS*PGSGSPFLVAVK.V |
| **Active BCR related Gene** | ABR | S33 | **10.3** | **45.3** | K.VLEDEDVFLLEECELGTPTS*PGSGSPFLVAVK.V |
| **Activity dependent neuroprotector** | ADNP | S953S955 | **100.0** | **64.8** | K.LMHNAS*DS*EVDQDDVVEWK.D |
| **Activity dependent neuroprotector** | ADNP | S1058 | | **57.7** | R.LS*NPQIEWQNSTIDSEDGEQFDNMTDGVAEPMHGSLAGVK.L |
| **Activity dependent neuroprotector** | ADNP | S1071 | **7.4** | **69.1** | R.LSNPQIEWQNSTIDS*EDGEQFDNMTDGVAEPMHGSLAGVK.L |
| **Activity dependent neuroprotector** | ADNP | S1067 | **7.7** | **23.4** | R.LSNPQIEWQNS*TIDSEDGEQFDNMTDGVAEPMHGSLAGVK.L |
| **Activity dependent neuroprotector** | ADNP | T1068S1071 | **3.5** | **42.7** | R.LSNPQIEWQNST*IDS*EDGEQFDNMTDGVAEPMHGSLAGVK.L |
| **Activity dependent neuroprotector** | ADNP | S1058S1067 | | **14.8** | R.LS*NPQIEWQNS*TIDSEDGEQFDNMTDGVAEPMHGSLAGVK.L |
| **Activity dependent neuroprotector** | ADNP | T1068 | **4.2** | **61.0** | R.LSNPQIEWQNST*IDSEDGEQFDNMTDGVAEPMHGSLAGVK.L |
| **ADAM17** | ADAM17 | S791 | **58.9** | **35.5** | K.S*FEDLTDHPVTR.S |
| | AP3D1 | S632S634S636 | **100.0** | **27.0** | K.KVPVPEGLDLDAWINEPLS*DS*ES*EDERPR.A |
| **Adaptin, delta** | AP3D1 | S758S759 | | **17.2** | R. HS*S*LPTESPEDIAPAQQVDIVTEEMPENALPSDEDDKDPNDPYR.A |
| **Adducin 1 alpha** | ADD2 | S757 | **13.9** | **23.5** | K.FRTPS*FLKK.S |
| **Adducin 1 alpha** | ADD1 | S431 | **15.8** | **69.7** | K.YSDVEVPASVTGYSFASDGDSGTCS*PLR.H |
| **Adducin 1 alpha** | ADD1 | S358 | **39.1** | **91.8** | R.SPGS*PVGEGTGSPPK.W |
| **Adducin 1 alpha** | ADD1 | T364 | **18.1** | **15.5** | R.SPGSPVGEGT*GSPPKWQIGEQEFEALMR.M |
| **Adducin 1 alpha** | ADD1 | S358S366 | **14.9** | **82.6** | R.SPGS*PVGEGTGS*PPKWQIGEQEFEALMR.M |
| **Adducin 1 alpha** | ADD1 | S465 | **17.0** | **48.3** | R.GDEASEEGQNGSS*PK.S |
| **Adducin 1 alpha** | ADD1 | S423S436 | **-0.2** | **21.7** | K.YSDVEVPASVTGYSFAS*DGDSGTCSPLRHS*FQK.Q |
| **Adducin 1 alpha** | ADD1 | S355S366 | **12.0** | **58.6** | R.S*PGSPVGEGTGS*PPKWQIGEQEFEALMR.M |
| **Adducin 1 alpha** | ADD1 | S355S358 | **17.2** | **48.3** | R.S*PGS*PVGEGTGSPPKWQIGEQEFEALMR.M |
| **Adducin 1 alpha** | ADD1 | S355T364 | **11.8** | **14.2** | R.S*PGSPVGEGT*GSPPKWQIGEQEFEALMR.M |
| **Adducin 1 alpha** | ADD1 | S512 | **6.5** | **22.1** | K.EDGHRTS*TSAVPNLFVPLNTNPK.E |
| **Adducin 1 alpha** | ADD1 | S700T702T706 | | **14.5** | K. EEEAHRPPS*PT*EAPT*EAS*PEPAPDPAPVAEEAAPSAVEEGAAAD |
| **Adducin 1 alpha** | ADD1 | S436 | **2.0** | **16.6** | K.YSDVEVPASVTGYSFASDGDSGTCSPLRHS*FQK.Q |
| **Adducin 1 alpha** | ADD1 | T511 | **-6.1** | **23.6** | K.EDGHRT*STSAVPNLFVPLNTNPK.E |
| **Adducin 1 alpha** | ADD1 | S366 | **6.6** | **45.8** | R.SPGSPVGEGTGS*PPKWQIGEQEFEALMR.M |
| **Adducin 1 alpha** | ADD1 | T364S366 | **2.3** | **59.4** | R.SPGSPVGEGT*GS*PPKWQIGEQEFEALMR.M |
| **Adducin 3** | ADD3 | S673S677S681 | **19.2** | **21.3** | K.IEEVLS*PEGS*PSKS*PSK.K |
| **Adducin 3** | ADD3 | S402S423 | **30.4** | **56.8** | R.HKS*DVEIPATVTAFSFEDDTVPLS*PLK.Y |
| **Adducin 3** | ADD3 | S423 | **23.1** | **55.1** | K.SDVEIPATVTAFSFEDDTVPLS*PLK.Y |
| **Adducin 3** | ADD3 | S677S681 | **6.9** | **10.9** | K.IEEVLSPEGS*PSKS*PSK.K |
| **Adducin 3** | ADD3 | S461 | **21.2** | **26.1** | R.NGETS*PR.T |
| **Adducin 3** | ADD3 | S42 | **100.0** | **21.1** | R.NMS*PDLR.Q |
| **Adducin 3** | ADD3 | T460 | | **13.7** | R.NGET*SPR.T |
| **Adducin 3** | ADD3 | S600 | **3.5** | **46.6** | R.KQQGLEDAEQELLSDDASSVSQIQSQTQS*PQNVPEK.L |
| **Adducin 3** | ADD3 | S585 | | **33.0** | R.KQQGLEDAEQELLS*DDASSVSQIQSQTQSPQNVPEK.L |
| **Adenosine deaminase, RNA specific** | ADAR | S825 | **100.0** | **34.1** | R.S*PEAQPK.T |
| **Adenosine deaminase, RNA specific** | ADAR | S614 | **37.8** | **75.0** | K.S*PVTTLLECMHK.L |
| **Adenosine monophosphate deaminase 2** | AMPD2 | S190 | **-0.3** | **32.1** | K.TDS*DSDLQLYK.E |
| **Adenylyl cyclase-associated protein** | CAP1 | S308 | **16.4** | **54.9** | R.SGPKPFSAPKPQTS*PSPK.R |

Peak Area: <-10, -5, -3, 0, 3, 5, >10

%CV: 0, 29, 43, 71, 86, >100

**White dots-** Significant change in peptide abundance at 5 %FDR compared to the timepoint with the minimum peak area for a given PSM

| Protein Name | Gene | Phosphosites | Ascor | MOWSE | Sequence |
|---|---|---|---|---|---|
| Adenylyl cyclase-associated protein | CAP1 | S308S310 | 17.7 | 22.0 | R.SGPKPFSAPKPQTS*PS*PK.R |
| Adenylyl cyclase-associated protein | CAP1 | T307 | 10.4 | 32.7 | R.SGPKPFSAPKPQT*SPSPK.R |
| Adenylyl cyclase-associated protein | CAP1 | T307S310 | 14.1 | 22.8 | R.SGPKPFSAPKPQT*SPS*PK.R |
| Adenylyl cyclase-associated protein | CAP1 | S310 | 2.4 | 12.7 | R.SGPKPFSAPKPQTSPS*PK.R |
| ADP ribosylation factor GTPase activating protein 1 | ARFGAP1 | S361 | -0.4 | 53.1 | R.SS*DSWEVWGSASTNR.N |
| ADP ribosylation factor GTPase activating protein 1 | ARFGAP1 | S363 | 6.1 | 31.0 | R.SSDS*WEVWGSASTNR.N |
| ADP ribosylation factor guanine nucleotide exchange factor 2 | ARFGEF2 | S277 | 11.9 | 86.4 | R.GSS*LSGTDDGAQEVVK.D |
| ADP ribosylation factor guanine nucleotide exchange factor 2 | ARFGEF2 | S1513S1514 | 20.0 | 46.3 | K.HLDVDLDRQSLS*S*IDKNPSER.G |
| ADP ribosylation factor guanine nucleotide exchange factor 2 | ARFGEF2 | S1513S1514S1 | 4.8 | 26.2 | K.HLDVDLDRQSLS*S*IDKNPS*ER.G |
| ADP ribosylation factor guanine nucleotide exchange factor 2 | ARFGEF2 | T626T628 | 7.2 | 69.0 | R.RCSVTSMESTVSSGT*QT*TVQDDPEQFEVIK.Q |
| ADP ribosylation factor guanine nucleotide exchange factor 2 | ARFGEF2 | S276 | 26.0 | 64.4 | R.GS*SLSGTDDGAQEVVK.D |
| ADP ribosylation factor guanine nucleotide exchange factor 2 | ARFGEF2 | S1511S1513S15 | 1.9 | 21.5 | K.HLDVDLDRQS*LS*S*IDKNPSER.G |
| ADP ribosylation factor guanine nucleotide exchange factor 2 | ARFGEF2 | S614S617 | 14.3 | 68.9 | R.RCS*VTS*MESTVSSGTQTTVQDDPEQFEVIK.Q |
| ADP ribosylation factor guanine nucleotide exchange factor 2 | ARFGEF2 | S617T628 | 5.1 | 15.7 | R.RCSVTS*MESTVSSGTQT*TVQDDPEQFEVIK.Q |
| ADP ribosylation factor like 3 | ARL3 | S5 | 100.0 | 38.7 | M.GLLS*ILR.K |
| ADP-ribosylation factor interacting protein 1 (arfaptin 1) | ARFIP1 | S39 | 54.5 | 87.5 | K.HSLPS*GLGLSETQITSHGFDNTK.E |
| ADP-ribosylation factor interacting protein 1 (arfaptin 1) | ARFIP1 | S44 | 13.6 | 49.8 | K.HSLPSGLGLS*ETQITSHGFDNTK.E |
| ADP-ribosylation factor interacting protein 1 (arfaptin 1) | ARFIP1 | S100 | 20.7 | 28.0 | R.KWS*LNTYK.C |
| ADP-ribosyltransferase like 1 | PARP4 | S1335 | 55.8 | 55.5 | R.AHS*PASLSFASYR.Q |
| AF10 | MLLT10 | S705 | 55.9 | 32.9 | R.S*PVSSLQIR.Y |
| AF1Q | MLLT11 | S84 | 28.4 | 48.7 | R.APIASIHS*FELDLL.- |
| AF5Q31 | AFF4 | S703S706 | 42.5 | 26.1 | K.ELLS*PLS*EPDDRYPLIVK.I |
| AF5Q31 | AFF4 | S1062 | 9.8 | 65.5 | K.LS*PGNSGNYSSGASSASASGSSVTIPQK.I |
| AF6 | AFDN | S1067 | 30.2 | 75.3 | R.TSS*VVTLEVAK.Q |
| AF6 | AFDN | S1185 | 29.7 | 65.4 | K.ITSVS*TGNLCTEEQTPPPRPEAYPIPTQTYTR.E |
| AF6 | AFDN | S1259 | 100.0 | 41.6 | R.S*QEELREDK.A |
| AF6 | AFDN | S1183 | 8.0 | 41.7 | K.ITS*VSTGNLCTEEQTPPPRPEAYPIPTQTYTR.E |
| AF6 | AFDN | S1185T1195 | 12.0 | 14.8 | K.ITSVS*TGNLCTEEQT*PPPRPEAYPIPTQTYTR.E |
| Aftiphilin | AFTPH | S411 | 7.3 | 70.8 | R.KFTNFQSPNIDPTEENDLDDSLS*VK.N |
| Aftiphilin | AFTPH | S409 | 48.1 | 81.0 | R.KFTNFQSPNIDPTEENDLDDS*LSVK.N |
| Aggrecan 1 | ACAN | S2403S2404 | 100.0 | 15.4 | K.RS*S*RHPR.R |
| AKAP95 | AKAP8 | S339 | 114.9 | 67.4 | R.S*GDEEFKGEDELCDSGR.Q |
| AKAP95 | AKAP8 | S328 | 47.8 | 105.6 | R.VDSEGDFS*ENDDAAGDFR.S |
| AKAP95 | AKAP8 | S323S328 | 100.0 | 156.6 | R.VDS*EGDFS*ENDDAAGDFR.S |
| AKT1 | AKT1 | S124 | 20.6 | 80.3 | R.SGS*PSDNSGAEEMEVSLAKPK.H |
| AKT1 | AKT1 | S122S129 | 9.7 | 98.5 | R.S*GSPSDNS*GAEEMEVSLAKPK.H |
| AKT1 | AKT1 | S124S129 | 6.6 | 77.0 | R.SGS*PSDNS*GAEEMEVSLAKPK.H |
| AKT1 | AKT1 | S124S126S129 | 35.1 | 66.7 | R.SGS*PS*DNS*GAEEMEVSLAKPK.H |
| AKT1 | AKT2 | Y315 | 5.6 | 61.0 | K.TFCGTPEY*LAPEVLEDNDYGR.A |
| AKT1 | AKT1 | Y474S477 | 9.7 | 68.2 | R.RPHFPQFSY*SAS*GTA.- |
| AKT1 | AKT1 | S473T479 | 3.2 | 66.9 | R.RPHFPQFS*YSASGT*A.- |
| AKT1 | AKT1 | S473Y474 | 10.1 | 25.2 | R.RPHFPQFS*Y*SASGTA.- |
| AKT1 | AKT1 | S477T479 | 6.9 | 62.5 | R.RPHFPQFSYSAS*GT*A.- |
| AKT1 | AKT1 | S475T479 | 9.2 | 30.8 | R.RPHFPQFSYS*ASGT*A.- |
| AKT1 | AKT1 | Y474S475 | -0.3 | 20.9 | R.RPHFPQFSY*S*ASGTA.- |
| AKT1 | AKT2 | T308 | 5.4 | 64.2 | K.T*FCGTPEYLAPEVLEDNDYGR.A |
| AKT1 substrate 1 | AKT1S1 | T246 | 19.1 | 33.9 | R.LNT*SDFQK.L |
| AKT1 substrate 1 | AKT1S1 | S88S92 | 25.4 | 23.6 | R.AATAARPPAPPPAPQPPS*PTPS*PPRPTLAR.E |
| AKT1 substrate 1 | AKT1S1 | S183 | 42.4 | 54.2 | K.S*LPVSVPVWGFK.E |
| AKT1 substrate 1 | AKT1S1 | S203S212 | 4.7 | 24.4 | R.SS*DEENGPPSS*PDLDR.I |
| AKT1 substrate 1 | AKT1S1 | S202S203S212 | 18.7 | 18.2 | R.TEARS*S*DEENGPPSS*PDLDR.I |
| AKT1 substrate 1 | AKT1S1 | S88T97 | 12.9 | 10.8 | R.AATAARPPAPPPAPQPPS*PTPSPPRPT*LAR.E |
| Aladin | AAAS | S495 | 100.0 | 26.7 | R.FS*PVLGR.A |
| AlkB, alkylation repair homolog 5 | ALKBH5 | S64S69 | 28.0 | 18.6 | K.YQEDS*DPERS*DYEEQQLQK.E |
| AlkB, alkylation repair homolog 5 | ALKBH5 | S361 | 44.4 | 71.5 | R.RGS*FSSENYWR.K |
| AlkB, alkylation repair homolog 5 | ALKBH5 | S64Y71 | 24.7 | 25.7 | R.KYQEDS*DPERSDY*EEQQLQK.E |
| AlkB, alkylation repair homolog 5 | ALKBH5 | S363 | -0.2 | 36.9 | R.RGSFS*SENYWR.K |
| AlkB, alkylation repair homolog 5 | ALKBH5 | Y60S69 | 7.4 | 11.1 | R.KY*QEDSDPERS*DYEEQQLQK.E |
| Alpha- and gamma-adaptin-binding protein p34 | AAGAB | S310S311 | 100.0 | 57.1 | R.DEIEGLS*S*DEEH.- |
| Alpha-2-HS glycoprotein | AHSG | S138 | 37.0 | 60.8 | K.CDSSPDS*AEDVRK.V |
| Alsin | ALS2 | S483S492 | 55.7 | 70.3 | R.RLS*LPGLLSQVS*PR.L |
| Alsin | ALS2 | S1464 | -0.7 | 57.6 | R.SES*PEPGYVVTSSGLLLPVLLPR.L |

Peak Area: <-10, -5, -3, 0, 3, 5, >10

%CV: 0, 29, 42, 71, 86, >100

**White dots-** Significant change in peptide abundance at **5** %FDR compared to the timepoint with the minimum peak area for a given PSM

| Protein Name | Gene | Phosphosites | CarT (0m 2m 5m) | RajiB (0m 2m 5m) | Ascor | MOWSE | Sequence |
|---|---|---|---|---|---|---|---|
| **Alsin** | ALS2 | S1462 | | | **13.1** | **35.3** | R.S*ESPEPGYVVTSSGLLLPVLLPR.L |
| **Alsin** | ALS2 | Y1469 | | | **0.2** | **28.0** | R.SESPEPGY*VVTSSGLLLPVLLPR.L |
| **Amida** | TFPT | S252 | | | **20.6** | **48.1** | K.LLPYPTLASPAS*D.- |
| **Amida** | TFPT | S249S252 | | | **31.0** | **44.2** | K.LLPYPTLAS*PAS*D.- |
| **Amida** | TFPT | T246S252 | | | **8.3** | **41.5** | K.LLPYPT*LASPAS*D.- |
| **Amida** | TFPT | Y244S252 | | | **8.9** | **47.2** | K.LLPY*PTLASPAS*D.- |
| **Amida** | TFPT | Y244T246 | | | **5.7** | **26.0** | K.LLPY*PT*LASPASD.- |
| **AMP deaminase 3** | AMPD3 | S112T130 | | | **9.8** | **12.4** | K.GPPAAS*PAMSPTTPVVTGATSLPT*PAPYAMPEFQR.V |
| **AMP deaminase 3** | AMPD3 | S112T119 | | | **1.2** | **44.8** | K.GPPAAS*PAMSPTT*PVVTGATSLPTPAPYAMPEFQR.V |
| **AMP deaminase 3** | AMPD3 | S116T118 | | | **6.1** | **40.6** | K.GPPAASPAMS*PT*TPVVTGATSLPTPAPYAMPEFQR.V |
| **AMP deaminase 3** | AMPD3 | S116T123 | | | **4.1** | **31.5** | K.GPPAASPAMS*PTTPVVT*GATSLPTPAPYAMPEFQR.V |
| **AMP deaminase 3** | AMPD3 | S116S127 | | | **-1.7** | **26.0** | K.GPPAASPAMS*PTTPVVTGATS*LPTPAPYAMPEFQR.V |
| **AMP deaminase 3** | AMPD3 | S112T118 | | | **4.9** | **40.7** | K.GPPAAS*PAMSPT*TPVVTGATSLPTPAPYAMPEFQR.V |
| **AMP deaminase 3** | AMPD3 | S112T123 | | | **6.2** | **38.4** | K.GPPAAS*PAMSPTTPVVT*GATSLPTPAPYAMPEFQR.V |
| **AMP deaminase 3** | AMPD3 | S112S116 | | | **0.5** | **32.8** | K.GPPAAS*PAMS*PTTPVVTGATSLPTPAPYAMPEFQR.V |
| **AMP deaminase 3** | AMPD3 | T119S127 | | | **-1.7** | **36.2** | K.GPPAASPAMSPTT*PVVTGATS*LPTPAPYAMPEFQR.V |
| **AMP deaminase 3** | AMPD3 | S112S127 | | | **1.9** | **23.6** | K.GPPAAS*PAM#SPTTPVVTGATS*LPTPAPYAMPEFQR.V |
| **AMP deaminase 3** | AMPD3 | T118T123 | | | **4.5** | **25.9** | K.GPPAASPAMSPT*TPVVT*GATSLPTPAPYAMPEFQR.V |
| **AMP deaminase 3** | AMPD3 | S116T119 | | | **12.2** | **38.2** | K.GPPAASPAMS*PTT*PVVTGATSLPTPAPYAMPEFQR.V |
| **Amphiphysin II** | BIN1 | S302S305 | | | **6.2** | **74.2** | K.SPS*QSS*LPAVVVETFPATVNGTVEGGSGAGR.L |
| **Amphiphysin II** | BIN1 | S300S305 | | | **12.6** | **70.3** | K.S*PSQSS*LPAVVVETFPATVNGTVEGGSGAGR.L |
| **Amphiphysin II** | BIN1 | S302S304 | | | **4.3** | **55.0** | K.SPS*QS*SLPAVVVETFPATVNGTVEGGSGAGR.L |
| **Amphiphysin II** | BIN1 | S304S305 | | | **5.4** | **50.4** | K.SPSQS*S*LPAVVVETFPATVNGTVEGGSGAGR.L |
| **Amphiphysin II** | BIN1 | S302 | | | **1.8** | **35.7** | K.SPS*QSSLPAVVVETFPATVNGTVEGGSGAGR.L |
| **Amphiphysin II** | BIN1 | S300S304 | | | **4.8** | **28.2** | K.S*PSQS*SLPAVVVETFPATVNGTVEGGSGAGR.L |
| **AMPK alpha 1** | PRKAA1 | T355 | | | **26.0** | **29.5** | K.DFYLAT*SPPDSFLDDHHLTRPHPER.V |
| **AMPK alpha 1** | PRKAA1 | S508 | | | **10.2** | **27.4** | R.SDS*DAEAQGK.S |
| **AMPK beta 2** | PRKAB2 | S183 | | | **12.6** | **61.5** | R.DLSS*SPPGPYGQEMYAFR.S |
| **AMPK beta1** | PRKAB1 | S108 | | | **137.2** | **42.4** | R.S*HNNFVAILDLPEGEHQYK.F |
| **Amyotrophic lateral sclerosis 2 chromosome region, candidate 13** | FAM117B | S170S173 | | | **8.3** | **86.7** | R.SQSVS*PTS*FLTISNEGSEESPCSADDLLVDPR.D |
| **Amyotrophic lateral sclerosis 2 chromosome region, candidate 13** | FAM117B | S170T176 | | | **6.7** | **50.0** | R.SQSVS*PTSFLT*ISNEGSEESPCSADDLLVDPR.D |
| **Anaphase promoting complex subunit 5** | ANAPC5 | S195 | | | **100.0** | **49.2** | K.EELDVS*VR.E |
| **Anaphase promoting complex, subunit 1** | ANAPC1 | S688 | | | **24.1** | **58.1** | R.NFDFEGSLS*PVIAPK.K |
| **Anaphase promoting complex, subunit 1** | ANAPC1 | S547S564 | | | **32.9** | **26.2** | K.LLGS*LDEVVLLSPVPELRDSS*K.L |
| **Anaphase promoting complex, subunit 1** | ANAPC1 | S51S60 | | | **25.4** | **63.5** | R.QLQPASELWSS*DGAAGLVGS*LQEVTIHEK.Q |
| **Anaphase promoting complex, subunit 1** | ANAPC1 | S50S60 | | | **25.9** | **47.5** | R.QLQPASELWS*SDGAAGLVGS*LQEVTIHEK.Q |
| **Anaphase promoting complex, subunit 1** | ANAPC1 | S547S563 | | | **42.3** | **31.7** | K.LLGS*LDEVVLLSPVPELRDS*SK.L |
| **Anaphase promoting complex, subunit 1** | ANAPC1 | S547S555 | | | **100.0** | **40.3** | K.LLGS*LDEVVLLS*PVPELR.D |
| **Anaphase promoting complex, subunit 3** | CDC27 | S364 | | | **52.0** | **78.0** | R.EVTPILAQTQSSGPQTSTTPQVLS*PTITSPPNALPR.R |
| **Anaphase promoting complex, subunit 3** | CDC27 | T343 | | | | **44.1** | R.EVT*PILAQTQSSGPQTSTTPQVLSPTITSPPNALPR.R |
| **Anaphase promoting complex, subunit 4** | ANAPC4 | S777 | | | | **34.6** | K.IKEEVLS*ESEAENQQAGAAALAPEIVIK.V |
| **Anaphase promoting complex, subunit 8** | CDC23 | S588 | | | **23.4** | **19.8** | R.RVS*PLNLSSVTP.- |
| **Androgen induced proliferation inhibitor** | PDS5B | S1358 | | | **45.3** | **105.4** | R.AES*PESSAIESTQSTPQK.G |
| **Androgen induced proliferation inhibitor** | PDS5B | S1358T1370 | | | **31.2** | **70.7** | R.AES*PESSAIESTQST*PQK.G |
| | PDS5B | S1283 | | | **100.0** | **51.4** | R.LKEDILENEDEQNS*PPKK.G |
| **Androgen induced proliferation inhibitor** | PDS5B | S1182 | | | **11.5** | **24.3** | R.LDSSEMDHS*ENEDYTMSSPLPGKK.S |
| **Androgen induced proliferation inhibitor** | PDS5B | S1176 | | | **-1.8** | **18.3** | R.LDS*SEMDHSENEDYTMSSPLPGK.K |
| **Androgen induced proliferation inhibitor** | PDS5B | T1381 | | | **20.3** | **21.5** | K.T*PSPSQPK.K |
| **Androgen induced proliferation inhibitor** | PDS5B | S1166 | | | **22.5** | **84.9** | R.METVSNASSSSNPSS*PGR.I |
| **Androgen induced proliferation inhibitor** | PDS5B | S1358S1369 | | | **24.2** | **19.5** | R.AES*PESSAIESTQS*TPQK.G |
| **Androgen induced proliferation inhibitor** | PDS5B | S1165 | | | **15.8** | **44.5** | R.METVSNASSSSNPS*SPGR.I |
| **Androgen induced proliferation inhibitor** | PDS5B | S1176S1177 | | | | **43.2** | R.LDS*S*EMDHSENEDYTMSSPLPGKK.S |
| **Androgen induced proliferation inhibitor** | PDS5B | S1383 | | | **39.8** | **33.2** | K.TPS*PSQPK.K |
| **Anillin actin binding protein** | ANLN | S97S99 | | | **9.3** | **30.7** | K.SCS*PS*PVSPQVQPQAADTISDSVAVPASLLGMR.R |
| **Anillin actin binding protein** | ANLN | S95S97 | | | **8.5** | **35.6** | K.S*CS*PSPVSPQVQPQAADTISDSVAVPASLLGMR.R |
| **Anillin actin binding protein** | ANLN | S97S102 | | | **10.8** | **20.9** | K.SCS*PSPVS*PQVQPQAADTISDSVAVPASLLGMR.R |
| **Anion exchange protein (Fragment)** | SLC4A7 | S258 | | | **13.9** | **41.2** | R.NGILASPQS*APGNLDNSK.S |
| **Anion exchange protein (Fragment)** | SLC4A7 | S743 | | | **33.2** | **33.8** | K.YSVDPS*IVNISDEMAK.T |
| **Ankyrin 2** | ANK2 | T3844 | | | **6.8** | **51.8** | R.KT*SLVIVESADNQPETCER.L |
| **Ankyrin 2** | ANK2 | S3845 | | | **8.0** | **18.5** | R.KTS*LVIVESADNQPETCER.L |
| **Ankyrin repeat and BTB/POZ domain-containing protein 2** | ABTB2 | S67 | | | **14.6** | **68.5** | R.HNS*WDTVNTVLPEDPEVADLFSR.C |
| **Ankyrin repeat and BTB/POZ domain-containing protein 2** | ABTB2 | S29 | | | **14.0** | **24.7** | R.S*LSLSSSK.S |

Peak Area: <-10, -5, -3, 0, 3, 5, >10

%CV: 0, 29, 42, 71, 86, >100

**White dots-** Significant change in peptide abundance at **5** %FDR compared to the timepoint with the minimum peak area for a given PSM

| Protein Name | Gene | Phosphosites | CarT 0m 2m 5m | RajiB 0m 2m 5m | Ascor | MOWSE | Sequence |
|---|---|---|---|---|---|---|---|
| Ankyrin repeat and KH domain containing 1 | ANKHD1 | S95 | | | 12.1 | 90.8 | R.TGGGGGASGS*DEDEVSEVESFILDQEDLDNPVLK.T |
| Ankyrin repeat and KH domain containing 1 | ANKHD1 | T86 | | | -1.2 | 13.2 | R.T*GGGGGASGSDEDEVSEVESFILDQEDLDNPVLK.T |
| Ankyrin repeat and KH domain containing 1 | ANKHD1 | S93 | | | 6.0 | 51.3 | R.TGGGGGAS*GSDEDEVSEVESFILDQEDLDNPVLK.T |
| Ankyrin repeat domain 11 | ANKRD11 | S1792 | | | 18.9 | 13.0 | R.SVS*VDIR.R |
| Ankyrin repeat domain 28 | ANKRD28 | S1048 | | | 5.2 | 43.7 | R.NEPSSYCSFNNIGGEQEYLYTDVDELNDS*DSETY.- |
| Ankyrin repeat domain 28 | ANKRD28 | T1040 | | | 0.7 | 27.1 | R.NEPSSYCSFNNIGGEQEYLYT*DVDELNDSDSETY.- |
| Ankyrin repeat domain 28 | ANKRD28 | Y1037 | | | 12.6 | 38.4 | R.NEPSSYCSFNNIGGEQEY*LYTDVDELNDSDSETY.- |
| Ankyrin repeat domain protein 17 | ANKRD17 | T5 | | | | 63.9 | K.AT*VPVAAATAAEGEGSPPAVAAVAGPPAAAEVGGGVGGSSR.A |
| Ankyrin repeat domain protein 17 | ANKRD17 | S19 | | | 9.9 | 35.8 | K.ATVPVAAATAAEGEGS*PPAVAAVAGPPAAAEVGGGVGGSSR.A |
| Ankyrin repeat domain protein 17 | ANKRD17 | S2041S2047 | | | 5.9 | 18.2 | K.EHYPVS*SPSSPS*PPAQPGGVSR.N |
| Anti silencing function 1B | ASF1B | S198 | | | 100.0 | 24.0 | K.GLGLPGCIPGLLPENS*MDCI.- |
| Anti-silencing function 1A | ASF1A | S166 | | | 25.9 | 63.3 | K.LEDAESS*NPNLQSLLSTDALPSASK.G |
| Anti-silencing function 1A | ASF1A | S165 | | | 48.3 | 57.1 | K.LEDAES*SNPNLQSLLSTDALPSASK.G |
| Antigen identified by monoclonal antibody Ki-67 | MKI67 | S2002 | | | 100.0 | 25.0 | R.LKIS*LGK.V |
| Antigen identified by monoclonal antibody Ki-67 | MKI67 | S584 | | | 29.9 | 84.7 | K.AQSLVISPPAPS*PR.K |
| Antigen identified by monoclonal antibody Ki-67 | MKI67 | S357 | | | 48.5 | 56.9 | K.TPVQYSQQQNS*PQK.H |
| Antigen identified by monoclonal antibody Ki-67 | MKI67 | S352 | | | 5.1 | 25.7 | K.TPVQYS*QQQNSPQK.H |
| Antigen identified by monoclonal antibody Ki-67 | MKI67 | S1131 | | | 20.0 | 30.3 | K.S*PPPESVDTPTSTK.Q |
| Antigen identified by monoclonal antibody Ki-67 | MKI67 | S2223 | | | 37.2 | 38.9 | R.S*PQPDPVGTPTIFKPQSK.R |
| Antisense ERCC1 | CD3EAP | S126S136 | | | 42.7 | 42.6 | R.ILEGPQQSLS*GSPLQPIPAS*PPPQIPPGLRPR.F |
| Antisense ERCC1 | CD3EAP | S124S136 | | | 26.1 | 16.9 | R.ILEGPQQS*LSGSPLQPIPAS*PPPQIPPGLRPR.F |
| Antisense ERCC1 | CD3EAP | S128S136 | | | 34.5 | 52.1 | R.ILEGPQQSLSGS*PLQPIPAS*PPPQIPPGLRPR.F |
| Antisense ERCC1 | CD3EAP | S124S126 | | | | 35.5 | R.ILEGPQQS*LS*GSPLQPIPASPPPQIPPGLRPR.F |
| AP2 associated kinase 1 | AAK1 | T620S624 | | | 12.5 | 25.9 | K.VGSLT*PPSS*PK.T |
| AP2 associated kinase 1 | AAK1 | S623S624 | | | 12.2 | 24.6 | K.VGSLTPPS*S*PK.T |
| AP2 associated kinase 1 | AAK1 | T640 | | | 13.8 | 48.5 | R.ILSDVT*HSAVFGVPASK.S |
| AP2 associated kinase 1 | AAK1 | T653 | | | -0.3 | 83.5 | K.ST*QLLQAAAAEASLNK.S |
| AP2 associated kinase 1 | AAK1 | S637 | | | 61.2 | 65.4 | R.ILS*DVTHSAVFGVPASK.S |
| AP47 | AP1M1 | T154 | | | 16.2 | 37.3 | K.LETGAPRPPATVT*NAVSWR.S |
| AP47 | AP1M1 | T152T154 | | | 34.9 | 29.3 | K.LETGAPRPPAT*VT*NAVSWR.S |
| APBB2 | APBB2 | S123 | | | 53.8 | 21.5 | K.NLS*PTAVINITSEK.L |
| APC | APC | S2837 | | | | 16.8 | K.RHSGS*YLVTSV.- |
| APC | APC | S780 | | | 36.4 | 41.0 | K.ALEAELDAQHLSETFDNIDNLS*PK.A |
| APG4 autophagy 4 homolog B | ATG4B | S383 | | | 60.5 | 48.6 | R.FFDS*EDEDFEILSL.- |
| Apoptosis antagonizing transcription factor | AATF | S203 | | | 107.9 | 48.8 | R.AGDRNS*EDDGVVMTFSSVK.V |
| Apoptosis antagonizing transcription factor | AATF | S320S321 | | | 10.1 | 31.4 | R.YLVDGTKPNAGSEEIS*S*EDDELVEEK.K |
| Apoptosis antagonizing transcription factor | AATF | T310S316S321 | | | 12.0 | 23.7 | R.YLVDGT*KPNAGS*EEISS*EDDELVEEK.K |
| Apoptosis antagonizing transcription factor | AATF | S316S320S321 | | | 28.0 | 66.1 | R.YLVDGTKPNAGS*EEIS*S*EDDELVEEK.K |
| Apoptosis antagonizing transcription factor | AATF | T310S316S320 | | | 9.7 | 17.1 | R.YLVDGT*KPNAGS*EEIS*SEDDELVEEK.K |
| Apoptosis antagonizing transcription factor | AATF | T310S320S321 | | | 3.9 | 22.9 | R.YLVDGT*KPNAGSEEIS*S*EDDELVEEK.K |
| Apoptosis antagonizing transcription factor | AATF | Y305T310S316 | | | 9.0 | 34.5 | R.Y*LVDGT*KPNAGS*EEISSEDDELVEEK.K |
| Apoptosis antagonizing transcription factor | AATF | S169S170S178 | | | 100.0 | 49.0 | K.GMDDLGS*S*EEEEDEES*GMEEGDDAEDS*QGES*EEDR.A |
| Apoptosis antagonizing transcription factor | AATF | T310S321 | | | 20.0 | 11.6 | R.YLVDGT*KPNAGSEEISS*EDDELVEEK.K |
| Apoptosis inhibitor 5 | API5 | S462 | | | 13.9 | 69.4 | R.ASEDTTS*GSPPKK.S |
| Apoptosis inhibitor 5 | API5 | S464 | | | 7.8 | 46.1 | R.ASEDTTSGS*PPKK.S |
| Apoptotic chromatin condensation inducer in the nucleus (Fragment) | ACIN1 | S125 | | | 3.4 | 30.2 | K.EAVVDLHADDS*RISEDETER.N |
| Apoptotic chromatin condensation inducer in the nucleus (Fragment) | ACIN1 | S128 | | | 9.2 | 24.2 | K.EAVVDLHADDSRIS*EDETER.N |
| Aprataxin | APTX | S132 | | | 22.2 | 42.2 | R.S*GNSDSIER.D |
| | ANP32B | T244 | | | 100.0 | 28.3 | R.ET*DDEGEDD.- |
| Arachidonate 12-oxidoreductase | ALOX12 | S246 | | | 10.2 | 27.2 | R.RSTS*LPSR.L |
| Arachidonate 12-oxidoreductase | ALOX12 | S244 | | | 15.4 | 13.3 | R.RS*TSLPSR.L |
| Arf-GAP with dual PH domain-containing protein 1 (Fragment) | ADAP1 | S71 | | | 100.0 | 14.7 | K.IAPS*ER.K |
| Arginine/serine-rich coiled-coil 2 | RSRC2 | S216S218T220 | | | 100.0 | 17.4 | R.S*LS*RT*PS*PPPFR.G |
| Arginine/serine-rich coiled-coil 2 | RSRC2 | S32 | | | 10.9 | 43.7 | K.EQSEVSVS*PR.A |
| Arginine/serine-rich coiled-coil 2 | RSRC2 | S41 | | | 6.5 | 11.7 | K.HHYS*R.S |
| Arginine/serine-rich coiled-coil 2 | RSRC2 | S30S32 | | | 10.5 | 14.7 | K.EQSEVS*VS*PR.A |
| Arginine/serine-rich coiled-coil 2 | RSRC2 | Y40 | | | | 17.0 | K.HHY*SR.S |
| Arginine/serine-rich splicing factor 10 | TRA2B | S39 | | | 26.3 | 27.8 | R.S*KEDSRR.S |
| Arginine/serine-rich splicing factor 10 | TRA2B | T201 | | | 32.0 | 33.5 | K.RPHT*PTPGIYMGRPTYGSSR.R |
| Arginine/serine-rich splicing factor 10 | TRA2B | T201Y213 | | | 17.1 | 23.8 | K.RPHT*PTPGIYMGRPTY*GSSR.R |
| Arginine/serine-rich splicing factor 10 | TRA2B | T201T212 | | | 8.5 | 22.2 | K.RPHT*PTPGIYMGRPT*YGSSR.R |
| Arginine/serine-rich splicing factor 10 | TRA2B | S29T33 | | | 100.0 | 17.6 | K.S*ARHT*PAR.S |

Peak Area: <-10, -5, -3, 0, 3, 5, >10

%CV: 0, 29, 42, 71, 86, >100

**White dots-** Significant change in peptide abundance at **5** %FDR compared to the timepoint with the minimum peak area for a given PSM

CarT: 0m 2m 5m — RajiB: 0m 2m 5m

| Protein Name | Gene | Phosphosites | Ascor | MOWSE | Sequence |
|---|---|---|---|---|---|
| **Arginine/serine-rich splicing factor 10** | TRA2B | T69 | **13.9** | **34.7** | R.RHYT*R.S |
| **Arginine/serine-rich splicing factor 10** | TRA2B | S95S97S99 | **33.8** | **34.5** | R.RHS*HS*HS*PMSTR.R |
| **Arginine/serine-rich splicing factor 10** | TRA2B | S95S97 | **14.9** | **38.8** | R.RHS*HS*HSPMSTR.R |
| **Arginine/serine-rich splicing factor 10** | TRA2B | S95S99 | **13.7** | **21.8** | R.RHS*HSHS*PMSTR.R |
| **Arginine/serine-rich splicing factor 10** | TRA2B | S97S99 | **7.3** | **25.6** | R.RHSHS*HS*PMSTR.R |
| **Arginine/serine-rich splicing factor 10** | TRA2B | T201Y207 | **16.1** | **25.4** | K.RPHT*PTPGIY*MGRPTYGSSR.R |
| **Arginine/serine-rich splicing factor 10** | TRA2B | T201S215 | **11.8** | **19.8** | K.RPHT*PTPGIYMGRPTYGS*SR.R |
| **Arginine/serine-rich splicing factor 10** | TRA2B | S97S102T103 | **15.3** | **18.6** | R.RHSHS*HSPMS*T*RR.R |
| **Arginine/serine-rich splicing factor 10** | TRA2B | T203Y207 | **19.0** | **25.8** | K.RPHTPT*PGIY*MGRPTYGSSR.R |
| **Arginine/serine-rich splicing factor 10** | TRA2B | T201T203 | **10.4** | **23.5** | K.RPHT*PT*PGIYMGRPTYGSSR.R |
| **Arginine/serine-rich splicing factor 10** | TRA2B | S43 | **-0.2** | **19.7** | R.SKEDS*RR.S |
| **Arginine/serine-rich splicing factor 10** | TRA2B | T203T212 | **3.9** | **17.4** | K.RPHTPT*PGIYMGRPT*YGSSR.R |
| **Arginine/serine-rich splicing factor 10** | TRA2B | S102T103 | **21.5** | **17.3** | R.RHSHSHSPMS*T*R.R |
| **Arginine/serine-rich splicing factor 10** | TRA2B | S97S99S102 | **3.3** | **12.6** | R.RRHSHS*HS*PMS*TR.R |
| **Arginine/serine-rich splicing factor 10** | TRA2B | S39S43 | **100.0** | **15.1** | R.S*KEDS*RR.S |
| **Arginine/serine-rich splicing factor 10** | TRA2B | Y207 | **8.9** | **11.9** | K.RPHTPTPGIY*MGRPTYGSSR.R |
| **Arginine/serine-rich splicing factor 10** | TRA2B | Y68 | | **24.6** | R.RHY*TR.S |
| **ARID1B** | ARID1B | S1550 | **32.1** | **52.4** | R.MS*PSKSPFLPSMK.M |
| **ARL6IP4** | ARL6IP4 | S252 | **90.9** | **108.6** | R.S*AGEEEDGPVLTDEQK.S |
| **Armadillo repeat containing 1** | ARMC1 | S120 | **24.9** | **61.9** | K.LLASEIYDILQSSNMADGDS*FNEMNSR.R |
| **ARPP-21** | ARPP21 | S383 | **15.1** | **76.2** | K.TAS*FGGITVLTR.G |
| **ARPP-21** | ARPP21 | S138 | **35.2** | **46.6** | K.DCS*QEYTDSTGIDLHEFLINTLK.N |
| **ARPP19** | ENSA | S62 | **35.2** | **66.6** | K.YFDS*GDYNMAK.A |
| **Arsenite resistance protein 2** | SRRT | S67 | **100.0** | **23.0** | R.ERFS*PPR.H |
| **Arsenite resistance protein 2** | SRRT | T544 | **26.7** | **58.1** | R.TQLWASEPGT*PPLPTSLPSQNPILK.N |
| **Arsenite resistance protein 2** | SRRT | S74 | **100.0** | **29.1** | R.HELS*PPQKR.M |
| **ASH1L** | ASH1L | S1162S1170 | **25.4** | **23.5** | R.RLS*PPTLLPNS*PSHLSELTSLK.E |
| **ASH2 like** | ASH2L | S623 | **100.0** | **28.0** | R.S*PPWEP.- |
| **ASK1** | MAP3K5 | S1029S1033 | **12.1** | **14.7** | R.TLFLGIPDENFEDHS*APPS*PEEK.D |
| **AT hook DNA binding motif containing 1** | AHDC1 | S846S849 | **29.5** | **53.6** | R.SLLDS*DDS*SDLLDFALSASRPESR.K |
| **AT hook DNA binding motif containing 1** | AHDC1 | S842S849 | **13.3** | **39.4** | R.S*LLDSDDS*SDLLDFALSASRPESR.K |
| **AT hook DNA binding motif containing 1** | AHDC1 | S842S846 | **5.0** | **15.4** | R.S*LLDS*DDSSDLLDFALSASRPESR.K |
| **AT hook transcription factor** | AKNA | S1170S1173 | **13.1** | **60.5** | R.LS*LSS*ESELPSLPLFSEK.S |
| **AT hook transcription factor** | AKNA | S534S537 | **9.7** | **44.4** | R.GDLS*PSS*LTSMPTLGWLPENR.D |
| **AT hook transcription factor** | AKNA | S534T539 | **2.2** | **23.3** | R.GDLS*PSSLT*SMPTLGWLPENR.D |
| **AT hook transcription factor** | AKNA | S1172S1173 | **16.5** | **79.9** | R.LSLS*S*ESELPSLPLFSEK.S |
| **AT hook transcription factor** | AKNA | T539 | **6.0** | **29.2** | R.GDLSPSSLT*SMPTLGWLPENR.D |
| **AT hook transcription factor** | AKNA | S1170S1175 | **3.5** | **19.7** | R.LS*LSSES*ELPSLPLFSEK.S |
| **AT hook transcription factor** | AKNA | S534 | **11.0** | **86.6** | R.GDLS*PSSLTSMPTLGWLPENR.D |
| **AT hook transcription factor** | AKNA | S540 | **4.8** | **38.7** | R.GDLSPSSLTS*MPTLGWLPENR.D |
| **AT hook transcription factor** | AKNA | S534S536 | **1.4** | **21.6** | R.GDLS*PS*SLTSMPTLGWLPENR.D |
| **AT hook transcription factor** | AKNA | S536 | **-0.2** | **42.5** | R.GDLSPS*SLTSMPTLGWLPENR.D |
| **AT rich interactive domain 1A** | ARID1A | S1600 | **16.7** | **52.4** | R.TS*PSKSPFLHSGMK.M |
| **AT rich interactive domain 1A** | ARID1A | S696 | **53.8** | **80.1** | R.GPS*PSPVGSPASVAQSR.S |
| **AT rich interactive domain 1A** | ARID1A | S1754 | | **78.3** | K.VS*SPAPMEGGEEEEELLGPK.L |
| **AT rich interactive domain 1A** | ARID1A | S1600S1602 | **12.8** | **24.1** | R.TS*PS*KSPFLHSGMK.M |
| **AT rich interactive domain 1A** | ARID1A | S363 | **31.6** | **74.8** | R.SHHAPMS*PGSSGGGGQPLAR.T |
| **AT rich interactive domain 1A** | ARID1A | S769 | **15.6** | **35.7** | R.NPQMPQYSSPQPGS*ALSPR.Q |
| **AT rich interactive domain 1A** | ARID1A | S772 | **3.5** | **25.9** | R.NPQMPQYSSPQPGSALS*PR.Q |
| **AT rich interactive domain 3A** | ARID3A | S77S81S88 | **100.0** | **58.3** | R.AAAAGLGHPAS*PGGS*EDGPPGS*EEEDAAR.E |
| **ATAD2** | ATAD2 | S327 | **45.3** | **61.5** | R.KPNIFYSGPAS*PARPR.Y |
| **ATAD2** | ATAD2 | Y750 | **4.3** | **70.8** | K.TLDSDISCPLLESDLAY*SDDDVPSVYENGLSQK.S |
| **ATAD2** | ATAD2 | S746Y750 | **51.4** | **47.3** | K.TLDSDISCPLLES*DLAY*SDDDVPSVYENGLSQK.S |
| **ATAD2** | ATAD2 | S746S757 | **7.3** | **51.5** | K.TLDSDISCPLLES*DLAYSDDDVPS*VYENGLSQK.S |
| **ATAD2** | ATAD2 | Y750S751 | **29.7** | **52.7** | K.TLDSDISCPLLESDLAY*S*DDDVPSVYENGLSQK.S |
| **ATAD2** | ATAD2 | Y750S757 | **8.2** | **33.4** | K.TLDSDISCPLLESDLAY*SDDDVPS*VYENGLSQK.S |
| **ATAD2** | ATAD2 | S757 | **13.8** | **45.8** | K.TLDSDISCPLLESDLAYSDDDVPS*VYENGLSQK.S |
| **ATAD2** | ATAD2 | S746 | **1.2** | **40.6** | K.TLDSDISCPLLES*DLAYSDDDVPSVYENGLSQK.S |
| **ATAD2** | ATAD2 | S751 | **4.6** | **40.5** | K.TLDSDISCPLLESDLAYS*DDDVPSVYENGLSQK.S |
| **Ataxin 10** | ATXN10 | S12 | **100.0** | **47.1** | R.LS*GVMVPAPIQDLEALR.A |
| **Ataxin 2** | ATXN2 | S848S850S865 | **5.8** | **14.3** | K.DSFIENS*SS*NCTSGSSKPNSPSIS*PSILSNTEHK.R |
| **Ataxin 2** | ATXN2 | S667 | **-0.3** | **82.2** | R.TS*PSGGTWSSVVSGVPR.L |

Peak Area: <-10, -5, -3, 0, 3, 5, >10

%CV: 0, 29, 42, 71, 86, >100

**White dots-** Significant change in peptide abundance at **5** %FDR compared to the timepoint with the minimum peak area for a given PSM

| Protein Name | Gene | Phosphosites | CarT 0m 2m 5m | RajiB 0m 2m 5m | Ascor | MOWSE | Sequence |
|---|---|---|---|---|---|---|---|
| **Ataxin 2** | ATXN2 | S667S684 | | | **16.2** | **28.1** | R.TS*PSGGTWSSVVSGVPRLS*PK.T |
| **Ataxin 2** | ATXN2 | S697 | | | | **36.0** | R.QNS*IGNTPSGPVLASPQAGIIPTEAVAMPIPAASPTPASPASNR.A |
| **Ataxin 2** | ATXN2 | S850S861S865 | | | **11.9** | **31.6** | K.DSFIENSSS*NCTSGSSKPNS*PSIS*PSILSNTEHK.R |
| **Ataxin 2 domain protein** | ATXN2L | S238 | | | **26.4** | **47.5** | R.WEGGDSNS*DDYDLESDMSNGWDPNEMFK.F |
| **Ataxin 2 domain protein** | ATXN2L | S236S238 | | | **24.7** | **64.4** | R.WEGGDS*NS*DDYDLESDMSNGWDPNEMFK.F |
| **Ataxin 2 domain protein** | ATXN2L | Y241S245 | | | **11.9** | **45.0** | R.WEGGDSNSDDY*DLES*DMSNGWDPNEMFK.F |
| **Ataxin 2 domain protein** | ATXN2L | S684 | | | **12.2** | **32.3** | K.STSTPTS*PGPR.T |
| **Ataxin 2 domain protein** | ATXN2L | S236Y241 | | | **7.5** | **36.2** | R.WEGGDS*NSDDY*DLESDMSNGWDPNEMFK.F |
| **Ataxin 2 domain protein** | ATXN2L | S594 | | | **5.1** | **15.6** | K.EVDGLLTSEPMGS*PVSSK.T |
| **Ataxin 2 domain protein** | ATXN2L | S452 | | | | **58.5** | K. [illegible] |
| **Ataxin 2 domain protein** | ATXN2L | S236 | | | **11.8** | **38.6** | R.WEGGDS*NSDDYDLESDMSNGWDPNEMFK.F |
| **Ataxin 2 domain protein** | ATXN2L | Y241S248 | | | **10.6** | **30.4** | R.WEGGDSNSDDY*DLESDMS*NGWDPNEMFK.F |
| **Ataxin 2 domain protein** | ATXN2L | S111 | | | **66.2** | **31.3** | K.GPPQS*PVFEGVYNNSR.M |
| **Ataxin 2 domain protein** | ATXN2L | S236S245 | | | **7.0** | **32.7** | R.WEGGDS*NSDDYDLES*DMSNGWDPNEMFK.F |
| **Ataxin 2 domain protein** | ATXN2L | T31 | | | | **19.6** | R. [illegible] |
| **Ataxin 2 domain protein** | ATXN2L | T31S32 | | | | **31.6** | R. [illegible] |
| **Ataxin 2 domain protein** | ATXN2L | S452S459 | | | | **32.5** | K. [illegible] |
| **Ataxin 2 domain protein** | ATXN2L | S236S248 | | | **7.6** | **37.5** | R.WEGGDS*NSDDYDLESDMS*NGWDPNEMFK.F |
| **ATH1, acid trehalase-like 1** | PGGHG | S552 | | | **10.7** | **43.2** | R.DPLQSPLWVTLGSSSPTES*LTVDPASE.- |
| **ATH1, acid trehalase-like 1** | PGGHG | T550 | | | **2.8** | **41.4** | R.DPLQSPLWVTLGSSSPT*ESLTVDPASE.- |
| **ATH1, acid trehalase-like 1** | PGGHG | S547 | | | **5.1** | **41.5** | R.DPLQSPLWVTLGSS*SPTESLTVDPASE.- |
| **ATH1, acid trehalase-like 1** | PGGHG | T543 | | | **7.4** | **19.5** | R.DPLQSPLWVT*LGSSSPTESLTVDPASE.- |
| **ATH1, acid trehalase-like 1** | PGGHG | S546 | | | **7.0** | **18.1** | R.DPLQSPLWVTLGS*SSPTESLTVDPASE.- |
| **ATH1, acid trehalase-like 1** | PGGHG | T554 | | | **7.3** | **18.7** | R.DPLQSPLWVTLGSSSPTESLT*VDPASE.- |
| **ATP binding cassette 50** | ABCF1 | S105 | | | **6.6** | **69.9** | K.LS*VPTSDEEDEVPAPKPR.G |
| **ATP binding cassette 50** | ABCF1 | T108 | | | **17.6** | **44.9** | K.LSVPT*SDEEDEVPAPKPR.G |
| **ATP binding cassette 50** | ABCF1 | S228 | | | **178.6** | **159.3** | K.AEQGS*EEEGEGEEEEEEGGESK.A |
| **ATP binding cassette 50** | ABCF1 | T108S109 | | | **20.8** | **19.9** | K.LSVPT*S*DEEDEVPAPKPR.G |
| **ATP binding cassette 50** | ABCF1 | S140 | | | **100.0** | **38.9** | K.GGNVFAALIQDQS*EEEEEEEKHPPKPAKPEK.N |
| **ATP binding cassette 50** | ABCF1 | S105T108S109 | | | **100.0** | **20.6** | K.KLS*VPT*S*DEEDEVPAPKPR.G |
| **ATP binding cassette 50** | ABCF1 | S109 | | | **5.5** | **36.8** | K.LSVPTS*DEEDEVPAPKPR.G |
| **ATP binding cassette 50** | ABCF1 | S105T108 | | | **7.3** | **38.5** | K.LS*VPT*SDEEDEVPAPKPR.G |
| **ATP binding cassette, subfamily C, member 4** | ABCC4 | T646 | | | **13.9** | **30.3** | K.DNEESEQPPVPGT*PTLR.N |
| **ATP binding cassette, subfamily C, member 4** | ABCC4 | S664S668 | | | **19.0** | **46.3** | R.TFSESSVWSQQS*SRPS*LK.D |
| **ATP binding cassette, subfamily C, member 4** | ABCC4 | S664S665 | | | **6.5** | **18.3** | R.TFSESSVWSQQS*S*RPSLK.D |
| **ATP citrate lyase** | ACLY | S481 | | | **28.4** | **45.6** | K.AKPAMPQDSVPS*PR.S |
| **ATP citrate lyase** | ACLY | S455 | | | **29.9** | **52.5** | R.TAS*FSESR.A |
| **ATP-dependent RNA helicase A** | DHX9 | S77 | | | | **42.3** | K.S*EEVPAFGVASPPPLTDTPDTTANAEGDLPTTMGGPLPPHLALK.^ |
| **ATP-dependent RNA helicase DDX3X** | DDX3X | S590 | | | **11.2** | **37.7** | K.S*RFSGGFGAR.D |
| **ATP-dependent RNA helicase DDX3X** | DDX3X | S593 | | | **13.6** | **35.7** | K.SRFS*GGFGAR.D |
| **ATP7A** | ATP7A | S272 | | | **-0.6** | **54.6** | R.SPS*YTNDSTATFIIDGMHCK.S |
| **ATPase classVI type11B** | ATP11B | S1154 | | | **13.8** | **52.4** | R.SWS*ASDPFYTNDR.S |
| **ATPase family AAA domain-containing protein 2B** | ATAD2B | S727T731 | | | **2.8** | **13.1** | K.EDIETLILEDSEDENALS*IFET*NCHSGSPK.K |
| **ATPase family AAA domain-containing protein 2B** | ATAD2B | S727S735 | | | **2.0** | **16.2** | K.EDIETLILEDSEDENALS*IFETNCHS*GSPK.K |
| **ATPase H+ transporting lysosomal V0 subunit a isoform** | ATP6V0A2 | S695 | | | **53.6** | **46.7** | R.KDS*EEEVSLLGSQDIEEGNHQVEDGCR.E |
| **ATPase II** | ATP8A1 | S25T28S29 | | | **38.4** | **78.1** | K.TDDVS*EKT*S*LADQEEVR.T |
| **ATPase II** | ATP8A1 | T21T28S29 | | | **11.3** | **49.5** | K.T*DDVSEKT*S*LADQEEVR.T |
| **ATPase II** | ATP8A1 | S9 | | | **20.5** | **36.8** | R.TVS*EIR.S |
| **ATPase II** | ATP8A1 | T21S25T28 | | | **5.6** | **27.7** | K.T*DDVS*EKT*SLADQEEVR.T |
| **ATPase, Class VI, type 11C** | ATP11C | S445 | | | **17.8** | **11.4** | K.GVTQEVDGLS*QTDGTLTYFDK.V |
| **Autoantigen La** | SSB | S366 | | | **137.7** | **75.6** | K.FAS*DDEHDEHDENGATGPVKR.A |
| **Autoantigen RCD8** | EDC4 | S879 | | | **25.3** | **77.2** | R.DSQDASAEQS*DHDDEVASLASASGGFGTK.V |
| **Autoantigen RCD8** | EDC4 | S723 | | | **6.6** | **79.0** | K.GPGQVPTATSALSLELQEVEPLGLPQAS*PSR.T |
| **Autoantigen RCD8** | EDC4 | S729S741 | | | **45.8** | **56.4** | R.TRS*PDVISSASTALS*QDIPEIASEALSR.G |
| **Autoantigen RCD8** | EDC4 | S729 | | | **7.6** | **93.4** | R.TRS*PDVISSASTALSQDIPEIASEALSR.G |
| **Autoantigen RCD8** | EDC4 | S708S723 | | | **34.2** | **67.8** | K.GPGQVPTATSALS*LELQEVEPLGLPQAS*PSR.T |
| **Autoantigen RCD8** | EDC4 | S875 | | | **12.4** | **36.2** | R.DSQDAS*AEQSDHDDEVASLASASGGFGTK.V |
| **Autoantigen RCD8** | EDC4 | T727 | | | **15.8** | **98.8** | R.T*RSPDVISSASTALSQDIPEIASEALSR.G |
| **Autoantigen RCD8** | EDC4 | T704S723 | | | **1.4** | **17.0** | K.GPGQVPTAT*SALSLELQEVEPLGLPQAS*PSR.T |
| **Autoantigen RCD8** | EDC4 | S729S737 | | | **26.1** | **49.2** | R.S*PDVISSAS*TALSQDIPEIASEALSR.G |
| **Autoantigen RCD8** | EDC4 | T702 | | | | **46.5** | K.GPGQVPT*ATSALSLELQEVEPLGLPQASPSR.T |
| **Autoantigen RCD8** | EDC4 | S705S723 | | | **12.1** | **34.1** | K.GPGQVPTATS*ALSLELQEVEPLGLPQAS*PSR.T |

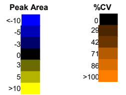


**White dots-** Significant change in peptide abundance at **5** %FDR compared to the timepoint with the minimum peak area for a given PSM

| Protein Name | Gene | Phosphosites | CarT 0m | CarT 2m | CarT 5m | RajiB 0m | RajiB 2m | RajiB 5m | Ascor | MOWSE | Sequence |
|---|---|---|---|---|---|---|---|---|---|---|---|
| Autoantigen RCD8 | EDC4 | S734 | | | | | | | 0.5 | 29.7 | R.SPDVIS*SASTALSQDIPEIASEALSR.G |
| Autoantigen RCD8 | EDC4 | S729S735 | | | | | | | 21.7 | 38.3 | R.S*PDVISS*ASTALSQDIPEIASEALSR.G |
| Autoantigen RCD8 | EDC4 | T727S741 | | | | | | | 6.4 | 18.7 | R.T*RSPDVISSASTALS*QDIPEIASEALSR.G |
| Autophagy related 9 homolog A | ATG9A | S761 | | | | | | | 7.6 | 67.9 | R.SAS*YPCAAPRPGAPETTALHGGFQR.R |
| Autophagy related 9 homolog A | ATG9A | S828 | | | | | | | 100.0 | 23.1 | R.HPEPVPEEGS*EDELPPQVHKV.- |
| B cell CLL/lymphoma 9 like | BCL9L | S1480 | | | | | | | 11.7 | 29.3 | R.SVS*LDSQMGYLPAPGGMANLPF.- |
| B cell CLL/lymphoma 9 like | BCL9L | S934 | | | | | | | 7.2 | 44.4 | K.S*PTLSQVHSPLVTSPSANLK.S |
| B cell CLL/lymphoma 9 like | BCL9L | S750 | | | | | | | 54.7 | 53.0 | R.GLLS*PPMGQSGLR.E |
| B cell CLL/lymphoma 9 like | BCL9L | S1478 | | | | | | | 10.0 | 32.8 | R.S*VSLDSQMGYLPAPGGMANLPF.- |
| B cell lymphoma 6 protein | BCL6 | S307S308 | | | | | | | 100.0 | 31.9 | K.EEERPS*S*EDEIALHFEPPNAPLNR.K |
| B-lymphocyte cell-surface antigen B1 | MS4A1 | S253 | | | | | | | 6.1 | 38.6 | K.EEVVGLTETS*SQPK.N |
| B-lymphocyte cell-surface antigen B1 | MS4A1 | S7 | | | | | | | 51.9 | 75.4 | R.NS*VNGTFPAEPMK.G |
| | MS4A1 | S225 | | | | | | | 101.1 | 64.7 | K.S*NIVLLSAEEKK.E |
| B-lymphocyte cell-surface antigen B1 | MS4A1 | T252S254 | | | | | | | 11.5 | 53.6 | K.EEVVGLTET*SS*QPK.N |
| | MS4A1 | S36 | | | | | | | -0.1 | 64.2 | R.MSS*LVGPTQSFFMR.E |
| B-lymphocyte cell-surface antigen B1 | MS4A1 | S35S36 | | | | | | | 68.1 | 86.9 | R.MS*S*LVGPTQSFFMR.E |
| B-lymphocyte cell-surface antigen B1 | MS4A1 | S295 | | | | | | | 30.5 | 26.4 | K.NEEDIEIIPIQEEEEEETETNFPEPPQDQESSPIENDS*SP.- |
| B-lymphocyte cell-surface antigen B1 | MS4A1 | S25 | | | | | | | 100.0 | 37.0 | K.GPIAMQS*GPKPLFR.R |
| | MS4A1 | S35 | | | | | | | 27.6 | 61.6 | R.RM#S*SLVGPTQSFFMR.E |
| B-lymphocyte cell-surface antigen B1 | MS4A1 | S288 | | | | | | | 4.6 | 18.6 | K.NEEDIEIIPIQEEEEEETETNFPEPPQDQES*SPIENDSSP.- |
| B-lymphocyte cell-surface antigen B1 | MS4A1 | T252S253 | | | | | | | 9.3 | 29.7 | K.EEVVGLTET*S*SQPK.N |
| B-lymphocyte cell-surface antigen B1 | MS4A1 | T275 | | | | | | | 6.1 | 23.1 | K.NEEDIEIIPIQEEEEEET*ETNFPEPPQDQESSPIENDSSP.- |
| B-lymphocyte cell-surface antigen B1 | MS4A1 | S254 | | | | | | | 7.2 | 35.6 | K.EEVVGLTETSS*QPK.N |
| B-lymphocyte cell-surface antigen B1 | MS4A1 | T250T252 | | | | | | | 14.6 | 33.9 | K.EEVVGLT*ET*SSQPK.N |
| B-lymphocyte cell-surface antigen B1 | MS4A1 | T252 | | | | | | | 0.5 | 30.7 | K.EEVVGLTET*SSQPK.N |
| B-lymphocyte cell-surface antigen B1 | MS4A1 | T250S254 | | | | | | | 3.5 | 26.1 | K.EEVVGLT*ETSS*QPK.N |
| B-lymphocyte cell-surface antigen B1 | MS4A1 | S253S254 | | | | | | | 8.9 | 39.1 | K.EEVVGLTETS*S*QPK.N |
| B-lymphocyte cell-surface antigen B1 | MS4A1 | S35S43 | | | | | | | -0.5 | 24.3 | R.RMS*SLVGPTQS*FFM#R.E |
| B-lymphocyte cell-surface antigen B1 | MS4A1 | S289 | | | | | | | 5.5 | 12.0 | K.NEEDIEIIPIQEEEEEETETNFPEPPQDQESS*PIENDSSP.- |
| B-Raf | BRAF | S446 | | | | | | | 92.2 | 50.0 | R.RDS*SDDWEIPDGQITVGQR.I |
| | BRAF | S365 | | | | | | | 23.6 | 77.7 | R.SSS*APNVHINTIEPVNIDDLIR.D |
| B-Raf | BRAF | S364 | | | | | | | -0.3 | 52.5 | R.SS*SAPNVHINTIEPVNIDDLIR.D |
| B-Raf | BRAF | S447 | | | | | | | 9.0 | 55.0 | R.RDSS*DDWEIPDGQITVGQR.I |
| B-Raf | BRAF | T753 | | | | | | | 70.5 | 53.2 | K.T*PIQAGGYGAFPVH.- |
| B-Raf | BRAF | S729 | | | | | | | 9.5 | 42.8 | R.SAS*EPSLNR.A |
| B-Raf | BRAF | S363 | | | | | | | 1.8 | 39.8 | R.DRS*SSAPNVHINTIEPVNIDDLIR.D |
| B-Raf | BRAF | T373 | | | | | | | -5.3 | 17.5 | R.SSSAPNVHINT*IEPVNIDDLIR.D |
| Baculoviral IAP repeat containing protein 6 | BIRC6 | S452 | | | | | | | 18.4 | 42.4 | K.LEGDSDDLLEDS*DSEEHSR.S |
| Baculoviral IAP repeat containing protein 6 | BIRC6 | T4689S4691T46 | | | | | | | 28.9 | 12.6 | R.GT*PS*GT*QSSR.E |
| Baculoviral IAP repeat containing protein 6 | BIRC6 | S421 | | | | | | | 12.0 | 80.7 | K.FEINAYDPAIVQQLILSGDPSSGVDS*R.R |
| Basic leucine zipper and W2 domains 1 | BZW1 | S345 | | | | | | | 22.6 | 92.9 | K.NAEEES*ESEAEEGD.- |
| Basic leucine zipper and W2 domains 1 | BZW1 | S345S347 | | | | | | | 100.0 | 62.9 | K.NAEEES*ES*EAEEGD.- |
| | BZW2 | S412S414 | | | | | | | 100.0 | 48.7 | K.FVEWLQNAEEES*ES*EGEEN.- |
| Basic transcription factor 3 | BTF3 | S158 | | | | | | | 15.8 | 52.9 | K.QLTEMLPSILNQLGADS*LTSLR.R |
| BAT2 domain containing 1 | PRRC2C | S878 | | | | | | | 25.7 | 47.1 | R.S*VEDVRPHHTDANNQSACFEAPDQK.T |
| BAT2 domain containing 1 | PRRC2C | S1248S1249 | | | | | | | 3.1 | 42.3 | R.SES*S*DFEVVPK.R |
| BAT2 domain containing 1 | PRRC2C | S1542 | | | | | | | 15.3 | 14.3 | R.S*FSSQRPVDR.Q |
| BAT2 domain containing 1 | PRRC2C | S1246S1249 | | | | | | | 7.4 | 35.0 | R.S*ESS*DFEVVPK.R |
| BAT2 domain containing 1 | PRRC2C | S1246S1248 | | | | | | | 5.5 | 37.7 | R.S*ES*SDFEVVPK.R |
| BAT2 domain containing 1 | PRRC2C | S2105 | | | | | | | 100.0 | 21.7 | K.LPDLS*PVENK.E |
| Bcl 2 related proline rich protein | BCL2L12 | S242 | | | | | | | 16.9 | 35.2 | R.LS*SDSFAR.L |
| BCL11A | BCL11A | S625S630 | | | | | | | 23.8 | 59.8 | K.LLLGS*PSSLS*PFSK.R |
| BCL2 antagonist of cell death | BAD | S118 | | | | | | | 53.1 | 28.4 | R.RMS*DEFVDSFKK.G |
| BCL2 antagonist of cell death | BAD | S99 | | | | | | | 100.0 | 39.6 | R.S*APPNLWAAQR.Y |
| BCL2 associated athanogene 2 | BAG2 | S19 | | | | | | | -0.4 | 34.4 | R.SS*SMADR.S |
| BCL2 associated athanogene 2 | BAG2 | S20 | | | | | | | 9.6 | 14.3 | R.SSS*MADR.S |
| BCL2 associated transcription factor 1 | BCLAF1 | S177 | | | | | | | 100.0 | 33.3 | K.AEGEPQEES*PLK.S |
| BCL2 associated transcription factor 1 | BCLAF1 | S285S287 | | | | | | | 16.4 | 36.3 | R.YS*PS*QNSPIHHIPSRR.S |
| BCL2 associated transcription factor 1 | BCLAF1 | S287S297 | | | | | | | 12.4 | 17.4 | R.YSPS*QNSPIHHIPS*RR.S |
| BCL2 associated transcription factor 1 | BCLAF1 | Y284S287 | | | | | | | 17.6 | 29.0 | R.Y*SPS*QNSPIHHIPSRR.S |
| BCL2 associated transcription factor 1 | BCLAF1 | S285S290 | | | | | | | 19.5 | 52.6 | R.YS*PSQNS*PIHHIPSRR.S |
| BCL2 associated transcription factor 1 | BCLAF1 | S397 | | | | | | | 55.7 | 87.2 | K.QKFNDS*EGDDTEETEDYR.Q |

Peak Area: <-10, -5, -3, 0, 3, 5, >10

%CV: 0, 29, 42, 71, 86, >100

**White dots-** Significant change in peptide abundance at **5** %FDR compared to the timepoint with the minimum peak area for a given PSM

| Protein Name | Gene | Phosphosites | CarT 0m | CarT 2m | CarT 5m | RajiB 0m | RajiB 2m | RajiB 5m | Ascor | MOWSE | Sequence |
|---|---|---|---|---|---|---|---|---|---|---|---|
| **BCL2 associated transcription factor 1** | BCLAF1 | Y284S290 | | | | | | | **13.4** | **64.1** | R.Y*SPSQNS*PIHHIPSR.R |
| | BCLAF1 | S285 | | | | | | | **22.0** | **78.3** | R.YS*PSQNSPIHHIPSR.R |
| **BCL2 associated transcription factor 1** | BCLAF1 | S397T402 | | | | | | | **63.3** | **125.1** | K.FNDS*EGDDT*EETEDYR.Q |
| **BCL2 associated transcription factor 1** | BCLAF1 | S658 | | | | | | | **32.3** | **61.5** | R.IDIS*PSTLR.K |
| **BCL2 associated transcription factor 1** | BCLAF1 | Y383 | | | | | | | **5.2** | **75.2** | R.AEGEWEDQEALDY*FSDKESGK.Q |
| **BCL2 associated transcription factor 1** | BCLAF1 | S385S389 | | | | | | | **6.0** | **47.8** | R.AEGEWEDQEALDYFS*DKES*GK.Q |
| **BCL2 associated transcription factor 1** | BCLAF1 | S222 | | | | | | | **39.4** | **55.8** | K.SSATSGDIWPGLSAYDNS*PR.S |
| **BCL2 associated transcription factor 1** | BCLAF1 | Y219 | | | | | | | **22.6** | **66.1** | K.SSATSGDIWPGLSAY*DNSPR.S |
| **BCL2 associated transcription factor 1** | BCLAF1 | S385 | | | | | | | **22.8** | **107.8** | R.AEGEWEDQEALDYFS*DK.E |
| **BCL2 associated transcription factor 1** | BCLAF1 | S121 | | | | | | | **13.8** | **29.5** | R.SVS*SQR.S |
| **BCL2 associated transcription factor 1** | BCLAF1 | S121S122 | | | | | | | **13.8** | **23.7** | R.SVS*S*QR.S |
| **BCL2 associated transcription factor 1** | BCLAF1 | S119S122 | | | | | | | **7.8** | **20.3** | R.S*VSS*QR.S |
| **BCL2 associated transcription factor 1** | BCLAF1 | S119S121 | | | | | | | **13.0** | **24.0** | R.S*VS*SQR.S |
| **BCL2 associated transcription factor 1** | BCLAF1 | S102S104 | | | | | | | **100.0** | **14.5** | R.RHS*RS*PR.R |
| | BCLAF1 | S496 | | | | | | | **27.7** | **45.3** | K.KETQS*PEQVK.S |
| **BCL2 associated transcription factor 1** | BCLAF1 | T257S268 | | | | | | | **35.6** | **40.9** | K.NT*PSQHSHSIQHS*PER.S |
| **BCL2 associated transcription factor 1** | BCLAF1 | S531 | | | | | | | **100.0** | **49.0** | R.EES*PLR.I |
| | BCLAF1 | Y284S285 | | | | | | | **12.3** | **16.8** | R.Y*S*PSQNSPIHHIPSRR.S |
| **BCL2 associated transcription factor 1** | BCLAF1 | S339 | | | | | | | **100.0** | **15.1** | K.FLKS*PPLHK.N |
| **BCL2 associated transcription factor 1** | BCLAF1 | S287 | | | | | | | **37.7** | **78.4** | R.YSPS*QNSPIHHIPSR.R |
| **BCL2 associated transcription factor 1** | BCLAF1 | S119S121S122 | | | | | | | **100.0** | **12.7** | R.S*VS*S*QRS*R.S |
| **BCL2 associated transcription factor 1** | BCLAF1 | T494 | | | | | | | **10.9** | **18.1** | K.ET*QSPEQVK.S |
| **BCL2 associated transcription factor 1** | BCLAF1 | S183 | | | | | | | **14.9** | **37.8** | K.S*QEEPKDTFEHDPSESIDEFNK.S |
| **BCL2 associated transcription factor 1** | BCLAF1 | Y383S385 | | | | | | | **13.8** | **61.9** | R.AEGEWEDQEALDY*FS*DKESGK.Q |
| **BCL2 associated transcription factor 1** | BCLAF1 | S217 | | | | | | | **7.3** | **61.6** | K.SSATSGDIWPGLS*AYDNSPR.S |
| **BCL2 associated transcription factor 1** | BCLAF1 | T257S264 | | | | | | | **30.1** | **18.0** | K.NT*PSQHSHS*IQHSPER.S |
| **BCL2 associated transcription factor 1** | BCLAF1 | S281S287 | | | | | | | **9.0** | **16.1** | R.SGSGSVGNGS*SRYSPS*QNSPIHHIPSR.R |
| **BCL2 associated transcription factor 1** | BCLAF1 | S132Y133S135 | | | | | | | **5.7** | **19.6** | R.RS*Y*RS*SR.S |
| **BCL2 associated transcription factor 1** | BCLAF1 | Y133S135S136 | | | | | | | **11.8** | **13.1** | R.RSY*RS*S*R.S |
| **BCL2 associated transcription factor 1** | BCLAF1 | Y383S389 | | | | | | | **4.5** | **48.3** | R.AEGEWEDQEALDY*FSDKES*GK.Q |
| **BCL2 associated transcription factor 1** | BCLAF1 | S259S268 | | | | | | | **16.6** | **16.5** | K.NTPS*QHSHSIQHS*PER.S |
| **BCL2 associated transcription factor 1** | BCLAF1 | S648 | | | | | | | **100.0** | **36.9** | R.QKS*PEIHR.R |
| **BCL2 associated transcription factor 1** | BCLAF1 | S268 | | | | | | | **6.5** | **32.9** | K.NTPSQHSHSIQHS*PER.S |
| **BCL2 associated transcription factor 1** | BCLAF1 | S276S281 | | | | | | | **3.8** | **12.0** | R.SGSGS*VGNGS*SRYSPSQNSPIHHIPSR.R |
| **BCL2 associated transcription factor 1** | BCLAF1 | S225S228 | | | | | | | **16.0** | **52.3** | R.S*PHS*PSPIATPPSQSSSCSDAPMLSTVHSAK.N |
| **BCL2 associated transcription factor 1** | BCLAF1 | S259 | | | | | | | **9.4** | **23.5** | K.NTPS*QHSHSIQHSPER.S |
| **BCL2 associated transcription factor 1** | BCLAF1 | S225T234 | | | | | | | **9.6** | **49.5** | R.S*PHSPSPIAT*PPSQSSSCSDAPMLSTVHSAK.N |
| **BCL2 associated transcription factor 1** | BCLAF1 | S36 | | | | | | | **16.9** | **16.3** | K.RYSS*R.S |
| **BCL2 associated transcription factor 1** | BCLAF1 | S276S287 | | | | | | | **3.8** | **14.1** | R.SGSGS*VGNGSSRYSPS*QNSPIHHIPSR.R |
| **BCL2 associated transcription factor 1** | BCLAF1 | S264 | | | | | | | **7.9** | **24.7** | K.NTPSQHSHS*IQHSPER.S |
| **BCL2 associated transcription factor 1** | BCLAF1 | S290 | | | | | | | **8.3** | **30.4** | R.YSPSQNS*PIHHIPSR.R |
| **BCL2 associated transcription factor 1** | BCLAF1 | Y284 | | | | | | | **26.0** | **75.0** | R.Y*SPSQNSPIHHIPSR.R |
| **BCL2 associated transcription factor 1** | BCLAF1 | S225S230 | | | | | | | **9.0** | **54.0** | R.S*PHSPS*PIATPPSQSSSCSDAPMLSTVHSAK.N |
| **BCL2 associated transcription factor 1** | BCLAF1 | S660 | | | | | | | **13.9** | **10.9** | R.RIDISPS*TLRK.H |
| **BCL2 associated transcription factor 1** | BCLAF1 | S512 | | | | | | | **7.4** | **20.3** | K.LKDLFDYS*PPLHK.N |
| **BCL2 associated transcription factor 1** | BCLAF1 | Y511 | | | | | | | | **12.9** | K.DLFDY*SPPLHK.N |
| **BCL2 associated transcription factor 1** | BCLAF1 | S276S290 | | | | | | | **21.6** | **11.9** | R.SGSGS*VGNGSSRYSPSQNS*PIHHIPSR.R |
| **BCL2 associated transcription factor 1** | BCLAF1 | S287S290 | | | | | | | **25.1** | **42.3** | R.YSPS*QNS*PIHHIPSR.R |
| **BCL2 associated transcription factor 1** | BCLAF1 | S119S121S122 | | | | | | | **100.0** | **13.0** | R.S*VS*S*QR.S |
| **BCL2-like 11 (apoptosis facilitator)** | BCL2L11 | S77 | | | | | | | **100.0** | **64.8** | R.S*PLFIFMR.R |
| **BCL2-like 11 (apoptosis facilitator)** | BCL2L11 | S86S87S90 | | | | | | | **100.0** | **20.8** | R.RS*S*LLS*R.S |
| **BCL2-like 11 (apoptosis facilitator)** | BCL2L11 | S94 | | | | | | | **7.6** | **33.7** | R.SSS*GYFSFDTDR.S |
| **BCL2L13** | BCL2L13 | S371 | | | | | | | **-0.3** | **74.5** | K.SS*PATSLFVELDEEEVK.A |
| **BCL6 corepressor** | BCOR | S747S749 | | | | | | | **100.0** | **11.2** | R.S*RS*HER.A |
| **BCR** | BCR | S1264 | | | | | | | **42.3** | **21.6** | K.RQS*ILFSTEV.- |
| **BCR** | BCR | S315Y316 | | | | | | | **10.9** | **13.5** | R.RS*Y*SPR.S |
| **Beta adaptin 3A** | AP3B1 | S276 | | | | | | | **39.3** | **30.1** | K.NFYES*DDDQKEK.T |
| **Beta adaptin 3A** | AP3B1 | Y274 | | | | | | | **1.0** | **19.1** | K.EGDELEDNGKNFY*ESDDDQKEK.T |
| **Beta adaptin 3A** | AP3B1 | S750S752 | | | | | | | **100.0** | **57.3** | K.GKS*DS*EDGEKENEK.S |
| **Beta cysteine string protein** | DNAJC5B | T16 | | | | | | | **10.3** | **28.1** | R.TLSTT*GEALYEILGLHK.G |
| **BHC110** | KDM1A | S166 | | | | | | | **58.7** | **39.7** | K.LPPPPPQAPPEEENES*EPEEPSGVEGAAFQSR.L |
| **BHC110** | KDM1A | S131S137 | | | | | | | **26.0** | **111.9** | R.EMDESLANLS*EDEYYS*EEER.N |

Peak Area: <-10, -5, -3, 0, 3, 5, >10

%CV: 0, 29, 42, 71, 86, >100

**White dots-** Significant change in peptide abundance at **5** %FDR compared to the timepoint with the minimum peak area for a given PSM

| Protein Name | Gene | Phosphosites | CarT (0m, 2m, 5m) | RajiB (0m, 2m, 5m) | Ascor | MOWSE | Sequence |
|---|---|---|---|---|---|---|---|
| **BHC110** | KDM1A | Y136S137 | | | 9.9 | 32.9 | R.EMDESLANLSEDEYY*S*EEER.N |
| **BHC110** | KDM1A | S126S137 | | | 12.5 | 50.2 | R.EMDES*LANLSEDEYYS*EEER.N |
| **BHC110** | KDM1A | S126S131 | | | 4.4 | 11.3 | R.EMDES*LANLS*EDEYYSEEER.N |
| **BHC110** | KDM1A | S849 | | | 14.7 | 19.3 | R.QATPGVPAQQS*PSM.- |
| **BHC80** | PHF21A | T446S455 | | | 19.3 | 25.6 | K.YNAVLGFGALTPT*SPQSSHPDS*PENEK.T |
| **BHC80** | PHF21A | T446S451 | | | 5.7 | 27.6 | K.YNAVLGFGALTPT*SPQSS*HPDSPENEK.T |
| **BHC80** | PHF21A | T446S450 | | | 4.3 | 25.5 | K.YNAVLGFGALTPT*SPQS*SHPDSPENEK.T |
| **BHC80** | PHF21A | S447S455 | | | 19.2 | 15.8 | K.YNAVLGFGALTPTS*PQSSHPDS*PENEK.T |
| **BIG1** | ARFGEF1 | S1569 | | | 73.2 | 39.7 | K.S*VDIHDSIQPR.S |
| **BIG1** | ARFGEF1 | Y662S670 | | | 4.3 | 67.9 | R.Y*GSLNSLES*TSSSGIGSYSTQMSGTDNPEQFEVLK.Q |
| **BIG1** | ARFGEF1 | S664S678 | | | 3.0 | 97.6 | R.YGS*LNSLESTSSSGIGS*YSTQMSGTDNPEQFEVLK.Q |
| **BIG1** | ARFGEF1 | S664S670 | | | 7.6 | 98.1 | R.YGS*LNSLES*TSSSGIGSYSTQMSGTDNPEQFEVLK.Q |
| **BIG1** | ARFGEF1 | S667S670 | | | 6.0 | 47.8 | R.YGSLNS*LES*TSSSGIGSYSTQMSGTDNPEQFEVLK.Q |
| **BIG1** | ARFGEF1 | S664S667 | | | 3.7 | 92.9 | R.YGS*LNS*LESTSSSGIGSYSTQMSGTDNPEQFEVLK.Q |
| **BIG1** | ARFGEF1 | S664S680 | | | 0.4 | 88.4 | R.YGS*LNSLESTSSSGIGSYS*TQMSGTDNPEQFEVLK.Q |
| **BIG1** | ARFGEF1 | S678Y679 | | | -4.5 | 33.2 | R.YGSLNSLESTSSSGIGS*Y*STQM#SGTDNPEQFEVLK.Q |
| **BIG1** | ARFGEF1 | S664S674 | | | 2.0 | 92.8 | R.YGS*LNSLESTSSS*GIGSYSTQMSGTDNPEQFEVLK.Q |
| **BIG1** | ARFGEF1 | S664Y679 | | | 4.7 | 66.8 | R.YGS*LNSLESTSSSGIGSY*STQMSGTDNPEQFEVLK.Q |
| **BIG1** | ARFGEF1 | S670S673T681 | | | 4.9 | 17.0 | R.YGSLNSLES*TSS*SGIGSYST*QMSGTDNPEQFEVLK.Q |
| **BIG1** | ARFGEF1 | Y662S667 | | | 3.6 | 53.0 | R.Y*GSLNS*LESTSSSGIGSYSTQMSGTDNPEQFEVLK.Q |
| **BIG1** | ARFGEF1 | S678S680 | | | 4.6 | 38.2 | R.YGSLNSLESTSSSGIGS*YS*TQMSGTDNPEQFEVLK.Q |
| **BIG1** | ARFGEF1 | S670T671 | | | 5.1 | 31.5 | R.YGSLNSLES*T*SSSGIGSYSTQMSGTDNPEQFEVLK.Q |
| **BIG1** | ARFGEF1 | S670S673S680 | | | 6.2 | 19.5 | R.YGSLNSLES*TSS*SGIGSYS*TQMSGTDNPEQFEVLK.Q |
| **BIG1** | ARFGEF1 | S670S672S678 | | | 6.2 | 36.2 | R.YGSLNSLES*TS*SSGIGS*YSTQMSGTDNPEQFEVLK.Q |
| **BIG1** | ARFGEF1 | S670T671S678 | | | 5.2 | 20.8 | R.YGSLNSLES*T*SSSGIGS*Y*STQMSGTDNPEQFEVLK.Q |
| **BIG1** | ARFGEF1 | S664T671 | | | 2.5 | 70.1 | R.YGS*LNSLEST*SSSGIGSYSTQMSGTDNPEQFEVLK.Q |
| **BIN2** | BIN2 | S458 | | | 7.8 | 48.4 | R.ASLGTGTAS*PR.T |
| **BIN2** | BIN2 | S357 | | | 24.4 | 35.4 | R.AKS*QEEVLPSSTTPSPGGALSPSGQPSSSATEVVLR.T |
| **BIN2** | BIN2 | S273S277 | | | 10.3 | 32.4 | R.TATVSS*PLTS*PTSPSTLSLK.S |
| **BIN2** | BIN2 | S259 | | | 67.1 | 35.7 | R.S*LVISPPVR.T |
| **BIN2** | BIN2 | T454 | | | 8.4 | 35.4 | R.ASLGT*GTASPR.T |
| **Bin3, bicoid-interacting 3, homolog** | MEPCE | S254 | | | 39.4 | 31.1 | K.DITDPLSLNTCTDEGHVVLAS*PLK.T |
| **Bin3, bicoid-interacting 3, homolog** | MEPCE | S330 | | | 12.4 | 71.1 | R.DEVVS*PLPSALQGPSGSLSAPPAASVISAPPSSSSR.H |
| **Bin3, bicoid-interacting 3, homolog** | MEPCE | S334 | | | 7.7 | 52.6 | R.DEVVSPLPS*ALQGPSGSLSAPPAASVISAPPSSSSR.H |
| **Bin3, bicoid-interacting 3, homolog** | MEPCE | S342 | | | -0.3 | 42.8 | R.DEVVSPLPSALQGPSGS*LSAPPAASVISAPPSSSSR.H |
| **Block of proliferation 1** | BOP1 | S126S127 | | | 91.4 | 114.9 | R.IGDEYAEDS*S*DEEDIR.N |
| **Block of proliferation 1** | BOP1 | Y122S126 | | | 6.3 | 27.6 | R.IGDEY*AEDS*SDEEDIR.N |
| **Block of proliferation 1** | BOP1 | S126T135 | | | 10.5 | 17.8 | R.IGDEYAEDS*SDEEDIRNT*VGNVPLEWYDDFPHVGYDLDGR.R |
| **BLOS3** | BLOC1S3 | T63S65 | | | 73.4 | 48.2 | R.VAGEAAET*DS*EPEPEPEPTAAPR.D |
| **BMP2 inducible protein kinase** | BMP2K | S949 | | | 35.1 | 68.7 | K.SES*NEDLFGLVPFDEITGSQQQK.V |
| **BMP2 inducible protein kinase** | BMP2K | S947 | | | 10.0 | 108.7 | K.S*ESNEDLFGLVPFDEITGSQQQK.V |
| | BMS1 | S639 | | | 3.1 | 33.4 | K.LGPQNFIDETS*DIENLLKEEEDYKEENNDSK.E |
| **BMS1 like ribosome assembly protein** | BMS1 | T638 | | | | 58.0 | K.LGPQNFIDET*SDIENLLK.E |
| **Bone marrow stromal cell derived ubiquitin like** | UBL7 | S230 | | | 123.4 | 59.2 | R.DMPGGFLFEGLS*DDEDDFHPNTR.S |
| **BPY2 interacting protein 1** | MAP1S | T782 | | | | 71.6 | R. [illegible] |
| **BPY2 interacting protein 1** | MAP1S | T638S640 | | | 36.2 | 38.8 | R.T*PS*PESHR.S |
| **BPY2 interacting protein 1** | MAP1S | S731 | | | -0.4 | 14.7 | R.SAS*PHDVDLCLVSPCEFEHRK.A |
| **BPY2 interacting protein 1** | MAP1S | S657 | | | 26.0 | 29.5 | R.LSLS*PLR.G |
| **BPY2 interacting protein 1** | MAP1S | S729 | | | 5.5 | 29.2 | R.S*ASPHDVDLCLVSPCEFEHR.K |
| **BPY2 interacting protein 1** | MAP1S | S759 | | | 25.4 | 34.7 | K.AVPMAPAPAS*PGSSNDSSAR.S |
| **Branched chain alpha-ketoacid dehydrogenase kinase** | BCKDK | T32 | | | -0.3 | 44.2 | R.ST*SATDTHHVEMAR.E |
| **Branched chain alpha-ketoacid dehydrogenase kinase** | BCKDK | S31 | | | 7.4 | 13.1 | R.S*TSATDTHHVEMAR.E |
| **Branched chain keto acid dehydrogenase E1, alpha polypeptide** | BCKDHA | Y353 | | | 7.1 | 11.1 | R.SVDEVNY*WDKQDHPISR.L |
| **Branched chain keto acid dehydrogenase E1, alpha polypeptide** | BCKDHA | S347 | | | 90.2 | 61.3 | R.S*VDEVNYWDK.Q |
| **Branched chain keto acid dehydrogenase E1, alpha polypeptide** | BCKDHA | S337 | | | 26.0 | 87.6 | R.IGHHS*TSDDSSAYR.S |
| **BRCA1** | BRCA1 | S1217S1218 | | | 36.9 | 79.7 | K.KLESSEENLS*S*EDEELPCFQHLLFGK.V |
| **BRCA1** | BRCA1 | S1189S1191 | | | 24.7 | 47.8 | R.S*PS*PFTHTHLAQGYR.R |
| **BRCA1** | BRCA1 | S1189T1194 | | | 19.1 | 30.9 | R.S*PSPFT*HTHLAQGYR.R |
| **BRCA1** | BRCA1 | S1212S1218 | | | 3.5 | 10.8 | K.LESS*EENLSS*EDEELPCFQHLLFGK.V |
| **BRCA1** | BRCA1 | S1189 | | | 20.5 | 51.1 | R.S*PSPFTHTHLAQGYR.R |
| **BRCA1** | BRCA1 | S1211S1212S12 | | | 100.0 | 30.1 | K.KLES*S*EENLS*S*EDEELPCFQHLLFGK.V |
| **BRCA2 and CDKN1A interacting protein** | BCCIP | S42 | | | 100.0 | 79.8 | R.DEEEEKEVENEDEDDDDS*DKEK.D |

Peak Area: <-10, -5, -3, 0, 3, 5, >10

%CV: 0, 29, 42, 71, 86, >100

**White dots-** Significant change in peptide abundance at **5** %FDR compared to the timepoint with the minimum peak area for a given PSM

| Protein Name | Gene | Phosphosites | CarT 0m | CarT 2m | CarT 5m | RajiB 0m | RajiB 2m | RajiB 5m | Ascor | MOWSE | Sequence |
|---|---|---|---|---|---|---|---|---|---|---|---|
| **BRCA2 and CDKN1A interacting protein** | BCCIP | S112S115 | | | | | | | **8.9** | **39.9** | K.QTDVS*EDS*NDDMDEDEVFGFISLLNLTER.K |
| **Brefeldin A-inhibited guanine nucleotide-exchange protein 2** | ARFGEF2 | S218S227 | | | | | | | **40.9** | **39.0** | R.ELEKPIQSKPQS*PVIQAAAVS*PK.F |
| **Brefeldin A-inhibited guanine nucleotide-exchange protein 2** | ARFGEF2 | S227 | | | | | | | **71.3** | **66.7** | R.ELEKPIQSKPQSPVIQAAAVS*PK.F |
| **BRG1 associated factor 180 KD** | PBRM1 | S10S14 | | | | | | | **15.1** | **58.3** | R.ATS*PSSS*VSGDFDDGHHSVSTPGPSR.K |
| **BRG1 associated factor 180 KD** | PBRM1 | S10S12S13S14 | | | | | | | **10.9** | **39.3** | R.ATS*PS*S*S*VSGDFDDGHHSVSTPGPSR.K |
| **BRG1 associated factor 180 KD** | PBRM1 | S371S375T377 | | | | | | | **7.4** | **26.6** | R.YEEGES*EAES*IT*SFMDVSNPFYQLYDTVR.S |
| **BRG1 associated factor 180 KD** | PBRM1 | S371S375S378 | | | | | | | **12.0** | **68.0** | R.YEEGES*EAES*ITS*FMDVSNPFYQLYDTVR.S |
| **BRG1 associated factor 180 KD** | PBRM1 | S375T377S378 | | | | | | | **6.8** | **10.9** | R.YEEGESEAES*IT*S*FMDVSNPFYQLYDTVR.S |
| **BRG1 associated factor 180 KD** | PBRM1 | S353S355 | | | | | | | **9.3** | **84.8** | R.LSAITMALQYGS*ES*EEDAALAAAR.Y |
| **BRG1 associated factor 180 KD** | PBRM1 | Y351S353 | | | | | | | **9.8** | **45.1** | R.LSAITMALQY*GS*ESEEDAALAAAR.Y |
| **BRG1 associated factor 180 KD** | PBRM1 | T9S10S12S14 | | | | | | | **5.0** | **14.5** | R.AT*S*PS*SS*VSGDFDDGHHSVSTPGPSR.K |
| **BRG1 associated factor 180 KD** | PBRM1 | S10S13S14S16 | | | | | | | **6.4** | **31.9** | R.ATS*PSS*S*VS*GDFDDGHHSVSTPGPSR.K |
| | PBRM1 | S375T377 | | | | | | | **2.7** | **63.3** | R.YEEGESEAES*IT*SFM#DVSNPFYQLYDTVR.S |
| | PBRM1 | S371S375 | | | | | | | **3.6** | **50.9** | R.YEEGES*EAES*ITSFM#DVSNPFYQLYDTVR.S |
| **BRG1 associated factor 180 KD** | PBRM1 | T346Y351 | | | | | | | **4.3** | **12.0** | R.LSAIT*MALQY*GSESEEDAALAAAR.Y |
| **BRG1 associated factor 180 KD** | PBRM1 | Y366S371 | | | | | | | **4.4** | **63.2** | R.Y*EEGES*EAESITSFMDVSNPFYQLYDTVR.S |
| **BRG1 associated factor 180 KD** | PBRM1 | Y366T377 | | | | | | | **3.9** | **13.2** | R.Y*EEGESEAESIT*SFMDVSNPFYQLYDTVR.S |
| **BRG1 associated factor 180 KD** | PBRM1 | S178 | | | | | | | **10.3** | **54.4** | K.GEADDEDDDEDGQDNQGTVTEGSS*PAYLK.E |
| **BRG1 associated factor 180 KD** | PBRM1 | S343T346 | | | | | | | **2.8** | **43.1** | R.LS*AIT*MALQYGSESEEDAALAAAR.Y |
| **BRG1 associated factor 180 KD** | PBRM1 | T346S353 | | | | | | | **6.7** | **23.4** | R.LSAIT*MALQYGS*ESEEDAALAAAR.Y |
| **BRG1 associated factor 180 KD** | PBRM1 | Y351S355 | | | | | | | **6.5** | **51.5** | R.LSAITMALQY*GSES*EEDAALAAAR.Y |
| **BRG1 associated factor 180 KD** | PBRM1 | S177 | | | | | | | **18.0** | **40.1** | K.GEADDEDDDEDGQDNQGTVTEGS*SPAYLK.E |
| **BRG1 associated factor 180 KD** | PBRM1 | S371T377 | | | | | | | **8.4** | **78.0** | R.YEEGES*EAESIT*SFM#DVSNPFYQLYDTVR.S |
| **BRG1 associated factor 180 KD** | PBRM1 | S371S383 | | | | | | | **4.5** | **41.8** | R.YEEGES*EAESITSFMDVS*NPFYQLYDTVR.S |
| **BRG1 associated factor 180 KD** | PBRM1 | S375S378 | | | | | | | **6.3** | **12.8** | R.YEEGESEAES*ITS*FMDVSNPFYQLYDTVR.S |
| **Bromo adjacent homology domain containing 1** | BAHD1 | S121 | | | | | | | **100.0** | **106.3** | R.LAS*LNAEALNNLLLER.E |
| **Bromodomain adjacent to zinc finger domain 1A** | BAZ1A | S820 | | | | | | | **58.5** | **76.8** | R.NSTADIS*IGEEER.E |
| **Bromodomain adjacent to zinc finger domain 1A** | BAZ1A | S420 | | | | | | | **45.1** | **40.5** | K.IAEQDFSYFFPDDPPTFIFS*PANR.R |
| **Bromodomain adjacent to zinc finger domain 1A** | BAZ1A | T731 | | | | | | | **28.9** | **22.4** | K.ELDQDMVT*EDEDDPGSHKR.G |
| **Bromodomain adjacent to zinc finger domain 1A** | BAZ1A | S1531 | | | | | | | **56.4** | **20.0** | R.KRQS*PEPSPVTLGR.R |
| **Bromodomain adjacent to zinc finger domain. 1B** | BAZ1B | S349 | | | | | | | **27.7** | **41.3** | K.SLSGS*PLK.V |
| **Bromodomain adjacent to zinc finger domain. 1B** | BAZ1B | S1468 | | | | | | | **113.7** | **145.4** | R.LAEDEGDS*EPEAVGQSR.G |
| **Bromodomain adjacent to zinc finger domain. 1B** | BAZ1B | S708T710 | | | | | | | **3.5** | **64.2** | R.SDVQEESEGS*DT*DDNKDSAAFEDNEVQDEFLEK.L |
| **Bromodomain adjacent to zinc finger domain. 1B** | BAZ1B | T710S716 | | | | | | | **4.0** | **64.3** | R.SDVQEESEGSDT*DDNKDS*AAFEDNEVQDEFLEK.L |
| **Bromodomain adjacent to zinc finger domain. 1B** | BAZ1B | S705S708 | | | | | | | **2.9** | **40.0** | R.SDVQEES*EGS*DTDDNKDSAAFEDNEVQDEFLEK.L |
| **Bromodomain and PHD finger containing 1** | BRPF1 | S238 | | | | | | | **14.1** | **36.5** | R.KTEGVS*PIPQEIFEYLMDR.L |
| **Bromodomain containing 1** | BRD1 | S1052S1055 | | | | | | | **17.4** | **36.6** | R.VHGEPTS*DLS*DID.- |
| **Bromodomain containing 4** | BRD4 | S1117 | | | | | | | **100.0** | **26.7** | K.IHS*PIIR.S |
| | BRD4 | S1064S1083 | | | | | | | **13.9** | **10.7** | R.EAPS*PLM#IHSPQMSQFQSLTHQS*PPQQNVQPK.K |
| **Bromodomain containing 4** | BRD4 | S1070S1083 | | | | | | | **24.3** | **23.2** | R.EAPSPLMIHS*PQMSQFQSLTHQS*PPQQNVQPK.K |
| **Bromodomain containing 4** | BRD4 | S1064S1070S1 | | | | | | | **35.3** | **18.8** | R.EAPS*PLMIHS*PQMSQFQSLTHQS*PPQQNVQPK.K |
| **Bromodomain containing 4** | BRD4 | T1080 | | | | | | | **21.4** | **27.0** | R.EAPSPLMIHSPQMSQFQSLT*HQSPPQQNVQPK.K |
| **Bromodomain containing 4** | BRD4 | S1078S1083 | | | | | | | **7.2** | **11.0** | R.EAPSPLMIHSPQMSQFQS*LTHQS*PPQQNVQPK.K |
| **Bromodomain containing 4** | BRD4 | S1083 | | | | | | | **14.2** | **12.4** | R.EAPSPLMIHSPQMSQFQSLTHQS*PPQQNVQPK.K |
| **Bromodomain containing 8** | BRD8 | T264S268 | | | | | | | **27.8** | **45.1** | K.AT*PPPS*PLLSELLK.K |
| **Bromodomain containing 8** | BRD8 | T264 | | | | | | | **41.9** | **52.1** | K.AT*PPPSPLLSELLK.K |
| **Bromodomain containing 8** | BRD8 | S268S272 | | | | | | | **1.6** | **38.8** | K.ATPPPS*PLLS*ELLK.K |
| **BTB POZ domain containing 14B** | NACC1 | S140S151 | | | | | | | **2.5** | **21.0** | K.VSSPSCDSQGLHAEEAPS*SEPQSPVAQTS*GWPACSTPLPLVSR.V |
| **BTB POZ domain containing 14B** | NACC1 | T150S151 | | | | | | | **6.9** | **21.7** | K.VSSPSCDSQGLHAEEAPSSEPQSPVAQT*S*GWPACSTPLPLVSR.V |
| **BTB POZ domain containing 14B** | NACC1 | S127S130 | | | | | | | **6.6** | **25.1** | K.VSSPS*CDS*QGLHAEEAPSSEPQSPVAQTSGWPACSTPLPLVSR.V |
| **BTB POZ domain containing 14B** | NACC1 | S124S125 | | | | | | | | **28.6** | K.VS*S*PSCDSQGLHAEEAPSSEPQSPVAQTSGWPACSTPLPLVSR.V |
| **BTB POZ domain containing 14B** | NACC1 | S124 | | | | | | | | **18.5** | K.VS*SPSCDSQGLHAEEAPSSEPQSPVAQTSGWPACSTPLPLVSR.V |
| **BTB POZ domain containing 14B** | NACC1 | S140S141 | | | | | | | **4.6** | **31.9** | K.VSSPSCDSQGLHAEEAPS*S*EPQSPVAQTSGWPACSTPLPLVSR.V |
| **BTKI** | IBTK | S1045 | | | | | | | **44.5** | **59.3** | R.DLQS*PDFTTGFHSDKIEAK.V |
| **BTKI** | IBTK | S1113S1116 | | | | | | | **9.0** | **24.5** | R.IDTTSSASWVAGSFS*PVS*PPVVDLR.T |
| **BTKI** | IBTK | Y996 | | | | | | | **6.5** | **19.2** | R.SDSSGGY*NLSDIIQSPSSTGLLK.S |
| **BTKI** | IBTK | S1111S1116 | | | | | | | **19.6** | **44.7** | R.IDTTSSASWVAGS*FSPVS*PPVVDLR.T |
| **BTKI** | IBTK | S992 | | | | | | | **5.1** | **22.6** | R.SDS*SGGYNLSDIIQSPSSTGLLK.S |
| **BTKI** | IBTK | S993 | | | | | | | **8.0** | **23.3** | R.SDSS*GGYNLSDIIQSPSSTGLLK.S |
| **BTKI** | IBTK | T1102S1116 | | | | | | | **2.4** | **12.1** | R.IDTT*SSASWVAGSFSPVS*PPVVDLR.T |
| **BUB1** | BUB1 | S602 | | | | | | | **0.9** | **83.2** | K.SPGDFTS*AAQLASTPFHK.L |
| **BUB1** | BUB1 | T601 | | | | | | | **32.1** | **81.5** | K.SPGDFT*SAAQLASTPFHK.L |

Peak Area: <-10, -5, -3, 0, 3, 5, >10

%CV: 0, 29, 42, 71, 86, >100

**White dots-** Significant change in peptide abundance at 5 %FDR compared to the timepoint with the minimum peak area for a given PSM

| Protein Name | Gene | Phosphosites | Ascor | MOWSE | Sequence |
|---|---|---|---|---|---|
| **BUB1** | BUB1 | S596 | 1.8 | 38.8 | K.S*PGDFTSAAQLASTPFHK.L |
| **Butyrate induced transcript 1** | HACD3 | S114 | 100.0 | 74.2 | R.WLDES*DAEMELR.A |
| **Bystin** | BYSL | S98 | 110.5 | 96.3 | R.MPQDGS*DDEDEEWPTLEK.A |
| **Bystin** | BYSL | S167S170 | 13.1 | 52.2 | K.QTEVETVMS*EVS*GFPMPQLDPR.V |
| **Bystin** | BYSL | S167 | 22.0 | 60.8 | K.QTEVETVMS*EVSGFPMPQLDPR.V |
| **Bystin** | BYSL | T164S170 | 7.0 | 15.0 | K.QTEVET*VMSEVS*GFPMPQLDPR.V |
| **Bystin** | BYSL | T164 | 12.8 | 23.7 | K.QTEVET*VMSEVSGFPMPQLDPR.V |
| **Bystin** | BYSL | T160 | 6.7 | 22.6 | K.QT*EVETVMSEVSGFPMPQLDPR.V |
| **c-Mpl binding protein** | LARP4 | T601 | 9.5 | 89.2 | R.AST*ASPCNNNINAATAVALQEPR.K |
| **c-Mpl binding protein** | LARP4 | S603 | 15.8 | 65.4 | R.ASTAS*PCNNNINAATAVALQEPR.K |
| **c-Mpl binding protein** | LARP4 | S589 | 37.7 | 38.4 | K.DGLNQTTIPVS*PPSTTKPSR.A |
| **c-Mpl binding protein** | LARP4 | S600 | 7.4 | 50.2 | R.AS*TASPCNNNINAATAVALQEPR.K |
| **c-Myb** | MYB | S11S12 | 13.7 | 29.1 | R.HSIYS*S*DEDDEDFEMCDHDYDGLLPK.S |
| **c-Myc** | MYC | T73S77 | 19.9 | 35.5 | K.KFELLPT*PPLS*PSRR.S |
| **c-Myc** | MYC | S86 | 22.3 | 53.4 | R.SGLCS*PSYVAVTPFSLR.G |
| **c-Myc** | MYC | T73S79 | 24.7 | 17.6 | K.FELLPT*PPLSPS*RR.S |
| **c-Src** | SRC | S17 | 125.2 | 55.9 | R.S*LEPAENVHGAGGGAFPASQTPSKPASADGHR.G |
| **C-X-C chemokine receptor type 5** | CXCR5 | S359 | 13.4 | 20.7 | R.RSS*LSESENATSLTTF.- |
| **C/EBP induced protein** | FAM117A | S213 | 100.0 | 71.6 | R.S*LEGLNQELEEVFVK.E |
| **C/EBP induced protein** | FAM117A | S305S317S319 | 8.7 | 12.2 | K.AS*SPGHPAFLEDGS*PS*PVLAFAAS*PRPNHSYIFK.R |
| **C10orf3 protein** | CEP55 | S428 | 14.0 | 115.0 | K.S*PTAALNESLVECPK.C |
| **C10orf3 protein** | CEP55 | T430 | 3.9 | 62.6 | K.SPT*AALNESLVECPK.C |
| **C10orf47 protein** | PROSER2 | S311 | 153.9 | 37.8 | R.AQVLATIHGHAGAFPAAGDAGEGAPGGGS*SPER.V |
| **C10orf69 protein** | ERLIN1 | S323 | 25.4 | 37.9 | R.ES*SLPSK.E |
| **C10orf69 protein** | ERLIN1 | S324 | 13.9 | 38.2 | R.ESS*LPSK.E |
| **C10orf86 protein** | NSMCE4A | S30S32 | 30.3 | 22.1 | R.S*RS*PLSPR.S |
| **C10orf86 protein** | NSMCE4A | S63S72 | 4.9 | 28.6 | R.EAPERPSLEDTEPSDS*GDEMMDPAS*LEAEADQGLCR.Q |
| **C10orf86 protein** | NSMCE4A | T58S61S72 | 5.1 | 33.6 | R.EAPERPSLEDT*EPS*DSGDEMMDPAS*LEAEADQGLCR.Q |
| **C11orf23 protein** | PPP6R3 | S537 | 100.0 | 131.5 | R.IQQFDDGGS*DEEDIWEEK.H |
| **C12orf5 protein** | TIGAR | S159 | 8.0 | 53.4 | K.EQFSQGSPS*NCLETSLAEIFPLGK.N |
| **C12orf5 protein** | TIGAR | S157 | 21.4 | 53.5 | K.EQFSQGS*PSNCLETSLAEIFPLGK.N |
| **C13orf3 protein** | SKA3 | S119 | 71.2 | 81.1 | K.NSVHEQEAINS*DPELSNCENFQK.T |
| **C13orf3 protein** | SKA3 | S155 | 67.1 | 66.1 | R.S*PQLSDFGLER.Y |
| **C13orf8 protein** | CHAMP1 | S476 | 100.0 | 49.3 | R.GGS*PDLWK.S |
| **C13orf8 protein** | CHAMP1 | S204S214 | 67.7 | 14.4 | K.LAPVPS*PEPQKPAPVS*PESVK.A |
| **C13orf8 protein** | CHAMP1 | S627 | 16.9 | 79.1 | K.DNQESS*DAELSSSEYIK.T |
| **C13orf8 protein** | CHAMP1 | S445S452 | 40.6 | 53.2 | R.KPSGS*PDLWKLS*PDQR.K |
| **C13orf8 protein** | CHAMP1 | S308S319 | 54.2 | 34.0 | R.RPAPAVS*PGSWKPGPPGS*PRPWK.S |
| **C13orf8 protein** | CHAMP1 | S286 | 29.7 | 68.2 | R.KPSPSES*PEPWKPFPAVSPEPR.R |
| **C13orf8 protein** | CHAMP1 | S282 | 18.4 | 52.0 | R.KPS*PSESPEPWKPFPAVSPEPR.R |
| **C13orf8 protein** | CHAMP1 | S282S297 | 56.8 | 29.6 | R.KPS*PSESPEPWKPFPAVS*PEPR.R |
| **C13orf8 protein** | CHAMP1 | S282S284 | 68.9 | 34.2 | R.KPS*PS*ESPEPWKPFPAVSPEPR.R |
| **C13orf8 protein** | CHAMP1 | S282S286 | 21.7 | 40.8 | R.KPS*PSES*PEPWKPFPAVSPEPR.R |
| **C13orf8 protein** | CHAMP1 | S652S653 | 7.6 | 67.1 | K.GQESS*S*DQEQVDVESIDFSK.E |
| **C13orf8 protein** | CHAMP1 | S651S653 | 49.6 | 130.4 | K.GQES*SS*DQEQVDVESIDFSK.E |
| **C13orf8 protein** | CHAMP1 | S432S436 | 100.0 | 41.1 | R.S*PAGS*PELR.K |
| **C13orf8 protein** | CHAMP1 | S443S452 | 63.3 | 45.3 | R.KPS*GSPDLWKLS*PDQR.K |
| **C13orf8 protein** | CHAMP1 | S284 | 20.3 | 70.1 | R.KPSPS*ESPEPWKPFPAVSPEPR.R |
| **C13orf8 protein** | CHAMP1 | S626 | 55.5 | 61.7 | K.DNQES*SDAELSSSEYIK.T |
| **C13orf8 protein** | CHAMP1 | S651S652 | 68.5 | 52.6 | K.GQES*S*SDQEQVDVESIDFSK.E |
| **C13orf8 protein** | CHAMP1 | S275 | 22.5 | 30.6 | R.TTS*PEPR.K |
| **C13orf8 protein** | CHAMP1 | S459 | 16.9 | 41.0 | R.KTS*PASLDFPESQK.S |
| **C13orf8 protein** | CHAMP1 | S445 | 5.9 | 38.5 | R.KPSGS*PDLWK.L |
| **C13orf8 protein** | CHAMP1 | S627S632 | 13.0 | 84.2 | K.DNQESS*DAELS*SSEYIK.T |
| **C13orf8 protein** | CHAMP1 | S427S432S436 | 100.0 | 13.0 | R.KPGPPLS*PEIRS*PAGS*PELR.K |
| **C13orf8 protein** | CHAMP1 | S284S286 | 20.1 | 34.7 | R.KPSPS*ES*PEPWKPFPAVSPEPR.R |
| **C13orf8 protein** | CHAMP1 | S282S286S297 | 24.8 | 19.4 | R.KPS*PSES*PEPWKPFPAVS*PEPR.R |
| **C13orf8 protein** | CHAMP1 | T274 | -0.2 | 28.9 | R.TT*SPEPR.K |
| **C13orf8 protein** | CHAMP1 | S627S633 | 8.7 | 79.5 | K.DNQESS*DAELSS*SEYIK.T |
| **C13orf8 protein** | CHAMP1 | S626S632 | 7.6 | 30.4 | K.DNQES*SDAELS*SSEYIK.T |
| **C13orf8 protein** | CHAMP1 | S311S319 | 24.9 | 29.0 | R.RPAPAVSPGS*WKPGPPGS*PRPWK.S |
| **C13orf8 protein** | CHAMP1 | S626S627 | 4.1 | 30.2 | K.DNQES*S*DAELSSSEYIK.T |

Peak Area: <-10, -5, -3, 0, 3, 5, >10

%CV: 0, 29, 42, 71, 86, >100

**White dots-** Significant change in peptide abundance at **5** %FDR compared to the timepoint with the minimum peak area for a given PSM

| Protein Name | Gene | Phosphosites | CarT 0m 2m 5m | RajiB 0m 2m 5m | Ascor | MOWSE | Sequence |
|---|---|---|---|---|---|---|---|
| **C13orf8 protein** | CHAMP1 | S286S297 | | | **49.6** | **26.6** | R.KPSPSES*PEPWKPFPAVS*PEPR.R |
| **C13orf8 protein** | CHAMP1 | S626S633 | | | **12.4** | **48.2** | K.DNQES*SDAELSS*SEYIK.T |
| **C14orf103 protein** | ATG2B | S1018T1022 | | | **7.4** | **27.0** | K.SAVHYDEESGS*EEET*LQYFSTVDPNYR.S |
| **C14orf103 protein** | ATG2B | S1016S1018 | | | **7.8** | **34.2** | K.SAVHYDEES*GS*EEETLQYFSTVDPNYR.S |
| **C14orf103 protein** | ATG2B | S1016T1022 | | | **13.5** | **25.8** | K.SAVHYDEES*GSEEET*LQYFSTVDPNYR.S |
| **C14orf106 protein** | MIS18BP1 | T821S824 | | | **27.6** | **27.5** | R.NTSNIPVILEPET*EES*ENEFYIK.Q |
| **C14orf11 protein** | EAPP | S109S111 | | | **48.4** | **67.9** | R.YYDDIYFDS*DS*EDEDR.A |
| **C14orf11 protein** | EAPP | Y106S111 | | | **21.1** | **28.7** | R.YYDDIY*FDSDS*EDEDR.A |
| **C14orf11 protein** | EAPP | Y106S109 | | | **9.7** | **62.9** | R.YYDDIY*FDS*DSEDEDR.A |
| **C14orf11 protein** | EAPP | S17S24S25S26 | | | **41.5** | **23.6** | R.LPDDYDPYAVEEPS*DEEPALS*S*S*EDEVDVLLHGTPDQK.R |
| | NGDN | S142S143 | | | **131.4** | **103.2** | K.LS*S*EDEEEDEAEDDQSEASGK.K |
| **C14orf133 protein** | VIPAS39 | S121 | | | **13.7** | **67.9** | R.TRPGS*FQSLSDALSDTPAK.S |
| **C14orf133 protein** | VIPAS39 | T117 | | | **22.6** | **67.1** | R.T*RPGSFQSLSDALSDTPAK.S |
| **C14orf133 protein** | VIPAS39 | S124 | | | **7.3** | **31.3** | R.TRPGSFQS*LSDALSDTPAK.S |
| **C14orf133 protein** | VIPAS39 | S93T104S105 | | | **8.2** | **12.8** | R.NS*FSSYAQLPKPT*S*TYSLSSFFR.G |
| **C14orf133 protein** | VIPAS39 | S95T104 | | | **13.1** | **16.7** | R.NSFS*SYAQLPKPT*STYSLSSFFR.G |
| **C14orf133 protein** | VIPAS39 | S93S95S96 | | | **15.2** | **16.4** | R.NS*FS*S*YAQLPKPTSTYSLSSFFR.G |
| **C14orf133 protein** | VIPAS39 | S93S105 | | | **4.4** | **14.1** | R.NS*FSSYAQLPKPTS*TYSLSSFFR.G |
| **C14orf133 protein** | VIPAS39 | S93S96Y97 | | | **12.9** | **13.0** | R.NS*FSS*Y*AQLPKPTSTYSLSSFFR.G |
| **C14orf133 protein** | VIPAS39 | S93T104 | | | **20.3** | **22.6** | R.NS*FSSYAQLPKPT*STYSLSSFFR.G |
| **C14orf133 protein** | VIPAS39 | S96T104S105 | | | **12.9** | **13.6** | R.NSFSS*YAQLPKPT*S*TYSLSSFFR.G |
| **C14orf171 protein** | TMEM63C | S77S80 | | | **17.4** | **46.6** | R.LALLIHNDS*LTS*LIYGEQSEK.T |
| **C14orf171 protein** | TMEM63C | S77T79 | | | **15.1** | **44.5** | R.LALLIHNDS*LT*SLIYGEQSEK.T |
| **C14orf35 protein** | NAA30 | S55 | | | **40.2** | **15.7** | R.S*PAGGESATVAAK.G |
| **C14orf4 protein** | IRF2BPL | S615S639 | | | **6.8** | **12.6** | R.TTPPES*APQNGPSPMAALMSVADTLGTAHS*PK.D |
| **C18orf25 protein** | C18orf25 | S66S69 | | | **29.4** | **31.7** | R.RDS*SES*QLASTESDKPTTGR.V |
| **C18orf25 protein** | C18orf25 | S67S69 | | | **8.6** | **11.4** | R.RDSS*ES*QLASTESDKPTTGR.V |
| **C19orf14 protein** | WDR62 | T1021S1022 | | | | **23.6** | R.FAT*S*LPHFPGCAGPTEDELSLPEGPSVPSSSLPQTPEQEK.F |
| **C19orf14 protein** | WDR62 | S1228 | | | **16.8** | **157.1** | R.SIS*LGDSEGPIVATLAQPLR.R |
| **C19orf14 protein** | WDR62 | S1249 | | | **5.8** | **79.9** | R.RPSS*VGELASLGQELQAITTATTPSLDSEGQEPALR.S |
| **C19orf14 protein** | WDR62 | S1248 | | | **29.8** | **55.1** | R.RPS*SVGELASLGQELQAITTATTPSLDSEGQEPALR.S |
| **C19orf14 protein** | WDR62 | S1048T1053 | | | **24.3** | **15.7** | R.FATSLPHFPGCAGPTEDELSLPEGPSVPSS*SLPQT*PEQEK.F |
| **C19orf14 protein** | WDR62 | S1248T1268 | | | **11.7** | **30.7** | R.RPS*SVGELASLGQELQAITTATT*PSLDSEGQEPALR.S |
| **C19orf14 protein** | WDR62 | S1249T1268 | | | **6.2** | **23.0** | R.RPSS*VGELASLGQELQAITTATT*PSLDSEGQEPALR.S |
| **C19orf14 protein** | WDR62 | S1255T1264 | | | **15.0** | **30.9** | R.RPSSVGELAS*LGQELQAIT*TATTPSLDSEGQEPALR.S |
| **C19orf14 protein** | WDR62 | S1248S1249 | | | **6.6** | **14.8** | R.RPS*S*VGELASLGQELQAITTATTPSLDSEGQEPALR.S |
| **C1orf16 protein** | SMG7 | S520 | | | **34.0** | **60.2** | K.ENLILQETSVIESLAADGS*PGLK.S |
| **C1orf24 protein** | FAM129A | S579T584 | | | **1.2** | **37.5** | K.HNLFEDNMALPSES*VSSLT*DLKPPTGSNQASPAR.R |
| **C1orf24 protein** | FAM129A | S577S581 | | | **2.9** | **37.7** | K.HNLFEDNMALPS*ESVS*SLTDLKPPTGSNQASPAR.R |
| **C1orf24 protein** | FAM129A | S577S592S596 | | | **7.9** | **24.5** | K.HNLFEDNMALPS*ESVSSLTDLKPPTGS*NQAS*PAR.R |
| **C1orf24 protein** | FAM129A | S579S582 | | | **4.9** | **70.2** | K.HNLFEDNMALPSES*VSS*LTDLKPPTGSNQASPAR.R |
| **C1orf24 protein** | FAM129A | S579S581 | | | **3.6** | **48.5** | K.HNLFEDNMALPSES*VS*SLTDLKPPTGSNQASPAR.R |
| **C1orf24 protein** | FAM129A | S581S592S596 | | | **10.2** | **30.8** | K.HNLFEDNMALPSESVS*SLTDLKPPTGS*NQAS*PAR.R |
| **C1orf24 protein** | FAM129A | S579T584T590 | | | **11.0** | **17.1** | K.HNLFEDNMALPSES*VSSLT*DLKPPT*GSNQAS*PAR.R |
| **C1orf24 protein** | FAM129A | S577S582 | | | **1.8** | **13.8** | K.KHNLFEDNMALPS*ESVSS*LTDLKPPTGSNQASPAR.R |
| **C1orf24 protein** | FAM129A | T584S592S596 | | | **9.9** | **45.6** | K.HNLFEDNMALPSESVSSLT*DLKPPTGS*NQAS*PAR.R |
| **C1orf24 protein** | FAM129A | S579T590S596 | | | **15.9** | **25.5** | K.HNLFEDNMALPSES*VSSLTDLKPPT*GSNQAS*PAR.R |
| **C1orf24 protein** | FAM129A | T584T590S596 | | | **4.2** | **14.3** | K.HNLFEDNMALPSESVSSLT*DLKPPT*GSNQAS*PAR.R |
| **C1orf24 protein** | FAM129A | S582T584T590 | | | **14.5** | **17.8** | K.HNLFEDNMALPSESVSS*LT*DLKPPT*GSNQAS*PAR.R |
| **C1orf24 protein** | FAM129A | S581S582 | | | **1.9** | **25.2** | K.HNLFEDNMALPSESVS*S*LTDLKPPTGSNQASPAR.R |
| **C1orf24 protein** | FAM129A | S577S579 | | | **5.7** | **46.1** | K.HNLFEDNMALPS*ES*VSSLTDLKPPTGSNQASPAR.R |
| **C1orf24 protein** | FAM129A | S579T590 | | | **2.7** | **15.8** | K.HNLFEDNMALPSES*VSSLTDLKPPT*GSNQASPAR.R |
| **C1orf24 protein** | FAM129A | S579S592S596 | | | **12.5** | **34.3** | K.HNLFEDNMALPSES*VSSLTDLKPPTGS*NQAS*PAR.R |
| **C1orf24 protein** | FAM129A | S579S581S592 | | | **2.0** | **17.2** | K.HNLFEDNMALPSES*VS*SLTDLKPPTGS*NQAS*PAR.R |
| **C1orf24 protein** | FAM129A | S581T584T590 | | | **2.1** | **29.5** | K.HNLFEDNMALPSESVS*SLT*DLKPPT*GSNQAS*PAR.R |
| **C1orf24 protein** | FAM129A | S577S579T590 | | | **1.2** | **14.7** | K.HNLFEDNMALPS*ES*VSSLTDLKPPT*GSNQAS*PAR.R |
| **C1orf24 protein** | FAM129A | S582T584 | | | **3.6** | **30.7** | K.HNLFEDNMALPSESVSS*LT*DLKPPTGSNQASPAR.R |
| **C1orf24 protein** | FAM129A | S577T584T590 | | | **16.8** | **19.2** | K.HNLFEDNMALPS*ESVSSLT*DLKPPT*GSNQAS*PAR.R |
| **C1orf24 protein** | FAM129A | S577T584S592 | | | **5.9** | **15.9** | K.HNLFEDNMALPS*ESVSSLT*DLKPPTGS*NQAS*PAR.R |
| **C1orf35 protein** | MMTAG2 | S217T219 | | | **-0.4** | **13.2** | R.RPAEATSS*PT*SPERPR.H |
| **C1orf35 protein** | MMTAG2 | S216T219 | | | **7.7** | **13.6** | R.RPAEATS*SPT*SPERPR.H |
| **C1orf52 protein** | C1orf52 | S158 | | | **4.3** | **65.7** | R.LLPEGEETLES*DDEKDEHTSK.K |

Peak Area: <-10, -5, -3, 0, 3, 5, >10

%CV: 0, 29, 42, 71, 86, >100

**White dots-** Significant change in peptide abundance at **5** %FDR compared to the timepoint with the minimum peak area for a given PSM

| Protein Name | Gene | Phosphosites | CarT (0m 2m 5m) | RajiB (0m 2m 5m) | Ascor | MOWSE | Sequence |
|---|---|---|---|---|---|---|---|
| **C1orf52 protein** | C1orf52 | T155 | | | 50.1 | 50.4 | R.LLPEGEET*LESDDEKDEHTSK.K |
| **C20ORF1 protein** | TPX2 | S486 | | | 100.0 | 39.3 | K.S*PAFALK.N |
| **C20ORF1 protein** | TPX2 | S738 | | | 32.7 | 35.0 | K.SSDQPLTVPVS*PK.F |
| **C20ORF121 protein** | TTPAL | S13S16 | | | 3.8 | 10.9 | R.TSPS*VAS*LSENELPPPPEPPGYVCSLTEDLVTK.A |
| **C20ORF121 protein** | TTPAL | S11S16 | | | 2.6 | 13.5 | R.TS*PSVAS*LSENELPPPPEPPGYVCSLTEDLVTK.A |
| **C20orf14 protein** | PRPF6 | T275S279 | | | 24.4 | 52.1 | K.GYLT*DLNS*MIPTHGGDINDIKK.A |
| **C20orf14 protein** | PRPF6 | T205T212T227T | | | 9.0 | 18.1 | R.QT*QFGGLNT*PYPGGLNTPYPGGMT*PGLMT*PGTGELDMR.K |
| **C20orf14 protein** | PRPF6 | T275T283 | | | 4.4 | 22.1 | K.GYLT*DLNSMIPT*HGGDINDIKK.A |
| **C20orf6 protein** | ESF1 | S153 | | | 18.8 | 49.4 | K.FKIDSNIS*PK.K |
| **C20orf6 protein** | ESF1 | S663 | | | 95.5 | 77.6 | K.ALAEEASEEELPS*DVDLNDPYFAEEVK.Q |
| **C20orf6 protein** | ESF1 | S657 | | | 40.8 | 63.0 | K.ALAEEAS*EEELPSDVDLNDPYFAEEVK.Q |
| **C20orf6 protein** | ESF1 | S657S663 | | | 98.3 | 79.4 | K.ALAEEAS*EEELPS*DVDLNDPYFAEEVK.Q |
| **C20orf6 protein** | ESF1 | T311S312T319 | | | 108.5 | 63.7 | R.GKGNIET*S*SEDEDDT*ADLFPEESGFEHAWR.E |
| **C20orf6 protein** | ESF1 | S312S313T319 | | | 6.9 | 49.0 | R.GKGNIETS*S*EDEDDT*ADLFPEESGFEHAWR.E |
| **C20orf6 protein** | ESF1 | T311S313T319 | | | 21.7 | 30.5 | K.GNIET*SS*EDEDDT*ADLFPEESGFEHAWR.E |
| **C20orf6 protein** | ESF1 | S75S77S79S82 | | | 21.9 | 51.1 | R.FYDLS*DS*DS*NLS*GEDSK.A |
| **C20orf6 protein** | ESF1 | T311S312S313 | | | 15.9 | 23.2 | R.GKGNIET*S*S*EDEDDTADLFPEESGFEHAWR.E |
| **C20orf6 protein** | ESF1 | T311T319 | | | | 43.9 | R.GKGNIET*SSEDEDDT*ADLFPEESGFEHAWR.E |
| **C20ORF77 protein** | RPRD1B | S164 | | | 42.4 | 76.2 | R.TFQQIQEEEDDDYPGS*YSPQDPSAGPLLTEELIK.A |
| **C20ORF77 protein** | RPRD1B | Y161 | | | 6.6 | 46.7 | R.TFQQIQEEEDDDY*PGSYSPQDPSAGPLLTEELIK.A |
| **C20ORF77 protein** | RPRD1B | Y165 | | | 13.9 | 83.5 | R.TFQQIQEEEDDDYPGSY*SPQDPSAGPLLTEELIK.A |
| **C20ORF77 protein** | RPRD1B | T149 | | | | 56.1 | R.T*FQQIQEEEDDDYPGSYSPQDPSAGPLLTEELIK.A |
| **C20ORF77 protein** | RPRD1B | S166 | | | | 51.8 | R.TFQQIQEEEDDDYPGSYS*PQDPSAGPLLTEELIK.A |
| **C21orf66 protein** | PAXBP1 | S557T563 | | | 13.5 | 83.5 | K.M#ADHLEGLS*SDDEET*STDITNFNLEK.D |
| **C21orf66 protein** | PAXBP1 | S557S558 | | | 22.5 | 97.5 | K.MADHLEGLS*S*DDEETSTDITNFNLEK.D |
| **C21orf66 protein** | PAXBP1 | S558T563 | | | 11.6 | 24.9 | K.MADHLEGLSS*DDEET*STDITNFNLEK.D |
| **C22orf5 protein** | TMEM184 | S402S403 | | | 37.8 | 20.8 | K.TLLLS*S*DDEF.- |
| **C22orf9 protein** | KIAA0930 | S309 | | | 35.2 | 63.4 | R.NNRPAFFS*PSLKR.K |
| **C22orf9 protein** | KIAA0930 | S367 | | | 40.6 | 39.6 | R.SLVGS*WLK.L |
| **C4orf9 protein** | NOP14 | T155S157 | | | 25.5 | 26.0 | K.HNDIVDSDSDAEDRGT*LS*AELTAAHFGGGGGLLHK.K |
| **C4orf9 protein** | NOP14 | S146T155 | | | 6.5 | 12.2 | K.HNDIVDS*DSDAEDRGT*LSAELTAAHFGGGGGLLHK.K |
| **C4orf9 protein** | NOP14 | S146T161 | | | 5.9 | 16.2 | K.HNDIVDS*DSDAEDRGTLSAELT*AAHFGGGGGLLHK.K |
| **C4orf9 protein** | NOP14 | S148T155 | | | 4.3 | 13.1 | K.HNDIVDSDS*DAEDRGT*LSAELTAAHFGGGGGLLHK.K |
| | NOP14 | S146S148 | | | | 35.3 | K.HNDIVDS*DS*DAEDRGTLSAELTAAHFGGGGGLLHKK.T |
| **C4orf9 protein** | NOP14 | S148S157 | | | 1.5 | 14.1 | K.HNDIVDSDS*DAEDRGTLS*AELTAAHFGGGGGLLHK.K |
| **C5orf3 protein** | FAM114A2 | S145 | | | 37.0 | 66.1 | K.ENENS*SPVAGAFGVFSTISTAVQSTGK.S |
| **C5orf6 protein** | FAM53C | S232S236 | | | 15.7 | 18.5 | R.RFS*LSPS*LGPQASR.F |
| **C5orf6 protein** | FAM53C | S232S234 | | | 31.3 | 54.2 | R.RFS*LS*PSLGPQASR.F |
| **C6orf106 protein** | C6orf106 | S209 | | | 9.1 | 14.9 | R.LSQNSVNLSPSS*HANNLSVVTYSK.G |
| **C6orf106 protein** | C6orf106 | S206 | | | 20.0 | 72.8 | R.LSQNSVNLS*PSSHANNLSVVTYSK.G |
| **C6orf106 protein** | C6orf106 | S202 | | | 1.2 | 14.2 | R.LSQNS*VNLSPSSHANNLSVVTYSK.G |
| **C6orf111 protein** | PNISR | S290S304T309 | | | 24.3 | 58.0 | R.SKFDS*DEEEEDTENVEAAS*SGKVT*R.S |
| **C6orf111 protein** | PNISR | T297S304T309 | | | 8.1 | 13.5 | R.SKFDSDEEEEDT*ENVEAAS*SGKVT*R.S |
| **C6orf111 protein** | PNISR | S290 | | | 18.8 | 61.5 | R.SKFDS*DEEEEDTENVEAASSGK.V |
| **C6orf111 protein** | PNISR | S290T309 | | | 11.8 | 55.2 | R.SKFDS*DEEEEDTENVEAASSGKVT*R.S |
| **C6orf111 protein** | PNISR | S290S305 | | | 2.6 | 15.6 | R.SKFDS*DEEEEDTENVEAASS*GKVTR.S |
| **C6orf111 protein** | PNISR | S211 | | | 100.0 | 44.5 | R.S*PIALPVK.Q |
| **C6orf111 protein** | PNISR | S393S396 | | | 4.3 | 27.3 | K.QLAQSSALASLTGLGGLGGYGS*GDS*EDER.S |
| **C6orf111 protein** | PNISR | S611S613 | | | 100.0 | 16.2 | R.NRS*PS*RER.R |
| **C6orf111 protein** | PNISR | S672 | | | 100.0 | 35.9 | R.S*IDKDR.K |
| **C6orf111 protein** | PNISR | S601 | | | 15.7 | 14.5 | R.SNRNS*IER.E |
| **C6orf111 protein** | PNISR | S726 | | | 25.8 | 57.5 | R.SGS*ISVK.I |
| **C6orf111 protein** | PNISR | S670S672 | | | 100.0 | 22.9 | R.S*RS*IDKDR.K |
| **C6orf111 protein** | PNISR | T383S396 | | | 4.0 | 47.8 | K.QLAQSSALASLT*GLGGLGGYGSGDS*EDER.S |
| **C6orf111 protein** | PNISR | S290S304 | | | 11.7 | 26.9 | R.SKFDS*DEEEEDTENVEAAS*SGK.V |
| **C6orf111 protein** | PNISR | Y391S396 | | | 23.8 | 77.8 | K.QLAQSSALASLTGLGGLGGY*GSGDS*EDER.S |
| **C6orf111 protein** | PNISR | S311S313T326 | | | 12.8 | 11.3 | R.S*PS*PVPQEEHSDPEMT*EEEKEYQMMLLTK.M |
| **C6orf111 protein** | PNISR | S290S304S305 | | | 25.9 | 47.6 | R.SKFDS*DEEEEDTENVEAAS*S*GKVT*R.S |
| | PNISR | S311S313S321 | | | 73.6 | 29.8 | R.S*PS*PVPQEEHS*DPEMTEEEKEYQMMLLTK.M |
| **C6orf111 protein** | PNISR | S724 | | | 20.3 | 47.3 | R.S*GSISVK.I |
| **C6orf113 protein** | ZUFSP | S253 | | | 20.2 | 16.1 | R.RSEES*R.Q |
| **C6orf32 protein** | RIPOR2 | T522 | | | 10.8 | 59.9 | R.LT*SAEVPMATDR.L |

**Peak Area**: <-10, -5, -3, 0, 3, 5, >10

**%CV**: 0, 29, 42, 71, 86, >100

**White dots-** Significant change in peptide abundance at **5** %FDR compared to the timepoint with the minimum peak area for a given PSM

CarT (0m, 2m, 5m) — RajiB (0m, 2m, 5m)

| Protein Name | Gene | Phosphosites | Ascor | MOWSE | Sequence |
|---|---|---|---|---|---|
| **C6orf32 protein** | RIPOR2 | S473 | | **35.9** | R. RQS*SCACAEHLELENDVAEALLQESEEASELKPVELDTSEGNITK.Q |
| **C6orf32 protein** | RIPOR2 | S19 | **18.0** | **61.4** | R.S*QSFAGFSGLQER.R |
| **C6orf32 protein** | RIPOR2 | S21 | **33.7** | **62.6** | R.SQS*FAGFSGLQER.R |
| **C6orf32 protein** | RIPOR2 | S535S538 | **45.1** | **64.1** | R.LLS*EGS*VGGESEGCR.S |
| **C6orf32 protein** | RIPOR2 | S344T350 | **15.7** | **16.0** | R.RMSMYS*QGTPET*PTFK.D |
| **C6orf32 protein** | RIPOR2 | S523 | **13.9** | **19.8** | R.LTS*AEVPMATDR.L |
| **C6orf32 protein** | RIPOR2 | Y343T350 | **12.2** | **23.1** | R.RMSMY*SQGTPET*PTFK.D |
| **C7orf25 protein** | C7orf25 | S208S210S212 | **65.4** | **18.3** | R.GDIVAVNALLDHPEELQPS*ES*ES*DDEGPELLQVTR.V |
| **C8orf20 protein** | REEP4 | S152 | **15.1** | **43.0** | R.SFS*MQDLR.S |
| **C8orf20 protein** | REEP4 | S194T196S202 | **21.8** | **13.5** | R.AGGLQDS*DT*EDECWS*DTEAVPR.A |
| **C8orf20 protein** | REEP4 | S150 | **10.5** | **33.3** | R.S*FSMQDLR.S |
| **C9orf25 protein** | FAM219A | T96S98 | **27.6** | **33.2** | K.GYSSLDQSPDEKPLVALDT*DS*DDDFDMSR.Y |
| **C9orf42 protein** | FAM122A | S143S147 | **21.9** | **52.3** | K.RIDFIPVS*PAPS*PTR.G |
| **C9orf42 protein** | FAM122A | T47 | **34.7** | **56.4** | R.SNSAPLIHGLSDT*SPVFQAEAPSAR.R |
| **C9orf42 protein** | FAM122A | S37 | **18.8** | **62.2** | R.SNS*APLIHGLSDTSPVFQAEAPSAR.R |
| **C9orf42 protein** | FAM122A | S45 | **16.9** | **51.9** | R.SNSAPLIHGLS*DTSPVFQAEAPSAR.R |
| **C9orf42 protein** | FAM122A | S143T149 | **20.9** | **48.2** | K.RIDFIPVS*PAPSPT*R.G |
| **C9orf42 protein** | FAM122A | S270 | **48.0** | **56.1** | K.VSTTTDSPVS*PAQAASPFIPLDELSSK.- |
| **C9orf42 protein** | FAM122A | S276 | **1.8** | **19.3** | K.VSTTTDSPVSPAQAAS*PFIPLDELSSK.- |
| **C9orf42 protein** | FAM122A | T263S270 | **4.2** | **46.9** | K.VST*TTDSPVS*PAQAASPFIPLDELSSK.- |
| **C9orf42 protein** | FAM122A | S267S270 | **-3.1** | **27.1** | K.VSTTTDS*PVS*PAQAASPFIPLDELSSK.- |
| **C9orf42 protein** | FAM122A | S76 | **100.0** | **36.4** | R.HGLLLPAS*PVR.M |
| **C9orf42 protein** | FAM122A | S189 | **-0.2** | **56.3** | R.SQS*PINCIRPSVLGPLK.R |
| **C9orf42 protein** | FAM122A | S35 | **26.2** | **49.4** | R.S*NSAPLIHGLSDTSPVFQAEAPSAR.R |
| **C9orf42 protein** | FAM122A | S270S276 | **41.4** | **34.0** | K.VSTTTDSPVS*PAQAAS*PFIPLDELSSK.- |
| **C9orf42 protein** | FAM122A | S267S276 | **-5.3** | **21.1** | K.VSTTTDS*PVSPAQAAS*PFIPLDELSSK.- |
| **C9orf42 protein** | FAM122A | S62 | **16.9** | **45.0** | R.RNS*TTFPSR.H |
| **C9orf42 protein** | FAM122A | S262S270 | **5.5** | **19.8** | K.VS*TTTDSPVS*PAQAASPFIPLDELSSK.- |
| **C9orf55 protein** | DENND4C | S1089 | **16.9** | **42.1** | R.STS*LSALVR.S |
| **C9orf55 protein** | DENND4C | S767S769 | **27.6** | **55.6** | R.KSSTGS*IS*NVLFSTQDPVEDAVFGEATNLKK.N |
| **C9orf55 protein** | DENND4C | S890 | **-0.3** | **35.7** | R.SS*PVPEMLEESQELLEPVVDDVPK.T |
| **C9orf55 protein** | DENND4C | S899 | **1.8** | **21.9** | R.SSPVPEMLEES*QELLEPVVDDVPK.T |
| **C9orf55 protein** | DENND4C | S769S774T775 | **8.1** | **35.6** | R.KSSTGSIS*NVLFS*T*QDPVEDAVFGEATNLK.K |
| **C9orf55 protein** | DENND4C | S764T765 | **8.8** | **57.6** | R.KSS*T*GSISNVLFSTQDPVEDAVFGEATNLKK.N |
| **C9orf55 protein** | DENND4C | S1404T1415 | **17.8** | **41.9** | R.SHS*VGGPLQNIDFT*QRPFHGISTVSLPNSLQEVVDPLGK.R |
| | DENND4C | S763S764T765 | **5.4** | **35.4** | R.KS*S*T*GSISNVLFSTQDPVEDAVFGEATNLKK.N |
| **C9orf55 protein** | DENND4C | S763S764S774 | **2.8** | **28.7** | R.KS*S*TGSISNVLFS*TQDPVEDAVFGEATNLKK.N |
| **C9orf55 protein** | DENND4C | S1404S1423S1 | **16.4** | **23.0** | R.SHS*VGGPLQNIDFTQRPFHGIS*TVSLPNS*LQEVVDPLGK.R |
| **C9orf55 protein** | DENND4C | S1404T1415T14 | **10.9** | **36.4** | R.SHS*VGGPLQNIDFT*QRPFHGIST*VSLPNSLQEVVDPLGK.R |
| **C9orf55 protein** | DENND4C | S763S769 | **16.5** | **64.5** | R.KS*STGSIS*NVLFSTQDPVEDAVFGEATNLK.K |
| **C9orf55 protein** | DENND4C | S763S764 | | **25.9** | R.KS*S*TGSISNVLFSTQDPVEDAVFGEATNLKK.N |
| **C9orf55 protein** | DENND4C | T765S769 | **6.6** | **15.8** | R.KSST*GSIS*NVLFSTQDPVEDAVFGEATNLKK.N |
| **C9orf55 protein** | DENND4C | T1088 | **-0.4** | **30.6** | R.ST*SLSALVR.S |
| **C9orf55 protein** | DENND4C | T765S769S774 | **9.0** | **31.7** | R.KSST*GSIS*NVLFS*TQDPVEDAVFGEATNLKK.N |
| **C9orf55 protein** | DENND4C | S774T775 | **28.3** | **31.0** | R.KSSTGSISNVLFS*T*QDPVEDAVFGEATNLKK.N |
| **C9orf55 protein** | DENND4C | S763S767 | **2.0** | **41.5** | R.KS*STGS*ISNVLFSTQDPVEDAVFGEATNLK.K |
| **C9orf55 protein** | DENND4C | T765S767S769 | **9.5** | **25.5** | R.KSST*GS*IS*NVLFSTQDPVEDAVFGEATNLK.K |
| **C9orf55 protein** | DENND4C | S732S737 | **24.9** | **20.0** | K.HSQPS*PEPHS*PTEPPAWGSSIVK.V |
| **C9orf55 protein** | DENND4C | S1404S1423 | **25.3** | **48.8** | R.SHS*VGGPLQNIDFTQRPFHGIS*TVSLPNSLQEVVDPLGK.R |
| **C9orf55 protein** | DENND4C | S1404T1424S1 | **10.3** | **48.3** | R.SHS*VGGPLQNIDFTQRPFHGIST*VS*LPNSLQEVVDPLGK.R |
| **C9orf55 protein** | DENND4C | T765S767 | **4.3** | **26.3** | R.KSST*GS*ISNVLFSTQDPVEDAVFGEATNLKK.N |
| **C9orf55 protein** | DENND4C | S764S769 | **9.3** | **57.2** | R.KSS*TGSIS*NVLFSTQDPVEDAVFGEATNLK.K |
| **C9orf55 protein** | DENND4C | T765S767S774 | **8.4** | **17.4** | R.KSST*GS*ISNVLFS*TQDPVEDAVFGEATNLK.K |
| **C9orf55 protein** | DENND4C | S1404S1426 | **9.8** | **42.7** | R.SHS*VGGPLQNIDFTQRPFHGISTVS*LPNSLQEVVDPLGK.R |
| **C9orf78** | C9orf78 | S261 | **92.1** | **23.6** | R.VGDTEKPEPERS*PPNR.K |
| **C9orf78** | C9orf78 | S15S17 | **68.5** | **89.7** | R.RRGDS*ES*EEDEQDSEEVR.L |
| **C9orf88 protein** | FAM129B | S691S696 | **24.0** | **37.3** | K.AAPEAS*SPPAS*PLQHLLPGK.A |
| **C9orf88 protein** | FAM129B | S641S646 | **8.7** | **48.1** | K.QVVSVVQDEEVGLPFEASPES*PPPAS*PDGVTEIR.G |
| **C9orf88 protein** | FAM129B | S692S696 | **26.1** | **33.5** | K.AAPEASS*PPAS*PLQHLLPGK.A |
| **Calcipressin 1** | RCAN1 | S163S167 | **100.0** | **70.3** | K.QFLIS*PPAS*PPVGWK.Q |
| **Calcium channel, voltage dependent, P/Q type, alpha 1A subunit** | CACNA1A | S2457 | **6.6** | **14.8** | R.S*PRTPR.A |
| **Calcium regulated heat stable protein 1** | CARHSP1 | S52 | **33.6** | **40.6** | R.TFS*ATVR.A |

Peak Area: <-10, -5, -3, 0, 3, 5, >10

%CV: 0, 29, 42, 71, 86, >100

**White dots-** Significant change in peptide abundance at **5** %FDR compared to the timepoint with the minimum peak area for a given PSM

| Protein Name | Gene | Phosphosites | CarT 0m | CarT 2m | CarT 5m | RajiB 0m | RajiB 2m | RajiB 5m | Ascor | MOWSE | Sequence |
|---|---|---|---|---|---|---|---|---|---|---|---|
| Calcium regulated heat stable protein 1 | CARHSP1 | S41 | | | | | | | 41.9 | 24.4 | R.GNVVPS*PLPTRR.T |
| Calcium regulated heat stable protein 1 | CARHSP1 | S146 | | | | | | | -0.4 | 28.0 | K.HETWSGHVIS*S.- |
| Calcium regulated heat stable protein 1 | CARHSP1 | S30S32S41 | | | | | | | 45.0 | 22.7 | R.ERS*PS*PLRGNVVPS*PLPTR.R |
| Calcium regulated heat stable protein 1 | CARHSP1 | S30S32 | | | | | | | 100.0 | 31.8 | R.ERS*PS*PLR.G |
| Calcium regulated heat stable protein 1 | CARHSP1 | T50 | | | | | | | 3.5 | 13.8 | R.TRT*FSATVR.A |
| Calcium regulated heat stable protein 1 | CARHSP1 | S30S41 | | | | | | | 23.1 | 27.5 | R.ERS*PSPLRGNVVPS*PLPTR.R |
| Calcium/calmodulin dependent protein kinase IV | CAMK4 | S360 | | | | | | | 100.0 | 47.9 | R.DPS*PIQDGNEDMK.A |
| Calmodulin regulated spectrin-associated protein 1 | CAMSAP1 | S447 | | | | | | | 16.3 | 74.3 | K.SPNS*HDSEPWTLLR.Q |
| Calmodulin regulated spectrin-associated protein 1 | CAMSAP1 | S297 | | | | | | | 39.0 | 34.5 | R.S*PQGQLDTSESKPDSFFLEPLMPAVLKPAK.E |
| Calmodulin regulated spectrin-associated protein 1 | CAMSAP1 | S311 | | | | | | | -2.7 | 36.1 | R.SPQGQLDTSESKPDS*FFLEPLMPAVLKPAK.E |
| Calmodulin regulated spectrin-associated protein 1 | CAMSAP1 | S307 | | | | | | | 8.2 | 45.1 | R.SPQGQLDTSES*KPDSFFLEPLMPAVLKPAK.E |
| Calmodulin regulated spectrin-associated protein 1 | CAMSAP1 | S444 | | | | | | | 15.0 | 80.1 | K.S*PNSHDSEPWTLLR.Q |
| Calmodulin regulated spectrin-associated protein 1 | CAMSAP1 | S450 | | | | | | | 0.2 | 29.1 | K.SPNSHDS*EPWTLLR.Q |
| Calmodulin regulated spectrin-associated protein 1 | CAMSAP1 | S460S462 | | | | | | | 183.2 | 72.5 | R.QDS*DS*DVVDIEEAEHDFMGEAHPVVFSR.Y |
| Calmodulin regulated spectrin-associated protein 1-like 1 | CAMSAP2 | S404 | | | | | | | -0.1 | 21.7 | R.SS*SMSYVDGFIGTWPK.E |
| Calnexin | CANX | S583 | | | | | | | 100.0 | 49.3 | K.AEEDEILNRS*PR.N |
| Calnexin | CANX | T562S564 | | | | | | | 15.5 | 66.7 | K.QKSDAEEDGGT*VS*QEEEDRKPK.A |
| | CANX | S554 | | | | | | | 28.6 | 66.4 | K.QKS*DAEEDGGTVSQEEEDR.K |
| Calnexin | CANX | S554T562 | | | | | | | 8.3 | 56.4 | K.QKS*DAEEDGGT*VSQEEEDRKPK.A |
| Calnexin | CANX | T562 | | | | | | | 6.5 | 37.7 | K.QKSDAEEDGGT*VSQEEEDR.K |
| Calnexin | CANX | S564 | | | | | | | 17.8 | 65.8 | K.SDAEEDGGTVS*QEEEDR.K |
| Calpastatin | CAST | S326 | | | | | | | 25.9 | 68.9 | K.KEGITGPPADSSKPIGPDDAIDALSSDFTCGS*PTAAGK.K |
| Calpastatin | CAST | T299 | | | | | | | | 63.0 | K.KEGIT*GPPADSSKPIGPDDAIDALSSDFTCGSPTAAGK.K |
| Calpastatin | CAST | S449 | | | | | | | -0.2 | 55.6 | R.SES*ELIDELSEDFDR.S |
| Calpastatin | CAST | T323 | | | | | | | 5.6 | 13.4 | K.EGITGPPADSSKPIGPDDAIDALSSDFT*CGSPTAAGK.K |
| CAP-GLY domain containing linker protein 1 | CLIP1 | S348 | | | | | | | 27.7 | 98.3 | R.KIS*GTTALQEALK.E |
| CAP-GLY domain containing linker protein 1 | CLIP1 | S147 | | | | | | | 1.4 | 77.2 | R.ATS*PLCTSTASMVSSSPSTPSNIPQKPSQPAAK.E |
| CAP-GLY domain containing linker protein 1 | CLIP1 | S195S200S204 | | | | | | | 20.3 | 55.8 | K.TASES*ISNLS*EAGS*IK.K |
| CAP-GLY domain containing linker protein 1 | CLIP1 | S200S204 | | | | | | | 26.7 | 67.7 | K.TASESISNLS*EAGS*IK.K |
| CAP-GLY domain containing linker protein 1 | CLIP1 | S147S160 | | | | | | | -0.5 | 18.3 | R.ATS*PLCTSTASMVSSS*PSTPSNIPQKPSQPAAK.E |
| CAP-GLY domain containing linker protein 1 | CLIP1 | S147S158 | | | | | | | 3.6 | 20.5 | R.ATS*PLCTSTASMVS*SSPSTPSNIPQKPSQPAAK.E |
| CAP-GLY domain containing linker protein 1 | CLIP1 | S197S204 | | | | | | | 19.3 | 42.0 | K.TASESIS*NLSEAGS*IK.K |
| CAP-GLY domain containing linker protein 1 | CLIP1 | S195S197S204 | | | | | | | 2.8 | 24.2 | K.TASES*IS*NLSEAGS*IK.K |
| CapZ-interacting protein | RCSD1 | S75 | | | | | | | 75.7 | 69.2 | K.LQANLTFDPAALLPGAS*PK.S |
| Carboxypeptidase D | CPD | T1368T1370 | | | | | | | 27.6 | 63.7 | K.SLLSHEFQDET*DT*EEETLYSSKH.- |
| Cardiac lineage protein 1 | HEXIM1 | T236S237S252 | | | | | | | 1.9 | 35.3 | K.SDDT*S*DDDFMEEGGEEDGGS*DGMGGDGSEFLQR.D |
| Cask-interacting protein 2 | CASKIN2 | S471 | | | | | | | 102.6 | 44.1 | R.S*GEQIFTQDVRPEQLLEGK.D |
| Cask-interacting protein 2 | CASKIN2 | T252 | | | | | | | 10.5 | 73.0 | R.NT*YNQTALDIVNQFTTSQASR.E |
| Cask-interacting protein 2 | CASKIN2 | S358 | | | | | | | 11.1 | 28.9 | R.LPS*APTPLRPGFSR.T |
| Cask-interacting protein 2 | CASKIN2 | Y253 | | | | | | | 16.9 | 68.8 | R.NTY*NQTALDIVNQFTTSQASR.E |
| CasL interacting molecule | MICAL1 | S872S875S876 | | | | | | | 66.5 | 63.9 | K.EEKES*PFS*S*EEEEEDVPLDSDVEQALQTFAK.T |
| CasL interacting molecule | MICAL1 | S876 | | | | | | | 5.3 | 75.0 | K.ESPFSS*EEEEEDVPLDSDVEQALQTFAK.T |
| CasL interacting molecule | MICAL1 | S875 | | | | | | | 12.3 | 81.1 | K.ESPFS*SEEEEEDVPLDSDVEQALQTFAK.T |
| CasL interacting molecule | MICAL1 | S875S876 | | | | | | | 9.4 | 86.6 | K.ESPFS*S*EEEEEDVPLDSDVEQALQTFAK.T |
| CasL interacting molecule | MICAL1 | S872S875 | | | | | | | 29.7 | 56.5 | K.EEKES*PFS*SEEEEEDVPLDSDVEQALQTFAK.T |
| Caspase 2 | CASP2 | S157 | | | | | | | 39.9 | 48.3 | K.LRLS*TDTVEHSLDNK.D |
| Caspase 2 | CASP2 | T160 | | | | | | | 4.5 | 32.2 | K.LRLSTDT*VEHSLDNK.D |
| CASPR4 | CNTNAP4 | S520 | | | | | | | 22.4 | 23.9 | R.LIS*ISGK.V |
| Catalase | CAT | S515 | | | | | | | 1.9 | 30.5 | K.NAIHTFVQS*GSHLAAR.E |
| Catenin alpha 1 | CTNNA1 | S641 | | | | | | | 49.2 | 74.8 | R.TPEELDDS*DFETEDFDVR.S |
| CBF1 interacting corepressor | CIR1 | S202 | | | | | | | 28.3 | 104.9 | R.NLTANDPSQEYVAS*EGEEDPEVEFLK.S |
| CBF1 interacting corepressor | CIR1 | Y199 | | | | | | | 9.2 | 42.3 | R.NLTANDPSQEY*VASEGEEDPEVEFLK.S |
| CBFA2 | RUNX1 | T41S48 | | | | | | | 41.4 | 45.1 | R.RFT*PPSTALS*PGK.M |
| CBFA2 | RUNX1 | S276S303 | | | | | | | 22.9 | 41.4 | R.QIQPS*PPWSYDQSYQYLGSIASPSVHPATPIS*PGR.A |
| CBFA2 | RUNX1 | S48 | | | | | | | 55.8 | 67.8 | R.RFTPPSTALS*PGK.M |
| CBFA2 | RUNX1 | S276S280 | | | | | | | | 19.0 | R.QIQPS*PPWS*YDQSYQYLGSIASPSVHPATPISPGR.A |
| CBFA2 | RUNX1 | S239 | | | | | | | 7.7 | 18.5 | R.TAMRVS*PHHPAPTPNPR.A |
| CBFA2 | RUNX1 | S303 | | | | | | | 11.4 | 26.8 | R.QIQPSPPWSYDQSYQYLGSIASPSVHPATPIS*PGR.A |
| CBL | CBL | S619 | | | | | | | 60.9 | 28.3 | R.HS*LPFSLPSQMEPRPDVPR.L |
| CBP80 | NCBP1 | S22 | | | | | | | 19.1 | 66.1 | R.KTS*DANETEDHLESLICK.V |
| CBP80 | NCBP1 | T27 | | | | | | | 11.9 | 28.8 | R.RKTSDANET*EDHLESLICK.V |
| CC chemokine receptor type 7 | CCR7 | S367 | | | | | | | 17.8 | 26.5 | R.RSSMS*VEAETTTTFSP.- |

Peak Area: <-10, -5, -3, 0, 3, 5, >10

%CV: 0, 29, 42, 71, 86, >100

**White dots-** Significant change in peptide abundance at **5** %FDR compared to the timepoint with the minimum peak area for a given PSM

CarT: 0m 2m 5m — RajiB: 0m 2m 5m

| Protein Name | Gene | Phosphosites | Ascor | MOWSE | Sequence |
|---|---|---|---|---|---|
| **CC chemokine receptor type 7** | CCR7 | S364 | **32.4** | **17.9** | R.RS*SMSVEAETTTTFSP.- |
| **CC chemokine receptor type 7** | CCR7 | S365 | **14.7** | **35.7** | R.RSS*MSVEAETTTTFSP.- |
| **CCAAT binding factor** | CEBPZ | S629 | **34.9** | **83.9** | R.SQLDDHPES*DDEENFIDANDDEDMEK.F |
| **CCAAT binding factor** | CEBPZ | T33 | **29.5** | **56.8** | K.RPWRPEEAVEDPDEEDEDNT*SEAENGFSLEEVLR.L |
| **CCAAT binding factor** | CEBPZ | S34 | **38.4** | **83.8** | K.RPWRPEEAVEDPDEEDEDNTS*EAENGFSLEEVLR.L |
| **CCAAT binding factor** | CEBPZ | S973 | **24.6** | **54.8** | R.NLNDS*SLFVSAEEFGHLLDENMGSK.F |
| **CCAAT/Enhancer binding protein, beta** | CEBPB | S229 | **12.1** | **26.7** | R.AYLGYQAVPSGSSGSLSTSSS*SSPPGTPSPADAK.A |
| **CCAAT/Enhancer binding protein, beta** | CEBPB | S228T235 | **10.9** | **15.5** | R.AYLGYQAVPSGSSGSLSTSS*SSSPPGT*PSPADAK.A |
| **CCAAT/Enhancer binding protein, beta** | CEBPB | S218T235 | **9.0** | **23.2** | R.AYLGYQAVPS*GSSGSLSTSSSSSPPGT*PSPADAK.A |
| | CTCF | S609S610S612 | | **18.4** | K.EDS*S*DS*ENAEPDLDDNEDEEEPAVEIEPEPEPQPVTPAPPPAK.K |
| **CCR4-NOT transcription complex, subunit 3** | CNOT3 | S473 | | **41.9** | K.[illegible] |
| **CCR4-NOT transcription complex, subunit 3** | CNOT3 | S506 | **14.6** | **34.0** | K.[illegible] |
| **CCR4-NOT transcription complex, subunit 3** | CNOT3 | S358 | | **13.1** | K.[illegible] |
| **CCR4-NOT transcription complex, subunit 3** | CNOT3 | S507 | **5.3** | **47.8** | K.[illegible] |
| **CD19** | CD19 | S499 | **13.9** | **106.5** | R.EATSLGSQS*YEDMR.G |
| **CD19** | CD19 | S494S499 | **13.2** | **80.5** | R.EATS*LGSQS*YEDMR.G |
| **CD19** | CD19 | T493S497 | **27.3** | **68.6** | R.EAT*SLGS*QSYEDMR.G |
| **CD19** | CD19 | Y531 | **-0.3** | **65.7** | R.GQPGPNHEEDADSY*ENMDNPDGPDPAWGGGGR.M |
| **CD19** | CD19 | T493S494S499 | **6.0** | **53.2** | R.EAT*S*LGSQS*YEDMR.G |
| **CD19** | CD19 | S530 | | **59.0** | R.GQPGPNHEEDADS*YENMDNPDGPDPAWGGGGR.M |
| **CD19** | CD19 | T493S497S499 | **11.3** | **56.2** | R.EAT*SLGS*QS*YEDMR.G |
| **CD19** | CD19 | S494S497 | **12.8** | **61.8** | R.EATS*LGS*QSYEDMR.G |
| **CD19** | CD19 | S494S497S499 | **9.7** | **55.8** | R.EATS*LGS*QS*YEDMR.G |
| **CD19** | CD19 | T493S499 | **20.8** | **78.3** | R.EAT*SLGSQS*YEDMR.G |
| **CD2 antigen binding protein 2** | CD2BP2 | S195 | **9.8** | **41.4** | R.KGPGQPSS*PQR.L |
| | CD2BP2 | S194 | | **11.3** | R.KGPGQPS*SPQRLDR.L |
| **CD2 antigen binding protein 2** | CD2BP2 | S49 | **36.0** | **124.2** | K.HSLDS*DEEEDDDDGGSSK.Y |
| **CD2 associated protein** | CD2AP | S233 | **7.9** | **51.8** | R.TSS*SETEEKKPEKPLILQSLGPK.T |
| **CD2 associated protein** | CD2AP | S458 | **78.4** | **69.9** | K.S*VDFDSLTVR.T |
| **CD2 associated protein** | CD2AP | S86 | **25.8** | **45.6** | R.IS*TYGLPAGGIQPHPQTK.N |
| **CD2 associated protein** | CD2AP | T87 | **12.6** | **50.0** | R.IST*YGLPAGGIQPHPQTK.N |
| **CD2 associated protein** | CD2AP | Y88 | **12.9** | **47.1** | R.ISTY*GLPAGGIQPHPQTK.N |
| **CD28** | | S189 | **7.9** | **22.4** | R.LLHS*DYMNMTPR.R |
| **CD28** | | Y191 | **11.1** | **26.5** | R.LLHSDY*MNMTPR.R |
| **CD3 gamma** | CD3G | T152 | **1.3** | **39.0** | R.ASDKQT*LLPNDQLYQPLK.D |
| **CD3 gamma** | CD3G | S148 | **4.4** | **22.2** | R.AS*DKQTLLPNDQLYQPLK.D |
| **CD40** | CD40 | S272 | **100.0** | **37.7** | R.IS*VQER.Q |
| **CD5** | CD5 | S428 | **14.9** | **89.0** | R.QWIGPTGMNQNMS*FHR.N |
| **CD5** | CD5 | S482S483S485 | **16.6** | **34.6** | R.SSMQPDNS*S*DS*DYDLHGAQR.L |
| **CD5** | CD5 | T421 | **2.6** | **84.9** | R.QWIGPT*GMNQNMSFHR.N |
| **CD5** | CD5 | S439 | **31.3** | **35.5** | R.S*HAENPTASHVDNEYSQPPR.N |
| **CD5** | CD5 | S476S482 | **3.3** | **50.9** | R.SS*MQPDNS*SDSDYDLHGAQR.L |
| **CD5** | CD5 | S476S482S483 | **2.8** | **34.0** | R.SS*MQPDNS*S*DS*DYDLHGAQR.L |
| **CD7** | CD7 | S228 | **10.7** | **52.6** | K.NSAACVVYEDMSHS*R.C |
| **CD7** | CD7 | S226 | **-0.4** | **50.1** | K.NSAACVVYEDMS*HSR.C |
| **CD74 antigen** | CD74 | S25 | **100.0** | **58.3** | R.DLIS*NNEQLPMLGR.R |
| **CD80 antigen** | CD80 | S284 | **100.0** | **20.5** | R.LRRES*VRPV.- |
| **CDC 25B** | CDC25B | S396 | **61.1** | **53.4** | K.S*LCHDEIENLLDSDHR.E |
| **CDC 25B** | CDC25B | S344 | **22.3** | **29.1** | R.SPS*MPCSVIRPILK.R |
| **CDC like kinase 3** | CDCA5 | S27 | **23.9** | **21.0** | R.RSYS*R.L |
| **CDC like kinase 3** | CDCA5 | S25Y26 | **13.9** | **17.8** | R.RS*Y*SR.L |
| **CDC like kinase 3** | CLK3 | S49S51 | **30.5** | **13.0** | R.S*RS*HDRLPYQR.R |
| **CDC2** | CDK1 | Y15 | **16.9** | **61.0** | K.IGEGTY*GVVYK.A |
| **CDC2** | CDK1 | T14Y15 | **50.2** | **60.6** | K.IGEGT*Y*GVVYK.A |
| **CDC2 related protein kinase 7** | CDK12 | S274S276 | **46.0** | **38.2** | R.RQS*VS*PPYKEPSAYQSSTR.S |
| **CDC2 related protein kinase 7** | CDK12 | S333S334 | **8.3** | **32.2** | R.SS*S*PFLSK.R |
| **CDC2 related protein kinase 7** | CDK12 | S382S385 | **21.3** | **43.5** | R.HS*SIS*PVRLPLNSSLGAELSR.K |
| **CDC2 related protein kinase 7** | CDK12 | S383S385 | **35.8** | **48.9** | R.HSS*IS*PVRLPLNSSLGAELSR.K |
| **CDC2 related protein kinase 7** | CDK12 | S681S685 | **126.6** | **39.6** | R.HLLTDLPLPPELPGGDLS*PPDS*PEPK.A |
| **CDC2 related protein kinase 7** | CDK12 | S301S303 | **38.3** | **25.5** | R.S*VS*PYSR.R |
| **CDC2 related protein kinase 7** | CDK12 | Y892 | **5.8** | **23.0** | R.LYNSEESRPY*TNK.V |
| **CDC2 related protein kinase 7** | CDK12 | S379S382S383 | **39.9** | **19.0** | R.S*RHS*S*IS*PVRLPLNSSLGAELSR.K |

Peak Area: <-10, -5, -3, 0, 3, 5, >10

%CV: 0, 29, 42, 71, 86, >100

**White dots-** Significant change in peptide abundance at **5** %FDR compared to the timepoint with the minimum peak area for a given PSM

CarT: 0m 2m 5m — RajiB: 0m 2m 5m

| Protein Name | Gene | Phosphosites | Ascor | MOWSE | Sequence |
|---|---|---|---|---|---|
| **CDC2 related protein kinase 7** | CDK12 | S77S78S80S84 | **7.9** | **26.2** | K.DMGLVTPEAASLGTVIKPLVEYDDIS*S*DS*DTFS*DDMAFK.L |
| **CDC2 related protein kinase 7** | CDK12 | Y73S78T82S84 | **5.6** | **15.2** | K.DMGLVTPEAASLGTVIKPLVEY*DDISS*DSDT*FS*DDMAFK.L |
| **CDC2 related protein kinase 7** | CDK12 | T57S62T65Y73 | **4.1** | **16.4** | K.DMGLVT*PEAAS*LGT*VIKPLVEY*DDISSDSDTFSDDMAFK.L |
| **CDC2 related protein kinase 7** | CDK12 | S332S333 | **10.0** | **39.0** | R.S*S*SPFLSK.R |
| **CDC2 related protein kinase 7** | CDK12 | S62T65S77S84 | **2.0** | **22.1** | K.DMGLVTPEAAS*LGT*VIKPLVEYDDIS*SDSDTFS*DDMAFK.L |
| **CDC2 related protein kinase 7** | CDK12 | S382S383 | **17.9** | **43.7** | R.HS*S*ISPVRLPLNSSLGAELSR.K |
| **CDC2 related protein kinase 7** | CDK12 | S77S78T82S84 | **6.6** | **11.4** | K.DMGLVTPEAASLGTVIKPLVEYDDIS*S*DSDT*FS*DDMAFK.L |
| **CDC2 related protein kinase 7** | CDK12 | Y73S78S80T82 | **4.4** | **23.0** | K.DMGLVTPEAASLGTVIKPLVEY*DDISS*DS*DT*FSDDMAFK.L |
| **CDC2 related protein kinase 7** | CDK12 | S1083 | **11.2** | **61.4** | K.NSS*PAPPQPAPGK.V |
| **CDC2 related protein kinase 7** | CDK12 | S383S393 | **12.2** | **29.3** | R.HSS*ISPVRLPLNS*SLGAELSR.K |
| **CDC2 related protein kinase 7** | CDK12 | S681 | **40.0** | **35.8** | R.HLLTDLPLPPELPGGDLS*PPDSPEPK.A |
| **CDC2 related protein kinase 7** | CDK12 | T893 | **12.1** | **32.0** | R.LYNSEESRPYT*NK.V |
| **CDC2 related protein kinase 7** | CDK12 | S276Y279 | **5.3** | **12.2** | R.RQSVS*PPY*KEPSAYQSSTR.S |
| **CDC2 related protein kinase 7** | CDK12 | Y73S78S80S84 | **7.1** | **22.7** | K.DMGLVTPEAASLGTVIKPLVEY*DDISS*DS*DTFS*DDMAFK.L |
| **CDC2 related protein kinase 7** | CDK12 | S62T65Y73S84 | **3.6** | **24.2** | K.DMGLVTPEAAS*LGT*VIKPLVEY*DDISSDSDTFS*DDMAFK.L |
| **CDC25C** | CDC25C | S263 | **6.6** | **64.7** | K.TVS*LCDITITQMLEEDSNQGHLIGDFSK.V |
| **CDC26** | CDC26 | S42 | **48.1** | **47.6** | K.QKEDVEVVGGS*DGEGAIGLSSDPK.S |
| **CDC42 effector protein 4** | CDC42EP | S9 | **20.1** | **15.4** | K.QLVS*SSVHSK.R |
| **CDC5** | CDC5L | T373T377T396 | **18.1** | **12.6** | R.ILQEAQNLMALT*NVDT*PLKGGLNTPLHESDFSGVT*PQR.Q |
| **CDCA5** | CDCA5 | S35 | **-0.2** | **36.2** | K.SGS*ELPSILPEIWPK.T |
| **Cdk5 and Abl enzyme substrate 2** | CABLES2 | S105 | **5.5** | **38.5** | R.TPAPQGLLS*PTQVPTGLGLDGQR.Q |
| **CDP-DAG synthase 2** | CDS2 | S33 | **49.2** | **65.4** | K.VDGETAS*DSESR.A |
| **Cell death regulator Aven** | AVEN | S94 | **6.9** | **76.2** | R.REPGGWGAGASAPVEDDS*DAETYGEENDEQGNYSK.R |
| **Cell death regulator Aven** | AVEN | T98 | **6.5** | **11.6** | R.REPGGWGAGASAPVEDDSDAET*YGEENDEQGNYSK.R |
| **Cell division cycle 2 like 1** | CDK11B | Y67 | **9.5** | **11.3** | -.MEITIRNSPY*R.R |
| **Cell division cycle 2 like 1** | CDK11B | T595 | **16.9** | **55.6** | K.AYT*PVVVTLWYR.A |
| **Cell division cycle 2 like 1** | CDK11B | S589T595 | **16.0** | **88.3** | R.EYGS*PLKAYT*PVVVTLWYR.A |
| **Cell division cycle 2 like 1** | CDK11B | Y587S589 | **24.9** | **67.4** | R.EY*GS*PLKAYTPVVVTLWYR.A |
| **Cell division cycle 2 like 1** | CDK11B | Y594T595 | **24.0** | **60.4** | R.EYGSPLKAY*T*PVVVTLWYR.A |
| **Cell division cycle 2 like 1** | CDK11B | Y587Y594 | **10.6** | **57.7** | R.EY*GSPLKAY*TPVVVTLWYR.A |
| **Cell division cycle 2 like 1** | CDK11B | S589Y594 | **14.5** | **34.4** | R.EYGS*PLKAY*TPVVVTLWYR.A |
| **Cell division cycle 2 like 1** | CDK11B | Y587T595 | **12.3** | **59.2** | R.EY*GSPLKAYT*PVVVTLWYR.A |
| **Cell division cycle 2 like 1** | CDK11B | S65 | **16.1** | **20.5** | -.MEITIRNS*PYR.R |
| **Cell division cycle 2 like 2** | CDK11B | S574 | **26.0** | **35.4** | R.EYGS*PLK.A |
| **Cell division cycle 2 like 2** | CDK11B | T736S737 | **73.0** | **26.9** | K.RGT*S*PRPPEGGLGYSQLGDDDLK.E |
| **Cell division cycle 2 like 2** | CDK11A | T580 | **16.8** | **55.4** | K.AYT*PVVVTQWYR.A |
| **Cell division cycle 2 like 2** | CDK11B | S271 | **16.8** | **59.3** | R.DLLSDLQDIS*DSER.K |
| **Cell division cycle 2 like 2** | CDK11A | Y579T580 | **16.6** | **76.2** | R.EYGSPLKAY*T*PVVVTQWYR.A |
| **Cell division cycle 2 like 2** | CDK11A | Y572Y579 | **17.4** | **61.4** | R.EY*GSPLKAY*TPVVVTQWYR.A |
| **Cell division cycle 2 like 2** | CDK11B | S265S271 | **38.0** | **89.0** | R.DLLS*DLQDIS*DSER.K |
| **Cell division cycle 2 like 2** | CDK11B | S363S366T377 | | **14.7** | R. [illegible] |
| **Cell division cycle 2 like 2** | CDK11B | S222 | **100.0** | **23.5** | R.S*PPRPPR.L |
| **Cell division cycle 2 like 2** | CDK11A | S574T580 | **11.2** | **74.9** | R.EYGS*PLKAYT*PVVVTQWYR.A |
| **Cell division cycle 2 like 2** | CDK11A | Y572T580 | **12.0** | **61.9** | R.EY*GSPLKAYT*PVVVTQWYR.A |
| **Cell division cycle 2 like 2** | CDK11A | Y572S574 | **13.5** | **20.6** | R.EY*GS*PLKAYTPVVVTQWYR.A |
| **Cell division cycle 2 like 5** | CDK13 | T1056T1058 | **0.3** | **39.1** | R.KDLSLGLDDSRT*NT*PQGVLPSSQLK.S |
| **Cell division cycle 2 like 5** | CDK13 | S437S439 | **20.0** | **57.7** | R.HSS*IS*PSTLTLK.S |
| **Cell division cycle 2 like 5** | CDK13 | S1054T1056 | **5.9** | **32.7** | R.KDLSLGLDDS*RT*NTPQGVLPSSQLK.S |
| **Cell division cycle 2 like 5** | CDK13 | S436S437 | **8.5** | **42.5** | R.HS*S*ISPSTLTLK.S |
| **Cell division cycle 2 like 5** | CDK13 | S315S317 | **100.0** | **29.8** | R.S*LS*PLGGR.D |
| **Cell division cycle 2 like 5** | CDK13 | S1054T1058 | **6.8** | **35.5** | R.KDLSLGLDDS*RTNT*PQGVLPSSQLK.S |
| **Cell division cycle 2 like 5** | CDK13 | S315S317S325 | **31.4** | **13.4** | R.S*LS*PLGGRDDS*PVSHR.A |
| **Cell division cycle 2 like 5** | CDK13 | S383 | **32.1** | **11.4** | R.GGDVS*PSPYSSSSWR.R |
| **Cell division cycle 2 like 5** | CDK13 | S436S439 | **13.4** | **45.5** | R.HS*SIS*PSTLTLK.S |
| **Cell division cycle and apoptosis regulator 1** | CCAR1 | S333 | **100.0** | **20.6** | R.ERS*PQR.K |
| **Centaurin beta 1** | ACAP1 | T739 | **46.2** | **33.9** | R.SHDLHT*L.- |
| **Centaurin beta 5** | ACAP3 | T454S456 | | **11.4** | R.KPT*AS*SSRQDK.E |
| **Centaurin delta 2** | ARAP1 | S229 | **39.3** | **91.2** | R.LFPEFDDS*DYDEVPEEGPGAPAR.V |
| **Centaurin delta 2** | ARAP1 | Y231 | **22.6** | **19.6** | R.LFPEFDDSDY*DEVPEEGPGAPAR.V |
| **Centaurin delta 2** | ARAP1 | S1195 | **81.4** | **71.2** | R.S*VAAFTADPLSLLR.N |
| **Centromeric protein a** | CENPA | S17S19 | **27.5** | **32.7** | R.RRS*PS*PTPTPGPSR.R |
| **Centromeric protein c1** | CENPC | S316 | **52.5** | **33.3** | R.S*WITIPR.K |

Peak Area: <-10, -5, -3, 0, 3, 5, >10

%CV: 0, 29, 42, 71, 86, >100

**White dots-** Significant change in peptide abundance at **5** %FDR compared to the timepoint with the minimum peak area for a given PSM

CarT: 0m 2m 5m — RajiB: 0m 2m 5m

| Protein Name | Gene | Phosphosites | Ascor | MOWSE | Sequence |
|---|---|---|---|---|---|
| **Centromeric protein E** | CENPE | S1259 | **17.8** | **22.5** | R.RSVS*EK.T |
| **Centromeric protein E** | CENPE | S1257 | **9.5** | **16.9** | R.RS*VSEK.T |
| **Centromeric protein F** | CENPF | S1651 | **8.9** | **29.9** | R.LQLQGLDLS*SR.S |
| **Centrosomal protein 72kDa** | CEP72 | S237 | **60.5** | **27.6** | R.HLLS*PQLVQYQCGDSGK.Q |
| **Centrosome protein 4** | CEP135 | S1125S1132 | **4.5** | **20.4** | R.HGLATPPLS*STLRSPS*HSPEHR.N |
| **Centrosome protein 4** | CEP135 | S439 | **9.6** | **14.6** | R.DRS*PSRLDTFLK.G |
| **Centrosome protein 4** | ELOA2 | S1132S1134 | **28.7** | **40.3** | R.SPS*HS*PEHR.N |
| **Centrosome protein 4** | CEP135 | S1125S1126 | **0.8** | **13.2** | R.HGLATPPLS*S*TLRSPSHSPEHR.N |
| **Centrosome protein 4** | CEP135 | T1127S1130 | **2.2** | **12.5** | R.HGLATPPLSST*LRS*PSHSPEHR.N |
| **Centrosome protein 4** | CEP135 | S439S441 | **100.0** | **14.7** | R.RDRS*PS*R.L |
| **Centrosome protein 4** | CEP135 | S441 | **7.9** | **13.3** | R.DRSPS*RLDTFLK.G |
| | CEP135 | S1130S1132 | **5.1** | **13.7** | R.HGLATPPLSSTLRS*PS*HSPEHR.N |
| **Centrosome protein 4** | CEP135 | T1127S1132 | **-0.2** | **16.6** | R.HGLATPPLSST*LRSPS*HSPEHR.N |
| **Ceramide synthase 4** | CERS4 | S291S296S298 | **100.0** | **55.3** | K.DIRS*DVEES*DS*S*EEAAAAQEPLQLK.N |
| **Ceramide synthase 4** | CERS4 | S291S296S298 | **14.7** | **37.9** | R.S*DVEES*DS*SEEAAAAQEPLQLK.N |
| **Ceramide synthase 4** | CERS4 | S291S299 | **24.7** | **18.4** | K.DIRS*DVEESDSS*EEAAAAQEPLQLK.N |
| **Ceramide synthase 4** | CERS4 | S291S296S299 | **8.5** | **42.6** | R.S*DVEES*DSS*EEAAAAQEPLQLK.N |
| **Ceramide synthase 4** | CERS4 | S291S298 | **14.4** | **13.2** | K.DIRS*DVEESDS*SEEAAAAQEPLQLK.N |
| **Ceramide synthase 4** | CERS4 | S291S296 | **14.1** | **11.7** | K.DIRS*DVEES*DSSEEAAAAQEPLQLK.N |
| **Ceramide synthase 4** | CERS4 | S291S298S299 | **14.6** | **52.5** | K.DIRS*DVEESDS*S*EEAAAAQEPLQLK.N |
| **Cerebellar degeneration-related autoantigen 2** | CDR2 | S311 | **10.9** | **53.4** | R.SSS*ETILSSLAGSDIVK.G |
| **Cerebellar degeneration-related autoantigen 2** | CDR2 | S310 | **-0.2** | **45.7** | R.SS*SETILSSLAGSDIVK.G |
| **Cerebral protein 11** | TMCC3 | S438 | **100.0** | **64.7** | R.NKFGS*ADNIAHLK.D |
| **CGI-07 protein** | NMD3 | S468T470 | **48.4** | **38.5** | R.DSAIPVES*DT*DDEGAPR.I |
| **CGI-07 protein** | NMD3 | T470 | **9.1** | **59.4** | R.DSAIPVESDT*DDEGAPR.I |
| **CGI-79 protein** | RBMX2 | S187 | **13.9** | **11.5** | R.EVQAEQPSSS*SPR.R |
| **CGI-79 protein** | RBMX2 | S188 | **9.8** | **24.6** | R.EVQAEQPSSSS*PR.R |
| **CGI115 protein** | RRP15 | S266S276S280 | **8.9** | **21.7** | K.DWDKES*DGPDDSRPES*ASDS*DT*.- |
| **CGI115 protein** | RRP15 | S272S276S280 | **0.6** | **18.1** | K.DWDKESDGPDDS*RPES*ASDS*DT*.- |
| **Chemokine, CXC motif, receptor 4** | CXCR4 | S321S324 | **4.0** | **14.5** | K.TSAQHALTSVS*RGS*SLK.I |
| **Chemokine, CXC motif, receptor 4** | CXCR4 | T318S319S324 | **12.1** | **15.5** | K.TSAQHALT*S*VSRGS*SLK.I |
| **Chemokine, CXC motif, receptor 4** | CXCR4 | S347 | **5.8** | **46.9** | K.RGGHSSVSTESESS*SFHSS.- |
| **Chemokine, CXC motif, receptor 4** | CXCR4 | S324S325 | **31.3** | **32.1** | K.TSAQHALTSVSRGS*S*LK.I |
| **Chemokine, CXC motif, receptor 4** | CXCR4 | T311S312 | **2.1** | **12.4** | K.T*S*AQHALTSVSRGSSLK.I |
| **CHERP** | CHERP | S813S815S817 | **47.5** | **83.2** | R.S*RS*RS*PT*PPSSAGLGSNSAPPIPDSR.L |
| **CHERP** | CHERP | S815S817T819 | **11.2** | **61.5** | R.S*RS*PT*PPSSAGLGSNSAPPIPDSR.L |
| **CHERP** | CHERP | S815S817S823 | **10.1** | **45.0** | R.S*RS*PTPPSS*AGLGSNSAPPIPDSR.L |
| **CHERP** | CHERP | T819S822S823 | **13.8** | **50.6** | R.SRSPT*PPS*S*AGLGSNSAPPIPDSR.L |
| **CHERP** | CHERP | S815T819S823 | **7.6** | **49.6** | R.S*RSPT*PPSS*AGLGSNSAPPIPDSR.L |
| **CHERP** | CHERP | S802S804S806 | **15.2** | **14.4** | R.S*KS*YS*PGRR.R |
| **CHERP** | CHERP | S815T819S822 | **9.1** | **37.7** | R.S*RSPT*PPS*SAGLGSNSAPPIPDSR.L |
| **CHERP** | CHERP | S817T819S823 | **9.3** | **36.3** | R.SRS*PT*PPSS*AGLGSNSAPPIPDSR.L |
| **CHERP** | CHERP | S815T819S822 | **7.7** | **17.0** | R.S*RSPT*PPS*S*AGLGSNSAPPIPDSR.L |
| **CHERP** | CHERP | S813S815S817 | **9.3** | **10.7** | R.S*RS*RS*PT*PPS*SAGLGSNSAPPIPDSR.L |
| **CHERP** | CHERP | S815S817T819 | **10.1** | **27.4** | R.S*RS*PT*PPS*SAGLGSNSAPPIPDSR.L |
| **CHERP** | CHERP | S813S815S817 | **11.8** | **22.1** | R.S*RS*RS*PTPPS*SAGLGSNSAPPIPDSR.L |
| **CHERP** | CHERP | S813S815S817 | **6.5** | **36.4** | R.S*RS*RS*PTPPSS*AGLGSNSAPPIPDSR.L |
| **CHERP** | CHERP | S817T819S822 | **2.0** | **47.9** | R.SRS*PT*PPS*SAGLGSNSAPPIPDSR.L |
| **CHERP** | CHERP | S815S817S822 | **2.6** | **61.0** | R.S*RS*PTPPS*SAGLGSNSAPPIPDSR.L |
| **CHL12** | CHTF18 | S871 | **65.8** | **102.3** | R.VENSPQVDGS*PPGLEGLLGGIGEK.G |
| **CHMP family, member 7** | CHMP7 | S417 | **100.0** | **80.7** | R.IS*DAELEAELEK.L |
| **Chondrocyte protein with a poly proline region** | MTFR1 | S119 | **84.9** | **68.5** | R.QIS*LPDLSQEEPQLK.T |
| **Chondroitin sulfate proteoglycan 6** | SMC3 | S1081 | **11.5** | **39.4** | R.GS*GSQSSVPSVDQFTGVGIR.V |
| **Chr2 synaptotagmin** | ESYT2 | S665 | **2.8** | **35.2** | K.SHMSGS*PGPGGSNTAPSTPVIGGSDKPGMEEK.A |
| **Chr2 synaptotagmin** | ESYT2 | T656S660S663 | **5.0** | **33.5** | R.KT*SIKS*HMS*GSPGPGGSNTAPSTPVIGGSDKPGMEEK.A |
| **Chr2 synaptotagmin** | ESYT2 | T656S657S660 | | **59.6** | R.KT*S*IKS*HMSGSPGPGGSNTAPSTPVIGGSDKPGMEEK.A |
| **Chr2 synaptotagmin** | ESYT2 | S710 | **23.5** | **91.8** | R.SSS*SLLASPGHISVK.E |
| **Chr2 synaptotagmin** | ESYT2 | S708 | **11.6** | **46.5** | R.S*SSSLLASPGHISVK.E |
| **Chr2 synaptotagmin** | ESYT2 | S710S711 | **23.3** | **66.1** | R.SSS*S*LLASPGHISVK.E |
| **Chr2 synaptotagmin** | ESYT2 | S733 | **9.1** | **36.0** | K.EPTPSIASDIS*LPIATQELR.Q |
| **Chr2 synaptotagmin** | ESYT2 | S730 | **37.7** | **118.8** | K.EPTPSIAS*DISLPIATQELR.Q |
| **Chr2 synaptotagmin** | ESYT2 | S730S733 | **9.1** | **72.1** | K.EPTPSIAS*DIS*LPIATQELR.Q |

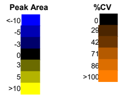


**White dots-** Significant change in peptide abundance at **5** %FDR compared to the timepoint with the minimum peak area for a given PSM

| Protein Name | Gene | Phosphosites | CarT 0m | CarT 2m | CarT 5m | RajiB 0m | RajiB 2m | RajiB 5m | Ascor | MOWSE | Sequence |
|---|---|---|---|---|---|---|---|---|---|---|---|
| Chr2 synaptotagmin | ESYT2 | S727S730 | | | | | | | 7.7 | 62.1 | K.EPTPS*IAS*DISLPIATQELR.Q |
| Chr2 synaptotagmin | ESYT2 | T725S730 | | | | | | | 9.9 | 43.1 | K.EPT*PSIAS*DISLPIATQELR.Q |
| Chr2 synaptotagmin | ESYT2 | S727S730S733 | | | | | | | 23.9 | 117.1 | K.EPTPS*IAS*DIS*LPIATQELR.Q |
| Chr2 synaptotagmin | ESYT2 | T725S727S730 | | | | | | | 9.8 | 37.9 | K.EPT*PS*IAS*DISLPIATQELR.Q |
| Chr2 synaptotagmin | ESYT2 | S648 | | | | | | | 27.7 | 29.7 | K.RPS*VSK.E |
| Chr2 synaptotagmin | ESYT2 | T656S660S665 | | | | | | | 5.1 | 58.6 | R.KT*SIKS*HMSGS*PGPGGSNTAPSTPVIGGSDKPGMEEK.A |
| Chr2 synaptotagmin | ESYT2 | T656S660S663 | | | | | | | 4.6 | 28.9 | R.KT*SIKS*HMS*GS*PGPGGSNTAPSTPVIGGSDKPGMEEK.A |
| Chr2 synaptotagmin | ESYT2 | T656S657S660 | | | | | | | | 27.8 | R.KT*S*IKS*HMS*GSPGPGGSNTAPSTPVIGGSDKPGMEEK.A |
| Chr2 synaptotagmin | ESYT2 | S711 | | | | | | | 12.6 | 60.8 | R.SSSS*LLASPGHISVK.E |
| Chr2 synaptotagmin | ESYT2 | S709S711 | | | | | | | 11.8 | 76.8 | R.SS*SS*LLASPGHISVK.E |
| Chr2 synaptotagmin | ESYT2 | S708S709 | | | | | | | 12.3 | 44.1 | R.S*S*SSLLASPGHISVK.E |
| Chr2 synaptotagmin | ESYT2 | S727S733 | | | | | | | 6.2 | 64.3 | K.EPTPS*IASDIS*LPIATQELR.Q |
| Chr2 synaptotagmin | ESYT2 | T725S727S733 | | | | | | | 6.6 | 65.0 | K.EPT*PS*IASDIS*LPIATQELR.Q |
| Chr2 synaptotagmin | ESYT2 | S709 | | | | | | | -0.3 | 61.8 | R.SS*SSLLASPGHISVK.E |
| Chr2 synaptotagmin | ESYT2 | S657S660S665 | | | | | | | 3.5 | 37.7 | R.KTS*IKS*HMSGS*PGPGGSNTAPSTPVIGGSDKPGMEEK.A |
| Chr2 synaptotagmin | ESYT2 | T725S733 | | | | | | | 12.3 | 35.9 | K.EPT*PSIASDIS*LPIATQELR.Q |
| Chr2 synaptotagmin | ESYT2 | S657S660S663 | | | | | | | 3.5 | 46.4 | R.KTS*IKS*HMS*GSPGPGGSNTAPSTPVIGGSDKPGMEEK.A |
| Chr2 synaptotagmin | ESYT2 | S663S665 | | | | | | | 17.6 | 43.1 | K.SHMS*GS*PGPGGSNTAPSTPVIGGSDKPGMEEK.A |
| Chr2 synaptotagmin | ESYT2 | S657S660S663 | | | | | | | 31.8 | 26.8 | R.KTS*IKS*HMS*GS*PGPGGSNTAPSTPVIGGSDKPGMEEK.A |
| Chr2 synaptotagmin | ESYT2 | T725 | | | | | | | 2.8 | 19.4 | K.EPT*PSIASDISLPIATQELR.Q |
| Chr2 synaptotagmin | ESYT2 | T725S727 | | | | | | | 9.0 | 48.3 | K.EPT*PS*IASDISLPIATQELR.Q |
| Chr2 synaptotagmin | ESYT2 | S660S663S665 | | | | | | | 30.2 | 25.7 | R.KTSIKS*HMS*GS*PGPGGSNTAPSTPVIGGSDKPGMEEK.A |
| Chr2 synaptotagmin | ESYT2 | T725S730S733 | | | | | | | 23.2 | 76.1 | K.EPT*PSIAS*DIS*LPIATQELR.Q |
| Chr2 synaptotagmin | ESYT2 | S663 | | | | | | | 3.8 | 11.9 | K.SHMS*GSPGPGGSNTAPSTPVIGGSDKPGMEEK.A |
| Chr2 synaptotagmin | ESYT2 | T656S657 | | | | | | | 7.3 | 13.2 | R.KT*S*IKSHMSGSPGPGGSNTAPSTPVIGGSDKPGMEEK.A |
| Chr2 synaptotagmin | ESYT2 | S709S710 | | | | | | | 5.0 | 40.0 | R.SS*S*SLLASPGHISVK.E |
| Chr2 synaptotagmin | ESYT2 | T656S657S665 | | | | | | | -0.6 | 23.6 | R.KT*S*IKSHMSGS*PGPGGSNTAPSTPVIGGSDKPGMEEK.A |
| Chr2 synaptotagmin | ESYT2 | S708S710 | | | | | | | 1.2 | 45.8 | R.S*SS*SLLASPGHISVK.E |
| Chromatin accessibility complex, subunit 1 | CHRAC1 | S124 | | | | | | | 20.4 | 41.8 | R.EEDEENDNDNES*DHDEADS.- |
| Chromatin assembly factor 1 subunit A | CHAF1A | S772 | | | | | | | 29.8 | 70.2 | R.GLLSNHTGS*PR.S |
| Chromatin assembly factor 1 subunit A | CHAF1A | S777 | | | | | | | 12.0 | 54.3 | R.SPS*TTYLHTPTPSEDAAIPSK.S |
| Chromatin assembly factor 1 subunit A | CHAF1A | S65 | | | | | | | 108.3 | 81.3 | K.S*PDLEASLDTLENNCHVGSDIDFRPK.L |
| Chromatin assembly factor 1 subunit A | CHAF1A | T770 | | | | | | | 22.0 | 44.8 | R.GLLSNHT*GSPR.S |
| Chromatin assembly factor 1 subunit A | CHAF1A | S775 | | | | | | | 10.1 | 50.6 | R.S*PSTTYLHTPTPSEDAAIPSK.S |
| Chromatin assembly factor 1 subunit A | CHAF1A | T778 | | | | | | | 7.2 | 45.3 | R.SPST*TYLHTPTPSEDAAIPSK.S |
| Chromatin assembly factor 1 subunit A | CHAF1A | S206 | | | | | | | 56.6 | 32.9 | R.S*CPELTSGPR.M |
| Chromatin assembly factor 1 subunit B | CHAF1B | S429 | | | | | | | 12.1 | 67.6 | R.TQDPSS*PGTTPPQAR.Q |
| Chromatin assembly factor 1 subunit B | CHAF1B | S428T433 | | | | | | | 23.4 | 44.5 | R.TQDPS*SPGTT*PPQAR.Q |
| Chromatin assembly factor 1 subunit B | CHAF1B | S538 | | | | | | | 42.1 | 33.6 | K.TDTPPSSVPTSVISTPSTEEIQSETPGDAQGS*PPELK.R |
| Chromatin assembly factor 1 subunit B | CHAF1B | S410 | | | | | | | 6.6 | 25.3 | R.GSS*PGPRPVEGTPASR.T |
| Chromatin assembly factor 1 subunit B | CHAF1B | S428 | | | | | | | 35.2 | 54.0 | R.TQDPS*SPGTTPPQAR.Q |
| Chromatin assembly factor 1 subunit B | CHAF1B | S429T433 | | | | | | | 17.7 | 10.9 | R.TQDPSS*PGTT*PPQAR.Q |
| Chromobox homolog 3 | CBX3 | S176 | | | | | | | 24.6 | 35.9 | R.LTWHS*CPEDEAQ.- |
| Chromobox homolog 3 | CBX3 | S93S95 | | | | | | | 21.8 | 58.9 | K.RKS*LS*DSESDDSK.S |
| | CBX3 | S95S97 | | | | | | | 13.6 | 38.9 | K.SLS*DS*ESDDSK.S |
| Chromobox homolog 3 | CBX3 | S95 | | | | | | | 28.9 | 62.3 | K.SLS*DSESDDSK.S |
| Chromobox homolog 3 | CBX3 | S95S97S99 | | | | | | | 12.2 | 38.1 | K.SLS*DS*ES*DDSK.S |
| Chromobox homolog 3 | CBX3 | S93 | | | | | | | 12.2 | 23.9 | K.S*LSDSESDDSK.S |
| Chromobox homolog 3 | CBX3 | S93S95S97 | | | | | | | 10.9 | 24.1 | K.S*LS*DS*ESDDSK.S |
| Chromobox homolog 7 | CBX7 | Y28 | | | | | | | 100.0 | 21.4 | R.KGKVEY*LVK.W |
| Chromodomain helicase DNA binding protein 1 | CHD1 | T250S252 | | | | | | | 109.3 | 31.4 | K.EDEEMKT*DS*DDLLEVCGEDVPQPEEEEFETIER.F |
| Chromodomain helicase DNA binding protein 2 | CHD2 | S1364 | | | | | | | | 32.7 | R.LKEEHGIELS*SPR.H |
| Chromodomain helicase DNA binding protein 3 | CHD3 | S1646S1650 | | | | | | | 27.6 | 20.4 | K.METEADAPS*PAPS*LGER.L |
| Chromodomain helicase DNA binding protein 4 | CHD4 | S512S528 | | | | | | | 38.4 | 20.7 | K.WGQPPS*PTPVPRPPDADPNTPS*PKPLEGRPER.Q |
| Chromodomain helicase DNA binding protein 4 | CHD4 | S306S307S316 | | | | | | | 19.5 | 95.7 | R.KRSS*S*EDDDLDVES*DFDDASINSYSVSDGSTSR.S |
| Chromodomain helicase DNA binding protein 4 | CHD4 | S305S306S307 | | | | | | | 49.4 | 119.1 | R.KRS*S*S*EDDDLDVES*DFDDASINSYSVSDGSTSR.S |
| Chromodomain helicase DNA binding protein 4 | CHD4 | S1360S1364S1 | | | | | | | 12.4 | 84.3 | R.DWQDDQS*DNQS*DYS*VAS*EEGDEDFDER.S |
| Chromodomain helicase DNA binding protein 4 | CHD4 | S1364Y1366S1 | | | | | | | 9.2 | 87.5 | R.DWQDDQSDNQS*DY*S*VAS*EEGDEDFDER.S |
| Chromodomain helicase DNA binding protein 4 | CHD4 | S307S322 | | | | | | | 2.3 | 61.0 | R.SSS*EDDDLDVESDFDDAS*INSYSVSDGSTSR.S |
| Chromodomain helicase DNA binding protein 4 | CHD4 | S307S316 | | | | | | | 6.3 | 20.5 | R.SSS*EDDDLDVES*DFDDASINSYSVSDGSTSR.S |
| Chromodomain helicase DNA binding protein 4 | CHD4 | S425 | | | | | | | 100.0 | 103.8 | K.EDNS*EGEEILEEVGGDLEEEDDHHMEFCR.V |
| Chromodomain helicase DNA binding protein 4 | CHD4 | S1560 | | | | | | | 13.9 | 38.7 | K.MSQPGS*PSPK.T |

Peak Area: <-10, -5, -3, 0, 3, 5, >10

%CV: 0, 29, 42, 71, 86, >100

**White dots-** Significant change in peptide abundance at **5** %FDR compared to the timepoint with the minimum peak area for a given PSM

| Protein Name | Gene | Phosphosites | CarT 0m | CarT 2m | CarT 5m | RajiB 0m | RajiB 2m | RajiB 5m | Ascor | MOWSE | Sequence |
|---|---|---|---|---|---|---|---|---|---|---|---|
| Chromodomain helicase DNA binding protein 4 | CHD4 | S1556S1560 | | | | | | | 16.4 | 43.2 | K.KMS*QPGS*PSPK.T |
| Chromodomain helicase DNA binding protein 4 | CHD4 | S1360S1364Y1 | | | | | | | 12.2 | 90.0 | R.DWQDDQS*DNQS*DY*SVAS*EEGDEDFDER.S |
| Chromodomain helicase DNA binding protein 4 | CHD4 | S305S307S316 | | | | | | | 18.7 | 62.6 | R.KRS*SS*EDDDLDVES*DFDDASINSYSVSDGSTSR.S |
| Chromodomain helicase DNA binding protein 4 | CHD4 | S306S307 | | | | | | | -0.4 | 12.5 | R.SS*S*EDDDLDVESDFDDASINSYSVSDGSTSR.S |
| Chromodomain helicase DNA binding protein 4 | CHD4 | T1574T1578 | | | | | | | 12.6 | 23.9 | K.TPTPSTPGDT*QPNT*PAPVPPAEDGIK.I |
| Chromodomain helicase DNA binding protein 4 | CHD4 | S1360Y1366S1 | | | | | | | 4.7 | 42.3 | R.DWQDDQS*DNQSDY*S*VAS*EEGDEDFDER.S |
| Chromodomain helicase DNA binding protein 4 | CHD4 | S1360S1364Y1 | | | | | | | 7.1 | 49.1 | R.DWQDDQS*DNQS*DY*S*VASEEGDEDFDER.S |
| Chromodomain helicase DNA binding protein 4 | CHD4 | S305S306 | | | | | | | 6.0 | 35.3 | R.S*S*SEDDDLDVESDFDDASINSYSVSDGSTSR.S |
| Chromodomain helicase DNA binding protein 4 | CHD4 | S305S307S316 | | | | | | | 8.7 | 63.0 | R.KRS*SS*EDDDLDVES*DFDDAS*INSYSVSDGSTSR.S |
| Chromodomain helicase DNA binding protein 4 | CHD4 | S1556 | | | | | | | 15.4 | 15.6 | K.MS*QPGSPSPK.T |
| Chromodomain helicase DNA binding protein 7 | CHD7 | S2559 | | | | | | | 49.0 | 27.6 | R.NIPS*PGQLDPDTRIPVINLEDGTR.L |
| Chromodomain helicase DNA binding protein 7 | CHD7 | S2956 | | | | | | | 35.7 | 44.0 | K.DGETLEGS*DAEESLDK.T |
| Chromodomain helicase DNA binding protein 7 | CHD7 | S2983 | | | | | | | 38.4 | 46.0 | K.TAESSLLEDEIAQGEELDS*LDGGDEIENNENDE.- |
| Chromodomain helicase DNA binding protein 7 | CHD7 | S1577S1581 | | | | | | | 1.5 | 12.6 | K.EDELMEFS*DLES*DSEEKPCAKPR.R |
| | CHD8 | S1141S1145 | | | | | | | 18.4 | 73.4 | R.HFSTLKDDDLVEFS*DLES*EDDERPR.S |
| Chromodomain helicase DNA binding protein 8 | CHD8 | T1714S1716 | | | | | | | | 28.4 | R.T*AS*PLPLRPDAPVEKSPEETATQVPSLESLTLK.L |
| Chromodomain helicase DNA binding protein 8 | CHD8 | S1716T1733 | | | | | | | 6.3 | 32.8 | R.TAS*PLPLRPDAPVEKSPEET*ATQVPSLESLTLK.L |
| Chromodomain helicase DNA binding protein 8 | CHD8 | S1940S1944 | | | | | | | 8.4 | 70.2 | R.SS*SAAS*MAEEEASAVSTAAAQFTK.L |
| Chromodomain helicase DNA binding protein 8 | CHD8 | S1940S1941 | | | | | | | 3.6 | 34.0 | R.SS*S*AASMAEEEASAVSTAAAQFTK.L |
| Chromodomain helicase DNA binding protein 8 | CHD8 | S1716S1729 | | | | | | | 11.4 | 57.1 | R.TAS*PLPLRPDAPVEKS*PEETATQVPSLESLTLK.L |
| Chromodomain helicase DNA binding protein 8 | CHD8 | T1925Y1929 | | | | | | | 10.7 | 17.9 | R.SQEMVTGGILGPGNHLLDSPSLT*PGEY*GDSPVPTPR.S |
| Chromodomain helicase DNA binding protein 8 | CHD8 | T1908S1932 | | | | | | | 13.9 | 27.2 | R.SQEMVT*GGILGPGNHLLDSPSLTPGEYGDS*PVPTPR.S |
| Chromodomain helicase DNA binding protein 8 | CHD8 | S1921S1932T1 | | | | | | | 25.6 | 24.9 | R.SQEMVTGGILGPGNHLLDS*PSLTPGEYGDS*PVPT*PR.S |
| Chromodomain helicase DNA binding protein 8 | CHD8 | S1941S1944 | | | | | | | 2.2 | 41.4 | R.SSS*AAS*MAEEEASAVSTAAAQFTK.L |
| Chromodomain helicase DNA binding protein 8 | CHD8 | S1697 | | | | | | | 37.8 | 66.7 | R.MNYMQNHQAGAPAPS*LSR.C |
| Chromodomain helicase DNA binding protein 8 | CHD8 | S1716T1735 | | | | | | | 9.4 | 24.1 | R.TAS*PLPLRPDAPVEKSPEETAT*QVPSLESLTLK.L |
| Chromodomain helicase DNA binding protein 8 | CHD8 | S1699 | | | | | | | 7.3 | 20.5 | R.MNYMQNHQAGAPAPSLS*R.C |
| Chromodomain helicase DNA binding protein 8 | CHD8 | S1939S1944 | | | | | | | 4.1 | 25.8 | R.S*SSAAS*MAEEEASAVSTAAAQFTK.L |
| Chromodomain-helicase-DNA-binding protein 3 (Fragment) | CHD3 | S371S375 | | | | | | | 35.8 | 63.4 | R.GRPPAQALGPAAS*PPPS*PPLGPSLG.- |
| Chromodomain-helicase-DNA-binding protein 3 (Fragment) | CHD3 | S371S381 | | | | | | | 14.9 | 26.1 | R.GRPPAQALGPAAS*PPPSPPLGPS*LG.- |
| Chromodomain-helicase-DNA-binding protein 4 | CHD4 | S96S98S101 | | | | | | | 6.0 | 22.7 | K.ELGDSSGEGPEFVEEEEEVALRS*DS*EGS*DYTPGK.K |
| Chromodomain-helicase-DNA-binding protein 4 | CHD4 | S78S96S98S10 | | | | | | | 6.6 | 17.2 | K.ELGDS*SGEGPEFVEEEEEVALRS*DS*EGS*DYTPGK.K |
| Chromosome 1 open reading frame 144 | SZRD1 | S37S39 | | | | | | | 62.2 | 60.4 | K.S*KS*PPKVPIVIQDDSLPAGPPPQIR.I |
| Chromosome 1 open reading frame 144 | SZRD1 | S39 | | | | | | | 48.4 | 72.2 | K.S*PPKVPIVIQDDSLPAGPPPQIR.I |
| Chromosome 1 open reading frame 144 | SZRD1 | S51 | | | | | | | 2.3 | 19.1 | K.SKSPPKVPIVIQDDS*LPAGPPPQIR.I |
| Chromosome 1 open reading frame 144 | SZRD1 | S105 | | | | | | | 2.8 | 39.8 | R.ILGS*ASPEEEQEKPILDRPTR.I |
| Chromosome 1 open reading frame 144 | SZRD1 | S37 | | | | | | | 6.2 | 15.6 | K.S*KSPPKVPIVIQDDSLPAGPPPQIR.I |
| Chromosome 1 open reading frame 218 | DENND1B | T270S275 | | | | | | | 9.5 | 45.5 | R.ET*LSQIS*DDLLIPGLGR.H |
| Chromosome 1 open reading frame 25 | TRMT1L | S31 | | | | | | | | 11.8 | R. [illegible] |
| Chromosome 1 open reading frame 9 | SUCO | S1377 | | | | | | | 10.2 | 60.9 | K.SGS*LPSLHDIIK.G |
| | FAM208B | S1243 | | | | | | | 25.3 | 16.4 | R.S*PLLVTVVESDPRPQGQPR.R |
| Chromosome 10 open reading frame 7 | CDC123 | T54S56 | | | | | | | | 15.4 | R.DDPPT*HS*QPDSDDEAEEIQWSDDENTATLTAPEFPEFATK.V |
| Chromosome 10 open reading frame 7 | CDC123 | S56T75 | | | | | | | 3.2 | 19.7 | R.DDPPTHS*QPDSDDEAEEIQWSDDENT*ATLTAPEFPEFATK.V |
| Chromosome 10 open reading frame 7 | CDC123 | T54S70 | | | | | | | 5.6 | 22.1 | R.DDPPT*HSQPDSDDEAEEIQWS*DDENTATLTAPEFPEFATK.V |
| Chromosome 10 open reading frame 7 | CDC123 | S60S70 | | | | | | | 5.8 | 32.9 | R.DDPPTHSQPDS*DDEAEEIQWS*DDENTATLTAPEFPEFATK.V |
| Chromosome 10 open reading frame 7 | CDC123 | S60T75 | | | | | | | 8.8 | 21.9 | R.DDPPTHSQPDS*DDEAEEIQWSDDENT*ATLTAPEFPEFATK.V |
| Chromosome 10 open reading frame 9 | CCNY | S83 | | | | | | | 100.0 | 31.3 | K.S*LFINHHPPGQIAR.K |
| Chromosome 10 open reading frame 9 | CCNY | S326 | | | | | | | 2.2 | 36.5 | R.SAS*ADNLTLPR.W |
| Chromosome 11 open reading frame 2 | VPS51 | Y660S663 | | | | | | | 6.0 | 35.3 | R.Y*APS*YTPSAPMDTNLLSNIQK.L |
| Chromosome 11 open reading frame 2 | VPS51 | S663Y664 | | | | | | | 3.0 | 42.4 | R.YAPS*Y*TPSAPMDTNLLSNIQK.L |
| Chromosome 11 open reading frame 2 | VPS51 | Y660S667 | | | | | | | 9.3 | 23.1 | R.Y*APSYTPS*APMDTNLLSNIQK.L |
| Chromosome 11 open reading frame 2 | VPS51 | Y664S667 | | | | | | | 4.8 | 34.7 | R.YAPSY*TPS*APMDTNLLSNIQK.L |
| Chromosome 12 open reading frame 41 | KANSL2 | S47 | | | | | | | 4.2 | 18.1 | R.ILDEDSWS*DGEQEPITVDQTWR.G |
| Chromosome 12 open reading frame 41 | KANSL2 | S45S47 | | | | | | | 25.2 | 28.8 | R.ILDEDS*WS*DGEQEPITVDQTWR.G |
| Chromosome 12 open reading frame 41 | KANSL2 | S66S70S73 | | | | | | | 100.0 | 62.9 | R.GDPDS*EADS*IDS*DQEDPLK.H |
| Chromosome 12 open reading frame 43 | C12orf43 | S179 | | | | | | | 19.2 | 47.2 | R.EAAVSASDILQESAIHS*PGTVEK.E |
| Chromosome 12 open reading frame 43 | C12orf43 | S138 | | | | | | | 100.0 | 35.9 | R.EKEES*PQPR.R |
| Chromosome 12 open reading frame 63 | CFAP54 | S278S279 | | | | | | | 12.2 | 11.8 | K.MLIS*S*EYSR.A |
| Chromosome 13 open reading frame 34 | C13orf34 | S270S273 | | | | | | | 12.0 | 59.9 | K.YSLGSITSPS*PIS*SPTFSPIEFQIGETPLSEQR.K |
| Chromosome 13 open reading frame 34 | C13orf34 | S270S274 | | | | | | | 7.9 | 60.4 | K.YSLGSITSPS*PISS*PTFSPIEFQIGETPLSEQR.K |
| Chromosome 13 open reading frame 34 | C13orf34 | S273S278 | | | | | | | 1.5 | 32.8 | K.YSLGSITSPSPIS*SPTFS*PIEFQIGETPLSEQR.K |
| Chromosome 14 open reading frame 173 | INF2 | S56 | | | | | | | 10.3 | 76.8 | R.EHNSMWASLSS*PDAEAVEPDFSSIER.L |

Peak Area: <-10, -5, -3, 0, 3, 5, >10

%CV: 0, 29, 42, 71, 86, >100

**White dots-** Significant change in peptide abundance at **5** %FDR compared to the timepoint with the minimum peak area for a given PSM

CarT (0m, 2m, 5m) | RajiB (0m, 2m, 5m)

| Protein Name | Gene | Phosphosites | Ascor | MOWSE | Sequence |
|---|---|---|---|---|---|
| Chromosome 14 open reading frame 173 | INF2 | S55 | 22.8 | 50.2 | R.EHNSMWASLS*SPDAEAVEPDFSSIER.L |
| Chromosome 14 open reading frame 173 | INF2 | S614 | 14.7 | 93.0 | K.DPTSLLGVLQAEADS*TSEGLEDAVHSR.G |
| Chromosome 14 open reading frame 43 | ELMSAN1 | Y654S661 | 39.3 | 25.2 | R.SFELPPY*TPPPILS*PVR.E |
| Chromosome 14 open reading frame 43 | ELMSAN1 | T655S661 | 30.6 | 28.8 | R.SFELPPYT*PPPILS*PVR.E |
| Chromosome 14 open reading frame 43 | ELMSAN1 | T704T715 | 26.9 | 63.8 | R.TNSAEVT*PPVLSVMGEAT*PVSIEPR.I |
| Chromosome 14 open reading frame 43 | ELMSAN1 | S700T704T715 | 10.9 | 36.2 | R.TNS*AEVT*PPVLSVMGEAT*PVSIEPR.I |
| Chromosome 14 open reading frame 43 | ELMSAN1 | S461 | 46.2 | 26.1 | R.RAS*QEANLLTLAQK.A |
| Chromosome 14 open reading frame 43 | ELMSAN1 | T704S709 | 6.5 | 23.4 | R.TNSAEVT*PPVLS*VMGEATPVSIEPR.I |
| Chromosome 14 open reading frame 43 | ELMSAN1 | S148 | 36.6 | 15.6 | K.GS*PHPGVGVPTYYNHPEALKR.E |
| Chromosome 14 open reading frame 46 | LIN52 | S28 | 75.2 | 42.1 | R.AS*PDLWPEQLPGVAEFAASFK.S |
| Chromosome 14 open reading frame 92 | TOX4 | S178S182 | 9.7 | 36.2 | R.LSTTPS*PTSS*LHEDGVEDFRR.Q |
| Chromosome 14 open reading frame 92 | TOX4 | S178S181 | 14.7 | 45.7 | R.LSTTPS*PTS*SLHEDGVEDFRR.Q |
| Chromosome 14 open reading frame 92 | TOX4 | T176S178S182 | 9.3 | 14.5 | R.LSTT*PS*PTSS*LHEDGVEDFRR.Q |
| Chromosome 14 open reading frame 92 | TOX4 | T176S178T180 | 23.2 | 19.1 | R.LSTT*PS*PT*SSLHEDGVEDFRR.Q |
| Chromosome 15 open reading frame 42 | TICRR | S1750 | 122.0 | 54.1 | R.TPILEDFELEGVCQLPDQS*PPR.N |
| Chromosome 16 open reading frame 53 | PAGR1 | T138S143S148 | 100.0 | 12.6 | R.RPPT*PEAQS*EEERS*DEEPEAK.E |
| Chromosome 17 open reading frame 49 | BAP18 | S96 | 29.2 | 38.7 | K.VYEDSGIPLPAES*PK.K |
| Chromosome 17 open reading frame 62 | C17orf62 | S178 | 7.3 | 34.5 | K.LITSFLELHCLESPTELSQSSDS*EAGDPASQS.- |
| Chromosome 17 open reading frame 62 | C17orf62 | S176 | 16.5 | 31.7 | K.LITSFLELHCLESPTELSQSS*DSEAGDPASQS.- |
| Chromosome 17 open reading frame 62 | C17orf62 | S173S176 | 9.8 | 21.0 | K.LITSFLELHCLESPTELS*QSS*DSEAGDPASQS.- |
| Chromosome 17 open reading frame 62 | C17orf62 | S175S176 | 7.5 | 30.4 | K.LITSFLELHCLESPTELSQS*S*DSEAGDPASQS.- |
| Chromosome 17 open reading frame 62 | C17orf62 | S173S175S176 | 19.7 | 24.8 | K.LITSFLELHCLESPTELS*QS*S*DSEAGDPASQS.- |
| Chromosome 17 open reading frame 62 | C17orf62 | S168T170S176 | 4.5 | 11.5 | K.LITSFLELHCLES*PT*ELSQSS*DSEAGDPASQS.- |
| Chromosome 17 open reading frame 62 | C17orf62 | T170S173S176 | 9.4 | 12.8 | K.LITSFLELHCLESPT*ELS*QSS*DSEAGDPASQS.- |
| Chromosome 17 open reading frame 62 | C17orf62 | S173S175S176 | 19.4 | 39.3 | K.LITSFLELHCLESPTELS*QS*S*DS*EAGDPASQS.- |
| Chromosome 17 open reading frame 62 | C17orf62 | T170S173S175 | 9.6 | 16.4 | K.LITSFLELHCLESPT*ELS*QS*S*DSEAGDPASQS.- |
| Chromosome 18 open reading frame 9 | CEP76 | S82 | 17.4 | 18.2 | K.ELNFVTDSVEQELPS*SPKQPICFDR.Q |
| Chromosome 19 open reading frame 13 | LSM14A | S183 | -0.3 | 33.4 | R.SS*PQLDPLR.K |
| Chromosome 19 open reading frame 13 | LSM14A | S192 | 31.8 | 87.4 | K.S*PTMEQAVQTASAHLPAPAAVGR.R |
| Chromosome 19 open reading frame 13 | LSM14A | S216 | 36.6 | 61.9 | R.S*PVSTRPLPSASQK.A |
| Chromosome 19 open reading frame 13 | LSM14A | S183S192 | 11.0 | 33.3 | R.SS*PQLDPLRKS*PTMEQAVQTASAHLPAPAAVGR.R |
| Chromosome 19 open reading frame 13 | LSM14A | S182S192 | 6.4 | 25.6 | R.S*SPQLDPLRKS*PTM#EQAVQTASAHLPAPAAVGR.R |
| Chromosome 19 open reading frame 13 | LSM14A | T194 | 12.8 | 57.2 | K.SPT*M#EQAVQTASAHLPAPAAVGR.R |
| Chromosome 19 open reading frame 13 | LSM14A | T201 | -1.6 | 41.2 | K.SPTMEQAVQT*ASAHLPAPAAVGR.R |
| Chromosome 19 open reading frame 13 | LSM14A | S178 | 26.9 | 26.0 | K.TQLS*QGR.S |
| Chromosome 19 open reading frame 13 | LSM14A | S182S183 | | 30.9 | R.S*S*PQLDPLRKSPTMEQAVQTASAHLPAPAAVGR.R |
| Chromosome 19 open reading frame 29 | CACTIN | S57S59 | 100.0 | 12.4 | R.RS*DS*EEER.W |
| Chromosome 19 open reading frame 43 | TRIR | S31S33T35 | | 84.7 | R. WAES*GS*GT*SPESGDEEVSGAGSSPVSGGVNLFANDGSFLELFK |
| Chromosome 19 open reading frame 43 | TRIR | S31 | | 16.9 | R.WAES*GSGTSPESGDEEVSGAGSSPVSGGVNLFANDGSFLELFK. R |
| Chromosome 19 open reading frame 43 | TRIR | S31S33T35S36 | | 38.1 | R. WAES*GS*GT*S*PESGDEEVSGAGSSPVSGGVNLFANDGSFLELFK |
| Chromosome 19 open reading frame 43 | TRIR | S31S33 | | 69.6 | R. WAES*GS*GTSPESGDEEVSGAGSSPVSGGVNLFANDGSFLELFK |
| Chromosome 19 open reading frame 43 | TRIR | S39 | 6.3 | 71.2 | R.WAESGSGTSPES*GDEEVSGAGSSPVSGGVNLFANDGSFLELFK. R |
| Chromosome 19 open reading frame 47 | C19orf47 | S213 | 34.2 | 110.6 | R.LGATPETDEDLAWDS*DNDSSSSVLQYAGVLK.K |
| Chromosome 19 open reading frame 47 | C19orf47 | S217 | | 11.4 | R.LGATPETDEDLAWDSDNDS*SSSVLQYAGVLK.K |
| chromosome 19 open reading frame 7 | ZC3H4 | S1269S1275 | 27.1 | 84.1 | K.TGSGS*PFAGNS*PAR.E |
| chromosome 19 open reading frame 7 | ZC3H4 | S1267S1275 | 21.7 | 74.4 | K.TGS*GSPFAGNS*PAR.E |
| chromosome 19 open reading frame 7 | ZC3H4 | T1106S1114 | 3.0 | 18.7 | R.AAKPGPAEAPSPT*ASPSGDAS*PPATAPYDPR.V |
| chromosome 19 open reading frame 7 | ZC3H4 | S1104S1110 | 19.2 | 35.4 | R.AAKPGPAEAPS*PTASPS*GDASPPATAPYDPR.V |
| chromosome 19 open reading frame 7 | ZC3H4 | S807S808 | 12.4 | 88.9 | R.ENEEGDTGNWYS*S*DEDEGGSSVTSILK.T |
| chromosome 19 open reading frame 7 | ZC3H4 | T802S808 | 2.9 | 27.7 | R.ENEEGDT*GNWYSS*DEDEGGSSVTSILK.T |
| chromosome 19 open reading frame 7 | ZC3H4 | S159 | 30.1 | 28.8 | R.EYS*PPYAPSHQQYPPSHATPLPK.K |
| chromosome 19 open reading frame 7 | ZC3H4 | S92S94 | 100.0 | 12.9 | K.HHS*DS*DEEK.S |
| chromosome 19 open reading frame 7 | ZC3H4 | S1104S1114 | 5.6 | 29.7 | R.AAKPGPAEAPS*PTASPSGDAS*PPATAPYDPR.V |
| chromosome 19 open reading frame 7 | ZC3H4 | Y806S807 | 12.9 | 88.1 | R.ENEEGDTGNWY*S*SDEDEGGSSVTSILK.T |
| chromosome 19 open reading frame 7 | ZC3H4 | T1106S1110 | 1.9 | 30.7 | R.AAKPGPAEAPSPT*ASPS*GDASPPATAPYDPR.V |
| chromosome 19 open reading frame 7 | ZC3H4 | T802Y806 | 9.2 | 17.2 | R.ENEEGDT*GNWY*SSDEDEGGSSVTSILK.T |
| chromosome 19 open reading frame 7 | ZC3H4 | T802S807 | 12.4 | 61.9 | R.ENEEGDT*GNWYS*SDEDEGGSSVTSILK.T |
| Chromosome 2 open reading frame 17 | RETREG2 | T279S281S283 | 37.6 | 29.1 | K.NAPPGGDEPLAET*ES*ES*EAELAGFSPVVDVK.K |
| Chromosome 2 open reading frame 17 | RETREG2 | S385 | 100.0 | 69.0 | R.QALDS*EEEEEDVAAK.E |
| Chromosome 2 open reading frame 49 | C2orf49 | S189S193 | 5.1 | 26.4 | R.KSPSGPVKS*PPLS*PVGTTPVK.L |
| Chromosome 2 open reading frame 49 | C2orf49 | S145 | 4.8 | 32.3 | R.KLSNS*SSSVSPLILSSNLPVNNK.T |
| Chromosome 20 open reading frame 172 protein | DSN1 | S81 | 16.3 | 47.7 | K.SLHLS*PQEQSASYQDR.R |

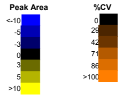


**White dots**- Significant change in peptide abundance at 5 %FDR compared to the timepoint with the minimum peak area for a given PSM

| Protein Name | Gene | Phosphosites | Ascor | MOWSE | Sequence |
|---|---|---|---|---|---|
| Chromosome 20 open reading frame 172 protein | DSN1 | S57 | 8.2 | 17.9 | R.IHLGS*SPK.K |
| Chromosome 20 open reading frame 172 protein | DSN1 | S27S28 | 12.5 | 36.3 | K.THDHQLES*S*LSPVEVFAK.T |
| Chromosome 20 open reading frame 74 | RALGAPA | S820S821 | 10.6 | 45.1 | R.RSS*S*PAELDLKDDLQQTQGK.C |
| Chromosome 20 open reading frame 74 | RALGAPA | S696 | 33.6 | 38.5 | R.SFS*LSWR.S |
| Chromosome 20 open reading frame 74 | RALGAPA | S373S375 | 10.1 | 33.8 | R.RLS*NS*SLCSIEEEHR.M |
| Chromosome 22 open reading frame 19 protein | THOC5 | T328 | 100.0 | 33.0 | R.RRPT*LGVQLDDK.R |
| Chromosome 22 open reading frame 19 protein | THOC5 | S307S312 | 2.0 | 16.1 | K.ALFKPPEDS*QDDES*DSDAEEEQTTKR.R |
| Chromosome 22 open reading frame 19 protein | THOC5 | S312S314 | 25.5 | 45.6 | K.ALFKPPEDSQDDES*DS*DAEEEQTTK.R |
| Chromosome 22 open reading frame 19 protein | THOC5 | S307S314 | 8.2 | 26.8 | K.ALFKPPEDS*QDDESDS*DAEEEQTTKR.R |
| Chromosome 5 open reading frame 22 | C5orf22 | Y435 | 11.9 | 43.4 | R.ALYGNLDLQVY*AAESPPS.- |
| Chromosome 5 open reading frame 22 | C5orf22 | Y427 | 0.1 | 16.4 | R.ALY*GNLDLQVYAAESPPS.- |
| Chromosome 5 open reading frame 37 | POC5 | S53 | 20.3 | 58.6 | R.ISTIHDILHSQGNNS*EVR.E |
| Chromosome 6 open reading frame 108 | DNPH1 | S169 | 32.7 | 56.7 | R.YFEADPPGQVAAS*PDPTT.- |
| Chromosome 6 open reading frame 108 | DNPH1 | S12 | 29.9 | 42.3 | R.SES*WER.G |
| Chromosome 6 open reading frame 203 | C6orf203 | S110S116 | 12.2 | 35.3 | K.VDEEDSDEES*HHDEMS*EQEEELEDDPTVVK.N |
| Chromosome 6 open reading frame 203 | C6orf203 | S106S116 | 24.3 | 43.7 | K.VDEEDS*DEESHHDEMS*EQEEELEDDPTVVK.N |
| Chromosome 6 open reading frame 206 | RSPH9 | T30 | 92.0 | 12.5 | R.ASLLT*SLMLVK.R |
| Chromosome 6 open reading frame 209 | LMBRD1 | S528S531 | 6.6 | 15.3 | K.SVIEGVDEDS*DIS*DDEPSVYSA.- |
| Chromosome 6 open reading frame 93 | LTV1 | S352S355 | 13.6 | 34.8 | K.EKWDCES*ICS*TYSNLYNHPQLIK.Y |
| Chromosome 6 open reading frame 93 | LTV1 | S352S355Y357 | 19.7 | 19.0 | K.EKWDCES*ICS*TY*S*NLYNHPQLIK.Y |
| Chromosome 6 open reading frame 93 | LTV1 | S331 | 12.7 | 23.3 | K.LNTLEPLEDQDLPM#NELDES*EEEEMITVVLEEAK.E |
| Chromosome 6 open reading frame 93 | LTV1 | T314 | 13.9 | 18.7 | K.LNT*LEPLEDQDLPM#NELDESEEEEMITVVLEEAK.E |
| Chromosome 6 open reading frame 93 | LTV1 | S355T356Y357 | 0.3 | 12.4 | K.EKWDCESICS*T*Y*S*NLYNHPQLIK.Y |
| Chromosome 7 open reading frame 47 | PPP1R35 | S52 | 42.6 | 28.2 | R.APVPEPGLDLSLSPRPDS*PQPR.H |
| Chromosome 7 open reading frame 47 | PPP1R35 | T84S87 | 100.0 | 22.4 | R.FRLT*PPS*PVR.S |
| Chromosome 7 open reading frame 50 | C7orf50 | S175 | 100.0 | 46.9 | R.ELDEEGS*DPPLPGR.A |
| Chromosome 9 open reading frame 142 | PAXX | S148 | 32.3 | 51.9 | R.LAAAEETAVS*PR.K |
| Chromosome 9 open reading frame 142 | PAXX | S152 | 100.0 | 35.8 | R.KS*PRPAGPQLFLPDPDPQR.G |
| Chromosome 9 open reading frame 82 protein | CAAP1 | S203 | 106.6 | 108.6 | K.ILEGDNGMDS*DMEEEADDGSK.M |
| Chromosome 9 open reading frame 82 protein | CAAP1 | S312 | 8.9 | 29.3 | K.SVNEILGLAESS*PNEPK.A |
| Chromosome 9 open reading frame 82 protein | CAAP1 | S311 | 111.6 | 38.6 | K.SVNEILGLAES*SPNEPK.A |
| Chromosome 9 open reading frame 86 | RABL6 | S597S598 | 54.3 | 80.9 | R.LKNDSDLFGLGLEEAGPKES*S*EEGK.E |
| Chromosome 9 open reading frame 86 | RABL6 | S382S384 | 100.0 | 84.7 | K.AAQQDS*DS*DGEALGGNPMVAGFQDDVDLEDQPR.G |
| Chromosome 9 open reading frame 86 | RABL6 | S411S421T425 | 8.1 | 42.6 | R.GS*PPLPAGPVPS*QDIT*LSSEEEAEVAAPTK.G |
| Chromosome 9 open reading frame 86 | RABL6 | S553T556 | 100.0 | 84.9 | R.DDPS*DVT*DEDEGPAEPPPPPK.L |
| Chromosome 9 open reading frame 86 | RABL6 | S411T425S427 | 0.2 | 35.3 | R.GS*PPLPAGPVPSQDIT*LS*SEEEAEVAAPTK.G |
| Chromosome 9 open reading frame 86 | RABL6 | S411S421S428 | 14.3 | 33.1 | R.GS*PPLPAGPVPS*QDITLSS*EEEAEVAAPTK.G |
| Chromosome 9 open reading frame 86 | RABL6 | S411S427S428 | 16.6 | 56.7 | R.GS*PPLPAGPVPSQDITLS*S*EEEAEVAAPTK.G |
| Chromosome 9 open reading frame 86 | RABL6 | S411T425S428 | 12.4 | 19.7 | R.GS*PPLPAGPVPSQDIT*LSS*EEEAEVAAPTK.G |
| Chromosome associated kinesin KIF4A | KIF4A | T1161 | 17.2 | 40.2 | K.LEDPTEVTPGLSFFNPVCAT*PNSK.I |
| Chromosome associated kinesin KIF4A | KIF4A | T799S801 | 30.0 | 24.6 | R.RRT*FS*LTEVR.G |
| Chromosome associated kinesin KIF4A | KIF4A | S801 | 17.4 | 47.3 | R.TFS*LTEVR.G |
| Chromosome condensation 1 | RCC1 | S11 | 100.0 | 13.0 | R.S*PPADAIPK.S |
| Chromosome condensation protein G | NCAPG | S674 | 39.1 | 57.4 | K.TLHCEGTEINS*DDEQESK.E |
| Chromosome condensation protein G | NCAPG | T968S973 | 21.0 | 29.7 | R.CQT*AEADS*ESDHEVPEPESEMK.M |
| Chromosome condensation protein G | NCAPG | S390 | 41.9 | 56.8 | R.GDFS*YIGNLMTK.E |
|  | NCAPG | S1015 | 100.0 | 69.0 | K.LNLAQFLNEDLS*.- |
| Chromosome condensation protein G | NCAPG | S1002 | 25.8 | 30.3 | K.S*KLNLAQFLNEDLS.- |
| Chromosome condensation protein G | NCAPG | S973S975 | 9.2 | 34.7 | R.CQTAEADS*ES*DHEVPEPESEMK.M |
| CIC | CIC | S1373S1382 | 17.5 | 31.0 | R.FAELPEFRPEEVLPS*PTLQSLATS*PR.A |
| CIC | CIC | S1373S1378 | 7.7 | 12.9 | R.FAELPEFRPEEVLPS*PTLQS*LATSPR.A |
| CIC | CIC | T1375S1382 | 7.7 | 18.2 | R.FAELPEFRPEEVLPSPT*LQSLATS*PR.A |
| CIC | CIC | S301 | -0.4 | 31.7 | R.SMS*ETGTAAAPGVSSELLSVAAQTLLSSDTK.A |
| CIC | CIC | S1373T1381 | 19.1 | 25.2 | R.FAELPEFRPEEVLPS*PTLQSLAT*SPR.A |
| Cingulin | CGN | S338 | 100.0 | 40.9 | R.KVS*LVLEK.M |
| Cingulin | CGN | S1182S1191 | 12.8 | 15.6 | K.NEGLSSDEEFDS*VYDPSSIAS*LLTESNLQTSSC.- |
| Cisplatin resistance associated overexpressed protein | LUC7L3 | T419S431 | 10.0 | 29.2 | K.NEVNGT*SEDIKSEGDTQS*N.- |
| Cisplatin resistance associated overexpressed protein | LUC7L3 | S425S431 | 10.8 | 22.2 | K.NEVNGTSEDIKS*EGDTQS*N.- |
| Cisplatin resistance associated overexpressed protein | LUC7L3 | S425T429 | 2.3 | 11.4 | K.NEVNGTSEDIKS*EGDT*QSN.- |
| Citron | CIT | S398 | 10.7 | 11.9 | K.NSWVS*SSPCQLSPSGFSGEELPFVGFSYSK.A |
| Claspin | CLSPN | S225 | 41.7 | 83.1 | K.DLFETGLEDENNS*PLEDEESLESIR.A |
| Claspin | CLSPN | T217 | 2.8 | 24.9 | K.DLFET*GLEDENNSPLEDEESLESIR.A |

**Peak Area**: <-10, -5, -3, 0, 3, 5, >10

**%CV**: 0, 29, 42, 71, 86, >100

**White dots-** Significant change in peptide abundance at **5** %FDR compared to the timepoint with the minimum peak area for a given PSM

CarT: 0m 2m 5m; RajiB: 0m 2m 5m

| Protein Name | Gene | Phosphosites | Ascor | MOWSE | Sequence |
|---|---|---|---|---|---|
| Clathrin, heavy polypeptide | CLTC | S1494 | 28.9 | 60.3 | R.TSIDAYDNFDNIS*LAQR.L |
| Claudin 12 | CLDN12 | S231 | 3.2 | 27.3 | R.SRLS*AIEIDIPVVSHTT.- |
| Cleavage and polyadenylation specificity factor 2 | CPSF2 | S419S420S423 | 114.1 | 83.3 | K.EADIDS*S*DES*DIEEDIDQPSAHK.T |
| Cleavage and polyadenylation specificity factor 6 | CPSF6 | T444 | 33.8 | 96.2 | R.T*PLSEAEFEEIMNR.N |
| Cleavage stimulation factor subunit 2 | CSTF2 | T507 | | 45.0 | R.QVPVMQGT*GMQGASIQGGSQPGGFSPGQNQVTPQDHEK.A |
| Cleavage stimulation factor subunit 2 | CSTF2 | S524 | 27.3 | 31.7 | R.QVPVMQGTGMQGASIQGGSQPGGFS*PGQNQVTPQDHEK.A |
| Cleavage stimulation factor subunit 2 | CSTF2 | S518 | 9.4 | 23.1 | R.QVPVMQGTGMQGASIQGGS*QPGGFSPGQNQVTPQDHEK.A |
| Cleavage stimulation factor subunit 2 | CSTF2 | T531 | -4.7 | 20.3 | R.QVPVMQGTGMQGASIQGGSQPGGFSPGQNQVT*PQDHEK.A |
| Cleavage stimulation factor, 3 Prime pre-RNA, subunit 3, 77kDa | CSTF3 | S691 | 100.0 | 33.2 | K.RPNEDS*DEDEEK.G |
| CLIP associated protein 1 | CLASP1 | S600 | 9.3 | 88.1 | R.SRS*DIDVNAAASAK.S |
| CLIP associated protein 1 | CLASP1 | S1196 | 100.0 | 32.2 | R.S*QEDLNEPIKR.D |
| CLIP associated protein 1 | CLASP1 | S572 | 10.5 | 36.8 | R.S*PTGSTTSR.A |
| CLN6 protein | CLN6 | S31 | 35.4 | 44.1 | R.HGS*VSADEAAR.T |
| CLNS1A | CLNS1A | S102 | 111.9 | 144.1 | K.EPVADEEEEDS*DDDVEPITEFR.F |
| CML2 | NAT8B | T83 | 100.0 | 11.1 | K.KPWT*R.Y |
| CND2 | NCAPH | T541 | | 31.3 | R. [illegible] |
| Coatomer protein complex, subunit beta 2 | COPB2 | T861 | 9.7 | 66.3 | R.STAQQELDGKPASPT*PVIVASHTANK.E |
| Coatomer protein complex, subunit beta 2 | COPB2 | S859 | 58.1 | 35.8 | R.STAQQELDGKPAS*PTPVIVASHTANKEEK.S |
| Coatomer protein complex, subunit epsilon | COPE | S45 | 12.1 | 40.1 | K.LSS*PER.D |
| Coatomer protein complex, subunit epsilon | COPE | S44 | 9.8 | 30.9 | K.LS*SPER.D |
| COBL like 1 | COBLL1 | S441 | 110.8 | 65.0 | K.S*QDIPFVSTDIINTLK.N |
| COG3 | COG3 | S536T543 | 6.5 | 40.8 | K.SGSTES*LNPRPQT*TISPADLHGMWYPTVR.R |
| COG3 | COG3 | S533T534 | -0.8 | 22.7 | K.SGS*T*ESLNPRPQTTISPADLHGMWYPTVR.R |
| Coiled coil domain containing 6 | CCDC6 | S240S244 | 100.0 | 58.9 | K.LDQPVS*APPS*PR.D |
| Coiled coil domain containing 6 | CCDC6 | S351S359S367 | 23.8 | 16.9 | R.TVS*SPIPYTPS*PSSSRPIS*PGLSYASHTVGFTPPTSLTR.A |
| Coiled coil domain containing 6 | CCDC6 | T427S431 | | 12.0 | K. [illegible] |
| Coiled coil domain containing 6 | CCDC6 | S352S359S367 | 13.4 | 11.5 | R.TVSS*PIPYTPS*PSSSRPIS*PGLSYASHTVGFTPPTSLTR.A |
| Coiled coil domain containing 6 | CCDC6 | S419 | 6.1 | 23.9 | R.RSNS*PDKFK.R |
| Coiled coil domain containing 6 | CCDC6 | S254 | 28.4 | 30.5 | R.DISMEIDS*PENMMR.H |
| Coiled coil domain containing 6 | CCDC6 | S351S352S363 | 7.6 | 11.8 | R.TVS*S*PIPYTPSPSSS*RPISPGLSYASHTVGFTPPTSLTR.A |
| Coiled coil domain containing 6 | CCDC6 | T349S351 | | 17.7 | R.T*VS*SPIPYTPSPSSSRPISPGLSYASHTVGFTPPTSLTR.A |
| Coiled coil domain containing 6 | CCDC6 | S351S359S363 | 14.5 | 13.5 | R.TVS*SPIPYTPS*PSSS*RPISPGLSYASHTVGFTPPTSLTR.A |
| Coiled coil domain containing 6 | CCDC6 | S352S361 | 1.1 | 16.5 | R.TVSS*PIPYTPSPS*SSRPISPGLSYASHTVGFTPPTSLTR.A |
| Coiled coil domain containing 6 | CCDC6 | S352S362 | 4.8 | 13.7 | R.TVSS*PIPYTPSPSS*SRPISPGLSYASHTVGFTPPTSLTR.A |
| Coiled coil domain containing 6 | CCDC6 | S351S363 | 7.6 | 22.7 | R.TVS*SPIPYTPSPSSS*RPISPGLSYASHTVGFTPPTSLTR.A |
| Coiled coil domain containing 9 | CCDC9 | S521 | 42.2 | 25.2 | R.TTHLAGALS*PGEAWPFESV.- |
| Coiled-coil domain containing 131 | ZFC3H1 | S1046 | 49.2 | 69.2 | R.S*FLESNYFTKPNLK.H |
| Coiled-coil domain containing 131 | ZFC3H1 | S28 | 60.5 | 91.1 | K.EEGELEDGEIS*DDDNNSQIR.S |
| Coiled-coil domain containing 131 | ZFC3H1 | S1301S1303 | 15.4 | 50.4 | R.KPISDNS*FS*SDEEQSTGPIK.Y |
| Coiled-coil domain containing 132 | VPS50 | Y96S98 | 19.7 | 29.3 | K.SAYQEY*DS*DSDVPEELKR.D |
| Coiled-coil domain containing 132 | VPS50 | Y96S100 | 14.1 | 29.3 | K.SAYQEY*DSDS*DVPEELKR.D |
| Coiled-coil domain containing 132 | VPS50 | Y93S98 | 3.2 | 26.2 | K.SAY*QEYDS*DSDVPEELKR.D |
| Coiled-coil domain containing 132 | VPS50 | S91 | 6.6 | 33.6 | K.S*AYQEYDSDSDVPEELKR.D |
| Coiled-coil domain containing 14 | CCDC14 | S598S602 | 17.3 | 17.9 | K.QDS*DEGS*ETMALIEDEHNLDNTIYIPFAR.S |
| Coiled-coil domain containing 144B | CCDC144 | S632 | 19.8 | 31.0 | K.DS*ASLPR.I |
| Coiled-coil domain containing 16 | ZNF830 | S351 | 97.8 | 71.2 | K.EEENADS*DDEGELQDLLSQDWR.V |
| | CCDC25 | S204 | 21.1 | 26.0 | K.VENMSSNQDGNDS*DEFM#.- |
| Coiled-coil domain containing 25 | CCDC25 | S196 | 10.0 | 28.2 | K.VENMS*SNQDGNDSDEFM#.- |
| Coiled-coil domain containing 43 | CCDC43 | T139 | 17.5 | 28.6 | K.AALLAQYADVT*DEEDEADEKDDSGATTMNIGSDK.L |
| Coiled-coil domain containing 52 | SPICE1 | S760S764 | 100.0 | 69.1 | K.LVGLNLS*PPMS*PVQLPLR.A |
| Coiled-coil domain containing 55 | NSRP1 | S248S254S255 | 91.4 | 78.4 | K.VEENPDADS*DFDAKS*S*ADDEIEETR.V |
| | NSRP1 | S33 | 5.8 | 58.0 | K.KTQQLHPVLQKPSVFGNDS*DDDDETSVSESLQR.E |
| Coiled-coil domain containing 55 | NSRP1 | S27 | 13.9 | 13.6 | K.TQQLHPVLQKPS*VFGNDSDDDDETSVSESLQR.E |
| Coiled-coil domain containing 55 | NSRP1 | T16 | | 46.8 | K.T*QQLHPVLQKPSVFGNDSDDDDETSVSESLQR.E |
| Coiled-coil domain containing 86 | CCDC86 | S91 | 122.6 | 45.8 | R.LQQGAGLESPQGQPEPGAAS*PQR.Q |
| Coiled-coil domain containing 86 | CCDC86 | S80S91 | 100.0 | 41.8 | R.LQQGAGLES*PQGQPEPGAAS*PQR.Q |
| Coiled-coil domain containing 86 | CCDC86 | S102S113 | 14.6 | 30.1 | R.QQDLHLES*PQRQPEYSPES*PR.C |
| Coiled-coil domain containing 86 | CCDC86 | S18 | 32.3 | 58.0 | R.LGGLRPES*PESLTSVSR.T |
| Coiled-coil domain containing 86 | CCDC86 | T42 | 6.4 | 22.2 | R.ALVEFESNPEET*REPGSPPSVQR.A |
| Coiled-coil domain containing 86 | CCDC86 | S47 | 6.0 | 19.7 | R.ALVEFESNPEETREPGS*PPSVQR.A |
| Coiled-coil domain containing 86 | CCDC86 | S58 | 100.0 | 20.5 | R.AGLGS*PERPPK.T |
| Coiled-coil domain containing 86 | CCDC86 | T65S66 | 21.8 | 15.9 | R.AGLGSPERPPKT*S*PGSPR.L |

Peak Area: <-10, -5, -3, 0, 3, 5, >10

%CV: 0, 29, 42, 71, 86, >100

**White dots-** Significant change in peptide abundance at **5** %FDR compared to the timepoint with the minimum peak area for a given PSM

| Protein Name | Gene | Phosphosites | CarT 0m | CarT 2m | CarT 5m | RajiB 0m | RajiB 2m | RajiB 5m | Ascor | MOWSE | Sequence |
|---|---|---|---|---|---|---|---|---|---|---|---|
| **Coiled-coil domain containing 86** | CCDC86 | S102Y109 | | | | | | | **2.7** | **21.5** | R.QQDLHLES*PQRQPEY*SPESPR.C |
| **Coiled-coil domain containing 86** | CCDC86 | S102S110 | | | | | | | **14.9** | **31.9** | R.QQDLHLES*PQRQPEYS*PESPR.C |
| **Coiled-coil domain containing 86** | CCDC86 | S217 | | | | | | | **8.9** | **48.8** | R.KGS*SSQAPASK.K |
| **Coiled-coil domain containing 86** | CCDC86 | T146 | | | | | | | | **15.6** | K.EELT*PGAPQHQLPPVPGSPEPYPGQQAPGPEPSQPLLELTPR.A |
| **Coiled-coil domain containing 94** | CCDC94 | S211S213 | | | | | | | **87.9** | **53.8** | R.LLEDS*DS*EDEAAPSPLQPALRPNPTAILDEAPKPK.R |
| **Coiled-coil domain containing 94** | CCDC94 | S305S306S316 | | | | | | | **6.7** | **17.7** | K.EANPTPLTPGAS*S*LSQLGAYLDS*DDSNGSN.- |
| | CCDC94 | S308Y313S316 | | | | | | | **15.7** | **36.6** | K.EANPTPLTPGASSLS*QLGAY*LDS*DDSNGSN.- |
| **Coiled-coil domain containing 94** | CCDC94 | Y313S316S322 | | | | | | | **6.5** | **15.6** | K.EANPTPLTPGASSLSQLGAY*LDS*DDSNGS*N.- |
| **Coiled-coil domain containing 94** | CCDC94 | S308Y313S319 | | | | | | | **1.3** | **23.0** | R.KEANPTPLTPGASSLS*QLGAY*LDSDDS*NGSN.- |
| **Coiled-coil domain containing 94** | CCDC94 | Y313S316 | | | | | | | **13.5** | **25.7** | K.EANPTPLTPGASSLSQLGAY*LDS*DDSNGSN.- |
| | CCDC94 | Y313S316S319 | | | | | | | **12.5** | **34.2** | K.EANPTPLTPGASSLSQLGAY*LDS*DDS*NGSN.- |
| **Coiled-coil domain containing 94** | CCDC94 | S211S213S220 | | | | | | | **57.9** | **14.1** | R.LLEDS*DS*EDEAAPS*PLQPALRPNPTAILDEAPKPK.R |
| **Coiled-coil domain containing 94** | CCDC94 | S319 | | | | | | | **-0.5** | **16.2** | K.EANPTPLTPGASSLSQLGAYLDSDDS*NGSN.- |
| **Coiled-coil domain containing 94** | CCDC94 | S308S316 | | | | | | | **7.8** | **27.2** | K.EANPTPLTPGASSLS*QLGAYLDS*DDSNGSN.- |
| **Coiled-coil domain containing 94** | CCDC94 | Y313S319S322 | | | | | | | **0.8** | **27.2** | R.KEANPTPLTPGASSLSQLGAY*LDSDDS*NGS*N.- |
| **Coiled-coil domain containing 94** | CCDC94 | S308Y313S322 | | | | | | | **3.7** | **25.1** | R.KEANPTPLTPGASSLS*QLGAY*LDSDDSNGS*N.- |
| **Coiled-coil domain containing 94** | CCDC94 | S308S316S319 | | | | | | | **1.3** | **18.1** | R.KEANPTPLTPGASSLS*QLGAYLDS*DDS*NGSN.- |
| **Coiled-coil domain containing 97** | CCDC97 | S257 | | | | | | | **100.0** | **49.9** | R.LLQQQEEEEACLEEEEEEEDS*DEEDQR.S |
| **Coiled-coil domain containing 97** | CCDC97 | S221 | | | | | | | **17.5** | **37.1** | R.TPTHQPPKPGSPGRPACPLS*NLLLQSYEER.E |
| **Coiled-coil domain containing 97** | CCDC97 | S212 | | | | | | | **22.9** | **37.2** | R.TPTHQPPKPGS*PGRPACPLSNLLLQSYEER.E |
| **Coiled-coil domain containing 97** | CCDC97 | T202T204 | | | | | | | **-2.9** | **30.1** | R.T*PT*HQPPKPGSPGRPACPLSNLLLQSYEER.E |
| **Coiled-coil domain containing 97** | CCDC97 | T204S212 | | | | | | | **8.4** | **37.3** | R.TPT*HQPPKPGS*PGRPACPLSNLLLQSYEER.E |
| **Coiled-coil domain containing 97** | CCDC97 | S337 | | | | | | | **98.3** | **22.6** | R.YFDEEEPEDAPS*PELDGD.- |
| **Coiled-coil domain containing 98** | ABRAXAS | S386S387T390 | | | | | | | **100.0** | **36.1** | K.MS*S*PET*DEEIEK.M |
| **Coiled-coil domain-containing protein 149** | CCDC149 | T337S341S354 | | | | | | | **1.6** | **11.5** | R.T*LEVS*GLWSLPGLSYNVS*VGFGRGK.D |
| **Coiled-coil-helix-coiled-coil-helix domain containing 3** | CHCHD3 | S50 | | | | | | | **13.9** | **111.3** | R.YS*GAYGASVSDEELK.R |
| **Coiled-coil-helix-coiled-coil-helix domain containing 3** | CHCHD3 | Y49 | | | | | | | **32.8** | **85.2** | R.Y*SGAYGASVSDEELK.R |
| **Coilin** | COIL | T122 | | | | | | | **100.0** | **35.0** | R.AFQLEEGEET*EPDCK.Y |
| **Cold shock domain protein A** | YBX3 | S201S203S204 | | | | | | | **54.1** | **45.5** | R.NYAGEEEEEGS*GS*S*EGFDPPATDR.Q |
| **Cold shock domain protein A** | YBX3 | S203S204 | | | | | | | **8.7** | **26.6** | R.NYAGEEEEEGSGS*S*EGFDPPATDR.Q |
| **Cold shock domain protein A** | YBX3 | S204 | | | | | | | **-0.2** | **24.8** | R.NYAGEEEEEGSGSS*EGFDPPATDR.Q |
| **Cold shock domain protein A** | YBX3 | S203 | | | | | | | | **19.7** | R.NYAGEEEEEGSGS*SEGFDPPATDR.Q |
| **Cold shock domain protein A** | YBX3 | Y192S201S203 | | | | | | | **50.7** | **15.8** | R.NY*AGEEEEEGS*GS*SEGFDPPATDR.Q |
| **Collagen type IV alpha 3 binding protein** | COL4A3B | S132S147 | | | | | | | **8.8** | **31.2** | R.HGS*MVSLVSGASGYSATS*TSSFKK.G |
| **Collagen type IV alpha 3 binding protein** | COL4A3B | S132T146 | | | | | | | **5.7** | **13.2** | R.HGS*MVSLVSGASGYSAT*STSSFKK.G |
| **Conserved nuclear protein NHN1** | ZC3H18 | S46 | | | | | | | **104.3** | **73.5** | R.AS*DLEDEESAAR.G |
| **Conserved nuclear protein NHN1** | ZC3H18 | S534 | | | | | | | **17.8** | **51.6** | K.LGVSVS*PSR.A |
| **Conserved nuclear protein NHN1** | ZC3H18 | S78 | | | | | | | **63.6** | **78.6** | K.S*QDQDSEVNELSR.G |
| **Conserved nuclear protein NHN1** | ZC3H18 | S532S534 | | | | | | | **15.5** | **31.7** | K.LGVS*VS*PSR.A |
| **Conserved nuclear protein NHN1** | ZC3H18 | S487 | | | | | | | **40.8** | **37.2** | R.S*PQPPSR.Q |
| **Conserved nuclear protein NHN1** | ZC3H18 | S868 | | | | | | | **100.0** | **46.1** | R.LGS*PKPER.Q |
| **Conserved nuclear protein NHN1** | ZC3H18 | S78S83 | | | | | | | **71.2** | **41.6** | K.S*QDQDS*EVNELSR.G |
| **Conserved nuclear protein NHN1** | ZC3H18 | S118 | | | | | | | **6.5** | **12.4** | R.DEASS*VTR.E |
| **Conserved nuclear protein NHN1** | ZC3H18 | S67S74 | | | | | | | **14.9** | **20.7** | R.GPSQEEEDNHS*DEEDRAS*EPK.S |
| **Conserved nuclear protein NHN1** | ZC3H18 | S59S67S74 | | | | | | | | **11.1** | R.GPS*QEEEDNHS*DEEDRAS*EPKSQDQDSEVNELSR.G |
| **Conserved nuclear protein NHN1** | ZC3H18 | S59S67S74S78 | | | | | | | | **16.7** | R.GPS*QEEEDNHS*DEEDRAS*EPKS*QDQDSEVNELSR.G |
| **Conserved nuclear protein NHN1** | ZC3H18 | S746 | | | | | | | **40.7** | **17.4** | R.S*PAPAQTR.K |
| **Conserved nuclear protein NHN1** | ZC3H18 | S117 | | | | | | | **13.8** | **23.8** | R.DEAS*SVTR.E |
| **Conserved nuclear protein NHN1** | ZC3H18 | S59S67S78 | | | | | | | **5.2** | **14.7** | R.GPS*QEEEDNHS*DEEDRASEPKS*QDQDSEVNELSR.G |
| **Conserved nuclear protein NHN1** | ZC3H18 | S532 | | | | | | | **-7.3** | **45.6** | K.KKLGVS*VSPSR.A |
| **Conserved nuclear protein NHN1** | ZC3H18 | S67S74S78S83 | | | | | | | **8.5** | **11.9** | R.GPSQEEEDNHS*DEEDRAS*EPKS*QDQDS*EVNELSR.G |
| **Conserved nuclear protein NHN1** | ZC3H18 | T851 | | | | | | | | **15.3** | K.RPNT*SPDR.G |
| **Conserved oligomeric Golgi complex subunit 1 (Fragment)** | COG1 | S113S114T115 | | | | | | | **100.0** | **13.5** | R.RS*S*T*AWLPR.S |
| **Conserved oligomeric Golgi complex subunit 1 (Fragment)** | COG1 | S113S114 | | | | | | | **5.2** | **23.7** | R.RS*S*TAWLPR.S |
| **Conserved oligomeric Golgi complex subunit 1 (Fragment)** | COG1 | S114T115 | | | | | | | **6.7** | **16.8** | R.RSS*T*AWLPR.S |
| **Copper homeostasis protein cutC homolog** | CUTC | S17 | | | | | | | **100.0** | **18.6** | R.ARIPS*GK.A |
| **Core binding factor, beta subunit** | CBFB | S173 | | | | | | | **46.9** | **89.0** | R.QQDPS*PGSNLGGGDDLK.L |
| **Coronin 7** | CORO7- | S21 | | | | | | | **20.7** | **18.5** | R.RES*WISDIR.A |
| **Coronin 7** | CORO7- | S462 | | | | | | | **39.8** | **82.2** | R.S*LQSLLGPSSK.F |
| **CRDBP** | IGF2BP1 | S181 | | | | | | | **100.0** | **79.9** | R.QGS*PVAAGAPAK.Q |
| **CRK** | CRK | T42 | | | | | | | **6.5** | **97.1** | R.DSST*SPGDYVLSVSENSR.V |
| **CRK** | CRK | S40 | | | | | | | **7.8** | **14.0** | R.DS*STSPGDYVLSVSENSR.V |

Peak Area: <-10, -5, -3, 0, 3, 5, >10

%CV: 0, 29, 42, 71, 86, >100

**White dots-** Significant change in peptide abundance at **5** %FDR compared to the timepoint with the minimum peak area for a given PSM

| Protein Name | Gene | Phosphosites | Ascor | MOWSE | Sequence |
|---|---|---|---|---|---|
| **CRK** | CRK | S41 | **9.5** | **74.5** | R.DSS*TSPGDYVLSVSENSR.V |
| **CRKL** | CRKL | S107 | **2.0** | **33.0** | R.YPS*PPMGSVSAPNLPTAEDNLEYVR.T |
| **CRKL** | CRKL | S112 | **10.6** | **49.2** | R.YPSPPMGS*VSAPNLPTAEDNLEYVR.T |
| **CRKL** | CRKL | S195 | | **13.8** | R. [illegible] |
| **Cross immune reaction antigen PCIA1** | DDA1 | S95 | **50.6** | **54.7** | R.TDS*PDMHEDT.- |
| **CRSP2** | MED14 | S995 | **12.0** | **16.1** | R.SVNEDDNPPSPIGGDMMDS*LISQLQPPPQQQPFPK.Q |
| **CRSP2** | MED14 | S998 | **37.2** | **17.8** | R.SVNEDDNPPSPIGGDMMDSLIS*QLQPPPQQQPFPK.Q |
| **CRSP2** | MED14 | S986 | **2.6** | **20.3** | R.SVNEDDNPPS*PIGGDMMDSLISQLQPPPQQQPFPK.Q |
| **CRSP2** | MED14 | S977 | | **10.9** | R.S*VNEDDNPPSPIGGDMMDSLISQLQPPPQQQPFPK.Q |
| **CRSP2** | MED14 | S977S995 | **2.2** | **22.1** | R.S*VNEDDNPPSPIGGDMMDS*LISQLQPPPQQQPFPK.Q |
| **CTD phosphatase, subunit 1** | CTDP1 | S869S872 | **24.7** | **39.7** | K.EVDDILGEGS*DDS*DSEK.R |
| **CTP synthetase 2** | CTPS2 | S568S571 | **13.4** | **27.2** | K.LSSSDRYS*DAS*DDSFSEPR.I |
| **CTP synthetase 2** | CTPS2 | S562S563S576 | **3.3** | **11.1** | K.LS*S*SDRYSDASDDSFS*EPR.I |
| **CTP synthetase 2** | CTPS2 | S563Y567S574 | **1.6** | **15.5** | K.LSS*SDRY*SDASDDS*FSEPR.I |
| **CTP synthetase 2** | CTPS2 | Y567S568S574 | **0.3** | **11.6** | K.LSSSDRY*S*DASDDS*FSEPR.I |
| **CTP synthetase 2** | CTPS2 | S571S574S576 | **15.0** | **21.7** | K.LSSSDRYSDAS*DDS*FS*EPR.I |
| **CTP synthetase 2** | CTPS2 | S563S571 | **2.4** | **34.9** | K.LSS*SDRYSDAS*DDSFSEPR.I |
| **CTP synthetase 2** | CTPS2 | S563Y567S568 | **5.3** | **14.9** | K.LSS*SDRY*S*DASDDSFSEPR.I |
| **CTP synthetase 2** | CTPS2 | S563Y567 | **6.3** | **11.8** | K.LSS*SDRY*SDASDDSFSEPR.I |
| **CTP synthetase 2** | CTPS2 | S563S564Y567 | **1.1** | **15.9** | K.LSS*S*DRY*SDASDDSFSEPR.I |
| **CTP synthetase 2** | CTPS2 | S564S571 | **2.5** | **17.6** | K.LSSS*DRYSDAS*DDSFSEPR.I |
| **CTP synthetase 2** | CTPS2 | S568S571S574 | **7.4** | **18.8** | K.LSSSDRYS*DAS*DDS*FSEPR.I |
| **CUGBP Elav-like family member 1 (Fragment)** | CELF1 | S22 | **19.1** | **24.0** | K.LDFLPEMMVDHCSLNSS*PVSK.K |
| **CUGBP Elav-like family member 1 (Fragment)** | CELF1 | S18 | **26.1** | **11.4** | K.LDFLPEMMVDHCS*LNSSPVSK.K |
| **Cullin 4A** | CUL4A | S10 | **29.8** | **70.3** | R.KGS*FSALVGR.T |
| **Cutaneous T cell lymphoma tumor antigen se70-2** | RBM26 | S127 | **20.7** | **11.7** | R.LNHS*PPQSSSR.Y |
| **Cutaneous T cell lymphoma tumor antigen se70-2** | RBM26 | T614 | **2.3** | **11.0** | R.EGSTQQLQT*TSPKPLVQQPILPVVK.Q |
| **CYBR** | CYTIP | S65 | **-0.3** | **59.7** | R.SS*SLSDFSWSQR.K |
| **CYBR** | CYTIP | S66 | **30.2** | **70.3** | R.SSS*LSDFSWSQR.K |
| **Cyclic GMP inhibited phosphodiesterase B** | PDE3B | S295S296 | **32.9** | **69.6** | R.RRS*S*CVSLGETAASYYGSCK.I |
| **Cyclic GMP inhibited phosphodiesterase B** | PDE3B | S981 | **-0.1** | **12.2** | R.SS*PQLAK.L |
| **Cyclic nucleotide gated channel beta 1** | CNGB1 | T1021 | **25.1** | **17.2** | K.SVLVT*LK.A |
| **Cyclin A1** | CCNA1 | T96T102 | **100.0** | **14.6** | R.RT*CGQGIT*R.I |
| **Cyclin B2** | CCNB2V | S92 | **32.3** | **59.7** | K.GPS*PTPEDVSMKEENLCQAFSDALLCK.I |
| **Cyclin dependent kinase 2** | CDK2 | T160 | **10.2** | **42.2** | R.TYT*HEVVTLWYR.A |
| **Cyclin dependent kinase 7** | CDK7 | S164 | **45.1** | **40.4** | K.SFGS*PNR.A |
| | CDKN3 | S14S15 | **21.4** | **77.0** | M.KPPSSIQTSEFDS*S*DEEPIEDEQTPIHISWLSLSR.V |
| **Cyclin dependent kinase inhibitor 3** | CDKN3 | S10S14 | **19.0** | **46.8** | M.KPPSSIQTS*EFDS*SDEEPIEDEQTPIHISWLSLSR.V |
| **Cyclin dependent kinase inhibitor 3** | CDKN3 | S5S6 | | **60.3** | M.KPPS*S*IQTSEFDSSDEEPIEDEQTPIHISWLSLSR.V |
| **Cyclin G associated kinase** | GAK | S826S829 | **25.8** | **52.4** | R.DES*EVS*DEGGSPISSEGQEPR.A |
| | CCNH | T315 | **67.5** | **56.6** | K.HEEEEWT*DDDLVESL.- |
| **Cyclin L1** | CCNL1 | S445 | **100.0** | **30.3** | R.HHNHGS*PHLK.A |
| **Cyclin L1** | CCNL1 | S352 | **27.8** | **67.6** | K.AEEKS*PISINVK.T |
| **Cyclin L1** | CCNL1 | S341 | | **23.7** | K.GLNPDGTPALSTLGGFSPASKPS*SPR.E |
| **Cyclin L1** | CCNL1 | S335S341 | **23.3** | **13.1** | K.GLNPDGTPALSTLGGFS*PASKPS*SPR.E |
| **Cyclin L1** | CCNL1 | S335S338 | **14.0** | **14.0** | K.GLNPDGTPALSTLGGFS*PAS*KPSSPR.E |
| **Cyclin L1** | CCNL1 | S335S342 | **18.6** | **19.4** | K.GLNPDGTPALSTLGGFS*PASKPSS*PR.E |
| **Cyclin L1** | CCNL1 | S338S341 | **15.0** | **11.0** | K.GLNPDGTPALSTLGGFSPAS*KPS*SPR.E |
| **Cyclin L2** | CCNL2 | S330 | **14.2** | **32.2** | R.GLLPGGTQVLDGTSGFS*PAPK.L |
| **Cyclin L2** | CCNL2 | S327 | **6.5** | **34.3** | R.GLLPGGTQVLDGTS*GFSPAPK.L |
| **Cyclin Y-like 1** | CCNYL1 | S274 | **25.1** | **85.8** | R.SFS*ADNFIGIQR.S |
| **Cyclin Y-like 1** | CCNYL1 | S272 | **14.0** | **73.7** | R.S*FSADNFIGIQR.S |
| **Cyclin-dependent kinase 9 (Fragment)** | CDK9 | S56 | **12.2** | **44.9** | R. [illegible] |
| **Cyclin-dependent kinase 9 (Fragment)** | CDK9 | S26 | | **64.1** | R. [illegible] |
| **Cyclin-dependent kinase 9 (Fragment)** | CDK9 | T54 | **12.2** | **68.7** | R. [illegible] |
| **Cylindromatosis gene protein** | CYLD | S418S422 | **37.2** | **26.0** | R.FHS*LPFS*LTK.M |
| **Cylindromatosis gene protein** | CYLD | S398 | **15.1** | **18.2** | K.SLTEISTDFDRS*SPPLQPPPVNSLTTENR.F |
| **Cylindromatosis gene protein** | CYLD | S399 | **4.0** | **35.5** | K.SLTEISTDFDRSS*PPLQPPPVNSLTTENR.F |
| **Cylindromatosis gene protein** | CYLD | S392 | **-0.1** | **25.6** | K.SLTEIS*TDFDRSSPPLQPPPVNSLTTENR.F |
| **Cysteine and glycine rich protein 1** | CSRP1 | S192 | **100.0** | **72.0** | K.GFGFGQGAGALVHS*E.- |
| **Cysteine string protein** | DNAJC5 | S12 | **1.4** | **84.6** | R.SLSTS*GESLYHVLGLDK.N |
| **Cysteine string protein** | DNAJC5 | S10 | **25.6** | **139.0** | R.SLS*TSGESLYHVLGLDK.N |

Peak Area: <-10, -5, -3, 0, 3, 5, >10

%CV: 0, 29, 42, 71, 86, >100

**White dots-** Significant change in peptide abundance at **5** %FDR compared to the timepoint with the minimum peak area for a given PSM

| Protein Name | Gene | Phosphosites | CarT (0m 2m 5m) | RajiB (0m 2m 5m) | Ascor | MOWSE | Sequence |
|---|---|---|---|---|---|---|---|
| Cysteine string protein | DNAJC5 | T11S12 | | | 8.9 | 77.1 | R.SLST*S*GESLYHVLGLDK.N |
| Cysteine string protein | DNAJC5 | S10S15 | | | 14.2 | 90.6 | R.SLS*TSGES*LYHVLGLDK.N |
| Cysteine string protein | DNAJC5 | T11 | | | -0.3 | 82.3 | R.SLST*SGESLYHVLGLDK.N |
| Cysteine string protein | DNAJC5 | T11S15 | | | 21.6 | 114.2 | R.SLST*SGES*LYHVLGLDK.N |
| Cysteine string protein | DNAJC5 | S12S15 | | | 12.9 | 104.8 | R.SLSTS*GES*LYHVLGLDK.N |
| Cysteine string protein | DNAJC5 | S10T11 | | | 7.9 | 64.8 | R.SLS*T*SGESLYHVLGLDK.N |
| Cysteine string protein | DNAJC5 | S10S12 | | | 6.6 | 37.2 | R.SLS*TS*GESLYHVLGLDK.N |
| Cysteine string protein | DNAJC5 | S8S15 | | | 4.0 | 44.1 | R.S*LSTSGES*LYHVLGLDK.N |
| Cytidine 5-prime triphosphate synthetase | CTPS1 | S574S575 | | | 14.3 | 108.2 | R.SGSS*S*PDSEITELKFPSINHD.- |
| Cytidine 5-prime triphosphate synthetase | CTPS1 | S571S574S575 | | | 15.6 | 63.9 | R.S*GSS*S*PDSEITELKFPSINHD.- |
| Cytidine 5-prime triphosphate synthetase | CTPS1 | S571S573S574 | | | 18.0 | 47.8 | R.DTYSDRS*GS*S*S*PDSEITELKFPSINHD.- |
| Cytidine 5-prime triphosphate synthetase | CTPS1 | S568S573S574 | | | -0.4 | 25.5 | R.DTYS*DRSGS*S*S*PDSEITELKFPSINHD.- |
| Cytidine 5-prime triphosphate synthetase | CTPS1 | S573S574 | | | 20.4 | 45.0 | R.SGS*S*SPDSEITELKFPSINHD.- |
| Cytidine 5-prime triphosphate synthetase | CTPS1 | S568 | | | 16.9 | 21.9 | R.DTYS*DR.S |
| Cytidine 5-prime triphosphate synthetase | CTPS1 | S571S574 | | | 6.0 | 24.8 | R.S*GSS*SPDSEITELKFPSINHD.- |
| Cytidine 5-prime triphosphate synthetase | CTPS1 | S571S578 | | | 4.4 | 18.2 | R.S*GSSSPDS*EITELKFPSINHD.- |
| Cytidine 5-prime triphosphate synthetase | CTPS1 | Y567S568S573 | | | 8.3 | 30.4 | R.DTY*S*DRSGS*SS*PDSEITELKFPSINHD.- |
| Cytidine 5-prime triphosphate synthetase | CTPS1 | Y567S568S571 | | | 13.2 | 46.7 | R.DTY*S*DRS*GSSS*PDSEITELKFPSINHD.- |
| Cytidine 5-prime triphosphate synthetase | CTPS1 | S571S573 | | | 9.1 | 87.0 | R.S*GS*SSPDSEITELKFPSINHD.- |
| Cytidine 5-prime triphosphate synthetase | CTPS1 | S571S573S574 | | | 9.4 | 63.6 | R.S*GS*S*SPDSEITELKFPSINHD.- |
| Cytidine 5-prime triphosphate synthetase | CTPS1 | Y567S571S574 | | | 6.7 | 11.6 | R.DTY*SDRS*GSS*S*PDSEITELKFPSINHD.- |
| Cytidine 5-prime triphosphate synthetase | CTPS1 | Y567S573S574 | | | 13.1 | 27.6 | R.DTY*SDRSGS*S*S*PDSEITELKFPSINHD.- |
| Cytidine 5-prime triphosphate synthetase | CTPS1 | S573S575 | | | 5.3 | 115.3 | R.SGS*SS*PDSEITELKFPSINHD.- |
| Cytidine 5-prime triphosphate synthetase | CTPS1 | S571S573S574 | | | 21.7 | 35.1 | R.DTYSDRS*GS*S*SPDS*EITELKFPSINHD.- |
| Cytidine 5-prime triphosphate synthetase | CTPS1 | S562 | | | 100.0 | 28.6 | K.GCRLS*PR.D |
| | CTPS1 | S575 | | | 10.8 | 64.2 | R.SGSSS*PDSEITELK.F |
| Cytidine 5-prime triphosphate synthetase | CTPS1 | S574 | | | 13.8 | 96.1 | R.SGSS*SPDSEITELKFPSINHD.- |
| Cytidine 5-prime triphosphate synthetase | CTPS1 | S573 | | | 10.6 | 70.7 | R.SGS*SSPDSEITELKFPSINHD.- |
| Cytidine 5-prime triphosphate synthetase | CTPS1 | S573S574S575 | | | 11.6 | 21.7 | R.DTYSDRSGS*S*S*PDS*EITELKFPSINHD.- |
| | CTPS1 | S573S574S575 | | | 9.2 | 59.3 | R.SGS*S*S*PDSEITELK.F |
| Cytidine 5-prime triphosphate synthetase | CTPS1 | S568S571S573 | | | 2.0 | 29.4 | R.DTYS*DRS*GS*S*SPDSEITELKFPSINHD.- |
| Cytidine 5-prime triphosphate synthetase | CTPS1 | S571S574S575 | | | 11.4 | 22.8 | R.S*GSS*S*PDS*EITELKFPSINHD.- |
| Cytoplasmic dynein 1 intermediate chain 2 (Fragment) | DYNC1I2 | S81 | | | 9.8 | 42.1 | R.EAEALLQSMGLTPESPIVPPPMS*PSSK.S |
| Cytoplasmic dynein 1 intermediate chain 2 (Fragment) | DYNC1I2 | S83 | | | 5.3 | 20.2 | R.EAEALLQSMGLTPESPIVPPPM#SPS*SK.S |
| Cytoplasmic linker associated protein 2 | CLASP2 | S603 | | | 9.3 | 84.3 | R.SRS*DIDVNAAAGAK.A |
| Cytoplasmic linker associated protein 2 | CLASP2 | S758S762 | | | 15.6 | 25.4 | R.IPRPSVS*QGCS*R.E |
| Cytoplasmic linker associated protein 2 | CLASP2 | S1224 | | | 26.5 | 43.5 | K.ASLLHSMPTHS*SPR.S |
| Cytoplasmic linker associated protein 2 | CLASP2 | S808 | | | 56.2 | 90.8 | R.VLNTGS*DVEEAVADALKKPAR.R |
| Cytoplasmic linker associated protein 2 | CLASP2 | S1164 | | | 54.8 | 66.7 | R.GVTEAIQNFS*FR.S |
| Cytoplasmic linker associated protein 2 | CLASP2 | T806 | | | 11.1 | 54.8 | R.VLNT*GSDVEEAVADALKKPAR.R |
| Cytoplasmic linker associated protein 2 | CLASP2 | S1113S1118S11 | | | 5.7 | 100.4 | R.S*PANWS*S*PLTSPTNTSQNTLSPSAFDYDTENMNSEDIYSSLR.G |
| Cytoplasmic linker associated protein 2 | CLASP2 | S1113S1118T11 | | | 9.7 | 100.4 | R.S*PANWS*SPLT*SPTNTSQNTLSPSAFDYDTENMNSEDIYSSLR.G |
| Cytoplasmic linker associated protein 2 | CLASP2 | S1113T1131S11 | | | 3.8 | 48.1 | R.S*PANWSSPLTSPTNTSQNT*LS*PSAFDYDTENMNSEDIYSSLR.G |
| Cytoplasmic linker associated protein 2 | CLASP2 | S1241 | | | 16.1 | 55.5 | R.DYNPYNYSDSIS*PFNK.S |
| Cytoplasmic linker associated protein 2 | CLASP2 | S1113T1125T11 | | | 2.9 | 94.3 | R.S*PANWSSPLTSPT*NTSQNT*LSPSAFDYDTENMNSEDIYSSLR.G |
| Cytoplasmic linker associated protein 2 | CLASP2 | S774 | | | 21.2 | 12.9 | R.DTS*PVR.S |
| Cytoplasmic linker associated protein 2 | CLASP2 | S756S758 | | | 6.8 | 17.5 | R.IPRPS*VS*QGCSR.E |
| Cytoplasmic linker associated protein 2 | CLASP2 | S1113S1128T11 | | | 2.6 | 52.2 | R.S*PANWSSPLTSPTNTS*QNT*LSPSAFDYDTENMNSEDIYSSLR.G |
| Cytoplasmic linker associated protein 2 | CLASP2 | S1113S1119T11 | | | 5.4 | 89.2 | R.S*PANWSS*PLT*SPTNTSQNTLSPSAFDYDTENMNSEDIYSSLR.G |
| Cytoplasmic linker associated protein 2 | CLASP2 | S1225 | | | | 19.6 | K.ASLLHSMPTHSS*PR.S |
| Cytoskeleton-like bicaudal D protein homolog 2 | BICD2 | S582 | | | 100.0 | 32.3 | R.S*PILLPK.G |
| D4, zinc and double PHD fingers family 2 | DPF2 | S142 | | | 66.8 | 92.8 | R.VDDDS*LGEFPVTNSR.A |
| D4, zinc and double PHD fingers family 2 | DPF2 | T176 | | | 27.3 | 48.2 | R.ILEPDDFLDDLDDEDYEEDT*PK.R |
| D4, zinc and double PHD fingers family 2 | DPF2 | S244T248 | | | 23.0 | 53.5 | K.NRPGLSYHYAHSHLAEEEGEDKEDS*QPPT*PVSQR.S |
| D4, zinc and double PHD fingers family 2 | DPF2 | S225Y226 | | | | 36.0 | K.NRPGLS*Y*HYAHSHLAEEEGEDKEDSQPPTPVSQR.S |
| Damage specific DNA binding protein 2 | DDB2 | S24 | | | 6.0 | 41.2 | R.S*RSPLELEPEAK.K |
| Damage specific DNA binding protein 2 | DDB2 | S24S26 | | | 100.0 | 36.3 | R.S*RS*PLELEPEAK.K |
| Damage specific DNA binding protein 2 | DDB2 | S26 | | | 100.0 | 77.5 | R.S*PLELEPEAK.K |
| Daxx | DAXX | S737S739 | | | 82.8 | 32.2 | K.TSVATQCDPEEIIVLS*DS*D.- |
| Daxx | DAXX | S668S671 | | | 18.1 | 50.4 | K.ICTLPS*PPS*PLASLAPVADSSTR.V |
| Daxx | DAXX | S668S675 | | | 9.0 | 43.9 | K.ICTLPS*PPSPLAS*LAPVADSSTR.V |
| Daxx | DAXX | S495 | | | 46.9 | 42.5 | K.DGDKS*PMSSLQISNEK.N |

Peak Area: <-10, -5, -3, 0, 3, 5, >10

%CV: 0, 29, 42, 71, 86, >100

**White dots-** Significant change in peptide abundance at **5** %FDR compared to the timepoint with the minimum peak area for a given PSM

| Protein Name | Gene | Phosphosites | CarT 0m | CarT 2m | CarT 5m | RajiB 0m | RajiB 2m | RajiB 5m | Ascor | MOWSE | Sequence |
|---|---|---|---|---|---|---|---|---|---|---|---|
| **Daxx** | DAXX | S671 | | | | | | | **4.5** | **11.6** | K.ICTLPSPPS*PLASLAPVADSSTR.V |
| **Daxx** | DAXX | S690S702 | | | | | | | **29.3** | **11.4** | R.VDSPS*HGLVTSSLCIPS*PAR.L |
| **DBC1** | CCAR2 | S124 | | | | | | | **100.0** | **85.8** | K.S*PAPPLLHVAALGQK.Q |
| **DBC1** | CCAR2 | S678S681 | | | | | | | **20.9** | **126.1** | R.SVAS*NQS*EMEFSSLQDMPK.E |
| **DBC1** | CCAR2 | S675S678 | | | | | | | **46.8** | **127.0** | R.S*VAS*NQSEMEFSSLQDMPK.E |
| **DBC1** | CCAR2 | S675S678S681 | | | | | | | **88.7** | **172.4** | R.S*VAS*NQS*EMEFSSLQDMPK.E |
| **DBC1** | CCAR2 | S678 | | | | | | | **7.9** | **35.3** | R.SVAS*NQSEMEFSSLQDMPK.E |
| **DBC1** | CCAR2 | S675S681 | | | | | | | **3.6** | **28.6** | R.S*VASNQS*EMEFSSLQDMPK.E |
| **DDHD domain containing 1** | DDHD1 | S723S727 | | | | | | | **16.8** | **22.9** | K.EPTSVSENEGISTIPS*PVTS*PVLSR.R |
| **DDX16** | DHX16 | S103S106 | | | | | | | **23.8** | **101.3** | R.LLEDS*EES*SEETVSR.A |
| **DDX16** | DHX16 | S103S106S107 | | | | | | | **38.3** | **76.9** | R.LLEDS*EES*S*EETVSR.A |
| **DDX18** | DDX18 | S86 | | | | | | | **29.8** | **19.0** | K.VTKS*PQK.S |
| **DDX21** | DDX21 | S121 | | | | | | | **100.0** | **79.8** | K.NEEPS*EEEIDAPKPK.K |
| **DDX21** | DDX21 | S164S171S173 | | | | | | | **2.2** | **14.8** | K.LKNGFPHPEPDCNPS*EAASEES*NS*EIEQEIPVEQK.E |
| | DDX21 | S168S171S173 | | | | | | | **41.3** | **63.1** | K.NGFPHPEPDCNPSEAAS*EES*NS*EIEQEIPVEQK.E |
| **DDX21** | DDX21 | S89 | | | | | | | **55.6** | **48.9** | K.KKEEPSQNDIS*PK.T |
| | DDX21 | S164S168S171 | | | | | | | **5.8** | **44.5** | K.NGFPHPEPDCNPS*EAAS*EES*NSEIEQEIPVEQK.E |
| **DDX23** | DDX23 | S14 | | | | | | | **32.2** | **50.2** | R.DAS*PSKEER.K |
| | DDX23 | S107S109 | | | | | | | **14.6** | **26.9** | K.RSS*LS*PGR.G |
| **DDX23** | DDX23 | S106S109 | | | | | | | **6.8** | **14.0** | R.KRS*SLS*PGR.G |
| **DDX23** | DDX23 | S106S107 | | | | | | | | **16.1** | R.KRS*S*LSPGR.G |
| **DDX23** | DDX23 | T25 | | | | | | | **100.0** | **15.3** | R.T*PDRER.D |
| **DDX24** | DDX24 | T302 | | | | | | | **9.4** | **67.4** | R.SPGKAEAESDALPDDT*VIESEALPSDIAAEAR.A |
| **DDX24** | DDX24 | S295 | | | | | | | **16.7** | **22.0** | R.SPGKAEAES*DALPDDTVIESEALPSDIAAEAR.A |
| **DDX24** | DDX24 | S82S94 | | | | | | | **27.2** | **23.7** | K.AQAVS*EEEEEEEGKSSS*PK.K |
| **DDX24** | DDX24 | S82 | | | | | | | **100.0** | **103.2** | K.AQAVS*EEEEEEEGK.S |
| **DDX24** | DDX24 | S82S93 | | | | | | | **26.7** | **35.6** | K.AQAVS*EEEEEEEGKSS*SPK.K |
| **DDX3** | DDX3X | S90 | | | | | | | **21.2** | **31.8** | K.SSFFSDRGS*GSR.G |
| **DDX3** | DDX3X | S594 | | | | | | | **100.0** | **48.6** | R.FS*GGFGAR.D |
| **DDX3** | DDX3X | S654 | | | | | | | **7.3** | **11.6** | R.GFGGGGYGGFYNSDGYGGNYNS*QGVDWWGN.- |
| **DDX39** | DDX39A | S426 | | | | | | | **3.2** | **42.2** | R.FEVNVAELPEEIDISTYIEQS*R.- |
| **DDX3Y** | DDX3Y | S588 | | | | | | | **16.5** | **19.4** | K.S*NRFSGGFGAR.D |
| **DDX3Y** | DDX3Y | S592 | | | | | | | **21.0** | **30.0** | K.SNRFS*GGFGAR.D |
| **DDX41** | DDX41 | S66S68 | | | | | | | **52.3** | **45.4** | K.GAAEEEQQDS*GS*EPRGDEDDIPLGPQSNVSLLDQHQHLK.E |
| **DDX41** | DDX41 | S23 | | | | | | | **107.2** | **135.8** | R.S*EAEDEDDEDYVPYVPLR.Q |
| **DDX41** | DDX41 | S21S23 | | | | | | | **33.8** | **35.2** | R.TDEVPAGGS*RS*EAEDEDDEDYVPYVPLR.Q |
| **DDX41** | DDX41 | S21 | | | | | | | **34.2** | **22.9** | R.TDEVPAGGS*R.S |
| **DDX42** | DDX42 | S104S111 | | | | | | | **16.0** | **72.0** | R.QQFHS*KPVDSDS*DDDPLEAFMAEVEDQAAR.D |
| **DDX42** | DDX42 | S751S754 | | | | | | | **11.3** | **10.8** | K.AGSSAAGASGWTSAGSLNSVPTNSAQQGHNS*PDS*PVTSAAK.C |
| **DDX42** | DDX42 | S109S111 | | | | | | | **26.5** | **71.3** | R.QQFHSKPVDS*DS*DDDPLEAFMAEVEDQAAR.D |
| **DDX42** | DDX42 | S104S109 | | | | | | | **7.0** | **44.9** | R.QQFHS*KPVDS*DSDDDPLEAFMAEVEDQAAR.D |
| **DDX42** | DDX42 | S754 | | | | | | | **12.0** | **67.4** | K.AGSSAAGASGWTSAGSLNSVPTNSAQQGHNSPDS*PVTSAAK.G |
| | DDX42 | S185 | | | | | | | **28.4** | **57.3** | R.YMAENPTAGVVQEEEEDNLEYDS*DGNPIAPTK.K |
| **DDX42** | DDX42 | S723 | | | | | | | | **16.4** | K.AGS*SAAGASGWTSAGSLNSVPTNSAQQGHNSPDSPVTSAAK.G |
| **DDX42** | DDX42 | S104 | | | | | | | **7.9** | **50.4** | R.QQFHS*KPVDSDSDDDPLEAFMAEVEDQAAR.D |
| **DDX42** | DDX42 | S751 | | | | | | | **8.6** | **24.7** | K.AGSSAAGASGWTSAGSLNSVPTNSAQQGHNS*PDSPVTSAAK.G |
| **DDX42** | DDX42 | T742S754 | | | | | | | **6.4** | **21.5** | K.AGSSAAGASGWTSAGSLNSVPT*NSAQQGHNSPDS*PVTSAAK.C |
| **DDX42** | DDX42 | S733S754 | | | | | | | **5.9** | **11.6** | K.AGSSAAGASGWTS*AGSLNSVPTNSAQQGHNSPDS*PVTSAAK.C |
| **DDX42** | DDX42 | T742S744 | | | | | | | **-0.4** | **16.6** | K.AGSSAAGASGWTSAGSLNSVPT*NS*AQQGHNSPDSPVTSAAK.C |
| **DDX46** | DDX46 | S24S26 | | | | | | | **18.7** | **20.1** | R.S*RS*PSDKR.S |
| **DDX46** | DDX46 | S804 | | | | | | | **145.3** | **89.5** | K.AALGLQDS*DDEDAAVDIDEQIESM#FNSK.K |
| **DDX46** | DDX46 | S295S296 | | | | | | | **10.6** | **48.4** | K.GELMENDQDAMEYS*S*EEEEVDLQTALTGYQTK.Q |
| **DDX46** | DDX46 | Y294S295 | | | | | | | **26.8** | **24.9** | K.GELMENDQDAMEY*S*SEEEEVDLQTALTGYQTK.Q |
| **DDX51** | DDX51 | S83 | | | | | | | **100.0** | **53.8** | R.VNDAEPGS*PEAPQGK.R |
| **DEAD (Asp-Glu-Ala-Asp) box polypeptide 17** | DDX17 | S599 | | | | | | | **14.7** | **22.9** | R.RDS*ASYR.D |
| **DEAD box polypeptide 55** | DDX55 | S544 | | | | | | | **144.0** | **87.8** | R.EEGS*DIEDEDMEELLNDTR.L |
| **DEAD-box protein 54** | DDX54 | S75 | | | | | | | **20.8** | **43.5** | K.LGPGRPLPTFFPTSECTS*DVEPDTR.E |
| **DEAD-box protein 54** | DDX54 | T74 | | | | | | | **42.1** | **35.0** | R.KLGPGRPLPTFFPTSECT*SDVEPDTR.E |
| **DEAD-box protein 54** | DDX54 | S39S41 | | | | | | | **100.0** | **114.8** | R.GS*DS*EDGEFEIQAEDDAR.A |
| **DEAD-box protein 54** | DDX54 | S782 | | | | | | | **73.4** | **28.8** | K.IDDRDS*DEEGASDR.R |
| **DEAH box polypeptide 57** | DHX57 | S127 | | | | | | | **100.0** | **70.7** | R.DLQEQDADAGS*ER.G |
| **Death associated protein** | DAP | S51 | | | | | | | **58.6** | **84.7** | K.DKDDQEWESPS*PPKPTVFISGVIAR.G |

Peak Area: <-10, -5, -3, 0, 3, 5, >10

%CV: 0, 29, 42, 71, 86, >100

**White dots-** Significant change in peptide abundance at **5** %FDR compared to the timepoint with the minimum peak area for a given PSM

CarT (0m, 2m, 5m) RajiB (0m, 2m, 5m)

| Protein Name | Gene | Phosphosites | Ascor | MOWSE | Sequence |
|---|---|---|---|---|---|
| **Death associated protein** | DAP | S49S51 | **84.1** | **39.4** | K.DKDDQEWES*PS*PPKPTVFISGVIAR.G |
| **Death associated protein** | DAP | T56 | **5.2** | **36.4** | K.DKDDQEWESPSPPKPT*VFISGVIAR.G |
| **Death associated protein** | DAP | S51T56 | **12.0** | **39.1** | K.DKDDQEWESPS*PPKPT*VFISGVIAR.G |
| | DIDO1 | T151S152S154 | **14.6** | **50.4** | K.VKGGDDHDDT*S*DS*DSDGLTLK.E |
| **Death associated transcription factor 1** | DIDO1 | S1040 | **38.5** | **70.4** | R.S*PPEGDTTLFLSR.L |
| **Death associated transcription factor 1** | DIDO1 | S1254S1260 | **4.9** | **13.8** | K.KPSKYPLCSADAAVS*TTPPGS*PPPPPPLPEPPVLK.V |
| | DIDO1 | T1256S1260 | **9.7** | **34.7** | K.YPLCSADAAVSTT*PPGS*PPPPPPLPEPPVLK.V |
| **Death associated transcription factor 1** | DIDO1 | S805S809 | **51.5** | **65.1** | R.QEAIPDLEDS*PPVS*DSEEQQESAR.A |
| **Death associated transcription factor 1** | DIDO1 | S1254T1255 | **2.2** | **25.6** | K.YPLCSADAAVS*T*TPPGSPPPPPPLPEPPVLK.V |
| | DIDO1 | T1255S1260 | **18.4** | **31.8** | K.YPLCSADAAVST*TPPGS*PPPPPPLPEPPVLK.V |
| **Death associated transcription factor 1** | DIDO1 | S1242Y1244 | | **40.1** | K.KPS*KY*PLCSADAAVSTTPPGSPPPPPPLPEPPVLK.V |
| **Death associated transcription factor 1** | DIDO1 | S1456 | **40.1** | **46.1** | R.RNS*VERPAEPVAGAATPSLVEQQK.M |
| **Death-inducing-protein** | GRAMD4 | S24S28 | **29.0** | **48.0** | R.DDFLDLAES*PNAS*DTECSDEIPLKVPR.T |
| **Decapping enzyme hDcp1b** | DCP1B | S276 | **16.9** | **36.4** | R.SLS*YEEPR.R |
| **Decapping enzyme hDcp2** | DCP2 | S246S247S249 | **45.3** | **49.1** | R.RFGDS*S*DS*DNGFSSTGSTPAKPTVEK.L |
| **Dedicator of cytokinesis 11** | DOCK11 | S440 | **31.0** | **60.0** | R.EMLWGSSTQLASDGS*PK.G |
| **Dedicator of cytokinesis 2** | DOCK2 | S1706 | **-0.4** | **62.1** | R.SS*VVFADEK.A |
| **Dedicator of cytokinesis 2** | DOCK2 | S587 | **13.6** | **46.5** | R.SSS*SVGGLSVSSR.D |
| **Dedicator of cytokinesis 2** | DOCK2 | S1731 | **27.8** | **57.8** | R.KHEFM#S*DTNLSEHAAIPLK.A |
| **Dedicator of cytokinesis 2** | DOCK2 | T1786 | **7.7** | **74.6** | R.LSQT*FLQLSDGDKK.T |
| **Dedicator of cytokinesis 2** | DOCK2 | S1685 | **32.3** | **68.1** | R.VEQEEPIS*PGSTLPEVK.L |
| **Dedicator of cytokinesis 2** | DOCK2 | T1733 | **2.8** | **11.8** | R.KHEFMSDT*NLSEHAAIPLK.A |
| **Dedicator of cytokinesis 2** | DOCK2 | S1784 | **4.4** | **64.4** | R.LS*QTFLQLSDGDKK.T |
| **Dedicator of cytokinesis 2** | DOCK2 | S1705 | | **37.5** | K.RS*SVVFADEK.A |
| **Dedicator of cytokinesis 2** | DOCK2 | S586 | **-0.3** | **29.3** | R.SS*SSVGGLSVSSR.D |
| **Dedicator of cytokinesis 7** | DOCK7 | S898 | **12.6** | **46.0** | R.SLS*NSNPDISGTPTSPDDEVR.S |
| **Dedicator of cytokinesis 7** | DOCK7 | S862 | **6.0** | **26.1** | R.LPNTYPNS*SSPGPGGLGGSVHYATMAR.S |
| **Dedicator of cytokinesis 7** | DOCK7 | S190 | **44.4** | **63.8** | R.GS*WACSIFDLK.N |
| **Dedicator of cytokinesis 7** | DOCK7 | S863 | **6.2** | **55.3** | R.LPNTYPNSS*SPGPGGLGGSVHYATMAR.S |
| **Dedicator of cytokinesis 7** | DOCK7 | S864 | **-0.5** | **23.1** | R.LPNTYPNSSS*PGPGGLGGSVHYATMAR.S |
| **Dedicator of cytokinesis 7** | DOCK7 | T935 | **7.3** | **38.0** | R.MSSHT*ETSSFLQTLTGR.L |
| **Dedicator of cytokinesis 7** | DOCK7 | S882 | **14.8** | **29.1** | R.S*AVRPASLNLNR.S |
| **Dedicator of cytokinesis 7** | DOCK7 | S900 | **7.0** | **59.0** | R.SLSNS*NPDISGTPTSPDDEVR.S |
| **Dedicator of cytokinesis 7** | DOCK7 | S180S190 | **27.7** | **21.7** | R.S*MSIDDTPRGS*WACSIFDLK.N |
| **Dedicator of cytokinesis 7** | DOCK7 | S180T186S190 | **4.1** | **11.3** | R.S*MSIDDT*PRGS*WACSIFDLK.N |
| **Dedicator of cytokinesis 7** | DOCK7 | S933 | **1.8** | **43.2** | R.MSS*HTETSSFLQTLTGR.L |
| **Dedicator of cytokinesis 7** | DOCK7 | S939 | **4.6** | **40.0** | R.MSSHTETSS*FLQTLTGR.L |
| **DEF6** | DEF6 | S590 | **100.0** | **23.5** | R.WGS*QGNR.T |
| **DEF6** | DEF6 | S387 | **100.0** | **62.2** | R.S*QHRELQQALEGQLR.E |
| **DEF6** | DEF6 | S580 | **13.1** | **23.8** | R.DS*SLKR.L |
| **DEF6** | DEF6 | S597 | **7.2** | **29.7** | R.TPS*PNSNEQQK.S |
| **DEF6** | DEF6 | S581 | **-0.2** | **15.9** | R.DSS*LKR.L |
| | DEK | S32 | **100.0** | **97.1** | R.EES*EEEEDEDDEEEEEEEKEK.S |
| **DEK Oncogene** | DEK | S301S303S306 | **100.0** | **48.7** | K.KES*ES*EDS*S*DDEPLIKK.L |
| **DEK Oncogene** | DEK | S243S244S251 | **100.0** | **45.6** | K.EES*S*DDEDKES*EEEPPKK.T |
| **DEK Oncogene** | DEK | T220S227S230 | **12.9** | **12.1** | R.T*KCPEILS*DES*SS*DEDEKK.N |
| **Delangin** | NIPBL | S306 | **14.3** | **58.7** | K.GSRPPLILQSQSLPCSS*PR.D |
| **Delangin** | NIPBL | S305 | **54.4** | **58.6** | K.GSRPPLILQSQSLPCS*SPR.D |
| **Delangin** | NIPBL | S2658 | **50.1** | **49.5** | K.AITSLLGGGS*PK.N |
| **Delangin** | NIPBL | S2482S2490S2 | **6.9** | **17.8** | R.KS*SPSKENES*S*DS*EEEVSRPR.K |
| **Delangin** | NIPBL | S1089S1090S1 | **55.8** | **58.1** | K.YAEIS*S*DEDNDS*DEAFESSR.K |
| **Deltex1** | DTX1 | S189 | **12.0** | **54.6** | K.SQS*WPVGASSGQPCSCQQCLLVNSTR.A |
| **DENN domain-containing protein 1B** | DENND1B | S653 | **20.1** | **99.0** | R.VSS*SGLTDSLFILK.E |
| **DENN domain-containing protein 1B** | DENND1B | S652S653 | **16.4** | **45.4** | K.RVS*S*SGLTDSLFILK.E |
| **DENN domain-containing protein 1B** | DENND1B | S652 | **10.8** | **102.2** | R.VS*SSGLTDSLFILK.E |
| **DENN domain-containing protein 1B** | DENND1B | S653S654 | **10.6** | **29.4** | K.RVSS*S*GLTDSLFILK.E |
| **Density regulated protein** | DENR | S73 | **36.8** | **20.3** | K.LTVENS*PK.Q |
| **Deoxycytidine kinase** | DCK | S74 | **18.8** | **45.8** | R.WCNVQSTQDEFEELTMS*QK.N |
| **DEP domain containing 1B** | DEPDC1B | S160 | **100.0** | **35.0** | K.RHS*IAIGEVPACR.L |
| **DEP domain containing 5** | DEPDC5 | S579S582 | **35.2** | **25.1** | R.FHVGS*AES*MLHVRPGGYTPQR.A |
| **Desmoglein 2** | DSG2 | S680 | **47.9** | **59.7** | K.VVPSFLPVDQGGS*LVGR.N |
| **DGCR14** | DGCR14 | T453 | **-0.9** | **44.5** | R.TPLT*QDPASITDNLLQLPAR.R |

Peak Area: <-10, -5, -3, 0, 3, 5, >10

%CV: 0, 29, 42, 71, 86, >100

**White dots-** Significant change in peptide abundance at **5** %FDR compared to the timepoint with the minimum peak area for a given PSM

| Protein Name | Gene | Phosphosites | CarT 0m | CarT 2m | CarT 5m | RajiB 0m | RajiB 2m | RajiB 5m | Ascor | MOWSE | Sequence |
|---|---|---|---|---|---|---|---|---|---|---|---|
| **Diacylglycerol kinase, eta** | | T26S27S31 | | | | | | | | **39.0** | -.[illegible] |
| **Diacylglycerol kinase, zeta** | DGKZ | S52S53 | | | | | | | **133.6** | **58.9** | R.RRS*S*AQLQGCLLSCGVR.A |
| **DiGeorge syndrome critical region gene 8** | DGCR8 | S35 | | | | | | | | **75.3** | R.[illegible] |
| **DiGeorge syndrome critical region gene 8** | DGCR8 | S275T279 | | | | | | | **21.5** | **12.9** | K.YGGDSDHPS*DGET*SVQPMMTK.I |
| **DiGeorge syndrome critical region gene 8** | DGCR8 | S377 | | | | | | | **6.8** | **27.7** | R.EQSSDLTPSGDVS*PVKPLSR.S |
| **DiGeorge syndrome critical region gene 8** | DGCR8 | S373 | | | | | | | **14.0** | **14.6** | R.EQSSDLTPS*GDVSPVKPLSR.S |
| **Dihydropyrimidinase-related protein 2** | DPYSL2 | S27 | | | | | | | **24.1** | **45.6** | K.NLGSGS*PKPR.Q |
| **Dihydrouridine synthase 3-like** | DUS3L | T273S276 | | | | | | | **19.3** | **39.5** | R.QENCGAQQVPAGPGTST*PPS*SPVR.T |
| **Dihydrouridine synthase 3-like** | DUS3L | S272S276 | | | | | | | **17.7** | **41.9** | R.QENCGAQQVPAGPGTS*TPPS*SPVR.T |
| **Dihydrouridine synthase 3-like** | DUS3L | T273S277 | | | | | | | **8.0** | **26.4** | R.QENCGAQQVPAGPGTST*PPSS*PVR.T |
| **DIS3** | DIS3 | S730 | | | | | | | **20.9** | **49.4** | K.SLAESLDQAES*PTFPYLNTLLR.I |
| **Disabled homolog 2-interacting protein** | DAB2IP | S35T37 | | | | | | | **100.0** | **16.1** | R.S*RT*RPAR.E |
| **Discs large associated protein 4** | DLGAP4 | S127 | | | | | | | **5.3** | **50.0** | R.KLSS*IGIQVDCIQPVPK.E |
| **Discs, large homolog 7** | DLGAP5 | S332 | | | | | | | **71.8** | **70.1** | R.S*ANAFLTPSYTWTPLK.T |
| **Discs, large homolog 7** | DLGAP5 | S806S812 | | | | | | | **44.0** | **61.9** | K.SLTTECHLLDS*PGLNCS*NPFTQLER.R |
| **Discs, large homolog 7** | DLGAP5 | S148S149 | | | | | | | **100.0** | **11.2** | K.AIPS*S*VR.I |
| **Disrupter of silencing 10** | UTP3 | S365S368 | | | | | | | **38.6** | **88.4** | K.TSAAACAVTDLS*DDS*DFDEK.A |
| **Disrupter of silencing 10** | UTP3 | T362S368 | | | | | | | **33.7** | **44.0** | K.TSAAACAVT*DLSDDS*DFDEK.A |
| **Disrupter of silencing 10** | UTP3 | T362S365 | | | | | | | **20.4** | **20.6** | K.TSAAACAVT*DLS*DDSDFDEK.A |
| **DKFZP434C212 protein** | GAPVD1 | S757T762 | | | | | | | **11.2** | **82.0** | R.EVS*SRPST*PGLSVVSGISATSEDIPNK.I |
| **DKFZP434C212 protein** | GAPVD1 | S757S758 | | | | | | | **9.8** | **43.0** | R.EVS*S*RPSTPGLSVVSGISATSEDIPNKIEDLR.S |
| **DKFZP434C212 protein** | GAPVD1 | T762S766 | | | | | | | **20.9** | **48.4** | R.EVSSRPST*PGLS*VVSGISATSEDIPNKIEDLR.S |
| **DKFZP434C212 protein** | GAPVD1 | S761T762 | | | | | | | **5.5** | **63.7** | R.EVSSRPS*T*PGLSVVSGISATSEDIPNK.I |
| **DKFZP434C212 protein** | GAPVD1 | S757S766 | | | | | | | **5.4** | **54.1** | R.EVS*SRPSTPGLS*VVSGISATSEDIPNKIEDLR.S |
| **DKFZP434C212 protein** | GAPVD1 | S757S761 | | | | | | | **16.6** | **85.5** | R.EVS*SRPS*TPGLSVVSGISATSEDIPNKIEDLR.S |
| **DKFZP434C212 protein** | GAPVD1 | S758T762 | | | | | | | **9.8** | **32.4** | R.EVSS*RPST*PGLSVVSGISATSEDIPNKIEDLR.S |
| **DKFZP434C212 protein** | GAPVD1 | S761S766 | | | | | | | **5.2** | **39.4** | R.EVSSRPS*TPGLS*VVSGISATSEDIPNKIEDLR.S |
| **DKFZP434C212 protein** | GAPVD1 | S930 | | | | | | | **-0.3** | **38.4** | R.SS*DIVSSVR.R |
| **DKFZP434C212 protein** | GAPVD1 | S929 | | | | | | | **17.0** | **36.0** | R.S*SDIVSSVR.R |
| **DKFZP434C212 protein** | GAPVD1 | S757S769 | | | | | | | **1.9** | **30.6** | R.EVS*SRPSTPGLSVVS*GISATSEDIPNK.I |
| **DKFZP434C212 protein** | GAPVD1 | S758S761 | | | | | | | **4.5** | **35.6** | R.EVSS*RPS*TPGLSVVSGISATSEDIPNKIEDLR.S |
| **DKFZP434C212 protein** | GAPVD1 | S974 | | | | | | | **4.8** | **15.2** | R.ELPPAAAIGATSLVAAPHSS*SSSPSK.D |
| **DKFZP434P1750 protein** | TBC1D10B | S383 | | | | | | | **22.5** | **60.8** | R.QQPPLGPSSS*LLSLPGLK.S |
| **DKFZP434P1750 protein** | TBC1D10B | S383S386 | | | | | | | **8.8** | **51.9** | R.QQPPLGPSSS*LLS*LPGLK.S |
| **DKFZP434P1750 protein** | TBC1D10B | S382S386 | | | | | | | **17.7** | **53.3** | R.QQPPLGPSS*SLLS*LPGLK.S |
| **DKFZP434P1750 protein** | TBC1D10B | S382S383 | | | | | | | **5.9** | **45.4** | R.QQPPLGPSS*S*LLSLPGLK.S |
| **DKFZP434P1750 protein** | TBC1D10B | T422 | | | | | | | **-3.2** | **20.9** | R.ASAGPAPGPVVT*AEGLHPSLPSPTGNSTPLGSSK.E |
| | TBC1D10B | S412 | | | | | | | **69.2** | **46.9** | R.AS*AGPAPGPVVTAEGLHPSLPSPTGNSTPLGSSK.E |
| **DKFZP434P1750 protein** | TBC1D10B | S381 | | | | | | | **11.1** | **30.1** | R.QQPPLGPS*SSLLSLPGLK.S |
| **DKFZP434P1750 protein** | TBC1D10B | S412S432 | | | | | | | **17.7** | **13.3** | R.RAS*AGPAPGPVVTAEGLHPSLPS*PTGNSTPLGSSK.E |
| **DKFZP434P1750 protein** | TBC1D10B | S381S386 | | | | | | | **6.8** | **11.6** | R.QQPPLGPS*SSLLS*LPGLK.S |
| **DKFZP564C186 protein** | NOC2L | S672S673 | | | | | | | **59.4** | **101.1** | K.DLFDLNS*S*EEDDTEGFSER.G |
| **DKFZP564C186 protein** | NOC2L | S22S26S30 | | | | | | | **16.5** | **39.9** | R.LAELTVDEFLAS*GFDS*ESES*ESENSPQAETR.E |
| **DKFZP564C186 protein** | NOC2L | S672 | | | | | | | **13.9** | **87.6** | K.DLFDLNS*SEEDDTEGFSER.G |
| **DKFZP564C186 protein** | NOC2L | S49 | | | | | | | **35.4** | **16.5** | R.S*PDKPGGSPSASR.R |
| **DKFZP564C186 protein** | NOC2L | S93S98S100S1 | | | | | | | | **55.1** | K.[illegible] |
| **DKFZP564C186 protein** | NOC2L | S49S56 | | | | | | | **18.3** | **28.8** | R.S*PDKPGGS*PSASR.R |
| **DKFZP564O123 protein** | CHMP2B | S199 | | | | | | | **11.0** | **42.0** | K.ATIS*DEEIER.Q |
| **DKFZp761A052 protein** | OTUD5 | S177 | | | | | | | **26.0** | **125.8** | R.EEVGAGYNS*EDEYEAAAAR.I |
| **DKFZp761A052 protein** | OTUD5 | S64 | | | | | | | **100.0** | **46.8** | R.AS*PPPQGPLPGPPGALHR.W |
| **DKFZp761A052 protein** | OTUD5 | T507 | | | | | | | **67.4** | **29.9** | R.AT*SPLVSLYPALECR.A |
| **DMAP1** | DMAP1 | T445 | | | | | | | **9.8** | **37.2** | K.DTIIDVVGAPLT*PNSR.K |
| **Dmx like 1** | DMXL1 | T573 | | | | | | | **-0.5** | **25.8** | R.ST*SMLISSGHNK.S |
| **DNA damage inducible protein 2** | DDI2 | T104 | | | | | | | **9.1** | **73.8** | R.IDFSSIAVPGT*SSPR.Q |
| **DNA damage inducible protein 2** | DDI2 | S194 | | | | | | | **100.0** | **72.9** | R.LFS*ADPFDLEAQAK.I |
| **DNA dependent protein kinase catalytic subunit** | PRKDC | T2609S2612 | | | | | | | **22.1** | **68.5** | R.STVLTPMFVET*QAS*QGTLQTR.T |
| **DNA dependent protein kinase catalytic subunit** | PRKDC | S2612 | | | | | | | **7.4** | **43.6** | R.STVLTPMFVETQAS*QGTLQTR.T |
| **DNA Ligase III** | LIG3 | S210 | | | | | | | **1.0** | **74.0** | K.LTTTGQVTS*PVKGASFVTSTNPR.K |
| **DNA Ligase III** | LIG3 | S241 | | | | | | | **7.8** | **14.1** | R.KFSGFSAKPNNSGEAPS*SPTPK.R |
| | LIG3 | T209 | | | | | | | **33.2** | **20.6** | K.LTTTGQVT*SPVK.G |
| **DNA methyltransferase 1** | DNMT1 | S714 | | | | | | | **100.0** | **86.5** | K.EADDDEEVDDNIPEMPS*PK.K |
| **DNA methyltransferase 1** | DNMT1 | S394 | | | | | | | **21.2** | **28.5** | K.LSIFDANES*GFESYEALPQHK.L |

**Peak Area**: <-10, -5, -3, 0, 3, 5, >10

**%CV**: 0, 29, 42, 71, 86, >100

**White dots-** Significant change in peptide abundance at **5** %FDR compared to the timepoint with the minimum peak area for a given PSM

| Protein Name | Gene | Phosphosites | CarT 0m | CarT 2m | CarT 5m | RajiB 0m | RajiB 2m | RajiB 5m | Ascor | MOWSE | Sequence |
|---|---|---|---|---|---|---|---|---|---|---|---|
| **DNA mismatch repair protein PMS2** | PMS2 | S523 | | | | | | | **-11.2** | **48.5** | K.DSGHGSTSVDSEGFSIPDTGSHCSSEYAASS*PGDR.G |
| **DNA mismatch repair protein PMS2** | PMS2 | S522 | | | | | | | **44.0** | **60.2** | K.DSGHGSTSVDSEGFSIPDTGSHCSSEYAAS*SPGDR.G |
| **DNA mismatch repair protein PMS2** | PMS2 | S517 | | | | | | | **8.3** | **23.9** | K.DSGHGSTSVDSEGFSIPDTGSHCSS*EYAASSPGDR.G |
| **DNA polymerase delta interacting protein 3** | POLDIP3 | S127 | | | | | | | **-0.3** | **42.7** | R.SS*PAAFINPPIGTVTPALK.L |
| **DNA polymerase subunit B** | | S141 | | | | | | | **35.2** | **70.5** | R.S*PHQLLSPSSFSPSATPSQK.Y |
| **DNA Polymerase, alpha** | POLA1 | S186 | | | | | | | **49.8** | **79.6** | R.S*IGASPNPFSVHTATAVPSGK.I |
| **DNA primase large subunit** | PRIM2 | Y504 | | | | | | | **8.8** | **33.3** | K.DASSALASLNSSLEMDMEGLEDY*FSEDS.- |
| | XRCC4 | S327S328 | | | | | | | **100.0** | **26.6** | R.NS*S*PEDLFDEI.- |
| **DNA replication licensing factor MCM4** | MCM4 | S131 | | | | | | | **58.4** | **86.5** | K.GLQVDLQS*DGAAAEDIVASEQSLGQK.L |
| **DNA topoisomerase II alpha** | TOP2A | S1469T1470S1 | | | | | | | **27.8** | **19.6** | R.KPS*T*S*DDS*DSNFEK.I |
| **DNA topoisomerase II alpha** | TOP2A | S1392 | | | | | | | **7.2** | **42.6** | K.GSVPLSS*SPPATHFPDETEITNPVPK.K |
| **DNA topoisomerase II alpha** | TOP2A | S1377 | | | | | | | **23.4** | **62.9** | K.SVVS*DLEADDVK.G |
| **DNA topoisomerase II alpha** | TOP2A | S1377S1391 | | | | | | | **6.5** | **40.4** | K.SVVS*DLEADDVKGSVPLS*SSPPATHFPDETEITNPVPK.K |
| **DNA topoisomerase II alpha** | TOP2A | S1525 | | | | | | | **35.3** | **35.7** | K.YLEES*DEDDLF.- |
| **DNA topoisomerase II alpha** | TOP2A | S1332S1337T1 | | | | | | | **22.2** | **36.5** | K.FTMDLDS*DEDFS*DFDEKT*DDEDFVPSDAS*PPK.T |
| **DNA topoisomerase II alpha** | TOP2A | S1332S1337T1 | | | | | | | **36.4** | **28.5** | K.FTMDLDS*DEDFS*DFDEKT*DDEDFVPS*DAS*PPK.T |
| **DNA topoisomerase II alpha** | TOP2A | T1112 | | | | | | | **3.9** | **71.1** | K.VPDEEENEESDNEKET*EKSDSVTDSGPTFNYLLDMPLWYLTK.E |
| **DNA topoisomerase II alpha** | TOP2A | S1115 | | | | | | | **4.3** | **71.0** | K.VPDEEENEESDNEKETEKS*DSVTDSGPTFNYLLDMPLWYLTK.E |
| **DNA topoisomerase II alpha** | TOP2A | T1124 | | | | | | | **-0.7** | **25.5** | K.VPDEEENEESDNEKETEKSDSVTDSGPT*FNYLLDMPLWYLTK.E |
| | TOP2A | S1106 | | | | | | | **40.3** | **65.3** | K.VPDEEENEES*DNEKETEK.S |
| **DNA topoisomerase II alpha** | TOP2A | S1247 | | | | | | | **22.8** | **107.9** | K.NENTEGS*PQEDGVELEGLK.Q |
| **DNA topoisomerase II alpha** | TOP2A | S1391 | | | | | | | **32.8** | **33.3** | K.GSVPLS*SSPPATHFPDETEITNPVPK.K |
| **DNA topoisomerase II alpha** | TOP2A | S1377S1392 | | | | | | | **7.4** | **36.3** | K.SVVS*DLEADDVKGSVPLSS*SPPATHFPDETEITNPVPK.K |
| **DNA topoisomerase II alpha** | TOP2A | S1121 | | | | | | | **7.5** | **51.9** | K.VPDEEENEESDNEKETEKSDSVTDS*GPTFNYLLDMPLWYLTK.E |
| **DNA topoisomerase II alpha** | TOP2A | S1393 | | | | | | | **5.3** | **24.7** | K.GSVPLSSS*PPATHFPDETEITNPVPK.K |
| **DNA topoisomerase II alpha** | TOP2A | S1332S1337 | | | | | | | **17.7** | **55.5** | K.FTMDLDS*DEDFS*DFDEK.T |
| **DNA topoisomerase II alpha** | TOP2A | S1332S1337T1 | | | | | | | **24.4** | **53.9** | K.FTMDLDS*DEDFS*DFDEKT*DDEDFVPSDASPPK.T |
| **DNA topoisomerase II alpha** | TOP2A | S1332S1337T1 | | | | | | | **16.3** | **31.8** | K.FTMDLDS*DEDFS*DFDEKT*DDEDFVPS*DASPPK.T |
| **DNA topoisomerase II alpha** | TOP2A | T1327S1332S1 | | | | | | | **18.2** | **13.7** | K.FT*MDLDS*DEDFS*DFDEKT*DDEDFVPS*DASPPK.T |
| **DNA topoisomerase II alpha** | TOP2A | S1117 | | | | | | | **0.8** | **67.4** | K.VPDEEENEESDNEKETEKSDS*VTDSGPTFNYLLDMPLWYLTK.E |
| **DNA topoisomerase II alpha** | TOP2A | S1387S1392 | | | | | | | **3.1** | **10.8** | K.SVVSDLEADDVKGS*VPLSS*SPPATHFPDETEITNPVPK.K |
| **DNA topoisomerase II alpha** | TOP2A | T1119 | | | | | | | **7.0** | **48.9** | K.VPDEEENEESDNEKETEKSDSVT*DSGPTFNYLLDMPLWYLTK.E |
| **DNA-binding protein RFX7** | RFX7 | T1025S1028 | | | | | | | **26.8** | **33.3** | R.HHDTHFGRLT*PVS*PVQHQGATVNNTNK.Q |
| **DNA-binding protein RFX7** | RFX7 | T1019T1025 | | | | | | | **5.3** | **10.8** | R.HHDT*HFGRLT*PVSPVQHQGATVNNTNK.Q |
| **DnaJ homology subfamily A member 5** | DNAJC21 | S283 | | | | | | | **100.0** | **26.5** | K.EFGDGS*DENEMEEHELK.D |
| **DOCK10** | DOCK10 | S318S322 | | | | | | | **18.3** | **32.1** | K.IPRPLS*LIGS*TLR.F |
| **DOCK10** | DOCK10 | S289S292 | | | | | | | **9.4** | **32.3** | R.AS*LAS*LDSNPSTNEK.S |
| **DOCK8** | DOCK8 | S1177 | | | | | | | **20.1** | **54.9** | R.TSGS*DEEQEGAGAINQNVALAIAGNNFNLK.T |
| **DOCK8** | DOCK8 | S1175S1177 | | | | | | | **16.2** | **65.3** | R.YRTS*GS*DEEQEGAGAINQNVALAIAGNNFNLK.T |
| **DOCK8** | DOCK8 | T1174S1175 | | | | | | | **14.0** | **50.9** | R.YRT*S*GSDEEQEGAGAINQNVALAIAGNNFNLK.T |
| **DOCK8** | DOCK8 | T1174S1177 | | | | | | | **11.2** | **64.3** | R.YRT*SGS*DEEQEGAGAINQNVALAIAGNNFNLK.T |
| **Docking protein 3** | DOK3 | S330 | | | | | | | **27.6** | **73.0** | R.ATS*LPSLDTPGELR.E |
| **Docking protein 3** | DOK3 | S425 | | | | | | | **13.1** | **69.6** | R.S*PTTSPIYHNGQDLSWPGPANDSTLEAQYR.R |
| **Docking protein 3** | DOK3 | S439 | | | | | | | **1.1** | **48.9** | R.SPTTSPIYHNGQDLS*WPGPANDSTLEAQYR.R |
| **Docking protein 3** | DOK3 | T427T428 | | | | | | | **5.8** | **40.4** | R.SPT*T*SPIYHNGQDLSWPGPANDSTLEAQYR.R |
| **Docking protein 3** | DOK3 | S429S439 | | | | | | | **1.6** | **18.7** | R.SPTTS*PIYHNGQDLS*WPGPANDSTLEAQYR.R |
| **Docking protein 3** | DOK3 | S429 | | | | | | | **1.8** | **14.2** | R.SPTTS*PIYHNGQDLSWPGPANDSTLEAQYR.R |
| **Docking protein 3** | DOK3 | S425Y432 | | | | | | | **7.2** | **69.2** | R.S*PTTSPIY*HNGQDLSWPGPANDSTLEAQYR.R |
| **Docking protein 4** | DOK4 | S35 | | | | | | | **16.9** | **25.9** | R.KSS*SKGPQR.L |
| **Double strand break repair protein MRE11A** | MRE11 | S688S689 | | | | | | | **141.5** | **138.0** | K.GVDFES*S*EDDDDDPFMNTSSLR.R |
| **Double strand break repair protein MRE11A** | MRE11 | S649 | | | | | | | **45.7** | **84.5** | K.NYSEVIEVDES*DVEEDIFPTTSK.T |
| **Down regulated in metastasis** | UTP20 | S2601 | | | | | | | **64.7** | **21.0** | K.AES*DGEEKEEVKEELGRPATLLWLIQK.L |
| **Downregulator of transcription 1** | DR1 | S157 | | | | | | | **2.2** | **73.6** | R. [illegible] |
| **Downregulator of transcription 1** | DR1 | S166S167 | | | | | | | **-0.6** | **32.2** | R. [illegible] |
| **Downregulator of transcription 1** | DR1 | S157S159 | | | | | | | | **35.2** | R. [illegible] |
| **Downregulator of transcription 1** | DR1 | S159S161 | | | | | | | **2.1** | **28.7** | R. [illegible] |
| **Drebrin E** | DBN1 | S341 | | | | | | | **4.0** | **90.8** | R.SPS*DSSTASTPVAEQIER.A |
| **Drebrin E** | DBN1 | S143 | | | | | | | **12.1** | **47.9** | R.LS*SPVLHR.L |
| **Drebrin E** | DBN1 | S339 | | | | | | | **21.2** | **96.2** | R.S*PSDSSTASTPVAEQIER.A |
| **Drebrin E** | DBN1 | S274 | | | | | | | **8.4** | **20.0** | K.S*ESEVEEAAAIIAQRPDNPR.E |
| **Dual adapter for phosphotyrosine and 3-phosphotyrosine and 3-phosphoinositide** | DAPP1 | S276 | | | | | | | **7.6** | **13.2** | R.SRS*FIFK.- |
| **Dual-specificity protein phosphatase 22** | DUSP22 | S86 | | | | | | | **13.8** | **32.7** | R.RWSS*FPALAPLTYDNYTTET.- |

Peak Area: <-10, -5, -3, 0, 3, 5, >10

%CV: 0, 29, 42, 71, 86, >100

**White dots-** Significant change in peptide abundance at **5** %FDR compared to the timepoint with the minimum peak area for a given PSM

CarT: 0m, 2m, 5m — RajiB: 0m, 2m, 5m

| Protein Name | Gene | Phosphosites | Ascor | MOWSE | Sequence |
|---|---|---|---|---|---|
| **dUTP pyrophosphatase** | DUT | S11 | **16.1** | **71.4** | M.PCSEETPAIS*PSKR.A |
| **Dynamin 2** | DNM2 | S736 | | **32.5** | K.EALNIIGDIS*TSTVSTPVPPPVDDTWLQSASSHSPTPQR.R |
| **Dynamin 2** | DNM2 | T762 | **8.8** | **36.6** | K.EALNIIGDISTSTVSTPVPPPVDDTWLQSASSHSPT*PQR.R |
| **Dynamin 2** | DNM2 | S760 | **12.2** | **31.1** | K.EALNIIGDISTSTVSTPVPPPVDDTWLQSASSHS*PTPQR.R |
| **Dynamin 2** | DNM2 | S757 | **4.7** | **12.5** | K.EALNIIGDISTSTVSTPVPPPVDDTWLQSAS*SHSPTPQR.R |
| **Dynamin 2** | DNM2 | S758 | | **13.5** | K.EALNIIGDISTSTVSTPVPPPVDDTWLQSASS*HSPTPQR.R |
| **Dynamin related protein 1** | DNM1L | S616 | **43.6** | **25.2** | K.SKPIPIMPAS*PQK.G |
| **Dynein light chain A** | DYNC1LI1 | S516 | **13.9** | **64.7** | R.KPVTVSPTTPTS*PTEGEAS.- |
| **Dynein light chain A** | DYNC1LI1 | T515 | **20.1** | **60.3** | R.KPVTVSPTTPT*SPTEGEAS.- |
| **Dynein light chain A** | DYNC1LI1 | S207 | **39.3** | **19.0** | R.DFQEYVEPGEDFPAS*PQRR.N |
| **Dynein light chain A** | DYNC1LI1 | T513T515 | **9.5** | **11.2** | R.KPVTVSPTT*PT*SPTEGEAS.- |
| **Dynein light chain A** | DYNC1LI1 | S510T515 | **8.5** | **14.8** | R.KPVTVS*PTTPT*SPTEGEAS.- |
| **Dynein, axonemal, heavy polypeptide 8** | DNAH8 | S4103T4106 | **100.0** | **11.2** | K.GVS*WNT*VR.Y |
| **Dynein, cytoplasmic, heavy polypeptide 1** | DYNC1H1 | T4369 | **6.5** | **44.1** | R.TDST*SDGRPAWMR.T |
| **Dynein, cytoplasmic, heavy polypeptide 1** | DYNC1H1 | S4368 | **-0.4** | **32.7** | R.TDS*TSDGRPAWMR.T |
| **Dynein, cytoplasmic, heavy polypeptide 2** | DYNC2H1 | T432 | **10.9** | **12.6** | K.RPT*ISK.E |
| **Dynein, cytoplasmic, light intermediate polypeptide 2** | DYNC1LI2 | S194 | **34.8** | **35.2** | K.DFQDYMEPEEGCQGS*PQRR.G |
| **DYRK1B** | DYRK1A | Y273 | **33.4** | **39.3** | R.IYQY*IQSR.F |
| **DYRK1B** | DYRK1A | Y271 | **15.1** | **47.6** | R.IY*QYIQSR.F |
| **Dyskerin** | DKC1 | S494 | **19.8** | **80.6** | K.AGLESGAEPGDGDS*DTTK.K |
| **Dyskerin** | DKC1 | S451S455 | **30.9** | **36.9** | R.KRES*ESES*DETPPAAPQLIK.K |
| | DKC1 | S513 | **100.0** | **42.9** | K.EVELVS*E.- |
| **Dyskerin** | DKC1 | S451S453S455 | **14.0** | **33.9** | R.KRES*ES*ES*DETPPAAPQLIK.K |
| **Dyskerin** | DKC1 | S451S455T458 | **8.8** | **28.9** | R.KRES*ESES*DET*PPAAPQLIK.K |
| **Dyskerin** | DKC1 | S21 | **126.1** | **49.7** | K.S*LPEEDVAEIQHAEEFLIKPESK.V |
| **Dyskerin** | DKC1 | S451S453T458 | **14.8** | **21.9** | R.KRES*ES*ESDET*PPAAPQLIK.K |
| **Dyskerin** | DKC1 | S451T458 | **1.2** | **15.7** | R.KRES*ESESDET*PPAAPQLIK.K |
| **Dyskerin** | DKC1 | S453S455 | **-0.1** | **26.4** | R.KRESES*ES*DETPPAAPQLIK.K |
| **Dyskerin** | DKC1 | S451S453S455 | **100.0** | **21.7** | R.KRES*ES*ES*DET*PPAAPQLIK.K |
| **Dyskerin** | DKC1 | S451S453 | **12.4** | **45.6** | R.ES*ES*ESDETPPAAPQLIK.K |
| **Dystrobrevin alpha** | DTNA | S564 | **-0.3** | **14.8** | R.SS*PSHTISRPIPMPIR.S |
| **Dystrophin** | DMD | S291 | **30.2** | **35.2** | R.TSS*PKPR.F |
| **Dystrophin** | DMD | T289 | **10.0** | **13.4** | R.T*SSPKPR.F |
| **E1A binding protein p300** | EP300 | T1906T1909 | **11.0** | **57.6** | K.AAGQVT*PPT*PPQTAQPPLPGPPPAAVEMAMQIQR.A |
| **E1A binding protein p300** | EP300 | T1909T1913 | **11.4** | **31.4** | K.AAGQVTPPT*PPQT*AQPPLPGPPPAAVEMAMQIQR.A |
| **E1A binding protein p400** | EP400 | S23S37 | | **50.3** | R. [illegible] |
| **E1A binding protein p400** | EP400 | S315T320 | **8.5** | **43.3** | R.TPGVLLPGAGGAAGFGMTS*PPPPT*SPSR.T |
| **E1A binding protein p400** | EP400 | T320 | **12.6** | **50.9** | R.TPGVLLPGAGGAAGFGMTSPPPPT*SPSR.T |
| **E1A binding protein p400** | EP400 | T314S321 | **6.0** | **18.7** | R.TPGVLLPGAGGAAGFGMT*SPPPPTS*PSR.T |
| **E1A binding protein p400** | EP400 | T314T320 | **25.3** | **38.1** | R.TPGVLLPGAGGAAGFGMT*SPPPPT*SPSR.T |
| **E1A binding protein p400** | EP400 | S904T908 | **23.0** | **14.4** | R.KAS*ISLT*DDEVDDEEETIEEEEANEGVVDHQTELSNLAK.E |
| **E2-230K** | UBE2O | S87S89S98S99 | **100.0** | **13.0** | R.LIHGEDS*DS*EGEEEGRGS*S*GCS*EAGGAGHEEGR.A |
| **E2-230K** | UBE2O | S839 | **17.8** | **71.5** | K.NMTVEQLLTGSPTS*PTVEPEKPTR.E |
| **E2-230K** | UBE2O | T838 | **19.3** | **47.5** | K.NMTVEQLLTGSPT*SPTVEPEKPTR.E |
| **E2-230K** | UBE2O | S836S839 | **11.5** | **54.9** | K.NMTVEQLLTGS*PTS*PTVEPEKPTR.E |
| **E2-230K** | UBE2O | S115 | **100.0** | **23.2** | R.AS*PLRR.G |
| **E2-230K** | UBE2O | S87S89 | **100.0** | **39.7** | R.LIHGEDS*DS*EGEEEGR.G |
| **E2-230K** | UBE2O | S836 | **15.4** | **55.0** | K.NMTVEQLLTGS*PTSPTVEPEKPTR.E |
| **E2-230K** | UBE2O | T838S839 | **7.0** | **14.7** | K.NMTVEQLLTGSPT*S*PTVEPEKPTR.E |
| **E2-230K** | UBE2O | T834T838 | **20.1** | **23.8** | K.NMTVEQLLT*GSPT*SPTVEPEKPTR.E |
| **E2-230K** | UBE2O | S836T838 | **1.9** | **25.4** | K.NMTVEQLLTGS*PT*SPTVEPEKPTR.E |
| **E2-230K** | UBE2O | S515 | **42.8** | **33.1** | R.KKS*IPLSIK.N |
| **E2-230K** | UBE2O | T834 | **7.8** | **15.4** | K.NMTVEQLLT*GSPTSPTVEPEKPTR.E |
| **E3 ubiquitin-protein ligase RNF213** | RNF213 | S1307 | **51.3** | **77.2** | K.EDQEAAELLS*EPEEESER.H |
| **(E3-independent) E2 ubiquitin-conjugating enzyme** | UBE2O | S1166S1167 | | **26.5** | K. [illegible] |
| **(E3-independent) E2 ubiquitin-conjugating enzyme** | UBE2O | S1166S1167S11 | | **16.6** | K. [illegible] |
| **E6 targeted protein 1** | SIPA1L1 | S258 | **-0.4** | **46.1** | K.GSGFS*LDVIDGPISQR.E |
| **E6 targeted protein 1** | SIPA1L1 | S208S211 | **1.8** | **58.4** | R.EYGS*TSS*IDKQGTSGESFFDLLK.G |
| **E6 targeted protein 1** | SIPA1L1 | T209S210 | **5.0** | **58.4** | R.EYGST*S*SIDKQGTSGESFFDLLK.G |
| **E6 targeted protein 1** | SIPA1L1 | Y206S208 | **13.6** | **12.6** | R.EY*GS*TSSIDKQGTSGESFFDLLK.G |
| **EBNA2 coactivator p100** | SND1 | S426 | **24.1** | **77.0** | K.VNVTVDYIRPAS*PATETVPAFSER.T |
| **Echinoderm microtubule associated protein like 3** | EML3 | S176 | **33.8** | **79.5** | K.AIS*SANLLVR.S |

Peak Area: <-10, -5, -3, 0, 3, 5, >10

%CV: 0, 29, 42, 71, 86, >100

**White dots-** Significant change in peptide abundance at **5** %FDR compared to the timepoint with the minimum peak area for a given PSM

CarT: 0m 2m 5m; RajiB: 0m 2m 5m

| Protein Name | Gene | Phosphosites | Ascor | MOWSE | Sequence |
|---|---|---|---|---|---|
| Echinoderm microtubule-associated protein-like 2 | EML2 | S136 | 44.7 | 25.0 | R.YLS*PER.L |
| EF hand calcium binding protein 1 | NECAB1 | Y121S125T132 | 100.0 | 12.0 | K.KDY*QEAS*NLEQFVT*RFLLK.E |
| EF hand domain family member D2 | EFHD2 | S74 | 6.9 | 22.2 | R.ADLNQGIGEPQS*PSR.R |
| EH domain binding protein 1-like 1 | EHBP1L1 | S173S176S180 | 0.8 | 10.8 | R.[illegible] |
| EH domain binding protein 1-like 1 | EHBP1L1 | S170S173S191 | 7.0 | 30.4 | R.[illegible] |
| EH domain binding protein 1-like 1 | EHBP1L1 | T164S170S173 | | 15.6 | R.[illegible] |
| EH domain containing 1 | EHD1 | S456 | 16.1 | 32.5 | K.DKPTYDEIFYTLS*PVNGK.I |
| eIF 4G1 | EIF4G1 | S1185 | 32.2 | 77.5 | R.S*FSKEVEER.S |
| eIF 4G1 | EIF4G1 | S1080 | 9.5 | 28.3 | K.ITKPGSIDS*NNQLFAPGGR.L |
| eIF 4G1 | EIF4G1 | S704 | 100.0 | 31.8 | R.S*QQGPR.K |
| | EIF4G1 | S1209 | 42.4 | 57.2 | K.AAS*LTEDR.D |
| eIF 4G1 | EIF4G1 | S1077 | 15.8 | 40.7 | K.ITKPGS*IDSNNQLFAPGGR.L |
| eIF 4G1 | EIF4G1 | S236S253S255 | | 17.4 | R.[illegible] |
| eIF 4G1 | EIF4G1 | S1185S1187 | 100.0 | 46.3 | R.S*FS*KEVEER.S |
| eIF 4G1 | EIF4G1 | S1596 | 100.0 | 37.9 | R.EAEEES*DHN.- |
| eIF 4G1 | EIF4G1 | T1073 | 8.5 | 93.4 | K.IT*KPGSIDSNNQLFAPGGR.L |
| eIF-2B epsilon | EIF2B5 | S544 | 100.0 | 62.1 | R.GGS*PQMDDIK.V |
| eIF-2B epsilon | EIF2B5 | S717S718 | 100.0 | 16.4 | K.EAEEES*S*EDD.- |
| eIF-2B epsilon | EIF2B5 | S429T432 | | 15.8 | R.[illegible] |
| eIF2beta | EIF2S2 | S105 | 20.8 | 68.7 | K.IES*DVQEPTEPEDDLDIMLGNK.K |
| eIF2beta | EIF2S2 | T111 | 48.0 | 71.6 | K.IESDVQEPT*EPEDDLDIMLGNK.K |
| EIF3S4 | EIF3G | T41 | 14.5 | 31.9 | K.GIPLATGDT*SPEPELLPGAPLPPPK.E |
| EIF3S4 | EIF3G | T38T41 | 5.3 | 19.3 | K.GIPLAT*GDT*SPEPELLPGAPLPPPK.E |
| EIF3S4 | EIF3G | S42 | 5.8 | 30.2 | K.GIPLATGDTS*PEPELLPGAPLPPPK.E |
| EIF3S4 | EIF3G | T41S42 | 14.7 | 39.9 | K.GIPLATGDT*S*PEPELLPGAPLPPPK.E |
| EIF3S4 | EIF3G | T38S42 | -1.4 | 21.4 | K.GIPLAT*GDTS*PEPELLPGAPLPPPK.E |
| EIF3S5 | EIF3F | S258 | 51.4 | 27.0 | K.TCFS*PNR.V |
| EIF3S5 | EIF3F | T255 | 9.2 | 28.4 | K.T*CFSPNR.V |
| EIF3S7 | EIF3D | S160 | 7.0 | 28.4 | R.DS*SVEVR.S |
| EIF3S8 | EIF3C | S39 | 67.6 | 57.6 | K.QPLLLS*EDEEDTKR.V |
| EIF3S8 | EIF3C | T7S9 | 7.9 | 15.0 | R.FFTT*GS*DSESESSLSGEELVTKPVGGNYGK.Q |
| EIF3S8 | EIF3C | S166S178S181 | 68.9 | 68.8 | K.QNPEQS*ADEDAEKNEEDS*EGS*S*DEDEDEDGVSAATFLK.K |
| EIF3S8 | EIF3C | S178S181S182 | 69.9 | 54.3 | K.NEEDS*EGS*S*DEDEDEDGVSAATFLK.K |
| EIF3S8 | EIF3C | S15S16 | 7.7 | 13.8 | R.FFTTGSDSESES*S*LSGEELVTKPVGGNYGK.Q |
| EIF3S8 | EIF3C | S9S11 | 3.6 | 77.2 | R.FFTTGS*DS*ESESSLSGEELVTKPVGGNYGK.Q |
| EIF3S8 | EIF3C | S166 | 100.0 | 49.0 | K.QNPEQS*ADEDAEK.N |
| EIF4G2 | EIF4G2 | S395 | 32.3 | 40.7 | R.FS*PTMGR.H |
| EIF4G2 | EIF4G2 | T896 | 6.9 | 25.2 | K.ALFQVNQWLTWLET*AEEEESEEEAD.- |
| EIF4G2 | EIF4G2 | T508 | 19.9 | 60.6 | R.TQT*PPLGQTPQLGLK.T |
| EIF4G2 | EIF4G2 | S381 | | 28.0 | K.MDRDPLGGLADMFGQMPGS*GIGTGPGVIQDRFSPTMGR.H |
| EIF4G2 | EIF4G2 | T397 | 28.6 | 39.0 | R.DPLGGLADMFGQMPGSGIGTGPGVIQDRFSPT*MGR.H |
| EIF4G2 | EIF4G2 | S902 | 4.3 | 17.0 | K.ALFQVNQWLTWLETAEEEES*EEEAD.- |
| EIF4G3 | EIF4G3 | S495 | 96.1 | 55.3 | R.S*PVPAQIAITVPK.T |
| ELAC | ELAC2 | S208S212S213 | 2.2 | 17.1 | R.LS*PERS*S*DS*ESNENEPHLPHGVSQR.R |
| ELAC | ELAC2 | S199 | 41.3 | 19.4 | K.HQPWQS*PERPLSR.L |
| ELAC | ELAC2 | S208S213S215 | 13.4 | 18.9 | R.LS*PERSS*DS*ES*NENEPHLPHGVSQR.R |
| ELAC | ELAC2 | S208S212S215 | 16.2 | 16.3 | R.LS*PERS*SDS*ES*NENEPHLPHGVSQR.R |
| ELAC | ELAC2 | S212S213S215 | -0.5 | 14.4 | R.LSPERS*S*DS*ES*NENEPHLPHGVSQR.R |
| ELF4 | ELF4 | S641S648 | 8.4 | 10.9 | R.S*PTPAPFS*PFNPTSLIK.M |
| ELL associated factor 1 | EAF1 | T157S165 | 22.6 | 40.8 | K.T*SPLKDNPS*PEPQLDDIKR.E |
| ELL associated factor 1 | EAF1 | S158S165 | 19.9 | 38.4 | K.TS*PLKDNPS*PEPQLDDIKR.E |
| Embryonic lethal abnormal vision like 1 | ELAVL1 | S202 | 35.4 | 96.9 | K.NVALLSQLYHS*PAR.R |
| Embryonic lethal abnormal vision like 1 | ELAVL1 | Y200 | 17.2 | 48.3 | K.NVALLSQLY*HSPAR.R |
| Embryonic lethal abnormal vision like 1 | ELAVL1 | S197 | 15.7 | 28.9 | K.NVALLS*QLYHSPAR.R |
| Emerin | EMD | Y167 | 4.8 | 39.8 | R.DSAYQSITHY*RPVSASR.S |
| Emerin | EMD | S141S143 | 8.5 | 13.9 | R.QSVTSFPDADAFHHQVHDDDLLS*SS*EEECKDR.E |
| Emerin | EMD | S120S141S142 | 3.1 | 13.8 | R.QS*VTSFPDADAFHHQVHDDDLLS*S*SEEECKDR.E |
| Emerin | EMD | T165 | 4.1 | 43.4 | R.DSAYQSIT*HYRPVSASR.S |
| Emerin | EMD | S159 | 4.2 | 29.0 | R.DS*AYQSITHYRPVSASR.S |
| Emerin | EMD | Y161 | 14.9 | 33.8 | R.DSAY*QSITHYRPVSASR.S |
| Emerin | EMD | S163 | 15.0 | 39.8 | R.DSAYQS*ITHYRPVSASR.S |
| Emerin | EMD | S141S142 | 24.7 | 22.4 | R.QSVTSFPDADAFHHQVHDDDLLS*S*SEEECKDR.E |

**Peak Area**: <-10, -5, -3, 0, 3, 5, >10

**%CV**: 0, 29, 42, 71, 86, >100

**White dots-** Significant change in peptide abundance at **5** %FDR compared to the timepoint with the minimum peak area for a given PSM

| Protein Name | Gene | Phosphosites | CarT 0m | CarT 2m | CarT 5m | RajiB 0m | RajiB 2m | RajiB 5m | Ascor | MOWSE | Sequence |
|---|---|---|---|---|---|---|---|---|---|---|---|
| Emerin | EMD | S53 | | | | | | | | 67.0 | R.RLSPPSS*SAASSYSFSDLNSTR.G |
| Emerin | EMD | S171 | | | | | | | 16.1 | 38.1 | R.DSAYQSITHYRPVS*ASR.S |
| Emerin | EMD | S60 | | | | | | | 6.8 | 50.5 | R.LSPPSSSAASSYS*FSDLNSTR.G |
| Emerin | EMD | Y59 | | | | | | | 6.5 | 68.1 | R.LSPPSSSAASSY*SFSDLNSTR.G |
| Emerin | EMD | S49 | | | | | | | 9.8 | 62.7 | R.RLS*PPSSSAASSYSFSDLNSTR.G |
| Emerin | EMD | S173 | | | | | | | 14.9 | 25.8 | R.DSAYQSITHYRPVSAS*R.S |
| Emerin | EMD | S123S141S143 | | | | | | | 4.0 | 12.5 | R.QSVTS*FPDADAFHHQVHDDDLLS*SS*EEECKDR.E |
| Emerin | EMD | S141S142S143 | | | | | | | 14.4 | 14.3 | R.QSVTSFPDADAFHHQVHDDDLLS*S*S*EEECKDR.E |
| Emerin | EMD | S176 | | | | | | | -0.4 | 54.3 | R.SS*LDLSYYPTSSSTSFMSSSSSSSSWLTR.R |
| EMK1 | MARK2 | S452 | | | | | | | 32.1 | 36.1 | R.SRNS*PLLER.A |
| EML4 | EML4 | S144S146 | | | | | | | 24.5 | 65.4 | R.AS*PS*PQPSSQPLQIHR.Q |
| EML4 | EML4 | S895T899 | | | | | | | 34.3 | 83.9 | K.APVSSTESVIQS*NTPT*PPPSQPLNETAEEESR.I |
| EML4 | EML4 | T897T899 | | | | | | | 37.1 | 88.9 | K.APVSSTESVIQSNT*PT*PPPSQPLNETAEEESR.I |
| EML4 | EML4 | S144 | | | | | | | 16.1 | 42.1 | R.AS*PSPQPSSQPLQIHR.Q |
| EML4 | EML4 | S919S934S938 | | | | | | | 4.2 | 55.6 | R. ISSS*PTLLENSLEQTVEPS*EDHS*EEESEEGSGDLGEPLYEEPCNEI |
| EML4 | EML4 | S917S918S919 | | | | | | | | 39.0 | R. IS*S*S*PTLLENSLEQTVEPSEDHSEEESEEGSGDLGEPLYEEPCNEI |
| EML4 | EML4 | S917S918S919 | | | | | | | | 21.3 | R. IS*S*S*PT*LLENSLEQTVEPSEDHSEEESEEGSGDLGEPLYEEPCN |
| EMSY Protein | EMSY | S168S173 | | | | | | | 13.8 | 24.7 | K.S*PRPAS*PASNVVVLPSGSTVYVK.S |
| EMSY Protein | EMSY | S173S176 | | | | | | | 6.1 | 20.9 | K.SPRPAS*PAS*NVVVLPSGSTVYVK.S |
| EMSY Protein | EMSY | S168S176 | | | | | | | 9.4 | 32.7 | K.S*PRPASPAS*NVVVLPSGSTVYVK.S |
| Ena vasodilator stimulated phosphoprotein like protein | EVL | S262 | | | | | | | 25.3 | 94.0 | K.SDANRASS*GGGGGGLMEEMNK.L |
| Ena vasodilator stimulated phosphoprotein like protein | EVL | S261 | | | | | | | 16.9 | 91.0 | R.AS*SGGGGGGLMEEMNK.L |
| Ena vasodilator stimulated phosphoprotein like protein | EVL | S333 | | | | | | | 20.4 | 75.2 | R.SNS*VEKPVSSILSR.T |
| Ena vasodilator stimulated phosphoprotein like protein | EVL | S333S343 | | | | | | | 9.3 | 54.9 | R.SNS*VEKPVSSILS*R.T |
| Ena vasodilator stimulated phosphoprotein like protein | EVL | S371 | | | | | | | 100.0 | 82.1 | R.MKPAGS*VNDMALDAFDLDR.M |
| Ena vasodilator stimulated phosphoprotein like protein | EVL | S347S351 | | | | | | | 33.2 | 36.2 | R.TPS*VAKS*PEAK.S |
| Ena vasodilator stimulated phosphoprotein like protein | EVL | S306 | | | | | | | 7.0 | 57.9 | K.KEDESQMEDPSTS*PSPGTR.A |
| Endonuclease III-like1 | NTHL1 | S71 | | | | | | | 12.4 | 50.7 | R.VAYEGS*DSEKGEGAEPLKVPVWEPQDWQQQLVNIR.A |
| Endonuclease III-like1 | NTHL1 | S73 | | | | | | | -0.4 | 27.9 | R.VAYEGSDS*EKGEGAEPLKVPVWEPQDWQQQLVNIR.A |
| | ENSA | S109 | | | | | | | 13.9 | 67.1 | R.KSS*LVTSK.L |
| Endothelial differentiation gene 1 | S1PR1 | S351 | | | | | | | 100.0 | 36.3 | K.RPIIAGMEFS*R.S |
| Endothelial differentiation gene 5 | S1PR2 | S330S331S332 | | | | | | | 39.5 | 18.7 | R.GGTPGHHLLPLRS*S*S*SLER.G |
| Endothelial differentiation gene 5 | S1PR2 | S331S332 | | | | | | | -0.3 | 11.8 | R.SS*S*SLER.G |
| Endothelial differentiation gene 5 | S1PR2 | S330S331 | | | | | | | 53.4 | 13.3 | R.GGTPGHHLLPLRS*S*SSLER.G |
| Endothelial differentiation gene 5 | S1PR2 | S330S333 | | | | | | | 9.5 | 15.4 | R.GGTPGHHLLPLRS*SSS*LER.G |
| Endothelial differentiation gene 5 | S1PR2 | S333 | | | | | | | 6.5 | 23.0 | R.SSSS*LER.G |
| Endothelial differentiation gene 5 | S1PR2 | S331S332S333 | | | | | | | 2.2 | 11.0 | R.GGTPGHHLLPLRSS*S*S*LER.G |
| Endothelial differentiation gene 5 | S1PR2 | S332S333 | | | | | | | 6.1 | 12.1 | R.SSS*S*LER.G |
| Endothelial differentiation gene 5 | S1PR2 | S332 | | | | | | | 12.6 | 12.9 | R.SSS*SLER.G |
| Endothelin converting enzyme 1 | ECE1 | T25S34 | | | | | | | 20.8 | 49.8 | R.AT*LDEEDLVDS*LSEGDAYPNGLQVNFHSPR.S |
| Endothelin converting enzyme 1 | ECE1 | T25S36 | | | | | | | 8.5 | 43.6 | R.AT*LDEEDLVDSLS*EGDAYPNGLQVNFHSPR.S |
| Endothelin converting enzyme 1 | ECE1 | S34 | | | | | | | 6.0 | 46.2 | R.ATLDEEDLVDS*LSEGDAYPNGLQVNFHSPR.S |
| Endothelin converting enzyme 1 | ECE1 | S36 | | | | | | | 4.2 | 65.9 | R.ATLDEEDLVDSLS*EGDAYPNGLQVNFHSPR.S |
| Engulfment and cell motility 1 | ELMO1 | S342 | | | | | | | 12.1 | 61.9 | R.IAFDAESEPNNSS*GSMEK.R |
| Enhancer of mRNA decapping 3 homolog | EDC3 | S131 | | | | | | | 53.2 | 80.4 | K.SQDVAVS*PQQQQCSK.S |
| Enolase 1 | ENO1 | Y257 | | | | | | | 2.1 | 47.6 | R.SGKY*DLDFK.S |
| Enolase 1 | ENO1 | S254 | | | | | | | 12.2 | 24.7 | R.S*GKYDLDFK.S |
| Enolase 1 | ENO1 | S419 | | | | | | | 100.0 | 25.9 | R.IEEELGS*K.A |
| Enolase 1 | ENO1 | T26 | | | | | | | 40.4 | 21.6 | R.GNPTVEVDLFT*SK.G |
| Enoyl CoA hydratase, peroxisomal | ECH1 | S139 | | | | | | | 22.4 | 16.2 | R.IS*WYLR.D |
| EP4 receptor | PTGER4 | S390T391 | | | | | | | 8.0 | 42.3 | K.EISS*T*SQTLLPDLSLPDLSENGLGGR.N |
| EP4 receptor | PTGER4 | S390S400 | | | | | | | 2.3 | 20.0 | K.EISS*TSQTLLPDLS*LPDLSENGLGGR.N |
| Eplin | LIMA1 | S686 | | | | | | | 50.3 | 117.3 | K.EGHSLEMENENLVENGADS*DEDNSFLK.Q |
| Eps15 | EPS15 | S324 | | | | | | | 9.8 | 104.2 | K.NIIGSS*PVADFSAIK.E |
| Eps15 | EPS15 | S851 | | | | | | | 49.2 | 67.7 | K.EADPSNFANFSAYPS*EEDMIEWAK.R |
| Eps15 | EPS15 | S323 | | | | | | | 55.8 | 59.6 | K.NIIGS*SPVADFSAIK.E |
| Eps15 | EPS15 | S796S814 | | | | | | | 59.4 | 22.6 | R.SINKLDS*PDPFKLNDPFQPFPGNDS*PK.E |
| Eps15 | EPS15 | S605 | | | | | | | | 50.7 | K.EEDPFNVDS*SSLTGPVADTNLDFFQSDPFVGSDPFKDDPFGK.I |
| Eps15 | EPS15 | S563 | | | | | | | -0.5 | 54.3 | R.SS*PELLPSGVTDENEVTTAVTEK.V |
| Eps15 | EPS15 | S569 | | | | | | | 13.7 | 21.9 | R.SSPELLPS*GVTDENEVTTAVTEK.V |
| Eps15 | EPS15 | S606 | | | | | | | -0.0 | 32.9 | K.EEDPFNVDSS*SLTGPVADTNLDFFQSDPFVGSDPFKDDPFGK.I |
| Eps15 | EPS15 | S562 | | | | | | | 11.5 | 76.5 | R.S*SPELLPSGVTDENEVTTAVTEK.V |

Peak Area: <-10, -5, -3, 0, 3, 5, >10

%CV: 0, 29, 42, 71, 86, >100

**White dots-** Significant change in peptide abundance at **5**%FDR compared to the timepoint with the minimum peak area for a given PSM

| Protein Name | Gene | Phosphosites | CarT 0m | CarT 2m | CarT 5m | RajiB 0m | RajiB 2m | RajiB 5m | Ascor | MOWSE | Sequence |
|---|---|---|---|---|---|---|---|---|---|---|---|
| **EPS15R** | EPS15L1 | S255 | | | | | | | **6.0** | **12.6** | R.STPSHGSVSSLNSTGSLS*PK.H |
| **EPS15R** | EPS15L1 | S246S255 | | | | | | | **7.8** | **44.5** | R.STPSHGSVS*SLNSTGSLS*PK.H |
| **EPS15R** | EPS15L1 | S575 | | | | | | | **6.0** | **29.7** | R.SLEQYDQVLDGAHGAS*LTDLANLSEGVSLAER.G |
| **EPS15R** | EPS15L1 | T577 | | | | | | | **8.9** | **60.1** | R.SLEQYDQVLDGAHGASLT*DLANLSEGVSLAER.G |
| **EPS15R** | EPS15L1 | S804 | | | | | | | **6.9** | **47.7** | K.STPVSQLGS*ADFPEAPDPFQPLGADSGDPFQSK.K |
| **EPS15R** | EPS15L1 | T797 | | | | | | | **-0.3** | **50.9** | K.ST*PVSQLGSADFPEAPDPFQPLGADSGDPFQSK.K |
| **EPS15R** | EPS15L1 | S253 | | | | | | | **25.4** | **18.6** | R.STPSHGSVSSLNSTGS*LSPK.H |
| **EPS15R** | EPS15L1 | S247S255 | | | | | | | **5.9** | **21.3** | R.STPSHGSVSS*LNSTGSLS*PK.H |
| **EPS15R** | EPS15L1 | S246S253 | | | | | | | **5.3** | **40.1** | R.STPSHGSVS*SLNSTGS*LSPK.H |
| **EPS15R** | EPS15L1 | S241S255 | | | | | | | **7.3** | **53.0** | R.STPS*HGSVSSLNSTGSLS*PK.H |
| **Epsin 1** | EPN1 | T444 | | | | | | | **53.6** | **61.0** | K.T*PESFLGPNAALVDLDSLVSRPGPTPPGAK.A |
| **Epsin 1** | EPN1 | T390 | | | | | | | **52.7** | **78.9** | R.TALPT*SGSSAGELELLAGEVPAR.S |
| **Epsin 1** | EPN1 | S391 | | | | | | | **24.2** | **34.4** | R.TALPTS*GSSAGELELLAGEVPAR.S |
| **Epsin 1** | EPN1 | S447 | | | | | | | **5.5** | **44.0** | K.TPES*FLGPNAALVDLDSLVSRPGPTPPGAK.A |
| **Epsin 1** | EPN1 | T444T468 | | | | | | | **31.5** | **14.9** | K.T*PESFLGPNAALVDLDSLVSRPGPT*PPGAK.A |
| **Epsin 1** | EPN1 | S409 | | | | | | | **99.0** | **64.1** | R.S*PGAFDMSGVR.G |
| **Epsin 1** | EPN1 | S393 | | | | | | | **10.7** | **28.7** | R.TALPTSGS*SAGELELLAGEVPAR.S |
| **Epsin 1** | EPN1 | S394 | | | | | | | **15.7** | **24.0** | R.TALPTSGSS*AGELELLAGEVPAR.S |
| **Epsin 4** | CLINT1 | S299 | | | | | | | **-3.9** | **12.8** | K.TIDLGAAAHYTGDKAS*PDQNASTHTPQSSVK.T |
| **Epsin 4** | CLINT1 | T294 | | | | | | | **6.5** | **19.3** | K.TIDLGAAAHYT*GDKASPDQNASTHTPQSSVK.T |
| **Epsin 4** | CLINT1 | S305 | | | | | | | **7.4** | **11.1** | K.TIDLGAAAHYTGDKASPDQNAS*THTPQSSVK.T |
| **Epstein Barr virus induced gene 2** | GPR183 | S328S330 | | | | | | | **24.2** | **63.1** | K.RQVS*VS*ISSAVK.S |
| **Epstein Barr virus induced gene 2** | GPR183 | S330S333 | | | | | | | **13.4** | **34.4** | R.QVSVS*ISS*AVK.S |
| **Epstein Barr virus induced gene 2** | GPR183 | S328S330S333 | | | | | | | **17.8** | **31.3** | R.QVS*VS*ISS*AVK.S |
| **Epstein Barr virus induced gene 2** | GPR183 | S337 | | | | | | | **27.8** | **22.1** | K.S*APEENSR.E |
| **Epstein Barr virus induced gene 2** | GPR183 | S328S333 | | | | | | | **9.9** | **61.5** | K.RQVS*VSISS*AVK.S |
| **Epstein Barr virus induced gene 2** | GPR183 | S333 | | | | | | | **8.2** | **53.6** | R.QVSVSISS*AVK.S |
| **Epstein Barr virus induced gene 2** | GPR183 | S328 | | | | | | | **35.4** | **83.4** | R.QVS*VSISSAVK.S |
| **Epstein Barr virus induced gene 2** | GPR183 | S343 | | | | | | | **9.5** | **17.2** | K.SAPEENS*R.E |
| **Epstein Barr virus induced gene 2** | GPR183 | S332S333 | | | | | | | **5.4** | **35.1** | R.QVSVSIS*S*AVK.S |
| **Epstein Barr virus induced gene 2** | GPR183 | S332 | | | | | | | **16.1** | **52.4** | R.QVSVSIS*SAVK.S |
| **Epstein Barr virus induced gene 2** | GPR183 | S328S330S332 | | | | | | | **10.4** | **31.9** | R.QVS*VS*IS*SAVK.S |
| **Epstein Barr virus induced gene 2** | GPR183 | S330S332S333 | | | | | | | **7.9** | **13.3** | R.QVSVS*IS*S*AVK.S |
| **Erbin** | ERBIN | S602S603 | | | | | | | **5.5** | **69.8** | K.HIVNHDDVFEESEELS*S*DEEMK.M |
| **Erbin** | ERBIN | S598S602 | | | | | | | **12.5** | **57.7** | K.HIVNHDDVFEES*EELS*SDEEMK.M |
| | ERBIN | T917S932 | | | | | | | **15.0** | **57.8** | R.SKSAT*LLYDQPLQVFTGSSS*SSDLISGTK.A |
| **Erbin** | ERBIN | S915S932 | | | | | | | **12.9** | **49.8** | K.S*ATLLYDQPLQVFTGSSS*SSDLISGTK.A |
| **Erbin** | ERBIN | S915 | | | | | | | **20.4** | **108.1** | K.S*ATLLYDQPLQVFTGSSSSSDLISGTK.A |
| **Erbin** | ERBIN | S915S933 | | | | | | | **14.1** | **54.8** | K.S*ATLLYDQPLQVFTGSSSS*SDLISGTK.A |
| **Erbin** | ERBIN | T917S933 | | | | | | | **13.6** | **49.2** | K.SAT*LLYDQPLQVFTGSSSS*SDLISGTK.A |
| **Erbin** | ERBIN | T917S930 | | | | | | | **12.2** | **39.5** | R.SKSAT*LLYDQPLQVFTGS*SSSSDLISGTK.A |
| **Erbin** | ERBIN | T917 | | | | | | | **12.2** | **41.4** | R.SKSAT*LLYDQPLQVFTGSSSSSDLISGTK.A |
| **Erbin** | ERBIN | S440 | | | | | | | **3.9** | **53.8** | R.TEDVMFIS*DNESFNPSLWEEQR.K |
| **Erbin** | ERBIN | S440S444 | | | | | | | **30.8** | **34.9** | R.TEDVMFIS*DNES*FNPSLWEEQR.K |
| **ERCC5** | BIVM- | S384 | | | | | | | **9.1** | **64.4** | R.NAPAAVDEGSIS*PR.T |
| **ERCC5** | BIVM- | S562S563 | | | | | | | **31.2** | **30.6** | K.FDSSLLS*S*DDETK.C |
| **ERCC5** | BIVM- | S156S157 | | | | | | | **41.7** | **16.9** | R.ENDLYVLPPLQEEEKHS*S*EEEDEKEWQER.M |
| **ERF** | ERF | Y16 | | | | | | | **6.0** | **34.9** | K.TPADTGFAFPDWAY*KPESSPGSR.Q |
| **ERF** | | T3S21 | | | | | | | **35.7** | **36.7** | -.MKT*PADTGFAFPDWAYKPESS*PGSR.Q |
| **ERF** | | T3S24 | | | | | | | **38.0** | **38.5** | -.MKT*PADTGFAFPDWAYKPESSPGS*R.Q |
| **ERF** | | T3S20 | | | | | | | **26.4** | **12.7** | -.MKT*PADTGFAFPDWAYKPES*SPGSR.Q |
| **ERF** | ERF | S20 | | | | | | | **7.1** | **23.9** | K.TPADTGFAFPDWAYKPES*SPGSR.Q |
| **ERK1** | MAPK3 | Y204 | | | | | | | **25.5** | **98.4** | R.IADPEHDHTGFLTEY*VATR.W |
| **ERK1** | MAPK3 | T202Y204 | | | | | | | **38.3** | **92.7** | R.IADPEHDHTGFLT*EY*VATR.W |
| **ERK1** | MAPK3 | T198Y204 | | | | | | | **11.9** | **33.6** | R.IADPEHDHT*GFLTEY*VATR.W |
| **ERK2** | MAPK1 | Y187 | | | | | | | **27.8** | **98.3** | R.VADPDHDHTGFLTEY*VATR.W |
| **ERK2** | MAPK1 | T185Y187 | | | | | | | **49.5** | **88.6** | R.VADPDHDHTGFLT*EY*VATR.W |
| **ERK2** | MAPK1 | T181Y187 | | | | | | | **13.2** | **18.6** | R.VADPDHDHT*GFLTEY*VATR.W |
| **Erythrocyte membrane protein band 4.1** | EPB41 | S684 | | | | | | | **20.7** | **40.8** | K.HHAS*ISELKK.N |
| **Erythrocyte membrane protein band 4.1** | EPB41 | S709 | | | | | | | **18.0** | **21.8** | K.RLS*THSPFR.T |
| **Erythrocyte membrane protein band 4.1** | EPB41 | S84 | | | | | | | **9.8** | **31.9** | R.LFS*SFLK.R |
| **Erythrocyte membrane protein band 4.1** | EPB41 | S84S85 | | | | | | | **100.0** | **34.2** | R.LFS*S*FLK.R |

Peak Area: <-10, -5, -3, 0, 3, 5, >10

%CV: 0, 29, 43, 71, 86, >100

**White dots-** Significant change in peptide abundance at 5 %FDR compared to the timepoint with the minimum peak area for a given PSM

| Protein Name | Gene | Phosphosites | CarT 0m | CarT 2m | CarT 5m | RajiB 0m | RajiB 2m | RajiB 5m | Ascor | MOWSE | Sequence |
|---|---|---|---|---|---|---|---|---|---|---|---|
| Erythrocyte membrane protein band 4.1 | EPB41 | T710 | | | | | | | 6.5 | 20.3 | K.RLST*HSPFR.T |
| Erythrocyte membrane protein band 4.1 | EPB41 | S510 | | | | | | | 8.9 | 14.6 | K.FRYS*GR.T |
| Erythrocyte membrane protein band 4.1 | EPB41 | Y509 | | | | | | | | 14.1 | K.FRY*SGR.T |
| Erythrocyte membrane protein band 4.1 | EPB41 | S85 | | | | | | | 9.8 | 32.5 | R.LFSS*FLK.R |
| Erythrocyte membrane protein band 4.1-like 2 | EPB41L2 | S87 | | | | | | | 16.9 | 46.8 | K.QKS*YTLVVAK.D |
| Erythrocyte membrane protein band 4.1-like 2 | EPB41L2 | T600 | | | | | | | 5.8 | 60.1 | R.SPT*KAPHLQLIEGK.K |
| Erythrocyte membrane protein band 4.1-like 2 | EPB41L2 | Y88 | | | | | | | -0.2 | 34.1 | K.SY*TLVVAK.D |
| Erythrocyte membrane protein band 4.1-like 2 | EPB41L2 | S598 | | | | | | | 22.5 | 65.8 | R.S*PTKAPHLQLIEGK.K |
| Essential meiotic endonuclease 1 | EME1 | S85S87 | | | | | | | 17.6 | 90.2 | R.LLSS*ES*EDEEEFIPLAQR.L |
| Essential meiotic endonuclease 1 | EME1 | S84S85S87 | | | | | | | 100.0 | 56.8 | R.LLS*S*ES*EDEEEFIPLAQR.L |
| Essential meiotic endonuclease 1 | EME1 | S84S87 | | | | | | | 12.1 | 20.2 | R.LLS*SES*EDEEEFIPLAQR.L |
| Estrogen receptor binding protein | DNTTIP2 | S141S148 | | | | | | | 13.0 | 63.8 | K.ESYTEEIVS*EAESHVS*GISR.I |
| Estrogen receptor binding protein | DNTTIP2 | S141S145S148 | | | | | | | 11.8 | 41.2 | K.ESYTEEIVS*EAES*HVS*GISR.I |
| Estrogen receptor binding protein | DNTTIP2 | S528S532S533 | | | | | | | 44.6 | 45.4 | K.EEEEDEKS*EEDS*S*DHDENEDEFS*DEEDFLNSTK.A |
| Estrogen receptor binding protein | DNTTIP2 | S141S145 | | | | | | | 21.8 | 60.3 | K.ESYTEEIVS*EAES*HVSGISR.I |
| Estrogen receptor binding protein | DNTTIP2 | Y135S145 | | | | | | | 12.9 | 42.4 | K.ESY*TEEIVSEAES*HVSGISR.I |
| Estrogen receptor binding protein | DNTTIP2 | S141 | | | | | | | 20.7 | 56.0 | K.ESYTEEIVS*EAESHVSGISR.I |
| Estrogen receptor binding protein | DNTTIP2 | S141S151 | | | | | | | 3.4 | 17.3 | K.ESYTEEIVS*EAESHVSGIS*R.I |
| Estrogen receptor binding protein | DNTTIP2 | Y135S145S148 | | | | | | | 8.2 | 38.5 | K.ESY*TEEIVSEAES*HVS*GISR.I |
| Estrogen receptor binding protein | DNTTIP2 | T136S145S148 | | | | | | | 13.0 | 29.7 | K.ESYT*EEIVSEAES*HVS*GISR.I |
| Estrogen receptor binding protein | DNTTIP2 | S134Y135S151 | | | | | | | -2.3 | 40.8 | K.ES*Y*TEEIVSEAESHVSGIS*R.I |
| Estrogen receptor binding protein | DNTTIP2 | T136S148 | | | | | | | 5.5 | 28.2 | K.ESYT*EEIVSEAESHVS*GISR.I |
| ETS translocation variant 6 | ETV6 | Y233S251 | | | | | | | 12.1 | 15.4 | R.AQGPRPHQENNHQESY*PLSVSPMENNHCPASSES*HPKPSSPR.[illegible] |
| ETS translocation variant 6 | ETV6 | S203 | | | | | | | 100.0 | 40.7 | R.S*PLDNMIR.R |
| ETS translocation variant 6 | ETV6 | T18S22 | | | | | | | 36.4 | 68.3 | R.ISYT*PPES*PVPSYASSTPLHVPVPR.A |
| ETS translocation variant 6 | ETV6 | S16S22 | | | | | | | 15.1 | 38.1 | R.IS*YTPPES*PVPSYASSTPLHVPVPR.A |
| ETS translocation variant 6 | ETV6 | Y17S22S29 | | | | | | | 22.9 | 54.0 | R.ISY*TPPES*PVPSYAS*STPLHVPVPR.A |
| ETS translocation variant 6 | ETV6 | Y17S22T31 | | | | | | | 22.5 | 41.3 | R.ISY*TPPES*PVPSYASST*PLHVPVPR.A |
| ETS translocation variant 6 | ETV6 | T161 | | | | | | | 3.7 | 28.5 | R.T*PRPSVDNVHHNPPTIELLHR.S |
| ETS translocation variant 6 | ETV6 | Y17S26 | | | | | | | 12.4 | 52.1 | R.ISY*TPPESPVPS*YASSTPLHVPVPR.A |
| ETS translocation variant 6 | ETV6 | Y17T18 | | | | | | | 2.7 | 21.9 | R.ISY*T*PPESPVPSYASSTPLHVPVPR.A |
| ETS translocation variant 6 | ETV6 | S16S26 | | | | | | | 23.4 | 49.1 | R.IS*YTPPESPVPS*YASSTPLHVPVPR.A |
| ETS translocation variant 6 | ETV6 | S22S26 | | | | | | | 11.5 | 38.1 | R.ISYTPPES*PVPS*YASSTPLHVPVPR.A |
| ETS translocation variant 6 | ETV6 | Y17Y27S29 | | | | | | | 10.3 | 34.6 | R.ISY*TPPESPVPSY*AS*STPLHVPVPR.A |
| ETS translocation variant 6 | ETV6 | Y17S22 | | | | | | | 26.9 | 44.2 | R.ISY*TPPES*PVPSYASSTPLHVPVPR.A |
| ETS translocation variant 6 | ETV6 | S26Y27S29 | | | | | | | 30.2 | 32.7 | R.ISYTPPESPVPS*Y*AS*STPLHVPVPR.A |
| ETS translocation variant 6 | ETV6 | Y17S22S30 | | | | | | | 17.9 | 36.0 | R.ISY*TPPES*PVPSYASS*TPLHVPVPR.A |
| ETS translocation variant 6 | ETV6 | S248S251 | | | | | | | 16.9 | 15.3 | R.AQGPRPHQENNHQESYPLSVSPMENNHCPAS*SES*HPKPSSPR.[illegible] |
| ETS translocation variant 6 | ETV6 | S232Y233 | | | | | | | | 13.8 | R.AQGPRPHQENNHQES*Y*PLSVSPMENNHCPASSESHPKPSSPR.[illegible] |
| ETS translocation variant 6 | ETV6 | S232S257 | | | | | | | 27.1 | 16.4 | R.AQGPRPHQENNHQES*YPLSVSPMENNHCPASSESHPKPSS*PR.[illegible] |
| ETS translocation variant 6 | ETV6 | S213 | | | | | | | 100.0 | 27.2 | R.RLS*PAER.A |
| ETS translocation variant 6 | ETV6 | S165 | | | | | | | 3.3 | 19.1 | R.TPRPS*VDNVHHNPPTIELLHR.S |
| ETS translocation variant 6 | ETV6 | T18S22S29 | | | | | | | 10.7 | 41.4 | R.ISYT*PPES*PVPSYAS*STPLHVPVPR.A |
| ETS translocation variant 6 | ETV6 | S232S251 | | | | | | | 15.6 | 14.4 | R.AQGPRPHQENNHQES*YPLSVSPMENNHCPASSES*HPKPSSPR.[illegible] |
| ETS translocation variant 6 | ETV6 | Y233S256 | | | | | | | 11.0 | 10.8 | R.AQGPRPHQENNHQESY*PLSVSPMENNHCPASSESHPKPS*SPR.[illegible] |
| ETS translocation variant 6 | ETV6 | Y233S257 | | | | | | | 13.2 | 11.6 | R.AQGPRPHQENNHQESY*PLSVSPMENNHCPASSESHPKPSS*PR.[illegible] |
| ETS translocation variant 6 | ETV6 | S182S184 | | | | | | | 3.2 | 11.9 | R.S*RS*PITTNHRPSPDPEQRPLR.S |
| ETS1 | ETS1 | Y283 | | | | | | | 9.8 | 34.0 | R.VPSY*DSFDSEDYPAALPNHKPK.G |
| ETS1 | ETS1 | T38 | | | | | | | 7.3 | 29.2 | K.VDLELFPSPDMECADVPLLT*PSSK.E |
| ETS1 | ETS1 | S26T38 | | | | | | | 43.2 | 32.0 | K.VDLELFPS*PDMECADVPLLT*PSSK.E |
| ETS1 | ETS1 | S285 | | | | | | | 7.3 | 34.8 | R.VPSYDS*FDSEDYPAALPNHKPK.G |
| ETS1 | ETS1 | S282S285 | | | | | | | 22.9 | 28.6 | R.VPS*YDS*FDSEDYPAALPNHKPK.G |
| ETS1 | ETS1 | S282 | | | | | | | 7.4 | 15.2 | R.VPS*YDSFDSEDYPAALPNHKPK.G |
| ETS1 | ETS1 | S41 | | | | | | | 7.8 | 23.2 | K.VDLELFPSPDMECADVPLLTPSS*K.E |
| ETS1 | ETS1 | S40 | | | | | | | 3.5 | 20.0 | K.VDLELFPSPDMECADVPLLTPS*SK.E |
| Eukaryotic translation elongation factor 1 beta 2 | EEF1B2 | Y79 | | | | | | | | 79.6 | K.Y*GPADVEDTTGSGATDSKDDDDIDLFGSDDEEESEEAK.R |
| Eukaryotic translation elongation factor 1 beta 2 | EEF1B2 | S106 | | | | | | | 35.4 | 46.7 | K.YGPADVEDTTGSGATDSKDDDDIDLFGS*DDEEESEEAK.R |
| Eukaryotic translation elongation factor 1 beta 2 | EEF1B2 | S90S95 | | | | | | | 6.2 | 58.4 | K.YGPADVEDTTGS*GATDS*KDDDDIDLFGSDDEEESEEAK.R |
| Eukaryotic translation elongation factor 1 beta 2 | EEF1B2 | T88S106 | | | | | | | 12.0 | 46.5 | K.YGPADVEDTT*GSGATDSKDDDDIDLFGS*DDEEESEEAK.R |
| Eukaryotic translation elongation factor 1 beta 2 | EEF1B2 | S90 | | | | | | | 4.3 | 39.0 | K.YGPADVEDTTGS*GATDSKDDDDIDLFGSDDEEESEEAK.R |
| Eukaryotic translation elongation factor 1 beta 2 | EEF1B2 | S90T93 | | | | | | | 16.8 | 69.3 | K.YGPADVEDTTGS*GAT*DSKDDDDIDLFGSDDEEESEEAK.R |
| Eukaryotic translation elongation factor 1 beta 2 | EEF1B2 | S90S106 | | | | | | | 19.0 | 78.9 | K.YGPADVEDTTGS*GATDSKDDDDIDLFGS*DDEEESEEAK.R |

Peak Area: <-10, -5, -3, 0, 3, 5, >10

%CV: 0, 29, 42, 71, 86, >100

**White dots-** Significant change in peptide abundance at **5** %FDR compared to the timepoint with the minimum peak area for a given PSM

CarT (0m, 2m, 5m) — RajiB (0m, 2m, 5m)

| Protein Name | Gene | Phosphosites | Ascor | MOWSE | Sequence |
|---|---|---|---|---|---|
| Eukaryotic translation elongation factor 1 beta 2 | EEF1B2 | S95S106 | 18.6 | 69.1 | K.YGPADVEDTTGSGATDS*KDDDDIDLFGS*DDEEESEEAK.R |
| Eukaryotic translation elongation factor 1 beta 2 | EEF1B2 | S95 | 3.5 | 61.0 | K.YGPADVEDTTGSGATDS*KDDDDIDLFGSDDEEESEEAKR.L |
| Eukaryotic translation elongation factor 1 beta 2 | EEF1B2 | T93 | 2.4 | 13.5 | K.YGPADVEDTTGSGAT*DSKDDDDIDLFGSDDEEESEEAKR.L |
| Eukaryotic translation elongation factor 1 beta 2 | EEF1B2 | T87 | -0.0 | 28.1 | K.YGPADVEDT*TGSGATDSKDDDDIDLFGSDDEEESEEAK.R |
| Eukaryotic translation elongation factor 1 gamma | EEF1G | T46 | 100.0 | 40.0 | R.T*PEFLR.K |
| Eukaryotic translation elongation factor 1, delta | EEF1D | S528 | 161.4 | 105.0 | K.KPATPAEDDEDDDIDLFGS*DNEEEDK.E |
| Eukaryotic translation elongation factor 1, delta | EEF1D | T513S528 | 100.0 | 71.5 | K.KPAT*PAEDDEDDDIDLFGS*DNEEEDKEAAQLR.E |
| Eukaryotic translation elongation factor 1, delta | EEF1D | S499 | 19.1 | 37.0 | R.ATAPQTQHVS*PMR.Q |
| Eukaryotic translation elongation factor 1, delta | EEF1D | T495 | 5.3 | 26.8 | R.ATAPQT*QHVSPMR.Q |
| Eukaryotic translation elongation factor 1, delta | EEF1D | T491 | 11.0 | 58.2 | R.AT*APQTQHVSPMR.Q |
| Eukaryotic translation elongation factor 2 | EEF2 | T57T59 | 100.0 | 21.3 | R.FT*DT*RKDEQER.C |
| Eukaryotic translation initiation factor 2A | EIF2A | S503 | 6.3 | 38.8 | R.S*DKSPDLAPTPAPQSTPR.N |
| Eukaryotic translation initiation factor 2A | EIF2A | S506 | 6.5 | 37.4 | R.SDKS*PDLAPTPAPQSTPR.N |
| Eukaryotic translation initiation factor 3 subunit k | EIF3K | S217 | 11.2 | 23.5 | K.IDFDSVSSIMASS*Q.- |
| Eukaryotic translation initiation factor 3 subunit k | EIF3K | S216 | 60.7 | 25.4 | K.IDFDSVSSIMAS*SQ.- |
| Eukaryotic translation initiation factor 3, subunit 10 | EIF3A | S882 | 12.1 | 45.0 | R.LGDSS*LSR.K |
| Eukaryotic translation initiation factor 3, subunit 10 | EIF3A | S881 | 2.3 | 12.6 | R.LGDS*SLSR.K |
| Eukaryotic translation initiation factor 3, subunit 6 48kDa | EIF3E | S399 | 27.8 | 27.2 | K.LGHVVMGNNAVS*PYQQVIEK.T |
| Eukaryotic translation initiation factor 3, subunit 9 eta, 116kDa | EIF3B | T68 |  | 38.5 | R. [illegible] |
| Eukaryotic translation initiation factor 3, subunit 9 eta, 116kDa | EIF3B | T68S78 |  | 36.4 | R. [illegible] |
| Eukaryotic translation initiation factor 3, subunit 9 eta, 116kDa | EIF3B | S152S164 | 13.8 | 43.9 | R.ALENGDADEPS*FSDPEDFVDDVS*EEELLGDVLK.D |
| Eukaryotic translation initiation factor 3, subunit 9 eta, 116kDa | EIF3B | S152 | 78.9 | 103.5 | R.ALENGDADEPS*FSDPEDFVDDVSEEELLGDVLK.D |
| Eukaryotic translation initiation factor 3, subunit 9 eta, 116kDa | EIF3B | T68S78S81S83 |  | 30.9 | R. [illegible] |
| Eukaryotic translation initiation factor 3, subunit 9 eta, 116kDa | EIF3B | S154S164 | 49.7 | 99.9 | R.ALENGDADEPSFS*DPEDFVDDVS*EEELLGDVLK.D |
| Eukaryotic translation initiation factor 3, subunit 9 eta, 116kDa | EIF3B | S152S154 | 82.9 | 132.5 | R.ALENGDADEPS*FS*DPEDFVDDVSEEELLGDVLK.D |
| Eukaryotic translation initiation factor 3, subunit 9 eta, 116kDa | EIF3B | S152S154S164 | 100.0 | 99.6 | R.ALENGDADEPS*FS*DPEDFVDDVS*EEELLGDVLK.D |
| Eukaryotic translation initiation factor 3, subunit 9 eta, 116kDa | EIF3B | S83S85 | 13.6 | 16.8 | R. [illegible] |
| Eukaryotic translation initiation factor 4B | EIF4B | S445 | 58.9 | 75.3 | K.S*LENETLNK.E |
| Eukaryotic translation initiation factor 4B | EIF4B | S597 | 37.6 | 73.8 | K.YAALS*VDGEDENEGEDYAE.- |
| Eukaryotic translation initiation factor 4B | EIF4B | S409 | 28.8 | 29.8 | R.S*EETQER.E |
| Eukaryotic translation initiation factor 4B | EIF4B | T97 | 5.8 | 75.6 | K.SPPYT*AFLGNLPYDVTEESIK.E |
| Eukaryotic translation initiation factor 4B | EIF4B | S93 | 30.1 | 135.8 | K.S*PPYTAFLGNLPYDVTEESIK.E |
| Eukaryotic translation initiation factor 4B | EIF4B | Y96 | 11.8 | 60.1 | K.SPPY*TAFLGNLPYDVTEESIK.E |
| Eukaryotic translation initiation factor 4B | EIF4B | S422 | 15.0 | 118.0 | R.TGS*ESSQTGTSTTSSR.N |
| Eukaryotic translation initiation factor 4B | EIF4B | S498S504 | 15.9 | 84.1 | R.SQSS*DTEQQS*PTSGGGK.V |
| Eukaryotic translation initiation factor 4B | EIF4B | T420S425 | 20.8 | 70.9 | R.T*GSESS*QTGTSTTSSR.N |
| Eukaryotic translation initiation factor 4B | EIF4B | S406 | 100.0 | 26.8 | R.HPS*WR.S |
| Eukaryotic translation initiation factor 4B | EIF4B | S504 | 38.7 | 100.4 | R.SQSSDTEQQS*PTSGGGK.V |
| Eukaryotic translation initiation factor 4B | EIF4B | T500S504 | 4.6 | 13.6 | R.SQSSDT*EQQS*PTSGGGK.V |
| Eukaryotic translation initiation factor 4B | EIF4B | S497 | 14.0 | 121.1 | R.SQS*SDTEQQSPTSGGGK.V |
| Eukaryotic translation initiation factor 4B | EIF4B | T500 | 9.8 | 59.2 | R.SQSSDT*EQQSPTSGGGK.V |
| Eukaryotic translation initiation factor 4B | EIF4B | S497T500 | 11.8 | 33.2 | R.SQS*SDT*EQQSPTSGGGK.V |
| Eukaryotic translation initiation factor 4B | EIF4B | S424S425 | 6.8 | 29.1 | R.TGSES*S*QTGTSTTSSR.N |
| Eukaryotic translation initiation factor 4B | EIF4B | S489 | 16.9 | 24.3 | K.RSS*NPPAR.S |
| Eukaryotic translation initiation factor 4B | EIF4B | S498 | 11.7 | 72.0 | R.SQSS*DTEQQSPTSGGGK.V |
| Eukaryotic translation initiation factor 4B | EIF4B | S422S425 | 5.9 | 35.7 | R.TGS*ESS*QTGTSTTSSR.N |
| Eukaryotic translation initiation factor 4B | EIF4B | S422S424 | -0.4 | 11.1 | R.TGS*ES*SQTGTSTTSSR.N |
| Eukaryotic translation initiation factor 4B | EIF4B | S424 | -0.4 | 54.4 | R.TGSES*SQTGTSTTSSR.N |
| Eukaryotic translation initiation factor 4B | EIF4B | S497S504 | 13.9 | 29.7 | R.SQS*SDTEQQS*PTSGGGK.V |
| Eukaryotic translation initiation factor 4E binding protein 1 | EIF4EBP1 | T70 | 14.0 | 61.7 | K.T*PPRDLPTIPGVTSPSSDEPPMEASQSHLR.N |
|  | EIF4EBP1 | T36S44 | 19.7 | 25.6 | R.RVVLGDGVQLPPGDYST*TPGGTLFS*TTPGGTR.I |
| Eukaryotic translation initiation factor 4E binding protein 1 | EIF4EBP1 | T36T45 | 14.4 | 22.3 | R.RVVLGDGVQLPPGDYST*TPGGTLFST*TPGGTR.I |
| Eukaryotic translation initiation factor 4E binding protein 1 | EIF4EBP1 | T37S44 | 20.7 | 23.2 | R.VVLGDGVQLPPGDYSTT*PGGTLFS*TTPGGTR.I |
| Eukaryotic translation initiation factor 4E binding protein 1 | EIF4EBP1 | T77 | 24.5 | 72.3 | K.TPPRDLPT*IPGVTSPSSDEPPMEASQSHLR.N |
| Eukaryotic translation initiation factor 4E binding protein 1 | EIF4EBP1 | S35S44 | 10.5 | 32.6 | R.VVLGDGVQLPPGDYS*TTPGGTLFS*TTPGGTR.I |
| Eukaryotic translation initiation factor 4E binding protein 1 | EIF4EBP1 | T36T41 | 16.1 | 13.8 | R.VVLGDGVQLPPGDYST*TPGGT*LFSTTPGGTR.I |
| Eukaryotic translation initiation factor 4E binding protein 1 | EIF4EBP1 | Y34S44 | 4.3 | 16.0 | R.VVLGDGVQLPPGDY*STTPGGTLFS*TTPGGTR.I |
| Eukaryotic translation initiation factor 4E binding protein 1 | EIF4EBP1 | S35T45 | 9.4 | 26.7 | R.RVVLGDGVQLPPGDYS*TTPGGTLFST*TPGGTR.I |
| Eukaryotic translation initiation factor 4E binding protein 2 | EIF4EBP2 | T37S44 | 17.8 | 34.2 | R.TVAISDAAQLPHDYCTT*PGGTLFS*TTPGGTR.I |
| Eukaryotic translation initiation factor 4E binding protein 2 | EIF4EBP2 | T36T45 | 24.7 | 34.0 | R.TVAISDAAQLPHDYCT*TPGGTLFST*TPGGTR.I |
| Eukaryotic translation initiation factor 4E binding protein 2 | EIF4EBP2 | T36T46 | 6.7 | 24.7 | R.TVAISDAAQLPHDYCT*TPGGTLFSTT*PGGTR.I |
| Eukaryotic translation initiation factor 4E binding protein 2 | EIF4EBP2 | S25S44 | 13.0 | 25.5 | R.TVAIS*DAAQLPHDYCTTPGGTLFS*TTPGGTR.I |

Peak Area: <-10, -5, -3, 0, 3, 5, >10

%CV: 0, 29, 42, 71, 86, >100

**White dots-** Significant change in peptide abundance at **5** %FDR compared to the timepoint with the minimum peak area for a given PSM

| Protein Name | Gene | Phosphosites | CarT 0m | CarT 2m | CarT 5m | RajiB 0m | RajiB 2m | RajiB 5m | Ascor | MOWSE | Sequence |
|---|---|---|---|---|---|---|---|---|---|---|---|
| **Eukaryotic translation initiation factor 4E binding protein 2** | EIF4EBP2 | T70 | | | | | | | **26.8** | **25.2** | R.NSPMAQT*PPCHLPNIPGVTSPGTLIEDSK.V |
| | EIF4EBP2 | S65T70 | | | | | | | **43.7** | **31.4** | R.NS*PM#AQT*PPCHLPNIPGVTSPGTLIEDSK.V |
| **Eukaryotic translation initiation factor 4E binding protein 2** | EIF4EBP2 | T37T45 | | | | | | | **21.6** | **35.8** | R.TVAISDAAQLPHDYCTT*PGGTLFST*TPGGTR.I |
| **Eukaryotic translation initiation factor 4E binding protein 2** | EIF4EBP2 | T36S44 | | | | | | | **6.2** | **22.3** | R.TVAISDAAQLPHDYCT*TPGGTLFS*TTPGGTR.I |
| **Eukaryotic translation initiation factor 4E binding protein 2** | EIF4EBP2 | S65 | | | | | | | **29.4** | **51.0** | R.NS*PMAQTPPCHLPNIPGVTSPGTLIEDSK.V |
| **Eukaryotic translation initiation factor 4E binding protein 2** | EIF4EBP2 | S25T45 | | | | | | | **11.4** | **26.0** | R.TVAIS*DAAQLPHDYCTTPGGTLFST*TPGGTR.I |
| **Eukaryotic translation initiation factor 4E binding protein 2** | EIF4EBP2 | T45 | | | | | | | **22.8** | **16.0** | R.TVAISDAAQLPHDYCTTPGGTLFST*TPGGTR.I |
| **Eukaryotic translation initiation factor 4E binding protein 2** | EIF4EBP2 | S25T46 | | | | | | | **2.3** | **22.9** | R.TVAIS*DAAQLPHDYCTTPGGTLFSTT*PGGTR.I |
| **Eukaryotic translation initiation factor 4E binding protein 2** | EIF4EBP2 | T36T50 | | | | | | | **4.9** | **14.2** | R.TVAISDAAQLPHDYCT*TPGGTLFSTTPGGT*R.I |
| **Eukaryotic translation initiation factor 4E binding protein 2** | EIF4EBP2 | S65T82 | | | | | | | **19.4** | **17.7** | R.RNS*PMAQTPPCHLPNIPGVT*SPGTLIEDSK.V |
| **Eukaryotic translation initiation factor 4E binding protein 2** | EIF4EBP2 | Y34T45 | | | | | | | **5.0** | **21.2** | R.TVAISDAAQLPHDY*CTTPGGTLFST*TPGGTR.I |
| **Eukaryotic translation initiation factor 4E binding protein 2** | EIF4EBP2 | Y34S44 | | | | | | | **3.5** | **17.4** | R.TVAISDAAQLPHDY*CTTPGGTLFS*TTPGGTR.I |
| **Eukaryotic translation initiation factor 4E nuclear import factor 1** | EIF4ENIF1 | S74 | | | | | | | **39.9** | **51.1** | K.WHASLYPAS*GR.S |
| **Eukaryotic translation initiation factor 4E nuclear import factor 1** | EIF4ENIF1 | S564 | | | | | | | **82.3** | **68.8** | R.APS*PPLSQVFQTR.A |
| **Eukaryotic translation initiation factor 4E nuclear import factor 1** | EIF4ENIF1 | S78 | | | | | | | **-0.3** | **34.3** | R.SS*PVESLKK.E |
| **Eukaryotic translation initiation factor 4E nuclear import factor 1** | EIF4ENIF1 | S951 | | | | | | | **30.2** | **61.6** | R.SSS*PVGLAK.W |
| **Eukaryotic translation initiation factor 4E nuclear import factor 1** | EIF4ENIF1 | S680S690 | | | | | | | **6.8** | **22.8** | R.GNSS*SPAPAASITS*MLSPSFTPTSVIR.K |
| **Eukaryotic translation initiation factor 4E nuclear import factor 1** | EIF4ENIF1 | S679S690 | | | | | | | **8.4** | **19.4** | R.GNS*SSPAPAASITS*MLSPSFTPTSVIR.K |
| **Eukaryotic translation initiation factor 4E nuclear import factor 1** | EIF4ENIF1 | S301 | | | | | | | **100.0** | **37.6** | R.DAVLPEQS*PGDFDFNEFFNLDK.V |
| **Eukaryotic translation initiation factor 4E nuclear import factor 1** | EIF4ENIF1 | S69 | | | | | | | **8.2** | **42.7** | K.YDSDGVWDPEKWHAS*LYPASGR.S |
| **Eukaryotic translation initiation factor 4E nuclear import factor 1** | EIF4ENIF1 | S680S693 | | | | | | | **9.1** | **28.2** | R.GNSS*SPAPAASITSMLS*PSFTPTSVIR.K |
| **Eukaryotic translation initiation factor 4E nuclear import factor 1** | EIF4ENIF1 | S345 | | | | | | | **9.5** | **15.1** | R.WFSNPS*R.S |
| **Eukaryotic translation initiation factor 4E nuclear import factor 1** | EIF4ENIF1 | Y71 | | | | | | | **-0.7** | **25.9** | K.YDSDGVWDPEKWHASLY*PASGR.S |
| **Eukaryotic translation initiation factor 4E nuclear import factor 1** | EIF4ENIF1 | S681S693 | | | | | | | **6.3** | **16.7** | R.GNSSS*PAPAASITSMLS*PSFTPTSVIR.K |
| **Eukaryotic translation initiation factor 4E nuclear import factor 1** | EIF4ENIF1 | S342 | | | | | | | | **13.1** | R.WFS*NPSR.S |
| **Eukaryotic translation initiation factor 4E nuclear import factor 1** | EIF4ENIF1 | Y382 | | | | | | | **4.1** | **14.0** | R.LAGLEQAILSPGQNSGNY*FAPIPLEDHAENKVDILEMLQK.A |
| **Eukaryotic translation initiation factor 4E nuclear import factor 1** | EIF4ENIF1 | S374 | | | | | | | | **23.5** | R.LAGLEQAILS*PGQNSGNYFAPIPLEDHAENKVDILEMLQK.A |
| **Eukaryotic translation initiation factor 4E nuclear import factor 1** | EIF4ENIF1 | S680S687 | | | | | | | **3.9** | **20.1** | R.GNSS*SPAPAAS*ITSMLSPSFTPTSVIR.K |
| **Eukaryotic translation initiation factor 4E nuclear import factor 1** | EIF4ENIF1 | S679S687 | | | | | | | **6.5** | **24.3** | R.GNS*SSPAPAAS*ITSMLSPSFTPTSVIR.K |
| **Eukaryotic translation initiation factor 4E nuclear import factor 1** | EIF4ENIF1 | T689 | | | | | | | **-10.9** | **22.1** | R.GNSSSPAPAASIT*SMLSPSFTPTSVIR.K |
| **Eukaryotic translation initiation factor 4E nuclear import factor 1** | EIF4ENIF1 | S687S690 | | | | | | | **4.5** | **18.4** | R.GNSSSPAPAAS*ITS*MLSPSFTPTSVIR.K |
| **Eukaryotic translation initiation factor 4E nuclear import factor 1** | EIF4ENIF1 | S950 | | | | | | | **-0.6** | **27.6** | R.SS*SPVGLAK.W |
| **Eukaryotic translation initiation factor 5** | EIF5 | S389S390 | | | | | | | **107.4** | **76.1** | K.EAEEES*S*GGEEEDEDENIEVVYSK.A |
| **Eukaryotic translation initiation factor 5B** | EIF5B | S113 | | | | | | | **43.8** | **110.0** | K.KQSFDDNDS*EELEDKDSK.S |
| **Eukaryotic translation initiation factor 5B** | EIF5B | S107S113 | | | | | | | **94.5** | **70.2** | K.KQS*FDDNDS*EELEDKDSK.S |
| **Eukaryotic translation initiation factor 5B** | EIF5B | S588S589S591 | | | | | | | **20.2** | **60.8** | K.EMS*S*DS*EYDS*DDDRTK.E |
| **Eukaryotic translation initiation factor 5B** | EIF5B | S214 | | | | | | | **147.1** | **111.7** | K.NKPGPNIES*GNEDDDASFK.I |
| **Eukaryotic translation initiation factor 5B** | EIF5B | S135S137 | | | | | | | **11.2** | **69.3** | K.VEMYS*GS*DDDDDFNKLPK.K |
| **Eukaryotic translation initiation factor 5B** | EIF5B | S182S183S186 | | | | | | | **112.4** | **114.3** | R.INS*S*GES*GDES*DEFLQSR.K |
| **Eukaryotic translation initiation factor 5B** | EIF5B | S511 | | | | | | | **34.6** | **63.3** | K.EETPPPVEPEEEEDTEDAGLDDWEAMAS*DEETEK.V |
| **Eukaryotic translation initiation factor 5B** | EIF5B | S164 | | | | | | | **115.1** | **74.4** | K.WDGS*EEDEDNSK.K |
| **Eukaryotic translation initiation factor 5B** | EIF5B | S588S589S591 | | | | | | | **8.6** | **36.5** | K.EMS*S*DS*EY*DSDDDRTK.E |
| **Eukaryotic translation initiation factor 5B** | EIF5B | S182S183S186 | | | | | | | **60.1** | **116.0** | R.INS*S*GES*GDESDEFLQSR.K |
| **Eukaryotic translation initiation factor 5B** | EIF5B | T498S511 | | | | | | | **43.0** | **43.4** | K.EETPPPVEPEEEEDT*EDAGLDDWEAMAS*DEETEK.V |
| **Eukaryotic translation initiation factor 5B** | EIF5B | S107 | | | | | | | **4.8** | **58.6** | K.QS*FDDNDSEELEDKDSK.S |
| **Eukaryotic translation initiation factor 5B** | EIF5B | S589S591Y593 | | | | | | | **14.9** | **13.1** | K.EMSS*DS*EY*DS*DDDRTK.E |
| **Eukaryotic translation initiation factor 5B** | EIF5B | Y134S135 | | | | | | | **5.7** | **37.4** | K.VEMY*S*GSDDDDDFNKLPK.K |
| **Eukaryotic translation initiation factor 6** | EIF6 | T245 | | | | | | | **16.9** | **30.2** | K.LNEAQPSTIATSMRDSLIDSLT*.- |
| **Eukaryotic translation initiation factor 6** | EIF6 | S239 | | | | | | | **6.6** | **29.5** | K.LNEAQPSTIATSMRDS*LIDSLT.- |
| **Eukaryotic translation initiation factor 6** | EIF6 | S243 | | | | | | | **13.0** | **13.3** | K.LNEAQPSTIATSMRDSLIDS*LT.- |
| **EVER1** | TMC6 | S62 | | | | | | | **9.5** | **29.5** | R.EVTGS*SQQTLWRPEGTQSTATLR.I |
| **EVER1** | TMC6 | T60 | | | | | | | **21.6** | **21.0** | R.EVT*GSSQQTLWRPEGTQSTATLR.I |
| **EVER1** | TMC6 | S63 | | | | | | | **6.5** | **43.0** | R.EVTGSS*QQTLWRPEGTQSTATLR.I |
| **EVER2 protein** | TMC8 | S521 | | | | | | | **12.1** | **28.1** | R.AS*SRPFR.A |
| **EVER2 protein** | TMC8 | S722 | | | | | | | **100.0** | **29.4** | R.FRFPS*GAEL.- |
| **EVER2 protein** | TMC8 | S522 | | | | | | | **8.2** | **31.0** | R.ASS*RPFR.A |
| **EVI2B** | EVI2B | S271 | | | | | | | **-1.0** | **21.5** | K.RTSIIS*LTPWKPSK.S |
| **EVI2B** | EVI2B | T267S268 | | | | | | | **6.8** | **43.6** | K.RT*S*IISLTPWKPSK.S |
| **EVI2B** | EVI2B | S268S271 | | | | | | | **16.5** | **26.9** | R.TS*IIS*LTPWKPSK.S |
| **EVI2B** | EVI2B | T267 | | | | | | | **9.1** | **23.6** | K.RT*SIISLTPWKPSK.S |
| **EVI2B** | EVI2B | S294 | | | | | | | **16.9** | **119.8** | K.LFES*SENIEDSNNPK.T |
| **EVI2B** | EVI2B | S268 | | | | | | | **-0.4** | **31.3** | R.TS*IISLTPWKPSK.S |

Peak Area: <-10, -5, -3, 0, 3, 5, >10

%CV: 0, 29, 42, 71, 86, >100

**White dots-** Significant change in peptide abundance at **5** %FDR compared to the timepoint with the minimum peak area for a given PSM

CarT (0m, 2m, 5m) — RajiB (0m, 2m, 5m)

| Protein Name | Gene | Phosphosites | Ascor | MOWSE | Sequence |
|---|---|---|---|---|---|
| Excision repair cross-complementing rodent repair deficiency, complementation | ERCC6 | S486S489 | 100.0 | 19.8 | K.LEDDS*EES*DAEFDEGFKVPGFLFK.K |
| Exocyst complex component 1 | EXOC1 | S470S473 | 15.3 | 41.1 | K.LTGS*TSS*LNK.L |
| Exocyst complex component 1 | EXOC1 | S448 | 19.1 | 65.0 | K.KFGLHGS*SGK.L |
| Exocyst complex component 1 | EXOC1 | S470T471 | 9.0 | 31.6 | K.LTGS*T*SSLNK.L |
| Exocyst complex component 1 | EXOC1 | S499 | 1.7 | 44.1 | R.SQSSSLLDMGNMS*ASDLDVADR.T |
| Exocyst complex component 1 | EXOC1 | T471S473 | 8.9 | 42.1 | K.LTGST*SS*LNK.L |
| Exocyst complex component 1 | EXOC1 | S472S473 | 10.0 | 26.0 | K.LTGSTS*S*LNK.L |
| Exocyst complex component 1 | EXOC1 | S490 | 17.7 | 35.8 | R.SQSS*SLLDMGNMSASDLDVADR.T |
| Exocyst complex component 1 | EXOC1 | S501 | 10.7 | 12.1 | R.SQSSSLLDMGNMSAS*DLDVADR.T |
| Exocyst complex component 1 | EXOC1 | T471S472 | 4.4 | 37.6 | K.LTGST*S*SLNK.L |
| Exocyst complex component 1 | EXOC1 | S470S472 | 7.6 | 32.4 | K.LTGS*TS*SLNK.L |
| Exocyst complex component 4 | EXOC4 | S226 | 67.3 | 61.3 | K.DAS*VPLIDVTNLPTPR.K |
| Exportin 6 | XPO6 | T210S211 | 24.2 | 99.6 | K.HSVTAATPPPSPT*S*GESGDLLSNLLQSPSSAK.L |
| Exportin 6 | XPO6 | T204S208 | 37.3 | 116.0 | K.HSVTAAT*PPPS*PTSGESGDLLSNLLQSPSSAK.L |
| Exportin 6 | XPO6 | S208S214 | 7.6 | 49.8 | K.HSVTAATPPPS*PTSGES*GDLLSNLLQSPSSAK.L |
| Exportin 6 | XPO6 | T204S211 | 13.5 | 101.9 | K.HSVTAAT*PPPSPTS*GESGDLLSNLLQSPSSAK.L |
| Exportin 6 | XPO6 | T210S214 | 1.2 | 40.9 | K.HSVTAATPPPSPT*SGES*GDLLSNLLQSPSSAK.L |
| Exportin 6 | XPO6 | T201T204 | 5.2 | 20.3 | K.HSVT*AAT*PPPSPTSGESGDLLSNLLQSPSSAK.L |
| Exportin 6 | XPO6 | S208S211 | 13.7 | 54.9 | K.HSVTAATPPPS*PTS*GESGDLLSNLLQSPSSAK.L |
| Exportin 6 | XPO6 | T204S219 | 10.2 | 46.0 | K.HSVTAAT*PPPSPTSGESGDLLS*NLLQSPSSAK.L |
| Exportin 6 | XPO6 | T204T210 | 6.7 | 26.3 | K.HSVTAAT*PPPSPT*SGESGDLLSNLLQSPSSAK.L |
| Exportin 6 | XPO6 | S199T201 |  | 31.9 | K.HS*VT*AATPPPSPTSGESGDLLSNLLQSPSSAK.L |
| Exportin 6 | XPO6 | S208S219 | 7.5 | 39.0 | K.HSVTAATPPPS*PTSGESGDLLS*NLLQSPSSAK.L |
| Exportin 6 | XPO6 | S208 | 4.2 | 25.3 | K.HSVTAATPPPS*PTSGESGDLLSNLLQSPSSAK.L |
| Exportin 6 | XPO6 | S214S219 | 2.3 | 31.2 | K.HSVTAATPPPSPTSGES*GDLLS*NLLQSPSSAK.L |
| Exportin 6 | XPO6 | T204S214 | 23.0 | 14.0 | K.HSVTAAT*PPPSPTSGES*GDLLSNLLQSPSSAK.L |
| EZH2 protein | EZH2 | S367S371 | 8.6 | 21.6 | R.GRLPNNS*SRPS*TPTINVLESK.D |
| EZH2 protein | EZH2 | S371T372 | 5.6 | 17.2 | R.GRLPNNSSRPS*T*PTINVLESK.D |
| EZH2 protein | EZH2 | T372T374 | 3.2 | 14.8 | R.GRLPNNSSRPST*PT*INVLESK.D |
| EZH2 protein | EZH2 | S367S368 | 0.8 | 16.3 | R.GRLPNNS*S*RPSTPTINVLESK.D |
| EZH2 protein | EZH2 | S368S371 | 6.1 | 13.0 | R.GRLPNNSS*RPS*TPTINVLESK.D |
| EZH2 protein | EZH2 | S367T372 | 4.0 | 24.0 | R.LPNNS*SRPST*PTINVLESK.D |
| F box and leucine rich repeat protein 11 | KDM2A | T713S718S731 | 22.0 | 11.7 | R.SCDEPLT*PPPHS*PTSMLQLIHDPVS*PR.G |
| F box and leucine rich repeat protein 11 | KDM2A | S869 | 100.0 | 98.5 | R.GDEEGLGGEEEEEEEEEEEDDS*AEEGGAAR.L |
| F box and leucine rich repeat protein 11 | KDM2A | T713S718 | 21.8 | 23.6 | R.SCDEPLT*PPPHS*PTSMLQLIHDPVSPR.G |
| F box and leucine rich repeat protein 11 | KDM2A | S718S731 | 12.8 | 13.1 | R.SCDEPLTPPPHS*PTSMLQLIHDPVS*PR.G |
| F box and leucine rich repeat protein 11 | KDM2A | S721S731 | 13.4 | 11.8 | R.SCDEPLTPPPHSPTS*MLQLIHDPVS*PR.G |
| F box and leucine rich repeat protein 11 | KDM2A | T713S721S731 | 18.5 | 12.3 | R.SCDEPLT*PPPHSPTS*MLQLIHDPVS*PR.G |
| F-box and leucine rich repeat protein 10 | KDM2B | Y490T493 | 10.9 | 24.4 | K.STTLAVDY*PKT*PTGSPATEVSAK.W |
| F-box and leucine rich repeat protein 10 | KDM2B | T493T495 | 20.3 | 17.8 | K.STTLAVDYPKT*PT*GSPATEVSAK.W |
| F-box and leucine rich repeat protein 10 | KDM2B | T493S497 | 14.6 | 24.0 | K.STTLAVDYPKT*PTGS*PATEVSAK.W |
| F-box and leucine-rich repeat protein 20, isoform CRA a | FBXL20 | S4 | 100.0 | 14.6 | M.APS*RDR.L |
| F-box protein 42 | FBXO42 | T539S545 | 17.0 | 47.7 | R.HPPEQTNGVHT*PPHVAS*ALAGAVSPGALR.R |
| F-box protein 42 | FBXO42 | T539S552 | 53.7 | 54.2 | R.HPPEQTNGVHT*PPHVASALAGAVS*PGALR.R |
| F-box protein 42 | FBXO42 | S365S373 | 24.4 | 43.5 | R.APLS*PSLNSRPS*PISATPPALVPETR.E |
| F-box protein 42 | FBXO42 | S365S376 | 13.5 | 12.8 | R.APLS*PSLNSRPSPIS*ATPPALVPETR.E |
| F-box protein 42 | FBXO42 | S365S367 | 13.3 | 36.1 | R.APLS*PS*LNSRPSPISATPPALVPETR.E |
| FACTp140 | SUPT16H | S979S982S986 | 100.0 | 25.1 | K.ES*LGS*EEES*GKDWDELEEEAR.K |
| FADD | FADD | S197 | -0.3 | 16.9 | R.SGAMSPMS*WNSDASTSEAS.- |
| FAM40A | STRIP1 | Y62 | 5.6 | 52.7 | R.KDSEGY*SESPDLEFEYADTDK.W |
| Family with sequence similarity 123C | AMER3 | S97 | 7.3 | 15.7 | R.KCKTHDS*M#SGAGR.A |
| Family with sequence similarity 126, member B | FAM126B | S430 | 100.0 | 17.1 | K.HLS*LPAGQVVPK.I |
| Family with sequence similarity 128, member B | MZT2B | S152 | 13.9 | 34.2 | K.GGGPGKS*PTR.G |
| Family with sequence similarity 133, member B | FAM133B | S191S192S194 | 28.8 | 75.0 | K.MYSEDKPLS*S*ES*LS*ESEYIEEVR.A |
| Family with sequence similarity 29, member A | HAUS6 | S406 | 15.6 | 36.0 | K.GWTPSVDLLPPMS*PLSFDPASEEVYAK.S |
| Family with sequence similarity 44, member A | BOD1L1 | S482S484 | 31.3 | 100.9 | K.YYS*DS*DDELTVEQR.R |
| Family with sequence similarity 44, member A | BOD1L1 | S1077 |  | 14.4 | R.RGS*LSQEMAK.G |
| Family with sequence similarity 44, member A | BOD1L1 | S2779S2780 | 10.8 | 45.7 | R.KQHYLS*S*EDEPDDNPDVLDSR.I |
| Family with sequence similarity 53, member B | FAM53B | S167 | 12.0 | 48.0 | R.SSS*FSLPSR.A |
| Family with sequence similarity 53, member B | FAM53B | S166 | -0.3 | 39.1 | R.SS*SFSLPSR.A |
| Family with sequence similarity 53, member B | FAM53B | S118 | 33.6 | 58.7 | R.SLS*FSDEMSSCR.T |
| Family with sequence similarity 53, member B | FAM53B | S165 | 17.0 | 35.5 | R.S*SSFSLPSR.A |

**Peak Area**: <-10, -5, -3, 0, 3, 5, >10

**%CV**: 0, 29, 42, 71, 86, >100

**White dots-** Significant change in peptide abundance at **5** %FDR compared to the timepoint with the minimum peak area for a given PSM

| Protein Name | Gene | Phosphosites | CarT 0m | CarT 2m | CarT 5m | RajiB 0m | RajiB 2m | RajiB 5m | Ascor | MOWSE | Sequence |
|---|---|---|---|---|---|---|---|---|---|---|---|
| **Family with sequence similarity 54, member B** | MTFR1L | S103 | | | | | | | **100.0** | **36.1** | R.NAS*VPNLR.G |
| **Family with sequence similarity 54, member B** | MTFR1L | S238 | | | | | | | **30.2** | **67.2** | K.ASS*FADMMGILK.D |
| **Family with sequence similarity 54, member B** | MTFR1L | S41 | | | | | | | **22.1** | **55.4** | R.AS*FETLPNISDLCLR.D |
| **Family with sequence similarity 54, member B** | MTFR1L | S237 | | | | | | | | **60.6** | K.AS*SFADMM#GILK.D |
| **Family with sequence similarity 65, member A** | RIPOR1 | S22 | | | | | | | **6.4** | **42.5** | R.SQS*FAGVLGSHER.G |
| **Family with sequence similarity 65, member A** | RIPOR1 | S20 | | | | | | | **12.2** | **87.8** | R.S*QSFAGVLGSHER.G |
| **Family with sequence similarity 76, member B** | FAM76B | S193 | | | | | | | **52.0** | **61.1** | K.ISNLS*PEEEQGLWK.Q |
| **Family with sequence similarity 82, member A** | RMDN2 | S128S129 | | | | | | | **100.0** | **13.3** | R.RRFS*S*R.K |
| **Family with sequence similarity 82, member C** | RMDN3 | S46 | | | | | | | **8.3** | **39.8** | R.SQS*LPNSLDYTQTSDPGR.H |
| **Family with sequence similarity 83, member G** | FAM83G | S127 | | | | | | | **51.9** | **57.8** | R.S*IPQLDLGWPDTIAYR.G |
| **Family with sequence similarity 91, member A1** | FAM91A1 | S829 | | | | | | | **25.6** | **75.4** | R.SPSS*LLIANLHLQ.- |
| **Family with sequence similarity 91, member A1** | FAM91A1 | S828 | | | | | | | **13.8** | **53.5** | R.SPS*SLLIANLHLQ.- |
| **Family with sequence similarity 91, member A1** | FAM91A1 | S671S674 | | | | | | | **100.0** | **21.2** | R.KLS*DAS*DER.G |
| **Far upstream element binding protein** | FUBP1 | S630 | | | | | | | **-0.4** | **21.7** | R.QQAAYYAQTS*PQGM#PQHPPAPQGQ.- |
| **Far upstream element binding protein** | FUBP1 | T629 | | | | | | | **30.1** | **72.4** | R.QQAAYYAQT*SPQGMPQHPPAPQGQ.- |
| **Far upstream element binding protein** | FUBP1 | Y626 | | | | | | | **6.5** | **11.2** | R.QQAAYY*AQTSPQGMPQHPPAPQGQ.- |
| **FAS associated factor 1** | FAF1 | S320 | | | | | | | **84.4** | **64.7** | K.S*PMMPENAENEGDALLQFTAEFSSR.Y |
| **FAS associated factor 1** | FAF1 | S278T283 | | | | | | | | **66.4** | R. EQS*EEQIT*DVHMVSDSDGDDFEDATEFGVDDGEVFGMASSALR.K |
| **Fas receptor** | FAS | S212 | | | | | | | **21.4** | **60.5** | K.ENQGSHES*PTLNPETVAINLSDVDLSK.Y |
| **Fas receptor** | FAS | S209 | | | | | | | **5.6** | **44.9** | K.ENQGS*HESPTLNPETVAINLSDVDLSK.Y |
| **Fas receptor** | FAS | T214 | | | | | | | **-2.2** | **62.9** | K.ENQGSHESPT*LNPETVAINLSDVDLSK.Y |
| **Fatty acid synthase** | FASN | S974 | | | | | | | **14.9** | **40.2** | R.LFDHPES*PTPNPTEPLFLAQAEVYK.E |
| **Fatty acid synthase** | FASN | T976 | | | | | | | **2.3** | **46.5** | R.LFDHPESPT*PNPTEPLFLAQAEVYK.E |
| **FBP3** | FUBP3 | S258 | | | | | | | **24.3** | **36.7** | R.QQVAFYGQTLGQAQAHS*QEQ.- |
| **FBP3** | FUBP3 | S538 | | | | | | | **24.4** | **65.5** | K.QSHAASAAPQAS*SPPDYTMAWAEYYR.Q |
| **FBP3** | FUBP3 | S539 | | | | | | | **13.8** | **60.7** | K.QSHAASAAPQASS*PPDYTMAWAEYYR.Q |
| **FCH and double SH3 domains 2** | FCHSD2 | S596S609S613 | | | | | | | **9.2** | **12.9** | R.EIQISPSPKPHAS*LPPLPLYDQPPSS*PYPS*PDKR.S |
| **FCH domain only 1** | FCHO1 | S523S529 | | | | | | | **59.1** | **66.9** | R.APPLPDS*PQPLAS*SPGPWGLEALAGGDLMPAPADPTAR.E |
| **FCH domain only 1** | FCHO1 | S523S530 | | | | | | | **43.1** | **79.8** | R.APPLPDS*PQPLASS*PGPWGLEALAGGDLMPAPADPTAR.E |
| **FCH domain only 1** | FCHO1 | S583S585S587 | | | | | | | **34.5** | **19.6** | R.S*LS*PS*PLGSSAASTALERPS*FLSQTGHGVSR.G |
| **Ferm, RhoGEF and pleckstrin domain protein 2** | FARP2 | S399Y407 | | | | | | | **19.8** | **55.8** | R.TPAS*PSSANAFY*SLSPSTLVPSGLPEFK.D |
| **Ferm, RhoGEF and pleckstrin domain protein 2** | FARP2 | S399S408 | | | | | | | **1.5** | **33.8** | R.TPAS*PSSANAFYS*LSPSTLVPSGLPEFK.D |
| **Ferm, RhoGEF and pleckstrin domain protein 2** | FARP2 | S399S410 | | | | | | | **1.7** | **42.1** | R.TPAS*PSSANAFYSLS*PSTLVPSGLPEFK.D |
| **Ferritin heavy chain 1** | FTH1 | S179 | | | | | | | **17.9** | **44.7** | K.HTLGDS*DNES.- |
| **Fetal Alzheimer antigen** | BPTF | S216 | | | | | | | **100.0** | **36.2** | R.S*PILEEK.D |
| **Fetal Alzheimer antigen** | BPTF | T938 | | | | | | | **38.7** | **22.6** | K.KSLEGT*K.N |
| **FGD1 family member 3** | FGD3 | S128 | | | | | | | **34.5** | **41.2** | K.VTPQEEADS*DVGEEPDSENTPQK.A |
| **FGD1 family member 3** | FGD3 | T121 | | | | | | | **5.9** | **12.5** | K.VT*PQEEADSDVGEEPDSENTPQK.A |
| **FGD1 family, member 2** | FGD2 | S654 | | | | | | | **30.9** | **34.5** | R.AASGWSPSWPNDGDLS*D.- |
| **FGFR1 oncogene partner** | FGFR1OP | S156S160 | | | | | | | **35.9** | **30.6** | K.EKGPTTGEGALDLSDVHS*PPKS*PEGK.T |
| **FGFR1 oncogene partner** | FGFR1OP | S152S160 | | | | | | | **18.8** | **14.7** | K.GPTTGEGALDLS*DVHSPPKS*PEGK.T |
| **Fibrillarin** | FBL | S124 | | | | | | | **29.7** | **67.2** | K.RVS*ISEGDDKIEYR.A |
| **Filamin A, alpha** | FLNA | S2144 | | | | | | | **51.1** | **41.8** | R.RAPS*VANVGSHCDLSLK.I |
| **FIP1 like 1** | FIP1L1 | T494S500 | | | | | | | **44.8** | **42.8** | R.ERDHSPT*PSVFNS*DEER.Y |
| **FIP1 like 1** | FIP1L1 | S492T494S500 | | | | | | | **23.9** | **37.2** | R.ERDHS*PT*PSVFNS*DEER.Y |
| **FIP1 like 1** | FIP1L1 | S492 | | | | | | | **53.8** | **87.0** | R.DHS*PTPSVFNSDEER.Y |
| **FIP1 like 1** | FIP1L1 | S492S496 | | | | | | | **42.0** | **78.2** | R.DHS*PTPS*VFNSDEER.Y |
| **FIP1 like 1** | FIP1L1 | T81S87S89 | | | | | | | **16.9** | **69.6** | K.VTET*EDDSDS*DS*DDDEDDVHVTIGDIK.T |
| **FIP1 like 1** | FIP1L1 | S85S87S89 | | | | | | | **39.3** | **59.8** | K.VTETEDDS*DS*DS*DDDEDDVHVTIGDIK.T |
| **FIP1 like 1** | FIP1L1 | S304 | | | | | | | **100.0** | **44.2** | R.AES*PDLR.R |
| **FIP1 like 1** | FIP1L1 | T494 | | | | | | | **5.0** | **95.7** | R.DHSPT*PSVFNSDEER.Y |
| **FIP1 like 1** | FIP1L1 | S492T494 | | | | | | | **34.3** | **84.9** | R.DHS*PT*PSVFNSDEER.Y |
| **FIP1 like 1** | FIP1L1 | T494S496S500 | | | | | | | **10.3** | **24.6** | R.DHSPT*PS*VFNS*DEER.Y |
| **FIP1 like 1** | FIP1L1 | T81S85S87 | | | | | | | **18.9** | **73.9** | K.VTET*EDDS*DS*DSDDDEDDVHVTIGDIK.T |
| **FIP1 like 1** | FIP1L1 | S496S500 | | | | | | | **25.1** | **54.0** | R.DHSPTPS*VFNS*DEER.Y |
| **FIP1 like 1** | FIP1L1 | T79T81S85S89 | | | | | | | **7.8** | **54.0** | K.VT*ET*EDDS*DSDS*DDDEDDVHVTIGDIK.T |
| **FIP1 like 1** | FIP1L1 | T494S496 | | | | | | | **13.3** | **48.1** | R.DHSPT*PS*VFNSDEER.Y |
| **FIP1 like 1** | FIP1L1 | T79T81S87S89 | | | | | | | **16.4** | **23.1** | K.VT*ET*EDDSDS*DS*DDDEDDVHVTIGDIK.T |
| **FIP1 like 1** | FIP1L1 | T81S85S89 | | | | | | | **8.8** | **51.5** | K.VTET*EDDS*DSDS*DDDEDDVHVTIGDIK.T |
| **FIP1 like 1** | FIP1L1 | S492S500 | | | | | | | **31.4** | **67.1** | R.DHS*PTPSVFNS*DEER.Y |
| **FIP1 like 1** | FIP1L1 | S492T494S496 | | | | | | | **15.9** | **47.5** | R.DHS*PT*PS*VFNSDEER.Y |
| **FIP1 like 1** | FIP1L1 | T79T81S85S87 | | | | | | | **12.7** | **39.6** | K.VT*ET*EDDS*DS*DSDDDEDDVHVTIGDIK.T |

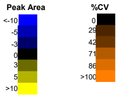


**White dots-** Significant change in peptide abundance at **5** %FDR compared to the timepoint with the minimum peak area for a given PSM

| Protein Name | Gene | Phosphosites | CarT 0m | CarT 2m | CarT 5m | RajiB 0m | RajiB 2m | RajiB 5m | Ascor | MOWSE | Sequence |
|---|---|---|---|---|---|---|---|---|---|---|---|
| FK506 binding protein 15, 133kDa | FKBP15 | S1164 | | | | | | | 19.8 | 87.9 | R.SSLS*GDEEDELFK.G |
| FK506 binding protein 15, 133kDa | FKBP15 | S1162S1164 | | | | | | | 9.6 | 92.1 | R.SS*LS*GDEEDELFK.G |
| FK506 binding protein 15, 133kDa | FKBP15 | S1114 | | | | | | | 112.5 | 64.0 | R.LSLTSDPEEGDPLALGPES*PGEPQPPQLK.K |
| FK506 binding protein 15, 133kDa | FKBP15 | S311 | | | | | | | 27.7 | 33.9 | R.DSAAPS*PIPGADNLSADPVVSPPTSIPFK.S |
| FK506 binding protein 15, 133kDa | FKBP15 | S307S311 | | | | | | | 5.0 | 25.2 | R.DS*AAPS*PIPGADNLSADPVVSPPTSIPFK.S |
| FK506 binding protein 15, 133kDa | FKBP15 | S295S297 | | | | | | | | 12.9 | R.DS*GS*DGHSVSSRDSAAPSPIPGADNLSADPVVSPPTSIPFK.S |
| FK506 binding protein 15, 133kDa | FKBP15 | S320 | | | | | | | 3.3 | 15.9 | R.DSAAPSPIPGADNLS*ADPVVSPPTSIPFK.S |
| FK506 binding protein 15, 133kDa | FKBP15 | S320S326 | | | | | | | 17.5 | 18.3 | R.DSAAPSPIPGADNLS*ADPVVS*PPTSIPFK.S |
| FK506 binding protein 15, 133kDa | FKBP15 | S939S940S941 | | | | | | | 10.4 | 22.1 | K.MVTLQLLNQQEQEKEES*S*S*EEEEEK.A |
| FK506 binding protein 15, 133kDa | FKBP15 | S956 | | | | | | | 22.8 | 64.5 | R.RPS*QEQSASASSGQPQAPLNR.E |
| FK506 binding protein 15, 133kDa | FKBP15 | S1162 | | | | | | | -0.4 | 60.7 | R.SS*LSGDEEDELFK.G |
| FK506 binding protein 15, 133kDa | FKBP15 | S1161S1162 | | | | | | | 11.6 | 51.0 | R.S*S*LSGDEEDELFK.G |
| FK506 binding protein 15, 133kDa | FKBP15 | S311S326 | | | | | | | 13.6 | 13.1 | R.DSAAPS*PIPGADNLSADPVVS*PPTSIPFK.S |
| FK506 binding protein 4 | FKBP4 | S453 | | | | | | | 29.8 | 75.3 | K.SNTAGSQS*QVETEA.- |
| FKHR | FOXO1 | S298S301 | | | | | | | 24.5 | 12.5 | K.WPAS*PGS*HSNDDFDNWSTFRPR.T |
| FKHR | FOXO1 | S298S303 | | | | | | | 6.4 | 16.1 | K.WPAS*PGSHS*NDDFDNWSTFRPR.T |
| FLI1 | FLI1 | S241 | | | | | | | 37.6 | 55.3 | R.GAWGNNMNSGLNKS*PPLGGAQTISK.N |
| FLI1 | FLI1 | S236 | | | | | | | 18.6 | 42.8 | R.GAWGNNMNS*GLNKSPPLGGAQTISK.N |
| Flightless 1 | FLII | S436 | | | | | | | 100.0 | 45.4 | R.RKDS*AQDDQAK.Q |
| Flightless 1 | FLII | S856 | | | | | | | 32.7 | 44.3 | R.NAEAVLQS*PGLSGK.V |
| FLJ10378 | LARP1B | T362 | | | | | | | 11.2 | 75.5 | R.GLST*SLPDLDSEPWIEVK.K |
| FLJ10378 | LARP1B | S363 | | | | | | | -0.2 | 82.5 | R.GLSTS*LPDLDSEPWIEVK.K |
| FLJ10378 | LARP1B | S340S343 | | | | | | | 100.0 | 27.4 | R.IGS*PLS*PK.K |
| FLJ12387 | KLC2 | S609 | | | | | | | 14.2 | 89.4 | R.TLSS*SSMDLSR.R |
| FLJ12387 | KLC2 | S582 | | | | | | | 30.2 | 70.1 | R.ASS*LNFLNK.S |
| FLJ12387 | KLC2 | S581 | | | | | | | 8.5 | 49.2 | R.AS*SLNFLNK.S |
| FLJ12387 | KLC2 | S610 | | | | | | | 12.1 | 84.7 | R.TLSSS*SMDLSR.R |
| FLJ12387 | KLC2 | S609S611 | | | | | | | 10.5 | 35.6 | R.TLSS*SS*MDLSR.R |
| FLJ12387 | KLC2 | S609S610 | | | | | | | 2.4 | 38.4 | R.TLSS*S*SMDLSR.R |
| | PRPF38A | S193S194 | | | | | | | 68.7 | 95.0 | R.VSALEEDMDDVES*S*EEEEEEDEK.L |
| FLJ14936 | PRPF38A | S92S94 | | | | | | | 100.0 | 22.1 | R.S*KS*PGHHR.S |
| FLJ14936 | PRPF38A | S22 | | | | | | | 100.0 | 36.2 | R.VPS*PDHR.R |
| FLJ14936 | PRPF38A | S182S194 | | | | | | | 13.7 | 83.2 | R.VS*ALEEDMDDVESS*EEEEEEDEKLER.V |
| FLJ14936 | PRPF38A | S182S193 | | | | | | | 3.6 | 11.4 | R.VS*ALEEDMDDVES*SEEEEEEDEKLER.V |
| FLJ14936 | PRPF38A | S73S75 | | | | | | | 100.0 | 10.8 | R.HRS*KS*PR.R |
| | PRPF38A | S105S107S109 | | | | | | | 100.0 | 11.2 | R.HRS*HS*KS*PER.S |
| FLJ14936 | PRPF38A | S39 | | | | | | | 26.0 | 20.5 | R.RS*PTLR.Y |
| FLJ20105 protein | ERCC6L | S820 | | | | | | | 45.4 | 15.2 | K.GFGS*VEELCTNSSLGMEK.S |
| FLJ20105 protein | ERCC6L | S946 | | | | | | | 7.4 | 97.9 | K.LEEEPSASS*PQYACDFNLFLEDSADNR.Q |
| FLJ20105 protein | ERCC6L | Y949 | | | | | | | 5.6 | 79.5 | K.LEEEPSASSPQY*ACDFNLFLEDSADNR.Q |
| FLJ20105 protein | ERCC6L | S1028 | | | | | | | 24.1 | 62.0 | R.IVS*DGEDEDDSFKDTSSINPFNTSLFQFSSVK.Q |
| FLJ20105 protein | ERCC6L | S1098 | | | | | | | 100.0 | 44.5 | R.S*LINMVLDHVEDMEER.L |
| FLJ20514 | GEMIN8 | T124S126 | | | | | | | 34.9 | 70.5 | K.EEEMET*ES*DAEVECDLSNMEITEELR.Q |
| FLJ20514 | GEMIN8 | T124S135 | | | | | | | 11.2 | 62.3 | K.EEEMET*ESDAEVECDLS*NMEITEELR.Q |
| FLJ20514 | GEMIN8 | S126S135 | | | | | | | 8.9 | 46.7 | K.EEEMETES*DAEVECDLS*NMEITEELR.Q |
| FLJ21924 | QSER1 | S991S992 | | | | | | | 0.9 | 31.3 | K.NLEHLSSFS*S*DEDDPGYSQDAYK.S |
| FLJ36874 protein | PATL1 | S36 | | | | | | | 13.8 | 47.0 | R.STS*PIIGSPPVR.A |
| FLJ36874 protein | PATL1 | T35S36 | | | | | | | 2.7 | 25.1 | R.ST*S*PIIGSPPVR.A |
| FLJ46354 protein | MROH7- | Y1166 | | | | | | | | 12.0 | K.RAY*SR.K |
| FLN29 protein | TRAFD1 | S415 | | | | | | | 19.1 | 38.2 | R.LDSQPQETS*PELPR.R |
| FLN29 protein | TRAFD1 | S327 | | | | | | | | 66.4 | R.ALPSLNTGSSS*PR.G |
| FLN29 protein | TRAFD1 | S325 | | | | | | | 13.9 | 44.6 | R.ALPSLNTGS*SSPR.G |
| FLN29 protein | TRAFD1 | T323 | | | | | | | 6.1 | 24.0 | R.ALPSLNT*GSSSPR.G |
| FLN29 protein | TRAFD1 | T414 | | | | | | | 35.7 | 12.3 | R.LDSQPQET*SPELPR.R |
| FMRP interacting protein, 82-kD | NUFIP2 | S652 | | | | | | | 55.8 | 57.7 | R.NDS*WGSFDLR.A |
| FMRP interacting protein, 82-kD | NUFIP2 | S629 | | | | | | | 26.0 | 99.2 | K.DYEIESQNPLAS*PTNTLLGSAK.E |
| FMRP interacting protein, 82-kD | NUFIP2 | S572 | | | | | | | 23.9 | 47.5 | K.RTS*PQVLGSILK.S |
| FMRP interacting protein, 82-kD | NUFIP2 | T571 | | | | | | | 37.7 | 38.6 | K.RT*SPQVLGSILK.S |
| | FNBP4 | S497S506 | | | | | | | 100.0 | 32.8 | K.IDENS*DKEMEVEES*PEKIK.V |
| FNBP4 | FNBP4 | Y111 | | | | | | | 36.4 | 113.5 | K.ATGGLCLLGAY*ADSDDDDNDVSEK.L |
| FNBP4 | FNBP4 | S114 | | | | | | | 33.4 | 102.8 | K.ATGGLCLLGAYADS*DDDDNDVSEK.L |
| FNBP4 | FNBP4 | S462 | | | | | | | 19.9 | 35.4 | R.ATS*PESTSR.S |

Peak Area: <-10, -5, -3, 0, 3, 5, >10

%CV: 0, 29, 42, 71, 86, >100

**White dots-** Significant change in peptide abundance at **5** %FDR compared to the timepoint with the minimum peak area for a given PSM

| Protein Name | Gene | Phosphosites | CarT 0m | CarT 2m | CarT 5m | RajiB 0m | RajiB 2m | RajiB 5m | Ascor | MOWSE | Sequence |
|---|---|---|---|---|---|---|---|---|---|---|---|
| **FNBP4** | FNBP4 | S429 | | | | | | | **12.3** | **33.2** | R.ALEEGDGSVSGS*SPR.S |
| **FNBP4** | FNBP4 | S961S962S963 | | | | | | | **14.5** | **25.7** | R.ELDEEDNS*S*S*S*EEDRESTAQKR.I |
| **FNBP4** | FNBP4 | T461 | | | | | | | **27.8** | **28.3** | R.AT*SPESTSR.S |
| **FNBP4** | FNBP4 | S430 | | | | | | | **7.0** | **32.7** | R.ALEEGDGSVSGSS*PR.S |
| **Fodrin beta** | SPTBN1 | S2160S2161S2 | | | | | | | **4.3** | **22.6** | R.TS*S*KES*SPIPS*PTSDRK.A |
| **Fodrin beta** | SPTBN1 | S2165S2169 | | | | | | | **18.0** | **38.1** | K.ESS*PIPS*PTSDR.K |
| **Fodrin beta** | SPTBN1 | S2138 | | | | | | | **78.4** | **68.3** | K.GEQVSQNGLPAEQGS*PR.M |
| **Fodrin beta** | SPTBN1 | S2341 | | | | | | | **9.1** | **69.3** | R.AQTLPTSVVTITSESS*PGKR.E |
| **Fodrin beta** | SPTBN1 | S2358 | | | | | | | **100.0** | **32.7** | K.RFS*LFGK.K |
| **Fodrin beta** | SPTBN1 | Y17 | | | | | | | **5.5** | **60.5** | R.TSSISGPLSPAY*TGQVPYNYNQLEGR.F |
| **Fodrin beta** | SPTBN1 | S10 | | | | | | | **16.7** | **72.6** | R.TSSIS*GPLSPAYTGQVPYNYNQLEGR.F |
| **Fodrin beta** | SPTBN1 | S8 | | | | | | | **13.4** | **89.4** | R.TSS*ISGPLSPAYTGQVPYNYNQLEGR.F |
| **Fodrin beta** | SPTBN1 | S14 | | | | | | | **7.4** | **88.3** | R.TSSISGPLS*PAYTGQVPYNYNQLEGR.F |
| **Fodrin beta** | SPTBN1 | S8Y17 | | | | | | | **8.0** | **88.0** | R.TSS*ISGPLSPAY*TGQVPYNYNQLEGR.F |
| **Fodrin beta** | SPTBN1 | S7S8 | | | | | | | **6.2** | **150.4** | R.TS*S*ISGPLSPAYTGQVPYNYNQLEGR.F |
| **Fodrin beta** | SPTBN1 | S7Y17 | | | | | | | **-0.3** | **60.1** | R.TS*SISGPLSPAY*TGQVPYNYNQLEGR.F |
| **Fodrin beta** | SPTBN1 | S2160S2164S2 | | | | | | | **2.5** | **19.5** | R.TS*SKES*S*PIPS*PTSDRK.A |
| **Fodrin beta** | SPTBN1 | S8S14 | | | | | | | **9.9** | **75.1** | R.TSS*ISGPLS*PAYTGQVPYNYNQLEGR.F |
| **Fodrin beta** | SPTBN1 | S2319 | | | | | | | **23.0** | **19.8** | K.DDEEMNTWIQAISSAISSDKHEVSASTQS*TPASSR.A |
| **Fodrin beta** | SPTBN1 | S2303T2320 | | | | | | | **13.6** | **26.3** | K.DDEEMNTWIQAIS*SAISSDKHEVSASTQST*PASSR.A |
| **Fodrin beta** | SPTBN1 | S2307T2317 | | | | | | | **9.1** | **15.4** | K.DDEEMNTWIQAISSAIS*SDKHEVSAST*QSTPASSR.A |
| **Fodrin beta** | SPTBN1 | S2307S2319 | | | | | | | **5.4** | **14.3** | K.DDEEMNTWIQAISSAIS*SDKHEVSASTQS*TPASSR.A |
| **Fodrin beta** | SPTBN1 | S2319T2320 | | | | | | | **3.4** | **13.9** | K.DDEEMNTWIQAISSAISSDKHEVSASTQS*T*PASSR.A |
| **Fodrin beta** | SPTBN1 | S2314S2319T2 | | | | | | | **13.2** | **11.3** | K.DDEEMNTWIQAISSAISSDKHEVS*ASTQS*T*PASSR.A |
| **Fodrin beta** | SPTBN1 | T2317S2319T23 | | | | | | | **4.6** | **15.1** | K.DDEEMNTWIQAISSAISSDKHEVSAST*QS*T*PASSR.A |
| **Fodrin beta** | SPTBN1 | S2161S2164S2 | | | | | | | **8.2** | **19.3** | R.TSS*KES*S*PIPS*PTSDRK.A |
| **Fodrin beta** | SPTBN1 | S2338 | | | | | | | **10.8** | **27.5** | R.AQTLPTSVVTITS*ESSPGKR.E |
| **Fodrin beta** | SPTBN1 | S8S10 | | | | | | | **10.4** | **115.4** | R.TSS*IS*GPLSPAYTGQVPYNYNQLEGR.F |
| **Fodrin beta** | SPTBN1 | S8T18 | | | | | | | **2.9** | **45.2** | R.TSS*ISGPLSPAYT*GQVPYNYNQLEGR.F |
| **Fodrin beta** | SPTBN1 | S2314T2320 | | | | | | | **5.0** | **15.2** | K.DDEEMNTWIQAISSAISSDKHEVS*ASTQST*PASSR.A |
| **Fodrin beta** | SPTBN1 | S2303S2319 | | | | | | | **8.7** | **22.5** | K.DDEEMNTWIQAIS*SAISSDKHEVSASTQS*TPASSR.A |
| **Fodrin beta** | SPTBN1 | S2316S2319 | | | | | | | **7.5** | **22.0** | K.DDEEMNTWIQAISSAISSDKHEVSAS*TQS*TPASSR.A |
| **Fodrin beta** | SPTBN1 | S2304S2307S2 | | | | | | | **-1.5** | **12.4** | K.DDEEMNTWIQAISS*AIS*S*DKHEVSASTQSTPASSR.A |
| **Fodrin beta** | SPTBN1 | S2303S2319T2 | | | | | | | **10.7** | **21.1** | K.DDEEMNTWIQAIS*SAISSDKHEVSASTQS*T*PASSR.A |
| **Fodrin beta** | SPTBN1 | T2317T2320 | | | | | | | **11.9** | **13.1** | K.DDEEMNTWIQAISSAISSDKHEVSAST*QST*PASSR.A |
| **Fodrin beta** | SPTBN1 | S2164S2165 | | | | | | | **31.0** | **19.9** | K.ES*S*PIPSPTSDRK.A |
| **Fodrin beta** | SPTBN1 | T2159S2160S2 | | | | | | | **8.8** | **13.4** | R.T*S*S*KESSPIPS*PTSDRK.A |
| **Fodrin beta** | SPTBN1 | S2316T2320 | | | | | | | **9.5** | **12.8** | K.DDEEMNTWIQAISSAISSDKHEVSAS*TQST*PASSR.A |
| **Fodrin beta** | SPTBN1 | S2303T2317 | | | | | | | **-1.1** | **13.2** | K.DDEEMNTWIQAIS*SAISSDKHEVSAST*QSTPASSR.A |
| **Fodrin beta** | SPTBN1 | S2303S2314 | | | | | | | **-0.1** | **11.3** | K.DDEEMNTWIQAIS*SAISSDKHEVS*ASTQSTPASSR.A |
| **Fodrin beta** | SPTBN1 | S2304T2320 | | | | | | | **7.5** | **19.2** | K.DDEEMNTWIQAISS*AISSDKHEVSASTQST*PASSR.A |
| **Fodrin beta** | SPTBN1 | S2160S2161S2 | | | | | | | **4.5** | **23.8** | R.TS*S*KESS*PIPS*PTSDRK.A |
| **Fodrin beta** | SPTBN1 | S2102 | | | | | | | **49.2** | **46.2** | R.RPPS*PEPSTK.V |
| **Fodrin beta** | SPTBN1 | S2340 | | | | | | | **11.1** | **54.2** | R.AQTLPTSVVTITSES*SPGKR.E |
| **Fodrin beta** | SPTBN1 | S7 | | | | | | | **2.5** | **88.0** | R.TS*SISGPLSPAYTGQVPYNYNQLEGR.F |
| **Fodrin beta** | SPTBN1 | T2297 | | | | | | | | **12.8** | K.DDEEMNT*WIQAISSAISSDKHEVSASTQSTPASSR.A |
| **Fodrin beta** | SPTBN1 | S2160S2161S2 | | | | | | | **9.1** | **11.7** | R.TS*S*KESSPIPS*PTSDRK.A |
| **Fodrin beta** | SPTBN1 | T2328S2340 | | | | | | | **15.6** | **42.9** | R.AQT*LPTSVVTITSES*SPGKR.E |
| **Fodrin beta** | SPTBN1 | T2320 | | | | | | | **6.8** | **36.8** | K.DDEEMNTWIQAISSAISSDKHEVSASTQST*PASSR.A |
| **Fodrin beta** | SPTBN1 | Y17T18 | | | | | | | **3.5** | **12.4** | R.TSSISGPLSPAY*T*GQVPYNYNQLEGR.F |
| **Fodrin beta** | SPTBN1 | T2328S2341 | | | | | | | **18.2** | **11.3** | R.AQT*LPTSVVTITSESS*PGKR.E |
| **Fodrin beta** | SPTBN1 | T2328 | | | | | | | **56.8** | **83.8** | R.AQT*LPTSVVTITSESSPGKR.E |
| **Fodrin beta** | SPTBN1 | S7S14 | | | | | | | **0.7** | **106.7** | R.TS*SISGPLS*PAYTGQVPYNYNQLEGR.F |
| **Fodrin beta** | SPTBN1 | T2317 | | | | | | | **7.3** | **23.1** | K.DDEEMNTWIQAISSAISSDKHEVSAST*QSTPASSR.A |
| **Fodrin beta** | SPTBN1 | S2164S2169 | | | | | | | **16.7** | **15.7** | K.ES*SPIPS*PTSDRK.A |
| **Forkhead box J3** | FOXJ3 | S223 | | | | | | | **10.9** | **29.8** | K.VTLYNTDQDGSDS*PR.S |
| **Forkhead box K1** | FOXK1 | T436S441 | | | | | | | **32.4** | **36.4** | R.SGGLQT*PECLS*R.E |
| **Forkhead box K1** | FOXK1 | S445 | | | | | | | **95.6** | **47.3** | R.EGS*PIPHDPEFGSK.L |
| **Forkhead box K1** | FOXK1 | S416S420S428 | | | | | | | **20.8** | **53.2** | R.S*APAS*PTHPGLMS*PR.S |
| **Forkhead box K1** | FOXK1 | S213S223 | | | | | | | **43.0** | **25.8** | K.EEAPAS*PLRPLYPQIS*PLK.I |
| **Forkhead box K1** | FOXK1 | S299 | | | | | | | **6.7** | **41.2** | K.AASEQQADTSGGDS*PKDESKPPFSYAQLIVQAISSAQDR.Q |
| **Forkhead box K1** | FOXK1 | S309 | | | | | | | **2.1** | **39.2** | K.AASEQQADTSGGDSPKDESKPPFS*YAQLIVQAISSAQDR.Q |

Peak Area: <-10, -5, -3, 0, 3, 5, >10

%CV: 0, 29, 42, 71, 86, >100

**White dots-** Significant change in peptide abundance at **5** %FDR compared to the timepoint with the minimum peak area for a given PSM

| Protein Name | Gene | Phosphosites | CarT (0m, 2m, 5m) | RajiB (0m, 2m, 5m) | Ascor | MOWSE | Sequence |
|---|---|---|---|---|---|---|---|
| **Forkhead box K1** | FOXK1 | S416S420T422 | | | **14.0** | **27.9** | R.S*APAS*PT*HPGLMSPR.S |
| **Forkhead box K1** | FOXK1 | S416S420 | | | **47.3** | **45.0** | R.S*APAS*PTHPGLMSPR.S |
| **Forkhead box K1** | FOXK1 | Y219S223 | | | **39.4** | **18.4** | K.EEAPASPLRPLY*PQIS*PLK.I |
| **Forkhead box K1** | FOXK1 | S295S299 | | | **5.7** | **51.0** | K.AASEQQADTS*GGDS*PKDESKPPFSYAQLIVQAISSAQDR.Q |
| **Forkhead box K1** | FOXK1 | T294S299 | | | **7.3** | **55.4** | K.AASEQQADT*SGGDS*PKDESKPPFSYAQLIVQAISSAQDR.Q |
| **Forkhead box K1** | FOXK1 | S309Y310 | | | **9.6** | **51.8** | K.AASEQQADTSGGDSPKDESKPPFS*Y*AQLIVQAISSAQDR.Q |
| **Forkhead box K1** | FOXK1 | S213 | | | **14.0** | **27.6** | K.EEAPAS*PLRPLYPQISPLK.I |
| **Forkhead box K1** | FOXK1 | S441 | | | **11.1** | **18.0** | R.SGGLQTPECLS*R.E |
| **Forkhead box K1** | FOXK1 | S416T422 | | | **17.3** | **33.1** | R.S*APASPT*HPGLMSPR.S |
| **Forkhead box K1** | FOXK1 | S420T422S428 | | | **7.4** | **30.1** | R.SAPAS*PT*HPGLMS*PR.S |
| **Forkhead box K2** | FOXK2 | S398 | | | **100.0** | **49.9** | R.EGS*PAPLEPEPGAAQPK.L |
| **Forkhead box K2** | FOXK2 | S424S431 | | | **13.2** | **65.1** | R.FAQS*APGSPLS*SQPVLITVQR.Q |
| **Forkhead box K2** | FOXK2 | S428S431 | | | **9.1** | **38.2** | R.FAQSAPGS*PLS*SQPVLITVQR.Q |
| **Forkhead box K2** | FOXK2 | S428 | | | **20.9** | **45.3** | R.FAQSAPGS*PLSSQPVLITVQR.Q |
| **Forkhead box K2** | FOXK2 | S424S428 | | | **22.0** | **39.0** | R.FAQS*APGS*PLSSQPVLITVQR.Q |
| **Forkhead box P4** | FOXP4 | S443 | | | **-0.3** | **45.0** | R.SS*DKFCSPISSELAQNHEFYK.N |
| **Forkhead box protein O3A** | ZNF286B | S7S12 | | | **18.1** | **13.8** | -.MAEAPAS*PAPLS*PLEVELDPEFEPQSRPR.S |
| **Formin binding protein 1** | FNBP1 | S497 | | | **19.2** | **61.8** | R.RQS*GLYDSQNPPTVNNCAQDR.E |
| **Formin binding protein 1** | FNBP1 | T507 | | | **-1.4** | **17.0** | R.RQSGLYDSQNPPT*VNNCAQDRESPDGSYTEEQSQESEMK.V |
| **Formin binding protein 1** | FNBP1 | S521 | | | **10.0** | **22.1** | R.RQSGLYDSQNPPTVNNCAQDRESPDGS*YTEEQSQESEMK.V |
| **Formin binding protein 1** | FNBP1 | S497T507 | | | **12.9** | **20.7** | R.RQS*GLYDSQNPPT*VNNCAQDRESPDGSYTEEQSQESEMK.V |
| **Formin binding protein 1** | FNBP1 | S497Y500 | | | | **28.8** | R.RQS*GLY*DSQNPPTVNNCAQDRESPDGSYTEEQSQESEMK.V |
| **Formin binding protein 1** | FNBP1 | S296S299 | | | **19.0** | **40.6** | R.TVS*DNS*LSNSR.G |
| **Formin binding protein 1** | FNBP1 | S296 | | | **25.3** | **63.3** | R.TVS*DNSLSNSR.G |
| **Formin binding protein 1** | FNBP1 | T294 | | | **9.8** | **27.9** | R.T*VSDNSLSNSR.G |
| **Formin binding protein 1** | FNBP1 | S517S521 | | | **14.6** | **95.3** | R.ES*PDGS*YTEEQSQESEMK.V |
| **Formin binding protein 1** | FNBP1 | T507S521 | | | **5.3** | **18.6** | R.RQSGLYDSQNPPT*VNNCAQDRESPDGS*YTEEQSQESEMK.V |
| **Formin binding protein 1** | FNBP1 | S497Y522 | | | **13.6** | **21.9** | R.RQS*GLYDSQNPPTVNNCAQDRESPDGSY*TEEQSQESEMK.V |
| **Formin binding protein 1** | FNBP1 | S497S521 | | | **13.0** | **23.3** | R.RQS*GLYDSQNPPTVNNCAQDRESPDGS*YTEEQSQESEMK.V |
| **Formin binding protein 1** | FNBP1 | S497S517S521 | | | **9.1** | **20.2** | R.RQS*GLYDSQNPPTVNNCAQDRES*PDGS*YTEEQSQESEMK.V |
| **Formin binding protein 1** | FNBP1 | T507S517 | | | **5.4** | **42.2** | R.QSGLYDSQNPPT*VNNCAQDRES*PDGSYTEEQSQESEMK.V |
| **Formin binding protein 1** | FNBP1 | T294S296 | | | **20.9** | **34.0** | R.T*VS*DNSLSNSR.G |
| **Formin binding protein 1** | FNBP1 | T294S299 | | | **20.1** | **29.8** | R.T*VSDNS*LSNSR.G |
| **Formin binding protein 1** | FNBP1 | Y500 | | | **5.6** | **31.8** | R.RQSGLY*DSQNPPTVNNCAQDR.E |
| | FMNL1 | S184 | | | **67.8** | **48.9** | K.NKPLEQS*VEDLSK.G |
| **Formin like** | FMNL1 | S624 | | | **100.0** | **56.2** | R.RDS*ELGPGVK.A |
| **Formin like 2** | FMNL3 | T622 | | | **100.0** | **22.2** | K.KPIKT*K.F |
| **Formin like 3** | FMNL3 | S174 | | | **38.8** | **67.0** | R.S*IEDLQPPSALSAPFTNSLAR.S |
| **Formin like 3** | FMNL3 | S182 | | | **3.1** | **24.0** | R.SIEDLQPPS*ALSAPFTNSLAR.S |
| **Formin-like protein 1** | FMNL1 | S681 | | | **48.8** | **85.8** | R.RPPGPPLQVTSDLS*L.- |
| **Formin-like protein 1** | FMNL1 | T677 | | | **2.2** | **27.8** | R.RRPPGPPLQVT*SDLSL.- |
| **Forty-two-three domain containing 1** | FYTTD1 | S23 | | | **78.7** | **84.0** | R.S*NENLDKIDMSLDDIIK.L |
| **FRAS1-related extracellular matrix protein 3** | FREM3 | Y1044 | | | **-0.3** | **19.6** | K.DSY*QWVVGNSIIEK.V |
| | WAPL | S264S269 | | | **22.3** | **34.3** | K.RPES*PSEIS*PIKGSVR.T |
| **Friend of EBNA2** | WAPL | S120 | | | **36.0** | **82.3** | K.VEEESTGDPFGFDS*DDESLPVSSK.N |
| **Friend of EBNA2** | WAPL | S502S504 | | | **54.6** | **79.8** | K.IKYFGFDDLS*ES*EDDEDDDCQVER.K |
| **Friend of EBNA2** | WAPL | S111 | | | **6.5** | **19.8** | K.VEEES*TGDPFGFDSDDESLPVSSK.N |
| **Friend of EBNA2** | WAPL | S266S269 | | | **13.1** | **18.9** | K.RPESPS*EIS*PIKGSVR.T |
| **Friend of EBNA2** | WAPL | S264S266 | | | **3.0** | **47.8** | K.RPES*PS*EISPIKGSVR.T |
| **FtsJ homolog 3** | FTSJ3 | S335S336 | | | **40.2** | **155.7** | K.ALDISLS*S*GEEDEGDEEDSTAGTTK.Q |
| **FtsJ homolog 3** | FTSJ3 | T467S468S471 | | | **13.2** | **55.1** | R.DDIYVSDVEDDGDDT*S*LDS*DLDPEELAGVR.G |
| **FtsJ homolog 3** | FTSJ3 | S458T467S468 | | | **24.4** | **94.1** | R.DDIYVS*DVEDDGDDT*S*LDS*DLDPEELAGVR.G |
| **FtsJ homolog 3** | FTSJ3 | Y456S458T467 | | | **11.6** | **85.1** | R.DDIY*VS*DVEDDGDDT*SLDS*DLDPEELAGVR.G |
| **FtsJ homolog 3** | FTSJ3 | S458T467S471 | | | **2.2** | **64.3** | R.DDIYVS*DVEDDGDDT*SLDS*DLDPEELAGVR.G |
| **FtsJ homolog 3** | FTSJ3 | S458T467S468 | | | **6.6** | **35.3** | R.DDIYVS*DVEDDGDDT*S*LDSDLDPEELAGVR.G |
| **FtsJ homolog 3** | FTSJ3 | T436S448 | | | **9.1** | **64.0** | R.GHQLLEEVT*QGDMSAADTFLS*DLPR.D |
| **FtsJ homolog 3** | FTSJ3 | S441S448 | | | **8.4** | **45.4** | R.GHQLLEEVTQGDMS*AADTFLS*DLPR.D |
| **FtsJ homolog 3** | FTSJ3 | S333S335S336 | | | **44.1** | **67.7** | K.ALDIS*LS*S*GEEDEGDEEDSTAGTTK.Q |
| **FtsJ homolog 3** | FTSJ3 | Y456S458S468 | | | **4.4** | **74.7** | R.DDIY*VS*DVEDDGDDTS*LDS*DLDPEELAGVR.G |
| **FtsJ homolog 3** | FTSJ3 | S333S335 | | | **25.1** | **52.9** | K.ALDIS*LS*SGEEDEGDEEDSTAGTTK.Q |
| **FtsJ homolog 3** | FTSJ3 | Y456S458T467 | | | **11.1** | **58.3** | R.DDIY*VS*DVEDDGDDT*S*LDSDLDPEELAGVR.G |
| **FtsJ homolog 3** | FTSJ3 | S335S336S347 | | | **7.5** | **33.3** | K.ALDISLS*S*GEEDEGDEEDS*TAGTTK.Q |

**Peak Area**: <-10, -5, -3, 0, 3, 5, >10

**%CV**: 0, 29, 42, 71, 86, >100

**White dots-** Significant change in peptide abundance at **5**%FDR compared to the timepoint with the minimum peak area for a given PSM

CarT (0m, 2m, 5m) — RajiB (0m, 2m, 5m)

| Protein Name | Gene | Phosphosites | Ascor | MOWSE | Sequence |
|---|---|---|---|---|---|
| **FUS** | FUS | S277 | 51.8 | 77.6 | R.HDS*EQDNSDNNTIFVQGLGENVTIESVADYFK.Q |
| **FUS interacting protein 1** | SRSF10 | T255S256 | 5.3 | 16.2 | R.SWT*S*PK.S |
| **FUS interacting protein 1** | SRSF10 | S133 | 28.6 | 27.8 | R.S*FDYNYRR.S |
| **FUS interacting protein 1** | SRSF10 | S131S133 | 44.5 | 38.7 | R.S*RS*FDYNYR.R |
| **FUS interacting protein 1** | SRSF10 | S156S158 | 24.4 | 18.1 | R.S*RS*HSDNDRPNCSWNTQYSSAYYTSR.K |
| **FUS interacting protein 1** | SRSF10 | S158 | 30.0 | 66.3 | R.S*HSDNDRPNCSWNTQYSSAYYTSR.K |
| **FUS interacting protein 1** | SRSF10 | S158S160 | 36.2 | 80.5 | R.S*HS*DNDRPNCSWNTQYSSAYYTSR.K |
| **FUS interacting protein 1** | SRSF10 | S156S158S160 | 60.5 | 49.3 | R.S*RS*HS*DNDRPNCSWNTQYSSAYYTSR.K |
| **FUS interacting protein 1** | SRSF10 | S171S173 | 100.0 | 12.2 | R.NRS*FS*R.S |
| **FUS interacting protein 1** | SRSF10 | S158S160 | 100.0 | 22.1 | R.S*HS*DNDRFK.H |
| **FUS interacting protein 1** | SRSF10 | S253S256 | 16.7 | 19.9 | R.S*WTS*PK.S |
| **FUS interacting protein 1** | SRSF10 | S168 | 9.0 | 26.7 | R.SHSDNDRPNCS*WNTQYSSAYYTSR.K |
| **FUS interacting protein 1** | SRSF10 | S156S158T171 | 14.5 | 32.4 | R.S*RS*HSDNDRPNCSWNT*QYSSAYYTSR.K |
| **FUS interacting protein 1** | SRSF10 | S160S168T171 | 9.4 | 24.4 | R.SRSHS*DNDRPNCS*WNT*QYSSAYYTSR.K |
| **FUS interacting protein 1** | SRSF10 | S156S168 | 9.7 | 41.8 | R.S*RSHSDNDRPNCS*WNTQYSSAYYTSR.K |
| **FUS interacting protein 1** | SRSF10 | S156T171 | 3.4 | 27.9 | R.S*RSHSDNDRPNCSWNT*QYSSAYYTSR.K |
| **FUS interacting protein 1** | SRSF10 | S160 | 1.4 | 40.9 | R.SHS*DNDRPNCSWNTQYSSAYYTSR.K |
| **FUS interacting protein 1** | SRSF10 | S156S158S160 | 100.0 | 17.6 | R.S*RS*HS*DNDRFK.H |
| **FUS interacting protein 1** | SRSF10 | S251S253T255 | 2.7 | 11.1 | R.S*RS*WT*SPK.S |
| **FUS interacting protein 1** | SRSF10 | S156S158S168 | 19.2 | 29.2 | R.S*RS*HSDNDRPNCS*WNTQYSSAYYTSR.K |
| **FUS interacting protein 1** | CCNL2 | S121S123 | 10.2 | 11.2 | R.S*RS*YER.T |
| **FUS interacting protein 1** | SRSF10 | S256 | 22.5 | 16.5 | R.SWTS*PK.S |
| **FUS interacting protein 1** | SRSF10 | S156S160S168 | 7.3 | 22.9 | R.S*RSHS*DNDRPNCS*WNTQYSSAYYTSR.K |
| **FUS interacting protein 1** | SRSF10 | Y136 | 1.9 | 19.4 | R.SFDY*NYRR.S |
| **FUS interacting protein 1** | SRSF10 | S156S168T171 | -3.3 | 41.7 | R.S*RSHSDNDRPNCS*WNT*QYSSAYYTSR.K |
| **FUS interacting protein 1** | SRSF10 | S160S168 | -0.2 | 20.5 | R.SHS*DNDRPNCS*WNTQYSSAYYTSR.K |
| **FXR1** | FXR1 | S409 | 15.7 | 32.5 | R.RGPNYTSGYGTNSELSNPS*ETESER.K |
| **FXR1** | FXR1 | S406S409 | 11.0 | 14.7 | R.RGPNYTSGYGTNSELS*NPS*ETESER.K |
| **FXR2** | FXR2 | S601S603 | 18.7 | 56.4 | R.TDGS*IS*GDRQPVTVADYISR.A |
| **FXR2** | FXR2 | S601 | 7.5 | 25.1 | R.TDGS*ISGDRQPVTVADYISR.A |
| **FYB** | FYB1 | T443 | -1.2 | 39.7 | K.SPVNEDNQDGVT*HSDGAGNLDEEQDSEGETYEDIEASK.E |
| **FYB** | FYB1 | S432 |  | 76.2 | K.S*PVNEDNQDGVTHSDGAGNLDEEQDSEGETYEDIEASK.E |
| **FYB** | FYB1 | S457 | 10.8 | 30.0 | K.SPVNEDNQDGVTHSDGAGNLDEEQDS*EGETYEDIEASK.E |
| **Fyn** | YES1 | Y420 | 12.1 | 38.5 | R.LIEDNEY*TAR.Q |
| **G patch domain containing 2** | GPATCH2 | S115S117 | 100.0 | 54.0 | K.DHS*DS*DDQMLVAK.R |
| **G patch domain containing 8** | GPATCH8 | S1033 | 14.0 | 22.8 | R.S*QSPHYFR.S |
| **G patch domain containing 8** | GPATCH8 | S1035 | 10.4 | 25.1 | R.SQS*PHYFR.S |
| **G protein coupled purinergic receptor P2Y8** | P2RY8 | S324 | 39.9 | 30.9 | R.RES*LFSAR.T |
| **G protein coupled purinergic receptor P2Y8** | P2RY8 | S324S327 | 100.0 | 44.6 | R.RES*LFS*AR.T |
| **G protein coupled purinergic receptor P2Y8** | P2RY8 | S335 | 45.6 | 16.1 | R.S*EAGAHPEGMEGATRPGLQR.Q |
| **G protein coupled receptor kinase 6** | GRK6 | S484 | 13.9 | 50.4 | K.DVLDIEQFS*TVK.G |
| **G protein coupled receptor kinase 6** | GRK6 | S484T485 | 36.4 | 46.3 | K.DVLDIEQFS*T*VKGVELEPTDQDFYQK.F |
| **G protein dependent receptor kinase 2** | GRK2 | S670 | 71.9 | 50.3 | R.S*PVVELSK.V |
| **G protein signalling modulator 3** | GPSM3 | S59 | 8.1 | 57.1 | R.SASLLS*LQTELLLDLVAEAQSR.R |
| **G protein signalling modulator 3** | GPSM3 | S54 | 10.1 | 51.7 | R.S*ASLLSLQTELLLDLVAEAQSR.R |
| **G protein signalling modulator 3** | GPSM3 | S54S59 | 12.3 | 68.9 | R.S*ASLLS*LQTELLLDLVAEAQSR.R |
| **G protein signalling modulator 3** | GPSM3 | S54S56 | 24.8 | 31.5 | R.S*AS*LLSLQTELLLDLVAEAQSR.R |
| **G protein signalling modulator 3** | GPSM3 | S54T62 | 7.2 | 84.5 | R.S*ASLLSLQT*ELLLDLVAEAQSR.R |
| **G protein signalling modulator 3** | GPSM3 | T62 | 8.3 | 73.7 | R.SASLLSLQT*ELLLDLVAEAQSR.R |
| **G protein signalling modulator 3** | GPSM3 | S59T62 | 6.7 | 69.4 | R.SASLLS*LQT*ELLLDLVAEAQSR.R |
| **G protein signalling modulator 3** | GPSM3 | S56S59 | 4.6 | 64.5 | R.SAS*LLS*LQTELLLDLVAEAQSR.R |
| **G protein signalling modulator 3** | GPSM3 | S56T62 | 9.1 | 76.6 | R.SAS*LLSLQT*ELLLDLVAEAQSR.R |
| **G protein signalling modulator 3** | GPSM3 | S56 | -0.1 | 26.7 | R.SAS*LLSLQTELLLDLVAEAQSR.R |
| **G-protein signalling modulator 1** | GPSM1 | S469 | 23.9 | 84.4 | R.APS*SDEECFFDLLTK.F |
| **G-protein signalling modulator 1** | GPSM1 | S470 | 10.2 | 44.0 | R.APSS*DEECFFDLLTK.F |
| **Ga55** | TACC1 | S276 | 9.2 | 80.4 | K.ASYHFSPEELDENTS*PLLGDAR.F |
| **Gamma synergin** | SYNRG | S854S855 | 15.5 | 33.3 | K.HVMSDS*S*LDLPTVSGQHPPAADIEDLK.Y |
| **Gamma synergin** | SYNRG | S852S854 | 19.0 | 21.1 | K.HVMS*DS*SLDLPTVSGQHPPAADIEDLK.Y |
| **Gamma synergin** | SYNRG | S752 | 12.4 | 13.6 | R.QLS*LEGSGLGVEDLKDNTPSGK.S |
| **Gamma synergin** | SYNRG | S935 | 12.1 | 56.3 | K.ETSFGS*SENITMTSLSK.V |
| **Gamma synergin** | SYNRG | S852S855 | 14.7 | 41.6 | K.HVMS*DSS*LDLPTVSGQHPPAADIEDLK.Y |
| **Gamma synergin** | SYNRG | S1075 | 10.6 | 12.8 | R.SLS*LGDK.E |

Peak Area: <-10, -5, -3, 0, 3, 5, >10

%CV: 0, 29, 42, 71, 86, >100

**White dots-** Significant change in peptide abundance at **5** %FDR compared to the timepoint with the minimum peak area for a given PSM

CarT (0m, 2m, 5m) RajiB (0m, 2m, 5m)

| Protein Name | Gene | Phosphosites | Ascor | MOWSE | Sequence |
|---|---|---|---|---|---|
| **Gamma synergin** | SYNRG | S812 | **46.8** | **86.9** | K.S*LDLPSIGGSSVGK.E |
| **Gamma-tubulin complex component 3** | TUBGCP3 | S896 | **27.8** | **23.7** | R.LRVS*LGTR.G |
| **Gasdermin domain containing 1** | GSDMD | S252 | **11.9** | **76.0** | R.STS*EGAWPQLPSGLSMMR.C |
| **Gasdermin domain containing 1** | GSDMD | T251 | **-0.3** | **75.6** | R.ST*SEGAWPQLPSGLSMMR.C |
| **GATA binding protein 3** | GATA3 | S110S115 | **13.0** | **35.9** | K.ALGSHHTAS*PWNLS*PFSK.T |
| **GATA binding protein 3** | GATA3 | S105S115 | **11.2** | **18.2** | K.ALGS*HHTASPWNLS*PFSK.T |
| **GATA zinc finger domain containing 2B** | GATAD2B | T120S135 | **5.0** | **47.1** | R.GRLT*PSPDIIVLSDNEASS*PR.S |
| **GATA zinc finger domain containing 2B** | GATAD2B | T120S122S129 | **67.5** | **38.9** | R.GRLT*PS*PDIIVLS*DNEASS*PR.S |
| **GATA zinc finger domain containing 2B** | GATAD2B | T120S122S135 | **50.0** | **12.0** | R.GRLT*PS*PDIIVLSDNEASS*PR.S |
| **GATA zinc finger domain containing 2B** | GATAD2B | T120S122S129 | **50.0** | **30.2** | R.GRLT*PS*PDIIVLS*DNEAS*SPR.S |
| **GATA zinc finger domain containing 2B** | GATAD2B | T120S122S134 | **37.1** | **10.7** | R.GRLT*PS*PDIIVLSDNEAS*SPR.S |
| **GCF** | GCFC2 | T213S214S217 | **60.6** | **82.7** | R.NEET*S*EES*QEDEKQDTWEQQQMR.K |
| **GCF** | GCFC2 | S429S430 | **10.2** | **65.9** | R.VLSGNCNHQEGTS*S*DDELPSAEMIDFQK.S |
| **GCF** | GCFC2 | S419T428 | **6.1** | **12.2** | R.VLS*GNCNHQEGT*SSDDELPSAEMIDFQK.S |
| **GCF** | GCFC2 | T428S430 | **0.8** | **16.1** | R.VLSGNCNHQEGT*SS*DDELPSAEMIDFQK.S |
| **GCF** | GCFC2 | S16S17S19 | **27.0** | **26.0** | R.AADS*S*DS*DGAEESPAEPGAPR.E |
| **GEM interacting protein** | GMIP | S437S440 | **11.3** | **15.5** | R.SLDS*PTS*SPGAGTR.Q |
| **GEM interacting protein** | GMIP | S19 | **20.1** | **25.6** | R.KRYS*DIFR.S |
| **GEM interacting protein** | GMIP | S437S441 | **12.6** | **65.2** | R.SLDS*PTSS*PGAGTR.Q |
| **GEM interacting protein** | GMIP | T439S441 | **12.6** | **46.4** | R.SLDSPT*SS*PGAGTR.Q |
| **GEM interacting protein** | GMIP | Y18 | | **31.0** | R.KRY*SDIFR.S |
| **Gemin 5** | GEMIN5 | S778 | **14.9** | **44.4** | K.ENSGPVENGVS*DQEGEEQAR.E |
| **General transcription factor 2 I** | GTF2I | S674 | **12.2** | **108.0** | R.S*PGSNSKVPEIEVTVEGPNNNNPQTSAVR.T |
| **General transcription factor 2 I** | GTF2I | S679 | **2.8** | **65.1** | R.SPGSNS*KVPEIEVTVEGPNNNNPQTSAVR.T |
| **General transcription factor 2 I** | GTF2I | S818 | | **33.4** | K.ES*TSSKSPPR.K |
| **General transcription factor 2 I** | GTF2I | S830 | **17.4** | **42.0** | K.INS*SPNVNTTASGVEDLNIIQVTIPDDDNER.L |
| **General transcription factor 2 I** | GTF2I | S677 | **8.7** | **80.2** | R.SPGS*NSKVPEIEVTVEGPNNNNPQTSAVR.T |
| **General transcription factor 2 I** | GTF2I | S823 | **6.8** | **50.8** | K.ESTSSKS*PPR.K |
| **General transcription factor 2 I** | GTF2I | S831 | **22.8** | **67.7** | R.KINSS*PNVNTTASGVEDLNIIQVTIPDDDNER.L |
| **General transcription factor IIA, 1, 19/37kDa** | GTF2A1 | S316S321 | **16.1** | **80.2** | K.DGAEDGQVEEEPLNS*EDDVS*DEEGQELFDTENVVVCQYDK.I |
| **General transcription factor IIIC, polypeptide 2, beta 110kDa** | GTF3C2 | S892 | **27.8** | **56.0** | R.AHFNAMFQPS*SPTR.R |
| **General transcription factor IIIC, polypeptide 2, beta 110kDa** | GTF3C2 | S893 | **21.2** | **55.9** | R.AHFNAMFQPSS*PTR.R |
| **General transcription factor IIIC, polypeptide 2, beta 110kDa** | GTF3C2 | S167 | **32.1** | **28.7** | K.DLDRPESQS*PK.R |
| **General transcription factor IIIC, polypeptide 2, beta 110kDa** | GTF3C2 | S220 | **7.0** | **14.4** | K.VSS*PTKPK.K |
| **GFAT** | GFPT1 | S243 | **23.9** | **92.6** | R.VDS*TTCLFPVEEK.A |
| **GIGYF1** | GIGYF1 | S862 | **17.0** | **27.7** | R.S*SPSLSDSYSHLSGRPIR.K |
| **Girdin** | CCDC88A | S1491T1509 | **8.1** | **51.4** | R.SMS*MNDLVQSMVLAGQWTGST*ENLEVPDDISTGKR.R |
| **Girdin** | CCDC88A | S1491T1506 | **15.2** | **38.7** | R.SMS*MNDLVQSMVLAGQWT*GSTENLEVPDDISTGKR.R |
| **Girdin** | CCDC88A | S1489S1491 | **3.2** | **20.3** | R.S*MS*MNDLVQSMVLAGQWTGSTENLEVPDDISTGK.R |
| **Girdin** | CCDC88A | S1491S1508 | **4.6** | **42.6** | R.SMS*MNDLVQSMVLAGQWTGS*TENLEVPDDISTGKR.R |
| **GIT1** | GIT1 | S385S388 | **13.4** | **94.5** | R.SQSDLDDQHDYDS*VAS*DEDTDQEPLR.S |
| **GIT1** | GIT1 | S362 | **30.2** | **93.9** | K.SLSS*PTDNLELSLR.S |
| **GIT1** | GIT1 | S361 | **11.4** | **83.3** | K.SLS*SPTDNLELSLR.S |
| **GIT1** | GIT1 | S414 | **5.8** | **63.0** | R.SMDSS*DLSDGAVTLQEYLELKK.A |
| **GIT1** | GIT1 | Y383S385 | **14.7** | **26.4** | R.SQSDLDDQHDY*DS*VASDEDTDQEPLR.S |
| **GIT1** | GIT1 | S410 | **37.7** | **81.5** | R.S*MDSSDLSDGAVTLQEYLELKK.A |
| **GIT1** | GIT1 | S592 | **7.4** | **13.4** | R.HGS*GADSDYENTQSGDPLLGLEGK.R |
| **GIT1** | GIT1 | S592S596 | **38.4** | **55.3** | R.HGS*GADS*DYENTQSGDPLLGLEGK.R |
| **GIT1** | GIT1 | Y383S388 | **27.5** | **92.3** | R.SQSDLDDQHDY*DSVAS*DEDTDQEPLR.S |
| **GIT1** | GIT1 | S413 | **-0.3** | **76.0** | R.SMDS*SDLSDGAVTLQEYLELK.K |
| **GIT1** | GIT1 | S417 | **2.1** | **31.7** | R.SMDSSDLS*DGAVTLQEYLELKK.A |
| **GL004 protein** | MFF | S157 | **35.1** | **34.6** | R.SMS*ENAVR.Q |
| **GL004 protein** | MFF | S155 | **9.2** | **28.3** | R.S*MSENAVR.Q |
| **Glucocorticoid induced transcript 1** | GLCCI1 | S76 | **7.4** | **32.9** | R.GS*QHSPTRPPVAAAAASLGSLPGPGAAR.G |
| **Glucocorticoid induced transcript 1** | GLCCI1 | T177 | **3.2** | **31.8** | R.TSSLDTIT*GPYLTGQWPR.D |
| **Glucocorticoid induced transcript 1** | GLCCI1 | S258 | **59.9** | **45.0** | K.DRQS*PLHGNHITISHTQATGSR.S |
| **Glucocorticoid induced transcript 1** | GLCCI1 | T175 | **3.0** | **45.9** | R.TSSLDT*ITGPYLTGQWPR.D |
| **Glucocorticoid induced transcript 1** | GLCCI1 | S223 | **41.2** | **62.4** | R.SAS*WGSADQLK.E |
| **Glucocorticoid induced transcript 1** | GLCCI1 | S79 | **3.5** | **41.8** | R.GSQHS*PTRPPVAAAAASLGSLPGPGAAR.G |
| **Glucocorticoid induced transcript 1** | GLCCI1 | S171 | **-0.4** | **61.2** | R.TS*SLDTITGPYLTGQWPR.D |
| **Glucocorticoid induced transcript 1** | GLCCI1 | S172 | **-0.4** | **57.5** | R.TSS*LDTITGPYLTGQWPR.D |
| **Glucocorticoid receptor DNA binding factor 1** | ARHGAP3 | S1150 | **46.9** | **53.5** | R.KVS*IVSKPVLYR.T |

Peak Area: <-10, -5, -3, 0, 3, 5, >10

%CV: 0, 29, 42, 71, 86, >100

**White dots-** Significant change in peptide abundance at **5** %FDR compared to the timepoint with the minimum peak area for a given PSM

CarT: 0m 2m 5m; RajiB: 0m 2m 5m

| Protein Name | Gene | Phosphosites | Ascor | MOWSE | Sequence |
|---|---|---|---|---|---|
| Glucocorticoid receptor DNA binding factor 1 | ARHGAP3 | S1179 | 32.3 | 54.4 | R.TSFSVGS*DDELGPIR.K |
| Glucocorticoid receptor DNA binding factor 1 | ARHGAP3 | S975 | 1.2 | 30.9 | R.AGS*PLCNSNLQDSEEDIEPSYSLFR.E |
| Glucocorticoid receptor DNA binding factor 1 | ARHGAP3 | S980S985 | 19.1 | 61.0 | R.AGSPLCNS*NLQDS*EEDIEPSYSLFR.E |
| Glucocorticoid receptor DNA binding factor 1 | ARHGAP3 | S1070 | 18.0 | 78.7 | K.S*VSSSPWLPQDGFDPSDYAEPMDAVVKPR.N |
| Glucocorticoid receptor DNA binding factor 1 | ARHGAP3 | S970 | 60.6 | 86.6 | K.NIIEATHMYDNAAEACSTTEEVFNS*PR.A |
| Glucocorticoid receptor DNA binding factor 1 | ARHGAP3 | S975S985 | 37.3 | 64.4 | R.AGS*PLCNSNLQDS*EEDIEPSYSLFR.E |
| Glutamate-rich WD repeat-containing protein 1 | GRWD1 | S119S122 | 63.6 | 73.0 | R.MHNLHGTKPPPS*EGS*DEEEEEEDEEDEEER.K |
| Glutamate-rich WD repeat-containing protein 1 | GRWD1 | T114S122 | 10.8 | 64.2 | R.MHNLHGT*KPPPSEGS*DEEEEEEDEEDEEER.K |
| Glutamate-rich WD repeat-containing protein 1 | GRWD1 | T114S119 | 13.5 | 74.5 | R.MHNLHGT*KPPPS*EGSDEEEEEEDEEDEEER.K |
|  | CAD | S1859 | 100.0 | 49.9 | R.AS*DPGLPAEEPK.E |
| Glutamine dependent carbamoyl phosphate synthase | CAD | S1406 | 36.8 | 13.4 | R.RLS*SFVTK.G |
| Glutamine--tRNA ligase (Fragment) | QARS | T47 | 100.0 | 22.5 | R.GLT*LAQGGVK.W |
| Glutamine-fructose-6-phosphate transaminase 2 | GFPT2 | Y479S494 | 3.6 | 12.7 | K.AY*TSQFISLVM#FGLM#M#S*EDR.I |
| Glutamine-rich protein 1 | QRICH1 | T347 | 2.3 | 19.9 | R.GDPQQQSITHIAIPQEAYNAVHVSGSPT*ALAAVK.L |
| Glutamine-rich protein 1 | QRICH1 | S343 | -0.5 | 15.2 | R.GDPQQQSITHIAIPQEAYNAVHVS*GSPTALAAVK.L |
| Glutamyl-prolyl-tRNA synthetase | EPRS | S885 | 16.1 | 45.6 | K.EYIPGQPPLSQSSDS*SPTR.N |
| Glutamyl-prolyl-tRNA synthetase | EPRS | S883 | 8.9 | 36.8 | K.EYIPGQPPLSQSS*DSSPTR.N |
| Glutamyl-prolyl-tRNA synthetase | EPRS | S882S885 | 13.6 | 31.4 | K.EYIPGQPPLSQS*SDS*SPTR.N |
| Glutamyl-prolyl-tRNA synthetase | EPRS | S882 | 14.7 | 33.4 | K.EYIPGQPPLSQS*SDSSPTR.N |
| Glutamyl-prolyl-tRNA synthetase | EPRS | S883S886 | 9.9 | 11.6 | K.EYIPGQPPLSQSS*DSS*PTR.N |
| Glutamyl-prolyl-tRNA synthetase | EPRS | S882S886 | 15.3 | 23.4 | K.EYIPGQPPLSQS*SDSS*PTR.N |
| Glutamyl-prolyl-tRNA synthetase | EPRS | S886 | 7.0 | 11.9 | K.EYIPGQPPLSQSSDSS*PTR.N |
| Glutamyl-prolyl-tRNA synthetase | EPRS | S880S885 | 7.3 | 14.0 | K.EYIPGQPPLS*QSSDS*SPTR.N |
| Glyceraldehyde 3 phosphate dehydrogenase | GAPDH | S210 | 8.9 | 26.3 | R.GALQNIIPAS*TGAAK.A |
| Glyceraldehyde 3 phosphate dehydrogenase | GAPDH | T75 | -4.2 | 20.9 | K.LVINGNPIT*IFQERDPSK.I |
| Glyceraldehyde 3 phosphate dehydrogenase | GAPDH | T182 | 17.0 | 66.3 | K.VIHDNFGIVEGLMTTVHAIT*ATQK.T |
| Glyceraldehyde 3 phosphate dehydrogenase | GAPDH | S83 | -3.3 | 27.0 | K.LVINGNPITIFQERDPS*K.I |
| Glycogen phosphorylase, brain type | PYGB | S15 | 100.0 | 31.9 | R.KQIS*VR.G |
| Glycogen synthase kinase 3 alpha | GSK3A | S21 | 22.2 | 104.0 | R.TSS*FAEPGGGGGGGGGPGGSASGPGGTGGGK.A |
| Glycogen synthase kinase 3 alpha | GSK3A | S20 | -0.4 | 94.4 | R.TS*SFAEPGGGGGGGGGPGGSASGPGGTGGGK.A |
| Glycogen synthase kinase 3 beta | GSK3B | Y216 | 16.9 | 43.5 | R.GEPNVSY*ICSR.Y |
| Glycogen synthase kinase 3 beta | GSK3B | S9 | 8.7 | 53.4 | R.TTS*FAESCKPVQQPSAFGSMK.V |
| Golgi autoantigen, golgin subfamily A, 4 | GOLGA4 | S41 | 11.7 | 93.5 | R.TSS*FTEQLDEGTPNR.E |
| Golgi autoantigen, golgin subfamily A, 4 | GOLGA4 | S71 | 55.8 | 57.2 | R.VPS*VESLFR.S |
| Golgi autoantigen, golgin subfamily A, 4 | GOLGA4 | S40 | -0.5 | 49.8 | R.TS*SFTEQLDEGTPNR.E |
| Golgi autoantigen, golgin subfamily A, 4 | GOLGA4 | S118S122 | 22.4 | 46.4 | R.LDLDSSTASFDPPS*DMDS*EAEDLVGNSDSLNK.E |
| Golgi reassembly stacking protein 2, 55Kda | GORASP2 | S449 | 6.6 | 15.6 | R.VGDSTPVSEKPVSAAVDANAS*ESP.- |
| Golgi reassembly stacking protein 2, 55Kda | GORASP2 | S451 | 17.5 | 37.4 | R.VGDSTPVSEKPVSAAVDANASES*P.- |
| Golgi specific brefeldin A resistance factor 1 | GBF1 | S1298 | 61.7 | 79.3 | R.ADAPDAGAQS*DSELPSYHQNDVSLDR.G |
| Golgi specific brefeldin A resistance factor 1 | GBF1 | T1337 | 19.9 | 98.2 | R.SAT*DADVVNSGWLVVGK.D |
| Golgi specific brefeldin A resistance factor 1 | GBF1 | T1317 |  | 20.9 | R.GYT*SDSEVYTDHGRPGK.I |
| Golgi specific brefeldin A resistance factor 1 | GBF1 | S1318 | 17.7 | 24.6 | R.GYTS*DSEVYTDHGRPGK.I |
| Golgi specific brefeldin A resistance factor 1 | GBF1 | S1320 | 8.3 | 19.9 | R.GYTSDS*EVYTDHGRPGK.I |
| Golgi specific brefeldin A resistance factor 1 | GBF1 | S1475 | 25.8 | 40.1 | R.GGQS*DDDEDEGVPASYHTVSLQVSQDLLDLMHTLHTR.A |
| Golgi specific brefeldin A resistance factor 1 | GBF1 | S349S352 | 14.4 | 41.8 | K.SQSAS*VES*IPEVLEECTSPADHSDSASVHDMDYVNPR.G |
| Golgi specific brefeldin A resistance factor 1 | GBF1 | S347S352 | 12.8 | 13.4 | K.SQS*ASVES*IPEVLEECTSPADHSDSASVHDMDYVNPR.G |
| Golgin 84 | GOLGA5 | S116 | 134.5 | 111.8 | K.S*EPDDELLFDFLNSSQK.E |
| GRAM domain containing 2 | GRAMD2A | S248 | 100.0 | 12.2 | R.KPPMS*EK.S |
| Grb2 | GRB2 | T159 |  | 15.4 | R.DIEQVPQQPT*YVQALFDFDPQEDGELGFR.R |
| GRB2 associated binding protein 3 | GAB3 | S173 | 8.5 | 22.5 | R.S*ESELLFLPDYLVLSNCETGR.L |
| GRB2 associated binding protein 3 | GAB3 | S175 | -0.3 | 12.3 | R.SES*ELLFLPDYLVLSNCETGR.L |
| GRB2 associated binding protein 3 | GAB3 | Y183 | 2.1 | 15.9 | R.SESELLFLPDY*LVLSNCETGR.L |
| GRB2 associated binding protein 3 | GAB3 | T191 | -3.2 | 13.7 | R.SESELLFLPDYLVLSNCET*GR.L |
| Grb4 | NCK2 | Y110 | 100.0 | 64.1 | R.IY*DLNIPAFVK.F |
| GRID | GRAP2 | Y222 | 100.0 | 26.0 | R.Y*LQHHHFHQER.R |
| GRID | GRAP2 | T262 | 100.0 | 54.5 | R.RHT*DPVQLQAAGR.V |
| GRID | GRAP2 | S164 | 59.7 | 47.8 | R.S*QGGPHLSGAVGEEIRPSMNR.K |
| GRID | GRAP2 | S187Y207 | 7.8 | 30.9 | R.KLS*DHPPTLPLQQHQHQPQPPQY*APAPQQLQQPPQQR.Y |
| GRID | GRAP2 | S187 | 15.9 | 44.4 | R.KLS*DHPPTLPLQQHQHQPQPPQYAPAPQQLQQPPQQR.Y |
| GRID | GRAP2 | S236 | 90.3 | 97.6 | R.GGS*LDINDGHCGTGLGSEMNAALMHR.R |
| GRID | GRAP2 | S159 | 100.0 | 27.7 | R.GNS*LDRR.S |
| GRID | GRAP2 | S181 | 19.9 | 26.7 | R.SQGGPHLSGAVGEEIRPS*MNR.K |

Peak Area: <-10, -5, -3, 0, 3, 5, >10

%CV: 0, 29, 42, 71, 86, >100

**White dots-** Significant change in peptide abundance at **5** %FDR compared to the timepoint with the minimum peak area for a given PSM

| Protein Name | Gene | Phosphosites | CarT 0m | CarT 2m | CarT 5m | RajiB 0m | RajiB 2m | RajiB 5m | Ascor | MOWSE | Sequence |
|---|---|---|---|---|---|---|---|---|---|---|---|
| **GRID** | GRAP2 | Y207 | | | | | | | **-0.3** | **18.5** | R.KLSDHPPTLPLQQHQHQPQPPQY*APAPQQLQQPPQQR.Y |
| **GRID** | GRAP2 | T192Y207 | | | | | | | **2.9** | **14.0** | R.KLSDHPPT*LPLQQHQHQPQPPQY*APAPQQLQQPPQQR.Y |
| **GRID** | GRAP2 | S187T192 | | | | | | | | **11.1** | R.KLS*DHPPT*LPLQQHQHQPQPPQYAPAPQQLQQPPQQR.Y |
| **GRINL1A downstream protein Gdown1** | GCOM1 | S179 | | | | | | | **15.0** | **110.2** | R.VSS*QAEDTSSSFDNLFIDR.L |
| **GRINL1A downstream protein Gdown1** | GCOM1 | S178 | | | | | | | **30.1** | **54.6** | R.VS*SQAEDTSSSFDNLFIDR.L |
| **GRINL1A downstream protein Gdown1** | GCOM1 | S364S365 | | | | | | | **100.0** | **32.1** | R.DEDDDWS*S*DEF.- |
| **GRIP1 associated protein 1** | GRIPAP1 | S691 | | | | | | | **17.7** | **56.5** | R.SLSS*SPQAQPPRPAELSDEEVAELFQR.L |
| **GRIP1 associated protein 1** | GRIPAP1 | S655 | | | | | | | **15.8** | **45.3** | R.SGLEELVLSEMNS*PSR.T |
| **GRIP1 associated protein 1** | GRIPAP1 | S688S691 | | | | | | | **17.4** | **55.8** | R.S*LSS*SPQAQPPRPAELSDEEVAELFQR.L |
| **GRIP1 associated protein 1** | GRIPAP1 | S690S692 | | | | | | | **13.6** | **56.5** | R.SLS*SS*PQAQPPRPAELSDEEVAELFQR.L |
| **GRIP1 associated protein 1** | GRIPAP1 | S692 | | | | | | | **5.8** | **53.5** | R.SLSSS*PQAQPPRPAELSDEEVAELFQR.L |
| **GRIP1 associated protein 1** | GRIPAP1 | S690S691 | | | | | | | **16.9** | **59.1** | R.SLS*S*SPQAQPPRPAELSDEEVAELFQR.L |
| **GRIP1 associated protein 1** | GRIPAP1 | S688S692 | | | | | | | **8.8** | **68.7** | R.S*LSSS*PQAQPPRPAELSDEEVAELFQR.L |
| **GRIP1 associated protein 1** | GRIPAP1 | S690S691S692 | | | | | | | **-0.4** | **28.0** | R.SLS*S*S*PQAQPPRPAELSDEEVAELFQR.L |
| **GRK-interacting protein 2** | GIT2 | Y392S397 | | | | | | | **26.7** | **110.6** | K.TINNQHSVESQDNDQPDY*DSVAS*DEDTDLETTASK.T |
| **GRK-interacting protein 2** | GIT2 | Y392S394 | | | | | | | **5.4** | **38.5** | K.TINNQHSVESQDNDQPDY*DS*VASDEDTDLETTASK.T |
| **GRK-interacting protein 2** | GIT2 | S418S421 | | | | | | | **7.4** | **41.7** | K.SLDS*DLS*DGPVTVQEFMEVK.N |
| **GRK-interacting protein 2** | GIT2 | S415S418 | | | | | | | **12.3** | **34.5** | K.S*LDS*DLSDGPVTVQEFMEVK.N |
| **GRK-interacting protein 2** | GIT2 | S415S421 | | | | | | | **5.8** | **41.2** | K.S*LDSDLS*DGPVTVQEFMEVK.N |
| **GRK-interacting protein 2** | GIT2 | S397 | | | | | | | **5.9** | **55.4** | K.TINNQHSVESQDNDQPDYDSVAS*DEDTDLETTASK.T |
| | GIT2 | S614 | | | | | | | **114.6** | **49.0** | R.S*MVWPGDGLVPDTAEPHVAPSPTLPSTEDVIR.K |
| **Growth arrest specific 2 like 1** | GAS2L1 | S316 | | | | | | | **4.2** | **15.0** | R.RGS*RPEMTPVSLR.S |
| **GTP binding protein 1** | GTPBP1 | S44S47 | | | | | | | **57.2** | **59.6** | R.LHGGFDS*DCS*EDGEALNGEPELDLTSK.L |
| **GTP binding protein 1** | GTPBP1 | S25 | | | | | | | **6.5** | **42.2** | R.SAMDSPVPASMFAPEPSS*PGAAR.A |
| **GTP binding protein 1** | GTPBP1 | S24 | | | | | | | **57.7** | **30.3** | R.SAMDSPVPASMFAPEPS*SPGAAR.A |
| **GTP binding protein 4** | GTPBP4 | S468S470S472 | | | | | | | **17.5** | **114.1** | R.TAAGEYDS*VS*ES*EDEEMLEIR.Q |
| **GTP binding protein 4** | GTPBP4 | Y466S470S472 | | | | | | | **5.1** | **81.2** | R.TAAGEY*DSVS*ES*EDEEMLEIR.Q |
| **GTP binding protein 4** | GTPBP4 | Y466S468S472 | | | | | | | **6.1** | **113.8** | R.TAAGEY*DS*VSES*EDEEMLEIR.Q |
| **GTP binding protein 4** | GTPBP4 | Y466S468S470 | | | | | | | **5.3** | **84.5** | R.TAAGEY*DS*VS*ESEDEEMLEIR.Q |
| **GTPase activating protein Ran 1** | RANGAP1 | S428S442 | | | | | | | **23.5** | **38.4** | K.ILDPNTGEPAPVLSS*PPPADVSTFLAFPS*PEK.L |
| **GTPase activating protein Ran 1** | RANGAP1 | S427S442 | | | | | | | **22.5** | **20.6** | K.ILDPNTGEPAPVLS*SPPPADVSTFLAFPS*PEK.L |
| **GTPase activating protein Ran 1** | RANGAP1 | S428 | | | | | | | **-0.4** | **18.2** | K.ILDPNTGEPAPVLSS*PPPADVSTFLAFPSPEK.L |
| **GTPase activating protein Ran 1** | RANGAP1 | S435S442 | | | | | | | **10.3** | **17.1** | K.ILDPNTGEPAPVLSSPPPADVS*TFLAFPS*PEK.L |
| **GTPase activating protein Ran 1** | RANGAP1 | T419S427 | | | | | | | | **31.4** | K.ILDPNT*GEPAPVLS*SPPPADVSTFLAFPSPEK.L |
| **GTPase activating Rap/RanGAP domain like 1** | RALGAPA | S860S861 | | | | | | | **33.1** | **12.4** | R.RGS*S*PGSLEIPK.D |
| **GTPase activating Rap/RanGAP domain like 1** | RALGAPA | S773 | | | | | | | **16.0** | **87.7** | R.HFS*QSEETGNEVFGALNEEQPLPR.S |
| **GTPase activating Rap/RanGAP domain like 1** | RALGAPA | S797 | | | | | | | **15.0** | **69.8** | R.SSS*TSDILEPFTVER.A |
| **GTPase activating Rap/RanGAP domain like 1** | RALGAPA | S831 | | | | | | | **23.9** | **46.0** | K.LPPLNSDIGGS*SANVPDLMDEFIAER.L |
| **GTPase activating Rap/RanGAP domain like 1** | RALGAPA | S775 | | | | | | | **3.5** | **42.2** | R.HFSQS*EETGNEVFGALNEEQPLPR.S |
| **GTPase activating Rap/RanGAP domain like 1** | RALGAPA | S1000T1002S1 | | | | | | | **12.3** | **31.6** | R.S*QT*PS*PSTLNIDHMEQK.D |
| **GTPase activating Rap/RanGAP domain like 1** | RALGAPA | T798 | | | | | | | **-0.4** | **26.2** | R.SSST*SDILEPFTVER.A |
| **GTPase activating Rap/RanGAP domain like 4** | RAP1GAP | S609S613 | | | | | | | **14.2** | **51.2** | K.SETSS*NPSS*PEICPNK.E |
| **GTPase activating Rap/RanGAP domain like 4** | RAP1GAP | T607S608S609 | | | | | | | **10.7** | **13.5** | K.SET*S*S*NPSSPEICPNK.E |
| | RAP1GAP | S45T49 | | | | | | | **34.7** | **24.4** | K.QELANSSDATLPDRPLS*PPLT*APPTMK.S |
| **GTPase activating Rap/RanGAP domain like 4** | RAP1GAP | S9 | | | | | | | **33.6** | **75.1** | R.SVS*FGGFGWIDK.T |
| **GTPase activating Rap/RanGAP domain like 4** | RAP1GAP | S680S685 | | | | | | | **9.1** | **80.7** | K.QEVFVYSPS*PSSES*PSLGAAATPIIMSR.S |
| **GTPase activating Rap/RanGAP domain like 4** | RAP1GAP | S678S685 | | | | | | | **14.5** | **35.2** | K.QEVFVYS*PSPSSES*PSLGAAATPIIMSR.S |
| **GTPase activating Rap/RanGAP domain like 4** | RAP1GAP | Y677S682 | | | | | | | **8.5** | **22.3** | K.QEVFVY*SPSPS*SESPSLGAAATPIIM#SR.S |
| **GTPase activating Rap/RanGAP domain like 4** | RAP1GAP | S680 | | | | | | | **5.6** | **76.0** | K.QEVFVYSPS*PSSESPSLGAAATPIIMSR.S |
| **GTPase activating Rap/RanGAP domain like 4** | RAP1GAP | S678S680 | | | | | | | **4.5** | **18.5** | K.QEVFVYS*PS*PSSESPSLGAAATPIIMSR.S |
| **GTPase activating Rap/RanGAP domain like 4** | RAP1GAP | S34T49 | | | | | | | **4.0** | **39.2** | K.QELANS*SDATLPDRPLSPPLT*APPTMK.S |
| **GTPase activating Rap/RanGAP domain like 4** | RAP1GAP | S682 | | | | | | | **2.8** | **80.8** | K.QEVFVYSPSPS*SESPSLGAAATPIIMSR.S |
| **GTPase activating Rap/RanGAP domain like 4** | RAP1GAP | S680S682 | | | | | | | **2.0** | **94.5** | K.QEVFVYSPS*PS*SESPSLGAAATPIIMSR.S |
| **GTPase activating Rap/RanGAP domain like 4** | RAP1GAP | S678S683T693 | | | | | | | **12.0** | **26.8** | K.QEVFVYS*PSPSS*ESPSLGAAAT*PIIMSR.S |
| **GTPase activating Rap/RanGAP domain like 4** | RAP1GAP | S609S612S613 | | | | | | | **5.1** | **28.5** | K.SETSS*NPS*S*PEICPNK.E |
| **GTPase activating Rap/RanGAP domain like 4** | RAP1GAP | S678T693 | | | | | | | **23.4** | **54.7** | K.QEVFVYS*PSPSSESPSLGAAAT*PIIMSR.S |
| **GTPase activating Rap/RanGAP domain like 4** | RAP1GAP | S680S682T693 | | | | | | | **9.2** | **22.7** | K.QEVFVYSPS*PS*SESPSLGAAAT*PIIMSR.S |
| **GTPase activating Rap/RanGAP domain like 4** | RAP1GAP | S608S612S613 | | | | | | | **18.1** | **22.3** | K.SETS*SNPS*S*PEICPNK.E |
| **GTPase activating Rap/RanGAP domain like 4** | RAP1GAP | S34S35 | | | | | | | **5.3** | **17.2** | K.QELANS*S*DATLPDRPLSPPLTAPPTMK.S |
| **GTPase activating Rap/RanGAP domain like 4** | RAP1GAP | S678S682 | | | | | | | **7.7** | **24.8** | K.QEVFVYS*PSPS*SESPSLGAAATPIIM#SR.S |
| **GTPase activating Rap/RanGAP domain like 4** | RAP1GAP | S678S683 | | | | | | | **5.0** | **50.2** | K.QEVFVYS*PSPSS*ESPSLGAAATPIIMSR.S |
| **GTPase activating Rap/RanGAP domain like 4** | RAP1GAP | S608S613 | | | | | | | **5.8** | **26.8** | K.SETS*SNPSS*PEICPNK.E |

Peak Area: <-10, -5, -3, 0, 3, 5, >10

%CV: 0, 29, 42, 71, 86, >100

**White dots-** Significant change in peptide abundance at **5** %FDR compared to the timepoint with the minimum peak area for a given PSM

CarT (0m, 2m, 5m) · RajiB (0m, 2m, 5m)

| Protein Name | Gene | Phosphosites | Ascor | MOWSE | Sequence |
|---|---|---|---|---|---|
| GTPase activating Rap/RanGAP domain like 4 | RAP1GAP | S683S685 | -3.2 | 29.9 | K.QEVFVYSPSPSS*ES*PSLGAAATPIIMSR.S |
| GTPase activating Rap/RanGAP domain like 4 | RAP1GAP | S683T693 | 7.6 | 20.3 | K.QEVFVYSPSPSS*ESPSLGAAAT*PIIMSR.S |
| Guanine nucleotide binding protein G(S), alpha subunit | GNAS | S572 | 13.3 | 11.6 | R.ALS*RSR.R |
| Guanine nucleotide binding protein like 2 | GNL2 | S580S581S582 | 108.2 | 61.4 | R.DDAEES*S*S*EPEEENVGNDTK.A |
| Guanine nucleotide binding protein like 2 | GNL2 | S580S581 | 82.9 | 45.6 | R.DDAEES*S*SEPEEENVGNDTK.A |
| Guanine nucleotide binding protein like1 | GNL1 | T48T50 | 16.1 | 71.5 | R.EEQT*DT*SDGESVTHHIR.R |
| Guanine nucleotide binding protein like1 | GNL1 | S51S55 | 8.9 | 22.4 | R.EEQTDTS*DGES*VTHHIR.R |
| | GNL1 | T48T50S51 | 59.6 | 65.5 | R.EEQT*DT*S*DGESVTHHIR.R |
| Guanine nucleotide binding protein like1 | GNL1 | S32S33S34S36 | 100.0 | 16.7 | R.GLQDGLRS*S*S*NS*R.S |
| Guanine nucleotide binding protein like1 | GNL1 | T50S51 | 12.7 | 17.1 | R.EEQTDT*S*DGESVTHHIR.R |
| Guanine nucleotide binding protein like1 | GNL1 | T48S51 | 6.0 | 31.0 | R.EEQT*DTS*DGESVTHHIR.R |
| Guanine nucleotide binding protein like1 | GNL1 | S561S562S563 | 100.0 | 71.5 | R.VGPAGDEEEEEEEELS*S*S*CEEEGEEDR.D |
| Guanine nucleotide binding protein like1 | GNL1 | T50 | 7.9 | 51.6 | R.EEQTDT*SDGESVTHHIR.R |
| Guanine nucleotide binding protein like1 | GNL1 | S51 | 7.8 | 26.1 | R.EEQTDTS*DGESVTHHIR.R |
| Guanine nucleotide releasing factor 2 | RAPGEF1 | S378 | 60.6 | 62.9 | K.S*DEQLSSLDR.D |
| Guanine nucleotide releasing factor 2 | RAPGEF1 | S329 | 13.9 | 54.9 | R.ATS*GSSLPVGINR.Q |
| Guanine nucleotide releasing factor 2 | RAPGEF1 | T328 | 20.3 | 26.0 | R.AT*SGSSLPVGINR.Q |
| H1 histone family, member T | HIST1H1T | T159T160 | 24.1 | 11.2 | R.AT*T*PKTVR.S |
| H2AX histone | H2AFX | S140 | 22.8 | 30.9 | K.ATQAS*QEY.- |
| | H2AFX | T137S140 | 4.9 | 12.5 | K.AT*QAS*QEY.- |
| H2B Histone family, member E | HIST1H2B | S37 | 21.2 | 33.3 | R.KES*YSVYVYK.V |
| H3 Histone, family 3B | H3F3A | S58 | 17.0 | 38.0 | K.S*TELLIR.K |
| H4 Histone family, member A | HIST1H4A | S48 | 74.6 | 46.6 | K.RIS*GLIYEETR.G |
| HBS1L | HBS1L | Y22 | -0.4 | 38.1 | R.GYNYDEDFEDDDLY*GQSVEDDYCISPSTAAQFIYSR.R |
| HBS1L | HBS1L | S33 | 18.2 | 76.4 | R.GYNYDEDFEDDDLYGQSVEDDYCIS*PSTAAQFIYSR.R |
| HBS1L | HBS1L | S25 | 2.3 | 60.5 | R.GYNYDEDFEDDDLYGQS*VEDDYCISPSTAAQFIYSR.R |
| HBXAP | RSF1 | S604 | 100.0 | 38.9 | R.LS*PIPEEVPK.S |
| HBXAP | RSF1 | S1345 | 100.0 | 89.0 | R.IES*DEEEDFENVGK.V |
| HBXAP | RSF1 | S1359S1375 | 32.2 | 40.3 | K.VGS*PLDYSLVDLPSTNGQS*PGK.A |
| HBXAP | RSF1 | S1277T1278S1 | 9.5 | 38.0 | R.GRS*T*DEYS*EADEEEEEEEGKPSR.K |
| HBXAP | RSF1 | S1370 | 8.2 | 18.1 | K.VGSPLDYSLVDLPS*TNGQSPGK.A |
| HBXAP | RSF1 | S622T628 | 25.9 | 38.4 | K.STLESEKPGS*PEAAET*SPPSNIIDHCEK.L |
| HBXAP | RSF1 | S622S629 | 5.0 | 25.7 | K.STLESEKPGS*PEAAETS*PPSNIIDHCEK.L |
| HDCMA18P protein | LARP7 | S258S261 | 9.8 | 39.3 | R.SRPTS*EGS*DIESTEPQK.Q |
| HDCMA18P protein | LARP7 | S337T338 | 69.1 | 57.2 | R.DIEIS*T*EEEKDTGDLKDSSLLK.T |
| HDCMA18P protein | LARP7 | S261 | 44.8 | 70.7 | R.SRPTSEGS*DIESTEPQK.Q |
| HDCMA18P protein | LARP7 | T257S258 | 10.7 | 16.7 | R.SRPT*S*EGSDIESTEPQK.Q |
| HDCMA18P protein | LARP7 | S300 | -0.4 | 36.4 | R.SSS*EDAESLAPR.S |
| Headcase protein homolog | HECA | S264S268 | 38.0 | 66.6 | R.S*PGGS*PGQSPPTGYSILSPAHFSGPR.S |
| Headcase protein homolog | HECA | S264S268S272 | 51.5 | 73.2 | R.S*PGGS*PGQS*PPTGYSILSPAHFSGPR.S |
| Headcase protein homolog | HECA | S272 | 16.8 | 17.8 | R.SPGGSPGQS*PPTGYSILSPAHFSGPR.S |
| Headcase protein homolog | HECA | S268S272 | 12.2 | 69.7 | R.SPGGS*PGQS*PPTGYSILSPAHFSGPR.S |
| Heat shock 40 kDa protein 3 | DNAJB2 | S247S250 | | 22.7 | R.SGGTQVQQTPASCPLDS*DLS*EDEDLQLAMAYSLSEMEAAGK.K |
| Heat shock 40 kDa protein 4 | DNAJA1 | S335 | 18.4 | 31.7 | K.VNFPENGFLS*PDKLSLLEK.L |
| Heat shock 60 KD protein 1 (chaperonin) | HSPD1 | S70 | 55.8 | 70.3 | R.TVIIEQSWGS*PK.V |
| Heat shock 60 KD protein 1 (chaperonin) | HSPD1 | S67 | 5.0 | 30.4 | R.TVIIEQS*WGSPK.V |
| Heat shock 70 kDa protein 8 | HSPA8 | S362 | 10.1 | 73.3 | K.S*INPDEAVAYGAAVQAAILSGDK.S |
| Heat shock 70kD protein 12B | HSPA12B | S291S294 | 100.0 | 15.0 | R.RS*RHS*R.T |
| HECT domain and ankyrin repeat containing, E3 ubiquitin protein ligase 1 | HACE1 | S385 | 62.4 | 54.0 | K.NKRDS*TEITSILLK.Q |
| Hect domain and RCC1 like domain 1 | HERC1 | S4857 | 20.6 | 13.3 | R.NVDNAEGS*DTDY.- |
| Helicase with zinc finger domain | HELZ | S1614 | 73.8 | 48.6 | R.S*PPAVPSPPSSTDHSSHFSNFNDNSR.D |
| Hematological and neurological expressed 1 | JPT1 | S88 | 16.9 | 36.0 | R.RNSS*EASSGDFLDLK.G |
| Hematological and neurological expressed 1 | JPT1 | S87 | 27.6 | 106.7 | R.NS*SEASSGDFLDLK.G |
| Hematological and neurological expressed 1 | JPT1 | S92 | 10.9 | 14.4 | R.RNSSEAS*GDFLDLK.G |
| Hematopoietic cell specific LYN substrate 1 | HCLS1 | S275 | 100.0 | 53.3 | R.S*PEAPQPVIAM#EEPAVPAPLPK.K |
| Hepatocyte cell adhesion molecule | HEPACAM | T374S376 | 7.2 | 15.0 | R.T*HS*SPPR.A |
| Hepatocyte cell adhesion molecule | HEPACAM | S376S377 | 11.1 | 16.4 | R.THS*S*PPR.A |
| Hepatoma derived growth factor | HDGF | S165 | 100.0 | 38.3 | R.RAGDLLEDS*PK.R |
| Hepatoma derived growth factor | HDGF | S132 | 9.0 | 21.4 | K.GNAEGS*SDEEGKLVIDEPAK.E |
| Hepatoma derived growth factor | HDGF | S132S133 | 100.0 | 132.7 | K.GNAEGS*S*DEEGKLVIDEPAK.E |
| Hepatoma derived growth factor | HDGF | S107S132 | | 15.1 | K.S*CVEEPEPEPEAAEGDGDKKGNAEGS*SDEEGK.L |
| Hepatoma-derived growth factor-related protein 2 | HDGFL2 | S232S234S236 | 7.1 | 24.3 | K.KAPSAS*DS*DS*KADS*DGAKPEPVAMAR.S |

Peak Area: <-10, -5, -3, 0, 3, 5, >10

%CV: 0, 29, 42, 71, 86, >100

**White dots-** Significant change in peptide abundance at 5 %FDR compared to the timepoint with the minimum peak area for a given PSM

| Protein Name | Gene | Phosphosites | CarT 0m | CarT 2m | CarT 5m | RajiB 0m | RajiB 2m | RajiB 5m | Ascor | MOWSE | Sequence |
|---|---|---|---|---|---|---|---|---|---|---|---|
| Hepatoma-derived growth factor-related protein 2 | HDGFL2 | S454 | | | | | | | 42.5 | 50.4 | R.S*EGFSMDR.K |
| Hepatoma-derived growth factor-related protein 2 | HDGFL2 | S137 | | | | | | | 9.8 | 80.4 | R.GVMAVTAVTATAAS*DR.M |
| Hepatoma-derived growth factor-related protein 2 | HDGFL2 | S652 | | | | | | | 100.0 | 25.8 | R.EGPDLDRPGS*DR.Q |
| Hepatoma-derived growth factor-related protein 2 | HDGFL2 | S366S369S370 | | | | | | | 100.0 | 20.4 | R.GEAERGS*GGS*S*GDELREDDEPVK.K |
| Hepatoma-derived growth factor-related protein 2 | HDGFL2 | S230S232S234 | | | | | | | 4.5 | 19.2 | K.KAPS*AS*DS*DSKADS*DGAKPEPVAMAR.S |
| Hepatoma-derived growth factor-related protein 2 | HDGFL2 | S671 | | | | | | | 8.9 | 16.8 | R.GDSEALDEES*.- |
| Hepatoma-derived growth factor-related protein 2 | HDGFL2 | S664 | | | | | | | 5.4 | 10.9 | R.GDS*EALDEES.- |
| Hepatoma-derived growth factor-related protein 2 | HDGFL2 | S395S396S397 | | | | | | | 100.0 | 21.7 | R.GRGPPS*S*S*DS*EPEAELER.E |
| Hepatoma-derived growth factor-related protein 2 | HDGFL2 | T132 | | | | | | | 18.8 | 14.0 | R.GVM#AVTAVT*ATAASDR.M |
| Hepatoma-derived growth factor-related protein 2 | HDGFL2 | T134 | | | | | | | 9.0 | 48.4 | R.GVMAVTAVTAT*AASDR.M |
| Hepatoma-derived growth factor-related protein 2 | HDGFL2 | S230S232S234 | | | | | | | -2.6 | 24.2 | K.KAPS*AS*DS*DS*KADSDGAKPEPVAMAR.S |
| Heterochromatin protein 1 alpha | CBX5 | S11S12S13S14 | | | | | | | 44.4 | 55.2 | K.RTADS*S*S*S*EDEEEYVVEK.V |
| Heterochromatin protein 1 alpha | CBX5 | T8S11S13S14 | | | | | | | 6.7 | 14.4 | R.T*ADS*SS*S*EDEEEYVVEK.V |
| Heterogeneous nuclear ribonucleoprotein A2 | HNRNPA2 | S259 | | | | | | | 40.6 | 129.0 | R.GFGDGYNGYGGGPGGGNFGGS*PGYGGGR.G |
| Heterogeneous nuclear ribonucleoprotein A2 | HNRNPA2 | S341 | | | | | | | 15.8 | 119.1 | R.NMGGPYGGGNYGPGGS*GGSGGYGGR.S |
| Heterogeneous nuclear ribonucleoprotein A2 | HNRNPA2 | Y331 | | | | | | | 13.3 | 37.0 | R.NMGGPY*GGGNYGPGGSGGSGGYGGR.S |
| Heterogeneous nuclear ribonucleoprotein A3 | HNRNPA3 | S358 | | | | | | | 25.4 | 158.8 | R.SSGS*PYGGGYGSGGGSGGYGSR.R |
| Heterogeneous nuclear ribonucleoprotein A3 | HNRNPA3 | S356 | | | | | | | -0.3 | 128.8 | R.SS*GSPYGGGYGSGGGSGGYGSR.R |
| Heterogeneous nuclear ribonucleoprotein A3 | HNRNPA3 | Y360 | | | | | | | -0.5 | 13.6 | R.SSGSPY*GGGYGSGGGSGGYGSR.R |
| Heterogeneous nuclear ribonucleoprotein C | HNRNPC | S253 | | | | | | | | 68.8 | K.MES*EGGADDSAEEGDLLDDDDNEDRGDDQLELIKDDEK.E |
| | HNRNPC | S253S260 | | | | | | | 100.0 | 81.6 | K.MES*EGGADDS*AEEGDLLDDDDNEDRGDDQLELIKDDEK.E |
| | HNRNPC | S260 | | | | | | | 98.8 | 107.2 | K.M#ESEGGADDS*AEEGDLLDDDDNEDRGDDQLELIK.D |
| Heterogeneous nuclear ribonucleoprotein C | HNRNPC | S299 | | | | | | | 58.5 | 71.3 | K.EAEEGEDDRDS*ANGEDDS.- |
| Heterogeneous nuclear ribonucleoprotein C | HNRNPC | S306 | | | | | | | 3.2 | 15.9 | K.DDEKEAEEGEDDRDSANGEDDS*.- |
| Heterogeneous nuclear ribonucleoprotein C | HNRNPC | S240S241 | | | | | | | 4.5 | 29.6 | K.SEEEQSSS*S*VKKDETNVK.M |
| Heterogeneous nuclear ribonucleoprotein C | HNRNPC | S233 | | | | | | | 56.4 | 65.6 | K.NDKS*EEEQSSSSVKK.D |
| Heterogeneous nuclear ribonucleoprotein C | HNRNPC | S238S240 | | | | | | | | 13.2 | K.SEEEQS*SS*SVKKDETNVK.M |
| Heterogeneous nuclear ribonucleoprotein C | HNRNPC | S238S241 | | | | | | | 7.0 | 38.0 | K.SEEEQS*SSS*VKKDETNVK.M |
| Heterogeneous nuclear ribonucleoprotein C | HNRNPC | S107 | | | | | | | 10.9 | 77.0 | R.SAAEMYGS*SFDLDYDFQR.D |
| Heterogeneous nuclear ribonucleoprotein C | HNRNPC | S238S239 | | | | | | | 18.1 | 19.5 | K.SEEEQS*S*SSVKKDETNVK.M |
| Heterogeneous nuclear ribonucleoprotein D | HNRNPD | S80S83 | | | | | | | 12.8 | 45.6 | K.NEEDEGHS*NSS*PR.H |
| | HNRNPD | S83 | | | | | | | 23.9 | 89.0 | K.NEEDEGHSNSS*PR.H |
| Heterogeneous nuclear ribonucleoprotein D | HNRNPD | S82 | | | | | | | 19.8 | 58.6 | K.NEEDEGHSNS*SPR.H |
| | HNRNPD | S80 | | | | | | | 62.3 | 66.4 | K.IDASKNEEDEGHS*NSSPR.H |
| Heterogeneous nuclear ribonucleoprotein D | HNRNPD | S80S82 | | | | | | | 6.0 | 41.2 | K.NEEDEGHS*NS*SPR.H |
| Heterogeneous nuclear ribonucleoprotein H1 | HNRNPH1 | S104 | | | | | | | 44.4 | 85.2 | K.HTGPNS*PDTANDGFVR.L |
| Heterogeneous nuclear ribonucleoprotein H1 | HNRNPH1 | T100 | | | | | | | 9.2 | 31.0 | K.HT*GPNSPDTANDGFVR.L |
| Heterogeneous nuclear ribonucleoprotein H1 | HNRNPH1 | S310 | | | | | | | 42.3 | 48.8 | K.ATENDIYNFFS*PLNPVR.V |
| Heterogeneous nuclear ribonucleoprotein H1 | HNRNPH1 | T107 | | | | | | | 12.2 | 18.0 | K.HTGPNSPDT*ANDGFVR.L |
| Heterogeneous nuclear ribonucleoprotein K | HNRNPK | S379 | | | | | | | 99.9 | 115.7 | R.GS*YGDLGGPIITTQVTIPK.D |
| Heterogeneous nuclear ribonucleoprotein K | HNRNPK | Y380 | | | | | | | -0.1 | 70.1 | R.GSY*GDLGGPIITTQVTIPK.D |
| Heterogeneous nuclear ribonucleoprotein K | HNRNPK | T389 | | | | | | | 4.3 | 55.0 | R.GSYGDLGGPIIT*TQVTIPK.D |
| Heterogeneous nuclear ribonucleoprotein K | HNRNPK | S216 | | | | | | | 26.0 | 64.4 | K.IILDLISES*PIK.G |
| Heterogeneous nuclear ribonucleoprotein K | HNRNPK | T118 | | | | | | | 3.8 | 25.7 | K.IIPTLEEGLQLPSPT*ATSQLPLESDAVECLNYQHYK.G |
| Heterogeneous nuclear ribonucleoprotein K | HNRNPK | T107 | | | | | | | | 55.8 | K.IIPT*LEEGLQLPSPTATSQLPLESDAVECLNYQHYK.G |
| Heterogeneous nuclear ribonucleoprotein K | HNRNPK | S116 | | | | | | | 28.4 | 62.6 | K.IIPTLEEGLQLPS*PTATSQLPLESDAVECLNYQHYK.G |
| Heterogeneous nuclear ribonucleoprotein K | HNRNPK | S284 | | | | | | | 32.7 | 37.5 | R.RDYDDMS*PR.R |
| Heterogeneous nuclear ribonucleoprotein K | HNRNPK | T120 | | | | | | | 5.0 | 16.3 | K.IIPTLEEGLQLPSPTAT*SQLPLESDAVECLNYQHYK.G |
| Heterogeneous nuclear ribonucleoprotein K | HNRNPK | S214 | | | | | | | | 21.6 | K.IILDLIS*ESPIKGR.A |
| Heterogeneous nuclear ribonucleoprotein M | HNRNPM | S633 | | | | | | | 52.3 | 57.8 | R.GNFGGS*FAGSFGGAGGHAPGVAR.K |
| Heterogeneous nuclear ribonucleoprotein U | HNRNPU | S59 | | | | | | | 54.1 | 55.4 | R.LQAALDDEEAGGRPAMEPGNGS*LDLGGDSAGR.S |
| | HNRNPU | S271 | | | | | | | 7.7 | 41.5 | K.S*PQPPVEEEDEHFDDTVVCLDTYNCDLHFK.I |
| Heterogeneous nuclear ribonucleoprotein U | HNRNPU | T286 | | | | | | | 0.8 | 15.2 | K.SPQPPVEEEDEHFDDT*VVCLDTYNCDLHFK.I |
| Heterogeneous nuclear ribonucleoprotein U like 1 | HNRNPUL | Y717 | | | | | | | 5.3 | 21.8 | R.APQQQPPPQQPPPPQPPPQQPPPPPSY*SPAR.N |
| Heterogeneous nuclear ribonucleoprotein U like 1 | HNRNPUL | S716 | | | | | | | | 26.1 | R.APQQQPPPQQPPPPQPPPQQPPPPPS*YSPAR.N |
| Heterogeneous nuclear ribonucleoprotein U like 1 | HNRNPUL | S718 | | | | | | | 5.8 | 19.5 | R.APQQQPPPQQPPPPQPPPQQPPPPPSYS*PAR.N |
| Heterogeneous nuclear ribonucleoprotein U like 1 | HNRNPUL | S194 | | | | | | | 11.9 | 21.3 | R.S*PQPPAEEDEDDFDDTLVAIDTYNCDLHFK.V |
| Heterogeneous nuclear ribonucleoprotein U-like 2 | HNRNPUL | S161T165 | | | | | | | 49.3 | 75.1 | R.S*GDET*PGSEVPGDK.A |
| Heterogeneous nuclear ribonucleoprotein U-like 2 | HNRNPUL | S161T165S168 | | | | | | | 100.0 | 73.1 | R.S*GDET*PGS*EVPGDK.A |
| Heterogeneous nuclear ribonucleoprotein U-like 2 | HNRNPUL | S193 | | | | | | | 10.4 | 27.7 | K.SKPAGS*DGER.R |
| Heterogeneous nuclear ribonucleoprotein U-like 2 | HNRNPUL | S185 | | | | | | | 100.0 | 58.9 | K.AAEEQGDDQDS*EK.S |
| Heterogeneous nuclear ribonucleoprotein U-like 2 | HNRNPUL | S228 | | | | | | | 39.8 | 52.7 | R.SKS*PLPPEEEAK.D |

**Peak Area**: <-10, -5, -3, 0, 3, 5, >10

**%CV**: 0, 29, 42, 71, 86, >100

**White dots-** Significant change in peptide abundance at **5**%FDR compared to the timepoint with the minimum peak area for a given PSM

| Protein Name | Gene | Phosphosites | CarT 0m | CarT 2m | CarT 5m | RajiB 0m | RajiB 2m | RajiB 5m | Ascor | MOWSE | Sequence |
|---|---|---|---|---|---|---|---|---|---|---|---|
| **Heterogeneous nuclear ribonucleoprotein U-like 2** | HNRNPUL | T165 | | | | | | | **37.7** | **87.1** | R.SGDET*PGSEVPGDK.A |
| | HNRNPUL | S161 | | | | | | | **11.8** | **21.3** | R.S*GDETPGSEVPGDKAAEEQGDDQDSEK.S |
| **Heterogeneous nuclear ribonucleoprotein U-like 2** | HNRNPUL | S226 | | | | | | | **9.2** | **40.6** | R.S*KSPLPPEEEAK.D |
| **Heterogeneous nuclear ribonucleoprotein U-like 2** | HNRNPUL | T165S168 | | | | | | | **34.8** | **57.2** | R.SGDET*PGS*EVPGDK.A |
| **Heterogeneous nuclear ribonucleoproteins C1/C2** | HNRNPC | S247 | | | | | | | **40.8** | **99.3** | K.MESEGGADDS*AEEGDLLDDDDNEDRGDDQVK.T |
| **Heterogeneous nuclear ribonucleoproteins C1/C2** | HNRNPC | S240S247 | | | | | | | **100.0** | **78.7** | K.MES*EGGADDS*AEEGDLLDDDDNEDRGDDQVK.T |
| **Heterogenous nuclear ribonucleoprotein A1** | HNRNPA1 | S4S6 | | | | | | | **100.0** | **29.7** | K.S*ES*PKEPEQLRK.L |
| **Heterogenous nuclear ribonucleoprotein A1** | HNRNPA1 | S6 | | | | | | | **12.0** | **48.9** | K.SES*PKEPEQLR.K |
| **Heterogenous nuclear ribonucleoprotein A1** | HNRNPA1 | S338 | | | | | | | **-0.3** | **91.6** | R.SS*GPYGGGGQYFAKPR.N |
| **Heterogenous nuclear ribonucleoprotein A1** | HNRNPA1 | S4 | | | | | | | **8.5** | **35.0** | K.S*ESPKEPEQLR.K |
| **Hexamthylene bis-acetamide inducible 2** | HEXIM2 | S51S53 | | | | | | | **100.7** | **45.7** | R.MES*HS*EDEDLAGAVGGLGWNSR.S |
| **HGAL** | GCSAM | S102 | | | | | | | **16.3** | **36.4** | R.VLCTRPSGNS*AEEYYENVPCK.A |
| **HGAL** | GCSAM | S120 | | | | | | | **20.3** | **34.3** | R.ES*LGGTETEYSLLHMPSTDPR.H |
| **HGAL** | GCSAM | S69 | | | | | | | **7.2** | **56.1** | R.MSS*TPIQDNVDQTYSEELCYTLINHR.V |
| **HGAL** | GCSAM | S68 | | | | | | | **9.2** | **86.1** | R.MS*STPIQDNVDQTYSEELCYTLINHR.V |
| **HGAL** | GCSAM | T70 | | | | | | | **5.3** | **22.1** | R.MSST*PIQDNVDQTYSEELCYTLINHR.V |
| **High mobility group 20A** | HMG20A | S105 | | | | | | | **48.9** | **46.0** | K.S*PLTGYVR.F |
| **High mobility group AT hook 1** | HMGA1 | S102 | | | | | | | **42.5** | **68.9** | K.EEEEGISQES*SEEEQ.- |
| | HMGA1 | S102S103 | | | | | | | **20.6** | **82.5** | K.EEEEGISQES*S*EEEQ.- |
| **High mobility group AT hook 1** | HMGA1 | S9 | | | | | | | **-0.2** | **54.8** | K.SS*QPLASK.Q |
| **High mobility group AT hook 1** | HMGA1 | S99S102 | | | | | | | **12.6** | **41.8** | K.EEEEGIS*QES*SEEEQ.- |
| | HMGA1 | S44 | | | | | | | **51.6** | **70.1** | R.KQPPVSPGTALVGS*QK.E |
| **High mobility group AT hook 1** | HMGA1 | S36 | | | | | | | **12.5** | **18.4** | R.KQPPVS*PGTALVGSQK.E |
| **High mobility group AT hook 1** | HMGA1 | T39 | | | | | | | **1.5** | **21.0** | R.KQPPVSPGT*ALVGSQKEPSEVPTPK.R |
| **High mobility group AT hook 1** | HMGA1 | S99 | | | | | | | | **13.2** | K.EEEEGIS*QESSEEEQ.- |
| **High mobility group AT hook 1** | HMGA1 | S99S102S103 | | | | | | | **100.0** | **54.5** | K.KLEKEEEEGIS*QES*S*EEEQ.- |
| **High mobility group AT hook 1** | HMGA1 | S103 | | | | | | | **7.0** | **14.4** | K.EEEEGISQESS*EEEQ.- |
| **High mobility group AT hook 1 like 4** | INO80B | S84T87 | | | | | | | | **19.7** | K.S*VPT*FTVIPEGPRSPSPLMVVDNEEEPMEGVPLEQYR.A |
| **High mobility group AT hook 1 like 4** | INO80B | S97S99 | | | | | | | **4.2** | **17.5** | K.SVPTFTVIPEGPRS*PS*PLMVVDNEEEPMEGVPLEQYR.A |
| **High mobility group protein 2 like 1** | HMGXB4 | S393S403 | | | | | | | **25.8** | **13.3** | K.S*PPTTMLLPAS*PAKAPETEPIDVAAHLQLLGESLSLIGHR.L |
| **High mobility group protein 2 like 1** | HMGXB4 | T396S403 | | | | | | | **11.5** | **18.8** | K.SPPT*TMLLPAS*PAKAPETEPIDVAAHLQLLGESLSLIGHR.L |
| **HIP55** | DBNL | S275 | | | | | | | **26.0** | **34.8** | R.AMSTTSIS*SPQPGK.L |
| **HIP55** | DBNL | S284 | | | | | | | **100.0** | **17.3** | K.LRS*PFLQK.Q |
| **HIRA interacting protein 3** | HIRIP3 | S289S291 | | | | | | | **100.0** | **66.8** | R.LLGDS*DS*EEEQK.E |
| **HIRA interacting protein 3** | HIRIP3 | T358S359S363 | | | | | | | **34.3** | **66.6** | R.LGST*S*GEES*DLER.E |
| **HIRA interacting protein 3** | HIRIP3 | S223S227 | | | | | | | **33.0** | **59.8** | K.SLKES*EQES*EEEILAQK.K |
| **HIRA interacting protein 3** | HIRIP3 | S530 | | | | | | | **28.3** | **23.5** | R.TLDS*DEERPRPAPPDWSHMR.G |
| **HIRA interacting protein 3** | HIRIP3 | S550S551S555 | | | | | | | **100.0** | **41.8** | R.GIIS*S*DGES*N.- |
| **HIRA interacting protein 3** | HIRIP3 | S357T358S359 | | | | | | | **100.0** | **65.2** | R.LGS*T*S*GEES*DLER.E |
| | HIRIP3 | S142S143 | | | | | | | **100.0** | **33.1** | K.AVEES*S*DEER.Q |
| **HIRA interacting protein 3** | HIRIP3 | S370 | | | | | | | **32.2** | **77.2** | R.EVS*DSEAGGGPQGER.K |
| **HIRA interacting protein 3** | HIRIP3 | S370S372 | | | | | | | **100.0** | **68.9** | R.EVS*DS*EAGGGPQGER.K |
| **HIRA interacting protein 3** | HIRIP3 | S330S332S333 | | | | | | | **16.7** | **25.7** | K.RLS*GS*S*EDEEDS*GKGEPTAK.G |
| **HIRA interacting protein 3** | HIRIP3 | S196S199 | | | | | | | **100.0** | **84.6** | R.EES*EES*EAEPVQR.T |
| **HIRA interacting protein 3** | HIRIP3 | T358S363 | | | | | | | **15.7** | **25.9** | R.LGST*SGEES*DLER.E |
| **HIRA interacting protein 3** | HIRIP3 | S551S555 | | | | | | | **8.6** | **42.3** | R.GIIS S*DGES*N.- |
| | HIRIP3 | S159S160 | | | | | | | **40.9** | **24.9** | R.GEES*S*EEEEKGYK.G |
| **HIRA interacting protein 3** | HIRIP3 | S289S291S300 | | | | | | | **100.0** | **16.9** | R.LLGDS*DS*EEEQKEAAS*S*GDDS*GRDR.E |
| | HIRIP3 | S300S301S305 | | | | | | | **100.0** | **23.4** | K.EAAS*S*GDDS*GRDR.E |
| **HIRA interacting protein 3** | HIRIP3 | T84S87 | | | | | | | **100.0** | **19.1** | K.RPPT*PCS*DPER.K |
| **HIRA interacting protein 3** | HIRIP3 | S330S332S333 | | | | | | | **47.9** | **34.2** | K.RLS*GS*S*EDEEDSGKGEPTAK.G |
| **HIRA interacting protein 3** | HIRIP3 | S550S551 | | | | | | | **4.6** | **41.9** | R.GIIS*S*DGESN.- |
| **HIRA interacting protein 3** | HIRIP3 | S227 | | | | | | | **26.5** | **54.3** | K.ESEQES*EEEILAQK.K |
| **HIRA interacting protein 3** | HIRIP3 | S550S555 | | | | | | | **5.6** | **42.4** | R.GIIS*SDGES*N.- |
| **HIRA interacting protein 3** | HIRIP3 | S196 | | | | | | | **28.4** | **13.9** | R.EES*EESEAEPVQR.T |
| **HIRA interacting protein 3** | HIRIP3 | S219S227 | | | | | | | **13.0** | **29.8** | K.S*LKESEQES*EEEILAQKK.E |
| **HIRA interacting protein 3** | HIRIP3 | S357S359S363 | | | | | | | **13.7** | **25.2** | R.LGS*TS*GEES*DLER.E |
| **Histamine receptor H1** | HRH1 | S396S398 | | | | | | | **100.0** | **12.5** | R.LRS*HS*R.Q |
| **Histidyl tRNA synthetase 2** | DTD1 | S197 | | | | | | | **9.1** | **77.1** | R.SASS*GAEGDVSSEREP.- |
| **Histidyl tRNA synthetase 2** | DTD1 | S196 | | | | | | | **5.5** | **78.3** | R.SAS*SGAEGDVSSEREP.- |
| **Histidyl tRNA synthetase 2** | DTD1 | S194 | | | | | | | **7.0** | **66.0** | R.S*ASSGAEGDVSSEREP.- |
| **Histidyl tRNA synthetase like** | HARS2 | S67 | | | | | | | **100.0** | **36.7** | R.DLS*PQHMVVR.E |

Peak Area: <-10, -5, -3, 0, 3, 5, >10

%CV: 0, 29, 42, 71, 86, >100

**White dots-** Significant change in peptide abundance at **5** %FDR compared to the timepoint with the minimum peak area for a given PSM

| Protein Name | Gene | Phosphosites | CarT 0m 2m 5m | RajiB 0m 2m 5m | Ascor | MOWSE | Sequence |
|---|---|---|---|---|---|---|---|
| **Histone 1 H1C** | HIST1H1C | T31 | | | **100.0** | **24.9** | K.KAGGT*PR.K |
| **Histone 1 H1C** | HIST1H1C | T146 | | | **100.0** | **48.1** | K.KAAGGAT*PK.K |
| **Histone 1 H1E** | HIST1H1E | T146 | | | **49.2** | **49.6** | K.KATGAAT*PK.K |
| **Histone 1 H1E** | HIST1H1E | S187 | | | **100.0** | **51.4** | K.KAPKS*PAK.A |
| **Histone 1 H1E** | HIST1H1E | T142 | | | **7.3** | **11.3** | K.KAT*GAATPK.K |
| **Histone 1, H1a** | HIST1H1A | S165S166 | | | **100.0** | **18.2** | R.KS*S*KNPK.K |
| **Histone 1, H1b** | HIST1H1B | S18 | | | **47.5** | **47.0** | M.SETAPAETATPAPVEKS*PAK.K |
| **Histone 1, H1b** | HIST1H1B | S173 | | | **100.0** | **40.3** | K.KVAKS*PK.K |
| | HIST1H1B | S189 | | | **26.0** | **31.0** | K.ATKS*PAKPK.A |
| **Histone 1, H1b** | HIST1H1B | T138 | | | **100.0** | **19.0** | K.AKKPAGAT*PK.K |
| **Histone acetyltransferase 1** | HAT1 | S361 | | | **10.9** | **21.1** | R.LIS*PYK.K |
| **Histone deacetylase 1** | HDAC1 | S421S423 | | | **100.0** | **88.9** | R.IACEEEFS*DS*EEEGEGGR.K |
| **Histone deacetylase 1** | HDAC1 | S393 | | | **100.0** | **44.5** | R.MLPHAPGVQMQAIPEDAIPEES*GDEDEDDPDKR.I |
| | HDAC2 | S516S518 | | | **100.0** | **82.1** | R.IACDEEFS*DS*EDEGEGGRR.N |
| **Histone deacetylase 2** | HDAC2 | S488 | | | **100.0** | **30.3** | R.MLPHAPGVQMQAIPEDAVHEDS*GDEDGEDPDKR.I |
| **Histone deacetylase 3** | HDAC3 | S424 | | | **49.9** | **30.1** | R.GPEENYSRPEAPNEFYDGDHDNDKES*DVEI.- |
| **Histone deacetylase 4** | HDAC4 | S632 | | | **35.8** | **110.8** | R.AQS*SPASATFPVSVQEPPTKPR.F |
| **Histone deacetylase 4** | HDAC4 | S467 | | | **12.1** | **45.6** | R.TQS*APLPQNAQALQHLVIQQQHQQFLEK.H |
| **Histone deacetylase 7A** | HDAC7 | S486 | | | **56.6** | **83.2** | R.AQS*SPAAPASLSAPEPASQAR.V |
| **Histone deacetylase 7A** | HDAC7 | S487 | | | **-0.4** | **32.8** | R.AQSS*PAAPASLSAPEPASQAR.V |
| **Histone H1x** | H1FX | S31 | | | **29.3** | **73.1** | K.AGGSAALS*PSK.K |
| **Histone H1x** | H1FX | S33 | | | **11.0** | **55.4** | K.AGGSAALSPS*KK.R |
| **Histone methyltransferase DOT1L** | DOT1L | S447 | | | **12.6** | **59.8** | K.KNQTALDALHAQTVSQTAAS*SPQDAYR.S |
| **Histone methyltransferase DOT1L** | DOT1L | S448 | | | **7.2** | **61.9** | K.KNQTALDALHAQTVSQTAASS*PQDAYR.S |
| **Histone1, H1D** | HIST1H1D | T147 | | | **100.0** | **38.7** | K.KVAGAAT*PK.K |
| **HIV 1 REV binding protein** | AGFG1 | T177S179 | | | **9.2** | **14.7** | K.SLLGDSAPTLHLNKGT*PS*QSPVVGR.S |
| **HIV 1 REV binding protein** | AGFG1 | T177 | | | **28.8** | **43.5** | K.GT*PSQSPVVGR.S |
| **HIV 1 REV binding protein** | AGFG1 | T177S181 | | | **16.0** | **21.1** | K.SLLGDSAPTLHLNKGT*PSQS*PVVGR.S |
| **HIV 1 REV binding protein** | AGFG1 | S162 | | | **78.9** | **54.1** | K.S*LLGDSAPTLHLNK.G |
| **HIV 1 REV binding protein** | AGFG1 | T170S181 | | | **4.1** | **14.5** | K.SLLGDSAPT*LHLNKGTPSQS*PVVGR.S |
| **HLA class I histocompatibility antigen, B-58 alpha chain** | HLA-B | S359 | | | **12.1** | **69.6** | K.GGSYSQAASSDSAQGSDVS*LTA.- |
| **HLA class I histocompatibility antigen, B-58 alpha chain** | HLA-B | S356 | | | **14.0** | **51.8** | K.GGSYSQAASSDSAQGS*DVSLTA.- |
| **HLA class I histocompatibility antigen, B-58 alpha chain** | HLA-B | S356S359 | | | **16.9** | **74.7** | K.GGSYSQAASSDSAQGS*DVS*LTA.- |
| **HLA class I histocompatibility antigen, B-58 alpha chain** | HLA-B | S352S356S359 | | | **11.0** | **38.4** | K.GGSYSQAASSDS*AQGS*DVS*LTA.- |
| **HLA class I histocompatibility antigen, B-58 alpha chain** | HLA-B | S350S352S359 | | | **10.2** | **16.6** | K.GGSYSQAASS*DS*AQGSDVS*LTA.- |
| **HLA class I histocompatibility antigen, B-58 alpha chain** | HLA-B | S350S352S356 | | | **-0.5** | **14.3** | K.GGSYSQAASS*DS*AQGS*DVSLTA.- |
| **HLA class I histocompatibility antigen, B-58 alpha chain** | HLA-B | S350S356S359 | | | **14.4** | **35.9** | K.GGSYSQAASS*DSAQGS*DVS*LTA.- |
| **HLA-A** | HLA-A | S359 | | | **28.2** | **114.2** | R.KGGSYTQAASSDSAQGSDVS*LTACKV.- |
| **HLA-A** | HLA-A | S352 | | | **33.2** | **107.3** | R.KGGSYTQAASSDS*AQGSDVSLTACKV.- |
| **HLA-A** | HLA-A | S350 | | | **10.9** | **68.8** | R.KGGSYTQAASS*DSAQGSDVSLTACKV.- |
| **HLA-A** | HLA-A | S352S359 | | | **26.1** | **70.0** | R.KGGSYTQAASSDS*AQGSDVS*LTACKV.- |
| **HLA-A** | HLA-A | S352S356S359 | | | **20.0** | **70.8** | R.KGGSYTQAASSDS*AQGS*DVS*LTACKV.- |
| **HLA-A** | HLA-A | Y344 | | | **17.7** | **54.7** | R.KGGSY*TQAASSDSAQGSDVSLTACKV.- |
| **HLA-A** | HLA-A | S356 | | | **15.9** | **96.9** | R.KGGSYTQAASSDSAQGS*DVSLTACKV.- |
| **HLA-A** | HLA-A | S356S359 | | | **19.4** | **50.0** | R.KGGSYTQAASSDSAQGS*DVS*LTACKV.- |
| **HLA-A** | HLA-A | S350S359 | | | **8.3** | **33.8** | R.KGGSYTQAASS*DSAQGSDVS*LTACKV.- |
| **HLA-A** | HLA-A | T345 | | | **9.0** | **27.2** | R.KGGSYT*QAASSDSAQGSDVSLTACKV.- |
| **HLA-A** | HLA-A | S352S356 | | | **8.9** | **39.6** | R.KGGSYTQAASSDS*AQGS*DVSLTACKV.- |
| **HLA-B associated transcript 2** | PRRC2A | S342S350 | | | **100.0** | **41.3** | K.LKFS*DEEDGRDS*DEEGAEGHR.D |
| **HLA-B associated transcript 2** | PRRC2A | S456 | | | **22.5** | **81.7** | R.KQS*SSEISLAVER.A |
| **HLA-B associated transcript 2** | PRRC2A | S1089S1092 | | | **52.2** | **83.2** | R.S*EGS*EYEEIPK.R |
| **HLA-B associated transcript 2** | PRRC2A | S1014 | | | **15.8** | **13.9** | R.DYS*YER.V |
| **HLA-B associated transcript 2** | PRRC2A | T825 | | | **-0.9** | **21.4** | R.SET*PPVPPPPPYLASYPGFPENGAPGPPISR.F |
| **HLA-B associated transcript 2** | PRRC2A | Y1013 | | | **18.1** | **21.2** | R.DY*SYER.V |
| **HLA-B associated transcript 2** | PRRC2A | S1168 | | | **100.0** | **11.1** | R.GVPS*RR.G |
| **HLA-B associated transcript 2** | PRRC2A | S1306S1310S1 | | | **11.9** | **18.4** | K.S*PDLS*NQNS*DQANEEWET*ASESSDFTSER.R |
| **HLA-B associated transcript 2** | PRRC2A | S1306S1314S1 | | | **17.8** | **15.8** | K.S*PDLSNQNS*DQANEEWETAS*ES*SDFTSER.R |
| **HLA-B associated transcript 2** | PRRC2A | S457 | | | **8.0** | **27.8** | R.KQSS*SEISLAVER.A |
| **HLA-B associated transcript 2** | PRRC2A | S1089 | | | **30.0** | **30.0** | R.S*EGSEYEEIPK.R |
| **HLA-B associated transcript 3** | BAT3 | S973 | | | **75.8** | **114.5** | R.ENAS*PAPGTTAEEAMSR.G |
| **HLA-B associated transcript 3** | BAT3 | S964S973 | | | **36.3** | **44.8** | R.AS*PEPQRENAS*PAPGTTAEEAMSR.G |
| **HLA-B associated transcript 3** | BAT3 | S964 | | | **100.0** | **27.7** | R.AS*PEPQR.E |

Peak Area: <-10, -5, -3, 0, 3, 5, >10

%CV: 0, 29, 42, 71, 86, >100

**White dots-** Significant change in peptide abundance at **5** %FDR compared to the timepoint with the minimum peak area for a given PSM

CarT: 0m 2m 5m; RajiB: 0m 2m 5m

| Protein Name | Gene | Phosphosites | Ascor | MOWSE | Sequence |
|---|---|---|---|---|---|
| **HLA-B associated transcript 3** | BAT3 | S113 | 7.7 | 40.4 | R.APPQTHLPSGASSGTGSASATHGGGS*PPGTR.G |
| **HLA-B associated transcript 3** | BAT3 | S964T978 | 5.9 | 19.9 | R.AS*PEPQRENASPAPGT*TAEEAMSR.G |
| **HLA-B associated transcript 3** | BAT3 | T92 | | 13.7 | R.APPQT*HLPSGASSGTGSASATHGGGSPPGTR.G |
| **HLA-B-associated transcript 1** | DDX39B | S38 | 30.0 | 32.9 | K.GS*YVSIHSSGFR.D |
| **HMG CoA reductase** | HMGCR | S872 | 120.6 | 84.5 | R.S*KINLQDLQGACTK.K |
| **HMG-BOX transcription factor BBX** | BBX | S478S479S481 | 19.4 | 35.3 | K.KRQS*S*ES*DIESVIYTIEAVAK.G |
| **HMGN1** | HMGN1 | S86S89 | 45.2 | 78.0 | K.TEES*PAS*DEAGEK.E |
| **HMGN1** | HMGN1 | S89 | 32.3 | 67.2 | K.TEESPAS*DEAGEK.E |
| **HNRPLL** | HNRNPLL | T46 | 100.0 | 12.0 | R.REAT*PR.G |
| **Host cell factor C1** | HCFC1 | S666 | 50.0 | 72.9 | K.S*PISVPGGSALISNLGK.V |
| **Host cell factor C1** | HCFC1 | S411 | 6.5 | 19.9 | K.YDIPATAATATS*PTPNPVPSVPANPPK.S |
| **Host cell factor C1** | HCFC1 | T413 | -0.3 | 31.8 | K.YDIPATAATATSPT*PNPVPSVPANPPK.S |
| **Host cell factor C1** | HCFC1 | S1205 | 113.9 | 56.1 | R.S*PAFVQLAPLSSK.V |
| **HP1 beta** | CBX1 | S89 | 39.0 | 51.5 | R.KADS*DSEDKGEESKPK.K |
| **HP1-BP74** | HP1BP3 | S441S442S446 | 84.1 | 40.7 | K.KEPDDSRDEDEDEDES*S*EEDS*EDEEPPPK.R |
| **HPK1** | MAP4K1 | S376S377Y381 | 6.7 | 16.9 | R.KQLSES*S*DDDY*DDVDIPTPAEDTPPPLPPKPK.F |
| **HPK1** | MAP4K1 | S374S376S377 | 10.7 | 14.8 | R.KQLS*ES*S*DDDYDDVDIPTPAEDTPPPLPPKPK.F |
| **HPRP3P** | PRPF3 | S619 | 100.0 | 64.2 | K.GDDDEES*DEEAVKK.T |
| **HPRP3P** | PRPF3 | T611 | 5.8 | 20.1 | R.IKWDEQTSNT*KGDDDEESDEEAVK.K |
| **Hsc70 interacting protein** | ST13 | S79 | 16.4 | 86.4 | K.KVEEDLKADEPSSEES*DLEIDK.E |
| **Hsc70 interacting protein** | ST13 | S76 | 9.1 | 83.0 | K.KVEEDLKADEPSS*EESDLEIDK.E |
| **Hsc70 interacting protein** | ST13 | S75S76S79 | 100.0 | 71.6 | K.KVEEDLKADEPS*S*EES*DLEIDK.E |
| **Hsc70 interacting protein** | ST13 | S75S79 | 28.4 | 56.5 | K.ADEPS*SEES*DLEIDK.E |
| | ST13 | S75S76 | | 39.6 | K.ADEPS*S*EESDLEIDK.E |
| | ST13 | S76S79 | 19.1 | 41.8 | K.ADEPSS*EES*DLEIDK.E |
| **Hsc70 interacting protein** | ST13 | S75 | 10.9 | 48.0 | K.KVEEDLKADEPS*SEESDLEIDK.E |
| **HSHIN1** | OTUD4 | S480 | 15.3 | 54.8 | K.RPEPSTLENITDDKYATVSS*PSK.S |
| **HSHIN1** | OTUD4 | S957S958 | 103.0 | 105.1 | K.EES*S*EDENEVSNILR.S |
| **HSHIN1** | OTUD4 | S827 | 33.4 | 19.1 | K.GELDLS*LENLDLSK.D |
| **HSHIN1** | OTUD4 | S479 | 10.7 | 69.2 | K.RPEPSTLENITDDKYATVS*SPSK.S |
| **HSP90A** | HSP90AA1 | S263 | 144.9 | 97.3 | K.ESEDKPEIEDVGS*DEEEEKK.D |
| **HSP90A** | HSP90AA1 | T725 | 76.3 | 74.7 | K.LGLGIDEDDPTADDTSAAVTEEMPPLEGDDDT*SR.M |
| | HSP90AA1 | S231 | 100.0 | 85.4 | R.DKEVS*DDEAEEKEDK.E |
| **HSP90A** | HSP90AA1 | S252 | | 44.4 | K.ES*EDKPEIEDVGSDEEEEKK.D |
| **HSP90B** | HSP90AB1 | S226 | 100.0 | 87.7 | K.EIS*DDEAEEEKGEK.E |
| **HSP90B** | HSP90AB1 | S255 | 71.8 | 131.8 | K.IEDVGS*DEEDDSGKDK.K |
| **HSP90B** | HSP90AB1 | S718 | 100.0 | 47.0 | K.LGLGIDEDEVAAEEPNAAVPDEIPPLEGDEDAS*R.M |
| **Hugl protein** | LLGL1 | S936 | 10.9 | 12.6 | R.FS*LSAR.N |
| **Human immunodeficiency virus type1 enhancer-binding protein2** | HIVEP2 | S2130 | 100.0 | 15.0 | R.RDLS*PR.R |
| **Huntingtin** | HTT | S421S434 | 15.5 | 90.1 | R.SGS*IVELIAGGGSSCS*PVLSR.K |
| **Huntingtin** | HTT | S432 | 9.1 | 52.0 | R.SGSIVELIAGGGSS*CSPVLSR.K |
| **Huntingtin** | HTT | S419S434 | 12.8 | 70.4 | R.S*GSIVELIAGGGSSCS*PVLSR.K |
| **Huntingtin** | HTT | S421S432 | 1.4 | 23.9 | R.SGS*IVELIAGGGSS*CSPVLSR.K |
| **Huntingtin interacting protein 1** | SETD2 | S121 | | 20.5 | R.LNDS*PTLK.K |
| **Huntingtin interacting protein 1** | SETD2 | T123 | 6.8 | 10.8 | R.LNDSPT*LK.K |
| **Hydroxymethylglutaryl-CoA synthase, cytoplasmic** | HMGCS1 | T471 | | 38.5 | R.RPT*PNDDTLDEGVGLVHSNIATEHIPSPAKK.V |
| | HMGCS1 | S495 | 49.7 | 24.6 | R.RPTPNDDTLDEGVGLVHSNIATEHIPS*PAKK.V |
| **Hyperpolarization activated cyclic nucleotide gated potassium channel** | HCN3 | T380 | 8.7 | 11.1 | K.YKQVEQYMSFHKLPADT*R.Q |
| **Hypothetical protein BC007540** | C11orf84 | S248S251 | 12.8 | 18.6 | K.NLDPDPEPPS*PDS*PTETFAAPAEVR.H |
| **Hypothetical protein BC008207** | NAF1 | S315 | 100.0 | 54.6 | K.NDQEPPPEALDFS*DDEKEK.E |
| **Hypothetical protein DKFZp762E1312** | HJURP | S473 | 68.0 | 40.8 | R.GGPAS*PGGLQGLETR.R |
| **Hypothetical protein DKFZp762E1312** | HJURP | T600 | -9.4 | 20.0 | K.SPGQMT*VPLCIGVSTDK.A |
| **Hypothetical protein FLJ10154** | ARGLU1 | S76 | 12.2 | 39.7 | R.AS*SPPDRIDIFGR.T |
| **Hypothetical protein FLJ10154** | ARGLU1 | S77 | 15.0 | 33.0 | R.ASS*PPDRIDIFGR.T |
| **Hypothetical protein FLJ10154** | ARGLU1 | S60 | 17.0 | 53.0 | R.S*TNTAVSR.R |
| **Hypothetical protein FLJ10154** | ARGLU1 | S58S60 | 16.3 | 18.6 | R.S*RS*TNTAVSR.R |
| **Hypothetical protein FLJ10154** | ARGLU1 | T61 | -0.1 | 56.0 | R.ST*NTAVSR.R |
| **Hypothetical protein FLJ20160** | MFSD6 | S644 | 5.3 | 59.4 | R.IPVPSS*PVPIATIDLVQQQTEDVMPR.I |
| **Hypothetical protein FLJ20309** | INO80D | S232 | 29.8 | 48.9 | K.S*PQPQNTSLPMQGVAPTTHTIAQAR.Q |
| | RASAL3 | S164S166 | 21.4 | 22.8 | R.VGS*AS*SEGSIHVAMGNFRDPDRMPGK.T |
| **Hypothetical protein FLJ21438** | RASAL3 | S166S167 | 2.6 | 78.4 | R.VGSAS*S*EGSIHVAMGNFR.D |
| **Hypothetical protein FLJ21438** | RASAL3 | S164S167 | 12.5 | 80.0 | R.VGS*ASS*EGSIHVAMGNFR.D |

Peak Area: <-10, -5, -3, 0, 3, 5, >10

%CV: 0, 29, 42, 71, 86, >100

**White dots**- Significant change in peptide abundance at **5** %FDR compared to the timepoint with the minimum peak area for a given PSM

CarT (0m, 2m, 5m) RajiB (0m, 2m, 5m)

| Protein Name | Gene | Phosphosites | Ascor | MOWSE | Sequence |
|---|---|---|---|---|---|
| Hypothetical protein FLJ21438 | RASAL3 | S228S231 | 12.8 | 75.4 | R.DGPPSALGS*RES*LATLSELDLGAER.D |
| Hypothetical protein FLJ21438 | RASAL3 | S72 | 29.9 | 40.5 | R.TQS*VPVR.R |
| Hypothetical protein FLJ21438 | RASAL3 | S164S166S167 | 6.7 | 44.2 | R.VGS*AS*S*EGSIHVAMGNFR.D |
| Hypothetical protein FLJ21438 | RASAL3 | S94 | 32.1 | 46.9 | K.GS*LSMGPAPR.A |
| Hypothetical protein FLJ21438 | RASAL3 | T70 | 12.2 | 35.7 | R.T*QSVPVR.R |
| Hypothetical protein FLJ21438 | RASAL3 | S247 | -0.2 | 21.1 | R.TRGS*WSQPQPLK.A |
| Hypothetical protein FLJ21438 | RASAL3 | S166S167S170 | 5.5 | 22.4 | R.VGSAS*S*EGS*IHVAMGNFR.D |
| Hypothetical protein FLJ25476 | ZNF362 | S404 | 51.2 | 62.5 | K.HTVVEHLVSHHS*PQR.T |
| Hypothetical protein FLJ25476 | ZNF362 | S401 | 7.1 | 60.8 | K.HTVVEHLVS*HHSPQR.T |
| Hypothetical protein KIAA0084 (HA2022) (Fragment) | | S32 | 100.0 | 14.4 | R.RPS*PPR.R |
| Hypothetical protein KIAA0826 | FRYL | T745 | -0.2 | 92.2 | K.ST*GQLNLSTSPINSSSYLGYNSNAR.S |
| Hypothetical protein KIAA0826 | FRYL | T752 | 5.8 | 43.7 | K.STGQLNLST*SPINSSSYLGYNSNAR.S |
| Hypothetical protein KIAA0826 | FRYL | T745S751 | 8.2 | 76.4 | K.ST*GQLNLS*TSPINSSSYLGYNSNAR.S |
| Hypothetical protein KIAA0826 | FRYL | S753 | 19.1 | 51.5 | K.STGQLNLSTS*PINSSSYLGYNSNAR.S |
| Hypothetical protein KIAA0826 | FRYL | S744S757 | 11.9 | 14.8 | K.S*TGQLNLSTSPINS*SSYLGYNSNAR.S |
| Hypothetical protein KIAA0826 | FRYL | T745S757 | 4.0 | 55.0 | K.ST*GQLNLSTSPINS*SSYLGYNSNAR.S |
| Hypothetical protein KIAA0826 | FRYL | S744 | 8.0 | 111.2 | K.S*TGQLNLSTSPINSSSYLGYNSNAR.S |
| Hypothetical protein KIAA0826 | FRYL | T745S753 | 1.2 | 52.0 | K.ST*GQLNLSTS*PINSSSYLGYNSNAR.S |
| Hypothetical protein KIAA0889 | SOGA1 | S64 | 11.5 | 68.9 | K.SVSSMSEFES*LLDCSPYLAGGDAR.G |
| Hypothetical protein KIAA0889 | SOGA1 | S57S64 | 0.6 | 62.1 | R.TKSVS*SMSEFES*LLDCSPYLAGGDAR.G |
| Hypothetical protein KIAA0889 | SOGA1 | S55S60 | -0.8 | 58.0 | R.TKS*VSSMS*EFESLLDCSPYLAGGDAR.G |
| Hypothetical protein KIAA0889 | SOGA1 | S57S58 | 1.8 | 59.9 | K.SVS*S*MSEFESLLDCSPYLAGGDAR.G |
| Hypothetical protein KIAA0889 | SOGA1 | S55S57 | 6.9 | 72.6 | K.S*VS*SMSEFESLLDCSPYLAGGDAR.G |
| Hypothetical protein KIAA0889 | SOGA1 | S60S64 | 1.1 | 40.2 | R.TKSVSSMS*EFES*LLDCSPYLAGGDAR.G |
| Hypothetical protein KIAA0889 | SOGA1 | S57 | 15.8 | 27.6 | K.SVS*SMSEFESLLDCSPYLAGGDAR.G |
| Hypothetical protein KIAA0889 | SOGA1 | S55S58 | 6.8 | 60.0 | K.S*VSS*MSEFESLLDCSPYLAGGDAR.G |
| Hypothetical protein KIAA0889 | SOGA1 | S60 | 1.2 | 70.5 | K.SVSSMS*EFESLLDCSPYLAGGDAR.G |
| Hypothetical protein KIAA0889 | SOGA1 | S58 | 9.8 | 19.6 | K.SVSS*MSEFESLLDCSPYLAGGDAR.G |
| Hypothetical protein KIAA0889 | SOGA1 | S55S64 | 8.0 | 75.5 | R.TKS*VSSMSEFES*LLDCSPYLAGGDAR.G |
| Hypothetical protein KIAA0889 | SOGA1 | S55 | 16.1 | 115.2 | K.S*VSSMSEFESLLDCSPYLAGGDAR.G |
| Hypothetical protein LOC137886 | UBXN2B | S17 | 10.9 | 16.0 | K.RSS*GPR.A |
| Hypothetical protein LOC137886 | UBXN2B | S16 | 23.9 | 27.3 | K.RS*SGPR.A |
| Hypothetical protein LOC137886 | UBXN2B | S235S242 | 5.6 | 54.9 | K.LGSLTPEIVSTPSS*PEEEDKS*ILNAVVLIDDSVPTTK.I |
| Hypothetical protein LOC137886 | UBXN2B | T232S242 | 2.6 | 52.0 | K.LGSLTPEIVST*PSSPEEEDKS*ILNAVVLIDDSVPTTK.I |
| Hypothetical protein LOC137886 | UBXN2B | T232S234 | 4.3 | 51.0 | K.LGSLTPEIVST*PS*SPEEEDKSILNAVVLIDDSVPTTK.I |
| Hypothetical protein LOC137886 | UBXN2B | S16S17 | 100.0 | 21.7 | K.RS*S*GPR.A |
| Hypothetical protein LOC137886 | UBXN2B | S234S235 | 2.9 | 31.3 | K.LGSLTPEIVSTPS*S*PEEEDKSILNAVVLIDDSVPTTK.I |
| Hypothetical protein LOC137886 | UBXN2B | S224S234 | 1.6 | 17.9 | K.LGS*LTPEIVSTPS*SPEEEDKSILNAVVLIDDSVPTTK.I |
| Hypothetical protein LOC137886 | UBXN2B | S234S242 | 11.4 | 23.4 | K.LGSLTPEIVSTPS*SPEEEDKS*ILNAVVLIDDSVPTTK.I |
| Hypothetical protein LOC137886 | UBXN2B | S231S234 | 4.3 | 35.1 | K.LGSLTPEIVS*TPS*SPEEEDKSILNAVVLIDDSVPTTK.I |
| Hypothetical protein LOC162427 | RETREG3 | S258S260 | 171.7 | 58.0 | R.AMDNHS*DS*EEELAAFCPQLDDSTVAR.E |
| Hypothetical protein LOC162427 | RETREG3 | T440 | 31.8 | 47.2 | R.SPSSDLDT*DAEGDDFELLDQSELSQLDPASSR.S |
| Hypothetical protein LOC162427 | RETREG3 | S435S436 | 8.4 | 90.0 | R.SPS*S*DLDTDAEGDDFELLDQSELSQLDPASSR.S |
| Hypothetical protein LOC162427 | RETREG3 | S433S436T440 | 3.9 | 67.6 | R.S*PSS*DLDT*DAEGDDFELLDQSELSQLDPASSR.S |
| Hypothetical protein LOC162427 | RETREG3 | S435S436T440 | 12.2 | 77.7 | R.SPS*S*DLDT*DAEGDDFELLDQSELSQLDPASSR.S |
| Hypothetical protein LOC162427 | RETREG3 | S350 | 12.1 | 70.9 | R.DLPDFPSINMDPAGLDDEDDTS*IGMPSLMYR.S |
| Hypothetical protein LOC162427 | RETREG3 | T349 | 8.2 | 36.4 | R.DLPDFPSINMDPAGLDDEDDT*SIGMPSLMYR.S |
| Hypothetical protein LOC162427 | RETREG3 | S435T440 | 19.3 | 89.5 | R.SPS*SDLDT*DAEGDDFELLDQSELSQLDPASSR.S |
| Hypothetical protein LOC162427 | RETREG3 | S433S435 | 7.9 | 11.1 | R.S*PS*SDLDTDAEGDDFELLDQSELSQLDPASSR.S |
| Hypothetical protein LOC162427 | RETREG3 | S335 | 8.9 | 32.7 | R.DLPDFPS*INMDPAGLDDEDDTSIGMPSLMYR.S |
| Hypothetical protein LOC162427 | RETREG3 | S435 | 17.1 | 96.6 | R.SPS*SDLDTDAEGDDFELLDQSELSQLDPASSR.S |
| Hypothetical protein LOC162427 | RETREG3 | T307T310S313 | 37.5 | 24.9 | R.GQT*PLT*EGS*EDLDGHS*DPEESFAR.D |
| Hypothetical protein LOC162427 | RETREG3 | S436T440 | 11.3 | 52.8 | R.SPSS*DLDT*DAEGDDFELLDQSELSQLDPASSR.S |
| Hypothetical protein LOC162427 | RETREG3 | S433S435T440 | 10.2 | 40.5 | R.S*PS*SDLDT*DAEGDDFELLDQSELSQLDPASSR.S |
| Hypothetical protein LOC162427 | RETREG3 | S436 | -1.2 | 21.9 | R.SPSS*DLDTDAEGDDFELLDQSELSQLDPASSR.S |
| Hypothetical protein LOC348180 | CTU2 | S508 | 7.1 | 17.0 | R.DCLIEDS*DDEAGQS.- |
| Hypothetical protein LOC348262 | MCRIP1 | S21 | 28.1 | 87.1 | R.S*PPSSSEIFTPAHEENVR.F |
| Hypothetical protein LOC348262 | MCRIP1 | S25 | 13.1 | 55.1 | R.SPPSS*SEIFTPAHEENVR.F |
| Hypothetical protein LOC348262 | MCRIP1 | S24 | | 14.2 | R.SPPS*SSEIFTPAHEENVR.F |
| Hypothetical protein LOC55580 | CCDC88A | S155S158 | 12.5 | 40.6 | R.DGLHFLPHASSS*AQS*PCGSPGMKR.T |
| Hypothetical protein LOC55580 | CCDC88A | S158 | 16.7 | 55.8 | R.DGLHFLPHASSSAQS*PCGSPGMK.R |
| Hypothetical protein LOC55580 | CCDC88A | S153S162 | 14.7 | 10.8 | R.DGLHFLPHAS*SSAQSPCGS*PGMKR.T |

**Peak Area**: <-10, -5, -3, 0, 3, 5, >10

**%CV**: 0, 29, 42, 71, 86, >100

**White dots-** Significant change in peptide abundance at **5** %FDR compared to the timepoint with the minimum peak area for a given PSM

CarT: 0m 2m 5m — RajiB: 0m 2m 5m

| Protein Name | Gene | Phosphosites | Ascor | MOWSE | Sequence |
|---|---|---|---|---|---|
| **Hypothetical protein LOC55580** | CCDC88A | S154 | **3.5** | **44.5** | R.DGLHFLPHASS*SAQSPCGSPGMK.R |
| **Hypothetical protein LOC55580** | CCDC88A | S158S162 | **37.5** | **40.8** | R.DGLHFLPHASSSAQS*PCGS*PGMK.R |
| **Hypothetical protein LOC729440** | CCDC61 | S392S395 | **57.5** | **86.8** | R.LGS*GGS*GDGPSVSWSR.Q |
| **Hypothetical protein MGC22014** | TET3 | T657S666 | **18.6** | **18.0** | K.AENPLT*PTLSGFLES*PLK.Y |
| **Hypoxia inducible factor 1 alpha subunit** | HIF3A | S22S28 | **100.0** | **14.1** | K.S*RDAARS*R.R |
| **Hypoxia inducible factor 1 alpha subunit** | HIF1A | S687 | **63.9** | **53.0** | R.S*PNVLSVALSQR.T |
| **Ig alpha** | CD79A | S215 | **15.2** | **30.5** | R.GLQGTYQDVGS*LNIGDVQLEKP.- |
| **IGF II mRNA binding protein 3** | IGF2BP3 | S184 | **22.0** | **51.9** | R.QGS*PGSVSK.Q |
| **IGF II receptor** | IGF2R | S2409 | **25.8** | **65.3** | K.ALSSLHGDDQDS*EDEVLTIPEVK.V |
| **IGF II receptor** | IGF2R | S2479 | **21.0** | **23.6** | K.LVS*FHDDSDEDLLHI.- |
| **IGF II receptor** | IGF2R | S2479S2484 | **100.0** | **45.4** | K.LVS*FHDDS*DEDLLHI.- |
| **IGF II receptor** | IGF2R | S2484 | **19.9** | **22.2** | K.LVSFHDDS*DEDLLHI.- |
| **IK factor** | IK | S460 | **49.0** | **76.3** | K.QLGDFFGMSNSYAECYPATMDDMAVDS*DEEVDYSK.M |
| **Immediate early response erythropoietin 4** | ZC3H15 | S328 | **11.1** | **74.2** | R.YTQGTGGDEVDDSVS*VNDIDLSLYIPR.D |
| **Immediate early response erythropoietin 4** | ZC3H15 | S326 | **20.4** | **42.7** | R.YTQGTGGDEVDDS*VSVNDIDLSLYIPR.D |
| **Immediate early response erythropoietin 4** | ZC3H15 | S381 | **100.0** | **52.3** | R.S*DLEEDNER.E |
| **Immediate early response erythropoietin 4** | ZC3H15 | T315 | **-0.4** | **28.5** | R.YT*QGTGGDEVDDSVSVNDIDLSLYIPR.D |
| **Immunoglobulin superfamily, member 3** | IGSF3 | T637 | **15.1** | **16.9** | R.TRT*AIEK.A |
| **IMP dehydrogenase 2** | IMPDH2 | S159 | **8.9** | **33.6** | R.LVGIIS*SR.D |
| **IMP dehydrogenase 2** | IMPDH2 | S416 | **100.0** | **45.4** | R.GMGS*LDAMDK.H |
| **Importin 7** | IPO7 | Y876S886 | | **84.0** | R. AY*ACHAEHENDS*DDDDEAEDDDETEELGSDEDDIDEDGQEYLEIL |
| **Importin 7** | IPO7 | S886S903 | **31.9** | **73.8** | R. AYACHAEHENDS*DDDDEAEDDDETEELGS*DEDDIDEDGQEYLEIL |
| **Importin 7** | IPO7 | S886T898 | **25.2** | **81.1** | R. AYACHAEHENDS*DDDDEAEDDDET*EELGSDEDDIDEDGQEYLEIL |
| **Importin 7** | IPO7 | S886Y915 | **15.2** | **67.6** | R. AYACHAEHENDS*DDDDEAEDDDETEELGSDEDDIDEDGQEY*LEIL |
| **Importin 7** | IPO7 | T898S903 | **12.6** | **29.7** | R. AYACHAEHENDSDDDDEAEDDDET*EELGS*DEDDIDEDGQEYLEIL |
| **Influenza virus NS1A binding protein** | IVNS1ABP | S338 | **22.5** | **68.9** | K.SLS*FEMQQDELIEKPMSPMQYAR.S |
| **Inner centromere protein** | INCENP | Y818S824T828 | **13.2** | **60.6** | K.INPDNY*GMDLNS*DDST*DDEAHPR.K |
| **Inner centromere protein** | INCENP | S142T145 | **8.0** | **50.5** | R.AAAAAAAATMALAAPS*SPT*PESPTMLTK.K |
| **Inner centromere protein** | INCENP | S142S148 | **25.3** | **55.2** | R.AAAAAAAATMALAAPS*SPTPES*PTMLTK.K |
| **Inner centromere protein** | INCENP | Y818S827T828 | **8.2** | **71.4** | K.INPDNY*GMDLNSDDS*T*DDEAHPR.K |
| **Inner centromere protein** | INCENP | S302S312 | **7.2** | **14.5** | R.TDSQS*VRHSPIAPSS*PSPQVLAQK.Y |
| **Inner centromere protein** | INCENP | S824S827T828 | **8.9** | **75.8** | K.INPDNYGMDLNS*DDS*T*DDEAHPR.K |
| **Inner centromere protein** | INCENP | S263 | **100.0** | **21.1** | R.IAQVS*PGPR.D |
| **Inner centromere protein** | INCENP | S300S311 | **15.4** | **12.0** | R.TDS*QSVRHSPIAPS*SPSPQVLAQK.Y |
| **Inner centromere protein** | INCENP | S142T150 | **5.5** | **21.2** | R.AAAAAAAATMALAAPS*SPTPESPT*MLTK.K |
| **Inorganic pyrophosphatase 2** | PPA2 | S317 | **14.3** | **57.4** | R.SLVESVSSS*PNKESNEEEQVWHFLGK.- |
| **Inorganic pyrophosphatase 2** | PPA2 | S315 | **18.4** | **24.7** | R.SLVESVS*SSPNKESNEEEQVWHFLGK.- |
| **Inorganic pyrophosphatase 2** | PPA2 | S316 | **15.3** | **44.2** | R.SLVESVSS*SPNKESNEEEQVWHFLGK.- |
| **Inorganic pyrophosphatase 2** | PPA2 | S313 | **19.4** | **15.5** | R.SLVES*VSSSPNKESNEEEQVWHFLGK.- |
| **Inositol 1,4,5-trisphosphate receptor type 3** | ITPR3 | S934 | **22.2** | **79.1** | R.KQS*VFSAPSLSAGASAAEPLDR.S |
| **Inositol 1,4,5-trisphosphate receptor type 3** | ITPR3 | S937 | **3.2** | **23.1** | R.KQSVFS*APSLSAGASAAEPLDR.S |
| **Inositol 1,4,5-trisphosphate receptor type 3** | ITPR3 | S2670 | **100.0** | **52.3** | R.LGFVDVQNCIS*R.- |
| **Inositol polyphosphate 5-phosphatase** | INPP5J | S545S547 | **31.9** | **17.9** | R.S*PS*PQSR.R |
| **Inositol-trisphosphate 3-kinase B** | ITPKB | S174S176 | **100.0** | **22.0** | R.ARS*PS*PCPFR.S |
| **Integrator complex subunit 1** | INTS1 | S134 | **20.8** | **37.2** | R.RPSAAAKPS*GHPPPGDFIALGSK.G |
| **Integrator complex subunit 1** | INTS1 | S166 | **-0.8** | **33.8** | K.SS*PEQPIGQGR.I |
| **Integrin alpha 4** | ITGA4 | S1021 | **17.8** | **41.8** | R.RDS*WSYINSK.S |
| **Intercellular adhesion molecule 1** | ICAM1 | T530 | **25.4** | **35.8** | K.GTPMKPNTQAT*PP.- |
| **Intercellular adhesion molecule 1** | ICAM1 | T527 | **-0.4** | **31.4** | K.GTPMKPNT*QATPP.- |
| **Interferon gamma inducible protein 16** | IFI16 | S153 | **58.0** | **90.8** | K.VSEEQTQPPS*PAGAGMSTAMGR.S |
| **Interferon gamma inducible protein 16** | IFI16 | T723 | **5.0** | **12.2** | K.DILNPDSSM#ET*SPDFFF.- |
| **Interferon induced protein 35** | IFI35 | S36 | **100.0** | **27.0** | R.KELGDS*PK.D |
| **Interferon regulatory factor 2 binding protein 2** | IRF2BP2 | S71 | **54.1** | **73.3** | R.S*PPGAAASAAAKPPPLSAK.D |
| **Interferon regulatory factor 2 binding protein 2** | IRF2BP2 | S406 | **5.1** | **25.5** | K.IPMTPTSSFVSPPPPTAS*PHSNR.T |
| **Interferon regulatory factor 2 binding protein 2** | IRF2BP2 | S423 | **27.0** | **68.8** | R.TTPPEAAQNGQS*PMAALILVADNAGGSHASK.D |
| **Interferon regulatory factor 2 binding protein 2** | IRF2BP2 | T413S423 | **2.6** | **36.4** | R.TT*PPEAAQNGQS*PMAALILVADNAGGSHASK.D |
| **Interferon regulatory factor 2 binding protein 2** | IRF2BP2 | T413 | **-0.5** | **21.1** | R.TT*PPEAAQNGQSPMAALILVADNAGGSHASK.D |
| **Interferon regulatory factor 2 binding protein 2** | IRF2BP2 | S175 | **100.0** | **23.5** | K.LEEPPELNRQS*PNPR.R |
| **Interferon regulatory factor 2 binding protein 2** | IRF2BP2 | T404 | **3.5** | **14.5** | K.IPMTPTSSFVSPPPPT*ASPHSNR.T |
| **Interferon regulatory factor 3** | IRF3 | S173 | **31.8** | **36.7** | R.S*PSLDNPTPFPNLGPSENPLKR.L |
| **Interferon regulatory factor 3** | IRF3 | S175 | **26.3** | **53.9** | R.SPS*LDNPTPFPNLGPSENPLKR.L |
| **Interleukin 17 receptor** | IL17RA | S726S736 | **18.1** | **45.3** | R.NSVLFLPVDPEDS*PLGSSTPMAS*PDLLPEDVR.E |

Peak Area: <-10, -5, -3, 0, 3, 5, >10

%CV: 0, 29, 42, 71, 86, >100

**White dots**- Significant change in peptide abundance at **5** %FDR compared to the timepoint with the minimum peak area for a given PSM

| Protein Name | Gene | Phosphosites | Ascor | MOWSE | Sequence |
|---|---|---|---|---|---|
| Interleukin enhancer binding factor 3 | ILF3 | S792 | 33.3 | 112.9 | K.GYNHGQGSYSYSNSYNS*PGGGGGSDYNYESK.F |
| Interleukin enhancer binding factor 3 | ILF3 | S787 | 9.1 | 21.9 | K.GYNHGQGSYSYS*NSYNSPGGGGGSDYNYESK.F |
| Interleukin enhancer binding factor 3 | ILF3 | S476 | | 51.5 | R. DS*SKGEDSAEETEAKPAVVAPAPVVEAVSTPSAAFPSDATAEQGPIL |
| Interleukin enhancer binding factor 3 | ILF3 | S482 | | 76.2 | K.GEDS*AEETEAKPAVVAPAPVVEAVSTPSAAFPSDATAEQGPILTK. H |
| Interleukin enhancer binding factor 3 | ILF3 | S382 | 28.4 | 73.6 | K.RPMEEDGEEKS*PSK.K |
| Interleukin enhancer binding factor 3 | ILF3 | S789 | 7.8 | 36.9 | K.GYNHGQGSYSYSNS*YNSPGGGGGSDYNYESK.F |
| Interleukin enhancer binding factor 3 | ILF3 | S476S477 | | 35.4 | R. DS*S*KGEDSAEETEAKPAVVAPAPVVEAVSTPSAAFPSDATAEQGP |
| Interleukin enhancer binding factor 3 | ILF3 | T469S476S477 | | 23.9 | K. VLQDMGLPT*GAEGRDS*S*KGEDSAEETEAKPAVVAPAPVVEAVST |
| Interleukin enhancer binding factor 3 | ILF3 | S860 | 7.4 | 108.9 | K.QGGYSQSNYNS*PGSGQNYSGPPSSYQSSQGGYGR.N |
| Interleukin enhancer binding factor 3 | ILF3 | S868 | 10.2 | 66.9 | K.QGGYSQSNYNSPGSGQNYS*GPPSSYQSSQGGYGR.N |
| Interleukin enhancer binding factor 3 | ILF3 | S476S477S482 | | 39.6 | R. DS*S*KGEDS*AEETEAKPAVVAPAPVVEAVSTPSAAFPSDATAEQGP |
| Interleukin enhancer binding factor 3 | ILF3 | Y858 | | 61.6 | K.QGGYSQSNY*NSPGSGQNYSGPPSSYQSSQGGYGR.N |
| Interleukin enhancer binding factor 3 | ILF3 | S854 | 9.0 | 31.0 | K.QGGYS*QSNYNSPGSGQNYSGPPSSYQSSQGGYGR.N |
| Interleukin enhancer binding factor 3 | ILF3 | T469S476 | | 15.3 | K. VLQDMGLPT*GAEGRDS*SKGEDSAEETEAKPAVVAPAPVVEAVSTP |
| Interleukin enhancer binding factor 3 | ILF3 | Y790 | -0.3 | 71.8 | K.GYNHGQGSYSYSNSY*NSPGGGGGSDYNYESK.F |
| Intersectin 1 | ITSN1 | S315 | -0.3 | 29.0 | R.SGS*GISVISSTSVDQR.L |
| Intersectin 2 | ITSN2 | S883S888 | 14.2 | 54.0 | R.TVS*PGSVS*PIHGQGQVVENLK.A |
| Intersectin 2 | ITSN2 | S886S888 | 12.6 | 33.2 | R.TVSPGS*VS*PIHGQGQVVENLK.A |
| Intersectin 2 | ITSN2 | S883S886 | 10.1 | 57.1 | R.TVS*PGS*VSPIHGQGQVVENLK.A |
| Intersectin 2 | ITSN2 | S888 | 0.4 | 41.6 | R.TVSPGSVS*PIHGQGQVVENLK.A |
| Intersectin 2 | ITSN2 | T881 | 10.1 | 26.4 | R.T*VSPGSVSPIHGQGQVVENLK.A |
| Intersectin 2 | ITSN2 | S886 | 22.2 | 28.4 | R.TVSPGS*VSPIHGQGQVVENLK.A |
| Intersectin 2 | ITSN2 | S883 | 7.6 | 34.5 | R.TVS*PGSVSPIHGQGQVVENLK.A |
| Intersectin 2 | ITSN2 | T881S888 | 6.3 | 37.9 | R.T*VSPGSVS*PIHGQGQVVENLK.A |
| Intraflagellar transport protein IFT20 | IFT20 | S72S76 | 100.0 | 12.6 | K.S*LAVS*PR.L |
| IQ motif and Sec7 domain 1 | IQSEC1 | S925 | 9.8 | 53.1 | R.SALSSS*LR.D |
| IQ motif and Sec7 domain 1 | IQSEC1 | S512 | 76.7 | 84.8 | R.NS*WDSPAFSNDVIR.K |
| IQ motif and Sec7 domain 1 | IQSEC1 | S87 | | 19.4 | K.LQHS*TSILR.K |
| IQ motif and SEC7 domain-containing protein 1 | IQSEC1 | S802 | -0.4 | 16.3 | R.SALSS*SLRDLSEAGVHH.- |
| IQ motif and SEC7 domain-containing protein 1 | IQSEC1 | S926S940 | 27.5 | 74.8 | R.RSS*AGSLESNVEGSIIS*SPHMR.R |
| IQ motif and SEC7 domain-containing protein 1 | IQSEC1 | S926S941 | 9.9 | 76.6 | R.RSS*AGSLESNVEGSIISS*PHMR.R |
| | IQSEC1 | S929S940 | 37.5 | 54.3 | R.SSAGS*LESNVEGSIIS*SPHMR.R |
| IQ motif and SEC7 domain-containing protein 1 | IQSEC1 | S929S941 | 9.0 | 50.1 | R.SSAGS*LESNVEGSIISS*PHMR.R |
| IQ motif and SEC7 domain-containing protein 1 | IQSEC1 | S929S932S941 | 8.1 | 12.4 | R.RSSAGS*LES*NVEGSIISS*PHMR.R |
| IQ motif and SEC7 domain-containing protein 1 | IQSEC1 | S932S940 | -2.2 | 46.7 | R.SSAGSLES*NVEGSIIS*SPHMR.R |
| IQGAP2 | IQGAP2 | S16 | 39.3 | 53.9 | R.YGS*IVDDER.L |
| IQGAP2 | IQGAP2 | Y14 | 20.1 | 34.0 | R.Y*GSIVDDER.L |
| | MPST | S15 | 7.3 | 37.8 | R.ARS*PSVAAMASPQLCR.A |
| Isoform 2 of Armadillo repeat-containing protein 10 | ARMC10 | S45 | 29.0 | 80.4 | K.S*AEDLTDGSYDDVLNAEQLQK.L |
| Isoform 2 of Chromatin complexes subunit BAP18 | BAP18 | S35 | -0.2 | 26.1 | K.LGELTMQLHPVADS*SPAGAQIK.A |
| Isoform 2 of EF-hand calcium-binding domain-containing protein 4B | CRACR2A | S473 | 120.5 | 60.6 | R.IIS*VEEDPLPQLLDGGFEQPLSK.C |
| Isoform 2 of Eomesodermin homolog | EOMES | T177S187 | 4.5 | 25.2 | R.DNYDSMYT*ASENDRLTPS*PTDSPR.S |
| Isoform 2 of Golgi-specific brefeldin A-resistance guanine nucleotide exchange | GBF1 | S1487 | 11.2 | 16.4 | R.GGQSDDDEDEGVPAS*YHTVSLQLLDLMHTLHTR.A |
| Isoform 2 of Golgi-specific brefeldin A-resistance guanine nucleotide exchange | GBF1 | S1476 | 26.2 | 55.3 | R.GGQS*DDDEDEGVPASYHTVSLQLLDLMHTLHTR.A |
| Isoform 2 of HBS1-like protein | HBS1L | S483 | 24.5 | 50.5 | R.S*PGIDSNIDLSVLIK.N |
| Isoform 2 of Hematopoietic lineage cell-specific protein | HCLS1 | T166S167 | 100.0 | 13.5 | R.RRNT*S*PR.E |
| Isoform 2 of Protein SOGA1 | SOGA1 | S141S142 | 46.7 | 64.6 | R.LLGLELALS*S*DAESAAGGPAGVR.T |
| Isoform 2 of Rho guanine nucleotide exchange factor 18 | ARHGEF1 | S153 | 8.7 | 32.5 | R.S*RSVPVSFYEIR.S |
| Isoform 2 of Rho guanine nucleotide exchange factor 18 | ARHGEF1 | S94 | 97.3 | 75.0 | R.RLS*LDASAVDEEPCLPR.T |
| Isoform 2 of Rho guanine nucleotide exchange factor 18 | ARHGEF1 | S159 | 13.3 | 38.3 | R.SVPVS*FYEIR.S |
| Isoform 2 of Rho guanine nucleotide exchange factor 18 | ARHGEF1 | S263 | 100.0 | 17.8 | R.RLS*CLR.S |
| Isoform 2 of Rho guanine nucleotide exchange factor 18 | ARHGEF1 | S71 | 36.8 | 63.9 | R.DSLFSSLAGS*QDLSR.R |
| | ARHGEF1 | S155 | -0.3 | 28.6 | R.SRS*VPVSFYEIR.S |
| Isoform 2 of Uncharacterized protein C15orf39 | C15orf39 | S496S497 | 100.8 | 76.2 | K.EGARPPS*S*PPMPVIDNVFSLAPYR.D |
| Isoform 3 of Proline-rich protein 12 | PRR12 | S651 | 73.0 | 34.4 | R.TEDEEFLIQHLLQAPS*PPR.T |
| Isoform 3 of UV excision repair protein RAD23 homolog A | RAD23A | Y197 | | 71.6 | R.AVEY*LLTGIPGSPEPEHGSVQESQVSEQPATEAGENPLEFLR.D |
| Isoform 3 of UV excision repair protein RAD23 homolog A | RAD23A | S219 | 3.0 | 26.2 | R.AVEYLLTGIPGSPEPEHGSVQESQVS*EQPATEAGENPLEFLR.D |
| Isoform 3 of UV excision repair protein RAD23 homolog A | RAD23A | S205 | 2.2 | 39.8 | R.AVEYLLTGIPGS*PEPEHGSVQESQVSEQPATEAGENPLEFLR.D |
| Isoform 4 of Interleukin enhancer-binding factor 3 | ILF3 | S476 | | 49.6 | R. DS*SKGEDSAEETEAKPAVVAPAPVVEAVSTPSAAFPSDATAENVK.Q |
| Isoform 4 of Interleukin enhancer-binding factor 3 | ILF3 | S482 | 60.3 | 70.0 | K.GEDS*AEETEAKPAVVAPAPVVEAVSTPSAAFPSDATAENVK.Q |
| Isoform 4 of Interleukin enhancer-binding factor 3 | ILF3 | T486 | 0.3 | 37.4 | K.GEDSAEET*EAKPAVVAPAPVVEAVSTPSAAFPSDATAENVK.Q |
| Isoform 4 of Interleukin enhancer-binding factor 3 | ILF3 | T469S476S477 | | 21.4 | K. VLQDMGLPT*GAEGRDS*S*KGEDSAEETEAKPAVVAPAPVVEAVST |

Peak Area: <-10, -5, -3, 0, 3, 5, >10

%CV: 0, 29, 42, 71, 86, >100

**White dots-** Significant change in peptide abundance at **5** %FDR compared to the timepoint with the minimum peak area for a given PSM

CarT (0m, 2m, 5m) — RajiB (0m, 2m, 5m)

| Protein Name | Gene | Phosphosites | Ascor | MOWSE | Sequence |
|---|---|---|---|---|---|
| **Isoform 4 of Interleukin enhancer-binding factor 3** | ILF3 | T469S476 | | **18.2** | K.VLQDMGLPT*GAEGRDS*SKGEDSAEETEAKPAVVAPAPVVEAVSTE |
| **Isoform 4 of Protein YIF1B** | YIF1B | T285S287 | **100.0** | **11.3** | R.IT*RS*GR.C |
| **Isoform 5 of Mitochondrial fission factor** | MFF | S146 | **100.0** | **24.0** | R.NDS*LPVLR.G |
| **ITK** | ITK | Y512 | **26.0** | **48.1** | R.FVLDDQY*TSSTGTKFPVK.W |
| **IWS1 homolog** | IWS1 | S302S304S313 | **8.9** | **29.6** | R.VSDS*ES*EGPQKGPAS*DS*ETEDASR.H |
| **IWS1 homolog** | IWS1 | S287S289 | **100.0** | **46.8** | R.NQAS*DS*ENEELPKPR.V |
| **IWS1 homolog** | IWS1 | S398S400 | **100.0** | **58.0** | K.AAVLS*DS*EDEEK.A |
| **IWS1 homolog** | IWS1 | S415S420S422 | **100.0** | **53.9** | R.VVS*DADDS*DS*DAVS*DK.S |
| **IWS1 homolog** | IWS1 | S300S302S304 | **17.4** | **38.3** | R.VS*DS*ES*EGPQKGPASDS*ET*EDASR.H |
| | IWS1 | S438S440 | **29.4** | **89.1** | K.TIAS*DS*EEEAGKELSDK.K |
| **IWS1 homolog** | IWS1 | S274S276S287 | **12.5** | **13.1** | R.IS*DS*ESEDPPRNQAS*DS*ENEELPKPR.V |
| **IWS1 homolog** | IWS1 | T67S69S80S82 | **56.4** | **43.6** | K.GHHVT*DS*ENDEPLNLNAS*DS*ESEELHR.Q |
| **IWS1 homolog** | IWS1 | T67S69S80S82 | **100.0** | **22.6** | K.GHHVT*DS*ENDEPLNLNAS*DS*ES*EELHR.Q |
| **IWS1 homolog** | IWS1 | S511S513 | **100.0** | **89.5** | K.EAEDS*DS*DDNIKR.G |
| **IWS1 homolog** | IWS1 | S248S250 | **28.3** | **53.4** | R.IS*DS*ESEDPPR.H |
| **IWS1 homolog** | IWS1 | S248S250S261 | **19.2** | **13.3** | R.IS*DS*ESEDPPRHQAS*DS*ENEELPKPR.I |
| **IWS1 homolog** | IWS1 | S93S95 | **5.7** | **13.3** | R.QKDS*DS*ESEER.A |
| **IWS1 homolog** | IWS1 | S300S304S313 | **12.6** | **17.8** | R.VS*DSES*EGPQKGPAS*DS*ETEDASR.H |
| **IWS1 homolog** | IWS1 | S415S420S422 | **52.3** | **75.8** | R.VVS*DADDS*DS*DAVSDK.S |
| **IWS1 homolog** | IWS1 | S300S302S304 | **5.8** | **11.5** | R.VS*DS*ES*EGPQKGPAS*DS*ETEDASR.H |
| **IWS1 homolog** | IWS1 | S196S198 | **100.0** | **26.2** | R.HQAS*DS*ENEEPPKPR.M |
| **IWS1 homolog** | IWS1 | S261S263 | **100.0** | **49.8** | R.HQAS*DS*ENEELPKPR.I |
| **IWS1 homolog** | IWS1 | S80S82 | **13.8** | **38.7** | K.GHHVTDSENDEPLNLNAS*DS*ESEELHR.Q |
| **IWS1 homolog** | IWS1 | S69S80S82 | **19.1** | **34.5** | K.GHHVTDS*ENDEPLNLNAS*DS*ESEELHR.Q |
| **IWS1 homolog** | IWS1 | T67S80S84 | **3.8** | **14.5** | K.GHHVT*DSENDEPLNLNAS*DSES*EELHR.Q |
| **IWS1 homolog** | IWS1 | S377 | **100.0** | **91.7** | K.MDS*DEDEKEGEEEK.V |
| **IWS1 homolog** | IWS1 | S313S315 | **25.1** | **33.1** | K.GPAS*DS*ETEDASR.H |
| **IWS1 homolog** | IWS1 | S400 | **24.1** | **32.2** | K.AAVLSDS*EDEEK.A |
| **IWS1 homolog** | IWS1 | T67S69S80S84 | **14.4** | **15.0** | K.GHHVT*DS*ENDEPLNLNAS*DSES*EELHR.Q |
| **IWS1 homolog** | IWS1 | S300S302S313 | **18.5** | **22.8** | R.VS*DS*ESEGPQKGPAS*DS*ETEDASR.H |
| **IWS1 homolog** | IWS1 | S276S278S287 | **3.9** | **13.0** | R.ISDS*ES*EDPPRNQAS*DS*ENEELPKPR.V |
| **Janus kinase 3** | JAK3 | S17S20 | **18.2** | **79.2** | R.SCS*LLS*TEAGALHVLLPAR.G |
| **Janus kinase and microtubule interacting protein 1** | JAKMIP1 | S382 | **30.2** | **40.0** | R.HTS*LNDLSLTR.D |
| **JAW1** | LRMP | S73 | **13.2** | **71.0** | R.S*ASPTIEAQGTSPAHDNIAFQDSTSKDK.T |
| **JAW1** | LRMP | S75 | **18.4** | **87.1** | R.SAS*PTIEAQGTSPAHDNIAFQDSTSK.D |
| **JAW1** | LRMP | S75T83 | **27.7** | **47.8** | R.SAS*PTIEAQGT*SPAHDNIAFQDSTSK.D |
| **JAW1** | LRMP | S36 | **5.1** | **29.0** | R.HTS*STDGTITSSDPGLEILNMASCDLDR.N |
| **JAW1** | LRMP | S403 | **10.4** | **27.6** | R.TRKPS*LSEK.K |
| **JAW1** | LRMP | S388S391 | **52.2** | **67.7** | K.TKDDS*EPS*GEETVER.T |
| **JAW1** | LRMP | S75S84 | **28.9** | **49.6** | R.SAS*PTIEAQGTS*PAHDNIAFQDSTSK.D |
| **JAW1** | LRMP | T319 | **26.4** | **29.0** | R.RVT*IASLPR.N |
| **JAW1** | LRMP | S322 | **48.6** | **55.2** | R.VTIAS*LPR.N |
| **JAW1** | LRMP | T41 | **12.4** | **56.4** | R.HTSSTDGT*ITSSDPGLEILNMASCDLDR.N |
| **JAW1** | LRMP | T77 | **5.0** | **23.7** | R.SASPT*IEAQGTSPAHDNIAFQDSTSKDK.T |
| **JAW1** | LRMP | S73T83 | **5.4** | **35.6** | R.S*ASPTIEAQGT*SPAHDNIAFQDSTSK.D |
| **Jouberin** | AHI1 | S267 | | **23.2** | K.KES*SVR.S |
| **Jumonji domain containing 1B** | KDM3B | S727S743 | **8.3** | **21.8** | R.SSS*PTSSLTQPIEMPTLSS*SPTEERPTVGPGQQDNPLLK.T |
| **Jumonji domain containing 1B** | KDM3B | S727S742 | **5.9** | **22.9** | R.SSS*PTSSLTQPIEMPTLS*SSPTEERPTVGPGQQDNPLLK.T |
| **Jumonji domain containing 1B** | KDM3B | S730S743 | **7.1** | **16.5** | R.SSSPTS*SLTQPIEMPTLSS*SPTEERPTVGPGQQDNPLLK.T |
| **Jumonji domain containing 1B** | KDM3B | S731S744 | **9.8** | **19.3** | R.SSSPTSS*LTQPIEMPTLSSS*PTEERPTVGPGQQDNPLLK.T |
| **Jumonji domain containing 1B** | KDM3B | S727S744 | **7.6** | **14.7** | R.SSS*PTSSLTQPIEMPTLSSS*PTEERPTVGPGQQDNPLLK.T |
| **Jumonji domain containing 1B** | KDM3B | S725S726 | | **14.9** | R.S*S*SPTSSLTQPIEMPTLSSSPTEERPTVGPGQQDNPLLK.T |
| **Jumonji domain containing 1B** | KDM3B | S730S744 | **7.5** | **18.5** | R.SSSPTS*SLTQPIEMPTLSSS*PTEERPTVGPGQQDNPLLK.T |
| **Jumonji domain containing 1B** | KDM3B | S725S744 | **7.6** | **10.9** | R.S*SSPTSSLTQPIEM#PTLSSS*PTEERPTVGPGQQDNPLLK.T |
| **Jumonji domain containing 1B** | KDM3B | S727T746 | **5.1** | **13.7** | R.SSS*PTSSLTQPIEMPTLSSSPT*EERPTVGPGQQDNPLLK.T |
| **Jumonji domain containing 1B** | KDM3B | S730S742 | **8.2** | **22.4** | R.SSSPTS*SLTQPIEMPTLS*SSPTEERPTVGPGQQDNPLLK.T |
| **Jumonji domain containing 1B** | KDM3B | T733S742 | **6.3** | **11.2** | R.SSSPTSSLT*QPIEMPTLS*SSPTEERPTVGPGQQDNPLLK.T |
| **Jumonji domain containing 2B** | KDM4B | S632S633 | **44.2** | **14.7** | K.QEAS*S*DEEASPFSGEEDVSDPDALRPLLSLQWK.N |
| **JUN-D** | JUND | S251S259 | **22.8** | **10.8** | K.DEPQTVPDVPS*FGESPPLS*PIDMDTQER.I |
| **KAISO** | ZBTB33 | T208 | | **11.6** | K.ET*LPSNNTVAQVQSNPGPVAISDVAPSASNNSPPLTNITPTQK.L |
| **Kanadaptin** | SLC4A1AP | S466 | **26.0** | **152.3** | K.NWEDEDFYDS*DDDTFLDR.T |
| **Kanadaptin** | SLC4A1AP | S258 | **118.6** | **115.5** | K.MLGEDS*DEEEEMDTSER.K |

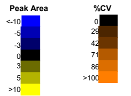


**White dots-** Significant change in peptide abundance at **5** %FDR compared to the timepoint with the minimum peak area for a given PSM

| Protein Name | Gene | Phosphosites | Ascor | MOWSE | Sequence |
|---|---|---|---|---|---|
| **Kanadaptin** | SLC4A1AP | Y464 | 10.9 | 96.5 | K.NWEDEDFY*DSDDDTFLDR.T |
| | CEP170 | S356S359 | 70.4 | 52.7 | K.S*IKS*DVPVYLK.R |
| **KARP 1 binding protein** | CEP170 | S1165 | 13.1 | 20.1 | R.S*DSEATISR.S |
| **KARP 1 binding protein** | CEP170 | S359 | 11.8 | 36.2 | K.SIKS*DVPVYLKR.L |
| **KARP 1 binding protein** | PLIN5 | S1160 | 13.9 | 13.6 | R.LGS*LSAR.S |
| **KARP 1 binding protein** | CEP170 | S356 | 4.2 | 12.7 | K.S*IKSDVPVYLKR.L |
| **KARP 1 binding protein** | CEP170 | S838 | 9.5 | 15.0 | R.QGS*FTIEKPSPNIPIELIPHINK.Q |
| **KARP 1 binding protein** | CEP170 | S1167 | -0.8 | 26.4 | R.SDS*EATISR.S |
| **Karyopherin alpha3** | KPNA3 | S60 | 51.5 | 75.1 | R.NVPQEESLEDS*DVDADFK.A |
| **Karyopherin alpha3** | KPNA3 | S56 | 9.4 | 43.6 | R.NVPQEES*LEDSDVDADFK.A |
| **Karyopherin beta 3** | IPO5 | S670 | 2.5 | 26.7 | K.TASIKPEVALLDTQDMENMS*DDDGWEFVNLGDQQSFGIK.T |
| **Kelch domain containing 4** | KLHDC4 | S413S418 | 76.0 | 117.2 | R.S*EDEDS*LEEAGSPAPGPCPR.S |
| **Kelch domain containing 4** | KLHDC4 | S413 | 11.6 | 45.7 | R.S*EDEDSLEEAGSPAPGPCPR.S |
| **Kelch domain containing 4** | KLHDC4 | S418 | -0.5 | 58.8 | R.SEDEDS*LEEAGSPAPGPCPR.S |
| **Keratin 13** | KRT13 | T319 | 12.3 | 11.4 | K.T*EITELR.R |
| **KH type splicing regulatory protein** | KHSRP | S181 | 25.4 | 41.1 | K.VQIS*PDSGGLPER.S |
| **KIAA0056** | NCAPD3 | S1382 | 18.4 | 36.1 | R.SLGVLPFTLNS*GSPEK.T |
| **KIAA0056** | NCAPD3 | S1384 | 16.1 | 45.5 | R.SLGVLPFTLNSGS*PEK.T |
| **KIAA0082** | CMTR1 | T48S49S55 | 6.0 | 58.7 | K.ASTT*S*LSGSDS*ETEGK.Q |
| **KIAA0082** | CMTR1 | S51S53S55 | 7.1 | 63.1 | K.ASTTSLS*GS*DS*ETEGK.Q |
| **KIAA0082** | CMTR1 | T47T48S55 | 9.1 | 47.8 | K.AST*T*SLSGSDS*ETEGK.Q |
| **KIAA0146** | SPIDR | S132 | 100.0 | 64.3 | R.DELQFIDWEIDS*DR.A |
| **KIAA0153 protein** | TTLL12 | S16 | 9.1 | 99.9 | R.SS*PGQTPEEGAQALAEFAALHGPALR.A |
| **KIAA0153 protein** | TTLL12 | S15 | 10.5 | 66.5 | R.S*SPGQTPEEGAQALAEFAALHGPALR.A |
| **KIAA0157** | ABRAXAS | S368S372Y377 | 9.6 | 46.4 | R.AAGDS*GEDS*DDSDY*ENLIDPTEPSNSEYSHSK.D |
| **KIAA0157** | ABRAXAS | S368S372S375 | 15.2 | 55.5 | R.AAGDS*GEDS*DDS*DYENLIDPTEPSNSEYSHSK.D |
| **KIAA0157** | ABRAXAS | S368S375Y377 | 4.6 | 46.6 | R.AAGDS*GEDSDDS*DY*ENLIDPTEPSNSEYSHSK.D |
| **KIAA0179** | RRP1B | S732S735 | 28.4 | 38.5 | K.TPTSS*PAS*SPLVAK.K |
| **KIAA0179** | RRP1B | T728S732 | 14.5 | 28.4 | R.VAFDPEQKPLHGVLKT*PTSS*PASSPLVAK.K |
| **KIAA0179** | RRP1B | S731S732S736 | 14.1 | 20.4 | R.VAFDPEQKPLHGVLKTPTS*S*PASS*PLVAK.K |
| **KIAA0179** | RRP1B | T728T730 | 7.1 | 18.3 | R.VAFDPEQKPLHGVLKT*PT*SSPASSPLVAK.K |
| **KIAA0179** | RRP1B | T728S731S736 | 14.6 | 16.3 | R.VAFDPEQKPLHGVLKT*PTS*SPASS*PLVAK.K |
| **KIAA0179** | RRP1B | S731S735 | 35.5 | 36.6 | K.TPTS*SPAS*SPLVAK.K |
| **KIAA0179** | RRP1B | T728S732S736 | 10.4 | 21.2 | R.VAFDPEQKPLHGVLKT*PTSS*PASS*PLVAK.K |
| **KIAA0179** | RRP1B | T728T730S736 | 11.9 | 23.7 | R.VAFDPEQKPLHGVLKT*PT*SSPASS*PLVAK.K |
| **KIAA0179** | RRP1B | T728S732S735 | 11.4 | 35.3 | R.VAFDPEQKPLHGVLKT*PTSS*PAS*SPLVAK.K |
| **KIAA0179** | RRP1B | T728S731 | 18.2 | 29.6 | R.VAFDPEQKPLHGVLKT*PTS*SPASSPLVAK.K |
| **KIAA0182** | GSE1 | S10 | 44.9 | 59.5 | K.S*PSLGMLSTATR.T |
| **KIAA0217 protein** | LARP4B | S516T518 | 14.5 | 24.7 | K.FTSSQTQS*PT*PPKPPSPSFELGLSSFPPLPGAAGNLK.T |
| **KIAA0217 protein** | LARP4B | S511S512 | 5.7 | 15.4 | K.FTS*S*QTQSPTPPKPPSPSFELGLSSFPPLPGAAGNLK.T |
| **KIAA0217 protein** | LARP4B | T514T518S526 | 7.8 | 37.7 | K.FTSSQT*QSPT*PPKPPSPS*FELGLSSFPPLPGAAGNLK.T |
| **KIAA0217 protein** | LARP4B | T510S511S516 | 5.9 | 14.2 | K.FT*S*SQTQS*PTPPKPPSPSFELGLSSFPPLPGAAGNLK.T |
| **KIAA0217 protein** | LARP4B | S498 | 34.4 | 37.4 | R.KNS*FGYR.K |
| **KIAA0217 protein** | LARP4B | T510S511 | 8.7 | 18.2 | K.FT*S*SQTQSPTPPKPPSPSFELGLSSFPPLPGAAGNLK.T |
| **KIAA0217 protein** | LARP4B | S601 | 11.4 | 30.0 | R.SPS*PAHLPDDPK.V |
| **KIAA0217 protein** | LARP4B | Y501 | 13.8 | 25.6 | R.KNSFGY*R.K |
| **KIAA0217 protein** | LARP4B | T518S532 | 2.9 | 17.6 | K.FTSSQTQSPT*PPKPPSPSFELGLS*SFPPLPGAAGNLK.T |
| **KIAA0217 protein** | LARP4B | S511T518S526 | 12.4 | 25.7 | K.FTS*SQTQSPT*PPKPPSPS*FELGLSSFPPLPGAAGNLK.T |
| **KIAA0217 protein** | LARP4B | S516S524S526 | 6.0 | 16.5 | K.FTSSQTQS*PTPPKPPS*PS*FELGLSSFPPLPGAAGNLK.T |
| **KIAA0217 protein** | LARP4B | S516T518S524 | 19.3 | 38.4 | K.FTSSQTQS*PT*PPKPPS*PSFELGLSSFPPLPGAAGNLK.T |
| **KIAA0217 protein** | LARP4B | T518S524 | 4.7 | 16.9 | K.FTSSQTQSPT*PPKPPS*PSFELGLSSFPPLPGAAGNLK.T |
| **KIAA0217 protein** | LARP4B | T514T518S524 | 8.5 | 16.9 | K.FTSSQT*QSPT*PPKPPS*PSFELGLSSFPPLPGAAGNLK.T |
| **KIAA0217 protein** | LARP4B | S568 | -0.5 | 14.4 | R.TLS*ADASVNTLPVVVSR.E |
| **KIAA0217 protein** | LARP4B | S516S524 | 7.0 | 12.4 | K.FTSSQTQS*PTPPKPPS*PSFELGLSSFPPLPGAAGNLK.T |
| **KIAA0284 protein** | CEP170B | S221 | 100.0 | 13.1 | K.FS*LRQR.R |
| **KIAA0409 protein** | RRP8 | S104S106 | 100.0 | 20.7 | K.QGPPCS*DS*EEEVER.K |
| **KIAA0409 protein** | RRP8 | S62S64 | 17.3 | 55.4 | R.ALEAASLSQHPPSLCIS*DS*EEEEEER.K |
| **KIAA0409 protein** | RRP8 | S58S62 | 8.3 | 11.6 | R.ALEAASLSQHPPS*LCIS*DSEEEEEER.K |
| **KIAA0409 protein** | RRP8 | S58S64 | 4.6 | 20.8 | R.ALEAASLSQHPPS*LCISDS*EEEEEER.K |
| **KIAA0433 protein** | PPIP5K2 | S38 | 100.0 | 59.1 | R.HFFHHADEDDEEEDDS*PPER.Q |
| **KIAA0433 protein** | PPIP5K2 | S492S493 | -0.4 | 22.1 | K.TS*S*EEEDSRR.E |
| **KIAA0433 protein** | PPIP5K2 | S1151 | -0.2 | 26.2 | R.SS*PIMR.K |

Peak Area: <-10, -5, -3, 0, 3, 5, >10

%CV: 0, 29, 42, 71, 86, >100

**White dots-** Significant change in peptide abundance at **5** %FDR compared to the timepoint with the minimum peak area for a given PSM

| Protein Name | Gene | Phosphosites | CarT 0m | CarT 2m | CarT 5m | RajiB 0m | RajiB 2m | RajiB 5m | Ascor | MOWSE | Sequence |
|---|---|---|---|---|---|---|---|---|---|---|---|
| **KIAA0460** | RPRD2 | S899S902 | | | | | | | | **41.9** | R.GNEPGS*DRS*PSPSKNDSFFTPDSNHNSLSQSTTGHLSLPQK.Q |
| **KIAA0460** | RPRD2 | S348 | | | | | | | **71.6** | **112.6** | R.DVEDMELS*DVEDDGSK.I |
| **KIAA0460** | RPRD2 | S459 | | | | | | | **15.2** | **20.5** | K.NTGVSPASRPSPGTPTS*PSNLTSGLK.T |
| **KIAA0460** | RPRD2 | S902S904 | | | | | | | **12.3** | **24.2** | R.GNEPGSDRS*PS*PSKNDSFFTPDSNHNSLSQSTTGHLSLPQK.Q |
| **KIAA0460** | RPRD2 | S902S906 | | | | | | | **17.4** | **36.7** | R.GNEPGSDRS*PSPS*KNDSFFTPDSNHNSLSQSTTGHLSLPQK.Q |
| **KIAA0460** | RPRD2 | T458 | | | | | | | **4.0** | **26.2** | K.NTGVSPASRPSPGTPT*SPSNLTSGLK.T |
| **KIAA0460** | RPRD2 | S330T332 | | | | | | | **53.2** | **20.0** | K.S*AT*PEPVTDNR.D |
| **KIAA0460** | RPRD2 | S902S910 | | | | | | | **15.7** | **14.6** | R.GNEPGSDRS*PSPSKNDS*FFTPDSNHNSLSQSTTGHLSLPQK.Q |
| **KIAA0467** | SZT2 | S920S926 | | | | | | | **23.0** | **57.6** | R.AEGIEGETLTAS*PQAPGS*PEDSEGVPLISLPR.V |
| **KIAA0515 protein** | PRRC2B | S388 | | | | | | | **100.0** | **36.1** | K.LKFS*DDEEEEEVVK.D |
| **KIAA0515 protein** | PRRC2B | T225 | | | | | | | **23.9** | **91.5** | R.HIISATSLST*SPTELGSR.N |
| **KIAA0515 protein** | PRRC2B | S226 | | | | | | | **17.2** | **36.0** | R.HIISATSLSTS*PTELGSR.N |
| **KIAA0515 protein** | PRRC2B | S224 | | | | | | | **0.5** | **38.1** | R.HIISATSLS*TSPTELGSR.N |
| **KIAA0528** | C2CD5 | S260 | | | | | | | **80.2** | **40.7** | K.LSSPAAFLPACNS*PSK.E |
| **KIAA0528** | C2CD5 | S817 | | | | | | | **12.2** | **42.7** | R.AS*TDNEELLQFPLELCSDSLPSHPFPPAK.A |
| **KIAA0556** | KIAA0556 | S753S758 | | | | | | | **5.8** | **10.8** | K.TPS*WLQPS*PTGK.D |
| **KIAA0592** | WASHC2C | S539 | | | | | | | **75.8** | **173.2** | K.GLFS*DEEDSEDLFSSQSASNLK.G |
| **KIAA0592** | WASHC2C | S544 | | | | | | | **5.1** | **33.2** | K.GLFSDEEDS*EDLFSSQSASNLK.G |
| **KIAA0592** | WASHC2A | S695S702 | | | | | | | **11.3** | **32.2** | R.VSLLFEDDVDS*GGSLFGS*PPTSVPPATK.K |
| **KIAA0592** | WASHC2A | S284S288 | | | | | | | **11.1** | **16.4** | R.S*RPTS*FADELAAR.I |
| **KIAA0592** | WASHC2A | S352 | | | | | | | **23.9** | **62.8** | K.LTDEDFS*PFGSGGGLFSGGK.G |
| **KIAA0592** | WASHC2A | S284T287 | | | | | | | **8.6** | **12.1** | R.S*RPT*SFADELAAR.I |
| **KIAA0592** | WASHC2A | S686S695 | | | | | | | **2.7** | **16.9** | R.VS*LLFEDDVDS*GGSLFGSPPTSVPPATK.K |
| **KIAA0592** | WASHC2A | S695S698S702 | | | | | | | **13.7** | **12.0** | R.VSLLFEDDVDS*GGS*LFGS*PPTSVPPATK.K |
| **KIAA0592** | WASHC2A | S698S702 | | | | | | | **2.4** | **34.5** | R.VSLLFEDDVDSGGS*LFGS*PPTSVPPATK.K |
| **KIAA0652 gene product** | ATG13 | S324 | | | | | | | **86.0** | **45.5** | R.AS*PHDVLETIFVR.K |
| **KIAA0683** | TELO2 | T632 | | | | | | | **5.6** | **48.8** | R.T*PQPGSPSPNTPCLPEAAVSQPGSAVASDWR.V |
| **KIAA0683** | TELO2 | S637 | | | | | | | **5.0** | **57.4** | R.TPQPGS*PSPNTPCLPEAAVSQPGSAVASDWR.V |
| **KIAA0683** | TELO2 | S639 | | | | | | | **2.3** | **55.1** | R.TPQPGSPS*PNTPCLPEAAVSQPGSAVASDWR.V |
| **KIAA0685** | PPP6R2 | S766 | | | | | | | **13.8** | **28.6** | K.AFS*PASPCAWNVCVTR.K |
| **KIAA0692** | ANKLE2 | S662 | | | | | | | **19.4** | **19.1** | R.NNS*PPTVGAFGHTR.C |
| **KIAA0701 protein** | UHRF1BP | S889S891 | | | | | | | **7.6** | **37.4** | K.SPVSES*VS*PVVPDYLPTENGDFLSSK.R |
| **KIAA0701 protein** | UHRF1BP | S887S889 | | | | | | | **23.8** | **46.0** | K.SPVS*ES*VSPVVPDYLPTENGDFLSSK.R |
| **KIAA0701 protein** | UHRF1BP | S935 | | | | | | | **7.6** | **27.7** | R.SMS*VDLSHIPLKDPLLFK.S |
| **KIAA0748** | TESPA1 | S454 | | | | | | | **61.4** | **77.6** | K.S*LDLSITQQK.W |
| **KIAA0776** | UFL1 | S458 | | | | | | | **69.6** | **98.6** | R.KDDDS*DDESQSSHTGK.K |
| **KIAA0776** | UFL1 | S790 | | | | | | | **12.1** | **23.7** | R.KSS*VTEE.- |
| **KIAA0912 protein** | CEP152 | S717 | | | | | | | **4.0** | **16.5** | R.THLQLRS*ELDK.L |
| **KIAA0912 protein** | CEP152 | S1405 | | | | | | | **100.0** | **47.9** | R.NVS*PEFVPCEGEGGFGLHK.K |
| **KIAA0999 protein** | SIK3 | S568 | | | | | | | **60.3** | **51.0** | R.RFS*DGAASIQAFK.A |
| **KIAA0999 protein** | SIK3 | T163 | | | | | | | **37.8** | **89.6** | K.T*WCGSPPYAAPELFEGK.E |
| **KIAA0999 protein** | SIK3 | S167 | | | | | | | **-0.5** | **47.9** | K.TWCGS*PPYAAPELFEGK.E |
| **KIAA1002 protein** | R3HDM2 | S10 | | | | | | | **10.2** | **66.6** | K.ASS*FSGISILTR.G |
| **KIAA1002 protein** | R3HDM2 | S9 | | | | | | | **12.2** | **74.3** | K.AS*SFSGISILTR.G |
| **KIAA1115 protein** | PPP6R1 | S540S541 | | | | | | | **12.7** | **56.0** | K.NMVDLVNTHHLHSS*S*DDEDDRLK.E |
| **KIAA1115 protein** | PPP6R1 | S539S540 | | | | | | | **11.0** | **38.0** | K.NMVDLVNTHHLHS*S*SDDEDDRLK.E |
| **KIAA1115 protein** | PPP6R1 | S541 | | | | | | | **11.3** | **41.8** | K.NMVDLVNTHHLHSSS*DDEDDRLK.E |
| **KIAA1115 protein** | PPP6R1 | S539S540S541 | | | | | | | **15.0** | **30.3** | K.NMVDLVNTHHLHS*S*S*DDEDDRLK.E |
| **KIAA1115 protein** | PPP6R1 | T735 | | | | | | | **11.0** | **22.1** | R.GGATPLSYPSPGPQPPGPSWTATFDPVPTDAPT*SPR.V |
| **KIAA1115 protein** | PPP6R1 | S712T735 | | | | | | | **18.0** | **30.5** | R.GGATPLSYPS*PGPQPPGPSWTATFDPVPTDAPT*SPR.V |
| **KIAA1115 protein** | PPP6R1 | S540 | | | | | | | **13.9** | **48.2** | K.NMVDLVNTHHLHSS*SDDEDDRLK.E |
| **KIAA1115 protein** | PPP6R1 | S677T678S680 | | | | | | | **6.2** | **68.3** | R.SGGS*T*DS*EDEEEEDEEEEEDEEGIGCAAR.G |
| **KIAA1115 protein** | PPP6R1 | T534S539S541 | | | | | | | **3.6** | **12.5** | K.NMVDLVNT*HHLHS*SS*DDEDDRLK.E |
| **KIAA1115 protein** | PPP6R1 | S674S677T678 | | | | | | | **4.3** | **85.8** | R.S*GGS*T*DSEDEEEEDEEEEEDEEGIGCAAR.G |
| **KIAA1115 protein** | PPP6R1 | T534 | | | | | | | | **16.7** | K.NM#VDLVNT*HHLHSSSDDEDDRLK.E |
| **KIAA1115 protein** | PPP6R1 | T534S540S541 | | | | | | | **8.3** | **22.7** | K.NMVDLVNT*HHLHSS*S*DDEDDR.L |
| **KIAA1115 protein** | PPP6R1 | T731 | | | | | | | **4.2** | **22.3** | R.GGATPLSYPSPGPQPPGPSWTATFDPVPT*DAPTSPR.V |
| **KIAA1115 protein** | PPP6R1 | S736 | | | | | | | **8.0** | **38.9** | R.GGATPLSYPSPGPQPPGPSWTATFDPVPTDAPTS*PR.V |
| **KIAA1115 protein** | PPP6R1 | S539S541 | | | | | | | **7.9** | **16.9** | K.NMVDLVNTHHLHS*SS*DDEDDRLK.E |
| **KIAA1115 protein** | PPP6R1 | T706 | | | | | | | | **22.5** | R.GGAT*PLSYPSPGPQPPGPSWTATFDPVPTDAPTSPR.V |
| **KIAA1115 protein** | PPP6R1 | T723T735 | | | | | | | **23.8** | **15.9** | R.GGATPLSYPSPGPQPPGPSWT*ATFDPVPTDAPT*SPR.V |
| **KIAA1115 protein** | PPP6R1 | T534S539 | | | | | | | **0.1** | **15.4** | K.NMVDLVNT*HHLHS*SSDDEDDRLK.E |

Peak Area: <-10, -5, -3, 0, 3, 5, >10

%CV: 0, 29, 42, 71, 86, >100

**White dots-** Significant change in peptide abundance at **5** %FDR compared to the timepoint with the minimum peak area for a given PSM

| Protein Name | Gene | Phosphosites | CarT 0m | CarT 2m | CarT 5m | RajiB 0m | RajiB 2m | RajiB 5m | Ascor | MOWSE | Sequence |
|---|---|---|---|---|---|---|---|---|---|---|---|
| **KIAA1115 protein** | PPP6R1 | T534S539S540 | | | | | | | 5.9 | 16.2 | K.NMVDLVNT*HHLHS*S*SDDEDDRLK.E |
| **KIAA1115 protein** | PPP6R1 | S887 | | | | | | | 16.7 | 18.0 | K.SPEPLGLPQSQSAQALTPPPIPNGSAPEGPAS*PGSQ.- |
| **KIAA1115 protein** | PPP6R1 | S645S648 | | | | | | | 75.2 | 101.4 | R.IQQFDDDEEEEDEEEAQGS*GES*DGEDGAWQGSQLAR.G |
| **KIAA1115 protein** | PPP6R1 | S712 | | | | | | | 12.2 | 31.1 | R.GGATPLSYPS*PGPQPPGPSWTATFDPVPTDAPTSPR.V |
| **KIAA1115 protein** | PPP6R1 | S712S736 | | | | | | | 6.5 | 22.1 | R.GGATPLSYPS*PGPQPPGPSWTATFDPVPTDAPTS*PR.V |
| **KIAA1115 protein** | PPP6R1 | S539 | | | | | | | -0.3 | 22.0 | K.NMVDLVNTHHLHS*SSDDEDDRLK.E |
| **KIAA1143 protein** | KIAA1143 | S50 | | | | | | | 100.0 | 22.6 | R.IQPQPPDEDGDHS*DKEDEQPQVVVLK.K |
| **KIAA1211 protein** | KIAA1211 | T968S975 | | | | | | | 5.0 | 25.6 | K.MPLAQKPALAPKPT*SQTPPAS*PLSK.L |
| **KIAA1211 protein** | KIAA1211 | T968S969 | | | | | | | 3.1 | 21.1 | K.MPLAQKPALAPKPT*S*QTPPASPLSK.L |
| **KIAA1211 protein** | KIAA1211 | T971S975 | | | | | | | 9.4 | 20.5 | K.MPLAQKPALAPKPTSQT*PPAS*PLSK.L |
| **KIAA1267** | KANSL1 | S268 | | | | | | | 31.8 | 100.3 | K.S*PLSSILFSALDSDTR.I |
| **KIAA1267** | KANSL1 | S271 | | | | | | | -2.8 | 38.3 | K.SPLS*SILFSALDSDTR.I |
| **KIAA1271 protein** | MAVS | S222 | | | | | | | 67.6 | 24.7 | R.GPVS*PSVSFQPLAR.S |
| **KIAA1370** | FAM214A | T32 | | | | | | | 49.3 | 10.8 | R.T*PECSVKGR.T |
| **KIAA1429** | VIRMA | S1579 | | | | | | | 9.8 | 23.9 | R.SFLSEPSS*PGR.T |
| **KIAA1429** | VIRMA | S1578 | | | | | | | 42.4 | 16.6 | R.SFLSEPS*SPGR.T |
| **KIAA1429** | VIRMA | T184 | | | | | | | 25.5 | 14.4 | R.T*PPGPPPPDDDEDDPVPLPVSGDK.E |
| **KIAA1432** | RIC1 | T913S916 | | | | | | | 7.0 | 43.1 | K.AIGSGESET*PPS*TPTAQEPSSSGGFEFFR.N |
| **KIAA1462 protein** | JCAD | S948S949S951 | | | | | | | 4.9 | 19.5 | K.EVS*S*VS*RM#RVLSFR.N |
| **KIAA1467** | FAM234B | S16T26S33 | | | | | | | 16.9 | 41.5 | K.S*PDLGEYDPLT*QADSDES*EDDLVLNLQK.N |
| **KIAA1467** | FAM234B | S16T26S30 | | | | | | | 8.4 | 16.3 | K.S*PDLGEYDPLT*QADS*DESEDDLVLNLQK.N |
| **KIAA1467** | FAM234B | T26S30 | | | | | | | 7.6 | 13.6 | K.SPDLGEYDPLT*QADS*DESEDDLVLNLQK.N |
| **KIAA1467** | FAM234B | S30S33 | | | | | | | 23.1 | 68.3 | K.SPDLGEYDPLTQADS*DES*EDDLVLNLQK.N |
| **KIAA1468** | RELCH | S45S51S54 | | | | | | | | 33.6 | R. [illegible] |
| **KIAA1542 Protein** | PHRF1 | S973 | | | | | | | 51.5 | 61.3 | R.TVTCVTVVEPEAPPS*PDVLQAATHR.V |
| **KIAA1542 Protein** | PHRF1 | S1032S1034 | | | | | | | 37.7 | 38.0 | R.S*AS*PSVGEERPR.R |
| **KIAA1542 Protein** | PHRF1 | S1114S1116 | | | | | | | 100.0 | 15.8 | R.RS*AS*RPR.G |
| **KIAA1542 Protein** | PHRF1 | S1032S1036 | | | | | | | 11.8 | 37.6 | R.S*ASPS*VGEERPR.R |
| **KIAA1542 Protein** | PHRF1 | S1359S1371 | | | | | | | 29.4 | 13.3 | K.AEAPS*SPDVAPAGKEDS*PSASGR.V |
| **KIAA1542 Protein** | PHRF1 | S1034S1036 | | | | | | | 6.7 | 39.2 | R.SAS*PS*VGEERPR.R |
| **KIAA1542 Protein** | PHRF1 | S101 | | | | | | | 20.4 | 29.2 | K.LEAAGSFNS*DDDAESCPICLNAFR.D |
| **KIAA1542 Protein** | PHRF1 | S98 | | | | | | | 4.3 | 19.6 | K.LEAAGS*FNSDDDAESCPICLNAFR.D |
| **KIAA1602** | NCKAP5L | S571S577 | | | | | | | 10.6 | 16.5 | R.GPS*PEPPPS*PLQVPTYPQLTLEVPQAPEVLR.S |
| **KIAA1604 protein** | CWC22 | S91S93 | | | | | | | 25.1 | 13.8 | R.SRKS*PS*PGR.R |
| **KIAA1604 protein** | CWC22 | S831 | | | | | | | 5.2 | 14.5 | R.RNSFS*ENEK.H |
| **KIAA1604 protein** | CWC22 | S829 | | | | | | | 13.4 | 23.0 | R.RNS*FSENEK.H |
| **KIAA1704 protein** | GPALPP1 | S105 | | | | | | | 100.0 | 47.3 | K.QDDS*PPRPIIGPALPPGFIK.S |
| | GPALPP1 | T138S140S141 | | | | | | | 100.0 | 30.1 | R.DDPGQQET*DS*S*EDEDIIGPMPAK.G |
| **KIAA1706 protein** | EEPD1 | S173 | | | | | | | 100.0 | 50.6 | R.S*VEDLVR.M |
| **KIAA1706 protein** | EEPD1 | S25S31 | | | | | | | 100.0 | 63.8 | R.KFS*AACNFS*NILVNQER.L |
| **KIAA1826 protein** | MSANTD4 | S152 | | | | | | | 78.4 | 37.9 | R.DPQS*PEFEIEEEEEMLSSVIPDSR.R |
| **KIAA1836 protein** | CC2D1B | T824 | | | | | | | 100.0 | 10.8 | R.KPT*GGKLEVK.V |
| **KIAA1949** | PPP1R18 | S224 | | | | | | | 67.0 | 50.0 | R.LS*PGESAYQK.L |
| **KIDINS220** | KIDINS220 | S1411 | | | | | | | -0.4 | 42.2 | R.SS*PHSTYYMGQSSSGGSIHSNLEQEK.G |
| **Kinase suppressor of ras 1** | KSR1 | T133T136 | | | | | | | 50.0 | 44.9 | R.ALHSFIT*PPT*TPQLR.R |
| **Kinase suppressor of ras 1** | KSR1 | T133T137 | | | | | | | 11.6 | 31.9 | R.ALHSFIT*PPTT*PQLR.R |
| **Kindlin 2** | FERMT2 | T188T190T192 | | | | | | | | 24.8 | K. [illegible] |
| **Kinectin** | KTN1 | S75 | | | | | | | 14.4 | 20.0 | K.EIQNGNLHES*DSESVPR.D |
| **Kinesin 2** | KLC1 | S521 | | | | | | | 18.7 | 53.9 | R.S*RESLNVDVVK.Y |
| **Kinesin 2** | KLC1 | S521S524 | | | | | | | 100.0 | 37.5 | R.S*RES*LNVDVVK.Y |
| **Kinesin 2** | KLC1 | S524 | | | | | | | 1.3 | 31.8 | R.SRES*LNVDVVK.Y |
| **Kinesin family member 13B** | KIF13B | S1778 | | | | | | | 12.1 | 17.3 | R.RS*TGLR.L |
| **Kinesin family member 1B** | KIF1B | S1612 | | | | | | | 15.2 | 52.3 | R.AS*SPCPEFEQFQIVPAVETPYLAR.A |
| **Kinesin family member 1B** | KIF1B | S1613 | | | | | | | 16.9 | 24.8 | R.ASS*PCPEFEQFQIVPAVETPYLAR.A |
| **Kinesin family member 1C** | KIF1C | S1033 | | | | | | | 100.0 | 34.3 | R.RNS*LDGGGR.S |
| **Kinesin family member 1C** | KIF1C | S674S676 | | | | | | | 37.5 | 65.6 | R.LYADS*DS*GDDSDKR.S |
| **Kinesin family member 21B** | KIF21B | S1167 | | | | | | | 31.4 | 12.3 | K.S*LASLVEIK.E |
| **Kinesin family member 23** | KIF23 | S684 | | | | | | | 5.1 | 73.4 | R.SVS*PSPVPLLFQPDQNAPPIR.L |
| **Kinesin family member 23** | KIF23 | S686 | | | | | | | -0.3 | 44.6 | R.SVSPS*PVPLLFQPDQNAPPIR.L |
| **Kinesin family member 23** | KIF23 | S684S686 | | | | | | | -0.4 | 31.4 | R.SVS*PS*PVPLLFQPDQNAPPIR.L |
| **Kinesin family member 23** | KIF23 | S682S684 | | | | | | | 6.2 | 47.2 | R.S*VS*PSPVPLLFQPDQNAPPIR.L |
| **Kinesin family member 3A** | KIF3A | T692 | | | | | | | 3.3 | 16.1 | R.SAKPET*VIDSLLQ.- |

Peak Area: <-10, -5, -3, 0, 3, 5, >10

%CV: 0, 29, 42, 71, 86, >100

**White dots-** Significant change in peptide abundance at **5** %FDR compared to the timepoint with the minimum peak area for a given PSM

| Protein Name | Gene | Phosphosites | CarT 0m | CarT 2m | CarT 5m | RajiB 0m | RajiB 2m | RajiB 5m | Ascor | MOWSE | Sequence |
|---|---|---|---|---|---|---|---|---|---|---|---|
| Kinesin family member 5B | KIF5B | S933 | | | | | | | 9.1 | 15.1 | R.GHSAQIAKPIRPGQHPAAS*PTHPSAIR.G |
| Kinesin heavy chain 2 | KIF2A | T78 | | | | | | | 24.1 | 13.0 | K.EIDLESIFSLNPDLVPDEEIEPSPET*PPPPASSAK.V |
| Kinesin heavy chain 2 | KIF2A | S75 | | | | | | | 8.3 | 21.9 | K.EIDLESIFSLNPDLVPDEEIEPS*PETPPPPASSAK.V |
| Kinesin light chain 1 | KLC1 | S590 | | | | | | | 9.8 | 59.4 | R.AS*SLNVLNVGGK.A |
| Kinesin light chain 1 | KLC1 | S591 | | | | | | | 25.6 | 75.4 | R.ASS*LNVLNVGGK.A |
| Kinesin light chain 1 (Fragment) | KLC1 | S262 | | | | | | | 29.8 | 73.4 | R.ALSAS*HTDLAH.- |
| Kinesin light chain 4 | KLC4 | S608 | | | | | | | 44.2 | 130.8 | R.AAS*LNYLNQPSAAPLQVSR.G |
| Kinesin light chain 4 | KLC4 | T630 | | | | | | | 7.0 | 18.5 | R.GLSAST*MDLSSSS.- |
| Kinesin light chain 4 | KLC4 | S629 | | | | | | | 16.1 | 38.2 | R.GLSAS*TMDLSSSS.- |
| KRI1 homolog | KRI1 | S177 | | | | | | | 110.9 | 154.1 | R.AFVEDS*EDEDGAGEGGSSLLQK.R |
| KRI1 homolog | KRI1 | S634S645 | | | | | | | 48.5 | 14.7 | R.QLPALDGSLMGPES*PPAQEEEAPVS*PHKKPAPQKR.R |
| KRI1 homolog | KRI1 | S99S100S101S | | | | | | | 17.7 | 61.2 | R.TAS*S*S*DS*EEDPEALEK.Q |
| Kruppel like factor 12 | KLF12 | S202 | | | | | | | 70.9 | 119.5 | R.S*PGNVNNTIVVPLLEDGR.G |
| Kruppel like factor 12 | KLF12 | S238S240 | | | | | | | 1.7 | 52.6 | R.QSKS*DS*DDDDLPNVTLDSVNETGSTALSIAR.A |
| Kruppel like factor 12 | KLF12 | S236S240 | | | | | | | 12.2 | 74.0 | R.QS*KSDS*DDDDLPNVTLDSVNETGSTALSIAR.A |
| Kruppel like factor 12 | KLF12 | S236T249 | | | | | | | 11.0 | 74.5 | R.QS*KSDSDDDDLPNVT*LDSVNETGSTALSIAR.A |
| Kruppel like factor 12 | KLF12 | S311 | | | | | | | 7.3 | 11.4 | R.SESPDS*RK.R |
| Kruppel like factor 2 | KLF2 | T244S248 | | | | | | | 100.0 | 48.7 | R.GLLT*PPAS*PLELLEAKPK.R |
| L3MBTL2 | L3MBTL2 | T66S73 | | | | | | | 25.1 | 22.9 | R.EAGELPT*SPLHLLS*PGTPR.S |
| L3MBTL2 | L3MBTL2 | S67S73 | | | | | | | 5.6 | 21.4 | R.EAGELPTS*PLHLLS*PGTPR.S |
| Lactate dehydrogenase A | LDHA | Y10 | | | | | | | 100.0 | 43.4 | K.DQLIY*NLLK.E |
| LAG1 longevity assurance homolog 5 | CERS5 | S350S354S355 | | | | | | | 15.6 | 27.6 | K.VSKDDRS*DVES*S*S*EEEDVTTCTK.S |
| LAG1 longevity assurance homolog 5 | CERS5 | S345S354S355 | | | | | | | 43.5 | 44.8 | K.VS*KDDRSDVES*S*S*EEEDVTTCTK.S |
| Lamin B receptor | LBR | S86 | | | | | | | 100.0 | 20.2 | R.S*PGRPPK.S |
| Lamin B receptor | LBR | S84S86 | | | | | | | 100.0 | 18.6 | R.S*RS*PGRPPK.S |
| Lamin B receptor | LBR | S99 | | | | | | | 11.1 | 76.7 | R.SAS*ASHQADIK.E |
| Lamin B receptor | LBR | S97 | | | | | | | 23.5 | 46.2 | R.S*ASASHQADIK.E |
| Lamin B receptor | LBR | S67T68 | | | | | | | 6.5 | 16.4 | R.KGGS*T*SSSPSRR.R |
| Lamin B1 | LMNB1 | S391 | | | | | | | 22.4 | 22.4 | R.LKLS*PSPSSR.V |
| Lamin B1 | LMNB1 | S391S393 | | | | | | | 20.6 | 39.2 | R.LKLS*PS*PSSR.V |
| Lamin B1 | LMNB1 | T20S23 | | | | | | | 15.2 | 24.0 | R.AGGPTT*PLS*PTR.L |
| Lamin B1 | LMNB1 | T575 | | | | | | | 114.9 | 29.0 | K.TTIPEEEEEEEEAAGVVVEEELFHQQGT*PR.A |
| Lamin B1 | LMNB1 | T19S23 | | | | | | | 9.0 | 18.4 | R.AGGPT*TPLS*PTR.L |
| | TOR1AIP1 | S156S157 | | | | | | | 10.4 | 73.5 | R.DSHS*S*EEDEASSQTDLSQTISKK.T |
| Lamina associated polypeptide 1B | TOR1AIP1 | T220 | | | | | | | 9.0 | 64.4 | K.VNFSEEGET*EEDDQDSSHSSVTTVK.A |
| Lamina associated polypeptide 1B | TOR1AIP1 | S143 | | | | | | | 24.0 | 45.4 | R.LQQQHSEQPPLQPS*PVMTR.R |
| Lamina associated polypeptide 1B | TOR1AIP1 | S154S156S157 | | | | | | | 64.6 | 66.5 | R.GLRDS*HS*S*EEDEASSQTDLSQTISK.K |
| Lamina associated polypeptide 1B | TOR1AIP1 | S154S157 | | | | | | | 8.9 | 41.2 | R.DS*HSS*EEDEASSQTDLSQTISK.K |
| Lamina associated polypeptide 1B | TOR1AIP1 | S154S156S163 | | | | | | | 9.4 | 26.5 | R.RGLRDS*HS*SEEDEAS*SQTDLSQTISK.K |
| Lamina associated polypeptide 1B | TOR1AIP1 | S154S156 | | | | | | | 24.1 | 17.3 | R.DS*HS*SEEDEASSQTDLSQTISK.K |
| LARP | LARP1 | S697 | | | | | | | 25.6 | 49.9 | R.SLPTTVPES*PNYR.N |
| LARP | LARP1 | S689 | | | | | | | 30.0 | 54.0 | R.S*LPTTVPESPNYR.N |
| LARP | LARP1 | S689S697 | | | | | | | 18.3 | 38.2 | R.S*LPTTVPES*PNYR.N |
| LARP | LARP1 | S770S776 | | | | | | | 3.4 | 64.3 | R.TAS*ISSSPS*EGTPTVGSYGCTPQSLPK.F |
| LARP | LARP1 | S90 | | | | | | | 105.7 | 139.4 | R.ESPRPLQLPGAEGPAIS*DGEEGGGEPGAGGGAAGAAGAGR.R |
| LARP | LARP1 | S75 | | | | | | | | 79.2 | R.ES*PRPLQLPGAEGPAISDGEEGGGEPGAGGGAAGAAGAGR.R |
| LARP | LARP1 | S747 | | | | | | | 15.3 | 83.4 | R.HSS*NPPLESHVGWVMDSR.E |
| LARP | LARP1 | S75S90 | | | | | | | 100.0 | 47.5 | R.ES*PRPLQLPGAEGPAIS*DGEEGGGEPGAGGGAAGAAGAGR.R |
| LARP | LARP1 | S550S554 | | | | | | | 37.8 | 137.9 | K.NTFTAWS*DEES*DYEIDDR.D |
| LARP | LARP1 | S471 | | | | | | | 23.9 | 104.4 | K.GLSAS*LPDLDSENWIEVK.K |
| LARP | LARP1 | S440S444 | | | | | | | 23.4 | 52.2 | K.ETES*APGS*PR.A |
| LARP | LARP1 | T449 | | | | | | | 26.5 | 40.7 | R.AVT*PVPTK.T |
| LARP | LARP1 | T693 | | | | | | | 23.6 | 49.7 | R.SLPTT*VPESPNYR.N |
| LARP | LARP1 | T570 | | | | | | | 11.4 | 29.2 | K.ILIVT*QTPHYMR.R |
| LARP | LARP1 | S770T779 | | | | | | | 8.0 | 56.2 | R.TAS*ISSSPSEGT*PTVGSYGCTPQSLPK.F |
| LARP | LARP1 | S774S776T779 | | | | | | | 3.5 | 38.8 | R.TASISSS*PS*EGT*PTVGSYGCTPQSLPK.F |
| LARP | LARP1 | S774 | | | | | | | 2.3 | 19.3 | R.TASISSS*PSEGTPTVGSYGCTPQSLPK.F |
| LARP | LARP1 | T545T547 | | | | | | | 10.8 | 13.0 | K.NT*FT*AWSDEESDYEIDDRDVNK.I |
| LARP | LARP1 | S550Y556 | | | | | | | 18.9 | 76.8 | K.NTFTAWS*DEESDY*EIDDR.D |
| LARP | LARP1 | T692 | | | | | | | -0.2 | 16.3 | R.SLPT*TVPESPNYR.N |
| LARP | LARP1 | S550 | | | | | | | 25.3 | 54.3 | K.NTFTAWS*DEESDYEIDDR.D |
| LARP | LARP1 | S444 | | | | | | | 19.3 | 27.3 | K.ETESAPGS*PR.A |

**Peak Area**: <-10, -5, -3, 0, 3, 5, >10

**%CV**: 0, 29, 42, 71, 86, >100

**White dots**- Significant change in peptide abundance at **5** %FDR compared to the timepoint with the minimum peak area for a given PSM

| Protein Name | Gene | Phosphosites | CarT 0m | CarT 2m | CarT 5m | RajiB 0m | RajiB 2m | RajiB 5m | Ascor | MOWSE | Sequence |
|---|---|---|---|---|---|---|---|---|---|---|---|
| **LARP** | LARP1 | S979 | | | | | | | **18.8** | **19.3** | R.CPS*QSSSRPAAMISQPPTPPTGQPVR.E |
| **LARP** | LARP1 | T779 | | | | | | | **9.1** | **73.3** | R.TASISSSPSEGT*PTVGSYGCTPQSLPK.F |
| **LARP** | LARP1 | S143 | | | | | | | **30.7** | **42.4** | K.NALPPVLTTVNGQS*PPEHSAPAK.V |
| **LARP** | LARP1 | S746 | | | | | | | **69.6** | **61.3** | R.HS*SNPPLESHVGWVMDSR.E |
| **LARP** | LARP1 | S772S773S776 | | | | | | | **6.0** | **17.3** | R.TASIS*S*SPS*EGTPTVGSYGCTPQSLPK.F |
| **LARP** | LARP1 | S469 | | | | | | | **37.2** | **66.9** | K.GLS*ASLPDLDSENWIEVK.K |
| **LARP** | LARP1 | S776 | | | | | | | **-0.2** | **48.0** | R.TASISSSPS*EGTPTVGSYGCTPQSLPK.F |
| **LARP** | LARP1 | S554Y556 | | | | | | | **10.3** | **43.7** | R.KNTFTAWSDEES*DY*EIDDR.D |
| **LARP** | LARP1 | S770 | | | | | | | **5.8** | **49.3** | R.TAS*ISSSPSEGTPTVGSYGCTPQSLPK.F |
| **LARP** | LARP1 | S770S772S773 | | | | | | | **8.7** | **33.3** | R.TAS*IS*S*SPSEGTPTVGSYGCTPQSLPK.F |
| **LARP** | LARP1 | S772T779 | | | | | | | **9.4** | **44.9** | R.TASIS*SSPSEGT*PTVGSYGCTPQSLPK.F |
| **LARP** | LARP1 | S770S774 | | | | | | | **1.6** | **25.1** | R.TAS*ISSS*PSEGTPTVGSYGCTPQSLPK.F |
| **LARP** | LARP1 | T693S697 | | | | | | | **5.1** | **14.2** | R.SLPTT*VPES*PNYR.N |
| **LARP** | LARP1 | S772S776 | | | | | | | **4.4** | **21.3** | R.TASIS*SSPS*EGTPTVGSYGCTPQSLPK.F |
| **LARP** | LARP1 | S774S776T781 | | | | | | | **4.6** | **24.3** | R.TASISSS*PS*EGTPT*VGSYGCTPQSLPK.F |
| **LARP** | LARP1 | T438S444 | | | | | | | **19.3** | **45.8** | K.ET*ESAPGS*PR.A |
| **LARP** | LARP1 | S770S773 | | | | | | | **3.8** | **48.7** | R.TAS*ISS*SPSEGTPTVGSYGCTPQSLPK.F |
| **LARP** | LARP1 | S770S772 | | | | | | | **0.5** | **43.1** | R.TAS*IS*SSPSEGTPTVGSYGCTPQSLPK.F |
| **LASS2** | CERS2 | S341T346S348 | | | | | | | **100.0** | **59.0** | K.LVEDERS*DREET*ES*S*EGEEAAAGGGAK.S |
| **LASS2** | CERS2 | S341 | | | | | | | **100.0** | **51.6** | K.LVEDERS*DR.E |
| **LASS2** | CERS2 | T346S348S349 | | | | | | | **100.0** | **112.3** | R.EET*ES*S*EGEEAAAGGGAK.S |
| **LATS1** | LATS1 | S278 | | | | | | | **25.6** | **53.2** | K.RYS*GNMEYVISR.I |
| **LATS1** | LATS1 | S1111 | | | | | | | **15.1** | **31.0** | R.FFDDNGYPYNYPKPIEYEYINSQGSEQQS*DEDDQNTGSEIK.N |
| **LBA** | LRBA | S2064 | | | | | | | **15.2** | **112.3** | R.SQALGNQNSENEILLEGDDDTLS*SVDEK.D |
| **LBA** | LRBA | S1787 | | | | | | | **11.1** | **80.6** | K.LPSVPTVDSVS*QDPVSNMSITER.L |
| **LBA** | LRBA | S1767 | | | | | | | **17.2** | **83.4** | K.SAVSPSTFNTSIPTNAVSVVSSVDSAQASDMGGES*PGSR.S |
| **LBA** | LRBA | S1785 | | | | | | | **8.0** | **45.5** | K.LPSVPTVDS*VSQDPVSNMSITER.L |
| **LBA** | LRBA | S1733 | | | | | | | | **38.4** | K.S*AVSPSTFNTSIPTNAVSVVSSVDSAQASDMGGESPGSR.S |
| **LBH limb bud and heart development homolog** | LBH | S63 | | | | | | | **89.5** | **31.5** | R.LPS*IVVEPTEGEVESGELR.W |
| **Lck** | LCK | Y192 | | | | | | | **32.1** | **58.9** | R.NLDNGGFY*ISPR.I |
| **Lck** | LCK | S42 | | | | | | | | **45.6** | R. NGS*EVRDPLVTYEGSNPPASPLQDNLVIALHSYEPSHDGDLGFEK |
| **Lck** | LCK | T50 | | | | | | | | **74.5** | R.DPLVT*YEGSNPPASPLQDNLVIALHSYEPSHDGDLGFEK.G |
| **Lck** | LCK | S59 | | | | | | | **48.9** | **53.0** | R.DPLVTYEGSNPPAS*PLQDNLVIALHSYEPSHDGDLGFEK.G |
| **Lck** | LCK | Y505 | | | | | | | **27.9** | **28.7** | R.SVLEDFFTATEGQY*QPQP.- |
| **Lck** | LCK | T501 | | | | | | | **13.4** | **19.1** | R.SVLEDFFTAT*EGQYQPQP.- |
| **Lck** | LCK | S42T50 | | | | | | | | **41.4** | R. NGS*EVRDPLVT*YEGSNPPASPLQDNLVIALHSYEPSHDGDLGFEK |
| **Lck-interacting transmembrane adapter 1** | LIME1 | S86 | | | | | | | **31.1** | **51.7** | R.AAPGPAQQQGPAACQHGS*PAPTLAGGVQGHHR.T |
| **LEM domain containing 2** | LEMD2 | S496 | | | | | | | **9.1** | **42.3** | R.WTKPS*SFSDSER.- |
| **Leucine rich repeat containing 47** | LRRC47 | S518 | | | | | | | **12.6** | **76.9** | K.EEGS*LSDTEADAVSGQLPDPTTNPSAGK.D |
| **Leucine rich repeat containing 47** | LRRC47 | S520 | | | | | | | **18.8** | **71.3** | K.EEGSLS*DTEADAVSGQLPDPTTNPSAGK.D |
| **Leucine rich repeat containing 47** | LRRC47 | S518S520 | | | | | | | **5.6** | **14.2** | K.EEGS*LS*DTEADAVSGQLPDPTTNPSAGK.D |
| **Leucine rich repeat containing 47** | LRRC47 | T522 | | | | | | | **18.4** | **73.8** | K.EEGSLSDT*EADAVSGQLPDPTTNPSAGK.D |
| **Leucine rich repeat in FLII interacting protein 1** | LRRFIP1 | S115 | | | | | | | **9.8** | **57.6** | R.NMPGLSAATLASLGGTS*SR.R |
| **Leucine rich repeat in FLII interacting protein 1** | LRRFIP1 | T114 | | | | | | | **29.3** | **51.3** | R.NMPGLSAATLASLGGT*SSR.R |
| **Leucine rich repeat in FLII interacting protein 1** | LRRFIP1 | S116 | | | | | | | **6.8** | **69.8** | R.NMPGLSAATLASLGGTSS*R.R |
| **Leucine rich repeat in FLII interacting protein 1** | LRRFIP1 | S110 | | | | | | | **6.2** | **60.4** | R.NMPGLSAATLAS*LGGTSSR.R |
| **Leucine rich repeats and calponin homology domain containing protein 1** | LRCH1 | S532S536 | | | | | | | **37.0** | **72.5** | R.ENS*PAVS*PTTNSTAPFGLKPR.S |
| **Leucine rich repeats and calponin homology domain containing protein 1** | LRCH1 | T538T539 | | | | | | | **17.2** | **26.4** | R.ENSPAVSPT*T*NSTAPFGLKPR.S |
| **Leucine zipper protein 1** | LUZP1 | S659 | | | | | | | **57.5** | **46.2** | R.EKPDS*DDDLDIASLVTAK.L |
| **Leucine-rich repeat flightless-interacting protein 1** | LRRFIP1 | S124 | | | | | | | **8.5** | **53.8** | R.RGSGDTS*ISIDTEASIR.E |
| **Leucine-rich repeat flightless-interacting protein 1** | LRRFIP1 | S120 | | | | | | | **68.1** | **81.4** | R.RGS*GDTSISIDTEASIR.E |
| **Leucine-rich repeat flightless-interacting protein 1** | LRRFIP1 | S126 | | | | | | | **6.5** | **55.0** | R.RGSGDTSIS*IDTEASIR.E |
| **Leucine-rich repeats and IQ motif containing 2** | CEP97 | S770 | | | | | | | **21.9** | **83.2** | K.ESSNNEQDNS*LLEQYLTSVQQLEDADER.T |
| **Leucine-rich repeats and IQ motif containing 2** | CEP97 | T811S813 | | | | | | | **2.4** | **44.5** | K.LHIACFPVQLDT*LS*DGASVDESHGISPPLQGEISQTQENSK.L |
| **Leucine-rich repeats and IQ motif containing 2** | CEP97 | S813S817 | | | | | | | **0.1** | **28.5** | K.LHIACFPVQLDTLS*DGAS*VDESHGISPPLQGEISQTQENSK.L |
| **Leucine-rich repeats and IQ motif containing 2** | CEP97 | S762 | | | | | | | **15.2** | **58.5** | K.ES*SNNEQDNSLLEQYLTSVQQLEDADER.T |
| **Leucine-rich repeats and IQ motif containing 2** | CEP97 | S763 | | | | | | | **-0.4** | **36.6** | K.ESS*NNEQDNSLLEQYLTSVQQLEDADER.T |
| **Leucine-rich repeats and IQ motif containing 2** | CEP97 | S763S770 | | | | | | | **4.1** | **83.8** | K.ESS*NNEQDNS*LLEQYLTSVQQLEDADER.T |
| **Leupaxin** | LPXN | S19 | | | | | | | **7.4** | **49.2** | R.STLQDS*DEYSNPAPLPLDQHSR.K |
| **Leupaxin** | LPXN | S81 | | | | | | | **58.0** | **33.3** | K.ES*PPPSK.T |
| **LIG1 DNA ligase I** | LIG1 | T195 | | | | | | | **32.3** | **63.0** | K.AET*PTESVSEPEVATK.Q |
| **LIG1 DNA ligase I** | LIG1 | S911T918 | | | | | | | **15.2** | **67.9** | R.KQSQIQNQQGEDS*GSDPEDT*Y.- |

Peak Area: <-10, -5, -3, 0, 3, 5, >10

%CV: 0, 29, 42, 71, 86, >100

**White dots**- Significant change in peptide abundance at 5 %FDR compared to the timepoint with the minimum peak area for a given PSM

| Protein Name | Gene | Phosphosites | CarT 0m 2m 5m | RajiB 0m 2m 5m | Ascor | MOWSE | Sequence |
|---|---|---|---|---|---|---|---|
| LIG1 DNA ligase I | LIG1 | T918Y919 | | | 7.6 | 18.3 | R.KQSQIQNQQGEDSGSDPEDT*Y*.- |
| LIG1 DNA ligase I | LIG1 | T195S201 | | | 10.6 | 70.4 | K.AET*PTESVS*EPEVATK.Q |
| | LIG1 | S911S913 | | | 33.9 | 37.5 | K.QSQIQNQQGEDS*GS*DPEDTY.- |
| | LIG1 | S51 | | | | 49.9 | K.EWNGVVSESDS*PVK.R |
| LIG1 DNA ligase I | LIG1 | S66S76 | | | 100.0 | 105.0 | R.VLGS*EGEEEDEALS*PAK.G |
| LIG1 DNA ligase I | LIG1 | S141 | | | 134.4 | 95.6 | R.TIQEVLEEQS*EDEDR.E |
| LIG1 DNA ligase I | LIG1 | S91S109 | | | 13.1 | 32.8 | K.GQKPALDCSQVS*PPRPATSPENNASLSDTS*PMDSSPSGIPK.R |
| LIG1 DNA ligase I | LIG1 | S91T108 | | | 6.1 | 30.2 | K.GQKPALDCSQVS*PPRPATSPENNASLSDT*SPMDSSPSGIPK.R |
| LIG1 DNA ligase I | LIG1 | S91T97 | | | 14.4 | 37.3 | K.GQKPALDCSQVS*PPRPAT*SPENNASLSDTSPMDSSPSGIPK.R |
| LIG1 DNA ligase I | LIG1 | S91 | | | 2.3 | 35.8 | K.GQKPALDCSQVS*PPRPATSPENNASLSDTSPMDSSPSGIPK.R |
| LIG1 DNA ligase I | LIG1 | S91S113 | | | 16.2 | 26.0 | K.GQKPALDCSQVS*PPRPATSPENNASLSDTSPMDS*SPSGIPK.R |
| LIG1 DNA ligase I | LIG1 | S76 | | | 84.2 | 99.6 | R.VLGSEGEEEDEALS*PAK.G |
| LIG1 DNA ligase I | LIG1 | S91S98 | | | 3.9 | 41.8 | K.GQKPALDCSQVS*PPRPATS*PENNASLSDTSPMDSSPSGIPK.R |
| LIG1 DNA ligase I | LIG1 | S88S91 | | | 0.1 | 20.0 | K.GQKPALDCS*QVS*PPRPATSPENNASLSDTSPMDSSPSGIPK.R |
| LIG1 DNA ligase I | LIG1 | S49 | | | 22.4 | 30.6 | K.EWNGVVSES*DSPVK.R |
| LIG1 DNA ligase I | LIG1 | T97 | | | -5.7 | 38.9 | K.GQKPALDCSQVSPPRPAT*SPENNASLSDTSPMDSSPSGIPK.R |
| LIG1 DNA ligase I | LIG1 | S47 | | | | 17.4 | K.EWNGVVS*ESDSPVK.R |
| LIG1 DNA ligase I | LIG1 | S911Y919 | | | 6.8 | 23.5 | R.KQSQIQNQQGEDS*GSDPEDTY*.- |
| LIG1 DNA ligase I | LIG1 | S88S109 | | | 2.2 | 18.1 | K.GQKPALDCS*QVSPPRPATSPENNASLSDTS*PMDSSPSGIPK.R |
| LIG1 DNA ligase I | LIG1 | S91S104 | | | 11.4 | 22.3 | K.GQKPALDCSQVS*PPRPATSPENNAS*LSDTSPMDSSPSGIPK.R |
| Lim and SH3 protein 1 | LASP1 | S146 | | | 41.3 | 50.8 | R.DS*QDGSSYR.R |
| LIM domain containing 2 | LIMD2 | S29 | | | 39.8 | 43.6 | R.SKS*FSLR.A |
| LIM domain only 7 | LMO7 | S1159 | | | 12.0 | 64.2 | R.SAS*VNKEPVSLPGIMR.R |
| LIM domain only 7 | LMO7 | S342 | | | 60.7 | 85.1 | R.S*WASPVYTEADGTFSR.L |
| LIM domain only 7 | LMO7 | S863 | | | 21.8 | 63.9 | R.S*QFFEQGSSDSVVPDLPVPTISAPSR.W |
| LIM domain only 7 | LMO7 | S531 | | | 35.4 | 51.0 | R.VTTEIQLPS*QSPVEEQSPASLSSLR.S |
| LIM domain only 7 | LMO7 | S1157 | | | 6.2 | 13.2 | R.S*ASVNKEPVSLPGIMR.R |
| LIMD1 | LIMD1 | S316 | | | 14.5 | 73.9 | R.SNS*GLGGEVSGVMSKPNVDPQPWFQDGPK.S |
| LIMD1 | LIMD1 | S424 | | | 12.2 | 144.3 | K.EGPLGWSSDGSLGSVLLDSPSS*PR.V |
| LIMD1 | LIMD1 | S421 | | | -0.3 | 82.2 | K.EGPLGWSSDGSLGSVLLDS*PSSPR.V |
| Limkain b1 | MARF1 | S1093 | | | 22.5 | 62.3 | R.SKS*PVGNPQLIQFSR.E |
| Limkain b1 | MARF1 | S1091 | | | 12.2 | 71.1 | R.S*KSPVGNPQLIQFSR.E |
| Limkain beta 2 | CCDC92 | S211 | | | 18.6 | 14.3 | K.SLS*APLHPEFEEVYR.F |
| Lin 9 homolog | LIN9 | S325S337 | | | 87.5 | 49.7 | R.LHYTPPLQS*PIIDNDPLLGQS*PWR.S |
| Lin 9 homolog | LIN9 | T320S325S337 | | | 52.0 | 34.7 | R.LHYT*PPLQS*PIIDNDPLLGQS*PWR.S |
| Lin 9 homolog | LIN9 | Y319S325S337 | | | 39.3 | 32.1 | R.LHY*TPPLQS*PIIDNDPLLGQS*PWR.S |
| Linker for activation of T cells | LAT | S224 | | | 67.0 | 71.5 | R.EYVNVS*QELHPGAAK.T |
| Linker for activation of T cells | LAT | S84 | | | 66.8 | 40.6 | R.S*PQPLGGSHR.T |
| Lipase hormone sensitive | LIPE | S950 | | | 10.5 | 27.9 | R.RS*SQGATQMPLYSSPIVK.N |
| Liprin beta 2 | PPFIBP2 | S387 | | | 29.8 | 33.1 | K.KLSCS*LEDLR.S |
| Liprin beta 2 | PPFIBP2 | S512 | | | 19.9 | 108.8 | R.TQS*GNFYTDTLGMAEFR.R |
| Liprin beta 2 | PPFIBP2 | S414 | | | 32.7 | 21.6 | K.DS*PFLAEHKYPTLPGK.L |
| Liver-specific bHLH-Zip transcription factor | LSR | S530 | | | 43.4 | 41.5 | R.S*RDDLYDQDDSRDFPR.S |
| Liver-specific bHLH-Zip transcription factor | LSR | S643S646 | | | 100.0 | 15.7 | K.NLALS*RES*LVV.- |
| Liver-specific bHLH-Zip transcription factor | LSR | T336 | | | 11.4 | 23.4 | K.T*PPPPAMIPMGPAYNGYPGGYPGDVDR.S |
| Liver-specific bHLH-Zip transcription factor | LSR | S643 | | | 100.0 | 13.1 | K.NLALS*R.E |
| LKB1 interacting protein 1 | STK11IP | T614 | | | 4.7 | 35.8 | R.SPRPT*GSDLLPGAPILSLR.F |
| LKB1 interacting protein 1 | STK11IP | S610 | | | 8.6 | 42.4 | R.S*PRPTGSDLLPGAPILSLR.F |
| LKB1 interacting protein 1 | STK11IP | S616 | | | 21.3 | 52.5 | R.SPRPTGS*DLLPGAPILSLR.F |
| LKB1 interacting protein 1 | STK11IP | S415S431 | | | 18.2 | 25.9 | R.TLNPS*PAGWFVQQHPELELMS*SFR.E |
| LNK | SH2B3 | S121 | | | -0.4 | 15.4 | R.SS*EELAPPRPPGPCSFQHFR.R |
| LNK | SH2B3 | S120 | | | 8.5 | 14.5 | R.S*SEELAPPRPPGPCSFQHFR.R |
| LOC115294 protein | PCMTD1 | S302 | | | 100.4 | 56.7 | R.INTYVFVGNQLIPQPLDS*EEDEK.M |
| LOC159090 | FAM122B | S58 | | | 100.0 | 66.7 | R.HS*LEEGLDMVNR.E |
| LOC159090 | SPACIA2 | S115S119 | | | 36.8 | 44.6 | K.RIDFTPVS*PAPS*PTR.G |
| LOC159090 | SPACIA2 | S50 | | | 8.9 | 32.7 | R.RNS*TTIMSR.H |
| LOC159090 | SPACIA2 | S33 | | | 15.5 | 24.4 | R.SSSAPLIHGLS*DLSQVFQPYTLR.T |
| LOC159090 | SPACIA2 | S25 | | | 6.6 | 24.7 | R.SSS*APLIHGLSDLSQVFQPYTLR.T |
| LOC388974 | RGPD2 | S520S525 | | | 14.9 | 17.0 | R.EDALDDNVSSSSVHDS*PLASS*PVRK.N |
| LOC388974 | RGPD2 | S990 | | | 28.4 | 35.4 | R.ESLLPVINTMLQLS*PEEK.G |
| LOC389677 protein | RBM12B | S638 | | | 24.9 | 47.7 | R.S*PTEDFR.Q |
| LOC389677 protein | RBM12B | S710S718 | | | 100.0 | 22.6 | R.RPPEEDFRHS*PEEDFRQS*PQEHFR.R |

Peak Area: <-10, -5, -3, 0, 3, 5, >10

%CV: 0, 29, 42, 71, 86, >100

**White dots-** Significant change in peptide abundance at **5** %FDR compared to the timepoint with the minimum peak area for a given PSM

| Protein Name | Gene | Phosphosites | Ascor | MOWSE | Sequence |
|---|---|---|---|---|---|
| **LOC389677 protein** | RBM12B | S278S280 | **50.8** | **28.8** | R.S*RS*PLGFYVHLK.N |
| **LOC389677 protein** | RBM12B | S250S254 | **100.0** | **29.1** | R.S*EEHS*PPR.G |
| **LOC389677 protein** | RBM12B | T640 | **16.9** | **50.6** | R.SPT*EDFR.Q |
| **LOC389677 protein** | RBM12B | S98T100 | **16.0** | **42.7** | R.GRPGS*GT*SGVDSLSNFIESVK.E |
| **LOC389677 protein** | RBM12B | T100S101 | **6.4** | **31.7** | R.GRPGSGT*S*GVDSLSNFIESVK.E |
| **LOC389677 protein** | RBM12B | T100 | **1.8** | **54.0** | R.GRPGSGT*SGVDSLSNFIESVK.E |
| **LRCH3** | LRCH3 | S324 | **56.6** | **81.8** | R.WS*GNEPTDEFSDLPLR.V |
| **LRRFIP2** | LRRFIP2 | S328 | **29.9** | **65.1** | R.RGS*GDTSSLIDPDTSLSELR.D |
| **LRRFIP2** | LRRFIP2 | S333 | **4.2** | **20.2** | R.RGSGDTSS*LIDPDTSLSELR.D |
| **LUC7 like** | LUC7L | S363 | **100.0** | **39.8** | R.S*EEKEAGEI.- |
| **LUC7 like 2** | LUC7L2 | T17 | | **39.0** | R.AMLDQLMGT*SR.D |
| **LUC7 like 2** | FMC1 | S383S384 | **14.2** | **24.1** | R.SEDRRS*S*EER.E |
| **LUC7 like 2** | FMC1 | S378S383 | **6.7** | **18.3** | R.S*EDRRS*SEER.E |
| **LUC7 like 2** | LUC7L2 | S18 | **6.5** | **48.3** | R.AMLDQLMGTS*R.D |
| **LUC7 like 2** | FMC1 | S281S283S285 | **100.0** | **10.9** | R.S*RS*MS*R.E |
| **Lymphocyte antigen 9** | LY9 | S648 | **9.1** | **24.7** | R.KPQVVPPPQQNDLEIPES*PTYENFT.- |
| **Lymphocyte antigen 9** | LY9 | T650 | **13.9** | **18.9** | R.KPQVVPPPQQNDLEIPESPT*YENFT.- |
| **Lymphocyte cytosolic protein 1** | LCP1 | S5 | **17.7** | **102.5** | R.GS*VSDEEMM#ELR.E |
| **Lymphocyte cytosolic protein 1** | LCP1 | S7 | **-0.1** | **84.6** | R.GSVS*DEEMM#ELR.E |
| **Lymphocyte cytosolic protein 1** | LCP1 | S257 | **100.0** | **75.0** | R.EGES*LEDLMK.L |
| **Lymphocyte cytosolic protein 1** | LCP1 | Y28 | **35.4** | **71.8** | K.VDTDGNGY*ISFNELNDLFK.A |
| **Lymphocyte cytosolic protein 1** | LCP1 | S406 | **100.0** | **45.0** | R.NWMNS*LGVNPR.V |
| **Lymphocyte specific protein** | LSP1 | S139 | **5.4** | **36.3** | K.EDSDEVHLEELS*LSK.E |
| **Lymphocyte specific protein** | LSP1 | S130 | **99.2** | **87.5** | K.EDS*DEVHLEELSLSK.E |
| **Lymphocyte specific protein** | LSP1 | S111S130 | **7.5** | **33.2** | R.S*PEGEQEDRPGLHAYEKEDS*DEVHLEELSLSK.E |
| **Lymphocyte specific protein** | LSP1 | S252 | **60.5** | **102.9** | R.QAS*IELPSMAVASTK.S |
| **Lymphocyte specific protein** | LSP1 | S189 | **9.2** | **94.0** | R.TPSPLVLEGTIEQSS*PPLSPTTK.L |
| **Lymphocyte specific protein** | LSP1 | T184 | **32.4** | **74.8** | R.TPSPLVLEGT*IEQSSPPLSPTTK.L |
| **Lymphocyte specific protein** | LSP1 | S177 | **21.0** | **77.5** | R.TPS*PLVLEGTIEQSSPPLSPTTK.L |
| **Lymphocyte specific protein** | LSP1 | S177S189 | **26.8** | **47.7** | R.TPS*PLVLEGTIEQSS*PPLSPTTK.L |
| **Lymphocyte specific protein** | LSP1 | S188S189 | **40.0** | **68.0** | R.TPSPLVLEGTIEQS*S*PPLSPTTK.L |
| **Lymphocyte specific protein** | LSP1 | S188T196 | **14.8** | **41.8** | R.TPSPLVLEGTIEQS*SPPLSPTT*K.L |
| **Lymphocyte specific protein** | LSP1 | T175S177 | **30.0** | **53.2** | R.T*PS*PLVLEGTIEQSSPPLSPTTK.L |
| **Lymphocyte specific protein** | LSP1 | S177S189S193 | **14.1** | **21.0** | R.TPS*PLVLEGTIEQSS*PPLS*PTTK.L |
| **Lymphocyte specific protein** | LSP1 | T244 | **33.1** | **59.1** | K.IDQWLEQYTQAIETAGRT*PK.L |
| **Lymphocyte specific protein** | LSP1 | T175T184 | **9.9** | **36.7** | R.T*PSPLVLEGT*IEQSSPPLSPTTK.L |
| **Lymphocyte specific protein** | LSP1 | T175S177S189 | **23.1** | **52.2** | R.T*PS*PLVLEGTIEQSS*PPLSPTTK.L |
| **Lymphocyte specific protein** | LSP1 | S177T184S189 | **15.1** | **45.9** | R.TPS*PLVLEGT*IEQSS*PPLSPTTK.L |
| **Lymphocyte specific protein** | LSP1 | T175S177S189 | **22.3** | **16.4** | R.T*PS*PLVLEGTIEQSS*PPLS*PTTK.L |
| **Lymphocyte specific protein** | LSP1 | T175S177S188 | **19.1** | **14.2** | R.T*PS*PLVLEGTIEQS*SPPLS*PTTK.L |
| **Lymphocyte specific protein** | LSP1 | S204 | **38.1** | **32.7** | R.TES*LNR.S |
| **Lymphocyte specific protein** | LSP1 | S188 | **62.3** | **88.9** | R.TPSPLVLEGTIEQS*SPPLSPTTK.L |
| **Lymphocyte specific protein** | LSP1 | T184S189 | **24.1** | **40.6** | R.TPSPLVLEGT*IEQSS*PPLSPTTK.L |
| **Lymphocyte specific protein** | LSP1 | T175S177T184 | **13.2** | **17.4** | R.T*PS*PLVLEGT*IEQSSPPLS*PTTK.L |
| **Lymphocyte specific protein** | LSP1 | S177T184 | **16.7** | **51.4** | R.TPS*PLVLEGT*IEQSSPPLSPTTK.L |
| **Lymphocyte specific protein** | LSP1 | S188S193 | **11.2** | **41.0** | R.TPSPLVLEGTIEQS*SPPLS*PTTK.L |
| **Lymphocyte specific protein** | LSP1 | S111 | **53.7** | **24.0** | R.S*PEGEQEDRPGLHAYEK.E |
| **Lymphocyte specific protein** | LSP1 | S24 | **5.7** | **56.3** | R.EELLGPTAQWS*VEDEEEAVHEQCHER.D |
| **Lymphocyte specific protein** | LSP1 | T20 | **4.3** | **16.3** | R.EELLGPT*AQWSVEDEEEAVHEQCHER.D |
| **Lymphocyte specific protein** | LSP1 | T175S177T184 | **6.6** | **14.6** | R.T*PS*PLVLEGT*IEQSSPPLSPT*TK.L |
| **Lymphocyte specific protein** | LSP1 | S141 | **22.5** | **113.1** | K.EDSDEVHLEELSLS*K.E |
| **Lymphocyte specific protein** | LSP1 | S177T184S188 | **18.5** | **11.2** | R.TPS*PLVLEGT*IEQS*SPPLSPTTK.L |
| **Lymphocyte specific protein** | LSP1 | S177S188S189 | **15.6** | **21.7** | R.TPS*PLVLEGTIEQS*S*PPLSPTTK.L |
| **Lymphocyte specific protein** | LSP1 | T184S188 | **8.0** | **56.4** | R.TPSPLVLEGT*IEQS*SPPLSPTTK.L |
| **Lymphocyte specific protein** | LSP1 | T175S177T184 | **5.5** | **11.5** | R.T*PS*PLVLEGT*IEQSSPPLSPTTK.L |
| **Lymphoid enhancer binding factor 1** | LEF1 | S132 | **2.6** | **21.4** | K.GPSYSSYSGYIMMPNMNNDPYMSNGSLS*PPIPR.T |
| | LYN | S13 | **25.3** | **80.3** | K.DSLS*DDGVDLK.T |
| **Lyn** | LYN | Y501 | **7.4** | **56.2** | K.AEERPTFDYLQSVLDDFY*TATEGQYQQQP.- |
| **Lyn** | LYN | T502 | **6.5** | **54.7** | K.AEERPTFDYLQSVLDDFYT*ATEGQYQQQP.- |
| **Lyn** | LYN | S11 | **13.3** | **62.9** | K.DS*LSDDGVDLK.T |
| **LysM, peptidoglycan-binding, domain containing 2** | LYSMD2 | S33 | **-0.2** | **77.1** | R.SGS*ESEEAELSLSLAR.T |
| **Lysosomal trafficking regulator** | LYST | S2105 | **61.9** | **80.4** | R.S*LPAFPTSSLLTQSQK.L |

Peak Area: <-10, -5, -3, 0, 3, 5, >10

%CV: 0, 29, 42, 71, 86, >100

**White dots-** Significant change in peptide abundance at 5 %FDR compared to the timepoint with the minimum peak area for a given PSM

CarT: 0m, 2m, 5m — RajiB: 0m, 2m, 5m

| Protein Name | Gene | Phosphosites | Ascor | MOWSE | Sequence |
|---|---|---|---|---|---|
| M phase phosphoprotein 9 | MPHOSPH | S854 | 61.2 | 19.7 | K.NES*SPIRFDILLDDLDTVPVSTLQR.T |
| M phase phosphoprotein, mpp8 | MPHOSPH | S51 | 100.0 | 112.8 | R.GAEAFGDS*EEDGEDVFEVEK.I |
| M phase phosphoprotein, mpp8 | MPHOSPH | S188S189 | 21.0 | 46.3 | K.SKPDLES*S*LESLVFDLR.T |
| M phase phosphoprotein, mpp8 | MPHOSPH | S188S192 | 12.8 | 33.9 | K.SKPDLES*SLES*LVFDLR.T |
| M phase phosphoprotein, mpp8 | MPHOSPH | S136S138 | 10.0 | 60.3 | R.LSLNNDIFEANS*DS*DQQSETK.E |
| M phase phosphoprotein, mpp8 | MPHOSPH | S189S192 | 7.3 | 60.9 | K.SKPDLESS*LES*LVFDLR.T |
| M phase phosphoprotein, mpp8 | MPHOSPH | S392 | 100.0 | 11.2 | R.RLS*GEER.G |
| M phase phosphoprotein, mpp8 | MPHOSPH | S188S189S192 | 3.5 | 28.8 | K.SKPDLES*S*LES*LVFDLR.T |
| Macrophin 1 | MACF1 | S2956 | 100.0 | 36.1 | K.AFLAELEQNS*PK.I |
| Macrophin 1 | MACF1 | S5829 | 7.2 | 21.5 | K.LKRPTPTFHSS*R.T |
| Macrophin 1 | MACF1 | S5828 | 29.8 | 26.0 | K.LKRPTPTFHS*SR.T |
| | ZCRB1 | S155 | 42.5 | 76.0 | K.APEPEEEIEEVEES*EDEGEDPALDSLSQAIAFQQAK.I |
| MADP 1 protein | ZCRB1 | S210S216 | 12.4 | 26.2 | K.STYFS*DEEELS*D.- |
| MADP 1 protein | ZCRB1 | Y208S216 | 7.7 | 18.7 | K.STY*FSDEEELS*D.- |
| Maestro heat-like repeat-containing protein family member 6 (Fragment) | MROH6 | T506 | 100.0 | 11.4 | R.HT*GLNPR.L |
| Major vault protein | MVP | S873 | 28.2 | 75.9 | R.VASGPSPGEGIS*PQSAQAPQAPGDNHVVPVLR.- |
| Major vault protein | MVP | S876 | 12.0 | 29.5 | R.VASGPSPGEGISPQS*AQAPQAPGDNHVVPVLR.- |
| Major vault protein | MVP | S864 | 3.5 | 24.9 | R.VAS*GPSPGEGISPQSAQAPQAPGDNHVVPVLR.- |
| Major vault protein | MVP | S867 | 5.6 | 29.6 | R.VASGPS*PGEGISPQSAQAPQAPGDNHVVPVLR.- |
| Male specific lethal 3 like 1 | MSL3 | S400T405 | 6.4 | 16.1 | R.SSS*PIPLT*PSKEGSAVFAGFEGR.R |
| Male-specific lethal 2-like 1 | MSL2 | S447 | 9.5 | 41.6 | K.EKIPSHHFMPGS*PTK.T |
| Male-specific lethal-1 homolog | MSL1 | S99 | 121.5 | 51.5 | K.HS*PIKEEPCGSLSETVCK.R |
| MAN antigen 1 | LEMD3 | S259S261 | 6.6 | 69.8 | R.ENYS*DS*EEEDDDDVASSR.Q |
| MAN antigen 1 | LEMD3 | Y258S259 | 6.5 | 102.2 | R.ENY*S*DSEEEDDDDVASSR.Q |
| MAN antigen 1 | LEMD3 | S144 | 9.1 | 45.6 | K.VLLGFSSDES*DVEASPR.D |
| MAN antigen 1 | LEMD3 | Y258S261 | 77.7 | 115.6 | R.ENY*SDS*EEEDDDDVASSR.Q |
| MAP kinase activating death domain protein | MADD | S1239 | 21.2 | 148.5 | R.GTLS*DSEIETNSATSTIFGK.A |
| MAP kinase activating death domain protein | MADD | S1241 | -0.2 | 73.9 | R.GTLSDS*EIETNSATSTIFGK.A |
| MAP kinase activating death domain protein | MADD | T1237 | 9.2 | 59.6 | R.GT*LSDSEIETNSATSTIFGK.A |
| MAP2K2 | MAP2K2 | T396 | 25.6 | 31.7 | R.LNQPGTPT*R.T |
| MAP2K2 | MAP2K2 | T394 | | 32.8 | R.LNQPGT*PTR.T |
| MAP3K2 | MAP3K2 | S163 | 77.8 | 93.2 | R.DRS*SPPPGYIPDELHQVAR.N |
| MAP3K2 | MAP3K2 | S239 | 10.3 | 35.8 | R.AQS*YPDNHQEFSDYDNPIFEK.F |
| MAP3K2 | MAP3K2 | S153 | 67.4 | 45.0 | K.RLS*IIGPTSR.D |
| MAP3K2 | MAP3K2 | S164 | 5.8 | 73.6 | R.DRSS*PPPGYIPDELHQVAR.N |
| MAP3K4 | MAP3K4 | S66S84 | 17.1 | 16.9 | K.SPES*DLEDFSDETNTENLYGTS*PPSTPR.Q |
| MAP3K4 | MAP3K4 | S66T77S84 | 11.3 | 13.6 | K.SPES*DLEDFSDETNT*ENLYGTS*PPSTPR.Q |
| MAP3K4 | MAP3K4 | S66T83 | 12.8 | 26.7 | K.SPES*DLEDFSDETNTENLYGT*SPPSTPR.Q |
| MAP3K4 | MAP3K4 | S499 | 2.0 | 14.4 | R.LES*EDDSLGWGAPDWSTEAGFSR.H |
| MAP3K4 | MAP3K4 | S503 | 2.0 | 24.6 | R.LESEDDS*LGWGAPDWSTEAGFSR.H |
| MAP3K7 | MAP3K7 | S439 | 39.3 | 68.8 | R.S*IQDLTVTGTEPGQVSSR.S |
| MAP3K7 | MAP3K7 | S417 | -0.9 | 53.4 | K.TAS*FGNILDVPEIVISGNGQPR.R |
| MAP4K4 | MAP4K4 | S981 | 79.5 | 20.4 | R.KGS*VVNVNPTNTRPQSDTPEIR.K |
| MAP4K4 | MAP4K4 | S787 | -0.2 | 42.4 | R.SS*SKSEGSPSQR.L |
| MAP4K4 | MAP4K4 | T705S708 | 5.7 | 15.1 | R.TT*SRS*PVLSR.R |
| MAP4K4 | MAP4K4 | S934 | -0.1 | 71.4 | K.SS*SSFTPFIDPR.L |
| MAP4K4 | MINK1 | T187 | 12.7 | 75.3 | R.NT*FIGTPYWMAPEVIACDENPDATYDYR.S |
| MAP4K4 | MAP4K4 | S933 | 22.8 | 29.8 | K.HKS*SSSFTPFIDPR.L |
| MAP4K4 | MAP4K4 | S936T938 | 4.4 | 16.9 | K.HKSSSS*FT*PFIDPR.L |
| MAP4K4 | MAP4K4 | S933S934 | 9.3 | 22.0 | K.HKS*S*SSFTPFIDPR.L |
| MAP4K4 | MAP4K4 | S933S936 | 14.2 | 63.6 | K.HKS*SSS*FTPFIDPR.L |
| MAP4K4 | MAP4K4 | T319S324 | 3.1 | 26.7 | K.DET*EYEYS*GSEEEEEEVPEQEGEPSSIVNVPGESTLR.R |
| MAP4K4 | MAP4K4 | S324S326 | 9.6 | 28.8 | K.DETEYEYS*GS*EEEEEEVPEQEGEPSSIVNVPGESTLR.R |
| MAP4K4 | MAP4K4 | Y323S324 | 6.8 | 44.6 | K.DETEYEY*S*GSEEEEEEVPEQEGEPSSIVNVPGESTLR.R |
| MAP4K4 | MAP4K4 | T705S706 | 8.6 | 11.0 | R.TT*S*RSPVLSR.R |
| MAP4K4 | MAP4K4 | S935S936 | 6.4 | 14.5 | K.HKSSS*S*FTPFIDPR.L |
| MAP4K4 | MAP4K4 | S934S935 | 4.3 | 11.8 | K.HKSS*S*SFTPFIDPR.L |
| MAP4K4 | MAP4K4 | T319S326 | 7.6 | 14.2 | K.DET*EYEYSGS*EEEEEEVPEQEGEPSSIVNVPGESTLR.R |
| MAP4K4 | MAP4K4 | Y323S326 | 12.4 | 31.2 | K.DETEYEY*SGS*EEEEEEVPEQEGEPSSIVNVPGESTLR.R |
| MAP4K4 | MAP4K4 | S788 | 11.7 | 28.1 | R.SSS*KSEGSPSQR.L |
| MAP4K4 | MAP4K4 | T704T705 | 8.5 | 14.8 | R.T*T*SRSPVLSR.R |
| MAP7 domain containing 1 | MAP7D1 | S544S548T554 | 5.1 | 22.4 | K.ESAAPAS*PAPS*PAPSPT*PAPPQK.E |

**Peak Area**: <-10, -5, -3, 0, 3, 5, >10

**%CV**: 0, 29, 42, 71, 86, >100

**White dots**- Significant change in peptide abundance at **5** %FDR compared to the timepoint with the minimum peak area for a given PSM

CarT: 0m 2m 5m — RajiB: 0m 2m 5m

| Protein Name | Gene | Phosphosites | Ascor | MOWSE | Sequence |
|---|---|---|---|---|---|
| **MAP7 domain containing 1** | MAP7D1 | S544S548S552 | 22.7 | 26.6 | K.ESAAPAS*PAPS*PAPS*PTPAPPQK.E |
| **MAP7 domain containing 1** | MAP7D1 | T813 | 26.8 | 65.8 | R.T*PETLLPFAEAEAFLK.K |
| **MAP7 domain containing 1** | MAP7D1 | T816 | 13.9 | 28.9 | R.TPET*LLPFAEAEAFLK.K |
| **MAP7 domain containing 1** | MAP7D1 | S442 | 48.2 | 15.0 | K.RQS*LPASPRAR.L |
| **MAP7 domain containing 1** | MAP7D1 | S539S544T554 | 7.3 | 12.3 | K.ES*AAPAS*PAPSPAPSPT*PAPPQK.E |
| **MAPK14** | MAPK14 | Y182 | 17.8 | 72.3 | R.HTDDEMTGY*VATR.W |
| **MAPK14** | MAPK14 | T180 | 10.2 | 54.9 | R.HTDDEMT*GYVATR.W |
| **MARCKS like protein** | MARCKSL | T122 | 8.1 | 87.3 | K.EGGGDSSASSPT*EEEQEQGEIGACSDEGTAQEGK.A |
| **MARCKS like protein** | MARCKSL | S119 | 3.0 | 55.2 | K.EGGGDSSAS*SPTEEEQEQGEIGACSDEGTAQEGK.A |
| **MARCKS like protein** | MARCKSL | S116S119 | 10.7 | 126.3 | K.EGGGDS*SAS*SPTEEEQEQGEIGACSDEGTAQEGK.A |
| **MARCKS like protein** | MARCKSL | S116S117 | 46.0 | 119.7 | K.EGGGDS*S*ASSPTEEEQEQGEIGACSDEGTAQEGK.A |
| **MARCKS like protein** | MARCKSL | S177 | 29.8 | 124.9 | K.GAEASAASEEEAGPQATEPS*TPSGPESGPTPASAEQNE.- |
| **MARCKS like protein** | MARCKSL | T178 | 6.8 | 90.7 | K.GAEASAASEEEAGPQATEPST*PSGPESGPTPASAEQNE.- |
| **MARCKS like protein** | MARCKSL | T174 | 6.6 | 116.6 | K.GAEASAASEEEAGPQAT*EPSTPSGPESGPTPASAEQNE.- |
| **MARCKS like protein** | MARCKSL | S93 | 100.0 | 33.0 | K.KFS*FKKPFK.L |
| **MARCKS like protein** | MARCKSL | S165 | 19.1 | 92.4 | K.GAEASAAS*EEEAGPQATEPSTPSGPESGPTPASAEQNE.- |
| **MARCKS like protein** | MARCKSL | S162S165 | 8.8 | 61.7 | K.GAEAS*AAS*EEEAGPQATEPSTPSGPESGPTPASAEQNE.- |
| **MARCKS like protein** | MARCKSL | S177T178 | -0.2 | 33.5 | K.GAEASAASEEEAGPQATEPS*T*PSGPESGPTPASAEQNE.- |
| **MARCKS like protein** | MARCKSL | S104 | 67.4 | 33.0 | K.LSGLS*FK.R |
| **MARCKS like protein** | MARCKSL | S101 | 14.9 | 36.2 | K.KPFKLS*GLSFK.R |
| **MARCKS like protein** | MARCKSL | S101S104 | 100.0 | 44.6 | K.KPFKLS*GLS*FK.R |
| **MARCKS like protein** | MARCKSL | S36 | 78.3 | 78.3 | K.S*NGDLSPK.G |
| **MARCKS like protein** | MARCKSL | S22 | 146.6 | 73.1 | R.GDVTAEEAAGAS*PAK.A |
| **MARCKS like protein** | MARCKSL | T174S177 | 2.2 | 73.2 | K.GAEASAASEEEAGPQAT*EPS*TPSGPESGPTPASAEQNE.- |
| **MARCKS like protein** | MARCKSL | S116S117S120 | 7.2 | 60.5 | K.EGGGDS*S*ASS*PTEEEQEQGEIGACSDEGTAQEGK.A |
| **MARCKS like protein** | MARCKSL | S162 | | 57.4 | K.GAEAS*AASEEEAGPQATEPSTPSGPESGPTPASAEQNE.- |
| **MARCKS like protein** | MARCKSL | S116S119T122 | 4.5 | 126.3 | K.EGGGDS*SAS*SPT*EEEQEQGEIGACSDEGTAQEGK.A |
| **MARCKS like protein** | MARCKSL | S177S184 | -0.2 | 45.5 | K.GAEASAASEEEAGPQATEPS*TPSGPES*GPTPASAEQNE.- |
| **MARCKS like protein** | MARCKSL | S117S119 | 3.5 | 107.4 | K.EGGGDSS*AS*SPTEEEQEQGEIGACSDEGTAQEGK.A |
| **MARCKS like protein** | MARCKSL | S116S135 | 11.1 | 45.5 | K.EGGGDS*SASSPTEEEQEQGEIGACS*DEGTAQEGK.A |
| **MARCKS like protein** | MARCKSL | S190 | 16.3 | 33.2 | K.GAEASAASEEEAGPQATEPSTPSGPESGPTPAS*AEQNE.- |
| **MARCKS like protein** | MARCKSL | S116S119S120 | 12.5 | 72.0 | K.EGGGDS*SAS*S*PTEEEQEQGEIGACSDEGTAQEGK.A |
| **MARCKS like protein** | MARCKSL | S116S117T122 | 6.7 | 76.2 | K.EGGGDS*S*ASSPT*EEEQEQGEIGACSDEGTAQEGK.A |
| **MARCKS like protein** | MARCKSL | S135 | 13.6 | 40.4 | K.EGGGDSSASSPTEEEQEQGEIGACS*DEGTAQEGK.A |
| **MARCKS like protein** | MARCKSL | S116S120 | 7.3 | 75.2 | K.EGGGDS*SASS*PTEEEQEQGEIGACSDEGTAQEGK.A |
| **MARCKS like protein** | MARCKSL | S180 | 2.4 | 82.5 | K.GAEASAASEEEAGPQATEPSTPS*GPESGPTPASAEQNE.- |
| **MARCKS like protein** | MARCKSL | S116T122 | 17.4 | 107.5 | K.EGGGDS*SASSPT*EEEQEQGEIGACSDEGTAQEGK.A |
| **MARCKS like protein** | MARCKSL | S117 | 4.3 | 59.7 | K.EGGGDSS*ASSPTEEEQEQGEIGACSDEGTAQEGK.A |
| **MARCKS like protein** | MARCKSL | S119S120 | 2.8 | 102.4 | K.EGGGDSSAS*S*PTEEEQEQGEIGACSDEGTAQEGK.A |
| **Mastermind like 1** | MAML1 | S1015 | 17.0 | 32.0 | R.TSEEWMSDLDDLLGS*Q.- |
| **MASTL** | MASTL | S370 | 43.9 | 28.8 | K.DLELALS*PIHNSSALPTTGR.S |
| **Maternal embryonic leucine zipper kinase** | MELK | S505 | 135.8 | 109.0 | R.S*VELDLNQAHMEETPK.R |
| **Matrin 3** | MATR3 | T741 | | 25.6 | K.[illegible] |
| | MATR3 | S598 | 30.2 | 38.6 | R.SYS*PDGK.E |
| **Matrin 3** | MATR3 | S596S598 | 9.8 | 29.7 | R.KRS*YS*PDGK.E |
| **Matrin 3** | MATR3 | S598S604 | 16.7 | 47.5 | R.SYS*PDGKES*PSDKK.S |
| **Matrin 3** | MATR3 | Y597S604 | 7.1 | 21.8 | R.SY*SPDGKES*PSDKK.S |
| **Matrin 3** | MATR3 | S195 | 10.5 | 33.9 | R.DSFDDRGPS*LNPVLDYDHGSR.S |
| **Matrin 3** | MATR3 | S188 | 100.0 | 29.1 | R.RDS*FDDR.G |
| **Matrin 3** | MATR3 | T644S654S671 | | 27.1 | K.[illegible] |
| **Matrin 3** | MATR3 | S654S673S674 | 6.1 | 44.1 | K.[illegible] |
| **Matrin 3** | MATR3 | S596Y597 | | 24.4 | R.KRS*Y*SPDGK.E |
| **MBC2** | ESYT1 | Y822 | 3.5 | 42.9 | K.HLSPY*ATLTVGDSSHK.T |
| **MBC2** | ESYT1 | S820 | 7.3 | 53.3 | K.HLS*PYATLTVGDSSHK.T |
| **MBD5** | MBD5 | S1493 | 100.0 | 19.7 | R.RKIS*R.- |
| | MCM2 | T39S40S41 | 35.6 | 27.8 | R.TDALT*S*S*PGR.D |
| **MCM2 minichromosome maintenance deficient 2, mitotin** | MCM2 | S27 | 12.1 | 53.5 | R.GNDPLTSS*PGR.S |
| **MCM2 minichromosome maintenance deficient 2, mitotin** | MCM2 | Y137 | | 24.5 | R.GLLY*DSDEEDEERPARK.R |
| **MCM2 minichromosome maintenance deficient 2, mitotin** | MCM2 | S139 | 32.7 | 65.2 | R.GLLYDS*DEEDEERPAR.K |
| **MCM2 minichromosome maintenance deficient 2, mitotin** | MCM2 | S108 | 20.7 | 98.9 | R.AIPELDAYEAEGLALDDEDVEELTAS*QR.E |
| **MCM2 minichromosome maintenance deficient 2, mitotin** | MCM2 | T106 | 14.0 | 44.4 | R.AIPELDAYEAEGLALDDEDVEELT*ASQR.E |
| **MCM2 minichromosome maintenance deficient 2, mitotin** | MCM2 | S53 | 20.0 | 93.7 | R.DLPPFEDES*EGLLGTEGPLEEEEDGEELIGDGMER.D |

Peak Area: <-10, -5, -3, 0, 3, 5, >10

%CV: 0, 29, 42, 71, 86, >100

**White dots-** Significant change in peptide abundance at **5** %FDR compared to the timepoint with the minimum peak area for a given PSM

| Protein Name | Gene | Phosphosites | CarT 0m | CarT 2m | CarT 5m | RajiB 0m | RajiB 2m | RajiB 5m | Ascor | MOWSE | Sequence |
|---|---|---|---|---|---|---|---|---|---|---|---|
| **MCM2 minichromosome maintenance deficient 2. mitotin** | MCM2 | T59 | | | | | | | **24.1** | **93.5** | R.DLPPFEDESEGLLGT*EGPLEEEEDGEELIGDGMER.D |
| **MCM2 minichromosome maintenance deficient 2. mitotin** | MCM2 | S53T59 | | | | | | | **100.0** | **81.9** | R.DLPPFEDES*EGLLGT*EGPLEEEEDGEELIGDGMER.D |
| **MCM2 minichromosome maintenance deficient 2. mitotin** | MCM2 | S41 | | | | | | | **20.1** | **56.7** | R.RTDALTSS*PGR.D |
| | MCM2 | S40 | | | | | | | **20.1** | **59.0** | R.TDALTS*SPGR.D |
| **MCM2 minichromosome maintenance deficient 2. mitotin** | MCM2 | S40S41 | | | | | | | **35.8** | **35.5** | R.TDALTS*S*PGR.D |
| **MCM2 minichromosome maintenance deficient 2. mitotin** | MCM2 | T39S40 | | | | | | | **25.9** | **20.3** | R.TDALT*S*SPGR.D |
| **MCM2 minichromosome maintenance deficient 2. mitotin** | MCM2 | Y90 | | | | | | | **12.9** | **15.4** | R.AIPELDAY*EAEGLALDDEDVEELTASQR.E |
| **MCM2 minichromosome maintenance deficient 2. mitotin** | MCM2 | S26S27 | | | | | | | **12.5** | **11.9** | R.GNDPLTS*S*PGR.S |
| **MCM2 minichromosome maintenance deficient 2. mitotin** | MCM2 | S26 | | | | | | | **10.8** | **11.6** | R.RGNDPLTS*SPGR.S |
| **MCM3** | MCM3 | Y708T722 | | | | | | | **23.3** | **24.9** | K.DGDSYDPY*DFSDTEEEM#PQVHT*PK.T |
| **MCM3** | MCM3 | S711T722 | | | | | | | **15.6** | **17.0** | K.DGDSYDPYDFS*DTEEEM#PQVHT*PK.T |
| **MCM3** | MCM3 | S711 | | | | | | | **14.5** | **64.4** | K.DGDSYDPYDFS*DTEEEMPQVHTPK.T |
| **MCM3** | MCM3 | Y708S711T722 | | | | | | | **47.3** | **17.3** | K.DGDSYDPY*DFS*DTEEEMPQVHT*PK.T |
| **MCM3** | MCM3 | S160 | | | | | | | **7.2** | **64.0** | R.RYS*DLTTLVAFPSSSVYPTK.D |
| | MCM3 | S672T674 | | | | | | | **34.8** | **63.3** | R.SEDES*ET*EDEEEKSQEDQEQKR.K |
| **MCM3** | MCM3 | S711T713T722 | | | | | | | **15.2** | **11.0** | K.DGDSYDPYDFS*DT*EEEM#PQVHT*PK.T |
| **MCM3** | MCM3 | Y708 | | | | | | | **26.8** | **55.7** | K.DGDSYDPY*DFSDTEEEMPQVHTPK.T |
| **MCM3** | MCM3 | Y708S711 | | | | | | | **8.6** | **26.1** | K.DGDSYDPY*DFS*DTEEEMPQVHTPK.T |
| **MCM3** | MCM3 | Y705T713T722 | | | | | | | **11.6** | **12.1** | K.DGDSY*DPYDFSDT*EEEMPQVHT*PK.T |
| | MCM3 | S711T713 | | | | | | | **3.2** | **11.3** | K.DGDSYDPYDFS*DT*EEEM#PQVHTPK.T |
| **MCM3** | MCM3 | Y708T713 | | | | | | | **9.4** | **38.2** | K.DGDSYDPY*DFSDT*EEEMPQVHTPK.T |
| **MCM3** | MCM3 | Y708T713T722 | | | | | | | **24.3** | **13.7** | K.DGDSYDPY*DFSDT*EEEMPQVHT*PK.T |
| **MCM6** | MCM6 | S804 | | | | | | | **4.6** | **25.4** | K.GSTEGSES*YEEDPYLVVNPNYLLED.- |
| **Mediator of DNA damage checkpoint 1** | MDC1 | S299T301 | | | | | | | **16.1** | **11.0** | R.SQPPGEDS*DT*DVDDDSRPPGRPAEVHLER.A |
| **Mediator of DNA damage checkpoint 1** | MDC1 | S453T455 | | | | | | | **35.4** | **65.3** | R.SQTTTERDS*DT*DVEEEELPVENR.E |
| **Mediator of DNA damage checkpoint 1** | MDC1 | S376 | | | | | | | **14.4** | **19.8** | R.GPGAPGLAHLQESQAGS*DTDVEEGKAPQAVPLEK.S |
| **Mediator of DNA damage checkpoint 1** | MDC1 | S376T378 | | | | | | | **19.6** | **33.9** | R.GPGAPGLAHLQESQAGS*DT*DVEEGKAPQAVPLEK.S |
| **Mediator of DNA damage checkpoint 1** | MDC1 | S329T331 | | | | | | | **79.3** | **24.3** | R.AQPFGFIDS*DT*DAEEERIPATPVVIPMK.K |
| **Mediator of DNA damage checkpoint 1** | MDC1 | S402T404 | | | | | | | **44.0** | **140.5** | K.SQASMVINS*DT*DDEEEVSAALTLAHLK.E |
| **Mediator of DNA damage checkpoint 1** | MDC1 | S168 | | | | | | | **121.5** | **125.9** | R.LLLAEDS*EEEVDFLSER.R |
| **Mediator of DNA damage checkpoint 1** | MDC1 | S1775 | | | | | | | **72.7** | **54.8** | R.AAESLTAIPEPAS*PQLLETPIHASQIQK.V |
| **Mediator of DNA damage checkpoint 1** | MDC1 | S329 | | | | | | | **6.3** | **16.2** | R.AQPFGFIDS*DTDAEEERIPATPVVIPMK.K |
| **Mediator of DNA damage checkpoint 1** | MDC1 | S372S376 | | | | | | | | **18.0** | R.GPGAPGLAHLQES*QAGS*DTDVEEGKAPQAVPLEK.S |
| **Mediator of DNA damage checkpoint 1** | MDC1 | S397T404 | | | | | | | **6.0** | **62.6** | K.SQAS*MVINSDT*DDEEEVSAALTLAHLK.E |
| **Mediator of DNA damage checkpoint 1** | MDC1 | S372T378 | | | | | | | **8.6** | **16.9** | R.GPGAPGLAHLQES*QAGSDT*DVEEGKAPQAVPLEK.S |
| **Mediator of DNA damage checkpoint 1** | MDC1 | S329T341 | | | | | | | **13.2** | **11.4** | R.AQPFGFIDS*DTDAEEERIPAT*PVVIPM#KK.R |
| **Mediator of DNA damage checkpoint 1** | MDC1 | T448S453 | | | | | | | **3.4** | **13.3** | R.SQTT*TERDS*DTDVEEEELPVENR.E |
| **Mediator of RNA polymerase II transcription subunit 19** | MED19 | S226 | | | | | | | **58.0** | **26.6** | K.NRHS*PDHPGMGSSQASSSSSLR.- |
| **MEK1** | MAP2K2 | S218S222 | | | | | | | **53.9** | **81.4** | K.LCDFGVSGQLIDS*MANS*FVGTR.S |
| **MEK1** | MAP2K1 | T292S304 | | | | | | | **8.6** | **36.5** | R.T*PGRPLSSYGMDS*RPPMAIFELLDYIVNEPPPK.L |
| **MEK1** | MAP2K1 | T292S298 | | | | | | | | **21.3** | R.T*PGRPLS*SYGMDSRPPMAIFELLDYIVNEPPPK.L |
| **Melanoma antigen family F 1** | MAGEF1 | S95 | | | | | | | **28.2** | **28.0** | K.KS*PITR.S |
| **Melanoma differentiation associated protein 5** | IFIH1 | S301 | | | | | | | **78.5** | **50.3** | R.AS*PEPELQLRPYQMEVAQPALEGK.N |
| **Melanoma inhibitory activity family, member 3** | MIA3 | S1739 | | | | | | | **62.9** | **46.6** | R.WSAEASGKPSPSDPGSGTATMMNS*SSR.G |
| **Melanoma inhibitory activity family, member 3** | MIA3 | S1741 | | | | | | | **9.1** | **53.3** | R.WSAEASGKPSPSDPGSGTATMMNSSS*R.G |
| **Melanoma inhibitory activity family, member 3** | MIA3 | S1740 | | | | | | | **9.0** | **43.7** | R.WSAEASGKPSPSDPGSGTATMMNSS*SR.G |
| **Melanoma inhibitory activity family, member 3** | MIA3 | S1721 | | | | | | | **12.9** | **12.9** | R.WSAEAS*GKPSPSDPGSGTATMMNSSSR.G |
| **Melanoregulin** | MREG | S213 | | | | | | | **75.8** | **34.4** | K.ELHYLPFPS*P.- |
| **Membrane component, chromosome 11, surface marker 1** | CAPRIN1 | T317 | | | | | | | | **12.4** | K. EQVDEWT*VETVEVVNSLQQQPQAASPSVPEPHSLTPVAQADPLVR |
| **Meningioma expressed antigen 5** | MGEA5 | S364 | | | | | | | **119.8** | **91.7** | K.LENEGS*DEDIETDVLYSPQMALK.L |
| **Mesoderm induction early response 1** | MIER1 | S507S512 | | | | | | | **22.0** | **78.1** | R.VNS*NGKES*PGSSEFFQEAVSHGK.F |
| **Mesoderm induction early response 1** | MIER1 | T182S190 | | | | | | | **12.9** | **60.4** | K.YFDT*NSEVEEES*EEDEDYIPSEDWK.K |
| **Mesoderm induction early response 1** | MIER1 | S184S190 | | | | | | | **23.7** | **55.0** | K.YFDTNS*EVEEES*EEDEDYIPSEDWK.K |
| **Mesoderm induction early response 1** | MIER1 | Y179S190 | | | | | | | **10.1** | **46.5** | K.Y*FDTNSEVEEES*EEDEDYIPSEDWK.K |
| **Mesoderm induction early response 1** | MIER1 | T182S184 | | | | | | | **4.4** | **29.3** | K.YFDT*NS*EVEEESEEDEDYIPSEDWK.K |
| **Metadherin** | MTDH | S426 | | | | | | | **79.4** | **52.0** | K.SQEPIPDDQKVS*DDDKEK.G |
| **Metadherin** | MTDH | S344S347 | | | | | | | **3.1** | **79.0** | R.SWSDRS*IFS*GIGSTAEPVSQSTTSDYQWDVSR.N |
| **Metastasis associated protein MTA1** | MTA1 | S562 | | | | | | | **20.1** | **31.2** | K.SVSSVLSS*LTPAK.V |
| **Metastasis associated protein MTA1** | MTA1 | T564 | | | | | | | **19.8** | **54.6** | K.SVSSVLSSLT*PAK.V |
| **Metastasis associated protein MTA1** | MTA1 | S449 | | | | | | | **3.9** | **19.4** | R.SNMS*PHGLPAR.S |
| **Metastasis associated protein MTA2** | MTA2 | S435 | | | | | | | **33.2** | **79.7** | R.GHLSRPEAQSLS*PYTTSANR.A |
| **Metastasis associated protein MTA2** | MTA2 | T505 | | | | | | | **100.0** | **35.7** | K.T*PLKIHPLVR.L |

Peak Area: <-10, -5, -3, 0, 3, 5, >10

%CV: 0, 29, 42, 71, 86, >100

**White dots-** Significant change in peptide abundance at 5 %FDR compared to the timepoint with the minimum peak area for a given PSM

CarT: 0m 2m 5m; RajiB: 0m 2m 5m

| Protein Name | Gene | Phosphosites | Ascor | MOWSE | Sequence |
|---|---|---|---|---|---|
| Metastasis associated protein MTA2 | MTA2 | Y437 | 25.6 | 27.9 | R.GHLSRPEAQSLSPY*TTSANR.A |
| Methionyl tRNA synthetase | MARS | S825 | 13.6 | 66.0 | K.TS*PKPAVVETVTTAKPQQIQALMDEVTK.Q |
| Methionyl tRNA synthetase | MARS | T824 | 7.4 | 49.4 | K.T*SPKPAVVETVTTAKPQQIQALMDEVTK.Q |
| Methyl CpG binding domain protein 1 | MBD1 | S408 | 20.2 | 18.2 | R.EGAS*SPVQVPGPVAASTEALLQAVDPGLPSVK.Q |
| Methyl CpG binding domain protein 2 | MBD2 | S407 | 93.2 | 25.2 | R.AADTEEMDIEMDS*GDEA.- |
| Methyltransferase 10 domain containing | METTL16 | S329 | 31.7 | 21.1 | K.ELSLKAS*PLR.S |
| Methyltransferase like 6 | METTL6 | S53T54 | 100.0 | 24.5 | K.RNS*T*NFFK.D |
| MGC13125 | BUD13 | T159S163 | 100.0 | 28.8 | R.HDT*PDPS*PLR.G |
| MGC13125 | BUD13 | S271 | 139.2 | 59.5 | R.ARHDS*PDLAPNVTYSLPR.T |
| MGC13125 | BUD13 | S248 | 32.3 | 11.1 | R.VHNNS*PDTSR.R |
| MGC13125 | BUD13 | T135S139 | 100.0 | 34.4 | R.HGT*PDPS*PR.K |
| MGC13125 | BUD13 | S364S370S372 | 100.0 | 13.7 | R.HKQS*PGHQDS*DS*DLS*PPR.N |
| MGC13125 | BUD13 | T147S151 | 100.0 | 29.4 | R.HDT*PDPS*PR.R |
| MGC13125 | BUD13 | T352S354S357 | 17.3 | 22.2 | K.AT*DS*DLS*SPR.H |
| MGC20460 | RBM33 | S293S295 | 11.5 | 13.2 | R.LAQQQQQLYAPPPPAEQEEQALS*PS*PTNGNPLLPFPGAQVR.Q |
| MGC20460 | RBM33 | S293T297 | 4.1 | 17.9 | R.LAQQQQQLYAPPPPAEQEEQALS*PSPT*NGNPLLPFPGAQVR.Q |
| MGC20460 | RBM33 | Y279 |  | 25.1 | R.LAQQQQQLY*APPPPAEQEEQALSPSPTNGNPLLPFPGAQVR.Q |
| MGC20460 | RBM33 | S293 | 12.7 | 20.6 | R.LAQQQQQLYAPPPPAEQEEQALS*PSPTNGNPLLPFPGAQVR.Q |
| MGC51029 | FUNDC1 | S13 | 12.1 | 48.1 | R.NPPPQDYES*DDDSYEVLDLTEYAR.R |
| MGC51029 | FUNDC1 | Y11 |  | 26.0 | R.NPPPQDY*ESDDDSYEVLDLTEYAR.R |
| MGC51029 | FUNDC1 | S17 | 0.5 | 40.4 | R.NPPPQDYESDDDS*YEVLDLTEYAR.R |
| MGC874 | PBDC1 | S181 | 49.5 | 19.2 | K.RADS*GEEENTK.N |
| MGC874 | PBDC1 | S197 | 100.0 | 23.4 | K.GADS*GEEKEEGINR.E |
| MGEA6 | CTAGE4 | S139 | 8.6 | 63.1 | R.SNS*ELEDEILCLEK.D |
| MICA like 1 | MICALL1 | T469 | 32.3 | 25.5 | K.SLHPWYGITPT*SSPK.T |
| MICA like 1 | MICALL1 | S471 | 8.2 | 36.7 | K.SLHPWYGITPTSS*PK.T |
| MICA like 1 | MICALL1 | S470 | 9.8 | 25.3 | K.SLHPWYGITPTS*SPK.T |
| MICA like 1 | MICALL1 | T467S471 | 23.8 | 18.9 | K.SLHPWYGIT*PTSS*PK.T |
| MICA like 1 | MICALL1 | T467S470 | 18.5 | 18.5 | K.SLHPWYGIT*PTS*SPK.T |
|  | MFAP1 | S132S133 | 100.0 | 145.3 | R.EDS*S*EEEEEEIDDEEIER.R |
| Microfibrillar associated protein 1 | MFAP1 | S52S53 | 65.4 | 86.4 | K.RPDYAPMES*S*DEEDEEFQFIKK.A |
| Microfibrillar associated protein 1 | MFAP1 | S116S118 | 100.0 | 68.0 | K.IVEPEVVGES*DS*EVEGDAWR.M |
| Microfibrillar associated protein 1 | MFAP1 | T267 | 105.2 | 114.8 | R.SLAALDALNT*DDENDEEEYEAWK.V |
| Microspherule protein 1 | MCRS1 | S354 | 84.8 | 66.4 | K.GDQVLNFS*DAEDLIDDSKLK.D |
| Microtubule associated protein 1A | MAP1A | S2019S2022 | 13.5 | 23.8 | K.ELSS*PIS*PK.S |
| Microtubule associated protein 1A | MAP1A | S2108 | 22.4 | 32.7 | R.SHWDDSTSDS*ELEK.G |
| Microtubule associated protein 1A | MAP1A | S2104T2105 | 14.5 | 55.3 | R.SHWDDS*T*SDSELEK.G |
| Microtubule associated protein 1A | MAP1A | S1172 | 53.9 | 63.7 | K.S*PCGLTEQYLHK.D |
| Microtubule associated protein 1A | MAP1A | S667 | 24.7 | 24.2 | K.AELEEMEEVHPS*DEEEEDATK.A |
| Microtubule associated protein 1A | MAP1A | S1654 | 13.9 | 64.1 | R.GQDVVQEWQETS*PTR.E |
| Microtubule associated protein 1A | MAP1A | S1675 | 10.8 | 63.8 | K.ELAPAWEDTS*PEQDNR.Y |
| Microtubule associated protein 1A | MAP1A | S1069 | 124.2 | 91.3 | K.VPPPRS*PQAQEAPVNIDEGLTGCTIQLLPAQDK.A |
| Microtubule associated protein 1A | MAP1A | S2449 | 15.2 | 72.8 | R.GELS*PSFLNPPLPPSIDDR.D |
| Microtubule associated protein 1A | MAP1A | S526S527 | 100.4 | 93.8 | R.ELVLS*S*PEDLTQDFEEMKR.E |
| Microtubule associated protein 1A | MAP1A | S1218 | 48.8 | 98.8 | K.QLS*PESLGTLQFGELNLGK.E |
| Microtubule associated protein 1A | MAP1A | S2617 | 100.0 | 13.6 | K.AKPAS*PAR.R |
| Microtubule associated protein 1A | MAP1A | T1834 | 22.0 | 31.4 | K.NEPT*TPSWLADIPPWVPK.D |
| Microtubule associated protein 1A | MAP1A | T1674 | 8.9 | 51.8 | K.ELAPAWEDT*SPEQDNR.Y |
| Microtubule associated protein 1A | MAP1A | S612T616 | 36.2 | 53.1 | R.GLDS*GAET*EEEKDTWEEK.K |
| Microtubule associated protein 1A | MAP1A | T1653 | 7.6 | 29.7 | R.GQDVVQEWQET*SPTR.E |
| Microtubule associated protein 1A | MAP1A | S2629T2631 | 100.0 | 29.4 | R.S*PT*PGKGPADR.A |
| Microtubule associated protein 1A | MAP1A | S1797S1801 | 36.1 | 22.4 | R.SPFEIIS*PPAS*PPEMVGQR.V |
| Microtubule associated protein 1A | MAP1A | S2018S2019 | 9.4 | 14.9 | K.ELS*S*PISPK.S |
| Microtubule associated protein 1A | MAP1A | T2105 | 8.2 | 39.0 | R.SHWDDST*SDSELEK.G |
| Microtubule associated protein 1A | MAP1A | S1203 | 3.2 | 33.0 | K.DRWPEVSPEDTQSLSLSEES*PSK.E |
| Microtubule associated protein 1A | MAP1A | T1835 | 9.1 | 36.5 | K.NEPTT*PSWLADIPPWVPK.D |
| Microtubule associated protein 1A | MAP1A | S2104S2106 | 8.6 | 52.1 | R.SHWDDS*TS*DSELEK.G |
| Microtubule associated protein 1A | MAP1A | S2451 | -0.4 | 48.2 | R.GELSPS*FLNPPLPPSIDDR.D |
| Microtubule associated protein 1B | MAP1B | T1270 | 100.0 | 12.3 | K.T*PLGER.S |
| Microtubule associated protein 2 | MAP2 | S1539S1540 | 100.0 | 18.5 | R.RS*S*LPR.R |
| Microtubule associated protein 4 | MAP4 | S928 | 13.9 | 75.5 | R.LATNTS*APDLK.N |
| Microtubule associated protein 4 | MAP4 | S825 | 26.2 | 62.9 | R.S*PSTLLPK.K |

Peak Area: <-10, -5, -3, 0, 3, 5, >10

%CV: 0, 29, 42, 71, 86, >100

**White dots-** Significant change in peptide abundance at **5** %FDR compared to the timepoint with the minimum peak area for a given PSM

CarT (0m, 2m, 5m) RajiB (0m, 2m, 5m)

| Protein Name | Gene | Phosphosites | Ascor | MOWSE | Sequence |
|---|---|---|---|---|---|
| **Microtubule associated protein 4** | MAP4 | S358 | **33.9** | **13.1** | R.AS*PIKMDLAPSK.D |
| **Microtubule associated protein 4** | MAP4 | S1073 | **100.0** | **109.3** | K.VGS*LDNVGHLPAGGAVK.T |
| **Microtubule associated protein 4** | MAP4 | S280 | **22.4** | **88.4** | K.DMES*PTKLDVTLAK.D |
| **Microtubule associated protein 4** | MAP4 | T521 | **56.6** | **38.9** | K.DVT*PPPETEVVLIK.N |
| **Microtubule associated protein 4** | MAP4 | S507 | **68.1** | **73.0** | K.DMS*PLSETEMALGK.D |
| **Microtubule associated protein 4** | MAP4 | S507S510 | **46.4** | **83.2** | K.DMS*PLS*ETEMALGK.D |
| **Microtubule associated protein 4** | MAP4 | S941 | **25.6** | **56.5** | R.SKVGS*TENIK.H |
| **Microtubule associated protein 4** | MAP4 | T942 | **8.9** | **52.0** | K.VGST*ENIK.H |
| **Microtubule associated protein 4** | MAP4 | Y113 | **27.3** | **32.7** | K.MAYQEY*PNSQNWPEDTNFCFQPEQVVDPIQTDPFK.M |
| **Microtubule associated protein 4** | MAP4 | S68 | | **42.6** | K.KKPCS*ETSQIEDTPSSKPTLLANGGHGVEGSDTTGSPTEFLEEK.M |
| **Microtubule associated protein 4** | MAP4 | S116 | **8.8** | **24.1** | K.MAYQEYPNS*QNWPEDTNFCFQPEQVVDPIQTDPFK.M |
| **Microtubule associated protein 4** | MAP4 | T123 | **0.1** | **22.7** | K.MAYQEYPNSQNWPEDT*NFCFQPEQVVDPIQTDPFK.M |
| **Microtubule associated protein 4** | MAP4 | T925 | **4.0** | **10.7** | R.LAT*NTSAPDLK.N |
| **Microtubule associated protein 4** | MAP4 | S827 | **2.6** | **50.2** | R.SPS*TLLPK.K |
| **Microtubule associated protein 4** | MAP4 | T927 | | **29.2** | R.LATNT*SAPDLK.N |
| **Microtubule associated protein 4** | MAP4 | S68T70 | | **14.1** | K.KKPCS*ET*SQIEDTPSSKPTLLANGGHGVEGSDTTGSPTEFLEEK.M |
| **Microtubule associated protein 4** | MAP4 | S507S510T521 | **14.2** | **12.0** | K.DMS*PLS*ETEMALGKDVT*PPPETEVVLIK.N |
| **Microtubule associated protein tau** | MAPT | S713S717T720 | **15.8** | **39.8** | K.TDHGAEIVYKS*PVVS*GDT*SPR.H |
| **Microtubule associated protein tau** | MAPT | S729S730 | **11.8** | **63.0** | R.HLSNVS*S*TGSIDMVDSPQLATLADEVSASLAK.Q |
| **Microtubule associated protein tau** | MAPT | S726S729S730 | **0.4** | **22.6** | R.HLS*NVS*S*TGSIDMVDS*PQLATLADEVSASLAK.Q |
| **Microtubule associated protein tau** | MAPT | S726S729 | **13.9** | **45.0** | R.HLS*NVS*STGSIDMVDSPQLATLADEVSASLAK.Q |
| **Microtubule associated protein tau** | MAPT | S717T720S721 | **15.4** | **22.0** | K.TDHGAEIVYKSPVVS*GDT*S*PR.H |
| **Microtubule associated protein tau** | MAPT | S733S739 | **5.3** | **24.5** | R.HLSNVSSTGS*IDMVDS*PQLATLADEVSASLAK.Q |
| **Microtubule associated serine/threonine kinase 2** | MAST2 | S148 | **37.7** | **44.7** | R.NQSLGQS*APSLTAGLK.E |
| **Microtubule associated serine/threonine kinase 3** | MAST3 | S88S100 | **11.3** | **40.8** | R.RWSLAS*LPSSGYGTNTPS*STLSSSSSSR.E |
| **Microtubule associated serine/threonine kinase 3** | MAST3 | S85S106 | **7.4** | **39.3** | R.RWS*LASLPSSGYGTNTPSSTLSSS*SSSR.E |
| **Microtubule associated serine/threonine kinase 3** | MAST3 | S85S100 | **11.6** | **58.8** | R.RWS*LASLPSSGYGTNTPS*STLSSSSSSR.E |
| **Microtubule associated serine/threonine kinase 3** | MAST3 | S85T98 | **9.1** | **36.9** | R.RWS*LASLPSSGYGTNT*PSSTLSSSSSSR.E |
| **Microtubule associated serine/threonine kinase 3** | MAST3 | S709S710 | **16.7** | **72.2** | K.VYS*S*SEFLAVQPTPTFAER.S |
| **Microtubule associated serine/threonine kinase 3** | MAST3 | S85S105 | **14.2** | **51.8** | R.RWS*LASLPSSGYGTNTPSSTLSS*SSSSR.E |
| **Microtubule associated serine/threonine kinase 3** | MAST3 | Y708S709 | **61.2** | **20.5** | K.VY*S*SSEFLAVQPTPTFAER.S |
| **Microtubule associated serine/threonine kinase 3** | MAST3 | S85S108 | **5.2** | **11.9** | R.RWS*LASLPSSGYGTNTPSSTLSSSSS*SR.E |
| **Microtubule associated serine/threonine kinase 3** | MAST3 | S92S106 | **10.8** | **26.4** | R.RWSLASLPSS*GYGTNTPSSTLSSS*SSSR.E |
| **Microtubule associated serine/threonine kinase 3** | MAST3 | S774 | **14.0** | **20.6** | R.GPS*PSLLNTISLDTMPK.F |
| **Microtubule associated serine/threonine kinase 3** | MAST3 | S88Y94 | **7.6** | **12.8** | R.RWSLAS*LPSSGY*GTNTPSSTLSSSSSSR.E |
| **Microtubule associated serine/threonine kinase 3** | MAST3 | Y708S711 | **12.0** | **23.5** | K.VY*SSS*EFLAVQPTPTFAER.S |
| **Microtubule-associated protein 1A** | MAP1A | S25 | **100.0** | **63.3** | R.S*PGAPLAQNPAELLCEAGAAVAAAR.W |
| **Microtubule-associated protein 9** | MAP9 | S175 | **100.0** | **22.3** | M.PS*PRPR.G |
| **Midasin** | MDN1 | S4752S4754 | **100.0** | **98.2** | K.MHDGELEEQEEDDEKS*DS*EGGDLDK.H |
| **Midasin** | MDN1 | S4538 | **56.6** | **61.8** | K.AEENTDQAS*PQEDYAGFER.L |
| **Midasin** | MDN1 | T5017 | **-0.2** | **32.8** | R.EDSDT*EEQVPEALER.K |
| **Midasin** | MDN1 | S5015 | **15.2** | **71.8** | R.EDS*DTEEQVPEALER.K |
| **Minichromosome maintenance protein 10** | MCM10 | T85 | **60.8** | **63.1** | K.ENLATLFGDMEDLT*DEEEVPASQSTENR.V |
| **Minor histocompatibility antigen HA1** | ARHGAP4 | S23 | **32.1** | **58.5** | R.AGS*PSPQPSGELPR.K |
| **Minor histocompatibility antigen HA1** | ARHGAP4 | S578 | **18.6** | **66.7** | R.KSS*FNVSDVARPEAAGSPPEEGGCTEGTPAK.D |
| **Minor histocompatibility antigen HA1** | ARHGAP4 | S564 | **8.2** | **60.7** | R.DQEPDVHYDFEPHVS*ANAWSPVMR.A |
| **Minor histocompatibility antigen HA1** | ARHGAP4 | S569 | **19.8** | **57.9** | R.DQEPDVHYDFEPHVSANAWS*PVMR.A |
| **Minor histocompatibility antigen HA1** | ARHGAP4 | S14 | **7.8** | **14.1** | K.TPS*ISKK.N |
| **Minor histocompatibility antigen HA1** | ARHGAP4 | S619 | **36.8** | **56.0** | K.S*WPLSISDSDSGLDPGPGAGDFKK.F |
| **Misshapen-like kinase 1 (zebrafish)** | MINK1 | S701 | **18.1** | **78.8** | R.SNS*AWQIYLQR.R |
| **Misshapen-like kinase 1 (zebrafish)** | MINK1 | S674 | **8.7** | **28.8** | R.TSS*IATALNTSGAGGSRPAQAVR.A |
| **Misshapen-like kinase 1 (zebrafish)** | MINK1 | S763 | **46.1** | **41.4** | R.SDSVLPASHGHLPQAGS*LER.N |
| **Misshapen-like kinase 1 (zebrafish)** | MINK1 | T681 | **-5.9** | **12.6** | R.TSSIATALNT*SGAGGSRPAQAVR.A |
| **Misshapen-like kinase 1 (zebrafish)** | MINK1 | T860 | **-0.8** | **44.6** | R.SDGDT*DSVSTMVVHDVEEITGTQPPYGGGTMVVQR.T |
| **Mitochondrial fission factor** | MFF | Y19 | **13.9** | **114.6** | R.NDSLY*GISNIDTTIEGTSDDLTVVDAASLR.R |
| **Mitochondrial fission factor** | MFF | T26 | **8.0** | **73.4** | R.NDSLYGISNIDT*TIEGTSDDLTVVDAASLR.R |
| **Mitochondrial fission factor** | MFF | S17 | **12.1** | **100.1** | R.NDS*LYGISNIDTTIEGTSDDLTVVDAASLR.R |
| **Mitochondrial isoleucine tRNA synthetase** | IARS2 | S186 | **100.0** | **25.0** | K.ARS*FAK.A |
| **Mitochondrial ribosomal protein S16** | MRPS16 | T130 | **25.6** | **65.0** | K.TDAEAT*DTEATET.- |
| **Mitogen activated protein kinase 8 interacting protein 3** | MAPK8IP3 | S278 | **22.8** | **67.1** | K.SNTPTS*SVPSAAVTPLNESLQPLGDYGVGSK.N |
| **Mitogen activated protein kinase 8 interacting protein 3** | MAPK8IP3 | T277 | **12.2** | **58.1** | K.SNTPT*SSVPSAAVTPLNESLQPLGDYGVGSK.N |
| **Mitogen activated protein kinase 8 interacting protein 3** | MAPK8IP3 | S365 | **9.1** | **55.7** | R.TGSS*PTQGIVNK.A |

Peak Area: <-10, -5, -3, 0, 3, 5, >10

%CV: 0, 29, 42, 71, 86, >100

**White dots-** Significant change in peptide abundance at **5** %FDR compared to the timepoint with the minimum peak area for a given PSM

CarT (0m, 2m, 5m) RajiB (0m, 2m, 5m)

| Protein Name | Gene | Phosphosites | Ascor | MOWSE | Sequence |
|---|---|---|---|---|---|
| **Mitogen activated protein kinase 8 interacting protein 3** | MAPK8IP3 | S273 | 7.5 | 66.7 | K.S*NTPTSSVPSAAVTPLNESLQPLGDYGVGSK.N |
| **Mitogen activated protein kinase 8 interacting protein 3** | MAPK8IP3 | S279 | 10.7 | 46.0 | K.SNTPTSS*VPSAAVTPLNESLQPLGDYGVGSK.N |
| **Mitogen-activated protein kinase associated protein 1** | MAPKAP1 | S510 | 12.3 | 14.1 | R.RTS*FSFQK.E |
| **Mitogen-activated protein kinase associated protein 1** | MAPKAP1 | S512 | 3.5 | 20.4 | R.RTSFS*FQK.E |
| **Mitogen-activated protein kinase associated protein 1** | MAPKAP1 | T509 | 8.9 | 13.4 | R.RT*SFSFQK.E |
| **Mitogen-activated protein kinase kinase kinase 7 interacting protein 1** | TAB1 | S7 | 34.5 | 39.0 | R.S*LLQSEQQPSWTDDLPLCHLSGVGSASNR.S |
| **MLF2** | MLF2 | S238 | 6.8 | 14.0 | R.LAIQGPEDS*PSR.Q |
| **MLL** | KMT2A | S1114S1115S11 | 6.5 | 43.2 | K.ILSSMGNDDKS*S*IAGS*EDAEPLAPPIKPIKPVTR.N |
| **MLL** | KMT2A | S2869 | 6.5 | 83.0 | K.NTPSMQALGESPES*SSSELLNLGEGLGLDSNR.E |
| **MLL** | KMT2A | S1114S1119 | 9.4 | 33.7 | K.ILSSMGNDDKS*SIAGS*EDAEPLAPPIKPIKPVTR.N |
| **MLL** | KMT2A | S2866 | 6.4 | 49.6 | K.NTPSMQALGES*PESSSSELLNLGEGLGLDSNR.E |
| **MLL** | KMT2A | S2872 | 8.0 | 53.4 | K.NTPSMQALGESPESSSS*ELLNLGEGLGLDSNR.E |
| **MLL** | KMT2A | S1107S1119 | 1.1 | 23.7 | K.ILSS*MGNDDKSSIAGS*EDAEPLAPPIKPIKPVTR.N |
| **MLL** | KMT2A | S2870 | 5.3 | 72.3 | K.NTPSMQALGESPESS*SSELLNLGEGLGLDSNR.E |
| **MLL** | KMT2A | S3036 | 25.4 | 30.2 | R.NSSTPGLQVPVS*PTVPIQNQK.Y |
| **MLL** | KMT2A | S2866S2871 | 10.4 | 34.2 | K.NTPSMQALGES*PESSS*SELLNLGEGLGLDSNR.E |
| **MLL** | KMT2A | S1115S1119 | 8.4 | 27.8 | K.ILSSMGNDDKSS*IAGS*EDAEPLAPPIKPIKPVTR.N |
| **MLL** | KMT2A | S1107S1114S11 | 6.7 | 25.5 | K.ILSS*MGNDDKS*S*IAGSEDAEPLAPPIKPIKPVTR.N |
| **MLL** | KMT2A | S153 | 43.3 | 53.2 | R.AVFGESGGGGGSGEDEQFLGFGS*DEEVR.V |
| **MLL** | KMT2A | S1106S1107S11 | | 27.3 | K.ILS*S*MGNDDKS*SIAGSEDAEPLAPPIKPIKPVTR.N |
| **MLL** | KMT2A | S1114S1115 | 1.9 | 20.0 | K.ILSSMGNDDKS*S*IAGSEDAEPLAPPIKPIKPVTR.N |
| **MLL** | KMT2A | S1107S1114S11 | 9.2 | 15.2 | K.ILSS*MGNDDKS*SIAGS*EDAEPLAPPIKPIKPVTR.N |
| **MLL2** | KMT2D | S4738 | 100.0 | 42.7 | R.ALS*PVIPLIPR.A |
| **MLL3** | KMT2C | S2937S2946 | 15.0 | 30.3 | K.SDNSDIRPSGS*PPPPTLPAS*PSNHVSSLPPFIAPPGR.V |
| **MLL3** | KMT2C | S2927S2930 | | 28.6 | K.S*DNS*DIRPSGSPPPPTLPASPSNHVSSLPPFIAPPGR.V |
| **MLLT2** | AFF1 | S203S206S212 | 20.0 | 28.7 | R.ELSPLIS*LPS*PVPPLS*PIHSNQQTLPR.T |
| **MLLT2** | AFF1 | S199S206S212 | 21.3 | 12.4 | R.ELS*PLISLPS*PVPPLS*PIHSNQQTLPR.T |
| **MLLT2** | AFF1 | S206S212 | 27.6 | 16.0 | R.ELSPLISLPS*PVPPLS*PIHSNQQTLPR.T |
| **MMRP19** | APIP | S87 | 13.9 | 34.7 | K.DISGPS*PSKK.L |
| **MMRP19** | APIP | S89 | 13.9 | 20.4 | K.DISGPSPS*KK.L |
| **MPP10** | MPHOSPH | S120S139 | 100.0 | 128.0 | R.EEDGS*EIEADDKEDLEDLEEEEVS*DMGNDDPEMGER.A |
| **MPP10** | MPHOSPH | S163S167S171 | 78.7 | 94.3 | K.S*PVFS*DEDS*DLDFDISK.L |
| **MPP10** | MPHOSPH | S242 | 92.1 | 122.5 | R.KDDNDEEEEDIDFFEDIDS*DEDEGGLFGSK.K |
| **MPP10** | MPHOSPH | S163S171 | 24.0 | 45.9 | K.S*PVFSDEDS*DLDFDISK.L |
| **MPP10** | MPHOSPH | S167S171 | 39.5 | 55.5 | K.SPVFS*DEDS*DLDFDISK.L |
| **MSH6** | MSH6 | S219S227 | 100.0 | 95.1 | K.S*EEDNEIES*EEEVQPK.T |
| **MSH6** | MSH6 | S252S254S256 | 41.0 | 50.2 | R.VIS*DS*ES*DIGGS*DVEFKPDTK.E |
| **MSH6** | MSH6 | S137 | 17.8 | 47.9 | R.VHVQFFDDS*PTR.G |
| **mSin3A-associated protein 130** | SAP130 | T853S855 | 12.8 | 56.7 | R.KQQHVISTEEGDMMET*NS*TDDEKSTAK.S |
| **mSin3A-associated protein 130** | SAP130 | S855T856 | 19.7 | 52.6 | R.KQQHVISTEEGDMMETNS*T*DDEK.S |
| **MST4** | STK26 | S300S304S306 | 45.4 | 48.6 | R.WKAEGHS*DDES*DS*EGS*DSESTSR.E |
| **MST4** | STK26 | T178 | 32.5 | 67.4 | R.NT*FVGTPFWMAPEVIQQSAYDSK.A |
| **MST4** | STK26 | T182 | -0.2 | 17.8 | R.NTFVGT*PFWMAPEVIQQSAYDSK.A |
| **mTOR** | MTOR | S1261 | 12.1 | 24.3 | K.LHVS*TINLQK.A |
| **Mucoepidermoid carcinoma translocated 1** | CRTC1 | Y60 | 1.9 | 54.9 | R.GQY*YGGSLPNVNQIGSGTMDLPFQTPFQSSGLDTSR.T |
| **Mucoepidermoid carcinoma translocated 1** | CRTC1 | S73 | -0.2 | 44.3 | R.GQYYGGSLPNVNQIGS*GTMDLPFQTPFQSSGLDTSR.T |
| **Mucoepidermoid carcinoma translocated 1** | CRTC1 | T75 | -0.1 | 51.8 | R.GQYYGGSLPNVNQIGSGT*MDLPFQTPFQSSGLDTSR.T |
| **Mucoepidermoid carcinoma translocated 1** | CRTC1 | S64 | 1.7 | 55.2 | R.GQYYGGS*LPNVNQIGSGTMDLPFQTPFQSSGLDTSR.T |
| **Mucoepidermoid carcinoma translocated 1** | CRTC1 | Y61 | 8.2 | 38.0 | R.GQYY*GGSLPNVNQIGSGTMDLPFQTPFQSSGLDTSR.T |
| **Multimerin 2** | MMRN2 | T561S562 | 100.0 | 12.9 | R.AAT*S*RLR.S |
| **Muscle-derived protein 77** | TXLNB | S552 | 6.2 | 53.7 | R.DS*ESPLPPLTPQAEAEGGSDAEPPSK.A |
| **Muscle-derived protein 77** | TXLNB | S554 | 3.1 | 39.7 | R.DSES*PLPPLTPQAEAEGGSDAEPPSK.A |
| **MYC binding protein 2** | MYCBP2 | S2749 | 20.2 | 66.3 | R.S*LSPNHNTLQTLK.S |
| **MYC binding protein 2** | MYCBP2 | S2749S2751 | 44.0 | 46.8 | R.S*LS*PNHNTLQTLK.S |
| **MYC binding protein 2** | MYCBP2 | S2833 | 39.9 | 65.7 | R.SKS*DSYTLDPDTLR.K |
| **MYC binding protein 2** | MYCBP2 | S3467 | 67.4 | 80.3 | R.VNS*GDTEVGSSLLR.H |
| **MYC binding protein 2** | MYCBP2 | T2645 | 71.7 | 95.6 | R.HEDEQALLDQNSQT*PPPSPFSVQAFNK.G |
| **MYC binding protein 2** | MYCBP2 | T2645S2649 | 22.2 | 93.8 | R.HEDEQALLDQNSQT*PPPS*PFSVQAFNK.G |
| **MYC binding protein 2** | MYCBP2 | S2882 | 68.5 | 33.8 | R.APS*PHVVQENLHSEVVEVCTSSTLK.T |
| **MYC binding protein 2** | MYCBP2 | S2831 | 10.5 | 40.9 | R.S*KSDSYTLDPDTLR.K |
| **MYC binding protein 2** | MYCBP2 | S2774 | 29.9 | 21.0 | R.AES*PGPGSR.L |
| **MYC binding protein 2** | MYCBP2 | S2751 | 9.2 | 64.2 | R.SLS*PNHNTLQTLK.S |

Peak Area: <-10, -5, -3, 0, 3, 5, >10

%CV: 0, 29, 42, 71, 86, >100

**White dots-** Significant change in peptide abundance at **5**%FDR compared to the timepoint with the minimum peak area for a given PSM

CarT: 0m, 2m, 5m — RajiB: 0m, 2m, 5m

| Protein Name | Gene | Phosphosites | Ascor | MOWSE | Sequence |
|---|---|---|---|---|---|
| **Myc Promoter binding protein** | DENND4A | S1589 | **10.9** | **84.2** | R.SIS*TSGPLDKEDTGR.Q |
| **Myc Promoter binding protein** | DENND4A | T1512 | **20.8** | **38.5** | K.SSPST*ENMHFPSSISSQTR.Q |
| **Myc Promoter binding protein** | DENND4A | S1128 | **22.4** | **59.9** | K.VDSPVIFDLEDLDS*ETDVSK.A |
| **Myc Promoter binding protein** | DENND4A | S1511 | **22.8** | **52.2** | K.SSPS*TENMHFPSSISSQTR.Q |
| **Myc Promoter binding protein** | DENND4A | T1220S1221 | **13.3** | **27.8** | R.RDLAEEIVMYMNNMSSPLT*S*R.T |
| | DENND4A | S1216T1220 | **12.3** | **15.4** | R.DLAEEIVMYMNNMS*SPLT*SR.T |
| **Myc Promoter binding protein** | DENND4A | S1216 | **21.7** | **66.2** | R.RDLAEEIVMYMNNMS*SPLTSR.T |
| **Myc Promoter binding protein** | DENND4A | S1216S1221 | **13.6** | **32.2** | R.RDLAEEIVMYMNNMS*SPLTS*R.T |
| **Myc Promoter binding protein** | DENND4A | S1217S1221 | **8.5** | **17.3** | R.DLAEEIVMYMNNMSS*PLTS*R.T |
| **Myc Promoter binding protein** | DENND4A | S1099 | **4.3** | **52.2** | K.ESS*DDDTPFDGSNYLADKVDSPVIFDLEDLDSETDVSK.A |
| **Myc Promoter binding protein** | DENND4A | S1015 | **22.0** | **63.6** | R.HKS*DNETNLQQQVVWGNR.N |
| **Myc Promoter binding protein** | DENND4A | S1217 | **12.3** | **56.5** | R.DLAEEIVMYMNNMSS*PLTSR.T |
| **Myc Promoter binding protein** | DENND4A | S1093S1098S1 | | **28.2** | R.DS*LEKES*S*DDDTPFDGSNYLADKVDSPVIFDLEDLDSETDVSK.^ |
| **Myc Promoter binding protein** | DENND4A | S1108Y1110S11 | **-1.4** | **19.2** | R.DSLEKESSDDDTPFDGS*NY*LADKVDS*PVIFDLEDLDSETDVSK.^ |
| **Myc Promoter binding protein** | DENND4A | S1509 | **-0.3** | **31.7** | K.SS*PSTENMHFPSSISSQTR.Q |
| **Myc Promoter binding protein** | DENND4A | T1019 | **1.2** | **20.6** | R.HKSDNET*NLQQQVVWGNR.N |
| **Myc proto-oncogene protein** | MYC | S6 | **26.0** | **40.3** | M.PLNVS*FTNR.N |
| **Myelin basic protein** | MBP | S40 | **9.6** | **82.5** | R.TTS*EDNEVFGEADANQNNGTSSQDTAVTDSKR.T |
| **Myelin basic protein** | MBP | S112 | **8.3** | **107.1** | K.DRPS*ESDELQTIQEDSAATSESLDVMASQK.R |
| **Myelin basic protein** | MBP | S36 | **100.0** | **17.2** | R.NLGELS*R.T |
| **Myelin expression factor 2** | MYEF2 | S17 | **15.5** | **29.8** | K.AEVPGATGGDS*PHLQPAEPPGEPR.R |
| **Myelin expression factor 2** | MYEF2 | T13 | **9.2** | **11.3** | K.AEVPGAT*GGDSPHLQPAEPPGEPR.R |
| **Myeloid/lymphoid or mixed-lineage leukemia translocated to. 1** | MLLT1 | S359S361S366 | **100.0** | **30.3** | K.ALEVEES*NS*EDEAS*FK.S |
| **Myeloid/lymphoid or mixed-lineage leukemia translocated to. 1** | MLLT1 | S475 | **46.9** | **34.4** | R.S*PESCSKPEK.I |
| **Myocardin related protein** | MKL1 | S446S454 | **9.0** | **13.4** | K.FGS*TGSTPPVS*PTPSER.S |
| **Myocardin related protein** | MKL1 | T447S449 | **8.5** | **20.4** | K.FGST*GS*TPPVSPTPSER.S |
| **Myocardin related protein** | MKL1 | S678S682 | | **14.5** | K. [illegible] |
| **Myocardin related transcription factor B** | MKL2 | S222 | **6.7** | **18.0** | K.VSESPS*PVTTNTPAQFASVSPTVPEFLK.T |
| **Myocardin related transcription factor B** | MKL2 | S222S236 | **5.1** | **14.1** | K.VSESPS*PVTTNTPAQFASVS*PTVPEFLK.T |
| **Myocardin related transcription factor B** | MKL2 | T228S234 | **7.0** | **22.7** | K.VSESPSPVTTNT*PAQFAS*VSPTVPEFLK.T |
| **Myocyte specific enhancer factor 2A** | MEF2C | S494 | **16.1** | **20.0** | R.ES*PSVKR.M |
| **Myocyte specific enhancer factor 2C** | MEF2C | S240 | **100.0** | **35.7** | K.S*PPPMNLGMNNR.K |
| **Myocyte specific enhancer factor 2C** | MEF2C | S222 | **90.1** | **72.1** | R.NS*PGLLVSPGNLNK.N |
| **Myosin 5A** | MYO5A | S1115S1119 | **7.1** | **71.8** | R.TDS*THSS*NESEYIFSSEIAEMEDIPSR.T |
| **Myosin 5A** | MYO5A | S1115T1116 | **8.2** | **95.2** | R.TDS*T*HSSNESEYIFSSEIAEMEDIPSR.T |
| **Myosin 5A** | MYO5A | S1652 | **11.9** | **85.6** | R.TSS*IADEGTYTLDSILR.Q |
| **Myosin 5A** | MYO5A | S1115S1118 | **5.7** | **74.3** | K.RTDS*THS*SNESEYIFSSEIAEMEDIPSR.T |
| **Myosin 5A** | MYO5A | T1116S1119 | **11.0** | **39.1** | R.TDST*HSS*NESEYIFSSEIAEMEDIPSR.T |
| **Myosin heavy chain 9, nonmuscle** | MYH9 | S1943 | **100.0** | **72.7** | R.KGAGDGS*DEEVDGKADGAEAKPAE.- |
| **Myosin heavy chain, nonmuscle type B** | MYH10 | T1960 | **20.9** | **25.2** | R.RQLHLEGASLELSDDDT*ESK.T |
| **Myosin heavy chain, nonmuscle type B** | MYH10 | S1952S1956 | **33.6** | **72.8** | R.RQLHLEGAS*LELS*DDDTESK.T |
| **Myosin heavy chain, nonmuscle type B** | MYH10 | S1956 | **29.7** | **97.6** | R.QLHLEGASLELS*DDDTESK.T |
| **Myosin IF** | MYO1F | S939 | **8.9** | **12.9** | R.RSS*QAPTR.A |
| **Myosin IG** | MYO1G | S583 | **85.6** | **36.0** | K.NS*MVALVENLASK.E |
| **Myosin IG** | MYO1G | S1010 | **32.1** | **62.6** | R.GS*FTLLWPSR.- |
| **Myosin phosphatase target subunit 1** | PPP1R12A | S507 | **12.1** | **52.0** | R.LAS*TSDIEEK.E |
| **Myosin phosphatase target subunit 1** | PPP1R12A | S862 | **0.1** | **58.7** | R.STGVSFWTQDS*DENEQEQQSDTEEGSNKK.E |
| **Myosin phosphatase target subunit 1** | PPP1R12A | S871 | **19.2** | **71.9** | R.STGVSFWTQDSDENEQEQQS*DTEEGSNKK.E |
| **Myosin phosphatase target subunit 1** | PPP1R12A | T859 | **5.9** | **15.1** | R.STGVSFWT*QDSDENEQEQQSDTEEGSNKK.E |
| **Myosin phosphatase target subunit 1** | PPP1R12A | S445 | **23.9** | **93.7** | K.TGS*YGALAEITASK.E |
| **Myosin phosphatase target subunit 1** | PPP1R12A | Y446 | **6.8** | **50.5** | R.KTGSY*GALAEITASK.E |
| **Myosin phosphatase target subunit 1** | PPP1R12A | S862S871 | **4.0** | **42.6** | R.STGVSFWTQDS*DENEQEQQS*DTEEGSNKK.E |
| **Myosin phosphatase target subunit 1** | PPP1R12A | T443 | **6.0** | **23.6** | R.KT*GSYGALAEITASK.E |
| | MYL12A | S20 | **16.9** | **127.5** | R.ATS*NVFAM#FDQSQIQEFK.E |
| **Myosin regulatory light chain MRLC2** | MYL12A | T19S20 | **58.1** | **86.9** | R.AT*S*NVFAMFDQSQIQEFK.E |
| **Myosin regulatory light chain MRLC2** | MYL12A | T19 | **15.7** | **100.4** | R.AT*SNVFAMFDQSQIQEFK.E |
| **Myosin XVIIIA** | MYO18A | S2041S2043 | **32.2** | **56.2** | R.YSHSYLS*DS*DTEAK.L |
| **Myosin XVIIIA** | MYO18A | S1067 | **24.3** | **50.4** | R.RVS*SSSELDLPSGDHCEAGLLQLDVPLLR.T |
| **Myosin XVIIIA** | MYO18A | S1942S1951S1 | **100.0** | **30.0** | R.IGDLQAAIEDEMES*DENEDLINS*EGDS*DVDS*ELEDRVDGVK.S |
| **Myosin XVIIIA** | MYO18A | Y2039S2041 | **10.0** | **21.7** | R.YSHSY*LS*DSDTEAK.L |
| **Myosin XVIIIA** | MYO18A | Y2039S2043 | **5.8** | **49.3** | R.YSHSY*LSDS*DTEAK.L |
| **Myosin XVIIIA** | MYO18A | S1942 | **79.0** | **105.1** | R.IGDLQAAIEDEMES*DENEDLINSLQDMVTK.Y |

Peak Area: <-10, -5, -3, 0, 3, 5, >10

%CV: 0, 29, 42, 71, 86, >100

**White dots-** Significant change in peptide abundance at 5 %FDR compared to the timepoint with the minimum peak area for a given PSM

| Protein Name | Gene | Phosphosites | CarT 0m | CarT 2m | CarT 5m | RajiB 0m | RajiB 2m | RajiB 5m | Ascor | MOWSE | Sequence |
|---|---|---|---|---|---|---|---|---|---|---|---|
| Myosin XVIIIA | MYO18A | S1069 | | | | | | | 35.1 | 43.0 | R.RVSSS*SELDLPSGDHCEAGLLQLDVPLLR.T |
| Myosin XVIIIA | MYO18A | S1068 | | | | | | | 6.5 | 18.5 | R.RVSS*SSELDLPSGDHCEAGLLQLDVPLLR.T |
| Myosin XVIIIA | MYO18A | S2007 | | | | | | | 27.6 | 44.3 | K.SSS*PTSYWK.S |
| Myosin XVIIIA | MYO18A | S140 | | | | | | | 24.1 | 15.2 | K.RFS*FSQR.S |
| Myosin XVIIIA | MYO18A | Y2039S2041S2 | | | | | | | -0.2 | 31.0 | R.YSHSY*LS*DS*DTEAK.L |
| Myosin XVIIIA | MYO18A | S2041S2043T2 | | | | | | | 9.0 | 21.7 | R.YSHSYLS*DS*DT*EAK.L |
| Myosin XVIIIA | MYO18A | S2038S2043 | | | | | | | 8.3 | 14.1 | R.YSHS*YLSDS*DTEAK.L |
| Myosin XVIIIB | MYO18B | T2549 | | | | | | | 100.0 | 19.6 | R.REPGT*GR.K |
| Myotubularin | MTM1 | S588S591 | | | | | | | 2.7 | 11.8 | K.LSDPPTS*PSS*PSQMMPHVQTHF.- |
| Myotubularin | MTM1 | S590S591 | | | | | | | 6.5 | 23.8 | K.LSDPPTSPS*S*PSQMMPHVQTHF.- |
| Myotubularin | MTM1 | T587S591 | | | | | | | 8.0 | 19.9 | K.LSDPPT*SPSS*PSQMMPHVQTHF.- |
| Myotubularin | MTM1 | S591S593 | | | | | | | 4.6 | 30.6 | K.LSDPPTSPSS*PS*QMMPHVQTHF.- |
| Myotubularin related protein 1 | MTMR1 | S652 | | | | | | | 13.8 | 14.8 | R.GS*SPSHSATSVHTSV.- |
| Myotubularin related protein 3 | MTMR3 | S647 | | | | | | | 43.2 | 107.0 | R.S*LELSSLAGPGEDPLSADSLGKPTR.V |
| MYST histone acetyltransferase 2 | KAT7 | T118S124 | | | | | | | | 30.9 | K.NT*ADHDES*PPRTPTGNAPSSESDIDISSPNVSHDESIAK.D |
| MYST histone acetyltransferase 2 | KAT7 | S99S100S102 | | | | | | | 39.7 | 93.1 | R.S*S*GS*ETEQVVDFSDR.E |
| MYST histone acetyltransferase 2 | KAT7 | S124T128 | | | | | | | 13.6 | 25.2 | K.NTADHDES*PPRT*PTGNAPSSESDIDISSPNVSHDESIAK.D |
| MYST histone acetyltransferase 2 | KAT7 | S56 | | | | | | | 9.8 | 51.7 | R.LSQSSQDS*SPVR.N |
| MYST histone acetyltransferase 2 | KAT7 | S100S102T104 | | | | | | | 9.8 | 14.3 | R.SS*GS*ET*EQVVDFSDR.E |
| MYST histone acetyltransferase 2 | KAT7 | S57 | | | | | | | 7.6 | 58.2 | R.LSQSSQDSS*PVR.N |
| N-acetylglucosamine kinase | NAGK | S76 | | | | | | | 10.9 | 82.5 | R.SLGLSLS*GGDQEDAGR.I |
| N-myristoyl transferase 1 | NMT1 | S47 | | | | | | | 59.6 | 82.9 | R.GGLS*PANDTGAK.K |
| NACHT and WD repeat domain-containing protein 2 | NWD2 | T695S698 | | | | | | | 100.0 | 11.7 | K.ENT*RPS*NPLR.V |
| NACSIN | EHBP1 | S335 | | | | | | | 21.0 | 14.6 | K.EVQTPQYLNPFDEPEAFVTIKDS*PPQSTK.R |
| NACSIN | EHBP1 | T165S171 | | | | | | | 20.2 | 20.9 | K.AT*DEDMQS*LASLMSMK.Q |
| NACSIN | EHBP1 | S171S174 | | | | | | | 8.5 | 16.4 | K.ATDEDMQS*LAS*LMSMK.Q |
| NACSIN | EHBP1 | T165S174 | | | | | | | 8.4 | 41.7 | K.AT*DEDMQSLAS*LMSMK.Q |
| NAD synthetase 1 | NADSYN1 | S280Y281 | | | | | | | 18.1 | 12.3 | R.S*Y*RAEISSR.N |
| NADH ubiquinone oxidoreductase flavoprotein 3 | NDUFV3 | S159S160S164 | | | | | | | 6.3 | 111.8 | R.KVTSPSSSSSSSSS*S*DSES*DDEADVSEVTPR.V |
| NADH ubiquinone oxidoreductase flavoprotein 3 | NDUFV3 | S162 | | | | | | | 6.0 | 58.1 | R.KVTSPSSSSSSSSSSDS*ESDDEADVSEVTPR.V |
| NADH ubiquinone oxidoreductase flavoprotein 3 | NDUFV3 | S159S162S164 | | | | | | | 10.6 | 116.7 | R.KVTSPSSSSSSSSS*SDS*ES*DDEADVSEVTPR.V |
| NADH ubiquinone oxidoreductase flavoprotein 3 | NDUFV3 | S158S159S160 | | | | | | | 7.6 | 74.7 | K.VTSPSSSSSSSS*S*S*DSESDDEADVSEVTPR.V |
| NADH ubiquinone oxidoreductase flavoprotein 3 | NDUFV3 | S160S162 | | | | | | | 1.3 | 66.0 | R.KVTSPSSSSSSSSSS*DS*ESDDEADVSEVTPR.V |
| NADH ubiquinone oxidoreductase flavoprotein 3 | NDUFV3 | S160S162S164 | | | | | | | 11.2 | 102.1 | R.KVTSPSSSSSSSSSS*DS*ES*DDEADVSEVTPR.V |
| NADH ubiquinone oxidoreductase flavoprotein 3 | NDUFV3 | S159S160 | | | | | | | 16.2 | 69.4 | R.KVTSPSSSSSSSSS*S*DSESDDEADVSEVTPR.V |
| NADH ubiquinone oxidoreductase flavoprotein 3 | NDUFV3 | S159S160S162 | | | | | | | | 62.0 | R.KVTSPSSSSSSSSS*S*DS*ESDDEADVSEVTPR.V |
| NADH ubiquinone oxidoreductase flavoprotein 3 | NDUFV3 | S158S159S162 | | | | | | | 5.4 | 62.5 | K.VTSPSSSSSSSS*S*SDS*ESDDEADVSEVTPR.V |
| NAG | NBAS | S473 | | | | | | | 13.9 | 91.2 | R.AGEEDEGEEDS*DSDYEISAK.A |
| NAG | NBAS | S473S475 | | | | | | | 28.8 | 29.7 | R.AGEEDEGEEDS*DS*DYEISAK.A |
| NaGLT1 | MFSD4B | S474 | | | | | | | 10.2 | 22.7 | R.SS*LTEPTAEVYNQYPSNALVFESSPFNTGSAHVK.H |
| NaGLT1 | MFSD4B | T476 | | | | | | | -0.8 | 26.2 | R.SSLT*EPTAEVYNQYPSNALVFESSPFNTGSAHVK.H |
| Nardilysin | NRDC | S94 | | | | | | | 17.8 | 40.8 | R.RGS*LSNAGDPEIVK.S |
| Nascent polypeptide associated complex, alpha polypeptide | NACA | S166 | | | | | | | 11.5 | 129.6 | K.VQGEAVSNIQENTQTPTVQEES*EEEEVDETGVEVK.D |
| Nascent polypeptide associated complex, alpha polypeptide | NACA | T161 | | | | | | | 44.9 | 83.9 | K.VQGEAVSNIQENTQTPT*VQEESEEEEVDETGVEVK.D |
| Nascent polypeptide associated complex, alpha polypeptide | NACA | T159 | | | | | | | 5.1 | 44.0 | K.VQGEAVSNIQENTQT*PTVQEESEEEEVDETGVEVK.D |
| Nascent polypeptide associated complex, alpha polypeptide | NACA | S151 | | | | | | | | 76.4 | K.VQGEAVS*NIQENTQTPTVQEESEEEEVDETGVEVK.D |
| Nascent polypeptide associated complex, alpha polypeptide | NACA | T157S166 | | | | | | | 13.3 | 74.5 | K.VQGEAVSNIQENT*QTPTVQEES*EEEEVDETGVEVK.D |
| NATH | NAA15 | S855 | | | | | | | 22.0 | 11.3 | K.ITVNGDS*SAEAEELANEI.- |
| NATH | NAA15 | S855S856 | | | | | | | 45.1 | 26.3 | K.ITVNGDS*S*AEAEELANEI.- |
| | TRA2B | S225Y226 | | | | | | | 17.5 | 17.8 | R.SRS*Y*SPR.R |
| Natural killer tumor recognition sequence | RBM26 | S699S701 | | | | | | | 17.4 | 23.1 | R.S*RS*YSR.D |
| Natural killer tumor recognition sequence | NKTR | S613 | | | | | | | 9.1 | 44.0 | R.ATMAQNENVVVQPVVAENIPVIPLSDS*PPPSR.W |
| Natural killer tumor recognition sequence | RBM26 | S701Y702 | | | | | | | 9.6 | 13.5 | R.SRS*Y*SR.D |
| Natural killer tumor recognition sequence | NKTR | S611 | | | | | | | 15.6 | 21.8 | R.ATMAQNENVVVQPVVAENIPVIPLS*DSPPPSR.W |
| NCK1 | NCK1 | Y105 | | | | | | | 190.9 | 66.4 | R.LY*DLNMPAYVK.F |
| NDRG family member 2 | NDRG2 | S332 | | | | | | | 12.3 | 65.0 | R.TAS*LTSAASVDGNR.S |
| NDRG family member 3 | NDRG3 | S331 | | | | | | | 19.9 | 73.1 | R.THS*TSSSLGSGESPFSR.S |
| NDRG family member 3 | NDRG3 | T332 | | | | | | | -0.3 | 26.6 | R.THST*SSSLGSGESPFSR.S |
| NDRG family member 3 | NDRG3 | S335 | | | | | | | 18.0 | 44.0 | R.THSTSSS*LGSGESPFSR.S |
| NDRG family member 3 | NDRG3 | T329S333 | | | | | | | 2.6 | 66.1 | R.SRT*HSTS*SSLGSGESPFSR.S |
| NDRG family member 3 | NDRG3 | T329T332 | | | | | | | 16.7 | 86.8 | R.SRT*HST*SSSLGSGESPFSR.S |
| | NDRG3 | S331T332 | | | | | | | 3.3 | 54.6 | R.SRTHS*T*SSSLGSGESPFSR.S |

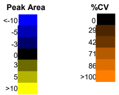


**White dots-** Significant change in peptide abundance at 5 %FDR compared to the timepoint with the minimum peak area for a given PSM

| Protein Name | Gene | Phosphosites | Ascor | MOWSE | Sequence |
|---|---|---|---|---|---|
| NDRG family member 3 | NDRG3 | T329S331 | 16.9 | 66.8 | R.SRT*HS*TSSSLGSGESPFSR.S |
| | NDRG3 | S331S334 | 19.5 | 60.5 | R.THS*TSS*SLGSGESPFSR.S |
| NDRG family member 3 | NDRG3 | T332S333 | 1.8 | 40.4 | R.THST*S*SSLGSGESPFSR.S |
| NECAP endocytosis associated 2 | NECAP2 | T182 | 25.4 | 39.5 | R.VRPAST*GGLSLLPPPPGGK.T |
| Nectin 1 | NECTIN1 | S422 | 31.5 | 23.1 | K.AGIPQHHPPMAQNLQYPDDS*DDEKK.A |
| Nectin 1 | NECTIN1 | S508S511 | 5.2 | 48.7 | R.TLGYQYDPEQLDLAENMVSQNDGS*FIS*KK.E |
| | NECTIN1 | S508 | 14.6 | 40.2 | R.TLGYQYDPEQLDLAENMVSQNDGS*FISKK.E |
| Nectin 1 | NECTIN1 | S503S511 | 10.4 | 57.3 | R.TLGYQYDPEQLDLAENMVS*QNDGSFIS*KK.E |
| | NECTIN1 | S434S435 | 17.2 | 74.8 | K.AGPLGGS*S*YEEEEEEEEGGGGGERK.V |
| Nectin 1 | NECTIN1 | S435 | 8.2 | 16.1 | K.AGPLGGSS*YEEEEEEEEGGGGGER.K |
| Nectin 1 | NECTIN1 | S434Y436 | 15.6 | 35.6 | K.AGPLGGS*SY*EEEEEEEEGGGGGER.K |
| Nectin 1 | NECTIN1 | S435Y436 | 10.4 | 80.2 | K.AGPLGGSS*Y*EEEEEEEEGGGGGER.K |
| Nectin 1 | NECTIN1 | S503S508 | 18.6 | 47.6 | R.TLGYQYDPEQLDLAENMVS*QNDGS*FISK.K |
| NEDD4 | NEDD4 | S675 | 24.1 | 44.5 | R.RGS*LQAYTFEEQPTLPVLLPTSSGLPPGWEEK.Q |
| NEDD4 | NEDD4 | Y679 | | 45.5 | R.RGSLQAY*TFEEQPTLPVLLPTSSGLPPGWEEK.Q |
| Negative elongation factor protein B | NELFB | S557 | 46.6 | 46.9 | R.KPS*PAQAAETPALELPLPSVPAPAPL.- |
| NELFE | NELFE | S115 | 26.9 | 75.8 | R.SIS*ADDDLQESSR.R |
| NELFE | NELFE | S113 | 10.5 | 62.4 | R.S*ISADDDLQESSR.R |
| NELFE | NELFE | S353 | 44.2 | 60.8 | K.SVWGSLAVQNS*PK.G |
| Nesprin 2 | SYNE2 | S4108 | 85.4 | 28.6 | R.RGS*MSYLAAVEEEVEESSVK.S |
| Nesprin 2 | SYNE2 | T6422S6428 | 9.6 | 18.2 | R. SCGETPVSVDSIPLEWDHT*GDVGGS*SSHEEDEEGPYYSALSGK.S |
| Nesprin 2 | SYNE2 | S4110 | 5.8 | 48.2 | R.RGSMS*YLAAVEEEVEESSVK.S |
| Nesprin 2 | SYNE2 | Y4111 | -0.2 | 31.6 | R.RGSMSY*LAAVEEEVEESSVK.S |
| Neurobeachin-like 2 | NBEAL2 | S770 | 4.3 | 94.3 | R.SQSVPASTGLGWGS*GLVAPLQEGSIDSTLAGTQDTR.W |
| Neurobeachin-like 2 | NBEAL2 | S763 | 12.0 | 118.6 | R.SQSVPAS*TGLGWGSGLVAPLQEGSIDSTLAGTQDTR.W |
| Neurobeachin-like 2 | NBEAL2 | T764 | 8.2 | 124.8 | R.SQSVPAST*GLGWGSGLVAPLQEGSIDSTLAGTQDTR.W |
| Neurobeachin-like 2 | NBEAL2 | S1364S1367 | 1.5 | 85.8 | R.SS*VGS*GNTAGGGGSSGTLTPASQPGTPSPLDGPRPFPAAPGR.[illegible] |
| Neurobeachin-like 2 | NBEAL2 | S1363S1364 | | 12.7 | R.S*S*VGSGNTAGGGGSSGTLTPASQPGTPSPLDGPRPFPAAPGR.[illegible] |
| Neurobeachin-like 2 | NBEAL2 | S1367T1370 | 2.3 | 24.5 | R.SSVGS*GNT*AGGGGSSGTLTPASQPGTPSPLDGPRPFPAAPGR.[illegible] |
| Neurobeachin-like 2 | NBEAL2 | S759 | 3.2 | 74.9 | R.SQS*VPASTGLGWGSGLVAPLQEGSIDSTLAGTQDTR.W |
| Neurobeachin-like 2 | NBEAL2 | S757 | | 73.9 | R.S*QSVPASTGLGWGSGLVAPLQEGSIDSTLAGTQDTR.W |
| Neurofibromatosis type I | NF1 | S2515 | 9.5 | 21.1 | K.GSEGYLAATYPTVGQTS*PR.A |
| Neurofibromatosis type I | NF1 | S1140 | 100.0 | 23.8 | R.RLAS*LR.H |
| Neurofibromatosis type I | NF1 | T2514 | 6.0 | 12.1 | K.GSEGYLAATYPTVGQT*SPR.A |
| Neurofibromin 2 | NF2 | S518 | 100.0 | 29.7 | K.RLS*MEIEK.E |
| Neutrophil cytosolic factor 2 | NCF2 | T233 | 100.0 | 18.3 | K.T*PEIFR.A |
| NF kappaB activating protein | NKAP | S149 | 100.0 | 71.9 | R.IGELGAPEVWGLS*PK.N |
| NF kappaB activating protein | NKAP | S15 | 2.5 | 14.7 | R.SPDREAS*GSGGR.R |
| NFAT1 | NFATC2 | T216S221 | 20.6 | 18.9 | R.T*SPIMS*PR.T |
| NFAT1 | NFATC2 | S268 | 142.1 | 38.9 | R.HSCAEALVALPPGAS*PQR.S |
| NFAT1 | NFATC2 | S755S757 | 4.0 | 79.5 | R.S*KS*LSPSLLGYQQPALMAAPLSLADAHR.S |
| NFAT1 | NFATC2 | S56 | 7.8 | 30.6 | K.VASPPS*GPAYPDDVLDYGLKPYSPLASLSGEPPGR.F |
| NFAT1 | NFATC2 | S757S759 | 14.0 | 72.4 | K.S*LS*PSLLGYQQPALMAAPLSLADAHR.S |
| NFAT1 | NFATC2 | S326 | -0.3 | 52.3 | K.TS*PDPSPVSAAPSK.A |
| NFAT1 | NFATC2 | S859 | 7.8 | 101.4 | R.LSPGS*YPTVIQQQNATSQR.A |
| NFAT1 | NFATC2 | T803Y813 | 2.6 | 20.3 | R.SVLVHAGSQGQSSALLHPSPT*NQQASPVIHY*SPTNQQLR.C |
| NFAT1 | NFATC2 | S808Y813 | 6.7 | 22.3 | R.SVLVHAGSQGQSSALLHPSPTNQQAS*PVIHY*SPTNQQLR.C |
| NFAT1 | NFATC2 | S53 | 14.6 | 32.1 | K.VAS*PPSGPAYPDDVLDYGLKPYSPLASLSGEPPGR.F |
| NFAT1 | NFATC2 | S272S274S276 | | 34.3 | R. S*RS*RS*PQRSSHVAPQDHCSPACYPRVACSAVIMDALNSLATDSR |
| NFAT1 | NFATC2 | Y294S300S309 | 5.0 | 13.9 | R. SRSRSPQRSSHVAPQDHCSPACY*PRVACS*AVIMDALNS*LATDSR |
| NFAT1 | NFATC2 | S272S274S276 | | 34.9 | R. S*RS*RS*PQRS*SHVAPQDHCSPACYPRVACSAVIMDALNSLATDSR |
| NFAT1 | NFATC2 | S272S290Y294 | 5.3 | 34.8 | R. S*RSRSPQRSSHVAPQDHCS*PACY*PRVACS*AVIMDALNSLATDSR |
| NFAT1 | NFATC2 | S56Y72 | 7.5 | 20.6 | K.VASPPS*GPAYPDDVLDYGLKPY*SPLASLSGEPPGR.F |
| NFAT1 | NFATC2 | S856 | 12.6 | 88.1 | R.LS*PGSYPTVIQQQNATSQR.A |
| NFAT1 | NFATC2 | S274S276Y294 | 1.0 | 11.4 | R. S*RS*PQRSSHVAPQDHCSPACY*PRVACSAVIMDALNSLATDSPCGI |
| NFAT1 | NFATC2 | S274S276S280 | | 15.6 | R. S*RS*PQRS*SHVAPQDHCSPACYPRVACSAVIMDALNSLATDSPCGI |
| NFAT1 | NFATC2 | S53S73 | 30.5 | 17.2 | K.VAS*PPSGPAYPDDVLDYGLKPYS*PLASLSGEPPGR.F |
| NFAT1 | NFATC2 | S53Y72 | 10.7 | 17.2 | K.VAS*PPSGPAYPDDVLDYGLKPY*SPLASLSGEPPGR.F |
| NFAT1 | NFATC2 | S236S243 | 15.1 | 24.1 | R.TSLAEDSCLGRHS*PVPRPAS*R.S |
| NFAT1 | NFATC2 | S255S268 | 100.0 | 31.0 | K.RRHS*CAEALVALPPGAS*PQR.S |
| NFAT1 | NFATC2 | S759S761 | 4.5 | 91.1 | R.SKSLS*PS*LLGYQQPALMAAPLSLADAHR.S |
| NFAT1 | NFATC2 | S217S221 | 14.3 | 14.1 | R.TS*PIMS*PR.T |

Peak Area: <-10, -5, -3, 0, 3, 5, >10

%CV: 0, 29, 42, 71, 86, >100

**White dots-** Significant change in peptide abundance at **5** %FDR compared to the timepoint with the minimum peak area for a given PSM

| Protein Name | Gene | Phosphosites | CarT 0m | CarT 2m | CarT 5m | RajiB 0m | RajiB 2m | RajiB 5m | Ascor | MOWSE | Sequence |
|---|---|---|---|---|---|---|---|---|---|---|---|
| **NFAT1** | NFATC2 | S757S761 | | | | | | | **6.6** | **75.3** | R.SKS*LSPS*LLGYQQPALMAAPLSLADAHR.S |
| **NFAT1** | NFATC2 | T862 | | | | | | | **20.5** | **76.0** | R.LSPGSYPT*VIQQQNATSQR.A |
| **NFAT1** | NFATC2 | S759Y765 | | | | | | | **1.6** | **57.3** | K.SLS*PSLLGY*QQPALMAAPLSLADAHR.S |
| **NFAT1** | NFATC2 | S761Y765 | | | | | | | **0.0** | **31.7** | K.SLSPS*LLGY*QQPALMAAPLSLADAHR.S |
| **NFAT1** | NFATC2 | S230S243 | | | | | | | **6.5** | **27.7** | R.TSLAEDS*CLGRHSPVPRPAS*R.S |
| **NFAT1** | NFATC2 | S274S280S309 | | | | | | | **6.6** | **14.9** | R. [illegible] |
| **NFAT3** | NFATC4 | S221T226 | | | | | | | **55.4** | **43.1** | R.AS*PRPWT*PEDPWSLYGPSPGGR.G |
| **NFKB1** | NFKB1 | S893 | | | | | | | **85.0** | **56.2** | R.QMGYTEAIEVIQAAS*SPVK.T |
| **NFKB1** | NFKB1 | S894 | | | | | | | **7.2** | **52.0** | R.QMGYTEAIEVIQAASS*PVK.T |
| **NHS** | GALR1 | S836 | | | | | | | **13.9** | **26.3** | R.RSSS*LR.V |
| **NHS** | GALR1 | S834 | | | | | | | | **17.0** | R.RS*SSLR.V |
| **Nibrin** | NBN | S432 | | | | | | | **25.4** | **87.5** | R.IPNYQLS*PTKLPSINK.S |
| **Nibrin** | NBN | Y429 | | | | | | | **9.2** | **60.3** | R.IPNY*QLSPTKLPSINK.S |
| **NICE 4 protein** | UBAP2L | S453 | | | | | | | **19.8** | **38.7** | K.SPAVATSTAAPPPS*SPLPSK.S |
| **NICE 4 protein** | UBAP2L | S604S609 | | | | | | | **5.9** | **40.2** | R.RYPS*SISSS*PQKDLTQAK.N |
| **NICE 4 protein** | UBAP2L | S609 | | | | | | | **13.9** | **35.2** | R.RYPSSISSS*PQK.D |
| **NICE 4 protein** | UBAP2L | S454 | | | | | | | **7.0** | **27.4** | K.SPAVATSTAAPPPPSS*PLPSK.S |
| **NICE 4 protein** | UBAP2L | S605S608 | | | | | | | **11.1** | **13.3** | R.RYPSS*ISS*SPQK.D |
| **NICE 4 protein** | UBAP2L | Y602S608 | | | | | | | **14.0** | **40.4** | R.RY*PSSISS*SPQKDLTQAK.N |
| **NICE 4 protein** | UBAP2L | S604S608 | | | | | | | **12.6** | **34.5** | R.RYPS*SISS*SPQKDLTQAK.N |
| **NICE 4 protein** | UBAP2L | S605S607 | | | | | | | **6.5** | **15.8** | R.RYPSS*IS*SSPQKDLTQAK.N |
| **NICE 4 protein** | UBAP2L | S608 | | | | | | | **29.7** | **18.2** | R.RYPSSISS*SPQKDLTQAK.N |
| **NICE 4 protein** | UBAP2L | S458 | | | | | | | **7.9** | **50.1** | K.SPAVATSTAAPPPPSSPLPS*K.S |
| **NifU-like N-terminal domain containing** | ISCU | S14 | | | | | | | **100.0** | **50.3** | R.AAS*ALLLR.S |
| **NifU-like N-terminal domain containing** | ISCU | S29 | | | | | | | **100.0** | **23.3** | R.ELS*APAR.L |
| **Nin one binding protein** | NOB1 | S184 | | | | | | | **100.0** | **98.3** | R.GEDVPS*EEEEEEENGFEDR.K |
| **Nin one binding protein** | NOB1 | S201 | | | | | | | **144.2** | **124.3** | R.KDDS*DDDGGGWITPSNIK.Q |
| **NIP2** | BNIP2 | S114 | | | | | | | **19.8** | **55.3** | R.KGS*ITEYTAAEEK.E |
| | ZC3HC1 | S354S370 | | | | | | | **24.3** | **21.4** | R.S*WDSSSPVDRPEPEAAS*PTTR.T |
| **NIPA** | ZC3HC1 | S62 | | | | | | | **1.9** | **62.6** | K.DTSATSQSVNGS*PQAEQPSLESTSK.E |
| **NIPA** | ZC3HC1 | S58 | | | | | | | **8.9** | **32.2** | K.DTSATSQS*VNGSPQAEQPSLESTSK.E |
| **NIPA-like domain containing 3** | NIPAL3 | S372Y375 | | | | | | | **1.2** | **12.5** | K.ASFSYGALENNDNIS*EIY*APATLPVMQEEHGSR.S |
| **NIPA-like domain containing 3** | NIPAL3 | S359S361 | | | | | | | **9.1** | **17.0** | K.AS*FS*YGALENNDNISEIYAPATLPVMQEEHGSR.S |
| **NIR2** | PITPNM1 | S593S600 | | | | | | | **100.0** | **27.5** | R.RGS*MNNELLS*PEFGPVRDPLADGVEGLGR.G |
| **NIR2** | PITPNM1 | S373 | | | | | | | **46.8** | **76.3** | K.WNS*NDFIDAFASPVEAEGTPEPGAEAAK.G |
| **NIR2** | PITPNM1 | S593 | | | | | | | **29.7** | **14.8** | R.RGS*MNNELLSPEFGPVRDPLADGVEGLGR.G |
| **NIR3** | PITPNM2 | S1277 | | | | | | | **100.0** | **48.5** | R.KGS*FGLPGQGDFLR.S |
| **NIX** | BNIP3L | S168 | | | | | | | **14.7** | **22.2** | R.SVSLS*MR.K |
| **NIX** | BNIP3L | S164 | | | | | | | **9.2** | **21.9** | R.S*VSLSMR.K |
| **NIX** | BNIP3L | S166 | | | | | | | **17.4** | **27.0** | R.SVS*LSMR.K |
| **NIX** | BNIP3L | S63S64 | | | | | | | **-0.3** | **15.9** | K.NGGLEHVPSS*S*SIHNGDMEK.I |
| **NmrA-like family domain-containing protein 1 (Fragment)** | NMRAL1 | S218 | | | | | | | **100.0** | **29.6** | R.REGVS*R.A |
| **NMYC downstream regulated gene 1** | NDRG1 | S330 | | | | | | | **-0.1** | **30.4** | R.TAS*GSSVTSLDGTR.S |
| **NMYC downstream regulated gene 1** | NDRG1 | T366 | | | | | | | **6.5** | **39.7** | R.SHT*SEGAHLDITPNSGAAGNSAGPK.S |
| **NMYC downstream regulated gene 1** | NDRG1 | T346 | | | | | | | **16.9** | **19.0** | R.SHT*SEGTR.S |
| **NMYC downstream regulated gene 1** | NDRG1 | S330S333S336 | | | | | | | **9.7** | **33.3** | R.TAS*GSS*VTS*LDGTR.S |
| **NMYC downstream regulated gene 1** | NDRG1 | S330S332S336 | | | | | | | **8.1** | **15.8** | R.TAS*GS*SVTS*LDGTR.S |
| **NOL1/NOP2/Sun domain family, member 2** | NSUN2 | S743 | | | | | | | **80.9** | **75.3** | R.AGEPNS*PDAEEANSPDVTAGCDPAGVHPPR.- |
| **NOL1/NOP2/Sun domain family, member 2** | NSUN2 | S743S751 | | | | | | | **67.4** | **57.0** | R.AGEPNS*PDAEEANS*PDVTAGCDPAGVHPPR.- |
| **Non pou domain containing octamer binding protein** | NONO | T15 | | | | | | | **100.0** | **20.7** | K.QNHT*PR.K |
| **Non pou domain containing octamer binding protein** | NONO | T450 | | | | | | | **79.9** | **83.5** | R.FGQAATMEGIGAIGGT*PPAFNR.A |
| **Non-SMC condensin II complex, subunit H2** | NCAPH2 | S232S246 | | | | | | | **12.5** | **35.5** | R. [illegible] |
| **Non-SMC condensin II complex, subunit H2** | NCAPH2 | S284 | | | | | | | **69.9** | **76.9** | R.S*PQQSAALPR.R |
| **Non-SMC condensin II complex, subunit H2** | NCAPH2 | S232S240 | | | | | | | | **21.1** | R. [illegible] |
| **Non-SMC condensin II complex, subunit H2** | NCAPH2 | S240S246 | | | | | | | **0.1** | **13.3** | R. [illegible] |
| **NOP56** | NOP56 | S569 | | | | | | | **15.0** | **75.0** | K.EEPVS*SGPEEAVGK.S |
| **NOP56** | NOP56 | S569S570 | | | | | | | **100.0** | **39.2** | K.EEPVS*S*GPEEAVGK.S |
| **NOP56** | NOP56 | S467T468 | | | | | | | **3.4** | **32.6** | R.LAALALASSENSSS*T*PEECEEMSEKPK.K |
| **NOP56** | NOP56 | S520 | | | | | | | **4.8** | **54.1** | K.EELMSS*DLEETAGSTSIPK.R |
| **NOP56** | NOP56 | S519 | | | | | | | **66.9** | **70.1** | K.EELMS*SDLEETAGSTSIPK.R |
| **NOP56** | NOP56 | S519S520 | | | | | | | **73.8** | **114.6** | K.EELMS*S*DLEETAGSTSIPK.R |
| **NOT2** | CNOT2 | S161S169 | | | | | | | **2.7** | **27.7** | R.TNSMSSS*GLGSPNRS*SPSIICMPK.Q |

Peak Area: <-10, -5, -3, 0, 3, 5, >10

%CV: 0, 29, 42, 71, 86, >100

**White dots-** Significant change in peptide abundance at **5** %FDR compared to the timepoint with the minimum peak area for a given PSM

| Protein Name | Gene | Phosphosites | CarT 0m | CarT 2m | CarT 5m | RajiB 0m | RajiB 2m | RajiB 5m | Ascor | MOWSE | Sequence |
|---|---|---|---|---|---|---|---|---|---|---|---|
| **NOT2** | CNOT2 | S165S169 | | | | | | | **4.0** | **26.3** | R.TNSMSSSGLGS*PNRS*SPSIICMPK.Q |
| **NOT2** | CNOT2 | S165 | | | | | | | **27.8** | **78.2** | R.TNSMSSSGLGS*PNR.S |
| **NOT2** | CNOT2 | S101S120 | | | | | | | **2.8** | **12.8** | R.SLS*QGTQLPSHVTPTTGVPTMS*LHTPPSPSR.G |
| **NOT2** | CNOT2 | S160S161 | | | | | | | **6.8** | **12.6** | R.TNSMSS*S*GLGSPNRSSPSIICMPK.Q |
| **NOT4** | CNOT4 | T328 | | | | | | | **18.4** | **60.5** | R.SPFEGAVT*ESQSLFSDNFR.H |
| **NOT4** | CNOT4 | S321 | | | | | | | **3.6** | **73.7** | R.S*PFEGAVTESQSLFSDNFR.H |
| **Notch2** | NOTCH2 | S1841 | | | | | | | **5.5** | **36.0** | R.GGS*SDLSDEDEDAEDSSANIITDLVYQGASLQAQTDR.T |
| **NPAT** | NPAT | S775S779 | | | | | | | **8.8** | **29.7** | K.VIISDDPFVSSDTELTSAVSSINGENLPTIILS*PTKS*PTK.N |
| **NPAT** | NPAT | S774S779 | | | | | | | **13.9** | **31.9** | K.VIISDDPFVSSDTELTSAVSSINGENLPTIILS*SPTKS*PTK.N |
| **NPAT** | NPAT | S774T781 | | | | | | | **10.1** | **17.5** | K.VIISDDPFVSSDTELTSAVSSINGENLPTIILS*SPTKSPT*K.N |
| **NPAT** | NPAT | S762T770 | | | | | | | **5.1** | **16.4** | K.VIISDDPFVSSDTELTSAVSS*INGENLPT*IILSSPTKSPTK.N |
| **NSFL1C** | NSFL1C | S140 | | | | | | | **80.8** | **23.6** | K.S*PGETSKPRPFAGGGYR.L |
| **NSFL1C** | NSFL1C | S114 | | | | | | | **100.0** | **100.6** | K.S*PNELVDDLFK.G |
| **NTBA** | SLAMF6 | S277 | | | | | | | **12.1** | **64.6** | R.NLEYVSVS*PTNNTVYASVTHSNR.E |
| **NTBA** | SLAMF6 | S275 | | | | | | | | **40.8** | R.NLEYVS*VSPTNNTVYASVTHSNR.E |
| **Nuclear autoantigenic sperm protein** | NASP | T479S482 | | | | | | | **52.2** | **52.8** | K.EGEET*EGS*EEDDKENDKTEEMPNDSVLENK.S |
| **Nuclear autoantigenic sperm protein** | NASP | S728 | | | | | | | **100.0** | **45.9** | R.KPEEES*PR.K |
| **Nuclear autoantigenic sperm protein** | NASP | S505 | | | | | | | **100.0** | **81.1** | K.S*LQENEEEEIGNLELAWDMLDLAK.I |
| | NASP | S141 | | | | | | | **34.5** | **57.5** | K.TEDESLVENNDNIDETEGS*EEDDKENDK.T |
| **Nuclear autoantigenic sperm protein** | NASP | S127 | | | | | | | **12.8** | **41.4** | K.TEDES*LVENNDNIDETEGSEEDDKENDK.T |
| | NASP | T138S141 | | | | | | | **9.5** | **42.5** | K.TEDESLVENNDNIDET*EGS*EEDDKENDK.T |
| **Nuclear autoantigenic sperm protein** | NASP | T123S127 | | | | | | | | **49.5** | K.T*EDES*LVENNDNIDETEGSEEDDKENDKTEEMPNDSVLENK.S |
| **Nuclear autoantigenic sperm protein** | NASP | S127T138 | | | | | | | **11.1** | **42.1** | K.TEDES*LVENNDNIDET*EGSEEDDKENDKTEEMPNDSVLENK.S |
| | NASP | T138 | | | | | | | **6.8** | **56.4** | K.TEDESLVENNDNIDET*EGSEEDDKENDK.T |
| | NASP | S127S141 | | | | | | | **10.0** | **47.2** | K.TEDES*LVENNDNIDETEGS*EEDDKENDK.T |
| **Nuclear autoantigenic sperm protein** | NASP | T123 | | | | | | | | **16.7** | K.T*EDESLVENNDNIDETEGSEEDDKENDKTEEMPNDSVLENK.S |
| **Nuclear envelope pore membrane protein POM 121** | POM121 | S127 | | | | | | | **-0.2** | **96.6** | R.SS*SMSSLTGAYASGIPSSSR.N |
| **Nuclear envelope pore membrane protein POM 121** | POM121 | S130 | | | | | | | **16.1** | **75.2** | R.SSSMS*SLTGAYASGIPSSSR.N |
| **Nuclear envelope pore membrane protein POM 121** | POM121 | S127S131 | | | | | | | **4.3** | **41.5** | R.SS*SMSS*LTGAYASGIPSSSR.N |
| **Nuclear envelope pore membrane protein POM 121** | POM121 | S131 | | | | | | | **0.3** | **58.7** | R.SSSMSS*LTGAYASGIPSSSR.N |
| **Nuclear factor I/A** | NFIA | S300 | | | | | | | **49.3** | **66.6** | R.S*PGSGSQSSGWHEVEPGMPSPTTLKK.S |
| **Nuclear factor I/A** | NFIA | S258 | | | | | | | **23.9** | **47.6** | R.VSQTPIAAGTGPNFSLSDLESSSYYSMS*PGAMR.R |
| **Nuclear factor of activated T-cells, cytoplasmic, calcineurin-dependent 1** | NFATC1 | S233 | | | | | | | **54.3** | **66.3** | R.GLGACTLLGS*PR.H |
| **Nuclear factor related to kappa B binding protein** | NFRKB | S372S376 | | | | | | | **52.1** | **40.7** | K.SEAEDLAEPLSSTEGVAPLS*QAPS*PLAIPAIK.E |
| **Nuclear factor related to kappa B binding protein** | NFRKB | S376 | | | | | | | **27.2** | **51.1** | K.SEAEDLAEPLSSTEGVAPLSQAPS*PLAIPAIK.E |
| **Nuclear factor related to kappa B binding protein** | NFRKB | T365S376 | | | | | | | **8.3** | **16.5** | K.SEAEDLAEPLSST*EGVAPLSQAPS*PLAIPAIK.E |
| **Nuclear FMRP interacting protein 1** | NUFIP1 | S338S342S351 | | | | | | | **13.3** | **31.5** | K.LEGPPEANADPLGVLINS*DSES*DKEEKPQHS*VIPK.E |
| **Nuclear FMRP interacting protein 1** | NUFIP1 | S338S340S342 | | | | | | | **28.7** | **24.6** | K.LEGPPEANADPLGVLINS*DS*ES*DKEEKPQHSVIPK.E |
| **Nuclear mitotic apparatus protein 1** | NUMA1 | S1757 | | | | | | | **42.5** | **64.1** | R.TQPDGTSVPGEPAS*PISQR.L |
| **Nuclear mitotic apparatus protein 1** | NUMA1 | T2055 | | | | | | | **41.9** | **89.9** | R.QSMAFSILNT*PK.K |
| **Nuclear mitotic apparatus protein 1** | NUMA1 | S1862 | | | | | | | **82.2** | **78.1** | R.LGS*PDYGNSALLSLPGYRPTTR.S |
| **Nuclear mitotic apparatus protein 1** | NUMA1 | S1721S1724 | | | | | | | **30.5** | **51.4** | K.SREPQAKPQLDLS*IDS*LDLSCEEGTPLSITSK.L |
| **Nuclear mitotic apparatus protein 1** | NUMA1 | S1225 | | | | | | | **9.3** | **54.9** | R.KNS*LISSLEEEVSILNR.Q |
| **Nuclear mitotic apparatus protein 1** | NUMA1 | S1760 | | | | | | | **-9.8** | **21.2** | R.TQPDGTSVPGEPASPIS*QR.L |
| **Nuclear mitotic apparatus protein 1** | NUMA1 | S1721S1724S1 | | | | | | | **21.4** | **36.0** | K.SREPQAKPQLDLS*IDS*LDLS*CEEGTPLSITSK.L |
| **Nuclear mitotic apparatus protein 1** | NUMA1 | S200S203T211 | | | | | | | **16.8** | **41.1** | K.VASSSSGNNFLSGS*PAS*PMGDILQT*PQFQMR.R |
| **Nuclear mitotic apparatus protein 1** | NUMA1 | S1228 | | | | | | | **6.9** | **41.4** | R.KNSLIS*SLEEEVSILNR.Q |
| **Nuclear mitotic apparatus protein 1** | NUMA1 | S1769T1776 | | | | | | | **26.1** | **54.4** | K.VES*LESLYFT*PIPAR.S |
| **Nuclear mitotic apparatus protein 1** | NUMA1 | S1792S1800 | | | | | | | **5.1** | **11.2** | R.SQAPLESSLDS*LGDVFLDS*GRK.T |
| **Nuclear mitotic apparatus protein 1** | NUMA1 | S1788S1789 | | | | | | | **0.0** | **11.2** | R.SQAPLES*S*LDSLGDVFLDSGRK.T |
| **Nuclear protein UKp68** | ZC3H14 | S515 | | | | | | | **100.0** | **43.9** | R.DLVQPDKPAS*PK.F |
| **Nuclear protein UKp68** | ZC3H14 | S274 | | | | | | | **9.0** | **57.0** | R.LCEPEVLNSLEETYS*PFFR.N |
| **Nuclear protein UKp68** | ZC3H14 | S527S533 | | | | | | | **33.0** | **32.3** | K.FIVTLDGVPS*PPGYMS*DQEEDMCFEGMKPVNQTAASNK.G |
| **Nuclear protein UKp68** | ZC3H14 | Y531S533 | | | | | | | **10.1** | **11.4** | K.FIVTLDGVPSPPGY*MS*DQEEDMCFEGMKPVNQTAASNK.G |
| **Nuclear protein UKp68** | ZC3H14 | T521S527 | | | | | | | | **21.1** | K.FIVT*LDGVPS*PPGYMSDQEEDMCFEGMKPVNQTAASNK.G |
| **Nuclear receptor coactivator 5** | NCOA5 | S126 | | | | | | | **16.9** | **29.9** | R.EGS*YDR.Y |
| **Nuclear receptor coactivator 5** | NCOA5 | S29S34 | | | | | | | **100.0** | **15.5** | R.DRS*PIRGS*PR.R |
| **Nuclear receptor coactivator 5** | NCOA5 | S29 | | | | | | | **100.0** | **14.9** | R.DRS*PIR.G |
| **Nuclear receptor coactivator 5** | NCOA5 | S378T379 | | | | | | | **-0.4** | **38.8** | R.SS*T*DSLPGPISR.Q |
| **Nuclear receptor coactivator 6 interacting protein** | TGS1 | S89 | | | | | | | **41.9** | **95.1** | K.GIGLDESELDS*EAELMR.S |
| **Nuclear receptor coactivator 6 interacting protein** | TGS1 | S85S89 | | | | | | | **100.0** | **90.8** | K.GIGLDES*ELDS*EAELMR.S |
| **Nuclear receptor coactivator 7** | NCOA7 | S376 | | | | | | | **12.1** | **13.1** | K.KLDS*SR.E |

Peak Area: <-10, -5, -3, 0, 3, 5, >10

%CV: 0, 29, 42, 71, 86, >100

**White dots-** Significant change in peptide abundance at **5** %FDR compared to the timepoint with the minimum peak area for a given PSM

| Protein Name | Gene | Phosphosites | CarT 0m | CarT 2m | CarT 5m | RajiB 0m | RajiB 2m | RajiB 5m | Ascor | MOWSE | Sequence |
|---|---|---|---|---|---|---|---|---|---|---|---|
| **Nuclear receptor corepressor 1** | NCOR1 | S2120 | | | | | | | **100.0** | **36.8** | R.VS*PENLVDK.S |
| **Nuclear receptor corepressor 1** | NCOR1 | S2337 | | | | | | | **69.6** | **21.1** | K.S*PIPGQGYLGTERPSSVSSVHSEGDYHR.Q |
| **Nuclear receptor corepressor 1** | NCOR1 | S2151 | | | | | | | **50.4** | **60.5** | R.SHVSSEPYEPIS*PPQVPVVHEK.Q |
| **Nuclear receptor corepressor 1** | NCOR1 | S992T996 | | | | | | | **11.8** | **30.3** | R.SSTS*PCGT*SKSPNREWEVLQPAPHQVITNLPEGVR.L |
| **Nuclear receptor corepressor 1** | NCOR1 | S999 | | | | | | | **104.6** | **59.7** | K.S*PNREWEVLQPAPHQVITNLPEGVR.L |
| **Nuclear receptor corepressor 1** | NCOR1 | S2436S2438 | | | | | | | **11.5** | **18.7** | R.EPAPLLSAQYETLS*DS*DD.- |
| **Nuclear receptor corepressor 1** | NCOR1 | S2184 | | | | | | | **48.4** | **98.7** | R.S*PGSISYLPSFFTK.L |
| **Nuclear receptor corepressor 1** | NCOR1 | S992S999 | | | | | | | **-0.3** | **26.3** | R.SSTS*PCGTSKS*PNREWEVLQPAPHQVITNLPEGVR.L |
| **Nuclear receptor corepressor 1** | NCOR1 | S1977S1981 | | | | | | | **14.5** | **34.2** | R.YETPSDAIEVIS*PASS*PAPPQEK.L |
| **Nuclear receptor corepressor 1** | NCOR1 | S2395T2399 | | | | | | | **20.6** | **68.9** | R.MLSS*TPPT*PIACAPSAVNQAAPHQQNR.I |
| **Nuclear receptor corepressor 2** | NCOR2 | S149S152 | | | | | | | **102.2** | **53.9** | K.LEPVS*PPS*PPHTDPELELVPPR.L |
| **Nuclear receptor corepressor 2** | NCOR2 | S2056S2064 | | | | | | | **20.9** | **16.9** | R.SLGYHGSSYS*PEGVEPVS*PVSSPSLTHDK.G |
| **Nuclear receptor corepressor 2** | NCOR2 | S2056S2064S2 | | | | | | | **12.3** | **20.6** | R.SLGYHGSSYS*PEGVEPVS*PVS*SPSLTHDK.G |
| **Nuclear receptor corepressor 2** | NCOR2 | S2268 | | | | | | | **33.2** | **54.0** | K.S*PGNTSQPPAFFSK.L |
| **Nuclear receptor corepressor 2** | NCOR2 | S2056S2067 | | | | | | | **10.7** | **28.4** | R.SLGYHGSSYS*PEGVEPVSPVS*SPSLTHDK.G |
| **Nuclear receptor corepressor 2** | NCOR2 | S2054S2064S2 | | | | | | | **10.6** | **11.0** | R.SLGYHGSS*YSPEGVEPVS*PVS*SPSLTHDK.G |
| **Nuclear receptor corepressor 2** | NCOR2 | Y2055S2064S2 | | | | | | | **14.0** | **14.1** | R.SLGYHGSSY*SPEGVEPVS*PVSS*PSLTHDK.G |
| **Nuclear ubiquitous casein kinase and cyclin dependent kinase substrate** | NUCKS1 | S181 | | | | | | | **35.4** | **80.3** | R.LKATVTPS*PVK.G |
| **Nuclear ubiquitous casein kinase and cyclin dependent kinase substrate** | NUCKS1 | T179S181 | | | | | | | **32.2** | **27.3** | R.LKATVT*PS*PVK.G |
| **Nuclear ubiquitous casein kinase and cyclin dependent kinase substrate** | NUCKS1 | S19 | | | | | | | **83.0** | **114.9** | K.VVDYSQFQES*DDADEDYGR.D |
| **Nuclear ubiquitous casein kinase and cyclin dependent kinase substrate** | NUCKS1 | Y13 | | | | | | | **15.9** | **76.8** | K.VVDY*SQFQESDDADEDYGR.D |
| **Nuclear ubiquitous casein kinase and cyclin dependent kinase substrate** | NUCKS1 | T202S204 | | | | | | | **100.0** | **26.2** | K.EKT*PS*PK.E |
| **Nuclear ubiquitous casein kinase and cyclin dependent kinase substrate** | NUCKS1 | T222 | | | | | | | **27.6** | **23.9** | K.TST*SPPPEK.S |
| **Nuclear ubiquitous casein kinase and cyclin dependent kinase substrate** | NUCKS1 | S75S79 | | | | | | | **28.0** | **55.5** | K.DDSHS*AEDS*EDEKEDHK.N |
| **Nuclear ubiquitous casein kinase and cyclin dependent kinase substrate** | NUCKS1 | S58 | | | | | | | **42.5** | **96.6** | K.NSQEDS*EDSEDKDVK.T |
| **Nuclear ubiquitous casein kinase and cyclin dependent kinase substrate** | NUCKS1 | S54S58S61 | | | | | | | **19.2** | **46.1** | R.SGKNS*QEDS*EDS*EDKDVK.T |
| **Nuclear ubiquitous casein kinase and cyclin dependent kinase substrate** | NUCKS1 | S73S75S79 | | | | | | | **100.0** | **40.4** | K.DDS*HS*AEDS*EDEKEDHKNVR.Q |
| **Nuclear ubiquitous casein kinase and cyclin dependent kinase substrate** | NUCKS1 | S58S61 | | | | | | | **70.3** | **78.7** | K.NSQEDS*EDS*EDKDVK.T |
| **Nuclear ubiquitous casein kinase and cyclin dependent kinase substrate** | NUCKS1 | S214 | | | | | | | **100.0** | **59.7** | K.EEDEEPES*PPEKK.T |
| **Nuclear ubiquitous casein kinase and cyclin dependent kinase substrate** | NUCKS1 | T177T179 | | | | | | | **10.3** | **11.8** | K.AT*VT*PSPVKGK.G |
| **Nuclear ubiquitous casein kinase and cyclin dependent kinase substrate** | NUCKS1 | S234S240 | | | | | | | **46.1** | **41.6** | K.SGDEGS*EDEAPS*GED.- |
| **Nuclear ubiquitous casein kinase and cyclin dependent kinase substrate** | NUCKS1 | T179 | | | | | | | **13.9** | **76.1** | R.LKATVT*PSPVKGK.G |
| **Nuclear ubiquitous casein kinase and cyclin dependent kinase substrate** | NUCKS1 | S229S234S240 | | | | | | | **100.0** | **47.2** | K.S*GDEGS*EDEAPS*GED.- |
| **Nuclear ubiquitous casein kinase and cyclin dependent kinase substrate** | NUCKS1 | S130S132S144 | | | | | | | **75.7** | **97.3** | K.DS*GS*DEDFLM#EDDDDS*DYGSSK.K |
| **Nuclear ubiquitous casein kinase and cyclin dependent kinase substrate** | NUCKS1 | S130S132 | | | | | | | **91.1** | **100.2** | K.DS*GS*DEDFLMEDDDDSDYGSSK.K |
| **Nuclear ubiquitous casein kinase and cyclin dependent kinase substrate** | NUCKS1 | S223 | | | | | | | **10.8** | **42.2** | K.TSTS*PPPEK.S |
| **Nuclear ubiquitous casein kinase and cyclin dependent kinase substrate** | NUCKS1 | S54S58 | | | | | | | **7.6** | **28.4** | K.NS*QEDS*EDSEDKDVK.T |
| **Nuclear ubiquitous casein kinase and cyclin dependent kinase substrate** | NUCKS1 | S14 | | | | | | | **14.3** | **89.9** | R.KVVDYS*QFQESDDADEDYGR.D |
| **Nuclear ubiquitous casein kinase and cyclin dependent kinase substrate** | NUCKS1 | Y13S14 | | | | | | | **6.9** | **22.2** | R.KVVDY*S*QFQESDDADEDYGR.D |
| **Nuclear ubiquitous casein kinase and cyclin dependent kinase substrate** | NUCKS1 | S204 | | | | | | | **14.7** | **27.4** | K.EKTPS*PK.E |
| **Nuclear ubiquitous casein kinase and cyclin dependent kinase substrate** | NUCKS1 | S54S61 | | | | | | | **7.5** | **28.8** | K.NS*QEDSEDS*EDKDVK.T |
| | NUCKS1 | S113S130S132 | | | | | | | | **39.6** | R. [illegible] |
| **Nuclear ubiquitous casein kinase and cyclin dependent kinase substrate** | NUCKS1 | S50S54S58 | | | | | | | **8.7** | **15.2** | R.S*GKNS*QEDS*EDSEDKDVK.T |
| **Nuclear ubiquitous casein kinase and cyclin dependent kinase substrate** | NUCKS1 | T177S181 | | | | | | | **23.0** | **17.1** | R.LKAT*VTPS*PVKGK.G |
| **Nucleolar phosphoprotein p130** | NOLC1 | S84S85S86S88 | | | | | | | | **55.3** | K.KKAS*S*S*DS*EDS*S*EEEEEVQGPPAK.K |
| **Nucleolar phosphoprotein p130** | NOLC1 | S699 | | | | | | | **78.2** | **69.9** | R.GGSISVQVNSIKFDS*E.- |
| **Nucleolar phosphoprotein p130** | NOLC1 | S539 | | | | | | | **100.0** | **40.4** | K.GKGS*PRPQAPK.A |
| **Nucleolar phosphoprotein p130** | NOLC1 | S624 | | | | | | | **25.6** | **43.5** | R.ASS*PFR.R |
| **Nucleolar phosphoprotein p130** | NOLC1 | S644 | | | | | | | **100.0** | **23.1** | R.VADNS*FDAK.R |
| **Nucleolar phosphoprotein p130** | NOLC1 | T608T611 | | | | | | | **100.0** | **42.7** | K.LQT*PNT*FPK.R |
| | NOLC1 | S564 | | | | | | | **100.0** | **22.0** | K.NS*EEEEEEKKK.A |
| **Nucleolar phosphoprotein p130** | NOLC1 | S694 | | | | | | | **10.4** | **20.5** | R.GGSISVQVNS*IKFDSE.- |
| **Nucleolar protein 1** | NOP2 | S177T181 | | | | | | | **108.6** | **77.4** | R.EAAAGIQWS*EEET*EDEEEEKEVTPESGPPK.V |
| **Nucleolar protein 1** | NOP2 | S728 | | | | | | | **6.9** | **64.0** | K.GTDTQTPAVLS*PSK.T |
| **Nucleolar protein 1** | NOP2 | S782 | | | | | | | **14.7** | **24.8** | K.GPQPPTVS*PIR.S |
| **Nucleolar protein 8** | NOL8 | S296S298T302 | | | | | | | **100.0** | **39.3** | K.KRNS*IS*DDDT*DS*EDELR.M |
| **Nucleolar protein 8** | NOL8 | T888S890 | | | | | | | **100.0** | **41.4** | R.FLET*DS*EEEQEEVNEKK.T |
| **Nucleolar protein 8** | NOL8 | S1082S1083S1 | | | | | | | **79.6** | **123.4** | R.LQDS*S*S*EEEDVTEETDHR.N |
| **Nucleolar protein 8** | NOL8 | S298T302S304 | | | | | | | **48.2** | **66.2** | R.NSIS*DDDT*DS*EDELR.M |
| **Nucleolar protein 8** | NOL8 | S837S838S843 | | | | | | | **40.8** | **51.5** | K.LFDS*S*DDDES*DS*EDDSNRFK.I |
| | NOL8 | Y376S378 | | | | | | | **6.0** | **13.3** | R.EY*DS*GDTDEIIAMKK.N |
| **Nucleolar protein NOP5** | NOP58 | S502S514 | | | | | | | **29.1** | **53.1** | K.EEPLS*EEEPCTSTAIAS*PEKK.K |

**Peak Area**: <-10, -5, -3, 0, 3, 5, >10

**%CV**: 0, 29, 42, 71, 86, >100

**White dots**- Significant change in peptide abundance at **5**%FDR compared to the timepoint with the minimum peak area for a given PSM

| Protein Name | Gene | Phosphosites | CarT 0m | CarT 2m | CarT 5m | RajiB 0m | RajiB 2m | RajiB 5m | Ascor | MOWSE | Sequence |
|---|---|---|---|---|---|---|---|---|---|---|---|
| | NOP58 | S502 | | | | | | | 101.3 | 101.5 | K.EEPLS*EEEPCTSTAIASPEK.K |
| Nucleolar protein with MIF4G domain 1 | NOM1 | S317S320S321 | | | | | | | 40.0 | 36.3 | R.FAEDEEKS*ENS*S*EDGDITDK.S |
| | NCL | S145S153 | | | | | | | 100.0 | 101.1 | K.KEDS*DEEEDDDS*EEDEEDDEDEDEDEDEIEPAAM#K.A |
| Nucleolin | NCL | S184S206 | | | | | | | 70.9 | 77.0 | K.AAAAAPAS*EDEDDEDDEDDEDDDDDEEDDS*EEEAMETTPAK.G |
| Nucleolin | NCL | S153 | | | | | | | 26.3 | 146.8 | K.KEDSDEEEDDDS*EEDEEDDEDEDEDEDEIEPAAMK.A |
| Nucleolin | NCL | S145 | | | | | | | 53.4 | 135.0 | K.KEDS*DEEEDDDSEEDEEDDEDEDEDEDEIEPAAMK.A |
| Nucleolin | NCL | S41S42 | | | | | | | 17.0 | 33.2 | K.EVEEDSEDEEMSEDEEDDS*S*GEEVVIPQKK.G |
| Nucleolin | NCL | S34S41 | | | | | | | 27.0 | 35.3 | K.EVEEDSEDEEMS*EDEEDDS*SGEEVVIPQKK.G |
| | NCL | S28S34S41S42 | | | | | | | 100.0 | 38.4 | K.EVEEDS*EDEEM#S*EDEEDDS*S*GEEVVIPQKK.G |
| Nucleolin | NCL | S28S34S41 | | | | | | | 19.0 | 41.3 | K.EVEEDS*EDEEMS*EDEEDDS*SGEEVVIPQKK.G |
| Nucleolin | NCL | S67 | | | | | | | 25.8 | 44.5 | K.VVVS*PTKK.V |
| Nucleolin | NCL | T69 | | | | | | | 22.5 | 41.3 | K.VVVSPT*KK.V |
| Nucleolin | NCL | Y495 | | | | | | | -0.2 | 39.5 | K.TLVLSNLSY*SATEETLQEVFEK.A |
| Nucleolin | NCL | S184 | | | | | | | 4.3 | 42.6 | K.AAAAAPAS*EDEDDEDDEDDEDDDDDEEDDSEEEAMETTPAK.G |
| Nucleolin | NCL | S28S41 | | | | | | | 8.2 | 20.0 | K.EVEEDS*EDEEMSEDEEDDS*SGEEVVIPQKK.G |
| Nucleolin | NCL | S206 | | | | | | | 18.7 | 45.9 | K.AAAAAPASEDEDDEDDEDDEDDDDDEEDDS*EEEAMETTPAK.G |
| Nucleolin | NCL | S28S34S42 | | | | | | | -2.0 | 16.8 | K.EVEEDS*EDEEMS*EDEEDDSS*GEEVVIPQKK.G |
| Nucleolin | NCL | S34S41S42 | | | | | | | 5.6 | 11.0 | K.EVEEDSEDEEMS*EDEEDDS*S*GEEVVIPQK.K |
| Nucleolin | NCL | S34S42 | | | | | | | 7.5 | 18.8 | K.EVEEDSEDEEM#S*EDEEDDSS*GEEVVIPQKK.G |
| Nucleolin | NCL | S491 | | | | | | | | 22.3 | K.TLVLS*NLSYSATEETLQEVFEK.A |
| | NCL | S28S41S42 | | | | | | | 13.1 | 31.1 | K.EVEEDS*EDEEMSEDEEDDS*S*GEEVVIPQK.K |
| Nucleolin | NCL | S28S34 | | | | | | | 4.6 | 16.2 | K.EVEEDS*EDEEMS*EDEEDDSSGEEVVIPQKK.G |
| Nucleophosmin 1 | NPM1 | S243 | | | | | | | 23.9 | 36.5 | K.GPSS*VEDIK.A |
| Nucleophosmin 1 | NPM1 | S125 | | | | | | | 118.9 | 75.9 | K.CGSGPVHISGQHLVAVEEDAES*EDEEEEDVK.L |
| Nucleophosmin 1 | NPM1 | S70 | | | | | | | 14.6 | 69.3 | K.DELHIVEAEAM#NYEGS*PIK.V |
| Nucleophosmin 1 | NPM1 | S218 | | | | | | | 15.5 | 20.6 | K.DSKPSS*TPR.S |
| Nucleophosmin 1 | NPM1 | S214 | | | | | | | 8.1 | 12.9 | K.DS*KPSSTPR.S |
| Nucleophosmin 1 | NPM1 | S254 | | | | | | | 100.0 | 28.3 | K.MQAS*IEK.G |
| | NPM1 | Y67 | | | | | | | 3.6 | 32.3 | K.DELHIVEAEAM#NY*EGSPIKVTLATLK.M |
| Nucleophosmin 1 | NPM1 | S106 | | | | | | | | 51.3 | K.CGS*GPVHISGQHLVAVEEDAESEDEEEEDVK.L |
| Nucleophosmin 1 | NPM1 | S137 | | | | | | | 27.7 | 41.7 | K.LLS*ISGKR.S |
| Nucleophosmin 1 | NPM1 | S217 | | | | | | | 7.6 | 11.1 | K.DSKPS*STPR.S |
| Nucleoporin 160kDa | NUP160 | Y1151 | | | | | | | 12.7 | 35.1 | R.LIRPEYAWIVQPVSGAVY*DRPGASPK.R |
| Nucleoporin 160kDa | NUP160 | S1157 | | | | | | | 37.3 | 28.9 | R.LIRPEYAWIVQPVSGAVYDRPGAS*PK.R |
| Nucleoporin 205kDa | NUP205 | S1165 | | | | | | | 20.0 | 83.8 | R.S*VSGFLHFDTATK.V |
| Nucleoporin 205kDa | NUP205 | S1167 | | | | | | | -0.4 | 105.3 | R.SVS*GFLHFDTATK.V |
| Nucleoporin 50kDa | NUP50 | T219 | | | | | | | 10.9 | 38.1 | R.NSESESNKVAAET*QSPSLFGSTK.L |
| Nucleoporin 50kDa | NUP50 | S221 | | | | | | | 26.0 | 100.4 | K.VAAETQS*PSLFGSTK.L |
| Nucleosome assembly protein 1 like 1 | NAP1L1 | S10 | | | | | | | 100.3 | 68.3 | K.EQS*ELDQDLDDVEEVEEEETGEETK.L |
| Nucleosome assembly protein 1 like 1 | NAP1L1 | T62 | | | | | | | 14.0 | 62.0 | R.LDGLVET*PTGYIESLPR.V |
| Nucleosome assembly protein 1 like 1 | NAP1L1 | S143 | | | | | | | 1.8 | 30.9 | R.FEIINAIYEPTEEECEWKPDEEDEIS*EELK.E |
| Nucleosome assembly protein 1 like 4 | NAP1L4 | S125 | | | | | | | 38.9 | 71.4 | R.EFITGDVEPTDAESEWHS*ENEEEEK.L |
| Nucleosome assembly protein 1 like 4 | NAP1L4 | S304 | | | | | | | 21.1 | 114.9 | K.ASGDGES*LDEDSEFTLASDFEIGHFFR.E |
| Nucleosome assembly protein 1 like 4 | NAP1L4 | S304S309 | | | | | | | 5.5 | 60.4 | K.ASGDGES*LDEDS*EFTLASDFEIGHFFR.E |
| Nucleosome assembly protein 1 like 4 | NAP1L4 | S299S309 | | | | | | | 4.7 | 44.7 | K.AS*GDGESLDEDS*EFTLASDFEIGHFFR.E |
| Nucleosome assembly protein 1 like 4 | NAP1L4 | S299 | | | | | | | 15.0 | 65.2 | K.AS*GDGESLDEDSEFTLASDFEIGHFFR.E |
| Nucleosome assembly protein 1 like 4 | NAP1L4 | S121 | | | | | | | | 30.7 | R.EFITGDVEPTDAES*EWHSENEEEEK.L |
| Numb homolog | NUMB | S228S229 | | | | | | | | 22.6 | K.IVVGS*S*VAPGNTAPSPSSPTSPTSDATTSLEMNNPHAIPR.R |
| NUP107 | NUP107 | S86 | | | | | | | 31.7 | 38.0 | R.QPDISCILGTGGKS*PR.L |
| NUP133 | NUP133 | S45S50 | | | | | | | 43.2 | 49.1 | K.GLPLGSAVSS*PVLFS*PVGR.R |
| NUP133 | NUP133 | S44S50 | | | | | | | 53.6 | 15.7 | K.GLPLGSAVS*SPVLFS*PVGR.R |
| NUP133 | NUP133 | S57 | | | | | | | 31.9 | 19.0 | R.RSS*LSSR.G |
| NUP153 | NUP153 | S192 | | | | | | | 41.4 | 51.4 | R.AS*DKDITVSK.N |
| NUP153 | NUP153 | T102 | | | | | | | 9.1 | 37.4 | R.IT*PEPAVSNTEEPSTTSTASNYPDVLTRPSLHR.S |
| NUP153 | NUP153 | S333 | | | | | | | 76.7 | 101.0 | R.IPSIVS*SPLNSPLDR.S |
| NUP153 | NUP153 | S334S338 | | | | | | | 11.9 | 18.1 | K.RIPSIVSS*PLNS*PLDR.S |
| NUP210 | NUP210 | S1852 | | | | | | | 3.2 | 51.4 | R.ASPGHS*PHYFAASSPTSPNALPPAR.K |
| NUP210 | NUP210 | S1848 | | | | | | | 6.2 | 75.8 | R.AS*PGHSPHYFAASSPTSPNALPPAR.K |
| NUP210 | NUP210 | S1874 | | | | | | | 14.9 | 48.8 | R.KAS*PPSGLWSPAYASH.- |
| NUP210 | NUP210 | S1877 | | | | | | | 4.6 | 43.9 | R.KASPPS*GLWSPAYASH.- |
| NUP210 | NUP210 | Y1855 | | | | | | | 0.7 | 58.9 | R.ASPGHSPHY*FAASSPTSPNALPPAR.K |
| NUP210 | NUP210 | S1859 | | | | | | | 13.9 | 16.0 | R.ASPGHSPHYFAAS*SPTSPNALPPAR.K |

Peak Area: <-10, -5, -3, 0, 3, 5, >10

%CV: 0, 29, 42, 71, 86, >100

**White dots-** Significant change in peptide abundance at **5** %FDR compared to the timepoint with the minimum peak area for a given PSM

CarT: 0m 2m 5m — RajiB: 0m 2m 5m

| Protein Name | Gene | Phosphosites | Ascor | MOWSE | Sequence |
|---|---|---|---|---|---|
| **NUP214** | NUP214 | S433T436 | **14.6** | **43.5** | K.SPGS*TPT*TPTSSQAPQK.L |
| **NUP214** | NUP214 | S1023 | **15.7** | **81.0** | R.TPS*IQPSLLPHAAPFAK.S |
| **NUP214** | NUP214 | S433T437 | **11.8** | **47.0** | K.SPGS*TPTT*PTSSQAPQK.L |
| **NUP214** | NUP214 | S1081 | **2.8** | **33.0** | K.HGAPS*PSHPISAPQAAAAAALR.R |
| **NUP88** | NUP88 | S35 | **39.9** | **43.2** | K.NQS*PTEAEKPASSSLPSSPPPQLLTR.N |
| **NUP98** | NUP98 | S871 | **64.3** | **45.6** | K.YGLQDS*DEEEEEHPSK.T |
| **NUP98** | NUP98 | S606 | **32.1** | **78.7** | R.DSENLAS*PSEYPENGER.F |
| **NUP98** | NUP98 | S595 | **67.0** | **83.3** | K.NLNNSNLFS*PVNR.D |
| **NUP98** | NUP98 | S591 | **61.4** | **70.2** | K.NLNNS*NLFSPVNR.D |
| **NUP98** | NUP98 | S608 | **10.9** | **27.8** | R.DSENLASPS*EYPENGER.F |
| **Odin** | ANKS1A | S620 | **100.0** | **28.7** | K.AELKLS*R.S |
| **Odin** | ANKS1A | S663 | **35.1** | **70.5** | K.SPS*FASEWDEIEK.I |
| **Odin** | ANKS1A | S647 | **9.9** | **26.0** | R.SES*LSNCSIGK.K |
| **OIP106** | TRAK1 | S919 | **30.0** | **101.3** | R.S*FPTMVGSSMQMK.A |
| **Opa interacting protein 5** | OIP5 | S48 | **-0.2** | **70.3** | K.GSS*PLGPAGLGAEEPAAGPQLPSWLQPER.C |
| **Opioid growth factor receptor** | OGFR | S315 | **52.0** | **64.6** | R.KVEEEGS*PGDPDHEASTQGR.T |
| **Opioid growth factor receptor** | OGFR | S378 | **55.8** | **41.9** | R.SQGDEAGGHGEDRPEPLS*PK.E |
| **OSBP related protein 10** | OSBPL10 | S199 | **26.0** | **45.4** | R.SLTLLPHGTPNS*ASPCSQR.H |
| **OSBP related protein 10** | OSBPL10 | S201 | **6.7** | **55.7** | R.SLTLLPHGTPNSAS*PCSQR.H |
| **OSBP related protein 11** | OSBPL11 | S189 | **75.4** | **76.6** | R.RPS*QNAISFFNVGHSK.L |
| **OSBP related protein 11** | OSBPL11 | S181 | **12.9** | **73.1** | R.SFSLASSSNS*PISQR.R |
| **OSBP related protein 11** | OSBPL11 | S179 | **5.2** | **16.3** | R.SFSLASSS*NSPISQR.R |
| **OSBP related protein 9** | OSBPL9 | S334 | **17.6** | **33.5** | R.LIDSSGSASVLTHS*SSGNSLK.R |
| **OSBP related protein 9** | OSBPL9 | S329 | **6.4** | **24.3** | R.LIDSSGSAS*VLTHSSSGNSLK.R |
| **OSBP related protein 9** | OSBPL9 | S335 | **9.1** | **20.6** | R.LIDSSGSASVLTHSS*SGNSLK.R |
| **Osmosis responsive factor** | TTC33 | S197 | **15.6** | **52.2** | K.SEAPAEVTHFS*PK.S |
| **Osteoclast stimulating factor 1** | OSTF1 | S213 | **53.5** | **94.1** | R.TLSNAEDYLDDEDS*D.- |
| **Osteoclast stimulating factor 1** | OSTF1 | S202S213 | **26.9** | **84.2** | R.TLS*NAEDYLDDEDS*D.- |
| **Osteoclast stimulating factor 1** | OSTF1 | Y207 | **21.7** | **17.6** | R.TLSNAEDY*LDDEDSD.- |
| **Osteoclast stimulating factor 1** | OSTF1 | S202Y207 | **9.9** | **34.3** | R.TLS*NAEDY*LDDEDSD.- |
| **Osteoclast stimulating factor 1** | OSTF1 | Y207S213 | **3.5** | **38.2** | R.TLSNAEDY*LDDEDS*D.- |
| **Osteoclast stimulating factor 1** | OSTF1 | T200S202 | **6.9** | **43.8** | R.T*LS*NAEDYLDDEDSD.- |
| **Osteoclast stimulating factor 1** | OSTF1 | T200S213 | **6.1** | **14.1** | R.T*LSNAEDYLDDEDS*D.- |
| **Osteopetrosis associated transmembrane protein 1** | OSTM1 | T324 | **21.2** | **21.9** | K.SST*SFANIQENSN.- |
| **Osteopetrosis associated transmembrane protein 1** | OSTM1 | S325 | **-0.3** | **23.5** | K.SSTS*FANIQENSN.- |
| **OTUBAIN 1** | OTUB1 | S18 | **13.8** | **87.1** | K.QEPLGSDS*EGVNCLAYDEAIMAQQDR.I |
| **OTUBAIN 1** | OTUB1 | S16 | **16.2** | **82.7** | K.QEPLGS*DSEGVNCLAYDEAIMAQQDR.I |
| **Oxidative stress responsive 1** | OXSR1 | S339 | **92.5** | **98.5** | K.TEDGGWEWS*DDEFDEESEEGK.A |
| **Oxidative stress responsive 1** | OXSR1 | S325 | **20.1** | **29.8** | R.RVPGSS*GR.L |
| **Oxysterol binding protein** | OSBP | S379 | **10.9** | **88.4** | R.TGS*NISGASSDISLDEQYK.H |
| **Oxysterol binding protein** | OSBP | T377S382S385 | **17.2** | **76.2** | R.T*GSNIS*GAS*SDISLDEQYKHQLEETKK.E |
| **Oxysterol binding protein** | OSBP | S190S193 | **60.8** | **98.5** | K.MLAES*DES*GDEESVSQTDKTELQNTLR.T |
| **Oxysterol binding protein** | OSBP | S351 | **91.6** | **82.3** | K.GDMS*DEDDENEFFDAPEIITMPENLGHK.R |
| **Oxysterol binding protein** | OSBP | S379S382S385 | **21.1** | **124.5** | R.TGS*NIS*GAS*SDISLDEQYK.H |
| **Oxysterol binding protein** | OSBP | T367 | **13.6** | **25.7** | K.GDMSDEDDENEFFDAPEIIT*MPENLGHK.R |
| **Oxysterol binding protein** | OSBP | S382S386 | **11.9** | **47.8** | R.TGSNIS*GASS*DISLDEQYKHQLEETKK.E |
| **Oxysterol binding protein** | OSBP | T377S382 | **9.5** | **51.1** | R.T*GSNIS*GASSDISLDEQYK.H |
| **Oxysterol binding protein** | OSBP | T377S379S389 | **16.0** | **20.0** | R.T*GS*NISGASSDIS*LDEQYKHQLEETKK.E |
| **Oxysterol binding protein** | OSBP | S379S382 | **3.4** | **51.8** | R.TGS*NIS*GASSDISLDEQYK.H |
| **Oxysterol binding protein like 8** | OSBPL8 | S328S331S342 | **19.2** | **35.2** | K.DQDMYS*DKS*DKENDQEHDES*DNEVMGK.S |
| **Oxysterol binding protein like 8** | OSBPL8 | S807S808S810 | **39.3** | **25.0** | K.GYSSPEPDIQDS*S*GS*EAQSVKPSTR.R |
| **Oxysterol binding protein like 8** | OSBPL8 | S799S807S810 | **13.4** | **30.3** | K.GYSS*PEPDIQDS*SGS*EAQSVKPSTR.R |
| **Oxysterol binding protein like 8** | OSBPL8 | S798S799S810 | **10.8** | **35.8** | K.GYS*S*PEPDIQDSSGS*EAQSVKPSTR.R |
| **Oxysterol binding protein like 8** | OSBPL8 | S799S810S814 | **15.3** | **20.6** | K.GYSS*PEPDIQDSSGS*EAQS*VKPSTR.R |
| **Oxysterol binding protein like 8** | OSBPL8 | S799S808S810 | **9.4** | **27.6** | K.GYSS*PEPDIQDSS*GS*EAQSVKPSTR.R |
| **Oxysterol binding protein like 8** | OSBPL8 | Y327S328S342 | **13.3** | **27.2** | K.DQDMY*S*DKSDKENDQEHDES*DNEVMGK.S |
| **Oxysterol binding protein like 8** | OSBPL8 | S798S799S807 | **8.9** | **24.7** | K.GYS*S*PEPDIQDS*SGSEAQSVKPSTR.R |
| **Oxysterol binding protein like 8** | OSBPL8 | S798S799S807 | **25.8** | **20.9** | K.GYS*S*PEPDIQDS*S*GS*EAQSVKPSTR.R |
| **Oxysterol binding protein like 8** | OSBPL8 | S807S810S814 | **16.2** | **24.0** | K.GYSSPEPDIQDS*SGS*EAQS*VKPSTR.R |
| **Oxysterol binding protein like 8** | OSBPL8 | Y327S328S331 | **4.9** | **18.1** | K.DQDMY*S*DKS*DKENDQEHDESDNEVMGK.S |
| **Oxysterol binding protein like 8** | OSBPL8 | S799S807S808 | **2.5** | **27.5** | K.GYSS*PEPDIQDS*S*GSEAQSVKPSTR.R |
| **Oxysterol binding protein-like 3** | OSBPL3 | S304 | **7.2** | **54.5** | R.LHSS*NPNLSTLDFGEEK.N |

Peak Area: <-10, -5, -3, 0, 3, 5, >10

%CV: 0, 29, 42, 71, 86, >100

**White dots-** Significant change in peptide abundance at **5** %FDR compared to the timepoint with the minimum peak area for a given PSM

CarT: 0m 2m 5m — RajiB: 0m 2m 5m

| Protein Name | Gene | Phosphosites | Ascor | MOWSE | Sequence |
|---|---|---|---|---|---|
| Oxysterol binding protein-like 3 | OSBPL3 | S303 | 56.7 | 42.9 | R.LHS*SNPNLSTLDFGEEK.N |
| Oxysterol binding protein-like 3 | OSBPL3 | S251 | 12.1 | 71.0 | R.TYS*APAINAIQGGSFESPKK.E |
| P160 | MYBBP1A | S1163 | 5.2 | 40.0 | K.EIPSATQS*PISK.K |
| P160 | MYBBP1A | S1267 | 25.1 | 43.0 | K.NQKPSQVNGAPGS*PTEPAGQK.Q |
| P160 | MYBBP1A | S11 | 59.4 | 23.2 | R.DPAQPMS*PGEATQSGARPADR.Y |
| P160 | MYBBP1A | S731S732S734 | 17.7 | 11.0 | K.SEEGEDNRS*S*ES*EEES*EGEES*EEER.D |
| P160 | MYBBP1A | T1161 | 8.1 | 28.1 | K.EIPSAT*QSPISK.K |
| p21 activated kinase 3 | PAK3 | T421 | -0.3 | 68.5 | R.ST*MVGTPYWMAPEVVTR.K |
| p21 activated kinase 4 | PAK4 | S474 | 48.7 | 76.9 | K.S*LVGTPYWMAPELISR.L |
| p21 activated kinase 4 | PAK4 | T478 | 5.3 | 19.9 | K.SLVGT*PYWMAPELISR.L |
| p21 activated kinase 4 | PAK4 | S181 | 28.9 | 39.2 | R.DKRPLS*GPDVGTPQPAGLASGAK.L |
| p21 activated kinase 4 | PAK4 | Y480 | 2.7 | 40.0 | K.SLVGTPY*WMAPELISR.L |
| p53 induced gene 8 | EI24 | S56 | 100.0 | 39.5 | R.AQS*IER.K |
| p53 induced gene 8 | EI24 | S320 | -0.2 | 23.3 | K.TVYLQSALSSSTS*AEKFPSPHPSPAK.L |
| p53 induced gene 8 | EI24 | S316 | -0.0 | 14.6 | K.TVYLQSALS*SSTSAEKFPSPHPSPAK.L |
| PA2G4 | PA2G4 | T386 | 39.9 | 56.1 | K.TAENATSGET*LEENEAGD.- |
| Paf1, RNA polymerase II associated factor, homolog | PAF1 | S456S460S466 | 100.0 | 43.8 | R.DKEEIFGS*DADS*EDDADS*DDEDR.G |
| PAG | PAG1 | S296 | 9.1 | 30.9 | R.FSS*LSYK.S |
| PAG | PAG1 | S295 | 9.8 | 23.5 | K.RFS*SLSYK.S |
| PAG | PAG1 | S295S296 | 5.4 | 19.8 | K.RFS*S*LSYK.S |
| PAG | PAG1 | S351 | 35.4 | 92.8 | R.S*PSSCNDLYATVK.D |
| PAG | PAG1 | S229 | 23.9 | 55.3 | K.AEFAEYAS*VDR.N |
| PAG | PAG1 | S288S295S298 | 4.7 | 16.0 | K.EGGEAEESATDTTS*ETNKRFS*SLS*YK.S |
| PAG | PAG1 | S351S354 | 25.4 | 19.7 | K.SGQSLTVPESTYTSIQGDPQRS*PSS*CNDLYATVK.D |
| PAG | PAG1 | S170 | 19.7 | 65.8 | K.DSS*SQENMVEDCLYETVK.E |
| PAG | PAG1 | S301 | 26.8 | 78.0 | K.S*REEDPTLTEEEISAMYSSVNKPGQLVNK.S |
| PAG | PAG1 | S288 | 18.4 | 59.1 | K.EGGEAEESATDTTS*ETNKR.F |
| PAG | PAG1 | T286 | -0.3 | 38.3 | K.EGGEAEESATDT*TSETNKR.F |
| PAG | PAG1 | S295S298 | 12.6 | 21.4 | K.RFS*SLS*YK.S |
| PAG | PAG1 | S354 | 33.6 | 81.4 | R.SPSS*CNDLYATVK.D |
| PAG | PAG1 | S351S353 | 6.0 | 15.2 | K.SGQSLTVPESTYTSIQGDPQRS*PS*SCNDLYATVK.D |
| PAG | PAG1 | S169 | 67.8 | 53.4 | K.DS*SSQENMVEDCLYETVK.E |
| PAG | PAG1 | S353 | 11.7 | 82.3 | R.SPS*SCNDLYATVK.D |
| PAG | PAG1 | Y417 | 38.2 | 69.2 | K.ENDY*ESISDLQQGR.D |
| PAG | PAG1 | S171 | -0.2 | 99.6 | K.DSSS*QENMVEDCLYETVK.E |
| PAG | PAG1 | S150 | 51.7 | 38.0 | R.S*VDGDQGLGMEGPYEVLK.D |
| PAG | PAG1 | S298 | 7.8 | 24.1 | R.FSSLS*YK.S |
| PAG | PAG1 | S353S354 | -5.9 | 13.0 | K.SGQSLTVPESTYTSIQGDPQRSPS*S*CNDLYATVK.D |
| PAK2 | PAK2 | S197 | 81.1 | 41.1 | R.S*VIDPVPAPVGDSHVDGAAK.S |
| PAK2 | PAK2 | S141 | 25.3 | 56.3 | K.YLS*FTPPEK.D |
| PAK2 | PAK2 | S141S152 | 18.6 | 36.2 | K.YLS*FTPPEKDGFPS*GTPALNAK.G |
| PAK2 | PAK2 | S141T154 | 4.9 | 22.2 | K.YLS*FTPPEKDGFPSGT*PALNAK.G |
| PAK2 | PAK2 | T143S152 | 14.9 | 21.3 | K.YLSFT*PPEKDGFPS*GTPALNAK.G |
| PAK2 | PAK2 | T143 | | 50.1 | K.YLSFT*PPEKDGFPSGTPALNAK.G |
| PAK2 | PAK2 | S141T143 | 6.2 | 18.2 | K.YLS*FT*PPEKDGFPSGTPALNAK.G |
| PAK2 | PAK2 | Y139 | 13.9 | 35.1 | K.Y*LSFTPPEK.D |
| PAK2 | PAK2 | S55 | 28.4 | 45.3 | K.IIS*IFSGTEK.G |
| Palladin | PALLD | S892S894S897 | 100.0 | 21.9 | R.S*RS*RDS*GDENEPIQER.F |
| Palladin | PALLD | S669 | 37.6 | 35.7 | R.IAS*DEEIQGTK.D |
| Palladin | PALLD | S763S766 | 14.3 | 11.9 | K.QFIAAQNLGPASGHGTPASSPSSSSLPS*PMS*PTPR.Q |
| Pannexin 1 | PANX1 | S182 | 3.5 | 28.6 | R.DGACSVPGVTENLGQS*LWEVSESHFK.Y |
| Pantothenate kinase 4 | PANK4 | S393 | 18.4 | 93.1 | K.GAEQDNPNQYSWGENYAGSSGLMSAS*PELGPAQR.A |
| Pantothenate kinase 4 | PANK4 | S404 | 16.7 | 58.1 | R.S*GTFDLLEMDR.L |
| Pantothenate kinase 4 | PANK4 | Y383 | 12.4 | 22.4 | K.GAEQDNPNQYSWGENY*AGSSGLMSASPELGPAQR.A |
| Pantothenate kinase 4 | PANK4 | T406 | -0.8 | 36.3 | R.SGT*FDLLEMDR.L |
| PARD3 | PARD3 | S174 | 12.1 | 25.8 | R.WS*TTAGFLK.Q |
| PARN | PARN | S619 | 42.4 | 59.6 | K.ELS*PAGSISK.N |
| PARN | PARN | S628 | 16.4 | 36.3 | K.NS*PATLFEVPDTW.- |
| PARN | PARN | S570T589 | 8.4 | 112.3 | R.NLS*PSQEEAGLEDGVSGEISDT*ELEQTDSCAEPLSEGR.K |
| PARN | PARN | S570S583 | 7.8 | 72.4 | R.NLS*PSQEEAGLEDGVS*GEISDTELEQTDSCAEPLSEGR.K |
| PARN | PARN | S570S587 | 8.3 | 41.3 | R.NLS*PSQEEAGLEDGVSGEIS*DTELEQTDSCAEPLSEGR.K |
| Parvin beta | PARVB | T9 | 14.8 | 22.6 | R.SPT*PRPR.R |

Peak Area: <-10, -5, -3, 0, 3, 5, >10

%CV: 0, 29, 42, 71, 86, >100

**White dots-** Significant change in peptide abundance at **5** %FDR compared to the timepoint with the minimum peak area for a given PSM

CarT: 0m, 2m, 5m — RajiB: 0m, 2m, 5m

| Protein Name | Gene | Phosphosites | Ascor | MOWSE | Sequence |
|---|---|---|---|---|---|
| **Parvin beta** | PARVB | S7 | -4.7 | 18.2 | R.S*PTPRPR.R |
| **PAS kinase** | PASK | S119 | -0.4 | 24.5 | R.GLSSGWS*SPLLPAPVCNPNK.A |
| **Patatin like phospholipase domain containina 2** | PNPLA2 | S404 | 55.8 | 58.3 | R.VQS*LPSVPLSCAAYR.E |
| **Patatin like phospholipase domain containina 2** | PNPLA2 | S428 | 100.0 | 37.6 | R.NNLS*LGDALAK.W |
| **Patatin like phospholipase domain containina 2** | PNPLA2 | S407 | 15.8 | 43.7 | R.VQSLPS*VPLSCAAYR.E |
| **Paxillin** | PXN | S336 | 8.0 | 61.7 | K.TGSSS*PPGGPPKPGSQLDSMLGSLQSDLNK.L |
| **Paxillin** | PXN | S346 | 6.8 | 45.0 | K.TGSSSPPGGPPKPGS*QLDSMLGSLQSDLNK.L |
| **Paxillin** | PXN | T332 | 6.7 | 12.4 | K.T*GSSSPPGGPPKPGSQLDSMLGSLQSDLNK.L |
| **PC2** | CBX4 | S432S434 | | 50.4 | R.S*IS*TPTCLGGSPAAERPADLPPAAALPQPEVILLDSDLDEPIDLR.C |
| **PC2** | CBX4 | S432S434T435 | | 34.7 | R.S*IS*T*PTCLGGSPAAERPADLPPAAALPQPEVILLDSDLDEPIDLR.C |
| **PC2** | CBX4 | S434T435 | -0.8 | 12.7 | R.SIS*T*PTCLGGSPAAERPADLPPAAALPQPEVILLDSDLDEPIDLR.C |
| **PC2** | CBX4 | T437S442 | 0.3 | 26.4 | R.SISTPT*CLGGS*PAAERPADLPPAAALPQPEVILLDSDLDEPIDLR.C |
| **PC4 and SFRS1 interacting protein 1** | PSIP1 | T272S273S275 | 16.8 | 95.9 | K.TGVTST*S*DS*EEEGDDQEGEK.K |
| **PC4 and SFRS1 interacting protein 1** | PSIP1 | S177 | 10.9 | 50.3 | K.QVETEEAGVVTTATASVNLKVS*PK.R |
| **PC4 and SFRS1 interacting protein 1** | PSIP1 | S106 | 16.9 | 96.5 | K.QSNASS*DVEVEEK.E |
| **PC4 and SFRS1 interacting protein 1** | PSIP1 | S273S275 | 22.9 | 105.8 | K.TGVTSTS*DS*EEEGDDQEGEK.K |
| **PC4 and SFRS1 interacting protein 1** | PSIP1 | T272S275 | 17.3 | 75.4 | K.TGVTST*SDS*EEEGDDQEGEK.K |
| | PSIP1 | T272S273 | 9.9 | 125.6 | K.TGVTST*S*DSEEEGDDQEGEK.K |
| **PC4 and SFRS1 interacting protein 1** | PSIP1 | T270S275 | 6.7 | 26.3 | K.TGVT*STSDS*EEEGDDQEGEK.K |
| **PC4 and SFRS1 interacting protein 1** | PSIP1 | T270S271 | 13.9 | 13.6 | K.TGVT*S*TSDSEEEGDDQEGEK.K |
| **PC4 and SFRS1 interacting protein 1** | PSIP1 | S118S129 | 7.0 | 17.0 | K.ETSVS*KEDTDHEEKAS*NEDVTK.A |
| **PC4 and SFRS1 interacting protein 1** | PSIP1 | S271S275 | 3.5 | 31.9 | K.TGVTS*TSDS*EEEGDDQEGEKKR.K |
| **PC4 and SFRS1 interacting protein 1** | PSIP1 | S271T272 | 7.5 | 42.0 | K.TGVTS*T*SDSEEEGDDQEGEK.K |
| **PC4 and SFRS1 interacting protein 1** | PSIP1 | S271T272S273 | 7.7 | 52.6 | K.TGVTS*T*S*DSEEEGDDQEGEK.K |
| **PC4 and SFRS1 interacting protein 1** | PSIP1 | S271T272S275 | 9.8 | 44.7 | K.TGVTS*T*SDS*EEEGDDQEGEK.K |
| **PCTAIRE protein kinase 1** | CDK16 | S119 | 100.0 | 43.7 | K.RLS*LPADIR.I |
| **PCTAIRE protein kinase 1** | CDK16 | S153 | 32.3 | 44.9 | R.RVS*LSEIGFGK.L |
| **PCTAIRE protein kinase 2** | CDK17 | S9 | 19.8 | 30.6 | R.RLS*LTLR.G |
| **PCTAIRE protein kinase 2** | CDK17 | S180 | 40.2 | 58.8 | R.RAS*LSEIGFGK.M |
| **PCTAIRE protein kinase 2** | CDK17 | S182 | 10.9 | 33.4 | R.RASLS*EIGFGK.M |
| **PCTAIRE protein kinases 3** | CDK18 | S132 | 38.2 | 61.1 | R.RAS*LSDIGFGK.L |
| **PCTAIRE protein kinases 3** | CDK18 | S98 | 100.0 | 48.1 | K.RLS*LPMDIR.L |
| **PCTAIRE protein kinases 3** | CDK18 | S14 | 19.8 | 37.7 | R.RFS*LSVPR.T |
| | PDAP1 | S60S63 | 45.3 | 83.5 | K.KSLDS*DES*EDEEDDYQQK.R |
| **PDGF alpha associated protein 1** | PDAP1 | S63 | 0.7 | 32.9 | K.SLDSDES*EDEEDDYQQK.R |
| **PDGF alpha associated protein 1** | PDAP1 | S60 | 23.6 | 44.0 | K.SLDS*DESEDEEDDYQQK.R |
| **PDGF alpha associated protein 1** | PDAP1 | S57S60 | 8.3 | 20.6 | K.KS*LDS*DESEDEEDDYQQKR.K |
| **PDGF alpha associated protein 1** | PDAP1 | S57S63 | 6.8 | 40.3 | K.KS*LDSDES*EDEEDDYQQK.R |
| **PDZ and LIM domain 5** | PDLIM5 | S360 | 24.0 | 33.0 | K.S*PSWQRPNQGVPSTGR.I |
| **PDZ binding kinase** | PBK | S23S32 | 25.1 | 30.8 | K.SVLCS*TPTINIPAS*PFMQK.L |
| **PDZ binding kinase** | PBK | T26S32 | 13.9 | 13.8 | K.SVLCSTPT*INIPAS*PFMQK.L |
| **PDZ binding kinase** | PBK | T24S32 | 17.9 | 19.3 | K.SVLCST*PTINIPAS*PFMQK.L |
| **PEPP2** | PLEKHA5 | S885 | 65.4 | 27.2 | R.S*AVEQLCLAESTRPR.M |
| **PEPP2** | PLEKHA5 | T857 | 5.0 | 64.3 | R.AKSPT*PESSTIASYVTLR.K |
| | PLEKHA5 | S410 | 28.9 | 76.3 | R.TNS*M#QQLEQWIK.I |
| **PEPP2** | PLEKHA5 | S382 | 100.0 | 60.1 | K.IVNVS*LADLR.G |
| **PEPP2** | PLEKHA5 | S355 | 9.6 | 23.8 | K.LNSLPSEYES*GSACPAQTVHYRPINLSSSENK.I |
| **PEPP2** | PLEKHA5 | S855 | 10.1 | 55.5 | K.S*PTPESSTIASYVTLR.K |
| **PEPP2** | PLEKHA5 | S933 | 15.5 | 35.5 | K.GLNVIGASDQS*PLQSPSNLR.D |
| **PEPP2** | PLEKHA5 | T408 | 9.2 | 67.4 | R.T*NSMQQLEQWIK.I |
| **PEPP2** | PLEKHA5 | S372 | 4.7 | 18.5 | K.LNSLPSEYESGSACPAQTVHYRPINLS*SSENK.I |
| **PEPP2** | PLEKHA5 | Y366 | | 14.3 | K.LNSLPSEYESGSACPAQTVHY*RPINLSSSENK.I |
| **PEPP2** | PLEKHA5 | S861 | 0.8 | 15.2 | R.AKSPTPESS*TIASYVTLR.K |
| **PEPP2** | PLEKHA5 | S809 | 6.4 | 48.8 | K.S*EPELTTVAEVDESNGEEK.S |
| **Peptidyl prolyl isomerase G** | PPIG | S356T358 | 100.0 | 53.6 | R.S*ET*PPHWR.Q |
| **Peptidyl prolyl isomerase G** | PPIG | S413S415 | 100.0 | 30.5 | R.NVS*ES*PNRK.N |
| | PPIG | S290 | 100.0 | 26.9 | R.KS*PPKADEK.E |
| **Peptidyl prolyl isomerase G** | PPIG | S687 | 27.6 | 29.8 | K.ADRDQS*PFSK.I |
| **Peptidyl prolyl isomerase G** | PPIG | S744S745T748 | 100.0 | 18.8 | K.FDHES*S*PGT*DEDKS*G.- |
| **Peptidyl prolyl isomerase G** | PPIG | S587 | 30.0 | 21.7 | R.S*KEYHR.Y |
| **Peptidyl prolyl isomerase G** | PPIG | S254S256S257 | 69.1 | 44.6 | K.S*AS*S*ES*EAENLEAQPQSTVRPEEIPPIPENR.F |
| **Peptidyl prolyl isomerase G** | PPIG | S546 | 37.5 | 25.2 | R.S*RECDITK.G |

Peak Area: <-10, -5, -3, 0, 3, 5, >10

%CV: 0, 29, 42, 71, 86, >100

**White dots-** Significant change in peptide abundance at **5** %FDR compared to the timepoint with the minimum peak area for a given PSM

CarT (0m, 2m, 5m) — RajiB (0m, 2m, 5m)

| Protein Name | Gene | Phosphosites | Ascor | MOWSE | Sequence |
|---|---|---|---|---|---|
| **Peptidyl prolyl isomerase G** | PPIG | S415 | 13.4 | 15.5 | R.NVSES*PNRK.N |
| **Peptidyl prolyl isomerase like 4** | PPIL4 | S178 | 11.3 | 40.0 | R.INHTVILDDPFDDPPDLLIPDRS*PEPTR.E |
| **Peptidyl prolyl isomerase like 4** | PPIL4 | T182 | -0.1 | 22.1 | R.INHTVILDDPFDDPPDLLIPDRSPEPT*R.E |
| **Pericentrin 2** | PCNT | S2177 | 91.9 | 16.6 | K.NWDSLIPDEMPDS*PIQEK.S |
| **Pericentriolar material 1** | PCM1 | S65 | 67.7 | 95.3 | R.VTNDIS*PESSPGVGR.R |
| **Pericentriolar material 1** | PCM1 | S1958 | 100.0 | 67.9 | K.S*DEEDFVK.V |
| **Pericentriolar material 1** | PCM1 | T531Y536S538 | 5.0 | 81.0 | R.KDEET*EESEY*DS*EHENSEPVTNIR.N |
| **Pericentriolar material 1** | PCM1 | S1765S1768 | 32.8 | 50.9 | K.NVRS*DIS*DQEEDEESEGCPVSINLSK.A |
| **Pericentriolar material 1** | PCM1 | S1765S1768S1 | 53.3 | 44.5 | K.NVRS*DIS*DQEEDEES*EGCPVSINLSK.A |
| **Pericentriolar material 1** | PCM1 | S1257S1260S1 | 52.3 | 48.6 | R.RQFDEES*LES*FS*SMPDPVDPTTVTK.T |
| **Pericentriolar material 1** | PCM1 | S1257S1260S1 | 7.2 | 18.1 | R.QFDEES*LES*FSS*MPDPVDPTTVTK.T |
| **Pericentriolar material 1** | PCM1 | S1485S1494 | | 83.4 | K.IS*EQNDADNAS*VLSVSSNFEPFATDDLGNTVIHLDQALAR.M |
| **Pericentriolar material 1** | PCM1 | S1257S1262S1 | 5.0 | 50.4 | R.RQFDEES*LESFS*S*MPDPVDPTTVTK.T |
| **Pericentriolar material 1** | PCM1 | T531S534S538 | 15.8 | 70.4 | R.KDEET*EES*EYDS*EHENSEPVTNIR.N |
| **Pericentriolar material 1** | PCM1 | S1494S1497S1 | 8.2 | 59.3 | K.ISEQNDADNAS*VLS*VS*SNFEPFATDDLGNTVIHLDQALAR.M |
| **Pericentriolar material 1** | PCM1 | S65S69 | 22.4 | 28.5 | R.VTNDIS*PESS*PGVGR.R |
| **Pericentriolar material 1** | PCM1 | S65S68S69 | 35.3 | 53.0 | R.VTNDIS*PES*S*PGVGR.R |
| **Pericentriolar material 1** | PCM1 | T531S534Y536 | 5.9 | 17.6 | R.KDEET*EES*EY*DSEHENS*EPVTNIR.N |
| **Pericentriolar material 1** | PCM1 | S777S780 | 15.8 | 16.2 | K.IQALQTACPDLQLS*AAS*VGNCPTKK.Y |
| **Pericentriolar material 1** | PCM1 | T1481S1485S1 | | 20.9 | R. [illegible] |
| **Pericentriolar material 1** | PCM1 | S862S867 | 19.7 | 41.8 | R.RQGLAETAS*PVAVS*LR.S |
| **Pericentriolar material 1** | PCM1 | S534Y536S538 | -2.4 | 16.1 | R.KDEETEES*EY*DS*EHENS*EPVTNIR.N |
| **Pericentriolar material 1** | PCM1 | S1494S1499 | 0.3 | 29.7 | K.ISEQNDADNAS*VLSVS*SNFEPFATDDLGNTVIHLDQALAR.M |
| **Pericentriolar material 1** | PCM1 | S65S68 | 30.4 | 38.7 | R.VTNDIS*PES*SPGVGR.R |
| **Pericentriolar material 1** | PCM1 | T531S534Y536 | 7.8 | 61.7 | K.DEET*EES*EY*DSEHENSEPVTNIR.N |
| **Pericentriolar material 1** | PCM1 | T860S867 | 26.9 | 27.5 | R.RQGLAET*ASPVAVS*LR.S |
| **Pericentriolar material 1** | PCM1 | S1485S1494S1 | 2.5 | 33.8 | R. [illegible] |
| **Pericentriolar material 1** | PCM1 | S1485S1494S1 | | 60.1 | K.IS*EQNDADNAS*VLS*VSSNFEPFATDDLGNTVIHLDQALAR.M |
| **Pericentriolar material 1** | PCM1 | S1497S1499 | 2.2 | 61.4 | K.ISEQNDADNASVLS*VS*SNFEPFATDDLGNTVIHLDQALAR.M |
| **Pericentriolar material 1** | PCM1 | S534Y536S538 | 8.9 | 50.7 | K.DEETEES*EY*DS*EHENSEPVTNIR.N |
| **Pericentriolar material 1** | PCM1 | S1185S1187S11 | 12.4 | 15.9 | K.TEYMAFPKPFES*SS*S*IGAEKPR.N |
| **Pericentriolar material 1** | PCM1 | S1260S1262S1 | 0.5 | 18.9 | R.QFDEESLES*FS*S*MPDPVDPTTVTK.T |
| **Periodic tryptophan protein 1 homolog** | PWP1 | S50 | 42.7 | 60.9 | K.EKLQEEGGGS*DEEETGSPSEDGMQSAR.T |
| **Periodic tryptophan protein 1 homolog** | PWP1 | S50S57 | 8.7 | 60.1 | K.EKLQEEGGGS*DEEETGS*PSEDGMQSAR.T |
| **Periodic tryptophan protein 1 homolog** | PWP1 | S50T55 | 5.4 | 34.5 | K.EKLQEEGGGS*DEEET*GSPSEDGMQSAR.T |
| **Periphilin 1** | PPHLN1 | S133 | 49.5 | 32.1 | R.DNTFFRES*PVGR.K |
| **Periphilin 1** | PPHLN1 | T204 | 36.0 | 101.6 | R.DT*SPSSGSAVSSSK.V |
| **Periphilin 1** | PPHLN1 | S205 | 30.2 | 100.0 | R.DTS*PSSGSAVSSSK.V |
| **Periphilin 1** | PPHLN1 | T128 | 2.1 | 35.6 | R.DNT*FFRESPVGR.K |
| **Peroxisomal acyl-coenzyme A oxidase 1** | ACOX1 | S217 | -0.2 | 14.3 | R.SS*FSTVLL.- |
| **Pescadillo homolog 1** | PES1 | S457S488 | 100.0 | 56.5 | R. [illegible] |
| **PEST containing nuclear protein** | PCNP | S108 | 15.1 | 66.1 | K.TLS*VAAAFNEDEDSEPEEMPPEAK.M |
| **PEST containing nuclear protein** | PCNP | S119 | 29.2 | 67.6 | K.TLSVAAAFNEDEDS*EPEEM#PPEAK.M |
| **PEST containing nuclear protein** | PCNP | T106 | 0.9 | 13.9 | K.T*LSVAAAFNEDEDSEPEEMPPEAK.M |
| **PH domain containing protein** | PLEKHO2 | S439 | 15.3 | 56.3 | K.VLGSEPAPVS*AETLLSQAVEQLR.Q |
| **PHC2 polyhomeotic homolog 2** | PHC2 | T619S621 | 9.3 | 52.8 | K.LPQQDHTTTT*DS*EMEEPYLQESK.E |
| **PHD finger protein 10** | PHF10 | S56 | 14.7 | 68.2 | R.S*CETSSQDLGFSYYPAENLIEYK.W |
| **PHD finger protein 10** | PHF10 | S50S53 | 15.3 | 24.9 | R.MGS*GDS*SR.S |
| **PHD finger protein 14** | PHF14 | S208 | 39.4 | 90.7 | R.NRPLLDFVSMEELNDMDDYDS*EDDNDWRPTVVK.R |
| **PHD finger protein 14** | PHF14 | S25S26S29 | 3.7 | 13.5 | R.RQVKPLAASLLEALDYDS*S*DDS*DFK.V |
| **PHD finger protein 14** | PHF14 | Y23S25S29 | 49.3 | 21.4 | R.RQVKPLAASLLEALDY*DS*SDDS*DFK.V |
| **PHD finger protein 14** | PHF14 | Y23S26S29 | 7.7 | 14.8 | R.QVKPLAASLLEALDY*DSS*DDS*DFK.V |
| **PHD finger protein 14** | PHF14 | S25S26 | 6.1 | 54.1 | R.QVKPLAASLLEALDYDS*S*DDSDFK.V |
| **PHD finger protein 14** | PHF14 | S25S29 | 7.1 | 27.7 | R.QVKPLAASLLEALDYDS*SDDS*DFK.V |
| **PHD finger protein 14** | PHF14 | Y23S25S26 | 56.0 | 36.4 | R.RQVKPLAASLLEALDY*DS*S*DDSDFK.V |
| **PHD finger protein 14** | PHF14 | Y206 | 2.4 | 34.8 | R.NRPLLDFVSMEELNDMDDY*DSEDDNDWRPTVVK.R |
| **PHD finger protein 14** | PHF14 | S835 | 100.0 | 32.2 | R.S*FVPEEEKHEER.V |
| **PHD finger protein 14** | PHF14 | T287S290 | 18.5 | 75.7 | R.NSADDEELT*NDS*LTLSQSK.S |
| **PHD finger protein 14** | PHF14 | T217 | 7.0 | 16.9 | R.NRPLLDFVSMEELNDMDDYDSEDDNDWRPT*VVK.R |
| **PHD finger protein 2** | PHF2 | T541 | 2.8 | 64.5 | R.ESASPT*IPNLDLLEAHTK.E |
| **PHD finger protein 2** | PHF2 | S879 | 16.4 | 23.8 | K.DSDYVYPS*LESDEDNPIFK.S |
| **PHD finger protein 2** | PHF2 | S537 | 3.8 | 64.3 | R.ES*ASPTIPNLDLLEAHTK.E |

Peak Area: <-10, -5, -3, 0, 3, 5, >10

%CV: 0, 29, 42, 71, 86, >100

**White dots-** Significant change in peptide abundance at **5** %FDR compared to the timepoint with the minimum peak area for a given PSM

| Protein Name | Gene | Phosphosites | CarT 0m 2m 5m | RajiB 0m 2m 5m | Ascor | MOWSE | Sequence |
|---|---|---|---|---|---|---|---|
| **PHD finger protein 2** | PHF2 | S882 | | | **12.9** | **52.6** | K.DSDYVYPSLES*DEDNPIFK.S |
| **PHD finger protein 2** | PHF2 | S539 | | | **5.6** | **75.1** | R.ESAS*PTIPNLDLLEAHTK.E |
| **PHD finger protein 3** | PHF3 | S682 | | | **10.9** | **80.9** | K.SFS*LDEPPLFIPDNIATIR.R |
| **PHD finger protein 3** | PHF3 | S680 | | | **17.0** | **65.4** | K.S*FSLDEPPLFIPDNIATIR.R |
| **PHD finger protein 3** | PHF3 | S680S682 | | | **11.2** | **34.0** | R.SSKS*FS*LDEPPLFIPDNIATIR.R |
| **PHD finger protein 3** | PHF3 | S678S680 | | | **4.8** | **21.6** | R.RSS*KS*FSLDEPPLFIPDNIATIR.R |
| **PHD finger protein 6** | PHF6 | S145S154 | | | **27.6** | **36.0** | K.TAHNSEADLEES*FNEHELEPS*SPK.S |
| **PHD finger protein 6** | PHF6 | S154 | | | **61.4** | **48.0** | K.TAHNSEADLEESFNEHELEPS*SPK.S |
| **PHD finger protein 6** | PHF6 | S146S155 | | | **33.8** | **30.9** | K.TAHNSEAADLEES*FNEHELEPS*SPK.S |
| **PHD finger protein 6** | PHF6 | S155 | | | **51.0** | **41.3** | K.TAHNSEAADLEESFNEHELEPS*SPK.S |
| **PHD finger protein 6** | PHF6 | S145S155 | | | **16.9** | **23.4** | K.TAHNSEADLEES*FNEHELEPSS*PK.S |
| **PHD finger protein 6** | PHF6 | S199S203 | | | **22.2** | **20.2** | R.S*PHRS*SPSDTRPK.C |
| **PHD finger protein 6** | PHF6 | S138S154 | | | **39.6** | **14.7** | K.TAHNS*EADLEESFNEHELEPS*SPK.S |
| **PHD finger protein 6** | PHF6 | S199S204 | | | **11.4** | **17.9** | R.S*PHRSS*PSDTRPK.C |
| **Phosducin like** | PDCL | S296 | | | **39.4** | **28.8** | R.NSATCHSEDS*DLEID.- |
| **Phosducin like** | PDCL | Y16Y17S19S25 | | | **11.8** | **20.3** | K.LQYY*Y*SS*SEDEDS*DHEDKDR.G |
| **Phosducin like** | PDCL | S293S296 | | | **21.8** | **37.0** | R.NSATCHS*EDS*DLEID.- |
| **Phosducin like** | PDCL | T290S296 | | | **17.2** | **12.8** | R.NSAT*CHSEDS*DLEID.- |
| **Phosducin like** | PDCL | Y16Y17S18S19 | | | **11.1** | **19.6** | K.LQYY*Y*S*S*SEDEDSDHEDKDR.G |
| **Phosducin like** | PDCL | Y17S18S19S25 | | | **5.5** | **40.9** | K.LQYYY*S*S*SEDEDS*DHEDKDR.G |
| **Phosophatidylinositol 3 kinase, class 2 alpha** | PIK3C2A | S338 | | | **33.6** | **46.6** | R.SQS*LNIR.T |
| **Phosphatase and actin regulator** | PHACTR2 | S70 | | | | **22.8** | R.VHIS*GSVDGLDK.A |
| **Phosphatase and actin regulator 4** | PHACTR4 | S127 | | | **-0.3** | **54.3** | R.SS*SPVQVEEEPVR.L |
| **Phosphatase and actin regulator 4** | PHACTR4 | S638 | | | **31.2** | **48.3** | R.KLS*QRPTVAELLAR.K |
| **Phosphatase and actin regulator 4** | PHACTR4 | S474 | | | **61.5** | **59.2** | R.S*LPITIEMLK.V |
| **Phosphatase and actin regulator 4** | PHACTR4 | S354T368 | | | **22.2** | **14.3** | R.SPS*PPLPTHIPPEPPRT*PPFPAK.T |
| **Phosphatase and actin regulator 4** | PHACTR4 | T436T442 | | | **32.4** | **37.4** | R.IQQALT*SPLPMT*PILEGSHR.A |
| **Phosphatase and actin regulator 4** | PHACTR4 | S128 | | | **10.2** | **47.7** | R.SSS*PVQVEEEPVR.L |
| **Phosphatase and actin regulator 4** | PHACTR4 | T436S437 | | | **10.3** | **18.4** | R.IQQALT*S*PLPMTPILEGSHR.A |
| **Phosphate cytidylyltransferase 1 choline alpha isoform** | PCYT1A | S343S347 | | | **5.0** | **14.0** | K.TS*PPCS*PANLSR.H |
| **Phosphate cytidylyltransferase 1 choline alpha isoform** | PCYT1A | S315 | | | **100.0** | **14.3** | R.MLQAIS*PK.Q |
| **Phosphate cytidylyltransferase 1 choline alpha isoform** | PCYT1A | S362 | | | **31.9** | **41.4** | K.AAAYDIS*EDEED.- |
| **Phosphate cytidylyltransferase 1 choline alpha isoform** | PCYT1A | S329S331 | | | **22.7** | **15.9** | R.ERS*PS*PSFR.W |
| **Phosphate cytidylyltransferase 1 choline alpha isoform** | PCYT1A | S319S321 | | | **10.0** | **12.4** | K.QS*PS*SSPTR.E |
| **Phosphate cytidylyltransferase 1 choline, beta** | PCYT1B | S335 | | | **10.6** | **66.3** | R.SPS*PTFSWLPLK.T |
| **Phosphatidylinositol 3 phosphate 3 phosphatase adaptor subunit** | MTMR12 | S716 | | | **20.0** | **62.4** | R.HSS*KPVLPTSGWK.A |
| **Phosphatidylinositol 3-kinase, regulatory 4** | PIK3R4 | S903 | | | **-1.5** | **37.9** | R.SESSAGICVPLS*TSSQVPEVTTVQNK.K |
| **Phosphatidylinositol 4 kinase catalytic beta polypeptide** | PI4KB | S440 | | | **92.3** | **56.6** | R.S*VENLPECGITHEQR.A |
| **Phosphatidylinositol 4 kinase catalytic beta polypeptide** | PI4KB | S523 | | | **37.4** | **19.6** | R.RLS*EQLAHTPTAFK.R |
| **Phosphatidylinositol 4 kinase type II** | PI4K2A | S462 | | | **11.7** | **69.5** | R.SSS*ESYTQSFQSR.K |
| **Phosphatidylinositol 4 kinase type II** | PI4K2A | S47S51 | | | **12.9** | **34.0** | R.VAAAAGSGPS*PPGS*PGHDR.E |
| **Phosphatidylinositol 4-kinase alpha** | PI4KA | S172 | | | **100.0** | **24.6** | R.S*FNDFR.S |
| **Phosphatidylinositol transfer protein, beta** | PITPNB | S262 | | | **100.0** | **14.6** | R.KRGS*VR.G |
| **Phosphatidylinositol transfer protein, cytoplasmic 1** | PITPNC1 | S270 | | | **5.1** | **26.6** | R.S*APSSAPSTPLSTDAPEFLSVPK.D |
| **Phosphatidylinositol transfer protein, cytoplasmic 1** | PITPNC1 | S274 | | | **8.0** | **49.2** | R.SAPSS*APSTPLSTDAPEFLSVPK.D |
| **Phosphatidylinositol-3-phosphate/phosphatidylinositol 5-kinase** | PIKFYVE | T22S23 | | | **4.2** | **44.0** | R.SPT*S*PSHLTHFKPLTPDQDEPPFK.S |
| **Phosphatidylinositol-3-phosphate/phosphatidylinositol 5-kinase** | PIKFYVE | S307 | | | **18.9** | **56.6** | R.SAS*ITNLSLDR.S |
| **Phosphatidylinositol-3-phosphate/phosphatidylinositol 5-kinase** | PIKFYVE | S20S25 | | | **6.4** | **45.1** | R.S*PTSPS*HLTHFKPLTPDQDEPPFK.S |
| **Phosphatidylinositol-4-phosphate 5-kinase type II beta** | PIP4K2B | T322S326 | | | **50.5** | **63.0** | R.AEDEECENDGVGGNLLCSYGT*PPDS*PGNLLSFPR.F |
| **Phosphatidylinositol-4-phosphate 5-kinase type II beta** | PIP4K2B | Y320 | | | **6.6** | **49.9** | R.AEDEECENDGVGGNLLCSY*GTPPDSPGNLLSFPR.F |
| **Phosphatidylinositol-4-phosphate 5-kinase type II beta** | PIP4K2B | S319S326 | | | **22.3** | **28.8** | R.AEDEECENDGVGGNLLCS*YGTPPDS*PGNLLSFPR.F |
| **Phosphatidylinositol-4-phosphate 5-kinase type II beta** | PIP4K2B | Y320S326 | | | **20.2** | **17.1** | R.AEDEECENDGVGGNLLCSY*GTPPDS*PGNLLSFPR.F |
| **Phosphatidylinositol-4-phosphate 5-kinase type II beta** | PIP4K2B | T322 | | | **2.2** | **53.0** | R.AEDEECENDGVGGNLLCSYGT*PPDSPGNLLSFPR.F |
| **Phosphatidylserine synthase 1** | PTDSS1 | S417 | | | **28.6** | **61.1** | K.TYS*ECEDGTYSPEISWHHR.K |
| **Phosphatidylserine synthase 1** | PTDSS1 | Y416 | | | **-0.5** | **22.5** | K.TY*SECEDGTYSPEISWHHR.K |
| **Phosphatidylserine synthase 1** | PTDSS1 | T423 | | | **-1.7** | **10.9** | K.TYSECEDGT*YSPEISWHHR.K |
| **Phosphodiesterase 7A** | PDE7A | Y61 | | | **19.7** | **34.5** | R.RGSHPY*IDFR.I |
| **Phosphofructokinase 1** | PFKM | S775 | | | **100.0** | **19.0** | R.KRS*GEAAV.- |
| | PFKP | S386 | | | **89.8** | **73.8** | R.S*FAGNLNTYKR.L |
| **Phosphofurin acidic cluster sorting protein 1** | PACS1 | S779 | | | **29.0** | **29.0** | K.VGLVEDSPSTAGDGDDSPVVSLTVPS*TSPPSSSGLSR.D |
| **Phosphofurin acidic cluster sorting protein 1** | PACS1 | S379 | | | **19.3** | **78.8** | R.EVEEDLDELYDSLEMYNPS*DSGPEMEETESILSTPKPK.L |
| **Phosphofurin acidic cluster sorting protein 1** | PACS1 | S379S381 | | | **42.7** | **84.6** | R.EVEEDLDELYDSLEMYNPS*DS*GPEMEETESILSTPKPK.L |

Peak Area: <-10, -5, -3, 0, 3, 5, >10

%CV: 0, 29, 42, 71, 86, >100

**White dots-** Significant change in peptide abundance at **5** %FDR compared to the timepoint with the minimum peak area for a given PSM

| Protein Name | Gene | Phosphosites | CarT 0m | CarT 2m | CarT 5m | RajiB 0m | RajiB 2m | RajiB 5m | Ascor | MOWSE | Sequence |
|---|---|---|---|---|---|---|---|---|---|---|---|
| **Phosphofurin acidic cluster sorting protein 1** | PACS1 | S529S531 | | | | | | | **11.0** | **45.2** | R.TNSS*DS*ERSPDLGHSTQIPR.K |
| **Phosphofurin acidic cluster sorting protein 1** | PACS1 | Y370S372 | | | | | | | | **50.0** | R.EVEEDLDELY*DS*LEMYNPSDSGPEMEETESILSTPKPK.L |
| **Phosphofurin acidic cluster sorting protein 1** | PACS1 | S410 | | | | | | | **6.8** | **12.6** | K.LKPFFEGMSQSS*SQTEIGSLNSK.G |
| **Phosphofurin acidic cluster sorting protein 1** | PACS1 | S781 | | | | | | | **9.0** | **35.5** | K.VGLVEDSPSTAGDGDDSPVVSLTVPSTS*PPSSSGLSR.D |
| **Phosphofurin acidic cluster sorting protein 1** | PACS1 | S529S534 | | | | | | | **3.1** | **45.2** | R.TNSS*DSERS*PDLGHSTQIPR.K |
| **Phosphoglucomutase 1** | PGM1 | S117 | | | | | | | **6.1** | **68.0** | K.AIGGIILTAS*HNPGGPNGDFGIK.F |
| **Phosphoglucomutase 1** | PGM1 | T115 | | | | | | | | **76.2** | K.AIGGIILT*ASHNPGGPNGDFGIK.F |
| **Phosphoglucomutase 2 like 1** | PGM2L1 | T173 | | | | | | | **10.9** | **61.8** | K.AVAGVMIT*ASHNR.K |
| **Phosphoglucomutase 2 like 1** | PGM2L1 | S175 | | | | | | | **6.7** | **39.0** | K.AVAGVMITAS*HNR.K |
| **Phosphoglucomutase 3** | PGM3 | T62 | | | | | | | **9.1** | **118.8** | K.STIGVMVT*ASHNPEEDNGVK.L |
| **Phosphoglucomutase 3** | PGM3 | S64 | | | | | | | **13.0** | **119.7** | K.STIGVMVTAS*HNPEEDNGVK.L |
| **Phosphoglycerate kinase 1** | PGK1 | S203 | | | | | | | **100.0** | **71.2** | K.ALES*PERPFLAILGGAK.V |
| **Phosphoglycerate mutase 1** | PGAM1 | S14 | | | | | | | **100.0** | **67.3** | R.HGES*AWNLENR.F |
| **Phosphoglycerate mutase 1** | PGAM1 | S23 | | | | | | | **45.2** | **57.0** | R.FS*GWYDADLSPAGHEEAK.R |
| **Phosphoglycerate mutase 1** | PGAM1 | Y119 | | | | | | | **-0.2** | **47.8** | R.SY*DVPPPPMEPDHPFYSNISK.D |
| **Phosphoinositide 3 kinase adaptor protein 1** | PIK3AP1 | S759 | | | | | | | **19.6** | **25.0** | R.SRS*PGPPQVDGTPTMSLERPPR.V |
| **Phosphoinositide 3 kinase adaptor protein 1** | PIK3AP1 | S757 | | | | | | | **16.5** | **20.7** | R.S*RSPGPPQVDGTPTMSLERPPR.V |
| | AGAP2 | S312 | | | | | | | **13.9** | **23.2** | R.RTS*LFANR.R |
| **Phosphoinositide 3 kinase enhancer** | AGAP2 | Y272S278 | | | | | | | **6.3** | **14.8** | K.SLPSSPSHSAASTPVAGQASNGGHTSDY*SSSLPS*SPNVGHR.E |
| **Phosphoinositide 3 kinase enhancer** | AGAP2 | S245S248 | | | | | | | | **18.5** | K.S*LPS*SPSHSAASTPVAGQASNGGHTSDYSSSLPSSPNVGHR.E |
| **Phosphoinositide 3 kinase enhancer** | AGAP2 | T311 | | | | | | | **8.2** | **39.1** | K.RRT*SLFANR.R |
| **Phosphoinositide 3 kinase enhancer** | AGAP2 | T269S279 | | | | | | | **8.2** | **11.7** | K.SLPSSPSHSAASTPVAGQASNGGHT*SDYSSSLPSS*PNVGHR.E |
| **Phosphoinositide 3 kinase enhancer** | AGAP2 | S274S279 | | | | | | | **7.1** | **15.2** | K.SLPSSPSHSAASTPVAGQASNGGHTSDYSS*SLPSS*PNVGHR.E |
| **Phospholipase C like 1** | PLCL1 | S762S764 | | | | | | | **100.0** | **20.8** | K.RS*LS*VR.M |
| **Phospholipase C, beta 3** | PLCB3 | S537 | | | | | | | **100.0** | **13.9** | K.S*LGDEGLNR.G |
| **Phospholipase C, gamma 1** | PLCG1 | Y783 | | | | | | | **158.2** | **71.6** | R.NPGFY*VEANPMPTFK.C |
| **Phospholipase C, gamma 1** | PLCG1 | Y775Y783 | | | | | | | **59.8** | **41.9** | K.IGTAEPDYGALY*EGRNPGFY*VEANPMPTFK.C |
| | PLCG1 | S1248 | | | | | | | **9.8** | **21.0** | R.AREGS*FESR.Y |
| **Phospholipase C, gamma 1** | PLCG1 | Y771Y775Y783 | | | | | | | **48.0** | **15.4** | K.IGTAEPDY*GALY*EGRNPGFY*VEANPMPTFK.C |
| **Phosphoribosylformylglycinamidine synthase** | PFAS | S569 | | | | | | | **93.9** | **35.3** | R.S*PNRDFLTHVSAR.E |
| **Phosphorybosylaminoimidazole carboxylase** | PAICS | S27 | | | | | | | **112.3** | **58.9** | K.TKEVYELLDS*PGK.V |
| **Phosphorylase B kinase, beta** | PHKB | S700S701 | | | | | | | **4.3** | **46.0** | K.RQS*S*TPSAPELGQQPDVNISEWK.D |
| **Phosphorylase kinase liver alpha 2 subunit** | PHKA2 | T1041S1044 | | | | | | | **3.4** | **24.2** | R.SST*PSS*PTGTSSSDSGGHHIGWGER.Q |
| **Phosphorylase kinase liver alpha 2 subunit** | PHKA2 | T1041S1043 | | | | | | | **6.0** | **11.4** | R.SST*PS*SPTGTSSSDSGGHHIGWGER.Q |
| **Phosphorylated CTD interacting factor 1** | PCIF1 | S144 | | | | | | | **12.1** | **27.0** | K.IEIPVTPTGQSVPSS*PSIPGTPTLK.M |
| **Phosphorylated CTD interacting factor 1** | PCIF1 | T137S144 | | | | | | | **5.5** | **10.8** | K.IEIPVTPT*GQSVPSS*PSIPGTPTLK.M |
| **Phosphorylated CTD interacting factor 1** | PCIF1 | S144T150 | | | | | | | **13.1** | **12.1** | K.IEIPVTPTGQSVPSS*PSIPGT*PTLK.M |
| **Phosphorylated CTD interacting factor 1** | PCIF1 | S140S144 | | | | | | | **6.2** | **21.5** | K.IEIPVTPTGQS*VPSS*PSIPGTPTLK.M |
| **Phosphorylated CTD interacting factor 1** | PCIF1 | T137S143 | | | | | | | **6.5** | **19.6** | K.IEIPVTPT*GQSVPS*SPSIPGTPTLK.M |
| **Phosphorylated CTD interacting factor 1** | PCIF1 | S143T150 | | | | | | | **9.5** | **11.6** | K.IEIPVTPTGQSVPS*SPSIPGT*PTLK.M |
| **Phosphorylated CTD interacting factor 1** | PCIF1 | S143S146 | | | | | | | **8.4** | **11.0** | K.IEIPVTPTGQSVPS*SPS*IPGTPTLK.M |
| **PI3 kinase related kinase SMG1** | SMG1 | T3569S3570 | | | | | | | **26.2** | **22.7** | K.NLAT*S*ADTPPSTVPGTGK.S |
| **Pinin** | PNN | S347 | | | | | | | **100.0** | **55.5** | K.EIAIVHS*DAEK.E |
| **Pinin** | PNN | S99 | | | | | | | **122.4** | **67.1** | R.QES*DPEDDDVKKPALQSSVVATSK.E |
| **Pinin** | PNN | S442 | | | | | | | **18.6** | **81.4** | K.SLS*PGKENVSALDMEK.E |
| **Pinin** | PNN | S718S720 | | | | | | | **27.7** | **36.1** | R.SIS*ES*SR.S |
| **Pinin** | PNN | S684 | | | | | | | **15.8** | **19.6** | R.DTS*GLER.S |
| **Pinin** | PNN | S720S721 | | | | | | | **8.0** | **16.9** | R.SISES*S*R.S |
| **Pinin** | PNN | S380 | | | | | | | **34.9** | **80.4** | R.ESEKQQDS*QPEEVMDVLEMVENVK.H |
| **Pinin** | PNN | S718S721 | | | | | | | **16.4** | **28.0** | R.SIS*ESS*R.S |
| **Pinin** | PNN | S440S442 | | | | | | | **30.5** | **10.8** | K.S*LS*PGKENVSALDMEK.E |
| **Pinin** | PNN | S449 | | | | | | | **-18.5** | **30.9** | K.SLSPGKENVS*ALDMEK.E |
| **Pinin** | PNN | S716S718 | | | | | | | **7.6** | **16.3** | R.S*IS*ESSR.S |
| **PKD1like** | KIAA0319L | S998S1003S10 | | | | | | | **6.4** | **28.8** | K.GLLLSS*SLMHS*ES*ELDSDDAIFTWPDR.E |
| **PKD1like** | KIAA0319L | S998S1003S10 | | | | | | | **8.8** | **16.2** | K.GLLLSS*SLM#HS*ESELDS*DDAIFTWPDR.E |
| **PKR protein activator** | PRKRA | S18 | | | | | | | **39.3** | **75.2** | R.EDS*GTFSLGK.M |
| **Plasma membrane Ca(2+) ATPase type 4** | ATP2B4 | S328 | | | | | | | **47.3** | **41.5** | K.TQDGVALEIQPLNS*QEGIDNEEK.D |
| **Plasma membrane Ca(2+) ATPase type 4** | ATP2B4 | T1145 | | | | | | | **100.0** | **65.0** | R.T*PLLDEEEEENPDK.A |
| **Plasma membrane Ca(2+) ATPase, type 1** | ATP2B1 | S1140 | | | | | | | **9.6** | **59.6** | R.SS*IHNFMTHPEFR.I |
| **Plasma protease C1 inhibitor** | SERPING1 | T47 | | | | | | | | **13.3** | K.VAT*TVISK.M |
| **Platelet-activating factor receptor** | PTAFR | S339 | | | | | | | **50.2** | **22.3** | R.ATTDTVTEVVVPFNQIPGNS*LKN.- |
| **PLC, gamma 2** | PLCG2 | S677 | | | | | | | **32.1** | **50.6** | R.EGS*DSYAITFR.A |

Peak Area: <-10, -5, -3, 0, 3, 5, >10

%CV: 0, 29, 42, 71, 86, >100

**White dots**- Significant change in peptide abundance at 5 %FDR compared to the timepoint with the minimum peak area for a given PSM

| Protein Name | Gene | Phosphosites | CarT 0m | CarT 2m | CarT 5m | RajiB 0m | RajiB 2m | RajiB 5m | Ascor | MOWSE | Sequence |
|---|---|---|---|---|---|---|---|---|---|---|---|
| Pleckstrin | PLEK | S199T204 | | | | | | | | 41.0 | K.S*AVDGT*AENPFLDNPDAFYYFPDSGFFCEENSSDDDVILK.E |
| Pleckstrin | PLEK | S222S231 | | | | | | | 5.3 | 32.6 | K.SAVDGTAENPFLDNPDAFYYFPDS*GFFCEENSS*DDDVILK.E |
| Pleckstrin | PLEK | S230S231 | | | | | | | 4.8 | 27.0 | K.SAVDGTAENPFLDNPDAFYYFPDSGFFCEENS*S*DDDVILK.E |
| Pleckstrin | PLEK | S222S230 | | | | | | | 4.8 | 11.1 | K.SAVDGTAENPFLDNPDAFYYFPDS*GFFCEENS*SDDDVILK.E |
| Pleckstrin and Sec7 domain containing 4 | PSD4 | S118 | | | | | | | 11.5 | 46.4 | R.QDAPPWGSGVELTHLGS*PSAQR.E |
| Pleckstrin and Sec7 domain containing 4 | PSD4 | T114 | | | | | | | 17.8 | 21.5 | R.QDAPPWGSGVELT*HLGSPSAQR.E |
| Pleckstrin and Sec7 domain containing 4 | PSD4 | S461S464S465 | | | | | | | 41.7 | 10.7 | R.GPGPRPS*PAS*S*QEGS*PQLQHHSSGILPK.W |
| Pleckstrin homology domain containing, family F (with FYVE domain) member 2 | PLEKHF2 | S239S248 | | | | | | | 36.6 | 100.6 | K.SPLNDMS*DDDDDDDSS*D.- |
| Pleckstrin homology domain containing, family F (with FYVE domain) member 2 | PLEKHF2 | S239S247S248 | | | | | | | 81.6 | 107.8 | K.SPLNDMS*DDDDDDDS*S*D.- |
| Pleckstrin homology domain containing, family F (with FYVE domain) member 2 | PLEKHF2 | S226 | | | | | | | 23.9 | 38.2 | R.SDS*YSQSLK.S |
| Pleckstrin homology domain containing, family F (with FYVE domain) member 2 | PLEKHF2 | S239S247 | | | | | | | 46.6 | 50.7 | K.SPLNDMS*DDDDDDDS*SD.- |
| Pleckstrin homology domain containing, family M (with RUN domain) member 2 | PLEKHM2 | S544 | | | | | | | 3.0 | 29.2 | R.TGS*PGDAPERPPLCDFSEGLSAPMDFYR.F |
| Pleckstrin homology domain containing, family M (with RUN domain) member 2 | PLEKHM2 | T542 | | | | | | | 10.1 | 55.4 | R.T*GSPGDAPERPPLCDFSEGLSAPMDFYR.F |
| Pleckstrin homology domain interacting protein | PHIP | S1783 | | | | | | | 60.5 | 87.4 | R.TAFYNEDDS*EEEQR.Q |
| Pleckstrin homology domain interacting protein | PHIP | S911 | | | | | | | 100.0 | 45.6 | K.DGPIS*PK.K |
| Pleckstrin homology domain interacting protein | PHIP | S674 | | | | | | | 8.9 | 55.8 | R.GS*ISSTSEVHSPPNVGLR.R |
| | PHIP | S1315 | | | | | | | 16.9 | 12.6 | R.AQS*YDIQAWKK.Q |
| Pleckstrin homology domain-containing family A member 2 | PLEKHA2 | S134 | | | | | | | 15.1 | 20.7 | R.SQS*YIPTSGCR.A |
| Pleckstrin homology domain-containing family A member 2 | PLEKHA2 | S351 | | | | | | | 100.0 | 20.9 | R.S*EPQHPK.E |
| Pleckstrin homology domain-containing family A member 2 | PLEKHA2 | S299 | | | | | | | 24.3 | 24.3 | K.APS*VASSWQPWTPVPQAGEK.L |
| Pleckstrin homology domain-containing family A member 2 | PLEKHA2 | S132 | | | | | | | 12.3 | 18.4 | R.S*QSYIPTSGCR.A |
| Plectin 1 | PLEC | T4623 | | | | | | | 27.6 | 78.6 | K.GYYSPYSVSGSGST*AGSR.T |
| Plectin 1 | PLEC | S4626 | | | | | | | 19.1 | 64.7 | K.GYYSPYSVSGSGSTAGS*R.T |
| Plectin 1 | PLEC | S4386 | | | | | | | 11.7 | 79.4 | R.SSS*VGSSSSYPISPAVSR.T |
| Plectin 1 | PLEC | S4620S4622 | | | | | | | 3.7 | 20.6 | K.GYYSPYSVSGS*GS*TAGSR.T |
| Plectin 1 | PLEC | S21 | | | | | | | 16.9 | 97.8 | R.TSS*EDNLYLAVLR.A |
| Plectin 1 | PLEC | S4385 | | | | | | | -0.2 | 82.2 | R.SS*SVGSSSSYPISPAVSR.T |
| Plectin 1 | PLEC | S20 | | | | | | | 10.8 | 90.3 | R.TS*SEDNLYLAVLR.A |
| Plectin 1 | PLEC | S4620 | | | | | | | 9.1 | 33.6 | K.GYYSPYSVSGS*GSTAGSR.T |
| Plectin 1 | PLEC | S4622 | | | | | | | 7.8 | 22.1 | K.GYYSPYSVSGSGS*TAGSR.T |
| Plectin 1 | PLEC | S4622T4623 | | | | | | | 1.5 | 53.5 | K.GYYSPYSVSGSGS*T*AGSR.T |
| Plectin 1 | PLEC | S4618 | | | | | | | 10.9 | 29.1 | K.GYYSPYSVS*GSGSTAGSR.T |
| Plectin 1 | PLEC | S4618T4623 | | | | | | | 0.6 | 43.7 | K.GYYSPYSVS*GSGST*AGSR.T |
| PLEKHA7 | PLEKHA7 | T992 | | | | | | | 8.3 | 107.4 | R.SYVSEPELAT*LSGDMAQPSLGLVGPESR.Y |
| PLEKHA7 | PLEKHA7 | S994 | | | | | | | 4.2 | 88.5 | R.SYVSEPELATLS*GDMAQPSLGLVGPESR.Y |
| PLEKHA7 | PLEKHA7 | S986 | | | | | | | 7.1 | 98.5 | R.SYVS*EPELATLSGDMAQPSLGLVGPESR.Y |
| PML | PML | S518S527 | | | | | | | 100.0 | 51.1 | K.AVS*PPHLDGPPS*PR.S |
| PML | PML | S530 | | | | | | | 82.5 | 76.0 | R.S*PVIGSEVFLPNSNHVASGAGEAGR.E |
| PML | PML | S530 | | | | | | | 108.4 | 76.0 | R.S*PVIGSEVFLPNSNHVASGAGEAEER.V |
| PML | PML | S505 | | | | | | | -0.3 | 39.0 | R.SS*PEQPRPSTSK.A |
| PML | PML | S518S527S530 | | | | | | | 18.4 | 20.5 | K.AVS*PPHLDGPPS*PRS*PVIGSEVFLPNSNHVASGAGEAEER.V |
| PML | PML | S518S527S530 | | | | | | | 12.5 | 25.2 | K.AVS*PPHLDGPPS*PRS*PVIGSEVFLPNSNHVASGAGEAGR.E |
| PML | PML | S504 | | | | | | | 12.4 | 33.6 | R.S*SPEQPRPSTSK.A |
| Poly A polymerase alpha | PAPOLA | S24 | | | | | | | 23.9 | 58.0 | K.HYGITS*PISLAAPK.E |
| Poly adenylate binding protein 2 | PABPN1 | S150 | | | | | | | 74.7 | 36.4 | K.QMNMS*PPPGNAGPVIMSIEEK.M |
| Poly(rC) binding protein 1 | PCBP1 | S189 | | | | | | | 29.8 | 57.9 | R.VMTIPYQPMPAS*SPVICAGGQDR.C |
| Poly(rC) binding protein 1 | PCBP1 | S262 | | | | | | | 53.2 | 42.0 | R.QQSHFAMMHGGTGFAGIDS*SSPEVK.G |
| Poly(rC) binding protein 1 | PCBP1 | S190 | | | | | | | 4.6 | 68.5 | R.VMTIPYQPMPASS*PVICAGGQDR.C |
| Poly(rC) binding protein 1 | PCBP1 | Y183 | | | | | | | 21.2 | 72.0 | R.VMTIPY*QPMPASSPVICAGGQDR.C |
| Poly(rC) binding protein 1 | PCBP1 | S264 | | | | | | | 9.1 | 26.4 | R.QQSHFAMMHGGTGFAGIDSSS*PEVK.G |
| Poly(rC) binding protein 2 | PCBP2 | S189 | | | | | | | 34.0 | 40.1 | K.GVTIPYRPKPSSS*PVIFAGGQDR.Y |
| Poly(rC) binding protein 2 | PCBP2 | Y182 | | | | | | | 13.1 | 28.3 | K.GVTIPY*RPKPSSSPVIFAGGQDR.Y |
| Poly(rC) binding protein 2 | PCBP2 | S187 | | | | | | | 12.2 | 21.1 | K.GVTIPYRPKPS*SSPVIFAGGQDR.Y |
| Poly(rC) binding protein 2 | PCBP2 | S270 | | | | | | | -1.4 | 16.0 | K.LHQLAMQQSHFPMTHGNTGFSGIES*SSPEVK.G |
| Poly(rC) binding protein 2 | PCBP2 | S272 | | | | | | | 3.6 | 21.2 | K.LHQLAMQQSHFPMTHGNTGFSGIESSS*PEVK.G |
| Poly(rC) binding protein 2 | PCBP2 | S188 | | | | | | | 16.9 | 39.3 | K.GVTIPYRPKPSS*SPVIFAGGQDR.Y |
| Polycomb protein SUZ12 | SUZ12 | S546 | | | | | | | 89.9 | 131.8 | K.ASMSEFLES*EDGEVEQQR.T |
| Polycomb protein SUZ12 | SUZ12 | S539 | | | | | | | 12.2 | 78.6 | K.AS*MSEFLESEDGEVEQQR.T |
| Polycomb protein SUZ12 | SUZ12 | S583 | | | | | | | 15.5 | 20.4 | R.LYFHSDTCLPLRPQEMEVDS*EDEKDPEWLR.E |
| Polyhomeotic like 3 | PHC3 | S773 | | | | | | | 32.7 | 22.1 | K.HADNS*SDTEMEDMIAEETLEEMDSELLK.C |
| Polyhomeotic like 3 | PHC3 | S774 | | | | | | | 7.8 | 71.4 | K.HADNSS*DTEMEDMIAEETLEEMDSELLK.C |
| Polymyositis/scleroderma autoantigen 1 | EXOSC9 | S306 | | | | | | | 25.6 | 61.8 | K.APIDTS*DVEEK.A |

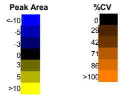


**White dots-** Significant change in peptide abundance at 5 %FDR compared to the timepoint with the minimum peak area for a given PSM

CarT: 0m 2m 5m | RajiB: 0m 2m 5m

| Protein Name | Gene | Phosphosites | Ascor | MOWSE | Sequence |
|---|---|---|---|---|---|
| Polymyositis/scleroderma autoantigen 1 | EXOSC9 | S392S394 | 11.5 | 30.5 | K.MDTGVEVSDIGSQDAPIILS*DS*EEEEMIILEPDKNPK.K |
| Polymyositis/scleroderma autoantigen 1 | EXOSC9 | S384S392 | 16.9 | 14.5 | K.MDTGVEVSDIGS*QDAPIILS*DSEEEEMIILEPDKNPK.K |
| Polynucleotide kinase 3 prime phosphatase | PNKP | T118T122 | 31.4 | 34.7 | R.TPESQPDT*PPGT*PLVSQDEKR.D |
| Polypyrimidine tract binding protein 1 | PTBP1 | S141 | 23.9 | 54.5 | K.TDSS*PNQAR.A |
| Polypyrimidine tract binding protein 1 | PTBP1 | S140 | 16.9 | 55.8 | K.TDS*SPNQAR.A |
| POM121 membrane glycoprotein-like | POM121C | S80S81S83 | | 15.7 | R.RHDS*S*GS*GHSAFEPLVASGVPASFVPKPGSLK.R |
| POM121 membrane glycoprotein-like | POM121C | S127 | -0.5 | 92.6 | R.SS*SMSSLTGAYTSGIPSSSR.N |
| POM121 membrane glycoprotein-like | POM121C | S81S83S86 | 4.5 | 24.2 | R.RHDSS*GS*GHS*AFEPLVASGVPASFVPKPGSLK.R |
| POM121 membrane glycoprotein-like | POM121C | S128S130 | 13.5 | 85.7 | R.SSS*MS*SLTGAYTSGIPSSSR.N |
| POM121 membrane glycoprotein-like | POM121C | S127S130 | 5.9 | 108.9 | R.SS*SMS*SLTGAYTSGIPSSSR.N |
| POM121 membrane glycoprotein-like | POM121C | S127S131 | 10.3 | 93.0 | R.SS*SMSS*LTGAYTSGIPSSSR.N |
| POM121 membrane glycoprotein-like | POM121C | S128S131 | 11.8 | 116.6 | R.SSS*MSS*LTGAYTSGIPSSSR.N |
| POM121 membrane glycoprotein-like | POM121C | S130 | 7.6 | 54.9 | R.SSSMS*SLTGAYTSGIPSSSR.N |
| POM121 membrane glycoprotein-like | POM121C | S128 | 9.6 | 57.8 | R.SSS*MSSLTGAYTSGIPSSSR.N |
| POP1 | POP1 | S729 | 28.4 | 20.5 | R.VQAYEEPSVAS*SPNGK.E |
| Positive cofactor 2 glutamine | MED15 | S505T511 | 9.3 | 52.7 | R.TPQNFSVPS*PGPLNT*PVNPSSVMSPAGSSQAEEQQYLDK.L |
| Positive cofactor 2 glutamine | MED15 | S505S517 | 7.7 | 27.6 | R.TPQNFSVPS*PGPLNTPVNPSS*VMSPAGSSQAEEQQYLDK.L |
| Potassium channel tetramerisation domain containing 15 | KCTD15 | S35 | 24.0 | 62.8 | R.S*PVSPLAAQGIPLPAQLTK.S |
| Potassium channel tetramerisation domain containing 15 | KCTD15 | S38 | 0.3 | 14.3 | R.SPVS*PLAAQGIPLPAQLTK.S |
| Potassium channel tetramerisation domain containing 15 | KCTD15 | S35S38 | 43.6 | 12.8 | R.S*PVS*PLAAQGIPLPAQLTK.S |
| Potassium channel, calcium activated, intermediate/small conductance subfamily | KCNN3 | S128 | 57.2 | 16.8 | R.QGS*QLNLNDHLLGHSPSSTATSGPGGGSR.H |
| Potassium channel, voltage gated, KQT like subfamily, member 4 | KCNQ5 | S484 | 17.3 | 18.0 | K.SWS*FNDR.T |
| Potassium channel, voltage gated, KQT like subfamily, member 4 | KCNQ5 | S482 | 12.2 | 17.5 | K.S*WSFNDR.T |
| Potassium voltage gated channel, shaker related subfamily, beta member 1 | KCNAB1 | S153 | 7.6 | 16.0 | R.RSS*LVITTK.I |
| Potassium voltage gated channel, shaker related subfamily, beta member 1 | KCNAB1 | S152 | 39.1 | 11.6 | R.RS*SLVITTK.I |
| Potassium voltage-gated channel, KQT-like subfamily, member 5 | KCNQ5 | S831 | 2.7 | 62.8 | K.SLS*VQNLIR.S |
| Potassium voltage-gated channel, KQT-like subfamily, member 5 | KCNQ5 | S829 | 9.3 | 51.0 | K.S*LSVQNLIR.S |
| Pre mRNA cleavage complex II protein Pcf11 | PCF11 | S705 | 100.0 | 52.9 | R.S*PFNDRFPLK.R |
| Pre mRNA cleavage complex II protein Pcf11 | PCF11 | S489S494 | 29.1 | 16.6 | K.SRS*PIIHS*PK.R |
| Pre mRNA splicing factor PRP17 | CDC40 | S46 | 15.9 | 100.2 | K.SPSS*KPSLAVAVDSAPEVAVK.E |
| Pre mRNA splicing factor PRP17 | CDC40 | S43 | 8.4 | 50.2 | K.S*PSSKPSLAVAVDSAPEVAVK.E |
| Pre mRNA splicing factor PRP17 | CDC40 | S45 | 11.5 | 68.8 | K.SPS*SKPSLAVAVDSAPEVAVK.E |
| Pre mRNA splicing factor PRP17 | CDC40 | S49 | 8.0 | 35.0 | K.SPSSKPS*LAVAVDSAPEVAVK.E |
| Pre-B-cell leukemia transcription factor interacting protein 1 | PBXIP1 | S146S147S148 | 100.0 | 13.5 | R.EEGRCS*S*S*DDDT*DVDMEGLRR.R |
| PREDICTED: lateral signaling target protein 2 homolog isoform X5 [Homo sapiens] | | S921 | -0.9 | 16.0 | K.SS*ARAAPR.S |
| PREDICTED: paired box protein Pax-2 isoform X8 [Homo sapiens] | | S337 | | 19.1 | K.RS*YPR.N |
| Prefoldin 4 | PFDN4 | S125 | 16.4 | 36.0 | K.FGS*NINLEADES.- |
| PRO1770 | PELO | S374S380S381 | 100.0 | 44.0 | R.FPVPELS*DQEGDS*S*S*EED.- |
| Probable ATP-dependent RNA helicase DDX27 | DDX27 | S23S25 | 26.3 | 69.5 | M.LADLGLIGTIGEDDEVPVEPES*DS*GDEEEEGPIVLGR.R |
| Processing of precursor 5, ribonuclease P/MRP subunit | POP5 | S154 | 108.1 | 112.0 | R.SCLLEEEEES*GEEAAEAME.- |
| Profilin 1 | PFN1 | T93 | -0.3 | 72.7 | K.ST*GGAPTFNVTVTK.T |
| Progesterone receptor membrane component 1 | PGRMC1 | S181 | 19.8 | 74.7 | K.EGEEPTVYS*DEEEPKDESAR.K |
| Progesterone receptor membrane component 1 | PGRMC1 | Y180 | 5.8 | 42.5 | K.EGEEPTVY*SDEEEPKDESAR.K |
| Progesterone receptor membrane component 1 | PGRMC1 | T178 | | 22.2 | K.EGEEPT*VYSDEEEPK.D |
| Progesterone receptor membrane component 2 | PGRMC2 | S208 | 15.7 | 43.0 | R.LLKPGEEPS*EYTDEEDTKDHNKQD.- |
| Progesterone receptor membrane component 2 | PGRMC2 | Y210 | 12.2 | 55.5 | R.LLKPGEEPSEY*TDEEDTKDHNKQD.- |
| Programmed cell death 4 | PDCD4 | S94 | 8.2 | 44.5 | R.SGLTVPTS*PK.G |
| Programmed cell death 4 | PDCD4 | T93 | 32.3 | 44.2 | R.SGLTVPT*SPK.G |
| Programmed cell death 4 | PDCD4 | S457 | 100.0 | 23.6 | R.FVS*EGDGGR.L |
| Programmed cell death 5 | PDCD5 | S119 | 63.8 | 53.0 | R.KVM#DS*DEDDDY.- |
| Programmed cell death 5 | PDCD5 | Y125 | 21.6 | 67.3 | R.RKVMDSDEDDDY*.- |
| Programmed cell death 8 | AIFM1 | S266 | 10.5 | 42.4 | R.S*LSAIDR.A |
| Programmed cell death 8 | AIFM1 | S268 | 15.1 | 32.6 | R.SLS*AIDR.A |
| Proline glutamic acid leucine-rich protein 1 | PELP1 | S481 | 35.4 | 70.2 | R.GS*PDGSLQTGKPSAPK.K |
| Proline-rich transmembrane protein 3 | PRRT3 | S924S928 | 15.9 | 35.3 | R.HGLS*SVDS*LPLDELPSTVQLLPAPTPAPDSTAAR.Q |
| Proline-rich transmembrane protein 3 | PRRT3 | S798S808 | 21.1 | 18.6 | R.NGVGPAPS*LSELDLRPPS*PINLSR.S |
| Proline-rich transmembrane protein 3 | PRRT3 | S925S928 | 5.1 | 38.9 | R.HGLSS*VDS*LPLDELPSTVQLLPAPTPAPDSTAAR.Q |
| Proline-rich transmembrane protein 3 | PRRT3 | S924S925 | 7.7 | 25.6 | R.HGLS*S*VDSLPLDELPSTVQLLPAPTPAPDSTAAR.Q |
| Proline-rich transmembrane protein 3 | PRRT3 | S798S800S808 | 38.3 | 13.8 | R.NGVGPAPS*LS*ELDLRPPS*PINLSR.S |
| Proline-rich transmembrane protein 3 | PRRT3 | S815 | 100.0 | 37.7 | R.S*IDAALFR.E |
| Proline/serine-rich coiled-coil 1 | PSRC1 | S65S70 | 100.0 | 17.6 | R.LS*LGPLS*PEKLEEILDEANR.L |
| Proteasome 26S subunit ATPase 4 | PSMC4 | S19S21T25 | | 28.1 | K.AQDEIPALS*VS*RPQT*GLSFLGPEPEDLEDLYSR.Y |

Peak Area: <-10, -5, -3, 0, 3, 5, >10

%CV: 0, 29, 42, 71, 86, >100

**White dots-** Significant change in peptide abundance at **5** %FDR compared to the timepoint with the minimum peak area for a given PSM

| Protein Name | Gene | Phosphosites | CarT (0m 2m 5m) | RajiB (0m 2m 5m) | Ascor | MOWSE | Sequence |
|---|---|---|---|---|---|---|---|
| **Proteasome 26S subunit ATPase 4** | PSMC4 | S19T25S28 | | | **29.6** | **37.2** | K.AQDEIPALS*VSRPQT*GLS*FLGPEPEDLEDLYSR.Y |
| **Proteasome 26S subunit ATPase 4** | PSMC4 | S21T25S28 | | | **13.6** | **52.8** | K.AQDEIPALSVS*RPQT*GLS*FLGPEPEDLEDLYSR.Y |
| **Proteasome 26S subunit ATPase 4** | PSMC4 | S19S21 | | | **10.0** | **25.9** | K.AQDEIPALS*VS*RPQTGLSFLGPEPEDLEDLYSR.Y |
| **Proteasome 26S subunit ATPase 4** | PSMC4 | S19S28 | | | **4.7** | **15.7** | K.AQDEIPALS*VSRPQTGLS*FLGPEPEDLEDLYSR.Y |
| **Proteasome 26S subunit, non ATPase, 2** | PSMD2 | S16 | | | **92.8** | **68.3** | R.DKAPVQPQQS*PAAAPGGTDEKPSGK.E |
| **Proteasome subunit alpha type 2** | PSMA2 | S7 | | | **10.8** | **77.3** | R.GYS*FSLTTFSPSGK.L |
| | PSMA3 | S250 | | | **100.0** | **10.8** | K.EEDES*DDDNM.- |
| **Protein ABHD11 (Fragment)** | ABHD11 | S106 | | | **44.9** | **14.6** | K.TAM#LLALQRIS*AQWK.A |
| **Protein arginine N methyltransferase 3** | PRMT3 | S25S27 | | | **100.0** | **86.9** | R.GAVENEEDLPELS*DS*GDEAAWEDEDDADLPHGK.Q |
| **Protein disulfide isomerase P5** | PDIA6 | S428 | | | **100.0** | **80.0** | R.DGELPVEDDIDLS*DVELDDLGKDEL.- |
| **Protein FAM117B** | FAM117B | S220 | | | **24.4** | **88.9** | R.TSS*LDTLAAPYLAGHWPR.D |
| **Protein FAM117B** | FAM117B | S219 | | | **-0.4** | **51.2** | R.TS*SLDTLAAPYLAGHWPR.D |
| **Protein FAM117B** | FAM117B | T149S152T158 | | | **25.1** | **62.2** | R.GSPPRPRPPPPPLLGT*VSS*PSSSPT*HLWTGEVSAAPPPAR.V |
| **Protein FAM117B** | FAM117B | T149S151T158 | | | **25.3** | **52.8** | R.GSPPRPRPPPPPLLGT*VS*SPSSSPT*HLWTGEVSAAPPPAR.V |
| **Protein FAM117B** | FAM117B | S136T149S156 | | | **11.9** | **17.6** | R.GS*PPRPRPPPPPLLGT*VSSPSSS*PTHLWTGEVSAAPPPAR.V |
| **Protein FAM117B** | FAM117B | S136T149S154 | | | **27.1** | **40.3** | R.GS*PPRPRPPPPPLLGT*VSSPS*SSPTHLWTGEVSAAPPPAR.V |
| **Protein FAM117B** | FAM117B | S136T149T158 | | | **29.2** | **26.0** | R.GS*PPRPRPPPPPLLGT*VSSPSSSPT*HLWTGEVSAAPPPAR.V |
| **Protein FAM117B** | FAM117B | S136S154T158 | | | **34.5** | **54.9** | R.GS*PPRPRPPPPPLLGTVSSPS*SSPT*HLWTGEVSAAPPPAR.V |
| **Protein FAM117B** | FAM117B | S136S151S156 | | | **27.4** | **40.6** | R.GS*PPRPRPPPPPLLGTVS*SPSSS*PTHLWTGEVSAAPPPAR.V |
| **Protein FAM117B** | FAM117B | S151S154 | | | **2.6** | **17.7** | R.GSPPRPRPPPPPLLGTVS*SPS*SSPTHLWTGEVSAAPPPAR.V |
| **Protein FAM117B** | FAM117B | S136S151S154 | | | **10.5** | **22.1** | R.GS*PPRPRPPPPPLLGTVS*SPS*SSPTHLWTGEVSAAPPPAR.V |
| **Protein FAM117B** | FAM117B | S136T149S151 | | | | **36.9** | R.GS*PPRPRPPPPPLLGT*VS*SPSSSPTHLWTGEVSAAPPPAR.V |
| **Protein FAM117B** | FAM117B | T223 | | | **7.4** | **44.0** | R.TSSLDT*LAAPYLAGHWPR.D |
| **Protein FAM117B** | FAM117B | S136S155T158 | | | **17.2** | **20.6** | R.GS*PPRPRPPPPPLLGTVSSPSS*SPT*HLWTGEVSAAPPPAR.V |
| **Protein FAM117B** | FAM117B | S136S154S156 | | | **18.1** | **18.9** | R.GS*PPRPRPPPPPLLGTVSSPS*SS*PTHLWTGEVSAAPPPAR.V |
| **Protein FAM117B** | FAM117B | S10 | | | **39.3** | **129.0** | R.NGS*PTPAGSLGGGAVATAGGPGSR.L |
| **Protein FAM117B** | FAM117B | S151S152T158 | | | **14.4** | **27.5** | R.GSPPRPRPPPPPLLGTVS*S*PSSSPT*HLWTGEVSAAPPPAR.V |
| **Protein FAM117B** | FAM117B | S136T158T162 | | | **27.6** | **39.9** | R.GS*PPRPRPPPPPLLGTVSSPSSSPT*HLWT*GEVSAAPPPAR.V |
| **Protein FAM189A1** | FAM189A1 | S18S20 | | | **100.0** | **22.2** | R.S*LS*RLR.E |
| **Protein HSPC148** | CWC15 | T110S121 | | | **45.3** | **78.3** | R.LDQIPAANLDADDPLT*DEEDEDFEEES*DDDDTAALLAELEK.I |
| **Protein inhibitor of activated STAT, 1** | PIAS1 | S510 | | | **4.5** | **46.6** | R.TPS*LPAVDTSYINTSLIQDYR.H |
| **Protein inhibitor of activated STAT, 1** | PIAS1 | T508 | | | **12.2** | **86.3** | R.T*PSLPAVDTSYINTSLIQDYR.H |
| **Protein inhibitor of activated STAT, 1** | PIAS1 | S503 | | | **18.8** | **66.7** | K.GILSLPHQAS*PVSR.T |
| **Protein inhibitor of activated STAT, 1** | PIAS1 | S517 | | | **-2.9** | **51.1** | R.TPSLPAVDTS*YINTSLIQDYR.H |
| **Protein kinase B, beta** | AKT2 | S447T449 | | | **14.8** | **15.5** | R.YFDDEFTAQS*IT*ITPPDRYDSLGLLELDQR.T |
| **Protein kinase C alpha** | PRKCA | S226 | | | **48.5** | **75.7** | R.STLNPQWNES*FTFK.L |
| **Protein kinase C alpha** | PRKCB | T497 | | | **19.5** | **92.0** | R.T*FCGTPDYIAPEIIAYQPYGK.S |
| **Protein kinase C alpha** | PRKCB | T501 | | | **1.8** | **39.4** | R.TFCGT*PDYIAPEIIAYQPYGK.S |
| **Protein kinase C and casein kinase substrate in neurons 2** | PACSIN2 | T321T325 | | | | **12.3** | K. [illegible] |
| **Protein kinase C binding protein 1** | ZMYND8 | S445 | | | **22.4** | **45.0** | R.RIS*LSDMPR.S |
| **Protein kinase C binding protein 1** | ZMYND8 | S426 | | | **35.4** | **60.2** | K.LNFDMTAS*PK.I |
| **Protein kinase C binding protein 1** | ZMYND8 | S510 | | | **20.7** | **65.8** | K.TGQAGSLSGS*PKPFSPQLSAPITTK.T |
| **Protein kinase C binding protein 1** | ZMYND8 | S508 | | | **14.9** | **58.1** | K.TGQAGSLS*GSPKPFSPQLSAPITTK.T |
| **Protein kinase C binding protein 1** | ZMYND8 | S506S515 | | | **9.2** | **24.4** | K.TGQAGS*LSGSPKPFS*PQLSAPITTK.T |
| **Protein kinase C binding protein 1** | ZMYND8 | S506S510 | | | **8.4** | **50.7** | K.TGQAGS*LSGS*PKPFSPQLSAPITTK.T |
| **Protein kinase C binding protein 1** | ZMYND8 | S510S515 | | | **18.1** | **35.6** | K.TGQAGSLSGS*PKPFS*PQLSAPITTK.T |
| **Protein kinase C binding protein 1** | ZMYND8 | S506 | | | **4.2** | **31.0** | K.TGQAGS*LSGSPKPFSPQLSAPITTK.T |
| **Protein kinase C binding protein 1** | ZMYND8 | S506S508 | | | **18.7** | **52.7** | K.TGQAGS*LS*GSPKPFSPQLSAPITTK.T |
| **Protein kinase C binding protein 1** | ZMYND8 | S515 | | | **20.4** | **30.8** | K.TGQAGSLSGSPKPFS*PQLSAPITTK.T |
| **Protein kinase C binding protein 1** | ZMYND8 | S508S515 | | | **13.6** | **24.8** | K.TGQAGSLS*GSPKPFS*PQLSAPITTK.T |
| **Protein kinase C delta** | PRKCD | S506 | | | **9.2** | **60.4** | R.AS*TFCGTPDYIAPEILQGLK.Y |
| **Protein kinase C delta** | PRKCD | T507 | | | **11.2** | **94.2** | R.AST*FCGTPDYIAPEILQGLK.Y |
| | PRKCD | S654 | | | **15.5** | **60.8** | R.LSYSDKNLIDS*MDQSAFAGFSFVNPK.F |
| **Protein kinase C delta** | PRKCD | Y646 | | | **5.3** | **73.8** | R.LSY*SDKNLIDSMDQSAFAGFSFVNPK.F |
| **Protein kinase C delta** | PRKCD | Y646S664 | | | **54.2** | **82.2** | R.LSY*SDKNLIDSMDQSAFAGFS*FVNPK.F |
| **Protein kinase C delta** | PRKCD | S645S664 | | | **33.4** | **79.9** | R.LS*YSDKNLIDSMDQSAFAGFS*FVNPK.F |
| **Protein kinase C delta** | PRKCD | S645 | | | **23.9** | **44.7** | R.LS*YSDK.N |
| **Protein kinase C delta** | PRKCD | T511 | | | **0.1** | **21.6** | R.ASTFCGT*PDYIAPEILQGLK.Y |
| **Protein kinase C delta** | PRKCD | S647 | | | **5.8** | **44.8** | R.LSYS*DKNLIDSMDQSAFAGFSFVNPK.F |
| **Protein kinase C delta** | PRKCD | S664 | | | **62.3** | **85.1** | K.NLIDSMDQSAFAGFS*FVNPK.F |
| **Protein kinase C delta** | PRKCD | S506T507 | | | **8.0** | **48.3** | R.AS*T*FCGTPDYIAPEILQGLK.Y |
| **Protein kinase C delta** | PRKCD | S654S664 | | | **25.7** | **69.6** | K.ARLSYSDKNLIDS*MDQSAFAGFS*FVNPK.F |

Peak Area: <-10, -5, -3, 0, 3, 5, >10

%CV: 0, 29, 42, 71, 86, >100

**White dots-** Significant change in peptide abundance at **5** %FDR compared to the timepoint with the minimum peak area for a given PSM

CarT (0m, 2m, 5m) — RajiB (0m, 2m, 5m)

| Protein Name | Gene | Phosphosites | Ascor | MOWSE | Sequence |
|---|---|---|---|---|---|
| **Protein kinase C delta** | PRKCD | S506Y514 | **6.7** | **27.6** | R.AS*TFCGTPDY*IAPEILQGLK.Y |
| **Protein kinase C delta** | PRKCD | S658S664 | **26.1** | **24.3** | R.LSYSDKNLIDSMDQS*AFAGFS*FVNPK.F |
| **Protein kinase C delta** | PRKCD | Y514 | **-0.6** | **17.7** | R.ASTFCGTPDY*IAPEILQGLK.Y |
| **Protein kinase C epsilon** | PRKCE | S729 | **21.2** | **59.1** | K.GFS*YFGEDLMP.- |
| **Protein kinase C like 2** | PKN2 | S583 | **10.2** | **76.3** | R.ASS*LGEIDESSELR.V |
| **Protein kinase C like 2** | PKN2 | T814 | **7.2** | **83.9** | R.T*STFCGTPEFLAPEVLTETSYTR.A |
| **Protein kinase C like 2** | PKN2 | T816 | **6.1** | **72.7** | R.TST*FCGTPEFLAPEVLTETSYTR.A |
| **Protein kinase C, beta 1** | PRKCB | S654 | **10.2** | **78.2** | R.NIDQS*EFEGFSFVNSEFLKPEVK.S |
| **Protein kinase C, beta 1** | PRKCB | S660 | **32.2** | **68.4** | R.NIDQSEFEGFS*FVNSEFLKPEVK.S |
| **Protein kinase C, beta 1** | PRKCB | S660S664 | **12.1** | **44.7** | R.NIDQSEFEGFS*FVNS*EFLKPEVK.S |
| **Protein kinase C, beta 1** | PRKCB | S654S660 | **40.0** | **64.0** | R.NIDQS*EFEGFS*FVNSEFLKPEVK.S |
| **Protein kinase C, beta 1** | PRKCB | S654S660S664 | **100.0** | **15.1** | R.NIDQS*EFEGFS*FVNS*EFLKPEVK.S |
| **Protein kinase C, beta 1** | PRKCB | S654S664 | **11.2** | **24.2** | R.NIDQS*EFEGFSFVNS*EFLKPEVK.S |
| **Protein kinase C, eta** | PRKCH | S28S32 | **100.0** | **20.5** | R.WS*LRHS*LFK.K |
| **Protein kinase C, eta** | PRKCH | S317 | **13.9** | **65.3** | K.TLAGMGLQPGNIS*PTSK.L |
| **Protein kinase C, eta** | PRKCH | S675 | **21.2** | **37.1** | R.NFS*YVSPELQP.- |
| **Protein kinase C, iota type** | PRKCI | T412 | **10.3** | **60.1** | K.EGLRPGDTTST*FCGTPNYIAPEILR.G |
| **Protein kinase C, iota type** | PRKCI | T416 | **-2.0** | **12.6** | K.EGLRPGDTTSTFCGT*PNYIAPEILR.G |
| **Protein kinase C, iota type** | PRKCI | S411T412 | **6.5** | **38.2** | K.EGLRPGDTTS*T*FCGTPNYIAPEILR.G |
| **Protein kinase C, mu** | PRKD1 | S742 | **43.6** | **53.5** | R.S*VVGTPAYLAPEVLR.S |
| **Protein kinase C, nu** | PRKD3 | S27S30 | **35.1** | **18.8** | K.SVLPTAIPAVLPAAS*PCS*SPK.T |
| **Protein kinase C, nu** | PRKD3 | S27S31 | **9.0** | **14.8** | K.SVLPTAIPAVLPAAS*PCSS*PK.T |
| **Protein kinase C, theta** | PRKCQ | S676 | **100.0** | **29.2** | R.LS*FADR.A |
| **Protein kinase C, theta** | PRKCQ | S695 | **100.0** | **51.5** | R.NFS*FMNPGMER.L |
| **Protein kinase C, theta** | PRKCQ | S685 | **100.0** | **69.0** | R.ALINS*MDQNMFR.N |
| **Protein kinase C, theta** | PRKCQ | T538 | **10.1** | **73.3** | K.TNT*FCGTPDYIAPEILLGQK.Y |
| **Protein kinase C, theta** | PRKCQ | S323 | **100.0** | **34.0** | R.EGPVEIGLPCS*IK.N |
| **Protein kinase C, theta** | PRKCQ | Y545 | **11.3** | **16.5** | K.TNTFCGTPDY*IAPEILLGQK.Y |
| **Protein kinase C, theta** | PRKCQ | T542 | **3.0** | **34.4** | K.TNTFCGT*PDYIAPEILLGQK.Y |
| **Protein kinase cAMP dependent, catalytic beta** | KIN27 | T245 | **19.9** | **95.7** | R.TWT*LCGTPEYLAPEIILSK.G |
| **Protein kinase D2** | PRKD2 | S710 | **19.6** | **105.5** | R.S*VVGTPAYLAPEVLLNQGYNR.S |
| **Protein kinase D2** | PRKD2 | S396 | **10.9** | **19.1** | R.KSS*TTLR.E |
| **Protein kinase D2** | PRKD2 | S197 | **31.1** | **53.5** | R.RLS*STSLASGHSVR.L |
| **Protein kinase D2** | PRKD2 | S375 | **15.0** | **38.7** | K.AQSS*LGYIPLMR.V |
| **Protein kinase D2** | PRKD2 | S200 | **3.0** | **27.9** | R.RLSSTS*LASGHSVR.L |
| **Protein kinase serine/arginine specific 2** | SRPK2 | S505T509 | **18.4** | **36.5** | R.TVS*ASST*GDLPK.A |
| **Protein kinase serine/arginine specific 2** | SRPK2 | T52 | | **32.1** | K.<br>APLVPPPPPPPPPPPPLPDPT*PPEPEEEILGSDDEEQEDPADYCK |
| **Protein kinase serine/arginine specific 2** | SRPK2 | S391 | **24.2** | **60.6** | K.DEDDVDQELANIDPTWIES*PK.T |
| **Protein kinase serine/arginine specific 2** | SRPK2 | S505S507 | **6.3** | **18.4** | R.TVS*AS*STGDLPK.A |
| **Protein kinase serine/arginine specific 2** | SRPK2 | T503S508 | **8.1** | **41.7** | R.T*VSASS*TGDLPK.A |
| **Protein kinase serine/arginine specific 2** | SRPK2 | S505S508 | **12.0** | **45.7** | R.TVS*ASS*TGDLPK.A |
| **Protein kinase serine/arginine specific 2** | SRPK2 | S507T509 | **8.3** | **22.5** | R.TVSAS*ST*GDLPK.A |
| **Protein kinase substrate MK2S4** | RCSD1 | S268 | **11.9** | **70.9** | R.SS*EEVDGQHPAQEEVPESPQTSGPEAENR.C |
| **Protein kinase substrate MK2S4** | RCSD1 | S216 | **56.2** | **95.4** | K.APGS*PLSSEGAAGEGVR.T |
| **Protein kinase substrate MK2S4** | RCSD1 | S179 | **35.1** | **85.6** | R.SQS*DCGELGDFR.A |
| **Protein kinase substrate MK2S4** | RCSD1 | S120S127 | **11.6** | **28.6** | K.AMVSPFHS*PPSTPSS*PGVR.S |
| **Protein kinase substrate MK2S4** | RCSD1 | S83 | **-0.5** | **21.2** | K.SS*PLIEK.L |
| **Protein kinase substrate MK2S4** | RCSD1 | S120 | **14.4** | **20.8** | K.AMVSPFHS*PPSTPSSPGVR.S |
| **Protein kinase substrate MK2S4** | RCSD1 | S116S120 | **43.5** | **45.3** | K.AM#VS*PFHS*PPSTPSSPGVR.S |
| **Protein kinase substrate MK2S4** | RCSD1 | S284 | **40.6** | **65.5** | R.SSEEVDGQHPAQEEVPES*PQTSGPEAENR.C |
| **Protein kinase substrate MK2S4** | RCSD1 | S116S126 | **17.2** | **38.7** | K.AMVS*PFHSPPSTPS*SPGVR.S |
| **Protein kinase substrate MK2S4** | RCSD1 | S177S179 | **100.0** | **60.3** | R.S*QS*DCGELGDFR.A |
| **Protein kinase substrate MK2S4** | RCSD1 | S116S120T124 | **7.9** | **20.8** | K.AMVS*PFHS*PPST*PS*SPGVR.S |
| **Protein kinase substrate MK2S4** | RCSD1 | S120S123 | **34.0** | **25.5** | K.AMVSPFHS*PPS*TPSSPGVR.S |
| **Protein kinase substrate MK2S4** | RCSD1 | S116S120S123 | **10.3** | **17.2** | K.AMVS*PFHS*PPS*TPSS*PGVR.S |
| **Protein kinase substrate MK2S4** | RCSD1 | S268S284 | **1.3** | **37.4** | R.SS*EEVDGQHPAQEEVPES*PQTSGPEAENR.C |
| **Protein kinase substrate MK2S4** | RCSD1 | S120S126 | **15.1** | **39.5** | K.AMVSPFHS*PPSTPS*SPGVR.S |
| **Protein kinase substrate MK2S4** | RCSD1 | S116S123T124 | **4.6** | **11.7** | K.AMVS*PFHSPPS*T*PSS*PGVR.S |
| **Protein kinase substrate MK2S4** | RCSD1 | S116S120S123 | **16.3** | **17.9** | K.AMVS*PFHS*PPS*TPS*SPGVR.S |
| **Protein kinase substrate MK2S4** | RCSD1 | S177 | **11.0** | **10.8** | R.S*QSDCGELGDFR.A |
| **Protein kinase substrate MK2S4** | RCSD1 | S126 | **22.6** | **22.4** | K.AMVSPFHSPPSTPS*SPGVR.S |
| **Protein kinase substrate MK2S4** | RCSD1 | S116T124 | **5.5** | **22.1** | K.AMVS*PFHSPPST*PSSPGVR.S |

Peak Area: <-10, -5, -3, 0, 3, 5, >10

%CV: 0, 29, 42, 71, 86, >100

**White dots-** Significant change in peptide abundance at **5** %FDR compared to the timepoint with the minimum peak area for a given PSM

| Protein Name | Gene | Phosphosites | Ascor | MOWSE | Sequence |
|---|---|---|---|---|---|
| **Protein kinase substrate MK2S4** | RCSD1 | S116S127 | **14.2** | **30.1** | K.AMVS*PFHSPPSTPSS*PGVR.S |
| **Protein kinase, cAMP dependent, regulatory type II, alpha** | PRKAR2A | S99 | **40.7** | **55.9** | R.RVS*VCAETYNPDEEEEDTDPR.V |
| | PRKAR2A | S78S80 | **85.8** | **51.4** | R.VADAKGDS*ES*EEDEDLEVPVPSR.F |
| | PRKAR1A | S83 | **100.0** | **20.2** | R.EDEIS*PPPPNPVVK.G |
| **Protein kinase, cAMP dependent, regulatory, type II, beta** | PRKAR2B | S83S85 | **100.0** | **50.4** | K.GVNFAEEPMQS*DS*EDGEEEEAAPADAGAFNAPVINR.F |
| **Protein kinase, cAMP dependent, regulatory, type II, beta** | PRKAR2B | S114 | **49.8** | **73.6** | R.RAS*VCAEAYNPDEEEDDAESR.I |
| **Protein kinase, lysine deficient 1** | WNK1 | S2011 | **26.0** | **21.3** | K.KEKPELSEPSHLNGPS*SDPEAAFLSR.D |
| **Protein kinase, lysine deficient 1** | WNK1 | S2027S2029S2 | **49.9** | **24.4** | R.DVDDGS*GS*PHS*PHQLSSK.S |
| **Protein kinase, lysine deficient 1** | WNK1 | S1978 | **100.0** | **50.3** | K.EGPVAS*PPFMDLEQAVLPAVIPK.K |
| **Protein kinase, lysine deficient 1** | WNK1 | S2011S2012 | **6.8** | **34.4** | K.EKPELSEPSHLNGPS*S*DPEAAFLSR.D |
| **Protein kinase, lysine deficient 4** | WNK1 | S331 | **100.0** | **21.8** | K.RAS*FAK.S |
| **Protein phosphatase 1 regulatory subunit 10** | PPP1R10 | S587 | **2.2** | **41.8** | K.GPQGPGGGGINVQEILTS*IMGSPNSHPSEELLK.Q |
| **Protein phosphatase 1 regulatory subunit 10** | PPP1R10 | S591 | **4.4** | **37.1** | K.GPQGPGGGGINVQEILTSIMGS*PNSHPSEELLK.Q |
| **Protein phosphatase 1 regulatory subunit 7** | PPP1R7 | S27 | **18.2** | **33.4** | R.VESEES*GDEEGKKHSSGIVADLSEQSLK.D |
| **Protein phosphatase 1 regulatory subunit 7** | PPP1R7 | S24S36 | **9.8** | **66.4** | R.VES*EESGDEEGKKHS*SGIVADLSEQSLK.D |
| **Protein phosphatase 1 regulatory subunit 7** | PPP1R7 | S27S36 | **6.0** | **55.2** | R.VESEES*GDEEGKKHS*SGIVADLSEQSLK.D |
| **Protein phosphatase 1 regulatory subunit 7** | PPP1R7 | S24S27 | **100.0** | **70.2** | R.VES*EES*GDEEGKK.H |
| **Protein phosphatase 1 regulatory subunit 7** | PPP1R7 | S27S37 | **-0.2** | **20.8** | R.VESEES*GDEEGKKHSS*GIVADLSEQSLK.D |
| **Protein phosphatase 1, regulatory subunit 11** | PPP1R11 | S73S74T75S77 | **125.5** | **68.4** | R.AFGES*S*T*ES*DEEEEEGCGHTHCVR.G |
| **Protein phosphatase 1, regulatory subunit 9B** | PPP1R9B | S192 | **32.0** | **67.1** | R.FNGS*TEALDKLDADAVSPTVSQLSAVFEK.A |
| **Protein phosphatase 1, regulatory subunit 9B** | PPP1R9B | T193S205 | **17.1** | **28.3** | R.FNGST*EALDKLDADAVS*PTVSQLSAVFEK.A |
| **Protein phosphatase 1, regulatory subunit 9B** | PPP1R9B | S205 | **3.7** | **23.3** | R.FNGSTEALDKLDADAVS*PTVSQLSAVFEK.A |
| **Protein phosphatase 4, regulatory subunit 2** | PPP4R2 | S226 | **12.1** | **46.0** | K.NHSDSSTSESEVSSVS*PLK.N |
| **Protein phosphatase 4, regulatory subunit 2** | PPP4R2 | S223 | **11.2** | **16.9** | K.NHSDSSTSESEVS*SVSPLK.N |
| **Protein phosphatase inhibitor 2** | PPP1R2 | S121 | **55.5** | **151.5** | R.IQEQES*SGEEDSDLSPEER.E |
| **Protein phosphatase inhibitor 2** | PPP1R2 | S121S122 | **73.3** | **155.0** | R.IQEQES*S*GEEDSDLSPEER.E |
| **Protein phosphatase inhibitor 2** | PPP1R2 | Y75 | **2.3** | **54.0** | K.IDEPSTPY*HSMM#GDDEDACSDTEATEAMAPDILAR.K |
| | PPP1R2 | S87 | **22.0** | **99.3** | K.IDEPSTPYHSMMGDDEDACS*DTEATEAMAPDILAR.K |
| **Protein phosphatase inhibitor 2** | PPP1R2 | S72 | **15.8** | **17.6** | K.IDEPS*TPYHSMMGDDEDACSDTEATEAM#APDILAR.K |
| | PPP1R2 | T89 | **18.7** | **79.6** | K.IDEPSTPYHSMM#GDDEDACSDT*EATEAMAPDILAR.K |
| **Protein phosphatase inhibitor 2** | PPP1R2 | S77 | **-2.2** | **31.9** | K.IDEPSTPYHS*MMGDDEDACSDTEATEAMAPDILAR.K |
| **Protein phosphatase methylesterase 1** | PPME1 | S42 | **80.2** | **99.6** | R.DFS*PVPWSQYFESMEDVEVENETGKDTFR.V |
| **Protein phosphatase methylesterase 1** | PPME1 | S47 | **2.3** | **16.2** | R.DFSPVPWS*QYFESMEDVEVENETGKDTFR.V |
| **Protein phosphatase methylesterase 1** | PPME1 | S52 | **11.1** | **30.6** | R.DFSPVPWSQYFES*MEDVEVENETGK.D |
| **Protein phosphatase methylesterase 1** | PPME1 | Y49 | **2.3** | **14.0** | R.DFSPVPWSQY*FESMEDVEVENETGK.D |
| **Protein RUFY3 (Fragment)** | RUFY3 | S50S55 | | **24.9** | R. [illegible] |
| **Protein SET** | SET | S7 | **100.0** | **36.7** | K.RQS*PLPPQK.K |
| **Protein strawberry notch homolog 1** | SBNO1 | S754S755 | **52.6** | **84.3** | K.NMS*S*GDDDDFNPFLDESNEDDENDPWLIR.K |
| **Protein strawberry notch homolog 1** | SBNO1 | S754S768 | **4.8** | **22.7** | K.NMS*SGDDDDFNPFLDES*NEDDENDPWLIR.K |
| **Protein transport protein SEC61 beta subunit** | SEC61B | S17 | **7.4** | **41.9** | M.PGPTPSGTNVGSSGRS*PSK.A |
| **Protein tyrosine phosphatase nonreceptor 7** | PTPN7 | S83 | **100.0** | **13.4** | R.RGS*NVALMLDVR.S |
| **Protein tyrosine phosphatase nonreceptor 7** | PTPN7 | S398 | **52.7** | **20.9** | R.GGMIQTAEQYQFLHHTLALYAGQLPEEPS*P.- |
| **Protein tyrosine phosphatase nonreceptor type 12** | PTPN12 | S435 | **100.0** | **22.9** | R.NLS*FEIK.K |
| **Protein tyrosine phosphatase receptor type C** | PTPRC | S992 | **100.0** | **66.9** | R.VPLKHELEMS*K.E |
| **Protein tyrosine phosphatase receptor type C** | PTPRC | S1297 | **16.8** | **23.5** | K.EQAEGSEPTSGTEGPEHSVNGPAS*PALNQGS.- |
| **Protein tyrosine phosphatase receptor type C** | PTPRC | S973 | **95.6** | **55.3** | R.NS*NVIPYDYNR.V |
| **Protein tyrosine phosphatase receptor type C** | PTPRC | S1291 | **7.4** | **11.4** | K.EQAEGSEPTSGTEGPEHS*VNGPASPALNQGS.- |
| **Protein tyrosine phosphatase receptor type C** | PTPRC | S992S995S999 | | **26.9** | K.HELEMS*KES*EHDS*DES*S*DDDS*DSEEPSK.Y |
| **Protein tyrosine phosphatase receptor type C** | PTPRC | S999S1002S10 | **16.2** | **26.3** | K.ESEHDS*DES*S*DDDS*DSEEPSK.Y |
| **Protein tyrosine phosphatase, non receptor type 2** | PTPN2 | S304 | **39.1** | **54.2** | K.EDLSPAFDHS*PNK.I |
| **PRP38 pre-mRNA processing factor 38 domain containing B** | PRPF38B | S266S268 | **100.0** | **16.0** | R.RS*LS*PR.R |
| **PRP38 pre-mRNA processing factor 38 domain containing B** | PRPF38B | S268 | **39.7** | **31.1** | R.RSLS*PR.R |
| | PRPF38B | S527S529 | **73.3** | **60.9** | R.S*QS*IEQESQEK.Q |
| **PRP38 pre-mRNA processing factor 38 domain containing B** | PRPF38B | S266 | **17.8** | **28.5** | R.RS*LSPR.R |
| **PRP38 pre-mRNA processing factor 38 domain containing B** | PRPF38B | S318S320 | **100.0** | **21.9** | R.S*RS*IDR.G |
| **PRP38 pre-mRNA processing factor 38 domain containing B** | PRPF38B | S529 | **19.9** | **47.9** | R.SQS*IEQESQEK.Q |
| **PRP38 pre-mRNA processing factor 38 domain containing B** | PRPF38B | S527 | **9.2** | **49.8** | R.S*QSIEQESQEK.Q |
| **PRP4 pre-mRNA processing factor 4 homolog B** | PRPF4B | S431S437 | **21.0** | **21.1** | K.DAS*PINRWS*PTR.R |
| **PRP4 pre-mRNA processing factor 4 homolog B** | PRPF4B | Y849 | **52.3** | **76.6** | K.LCDFGSASHVADNDITPY*LVSR.F |
| **PRP4 pre-mRNA processing factor 4 homolog B** | PRPF4B | T847 | **9.3** | **68.5** | K.LCDFGSASHVADNDIT*PYLVSR.F |
| **PRP4 pre-mRNA processing factor 4 homolog B** | PRPF4B | S518S519S520 | **100.0** | **51.5** | K.VEQES*S*S*DDNLEDFDVEEEDEEALIEQR.R |
| **PRP4 pre-mRNA processing factor 4 homolog B** | PRPF4B | S257 | **11.1** | **11.4** | K.ARS*PTDDKVK.I |

Peak Area: <-10, -5, -3, 0, 3, 5, >10

%CV: 0, 29, 42, 71, 86, >100

**White dots-** Significant change in peptide abundance at **5** %FDR compared to the timepoint with the minimum peak area for a given PSM

CarT: 0m, 2m, 5m — RajiB: 0m, 2m, 5m

| Protein Name | Gene | Phosphosites | Ascor | MOWSE | Sequence |
|---|---|---|---|---|---|
| PRP4 pre-mRNA processing factor 4 homolog B | PRPF4B | S354S356 | 100.0 | 38.0 | R.S*LS*PKPR.D |
| PRP4 pre-mRNA processing factor 4 homolog B | PRPF4B | S277 | 91.7 | 54.5 | R.KKS*PIINESR.S |
| PRP4 pre-mRNA processing factor 4 homolog B | PRPF4B | S294 | 100.0 | 35.0 | R.S*PVDLR.G |
| PRP4 pre-mRNA processing factor 4 homolog B | PRPF4B | S292S294 | 100.0 | 26.1 | K.S*RS*PVDLR.G |
| PRP4 pre-mRNA processing factor 4 homolog B | PRPF4B | T576S578S580 | 100.0 | 27.1 | R.T*RS*PS*PDDILER.V |
| PRP4 pre-mRNA processing factor 4 homolog B | PRPF4B | S239S241 | 29.0 | 24.7 | K.S*KS*PTLR.R |
| PRP4 pre-mRNA processing factor 4 homolog B | PRPF4B | S427S431S437 | 18.5 | 14.4 | R.S*KDAS*PINRWS*PTR.R |
| PRP4 pre-mRNA processing factor 4 homolog B | PRPF4B | S636 | 5.4 | 24.8 | K.LLAPDMFTES*DDMFAAYFDSAR.L |
| PRP4 pre-mRNA processing factor 4 homolog B | PRPF4B | S20S23S32 | 47.4 | 19.7 | R.EQPEMEDANS*EKS*INEENGEVS*EDQSQNK.H |
| PRP4 pre-mRNA processing factor 4 homolog B | PRPF4B | S427S431T439 | 14.6 | 13.9 | R.S*KDAS*PINRWSPT*R.R |
| PRP4 pre-mRNA processing factor 4 homolog B | PRPF4B | S427S431 | 100.0 | 27.7 | R.S*KDAS*PINR.W |
| PRP4 pre-mRNA processing factor 4 homolog B | PRPF4B | S131Y140 | 6.4 | 13.1 | K.VQS*GMGLILQGY*ESGSEEEGEIHEK.A |
| PRP4 pre-mRNA processing factor 4 homolog B | PRPF4B | S131S144 | 5.3 | 14.2 | K.VQS*GMGLILQGYESGS*EEEGEIHEK.A |
| PRP4 pre-mRNA processing factor 4 homolog B | PRPF4B | Y140S142 | 4.8 | 27.0 | K.VQSGMGLILQGY*ES*GSEEEGEIHEK.A |
| PRP4 pre-mRNA processing factor 4 homolog B | PRPF4B | T259 | 16.6 | 12.3 | R.SPT*DDKVK.I |
| PRP4 pre-mRNA processing factor 4 homolog B | PRPF4B | S431 | 100.0 | 18.9 | K.DAS*PINR.W |
| | PRPF4B | S87S93 | 100.0 | 38.6 | K.EIIDAS*DKEGMS*PAKR.T |
| PRP4 pre-mRNA processing factor 4 homolog B | PRPF4B | S578S580 | 31.8 | 28.3 | R.TRS*PS*PDDILER.V |
| PRP4 pre-mRNA processing factor 4 homolog B | PRPF4B | S20S23 | 34.5 | 21.5 | R.EQPEMEDANS*EKS*INEENGEVSEDQSQNK.H |
| PRP4 pre-mRNA processing factor 4 homolog B | PRPF4B | Y140S144 | 4.5 | 21.8 | K.VQSGM#GLILQGY*ESGS*EEEGEIHEK.A |
| PRP4 pre-mRNA processing factor 4 homolog B | PRPF4B | T634 | 45.0 | 44.5 | K.LLAPDMFT*ESDDMFAAYFDSAR.L |
| PRP4 pre-mRNA processing factor 4 homolog B | PRPF4B | S142S144 | 9.4 | 11.3 | K.VQSGMGLILQGYES*GS*EEEGEIHEK.A |
| | PRPF40A | T190S191S193 | 35.1 | 83.3 | K.DSGNWDT*S*GS*ELS*EGELEK.R |
| PRP40 pre-mRNA processing factor 40 homolog A | PRPF40A | S141S143S146 | 100.0 | 26.0 | R.HKS*DS*PES*DAEREK.D |
| PRP40 pre-mRNA processing factor 40 homolog A | PRPF40A | S143S146 | 5.7 | 13.9 | R.HKSDS*PES*DAER.E |
| PRPF31 | PRPF31 | T455 | 17.6 | 67.0 | R.SSGTASSVAFT*PLQGLEIVNPQAAEK.K |
| PRPF31 | PRPF31 | T448T455 | 12.2 | 92.9 | R.SSGT*ASSVAFT*PLQGLEIVNPQAAEK.K |
| PRPF31 | PRPF31 | T448 | -0.5 | 24.8 | R.SSGT*ASSVAFTPLQGLEIVNPQAAEKK.V |
| PRPF31 | PRPF31 | S450T455 | 14.6 | 77.7 | R.SSGTAS*SVAFT*PLQGLEIVNPQAAEK.K |
| PRPF31 | PRPF31 | S445T448 | 4.9 | 37.4 | R.S*SGT*ASSVAFTPLQGLEIVNPQAAEK.K |
| PRPF31 | PRPF31 | S450 | 7.3 | 77.6 | R.SSGTAS*SVAFTPLQGLEIVNPQAAEKK.V |
| PRPF31 | PRPF31 | S445T455 | 3.6 | 40.1 | R.S*SGTASSVAFT*PLQGLEIVNPQAAEK.K |
| PRPF31 | PRPF31 | S451T455 | 11.9 | 51.1 | R.SSGTASS*VAFT*PLQGLEIVNPQAAEK.K |
| PRPF31 | PRPF31 | S450S451 | 11.5 | 65.9 | R.SSGTAS*S*VAFTPLQGLEIVNPQAAEK.K |
| PRPF31 | PRPF31 | S446T455 | -0.7 | 27.6 | R.SS*GTASSVAFT*PLQGLEIVNPQAAEK.K |
| PRPF31 | PRPF31 | T448S450 | 3.1 | 45.7 | R.SSGT*AS*SVAFTPLQGLEIVNPQAAEK.K |
| PRPF31 | PRPF31 | T448S451T455 | -2.3 | 12.0 | R.SSGT*ASS*VAFT*PLQGLEIVNPQAAEK.K |
| PRPF31 | PRPF31 | S445S446T448 | -6.1 | 12.7 | R.S*S*GT*ASSVAFTPLQGLEIVNPQAAEK.K |
| PRPF31 | PRPF31 | T448S450S451 | 5.0 | 20.6 | R.SSGT*AS*S*VAFTPLQGLEIVNPQAAEK.K |
| PRPF31 | PRPF31 | S445S446 | 3.4 | 55.4 | R.S*S*GTASSVAFTPLQGLEIVNPQAAEK.K |
| PRPP synthetase associated protein 2 | PRPSAP2 | S227 | 16.4 | 61.6 | R.LGIAVIHGEAQDAESDLVDGRHS*PPMVR.S |
| PRPP synthetase associated protein 2 | PRPSAP2 | S219 | 4.6 | 15.7 | R.LGIAVIHGEAQDAES*DLVDGRHSPPMVR.S |
| PSD3 | PSD3 | S337 | 23.9 | 23.8 | K.ES*SKVPR.H |
| PSD3 | PSD3 | S338 | 8.9 | 22.7 | K.ESS*KVPR.H |
| Pseudoautosomal gene XE7 | AKAP17A | T639S640 | -0.2 | 13.1 | R.ST*S*PDHTR.S |
| Pseudouridine synthase 1 | PUS1 | T426 | 47.1 | 72.7 | K.VPSPLEGSEGDGDT*D.- |
| Pseudouridine synthase 1 | PUS1 | S420T426 | 22.7 | 46.7 | K.VPSPLEGS*EGDGDT*D.- |
| PSPC1 | PSPC1 | S409 | 100.0 | 58.0 | R.GAINMGDAFS*PAPAGNQGPPPMMGMNMNNR.A |
| PTK2B protein tyrosine kinase 2 beta | PTK2B | S375 | 100.0 | 58.6 | K.RNS*LPQIPMLNLEAR.R |
| PTK2B protein tyrosine kinase 2 beta | PTK2B | S839T842 | 10.2 | 54.5 | K.SLDPMVYMNDKS*PLT*PEKEVGYLEFTGPPQKPPR.L |
| PTK2B protein tyrosine kinase 2 beta | PTK2B | Y834T842 | 20.1 | 60.8 | K.SLDPMVY*MNDKSPLT*PEKEVGYLEFTGPPQKPPR.L |
| PTK2B protein tyrosine kinase 2 beta | PTK2B | S746T765 | 18.7 | 18.7 | K.LQFQVPEGLCAS*SPTLTSPMEYPSPVNSLHT*PPLHR.H |
| PTK2B protein tyrosine kinase 2 beta | PTK2B | S778 | 100.0 | 18.8 | K.RHS*MR.E |
| PTK2B protein tyrosine kinase 2 beta | PTK2B | S746S762 | 10.0 | 12.1 | K.LQFQVPEGLCAS*SPTLTSPM#EYPSPVNS*LHTPPLHR.H |
| PTK2B protein tyrosine kinase 2 beta | PTK2B | S746 | 13.6 | 42.2 | K.LQFQVPEGLCAS*SPTLTSPMEYPSPVNSLHTPPLHR.H |
| PTPRC-associated protein | PTPRCAP | S99 | 22.5 | 91.0 | R.AELGS*TDNDLER.Q |
| PTPRC-associated protein | PTPRCAP | S153 | | 57.8 | R. AEEARDS*DTEGDLVLGSPGPASAGGSAEALLSDLHAFAGSAAWDD |
| PTPRC-associated protein | PTPRCAP | S153T155 | | 58.7 | R. AEEARDS*DT*EGDLVLGSPGPASAGGSAEALLSDLHAFAGSAAWD |
| PTPRC-associated protein | PTPRCAP | S163S172 | 6.1 | 39.9 | R. DSDTEGDLVLGS*PGPASAGGS*AEALLSDLHAFAGSAAWDDSAR.A |
| PTPRC-associated protein | PTPRCAP | T155 | -0.3 | 90.4 | R.DSDT*EGDLVLGSPGPASAGGSAEALLSDLHAFAGSAAWDDSAR. A |
| PTPRC-associated protein | PTPRCAP | S153T155S163 | | 42.7 | R. AEEARDS*DT*EGDLVLGS*PGPASAGGSAEALLSDLHAFAGSAAWD |
| PTPRC-associated protein | PTPRCAP | T155S172 | 1.4 | 28.8 | R. DSDT*EGDLVLGSPGPASAGGS*AEALLSDLHAFAGSAAWDDSAR.A |

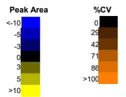


**White dots-** Significant change in peptide abundance at **5** %FDR compared to the timepoint with the minimum peak area for a given PSM

| Protein Name | Gene | Phosphosites | CarT 0m | CarT 2m | CarT 5m | RajiB 0m | RajiB 2m | RajiB 5m | Ascor | MOWSE | Sequence |
|---|---|---|---|---|---|---|---|---|---|---|---|
| **PTPRC-associated protein** | PTPRCAP | T155S168 | | | | | | | **2.9** | **88.7** | R.DSDT*EGDLVLGSPGPAS*AGGSAEALLSDLHAFAGSAAWDDSAR.A |
| **PTPRC-associated protein** | PTPRCAP | S163 | | | | | | | **45.8** | **80.4** | R.DSDTEGDLVLGS*PGPASAGGSAEALLSDLHAFAGSAAWDDSAR.A |
| **PTPRC-associated protein** | PTPRCAP | S163S168 | | | | | | | **2.9** | **33.3** | R.DSDTEGDLVLGS*PGPAS*AGGSAEALLSDLHAFAGSAAWDDSAR.A |
| **PTPRF interacting protein alpha 1** | PPFIA1 | S239S242 | | | | | | | **1.1** | **37.7** | R.SS*DGS*LSHEEDLAK.V |
| **PTPRF interacting protein alpha 1** | PPFIA1 | S238S239 | | | | | | | **9.3** | **15.4** | R.S*S*DGSLSHEEDLAK.V |
| **PTPRF interacting protein binding protein 1** | PPFIBP1 | S448 | | | | | | | **-0.6** | **25.3** | K.SS*SLGNLKK.E |
| **PTPRF interacting protein binding protein 1** | PPFIBP1 | S346S347 | | | | | | | **7.6** | **45.9** | K.GKDGEYEELLNSSSIS*S*LLDAQGFSDLEK.S |
| **PTPRF interacting protein binding protein 1** | PPFIBP1 | S342S347 | | | | | | | **0.2** | **49.3** | K.GKDGEYEELLNS*SSISS*LLDAQGFSDLEK.S |
| **PTPRF interacting protein binding protein 1** | PPFIBP1 | S343S347 | | | | | | | **10.1** | **78.5** | K.DGEYEELLNSS*SISS*LLDAQGFSDLEK.S |
| **PTPRF interacting protein binding protein 1** | PPFIBP1 | S342S346 | | | | | | | **9.3** | **68.2** | K.GKDGEYEELLNS*SSIS*SLLDAQGFSDLEK.S |
| **PTPRF interacting protein binding protein 1** | PPFIBP1 | S342S343 | | | | | | | **-0.0** | **55.1** | K.GKDGEYEELLNS*S*SISSLLDAQGFSDLEK.S |
| **PTPRF interacting protein binding protein 1** | PPFIBP1 | S344S347 | | | | | | | **4.5** | **17.9** | K.GKDGEYEELLNSSS*ISS*LLDAQGFSDLEK.S |
| **PTPRF interacting protein binding protein 1** | PPFIBP1 | S37S40 | | | | | | | **7.3** | **13.3** | K.ALEYSNGIFDCQS*PTS*PFMGSLR.A |
| **PTPRF interacting protein binding protein 1** | PPFIBP1 | S344S346 | | | | | | | **4.7** | **67.1** | K.DGEYEELLNSSS*IS*SLLDAQGFSDLEK.S |
| **PTPRF interacting protein binding protein 1** | PPFIBP1 | S343S344 | | | | | | | **10.7** | **68.2** | K.DGEYEELLNSS*S*ISSLLDAQGFSDLEK.S |
| **Pumilio homolog 1** | PUM1 | S709 | | | | | | | **12.8** | **67.1** | R.RDS*LTGSSDLYKR.T |
| **Pumilio homolog 1** | PUM1 | S228 | | | | | | | **8.0** | **30.3** | R.SESGGLGVSMVEYVLSS*SPGDSCLR.K |
| **Pumilio homolog 1** | PUM1 | S229 | | | | | | | **9.0** | **57.8** | R.SESGGLGVSMVEYVLSSS*PGDSCLR.K |
| **Pumilio homolog 1** | PUM1 | S220S228 | | | | | | | **5.8** | **21.6** | R.SESGGLGVS*MVEYVLSS*SPGDSCLR.K |
| **Pumilio homolog 1** | PUM1 | T112 | | | | | | | **100.0** | **27.5** | K.HRWPT*GDNIHAEHQVR.S |
| **Pumilio homolog 1** | PUM1 | S214S229 | | | | | | | **2.9** | **23.4** | R.SES*GGLGVSMVEYVLSSS*PGDSCLR.K |
| **Pumilio homolog 1** | PUM1 | S209 | | | | | | | | **55.6** | R.RPGQSFHVNSEVNSVLS*PR.S |
| **Pumilio homolog 1** | PUM1 | S227 | | | | | | | **10.3** | **21.9** | R.SESGGLGVSMVEYVLS*SSPGDSCLR.K |
| **Pumilio homolog 1** | PUM1 | S124 | | | | | | | **100.0** | **52.2** | R.S*MDELNHDFQALALEGR.A |
| **Pumilio2** | PUM2 | S82 | | | | | | | **59.9** | **32.3** | R.SGQGFHGNSEVNAILS*PR.S |
| **Pumilio2** | PUM2 | S136 | | | | | | | **100.0** | **50.6** | K.GKAS*PFEEDQNR.D |
| **PUR alpha** | PURA | S182 | | | | | | | **16.3** | **61.8** | R.GPGLGS*TQGQTIALPAQGLIEFR.D |
| **Purine rich element binding protein B** | PURB | S101 | | | | | | | **22.4** | **79.6** | R.DSLGDFIEHYAQLGPSS*PEQLAAGAEEGGGPR.R |
| **Purine rich element binding protein B** | PURB | S298S304 | | | | | | | **100.0** | **53.7** | R.RGGGS*GGGEES*EGEEVDED.- |
| | PURB | S304 | | | | | | | **25.8** | **60.7** | R.GGGSGGGEES*EGEEVDED.- |
| **Purine rich element binding protein B** | PURB | S100 | | | | | | | **6.9** | **57.0** | R.DSLGDFIEHYAQLGPS*SPEQLAAGAEEGGGPR.R |
| **Purine rich element binding protein B** | PURB | S86 | | | | | | | | **56.8** | R.DS*LGDFIEHYAQLGPSSPEQLAAGAEEGGGPR.R |
| **PWP2** | PWP2 | Y613S614S619 | | | | | | | **3.5** | **13.9** | K.GKAFTALCY*S*ADGHS*ILAGGMS*K.F |
| | PWP2 | S891S898S902 | | | | | | | **37.9** | **13.9** | R.SLDPLGS*EEEAEAS*EDDS*LHLLGGGGRDS*EEEMLA.- |
| **Pyruvate dehydrogenase complex, E1-alpha polypeptide 1** | PDHA1 | S232 | | | | | | | **8.9** | **16.2** | R.YGMGTS*VER.A |
| **Pyruvate dehydrogenase E1 alpha subunit, testis specific form** | PDHA1 | S291 | | | | | | | **10.9** | **16.2** | R.YHGHS*MSDPGVSYR.T |
| **Pyruvate dehydrogenase E1 alpha subunit, testis specific form** | PDHA1 | S291S298 | | | | | | | **20.5** | **51.3** | R.YHGHS*MSDPGVS*YR.T |
| | PDHA1 | S293S298 | | | | | | | **3.8** | **28.1** | R.YHGHSM#S*DPGVS*YR.T |
| **Pyruvate dehydrogenase E1 alpha subunit, testis specific form** | PDHA1 | S291Y299 | | | | | | | **9.1** | **28.3** | R.YHGHS*MSDPGVSY*R.T |
| **Pyruvate dehydrogenase E1 alpha subunit, testis specific form** | PDHA1 | S298 | | | | | | | **16.4** | **25.1** | R.YHGHSMSDPGVS*YR.T |
| **Pyruvate dehydrogenase E1 alpha subunit, testis specific form** | PDHA1 | S293 | | | | | | | **-1.0** | **38.5** | R.YHGHSMS*DPGVSYR.T |
| **R Ras2** | RRAS2 | S186 | | | | | | | **27.8** | **52.8** | K.FQEQECPPS*PEPTR.K |
| **R3H domain protein 1** | R3HDM1 | S381 | | | | | | | **-0.3** | **27.8** | K.ASS*FSGISVLTR.G |
| **R3H domain protein 1** | R3HDM2 | S141 | | | | | | | **10.5** | **38.3** | R.DS*SQEYTDSTGIDLHEFLVNTLK.N |
| **R3H domain protein 1** | R3HDM2 | S142 | | | | | | | **10.6** | **82.0** | R.DSS*QEYTDSTGIDLHEFLVNTLK.N |
| **R3H domain protein 1** | R3HDM1 | S380 | | | | | | | **26.4** | **44.7** | K.AS*SFSGISVLTR.G |
| **RA regulated nuclear matrix associated protein** | DTL | S655 | | | | | | | **5.8** | **52.5** | K.ENS*SPENKNWLLAMAAK.R |
| **RA regulated nuclear matrix associated protein** | DTL | S623S626 | | | | | | | **87.7** | **38.9** | K.IEGAGTSISEPPS*PIS*PYASESCGTLPLPLRPCGEGSEMVGK.E |
| **RA regulated nuclear matrix associated protein** | DTL | T616S617 | | | | | | | | **21.4** | K.IEGAGT*S*ISEPPSPISPYASESCGTLPLPLRPCGEGSEMVGK.E |
| **RA regulated nuclear matrix associated protein** | DTL | S619S623S626 | | | | | | | **21.5** | **41.5** | K.IEGAGTSIS*EPPS*PIS*PYASESCGTLPLPLRPCGEGSEMVGK.E |
| **RA regulated nuclear matrix associated protein** | DTL | T616S617S619 | | | | | | | | **46.7** | K.IEGAGT*S*IS*EPPSPISPYASESCGTLPLPLRPCGEGSEMVGK.E |
| **RA regulated nuclear matrix associated protein** | DTL | S656 | | | | | | | **-0.3** | **82.4** | K.ENSS*PENKNWLLAMAAK.R |
| **RA regulated nuclear matrix associated protein** | DTL | S485S487 | | | | | | | **8.1** | **25.2** | R.RGS*VS*SVSPKPPSSFK.M |
| **RA regulated nuclear matrix associated protein** | DTL | S679 | | | | | | | **12.2** | **59.2** | R.S*PSSQTPNSR.R |
| **RA regulated nuclear matrix associated protein** | DTL | S443 | | | | | | | **10.7** | **35.5** | K.CNPSNS*SPSSAACAPSCAGDLPLPSNTPTFSIK.T |
| **RA regulated nuclear matrix associated protein** | DTL | S441 | | | | | | | **9.1** | **16.4** | K.CNPS*NSSPSSAACAPSCAGDLPLPSNTPTFSIK.T |
| **RAB GTPase activating protein 1 like** | RABGAP1 | S464 | | | | | | | | **71.6** | R.ES*DKEEPVTPTSGGGPMSPQDDEAEEESDNELSSGTGDVSK.D |
| **RAB GTPase activating protein 1 like** | RABGAP1 | S490 | | | | | | | **4.5** | **43.0** | R.ESDKEEPVTPTSGGGPMSPQDDEAEEES*DNELSSGTGDVSK.D |
| **RAB GTPase activating protein 1 like** | RABGAP1 | S480 | | | | | | | **-4.8** | **34.0** | R.ESDKEEPVTPTSGGGPMS*PQDDEAEEESDNELSSGTGDVSK.D |
| **Rab GTPase binding effector protein 2** | RABEP2 | S193 | | | | | | | **28.7** | **61.0** | R.HAPSLHGS*TELLPLSR.D |
| **Rab GTPase binding effector protein 2** | RABEP2 | S193T194 | | | | | | | **11.4** | **45.5** | R.HAPSLHGS*T*ELLPLSR.D |
| **Rab GTPase binding effector protein 2** | RABEP2 | T194 | | | | | | | **9.1** | **41.1** | R.HAPSLHGST*ELLPLSR.D |

Peak Area: <-10, -5, -3, 0, 3, 5, >10

%CV: 0, 29, 42, 71, 86, >100

**White dots-** Significant change in peptide abundance at **5** %FDR compared to the timepoint with the minimum peak area for a given PSM

| Protein Name | Gene | Phosphosites | CarT 0m | CarT 2m | CarT 5m | RajiB 0m | RajiB 2m | RajiB 5m | Ascor | MOWSE | Sequence |
|---|---|---|---|---|---|---|---|---|---|---|---|
| **Rab GTPase binding effector protein 2** | RABEP2 | S200 | | | | | | | **12.8** | **59.4** | R.HAPSLHGSTELLPLS*R.D |
| **Rab GTPase binding effector protein 2** | RABEP2 | T194S200 | | | | | | | **11.4** | **60.6** | R.HAPSLHGST*ELLPLS*R.D |
| **Rab GTPase binding effector protein 2** | RABEP2 | S193S200 | | | | | | | **12.3** | **15.7** | R.HAPSLHGS*TELLPLS*R.D |
| **Rab-coupling protein** | RAB11FIP | S315 | | | | | | | **73.8** | **50.0** | R.S*NVCINGNHVYLEQPEAK.G |
| **Rab-coupling protein** | RAB11FIP | S202 | | | | | | | **19.1** | **49.4** | K.NKDSGSDTASAIIPSTTPSVDS*DDESVVK.D |
| | RAB11FIP | T154 | | | | | | | **64.0** | **76.3** | R.NNMT*ASMFDLSMKDK.S |
| **Rab-coupling protein** | RAB11FIP | S356 | | | | | | | **9.5** | **119.0** | K.HLFS*STENLAAGSWK.E |
| **Rab-coupling protein** | RAB11FIP | T358 | | | | | | | **25.4** | **62.7** | K.HLFSST*ENLAAGSWK.E |
| **Rab-coupling protein** | RAB11FIP | S357 | | | | | | | **20.1** | **102.1** | K.HLFSS*TENLAAGSWK.E |
| **Rab-coupling protein** | RAB11FIP | S156 | | | | | | | **27.6** | **79.7** | R.NNMTAS*MFDLSMK.D |
| **Rab-coupling protein** | RAB11FIP | S296 | | | | | | | **26.9** | **76.3** | K.SPIMADLNLSLPSIPEVAS*DDER.I |
| **Rab-coupling protein** | RAB11FIP | S234 | | | | | | | **6.7** | **38.6** | K.TPLSQS*MSVLPTSKPEK.V |
| **Rab-coupling protein** | RAB11FIP | S287 | | | | | | | **12.9** | **69.5** | K.SPIMADLNLS*LPSIPEVASDDER.I |
| **Rab-coupling protein** | RAB11FIP | S290 | | | | | | | **5.3** | **43.1** | K.SPIMADLNLSLPS*IPEVASDDER.I |
| **Rab-coupling protein** | RAB11FIP | S280 | | | | | | | **12.2** | **15.4** | R.TAS*TDLK.Q |
| **Rab-coupling protein** | RAB11FIP | S190S202 | | | | | | | **13.3** | **21.9** | K.NKDSGSDTAS*AIIPSTTPSVDS*DDESVVK.D |
| **Rab-coupling protein** | RAB11FIP | S435 | | | | | | | **9.8** | **26.7** | R.RSS*LLSLMTGK.K |
| **Rab-coupling protein** | RAB11FIP | S435S438 | | | | | | | **13.4** | **38.7** | R.RSS*LLS*LMTGK.K |
| **RAB11 interacting protein** | RAB11FIP | S174 | | | | | | | **41.9** | **34.6** | R.NNLS*ASMFDLSMK.D |
| **RAB11 interacting protein** | RAB11FIP | S176 | | | | | | | **7.8** | **46.0** | R.NNLSAS*MFDLSMK.D |
| **RAB14** | RAB14 | Y99 | | | | | | | **-0.4** | **17.8** | R.STY*NHLSSWLTDAR.N |
| **RAB3A interacting protein** | RAB3IP | T289 | | | | | | | **15.2** | **44.7** | R.NKST*SSAMSGSHQDLSVIQPIVK.D |
| **RAB3A interacting protein** | RAB3IP | S288 | | | | | | | **15.3** | **14.0** | R.NKS*TSSAMSGSHQDLSVIQPIVK.D |
| **RAB7** | RAB7A | S72 | | | | | | | **75.4** | **63.7** | R.FQS*LGVAFYR.G |
| **Rabaptin 5** | RABEP1 | T486 | | | | | | | **22.0** | **47.9** | K.AMTPEQEET*ASLLSSVTQGMESAYVSPSGYR.L |
| **Rabaptin 5** | RABEP1 | S374S377 | | | | | | | **41.0** | **25.7** | R.GS*VHS*LDAGLLLPSGDPFSK.S |
| **Rabaptin 5** | RABEP1 | S407S410 | | | | | | | **22.6** | **54.5** | R.AQS*TDS*LGTSGSLQSK.A |
| **Rabaptin 5** | RABEP1 | T480 | | | | | | | **2.7** | **19.4** | K.AMT*PEQEETASLLSSVTQGMESAYVSPSGYR.L |
| **Rabaptin 5** | RABEP1 | S437T454S456 | | | | | | | **1.4** | **27.4** | K.SAGNLDES*DFGPLVGADSVSENFDT*AS*LGS*LQMPSGFMLTK.D |
| **Rabaptin 5** | RABEP1 | S488 | | | | | | | **6.0** | **24.0** | K.AMTPEQEETAS*LLSSVTQGMESAYVSPSGYR.L |
| **Rabaptin 5** | RABEP1 | S430S437S447 | | | | | | | **5.7** | **20.6** | K.S*AGNLDES*DFGPLVGADS*VSENFDT*ASLGSLQMPSGFMLTK.D |
| **Rabconnectin-3** | DMXL2 | S1857 | | | | | | | **49.2** | **78.9** | R.NLAS*PEGTLATLGLK.T |
| **Rabconnectin-3 beta** | WDR7 | S1153 | | | | | | | **-0.4** | **52.3** | R.SS*SQIPEGFGLTSGGSNYSLAR.H |
| **Rabconnectin-3 beta** | WDR7 | S1154 | | | | | | | **7.2** | **32.4** | R.SSS*QIPEGFGLTSGGSNYSLAR.H |
| **Rabphilin 3A** | RPH3A | S259 | | | | | | | **100.0** | **17.0** | R.S*PAGLRR.A |
| **RAD18** | RAD18 | S99S103 | | | | | | | **15.8** | **45.3** | R.NHLLQFALES*PAKS*PASSSSK.N |
| **RAD18** | RAD18 | S99 | | | | | | | **100.0** | **64.6** | R.NHLLQFALES*PAK.S |
| **RAD18** | RAD18 | S471 | | | | | | | **39.7** | **55.9** | R.DLLEEEEAWEASHKNDLQDTEIS*PR.Q |
| **RAD21** | RAD21 | S153 | | | | | | | **100.0** | **62.2** | R.EEVGNIS*ILQENDFGDFGMDDR.E |
| **RAD21** | RAD21 | Y105 | | | | | | | | **21.4** | R.EAAY*NAITLPEEFHDFDQPLPDLDDIDVAQQFSLNQSR.V |
| **RAD23A** | RAD23A | Y197 | | | | | | | | **72.7** | R.AVEY*LLTGIPGSPEPEHGSVQESQVSEQPATEAAGENPLEFLR.D |
| **RAD23A** | RAD23A | S295 | | | | | | | | **46.1** | R. [illegible] |
| **RAD23A** | RAD23A | S128 | | | | | | | **-2.8** | **17.8** | R.EDKSPSEES*APTTSPESVSGSVPSSGSSGR.E |
| **RAD23A** | RAD23A | S123 | | | | | | | **11.8** | **34.2** | R.EDKS*PSEESAPTTSPESVSGSVPSSGSSGR.E |
| **RAD23A** | RAD23A | S205 | | | | | | | **7.4** | **28.6** | R.AVEYLLTGIPGS*PEPEHGSVQESQVSEQPATEAAGENPLEFLR.D |
| **RAD23A** | RAD23A | Y313 | | | | | | | **2.7** | **14.6** | R. [illegible] |
| **RAD23B** | RAD23B | S160 | | | | | | | **11.3** | **59.0** | K.QEKPAEKPAETPVATS*PTATDSTSGDSSR.S |
| **RAD23B** | RAD23B | T155 | | | | | | | | **22.0** | K.QEKPAEKPAET*PVATSPTATDSTSGDSSR.S |
| **RAD23B** | RAD23B | T159 | | | | | | | **15.6** | **25.2** | K.QEKPAEKPAETPVAT*SPTATDSTSGDSSR.S |
| **RAD51 interacting protein** | RAD51AP1 | S21S27 | | | | | | | **7.1** | **16.7** | R.HKKPVNYSQFDHSDS*DDDFVS*ATVPLNK.K |
| **RAD51 interacting protein** | RAD51AP1 | S14S19 | | | | | | | **9.5** | **13.9** | R.HKKPVNYS*QFDHS*DSDDDFVSATVPLNKK.S |
| | RAD51AP1 | S19S21 | | | | | | | **27.2** | **28.7** | K.KPVNYSQFDHS*DS*DDDFVSATVPLNKK.S |
| **RAD51 interacting protein** | RAD51AP1 | Y13S19 | | | | | | | **4.6** | **13.8** | R.HKKPVNY*SQFDHS*DSDDDFVSATVPLNKK.S |
| **RAD51 interacting protein** | RAD51AP1 | Y13S21 | | | | | | | **3.0** | **27.8** | R.HKKPVNY*SQFDHSDS*DDDFVSATVPLNKK.S |
| **RAD51 interacting protein** | RAD51AP1 | S19S27 | | | | | | | **6.7** | **16.4** | R.HKKPVNYSQFDHS*DSDDDFVS*ATVPLNK.K |
| **RAD54-like 2** | RAD54L2 | S1061S1064 | | | | | | | **66.2** | **116.5** | K.APDPEGLARPVS*PDS*PEIISELQQYADVAAAR.E |
| **RAD54-like 2** | RAD54L2 | S1089 | | | | | | | **12.2** | **45.5** | R.QSSPS*TNAALPGPPAQLMDSSAVPGTALGTEPR.L |
| **RAD9** | RAD9A | S375S387 | | | | | | | **19.6** | **56.0** | R.S*PQGPSPVLAEDS*EGEG.- |
| **RAD9** | RAD9A | S277 | | | | | | | **25.0** | **90.1** | K.DSLLDGHFVLATLSDTDSHSQDLGS*PER.H |
| **RAD9** | RAD9A | S272 | | | | | | | | **67.8** | K.DSLLDGHFVLATLSDTDSHS*QDLGSPER.H |
| **RAD9** | RAD9A | S387 | | | | | | | **66.8** | **48.6** | R.SPQGPSPVLAEDS*EGEG.- |
| **RAD9** | RAD9A | S266 | | | | | | | **4.3** | **15.7** | K.DSLLDGHFVLATLS*DTDSHSQDLGSPER.H |

Peak Area: <-10, -5, -3, 0, 3, 5, >10

%CV: 0, 29, 42, 71, 86, >100

**White dots-** Significant change in peptide abundance at **5** %FDR compared to the timepoint with the minimum peak area for a given PSM

CarT: 0m, 2m, 5m — RajiB: 0m, 2m, 5m

| Protein Name | Gene | Phosphosites | Ascor | MOWSE | Sequence |
|---|---|---|---|---|---|
| Radial spokehead like 2 | RSPH3 | S530 | 15.0 | 22.6 | R.RSS*QER.K |
| Raf kinase inhibitor protein | PEBP1 | S52 | 22.4 | 75.4 | K.NRPTS*ISWDGLDSGK.L |
| Raf kinase inhibitor protein | PEBP1 | T51 | 2.8 | 18.4 | K.NRPT*SISWDGLDSGK.L |
| RAF1 | RAF1 | S259 | 9.6 | 71.2 | R.STS*TPNVHMVSTTLPVDSR.M |
| RAF1 | RAF1 | S29 | 6.5 | 104.1 | K.DAVFDGSSCIS*PTIVQQFGYQR.R |
| RAF1 | RAF1 | S25 | 5.3 | 58.1 | K.DAVFDGS*SCISPTIVQQFGYQR.R |
| RAF1 | ARAF | S621 | 16.6 | 35.6 | R.SAS*EPSLHR.T |
| RAF1 | RAF1 | T258 | 8.2 | 76.0 | R.ST*STPNVHMVSTTLPVDSR.M |
| RAF1 | RAF1 | S296 | 6.8 | 91.1 | R.SHSESASPSALSSS*PNNLSPTGWSQPK.T |
| RAF1 | RAF1 | S295 | 17.4 | 62.8 | R.SHSESASPSALSS*SPNNLSPTGWSQPK.T |
| RAF1 | RAF1 | S26 | 10.3 | 38.3 | K.DAVFDGSS*CISPTIVQQFGYQR.R |
| RAF1 | RAF1 | T260 | 7.4 | 25.8 | R.STST*PNVHMVSTTLPVDSR.M |
| RAF1 | RAF1 | S295S301 | 18.6 | 21.8 | R.SHSESASPSALSS*SPNNLS*PTGWSQPK.T |
| RAF1 | ARAF | S619 | 7.2 | 45.3 | R.S*ASEPSLHR.T |
| Raft linking protein | RFTN1 | S199 | 80.7 | 59.6 | R.GDHAS*LENEKPGTGDVCSAPAGR.N |
| Raft linking protein | RFTN1 | S220 | 76.1 | 35.5 | R.NQS*PEPSSGPR.G |
| Ral guanine nucleotide dissociation stimulator-like 3 | RGL3 | S361S366 | 100.0 | 11.7 | K.RS*WGAVS*R.E |
| RalA binding protein 1 | RALBP1 | S92S93 | 70.9 | 122.0 | R.TEGYAAFQEDS*S*GDEAESPSK.M |
| RalA binding protein 1 | RALBP1 | S30 | 17.7 | 57.3 | R.TPSS*EEISPTKFPGLYR.T |
| RalA binding protein 1 | RALBP1 | S29S34 | 17.3 | 59.5 | R.TPS*SEEIS*PTKFPGLYR.T |
| RalA binding protein 1 | RALBP1 | S29 | 13.4 | 31.1 | R.TPS*SEEISPTKFPGLYR.T |
| RalA binding protein 1 | RALBP1 | S29S30 | 6.1 | 43.7 | R.TPS*S*EEISPTKFPGLYR.T |
| RalA binding protein 1 | RALBP1 | S30S34 | 13.4 | 56.9 | R.TPSS*EEIS*PTKFPGLYR.T |
| RalA binding protein 1 | RALBP1 | Y85S92 | 13.1 | 29.4 | R.TEGY*AAFQEDS*SGDEAESPSK.M |
| RalA binding protein 1 | RALBP1 | T27S29 | 6.6 | 33.3 | R.T*PS*SEEISPTKFPGLYR.T |
| RALBP1 | REPS1 | S656 | 30.0 | 18.5 | R.LKS*EDELRPEVDEHTQK.T |
| RALBP1 | REPS1 | S220 | 6.1 | 43.3 | R.RQS*SSYDDPWK.I |
| RALBP1 | REPS1 | S110S114S118S | 42.3 | 22.6 | R.TSADAQEPAS*PVVS*PQQS*PPTS*PHTWR.K |
| RALBP1 | REPS1 | T429 | 51.2 | 26.3 | R.TGSDHTNPT*SPLLVKPSDLLEENK.I |
| RALBP1 | REPS1 | S340 | 28.4 | 50.0 | K.LIDLEDSADVGDQPGEVGYSGS*PAEAPPSK.S |
| RALBP1 | REPS1 | S508 | -0.3 | 35.5 | R.SS*SLDMNR.T |
| RALBP1 | REPS1 | S378 | 19.7 | 54.6 | R.SSS*SQTLTQFDSNIAPADPDTAIVHPVPIR.M |
| RALBP1 | REPS1 | S378T383 | 13.1 | 15.6 | R.SSS*SQTLT*QFDSNIAPADPDTAIVHPVPIR.M |
| RALBP1 | REPS1 | S484S505 | 8.9 | 28.9 | R.SHS*GTSPDNTAPPPPPPRPQPSHS*R.S |
| RALBP1 | REPS1 | S484T491 | 13.2 | 25.2 | R.SHS*GTSPDNT*APPPPPPRPQPSHSR.S |
| RALBP1 | REPS1 | S376 | -1.7 | 54.6 | R.S*SSSQTLTQFDSNIAPADPDTAIVHPVPIR.M |
| RALBP1 | REPS1 | S378S379 | 7.6 | 63.0 | R.SSS*S*QTLTQFDSNIAPADPDTAIVHPVPIR.M |
| RALBP1 | REPS1 | T358 | 6.1 | 62.2 | K.SPSMPSLNQT*WPELNQSSEQWETFSER.S |
| RALBP1 | REPS1 | S349 | 7.2 | 13.4 | K.S*PSMPSLNQTWPELNQSSEQWETFSER.S |
| RALBP1 | REPS1 | T486S487 | 27.6 | 23.2 | R.SHSGT*S*PDNTAPPPPPPRPQPSHSR.S |
| RALBP1 | REPS1 | S484T486 | 5.1 | 15.7 | R.SHS*GT*SPDNTAPPPPPPRPQPSHSR.S |
| RALBP1 | REPS1 | S484S487S505 | 8.3 | 11.8 | R.SHS*GTS*PDNTAPPPPPPRPQPSHS*R.S |
| RALBP1 | REPS1 | S484T486S505 | 13.2 | 10.9 | R.SHS*GT*SPDNTAPPPPPPRPQPSHS*R.S |
| RALBP1 | REPS1 | S110S118S122 | 12.5 | 15.8 | R.TSADAQEPAS*PVVSPQQS*PPTS*PHTWR.K |
| RALBP1 | REPS1 | S110S114S118T | 26.2 | 22.0 | R.TSADAQEPAS*PVVS*PQQS*PPT*SPHTWR.K |
| RALBP1 | REPS1 | S378S387 | 2.0 | 23.4 | R.SSS*SQTLTQFDS*NIAPADPDTAIVHPVPIR.M |
| RALBP1 | REPS1 | S354 | 21.6 | 57.7 | K.SPSMPS*LNQTWPELNQSSEQWETFSER.S |
| RALBP1 | REPS1 | S387 | 8.4 | 27.9 | R.SSSSQTLTQFDS*NIAPADPDTAIVHPVPIR.M |
| RALBP1 | REPS1 | S487S505 | 10.7 | 13.0 | R.SHSGTS*PDNTAPPPPPPRPQPSHS*R.S |
| RALBP1 | REPS1 | S68 | -5.2 | 48.5 | R.HAASYSSDS*ENQGSYSGVIPPPPGR.G |
| RALBP1 | REPS1 | S110S118T121 | 15.4 | 15.2 | R.TSADAQEPAS*PVVSPQQS*PPT*SPHTWR.K |
| RALBP1 | REPS1 | Y223 | 5.9 | 18.4 | R.RQSSSY*DDPWK.I |
| RALBP1 | REPS1 | S66 | 9.5 | 25.4 | R.HAASYSS*DSENQGSYSGVIPPPPGR.G |
| RALBP1 | REPS1 | S430 | 7.2 | 29.9 | R.TGSDHTNPTS*PLLVKPSDLLEENK.I |
| RALBP1 | REPS1 | S114S118T121 | 15.6 | 12.8 | R.TSADAQEPASPVVS*PQQS*PPT*SPHTWR.K |
| RALBP1 | REPS1 | S509 | 9.1 | 19.9 | R.SSS*LDMNR.T |
| RAM2 | CDCA7L | S21 | 100.0 | 95.2 | K.EVADIFNAPS*DDEEFVGFR.D |
| RAM2 | CDCA7L | T77T81 | 19.4 | 57.1 | R.IFIEDT*DSET*EDFAGFTQSDLNGK.T |
| RAM2 | CDCA7L | S195T196S197 | 10.1 | 31.7 | R.EDS*T*S*ES*EDDSRDESQESSDALLKR.T |
| RAM2 | CDCA7L | S79T81 | 9.4 | 52.4 | R.IFIEDTDS*ET*EDFAGFTQSDLNGK.T |
| RAM2 | CDCA7L | T77S79 | 31.4 | 50.5 | R.IFIEDT*DS*ETEDFAGFTQSDLNGK.T |
| Ran binding protein 1 | RANBP1 | T13S14 | 11.2 | 20.9 | K.DTHEDHDT*S*TENTDESNHDPQFEPIVSLPEQEIK.T |

Peak Area: <-10, -5, -3, 0, 3, 5, >10

%CV: 0, 29, 42, 71, 86, >100

**White dots-** Significant change in peptide abundance at **5** %FDR compared to the timepoint with the minimum peak area for a given PSM

CarT: 0m 2m 5m | RajiB: 0m 2m 5m

| Protein Name | Gene | Phosphosites | Ascor | MOWSE | Sequence |
|---|---|---|---|---|---|
| Ran binding protein 1 | RANBP1 | T18 | 8.4 | 12.1 | K.DTHEDHDTSTENT*DESNHDPQFEPIVSLPEQEIK.T |
| Ran binding protein 1 | RANBP1 | T18S21 | -1.7 | 10.8 | K.DTHEDHDTSTENT*DES*NHDPQFEPIVSLPEQEIK.T |
| Ran binding protein 1 | RANBP1 | S14T15 | 5.6 | 21.8 | K.DTHEDHDTS*T*ENTDESNHDPQFEPIVSLPEQEIK.T |
| Ran binding protein 1 | RANBP1 | T15S21 | 7.1 | 14.3 | K.DTHEDHDTST*ENTDES*NHDPQFEPIVSLPEQEIK.T |
| Ran binding protein 2 | RANBP2 | T1156 | 4.0 | 20.1 | K.NHET*DGGSAHGDDDDDGPHFEPVVPLPDKIEVK.T |
| Ran binding protein 2 | RANBP2 | S2900 | 64.1 | 38.3 | K.VGEDEDGS*DEEVVHNEDIHFEPIVSLPEVEVK.S |
| Ran binding protein 2 | RANBP2 | S1509 | 13.8 | 24.9 | R.KQS*LPATSIPTPASFK.F |
| Ran binding protein 2 | RANBP2 | S788 | 16.1 | 21.3 | K.YSLS*PSK.S |
| Ran binding protein 2 | RANBP2 | S1400 | -0.2 | 47.2 | K.TS*PENVQDR.F |
| Ran binding protein 3 | RANBP3 | S539 | 24.2 | 49.8 | K. MPAPEPGAAPSNEEDDS*DDDDVLAPSGATAAGAGDEGDGQTTGS |
| Ran binding protein 3 | RANBP3 | S548 | 9.6 | 40.8 | K. MPAPEPGAAPSNEEDDSDDDDVLAPS*GATAAGAGDEGDGQTTGS |
| Ran binding protein 3 | RANBP3 | S211 | 18.2 | 15.5 | K.ALSQTVPSSGTNGVSLPADCTGAVPAAS*PDTAAWR.S |
| Ran binding protein 3 | RANBP3 | S101S108 | 4.5 | 22.1 | R.SAGGSS*PEGGEDS*DREDGNYCPPVKR.E |
| Ran binding protein 3 | RANBP3 | S333 | 62.7 | 51.6 | R.VLS*PPKLNEVSSDANR.E |
| Ran binding protein 3 | RANBP3 | S96S101S108 | 3.7 | 13.3 | R.S*AGGSS*PEGGEDS*DREDGNYCPPVK.R |
| Ran binding protein 3 | RANBP3 | S533S539 | 15.5 | 12.9 | K. MPAPEPGAAPS*NEEDDS*DDDDVLAPSGATAAGAGDEGDGQTTG |
| Ran binding protein 3 | RANBP3 | S100S108 | 16.2 | 12.0 | R.SAGGS*SPEGGEDS*DREDGNYCPPVKR.E |
| Ran binding protein 3 | RANBP3 | S100S101S108 | 13.8 | 12.1 | R.SAGGS*S*PEGGEDS*DREDGNYCPPVK.R |
| Ran binding protein 3 | RANBP3 | S533 | | 35.5 | K. MPAPEPGAAPS*NEEDDSDDDDVLAPSGATAAGAGDEGDGQTTGS |
| Ran binding protein 3 | RANBP3 | S96S100S108 | 10.2 | 11.1 | R.S*AGGS*SPEGGEDS*DREDGNYCPPVK.R |
| RAN binding protein 3-like | RANBP3L | T142 | 100.0 | 14.2 | R.KT*FGHK.A |
| RAN binding protein 9 | RANBP9 | S470S485 | -0.7 | 15.3 | K.S*QDSYPVSPRPFSSPS*MSPSHGMNIHNLASGK.G |
| | RANBP9 | S477S482S487 | 19.0 | 11.3 | K.SQDSYPVS*PRPFS*SPSM#S*PSHGMNIHNLASGK.G |
| | RANBP9 | S477S482S485 | 23.4 | 21.2 | K.SQDSYPVS*PRPFS*SPS*M#SPSHGMNIHNLASGK.G |
| RAN binding protein 9 | RANBP9 | S477S487 | 26.2 | 26.5 | K.SQDSYPVS*PRPFSSPSMS*PSHGMNIHNLASGK.G |
| RAN binding protein 9 | RANBP9 | S477S483S487 | 27.8 | 37.2 | K.SQDSYPVS*PRPFSS*PSMS*PSHGMNIHNLASGK.G |
| RAN binding protein 9 | RANBP9 | S477S487S489 | 15.1 | 11.9 | K.SQDSYPVS*PRPFSSPSM#S*PS*HGMNIHNLASGK.G |
| RAN binding protein 9 | RANBP9 | S482S483S485 | 12.6 | 18.0 | K.SQDSYPVSPRPFS*S*PS*M#SPSHGMNIHNLASGK.G |
| RAN binding protein 9 | RANBP9 | S477S482 | 15.6 | 15.4 | K.SQDSYPVS*PRPFS*SPSMSPSHGMNIHNLASGK.G |
| RAN binding protein 9 | RANBP9 | S477S482S483 | 16.8 | 18.6 | K.SQDSYPVS*PRPFS*S*PSMSPSHGMNIHNLASGK.G |
| RAN binding protein 9 | RANBP9 | Y474S487 | 9.8 | 16.9 | K.SQDSY*PVSPRPFSSPSMS*PSHGMNIHNLASGK.G |
| RAN binding protein 9 | RANBP9 | S477S485 | 22.1 | 25.6 | K.SQDSYPVS*PRPFSSPS*MSPSHGMNIHNLASGK.G |
| RAN binding protein 9 | RANBP9 | S477S485S487 | 19.2 | 18.5 | K.SQDSYPVS*PRPFSSPS*MS*PSHGMNIHNLASGK.G |
| RAN binding protein 9 | RANBP9 | Y474S485 | 11.1 | 23.2 | K.SQDSY*PVSPRPFSSPS*MSPSHGMNIHNLASGK.G |
| RAN binding protein 9 | RANBP9 | S482S485 | -1.1 | 13.2 | K.SQDSYPVSPRPFS*SPS*MSPSHGMNIHNLASGK.G |
| RAN binding protein 9 | RANBP9 | S483S485 | 4.1 | 11.9 | K.SQDSYPVSPRPFSS*PS*MSPSHGMNIHNLASGK.G |
| RAN binding protein 9 | RANBP9 | S477S483 | 18.5 | 12.0 | K.SQDSYPVS*PRPFSS*PSMSPSHGMNIHNLASGK.G |
| RAN binding protein 9 | RANBP9 | S482S483 | 3.1 | 10.9 | K.SQDSYPVSPRPFS*S*PSMSPSHGMNIHNLASGK.G |
| RAN binding protein 9 | RANBP9 | S470S473Y474 | | 22.2 | K.S*QDS*Y*PVSPRPFSSPSM#SPSHGMNIHNLASGK.G |
| RAN binding protein 9 | RANBP9 | S485S487 | 6.0 | 11.6 | K.SQDSYPVSPRPFSSPS*MS*PSHGMNIHNLASGK.G |
| RAN binding protein 9 | RANBP9 | S477S483S489 | 13.4 | 23.7 | K.SQDSYPVS*PRPFSS*PSMSPS*HGMNIHNLASGK.G |
| | RANBP9 | S477S482S489 | 9.9 | 15.1 | K.SQDSYPVS*PRPFS*SPSM#SPS*HGMNIHNLASGK.G |
| RAN, member RAS oncogene family | RAN | S135 | 100.0 | 27.4 | K.S*IVFHR.K |
| Rap guanine nucleotide exchange factor 6 | RAPGEF6 | S1590 | 8.4 | 19.9 | K.LGDVTDADS*EADENEQVSAV.- |
| Rap guanine nucleotide exchange factor GEF 6 | RAPGEF6 | S1094 | 9.8 | 22.4 | R.RSS*LLNAK.K |
| Rap guanine nucleotide exchange factor GEF 6 | RAPGEF6 | S1494 | 16.9 | 27.9 | K.GLIVYCVTS*PK.K |
| RAP1 GTPase activating protein 1 | RAP1GAP | S484 | 13.9 | 35.1 | K.AAGISLIVPGKS*PTR.K |
| Rap1 interacting factor 1 | RIF1 | S1579 | 6.9 | 39.9 | K.SNES*VDIQDQEEK.V |
| Rap1 interacting factor 1 | RIF1 | S2196 | 12.1 | 47.2 | R.SQEDEISS*PVNK.V |
| Rap1 interacting factor 1 | RIF1 | S2144 | 60.1 | 76.6 | K.VEEPSQCLASGTAISELIIEDNNAS*PQK.L |
| Rap1 interacting factor 1 | RIF1 | S2195 | 47.7 | 19.7 | R.SQEDEIS*SPVNK.V |
| Rap1 interacting factor 1 | RIF1 | S1454 | 100.0 | 60.5 | K.S*PLHIKDDVLPK.Q |
| | APBB1IP | S526 | -0.4 | 59.6 | R.SS*DTSGSPATPLK.A |
| RAP80 | UIMC1 | S677 | 100.0 | 23.1 | R.DLNES*PVK.S |
| RAP80 | UIMC1 | S463 | 18.8 | 17.6 | R.EVS*PGSR.D |
| Raptor | RPTOR | S722 | 15.0 | 35.5 | R.SVSS*YGNIR.A |
| Raptor | RPTOR | S877 | 62.5 | 108.0 | K.GVHIHQAGGS*PPASSTSSSSLTNDVAK.Q |
| Raptor | RPTOR | T857S863 | 15.6 | 56.6 | R.VLDTSSLT*QSAPAS*PTNK.G |
| Raptor | RPTOR | S859S863 | 24.3 | 72.4 | R.VLDTSSLTQS*APAS*PTNK.G |
| Raptor | RPTOR | S863 | 19.3 | 71.8 | R.VLDTSSLTQSAPAS*PTNK.G |
| Raptor | RPTOR | S721 | 10.6 | 12.3 | R.SVS*SYGNIR.A |
| Ras association (RalGDS/AF-6) domain family 4 | RASSF4 | S91 | 15.1 | 34.3 | R.RPS*CPLKEPSPQNGNITAQGPSIQPVHK.A |

Peak Area: <-10, -5, -3, 0, 3, 5, >10

%CV: 0, 29, 42, 71, 86, >100

**White dots-** Significant change in peptide abundance at **5** %FDR compared to the timepoint with the minimum peak area for a given PSM

| Protein Name | Gene | Phosphosites | CarT 0m 2m 5m | RajiB 0m 2m 5m | Ascor | MOWSE | Sequence |
|---|---|---|---|---|---|---|---|
| **RAS family, member RAB11B** | RAB11A | S42 | | | **17.0** | **64.5** | K.S*TIGVEFATR.S |
| **RAS guanyl nucleotide releasing protein 1** | RASGRP1 | T710S715 | | | **19.6** | **18.6** | K.TAQDTLYVLPSPT*SPCPS*PVLVR.K |
| **RAS guanyl nucleotide releasing protein 2** | RASGRP2 | S616S625S627 | | | **16.7** | **21.2** | R.AQS*VSLEGSAPS*PS*PMHSHHHR.A |
| **RAS guanyl nucleotide releasing protein 2** | RASGRP2 | S616S625 | | | **25.4** | **65.3** | R.AQS*VSLEGSAPS*PSPMHSHHHR.A |
| **RAS guanyl nucleotide releasing protein 2** | RASGRP2 | S616 | | | **43.4** | **92.5** | R.AQS*VSLEGSAPSPSPMHSHHHR.A |
| **RAS guanyl nucleotide releasing protein 2** | RASGRP2 | S178 | | | **45.4** | **45.8** | R.RHS*SLIDIDSVPTYK.W |
| **RAS guanyl nucleotide releasing protein 2** | RASGRP2 | S638 | | | **26.0** | **32.1** | R.AFS*FSLPRPGR.R |
| **RAS guanyl nucleotide releasing protein 2** | RASGRP2 | S618S627 | | | **5.4** | **34.1** | R.AQSVS*LEGSAPSPS*PMHSHHHR.A |
| **RAS guanyl nucleotide releasing protein 2** | RASGRP2 | S618 | | | **5.3** | **60.9** | R.AQSVS*LEGSAPSPSPMHSHHHR.A |
| **RAS guanyl nucleotide releasing protein 2** | RASGRP2 | S178S179 | | | **39.0** | **14.6** | R.RHS*S*LIDIDSVPTYK.W |
| **RAS guanyl nucleotide releasing protein 2** | RASGRP2 | S616S627 | | | **8.0** | **45.4** | R.AQS*VSLEGSAPSPS*PMHSHHHR.A |
| **RAS guanyl nucleotide releasing protein 2** | RASGRP2 | S622S625 | | | **16.0** | **33.2** | R.AQSVSLEGS*APS*PSPMHSHHHR.A |
| **RAS guanyl nucleotide releasing protein 2** | RASGRP2 | S622S627 | | | **13.9** | **20.1** | R.AQSVSLEGS*APSPS*PMHSHHHR.A |
| **RAS guanyl releasing protein 3** | RASGRP3 | S559T580 | | | **10.8** | **10.8** | R.APSLSSGHGS*LPGSPSLPPAQDEVFEFPGVT*AGHR.D |
| **Ras-GTPase activating protein binding protein 2** | G3BP2 | T227 | | | **18.6** | **68.3** | K.STT*PPPAEPVSLPQEPPKPR.V |
| **Ras-GTPase activating protein binding protein 2** | G3BP2 | S141S149 | | | **64.3** | **122.5** | R.YEDEVFGDS*EPELDEES*EDEVEEEQEER.Q |
| **Ras-GTPase activating protein binding protein 2** | G3BP2 | S149 | | | **23.5** | **34.9** | R.YEDEVFGDSEPELDEES*EDEVEEEQEER.Q |
| **Ras-GTPase activating protein binding protein 2** | G3BP2 | S141 | | | **13.8** | **32.8** | R.YEDEVFGDS*EPELDEESEDEVEEEQEER.Q |
| **Ras-GTPase activating protein binding protein 2** | G3BP2 | S225 | | | **8.4** | **23.9** | K.S*TTPPPAEPVSLPQEPPKPR.V |
| **Ras-GTPase-activating protein SH3-domain-binding protein** | G3BP1 | S232 | | | **22.2** | **130.8** | K.SSS*PAPADIAQTVQEDLR.T |
| **Ras-GTPase-activating protein SH3-domain-binding protein** | G3BP1 | S149 | | | **62.9** | **84.4** | R.YQDEVFGGFVTEPQEES*EEEVEEPEER.Q |
| **Ras-GTPase-activating protein SH3-domain-binding protein** | G3BP1 | S231 | | | **-0.3** | **117.9** | K.SS*SPAPADIAQTVQEDLR.T |
| **Ras-GTPase-activating protein SH3-domain-binding protein** | G3BP1 | S230 | | | **5.5** | **12.7** | K.S*SSPAPADIAQTVQEDLR.T |
| **RB1 inducible coiled coil 1** | RB1CC1 | T238 | | | **-0.2** | **94.1** | R.ST*ELVLSPDMPR.T |
| **RBM15B** | RBM15B | S552 | | | **12.1** | **37.3** | R.TFLEGDWTS*PSK.S |
| **RBM15B** | RBM15B | S231S232S233 | | | **12.6** | **12.3** | R.RSSS*S*S*AAASTPPPGPPAPADPLGYLPLHGGYQYK.Q |
| **RBM15B** | RBM15B | S265S267 | | | **100.0** | **18.8** | R.S*LS*PVAAPPLREPR.A |
| **RBMX** | RBMX | S208 | | | **39.3** | **26.3** | R.DVYLS*PR.D |
| **RCAS1** | EBAG9 | S36 | | | **84.9** | **120.9** | R.KLS*GDQITLPTTVDYSSVPK.Q |
| **RCAS1** | EBAG9 | T41 | | | **8.7** | **39.3** | R.KLSGDQIT*LPTTVDYSSVPK.Q |
| **RE repeats encoding gene** | RERE | T144 | | | **-5.5** | **11.4** | R.SPT*PALCDPPACSLPVASQPPQHLSEAGR.G |
| **RE repeats encoding gene** | RERE | S600 | | | **-0.4** | **23.1** | R.TS*PINEDIR.S |
| **RE repeats encoding gene** | RERE | S1106S1113S11 | | | **27.1** | **18.5** | K.EEALDDAEEPES*PPPPPRS*PS*PEPTVVDTPSHASQSAR.F |
| **Receptor expressed in lymphoid tissues like 1** | RELL1 | S109 | | | **9.5** | **47.2** | K.IELNDS*VNENSDTVGQIVHYIMK.N |
| **Receptor expressed in lymphoid tissues like 1** | RELL1 | S109T116 | | | **4.4** | **20.3** | K.IELNDS*VNENSDT*VGQIVHYIMK.N |
| **Receptor expressed in lymphoid tissues like 1** | RELL1 | S109S114 | | | **15.4** | **34.2** | K.IELNDS*VNENS*DTVGQIVHYIMK.N |
| **Receptor expressed in lymphoid tissues like 1** | RELL1 | S114T116 | | | **5.4** | **17.1** | K.IELNDSVNENS*DT*VGQIVHYIMK.N |
| **Receptor TNFRSF-interacting serine-threonine kinase 1** | RIPK1 | S320 | | | **117.6** | **101.9** | R.MQS*LQLDCVAVPSSR.S |
| **RecQ protein like 3** | BLM | S147 | | | **1.7** | **43.4** | K.KLEFSSSPDS*LSTINDWDDMDDFDTSETSK.S |
| **Regulating synaptic membrane exocytosis 3** | RIMS3 | S295S298 | | | **19.7** | **18.5** | R.RLS*QSS*LESATSPSCS.- |
| **Regulating synaptic membrane exocytosis 4** | RIMS4 | S6 | | | **10.9** | **15.1** | M.ERSQS*R.L |
| **Regulator of G protein signaling 14** | RGS14 | S218 | | | **100.0** | **38.5** | K.S*LPLGVEELGQLPPVEGPGGRPLRK.S |
| **Regulator of G protein signaling 14** | RGS14 | S40S45 | | | **1.5** | **48.0** | R.GSS*LSIHS*LPSGPSSPFPTEEQPVASWALSFER.L |
| **Regulator of G protein signaling 14** | RGS14 | S40S48 | | | **3.7** | **41.2** | R.GSS*LSIHSLPS*GPSSPFPTEEQPVASWALSFER.L |
| **Regulator of G protein signaling 14** | RGS14 | S48S51S52 | | | **1.6** | **31.7** | R.GSSLSIHSLPS*GPS*S*PFPTEEQPVASWALSFER.L |
| **Regulator of G protein signaling 14** | RGS14 | S39S40S42 | | | **11.1** | **49.5** | R.GS*S*LS*IHSLPSGPSSPFPTEEQPVASWALSFER.L |
| **Regulator of G protein signaling 14** | RGS14 | S39S40S48 | | | **4.0** | **29.7** | R.GS*S*LSIHSLPS*GPSSPFPTEEQPVASWALSFER.L |
| **Regulator of G protein signaling 14** | RGS14 | S39S42S48 | | | **7.1** | **40.5** | R.GS*SLS*IHSLPS*GPSSPFPTEEQPVASWALSFER.L |
| **Regulator of G protein signaling 3** | RGS3 | S943 | | | **33.7** | **21.7** | R.THS*EGSLLQEPR.G |
| **Regulator of G protein signaling 3** | RGS3 | S943S946 | | | **45.4** | **28.1** | R.THS*EGS*LLQEPR.G |
| **Regulator of G protein signaling 3** | RGS3 | S916 | | | **3.1** | **27.4** | K.RS*SMIETGQGAEGGLSLR.V |
| **Regulator of nonsense transcripts 1** | UPF1 | S1107 | | | **37.7** | **124.5** | K.SQIDVALS*QDSTYQGER.A |
| **Regulator of nonsense transcripts 1** | UPF1 | S1127 | | | **28.2** | **72.4** | R.AYQHGGVTGLS*QY.- |
| **Regulator of nonsense transcripts 1** | UPF1 | S1107S1110 | | | **20.4** | **98.5** | K.SQIDVALS*QDS*TYQGER.A |
| **Regulator of nonsense transcripts 1** | UPF1 | Y1129 | | | **12.1** | **71.6** | R.AYQHGGVTGLSQY*.- |
| **Regulatory factor X 1** | RFX1 | S117 | | | **20.7** | **55.9** | R.ASETVS*EASPGSTASQTGVPTQVVQQVQGTQQR.L |
| **Regulatory factor X 1** | RFX1 | S120 | | | **4.6** | **36.8** | R.ASETVSEAS*PGSTASQTGVPTQVVQQVQGTQQR.L |
| **Regulatory factor X 3 influences HLA class II expression** | RFX3 | S664 | | | **54.3** | **85.6** | R.VAQATGETPIAVMGEFGDLNAVS*PGNLDK.D |
| **Regulatory factor X domain containing 2** | RFX7 | S1178 | | | **35.4** | **76.4** | R.NLS*GSTLYPVSNIPR.S |
| **Renal cell carcinoma, papillary 1** | PRCC | S157S159 | | | **45.9** | **43.8** | K.IAAPELHKGDS*DS*EEDEPTK.K |
| **Renal cell carcinoma, papillary 1** | PRCC | S267 | | | **43.5** | **77.0** | K.QITQEEDDS*DEEVAPENFFSLPEK.A |
| **Replication factor C1** | RFC1 | S108T110 | | | **-0.1** | **47.8** | R.QDPVTYIS*ET*DEEDDFMCKK.A |

Peak Area: <-10, -5, -3, 0, 3, 5, >10

%CV: 0, 29, 42, 71, 86, >100

**White dots-** Significant change in peptide abundance at **5** %FDR compared to the timepoint with the minimum peak area for a given PSM

CarT (0m, 2m, 5m) RajiB (0m, 2m, 5m)

| Protein Name | Gene | Phosphosites | Ascor | MOWSE | Sequence |
|---|---|---|---|---|---|
| Replication factor C1 | RFC1 | S156S164T171 | 10.2 | 37.5 | K.NKPLS*PIKLTPTS*VLDYFGT*GSVQR.S |
| Replication factor C1 | RFC1 | S281S283 | 15.6 | 23.5 | R.S*YS*PR.S |
| Replication factor C1 | RFC1 | S368 | 28.9 | 52.1 | K.ESVS*PEDSEK.K |
| Replication factor C1 | RFC1 | S69S71 | 43.7 | 84.3 | R.IIYDS*DS*ESEETLQVK.N |
| Replication factor C1 | RFC1 | Y67S71 | 19.1 | 85.2 | R.IIY*DSDS*ESEETLQVK.N |
| Replication factor C1 | RFC1 | S156T163 | 14.5 | 32.3 | K.NKPLS*PIKLTPT*SVLDYFGTGSVQR.S |
| Replication factor C1 | RFC1 | S156T161 | 26.8 | 65.7 | K.NKPLS*PIKLT*PTSVLDYFGTGSVQR.S |
| Replication factor C1 | RFC1 | S156T161T171 | 14.3 | 40.4 | K.NKPLS*PIKLT*PTSVLDYFGT*GSVQR.S |
| Replication factor C1 | RFC1 | S281Y282 | 14.0 | 17.3 | R.S*Y*SPR.S |
| Replication factor C1 | RFC1 | Y282S283 | 10.6 | 26.2 | R.SY*S*PR.S |
| Replication factor C1 | RFC1 | T161T171 | 18.2 | 40.7 | K.LT*PTSVLDYFGT*GSVQR.S |
| Replication factor C1 | RFC1 | T163T171 | 9.0 | 44.1 | K.LTPT*SVLDYFGT*GSVQR.S |
| Replication factor C1 | RFC1 | S156S164S173 | 6.3 | 51.9 | K.NKPLS*PIKLTPTS*VLDYFGTGS*VQR.S |
| Replication factor C1 | RFC1 | T163S164T171 | 5.3 | 41.8 | K.NKPLSPIKLTPT*S*VLDYFGT*GSVQR.S |
| REST corepressor | RCOR1 | S457 | 24.1 | 16.1 | K.S*PDNSIK.M |
| REST corepressor | RCOR1 | S257 | 100.0 | 46.8 | R.EREES*EDELEEANGNNPIDIEVDQNK.E |
| REST corepressor 3 | RCOR3 | S156S171 | 24.5 | 31.1 | R.HNQGDS*DDDVEETHPMDGNDS*DYDPKK.E |
| Reticulon 2 | RTN2 | S94 | 10.6 | 12.1 | R.SVS*EPR.D |
| | RTN4 | S182 | 10.3 | 41.3 | K.RRGSS*GSVDETLFALPAASEPVIR.S |
| | RTN4 | S184 | 12.2 | 48.5 | K.RRGSSGS*VDETLFALPAASEPVIR.S |
| Reticulon 4 | RTN4 | T188 | 10.5 | 17.3 | R.RGSSGSVDET*LFALPAASEPVIR.S |
| Reticulon 4 | RTN4 | S181S182 | 16.4 | 48.9 | R.RGS*S*GSVDETLFALPAASEPVIR.S |
| | RTN4 | S181 | 9.1 | 43.8 | K.RRGS*SGSVDETLFALPAASEPVIR.S |
| Reticulon 4 | RTN4 | S184T188 | 5.5 | 12.5 | R.RGSSGS*VDET*LFALPAASEPVIR.S |
| Reticulon 4 | RTN4 | S181S184 | 10.6 | 41.2 | R.RGS*SGS*VDETLFALPAASEPVIR.S |
| | RTN4 | S182S184 | 3.1 | 21.9 | R.GSS*GS*VDETLFALPAASEPVIR.S |
| Reticulon 4 | RTN4 | S181T188 | 21.8 | 13.6 | K.RRGS*SGSVDET*LFALPAASEPVIR.S |
| Retinoblastoma 1 | RB1 | S249 | 82.7 | 50.4 | K.TAVIPINGS*PR.T |
| Retinoblastoma 1 | RB1 | T823T826 | 21.5 | 34.6 | K.ISEGLPTPT*KMT*PR.S |
| Retinoblastoma 1 | RB1 | T373 | 124.8 | 44.3 | R.KSNLDEEVNVIPPHT*PVR.T |
| Retinoblastoma 1 | RB1 | T821T826 | 18.9 | 20.0 | K.ISEGLPT*PTKMT*PR.S |
| Retinoblastoma 1 | RB1 | T353 | 17.2 | 24.3 | K.TLQTDSIDSFET*QRTPR.K |
| Retinoblastoma 1 | RB1 | S788 | 18.0 | 51.7 | R.S*PYKFPSSPLR.I |
| Retinoblastoma 1 | RB1 | S788S794 | 26.8 | 36.3 | R.S*PYKFPS*SPLR.I |
| Retinoblastoma 1 | RB1 | S788S795 | 10.0 | 25.2 | R.S*PYKFPSS*PLR.I |
| Retinoblastoma 1 | RB1 | Y790S794 | 24.3 | 39.9 | R.SPY*KFPS*SPLR.I |
| | RB1 | T9 | | 58.4 | R.KT*AATAAAAAAEPPAPPPPPPPEEDPEQDSGPEDLPLVR.L |
| Retinoblastoma 1 | RB1 | S807S811 | 55.8 | 55.3 | R.IPGGNIYIS*PLKS*PYK.I |
| | RB1 | S37 | 80.2 | 47.6 | R.KTAATAAAAAAEPPAPPPPPPPEEDPEQDS*GPEDLPLVR.L |
| Retinoblastoma 1 | RB1 | Y606S612 | 7.3 | 17.9 | K.DREGPTDHLESACPLNLPLQNNHTAADMY*LSPVRS*PK.K |
| Retinoblastoma 1 | RB1 | S608S612 | 20.9 | 31.7 | K.DREGPTDHLESACPLNLPLQNNHTAADMYLS*PVRS*PK.K |
| Retinoblastoma 1 | RB1 | T601 | 8.2 | 38.4 | K.DREGPTDHLESACPLNLPLQNNHT*AADMYLSPVR.S |
| Retinoblastoma 1 | RB1 | S788S794S807 | 37.8 | 23.9 | R.S*PYKFPS*SPLRIPGGNIYIS*PLK.S |
| Retinoblastoma 1 | RB1 | S838 | 33.2 | 34.1 | R.ILVSIGES*FGTSEK.F |
| Retinoblastoma 1 | RB1 | S855 | 100.0 | 58.7 | K.INQMVCNS*DR.V |
| Retinoblastoma 1 | RB1 | S807S811T821 | 13.1 | 29.3 | R.IPGGNIYIS*PLKS*PYKISEGLPT*PTK.M |
| Retinoblastoma 1 | RB1 | S807 | 44.9 | 65.7 | R.IPGGNIYIS*PLK.S |
| Retinoblastoma 1 | RB1 | T583S588 | | 16.1 | K.DREGPT*DHLES*ACPLNLPLQNNHTAADMYLSPVRSPK.K |
| Retinoblastoma 1 | RB1 | S794 | 71.6 | 30.4 | R.SPYKFPS*SPLR.I |
| Retinoblastoma 1 | RB1 | S811Y813 | 5.9 | 13.4 | R.IPGGNIYISPLKS*PY*K.I |
| Retinoblastoma 1 | RB1 | T601S612 | 5.9 | 14.9 | R.EGPTDHLESACPLNLPLQNNHT*AADM#YLSPVRS*PK.K |
| Retinoblastoma 1 | RB1 | S807Y813 | 8.9 | 12.7 | R.IPGGNIYIS*PLKSPY*K.I |
| Retinoblastoma 1 | RB1 | S608 | 3.8 | 12.2 | R.EGPTDHLESACPLNLPLQNNHTAADMYLS*PVR.S |
| Retinoblastoma 1 | RB1 | S624 | 6.5 | 19.6 | R.VNS*TANAETQATSAFQTQKPLK.S |
| Retinoblastoma 1 | RB1 | S350 | 28.3 | 15.3 | K.TLQTDSIDS*FETQRTPR.K |
| Retinoblastoma 1 | RB1 | Y790 | 9.2 | 47.2 | R.SPY*KFPSSPLR.I |
| Retinoblastoma 1 | RB1 | Y606S608S612 | 1.1 | 16.0 | R.EGPTDHLESACPLNLPLQNNHTAADMY*LS*PVRS*PK.K |
| Retinoblastoma 1 | RB1 | T841 | 13.8 | 21.1 | R.ILVSIGESFGT*SEK.F |
| Retinoblastoma 1 | RB1 | Y790S795 | 5.8 | 32.3 | R.SPY*KFPSS*PLR.I |
| Retinoblastoma 1 | RB1 | Y606S608 | 21.5 | 10.7 | K.DREGPTDHLESACPLNLPLQNNHTAADMY*LS*PVRSPK.K |
| Retinoblastoma 1 | RB1 | S788S795S807 | 9.6 | 11.6 | R.S*PYKFPSS*PLRIPGGNIYIS*PLK.S |
| Retinoblastoma associated factor 600 | UBR4 | S2718 | 40.2 | 30.8 | R.HVTLPS*SPR.S |

Peak Area: <-10, -5, -3, 0, 3, 5, >10

%CV: 0, 29, 42, 71, 86, >100

**White dots-** Significant change in peptide abundance at **5** %FDR compared to the timepoint with the minimum peak area for a given PSM

CarT (0m, 2m, 5m) — RajiB (0m, 2m, 5m)

| Protein Name | Gene | Phosphosites | Ascor | MOWSE | Sequence |
|---|---|---|---|---|---|
| **Retinoblastoma associated factor 600** | UBR4 | S2719 | **8.2** | **26.3** | R.HVTLPSS*PR.S |
| **Retinoblastoma binding protein 2** | KDM5A | S1111 | **109.0** | **101.6** | K.DLDLEPLS*DLEEGLEETR.D |
| **Retinoblastoma binding protein 5** | RBBP5 | S350 | **100.0** | **66.8** | R.ES*EFDIEDEDK.S |
| **Retinoblastoma binding protein 5** | RBBP5 | S525 | **29.3** | **28.2** | K.VQAELS*QPLTAGGAISELL.- |
| **Retinoblastoma binding protein 6** | RBBP6 | S1277 | **11.8** | **33.1** | K.VTGTEGSSSTLVDYTSTSSTGGS*PVRK.S |
| **Retinoblastoma binding protein 6** | RBBP6 | S1179 | **65.5** | **71.4** | K.LEVTEIVKPS*PK.R |
| **Retinoblastoma binding protein 6** | RBBP6 | S1328 | **100.0** | **79.5** | K.WDKDDFES*EEEDVK.S |
| **Retinoblastoma binding protein 6** | RBBP6 | S873 | **46.9** | **34.3** | R.NS*PFTR.G |
| **Retinoblastoma binding protein 6** | RBBP6 | S770S772 | **100.0** | **33.4** | R.S*RS*PQAFR.G |
| **Retinoblastoma binding protein 6** | RBBP6 | S861 | **100.0** | **21.3** | R.ENFS*PER.F |
| **Retinoblastoma like 1** | RBL1 | S988T997 | **16.2** | **14.9** | R.ISQQHSIYIS*PHKNGSGLT*PR.S |
| **Retinoblastoma like 1** | RBL1 | S749S762 | **25.7** | **36.6** | K.VKS*PVSLTAHSLIGAS*PK.Q |
| **Retinoblastoma like 2** | RBL2 | T1065S1068 | **11.7** | **31.8** | R.IQLSQNHPVYISPHKNET*MLS*PR.E |
| **Retinoblastoma like 2** | RBL2 | S1059S1068 | **9.8** | **21.2** | R.IQLSQNHPVYIS*PHKNETMLS*PR.E |
| **Retinoblastoma like 2** | RBL2 | S1059T1065 | **13.9** | **32.9** | R.IQLSQNHPVYIS*PHKNET*MLSPR.E |
| **RFFL ring finger and FYVE-like domain containing 1** | hCG_2039 | S240 | **19.2** | **75.9** | R.RAS*LSDLTDLEDIEGLTVR.Q |
| | hCG_2039 | S242 | **6.1** | **137.1** | R.ASLS*DLTDLEDIEGLTVR.Q |
| **RFFL ring finger and FYVE-like domain containing 1** | RFFL | S226S229S232 | **18.2** | **42.3** | R.VPAEDETQS*IDS*EDS*FVPGR.R |
| **RFFL ring finger and FYVE-like domain containing 1** | hCG_2039 | T245 | **-0.2** | **15.2** | R.ASLSDLT*DLEDIEGLTVR.Q |
| **RGD, leucine-rich repeat, tropomodulin and proline-rich containing protein** | CARMIL2 | S1362 | **100.0** | **23.2** | R.AVS*VHEDQLQAPAERPLR.L |
| **Rho family-interacting cell polarization regulator 1** | RIPOR1 | S20 | **19.6** | **19.8** | R.THS*MMSLSVRPQR.R |
| **RHO GDP dissociation inhibitor beta** | ARHGDIB | S145 | **9.0** | **30.6** | K.ATFMVGS*YGPRPEEYEFLTPVEEAPK.G |
| **Rho GTPase activating protein 11A** | ARHGAP1 | S285 | **60.5** | **42.7** | R.RQS*VGDFVSGALNK.F |
| **Rho GTPase activating protein 12** | ARHGAP1 | T231S240 | **19.1** | **33.5** | R.ATT*PPNQGRPDS*PVYANLQELK.I |
| **Rho GTPase activating protein 15** | ARHGAP1 | S43 | **61.5** | **58.7** | K.S*MILTDVGK.V |
| **Rho GTPase activating protein 15** | ARHGAP1 | S212 | **12.0** | **51.1** | R.SSS*TELLSHYDSDIK.E |
| **Rho GTPase activating protein 15** | ARHGAP1 | S64 | **100.0** | **21.5** | R.NHS*QHILK.D |
| **Rho GTPase activating protein 15** | ARHGAP1 | T213 | **12.6** | **39.8** | R.SSST*ELLSHYDSDIK.E |
| **Rho GTPase activating protein 17** | ARHGAP1 | S598 | **18.6** | **61.1** | R.SPS*PPTQHTGQPPGQPSAPSQLSAPR.R |
| **Rho GTPase activating protein 17** | ARHGAP1 | S517 | | **32.7** | R.NNS*QIASGQNQPQAAAGSHQLSMGQPHNAAGPSPHTLR.R |
| **Rho GTPase activating protein 17** | ARHGAP1 | S596S598 | **40.0** | **46.0** | R.S*PS*PPTQHTGQPPGQPSAPSQLSAPR.R |
| **Rho GTPase activating protein 17** | ARHGAP1 | S596T601 | **11.6** | **33.7** | R.S*PSPPT*QHTGQPPGQPSAPSQLSAPR.R |
| **Rho GTPase activating protein 17** | ARHGAP1 | S547 | **15.6** | **31.0** | R.NNSQIASGQNQPQAAAGSHQLSMGQPHNAAGPS*PHTLR.R |
| **Rho GTPase activating protein 17** | ARHGAP1 | S598T601 | **16.7** | **44.4** | R.SPS*PPT*QHTGQPPGQPSAPSQLSAPR.R |
| **Rho GTPase activating protein 17** | ARHGAP1 | S598T604 | **3.4** | **39.2** | R.SPS*PPTQHT*GQPPGQPSAPSQLSAPR.R |
| **Rho GTPase activating protein 17** | ARHGAP1 | T604 | **24.1** | **35.2** | R.SPSPPTQHT*GQPPGQPSAPSQLSAPR.R |
| **Rho GTPase activating protein 17** | ARHGAP1 | S596T604 | **-1.9** | **16.7** | R.S*PSPPTQHT*GQPPGQPSAPSQLSAPR.R |
| **Rho GTPase activating protein 25** | ARHGAP2 | S487 | **13.8** | **28.8** | R.TMS*QDLR.Q |
| **Rho GTPase activating protein 30** | ARHGAP3 | S630 | **26.0** | **83.0** | K.GS*GSLEGEAAGCGR.Q |
| **Rho GTPase activating protein 30** | ARHGAP3 | S630S632 | **100.0** | **71.4** | K.GS*GS*LEGEAAGCGR.Q |
| **Rho GTPase activating protein 30** | ARHGAP3 | S875 | **17.0** | **76.9** | R.S*YAFETQANPGK.G |
| **Rho GTPase activating protein 30** | ARHGAP3 | S632 | **-0.1** | **86.0** | K.GSGS*LEGEAAGCGR.Q |
| **Rho GTPase activating protein 30** | ARHGAP3 | Y876 | **-0.3** | **79.5** | R.SY*AFETQANPGK.G |
| **Rho GTPase activating protein 30** | ARHGAP3 | S785 | **13.9** | **68.6** | R.NGGSLS*FDAAVALAR.D |
| **Rho GTPase activating protein 30** | ARHGAP3 | S291 | **100.0** | **33.6** | R.S*IFNLGR.S |
| **Rho GTPase activating protein 9** | ARHGAP9 | S456 | **54.2** | **65.9** | R.LSGSGPAELS*AGEDEEEESELVSKPLLR.L |
| **Rho guanine nucleotide exchange factor 1** | ARHGEF1 | S384S388 | **27.4** | **35.9** | R.SES*LRVS*DR.R |
| **Rho guanine nucleotide exchange factor 1** | ARHGEF1 | S25 | | **102.6** | R. [illegible] |
| **Rho guanine nucleotide exchange factor 1** | ARHGEF1 | S410 | **-0.4** | **62.3** | R.SRS*DVDMDPSSATAVLGPAR.R |
| **Rho guanine nucleotide exchange factor 1** | ARHGEF1 | S29 | **-0.5** | **26.5** | R. [illegible] |
| **Rho guanine nucleotide exchange factor 2** | ARHGEF2 | S174 | **32.1** | **82.2** | R.ILSQS*TDSLNMR.N |
| **Rho guanine nucleotide exchange factor 2** | ARHGEF2 | S172 | **13.4** | **98.4** | R.ILS*QSTDSLNMR.N |
| **Rho guanine nucleotide exchange factor 2** | ARHGEF2 | S122 | **25.6** | **52.2** | R.ERPSS*AIYPSDSFR.Q |
| **Rho guanine nucleotide exchange factor 2** | ARHGEF2 | S151 | **12.6** | **155.4** | K.SVS*TTNIAGHFNDESPLGLR.R |
| | ARHGEF2 | S151S163 | **33.2** | **56.4** | K.SVS*TTNIAGHFNDES*PLGLR.R |
| **Rho guanine nucleotide exchange factor 2** | ARHGEF2 | S885 | **66.7** | **125.0** | R.S*LPAGDALYLSFNPPQPSR.G |
| **Rho guanine nucleotide exchange factor 2** | ARHGEF2 | S955S959 | **100.0** | **22.4** | R.LS*PPHS*PR.D |
| **Rho guanine nucleotide exchange factor 2** | ARHGEF2 | S174S177 | **16.9** | **50.7** | R.ILSQS*TDS*LNMR.N |
| **Rho guanine nucleotide exchange factor 2** | ARHGEF2 | S163 | **17.6** | **21.9** | K.SVSTTNIAGHFNDES*PLGLR.R |
| **Rho guanine nucleotide exchange factor 2** | ARHGEF2 | Y893 | **25.4** | **66.2** | R.SLPAGDALY*LSFNPPQPSR.G |
| **Rho guanine nucleotide exchange factor 2** | ARHGEF2 | S174T175 | **13.0** | **48.2** | R.ILSQS*T*DSLNMR.N |
| **Rho guanine nucleotide exchange factor 2** | ARHGEF2 | S695 | **6.8** | **30.7** | R.EPALPLEPDSGGNTS*PGVTANGEAR.T |

Peak Area: <-10, -5, -3, 0, 3, 5, >10

%CV: 0, 29, 42, 71, 86, >100

**White dots-** Significant change in peptide abundance at 5 %FDR compared to the timepoint with the minimum peak area for a given PSM

CarT: 0m 2m 5m — RajiB: 0m 2m 5m

| Protein Name | Gene | Phosphosites | Ascor | MOWSE | Sequence |
|---|---|---|---|---|---|
| Rho guanine nucleotide exchange factor 2 | ARHGEF2 | S690 | 4.8 | 25.3 | R.EPALPLEPDS*GGNTSPGVTANGEAR.T |
| Rho guanine nucleotide exchange factor 2 | ARHGEF2 | S151T153 | 9.4 | 15.9 | K.SVS*TT*NIAGHFNDESPLGLRR.I |
| Rho guanine nucleotide exchange factor 2 | ARHGEF2 | T678 | 100.0 | 48.5 | K.DLLVGPGVELLLT*PR.E |
| Rho guanine nucleotide exchange factor 2 | ARHGEF2 | S172T175 | 4.1 | 11.3 | R.ILS*QST*DSLNMR.N |
| Rho guanine nucleotide exchange factor 2 | ARHGEF2 | S931 | 100.0 | 48.0 | R.QELGS*PEER.L |
| Rho guanine nucleotide exchange factor 2 | ARHGEF2 | S151T152 | 5.9 | 18.3 | K.SVS*T*TNIAGHFNDESPLGLRR.I |
| Rho guanine nucleotide exchange factor 2 | ARHGEF2 | S122S127 | 18.9 | 11.2 | R.ERPSS*AIYPS*DSFRQSLLGSR.R |
| Rho guanine nucleotide exchange factor 2 | ARHGEF2 | T152 | 8.0 | 88.2 | K.SVST*TNIAGHFNDESPLGLR.R |
| Rho guanine nucleotide exchange factor 2 | ARHGEF2 | T153S163 | 14.3 | 43.8 | K.SVSTT*NIAGHFNDES*PLGLR.R |
| Rho guanine nucleotide exchange factor 2 | ARHGEF2 | S121 | 41.9 | 18.6 | R.ERPS*SAIYPSDSFR.Q |
| Rho guanine nucleotide exchange factor 2 | ARHGEF2 | S172S177 | 10.0 | 21.2 | R.ILS*QSTDS*LNMR.N |
| Rho guanine nucleotide exchange factor 38 | ARHGEF3 | S553 | 100.0 | 17.6 | R.KLS*FEKK.K |
| Rho guanine nucleotide exchange factor 6 | ARHGEF6 | S640 | 49.2 | 25.6 | R.KPS*EEEYVIR.K |
| Rho guanine nucleotide exchange factor 6 | ARHGEF7 | S488 | 45.8 | 46.7 | R.MS*GFIYQGK.I |
| Rho guanine nucleotide exchange factor 6 | ARHGEF6 | T133 | 45.7 | 105.0 | R.SSSLSAANTSQT*NPQGAVSSTVSGLQR.Q |
| Rho guanine nucleotide exchange factor 6 | ARHGEF6 | S684 | 100.0 | 85.0 | R.KDS*IPQVLLPEEEK.L |
| Rho guanine nucleotide exchange factor 6 | ARHGEF7 | S649 | 11.0 | 54.4 | K.S*TAALEEDAQILK.V |
| Rho guanine nucleotide exchange factor 6 | ARHGEF6 | S123 | 5.3 | 76.4 | R.SS*SLSAANTSQTNPQGAVSSTVSGLQR.Q |
| Rho guanine nucleotide exchange factor 6 | ARHGEF6 | S124 | 7.9 | 47.1 | R.SSS*LSAANTSQTNPQGAVSSTVSGLQR.Q |
| Rho guanine nucleotide exchange factor 6 | ARHGEF6 | S124S126 | 8.4 | 87.9 | R.SSS*LS*AANTSQTNPQGAVSSTVSGLQR.Q |
| Rho guanine nucleotide exchange factor 6 | ARHGEF6 | S124T133 | 4.2 | 72.0 | R.SSS*LSAANTSQT*NPQGAVSSTVSGLQR.Q |
| Rho guanine nucleotide exchange factor 6 | ARHGEF7 | T650 | -0.4 | 34.9 | K.ST*AALEEDAQILK.V |
| Rho guanine nucleotide exchange factor 6 | ARHGEF6 | S126 | 14.9 | 205.3 | R.SSSLS*AANTSQTNPQGAVSSTVSGLQR.Q |
| Rho guanine nucleotide exchange factor 6 | ARHGEF6 | S124T130 | 7.7 | 84.1 | R.SSS*LSAANT*SQTNPQGAVSSTVSGLQR.Q |
| Rho guanine nucleotide exchange factor 6 | ARHGEF6 | S126T133 | 1.0 | 71.8 | R.SSSLS*AANTSQT*NPQGAVSSTVSGLQR.Q |
| Rho guanine nucleotide exchange factor 7 | HIVEP3 | S739S740S742 | 100.0 | 13.0 | R.RS*S*LS*R.L |
| Rho guanine nucleotide exchange factor 7 | HIVEP3 | S739S740 | 28.6 | 15.5 | R.RS*S*LSR.L |
| Rho guanine nucleotide exchange factor 7 | HIVEP3 | S739 | 14.9 | 24.9 | R.RS*SLSR.L |
| Rho guanine nucleotide exchange factor 7 | HIVEP3 | S739S742 | 10.3 | 13.3 | R.RS*SLS*R.L |
| Rho specific guanine nucleotide exchange factor p114 | ARHGEF1 | S994S1002 | 18.6 | 26.1 | K.AGGTALLPGPPAPS*PLPATPLS*AK.E |
| Rho specific guanine nucleotide exchange factor p114 | ARHGEF1 | S943S945 | | 21.8 | R.S*LS*PILPGRHSPAPPPDPGFPAPSPPPADSPSEGFSLK.A |
| Rho specific guanine nucleotide exchange factor p114 | ARHGEF1 | S943S945S953 | | 15.3 | R.S*LS*PILPGRHS*PAPPPDPGFPAPSPPPADSPSEGFSLK.A |
| Rho specific guanine nucleotide exchange factor p114 | ARHGEF1 | S1002 | 6.9 | 43.9 | K.AGGTALLPGPPAPSPLPATPLS*AK.E |
| Rho specific guanine nucleotide exchange factor p114 | ARHGEF1 | T999S1002 | 28.3 | 26.7 | K.AGGTALLPGPPAPSPLPAT*PLS*AK.E |
| Rho specific guanine nucleotide exchange factor p114 | ARHGEF1 | S945 | 16.9 | 21.6 | R.SLS*PILPGR.H |
| Rho specific guanine nucleotide exchange factor p114 | ARHGEF1 | T999 | 31.1 | 21.9 | K.AGGTALLPGPPAPSPLPAT*PLSAK.E |
| RhoGAP1 | ARHGAP1 | S51 | 12.6 | 51.0 | K.SSS*PELVTHLK.W |
| Ribonuclease P, 29 KD | POP4 | S38T39 | 100.0 | 30.9 | R.RS*T*PR.S |
| Ribonucleotide reductase M2 subunit | RRM2 | S20 | 137.3 | 71.7 | R.VPLAPITDPQQLQLS*PLK.G |
| Ribosomal L1 domain containing 1 | RSL1D1 | S361 | 25.8 | 80.4 | K.ATNES*EDEIPQLVPIGK.K |
| Ribosomal L1 domain containing 1 | RSL1D1 | T358 | 6.1 | 31.3 | K.AT*NESEDEIPQLVPIGK.K |
| Ribosomal phosphoprotein large P2 | RPLP2 | S102S105 | 100.0 | 68.8 | K.DEKKEES*EES*DDDMGFGLFD.- |
| Ribosomal phosphoprotein large P2 | RPLP2 | S17 | 7.6 | 43.9 | R.YVASYLLAALGGNSS*PSAK.D |
| Ribosomal phosphoprotein large P2 | RPLP2 | S16 | 13.9 | 70.5 | R.YVASYLLAALGGNS*SPSAK.D |
| Ribosomal protein L14 | RPL14 | S139 | 100.0 | 47.3 | K.AALLKAS*PK.K |
| Ribosomal protein L17 | RPL17- | S142 | 14.9 | 45.1 | R.INPYMSS*PCHIEMILTEK.E |
| Ribosomal protein L17 | RPL17- | S141 | 8.5 | 40.2 | R.INPYMS*SPCHIEMILTEK.E |
| Ribosomal protein L23a | RPL23A | T42 | 8.0 | 18.4 | K.IRT*SPTFR.R |
| Ribosomal protein L23a | RPL23A | S43 | 8.9 | 22.0 | K.IRTS*PTFR.R |
| Ribosomal protein L31 | RPL31 | S98 | 15.5 | 55.1 | R.NEDEDS*PNKLYTLVTYVPVTTFK.N |
| Ribosomal protein L31 | RPL31 | Y103 | 5.5 | 33.7 | R.NEDEDSPNKLY*TLVTYVPVTTFK.N |
| Ribosomal protein S17 | RPS17 | S115 | -0.3 | 71.7 | K.LLDFGSLS*NLQVTQPTVGMNFK.T |
| Ribosomal protein S17 | RPS17 | S113 | 9.1 | 68.1 | K.LLDFGS*LSNLQVTQPTVGMNFK.T |
| Ribosomal protein S2 | RPS2 | S264 | 17.2 | 50.7 | K.S*PYQEFTDHLVK.T |
| Ribosomal protein S3 | RPS3 | T221 | 13.9 | 41.8 | K.DEILPTT*PISEQK.G |
| Ribosomal protein S3 | RPS3 | T220 | 25.4 | 31.3 | K.DEILPT*TPISEQK.G |
| Ribosomal protein S3a | RPS3A | S263 | 102.5 | 36.0 | R.ADGYEPPVQES*V.- |
| Ribosomal protein S3a | RPS3A | S237 | 12.1 | 28.8 | K.LMELHGEGS*SGK.A |
| Ribosomal protein S3a | RPS3A | S238 | 9.8 | 44.4 | K.LMELHGEGSS*GK.A |
| | RPS6 | S236S240 | 17.7 | 43.2 | R.RLSS*LRAS*TSK.S |
| Ribosomal protein S6 | RPS6 | S235S236 | 100.0 | 25.2 | R.RRLS*S*LR.A |
| Ribosomal protein S6 | RPS6 | S235 | 8.9 | 23.6 | R.RRLS*SLR.A |

Peak Area: <-10, -5, -3, 0, 3, 5, >10

%CV: 0, 29, 42, 71, 86, >100

**White dots-** Significant change in peptide abundance at **5** %FDR compared to the timepoint with the minimum peak area for a given PSM

CarT: 0m, 2m, 5m — RajiB: 0m, 2m, 5m

| Protein Name | Gene | Phosphosites | Ascor | MOWSE | Sequence |
|---|---|---|---|---|---|
| **Ribosomal protein S6** | RPS6 | S236S240T241 | **0.7** | **12.3** | R.RLSS*LRAS*T*SK.S |
| **Ribosomal protein S6** | RPS6 | S236T241 | **11.0** | **41.1** | R.LSS*LRAST*SK.S |
| **Ribosomal protein S6** | RPS6 | S235S236S240 | **1.1** | **12.6** | R.RLS*S*LRAS*TSK.S |
| **Ribosomal protein S6** | RPS6 | S236 | **9.8** | **26.7** | R.RRLSS*LR.A |
| **Ribosomal protein S6 kinase alpha 3** | RPS6KA1 | Y226 | **8.8** | **66.2** | K.AY*SFCGTVEYMAPEVVNR.Q |
| **Ribosomal protein S6 kinase alpha 3** | RPS6KA1 | T231 | **-0.2** | **26.0** | K.AYSFCGT*VEYMAPEVVNR.Q |
| **Ribosomal protein S6 kinase alpha 3** | RPS6KA3 | S369 | **43.9** | **25.6** | K.DS*PGIPPSANAHQLFR.G |
| **Ribosomal protein S6 kinase alpha 3** | RPS6KA1 | S227 | **5.8** | **31.8** | K.AYS*FCGTVEYMAPEVVNR.Q |
| **Ribosomal protein S6 kinase alpha 5** | RPS6KA5 | Y375S376 | **19.9** | **34.4** | K.LFQGY*S*FVAPSILFK.R |
| **Ribosomal protein S6 kinase alpha 5** | RPS6KA5 | Y375S381 | **12.6** | **55.7** | K.LFQGY*SFVAPS*ILFK.R |
| **Ribosomal protein S6 kinase, 70kDa, polypeptide 1** | RPS6KB1 | T444S447 | **75.3** | **79.5** | R.T*PVS*PVKFSPGDFWGR.G |
| **Ribosomal protein S6 kinase, 70kDa, polypeptide 1** | RPS6KB1 | S447S452 | **30.3** | **11.0** | R.TPVS*PVKFS*PGDFWGR.G |
| **Ribosomal protein, large, P0** | RPLP0 | S307 | **28.1** | **49.7** | K.EESEES*DEDMGFGLFD.- |
| **Ribosomal protein, large, P0** | RPLP0 | S304 | **16.1** | **41.8** | K.EES*EESDEDMGFGLFD.- |
| **Ribosomal protein, large, P0** | RPLP0 | S304S307 | **100.0** | **53.2** | K.EES*EES*DEDM#GFGLFD.- |
| | RPLP1 | S101 | **24.6** | **74.8** | K.KEES*EESDDDMGFGLFD.- |
| | RPLP1 | S101S104 | **100.0** | **69.2** | K.KEES*EES*DDDMGFGLFD.- |
| | RPLP1 | S104 | **22.8** | **27.1** | K.EESEES*DDDMGFGLFD.- |
| **Ribosomal RNA processing 12 homolog** | RRP12 | S1080 | **40.6** | **91.3** | K.GDSIEEILADS*EDEEDNEEEER.S |
| **Ribosomal RNA processing 12 homolog** | RRP12 | S1072 | **5.2** | **23.9** | K.GDS*IEEILADSEDEEDNEEEER.S |
| **Rictor** | RICTOR | S1385Y1386 | **21.8** | **72.1** | K.ALS*Y*ASLDKEDLLSPINQNTLQR.S |
| **Rictor** | RICTOR | S1385S1388 | **16.2** | **43.5** | K.ALS*YAS*LDKEDLLSPINQNTLQR.S |
| **Rictor** | RICTOR | S1282 | **13.1** | **56.1** | K.S*NSVSLVPPGSSHTLPR.R |
| **RIMS1** | RIMS1 | S989T991 | **21.6** | **17.0** | R.RSRS*PT*R.H |
| **Ring Finger Protein 1** | RING1 | S163T166T167 | | **94.3** | R. [illegible] |
| **Ring Finger Protein 1** | RING1 | T238T243 | | **24.2** | R.GGT*LGGGT*LGPPSPPGAPSPPEPGGEIELVFRPHPLLVEK.G |
| **Ring Finger Protein 1** | RING1 | S170S187S188 | **12.9** | **40.1** | R. [illegible] |
| **Ring Finger Protein 1** | RING1 | S248S254 | **69.3** | **12.1** | R.GGTLGGGTLGPPS*PPGAPS*PPEPGGEIELVFRPHPLLVEK.G |
| **Ring Finger Protein 1** | RING1 | S163T166T167 | | **16.8** | R. [illegible] |
| **Ring finger protein 126** | RNF126 | S301 | **12.2** | **42.0** | K.SLTGQNTATNPPGLTGVSFSSSSSSSSSS*PSNENATSNS.- |
| **Ring finger protein 126** | RNF126 | S297 | **3.7** | **22.9** | K.SLTGQNTATNPPGLTGVSFSSSSSSS*SSSSPSNENATSNS.- |
| **Ring finger protein 126** | RNF126 | S303 | **8.0** | **19.9** | K.SLTGQNTATNPPGLTGVSFSSSSSSSSSSSPS*NENATSNS.- |
| **Ring finger protein 126** | RNF126 | S298 | **5.9** | **23.8** | K.SLTGQNTATNPPGLTGVSFSSSSSSSS*SSSPSNENATSNS.- |
| **Ring finger protein 146** | RNF146 | T288S290S293 | **6.0** | **64.0** | R.VPAPDTSIEET*ES*DAS*SDSEDVSAVVAQHSLTQQR.L |
| **Ring finger protein 169** | RNF169 | S403 | **100.0** | **49.4** | R.VLS*PLIIK.S |
| **Ring finger protein 2** | RNF2 | T197T199 | | **10.9** | R.T*KT*SDDSGLELDNNNAAMAIDPVMDGASEIELVFRPHPTLMEK.[illegible] |
| **RING finger protein 20** | RNF20 | S138 | **7.8** | **17.6** | K.ALVVPEPEPDSDS*NQER.K |
| **RING finger protein 29** | TRIM55 | S292 | **37.7** | **40.8** | K.KISEAS*K.A |
| **Ring finger protein 4** | RNF4 | S94S95 | **36.1** | **50.3** | R.RLPQDHADSCVVS*S*DDEELSR.D |
| **RIO kinase 1** | RIOK1 | S22 | **7.0** | **71.6** | R.VVPGQFDDADSS*DSENR.D |
| **RIO kinase 1** | RIOK1 | S21 | **22.0** | **42.0** | R.VVPGQFDDADS*SDSENR.D |
| **RIO kinase 2** | RIOK2 | S380S382S385 | **22.3** | **58.5** | R.SSGDPEQIKEDS*LS*EES*ADAR.S |
| **RIO kinase 2** | RIOK2 | S337 | **10.2** | **43.8** | K.EGSEFSFS*DGEVAEK.A |
| **RIO kinase 2** | RIOK2 | S442 | **52.5** | **50.8** | R.VQGGVPAGS*DEYEDECPHLIALSSLNR.E |
| **RIO kinase 2** | RIOK2 | S332S335S337 | **100.0** | **80.6** | K.EGS*EFS*FS*DGEVAEK.A |
| **RIO kinase 2** | RIOK2 | S390 | **71.4** | **99.8** | R.S*FEMTEFNQALEEIK.G |
| **RIO kinase 2** | RIOK2 | S370S382S385 | **13.8** | **24.2** | R.SS*GDPEQIKEDSLS*EES*ADAR.S |
| **RIO kinase 2** | RIOK2 | S335 | **13.4** | **52.7** | K.EGSEFS*FSDGEVAEK.A |
| **RIO kinase 2** | RIOK2 | S380S382 | **13.2** | **24.0** | R.SSGDPEQIKEDS*LS*EESADAR.S |
| **RIO kinase 2** | RIOK2 | Y445 | **16.4** | **15.1** | R.VQGGVPAGSDEY*EDECPHLIALSSLNR.E |
| **RIP2** | RIPK2 | S393 | **24.5** | **65.9** | K.LHHCPGNHS*WDSTISGSQR.A |
| **RIP2** | RIPK2 | S527 | **29.8** | **56.0** | K.QMGLQPYPEILVVS*R.S |
| **RIP2** | RIPK2 | S363 | **56.8** | **28.2** | R.S*LPAPQDNDFLSR.K |
| **RIP2** | RIPK2 | S176 | **35.4** | **20.8** | R.MMSLS*QSR.S |
| **RIP2** | RIPK2 | S531 | **1.4** | **56.9** | R.SPS*LNLLQNK.S |
| **RIP2** | RIPK2 | S396 | **13.7** | **61.8** | K.LHHCPGNHSWDS*TISGSQR.A |
| **RNA binding motif protein 10** | RBM10 | S89 | **42.3** | **42.2** | R.HRHS*PTGPPGFPR.D |
| **RNA binding motif protein 10** | RBM10 | S736S738 | **36.7** | **88.9** | R.GLVAAYSGES*DS*EEEQER.G |
| **RNA binding motif protein 10** | RBM10 | S733S736S738 | **35.1** | **89.3** | R.GLVAAYS*GES*DS*EEEQER.G |
| **RNA binding motif protein 10** | RBM5 | S661 | **100.0** | **13.0** | R.S*LNKQK.E |
| **RNA binding motif protein 10** | RBM10 | S733S738 | **22.1** | **30.5** | R.GLVAAYS*GESDS*EEEQER.G |
| **RNA binding motif protein 10** | RBM10 | Y732S736S738 | **17.6** | **62.0** | R.GLVAAY*SGES*DS*EEEQER.G |

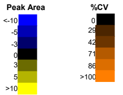


**White dots-** Significant change in peptide abundance at **5** %FDR compared to the timepoint with the minimum peak area for a given PSM

| Protein Name | Gene | Phosphosites | Ascor | MOWSE | Sequence |
|---|---|---|---|---|---|
| **RNA binding motif protein 10** | RBM10 | T91 | **10.9** | **19.0** | R.HRHSPT*GPPGFPR.D |
| **RNA binding motif protein 10** | RBM10 | Y732S733S738 | **5.6** | **26.7** | R.GLVAAY*S*GESDS*EEEQER.G |
| **RNA binding motif protein 12** | RBM12 | S411 | **12.2** | **20.1** | R.S*KSPSGQKR.S |
| **RNA binding motif protein 12** | RBM12 | S413 | **39.8** | **32.1** | R.SKS*PSGQKR.S |
| **RNA binding motif protein 13** | MAK16 | S199S200 | **1.9** | **80.6** | K.ALEQQEAESDS*S*DTEEKDDDDDDEEDVGKR.E |
| **RNA binding motif protein 13** | MAK16 | S197S199 | **22.9** | **57.0** | K.ALEQQEAES*DS*SDTEEKDDDDDDEEDVGKR.E |
| **RNA binding motif protein 13** | MAK16 | S197S199S200 | **20.1** | **55.9** | K.ALEQQEAES*DS*S*DTEEKDDDDDDEEDVGKR.E |
| **RNA binding motif protein 13** | MAK16 | S229S232 | **100.0** | **82.9** | R.EFVEDGEVDES*DIS*DFEDMDK.L |
| **RNA binding motif protein 13** | MAK16 | S197S199S200 | **100.0** | **58.1** | K.ALEQQEAES*DS*S*DT*EEKDDDDDDEEDVGKR.E |
| **RNA binding motif protein 13** | MAK16 | S197S200 | **8.3** | **53.5** | K.ALEQQEAES*DSS*DTEEKDDDDDDEEDVGKR.E |
| **RNA binding motif protein 13** | MAK16 | S199T202 | **7.8** | **19.8** | K.ALEQQEAESDS*SDT*EEKDDDDDDEEDVGKR.E |
| **RNA binding motif protein 13** | MAK16 | S199S200T202 | **-0.2** | **19.7** | K.ALEQQEAESDS*S*DT*EEKDDDDDDEEDVGKR.E |
| **RNA binding motif protein 14** | RBM14 | S520 | **28.6** | **93.7** | R.TQS*SASLAASYAAQQHPQAAASYR.G |
| **RNA binding motif protein 14** | RBM14 | T572 | **19.1** | **92.0** | R.GQPGNAYDGAGQPSAAYLSMSQGAVANANST*PPPYER.T |
| **RNA binding motif protein 14** | RBM14 | S555 | **5.3** | **18.5** | R.GQPGNAYDGAGQPS*AAYLSMSQGAVANANSTPPPYER.T |
| **RNA binding motif protein 14** | RBM14 | S291 | | **78.9** | R. [illegible] |
| **RNA binding motif protein 14** | RBM14 | S582 | **32.3** | **28.0** | R.TRLS*PPR.A |
| **RNA binding motif protein 14** | RBM14 | S520S521 | **10.6** | **90.7** | R.TQS*S*ASLAASYAAQQHPQAAASYR.G |
| **RNA binding motif protein 14** | RBM14 | T206 | **100.0** | **30.4** | R.QPT*PPFFGR.D |
| **RNA binding motif protein 14** | RBM14 | S571 | **33.7** | **62.4** | R.GQPGNAYDGAGQPSAAYLSMSQGAVANANS*TPPPYER.T |
| **RNA binding motif protein 14** | RBM14 | S618 | **17.0** | **21.9** | R.RLS*ESQLSFR.R |
| **RNA binding motif protein 14** | RBM14 | Y548 | | **39.5** | R.GQPGNAY*DGAGQPSAAYLSMSQGAVANANSTPPPYER.T |
| **RNA binding motif protein 14** | RBM14 | T579 | **14.9** | **20.9** | R.T*RLSPPR.A |
| **RNA binding motif protein 15** | RBM15 | T568 | **56.6** | **29.9** | R.DRT*PPLLYR.D |
| **RNA binding motif protein 15** | RBM15 | T265Y266 | **16.2** | **61.5** | R.SRSPLDKDT*Y*PPSASVVGASVGGHR.H |
| **RNA binding motif protein 15** | RBM15 | S257Y266 | **5.1** | **59.4** | R.S*RSPLDKDTY*PPSASVVGASVGGHR.H |
| **RNA binding motif protein 15** | RBM15 | S259T265 | **4.9** | **59.9** | R.SRS*PLDKDT*YPPSASVVGASVGGHR.H |
| **RNA binding motif protein 15** | RBM15 | S292 | **10.6** | **60.5** | R.S*LSPGGAALGYR.D |
| **RNA binding motif protein 15** | RBM15 | S51 | **100.0** | **13.3** | K.ERS*PVK.A |
| **RNA binding motif protein 15** | RBM15 | S708 | **100.0** | **16.4** | R.RGS*LEK.S |
| **RNA binding motif protein 15** | RBM15 | S259Y266 | **6.8** | **45.7** | R.SRS*PLDKDTY*PPSASVVGASVGGHR.H |
| **RNA binding motif protein 15** | RBM15 | S294 | **16.7** | **69.7** | R.SLS*PGGAALGYR.D |
| **RNA binding motif protein 15** | RBM15 | S670S674 | **39.0** | **18.1** | R.HCAPS*PDRS*PELSSSR.D |
| **RNA binding motif protein 15** | RBM15 | S257S259 | **2.8** | **36.7** | R.S*RS*PLDKDTYPPSASVVGASVGGHR.H |
| **RNA binding motif protein 15** | RBM15 | S259 | **3.9** | **60.7** | R.S*PLDKDTYPPSASVVGASVGGHR.H |
| **RNA binding motif protein 15** | RBM15 | T265 | **2.0** | **18.9** | R.SPLDKDT*YPPSASVVGASVGGHR.H |
| **RNA binding motif protein 15** | RBM15 | S700 | **100.0** | **17.3** | R.LLLERPS*PIRDR.R |
| **RNA binding motif protein 15** | RBM15 | S622 | **100.0** | **19.0** | R.DGWS*LDR.D |
| **RNA binding motif protein 15** | RBM15 | S257T265 | **7.0** | **42.4** | R.S*RSPLDKDT*YPPSASVVGASVGGHR.H |
| **RNA binding motif protein 15** | RBM15 | S656 | **12.2** | **31.0** | R.S*PESDRPR.K |
| **RNA binding motif protein 19** | RBM19 | S949T957 | **9.9** | **29.1** | R.SVVLDEILEQLEGS*DSDSEEQT*LQL.- |
| **RNA binding motif protein 19** | RBM19 | S953T957 | **13.4** | **19.2** | R.SVVLDEILEQLEGSDSDS*EEQT*LQL.- |
| **RNA binding motif protein 19** | RBM19 | S949S951 | **6.2** | **20.1** | R.SVVLDEILEQLEGS*DS*DSEEQTLQL.- |
| **RNA binding motif protein 19** | RBM19 | S949S953T957 | **19.5** | **17.3** | R.SVVLDEILEQLEGS*DSDS*EEQT*LQL.- |
| **RNA binding motif protein 25** | RBM25 | S677 | **55.2** | **101.9** | K.LGASNS*PGQPNSVK.R |
| **RNA binding motif protein 25** | RBM25 | S703 | **100.0** | **60.8** | K.FEDEDS*DDVPR.K |
| | RBM33 | S41 | **206.3** | **153.6** | R.AADEDWDS*ELEDDLLGEDLLSGK.K |
| **RNA binding motif protein 33** | RBM33 | S205 | **119.4** | **79.0** | K.EES*DEEEEDDEESGR.L |
| **RNA binding motif protein 34** | RBM34 | S9 | **100.0** | **57.4** | R.S*VQEGENPDDGVR.G |
| **RNA binding motif protein 5** | RBM5 | S624 | **39.4** | **130.0** | R.GLVAAYSGDS*DNEEELVER.L |
| **RNA binding motif protein 6** | RBM6 | S362 | **20.3** | **73.6** | R.EGETQGVAFEHESPADFQNSQS*PVQDQDK.S |
| **RNA binding motif protein 6** | RBM6 | Y908 | **17.6** | **66.7** | K.VVNPLIGLLGEY*GGDSDYEEEEEEEQTPPPQPR.T |
| **RNA binding motif protein 6** | RBM6 | Y914 | **5.5** | **48.4** | K.VVNPLIGLLGEYGGDSDY*EEEEEEEQTPPPQPR.T |
| **RNA binding motif protein 6** | RBM6 | S360 | **7.3** | **22.5** | R.EGETQGVAFEHESPADFQNS*QSPVQDQDK.S |
| **RNA binding motif protein 6** | RBM6 | T923 | **-0.0** | **19.4** | K.VVNPLIGLLGEYGGDSDYEEEEEEEQT*PPPQPR.T |
| **RNA binding motif protein 6** | RBM6 | S912 | **7.0** | **41.6** | K.VVNPLIGLLGEYGGDS*DYEEEEEEEQTPPPQPR.T |
| **RNA binding motif protein 7** | RBM7 | S136 | **16.9** | **14.9** | R.SFS*SPENFQR.Q |
| **RNA binding motif protein 8A** | RBM8A | S42 | **41.9** | **30.3** | R.GFGS*EEGSR.A |
| **RNA binding motif protein 8A** | RBM8A | S166S168 | **100.0** | **15.6** | R.S*RS*PDRR.R |
| **RNA binding protein 4** | RBM4 | S309 | **100.0** | **16.6** | R.DRS*PLRR.S |
| **RNA binding protein 4** | RBM4 | S86 | **10.9** | **38.8** | K.LHVGNIS*PTCTNK.E |
| **RNA binding protein S1, serine-rich domain** | RNPS1 | S27 | **22.6** | **28.3** | R.APS*PTKR.K |

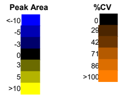


**White dots-** Significant change in peptide abundance at **5** %FDR compared to the timepoint with the minimum peak area for a given PSM

| Protein Name | Gene | Phosphosites | Ascor | MOWSE | Sequence |
|---|---|---|---|---|---|
| RNA binding protein S1, serine-rich domain | RNPS1 | S139S141 | 100.0 | 15.0 | R.S*KS*KPPKR.D |
| RNA binding protein S1, serine-rich domain | RNPS1 | S141 | 100.0 | 20.4 | K.S*KPPKRDEK.E |
| | RNPS1 | S155S157 | 20.9 | 11.0 | K.RRS*PS*PKPTK.V |
| RNA polymerase I transcription factor RRN3 | RRN3 | S170S172 | 76.4 | 41.9 | K.EGDVDVS*DS*DDEDDNLPANFDTCHR.A |
| RNA polymerase II associated protein 3 | RPAP3 | S116S119S121 | 21.7 | 34.3 | R.ILDELDKDDSTHESLS*QES*ES*EEDGIHVDSQK.A |
| RNA polymerase II elongation factor ELL | ELL | S437 | 0.6 | 21.3 | R.LGLPLLTDCAQPS*RPHGSPSR.S |
| RNA polymerase II elongation factor ELL | ELL | S442 | 12.2 | 14.0 | R.LGLPLLTDCAQPSRPHGS*PSR.S |
| RNA polymerase IIA 220 kd subunit | POLR2A | S1913T1919 | 14.1 | 19.9 | K.YSPTS*PTYSPT*SPK.Y |
| RNA polymerase IIA 220 kd subunit | POLR2A | S1962 | 19.7 | 27.4 | K.GSTYSPTSPGYSPTSPTYSLTSPAIS*PDDSDEEN.- |
| RNA polymerase IIA 220 kd subunit | POLR2A | S1913S1920 | 15.2 | 15.9 | K.YSPTS*PTYSPTS*PK.Y |
| RNA polymerase IIA 220 kd subunit | POLR2A | Y1909T1919 | 5.5 | 21.1 | K.Y*SPTSPTYSPT*SPK.Y |
| RNA polymerase III subunit 5 | POLR3E | S161S162 | 100.0 | 92.7 | R.EAANEAGDS*S*QDEAEDDVK.Q |
| RNA polymerase-associated protein RTF1 homolog | RTF1 | S53S58S60 | 14.6 | 36.1 | R.VVIDS*DTEDS*GS*DENLDQELLSLAK.R |
| RNA polymerase-associated protein RTF1 homolog | RTF1 | S53T55S58S60 | 74.9 | 80.5 | R.VVIDS*DT*EDS*GS*DENLDQELLSLAK.R |
| RNA polymerase-associated protein RTF1 homolog | RTF1 | T55S58S60 | 1.7 | 29.4 | R.VVIDSDT*EDS*GS*DENLDQELLSLAK.R |
| RNA polymerase-associated protein RTF1 homolog | RTF1 | S53T55S58 | 10.2 | 72.7 | R.VVIDS*DT*EDS*GSDENLDQELLSLAK.R |
| RNA U3 small nucleolar interacting protein 2 | RRP9 | S50S51S53 | 37.1 | 108.6 | K.MNEEIS*S*DS*ESESLAPR.K |
| RNA U3 small nucleolar interacting protein 2 | RRP9 | S50S51S53S55 | 14.0 | 46.0 | K.MNEEIS*S*DS*ES*ESLAPR.K |
| RNA U3 small nucleolar interacting protein 2 | RRP9 | S50S51S53S57 | 15.8 | 58.1 | K.MNEEIS*S*DS*ESES*LAPR.K |
| RNA-binding protein 7 | RBM7 | S107 | 12.2 | 22.8 | R.SGSSHAPQDVSLSYPQHHVGNS*SPTSTSPSSR.Y |
| RNA-binding protein Raly | RALY | S288 | 29.3 | 52.6 | R.TRDDGDEEGLLTHS*EEELEHSQDTDADDGALQ.- |
| RNA-binding protein Raly | RALY | S288S295 | 22.8 | 40.0 | R.TRDDGDEEGLLTHS*EEELEHS*QDTDADDGALQ.- |
| RNA-binding protein Raly | RALY | T286S295T298 | 28.3 | 39.7 | R.TRDDGDEEGLLT*HSEEELEHS*QDT*DADDGALQ.- |
| RNA-binding protein Raly | RALY | T298 | 27.8 | 52.0 | R.TRDDGDEEGLLTHSEEELEHSQDT*DADDGALQ.- |
| RNA-binding protein Raly | RALY | T286S288S295 | 55.5 | 34.1 | R.TRDDGDEEGLLT*HS*EEELEHS*QDT*DADDGALQ.- |
| RNA-binding protein Raly | RALY | S295T298 | 10.5 | 37.8 | R.TRDDGDEEGLLTHSEEELEHS*QDT*DADDGALQ.- |
| RNA-binding protein Raly | RALY | T286S288 | 20.3 | 33.2 | R.TRDDGDEEGLLT*HS*EEELEHSQDTDADDGALQ.- |
| RNA-binding protein Raly | RALY | T286T298 | 13.5 | 38.6 | R.TRDDGDEEGLLT*HSEEELEHSQDT*DADDGALQ.- |
| | RALY | S135 | 100.0 | 20.9 | R.LS*PVPVPR.A |
| RNA-binding protein Raly | RALY | S288S295T298 | 7.9 | 18.3 | R.TRDDGDEEGLLTHS*EEELEHS*QDT*DADDGALQ.- |
| RNA-binding protein Raly | RALY | S288T298 | 15.6 | 34.6 | R.TRDDGDEEGLLTHS*EEELEHSQDT*DADDGALQ.- |
| RNA-binding protein Raly | RALY | S106 | 28.6 | 69.6 | R.AAS*AIYSGYIFDYDYYR.D |
| RNA-binding protein Raly | RALY | S295 | 2.5 | 11.5 | R.TRDDGDEEGLLTHSEEELEHS*QDTDADDGALQ.- |
| RNA-binding protein Raly | RALY | T286S295 | 9.3 | 17.1 | R.TRDDGDEEGLLT*HSEEELEHS*QDTDADDGALQ.- |
| Round spermatid basic protein 1-like | RSBN1L | Y80S96 | 15.7 | 25.9 | R.QLQPPAAPSPQSY*GSPASWSFAPLSAAPS*PSSSR.S |
| RUN domain containing 2A | SNX29 | S344 | 114.7 | 97.4 | K.S*IDDEDVDENEDDVYGNSSGR.K |
| RY1 | SNRNP27 | S59T60S61 | 5.9 | 16.0 | R.S*T*S*PSPSR.L |
| Ryanodine receptor 1 | RYR1 | T952 | 100.0 | 21.9 | K.KT*KLPK.T |
| S phase kinase associated protein 2 | SKP2 | S64 | 30.2 | 28.9 | K.EEPDSENIPQELLSNLGHPES*PPR.K |
| SAFB-like, transcription modulator | SLTM | S553 | 116.5 | 94.6 | K.S*PGHMVILDQTK.G |
| SAFB-like, transcription modulator | SLTM | S551S553 | 17.8 | 130.8 | R.ISS*KS*PGHMVILDQTK.G |
| SAFB-like, transcription modulator | SLTM | S550S553 | 61.8 | 64.1 | R.IS*SKS*PGHMVILDQTK.G |
| SAFB-like, transcription modulator | SLTM | S289 | 39.1 | 34.2 | K.DGQDAIAQS*PEKESK.D |
| SAFB-like, transcription modulator | SLTM | S550S551 | 68.5 | 53.4 | R.IS*S*KSPGHMVILDQTK.G |
| SAFB-like, transcription modulator | SLTM | S1002 | -0.2 | 36.9 | R.AGAGMITQHSSNAS*PINR.I |
| Salt inducible kinase 2 | SIK2 | S587 | 44.7 | 56.0 | R.RAS*DTSLTQGIVAFR.Q |
| Salt inducible kinase 2 | SIK2 | S534 | 95.4 | 56.9 | R.DLNFLEDNPS*LKDIMLANQPSPR.M |
| SAM domain and HD domain containing protein 1 | SAMHD1 | S33 | 16.7 | 58.2 | R.TPSNTPSAEADWS*PGLELHPDYK.T |
| SAM domain and HD domain containing protein 1 | SAMHD1 | T25 | 1.8 | 63.0 | R.TPSNT*PSAEADWSPGLELHPDYK.T |
| SAM domain and HD domain containing protein 1 | SAMHD1 | T21S33 | 16.3 | 61.9 | R.T*PSNTPSAEADWS*PGLELHPDYK.T |
| SAM domain and HD domain containing protein 1 | SAMHD1 | S27 | -0.5 | 55.3 | R.TPSNTPS*AEADWSPGLELHPDYK.T |
| SAM domain and HD domain containing protein 1 | SAMHD1 | T25S33 | 5.4 | 23.0 | R.TPSNT*PSAEADWS*PGLELHPDYK.T |
| SAM domain and HD domain containing protein 1 | SAMHD1 | S23S33 | 4.9 | 35.5 | R.TPS*NTPSAEADWS*PGLELHPDYK.T |
| SAM domain and HD domain containing protein 1 | SAMHD1 | S23 | -0.9 | 15.0 | R.TPS*NTPSAEADWSPGLELHPDYK.T |
| SAM domain protein SAMSN1 | SAMSN1 | S134 | 13.9 | 89.9 | K.ASDSMDSLYSGQS*SSSGITSCSDGTSNR.D |
| SAM domain protein SAMSN1 | SAMSN1 | S23 | 16.9 | 59.2 | R.SSS*FGNFDR.F |
| SAM domain protein SAMSN1 | SAMSN1 | S125S128 | 1.6 | 75.8 | K.ASDS*MDS*LYSGQSSSSGITSCSDGTSNR.D |
| SAM domain protein SAMSN1 | SAMSN1 | S125 | -0.5 | 75.4 | K.ASDS*MDSLYSGQSSSSGITSCSDGTSNR.D |
| SAM domain protein SAMSN1 | SAMSN1 | Y130 | 5.2 | 71.2 | K.ASDSMDSLY*SGQSSSSGITSCSDGTSNR.D |
| SAM domain protein SAMSN1 | SAMSN1 | S134S135 | 9.1 | 48.2 | K.ASDSMDSLYSGQS*S*SSGITSCSDGTSNR.D |
| SAM domain protein SAMSN1 | SAMSN1 | S22 | -0.5 | 40.9 | R.SS*SFGNFDR.F |
| SAM domain protein SAMSN1 | SAMSN1 | S135 | 13.9 | 15.3 | K.ASDSMDSLYSGQSS*SSGITSCSDGTSNR.D |

Peak Area: <-10, -5, -3, 0, 3, 5, >10

%CV: 0, 29, 42, 71, 86, >100

**White dots-** Significant change in peptide abundance at 5 %FDR compared to the timepoint with the minimum peak area for a given PSM

| Protein Name | Gene | Phosphosites | CarT (0m 2m 5m) | RajiB (0m 2m 5m) | Ascor | MOWSE | Sequence |
|---|---|---|---|---|---|---|---|
| SAM domain protein SAMSN1 | SAMSN1 | S123S125 | | | 3.3 | 44.9 | K.AS*DS*MDSLYSGQSSSSGITSCSDGTSNR.D |
| SAM domain protein SAMSN1 | SAMSN1 | S134S136 | | | 6.3 | 66.8 | K.ASDSMDSLYSGQS*SS*SGITSCSDGTSNR.D |
| SAM domain protein SAMSN1 | SAMSN1 | S131 | | | 10.0 | 44.6 | K.ASDSMDSLYS*GQSSSSGITSCSDGTSNR.D |
| Sam68 | KHDRBS1 | S58 | | | | 30.2 | R.AS*PATQPPPLLPPSATGPDATVGGPAPTPLLPPSATASVK.M |
| Sam68 | KHDRBS1 | S20 | | | 9.1 | 36.5 | R.SGS*MDPSGAHPSVR.Q |
| Sam68 | KHDRBS1 | T61 | | | 5.9 | 24.5 | R.ASPAT*QPPPLLPPSATGPDATVGGPAPTPLLPPSATASVK.M |
| Sam68 | KHDRBS1 | S18 | | | 10.1 | 15.7 | R.S*GSMDPSGAHPSVR.Q |
| SAP30-like | SAP30L | S93S99 | | | 22.0 | 72.0 | R.KTS*DDGGDS*PEHDTDIPEVDLFQLQVNTLR.R |
| SAP30-like | SAP30L | S93T104 | | | 7.1 | 31.5 | R.KTS*DDGGDSPEHDT*DIPEVDLFQLQVNTLR.R |
| SAP30-like | SAP30L | T92T104 | | | 2.4 | 39.1 | R.KT*SDDGGDSPEHDT*DIPEVDLFQLQVNTLR.R |
| SAP30-like | SAP30L | T92S99 | | | 4.8 | 22.6 | R.KT*SDDGGDS*PEHDTDIPEVDLFQLQVNTLR.R |
| SAPK substrate protein 1 | UBXN1 | S200 | | | 9.2 | 13.0 | K.YGGSVGSQPPPVAPEPGPVPSS*PSQEPPTKR.E |
| Sarcoma antigen NY-SAR-91 | MEAF6 | S118T120S122 | | | | 12.8 | K.REPGS*GT*ES*DTSPDFHNQENEPSQEDPEDLDGSVQGVKPQK.^ |
| Sarcoma antigen NY-SAR-91 | MEAF6 | S118T120T124 | | | 3.3 | 20.7 | K.REPGS*GT*ESDT*SPDFHNQENEPSQEDPEDLDGSVQGVKPQK.^ |
| Sarcoma antigen NY-SAR-91 | MEAF6 | S118S122 | | | 2.2 | 18.4 | K.REPGS*GTES*DTSPDFHNQENEPSQEDPEDLDGSVQGVKPQK.A |
| Sarcoma antigen NY-SAR-91 | MEAF6 | S118S125 | | | -1.0 | 13.4 | R.EPGS*GTESDTS*PDFHNQENEPSQEDPEDLDGSVQGVKPQK.A |
| Sarcoma antigen NY-SAR-91 | MEAF6 | S122T124S125 | | | 13.6 | 25.9 | K.REPGSGTES*DT*S*PDFHNQENEPSQEDPEDLDGSVQGVKPQK.^ |
| Sarcoma antigen NY-SAR-91 | MEAF6 | S118T120S125 | | | 0.0 | 27.9 | K.REPGS*GT*ESDTS*PDFHNQENEPSQEDPEDLDGSVQGVKPQK.^ |
| SART1 | SART1 | S448 | | | 112.6 | 30.1 | R.RVS*EVEEEKEPVPQPLPSDDTR.V |
| SART1 | SART1 | S474S486 | | | 100.0 | 72.2 | R.VENMDIS*DEEEGGAPPPGS*PQVLEEDEAELELQK.Q |
| SART1 | SART1 | S596S607T608 | | | 11.8 | 25.0 | R.SANGGS*ESDGEENIGWS*T*VNLDEEK.Q |
| SART1 | SART1 | S596S598S607 | | | 17.2 | 33.3 | R.SANGGS*ES*DGEENIGWS*TVNLDEEK.Q |
| SART1 | SART1 | S591S596S598 | | | 26.3 | 37.7 | R.S*ANGGS*ES*DGEENIGWSTVNLDEEK.Q |
| SART1 | SART1 | S598S607T608 | | | 8.8 | 12.6 | R.SANGGSES*DGEENIGWS*T*VNLDEEK.Q |
| Scaffold Attachment Factor B | SAFB | S604 | | | 44.5 | 32.5 | R.SVVS*FDK.V |
| Scaffold Attachment Factor B | SAFB2 | T307 | | | | 97.7 | R.T*DCEPVGLEPAVEQSSAASELAEASSEELAEAPTEAPSPEAR.D |
| Scaffold Attachment Factor B | SAFB2 | S344 | | | 18.0 | 101.6 | R.TDCEPVGLEPAVEQSSAASELAEASSEELAEAPTEAPS*PEAR.D |
| Scaffold Attachment Factor B | SAFB | S197 | | | 10.4 | 69.4 | K.ETINNLDTSSS*DFTILQEIEEPSLEPENEK.I |
| Scaffold Attachment Factor B | SAFB | S195 | | | 29.0 | 79.0 | K.ETINNLDTS*SSDFTILQEIEEPSLEPENEK.I |
| Scaffold Attachment Factor B | SAFB | S195S196 | | | 10.6 | 46.4 | K.ETINNLDTS*S*SDFTILQEIEEPSLEPENEK.I |
| Scaffold Attachment Factor B | SAFB | T194S195S197 | | | 13.2 | 42.5 | K.ETINNLDT*S*SS*DFTILQEIEEPSLEPENEK.I |
| Scaffold Attachment Factor B | SAFB | S195S196S197 | | | 9.5 | 36.1 | K.ETINNLDTS*S*S*DFTILQEIEEPSLEPENEK.I |
| Scaffold Attachment Factor B | SAFB | S373S384 | | | 14.1 | 58.1 | K.ESS*TSEGADQKMSS*PEDDSDTKR.L |
| Scaffold Attachment Factor B | SAFB2 | T340 | | | 6.8 | 35.3 | R.TDCEPVGLEPAVEQSSAASELAEASSEELAEAPT*EAPSPEAR.D |
| Scaffold Attachment Factor B | SAFB | S196 | | | 8.0 | 69.1 | K.ETINNLDTSS*SDFTILQEIEEPSLEPENEK.I |
| Scaffold Attachment Factor B | SAFB | T374S383 | | | 0.4 | 40.4 | K.ESST*SEGADQKMS*SPEDDSDTKR.L |
| Scaffold Attachment Factor B | SAFB | T194 | | | | 16.9 | K.ELPEQLQEHAIEDKETINNLDT*SSSDFTILQEIEEPSLEPENEK.I |
| Scaffold Attachment Factor B | SAFB | S373S383 | | | 13.7 | 47.9 | K.ESS*TSEGADQKMS*SPEDDSDTKR.L |
| Scaffold Attachment Factor B | SAFB | T194S195 | | | 7.5 | 49.1 | K.ETINNLDT*S*SSDFTILQEIEEPSLEPENEK.I |
| Scaffold Attachment Factor B | SAFB | T374S384 | | | 10.8 | 52.8 | K.ESST*SEGADQKMSS*PEDDSDTKR.L |
| Scaffold Attachment Factor B | SAFB | S375S383 | | | 4.5 | 25.5 | K.ESSTS*EGADQKMS*SPEDDSDTKR.L |
| Scaffold Attachment Factor B | SAFB | T194S195S196 | | | -1.1 | 27.8 | K.ETINNLDT*S*S*SDFTILQEIEEPSLEPENEK.I |
| Scaffold Attachment Factor B | SAFB | S373S375 | | | 0.7 | 31.4 | K.ESS*TS*EGADQKMSSPEDDSDTKR.L |
| Scaffold Attachment Factor B | SAFB | S195T200 | | | 11.3 | 23.0 | K.ETINNLDTS*SSDFT*ILQEIEEPSLEPENEK.I |
| Scaffold Attachment Factor B | SAFB | S375S384 | | | 7.1 | 21.1 | K.ESSTS*EGADQKMSS*PEDDSDTKR.L |
| Scaffold Attachment Factor B | SAFB2 | T245 | | | 31.6 | 25.4 | K.SEPVKEESSELEQPFAQDT*SSVGPDRK.L |
| Scaffold Attachment Factor B | SAFB2 | S321S344 | | | 10.2 | 16.7 | R.TDCEPVGLEPAVEQS*SAASELAEASSEELAEAPTEAPS*PEAR.D |
| Scaffold Attachment Factor B | SAFB | S195S197 | | | 4.8 | 43.4 | K.ETINNLDTS*SS*DFTILQEIEEPSLEPENEK.I |
| Scaffold Attachment Factor B | SAFB | S197T200 | | | 7.5 | 46.8 | K.ETINNLDTSSS*DFT*ILQEIEEPSLEPENEK.I |
| Scaffold Attachment Factor B | SAFB | T188 | | | | 32.8 | K.ET*INNLDTSSSDFTILQEIEEPSLEPENEK.I |
| Scaffold Attachment Factor B | SAFB | T200 | | | 1.2 | 12.4 | K.ETINNLDTSSSDFT*ILQEIEEPSLEPENEK.I |
| Scaffold Attachment Factor B | SAFB | T194S195T200 | | | 3.7 | 33.4 | K.ETINNLDT*S*SSDFT*ILQEIEEPSLEPENEK.I |
| Scaffold Attachment Factor B | SAFB | T188S195S197 | | | 8.7 | 11.1 | K.ET*INNLDTS*SS*DFTILQEIEEPSLEPENEK.I |
| Scaffold attachment factor B2 | SAFB2 | S109 | | | 59.8 | 45.9 | K.MEEEGTEDNGLEDDS*R.D |
| Scaffold attachment factor B2 | SAFB2 | S832 | | | 100.0 | 15.9 | K.RLS*EGR.G |
| SCC112 protein | PDS5A | S1265 | | | 100.0 | 73.9 | R.AAVGQES*PGGLEAGNAK.A |
| Scribble | SCRIB | S688T689 | | | 96.4 | 90.1 | R.AEEEEAS*T*EEEDKEGAVVSAPSVK.G |
| Scribble | SCRIB | S1220 | | | 57.3 | 52.2 | R.NS*LESISSIDR.E |
| Scribble | SCRIB | S1559S1566 | | | 13.2 | 32.7 | R.LAEAPSPAPTPSPTPVEDLGPQTS*TSPGRLS*PDFAEELR.S |
| Scribble | SCRIB | T1558S1566 | | | 6.0 | 31.9 | R.LAEAPSPAPTPSPTPVEDLGPQT*STSPGRLS*PDFAEELR.S |
| Scribble | SCRIB | T1342S1348 | | | 30.4 | 37.3 | R.AFAAVPTSHPPEDAPAQPPT*PGPAAS*PEQLSFR.E |
| Scribble | SCRIB | T1560S1566 | | | 17.9 | 32.0 | R.LAEAPSPAPTPSPTPVEDLGPQTST*SPGRLS*PDFAEELR.S |

Peak Area: <-10, -5, -3, 0, 3, 5, >10

%CV: 0, 29, 42, 71, 86, >100

**White dots-** Significant change in peptide abundance at **5** %FDR compared to the timepoint with the minimum peak area for a given PSM

| Protein Name | Gene | Phosphosites | CarT 0m 2m 5m | RajiB 0m 2m 5m | Ascor | MOWSE | Sequence |
|---|---|---|---|---|---|---|---|
| Scribble | SCRIB | T1549S1566 | | | 7.2 | 11.4 | R.LAEAPSPAPTPSPT*PVEDLGPQTSTSPGRLS*PDFAEELR.S |
| Scribble | SCRIB | S1547S1566 | | | 8.0 | 21.0 | R.LAEAPSPAPTPS*PTPVEDLGPQTSTSPGRLS*PDFAEELR.S |
| SCY1 like 1 | SCYL1 | S651 | | | 7.2 | 87.5 | R.WDDEDWGS*LEQEAESVLAQQDDWSTGGQVSR.A |
| SCY1-like 2 | SCYL2 | T679 | | | 10.9 | 28.4 | K.RASLT*LEEK.Q |
| SCY1-like 2 | SCYL2 | S677 | | | 20.1 | 35.3 | K.RAS*LTLEEK.Q |
| SDAD1 | SDAD1 | S525 | | | 84.9 | 73.3 | K.YIEIDS*DEEPR.G |
| SDS3 | SUDS3 | S234S236 | | | 47.5 | 37.3 | K.RPAS*PS*SPEHLPATPAESPAQR.F |
| SDS3 | SUDS3 | S236S237 | | | 8.2 | 30.6 | K.RPASPS*S*PEHLPATPAESPAQR.F |
| SDS3 | SUDS3 | S45T49T55 | | | 19.0 | 24.7 | R.GRES*DEDT*EDASET*DLAKHDEEDYVEMK.E |
| SDS3 | SUDS3 | S45T49S53 | | | 24.3 | 31.4 | R.GRES*DEDT*EDAS*ETDLAK.H |
| SEC16 homolog A | SEC16A | S587S595T600 | | | 13.3 | 35.4 | R.GS*VSQPSTPS*PPKPT*GIFQTSANSSFEPVK.S |
| SEC16 homolog A | SEC16A | S587T593 | | | 5.8 | 36.5 | R.GS*VSQPST*PSPPKPTGIFQTSANSSFEPVK.S |
| SEC16 homolog A | SEC16A | S587T593S595 | | | 9.1 | 20.6 | R.GS*VSQPST*PS*PPKPTGIFQTSANSSFEPVK.S |
| SEC16 homolog A | SEC16A | S342 | | | 100.0 | 18.3 | R.GDS*PENR.T |
| SEC16 homolog A | SEC16A | S589S592 | | | 8.5 | 74.3 | R.GSVS*QPS*TPSPPKPTGIFQTSANSSFEPVK.S |
| SEC16 homolog A | SEC16A | T197S201 | | | | 36.1 | R. QNPHDCVVT*PAAS*PSLPQPGLQMPGQWGPVQGGPQPSGQHR.S |
| SEC16 homolog A | SEC16A | S592T593S595 | | | 18.7 | 44.0 | R.GSVSQPS*T*PS*PPKPTGIFQTSANSSFEPVK.S |
| SEC16 homolog A | SEC16A | S589S592T593 | | | 13.4 | 36.0 | R.GSVS*QPS*T*PS*PPKPTGIFQTSANSSFEPVK.S |
| SEC16 homolog A | SEC16A | S592T593 | | | 19.7 | 55.2 | R.GSVSQPS*T*PSPPKPTGIFQTSANSSFEPVK.S |
| SEC16 homolog A | SEC16A | S589T593S595 | | | 16.0 | 43.6 | R.GSVS*QPST*PS*PPKPTGIFQTSANSSFEPVK.S |
| SEC16 homolog A | SEC16A | S589S592S595 | | | 14.0 | 77.2 | R.GSVS*QPS*TPS*PPKPTGIFQTSANSSFEPVK.S |
| SEC16 homolog A | SEC16A | S587S592S595 | | | 12.2 | 41.6 | R.GS*VSQPS*TPS*PPKPTGIFQTSANSSFEPVK.S |
| SEC16 homolog A | SEC16A | S587S592T600 | | | 10.7 | 29.3 | R.GS*VSQPS*TPSPPKPT*GIFQTSANSSFEPVK.S |
| SEC16 homolog A | SEC16A | S2022 | | | 68.3 | 61.2 | R.S*PDPGIVPQEAPVGNSLSELSEENFDGK.F |
| SEC16 homolog A | SEC16A | S587S592T593 | | | 13.9 | 35.8 | R.GS*VSQPS*T*PS*PPKPTGIFQTSANSSFEPVK.S |
| SEC16 homolog A | SEC16A | S587T593T600 | | | 10.3 | 44.2 | R.GS*VSQPST*PSPPKPT*GIFQTSANSSFEPVK.S |
| SEC16 homolog A | SEC16A | S587S589T593 | | | 11.3 | 33.4 | R.GS*VS*QPST*PS*PPKPTGIFQTSANSSFEPVK.S |
| SEC22B | SEC22B | S137 | | | 46.9 | 104.7 | R.NLGS*INTELQDVQR.I |
| SEC31 like 1 | SEC31A | T1161 | | | 30.5 | 64.9 | R.EQT*LSPTITSGLHNIAR.S |
| SEC31 like 1 | SEC31A | S527 | | | 17.1 | 54.9 | K.DSDQVAQS*DGEESPAAEEQLLGEHIK.E |
| SEC31 like 1 | SEC31A | S1163 | | | 10.7 | 57.6 | R.EQTLS*PTITSGLHNIAR.S |
| SEC31 like 1 | SEC31A | S527S532 | | | 21.5 | 13.9 | K.DSDQVAQS*DGEES*PAAEEQLLGEHIK.E |
| SEC63 | SEC63 | S593S597 | | | 100.0 | 39.7 | K.DDGS*DRDS*DREQDEK.Q |
| Second step splicing factor 1 | PPAN- | S359 | | | 79.1 | 121.0 | R.VGGS*DEEASGIPSR.T |
| Second step splicing factor 1 | PPAN- | S238S240 | | | 4.8 | 78.9 | R.LQDISELLATGAGLS*ES*EAEPDGDHNITELPQAVAGR.G |
| Second step splicing factor 1 | PPAN- | S228T233 | | | 15.0 | 96.9 | R.LQDIS*ELLAT*GAGLSESEAEPDGDHNITELPQAVAGR.G |
| Second step splicing factor 1 | PPAN- | S238 | | | 17.2 | 64.0 | R.LQDISELLATGAGLS*ESEAEPDGDHNITELPQAVAGR.G |
| Second step splicing factor 1 | PPAN- | T233 | | | 5.4 | 69.1 | R.LQDISELLAT*GAGLSESEAEPDGDHNITELPQAVAGR.G |
| Second step splicing factor 1 | PPAN- | T233S240 | | | 20.2 | 69.8 | R.LQDISELLAT*GAGLSES*EAEPDGDHNITELPQAVAGR.G |
| Second step splicing factor 1 | PPAN- | T233S238 | | | 26.1 | 49.8 | R.LQDISELLAT*GAGLS*ESEAEPDGDHNITELPQAVAGR.G |
| Secreted frizzled related protein 2 | SFRP2 | S287S289 | | | 100.0 | 16.9 | K.RIS*RS*IR.K |
| Secreted frizzled related protein 2 | SFRP2 | S289 | | | 29.8 | 23.4 | K.RISRS*IR.K |
| Secretory carrier membrane protein 2 | SCAMP2 | S320 | | | -0.2 | 59.5 | R.AASS*AAQGAFQGN.- |
| Secretory carrier membrane protein 2 | SCAMP2 | S319 | | | | 53.6 | R.AAS*SAAQGAFQGN.- |
| Secretory carrier membrane protein 3 | SCAMP3 | S32 | | | 111.4 | 36.7 | R.DGGNPFAEPSELDNPFQDPAVIQHRPS*R.Q |
| Secretory carrier membrane protein 3 | SCAMP3 | S76 | | | 25.8 | 36.3 | R.KLS*PTEPK.N |
| Selectin P ligand | SELPLG | S358 | | | 7.2 | 70.9 | R.NYS*PTEMVCISSLLPDGGEGPSATANGGLSK.A |
| Selectin P ligand | SELPLG | Y357 | | | 5.7 | 33.4 | R.NY*SPTEMVCISSLLPDGGEGPSATANGGLSK.A |
| Selectin P ligand | SELPLG | T360 | | | 9.1 | 31.2 | R.NYSPT*EMVCISSLLPDGGEGPSATANGGLSK.A |
| Senataxin | SETX | S1017S1019 | | | 100.0 | 82.5 | R.GQVIIIS*DS*DDDDDER.I |
| Senescence downregulated leo1 like | LEO1 | T188S197 | | | 100.0 | 31.5 | K.MQNT*DDEERPQLS*DDER.Q |
| Senescence downregulated leo1 like | LEO1 | S212S220S229 | | | 100.0 | 35.1 | K.ANS*DDERPVAS*DNDDEKQNS*DDEEQPQLS*DEEK.M |
| Senescence downregulated leo1 like | LEO1 | S294S296S300 | | | 47.4 | 77.9 | K.NAIAS*DS*EADS*DTEVPK.D |
| Senescence downregulated leo1 like | LEO1 | S658 | | | 12.3 | 44.8 | K.YVIS*DEEEEDDD.- |
| Senescence downregulated leo1 like | LEO1 | S171S179 | | | 100.0 | 90.0 | R.AQGS*DEDKLQNS*DDDEK.M |
| Senescence downregulated leo1 like | LEO1 | S151S154S162 | | | 100.0 | 31.8 | R.EDKS*DQS*DDEKIQNS*DDEER.A |
| Senescence downregulated leo1 like | LEO1 | S171S179T188 | | | 100.0 | 21.7 | R.AQGS*DEDKLQNS*DDDEKMQNT*DDEERPQLS*DDER.Q |
| Senescence downregulated leo1 like | LEO1 | S271S273S277 | | | 100.0 | 15.1 | K.S*ES*ARGS*DS*EDEVLR.M |
| Senescence downregulated leo1 like | LEO1 | S162 | | | 100.0 | 40.2 | K.IQNS*DDEER.A |
| Senescence downregulated leo1 like | LEO1 | S205 | | | 100.0 | 40.4 | R.QQLS*EEEK.A |
| Senescence downregulated leo1 like | LEO1 | S66S72S75S77 | | | 100.0 | 13.9 | K.ELFGDDS*EDEGAS*HHS*GS*DNHS*ER.S |
| Senescence downregulated leo1 like | LEO1 | S84S88S91 | | | 100.0 | 17.5 | R.S*DNRS*EAS*ER.S |

**Peak Area**: <-10, -5, -3, 0, 3, 5, >10

**%CV**: 0, 29, 42, 71, 86, >100

**White dots-** Significant change in peptide abundance at **5** %FDR compared to the timepoint with the minimum peak area for a given PSM

| Protein Name | Gene | Phosphosites | CarT 0m 2m 5m | RajiB 0m 2m 5m | Ascor | MOWSE | Sequence |
|---|---|---|---|---|---|---|---|
| **Sentrin specific protease 3** | SENP3 | S232 | | | **14.8** | **60.2** | K.S*PLDPDSGLLSCTLPNGFGGQSGPEGER.S |
| **Sentrin specific protease 3** | SENP3 | S169 | | | **67.4** | **51.4** | K.NHLS*PQQGGATPQVPSPCCR.F |
| **Sentrin specific protease 7** | SENP7 | S443S444 | | | **87.9** | **85.2** | R.GHNEGNQSLISAEPIVVS*S*DEEGPVEHK.S |
| | SEPT1 | S206 | | | **19.2** | **22.9** | K.EEEIHIYQFPECDS*DEDEDFKRQDAEMK.E |
| **septin 1** | SEPT1 | S248 | | | **10.4** | **63.8** | R.YS*WGTVEVENPHHCDFLNLR.R |
| **septin 1** | SEPT1 | S315 | | | **84.3** | **75.4** | R.QS*ATEIPLPMLPLADTEK.L |
| **septin 1** | SEPT1 | T251 | | | **15.4** | **43.3** | R.RYSWGT*VEVENPHHCDFLNLR.R |
| **septin 1** | SEPT1 | T317 | | | **21.2** | **40.9** | R.QSAT*EIPLPMLPLADTEK.L |
| **septin 1** | SEPT1 | Y247 | | | **7.8** | **33.1** | R.RY*SWGTVEVENPHHCDFLNLR.R |
| **Septin 2** | SEPT2 | S218 | | | **63.9** | **109.8** | K.IYHLPDAES*DEDEDFKEQTR.L |
| **Septin 5** | SEPT5 | S225 | | | **72.9** | **72.1** | K.FGIHVYQFPECDS*DEDEDFKQQDR.E |
| **Sequestosome 1** | SQSTM1 | S365 | | | **16.5** | **35.6** | K.EVDPSTGELQSLQMPESEGPS*SLDPSQEGPTGLK.E |
| **Sequestosome 1** | SQSTM1 | S366 | | | **9.0** | **33.8** | K.EVDPSTGELQSLQMPESEGPSS*LDPSQEGPTGLK.E |
| **Serine arginine repetitive matrix protein 1** | SRRM1 | S597 | | | **27.6** | **31.3** | R.RYS*PPIQR.R |
| **Serine arginine repetitive matrix protein 1** | SRRM1 | T220 | | | **40.7** | **36.9** | K.EKT*PELPEPSVK.V |
| **Serine arginine repetitive matrix protein 1** | SRRM1 | S874 | | | **35.4** | **149.9** | K.KETES*EAEDNLDDLEK.H |
| | SRRM1 | S769S775 | | | **45.6** | **60.2** | K.KPPAPPS*PVQSQS*PSTNWSPAVPVKK.A |
| | SRRM1 | S769S777 | | | **52.8** | **65.9** | K.KPPAPPS*PVQSQSPS*TNWSPAVPVKK.A |
| **Serine arginine repetitive matrix protein 1** | SRRM1 | S769S775S781 | | | **22.4** | **29.2** | K.KPPAPPS*PVQSQS*PSTNWS*PAVPVKK.A |
| **Serine arginine repetitive matrix protein 1** | SRRM1 | S769S777S781 | | | **14.3** | **53.6** | K.KPPAPPS*PVQSQSPS*TNWS*PAVPVKK.A |
| **Serine arginine repetitive matrix protein 1** | SRRM1 | T872S874 | | | **100.0** | **108.3** | K.KET*ES*EAEDNLDDLEK.H |
| **Serine arginine repetitive matrix protein 1** | SRRM1 | S769S777T778 | | | **8.2** | **13.9** | K.KPPAPPS*PVQSQSPS*T*NWSPAVPVKK.A |
| **Serine arginine repetitive matrix protein 1** | SRRM1 | S769S773S781 | | | **25.4** | **29.1** | K.KPPAPPS*PVQS*QSPSTNWS*PAVPVKK.A |
| **Serine arginine repetitive matrix protein 1** | SRRM1 | S769S773S775 | | | **31.7** | **36.1** | K.KPPAPPS*PVQS*QS*PSTNWSPAVPVKK.A |
| **Serine arginine repetitive matrix protein 1** | SRRM1 | S769S773S775 | | | **53.6** | **33.5** | K.KPPAPPS*PVQS*QS*PSTNWS*PAVPVKK.A |
| **Serine arginine repetitive matrix protein 1** | SRRM1 | S705S707 | | | **13.4** | **19.6** | R.RGAS*SS*PQR.R |
| **Serine arginine repetitive matrix protein 1** | SRRM1 | S524S526 | | | **100.0** | **27.1** | R.RHS*PS*R.S |
| **Serine arginine repetitive matrix protein 1** | SRRM1 | S748 | | | **9.8** | **26.4** | R.RVS*SSR.S |
| **Serine arginine repetitive matrix protein 1** | SRRM1 | S541 | | | **7.6** | **31.5** | R.QKETS*PR.G |
| **Serine arginine repetitive matrix protein 1** | SRRM1 | T540 | | | **8.9** | **21.2** | R.QKET*SPR.G |
| **Serine arginine repetitive matrix protein 1** | SRRM1 | S260 | | | **100.0** | **44.9** | K.EPS*PEK.N |
| **Serine arginine repetitive matrix protein 1** | SRRM1 | S292S294 | | | **30.2** | **29.7** | R.S*PS*HTRPR.R |
| **Serine arginine repetitive matrix protein 1** | SRRM1 | S209S211 | | | **100.0** | **34.2** | R.S*PS*PAPEKK.E |
| **Serine arginine repetitive matrix protein 1** | SRRM1 | S207S209S211 | | | **100.0** | **21.0** | K.S*RS*PS*PAPEKK.E |
| | SRRM1 | S626S628 | | | **100.0** | **31.1** | R.AS*PS*PPPKR.R |
| **Serine arginine repetitive matrix protein 1** | SRRM1 | T411T416 | | | **14.2** | **15.1** | K.T*RHSPT*PQQSNR.T |
| | SRRM1 | S186S187 | | | **100.0** | **20.8** | R.RS*S*PVRR.E |
| **Serine arginine repetitive matrix protein 1** | SRRM1 | S683S685 | | | **100.0** | **19.2** | R.HS*PS*PRPR.A |
| **Serine arginine repetitive matrix protein 1** | SRRM1 | S904 | | | **16.0** | **40.8** | R.KAQVSPQS*.- |
| **Serine arginine repetitive matrix protein 1** | SRRM1 | S414 | | | **17.9** | **21.8** | K.TRHS*PTPQQSNR.T |
| **Serine arginine repetitive matrix protein 1** | SRRM1 | S316 | | | **100.0** | **23.1** | R.RRPS*PR.R |
| **Serine arginine repetitive matrix protein 1** | SRRM1 | S450S452 | | | **100.0** | **35.4** | K.RES*PS*PAPKPR.K |
| **Serine arginine repetitive matrix protein 1** | SRRM1 | S725T727 | | | **100.0** | **22.5** | R.RVS*RT*PEPK.K |
| **Serine arginine repetitive matrix protein 1** | SRRM1 | S369 | | | **100.0** | **20.7** | R.S*PPKKPPKR.T |
| **Serine arginine repetitive matrix protein 1** | SRRM1 | S549T555 | | | **17.4** | **20.8** | R.RRS*PSPPPT*R.R |
| **Serine arginine repetitive matrix protein 1** | USP24 | S653 | | | **26.1** | **54.4** | R.S*PSLSSK.H |
| **Serine arginine repetitive matrix protein 1** | SRRM1 | T572T574 | | | **100.0** | **37.0** | R.T*PT*PPPR.R |
| **Serine arginine repetitive matrix protein 1** | SRRM1 | S450 | | | **18.6** | **39.3** | K.RES*PSPAPKPR.K |
| | SRRM1 | T581S583 | | | **100.0** | **28.7** | R.RRT*PS*PPPR.R |
| | SRRM1 | S560S562 | | | **100.0** | **13.7** | R.RRS*PS*PAPPPR.R |
| **Serine arginine repetitive matrix protein 1** | SRRM1 | S653S655 | | | **14.2** | **20.3** | K.RRS*PS*LSSK.H |
| **Serine arginine repetitive matrix protein 1** | SRRM1 | S653S657 | | | **14.1** | **17.5** | K.RRS*PSLS*SK.H |
| | SRRM1 | T614S616 | | | **100.0** | **38.1** | R.T*AS*PPPPPKR.R |
| **Serine arginine repetitive matrix protein 1** | SRRM1 | S616 | | | **8.7** | **13.2** | R.TAS*PPPPPKR.R |
| **Serine arginine repetitive matrix protein 1** | SRRM1 | S752S754S756 | | | **100.0** | **43.9** | R.S*VS*GS*PEPAAK.K |
| **Serine arginine repetitive matrix protein 1** | SRRM1 | S431 | | | **26.0** | **47.4** | R.VSVS*PGR.T |
| **Serine arginine repetitive matrix protein 1** | SRRM1 | S738S740S743 | | | **100.0** | **26.2** | K.AAS*PS*PQS*VR.R |
| **Serine arginine repetitive matrix protein 1** | SRRM1 | S695 | | | **-0.3** | **26.7** | R.APQTSS*SPPPVRR.G |
| **Serine arginine repetitive matrix protein 1** | SRRM1 | S426S431 | | | **9.1** | **28.4** | R.KS*RVSVS*PGR.T |
| **Serine arginine repetitive matrix protein 1** | SRRM1 | S402T406 | | | **100.0** | **44.2** | R.HRPS*PPAT*PPPK.T |
| **Serine arginine repetitive matrix protein 1** | SRRM1 | S713S717 | | | **29.1** | **27.0** | R.RQS*PSPS*TRPIR.R |
| **Serine arginine repetitive matrix protein 1** | SRRM1 | S713T718 | | | **23.2** | **21.6** | R.RQS*PSPST*RPIR.R |

Peak Area: <-10, -5, -3, 0, 3, 5, >10

%CV: 0, 29, 42, 71, 86, >100

**White dots-** Significant change in peptide abundance at 5 %FDR compared to the timepoint with the minimum peak area for a given PSM

CarT: 0m 2m 5m — RajiB: 0m 2m 5m

| Protein Name | Gene | Phosphosites | Ascor | MOWSE | Sequence |
|---|---|---|---|---|---|
| Serine arginine repetitive matrix protein 1 | SRRM1 | S463S465 | 100.0 | 66.7 | R.KVELS*ES*EEDKGGK.M |
| Serine arginine repetitive matrix protein 1 | SRRM1 | S389S391 | 19.3 | 39.6 | R.RLS*PS*ASPPR.R |
| Serine arginine repetitive matrix protein 1 | SRRM1 | S389S391S393 | 100.0 | 24.9 | R.RLS*PS*AS*PPR.R |
| Serine arginine repetitive matrix protein 1 | SRRM1 | T872 | 7.3 | 42.9 | K.KET*ESEAEDNLDDLEK.H |
| Serine arginine repetitive matrix protein 1 | SRRM1 | S781 | 9.5 | 13.0 | K.KPPAPPSPVQSQSPSTNWS*PAVPVKK.A |
| Serine arginine repetitive matrix protein 1 | SRRM1 | S769S773T778 | 13.3 | 26.6 | K.KPPAPPS*PVQS*QSPST*NWSPAVPVKK.A |
| Serine arginine repetitive matrix protein 1 | SRRM1 | S901 | 12.2 | 46.1 | R.KAQVS*PQS.- |
| Serine arginine repetitive matrix protein 1 | SRRM1 | T416 | 18.4 | 12.6 | K.TRHSPT*PQQSNR.T |
| Serine arginine repetitive matrix protein 1 | USP24 | S655 | 2.7 | 54.2 | R.SPS*LSSK.H |
| Serine arginine repetitive matrix protein 1 | SRRM1 | S738 | 32.1 | 31.9 | K.AAS*PSPQSVR.R |
| Serine arginine repetitive matrix protein 1 | SRRM1 | S769 | 16.5 | 16.7 | K.KPPAPPS*PVQSQSPSTNWSPAVPVKK.A |
| Serine arginine repetitive matrix protein 1 | SRRM1 | S769T778 | 36.6 | 33.1 | K.KPPAPPS*PVQSQSPST*NWSPAVPVKK.A |
| Serine arginine repetitive matrix protein 1 | SRRM1 | S549S551 | 44.7 | 28.0 | R.RRS*PS*PPPTR.R |
| Serine arginine repetitive matrix protein 1 | SRRM1 | S426S429 | 10.5 | 25.4 | R.KS*RVS*VSPGR.T |
| Serine arginine repetitive matrix protein 1 | SRRM1 | S713S715 | 18.1 | 23.7 | R.RQS*PS*PSTRPIR.R |
| Serine arginine repetitive matrix protein 1 | SRRM1 | Y604S605 | 7.4 | 18.2 | R.RY*S*PSPPPK.R |
| Serine arginine repetitive matrix protein 1 | SRRM1 | S605S607 | 11.6 | 19.8 | R.RYS*PS*PPPK.R |
| Serine arginine repetitive matrix protein 1 | SRRM1 | S769S781 | 21.7 | 53.8 | K.KPPAPPS*PVQSQSPSTNWS*PAVPVK.K |
| Serine arginine repetitive matrix protein 1 | SRRM1 | S769S773 | 7.0 | 43.8 | K.KPPAPPS*PVQS*QSPSTNWSPAVPVK.K |
| Serine arginine repetitive matrix protein 1 | SRRM1 | S290S292 | 11.2 | 10.8 | R.S*RS*PSHTRPR.R |
| Serine arginine repetitive matrix protein 1 | SRRM1 | S715T718 | 16.0 | 17.4 | R.RQSPS*PST*RPIR.R |
| Serine arginine repetitive matrix protein 1 | SRRM1 | S528S530S532 | 100.0 | 18.1 | R.S*AS*PS*PR.K |
| Serine arginine repetitive matrix protein 1 | SRRM1 | S675 | 100.0 | 21.7 | R.S*PQPNKR.H |
| Serine arginine repetitive matrix protein 1 | SRRM1 | S754S756 | 17.3 | 32.4 | R.SVS*GS*PEPAAK.K |
| Serine arginine repetitive matrix protein 1 | SRRM1 | S773S775S781 | 17.6 | 34.2 | K.KPPAPPSPVQS*QS*PSTNWS*PAVPVKK.A |
| Serine arginine repetitive matrix protein 1 | SRRM1 | S769S775S777 | 16.0 | 29.8 | K.KPPAPPS*PVQSQS*PS*TNWSPAVPVKK.A |
| Serine arginine repetitive matrix protein 1 | SRRM1 | S294 | 3.9 | 31.0 | R.SPS*HTRPR.R |
| Serine arginine repetitive matrix protein 1 | SRRM1 | T693 | 7.2 | 22.9 | R.APQT*SSSPPPVRR.G |
| Serine arginine repetitive matrix protein 1 | SRRM1 | S463 | 23.9 | 29.3 | R.KVELS*ESEEDKGGK.M |
| Serine arginine repetitive matrix protein 1 | SRRM1 | S769S775T778 | 9.6 | 15.5 | K.KPPAPPS*PVQSQS*PST*NWSPAVPVKK.A |
| Serine arginine repetitive matrix protein 1 | SRRM1 | S696 | -0.4 | 27.9 | R.APQTSSS*PPPVRR.G |
| Serine arginine repetitive matrix protein 1 | SRRM1 | S738S740 | 10.5 | 12.4 | K.KAAS*PS*PQSVR.R |
| Serine arginine repetitive matrix protein 1 | SRRM1 | S429S431 | 0.4 | 24.3 | R.KSRVS*VS*PGR.T |
| Serine arginine repetitive matrix protein 1 | SRRM1 | S694 | 20.1 | 11.4 | R.APQTS*SSPPPVR.R |
| Serine arginine repetitive matrix protein 1 | SRRM1 | S713 | 12.2 | 24.2 | R.RQS*PSPSTRPIR.R |
| Serine arginine repetitive matrix protein 1 | SRRM1 | S769S773S777 | 14.0 | 36.1 | K.KPPAPPS*PVQS*QSPS*TNWSPAVPVKK.A |
| Serine arginine repetitive matrix protein 1 | SRRM1 | Y604S607 | 9.3 | 26.5 | R.RY*SPS*PPPK.R |
| Serine arginine repetitive matrix protein 1 | SRRM1 | S773 | 3.0 | 13.0 | K.KPPAPPSPVQS*QSPSTNWSPAVPVKK.A |
| Serine arginine repetitive matrix protein 1 | SRRM1 | S775S777 | 0.4 | 11.2 | K.KPPAPPSPVQSQS*PS*TNWSPAVPVKK.A |
| Serine arginine repetitive matrix protein 1 | SRRM1 | S769S773S775 | 12.8 | 21.1 | K.KPPAPPS*PVQS*QS*PS*TNWSPAVPVKK.A |
| Serine arginine repetitive matrix protein 1 | SRRM1 | S705S706 | | 22.1 | R.RGAS*S*SPQR.R |
| Serine arginine repetitive matrix protein 1 | SRRM1 | S706S707 | 3.3 | 17.0 | R.GASS*S*PQRR.Q |
| | SRRM1 | S414T416 | 20.6 | 23.5 | K.TRHS*PT*PQQSNR.T |
| Serine arginine repetitive matrix protein 1 | SRRM1 | S748S749 | | 14.5 | R.RVS*S*SR.S |
| Serine arginine repetitive matrix protein 1 | SPCS1 | S177S179S181 | 100.0 | 12.8 | R.S*RS*PS*PR.R |
| Serine arginine repetitive matrix protein 1 | SRRM1 | S389S393 | 11.3 | 37.4 | R.RLS*PSAS*PPR.R |
| Serine arginine rich pre-mRNA splicing factor SR-A1 | SCAF1 | S874 | 100.0 | 41.7 | R.S*PFLKPDER.A |
| Serine arginine rich pre-mRNA splicing factor SR-A1 | SCAF1 | S498S500 | 76.4 | 62.8 | R.S*PS*PAPAPAPAAAAGPPTR.K |
| | SCAF1 | S734S738 | 29.1 | 60.9 | R.EVLYDS*EGLS*GEER.G |
| Serine arginine rich pre-mRNA splicing factor SR-A1 | SCAF1 | Y238 | -0.3 | 14.1 | R.FDIYDPFHPTDEAY*SPPPAPEQK.Y |
| Serine arginine rich pre-mRNA splicing factor SR-A1 | SCAF1 | Y732S738 | 14.2 | 46.0 | R.EVLY*DSEGLS*GEER.G |
| Serine arginine rich pre-mRNA splicing factor SR-A1 | SCAF1 | T234 | 20.2 | 13.3 | R.FDIYDPFHPT*DEAYSPPPAPEQK.Y |
| Serine arginine rich pre-mRNA splicing factor SR-A1 | SCAF1 | S239 | 6.2 | 28.8 | R.FDIYDPFHPTDEAYS*PPPAPEQK.Y |
| Serine hydrolase-like protein 2 | SERHL2 | S103S113 | 25.1 | 17.0 | R.FS*ILGHSFALLS*PR.W |
| Serine protease 23 | PRSS23 | S138 | 100.0 | 22.0 | R.FS*IFGK.D |
| Serine threonine protein kinase N | PKN1 | S562 | 8.9 | 62.8 | K.LNLGTDSDSS*PQK.S |
| Serine threonine protein kinase N | PKN1 | S916 | 15.7 | 69.4 | R.TDVSNFDEEFTGEAPTLS*PPR.D |
| Serine threonine protein kinase N | PKN1 | T774 | 7.2 | 88.1 | R.TST*FCGTPEFLAPEVLTDTSYTR.A |
| Serine threonine protein kinase N | PKN1 | S773 | -0.3 | 61.3 | R.TS*TFCGTPEFLAPEVLTDTSYTR.A |
| Serine threonine protein kinase N | PKN1 | S578 | 5.8 | 43.9 | R.DPPSSPSSLS*SPIQESTAPELPSETQETPGPALCSPLR.K |
| Serine threonine protein kinase N | PKN1 | S573 | 19.1 | 65.5 | R.DPPSS*PSSLSSPIQESTAPELPSETQETPGPALCSPLR.K |
| Serine threonine protein kinase N | PKN1 | S575 | 7.3 | 36.6 | R.DPPSSPS*SLSSPIQESTAPELPSETQETPGPALCSPLR.K |

Peak Area: <-10, -5, -3, 0, 3, 5, >10

%CV: 0, 29, 42, 71, 86, >100

**White dots-** Significant change in peptide abundance at **5** %FDR compared to the timepoint with the minimum peak area for a given PSM

CarT: 0m 2m 5m — RajiB: 0m 2m 5m

| Protein Name | Gene | Phosphosites | Ascor | MOWSE | Sequence |
|---|---|---|---|---|---|
| Serine threonine protein kinase N | PKN1 | T778 | -0.8 | 20.0 | R.TSTFCGT*PEFLAPEVLTDTSYTR.A |
| Serine threonine protein kinase N | PKN1 | S561 | 17.8 | 37.3 | K.LNLGTDSDS*SPQK.S |
| Serine threonine protein kinase N | PKN1 | S572 | | 18.5 | R.DPPS*SPSSLSSPIQESTAPELPSETQETPGPALCSPLR.K |
| Serine/arginine repetitive matrix protein 1 | SRRM1 | S547 | 30.7 | 20.9 | R.WQS*PVTK.S |
| Serine/threonine kinase 24 (STE20 homolog) | STK24 | S315S316S317 | 6.8 | 32.2 | K.AEQSHDDS*S*S*EDSDAET*DGQASGGSDSGDWIFTIR.E |
| Serine/threonine kinase 24 (STE20 homolog) | STK24 | S316S317S320 | 12.3 | 87.9 | K.AEQSHDDSS*S*EDS*DAET*DGQASGGSDSGDWIFTIR.E |
| Serine/threonine kinase 24 (STE20 homolog) | STK24 | T190 | 12.7 | 18.0 | R.NT*FVGTPFWMAPEVIKQSAYDSK.A |
| Serine/threonine kinase 24 (STE20 homolog) | STK24 | S315S316S317 | 4.4 | 38.5 | K.AEQSHDDS*S*S*EDS*DAETDGQASGGSDSGDWIFTIR.E |
| Serine/threonine kinase 35 | STK35 | S281Y288 | 15.6 | 43.0 | K.YWLSS*ACGSDFY*MAPEVWEGHYTAK.A |
| Serine/threonine kinase 35 | STK35 | S280S281 | 14.1 | 60.3 | K.YWLS*S*ACGSDFYMAPEVWEGHYTAK.A |
| Serine/threonine kinase 35 | STK35 | S280S285 | 4.5 | 58.5 | K.YWLS*SACGS*DFYMAPEVWEGHYTAK.A |
| Serine/threonine kinase 35 | STK35 | S285Y288 | 20.4 | 56.4 | K.YWLSSACGS*DFY*MAPEVWEGHYTAK.A |
| Serine/threonine kinase 35 | STK35 | S281S285 | 1.8 | 34.4 | K.YWLSS*ACGS*DFYMAPEVWEGHYTAK.A |
| Serine/threonine kinase 35 | STK35 | S280Y288 | 5.1 | 32.7 | K.YWLS*SACGSDFY*MAPEVWEGHYTAK.A |
| Serine/threonine kinase 38 like | STK38L | Y281 | 70.1 | 66.3 | R.QLAY*STVGTPDYIAPEVFMQTGYNK.L |
| Serine/threonine kinase 38 like | STK38L | Y289 | 0.7 | 53.1 | R.QLAYSTVGTPDY*IAPEVFMQTGYNK.L |
| Serine/threonine kinase 38 like | STK38L | T286 | 1.0 | 54.9 | R.QLAYSTVGT*PDYIAPEVFMQTGYNK.L |
| Serine/threonine kinase 38 like | STK38L | S282 | 6.5 | 56.1 | R.QLAYS*TVGTPDYIAPEVFMQTGYNK.L |
| serine/threonine kinase receptor associated protein | STRAP | S335 | 14.8 | 37.7 | K.IGFPETTEEELEEIAS*ENSDCIFPSAPDVK.A |
| serine/threonine kinase receptor associated protein | STRAP | S335S338 | 58.8 | 68.5 | K.IGFPETTEEELEEIAS*ENS*DCIFPSAPDVK.A |
| Serine/Threonine protein kinase 10 | STK10 | S438 | 100.0 | 63.7 | K.QVAEQGGDLS*PAANR.S |
| Serine/Threonine protein kinase 10 | STK10 | S514 | 17.8 | 36.0 | K.EMGS*LSIKDPK.L |
| Serine/Threonine protein kinase 10 | STK10 | Y197 | 18.4 | 56.8 | R.DSFIGTPY*WMAPEVVMCETMK.D |
| Serine/Threonine protein kinase 10 | STK10 | T185 | 100.0 | 28.5 | K.NLKT*LQK.R |
| | STK10 | S191 | 1.3 | 13.9 | R.DS*FIGTPYWMAPEVVMCETMK.D |
| Serine/Threonine protein kinase 10 | STK10 | S450S454 | 4.8 | 48.0 | K.ASQS*RPNS*SALETLGGEK.L |
| Serine/Threonine protein kinase 10 | STK10 | T195 | 2.8 | 34.9 | R.DSFIGT*PYWMAPEVVMCETMK.D |
| Serine/Threonine protein kinase 10 | STK10 | S448S454 | 13.8 | 19.9 | K.AS*QSRPNS*SALETLGGEK.L |
| Serine/Threonine protein kinase 10 | STK10 | S13 | 8.9 | 13.2 | R.ILRLS*TFEK.R |
| Serine/threonine protein kinase 25 | STK24 | T174 | 80.7 | 58.8 | R.NT*FVGTPFWMAPEVIK.Q |
| Serine/threonine protein kinase 3 | STK3 | S316 | 75.5 | 61.8 | R.ELEEEEENS*DEDELDSHTMVK.T |
| Serine/threonine protein kinase 3 | STK3 | T384 | 81.0 | 24.9 | R.NAT*SPQVQRPSFMDYFDK.Q |
| Serine/threonine protein kinase 3 | STK3 | S385 | 19.1 | 28.8 | R.NATS*PQVQRPSFMDYFDKQDFK.N |
| Serine/threonine protein kinase 38 | STK38 | S281 | 9.0 | 37.2 | R.QLAFS*TVGTPDYIAPEVFMQTGYNK.L |
| Serine/threonine protein kinase 38 | STK38 | T285 | -0.7 | 57.8 | R.QLAFSTVGT*PDYIAPEVFMQTGYNK.L |
| Serine/threonine protein kinase Nek6 | NEK6 | T201S206 | 3.7 | 37.4 | R.FFSSET*TAAHS*LVGTPYYMSPER.I |
| Serine/threonine protein kinase Nek6 | NEK6 | S199S206 | 9.8 | 69.5 | R.FFSS*ETTAAHS*LVGTPYYMSPER.I |
| Serine/threonine protein kinase Nek6 | NEK6 | S199T202 | 5.6 | 57.7 | R.FFSS*ETT*AAHSLVGTPYYMSPER.I |
| Serine/threonine protein kinase Nek6 | NEK6 | S199T201T202 | 10.3 | 48.5 | R.FFSS*ET*T*AAHSLVGTPYYMSPER.I |
| Serine/threonine protein kinase Nek6 | NEK6 | S199T201 | 1.8 | 14.9 | R.FFSS*ET*TAAHSLVGTPYYMSPER.I |
| Serine/threonine protein kinase Nek6 | NEK6 | T202S206 | 8.2 | 18.5 | R.FFSSETT*AAHS*LVGTPYYMSPER.I |
| Serine/threonine-protein kinase B-raf | BRAF | T793 | 73.5 | 115.0 | K.T*PIQAGGYGEFAAFK.- |
| Serine/threonine-protein kinase RIO2 | RIOK2 | S350 | 10.9 | 28.9 | K.AEVYGS*ENESER.N |
| Serine/threonine-protein kinase Sgk1 | DKFZp686 | S118 | 12.2 | 12.4 | K.ILGLCFSCIS*AR.V |
| Serine/threonine-protein phosphatase 6 regulatory subunit 3 | PPP6R3 | T285S291S292 | 15.8 | 22.2 | R.NTVDLVT*TCHIHS*S*SDDEIDFK.E |
| Serine/threonine-protein phosphatase 6 regulatory subunit 3 | PPP6R3 | T286S292S293 | 7.5 | 22.6 | R.NTVDLVTT*CHIHSS*S*DDEIDFK.E |
| Serine/threonine-protein phosphatase 6 regulatory subunit 3 | PPP6R3 | S291S292S293 | 2.4 | 19.5 | R.NTVDLVTTCHIHS*S*S*DDEIDFK.E |
| Serine/threonine-protein phosphatase 6 regulatory subunit 3 | PPP6R3 | T286S291S292 | 0.4 | 15.7 | R.NTVDLVTT*CHIHS*S*SDDEIDFK.E |
| Serologically defined colon cancer antigen 1 | NEMF | S747S748 | 47.5 | 62.3 | R.DELNEELIQEES*S*EDEGEYEEVRK.D |
| Serologically defined colon cancer antigen 3 | SDCCAG3 | S217S224 | 12.2 | 72.1 | R.VS*PASPAGS*PSADFAVHGESLGDR.H |
| Serologically defined colon cancer antigen 3 | SDCCAG3 | S220S224 | 12.2 | 42.2 | R.VSPAS*PAGS*PSADFAVHGESLGDR.H |
| SERPINE1 mRNA binding protein 1 | SERBP1 | S203 | -0.2 | 65.8 | R.SS*FSHYSGLK.H |
| SERPINE1 mRNA binding protein 1 | SERBP1 | S330 | 11.8 | 61.8 | K.SKS*EEAHAEDSVMDHHFR.K |
| SERPINE1 mRNA binding protein 1 | SERBP1 | S328 | 4.6 | 30.4 | K.S*KSEEAHAEDSVMDHHFR.K |
| SERPINE1 mRNA binding protein 1 | SERBP1 | S25 | 100.0 | 45.7 | R.FDQLFDDES*DPFEVLK.A |
| | SERBP1 | S234 | 22.6 | 29.1 | K.DELTES*PK.Y |
| SERPINE1 mRNA binding protein 1 | SERBP1 | S237 | 14.5 | 13.0 | K.DELTDLDQS*NVTEETPEGEEHHPVADTENK.E |
| SERPINE1 mRNA binding protein 1 | SERBP1 | T232 | | 15.2 | K.DELT*ESPK.Y |
| Serum amyloid A-like 1 | SAAL1 | S56 | 25.4 | 119.3 | K.SSS*DDEEQLTELDEEMENEICR.V |
| Serum amyloid A-like 1 | SAAL1 | S55 | -0.2 | 88.0 | K.SS*SDDEEQLTELDEEMENEICR.V |
| Serum amyloid A-like 1 | SAAL1 | T63 | | 58.2 | K.SSSDDEEQLT*ELDEEMENEICR.V |
| Serum response factor | SRF | T66T67S77S85 | 5.6 | 29.1 | R.EAAAAAAT*T*PAPTAGALYS*GSEGDSES*GEEEELGAER.R |

Peak Area: <-10, -5, -3, 0, 3, 5, >10

%CV: 0, 29, 42, 71, 86, >100

**White dots-** Significant change in peptide abundance at **5** %FDR compared to the timepoint with the minimum peak area for a given PSM

| Protein Name | Gene | Phosphosites | CarT 0m | CarT 2m | CarT 5m | RajiB 0m | RajiB 2m | RajiB 5m | Ascor | MOWSE | Sequence |
|---|---|---|---|---|---|---|---|---|---|---|---|
| **Serum response factor binding protein 1** | SRFBP1 | S264 | | | | | | | **16.8** | **41.9** | K.EYFDDS*TEER.F |
| **SET binding factor 1** | SBF1 | T1747 | | | | | | | **-0.3** | **72.3** | R.ST*STLYSQFQTAESENR.S |
| **SET binding factor 1** | SBF1 | S1141 | | | | | | | **7.0** | **74.3** | R.LGLGTLSSS*LSR.A |
| **SET binding factor 1** | SBF1 | S1140 | | | | | | | **10.8** | **54.8** | R.LGLGTLSS*SLSR.A |
| **SET binding factor 1** | SBF1 | S1139 | | | | | | | **9.5** | **29.3** | R.LGLGTLS*SSLSR.A |
| **SET domain containing 1A** | SETD1A | S468S470 | | | | | | | **34.2** | **52.6** | R.S*GS*PAPETTNESVPFAQHSSLDSR.I |
| **SET domain-containing protein 5** | SETD5 | T70S74 | | | | | | | **33.3** | **18.4** | R.GLPYADHNYGAPPPPT*PPAS*PPVQTIIPR.S |
| **SET protein** | SET | T23 | | | | | | | **6.0** | **50.5** | K.ELNSNHDGADET*SEKEQQEAIEHIDEVQNEIDR.L |
| **Sex comb on midleg like 2** | SCML2 | S499S511 | | | | | | | **43.5** | **50.9** | R.S*PQQTVPYVVPLS*PK.L |
| **Sex comb on midleg like 2** | SCML2 | S511 | | | | | | | **17.6** | **11.8** | K.RSPQQTVPYVVPLS*PK.L |
| **Sex comb on midleg like 2** | SCML2 | S499 | | | | | | | **43.6** | **94.9** | R.S*PQQTVPYVVPLSPK.L |
| **Sex comb on midleg like 2** | SCML2 | T503S511 | | | | | | | **36.2** | **16.3** | R.SPQQT*VPYVVPLS*PK.L |
| **SFRS protein kinase 1** | SRPK1 | S51 | | | | | | | **76.1** | **95.7** | R.GSAPHSESDLPEQEEEILGS*DDDEQEDPNDYCK.G |
| **SFRS15** | SCAF4 | T138 | | | | | | | | **65.0** | K.IEIIQPLLDMAAGT*SNAAPVAENVTNNEGSPPPPVK.V |
| **SFRS15** | SCAF4 | S139 | | | | | | | **3.6** | **41.1** | K.IEIIQPLLDMAAGTS*NAAPVAENVTNNEGSPPPPVK.V |
| **SFRS15** | SCAF4 | S154 | | | | | | | **19.5** | **29.0** | K.IEIIQPLLDMAAGTSNAAPVAENVTNNEGS*PPPPVK.V |
| **SGEF** | ARHGEF2 | S222 | | | | | | | **62.9** | **22.8** | R.LPS*QENELLENPSVVLSTNSPAALK.V |
| **SGT** | SGTA | S305 | | | | | | | **10.9** | **71.2** | R.SRTPS*ASNDDQQE.- |
| **SGT** | SGTA | S77T81 | | | | | | | **40.9** | **33.8** | R.S*PART*PPSEEDSAEAER.L |
| **SGT** | SGTA | T303 | | | | | | | **13.9** | **46.7** | R.SRT*PSASNDDQQE.- |
| **SGT** | SGTA | T81S84 | | | | | | | **0.9** | **16.4** | R.SPART*PPS*EEDSAEAER.L |
| **SGT1 protein** | ECD | S503S505S518 | | | | | | | **66.4** | **32.9** | K.ILGPRPNES*DS*DDLDDEDFECLDS*DDDLDFETHEPGEEASLK.G |
| **SGTB** | SGTB | S297 | | | | | | | **13.9** | **27.2** | R.SFS*SSAEEHS.- |
| **SH2 B homolog** | SH2B1 | S88S96 | | | | | | | **19.4** | **42.0** | R.ASGSLS*PPILAPLS*PGAEISPHDLSLESCR.V |
| **SH2 domain binding protein 1** | CTR9 | T925 | | | | | | | **67.7** | **89.8** | K.KGGEFDEFVNDDT*DDDLPISK.K |
| **SH2 domain binding protein 1** | CTR9 | S1015S1016S1 | | | | | | | **100.0** | **15.0** | K.AIIS*S*S*DDS*S*DEDKLK.I |
| **SH2D3C** | SH2D3C | S343S346 | | | | | | | **17.4** | **14.7** | R.YLEASYGLGQGSSKPAS*PVS*PSGPK.G |
| **SH3 domain binding protein 1** | BARGIN | S550 | | | | | | | **42.4** | **50.4** | R.S*PPETAAPVEDMAR.R |
| **SH3 domain binding protein 1** | BARGIN | S262 | | | | | | | **15.6** | **61.3** | R.ENHGQADHS*PSMTATHFPR.V |
| **SH3 domain kinase binding protein 1** | SH3KBP1 | S511 | | | | | | | **11.6** | **23.0** | R.RPPSQSLTSSSLSSPDIFDSPS*PEEDKEEHISLAHR.G |
| **SH3 domain kinase binding protein 1** | SH3KBP1 | S509S511 | | | | | | | **40.7** | **43.4** | R.RPPSQSLTSSSLSSPDIFDS*PS*PEEDKEEHISLAHR.G |
| **SH3 domain kinase binding protein 1** | SH3KBP1 | S493 | | | | | | | **9.4** | **45.6** | R.RPPS*QSLTSSSLSSPDIFDSPSPEEDKEEHISLAHR.G |
| **SH3 domain kinase binding protein 1** | SH3KBP1 | S79 | | | | | | | **40.7** | **90.4** | K.APEKPLHEVPS*GNSLLSSETILR.T |
| **SH3 domain kinase binding protein 1** | SH3KBP1 | S230 | | | | | | | **100.0** | **97.6** | R.S*IEVENDFLPVEK.T |
| **SH3 domain kinase binding protein 1** | SH3KBP1 | T179 | | | | | | | **7.9** | **93.7** | R.ETT*GSESDGGDSSSTK.S |
| **SH3 domain kinase binding protein 1** | SH3KBP1 | S183 | | | | | | | **14.7** | **113.6** | R.ETTGSES*DGGDSSSTK.S |
| **SH3 domain kinase binding protein 1** | SH3KBP1 | S503S511 | | | | | | | **25.0** | **36.1** | R.RPPSQSLTSSSLSS*PDIFDSPS*PEEDKEEHISLAHR.G |
| **SH3 domain kinase binding protein 1** | SH3KBP1 | T178 | | | | | | | **7.4** | **84.9** | R.ET*TGSESDGGDSSSTK.S |
| **SH3 domain kinase binding protein 1** | SH3KBP1 | S493S495 | | | | | | | | **31.4** | R.RPPS*QS*LTSSSLSSPDIFDSPSPEEDKEEHISLAHR.G |
| **SH3 domain kinase binding protein 1** | SH3KBP1 | S495 | | | | | | | **0.0** | **43.9** | R.RPPSQS*LTSSSLSSPDIFDSPSPEEDKEEHISLAHR.G |
| **SH3 domain kinase binding protein 1** | SH3KBP1 | T497S503 | | | | | | | **-0.3** | **14.9** | R.RPPSQSLT*SSSLSS*PDIFDSPSPEEDKEEHISLAHR.G |
| **SH3 domain kinase binding protein 1** | SH3KBP1 | S181 | | | | | | | **10.9** | **107.6** | R.ETTGS*ESDGGDSSSTK.S |
| **SH3 multiple domains 2** | SH3RF1 | S735S739 | | | | | | | **15.1** | **33.2** | R.VS*PPAS*PTLEVELGSAELPLQGAVGPELPPGGGHGR.A |
| **SH3 protein expressed in lymphocytes** | SASH3 | S27 | | | | | | | **21.2** | **59.8** | R.SSS*FKDFAK.S |
| **SH3 protein expressed in lymphocytes** | SASH3 | S26S27 | | | | | | | **6.7** | **48.1** | R.SS*S*FKDFAK.S |
| **SH3 protein expressed in lymphocytes** | SASH3 | S108 | | | | | | | **14.4** | **88.3** | K.ALSEEMADTLEEGS*ASPTSPDYSLDSPGPEK.M |
| **SH3 protein expressed in lymphocytes** | SASH3 | S120 | | | | | | | **23.9** | **63.2** | K.ALSEEMADTLEEGSASPTSPDYSLDS*PGPEK.M |
| **SH3 protein expressed in lymphocytes** | SASH3 | T103 | | | | | | | **9.2** | **83.3** | K.ALSEEMADT*LEEGSASPTSPDYSLDSPGPEK.M |
| **SH3 protein expressed in lymphocytes** | SASH3 | S97 | | | | | | | **4.3** | **13.4** | K.ALS*EEMADTLEEGSASPTSPDYSLDSPGPEK.M |
| **SH3 protein expressed in lymphocytes** | SASH3 | S108S113 | | | | | | | **15.9** | **30.6** | K.ALSEEMADTLEEGS*ASPTS*PDYSLDSPGPEK.M |
| **SH3 protein expressed in lymphocytes** | SASH3 | T112S120 | | | | | | | **20.8** | **38.4** | K.ALSEEMADTLEEGSASPT*SPDYSLDS*PGPEK.M |
| **SH3 protein expressed in lymphocytes** | SASH3 | S113S120 | | | | | | | **16.6** | **48.5** | K.ALSEEMADTLEEGSASPTS*PDYSLDS*PGPEK.M |
| **SH3 protein expressed in lymphocytes** | SASH3 | S108S120 | | | | | | | **18.9** | **46.3** | K.ALSEEMADTLEEGS*ASPTSPDYSLDS*PGPEK.M |
| **SH3 protein expressed in lymphocytes** | SASH3 | T318S320 | | | | | | | **14.5** | **74.3** | K.LLTAAELLLDYDT*GS*EEAEEGAESSQEPVAHTVSEPK.V |
| **SH3 protein expressed in lymphocytes** | SASH3 | T308T318 | | | | | | | **12.0** | **70.7** | K.LLT*AAELLLDYDT*GSEEAEEGAESSQEPVAHTVSEPK.V |
| **SH3 protein expressed in lymphocytes** | SASH3 | Y316S320 | | | | | | | **24.6** | **72.6** | K.LLTAAELLLDY*DTGS*EEAEEGAESSQEPVAHTVSEPK.V |
| **SH3 protein expressed in lymphocytes** | SASH3 | S7 | | | | | | | **26.7** | **32.3** | R.RKPS*NASEKEPTQK.K |
| **SH3 protein expressed in lymphocytes** | SASH3 | S38S42 | | | | | | | **16.9** | **28.6** | K.SKPSS*PVVS*EK.E |
| **SH3 protein expressed in lymphocytes** | SASH3 | S37S42 | | | | | | | **11.5** | **13.9** | K.SKPS*SPVVS*EK.E |
| **SH3 protein expressed in lymphocytes** | SASH3 | Y316T318 | | | | | | | **25.3** | **79.7** | K.LLTAAELLLDY*DT*GSEEAEEGAESSQEPVAHTVSEPK.V |
| **SH3 protein expressed in lymphocytes** | SASH3 | S26 | | | | | | | **-0.6** | **58.4** | R.SS*SFKDFAK.S |
| **SH3 protein expressed in lymphocytes** | SASH3 | S110S120 | | | | | | | **21.1** | **56.5** | K.ALSEEMADTLEEGSAS*PTSPDYSLDS*PGPEK.M |

Peak Area: <-10, -5, -3, 0, 3, 5, >10

%CV: 0, 29, 43, 71, 86, >100

**White dots-** Significant change in peptide abundance at **5** %FDR compared to the timepoint with the minimum peak area for a given PSM

CarT: 0m 2m 5m — RajiB: 0m 2m 5m

| Protein Name | Gene | Phosphosites | Ascor | MOWSE | Sequence |
|---|---|---|---|---|---|
| SH3 protein expressed in lymphocytes | SASH3 | S110T112 | 12.4 | 20.0 | K.ALSEEMADTLEEGSAS*PT*SPDYSLDSPGPEK.M |
| SH3 protein expressed in lymphocytes | SASH3 | S108T112 | 6.7 | 42.9 | K.ALSEEMADTLEEGS*ASPT*SPDYSLDSPGPEK.M |
| SH3 protein expressed in lymphocytes | SASH3 | T308Y316 | | 36.9 | K.LLT*AAELLLDY*DTGSEEAEEGAESSQEPVAHTVSEPK.V |
| SH3 protein expressed in lymphocytes | SASH3 | T103S120 | 13.7 | 73.8 | K.ALSEEMADT*LEEGSASPTSPDYSLDS*PGPEK.M |
| SH3 protein expressed in lymphocytes | SASH3 | S97S113S120 | 6.7 | 16.6 | K.ALS*EEMADTLEEGSASPTS*PDYSLDS*PGPEK.M |
| SH3 protein expressed in lymphocytes | SASH3 | S97S120 | 15.8 | 66.0 | K.ALS*EEMADTLEEGSASPTSPDYSLDS*PGPEK.M |
| SH3-domain GRB2-like 1 | SH3GL1 | S288 | 8.9 | 37.7 | K.IAASSS*FR.S |
| SH3-domain GRB2-like 1 | SH3GL1 | S287 | 8.2 | 34.2 | K.IAASS*SFR.S |
| SH3-domain GRB2-like 1 | SH3GL1 | S286 | | 22.7 | K.IAAS*SSFR.S |
| SH3KBP1 binding protein 1 | SHKBP1 | S631S636 | 1.7 | 19.4 | R.ISLTS*LHSAS*SNTSLSGHR.G |
| SH3KBP1 binding protein 1 | SHKBP1 | S636S637 | 0.6 | 16.9 | R.ISLTSLHSAS*S*NTSLSGHR.G |
| SHARP | SPEN | S2120S2124 | 13.7 | 24.4 | R.ESGVVAVS*PEKS*ESPQKEDGLSSQLK.S |
| SHARP | SPEN | S312 | 4.9 | 61.2 | R.SVQS*AAVPAPTSQLLSSLEK.D |
| SHARP | SPEN | S309 | 22.2 | 40.4 | R.S*VQSAAVPAPTSQLLSSLEK.D |
| SHARP | SPEN | S725S727 | 100.0 | 17.5 | R.S*QS*PVHLR.R |
| Shaw related voltage gated potassium channel protein 4 | KCNC4 | S577T578 | 19.2 | 33.2 | R.RS*T*TRDR.N |
| SHD1 | SAC3D1 | S356 | 16.2 | 51.9 | R.TLEEVVMAEEEDEGTDRPGS*PA.- |
| SHIP1 | INPP5D | S970 | 39.2 | 32.7 | R.GESPPTPPGQPPIS*PK.K |
| SHIP1 | INPP5D | T962S970 | 47.7 | 15.1 | R.GESPPT*PPGQPPIS*PK.K |
| SHIP1 | INPP5D | Y864 | 100.0 | 37.5 | R.EKLY*DFVK.T |
| SHIP1 | INPP5D | S959T962 | 44.7 | 20.4 | R.GES*PPT*PPGQPPISPK.K |
| SHIP1 | INPP5D | Y1021 | 11.1 | 12.2 | K.NAGDTLPQEDLPLTKPEMFENPLY*GSLSSFPKPAPR.K |
| SHIP1 | INPP5D | S959S970 | 12.5 | 17.9 | R.GES*PPTPPGQPPIS*PK.K |
| Sialophorin | SPN | S336 | 27.5 | 33.1 | K.GSGFPDGEGS*SR.R |
| Sialophorin | SPN | S291 | 99.3 | 49.0 | R.TGALVLS*R.G |
| Sialophorin | SPN | S351S355 | 100.0 | 45.8 | K.S*RQGS*LAMEELK.S |
| Sialophorin | SPN | S355 | 100.0 | 64.8 | R.QGS*LAMEELK.S |
| Sialophorin | SPN | T341 | 32.3 | 54.6 | R.RPT*LTTFFGR.R |
| Sialophorin | SPN | S368 | 25.4 | 58.3 | K.SGSGPS*LK.G |
| Sialophorin | SPN | S351 | 35.9 | 52.1 | K.S*RQGSLAMEELK.S |
| Sialophorin | SPN | S365 | 20.6 | 28.8 | K.SGS*GPSLK.G |
| SID1 transmembrane family, member 2 | SIDT2 | S422S424S425 | 33.1 | 44.9 | R.VDS*MS*S*VEEDDYDTLTDIDSDKNVIR.T |
| SIGLEC2 | CD22 | S725 | 19.0 | 34.0 | R.TQSQQGLQENS*SGQSFFVR.N |
| SIGLEC2 | CD22 | S717 | 8.9 | 35.1 | R.TQS*QQGLQENSSGQSFFVR.N |
| Signal induced proliferation associated protein 1 | SIPA1 | S839 | 7.7 | 32.7 | R.TEFLHSQNSLS*PR.S |
| Signal induced proliferation associated protein 1 | SIPA1 | S834 | 35.2 | 22.2 | R.TEFLHS*QNSLSPR.S |
| Signal recognition particle receptor | SRPRA | S296S297S298 | 44.1 | 87.6 | R.GTGSGGQLQDLDCS*S*S*DDEGAAQNSTKPSATK.G |
| Signal sequence receptor alpha | SSR1 | S246 | 14.3 | 31.0 | K.VEMGTS*SQNDVDMSWIPQETLNQINK.A |
| Signal sequence receptor alpha | SSR1 | S247 | 14.3 | 30.0 | K.VEMGTSS*QNDVDMSWIPQETLNQINK.A |
| Signal sequence receptor alpha | SSR1 | T245 | 28.1 | 19.9 | K.VEMGT*SSQNDVDMSWIPQETLNQINK.A |
| Signal sequence receptor alpha | SSR1 | S268 | 31.5 | 16.9 | K.VEM#GTSSQNDVDMSWIPQETLNQINKAS*PR.R |
| Signal sequence receptor alpha | SSR1 | S247S268 | 19.0 | 20.7 | K.VEMGTSS*QNDVDMSWIPQETLNQINKAS*PR.R |
| Signaling lymphocytic activation molecule | SLAMF1 | Y281 | 42.9 | 31.8 | K.SLTIY*AQVQKPGPLQK.K |
| Similar to BMP2 inducible kinase | AAK1 | S36 | 21.6 | 104.0 | K.S*MEELTVIQCTSQELPAQTGLLSQTGDVPLPAGR.E |
| Similar to BMP2 inducible kinase | AAK1 | T46 | 2.3 | 54.9 | K.SM#EELTVIQCT*SQELPAQTGLLSQTGDVPLPAGR.E |
| Similar to KIAA0592 protein | WASHC2A | S250S265 | 20.8 | 22.3 | R.HTTQMS*DEEEDDDGCDLFADS*EKEEEDIEDIEENTRPK.R |
| Similar to KIAA0592 protein | WASHC2A | S539 | 75.8 | 142.8 | K.GLFS*DEEDSEDLFSSQSASK.L |
| Similar to KIAA0592 protein | WASHC2A | S619S620 | 47.0 | 85.3 | K.ASALLFS*S*DEEDQWNIPASQTHLASDSR.S |
| Similar to KIAA0592 protein | WASHC2A | S619 | 35.8 | 61.5 | K.ASALLFS*SDEEDQWNIPASQTHLASDSR.S |
| Similar to KIAA0592 protein | WASHC2A | S230S238 | 100.0 | 32.3 | K.EEEES*DEDFAHHS*DNEQNR.H |
| Similar to KIAA0592 protein | WASHC2A | S620 | 5.8 | 32.2 | K.ASALLFSS*DEEDQWNIPASQTHLASDSR.S |
| Similar to RIKEN cDNA 4933437K13 | SNX29 | S86S92 | 27.1 | 66.2 | R.YSVEAS*SPGHGS*PLSSLLPSASVPESMTISELR.Q |
| Similar to RIKEN cDNA 4933437K13 | SNX29 | S86S87 | 12.5 | 52.2 | R.YSVEAS*S*PGHGSPLSSLLPSASVPESMTISELR.Q |
| Similar to RIKEN cDNA 4933437K13 | SNX29 | S92S96 | 0.1 | 34.9 | R.YSVEASSPGHGS*PLSS*LLPSASVPESMTISELR.Q |
| Similar to RIKEN cDNA 4933437K13 | SNX29 | S86 | 9.8 | 25.5 | R.YSVEAS*SPGHGSPLSSLLPSASVPESMTISELR.Q |
| Similar to RIKEN cDNA 4933437K13 | SNX29 | S92S100 | 3.4 | 20.5 | R.YSVEASSPGHGS*PLSSLLPS*ASVPESMTISELR.Q |
| Similar to RIKEN cDNA 4933437K13 | SNX29 | S92 | 2.2 | 17.1 | R.YSVEASSPGHGS*PLSSLLPSASVPESMTISELR.Q |
| Similar to S-100 protein, alpha chain | SNTN | S39 | 12.3 | 20.2 | K.RIS*ISK.Q |
| Similar to TSG118.1 | KNOP1 | S42 | 20.5 | 12.9 | R.YSVLNNDDYFADVS*PLR.A |
| Sin3 associated polypeptide 30KD | SAP30 | S131 | 25.9 | 78.0 | R.KGS*DDDGGDSPVQDIDTPEVDLYQLQVNTLR.R |
| Sin3 associated polypeptide 30KD | SAP30 | S138 | 10.6 | 28.1 | R.KGSDDDGGDS*PVQDIDTPEVDLYQLQVNTLR.R |
| Sin3 associated polypeptide 30KD | SAP30 | S131S138 | 101.4 | 89.3 | R.KGS*DDDGGDS*PVQDIDTPEVDLYQLQVNTLR.R |

Peak Area: <-10, -5, -3, 0, 3, 5, >10

%CV: 0, 29, 42, 71, 86, >100

**White dots-** Significant change in peptide abundance at **5** %FDR compared to the timepoint with the minimum peak area for a given PSM

CarT: 0m 2m 5m — RajiB: 0m 2m 5m

| Protein Name | Gene | Phosphosites | Ascor | MOWSE | Sequence |
|---|---|---|---|---|---|
| **Sin3 associated polypeptide 30KD** | SAP30 | S131T145 | -0.1 | 37.6 | R.KGS*DDDGGDSPVQDIDT*PEVDLYQLQVNTLR.R |
| | SAP30 | S138T145 | 20.0 | 39.6 | R.KGSDDDGGDS*PVQDIDT*PEVDLYQLQVNTLR.R |
| **SIN3A** | SIN3A | T284 | 13.6 | 57.2 | R.SPPVQPHT*PVTISLGTAPSLQNNQPVEFNHAINYVNK.I |
| **SIN3A** | SIN3A | S277 | 1.5 | 53.9 | R.S*PPVQPHTPVTISLGTAPSLQNNQPVEFNHAINYVNK.I |
| **SIN3A** | SIN3A | S277T284 | 3.4 | 59.1 | R.S*PPVQPHT*PVTISLGTAPSLQNNQPVEFNHAINYVNK.I |
| **SIN3A** | SIN3A | S832 | 157.7 | 120.2 | R.GDLS*DVEEEEEEEMDVDEATGAVK.K |
| **SIN3A** | SIN3A | T287 | 1.0 | 39.2 | R.SPPVQPHTPVT*ISLGTAPSLQNNQPVEFNHAINYVNK.I |
| **SIN3A** | SIN3A | S277S289 | 6.3 | 39.6 | R.S*PPVQPHTPVTIS*LGTAPSLQNNQPVEFNHAINYVNK.I |
| **SIN3A** | SIN3A | T287S289 | 17.4 | 43.3 | R.SPPVQPHTPVT*IS*LGTAPSLQNNQPVEFNHAINYVNK.I |
| **SIN3A** | SIN3A | S277T287 | 7.3 | 54.7 | R.S*PPVQPHTPVT*ISLGTAPSLQNNQPVEFNHAINYVNK.I |
| **SIN3A** | SIN3A | S289T292 | -0.5 | 12.9 | R.SPPVQPHTPVTIS*LGT*APSLQNNQPVEFNHAINYVNK.I |
| **SIN3A** | SIN3A | T284T287 | 7.0 | 46.4 | R.SPPVQPHT*PVT*ISLGTAPSLQNNQPVEFNHAINYVNK.I |
| **SIN3A** | SIN3A | T292S295 | 14.8 | 37.8 | R.SPPVQPHTPVTISLGT*APS*LQNNQPVEFNHAINYVNK.I |
| **SIN3A** | SIN3A | T287T292 | 2.0 | 26.4 | R.SPPVQPHTPVT*ISLGT*APSLQNNQPVEFNHAINYVNK.I |
| **SIN3A** | SIN3A | Y272 | 6.0 | 20.4 | K.VSKPSQLQAHTPASQQTPPLPPY*ASPR.S |
| **SIN3A** | SIN3A | T292 | 4.5 | 41.5 | R.SPPVQPHTPVTISLGT*APSLQNNQPVEFNHAINYVNK.I |
| **Single Ig IL 1R related molecule** | SIGIRR | S383 | -0.3 | 20.0 | R.SS*EVDVSDLGSR.N |
| **Single stranded DNA binding protein 2** | SSBP2 | S354 | 7.6 | 37.4 | R.DDGEMGGNFLNPFQSESYS*PSMTMSV.- |
| **Single stranded DNA binding protein 2** | SSBP2 | S350 | -0.3 | 26.1 | R.DDGEMGGNFLNPFQS*ESYSPSMTMSV.- |
| **Single stranded DNA binding protein 2** | SSBP2 | Y353 | 5.3 | 13.1 | R.DDGEM#GGNFLNPFQSESY*SPSMTMSV.- |
| **Single stranded DNA binding protein 2** | SSBP2 | S360 | 27.6 | 37.6 | R.DDGEMGGNFLNPFQSESYSPSMTMS*V.- |
| **Single stranded DNA binding protein 2** | SSBP2 | S352 | 12.8 | 34.3 | R.DDGEMGGNFLNPFQSES*YSPSMTMSV.- |
| **Single stranded DNA binding protein 2** | SSBP2 | S356 | 12.1 | 16.4 | R.DDGEMGGNFLNPFQSESYSPS*MTMSV.- |
| **Single stranded DNA binding protein 3** | SSBP3 | S387 | 18.6 | 46.8 | R.DDGELGGNFLHSFQNDNYSPSMTMS*V.- |
| **Single stranded DNA binding protein 3** | SSBP3 | S381 | 14.0 | 56.1 | R.DDGELGGNFLHSFQNDNYS*PSMTMSV.- |
| **Single stranded DNA binding protein 3** | SSBP3 | S381S387 | 12.9 | 49.5 | R.DDGELGGNFLHSFQNDNYS*PSMTMS*V.- |
| **Single stranded DNA binding protein 3** | SSBP3 | S374S387 | 3.7 | 15.0 | R.DDGELGGNFLHS*FQNDNYSPSMTMS*V.- |
| **Single stranded DNA binding protein 3** | SSBP3 | Y380S387 | 19.5 | 49.7 | R.DDGELGGNFLHSFQNDNY*SPSMTMS*V.- |
| **Single stranded DNA binding protein 3** | SSBP3 | Y380 | 10.3 | 16.1 | R.DDGELGGNFLHSFQNDNY*SPSM#TMSV.- |
| **Single stranded DNA binding protein 3** | SSBP3 | S383S387 | 7.4 | 37.7 | R.DDGELGGNFLHSFQNDNYSPS*MTMS*V.- |
| **Single stranded DNA binding protein 3** | SSBP3 | S383 | 7.3 | 19.8 | R.DDGELGGNFLHSFQNDNYSPS*MTMSV.- |
| **Single stranded DNA binding protein 3** | SSBP3 | T360 | 7.7 | 72.5 | K.NSPNNISGISNPPGT*PR.D |
| **Single stranded DNA binding protein 3** | SSBP3 | S347T360 | 43.3 | 58.5 | K.NS*PNNISGISNPPGT*PR.D |
| **Sirtuin 1** | SIRT1 | S47 | 100.0 | 75.3 | R.S*PGEPGGAAPER.E |
| **Sirtuin 2** | SIRT2 | T365 | 8.0 | 58.6 | R.EHASIDAQSGAGVPNPST*SASPK.K |
| **SIT** | SIT1 | S182 | 25.1 | 33.2 | R.ARAS*FPDQAYANSQPAAS.- |
| **SIT** | SIT1 | Y188 | 15.4 | 58.7 | R.ASFPDQAY*ANSQPAAS.- |
| **SIT** | SIT1 | S191 | 3.5 | 38.2 | R.ASFPDQAYANS*QPAAS.- |
| **SIT** | SIT1 | S83 | 22.3 | 73.3 | R.SGES*VEEVPLYGNLHYLQTGR.L |
| **SIT** | SIT1 | S80S83 | 84.0 | 63.3 | R.S*GES*VEEVPLYGNLHYLQTGR.L |
| **SIT** | SIT1 | S80Y90 | 19.6 | 24.2 | R.S*GESVEEVPLY*GNLHYLQTGR.L |
| **SIT** | SIT1 | S80 | 7.5 | 41.6 | R.S*GESVEEVPLYGNLHYLQTGR.L |
| **SIT** | SIT1 | S102 | 122.6 | 81.7 | R.LS*QDPEPDQQDPTLGGPAR.A |
| **SKI2W** | SKIV2L | S256 | 33.8 | 64.9 | R.ASS*LEDLVLK.E |
| **SKI2W** | SKIV2L | S270T271 | 13.3 | 47.3 | K.EASTAVS*T*PEAPEPPSQEQWAIPVDATSPVGDFYR.L |
| **Skip** | SNW1 | S224S232 | 36.1 | 28.1 | R.GPPS*PPAPVMHS*PSRK.M |
| **SLAIN motif family, member 2** | SLAIN2 | S349S353 | 100.0 | 18.8 | R.NS*PRPS*PK.Q |
| **Slit 3** | SLIT3 | S685 | 100.0 | 28.1 | R.IVS*GNPR.C |
| **SLIT-ROBO Rho GTPase activating protein 2** | SRGAP2 | S1013 | 4.9 | 64.9 | R.SAS*TAGDIACAFRPVK.S |
| **Smad nuclear-interacting protein 1** | SNIP1 | S35 | 100.0 | 32.4 | R.LS*PEVAPPAHR.R |
| | SNIP1 | S394 | 9.2 | 79.7 | K.DDEDEEEEEEVS*DS.- |
| **Smad nuclear-interacting protein 1** | SNIP1 | S74S76 | 100.0 | 14.0 | R.GVS*RS*PPK.K |
| **SMAD4 interacting transcription factor** | DCP1A | S315S319 | 33.6 | 70.4 | K.HAPTYTIPLS*PVLS*PTLPAEAPTAQVPPSLPR.N |
| **SMAD4 interacting transcription factor** | DCP1A | S319T321 | 3.2 | 28.4 | K.HAPTYTIPLSPVLS*PT*LPAEAPTAQVPPSLPR.N |
| **SMAD4 interacting transcription factor** | DCP1A | S144 | 10.0 | 27.7 | R.DKQSPS*QANGCSDHRPIDILEMLSR.A |
| **SMAD4 interacting transcription factor** | DCP1A | T309Y310 | | 40.5 | K.HAPT*Y*TIPLSPVLSPTLPAEAPTAQVPPSLPR.N |
| **SMAD4 interacting transcription factor** | DCP1A | S62 | 8.6 | 34.1 | R.SAS*PYHGFTIVNR.L |
| **SMAD4 interacting transcription factor** | DCP1A | S176 | | 40.5 | R.NQMGDS*NISSPGLQPSTQLSNLGSTETLEEMPSGSQDK.S |
| **SMAD4 interacting transcription factor** | DCP1A | S180 | | 108.9 | R.NQMGDSNISS*PGLQPSTQLSNLGSTETLEEMPSGSQDK.S |
| **SMAD4 interacting transcription factor** | DCP1A | Y310T311 | 5.3 | 29.6 | K.HAPTY*T*IPLSPVLSPTLPAEAPTAQVPPSLPR.N |
| **SMAD4 interacting transcription factor** | DCP1A | S353 | 72.1 | 32.1 | R.S*PLLNQPVPELSHASLIANQSPFR.A |
| **SMAD4 interacting transcription factor** | DCP1A | S315T321 | 4.8 | 16.2 | K.HAPTYTIPLS*PVLSPT*LPAEAPTAQVPPSLPR.N |

Peak Area: <-10, -5, -3, 0, 3, 5, >10

%CV: 0, 29, 42, 71, 86, >100

**White dots-** Significant change in peptide abundance at 5 %FDR compared to the timepoint with the minimum peak area for a given PSM

CarT: 0m 2m 5m — RajiB: 0m 2m 5m

| Protein Name | Gene | Phosphosites | Ascor | MOWSE | Sequence |
|---|---|---|---|---|---|
| Small integral membrane protein 13 | SMIM13 | S50S60 | 17.6 | 44.2 | R.ELVGDTGS*QEGDHEPSGS*ETEEDTSSSPHR.I |
| Small integral membrane protein 13 | SMIM13 | S60T62 | -0.4 | 16.2 | R.ELVGDTGSQEGDHEPSGS*ET*EEDTSSSPHR.I |
| Small integral membrane protein 13 | SMIM13 | T48T62 | 12.0 | 27.9 | R.ELVGDT*GSQEGDHEPSGSET*EEDTSSSPHR.I |
| Small integral membrane protein 13 | SMIM13 | T48S60 | 20.3 | 53.9 | R.ELVGDT*GSQEGDHEPSGS*ETEEDTSSSPHR.I |
| Small integral membrane protein 13 | SMIM13 | T48S50 | 1.0 | 27.7 | R.ELVGDT*GS*QEGDHEPSGSETEEDTSSSPHR.I |
| Small integral membrane protein 13 | SMIM13 | T48S58 | 8.1 | 24.1 | R.ELVGDT*GSQEGDHEPS*GSETEEDTSSSPHR.I |
| Small integral membrane protein 13 | SMIM13 | S58S60 | 5.8 | 25.4 | R.ELVGDTGSQEGDHEPS*GS*ETEEDTSSSPHR.I |
| Small nuclear ribonucleoprotein 70 kD | SNRNP70 | S226 | 35.9 | 31.2 | R.YDERPGPS*PLPHR.D |
| Small nuclear ribonucleoprotein 70 kD | SNRNP70 | S268 | 100.0 | 17.3 | R.S*RDKEER.R |
| Small nuclear ribonucleoprotein 70 kD | SNRNP70 | S410 | 100.0 | 101.8 | R.GGGGGQDNGLEGLGNDS*R.D |
| Small nuclear ribonucleoprotein 70 kD | SNRNP70 | S320 | 100.0 | 18.9 | R.GGGGDMAEPS*EAGDAPPDDGPPGELGPDGPDGPEEK.G |
| SMAP | SMAP | S17 | 16.8 | 110.5 | R.SAS*PDDDLGSSNWEAADLGNEER.K |
| SMAP | SMAP | S147 | 100.0 | 36.1 | K.EES*AEELQAAEHPDEVEDPK.N |
| SMAP | SMAP | S17S24 | 1.8 | 24.9 | R.SAS*PDDDLGS*SNWEAADLGNEER.K |
| SMAP | SMAP | S17S25 | 8.9 | 52.3 | R.SAS*PDDDLGSS*NWEAADLGNEER.K |
| SMAP | SMAP | S15S17 | 6.5 | 16.8 | R.S*AS*PDDDLGSSNWEAADLGNEER.K |
| SMAP | SMAP | S24 | 6.2 | 76.6 | R.SASPDDDLGS*SNWEAADLGNEER.K |
| SMAP | SMAP | S25 | 3.2 | 52.8 | R.SASPDDDLGSS*NWEAADLGNEER.K |
| SMARCA2 | SMARCA2 | S1512S1516S1 | 41.2 | 53.1 | K.EEES*EDES*NEEEEEEDEEES*ESEAK.S |
| SMARCA2 | SMARCA2 | S1568S1572 | 100.0 | 53.6 | K.AKPVVS*DFDS*DEEQDER.E |
| SMARCA2 | SMARCA2 | S1572 | 49.2 | 54.9 | K.AKPVVSDFDS*DEEQDER.E |
| SMARCA4 | SMARCA4 | S1627S1631 | 100.0 | 74.8 | R.AKPVVS*DDDS*EEEQEEDR.S |
| SMARCA4 | SMARCA4 | S1570S1575S1 | 33.2 | 74.3 | K.EDDS*EGEES*EEEEEGEEEGS*ESESR.S |
| SMARCA4 | SMARCA4 | S1627S1631S1 | 100.0 | 28.6 | R.AKPVVS*DDDS*EEEQEEDRS*GS*GS*EED.- |
| SMARCA4 | SMARCA4 | T609S610 | 12.7 | 32.6 | K.AENAEGQTPAIGPDGEPLDET*S*QMSDLPVK.V |
| SMARCA4 | SMARCA4 | S655S657S660 | 68.8 | 38.4 | R.S*DS*EES*GS*EEEEEEEEEEQPQAAQPPTLPVEEK.K |
| SMARCA4 | SMARCA4 | S610S613 | 10.8 | 53.3 | K.KAENAEGQTPAIGPDGEPLDETS*QMS*DLPVK.V |
| SMARCA4 | SMARCA4 | S1382 | 19.8 | 37.7 | K.EVDYSDS*LTEK.Q |
| SMARCA4 | SMARCA4 | S1380 | 13.9 | 25.1 | K.EVDYS*DSLTEK.Q |
| SMARCA4 | SMARCA4 | S1422 | 8.2 | 35.6 | R.DSDAGSS*TPTTSTR.S |
| | SMARCA4 | T609S613 | 8.4 | 16.4 | K.AENAEGQTPAIGPDGEPLDET*SQMS*DLPVK.V |
| SMARCA5 | SMARCA5 | T113T115 | 35.0 | 52.3 | K.QTELFAHFIQPAAQKT*PT*SPLK.M |
| SMARCA5 | SMARCA5 | T113S116 | 9.0 | 36.0 | K.QTELFAHFIQPAAQKT*PTS*PLK.M |
| SMARCA5 | SMARCA5 | S116 | 12.1 | 28.7 | K.TPTS*PLK.M |
| SMARCA5 | SMARCA5 | T115S116 | 8.0 | 23.4 | K.QTELFAHFIQPAAQKTPT*S*PLK.M |
| SMARCAD1 | SMARCAD | S211S214 | 12.5 | 54.8 | R.KLS*SSS*EPYEEDEFNDDQSIKK.T |
| SMARCAD1 | SMARCAD | S211S212S214 | 15.2 | 64.6 | R.KLS*S*SS*EPYEEDEFNDDQSIKK.T |
| SMARCAD1 | SMARCAD | S96 | 10.3 | 48.7 | R.GIQYIDLSS*DSEDVVSPNCSNTVQEK.T |
| | SMARCAD | S124S127 | 46.7 | 105.8 | K.TFNKDTVIIVS*EPS*EDEESQGLPTMAR.R |
| SMARCAD1 | SMARCAD | S95S96 | 41.9 | 53.4 | R.GIQYIDLS*S*DSEDVVSPNCSNTVQEK.T |
| SMARCAD1 | SMARCAD | S152 | 58.1 | 64.2 | R.RNDDISELEDLS*ELEDLKDAK.L |
| SMARCAD1 | SMARCAD | S146S152 | 100.0 | 80.4 | R.RNDDIS*ELEDLS*ELEDLKDAK.L |
| SMARCAD1 | SMARCAD | S124S127S132 | 40.8 | 62.4 | K.DTVIIVS*EPS*EDEES*QGLPTMAR.R |
| SMARCAD1 | SMARCAD | S211S212S213 | 14.8 | 65.2 | R.KLS*S*S*SEPYEEDEFNDDQSIKK.T |
| SMARCAD1 | SMARCAD | T119S124 | 29.5 | 43.5 | K.DT*VIIVS*EPSEDEESQGLPTMAR.R |
| SMARCAD1 | SMARCAD | T54S57 | 29.2 | 32.0 | R.ANT*PDS*DITEK.T |
| SMARCAD1 | SMARCAD | S211S212 | 12.8 | 29.4 | R.KLS*S*SSEPYEEDEFNDDQSIKK.T |
| SMARCAD1 | SMARCAD | T54S57T60S66 | 24.5 | 25.6 | R.ANT*PDS*DIT*EKTEDS*S*VPETPDNER.K |
| SMARCAD1 | SMARCAD | S79 | 16.1 | 38.1 | R.KAS*ISYFK.N |
| SMARCAD1 | SMARCAD | S127 | 32.8 | 54.4 | K.DTVIIVSEPS*EDEESQGLPTMAR.R |
| SMARCAD1 | SMARCAD | S95S98S103 | 12.4 | 18.4 | R.GIQYIDLS*SDS*EDVVS*PNCSNTVQEK.T |
| SMARCAD1 | SMARCAD | S95S98 | 19.2 | 25.3 | R.GIQYIDLS*SDS*EDVVSPNCSNTVQEK.T |
| SMARCC1 | SMARCC1 | T404 | 1.0 | 18.3 | K.DSENTPVKGGT*VADLDEQDEETVTAGGK.E |
| SMARCC1 | SMARCC1 | S310 | 100.0 | 18.4 | K.NEEPVRS*PERR.D |
| SMARCC1 | SMARCC1 | S328S330 | 132.3 | 59.5 | R.KHS*PS*PPPPTPTESR.K |
| SMARCC1 | SMARCC1 | T398 | 32.3 | 37.0 | K.DSENT*PVK.G |
| SMARCC1 | SMARCC1 | S328S330T335 | 66.3 | 29.3 | R.KHS*PS*PPPPT*PTESR.K |
| SMARCC1 | SMARCC1 | S573 | 100.0 | 19.8 | R.S*PQVPAAQQMLNFPEK.N |
| SMARCC2 | SMARCC2 | S347 | 71.0 | 51.3 | K.DMDEPS*PVPNVEEVTLPK.T |
| SMARCC2 | SMARCC2 | S302S304 | 22.8 | 30.3 | R.KRS*PS*PSPTPEAK.K |
| SMARCC2 | SMARCC2 | S302T308 | 8.7 | 12.6 | R.KRS*PSPSPT*PEAK.K |
| SMC4 structural maintenance of chromosomes 4 | SMC4 | S22S28 | 40.3 | 35.7 | R.RREEGPPPPS*PDGASS*DAEPEPPSGR.T |

Peak Area: <-10, -5, -3, 0, 3, 5, >10

%CV: 0, 29, 42, 71, 86, >100

**White dots-** Significant change in peptide abundance at **5** %FDR compared to the timepoint with the minimum peak area for a given PSM

| Protein Name | Gene | Phosphosites | CarT 0m | CarT 2m | CarT 5m | RajiB 0m | RajiB 2m | RajiB 5m | Ascor | MOWSE | Sequence |
|---|---|---|---|---|---|---|---|---|---|---|---|
| **SMC4 structural maintenance of chromosomes 4** | SMC4 | S28 | | | | | | | **10.3** | **19.7** | R.RREEGPPPPSPDGASS*DAEPEPPSGR.T |
| **SMC4 structural maintenance of chromosomes 4** | SMC4 | S22S27 | | | | | | | **33.0** | **38.9** | R.EEGPPPPS*PDGAS*SDAEPEPPSGR.T |
| **SMC4 structural maintenance of chromosomes 4** | SMC4 | S27S28 | | | | | | | **4.8** | **33.6** | R.EEGPPPPSPDGAS*S*DAEPEPPSGR.T |
| **SMC4 structural maintenance of chromosomes 4** | SMC4 | S41 | | | | | | | **13.6** | **121.3** | R.TES*PATAAETASEELDNR.S |
| **SMC4 structural maintenance of chromosomes 4** | SMC4 | T44 | | | | | | | **19.1** | **64.2** | R.TESPAT*AAETASEELDNR.S |
| **SMC4 structural maintenance of chromosomes 4** | SMC4 | S27 | | | | | | | **20.6** | **24.4** | R.RREEGPPPPSPDGAS*SDAEPEPPSGR.T |
| **SMCR7L** | MIEF1 | S94 | | | | | | | **33.4** | **52.6** | R.S*LQTLPTDSSTFDTDTFCPPRPKPVAR.K |
| **SMCR7L** | MIEF1 | T97 | | | | | | | **-0.4** | **60.5** | R.SLQT*LPTDSSTFDTDTFCPPRPKPVAR.K |
| **SMEK homolog 1, suppressor of mek1** | PPP4R3A | S771S777 | | | | | | | **16.9** | **15.5** | K.TNLTSQSSTTNLPGSPGS*PGSPGS*PGSPGSVPK.N |
| **SMEK homolog 1, suppressor of mek1** | PPP4R3A | S774S780 | | | | | | | **16.3** | **10.8** | K.TNLTSQSSTTNLPGSPGSPGS*PGSPGS*PGSVPK.N |
| **SMEK homolog 1, suppressor of mek1** | PPP4R3A | S117S126 | | | | | | | **18.3** | **19.7** | K.DPSVDITQDLVDES*EEERFDDMS*SPGLELPSCELSR.L |
| **SMEK homolog 2, suppressor of mek1** | PPP4R3B | S117 | | | | | | | **82.1** | **78.1** | K.DPSVEVTQDLIDES*EEER.F |
| **Smith Magenis syndrome chromosome region candidate 8 protein** | SMCR8 | S417 | | | | | | | **13.9** | **36.5** | K.VLISVGS*YK.S |
| **SNAP190** | SNAPC4 | S1398S1400 | | | | | | | **46.1** | **41.7** | R.VGS*ES*EDEDLLSELELADR.D |
| **SNAP190** | SNAPC4 | S1398S1400S1 | | | | | | | **100.0** | **121.9** | R.VGS*ES*EDEDLLS*ELELADR.D |
| | SLK | S189 | | | | | | | **9.8** | **67.3** | R.RDS*FIGTPYWMAPEVVMCETSK.D |
| **SNF1 sucrose nonfermenting like kinase** | SLK | Y195 | | | | | | | **22.4** | **61.3** | R.DSFIGTPY*WMAPEVVMCETSK.D |
| **SNF1 sucrose nonfermenting like kinase** | SLK | S779 | | | | | | | **4.0** | **37.2** | K.DSGS*ISLQETR.R |
| **Sno** | SBNO1 | S688S692 | | | | | | | **10.1** | **28.9** | K.LYSLLGIDLTAPS*NNSS*PR.D |
| **Sno** | SBNO1 | S793 | | | | | | | **42.4** | **135.6** | K.S*IDPDSIQSALLASGLGSK.R |
| **Sno** | SBNO1 | S798 | | | | | | | **2.8** | **14.5** | K.SIDPDS*IQSALLASGLGSK.R |
| **SNX17** | SNX17 | S437S440 | | | | | | | **35.0** | **33.8** | K.LSSKLS*AVS*LR.G |
| **SNX5** | SNX5 | S22 | | | | | | | **4.5** | **46.1** | R.SVS*VDLNVDPSLQIDIPDALSER.D |
| **SODD** | BAG4 | Y255S259 | | | | | | | | **26.2** | R.Y*PWPS*SAPSAPPGNLYMTESTSPWPSSGSPQSPPSPPVQQPK.D |
| **Soluble lamin-associated protein of 75 kDa** | FAM169A | S575S576 | | | | | | | **13.0** | **60.3** | K.AVDS*S*SEEIEVEVPVVDRR.N |
| **Soluble scavenger receptor cysteine-rich domain-containing protein SSC5D** | SSC5D | S174 | | | | | | | **100.0** | **26.3** | K.KS*PRPK.Q |
| **Solute carrier family 16 (monocarboxylic acid transporters), member 1** | SLC16A1 | S461S467 | | | | | | | **9.4** | **53.0** | K.KES*KEEETS*IDVAGKPNEVTK.A |
| **Solute carrier family 16 (monocarboxylic acid transporters), member 1** | SLC16A1 | S461 | | | | | | | **68.5** | **65.6** | K.KES*KEEETSIDVAGKPNEVTK.A |
| **Solute carrier family 16 (monocarboxylic acid transporters), member 1** | SLC16A1 | S467 | | | | | | | **5.9** | **20.1** | K.EEETS*IDVAGKPNEVTK.A |
| **Solute carrier family 16 (monocarboxylic acid transporters), member 1** | SLC16A1 | S461T466 | | | | | | | **44.5** | **34.8** | K.KES*KEEET*SIDVAGKPNEVTK.A |
| **Solute carrier family 16 (monocarboxylic acid transporters), member 1** | SLC16A1 | T466 | | | | | | | **69.7** | **41.0** | K.EEET*SIDVAGKPNEVTK.A |
| **Solute carrier family 16 (monocarboxylic acid transporters), member 1** | SLC16A1 | S461T466S467 | | | | | | | **35.9** | **14.4** | K.KES*KEEET*S*IDVAGKPNEVTK.A |
| **Solute carrier family 16 (monocarboxylic acid transporters), member 1** | SLC16A1 | S213 | | | | | | | **23.6** | **38.8** | K.SKAS*LEK.A |
| **Solute carrier family 20 member 1** | SLC20A1 | Y421 | | | | | | | **14.3** | **64.9** | R.NNSYTSY*TMAICGMPLDSFR.A |
| **Solute carrier family 20 member 1** | SLC20A1 | T419 | | | | | | | **9.1** | **80.5** | R.NNSYT*SYTMAICGMPLDSFR.A |
| **Solute carrier family 20 member 1** | SLC20A1 | Y418 | | | | | | | **-0.3** | **17.5** | R.NNSY*TSYTMAICGMPLDSFR.A |
| **Solute carrier family 20 member 1** | SLC20A1 | T422 | | | | | | | **0.8** | **76.4** | R.NNSYTSYT*MAICGMPLDSFR.A |
| **Solute carrier family 20 member 1** | SLC20A1 | S417 | | | | | | | **5.2** | **22.7** | R.NNS*YTSYTMAICGMPLDSFR.A |
| **Solute carrier family 20, member 2** | SLC20A2 | S385 | | | | | | | | **68.5** | R.NNS*YTCYTAAICGLPVHATFR.A |
| **Solute carrier family 20, member 2** | SLC20A2 | Y389 | | | | | | | **5.5** | **10.7** | R.NNSYTCY*TAAICGLPVHATFR.A |
| **Solute carrier family 20, member 2** | SLC20A2 | S324 | | | | | | | **28.8** | **11.0** | K.S*PISNGTFGFDGHTR.S |
| **Solute carrier family 25, member 46** | SLC25A46 | T35 | | | | | | | **12.2** | **61.8** | R.SFST*GSDLGHWVTTPPDIPGSR.N |
| **Solute carrier family 25, member 46** | SLC25A46 | S34T35 | | | | | | | **10.9** | **60.9** | R.SFS*T*GSDLGHWVTTPPDIPGSR.N |
| **Solute carrier family 25, member 46** | SLC25A46 | S34S37 | | | | | | | **9.4** | **54.3** | R.SFS*TGS*DLGHWVTTPPDIPGSR.N |
| **Solute carrier family 25, member 46** | SLC25A46 | S37 | | | | | | | **14.7** | **63.0** | R.SFSTGS*DLGHWVTTPPDIPGSR.N |
| **Solute carrier family 25, member 46** | SLC25A46 | S34 | | | | | | | **12.2** | **59.3** | R.SFS*TGSDLGHWVTTPPDIPGSR.N |
| **Solute carrier family 26 (Sulfate transporter), member 2** | SLC26A2 | S16 | | | | | | | **70.4** | **49.6** | R.DS*AEGNDSYPSGIHLELQR.E |
| **Solute carrier family 35 member C2** | SLC35C2 | S335 | | | | | | | **21.2** | **86.4** | K.GLGS*SPDLELLLR.S |
| **Solute carrier family 35, member F2** | SLC35F2 | S371 | | | | | | | **17.8** | **38.0** | K.LEENLQETHS*AVL.- |
| **Solute carrier family 38 member 1** | SLC38A1 | S52 | | | | | | | **9.2** | **16.0** | R.S*LTNSHLEK.K |
| **Solute carrier family 38 member 1** | SLC38A1 | S52T54 | | | | | | | **9.6** | **31.1** | R.S*LT*NSHLEK.K |
| **Solute carrier family 38 member 1** | SLC38A1 | T54S56 | | | | | | | **10.6** | **40.9** | R.SLT*NS*HLEK.K |
| **Solute carrier family 38 member 1** | SLC38A1 | S52S56 | | | | | | | **17.5** | **39.2** | R.S*LTNS*HLEK.K |
| **Solute carrier family 38 member 1** | SLC38A1 | T17S28 | | | | | | | **2.1** | **41.9** | K.SGLELTELQNMT*VPEDDNISNDS*NDFTEVENGQINSK.F |
| **Solute carrier family 38 member 1** | SLC38A1 | T54 | | | | | | | **-0.2** | **60.4** | R.SLT*NSHLEK.K |
| **Solute carrier family 38 member 1** | SLC38A1 | S6T11 | | | | | | | | **30.5** | K.S*GLELT*ELQNMTVPEDDNISNDSNDFTEVENGQINSK.F |
| **Solute carrier family 38 member 1** | SLC38A1 | S25S28 | | | | | | | **12.2** | **60.6** | K.SGLELTELQNMTVPEDDNIS*NDS*NDFTEVENGQINSK.F |
| **Solute carrier family 39 (zinc transporter), member 3** | SLC39A3 | T120S129 | | | | | | | **7.6** | **70.4** | K.EKPSFIDLET*FNAGSDVGS*DSEYESPFMGGAR.G |
| **Solute carrier family 39 (zinc transporter), member 3** | SLC39A3 | S125S129 | | | | | | | **28.6** | **103.0** | K.EKPSFIDLETFNAGS*DVGS*DSEYESPFMGGAR.G |
| **Solute carrier family 39 (zinc transporter), member 3** | SLC39A3 | S125S129S131 | | | | | | | **18.4** | **98.3** | K.EKPSFIDLETFNAGS*DVGS*DS*EYESPFMGGAR.G |
| **Solute carrier family 39 (zinc transporter), member 3** | SLC39A3 | S125S129Y133 | | | | | | | **9.1** | **72.6** | K.EKPSFIDLETFNAGS*DVGS*DSEY*ESPFMGGAR.G |
| **Solute carrier family 39 (zinc transporter), member 3** | SLC39A3 | T120S129S131 | | | | | | | **1.2** | **22.0** | K.EKPSFIDLET*FNAGSDVGS*DS*EYESPFMGGAR.G |

**Peak Area**: <-10, -5, -3, 0, 3, 5, >10

**%CV**: 0, 29, 42, 71, 86, >100

**White dots**- Significant change in peptide abundance at 5 %FDR compared to the timepoint with the minimum peak area for a given PSM

| Protein Name | Gene | Phosphosites | CarT 0m | CarT 2m | CarT 5m | RajiB 0m | RajiB 2m | RajiB 5m | Ascor | MOWSE | Sequence |
|---|---|---|---|---|---|---|---|---|---|---|---|
| Solute carrier family 39 (zinc transporter), member 3 | SLC39A3 | T120S125 | | | | | | | 17.2 | 65.5 | K.EKPSFIDLET*FNAGS*DVGSDSEYESPFMGGAR.G |
| Solute carrier family 39 (zinc transporter), member 3 | SLC39A3 | T120S125S129 | | | | | | | 7.5 | 45.9 | K.EKPSFIDLET*FNAGS*DVGS*DSEYESPFMGGAR.G |
| Solute carrier family 39 (zinc transporter), member 3 | SLC39A3 | T120S129Y133 | | | | | | | 2.9 | 27.7 | K.EKPSFIDLET*FNAGSDVGS*DSEY*ESPFMGGAR.G |
| Solute carrier family 39 (zinc transporter), member 3 | SLC39A3 | S125S131 | | | | | | | 8.0 | 43.2 | K.EKPSFIDLETFNAGS*DVGSDS*EYESPFMGGAR.G |
| Solute carrier family 39 (zinc transporter), member 3 | SLC39A3 | S125Y133 | | | | | | | 5.8 | 62.0 | K.EKPSFIDLETFNAGS*DVGSDSEY*ESPFMGGAR.G |
| Solute carrier family 39 (zinc transporter), member 3 | SLC39A3 | S129S131Y133 | | | | | | | 5.5 | 41.9 | K.EKPSFIDLETFNAGSDVGS*DS*EY*ESPFMGGAR.G |
| Solute carrier family 39 (zinc transporter), member 3 | SLC39A3 | S131Y133 | | | | | | | 1.9 | 37.5 | K.EKPSFIDLETFNAGSDVGSDS*EY*ESPFMGGAR.G |
| Solute carrier family 39 member 7 | SLC39A7 | S275S276 | | | | | | | 72.8 | 42.2 | K.EKQS*S*EEEEKETR.G |
| Solute carrier family 4, member 7 | SLC4A7 | S407 | | | | | | | 16.9 | 61.4 | R.ENS*TVDFSK.G |
| Solute carrier family 4, member 7 | SLC4A7 | S242 | | | | | | | 100.0 | 36.5 | K.KHS*DPHLLER.N |
| Solute carrier family 4, member 7 | SLC4A10 | S233 | | | | | | | 100.0 | 46.5 | R.S*FADIGK.K |
| Solute carrier family 4, member 7 | SLC4A7 | S1176 | | | | | | | 61.7 | 33.0 | R.MLQDDDDTVHLPFEGGS*LLQIPVK.A |
| Solute carrier family 4, member 7 | SLC4A7 | T1167S1176 | | | | | | | 100.0 | 24.7 | R.MLQDDDDT*VHLPFEGGS*LLQIPVK.A |
| Solute carrier family 4, member 7 | SLC4A7 | S1213 | | | | | | | 12.1 | 43.3 | R.KKYVDAETS*L.- |
| Solute carrier family 4, member 7 | SLC4A7 | S556 | | | | | | | 13.9 | 31.3 | R.KIPVFHNGS*TPTLGETPK.E |
| Solute carrier family 4, sodium bicarbonate cotransporter, member 4 | SLC4A4 | S65S68 | | | | | | | 20.2 | 54.8 | R.ISENYS*DKS*DIENADESSSSILKPLISPAAER.I |
| Solute carrier family 4, sodium bicarbonate cotransporter, member 4 | SLC4A4 | Y64S65 | | | | | | | 9.8 | 12.5 | R.ISENY*S*DKSDIENADESSSSILKPLISPAAER.I |
| Solute carrier family 43 member 2 | SLC43A2 | S274 | | | | | | | 28.4 | 37.1 | R.RLS*VGSSMR.S |
| Solute carrier family 43 member 2 | SLC43A2 | S297 | | | | | | | | 47.5 | K.LCLS*TVDLEVK.C |
| Solute carrier family 5 member 1 | | S4T5 | | | | | | | 12.5 | 16.1 | -.MDSS*T*WSPK.T |
| Solute carrier family 7, member 6 opposite strand | SLC7A6O | Y300S302 | | | | | | | 23.8 | 36.7 | K.EFGY*DS*PHDLDSD.- |
| Solute carrier family 7, member 6 opposite strand | SLC7A6O | Y300 | | | | | | | 12.0 | 32.1 | K.EFGY*DSPHDLDSD.- |
| Solute carrier family 7, member 6 opposite strand | SLC7A6O | S302S308 | | | | | | | 2.7 | 12.8 | K.EFGYDS*PHDLDS*D.- |
| Solute carrier family 9 isoform A3 | SLC9A3 | S663 | | | | | | | 100.0 | 18.8 | R.KRLES*FK.S |
| Solute carrier family 9, isoform A1 | SLC9A1 | S599 | | | | | | | 15.6 | 45.8 | K.IPS*AVSTVSMQNIHPK.S |
| Solute carrier family 9, isoform A1 | SLC9A1 | S602 | | | | | | | 4.1 | 44.0 | K.IPSAVS*TVSMQNIHPK.S |
| Solute carrier family 9, isoform A1 | SLC9A1 | Y683T685 | | | | | | | 6.7 | 13.7 | K.INNY*LT*VPAHKLDSPTMSR.A |
| Solute carrier family 9, isoform A1 | SLC9A1 | T695S697 | | | | | | | 10.0 | 18.8 | K.INNYLTVPAHKLDSPT*MS*R.A |
| Solute carrier family 9, isoform A1 | SLC9A1 | S602S605 | | | | | | | 10.6 | 50.8 | K.IPSAVS*TVS*MQNIHPK.S |
| Solute carrier family 9, isoform A1 | SLC9A1 | Y649T653 | | | | | | | -0.3 | 42.9 | R.SY*NRHT*LVADPYEEAWNQMLLR.R |
| Solute carrier family 9, isoform A1 | SLC9A1 | T603S605 | | | | | | | 11.4 | 32.1 | K.IPSAVST*VS*MQNIHPK.S |
| Solute carrier family 9, isoform A3, regulatory factor 1 | SLC9A3R1 | S290 | | | | | | | 16.9 | 138.0 | R.SAS*SDTSEELNSQDSPPK.Q |
| Solute carrier family 9, isoform A3, regulatory factor 1 | SLC9A3R1 | S280 | | | | | | | 100.0 | 61.9 | R.EALAEAALES*PRPALVR.S |
| Solute carrier family 9, isoform A3, regulatory factor 1 | SLC9A3R1 | S288S291T309 | | | | | | | 8.0 | 41.5 | R.S*ASS*DTSEELNSQDSPPKQDST*APSSTSSSDPILDFNISLAMAK.E |
| Solute carrier family 9, isoform A3, regulatory factor 1 | SLC9A3R1 | S290T293S302 | | | | | | | 0.8 | 55.2 | R.SAS*SDT*SEELNSQDS*PPKQDSTAPSSTSSSDPILDFNISLAMAK.E |
| Solute carrier family 9, isoform A3, regulatory factor 1 | SLC9A3R1 | S288T293S308 | | | | | | | 6.3 | 22.4 | R.S*ASSDT*SEELNSQDSPPKQDS*TAPSSTSSSDPILDFNISLAMAK.E |
| Solute carrier family 9, isoform A3, regulatory factor 1 | SLC9A3R1 | S288S290 | | | | | | | | 41.5 | R.S*AS*SDTSEELNSQDSPPKQDSTAPSSTSSSDPILDFNISLAMAK.E |
| Solute carrier family 9, isoform A3, regulatory factor 1 | SLC9A3R1 | S294S302 | | | | | | | 15.3 | 18.4 | R.SASSDTS*EELNSQDS*PPKQDSTAPSSTSSSDPILDFNISLAMAK.E |
| Solute carrier family 9, isoform A3, regulatory factor 1 | SLC9A3R1 | S288T293T309 | | | | | | | 4.2 | 28.8 | R.S*ASSDT*SEELNSQDSPPKQDST*APSSTSSSDPILDFNISLAMAK.E |
| Solute carrier family 9, isoform A3, regulatory factor 1 | SLC9A3R1 | S288T293S299 | | | | | | | 8.2 | 34.8 | R.S*ASSDT*SEELNS*QDSPPKQDSTAPSSTSSSDPILDFNISLAMAK.E |
| Solute carrier family 9, isoform A3, regulatory factor 1 | SLC9A3R1 | S288T293S317 | | | | | | | 7.0 | 22.2 | R.S*ASSDT*SEELNSQDSPPKQDSTAPSSTSSS*DPILDFNISLAMAK.E |
| Solute carrier family 9, isoform A3, regulatory factor 1 | SLC9A3R1 | S299S302 | | | | | | | 2.1 | 10.9 | R.SASSDTSEELNS*QDS*PPKQDSTAPSSTSSSDPILDFNISLAMAK.E |
| Solute carrier family 9, isoform A3, regulatory factor 1 | SLC9A3R1 | S288S290S291 | | | | | | | | 20.3 | R.S*AS*S*DTSEELNSQDSPPKQDSTAPSSTSSSDPILDFNISLAMAK.E |
| Solute carrier family 9, isoform A3, regulatory factor 1 | SLC9A3R1 | S288 | | | | | | | | 20.4 | R.S*ASSDTSEELNSQDSPPKQDSTAPSSTSSSDPILDFNISLAMAK.E |
| Solute carrier family 9, isoform A3, regulatory factor 1 | SLC9A3R1 | S312S317 | | | | | | | 0.7 | 20.2 | R.SASSDTSEELNSQDSPPKQDSTAPS*STSSS*DPILDFNISLAMAK.E |
| Solute carrier family, member 6 | SLC2A6 | S496 | | | | | | | 70.5 | 44.5 | R.SLEQIES*FFR.T |
| SON DNA binding protein | SON | S1769 | | | | | | | 7.6 | 52.6 | R.SAAS*PVVSSMPER.A |
| SON DNA binding protein | SON | S1556 | | | | | | | 5.2 | 58.9 | K.EMEHNTVCAAGTS*PVGEIGEEK.I |
| | SON | S1948S1950S1 | | | | | | | 35.6 | 25.7 | R.S*FS*IS*PSR.R |
| SON DNA binding protein | SON | S2129 | | | | | | | 50.8 | 38.2 | K.EDDDVIVNKPHVS*DEEEEEPPFYHHPFK.L |
| SON DNA binding protein | SON | S1774S1780S1 | | | | | | | 4.3 | 14.6 | R.SAASPVVSS*MPERAS*ESS*SEEKDDYEIFVK.V |
| SON DNA binding protein | SON | S1697 | | | | | | | 72.2 | 58.8 | K.ESDQTLAALLS*PK.E |
| SON DNA binding protein | SON | S1769S1780S1 | | | | | | | 7.0 | 24.9 | R.SAAS*PVVSSMPERAS*ES*S*SEEKDDYEIFVK.V |
| SON DNA binding protein | SON | S1766S1769S1 | | | | | | | 6.0 | 11.1 | R.S*AAS*PVVS*S*MPERASESSSEEKDDYEIFVK.V |
| SON DNA binding protein | SON | S1769S1780S1 | | | | | | | 1.4 | 20.3 | R.SAAS*PVVSSMPERAS*ES*SS*EEKDDYEIFVK.V |
| SON DNA binding protein | SON | S1922T1924 | | | | | | | 12.7 | 14.5 | R.S*RT*PSRR.S |
| SON DNA binding protein | SON | S1929S1931T1 | | | | | | | 8.6 | 12.0 | R.S*RS*HT*PSRR.R |
| SON DNA binding protein | SON | S910 | | | | | | | 42.4 | 40.7 | K.S*PDPYR.L |
| SON DNA binding protein | SON | S2001S2003 | | | | | | | 100.0 | 16.5 | R.S*RS*VVR.R |
| SON DNA binding protein | SON | T1924S1926 | | | | | | | 13.9 | 15.5 | R.RSRT*PS*R.R |
| SON DNA binding protein | SON | T1555 | | | | | | | 40.4 | 60.0 | K.EMEHNTVCAAGT*SPVGEIGEEK.I |
| SON DNA binding protein | SON | S1780S1782 | | | | | | | 10.6 | 47.3 | R.AS*ES*SSEEKDDYEIFVK.V |

Peak Area: <-10, -5, -3, 0, 3, 5, >10

%CV: 0, 29, 42, 71, 86, >100

**White dots-** Significant change in peptide abundance at **5** %FDR compared to the timepoint with the minimum peak area for a given PSM

| Protein Name | Gene | Phosphosites | CarT 0m | CarT 2m | CarT 5m | RajiB 0m | RajiB 2m | RajiB 5m | Ascor | MOWSE | Sequence |
|---|---|---|---|---|---|---|---|---|---|---|---|
| SON DNA binding protein | SON | S1782S1783 | | | | | | | -0.3 | 40.7 | R.ASES*S*SEEKDDYEIFVK.V |
| SON DNA binding protein | SON | S1774S1782S1 | | | | | | | 4.4 | 19.1 | R.SAASPVVSS*MPERASES*S*SEEKDDYEIFVK.V |
| SON DNA binding protein | SON | S1769S1782S1 | | | | | | | 3.3 | 24.9 | R.SAAS*PVVSSMPERASES*S*S*EEKDDYEIFVK.V |
| SON DNA binding protein | SON | S1769S1773S1 | | | | | | | -4.0 | 19.1 | R.SAAS*PVVS*S*MPERAS*ESSSEEKDDYEIFVK.V |
| SON DNA binding protein | SON | S2020T2022 | | | | | | | 100.0 | 16.0 | R.S*RT*PLRR.R |
| SON DNA binding protein | SON | S1766S1769S1 | | | | | | | 5.7 | 12.4 | R.S*AAS*PVVSS*MPERASESSS*EEKDDYEIFVK.V |
| SON DNA binding protein | SON | S1766S1769S1 | | | | | | | 6.4 | 19.7 | R.S*AAS*PVVS*SMPERASESSS*EEKDDYEIFVK.V |
| SON DNA binding protein | SON | S1769S1773S1 | | | | | | | 6.7 | 21.9 | R.SAAS*PVVS*S*MPERASESSS*EEKDDYEIFVK.V |
| SON DNA binding protein | SON | S1769S1773S1 | | | | | | | 1.7 | 23.5 | R.SAAS*PVVS*SMPERAS*ES*SSEEKDDYEIFVK.V |
| Sortilin | SORT1 | S825 | | | | | | | 27.8 | 41.3 | K.SGYHDDS*DEDLLE.- |
| Sorting nexin 1 | SNX1 | S32S39 | | | | | | | 34.4 | 93.0 | R.LPPPFPGLEPESEGAAGGS*EPEAGDS*DTEGEDIFTGAAVVSK.H |
| Sorting nexin 1 | SNX1 | S39T41 | | | | | | | 3.1 | 31.7 | R.LPPPFPGLEPESEGAAGGSEPEAGDS*DT*EGEDIFTGAAVVSK.H |
| Sorting nexin 1 | SNX1 | S25S32S39 | | | | | | | | 91.2 | R.LPPPFPGLEPES*EGAAGGS*EPEAGDS*DTEGEDIFTGAAVVSK. |
| Sorting nexin 1 | SNX1 | S25S39T41 | | | | | | | 6.4 | 52.1 | R.LPPPFPGLEPES*EGAAGGSEPEAGDS*DT*EGEDIFTGAAVVSK. |
| Sorting nexin 1 | SNX1 | S32S39T41 | | | | | | | 29.5 | 47.0 | R.LPPPFPGLEPESEGAAGGS*EPEAGDS*DT*EGEDIFTGAAVVSK. |
| Sorting nexin 1 | SNX1 | S25S32 | | | | | | | | 81.8 | R.LPPPFPGLEPES*EGAAGGS*EPEAGDSDTEGEDIFTGAAVVSK.H |
| Sorting nexin 1 | SNX1 | S32T41 | | | | | | | 5.2 | 56.5 | R.LPPPFPGLEPESEGAAGGS*EPEAGDSDT*EGEDIFTGAAVVSK.H |
| Sorting nexin 11 | SNX11 | S191S192 | | | | | | | | 14.7 | R. RS*S*PSPPPSEEKDHLEVWAPVVDSEVPSLESPTLPPLSSPLCCDE |
| Sorting nexin 15 | SNX15 | S179S180 | | | | | | | | 20.3 | R.RGLEELEVPVDPPPS*S*PAQEALDLLFNCESTEEASGSPAR.G |
| Sorting nexin 15 | SNX15 | S179S201 | | | | | | | 6.5 | 14.4 | R.RGLEELEVPVDPPPS*SPAQEALDLLFNCESTEEASGS*PAR.G |
| Sorting nexin 2 | SNX2 | S185 | | | | | | | 55.8 | 32.6 | R.RFS*DFLGLHSK.L |
| Sorting nexin 2 | SNX2 | S119 | | | | | | | 33.7 | 58.4 | K.SMS*APVIFDR.S |
| Sorting nexin 3 | SNX3 | Y71 | | | | | | | 8.9 | 39.7 | R.RY*SDFEWLR.S |
| Sorting nexin 3 | SNX3 | S72 | | | | | | | 27.6 | 46.7 | R.RYS*DFEWLR.S |
| Source of immunodominant MHC associated peptides | STT3B | S498S499 | | | | | | | 77.9 | 32.0 | R.ENPPVEDS*S*DEDDKRNQGNLYDK.A |
| SP100 | SP100 | S407S409S410 | | | | | | | 101.6 | 43.9 | R.VIGQDHDFS*ES*S*EEEAPAEASSGALR.S |
| SP100 | SP100 | S157 | | | | | | | 100.0 | 45.8 | K.GFENVIHDKLPLQES*EEEER.E |
| SP100 | SP100 | S451S452 | | | | | | | 25.5 | 41.9 | R.RFS*S*SDFSDLSNGEELQETCSSSLR.R |
| SP100 | SP100 | S18 | | | | | | | 72.0 | 47.0 | R.LNECIS*PVANEMNHLPAHSHDLQR.M |
| SP100 | SP100 | S327S328S331 | | | | | | | 27.4 | 41.7 | R.THHNQASDIIVIS*S*EDS*EGS*T*DVDEPLEVFISAPR.S |
| SP110 nuclear body protein | SP110 | S256 | | | | | | | 103.8 | 37.5 | R.DNS*PEPNDPEEPQEVSSTPSDKK.G |
| SP110 nuclear body protein | SP110 | S244 | | | | | | | 39.1 | 39.1 | R.DKEDPQEMPHS*PLGSMPEIR.D |
| SPAK | STK39 | S385 | | | | | | | 161.4 | 104.7 | K.TEDGDWEWS*DDEMDEK.S |
| SPAK | STK39 | S370 | | | | | | | 17.8 | 36.5 | R.RVPGS*SGHLHK.T |
| Spatacsin | SPG11 | T1956S1969S1 | | | | | | | 3.9 | 53.9 | R.VHST*SSLDSQKFVTVPS*S*NEVVTNLEVLTSK.C |
| Spatacsin | SPG11 | S1955T1966S1 | | | | | | | 9.6 | 51.5 | R.VHS*TSSLDSQKFVT*VPS*SNEVVTNLEVLTSK.C |
| Spatacsin | SPG11 | S1955S1958S1 | | | | | | | 9.6 | 96.7 | R.VHS*TSS*LDS*QKFVTVPSSNEVVTNLEVLTSK.C |
| Spatacsin | SPG11 | T1956S1958S1 | | | | | | | 3.6 | 35.3 | R.VHST*SS*LDS*QKFVTVPSSNEVVTNLEVLTSK.C |
| Spatacsin | SPG11 | T1956S1957S1 | | | | | | | 0.4 | 24.5 | R.VHST*S*SLDS*QKFVTVPSSNEVVTNLEVLTSK.C |
| Spatacsin | SPG11 | S1969S1970T1 | | | | | | | -0.2 | 23.6 | R.VHSTSSLDSQKFVTVPS*S*NEVVT*NLEVLTSK.C |
| Spatacsin | SPG11 | S1955S1958S1 | | | | | | | 6.4 | 64.4 | R.VHS*TSS*LDSQKFVTVPS*SNEVVTNLEVLTSK.C |
| Spatacsin | SPG11 | S1955T1956T19 | | | | | | | 0.1 | 14.7 | R.VHS*T*SSLDSQKFVT*VPSSNEVVTNLEVLTSK.C |
| SPECC1-like | SPECC1L | S384 | | | | | | | 33.8 | 22.4 | R.KGS*SGNASEVSVACLTER.I |
| SPECC1-like | SPECC1L | S389 | | | | | | | 9.1 | 58.9 | R.KGSSGNAS*EVSVACLTER.I |
| SPECC1-like | SPECC1L | S973S981 | | | | | | | 6.9 | 24.7 | R.DISAQEGASPASLMAMGTTS*PQLSLSSS*PTASVTPTTR.S |
| SPECC1-like | SPECC1L | S887 | | | | | | | 20.7 | 22.0 | R.HS*ISGPISTSKPLTALSDK.R |
| Spectrin, alpha, non-erythrocytic 1 (alpha-fodrin) | SPTAN1 | S1031 | | | | | | | 10.9 | 32.9 | K.KLDPAQSAS*R.E |
| Spectrin, alpha, non-erythrocytic 1 (alpha-fodrin) | SPTAN1 | S1190 | | | | | | | 45.2 | 33.8 | K.TAS*PWK.S |
| Spectrin, alpha, non-erythrocytic 1 (alpha-fodrin) | SPTAN1 | S1217 | | | | | | | 100.0 | 53.3 | R.S*LQQLAEER.S |
| Spectrin, alpha, non-erythrocytic 1 (alpha-fodrin) | SPTAN1 | S1029 | | | | | | | | 18.1 | K.KLDPAQS*ASR.E |
| Spectrin, beta, non erythrocytic 5 | SPTBN5 | S766S767 | | | | | | | 100.0 | 36.4 | R.RS*S*LER.A |
| Sperm associated antigen 9 | SPAG9 | S716S719 | | | | | | | 17.2 | 87.2 | R.SAS*QSS*LDKLDQELK.E |
| Sperm associated antigen 9 | SPAG9 | S203T217 | | | | | | | 100.0 | 49.7 | K.ERPIS*LGIFPLPAGDGLLT*PDAQK.G |
| Sperm associated antigen 9 | SPAG9 | T217 | | | | | | | 82.2 | 30.2 | K.ERPISLGIFPLPAGDGLLT*PDAQK.G |
| Sperm associated antigen 9 | SPAG9 | S718S719 | | | | | | | 6.1 | 22.1 | R.SASQS*S*LDKLDQELK.E |
| Sperm associated antigen 9 | SPAG9 | S718 | | | | | | | 20.1 | 51.8 | R.SASQS*SLDKLDQELK.E |
| Sperm associated antigen 9 | SPAG9 | S716S718 | | | | | | | 3.2 | 22.6 | R.SAS*QS*SLDKLDQELK.E |
| Sperm associated antigen 9 | SPAG9 | S536S537 | | | | | | | 19.4 | 11.6 | K.RS*S*IWQFFSR.L |
| Sperm associated antigen 9 | SPAG9 | S203 | | | | | | | 100.5 | 53.1 | K.ERPIS*LGIFPLPAGDGLLTPDAQK.G |
| Sperm associated antigen 9 | SPAG9 | S537S543 | | | | | | | 15.0 | 13.0 | K.RSS*IWQFFS*R.L |
| Sperm specific antigen 2 | SSFA2 | S739 | | | | | | | 28.9 | 82.2 | R.SQS*LPTTLLSPVR.V |
| Sperm specific antigen 2 | SSFA2 | S737 | | | | | | | 12.2 | 69.2 | R.S*QSLPTTLLSPVR.V |

Peak Area: <-10, -5, -3, 0, 3, 5, >10

%CV: 0, 29, 42, 71, 86, >100

**White dots-** Significant change in peptide abundance at **5** %FDR compared to the timepoint with the minimum peak area for a given PSM

CarT (0m, 2m, 5m) RajiB (0m, 2m, 5m)

| Protein Name | Gene | Phosphosites | Ascor | MOWSE | Sequence |
|---|---|---|---|---|---|
| **Sperm specific antigen 2** | SSFA2 | S1055 | **8.4** | **18.9** | R.S*ADNLSCPSPLNVMEPVTELMQEQSYLK.S |
| **Spermatid perinuclear RNA binding protein** | STRBP | S465S466 | **4.6** | **36.0** | K.VLQAMGYPTGFDADIECMS*S*DEKSDNESK.N |
| **Spermatid perinuclear RNA binding protein** | STRBP | T455S470 | **11.8** | **11.1** | K.VLQAMGYPT*GFDADIECMSSDEKS*DNESK.N |
| **Spermatid perinuclear RNA binding protein** | STRBP | Y453T455 | **4.8** | **26.3** | K.VLQAMGY*PT*GFDADIECMSSDEKSDNESK.N |
| **Spermatogenesis associated 5 like 1** | SPATA5L1 | S93S95 | **100.0** | **11.4** | R.RS*LS*LNR.L |
| **Sphingosine kinase 2** | SPHK2 | T402S414 | **41.7** | **29.0** | R.AKSELT*LTPDPAPPMAHS*PLHR.S |
| **Sphingosine kinase 2** | SPHK2 | S399S414 | **48.4** | **12.2** | R.AKS*ELTLTPDPAPPMAHS*PLHR.S |
| **Sphingosine-1-phosphate phosphatase 1** | SGPP1 | S112 | **35.4** | **80.2** | R.NS*LTGEEGQLAR.V |
| **Spliceosome associated protein 145** | SF3B2 | S307S309 | **12.8** | **50.7** | R.SSLGQS*AS*ETEEDTVSVSK.K |
| **Spliceosome associated protein 145** | SF3B2 | T780 | **5.9** | **75.2** | K.IEEAMDGSET*PQLFTVLPEKR.T |
| **Spliceosome associated protein 145** | SF3B2 | S360 | **13.1** | **74.7** | R.GS*DSPAADVEIEYVTEEPEIYEPNFIFFK.R |
| **Spliceosome associated protein 145** | SF3B2 | S431S435S436 | **100.0** | **50.8** | K.GFEEEHKDS*DDDS*S*DDEQEKKPEAPK.L |
| **Spliceosome associated protein 145** | SF3B2 | S360S362 | **2.7** | **31.3** | R.SRGS*DS*PAADVEIEYVTEEPEIYEPNFIFFK.R |
| **Spliceosome associated protein 145** | SF3B2 | S357S362 | **5.5** | **26.1** | R.S*RGSDS*PAADVEIEYVTEEPEIYEPNFIFFK.R |
| **Spliceosome associated protein 145** | SF3B2 | T706S714S718 | | **15.8** | R. T*PWCELEPS*DEES*S*EEEEEEESDEDKPDETGEITRADSCLITPG |
| **Spliceosome associated protein 145** | SF3B2 | T706S714S718 | | **17.9** | R. T*PWCELEPS*DEES*S*EEEEEEES*DEDKPDETGEITRADSCLITPG |
| **Spliceosome associated protein 145** | SF3B2 | S302S309 | **8.5** | **18.4** | R.S*SLGQSAS*ETEEDTVSVSKK.E |
| **Spliceosome associated protein 145** | SF3B2 | S309 | **10.9** | **47.1** | R.SSLGQSAS*ETEEDTVSVSK.K |
| **Spliceosome associated protein 145** | SF3B2 | S778 | **64.0** | **96.9** | K.IEEAMDGS*ETPQLFTVLPEKR.T |
| **Spliceosome associated protein 145** | SF3B2 | S362 | **-0.1** | **11.3** | R.GSDS*PAADVEIEYVTEEPEIYEPNFIFFK.R |
| **Splicing factor 1** | SF1 | S80S82 | **32.3** | **15.7** | R.TGDLGIPPNPEDRS*PS*PEPIYNSEGK.R |
| | SF3A1 | S329 | **7.8** | **49.5** | K.FGESEEVEM#EVES*DEEDDKQEK.A |
| **Splicing factor 3 subunit 1** | SF3A1 | S344 | | **25.5** | K. AEEPPS*QLDQDTQVQDMDEGSDDEEEGQKVPPPPETPMPPPLPP |
| **Splicing factor 3 subunit 1** | SF3A1 | S359 | **57.0** | **68.0** | K.AEEPPSQLDQDTQVQDMDEGS*DDEEEGQK.V |
| **Splicing factor 3A, subunit 3** | SF3A3 | S365S367S369 | **102.2** | **76.2** | R.EEEEEEQIS*ES*ES*EDEENEIIYNPK.N |
| **Splicing factor 3B subunit 1** | SF3B1 | S308T313 | **20.8** | **52.4** | R.DTPGHGS*GWAET*PR.T |
| **Splicing factor 3B subunit 1** | SF3B1 | T244T248 | **19.1** | **53.3** | K.GSET*PGAT*PGSK.I |
| **Splicing factor 3B subunit 1** | SF3B1 | T235 | **100.0** | **20.3** | R.WDET*PGR.A |
| **Splicing factor 3B subunit 1** | SF3B1 | T434T436 | **27.5** | **106.6** | R.KLTAT*PT*PLGGMTGFHMQTEDR.T |
| **Splicing factor 3B subunit 1** | SF3B1 | T303T313 | **20.5** | **23.2** | R.DT*PGHGSGWAET*PR.T |
| **Splicing factor 3B subunit 1** | SF3B1 | S129T142 | **12.6** | **16.4** | R.TMIIS*PERLDPFADGGKT*PDPK.M |
| **Splicing factor 3B subunit 1** | SF3B1 | T211 | **12.7** | **23.6** | R.WDQTADQTPGAT*PK.K |
| **Splicing factor 3B subunit 1** | SF3B1 | T267T273 | **17.3** | **16.2** | K.IWDPTPSHTPAGAAT*PGRGDT*PGHATPGHGGATSSAR.K |
| **Splicing factor 3B subunit 1** | SF3B1 | T257S259 | | **12.0** | K.IWDPT*PS*HTPAGAATPGRGDTPGHATPGHGGATSSAR.K |
| **Splicing factor 3B subunit 1** | SF3B1 | T223S229 | **9.3** | **16.3** | K.KLSSWDQAET*PGHTPS*LR.W |
| **Splicing factor 3B subunit 1** | SF3B1 | T303S308 | **7.8** | **10.8** | R.DT*PGHGS*GWAETPR.T |
| **Splicing factor 3B subunit 1** | SF3B1 | T223T227 | **10.6** | **17.3** | K.LSSWDQAET*PGHT*PSLR.W |
| **Splicing factor 4** | SUGP1 | S485 | **31.5** | **99.4** | K.AVQQHQHGYDS*DEEVDSELGTWEHQLR.R |
| **Splicing factor 4** | SUGP1 | S491 | **5.8** | **45.7** | K.AVQQHQHGYDSDEEVDS*ELGTWEHQLR.R |
| **Splicing factor arginine/serine rich 5** | SRSF5 | S248S250S253 | **100.0** | **34.7** | R.S*KS*PAS*VDR.Q |
| **Splicing factor arginine/serine rich 5** | SRSF5 | S250S253 | **100.0** | **21.8** | K.S*PAS*VDR.Q |
| **Splicing factor arginine/serine rich 5** | SRSF5 | S248S250 | **18.5** | **36.4** | R.S*KS*PASVDR.Q |
| **Splicing factor arginine/serine rich 5** | SRSF5 | S116 | | **20.9** | R.LIVENLS*SR.C |
| **Splicing factor HCC1** | RBM39 | S136 | **212.6** | **64.8** | K.DKS*PVREPIDNLTPEER.D |
| **Splicing factor HCC1** | RBM39 | Y95 | **42.3** | **19.9** | R.Y*RSPYSGPK.F |
| **Splicing factor HCC1** | RBM39 | S127S129 | **100.0** | **17.6** | R.S*KS*PFRK.D |
| **Splicing factor HCC1** | RBM39 | S125S127S129 | **100.0** | **11.9** | R.S*RS*KS*PFR.K |
| **Splicing factor HCC1** | RBM39 | S97 | **-10.2** | **29.8** | R.YRS*PYSGPK.F |
| **Splicing factor YT521-B** | YTHDC1 | S424 | **7.7** | **54.5** | R.LSSESHHGGS*PIHWVLPAGMSAK.M |
| **Splicing factor YT521-B** | YTHDC1 | S308 | **100.0** | **40.9** | R.GIS*PIVFDR.S |
| **Splicing factor YT521-B** | YTHDC1 | S146T148 | **24.6** | **21.1** | R.AKS*PT*PDGSER.I |
| **Splicing factor YT521-B** | YTHDC1 | S119 | **2.8** | **16.2** | R.LSS*SASR.E |
| **Splicing factor YT521-B** | YTHDC1 | S419 | **-0.4** | **33.4** | R.LSSES*HHGGSPIHWVLPAGMSAK.M |
| **Splicing factor YT521-B** | YTHDC1 | S417 | **3.0** | **35.2** | R.LSS*ESHHGGSPIHWVLPAGMSAK.M |
| **Splicing factor, 45 kd** | RBM17 | S222 | **10.1** | **119.7** | R.S*PTGPSNSFLANMGGTVAHK.I |
| **Splicing factor, 45 kd** | RBM17 | S155 | **76.7** | **77.7** | R.RPDPDS*DEDEDYER.E |
| | RBM17 | S229 | **4.3** | **68.3** | R.SPTGPSNS*FLANM#GGTVAHK.I |
| **Splicing factor, 45 kd** | RBM17 | S227 | **0.5** | **74.4** | R.SPTGPS*NSFLANMGGTVAHK.I |
| **Splicing factor, 45 kd** | RBM17 | T224 | **-0.0** | **16.3** | R.SPT*GPSNSFLANMGGTVAHK.I |
| **Splicing factor, arginine/serine rich 1** | SRSF1 | Y237S238 | **6.1** | **15.6** | R.Y*S*PRHSR.S |
| **Splicing factor, arginine/serine rich 1** | SRSF1 | S199 | **14.0** | **42.8** | R.S*PSYGR.S |
| **Splicing factor, arginine/serine rich 1** | TRA2B | Y226S227 | **15.0** | **15.9** | R.SY*S*PRR.S |

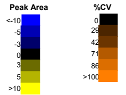


**White dots-** Significant change in peptide abundance at **5** %FDR compared to the timepoint with the minimum peak area for a given PSM

| Protein Name | Gene | Phosphosites | CarT 0m 2m 5m | RajiB 0m 2m 5m | Ascor | MOWSE | Sequence |
|---|---|---|---|---|---|---|---|
| **Splicing factor, arginine/serine rich 1** | TRA2B | S225Y226S227 | | | **9.7** | **16.7** | R.SRS*Y*S*PR.R |
| **Splicing factor, arginine/serine rich 1** | SRSF1 | S199S201 | | | **14.5** | **14.1** | R.VKVDGPRS*PS*YGR.S |
| **Splicing factor, arginine/serine rich 1** | SRSF1 | S231S234 | | | **100.0** | **18.0** | R.RS*RGS*PR.Y |
| **Splicing factor, arginine/serine rich 1** | SRSF1 | Y237S242 | | | **10.8** | **12.5** | R.Y*SPRHS*R.S |
| **Splicing factor, arginine/serine rich 1** | TRA2B | S223S225Y226 | | | **8.3** | **13.4** | R.S*RS*Y*SPR.R |
| **Splicing factor, arginine/serine rich 1** | SRSF1 | S201Y202 | | | **5.5** | **14.8** | R.VKVDGPRSPS*Y*GR.S |
| **Splicing factor, arginine/serine rich 1** | TRA2B | S223S225S227 | | | **9.0** | **17.7** | R.S*RS*YS*PR.R |
| **Splicing factor, arginine/serine rich 11** | SRSF11 | S483 | | | **100.0** | **30.4** | K.VNGDDHHEEDMDMS*D.- |
| **Splicing factor, arginine/serine rich 11** | SRSF11 | S207 | | | **27.7** | **71.5** | K.LNHVAAGLVS*PSLK.S |
| **Splicing factor, arginine/serine rich 11** | SRSF11 | S323T325 | | | **100.0** | **17.9** | R.S*KT*PPK.S |
| **Splicing factor, arginine/serine rich 11** | SRSF11 | S449 | | | **29.8** | **36.8** | K.KPIETGS*PK.T |
| **Splicing factor, arginine/serine rich 11** | SRSF11 | T260S262 | | | **13.1** | **25.0** | R.RRT*PS*SSR.H |
| **Splicing factor, arginine/serine rich 11** | SRSF11 | Y432 | | | **9.1** | **34.2** | R.DYDEEEQGY*DSEKEK.K |
| **Splicing factor, arginine/serine rich 11** | SRSF11 | S434 | | | **9.7** | **31.6** | R.DYDEEEQGYDS*EKEK.K |
| **Splicing factor, arginine/serine rich 11** | SRSF11 | T260S263 | | | **4.0** | **11.2** | R.RRT*PSS*SR.H |
| **Splicing factor, arginine/serine rich 11** | SRSF11 | T447 | | | **-19.1** | **40.0** | K.KPIET*GSPK.T |
| **Splicing factor, arginine/serine rich 11** | SRSF11 | S263S264 | | | **6.3** | **15.0** | R.RRTPSS*S*R.H |
| **Splicing factor, arginine/serine rich 2** | RTBDN | S119S121 | | | **100.0** | **19.0** | R.RS*RS*PR.R |
| **Splicing factor, arginine/serine rich 2** | SRSF2 | S145 | | | **19.1** | **21.7** | R.SRYS*R.S |
| **Splicing factor, arginine/serine rich 2** | SRSF2 | S191 | | | **45.2** | **47.3** | R.S*PPPVSK.R |
| **Splicing factor, arginine/serine rich 2** | SRSF2 | S189S191 | | | **61.7** | **31.0** | R.S*RS*PPPVSK.R |
| **Splicing factor, arginine/serine rich 2** | SRSF2 | S187S189S191 | | | **70.4** | **21.7** | R.S*RS*RS*PPPVSK.R |
| | SRSF2 | S26 | | | **15.0** | **35.2** | R.TS*PDTLR.R |
| **Splicing factor, arginine/serine rich 2** | SRSF2 | S206S208 | | | **45.0** | **28.6** | R.S*KS*PPKSPEEEGAVSS.- |
| **Splicing factor, arginine/serine rich 2** | SRSF2 | S206S208S220 | | | **14.3** | **38.8** | R.S*KS*PPKSPEEEGAVS*S.- |
| **Splicing factor, arginine/serine rich 2** | SRSF2 | S206S208S212 | | | **41.8** | **12.1** | R.S*KS*PPKS*PEEEGAVSS.- |
| **Splicing factor, arginine/serine rich 2** | SRSF2 | S206S212S220 | | | **4.7** | **22.1** | R.S*KSPPKS*PEEEGAVS*S.- |
| **Splicing factor, arginine/serine-rich 2, interacting protein** | SCAF11 | S796S802 | | | **18.3** | **32.2** | R.FHS*PSTTWS*PNKDTPQEK.K |
| **Splicing factor, arginine/serine-rich 2, interacting protein** | SCAF11 | S796S798 | | | **33.6** | **44.6** | R.FHS*PS*TTWSPNKDTPQEK.K |
| **Splicing factor, arginine/serine-rich 2, interacting protein** | SCAF11 | S338 | | | **11.0** | **46.4** | R.S*PISDNSGCDAPGNSNPSLSVPSSAESEK.Q |
| **Splicing factor, arginine/serine-rich 2, interacting protein** | SCAF11 | S901 | | | | **44.5** | R.VKDS*SPGEK.S |
| **Splicing factor, arginine/serine-rich 2, interacting protein** | SCAF11 | S816S818 | | | **100.0** | **13.8** | K.KRPQS*PS*PR.R |
| **Splicing factor, arginine/serine-rich 2, interacting protein** | SCAF11 | S878S880S882 | | | **100.0** | **12.4** | R.RS*ES*LS*PR.R |
| **Splicing factor, arginine/serine-rich 2, interacting protein** | SCAF11 | S796T800 | | | **18.9** | **33.2** | R.FHS*PSTT*WSPNKDTPQEK.K |
| **Splicing factor, arginine/serine-rich 2, interacting protein** | SCAF11 | S341 | | | **7.9** | **27.4** | R.SPIS*DNSGCDAPGNSNPSLSVPSSAESEK.Q |
| **Splicing factor, arginine/serine-rich 4** | SRSF4 | S364S366 | | | **24.4** | **20.1** | R.SREES*RS*R.S |
| **Splicing factor, arginine/serine-rich 4** | SRSF4 | S316 | | | **10.2** | **32.1** | K.RGS*VSR.G |
| **Splicing factor, arginine/serine-rich 4** | SRSF4 | S330 | | | **16.0** | **21.2** | K.SLRQS*R.S |
| **Splicing factor, arginine/serine-rich 4** | SRSF4 | S318 | | | **9.1** | **26.8** | K.RGSVS*R.G |
| **Splicing factor, arginine/serine-rich 4** | SRSF4 | S267S269 | | | **100.0** | **36.4** | R.S*KS*KDQAEEK.I |
| **Splicing factor, arginine/serine-rich 4** | SRSF4 | S410S412 | | | **100.0** | **16.9** | R.S*VS*KER.E |
| **Splicing factor, arginine/serine-rich 4** | SRSF4 | S456 | | | **12.8** | **20.0** | K.SKPNLPS*ESR.S |
| **Splicing factor, arginine/serine-rich 4** | SRSF4 | S458 | | | **5.9** | **30.9** | K.SKPNLPSES*R.S |
| **Splicing factor, arginine/serine-rich 6** | SRSF6 | S303 | | | **33.6** | **63.0** | R.SNS*PLPVPPSK.A |
| **Splicing factor, arginine/serine-rich 6** | SRSF6 | S218S220S222 | | | **100.0** | **17.5** | R.S*IS*KS*R.S |
| **Splicing factor, arginine/serine-rich 6** | SRSF6 | S314S316 | | | **100.0** | **29.5** | R.S*VS*PPPKR.A |
| **Splicing factor, arginine/serine-rich 6** | SRSF6 | S265 | | | **49.2** | **47.4** | R.S*KDEYEK.S |
| **Splicing factor, arginine/serine-rich 6** | SRSF6 | S263S265 | | | **39.9** | **18.3** | R.S*RS*KDEYEK.S |
| **Splicing factor, arginine/serine-rich 6** | SRSF6 | S257S259S261 | | | **18.5** | **28.4** | K.SDRGS*HS*HS*R.S |
| **Splicing factor, arginine/serine-rich 6** | SRSF6 | S301S303 | | | **50.0** | **18.8** | R.S*NS*PLPVPPSK.A |
| **Splicing factor, arginine/serine-rich 6** | SRSF6 | S45 | | | **36.9** | **49.3** | K.NGYGFVEFEDS*R.D |
| **Splicing factor, arginine/serine-rich 6** | SRSF6 | S265Y269 | | | **-2.4** | **20.2** | R.S*KDEY*EKSR.S |
| **Splicing factor, arginine/serine-rich 6** | SRSF6 | S295S297S299 | | | **22.5** | **10.9** | R.S*RS*QS*RS*NS*PLPVPPSK.A |
| **Splicing factor, arginine/serine-rich 6** | SRSF6 | S265S272 | | | **14.1** | **26.4** | R.S*KDEYEKS*R.S |
| **Splicing factor, arginine/serine-rich 7,35kDa** | SRSF7 | S165S167 | | | **100.0** | **19.4** | R.S*IS*LRR.S |
| **Splicing factor, arginine/serine-rich 7,35kDa** | SRSF7 | S163S165S167 | | | **100.0** | **13.3** | R.S*RS*IS*LR.R |
| **Splicing factor, arginine/serine-rich 7,35kDa** | SRSF7 | Y189S192 | | | **12.0** | **22.9** | R.Y*FQS*PSR.S |
| **Splicing factor, arginine/serine-rich 7,35kDa** | SRSF7 | S223S225S227 | | | **100.0** | **24.1** | R.S*PS*GS*PRR.S |
| **Splicing factor, arginine/serine-rich 7,35kDa** | SRSF7 | S215S217 | | | **100.0** | **27.0** | R.S*PS*PKR.S |
| | SRSF7 | S231S233 | | | **100.0** | **16.2** | R.RS*AS*PER.M |
| **Splicing factor, arginine/serine-rich 7,35kDa** | SRSF7 | S181S183 | | | **34.8** | **31.7** | R.S*GS*IKGSR.Y |
| **Splicing factor, arginine/serine-rich 7,35kDa** | SRSF7 | S179S181S183 | | | **34.7** | **16.6** | R.S*RS*GS*IKGSR.Y |

**Peak Area**: <-10, -5, -3, 0, 3, 5, >10

**%CV**: 0, 29, 42, 71, 86, >100

**White dots-** Significant change in peptide abundance at **5**%FDR compared to the timepoint with the minimum peak area for a given PSM

| Protein Name | Gene | Phosphosites | Ascor | MOWSE | Sequence |
|---|---|---|---|---|---|
| **Splicing factor, arginine/serine-rich 7,35kDa** | SRSF7 | S173S175 | **100.0** | **18.4** | R.S*AS*LRR.S |
| **Splicing factor, arginine/serine-rich 7,35kDa** | SRSF7 | S202S204 | **100.0** | **19.7** | R.S*IS*RPR.S |
| **Splicing factor, arginine/serine-rich 7,35kDa** | SRSF7 | Y189S192S196 | **10.8** | **14.2** | R.Y*FQS*PSRS*R.S |
| **Splicing factor, arginine/serine-rich 7,35kDa** | SRSF7 | Y189S194 | **11.0** | **32.4** | R.Y*FQSPS*R.S |
| **Splicing factor, arginine/serine-rich 7,35kDa** | SRSF7 | S192S194 | **42.3** | **32.4** | R.YFQS*PS*R.S |
| **Splicing factor, arginine/serine-rich 7,35kDa** | SRSF7 | S155S157S159 | **100.0** | **11.1** | R.S*RS*AS*PR.R |
| **Splicing factor, arginine/serine-rich 7,35kDa** | SRSF7 | Y189S192S194 | **6.8** | **11.8** | R.Y*FQS*PS*RSR.S |
| **Splicing factor, arginine/serine-rich 7,35kDa** | SRSF7 | S213S215S217 | **100.0** | **12.9** | K.S*RS*PS*PKR.S |
| **Splicing factor, arginine/serine-rich 7,35kDa** | SRSF7 | S181S183S187 | **100.0** | **19.9** | R.S*GS*IKGS*R.Y |
| **Splicing factor, arginine/serine-rich 7,35kDa** | SRSF7 | S165 | **11.0** | **19.4** | R.S*ISLRR.S |
| **Splicing factor, arginine/serine-rich 7,35kDa** | SRSF7 | S192S194S196 | **10.8** | **22.4** | R.YFQS*PS*RS*R.S |
| **Splicing factor, arginine/serine-rich 7,35kDa** | SRSF7 | S171S173S175 | **100.0** | **12.5** | R.RS*RS*AS*LR.R |
| **Splicing factor, arginine/serine-rich 7,35kDa** | SRSF7 | Y189S194S196 | **8.9** | **11.3** | R.Y*FQSPS*RS*R.S |
| **Splicing factor, arginine/serine-rich 8** | SFSWAP | S604 | **8.1** | **12.7** | R.VKLDDDS*DDDEESKEGQESSSSAANTNPAVAPPCVVVEEK.K |
| **Splicing factor, arginine/serine-rich 8** | SFSWAP | S283 | **12.1** | **76.8** | K.SGVSS*DNEDDDDEEDGNYLHPSLFASK.K |
| **Splicing factor, arginine/serine-rich 8** | SFSWAP | S282 | **24.9** | **72.3** | K.SGVS*SDNEDDDDEEDGNYLHPSLFASK.K |
| **Splicing factor, arginine/serine-rich 8** | SFSWAP | S815 | **100.0** | **40.3** | R.AHS*PER.R |
| **SPT2, Suppressor of Ty, domain containing 1** | SPTY2D1 | Y597 | **52.0** | **51.0** | R.EYEEEDDDDDEY*DSEMEDFIEDEGEPQEEISK.H |
| **SPT2, Suppressor of Ty, domain containing 1** | SPTY2D1 | S599 | **13.0** | **69.3** | R.EYEEEDDDDDEYDS*EMEDFIEDEGEPQEEISK.H |
| **SPT6** | SUPT6H | S73S75S78 | **100.0** | **26.0** | K.GFINDDDDEDEGEEDEGS*DS*GDS*EDDVGHKK.R |
| **SPT6** | SUPT6H | S1703S1712 | **11.4** | **29.3** | R.LTPRPSPS*PMIESTPMS*IAGDATPLLDEMDR.- |
| **SPT6** | SUPT6H | T1532S1535 | **20.9** | **65.7** | R.TRT*PAS*INATPANINLADLTR.A |
| **SPT6** | SUPT6H | S1703T1718 | **20.1** | **48.8** | R.LTPRPSPS*PMIESTPMSIAGDAT*PLLDEMDR.- |
| **SPT6** | SUPT6H | S1701T1718 | **15.0** | **36.8** | R.LTPRPS*PSPMIESTPMSIAGDAT*PLLDEMDR.- |
| **SPT6** | SUPT6H | S1701S1712 | **6.5** | **32.4** | R.LTPRPS*PSPMIESTPMS*IAGDATPLLDEMDR.- |
| **SPT6** | SUPT6H | S1535 | **6.1** | **25.0** | R.TPAS*INATPANINLADLTR.A |
| **SPT6** | SUPT6H | T1532S1535T15 | **11.6** | **38.5** | R.TRT*PAS*INAT*PANINLADLTR.A |
| **SPT6** | SUPT6H | T1530S1535T15 | **13.1** | **13.8** | R.T*RTPAS*INAT*PANINLADLTR.A |
| **SPT6** | SUPT6H | T1530S1535 | **12.6** | **46.4** | R.T*RTPAS*INATPANINLADLTR.A |
| **Src like adapter protein 2** | SLA2 | S129 | **16.9** | **31.6** | R.RGS*YSLSVR.L |
| **SRP20** | SRSF3 | S148 | **100.0** | **33.2** | R.NHKPS*R.S |
| **SRP20** | SRSF3 | S128S130 | **100.0** | **14.4** | R.S*LS*RDR.R |
| **SRP20** | TXLNG | S115 | **100.0** | **21.7** | R.RRS*PR.R |
| **SRP20** | SRSF3 | S126S128S130 | **100.0** | **12.8** | R.S*RS*LS*RDR.R |
| **SRP20** | SRSF3 | S138S140 | **100.0** | **16.6** | R.ERS*LS*RER.N |
| **SRP20** | KMT2E | S120S122S124 | **100.0** | **16.6** | K.S*FS*RS*R.T |
| **SRp30c** | SRSF9 | S211 | **21.8** | **63.1** | R.GS*PHYFSPFRPY.- |
| **SRp30c** | SRSF9 | S204S208S211 | **10.5** | **22.0** | R.GRDS*PYQS*RGS*PHYFSPFRPY.- |
| **SRp30c** | SRSF9 | S211S216 | **25.3** | **35.7** | R.GS*PHYFS*PFRPY.- |
| **SRp30c** | SRSF9 | S204 | **37.8** | **43.9** | R.GRDS*PYQSR.G |
| **SRp30c** | SRSF9 | S189Y190 | **10.3** | **17.2** | R.STS*Y*GYSR.S |
| **SRp30c** | SRSF9 | S187T188 | **24.7** | **21.9** | R.S*T*SYGYSR.S |
| **SRp30c** | SRSF9 | Y214 | **7.7** | **46.3** | R.GSPHY*FSPFRPY.- |
| **SRp30c** | SRSF9 | T188 | **-0.1** | **31.4** | R.ST*SYGYSR.S |
| **SRp30c** | SRSF9 | S189Y192 | **15.4** | **23.9** | R.STS*YGY*SR.S |
| **SRp30c** | SRSF9 | S204Y206S211 | **8.0** | **12.6** | R.GRDS*PY*QSRGS*PHYFSPFRPY.- |
| **SRp30c** | SRSF9 | S216Y221 | **26.6** | **22.6** | R.GSPHYFS*PFRPY*.- |
| **SRp30c** | SRSF9 | S211Y214 | **23.7** | **33.0** | R.GS*PHY*FSPFRPY.- |
| **SRp30c** | SRSF9 | S189 | **16.9** | **35.1** | R.STS*YGYSR.S |
| **SRp30c** | SRSF9 | Y192 | **11.5** | **12.0** | R.STSYGY*SR.S |
| **SRp30c** | SRSF9 | Y214S216 | **-2.6** | **28.9** | R.GSPHY*FS*PFRPY.- |
| **SRp46** | SRSF8 | S100 | **100.0** | **21.2** | R.RDLPRS*R.Q |
| **SRP72** | SRP72 | S620 | **22.5** | **63.2** | K.TVS*SPPTSPRPGSAATVSASTSNIIPPR.H |
| **SRP72** | SRP72 | S621T624 | **16.6** | **92.1** | K.TVSS*PPT*SPRPGSAATVSASTSNIIPPR.H |
| **SRP72** | SRP72 | S621S625 | **13.7** | **92.5** | K.TVSS*PPTS*PRPGSAATVSASTSNIIPPR.H |
| **SRP72** | SRP72 | S621 | **8.0** | **69.6** | K.TVSS*PPTSPRPGSAATVSASTSNIIPPR.H |
| **SRP72** | SRP72 | S620S625 | **13.8** | **77.6** | K.TVS*SPPTS*PRPGSAATVSASTSNIIPPR.H |
| **SRP72** | SRP72 | S621S630 | **11.2** | **89.5** | K.TVSS*PPTSPRPGS*AATVSASTSNIIPPR.H |
| **SRP72** | SRP72 | S620S630 | **1.9** | **28.4** | K.TVS*SPPTSPRPGS*AATVSASTSNIIPPR.H |
| **SRP72** | SRP72 | T618S620 | **8.1** | **49.6** | K.T*VS*SPPTSPRPGSAATVSASTSNIIPPR.H |
| **SRP72** | SRP72 | S621S635 | **4.5** | **48.5** | K.TVSS*PPTSPRPGSAATVS*ASTSNIIPPR.H |
| **SRP72** | SRP72 | S630T633 | **8.0** | **31.4** | K.TVSSPPTSPRPGS*AAT*VSASTSNIIPPR.H |

Peak Area: <-10, -5, -3, 0, 3, 5, >10

%CV: 0, 29, 42, 71, 86, >100

**White dots-** Significant change in peptide abundance at **5** %FDR compared to the timepoint with the minimum peak area for a given PSM

| Protein Name | Gene | Phosphosites | CarT 0m | CarT 2m | CarT 5m | RajiB 0m | RajiB 2m | RajiB 5m | Ascor | MOWSE | Sequence |
|---|---|---|---|---|---|---|---|---|---|---|---|
| SRP72 | SRP72 | S621T633 | | | | | | | -1.7 | 24.5 | K.TVSS*PPTSPRPGSAAT*VSASTSNIIPPR.H |
| SRP72 | SRP72 | S625S630 | | | | | | | 7.9 | 36.6 | K.TVSSPPTS*PRPGS*AATVSASTSNIIPPR.H |
| SRP72 | SRP72 | T624 | | | | | | | 4.5 | 12.1 | K.TVSSPPT*SPRPGSAATVSASTSNIIPPR.H |
| SRP72 | SRP72 | S620S621 | | | | | | | -0.3 | 30.6 | K.TVS*S*PPTSPRPGSAATVSASTSNIIPPR.H |
| SRP72 | SRP72 | T624S625 | | | | | | | 1.6 | 13.7 | K.TVSSPPT*S*PRPGSAATVSASTSNIIPPR.H |
| SRRM2 | SRRM2 | S1876S1878T1 | | | | | | | 35.3 | 11.3 | R.S*RS*RT*PLISR.R |
| SRRM2 | SRRM2 | S2100S2102 | | | | | | | 23.8 | 13.8 | R.NHS*GS*RTPPVALNSSR.M |
| SRRM2 | SRRM2 | S771 | | | | | | | 8.2 | 36.4 | K.S*RLSLR.R |
| SRRM2 | SRRM2 | S1727S1729S1 | | | | | | | 100.0 | 36.0 | R.S*PS*VS*S*PEPAEK.S |
| | SRRM2 | S1320 | | | | | | | 29.8 | 42.5 | K.ELSNS*PLR.E |
| SRRM2 | SRRM2 | T1003 | | | | | | | 75.8 | 40.6 | K.AQT*PPGPSLSGSK.S |
| | SRRM2 | T1003S1014 | | | | | | | 29.4 | 31.6 | K.VKAQT*PPGPSLSGSKS*PCPQEK.S |
| SRRM2 | SRRM2 | T2104 | | | | | | | 66.3 | 53.5 | R.T*PPVALNSSR.M |
| SRRM2 | SRRM2 | T856S864 | | | | | | | 10.3 | 30.1 | R.QGSIT*SPQANEQS*VTPQRR.S |
| SRRM2 | SRRM2 | T856T866 | | | | | | | 11.6 | 29.2 | R.QGSIT*SPQANEQSVT*PQRR.S |
| SRRM2 | SRRM2 | S857S864 | | | | | | | 26.8 | 51.6 | R.QGSITS*PQANEQS*VTPQRR.S |
| | SRRM2 | T1856S1857 | | | | | | | 10.2 | 21.4 | R.SRT*S*PAPWK.R |
| SRRM2 | SRRM2 | S1857 | | | | | | | -0.2 | 41.9 | R.TS*PAPWK.R |
| SRRM2 | SRRM2 | S1318S1320 | | | | | | | 100.0 | 34.0 | K.ELS*NS*PLR.E |
| SRRM2 | SRRM2 | S1382S1387 | | | | | | | 24.6 | 53.3 | R.SSGHS*SSELS*PDAVEK.A |
| SRRM2 | SRRM2 | S2044S2046 | | | | | | | 100.0 | 30.3 | R.S*RS*PLAIR.R |
| SRRM2 | SRRM2 | S322 | | | | | | | 9.2 | 27.1 | R.GEGDAPFSEPGTTSTQRPS*SPETATK.Q |
| SRRM2 | SRRM2 | S2032T2034 | | | | | | | 100.0 | 22.0 | R.S*RT*PLLPR.K |
| SRRM2 | SRRM2 | S1541S1542S1 | | | | | | | 8.6 | 61.4 | R.SGS*S*QELDVKPSAS*PQER.S |
| SRRM2 | SRRM2 | S1539S1542S1 | | | | | | | 12.4 | 51.7 | R.S*GSS*QELDVKPSAS*PQER.S |
| SRRM2 | SRRM2 | S1102 | | | | | | | -0.2 | 52.2 | R.SS*SPVTELASR.S |
| SRRM2 | SRRM2 | S2398 | | | | | | | -0.2 | 55.2 | R.TS*PPLLDR.A |
| SRRM2 | SRRM2 | S322S323 | | | | | | | 4.5 | 16.8 | R.GEGDAPFSEPGTTSTQRPS*S*PETATK.Q |
| SRRM2 | SRRM2 | S2692S2694 | | | | | | | 12.3 | 43.3 | R.SLS*YS*PVER.R |
| SRRM2 | SRRM2 | S876 | | | | | | | 10.8 | 37.3 | R.SCFESS*PDPELK.S |
| SRRM2 | SRRM2 | T2289 | | | | | | | 55.4 | 70.2 | R.T*PTAPAVNLAGAR.T |
| SRRM2 | SRRM2 | S1099S1101S11 | | | | | | | 17.8 | 32.2 | R.S*RS*SS*PVTELASR.S |
| SRRM2 | SRRM2 | S1102S1103 | | | | | | | 12.2 | 61.0 | R.SS*S*PVTELASR.S |
| SRRM2 | SRRM2 | S2272 | | | | | | | 171.2 | 119.7 | R.TPAAAAAMNLAS*PR.T |
| SRRM2 | SRRM2 | S2690S2692S2 | | | | | | | 20.9 | 33.1 | R.S*LS*YS*PVER.R |
| SRRM2 | SRRM2 | S875S876 | | | | | | | 58.5 | 36.2 | R.SCFES*S*PDPELK.S |
| SRRM2 | SRRM2 | S2132 | | | | | | | 105.4 | 56.3 | R.S*PGMLEPLGSSR.T |
| SRRM2 | SRRM2 | S2118S2121 | | | | | | | 10.3 | 23.5 | R.MSCFS*RPS*MSPTPLDR.C |
| SRRM2 | SRRM2 | S2118S2123 | | | | | | | 9.3 | 25.4 | R.MSCFS*RPSMS*PTPLDR.C |
| SRRM2 | SRRM2 | S377T384S395 | | | | | | | 23.4 | 22.2 | R.HGGS*PQPLATT*PLSQEPVNPPS*EASPTR.D |
| SRRM2 | SRRM2 | S377S398 | | | | | | | 27.2 | 44.4 | R.HGGS*PQPLATTPLSQEPVNPPSEAS*PTR.D |
| SRRM2 | SRRM2 | S351S353S357 | | | | | | | 32.9 | 42.2 | K.SATRPS*PS*PERS*STGPEPPAPTPLLAER.H |
| | SRRM2 | S2449 | | | | | | | 18.0 | 28.5 | R.MGQAPSQSLLPPAQDQPRS*PVPSAFSDQSR.C |
| SRRM2 | SRRM2 | S1404 | | | | | | | 9.1 | 82.5 | K.AGMSSNQSISS*PVLDAVPR.T |
| SRRM2 | SRRM2 | S1403 | | | | | | | 45.8 | 70.7 | K.AGMSSNQSIS*SPVLDAVPR.T |
| SRRM2 | SRRM2 | S1329 | | | | | | | 37.7 | 66.0 | R.ENSFGS*PLEFR.N |
| SRRM2 | SRRM2 | S1398S1401 | | | | | | | 13.8 | 62.7 | K.AGMSS*NQS*ISSPVLDAVPR.T |
| SRRM2 | SRRM2 | S1401S1404 | | | | | | | 23.0 | 103.6 | K.AGMSSNQS*ISS*PVLDAVPR.T |
| SRRM2 | SRRM2 | S1318S1329 | | | | | | | 8.5 | 53.5 | K.ELS*NSPLRENSFGS*PLEFR.N |
| SRRM2 | SRRM2 | T1043 | | | | | | | 11.9 | 99.0 | K.SST*PPGESYFGVSSLQLK.G |
| SRRM2 | SRRM2 | S1042 | | | | | | | -0.2 | 90.1 | K.SS*TPPGESYFGVSSLQLK.G |
| SRRM2 | SRRM2 | S484S486 | | | | | | | 16.6 | 15.0 | R.S*RS*PATAK.R |
| SRRM2 | SRRM2 | S818S819 | | | | | | | 5.0 | 19.6 | R.SRSS*S*SPPPK.Q |
| SRRM2 | SRRM2 | S713S715T717 | | | | | | | 100.0 | 13.0 | R.S*HS*RT*PQR.R |
| SRRM2 | SRRM2 | S1219 | | | | | | | 23.9 | 36.7 | R.SGAGSS*PETK.E |
| SRRM2 | SRRM2 | S562S564S566 | | | | | | | 47.7 | 20.8 | R.S*HS*RS*PATR.G |
| SRRM2 | SRRM2 | T1231 | | | | | | | 34.7 | 72.1 | K.EQNSALPT*SSQDEELMEVVEK.S |
| | SRRM2 | S1818Y1820 | | | | | | | 6.4 | 46.7 | R.GGS*GY*HSR.S |
| SRRM2 | SRRM2 | S1232 | | | | | | | 5.2 | 55.7 | K.EQNSALPTS*SQDEELMEVVEK.S |
| SRRM2 | SRRM2 | S1318S1320S1 | | | | | | | 46.6 | 53.8 | K.ELS*NS*PLRENS*FGSPLEFR.N |
| SRRM2 | SRRM2 | S1318S1320S1 | | | | | | | 18.2 | 46.7 | K.ELS*NS*PLRENSFGS*PLEFR.N |
| SRRM2 | SRRM2 | S1014 | | | | | | | 100.0 | 25.3 | K.S*PCPQEK.S |

Peak Area: <-10, -5, -3, 0, 3, 5, >10

%CV: 0, 29, 42, 71, 86, >100

**White dots-** Significant change in peptide abundance at **5** %FDR compared to the timepoint with the minimum peak area for a given PSM

| Protein Name | Gene | Phosphosites | CarT 0m | CarT 2m | CarT 5m | RajiB 0m | RajiB 2m | RajiB 5m | Ascor | MOWSE | Sequence |
|---|---|---|---|---|---|---|---|---|---|---|---|
| **SRRM2** | SRRM2 | T627S629 | | | | | | | **100.0** | **12.1** | R.T*RS*PVR.R |
| **SRRM2** | SRRM2 | S1937T1939 | | | | | | | **9.8** | **18.6** | R.S*RT*PTTR.R |
| **SRRM2** | SRRM2 | S2426 | | | | | | | **39.3** | **31.5** | R.APS*PSSR.M |
| **SRRM2** | SRRM2 | S836S839 | | | | | | | **24.9** | **15.1** | R.QSHS*SSS*PHPK.V |
| **SRRM2** | SRRM2 | S1064 | | | | | | | **9.8** | **27.5** | K.GQSQTS*PDHR.S |
| **SRRM2** | SRRM2 | S295S297 | | | | | | | **34.5** | **20.3** | R.S*PS*PASGRR.G |
| **SRRM2** | SRRM2 | S1866S1869 | | | | | | | **31.0** | **37.4** | R.S*RAS*PATHR.R |
| **SRRM2** | SRRM2 | T1986 | | | | | | | **12.8** | **22.4** | R.SRT*SPVTR.R |
| **SRRM2** | SRRM2 | S970S973 | | | | | | | **5.9** | **13.5** | R.YSHS*GSS*SPDTK.V |
| **SRRM2** | SRRM2 | S902S908 | | | | | | | **8.7** | **24.9** | K.SS*TPPRQS*PSR.S |
| **SRRM2** | SRRM2 | S1842S1848 | | | | | | | **13.1** | **15.5** | R.S*RTPPTS*R.K |
| **SRRM2** | SRRM2 | S1682T1684 | | | | | | | **16.4** | **18.3** | K.TKS*RT*PPR.R |
| **SRRM2** | SRRM2 | S2727S2729 | | | | | | | **19.9** | **27.9** | R.GDS*RS*PSHKR.R |
| **SRRM2** | SRRM2 | S1913S1916 | | | | | | | **55.0** | **32.0** | R.S*RAS*PVSR.R |
| **SRRM2** | SRRM2 | T1892S1893 | | | | | | | **20.5** | **26.4** | R.SRT*S*PVSR.R |
| **SRRM2** | SRRM2 | S1112 | | | | | | | **100.0** | **21.4** | R.S*PIRQDR.G |
| **SRRM2** | SRRM2 | S1984S1987 | | | | | | | **16.6** | **19.5** | R.S*RTS*PVTR.R |
| **SRRM2** | SRRM2 | S1600S1601 | | | | | | | **20.0** | **55.9** | R.SGS*S*PEVK.D |
| **SRRM2** | SRRM2 | S454S456 | | | | | | | **19.9** | **36.6** | R.EIS*SS*PTSK.N |
| | SRRM2 | S1987 | | | | | | | **9.8** | **22.8** | R.SRTS*PVTR.R |
| **SRRM2** | SRRM2 | S454S455 | | | | | | | **13.0** | **29.2** | R.EIS*S*SPTSK.N |
| **SRRM2** | SRRM2 | S2702S2706 | | | | | | | **100.0** | **26.0** | R.RRPS*PQPS*PR.D |
| **SRRM2** | SRRM2 | S351S353 | | | | | | | **30.2** | **21.0** | K.SATRPS*PS*PER.S |
| **SRRM2** | SRRM2 | S1975 | | | | | | | **-0.1** | **30.4** | R.TS*PITR.R |
| **SRRM2** | SRRM2 | S454S455S456 | | | | | | | **24.0** | **34.5** | R.EIS*S*S*PTSK.N |
| **SRRM2** | SRRM2 | T1531 | | | | | | | **100.0** | **29.6** | R.T*PLGQR.S |
| **SRRM2** | SRRM2 | S1598S1600S1 | | | | | | | **100.0** | **27.5** | R.S*GS*S*PEVK.D |
| **SRRM2** | SRRM2 | S1577S1579S1 | | | | | | | **62.5** | **24.1** | R.S*RS*GS*S*PEVDSK.S |
| **SRRM2** | SRRM2 | S1923S1925T1 | | | | | | | **31.1** | **13.8** | R.S*RS*RT*PPVTR.R |
| **SRRM2** | SRRM2 | S1132 | | | | | | | **35.2** | **55.2** | K.SGMS*PEQSR.F |
| **SRRM2** | SRRM2 | S1925T1927 | | | | | | | **24.3** | **19.0** | R.S*RT*PPVTR.R |
| **SRRM2** | SRRM2 | S2407T2409 | | | | | | | **22.2** | **41.6** | R.S*RT*PPSAPSQSR.M |
| **SRRM2** | SRRM2 | S2407S2412 | | | | | | | **20.9** | **39.1** | R.S*RTPPS*APSQSR.M |
| **SRRM2** | SRRM2 | S1444 | | | | | | | **19.1** | **52.0** | R.SGSS*PGLR.D |
| **SRRM2** | SRRM2 | S759S761S763 | | | | | | | **100.0** | **17.4** | R.S*RS*LS*S*PR.S |
| **SRRM2** | SRRM2 | S1616S1618S1 | | | | | | | **100.0** | **23.6** | R.AQS*GS*DS*S*PEPK.A |
| **SRRM2** | SRRM2 | S1083 | | | | | | | **24.1** | **58.9** | R.QSHSES*PSLQSK.S |
| **SRRM2** | SRRM2 | S435S436S437 | | | | | | | **103.8** | **33.0** | R.HAS*S*S*PES*PKPAPAPGSHR.E |
| **SRRM2** | SRRM2 | S435S436S437 | | | | | | | **100.0** | **18.6** | R.HAS*S*S*PES*PKPAPAPGS*HR.E |
| **SRRM2** | SRRM2 | S1443S1444 | | | | | | | **27.9** | **43.3** | R.SGS*S*PGLR.D |
| **SRRM2** | SRRM2 | S1694 | | | | | | | **-0.1** | **34.4** | R.SS*PELTR.K |
| **SRRM2** | SRRM2 | S2581 | | | | | | | **19.8** | **41.7** | K.RVPS*PTPAPK.E |
| **SRRM2** | SRRM2 | S1064S1069T1 | | | | | | | **8.5** | **11.5** | K.GQSQTS*PDHRS*DT*SS*PEVR.Q |
| **SRRM2** | SRRM2 | S1691S1693 | | | | | | | **7.4** | **20.6** | R.SS*RS*SPELTR.K |
| **SRRM2** | SRRM2 | S1497S1499S1 | | | | | | | **13.7** | **24.6** | R.S*RS*PSS*PELNNK.C |
| **SRRM2** | SRRM2 | S778S783 | | | | | | | **10.1** | **38.3** | R.S*LSGSS*PCPK.Q |
| **SRRM2** | SRRM2 | S1064T1071S1 | | | | | | | **2.9** | **10.8** | K.GQSQTS*PDHRSDT*S*S*PEVR.Q |
| **SRRM2** | SRRM2 | S778S780S783 | | | | | | | **20.9** | **40.0** | R.S*LS*GSS*PCPK.Q |
| **SRRM2** | SRRM2 | T289S297 | | | | | | | **3.4** | **13.6** | K.THTT*ALAGRSPS*PASGR.R |
| **SRRM2** | SRRM2 | S2018S2020T2 | | | | | | | **100.0** | **15.5** | R.S*RS*RT*PPAIR.R |
| **SRRM2** | SRRM2 | S2020T2022 | | | | | | | **100.0** | **19.0** | R.S*RT*PPAIR.R |
| **SRRM2** | SRRM2 | T1453S1458S1 | | | | | | | **10.8** | **21.8** | R.DGSGT*PSRHS*LSGS*SPGMK.D |
| **SRRM2** | SRRM2 | S2688 | | | | | | | **40.2** | **33.7** | R.KPIDSLRDS*R.S |
| **SRRM2** | SRRM2 | S2684 | | | | | | | **5.1** | **27.6** | R.KPIDS*LRDSR.S |
| **SRRM2** | SRRM2 | S508S510 | | | | | | | **100.0** | **18.0** | R.S*RS*PQWR.R |
| **SRRM2** | SRRM2 | S950S952S954 | | | | | | | **33.6** | **18.7** | R.S*RS*VS*PCSNVESR.L |
| **SRRM2** | SRRM2 | S536 | | | | | | | **16.3** | **22.5** | R.S*PQRPGWSR.S |
| **SRRM2** | SRRM2 | S534S536 | | | | | | | **42.1** | **18.4** | R.S*RS*PQRPGWSR.S |
| **SRRM2** | SRRM2 | S952S954 | | | | | | | **46.7** | **48.6** | R.S*VS*PCSNVESR.L |
| **SRRM2** | SRRM2 | S1854T1856 | | | | | | | **24.3** | **16.0** | R.S*RT*SPAPWKR.S |
| **SRRM2** | SRRM2 | T2397 | | | | | | | **10.2** | **37.1** | R.T*SPPLLDR.A |
| **SRRM2** | SRRM2 | T315S322 | | | | | | | **27.8** | **18.1** | R.GEGDAPFSEPGT*TSTQRPS*SPETATK.Q |

Peak Area: <-10, -5, -3, 0, 3, 5, >10

%CV: 0, 29, 42, 71, 86, >100

**White dots-** Significant change in peptide abundance at **5** %FDR compared to the timepoint with the minimum peak area for a given PSM

CarT (0m, 2m, 5m) — RajiB (0m, 2m, 5m)

| Protein Name | Gene | Phosphosites | Ascor | MOWSE | Sequence |
|---|---|---|---|---|---|
| SRRM2 | SRRM2 | S1099S1101S11 | 10.2 | 27.7 | R.S*RS*S*SPVTELASR.S |
| SRRM2 | SRRM2 | S1188 | 87.4 | 21.9 | K.DKFS*PFPVQDRPESSLVFK.D |
| SRRM2 | SRRM2 | S817S818 | 6.8 | 11.1 | R.SRS*S*SSPPPK.Q |
| SRRM2 | SRRM2 | S2310S2314 | 11.5 | 57.7 | R.TPAALAALS*LTGS*GTPPTAANYPSSSR.T |
| SRRM2 | SRRM2 | S525S527 | 100.0 | 15.7 | R.S*RS*PQRR.G |
| SRRM2 | SRRM2 | S1866T1872 | 21.5 | 34.8 | R.S*RASPAT*HR.R |
| SRRM2 | SRRM2 | S1762S1764 | 100.0 | 17.7 | R.GRS*PS*PKPR.G |
| SRRM2 | SRRM2 | S472S474T476 | 22.2 | 21.9 | R.DKS*HS*HT*PSRR.M |
| SRRM2 | SRRM2 | S1984T1986S1 | 34.3 | 19.3 | R.S*RT*S*PVTR.R |
| SRRM2 | SRRM2 | T2409S2412 | 17.3 | 40.3 | R.SRT*PPS*APSQSR.M |
| SRRM2 | SRRM2 | S1972T1974S1 | 57.6 | 19.2 | R.S*RT*S*PITR.R |
| SRRM2 | SRRM2 | T1974S1975 | 12.8 | 18.9 | R.SRT*S*PITR.R |
| SRRM2 | SRRM2 | S1499S1501S1 | 17.7 | 26.0 | R.SRS*PS*S*PELNNK.C |
| SRRM2 | SRRM2 | S1497S1499S1 | 100.0 | 12.6 | R.S*RS*PS*S*PELNNK.C |
| SRRM2 | SRRM2 | S1690S1693S1 | 15.3 | 24.0 | R.S*SRS*S*PELTR.K |
| SRRM2 | SRRM2 | S774 | 9.9 | 41.3 | K.SRLS*LR.R |
| SRRM2 | SRRM2 | S2100S2102T2 | 49.0 | 35.8 | R.NHS*GS*RT*PPVALNSSR.M |
| SRRM2 | SRRM2 | S2067T2069S2 | 100.0 | 22.7 | R.S*LT*RS*PPAIR.R |
| SRRM2 | SRRM2 | S1101S1102 | 10.0 | 45.9 | R.S*S*SPVTELASR.S |
| SRRM2 | SRRM2 | S377T383S398 | 20.3 | 24.0 | R.HGGS*PQPLAT*TPLSQEPVNPPSEAS*PTR.D |
| SRRM2 | SRRM2 | S351S353S358 | 21.1 | 29.5 | K.SATRPS*PS*PERSS*TGPEPPAPTPLLAER.H |
| SRRM2 | SRRM2 | Y1049 | -0.3 | 52.1 | K.SSTPPGESY*FGVSSLQLK.G |
| SRRM2 | SRRM2 | T2302T2312 | 10.2 | 66.9 | R.T*PAALAALSLT*GSGTPPTAANYPSSSR.T |
| SRRM2 | SRRM2 | S817S819 | 8.1 | 16.8 | R.SRS*SS*SPPPK.Q |
| SRRM2 | SRRM2 | T1680T1684 | 5.1 | 18.5 | K.T*KSRT*PPR.R |
| SRRM2 | SRRM2 | T1844S1848 | 12.9 | 18.8 | R.SRT*PPTS*R.K |
| SRRM2 | SRRM2 | T1847S1848 | 23.1 | 29.2 | R.SRTPPT*S*R.K |
| SRRM2 | SRRM2 | T1986S1987 | 18.2 | 22.3 | R.SRT*S*PVTR.R |
| SRRM2 | SRRM2 | S1982S1984T1 | 6.5 | 19.1 | R.S*RS*RT*SPVTR.R |
| SRRM2 | SRRM2 | S1972T1974 | 16.0 | 15.5 | R.S*RT*SPITR.R |
| SRRM2 | SRRM2 | S435S436S437 | 4.1 | 12.5 | R.HAS*S*S*PESPKPAPAPGSHR.E |
| SRRM2 | SRRM2 | S1691S1694 | 4.3 | 26.9 | R.SS*RSS*PELTR.K |
| SRRM2 | SRRM2 | T288T289 | 7.8 | 14.4 | K.THT*T*ALAGRSPSPASGR.R |
| SRRM2 | SRRM2 | S2102T2104 | 8.8 | 16.4 | R.NHSGS*RT*PPVALNSSR.M |
| SRRM2 | SRRM2 | S1541S1550 | 9.3 | 14.0 | R.SGS*SQELDVKPS*ASPQER.S |
| SRRM2 | SRRM2 | S1539S1541S1 | 8.3 | 13.1 | R.SRS*GS*S*QELDVKPSAS*PQER.S |
| SRRM2 | SRRM2 | S351S353T359 | 19.7 | 18.5 | K.SATRPS*PS*PERSST*GPEPPAPTPLLAER.H |
| SRRM2 | SRRM2 | S1320S1329 | 9.3 | 61.6 | K.ELSNS*PLRENSFGS*PLEFR.N |
| SRRM2 | SRRM2 | S2314T2316 | 0.1 | 29.3 | R.TPAALAALSLTGS*GT*PPTAANYPSSSR.T |
| SRRM2 | SRRM2 | S837S838 | 15.2 | 16.2 | R.QSHSS*S*SPHPK.V |
| SRRM2 | SRRM2 | S1890T1892 | 14.6 | 24.5 | R.S*RT*SPVSR.R |
| SRRM2 | SRRM2 | S2090T2092 | 47.1 | 23.3 | R.S*AT*PPATR.N |
| SRRM2 | SRRM2 | S455S456 | 16.2 | 32.7 | R.EISS*S*PTSK.N |
| SRRM2 | SRRM2 | S1972S1975 | 7.7 | 22.1 | R.S*RTS*PITR.R |
| SRRM2 | SRRM2 | S1970T1974S1 | 5.1 | 15.7 | R.S*RSRT*S*PITR.R |
| SRRM2 | SRRM2 | T2738S2740 | 100.0 | 15.7 | R.RET*PS*PRPMR.H |
| SRRM2 | SRRM2 | S778S780 | 38.3 | 73.0 | R.S*LS*GSSPCPK.Q |
| SRRM2 | SRRM2 | S1441S1443S1 | 100.0 | 34.7 | R.S*GS*S*PGLR.D |
| SRRM2 | SRRM2 | T1453S1458S1 | 13.5 | 13.8 | R.DGSGT*PSRHS*LS*GSSPGMK.D |
| SRRM2 | SRRM2 | S1878T1880 | 45.0 | 13.6 | R.S*RT*PLISR.R |
| SRRM2 | SRRM2 | S1542S1550 | 14.7 | 11.5 | R.SGSS*QELDVKPS*ASPQER.S |
| SRRM2 | SRRM2 | S323 | 14.3 | 28.7 | R.GEGDAPFSEPGTTSTQRPSS*PETATK.Q |
| SRRM2 | SRRM2 | S2729S2731 | 7.7 | 17.0 | R.GDSRS*PS*HKR.R |
| SRRM2 | SRRM2 | T318S322 | 4.4 | 18.2 | R.GEGDAPFSEPGTTST*QRPS*SPETATK.Q |
| SRRM2 | SRRM2 | S1101S1102S11 | 5.0 | 32.1 | R.SRS*S*S*PVTELASR.S |
| SRRM2 | SRRM2 | S377S387T400 | 13.9 | 12.8 | R.HGGS*PQPLATTPLS*QEPVNPPSEASPT*R.D |
| SRRM2 | SRRM2 | S377S387S398 | 24.2 | 14.9 | R.HGGS*PQPLATTPLS*QEPVNPPSEAS*PTR.D |
| SRRM2 | SRRM2 | S1403T1413 | 10.8 | 22.8 | K.AGMSSNQSIS*SPVLDAVPRT*PSR.E |
| SRRM2 | SRRM2 | S1318S1326 | 12.1 | 48.2 | K.ELS*NSPLRENS*FGSPLEFR.N |
| SRRM2 | SRRM2 | T2302T2316 | 9.5 | 61.9 | R.T*PAALAALSLTGSGT*PPTAANYPSSSR.T |
| SRRM2 | SRRM2 | S1320S1326S1 | 11.2 | 34.3 | K.ELSNS*PLRENS*FGS*PLEFR.N |
| SRRM2 | SRRM2 | S837S839 | 12.8 | 12.2 | R.QSHSS*SS*PHPK.V |

Peak Area
<-10
-5
-3
0
3
5
>10

%CV
0
29
42
71
86
>100

**White dots-** Significant change in peptide abundance at **5** %FDR compared to the timepoint with the minimum peak area for a given PSM

CarT 0m 2m 5m | RajiB 0m 2m 5m

| Protein Name | Gene | Phosphosites | Ascor | MOWSE | Sequence |
|---|---|---|---|---|---|
| **SRRM2** | SRRM2 | S248S250T251 | **6.8** | **11.1** | R.S*RS*T*TPAPK.S |
| **SRRM2** | SRRM2 | S1984 | **8.8** | **17.3** | R.S*RTSPVTR.R |
| **SRRM2** | SRRM2 | S2749S2751 | **25.6** | **14.8** | R.HRSS*RS*P.- |
| **SRRM2** | SRRM2 | S1069T1071S1 | **10.5** | **12.9** | K.GQSQTSPDHRS*DT*S*S*PEVR.Q |
| **SRRM2** | SRRM2 | T315 | **0.7** | **16.9** | R.GEGDAPFSEPGT*TSTQRPSSPETATK.Q |
| **SRRM2** | SRRM2 | S1403S1404 | **2.1** | **52.3** | K.AGMSSNQSIS*S*PVLDAVPR.T |
| **SRRM2** | SRRM2 | S1320S1326 | **13.8** | **64.2** | K.ELSNS*PLRENS*FGSPLEFR.N |
| **SRRM2** | SRRM2 | S1233 | **1.0** | **67.8** | K.EQNSALPTSS*QDEELMEVVEK.S |
| **SRRM2** | SRRM2 | Y1820S1822 | **1.2** | **36.7** | R.GGSGY*HS*R.S |
| **SRRM2** | SRRM2 | Y1820 | **-0.3** | **15.7** | R.GGSGY*HSR.S |
| | SRRM2 | S790T792 | **100.0** | **14.0** | K.QKS*QT*PPRR.S |
| **SRRM2** | SRRM2 | S1672S1675 | **25.6** | **25.8** | R.RGS*RSS*PEPK.T |
| | SRRM2 | S1818S1822 | **16.1** | **38.9** | R.RGGS*GYHS*R.S |
| **SRRM2** | SRRM2 | S472S474S478 | **15.7** | **13.5** | R.DKS*HS*HTPS*RR.M |
| **SRRM2** | SRRM2 | S972S973 | **8.0** | **16.7** | R.YSHSGS*S*SPDTK.V |
| **SRRM2** | SRRM2 | T1892 | **-0.4** | **14.4** | R.SRT*SPVSR.R |
| **SRRM2** | SRRM2 | T1927 | **25.5** | **24.8** | R.T*PPVTR.R |
| **SRRM2** | SRRM2 | S759S761S763 | **-0.1** | **12.1** | R.S*RS*LS*SPR.S |
| **SRRM2** | SRRM2 | S780S783 | **14.1** | **41.4** | R.SLS*GSS*PCPK.Q |
| **SRRM2** | SRRM2 | S1451S1458S1 | **10.2** | **16.6** | R.DGS*GTPSRHS*LS*GSSPGMK.D |
| **SRRM2** | SRRM2 | S543 | **18.5** | **18.4** | R.SPQRPGWS*R.S |
| **SRRM2** | SRRM2 | T1856 | **5.4** | **13.7** | R.SRT*SPAPWKR.S |
| **SRRM2** | SRRM2 | T2034 | **100.0** | **37.5** | R.T*PLLPR.K |
| **SRRM2** | SRRM2 | S317S322 | **0.8** | **18.6** | R.GEGDAPFSEPGTTS*TQRPS*SPETATK.Q |
| **SRRM2** | SRRM2 | S2123 | **14.0** | **26.2** | R.MSCFSRPSMS*PTPLDR.C |
| **SRRM2** | SRRM2 | S377T384T400 | **21.5** | **15.6** | R.HGGS*PQPLATT*PLSQEPVNPPSEASPT*R.D |
| **SRRM2** | SRRM2 | S819S820 | **4.6** | **22.4** | R.SRSSS*S*PPPK.Q |
| **SRRM2** | SRRM2 | S486T489 | **3.2** | **14.4** | R.SRS*PAT*AKR.G |
| **SRRM2** | SRRM2 | S970S974 | **6.8** | **12.7** | R.YSHS*GSSS*PDTK.V |
| **SRRM2** | SRRM2 | S1497S1499 | **15.4** | **33.5** | R.S*RS*PSSPELNNK.C |
| **SRRM2** | SRRM2 | S2100T2104 | **7.7** | **21.5** | R.NHS*GSRT*PPVALNSSR.M |
| **SRRM2** | SRRM2 | S1103 | **12.6** | **55.9** | R.SSS*PVTELASR.S |
| **SRRM2** | SRRM2 | S875 | **10.8** | **34.6** | R.SCFES*SPDPELK.S |
| **SRRM2** | SRRM2 | S1124 | **7.8** | **13.7** | R.GEFSAS*PMLK.S |
| **SRRM2** | SRRM2 | S1401S1403 | **15.5** | **58.8** | K.AGMSSNQS*IS*SPVLDAVPR.T |
| **SRRM2** | SRRM2 | T1063 | **22.4** | **31.7** | K.GQSQT*SPDHR.S |
| **SRRM2** | SRRM2 | S1916 | **10.4** | **28.7** | R.SRAS*PVSR.R |
| **SRRM2** | SRRM2 | S1690S1691S1 | **13.5** | **16.4** | R.S*S*RSS*PELTR.K |
| **SRRM2** | SRRM2 | S1397 | **-0.0** | **17.1** | K.AGMS*SNQSISSPVLDAVPR.T |
| **SRRM2** | SRRM2 | S2310 | **13.6** | **61.9** | R.TPAALAALS*LTGSGTPPTAANYPSSSR.T |
| **SRRM2** | SRRM2 | T1844T1847 | **7.1** | **14.3** | R.SRT*PPT*SR.K |
| **SRRM2** | SRRM2 | S895 | **6.0** | **15.8** | R.HSCSGSS*PPR.V |
| **SRRM2** | SRRM2 | S1923S1925T1 | **13.8** | **13.3** | R.S*RS*RTPPVT*R.R |
| **SRRM2** | SRRM2 | T1974 | **8.2** | **12.6** | R.SRT*SPITR.R |
| **SRRM2** | SRRM2 | S1541S1552 | **4.4** | **29.2** | R.SGS*SQELDVKPSAS*PQER.S |
| **SRRM2** | SRRM2 | S377T384S398 | **23.4** | **19.5** | R.HGGS*PQPLATT*PLSQEPVNPPSEAS*PTR.D |
| **SRRM2** | SRRM2 | S377S395 | **11.7** | **19.2** | R.HGGS*PQPLATTPLSQEPVNPPS*EASPTR.D |
| **SRRM2** | SRRM2 | S1691S1693S1 | **7.5** | **13.5** | R.SS*RS*S*PELTR.K |
| **SRRM2** | SRRM2 | S1537S1539S1 | **25.7** | **11.1** | R.S*RS*GS*SQELDVKPS*ASPQER.S |
| **SRRM2** | SRRM2 | S1541S1542S1 | **7.7** | **44.8** | R.SGS*S*QELDVKPS*ASPQER.S |
| **SRRM2** | SRRM2 | S2115S2118 | **3.7** | **10.7** | R.MS*CFS*RPSM#SPTPLDR.C |
| **SRRM2** | SRRM2 | T2312S2314 | **9.9** | **36.7** | R.TPAALAALSLT*GS*GTPPTAANYPSSSR.T |
| **SRRM2** | SRRM2 | S596T598 | **8.8** | **10.8** | R.S*RT*PTR.R |
| **SRRM2** | SRRM2 | S295S300 | **8.6** | **12.8** | R.S*PSPAS*GRR.G |
| **SRRM2** | SRRM2 | T903S908 | **18.1** | **21.7** | K.SST*PPRQS*PSR.S |
| **SRRM2** | SRRM2 | S1674S1675 | **5.0** | **11.1** | R.RGSRS*S*PEPK.T |
| **SRRM2** | SRRM2 | S1890S1893 | **9.1** | **20.5** | R.S*RTS*PVSR.R |
| **SRRM2** | SRRM2 | S1984T1986 | **10.8** | **17.0** | R.S*RT*SPVTR.R |
| **SRRM2** | SRRM2 | S857T866 | **13.1** | **45.0** | R.QGSITS*PQANEQSVT*PQRR.S |
| **SRRM2** | SRRM2 | S1539S1541S1 | **7.6** | **13.3** | R.SRS*GS*S*QELDVKPS*ASPQER.S |
| **SRRM2** | SRRM2 | S1404T1413 | **14.0** | **35.4** | K.AGMSSNQSISS*PVLDAVPRT*PSR.E |
| **SRRM2** | SRRM2 | S815S817 | **3.6** | **12.2** | R.S*RS*SSSPPPK.Q |

Peak Area: <-10, -5, -3, 0, 3, 5, >10

%CV: 0, 29, 42, 71, 86, >100

**White dots-** Significant change in peptide abundance at **5** %FDR compared to the timepoint with the minimum peak area for a given PSM

| Protein Name | Gene | Phosphosites | CarT 0m | CarT 2m | CarT 5m | RajiB 0m | RajiB 2m | RajiB 5m | Ascor | MOWSE | Sequence |
|---|---|---|---|---|---|---|---|---|---|---|---|
| **SRRM2** | SRRM2 | S1218S1219 | | | | | | | **22.5** | **23.1** | R.SGAGS*S*PETK.E |
| **SRRM2** | SRRM2 | S1061 | | | | | | | | **15.7** | K.GQS*QTSPDHR.S |
| **SRRM2** | SRRM2 | S1443 | | | | | | | **8.9** | **24.5** | R.SGS*SPGLR.D |
| **SRRM2** | SRRM2 | S377S395S398 | | | | | | | **14.4** | **11.2** | R.HGGS*PQPLATTPLSQEPVNPPS*EAS*PTR.D |
| **SRRM2** | SRRM2 | S1048 | | | | | | | **8.2** | **48.0** | K.SSTPPGES*YFGVSSLQLK.G |
| **SRRM2** | SRRM2 | S972S974 | | | | | | | **8.2** | **10.8** | R.YSHSGS*SS*PDTK.V |
| **SRRM2** | SRRM2 | S1869T1872 | | | | | | | **5.1** | **35.0** | R.SRAS*PAT*HR.R |
| **SRRM2** | SRRM2 | S902T903 | | | | | | | **-0.1** | **15.3** | K.SS*T*PPRQSPSR.S |
| **SRRM2** | SRRM2 | S1842T1844 | | | | | | | **13.9** | **19.3** | R.S*RT*PPTSR.K |
| **SRRM2** | SRRM2 | S1598S1600 | | | | | | | **13.5** | **41.2** | R.S*GS*SPEVK.D |
| **SRRM2** | SRRM2 | S1982S1984S1 | | | | | | | **2.3** | **11.4** | R.S*RS*RTS*PVTR.R |
| **SRRM2** | SRRM2 | S780S782S783 | | | | | | | **0.6** | **11.5** | R.SLS*GS*S*PCPK.Q |
| **SRRM2** | SRRM2 | S1398S1403 | | | | | | | **4.7** | **48.2** | K.AGMSS*NQSIS*SPVLDAVPR.T |
| **SRRM2** | SRRM2 | S1318 | | | | | | | **10.2** | **14.6** | K.ELS*NSPLRENSFGSPLEFR.N |
| **SRRM2** | SRRM2 | S575T577 | | | | | | | **100.0** | **12.4** | R.S*RT*PAR.T |
| **SRRM2** | SRRM2 | S818S820 | | | | | | | **3.0** | **15.7** | R.SRSS*SS*PPPK.Q |
| **SRRM2** | SRRM2 | T1939T1941 | | | | | | | **3.5** | **12.7** | R.SRT*PT*TR.R |
| **SRRM2** | SRRM2 | T348S353 | | | | | | | **13.3** | **15.6** | K.SAT*RPSPS*PER.S |
| **SRRM2** | SRRM2 | S377T383T400 | | | | | | | **18.2** | **16.4** | R.HGGS*PQPLAT*TPLSQEPVNPPSEASPT*R.D |
| **SRRM2** | SRRM2 | S1318S1326S1 | | | | | | | **9.4** | **21.5** | K.ELS*NSPLRENS*FGS*PLEFR.N |
| **SRRM2** | SRRM2 | S836S838 | | | | | | | **29.7** | **13.6** | R.QSHS*SS*SPHPK.V |
| **SRRM2** | SRRM2 | S817S820 | | | | | | | **-0.5** | **12.3** | R.SRS*SSS*PPPKQK.S |
| **SRRM2** | SRRM2 | S1902S1905 | | | | | | | **7.6** | **15.2** | R.S*RTS*VTR.R |
| **SRRM2** | SRRM2 | S1818S1822S1 | | | | | | | **15.4** | **11.2** | R.GGS*GYHS*RS*PAR.Q |
| **SRRM2** | SRRM2 | S892 | | | | | | | **19.8** | **18.9** | R.HSCS*GSSPPR.V |
| **SRRM2** | SRRM2 | S761S763S764 | | | | | | | **-0.1** | **16.5** | R.SRS*LS*S*PR.S |
| **SRRM2** | SRRM2 | S1081 | | | | | | | **13.4** | **14.9** | R.QSHS*ESPSLQSK.S |
| **SRRM2** | SRRM2 | S1690S1691 | | | | | | | **11.5** | **32.9** | R.S*S*RSSPELTR.K |
| **SRRM2** | SRRM2 | S1854S1857 | | | | | | | **14.2** | **16.1** | R.S*RTS*PAPWKR.S |
| **STAM binding protein** | STAMBP | Y114 | | | | | | | | **14.5** | K.AELLKRY*TK.E |
| **STARD3 N terminal like** | STARD3NL | S193 | | | | | | | **100.0** | **24.4** | R.LLIVQDAS*ER.A |
| **START domain containing 10** | STARD10 | S284 | | | | | | | **27.8** | **37.2** | R.MGGAGGEGS*DDDTSLT.- |
| **STAT3** | STAT3 | T716S727 | | | | | | | **25.1** | **29.7** | K.FICVTPT*TCSNTIDLPMS*PR.T |
| **STAT3** | STAT3 | S719S727 | | | | | | | **26.1** | **40.3** | K.FICVTPTTCS*NTIDLPMS*PR.T |
| **STAT3** | STAT3 | T714S727 | | | | | | | **34.2** | **50.4** | K.FICVT*PTTCSNTIDLPMS*PR.T |
| **STAT3** | STAT3 | S727 | | | | | | | **31.6** | **21.0** | K.FICVTPTTCSNTIDLPMS*PR.T |
| **STAT3** | STAT3 | T721S727 | | | | | | | **19.0** | **19.6** | K.FICVTPTTCSNT*IDLPMS*PR.T |
| **STAT4** | STAT4 | S708S721 | | | | | | | **23.2** | **21.6** | R.SDS*TEPHSPSDLLPMS*PSVYAVLR.E |
| **STAT4** | STAT4 | S713S721 | | | | | | | **21.3** | **40.8** | R.SDSTEPHS*PSDLLPMS*PSVYAVLR.E |
| **STAT5B** | STAT5B | S193 | | | | | | | **100.0** | **11.0** | R.IQAQFGPLAQLS*PQER.L |
| **Stathmin 1** | STMN1 | S38 | | | | | | | **38.8** | **38.3** | K.ESVPEFPLS*PPK.K |
| **Stathmin 1** | STMN1 | S25 | | | | | | | **114.5** | **72.1** | R.ASGQAFELILS*PR.S |
| **Stathmin 1** | STMN1 | S16 | | | | | | | **72.0** | **83.7** | R.AS*GQAFELILSPR.S |
| **Stathmin 1** | STMN1 | S16S25 | | | | | | | **100.0** | **65.8** | R.AS*GQAFELILS*PR.S |
| **Stathmin 1** | STMN1 | S31 | | | | | | | **25.0** | **20.8** | R.SKES*VPEFPLSPPK.K |
| **Stathmin 1** | STMN2 | S46 | | | | | | | **100.0** | **19.5** | K.DLS*LEEIQK.K |
| **Stathmin 1** | STMN1 | S63 | | | | | | | **100.0** | **30.4** | K.S*HEAEVLK.Q |
| **Staufen** | STAU1 | T277 | | | | | | | **19.1** | **22.5** | K.TKPIVKPQT*SPEYGQGINPISR.L |
| **Staufen RNA binding protein homolog 2** | STAU2 | S416 | | | | | | | **6.5** | **53.1** | K.DMNQPS*SSFFSISPTSNSSATIAR.E |
| **Staufen RNA binding protein homolog 2** | STAU2 | S453S460 | | | | | | | **16.7** | **13.8** | K.GS*SPTPPCS*PVQPSK.Q |
| **Staufen RNA binding protein homolog 2** | STAU2 | S423 | | | | | | | **-0.6** | **59.1** | K.DMNQPSSSFFSIS*PTSNSSATIAR.E |
| **STE20 like kinase MST1** | STK4 | S320 | | | | | | | **99.3** | **90.1** | R.EVDQDDEENS*EEDEMDSGTMVR.A |
| **STE20 like kinase MST1** | STK4 | S320T329 | | | | | | | **32.6** | **50.6** | R.EVDQDDEENS*EEDEMDSGT*MVR.A |
| **Step II splicing factor SLU7** | SLU7 | S235 | | | | | | | **19.5** | **53.6** | K.DHNS*EDEDEDKYADDIDMPGQNFDSK.R |
| **Stress induced phosphoprotein 1** | STIP1 | S16 | | | | | | | **113.5** | **105.8** | K.ALS*VGNIDDALQCYSEAIK.L |
| **Striatin calmodulin binding protein** | STRN | S134 | | | | | | | **0.1** | **32.9** | K.YGTELNQGDMKPPS*YDSDEGNETEVQPQQNSQLMWK.Q |
| **Striatin calmodulin binding protein** | STRN | T143 | | | | | | | **5.9** | **30.8** | K.YGTELNQGDMKPPSYDSDEGNET*EVQPQQNSQLMWK.Q |
| **Striatin calmodulin binding protein** | STRN | S137 | | | | | | | **16.2** | **35.3** | K.YGTELNQGDMKPPSYDS*DEGNETEVQPQQNSQLMWK.Q |
| **Striatin calmodulin binding protein** | STRN | S245 | | | | | | | **71.3** | **150.2** | K.FLESAAADFS*DEDEDDDVDGR.E |
| **Striatin calmodulin binding protein 4** | STRN4 | S276 | | | | | | | **28.8** | **18.6** | R.LGGSVLGQIPFLQNCEDEDS*DEDDELDSVQHKK.Q |
| **Striatin, calmodulin binding protein 3** | STRN3 | S229 | | | | | | | **100.0** | **51.2** | K.NLEQILNGGES*PK.Q |
| **Striatin, calmodulin binding protein 3** | STRN3 | S257 | | | | | | | **36.8** | **108.3** | R.SSGDVLETFNFLENADDS*DEDEENDMIEGIPEGK.D |

Peak Area: <-10, -5, -3, 0, 3, 5, >10

%CV: 0, 29, 42, 71, 86, >100

**White dots-** Significant change in peptide abundance at **5** %FDR compared to the timepoint with the minimum peak area for a given PSM

| Protein Name | Gene | Phosphosites | CarT 0m | CarT 2m | CarT 5m | RajiB 0m | RajiB 2m | RajiB 5m | Ascor | MOWSE | Sequence |
|---|---|---|---|---|---|---|---|---|---|---|---|
| Stromal antigen 2 | STAG2 | S1058S1061S1 | | | | | | | 10.7 | 69.4 | R.NSLLAGGDDDTMS*VIS*GIS*SR.G |
| Stromal antigen 2 | STAG2 | S1061S1064 | | | | | | | 7.1 | 26.5 | R.NSLLAGGDDDTMSVIS*GIS*SR.G |
| Stromal antigen 2 | STAG2 | S1047S1061S1 | | | | | | | 11.9 | 26.3 | R.NS*LLAGGDDDTMSVIS*GIS*SR.G |
| Stromal antigen 2 | STAG2 | T1056S1061S1 | | | | | | | 12.6 | 80.1 | R.NSLLAGGDDDT*MSVIS*GIS*SR.G |
| Stromal antigen 2 | STAG2 | S1058 | | | | | | | 16.1 | 34.7 | R.NSLLAGGDDDTMS*VISGISSR.G |
| Stromal antigen 2 | STAG2 | S1058S1061 | | | | | | | 13.1 | 57.5 | R.NSLLAGGDDDTMS*VIS*GISSR.G |
| Stromal antigen 2 | STAG2 | T1056S1058S1 | | | | | | | 13.5 | 37.4 | R.NSLLAGGDDDT*MS*VIS*GISSR.G |
| Stromal antigen 2 | STAG2 | T1056S1058S1 | | | | | | | 11.7 | 32.7 | R.NSLLAGGDDDT*MS*VISGIS*SR.G |
| Stromal antigen 2 | STAG2 | T1056S1058 | | | | | | | 13.5 | 109.4 | R.NSLLAGGDDDT*MS*VISGISSR.G |
| Stromal antigen 2 | STAG2 | S1047S1064 | | | | | | | 10.9 | 28.0 | R.NS*LLAGGDDDTMSVISGIS*SR.G |
| Stromal interaction molecule 1 | STIM1 | S618 | | | | | | | 7.3 | 62.4 | R.SHS*PSSPDPDTPSPVGDSR.A |
| Stromal interaction molecule 1 | STIM1 | S521 | | | | | | | 18.7 | 65.4 | R.DLTHSDS*ESSLHMSDR.Q |
| Stromal interaction molecule 1 | STIM1 | S575 | | | | | | | 82.3 | 28.4 | R.LIEGVHPGSLVEKLPDS*PALAK.K |
| Stromal interaction molecule 1 | STIM1 | S519 | | | | | | | 17.2 | 54.7 | R.DLTHS*DSESSLHMSDR.Q |
| Stromal interaction molecule 1 | STIM1 | S521S523 | | | | | | | 6.5 | 26.7 | R.DLTHSDS*ES*SLHMSDR.Q |
| Stromal interaction molecule 1 | STIM1 | S621 | | | | | | | 5.8 | 24.8 | R.SHSPSS*PDPDTPSPVGDSR.A |
| Stromal interaction molecule 1 | STIM1 | S519S521S524 | | | | | | | 5.3 | 30.5 | R.DLTHS*DS*ESS*LHMSDR.Q |
| Stromal interaction molecule 1 | STIM1 | S257 | | | | | | | 100.0 | 71.8 | R.AEQS*LHDLQER.L |
| Stromal interaction molecule 1 | STIM1 | S519S521 | | | | | | | 0.6 | 17.1 | R.DLTHS*DS*ESSLHMSDR.Q |
| Stromal interaction molecule 1 | STIM1 | S512 | | | | | | | 4.4 | 11.9 | R.LTEPQHGLGS*QR.D |
| Stromal interaction molecule 1 | STIM1 | T504 | | | | | | | 5.1 | 30.7 | R.LT*EPQHGLGSQR.D |
| Stromal interaction molecule 1 | STIM1 | S519S521S523 | | | | | | | 14.4 | 16.8 | R.DLTHS*DS*ES*SLHMSDR.Q |
| Stromal interaction molecule 1 | STIM1 | S519S523S524 | | | | | | | 2.3 | 15.4 | R.DLTHS*DSES*S*LHMSDR.Q |
| Stromal interaction molecule 1 | STIM1 | S620 | | | | | | | 5.8 | 43.3 | R.SHSPS*SPDPDTPSPVGDSR.A |
| Stromal interaction molecule 2 | STIM2 | S767 | | | | | | | 35.2 | 67.0 | K.SCS*MNQLSSGIPVPKPR.H |
| Stromal membrane associated protein 1 | SMAP1 | S152 | | | | | | | 15.3 | 97.2 | K.NAIAITNISSSDAPLQPLVSS*PSLQAAVDK.N |
| Stromal membrane associated protein 1 | SMAP1 | S151 | | | | | | | | 11.4 | K.NAIAITNISSSDAPLQPLVS*SPSLQAAVDK.N |
| Stromal membrane-associated protein 1-like | SMAP2 | S240 | | | | | | | 43.7 | 69.5 | R.KVVGSMPTAGSAGS*VPENLNLFPEPGSK.S |
| Stromal membrane-associated protein 1-like | SMAP2 | S219 | | | | | | | 12.2 | 41.7 | K.DLDLLASVPS*PSSSGSR.K |
| | SSRP1 | S667S668S671 | | | | | | | 16.7 | 68.3 | K.SKEFVS*S*DES*S*SGENK.S |
| Structure specific recognition protein 1 | SSRP1 | S667S668S671 | | | | | | | 29.3 | 24.1 | K.EFVS*S*DES*S*S*GENKSK.K |
| Structure specific recognition protein 1 | SSRP1 | S444 | | | | | | | 62.9 | 82.2 | K.EGMNPSYDEYADS*DEDQHDAYLER.M |
| Structure specific recognition protein 1 | SSRP1 | S437S444 | | | | | | | 25.4 | 47.1 | K.EGMNPS*YDEYADS*DEDQHDAYLER.M |
| Structure specific recognition protein 1 | SSRP1 | Y441 | | | | | | | 3.5 | 61.3 | K.EGM#NPSYDEY*ADSDEDQHDAYLER.M |
| Structure specific recognition protein 1 | SSRP1 | S667S668S671 | | | | | | | 7.9 | 33.3 | K.EFVS*S*DES*SS*GENK.S |
| SUDD | RIOK3 | S125S127S128 | | | | | | | 3.2 | 40.7 | R.KVHPYEDS*DS*S*EDEVDWQDTR.D |
| SUDD | RIOK3 | Y122S127S128 | | | | | | | 6.3 | 15.3 | R.KVHPY*EDSDS*S*EDEVDWQDTR.D |
| SUMO 1 specific protease 1 | SENP6 | S335S336 | | | | | | | 64.7 | 66.8 | R.KTSLSDLNDPIILS*S*DDDDDNDR.T |
| SUMO1 activating enzyme subunit 2 | UBA2 | S565 | | | | | | | | 70.5 | K.S*ITNGSDDGAQPSTSTAQEQDDVLIVDSDEEDSSNNADVSEEER.S |
| Supervillin | SVIL | S1000 | | | | | | | 100.0 | 20.6 | R.RGS*LER.A |
| SUPT5H | SUPT5H | S32S36 | | | | | | | 37.8 | 57.3 | R.S*AAGS*EKEEEPEDEEEEEEEEEYDEEEEEEDDDRPPKKPR.H |
| | SUPT5H | S666 | | | | | | | 76.7 | 57.8 | R.DVTNFTVGGFAPMS*PR.I |
| Surfeit 2 | SURF2 | S155S156S163 | | | | | | | 14.1 | 45.0 | R.EAFWEPTS*S*DEGGAAS*DDS*MTDLYPPELFTR.K |
| Surfeit 2 | SURF2 | S155S163S166 | | | | | | | 8.6 | 27.1 | R.EAFWEPTS*SDEGGAAS*DDS*MT*DLYPPELFTR.K |
| Surfeit 2 | SURF2 | S163S166 | | | | | | | 14.6 | 58.5 | R.EAFWEPTSSDEGGAAS*DDS*MTDLYPPELFTR.K |
| Surfeit 2 | SURF2 | S155S156 | | | | | | | 10.6 | 37.6 | R.EAFWEPTS*S*DEGGAASDDSMTDLYPPELFTR.K |
| Surfeit 2 | SURF2 | T154S155 | | | | | | | 8.8 | 24.7 | R.EAFWEPT*S*SDEGGAASDDSMTDLYPPELFTR.K |
| Surfeit 2 | SURF2 | S155S166 | | | | | | | 6.3 | 22.5 | R.EAFWEPTS*SDEGGAASDDS*MTDLYPPELFTR.K |
| Surfeit 2 | SURF2 | T154S155S156 | | | | | | | 5.9 | 35.7 | R.EAFWEPT*S*S*DEGGAAS*DDSMTDLYPPELFTR.K |
| Surfeit 2 | SURF2 | T154S155S156 | | | | | | | -0.8 | 21.7 | R.EAFWEPT*S*S*DEGGAASDDSMT*DLYPPELFTR.K |
| Surfeit 2 | SURF2 | S155S156S166 | | | | | | | 1.5 | 13.2 | R.EAFWEPTS*S*DEGGAASDDS*MT*DLYPPELFTR.K |
| Surfeit 2 | SURF2 | S166T168 | | | | | | | 2.9 | 13.4 | R.EAFWEPTSSDEGGAASDDS*MT*DLYPPELFTR.K |
| Surfeit 2 | SURF2 | T190T195 | | | | | | | 23.9 | 62.5 | R.KDLGSTEDGDGT*DDFLT*DKEDEK.A |
| Surfeit 2 | SURF2 | S156S166 | | | | | | | 7.2 | 20.3 | R.EAFWEPTSS*DEGGAASDDS*MTDLYPPELFTR.K |
| Surfeit 2 | SURF2 | S156S163 | | | | | | | 5.7 | 14.1 | R.EAFWEPTSS*DEGGAAS*DDSMTDLYPPELFTR.K |
| Surfeit 2 | SURF2 | S155S156S163 | | | | | | | 5.8 | 32.8 | R.EAFWEPTS*S*DEGGAAS*DDSMT*DLYPPELFTR.K |
| Surfeit 2 | SURF2 | T154S155S163 | | | | | | | 6.0 | 38.1 | R.EAFWEPT*S*SDEGGAAS*DDS*MTDLYPPELFTR.K |
| Survival of motor neuron 1, telomeric | SMN2 | T25 | | | | | | | 7.4 | 36.5 | R.RGT*GQSDDSDIWDDTALIK.A |
| Survival of motor neuron 1, telomeric | SMN2 | T25S31 | | | | | | | 28.5 | 53.4 | R.RGT*GQSDDS*DIWDDTALIK.A |
| Survival of motor neuron 1, telomeric | SMN2 | S28 | | | | | | | 19.8 | 150.8 | R.GTGQS*DDSDIWDDTALIK.A |
| Survival of motor neuron 1, telomeric | SMN2 | T25S28S31 | | | | | | | 38.7 | 19.8 | R.RGT*GQS*DDS*DIWDDTALIK.A |
| Survival of motor neuron 1, telomeric | SMN2 | S28S31 | | | | | | | 47.3 | 166.2 | R.GTGQS*DDS*DIWDDTALIK.A |

Peak Area: <-10, -5, -3, 0, 3, 5, >10

%CV: 0, 29, 42, 71, 86, >100

**White dots-** Significant change in peptide abundance at **5** %FDR compared to the timepoint with the minimum peak area for a given PSM

| Protein Name | Gene | Phosphosites | CarT 0m | CarT 2m | CarT 5m | RajiB 0m | RajiB 2m | RajiB 5m | Ascor | MOWSE | Sequence |
|---|---|---|---|---|---|---|---|---|---|---|---|
| **Survival of motor neuron 1, telomeric** | SMN2 | S166 | | | | | | | 12.0 | 48.1 | R.EEQNLSDLLSPICEVANNIEQNAQENENESQVS*TDESENSR.S |
| **Survival of motor neuron 1, telomeric** | SMN2 | S139 | | | | | | | | 22.5 | R.EEQNLS*DLLSPICEVANNIEQNAQENENESQVSTDESENSR.S |
| **Survival of motor neuron 1, telomeric** | SMN2 | S163S166T167 | | | | | | | 34.7 | 30.7 | R.EEQNLSDLLSPICEVANNIEQNAQENENES*QVS*T*DES*ENSR.S |
| **Survival of motor neuron 1, telomeric** | SMN2 | S139S143S163 | | | | | | | | 23.8 | R.EEQNLS*DLLS*PICEVANNIEQNAQENENES*QVS*TDESENSR.S |
| **Survival of motor neuron 1, telomeric** | SMN2 | T167 | | | | | | | | 33.6 | R.EEQNLSDLLSPICEVANNIEQNAQENENESQVST*DESENSR.S |
| **Survival of motor neuron 1, telomeric** | SMN2 | T25S28 | | | | | | | 3.0 | 35.7 | R.GT*GQS*DDSDIWDDTALIK.A |
| **Survival of motor neuron 1, telomeric** | SMN2 | S139S143S163 | | | | | | | | 13.6 | R.EEQNLS*DLLS*PICEVANNIEQNAQENENES*QVS*T*DESENSR.S |
| **SVH protein** | ARMC10 | S45 | | | | | | | 74.9 | 82.6 | K.S*AGALEEGTSEGQLCGR.S |
| **SWI/SNF related matrix associated actin dependent regulator of chromatin subfamily** | HLTF | S398S400T405 | | | | | | | 10.3 | 38.3 | K.TAVQYIESS*DS*EEIET*SELPQK.M |
| **SWI/SNF related matrix associated actin dependent regulator of chromatin subfamily** | HLTF | S397S398S400 | | | | | | | 20.8 | 26.2 | K.TAVQYIES*S*DS*EEIETSELPQK.M |
| **SWI/SNF-related matrix-associated actin-dependent regulator of chromatin subfamily** | SMARCA5 | S47 | | | | | | | | 66.5 | K.GGPEGVAAQAVAS*AASAGPADAEMEEIFDDASPGKQK.E |
| **SWI/SNF-related matrix-associated actin-dependent regulator of chromatin subfamily** | SMARCA5 | S66 | | | | | | | 134.8 | 128.0 | K.GGPEGVAAQAVASAASAGPADAEMEEIFDDAS*PGK.Q |
| **SWI/SNF-related matrix-associated actin-dependent regulator of chromatin subfamily** | SMARCA5 | S50 | | | | | | | 6.9 | 27.6 | K.GGPEGVAAQAVASAAS*AGPADAEMEEIFDDASPGKQK.E |
| **SYK** | SYK | S307 | | | | | | | -0.1 | 11.5 | R.KSS*PAQGNR.Q |
| **Symplekin** | SYMPK | S1243 | | | | | | | 25.6 | 79.2 | R.S*PQTLAPVGEDAMK.T |
| **Symplekin** | SYMPK | S494 | | | | | | | 53.1 | 98.9 | R.RLS*AQGQAISVVGSLSSMSPLEEEAPQAK.R |
| **Synapse associated protein 1** | SYAP1 | T248 | | | | | | | 100.0 | 49.4 | K.T*PPVVIK.S |
| **Synapse associated protein 1** | SYAP1 | S273 | | | | | | | 6.5 | 93.4 | K.TQEDEEEISTSPGVS*EFVSDAFDACNLNQEDLR.K |
| **Synapse associated protein 1** | SYAP1 | T268 | | | | | | | 12.1 | 113.6 | K.TQEDEEEIST*SPGVSEFVSDAFDACNLNQEDLR.K |
| **Synaptojanin 1** | SYNJ1 | S1049S1053 | | | | | | | 13.8 | 42.5 | R.TS*PCQS*PTISEGPVPSLPIRPSR.A |
| **Synaptojanin 2** | SYNJ2 | S1124 | | | | | | | 15.7 | 47.3 | K.SAS*DASISSGTHGQYSILQTAR.L |
| **Synaptosomal associated protein, 23 kd** | SNAP23 | S110 | | | | | | | 73.1 | 82.8 | K.TTWGDGGENS*PCNVVSK.Q |
| **Synaptosomal associated protein, 23 kd** | SNAP23 | S161 | | | | | | | 58.0 | 107.1 | R.EDEMEENLTQVGS*ILGNLK.D |
| **Synaptotagmin like 3** | SYTL3 | S452 | | | | | | | 100.0 | 15.9 | K.S*PVLRK.Q |
| **Synembryn** | RIC8A | Y441T447 | | | | | | | 32.4 | 76.9 | R.GLMAGGRPEGQY*SEDEDT*DTDEYKEAK.A |
| **Synembryn** | RIC8A | Y441S442 | | | | | | | 3.9 | 66.9 | R.GLMAGGRPEGQY*S*EDEDTDTDEYKEAK.A |
| **Synembryn** | RIC8A | S442T447 | | | | | | | 29.0 | 56.9 | R.GLMAGGRPEGQYS*EDEDT*DTDEYK.E |
| **Synembryn** | RIC8A | S529S530S534 | | | | | | | 68.2 | 109.9 | R.GHLTSLQDAMCETMEQQLS*S*DPDS*DPD.- |
| **Synembryn** | RIC8A | S530S534 | | | | | | | 16.9 | 49.7 | R.GHLTSLQDAMCETMEQQLSS*DPDS*DPD.- |
| **Synembryn** | RIC8A | S529S534 | | | | | | | 24.3 | 37.1 | R.GHLTSLQDAMCETMEQQLS*SDPDS*DPD.- |
| **Synembryn** | RIC8A | T523S529S534 | | | | | | | 15.6 | 50.1 | R.GHLTSLQDAMCET*MEQQLS*SDPDS*DPD.- |
| **Synembryn** | RIC8A | S529S530 | | | | | | | 8.7 | 28.2 | R.GHLTSLQDAMCETMEQQLS*S*DPDSDPD.- |
| **Syntaxin 4A** | STX4 | S14S15 | | | | | | | 100.0 | 26.5 | R.QGDDS*S*DEEDKER.V |
| **Syntaxin 7** | STX7 | S205 | | | | | | | 65.7 | 59.4 | K.DLGMMIHEQGDVIDS*IEANVENAEVHVQQANQQLSR.A |
| **Syntrophin beta 2** | SNTB2 | S393 | | | | | | | 44.8 | 75.5 | R.S*PSLGSDLTFATR.T |
| **Syntrophin beta 2** | SNTB2 | S233 | | | | | | | 13.9 | 44.1 | K.KPSLVSDLPWEGAAPQSPSFSGSEDSGS*PK.H |
| **Syntrophin beta 2** | SNTB2 | S231 | | | | | | | 12.6 | 58.0 | K.KPSLVSDLPWEGAAPQSPSFSGSEDS*GSPK.H |
| **Syntrophin beta 2** | SNTB2 | S228 | | | | | | | 7.3 | 20.9 | K.KPSLVSDLPWEGAAPQSPSFSGS*EDSGSPK.H |
| **Syntrophin beta 2** | SNTB2 | S222S231 | | | | | | | 12.4 | 22.4 | K.KPSLVSDLPWEGAAPQS*PSFSGSEDS*GSPK.H |
| **Syntrophin beta 2** | SNTB2 | S222S233 | | | | | | | 17.1 | 24.0 | K.KPSLVSDLPWEGAAPQS*PSFSGSEDSGS*PK.H |
| **T box brain 2** | EOMES | S596 | | | | | | | -0.3 | 40.8 | R.TS*PTVFSEDQLSK.E |
| **T box brain 2** | EOMES | S107 | | | | | | | 65.4 | 90.4 | R.KGS*PCGEEELPSAAAAAAAAAAAAAAATAR.Y |
| **T cell antigen receptor, zeta** | | Y72 | | | | | | | 33.9 | 50.9 | R.SADAPAYQQGQNQLY*NELNLGR.R |
| **T cell lymphoma invasion and metastasis 2** | TIAM2 | S1583 | | | | | | | 55.2 | 89.7 | K.QGS*PTKDIEIQFQR.L |
| **T Complex 1** | TCP1 | S544 | | | | | | | 12.1 | 38.0 | K.HGS*YEDAVHSGALND.- |
| **T Complex 1** | TCP1 | S551 | | | | | | | 42.0 | 14.5 | K.HGSYEDAVHS*GALND.- |
| **T Complex 1** | TCP1 | Y545 | | | | | | | 9.8 | 14.0 | K.HGSY*EDAVHSGALND.- |
| **TAB2** | TAB2 | S524 | | | | | | | -0.0 | 20.3 | R.KLS*MGSDDAAYTQALLVHQK.A |
| **TAB2** | TAB2 | S527 | | | | | | | 2.6 | 15.7 | R.KLSMGS*DDAAYTQALLVHQK.A |
| **TACC3** | TACC3 | S402 | | | | | | | | 36.5 | R.GS*YHLDWDK.M |
| **TAF15** | TAF15 | S226 | | | | | | | 24.0 | 76.1 | R.TDADS*ESDNSDNNTIFVQGLGEGVSTDQVGEFFK.Q |
| **TAF15** | TAF15 | T235 | | | | | | | 6.1 | 119.2 | R.TDADSESDNSDNNT*IFVQGLGEGVSTDQVGEFFK.Q |
| **TAF15** | TAF15 | T222 | | | | | | | | 55.7 | R.T*DADSESDNSDNNTIFVQGLGEGVSTDQVGEFFK.Q |
| **TAF15** | TAF15 | S231 | | | | | | | 19.8 | 110.0 | R.TDADSESDNS*DNNTIFVQGLGEGVSTDQVGEFFK.Q |
| **TAF15** | TAF15 | S228 | | | | | | | 2.0 | 68.6 | R.TDADSES*DNSDNNTIFVQGLGEGVSTDQVGEFFK.Q |
| **TAF15** | TAF15 | S226T235 | | | | | | | 1.0 | 15.7 | R.TDADS*ESDNSDNNT*IFVQGLGEGVSTDQVGEFFK.Q |
| **TAF15** | TAF15 | S231T235 | | | | | | | 3.3 | 53.6 | R.TDADSESDNS*DNNT*IFVQGLGEGVSTDQVGEFFK.Q |
| **TAF15** | TAF15 | S226S231T235 | | | | | | | -0.5 | 60.1 | R.TDADS*ESDNS*DNNT*IFVQGLGEGVSTDQVGEFFK.Q |
| **TAF15** | TAF15 | T222S226S228 | | | | | | | 3.3 | 60.5 | R.T*DADS*ES*DNSDNNTIFVQGLGEGVSTDQVGEFFK.Q |
| **TAF15** | TAF15 | T222S226 | | | | | | | 2.2 | 25.9 | R.T*DADS*ESDNSDNNTIFVQGLGEGVSTDQVGEFFK.Q |
| **TAF15** | TAF15 | S226S231 | | | | | | | 4.4 | 47.1 | R.TDADS*ESDNS*DNNTIFVQGLGEGVSTDQVGEFFK.Q |
| **TAF15** | TAF15 | S226S228S231 | | | | | | | 0.7 | 29.5 | R.TDADS*ES*DNS*DNNTIFVQGLGEGVSTDQVGEFFK.Q |

**Peak Area**: <-10, -5, -3, 0, 3, 5, >10

**%CV**: 0, 29, 42, 71, 86, >100

**White dots-** Significant change in peptide abundance at **5** %FDR compared to the timepoint with the minimum peak area for a given PSM

| Protein Name | Gene | Phosphosites | CarT 0m | CarT 2m | CarT 5m | RajiB 0m | RajiB 2m | RajiB 5m | Ascor | MOWSE | Sequence |
|---|---|---|---|---|---|---|---|---|---|---|---|
| **TAF15** | TAF15 | S226S228T235 | | | | | | | **1.5** | **13.5** | R.TDADS*ES*DNSDNNT*IFVQGLGEGVSTDQVGEFFK.Q |
| **TAF15** | TAF15 | S226S228 | | | | | | | **0.7** | **24.7** | R.TDADS*ES*DNSDNNTIFVQGLGEGVSTDQVGEFFK.Q |
| **TAF3 RNA polymerase II, TATA box binding protein (TBP)-associated factor, 140kDa** | TAF3 | S236S243 | | | | | | | **53.0** | **11.6** | K.IPPMLSPVHVQDS*TDLAPPS*PEPPMLAPVAK.S |
| **TAF3 RNA polymerase II, TATA box binding protein (TBP)-associated factor, 140kDa** | TAF3 | T237S243 | | | | | | | **24.8** | **27.2** | K.IPPMLSPVHVQDST*DLAPPS*PEPPMLAPVAK.S |
| **TAF6 like RNA polymerase II** | TAF6L | S495S501 | | | | | | | **7.6** | **19.3** | R.KMPQLTASAIVS*PHGDES*PR.G |
| **TAF9 RNA polymerase II** | TAF9 | T154S158 | | | | | | | **17.9** | **61.7** | R.LSVGSVT*SRPS*TPTLGTPTPQTMSVSTK.V |
| **TAF9 RNA polymerase II** | TAF9 | S152T154 | | | | | | | **14.8** | **52.6** | R.LSVGS*VT*SRPSTPTLGTPTPQTMSVSTK.V |
| **TAF9 RNA polymerase II** | TAF9 | S152T159 | | | | | | | **7.0** | **75.1** | R.LSVGS*VTSRPST*PTLGTPTPQTMSVSTK.V |
| **TAF9 RNA polymerase II** | TAF9 | S155T159 | | | | | | | **10.1** | **74.7** | R.LSVGSVTS*RPST*PTLGTPTPQTMSVSTK.V |
| **TAF9 RNA polymerase II** | TAF9 | T154S155 | | | | | | | **-5.2** | **31.3** | R.LSVGSVT*S*RPSTPTLGTPTPQTM#SVSTK.V |
| **TAF9 RNA polymerase II** | TAF9 | T154T159 | | | | | | | **12.8** | **77.6** | R.LSVGSVT*SRPST*PTLGTPTPQTMSVSTK.V |
| **TAF9 RNA polymerase II** | TAF9 | S152T161 | | | | | | | **13.1** | **50.7** | R.LSVGS*VTSRPSTPT*LGTPTPQTMSVSTK.V |
| **TAF9 RNA polymerase II** | TAF9 | S152T159T164 | | | | | | | **11.9** | **36.7** | R.LSVGS*VTSRPST*PTLGT*PTPQTMSVSTK.V |
| **TAF9 RNA polymerase II** | TAF9 | S152S158T164 | | | | | | | **8.7** | **50.7** | R.LSVGS*VTSRPS*TPTLGT*PTPQTMSVSTK.V |
| **TAF9 RNA polymerase II** | TAF9 | S152S158T161 | | | | | | | **7.7** | **31.3** | R.LSVGS*VTSRPS*TPT*LGTPTPQTMSVSTK.V |
| **TAF9 RNA polymerase II** | TAF9 | S158T159T161 | | | | | | | **4.7** | **18.0** | R.LSVGSVTSRPS*T*PT*LGTPTPQTMSVSTK.V |
| **TAF9 RNA polymerase II** | TAF9 | S149S152T154 | | | | | | | **7.5** | **58.5** | R.LS*VGS*VT*S*RPSTPTLGTPTPQTMSVSTK.V |
| **TAF9 RNA polymerase II** | TAF9 | T154T161 | | | | | | | **3.5** | **70.8** | R.LSVGSVT*SRPSTPT*LGTPTPQTMSVSTK.V |
| **TAF9 RNA polymerase II** | TAF9 | S152S158 | | | | | | | **4.0** | **54.8** | R.LSVGS*VTSRPS*TPTLGTPTPQTMSVSTK.V |
| **TAF9 RNA polymerase II** | TAF9 | S149T161 | | | | | | | **5.7** | **45.6** | R.LS*VGSVTSRPSTPT*LGTPTPQTMSVSTK.V |
| **TAF9 RNA polymerase II** | TAF9 | S149T154S155 | | | | | | | **6.3** | **44.3** | R.LS*VGSVT*S*RPSTPTLGTPTPQTMSVSTK.V |
| **TAF9 RNA polymerase II** | TAF9 | S155T161 | | | | | | | **18.8** | **47.2** | R.LSVGSVTS*RPSTPT*LGTPTPQTMSVSTK.V |
| **TAF9 RNA polymerase II** | TAF9 | S149S158 | | | | | | | **6.3** | **33.7** | R.LS*VGSVTSRPS*TPTLGTPTPQTMSVSTK.V |
| **TAF9 RNA polymerase II** | TAF9 | S158T159 | | | | | | | **10.3** | **54.8** | R.LSVGSVTSRPS*T*PTLGTPTPQTMSVSTK.V |
| **TAF9 RNA polymerase II** | TAF9 | S158T159T164 | | | | | | | **10.4** | **32.8** | R.LSVGSVTSRPS*T*PTLGT*PTPQTMSVSTK.V |
| **TAF9 RNA polymerase II** | TAF9 | S149S158T159 | | | | | | | **5.7** | **45.9** | R.LS*VGSVTSRPS*T*PTLGTPTPQTMSVSTK.V |
| **TAF9 RNA polymerase II** | TAF9 | S155S158T159 | | | | | | | **9.0** | **23.2** | R.LSVGSVTS*RPS*T*PTLGTPTPQTMSVSTK.V |
| **TAF9 RNA polymerase II** | TAF9 | T154S155S158 | | | | | | | **9.5** | **31.2** | R.LSVGSVT*S*RPS*T*PTLGTPTPQTMSVSTK.V |
| **TAF9 RNA polymerase II** | TAF9 | S155S158 | | | | | | | **13.6** | **52.6** | R.LSVGSVTS*RPS*TPTLGTPTPQTMSVSTK.V |
| **TAF9L** | TAF9B | S152S153 | | | | | | | **9.6** | **33.5** | R.LSVGAVS*S*KPTTPTIATPQTVSVPNK.V |
| **TAF9L** | TAF9B | S152T156 | | | | | | | **9.6** | **51.3** | R.LSVGAVS*SKPT*TPTIATPQTVSVPNK.V |
| **Talin** | TLN1 | S425 | | | | | | | **9.1** | **55.4** | K.SKDHFGLEGDEESTMLEDSVS*PK.K |
| **TANC** | TANC1 | S63S66 | | | | | | | **29.7** | **67.3** | K.GVSMS*LPS*SPLLPR.Q |
| **TANC** | TANC1 | S63S67 | | | | | | | **16.4** | **52.0** | K.GVSMS*LPSS*PLLPR.Q |
| **Tankyrase 1 binding protein 182KD** | TNKS1BP1 | S1620S1621 | | | | | | | **151.0** | **76.3** | R.VPS*S*DEEVVEEPQSR.R |
| **Tankyrase 1 binding protein 182KD** | TNKS1BP1 | S494S498 | | | | | | | **20.1** | **81.2** | R.LDS*PPPS*PITEASEAAEAAEAGNLAVSSR.E |
| **Tankyrase 1 binding protein 182KD** | TNKS1BP1 | S1554 | | | | | | | **17.1** | **75.2** | R.SPS*QDFSFIEDTEILDSAMYR.S |
| **Tankyrase 1 binding protein 182KD** | TNKS1BP1 | S672 | | | | | | | **79.7** | **39.7** | R.AS*PEPPGPESSSR.W |
| **Tankyrase 1 binding protein 182KD** | TNKS1BP1 | S1024 | | | | | | | **31.2** | **36.7** | R.GS*GGLFSPSTAHVPDGALGQR.D |
| **Tankyrase 1 binding protein 182KD** | TNKS1BP1 | T501 | | | | | | | **-0.1** | **17.2** | R.LDSPPPSPIT*EASEAAEAAEAGNLAVSSR.E |
| **Tankyrase 1 binding protein 182KD** | TNKS1BP1 | T501S504 | | | | | | | **5.5** | **18.8** | R.LDSPPPSPIT*EAS*EAAEAAEAGNLAVSSR.E |
| **Tankyrase 1 binding protein 182KD** | TNKS1BP1 | S872 | | | | | | | **43.4** | **11.6** | R.DS*LGTYSSR.D |
| **Tankyrase 1 binding protein 182KD** | TNKS1BP1 | S1552 | | | | | | | **-2.7** | **44.9** | R.S*PSQDFSFIEDTEILDSAMYR.S |
| **TAO kinase 1** | TAOK1 | S965 | | | | | | | **100.0** | **25.6** | R.NS*PQALR.R |
| **TAO kinase 1** | TAOK1 | S9 | | | | | | | **100.0** | **30.9** | R.AGS*LKDPEIAELFFK.E |
| **TAPP1** | PLEKHA1 | S332 | | | | | | | **35.1** | **74.9** | R.SNS*LVSTFTMEK.R |
| **TAPP1** | PLEKHA1 | S330 | | | | | | | **12.2** | **66.8** | R.S*NSLVSTFTMEK.R |
| **TAR (HIV) RNA binding protein 2** | TARBP2 | S156 | | | | | | | **-0.2** | **48.6** | R.SPPMELQPPVSPQQS*ECNPVGALQELVVQK.G |
| **TAR (HIV) RNA binding protein 2** | TARBP2 | S142S156 | | | | | | | **6.4** | **36.9** | R.S*PPMELQPPVSPQQS*ECNPVGALQELVVQK.G |
| **TAR (HIV) RNA binding protein 2** | TARBP2 | S152S156 | | | | | | | **2.6** | **38.5** | R.SPPMELQPPVS*PQQS*ECNPVGALQELVVQK.G |
| **TAR (HIV) RNA binding protein 2** | TARBP2 | S142S152 | | | | | | | **6.5** | **38.9** | R.S*PPMELQPPVS*PQQSECNPVGALQELVVQK.G |
| **TAR (HIV) RNA binding protein 2** | TARBP2 | S152 | | | | | | | **6.0** | **59.4** | R.SPPMELQPPVS*PQQSECNPVGALQELVVQK.G |
| **Tara like protein** | TRIOBP | T248 | | | | | | | **100.0** | **13.6** | R.ART*PAR.T |
| **Target of EGR1** | TOE1 | S5 | | | | | | | **46.0** | **79.8** | M.AADS*DDGAVSAPAASDGGVSK.S |
| **Target of myb1 like 2** | TOM1L2 | S378 | | | | | | | **77.6** | **52.3** | R.KQS*SEGIPVAQPSVMDDIEVWLR.T |
| **TASK 5** | KCNK15 | S5 | | | | | | | **100.0** | **13.5** | M.RRPS*VR.A |
| **Taste receptor type 2 member 7** | TAS2R7 | S305 | | | | | | | **100.0** | **10.9** | K.VM#S*ILK.G |
| **TATA element modulatory factor 1** | TMF1 | S344 | | | | | | | **51.5** | **69.0** | R.SVSEINS*DDELSGK.G |
| **TatD DNase domain containing 2** | TATDN2 | S328S329S331 | | | | | | | **57.4** | **35.7** | R.EVVMEHPS*S*GS*DWS*DVEEISTVR.F |
| **TATSF1** | HTATSF1 | S453 | | | | | | | **-0.2** | **17.4** | K.ESS*PEKEAEEGCPEKESEEGCPK.R |
| **TATSF1** | HTATSF1 | S702 | | | | | | | **100.0** | **52.8** | K.LFEDDDS*NEK.L |
| **TATSF1** | HTATSF1 | S721 | | | | | | | **100.0** | **62.5** | K.LFDDS*DER.G |

Peak Area: <-10, -5, -3, 0, 3, 5, >10

%CV: 0, 29, 42, 71, 86, >100

**White dots-** Significant change in peptide abundance at **5** %FDR compared to the timepoint with the minimum peak area for a given PSM

CarT (0m, 2m, 5m) — RajiB (0m, 2m, 5m)

| Protein Name | Gene | Phosphosites | Ascor | MOWSE | Sequence |
|---|---|---|---|---|---|
| **TATSF1** | HTATSF1 | S624 | **110.5** | **123.0** | R.EFDEDS*DEKEEEEDTYEK.V |
| **TATSF1** | HTATSF1 | S452S453 | **8.1** | **34.5** | K.TEDGGEFEEGASENNAKES*S*PEKEAEEGCPEK.E |
| **TATSF1** | HTATSF1 | S642 | **83.7** | **62.0** | K.VFDDES*DEKEDEEYADEK.G |
| **TATSF1** | HTATSF1 | S676 | **100.0** | **70.5** | K.LFEES*DDKEDEDADGKEVEDADEK.L |
| **TATSF1** | HTATSF1 | S597S600 | **119.5** | **87.2** | K.ELEENDS*ENS*EFEDDGSEK.V |
| **TATSF1** | HTATSF1 | S579 | **100.0** | **35.2** | K.DLDEEGS*EKELHENVLDK.E |
| **TATSF1** | HTATSF1 | S597S600S607 | **100.0** | **86.7** | K.ELEENDS*ENS*EFEDDGS*EK.V |
| **TATSF1** | HTATSF1 | S597S600S607 | **100.0** | **70.5** | K.ELEENDS*ENS*EFEDDGS*EKVLDEEGS*ER.E |
| **TATSF1** | HTATSF1 | S713S714 | **63.7** | **75.6** | K.LFDEEEDS*S*EKLFDDSDER.G |
| **TATSF1** | HTATSF1 | S713S714S721 | **100.0** | **57.7** | K.LFDEEEDS*S*EKLFDDS*DER.G |
| **TATSF1** | HTATSF1 | S702S713S714 | **65.6** | **59.2** | K.LFEDDDS*NEKLFDEEEDS*S*EKLFDDSDER.G |
| **TATSF1** | HTATSF1 | S702S713S714 | **100.0** | **49.0** | K.LFEDDDS*NEKLFDEEEDS*S*EKLFDDS*DER.G |
| **TATSF1** | HTATSF1 | S387 | **17.4** | **45.0** | R.SDS*VSASER.A |
| **TATSF1** | HTATSF1 | S616 | **100.0** | **55.5** | K.VLDEEGS*ER.E |
| **TATSF1** | HTATSF1 | S702S713 | **14.9** | **48.9** | K.LFEDDDS*NEKLFDEEEDS*SEK.L |
| **TATSF1** | HTATSF1 | S498 | **29.7** | **23.3** | R.GSEEDS*PKK.E |
| **TATSF1** | HTATSF1 | S385 | **9.2** | **30.1** | R.S*DSVSASER.A |
| **TATSF1** | HTATSF1 | T408 | **8.9** | **16.4** | R.HFSEHPST*SK.M |
| **TATSF1** | HTATSF1 | S714S721 | **6.8** | **16.2** | K.LFDEEEDSS*EKLFDDS*DER.G |
| **TATSF1** | HTATSF1 | S385S387 | **30.2** | **12.6** | R.S*DS*VSASER.A |
| **TATSF1** | HTATSF1 | T726S732S748 | **3.8** | **19.6** | R.GT*LGGFGS*VEEGPLSTGSSFILSS*DDDDDDI.- |
| **Taxilin** | TXLNA | S515 | **21.2** | **66.0** | R.RPEGPGAQAPSS*PR.V |
| **Taxilin** | TXLNA | S514 | | **41.0** | R.RPEGPGAQAPS*SPR.V |
| **TBC1 domain family member 10B** | TBC1D10B | T148T150 | **16.2** | **23.5** | K.TEEARPSPAPGPGT*PT*GTPTR.T |
| **TBC1 domain family member 15 (Fragment)** | TBC1D15 | S104 | **14.6** | **53.3** | K.SLS*QSFENLLDEPAYGLIQK.I |
| **TBC1 domain family member 15 (Fragment)** | TBC1D15 | S106 | **19.1** | **115.1** | K.SLSQS*FENLLDEPAYGLIQK.I |
| **TBC1 domain family member 15 (Fragment)** | TBC1D15 | S102 | **8.4** | **51.0** | K.S*LSQSFENLLDEPAYGLIQK.I |
| **TBC1 domain family member 15 (Fragment)** | TBC1D15 | S102S104 | **14.5** | **23.8** | K.S*LS*QSFENLLDEPAYGLIQK.I |
| **TBC1 domain family member 22A** | TBC1D22A | S167 | **15.1** | **121.5** | R.SQS*LPHSATVTLGGTSDPSTLSSSALSER.E |
| **TBC1 domain family member 22A** | TBC1D22A | T173 | **9.1** | **120.2** | R.SQSLPHSAT*VTLGGTSDPSTLSSSALSER.E |
| **TBC1 domain family member 22A** | TBC1D22A | S171 | **10.3** | **121.5** | R.SQSLPHS*ATVTLGGTSDPSTLSSSALSER.E |
| **TBC1 domain family member 22A** | TBC1D22A | S180 | **2.0** | **35.4** | R.SQSLPHSATVTLGGTS*DPSTLSSSALSER.E |
| **TBC1 domain family member 23** | TBC1D23 | S383 | **100.0** | **29.5** | R.MS*FNLPWPDR.S |
| **TBC1 domain family member 2B** | TBC1D2B | S957 | **34.0** | **47.8** | R.DTSPDKGELVS*DEEEDT.- |
| **TBC1 domain family member 4** | TBC1D4 | S570 | **13.9** | **87.0** | R.SLTSS*LENIFSR.G |
| **TBC1 domain family member 4** | TBC1D4 | S588 | **37.7** | **32.8** | R.LGS*VDSFER.S |
| **TBC1 domain family member 4** | TBC1D4 | S588S591 | **100.0** | **36.9** | R.LGS*VDS*FER.S |
| **TBC1 domain family member 5** | TBC1D5 | S43 | **-0.5** | **82.4** | R.TS*STLDSEGTFNSYR.K |
| **TBC1 domain family member 5** | TBC1D5 | S522 | **28.9** | **51.4** | K.SES*MPVQLNK.G |
| **TBC1 domain family member 5** | TBC1D5 | S44 | **-0.3** | **66.7** | R.TSS*TLDSEGTFNSYR.K |
| **TBC1 domain family, member 10C** | TBC1D10C | S438 | **15.2** | **41.0** | R.GS*TSFLDTRF.- |
| **TBC1 domain family, member 15** | TBC1D15 | S675 | **14.0** | **36.3** | R.NDS*PTQIPVSSDVCR.L |
| **TBC1 domain family, member 15** | TBC1D15 | S633 | | **53.2** | K. [illegible] |
| **TBC1 domain family, member 17** | TBC1D17 | S602S604S608 | **20.2** | **15.4** | R.QLTACPELPHNVQEILGLAPPAEPHS*PS*PTAS*PLPLS*PTR.A |
| **TBC1 domain family, member 9 (with GRAM domain)** | TBC1D9 | T1008 | **86.6** | **11.6** | R.NYLRLWT*PENKSK.S |
| **TBC1D1** | TBC1D1 | S237 | **35.1** | **30.3** | K.SFS*QPGLR.S |
| **TBC1D1** | TBC1D1 | S565S585 | **18.2** | **23.7** | K.LLGS*SEDLSSDSESHLPEEPAPLS*PQQAFR.R |
| **TBC1D1** | TBC1D1 | T505 | **4.1** | **51.2** | R.SLT*ESLESILSR.G |
| **TBC1D1** | TBC1D1 | S565S570 | **5.0** | **23.8** | K.LLGS*SEDLS*SDSESHLPEEPAPLSPQQAFR.R |
| **TBP-interacting protein** | CAND1 | Y304T306 | | **43.9** | K. [illegible] |
| **TCR interacting molecule** | TRAT1 | S116 | **18.0** | **21.5** | K.MQEATPSAQATNETQMCYASLDHS*VK.G |
| **Telomerase subunit EST1A** | SMG6 | T479S484 | **20.8** | **71.3** | K.TQTPQLHFLDT*DDEVS*PTSWGDSR.Q |
| **Telomeric repeat binding factor 2** | TERF2 | S323 | **47.1** | **67.4** | K.DLVLPTQALPAS*PALK.N |
| **Tensin 3** | TNS3 | S332 | **16.9** | **63.0** | R.WDS*YENLSADGEVLHTQGPVDGSLYAK.V |
| **Tensin 3** | TNS3 | S776 | **61.3** | **120.6** | R.KLS*LGQYDNDAGGQLPFSK.C |
| **Tensin 3** | TNS3 | S361 | | **27.9** | K.S*SSDPGIPGGPQAIPATNSPDHSDHTLSVSSDSGHSTASAR.T |
| **Tensin 3** | TNS3 | S690 | **20.0** | **50.7** | R.FPGDQVVNGAGPELSTGPSPGS*PTLDIDQSIEQLNR.L |
| **TERF2IP** | TERF2IP | S36 | **38.7** | **54.3** | R.DDGSSMSFYVRPS*PAK.R |
| **TERF2IP** | TERF2IP | S203 | **44.0** | **63.8** | K.YLLGDAPVS*PSSQK.L |
| **TERF2IP** | TERF2IP | S154 | **39.4** | **39.7** | R.S*PSSVTGNALWK.A |
| **Testin** | TES | S105 | **1.7** | **12.5** | K.NVS*INTVTYEWAPPVQNQALAR.Q |
| **Testin** | TES | T108 | **9.5** | **15.6** | K.NVSINT*VTYEWAPPVQNQALAR.Q |

Peak Area: <-10, -5, -3, 0, 3, 5, >10

%CV: 0, 29, 43, 71, 86, >100

**White dots**- Significant change in peptide abundance at 5 %FDR compared to the timepoint with the minimum peak area for a given PSM

| Protein Name | Gene | Phosphosites | Ascor | MOWSE | Sequence |
|---|---|---|---|---|---|
| Testis specific histone H2B | HIST1H2B | S89T90 | | 11.1 | K.RS*T*ISSR.E |
| Testis specific histone H2B | HIST1H2B | S89S92 | 7.6 | 17.5 | K.RS*TIS*SR.E |
| Testis specific protein kinase 2 | TESK2 | S456 | 56.5 | 48.5 | R.S*LPGSPEFLHQEACPFVGR.E |
| Tetratricopeptide repeat protein 16 | TTC16 | T765S767 | 6.9 | 17.0 | K.TT*RS*PR.Q |
| TFIIF beta | GTF2F2 | S248 | 52.9 | 22.8 | R.HYQGEEKS*D.- |
| Thioredoxin domain containing 13 | TMX4 | S251S259 | 100.0 | 29.4 | R.AEQLQDAEEEKDDS*NEEENKDS*LVDDEEEK.E |
| Thioredoxin domain containing protein 1 | TMX1 | S247 | 57.3 | 143.7 | K.VEEEQEADEEDVS*EEEAESK.E |
| Thioredoxin interacting protein | TXNIP | S314S315S318 | -0.1 | 24.8 | R.TS*S*EMS*WVDLNIPDTPEAPPCYMDVIPEDHR.L |
| Thioredoxin interacting protein | TXNIP | T348S358 | -0.2 | 17.4 | R.LESPT*TPLLDDMDGS*QDSPIFMYAPEFK.F |
| Thioredoxin interacting protein | TXNIP | S361 | -0.1 | 26.4 | R.LESPTTPLLDDMDGSQDS*PIFMYAPEFK.F |
| THRAP2 | MED13L | S806S826 | 26.4 | 61.7 | R.AGSSSLTQVTDLAPS*LHDLDNIFDNSDDDELGAVS*PALR.S |
| THRAP2 | MED13L | S796T798 | 11.3 | 31.3 | R.AGSSS*LT*QVTDLAPSLHDLDNIFDNSDDDELGAVSPALR.S |
| THRAP2 | MED13L | T801S806 | 5.6 | 41.1 | R.AGSSSLTQVT*DLAPS*LHDLDNIFDNSDDDELGAVSPALR.S |
| THRAP2 | MED13L | S817S826 | 19.5 | 53.3 | R.AGSSSLTQVTDLAPSLHDLDNIFDNS*DDDELGAVS*PALR.S |
| THRAP2 | MED13L | S817 | 7.6 | 24.2 | R.AGSSSLTQVTDLAPSLHDLDNIFDNS*DDDELGAVSPALR.S |
| THRAP2 | MED13L | S806 | 1.1 | 39.6 | R.AGSSSLTQVTDLAPS*LHDLDNIFDNSDDDELGAVSPALR.S |
| THRAP2 | MED13L | S794 | | 26.1 | R.AGS*SSLTQVTDLAPSLHDLDNIFDNSDDDELGAVSPALR.S |
| THRAP2 | MED13L | S794S795 | | 40.3 | R.AGS*S*SLTQVTDLAPSLHDLDNIFDNSDDDELGAVSPALR.S |
| Thyroid hormone receptor interactor 11 | TRIP11 | T1846S1854 | 7.6 | 21.8 | K.SVPNT*PLRPNQQS*VVNSSFSELFVK.F |
| Thyroid hormone receptor interactor 11 | TRIP11 | S1842T1846 | 46.5 | 37.2 | K.S*VPNT*PLRPNQQSVVNSSFSELFVK.F |
| Thyroid hormone receptor interactor 11 | TRIP11 | S1891 | 46.8 | 32.3 | K.LSVHDMKPLDS*PGRR.K |
| | THUMPD1 | S86S88 | 100.0 | 54.2 | K.DQQPS*GS*EGEDDDAEAALKK.E |
| Thymidine kinase soluble | TK1 | S231 | 100.0 | 15.3 | K.LFAPQQILQCS*PAN.- |
| Thymopoietin | TMPO | Y183 | 5.6 | 36.9 | R.QNGSNDSDRY*SDNEEDSKIELK.L |
| Thymopoietin | TMPO | S177S180 | 25.9 | 42.3 | R.QNGS*NDS*DRYSDNEEDSKIELK.L |
| Thymopoietin | TMPO | S177Y183 | 15.0 | 34.7 | R.QNGS*NDSDRY*SDNEEDSKIELK.L |
| Thymopoietin | TMPO | S159 | -0.3 | 70.7 | R.SS*TPLPTISSSAENTR.Q |
| Thymopoietin | TMPO | S158S159T160 | 30.7 | 37.9 | R.EQGTESRS*S*T*PLPTISSSAENTR.Q |
| Thymopoietin | TMPO | S66 | 5.8 | 22.2 | K.GPPDFS*SDEEREPTPVLGSGAAAAGR.S |
| Thymopoietin | TMPO | S310 | 12.2 | 48.9 | K.SSSSSSQPEHSAMLVSTAAS*PSLIK.E |
| Thymopoietin | TMPO | S66S67 | 42.6 | 92.3 | K.GPPDFS*S*DEEREPTPVLGSGAAAAGR.S |
| Thymopoietin | TMPO | S306 | 112.4 | 69.8 | K.HAS*PILPITEFSDIPR.R |
| Thymopoietin | TMPO | S156 | 26.0 | 28.7 | K.LREQGTES*R.S |
| Thymopoietin | TMPO | T160 | 10.2 | 70.6 | R.SST*PLPTISSSAENTR.Q |
| Thymopoietin | TMPO | S159T160 | -0.2 | 52.8 | R.SS*T*PLPTISSSAENTR.Q |
| Thymopoietin | TMPO | S66T74 | 10.4 | 22.3 | K.GPPDFS*SDEEREPT*PVLGSGAAAAGR.S |
| Thymopoietin | TMPO | S444 | -0.4 | 62.5 | R.DSGS*FVAFQNIPGSELMSSFAK.T |
| Thymopoietin | TMPO | T154S156S158 | 24.9 | 38.3 | R.EQGT*ES*RS*STPLPTISSSAENTR.Q |
| Thymopoietin | TMPO | S184 | 16.4 | 35.7 | R.QNGSNDSDRYS*DNEEDSKIELK.L |
| Thymopoietin | TMPO | S156S158S159 | 6.5 | 14.7 | K.LREQGTES*RS*S*TPLPTISSSAENTR.Q |
| Thymopoietin | TMPO | S312 | 10.7 | 22.6 | K.SSSSSSQPEHSAMLVSTAASPS*LIK.E |
| Thymopoietin | TMPO | S177S184 | 13.3 | 17.2 | R.QNGS*NDSDRYS*DNEEDSKIELK.L |
| Thymopoietin | TMPO | S180Y183 | 5.1 | 12.3 | R.QNGSNDS*DRY*SDNEEDSKIELK.L |
| Thymopoietin | TMPO | S67T74 | 6.9 | 30.6 | K.GPPDFSS*DEEREPT*PVLGSGAAAAGR.S |
| Thymopoietin | TMPO | T154S156T160 | 4.3 | 35.9 | R.EQGT*ES*RSST*PLPTISSSAENTR.Q |
| Thyroid hormone receptor interactor 12 | TRIP12 | S1317S1322 | 22.1 | 64.3 | R.VREDDEDS*DDDGS*DEEIDESLAAQFLNSGNVR.H |
| Thyroid hormone receptor interactor 12 | TRIP12 | S1322S1329 | 23.9 | 47.5 | R.VREDDEDSDDDGS*DEEIDES*LAAQFLNSGNVR.H |
| Thyroid hormone receptor interactor 12 | TRIP12 | S1317S1329 | 10.6 | 45.7 | R.VREDDEDS*DDDGSDEEIDES*LAAQFLNSGNVR.H |
| Thyroid hormone receptor interactor 3 | ZNHIT3 | S80 | 46.3 | 60.0 | K.TVKPVENKDDDDSIADFLNS*DEEEDR.V |
| Thyroid hormone receptor interactor 8 | JMJD1C | S966 | 9.2 | 21.4 | R.S*PTHLTVSSTNTLR.S |
| TIF2 | NCOA2 | S493 | 34.0 | 11.1 | R.MS*PGVAGSPR.I |
| Tight junction protein 2 | TJP2 | S1159 | 32.3 | 78.6 | R.GSYGS*DAEEEEYR.Q |
| Tight junction protein 2 | TJP2 | S398S400 | 8.1 | 61.7 | R.DSQQTLINIPSLNDS*DS*EIEDISEIESNR.S |
| Tight junction protein 2 | TJP2 | S394S398 | 28.0 | 78.4 | R.DSQQTLINIPS*LNDS*DSEIEDISEIESNR.S |
| Tight junction protein 2 | TJP2 | S920Y932T933 | 13.3 | 82.3 | R.LIS*DFEDTDGEGGAY*T*DNELDEPAEEPLVSSITR.S |
| Tight junction protein 2 | TJP2 | S920T925Y932 | 42.0 | 73.3 | R.LIS*DFEDT*DGEGGAY*TDNELDEPAEEPLVSSITR.S |
| Tight junction protein 2 | TJP2 | S244 | 56.7 | 29.7 | R.S*IDQDYER.A |
| Tight junction protein 2 | TJP2 | S394S400 | 10.6 | 50.9 | R.DSQQTLINIPS*LNDSDS*EIEDISEIESNR.S |
| Tight junction protein 2 | TJP2 | S170S174 | 30.2 | 19.6 | R.SRS*WEDS*PER.G |
| Tight junction protein 2 | TJP2 | S1156 | 5.0 | 48.6 | R.GS*YGSDAEEEEYR.Q |
| Tight junction protein 2 | TJP2 | T925Y932T933 | 0.1 | 41.1 | R.LISDFEDT*DGEGGAY*T*DNELDEPAEEPLVSSITR.S |
| Tight junction protein 2 | TJP2 | S920T925T933 | 6.2 | 76.1 | R.LIS*DFEDT*DGEGGAYT*DNELDEPAEEPLVSSITR.S |

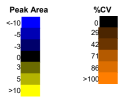


**White dots-** Significant change in peptide abundance at **5** %FDR compared to the timepoint with the minimum peak area for a given PSM

| Protein Name | Gene | Phosphosites | CarT 0m 2m 5m | RajiB 0m 2m 5m | Ascor | MOWSE | Sequence |
|---|---|---|---|---|---|---|---|
| **Tight junction protein 2** | TJP2 | S400 | | | **11.5** | **25.1** | R.DSQQTLINIPSLNDSDS*EIEDISEIESNR.S |
| **Tight junction protein 2** | TJP2 | T925Y932 | | | **6.5** | **44.7** | R.LISDFEDT*DGEGGAY*TDNELDEPAEEPLVSSITR.S |
| **Tight junction protein 2** | TJP2 | S394 | | | **2.4** | **54.6** | R.DSQQTLINIPS*LNDSDSEIEDISEIESNR.S |
| **Tight junction protein 3** | TJP3 | S183 | | | **34.4** | **58.5** | R.S*PGGGSEANGLALVSGFK.R |
| **Tight junction protein 3** | TJP3 | S371S375 | | | **12.2** | **11.3** | R.RES*SVDS*R.T |
| **Tight junction protein 4, peripheral** | TJAP1 | S535 | | | **8.0** | **61.1** | R.KDS*LTQAQEQGNLLN.- |
| **TIM1** | TIMELESS | S1173 | | | **100.0** | **86.7** | R.QLLDS*DEEQEEDEGR.N |
| **Titin** | TTN | Y15335S15339 | | | **17.6** | **13.6** | K.IRIY*AM#NS*EGLGEPALVPGTPKAEDR.M |
| **TLOC1** | SEC62 | T375 | | | **98.5** | **101.5** | K.EELEQQT*DGDCEEDEEEENDGETPK.S |
| **TLOC1** | SEC62 | S117 | | | **100.0** | **33.5** | K.AES*GKEEDKK.S |
| **TNFAIP3 interacting protein 1** | TNIP1 | T431S434 | | | **32.7** | **73.9** | K.ALEEALSIQT*PPS*SPPTAFGSPEGAGALLR.K |
| **TNFAIP3 interacting protein 1** | TNIP1 | S434S435 | | | **6.6** | **43.1** | K.ALEEALSIQTPPS*S*PPTAFGSPEGAGALLR.K |
| **TNFAIP3 interacting protein 1** | TNIP1 | S428S434 | | | **14.5** | **55.3** | K.ALEEALS*IQTPPS*SPPTAFGSPEGAGALLR.K |
| **TNFAIP3 interacting protein 1** | TNIP1 | T431S435S442 | | | **18.3** | **59.2** | K.ALEEALSIQT*PPSS*PPTAFGS*PEGAGALLR.K |
| **TNFAIP3 interacting protein 1** | TNIP1 | S428S435S442 | | | **9.6** | **49.7** | K.ALEEALS*IQTPPSS*PPTAFGS*PEGAGALLR.K |
| **TNFAIP3 interacting protein 1** | TNIP1 | S77 | | | **16.2** | **49.5** | K.DNELLPPPS*PSLGSFDPLAELTGK.D |
| **TNFAIP3 interacting protein 1** | TNIP1 | T431S434S442 | | | **11.2** | **30.4** | K.ALEEALSIQT*PPS*SPPTAFGS*PEGAGALLR.K |
| **TNFAIP3 interacting protein 1** | TNIP1 | T431S435 | | | **7.8** | **60.7** | K.ALEEALSIQT*PPSS*PPTAFGSPEGAGALLR.K |
| **TNFAIP3 interacting protein 1** | TNIP1 | S428S435 | | | **1.0** | **38.4** | K.ALEEALS*IQTPPSS*PPTAFGSPEGAGALLR.K |
| **TNFAIP3 interacting protein 1** | TNIP1 | S79 | | | **7.9** | **30.8** | K.DNELLPPPSPS*LGSFDPLAELTGK.D |
| **TNFAIP3 interacting protein 1** | TNIP1 | T431S435T438 | | | **6.2** | **29.6** | K.ALEEALSIQT*PPSS*PPT*AFGSPEGAGALLR.K |
| **TNFAIP3 interacting protein 1** | TNIP1 | S82 | | | **3.9** | **14.3** | K.DNELLPPPSPSLGS*FDPLAELTGK.D |
| **TNFAIP3 interacting protein 1** | TNIP1 | S428S435T438 | | | **4.2** | **20.6** | K.ALEEALS*IQTPPSS*PPT*AFGSPEGAGALLR.K |
| **TNFAIP3 interacting protein 1** | TNIP1 | S428S434S442 | | | **5.6** | **18.6** | K.ALEEALS*IQTPPS*SPPTAFGS*PEGAGALLR.K |
| **TNFAIP3 interacting protein 2** | TNIP2 | S333S339 | | | **100.0** | **59.2** | K.VAS*LLHQVS*WR.Q |
| **TOCA1** | FNBP1L | S443 | | | **42.4** | **83.5** | R.ES*PEGSYTDDANQEVR.G |
| **TOCA1** | FNBP1L | S430 | | | **48.7** | **39.6** | R.RHS*SDINHLVTQGR.E |
| **TOM1** | TOM1 | T154T164 | | | **25.6** | **19.0** | K.GLEFPMT*DLDMLSPIHT*PQR.T |
| **TOM1** | TOM1 | S160T164 | | | **41.7** | **24.9** | R.KGLEFPMTDLDMLS*PIHT*PQR.T |
| **TOM1** | TOM1 | S160 | | | **9.4** | **18.2** | K.GLEFPMTDLDMLS*PIHTPQR.T |
| **TOM20** | TOMM20 | S135 | | | **9.5** | **20.0** | R.IVS*AQSLAEDDVE.- |
| **TOMM70A** | TOMM70 | S91 | | | **64.4** | **59.0** | R.AS*PAPGSGHPEGPGAHLDMNSLDR.A |
| **TOMM70A** | TOMM70 | S96 | | | **7.8** | **51.6** | R.ASPAPGS*GHPEGPGAHLDMNSLDR.A |
| **Tomosyn** | STXBP5 | S723 | | | **12.9** | **52.9** | R.KLS*LPTDLKPDLDVK.D |
| **Topoisomerase DNA II, beta** | TOP2B | S1517S1519 | | | **40.2** | **72.0** | K.VVEAVNS*DS*DSEFGIPK.K |
| **Topoisomerase DNA II, beta** | TOP2B | S1395S1419 | | | **24.1** | **31.4** | K.VKAS*PITNDGEDEFVPSDGLDKDEYTFS*PGK.S |
| **Topoisomerase DNA II, beta** | TOP2B | S1395S1408 | | | **38.1** | **50.2** | K.VKAS*PITNDGEDEFVPS*DGLDKDEYTFSPGK.S |
| **Topoisomerase DNA II, beta** | TOP2B | S1395 | | | **39.0** | **63.7** | K.AS*PITNDGEDEFVPSDGLDKDEYTFSPGK.S |
| **Topoisomerase DNA II, beta** | TOP2B | S1395S1408S1 | | | **29.5** | **39.4** | K.VKAS*PITNDGEDEFVPS*DGLDKDEYTFS*PGK.S |
| | TOP2B | S1461 | | | **32.2** | **91.5** | K.SEDDSAKFDS*NEEDSASVFSPSFGLK.Q |
| | TOP2B | S1517S1519S1 | | | **100.0** | **58.7** | K.VVEAVNS*DS*DS*EFGIPK.K |
| | TOP2B | T1398S1408 | | | **14.9** | **35.0** | K.ASPIT*NDGEDEFVPS*DGLDKDEYTFSPGK.S |
| **Topoisomerase DNA II, beta** | TOP2B | T1366 | | | **-0.2** | **48.7** | K.YT*FDFSEEEDDDADDDDDDNNDLEELK.V |
| **Topoisomerase DNA II, beta** | TOP2B | S1370 | | | **17.5** | **142.9** | K.YTFDFS*EEEDDDADDDDDDNNDLEELK.V |
| **Topoisomerase DNA II, beta** | TOP2B | S1456S1461 | | | **18.3** | **61.4** | K.SEDDS*AKFDS*NEEDSASVFSPSFGLK.Q |
| **Topoisomerase DNA II, beta** | TOP2B | S1461S1466 | | | **4.3** | **22.4** | K.SEDDSAKFDS*NEEDS*ASVFSPSFGLK.Q |
| **Topoisomerase DNA II, beta** | TOP2B | S1331S1335S1 | | | **67.3** | **105.5** | R.NPWS*DDES*KS*ES*DLEETEPVVIPR.D |
| **Topoisomerase DNA II, beta** | TOP2B | S1456S1461S1 | | | **3.1** | **55.2** | K.SEDDS*AKFDS*NEEDS*ASVFSPSFGLK.Q |
| **Topoisomerase DNA II, beta** | TOP2B | T1570S1576 | | | **16.8** | **44.5** | K.KT*SFDQDS*DVDIFPSDFPTEPPSLPR.T |
| **Topoisomerase DNA II, beta** | TOP2B | S1449 | | | **13.9** | **87.6** | K.SQDFGNLFSFPSYS*QK.S |
| **Topoisomerase DNA II, beta** | TOP2B | S1576S1591 | | | **21.9** | **49.0** | K.KTSFDQDS*DVDIFPSDFPTEPPS*LPR.T |
| **Topoisomerase DNA II, beta** | TOP2B | S1576 | | | **11.5** | **65.5** | K.TSFDQDS*DVDIFPSDFPTEPPSLPR.T |
| **Topoisomerase DNA II, beta** | TOP2B | S1436S1449 | | | **39.9** | **100.5** | K.S*QDFGNLFSFPSYS*QK.S |
| **Topoisomerase DNA II, beta** | TOP2B | S1608 | | | **26.0** | **48.5** | K.YFAES*DEEEDDVDFAMFN.- |
| **Topoisomerase DNA II, beta** | TOP2B | S1517S1521 | | | **30.3** | **72.6** | K.VVEAVNS*DSDS*EFGIPK.K |
| | TOP2B | S1395T1398 | | | **2.4** | **24.4** | K.AS*PIT*NDGEDEFVPSDGLDKDEYTFSPGK.S |
| **Topoisomerase DNA II, beta** | TOP2B | T1570S1571 | | | **15.4** | **25.1** | K.KT*S*FDQDSDVDIFPSDFPTEPPSLPR.T |
| **Topoisomerase DNA II, beta** | TOP2B | S1571 | | | **-0.2** | **68.9** | K.TS*FDQDSDVDIFPSDFPTEPPSLPR.T |
| **Topoisomerase DNA II, beta** | TOP2B | S1571S1576 | | | **11.2** | **25.1** | K.KTS*FDQDS*DVDIFPSDFPTEPPSLPR.T |
| **Topoisomerase DNA II, beta** | TOP2B | T1570S1591 | | | **21.7** | **16.6** | K.KT*SFDQDSDVDIFPSDFPTEPPS*LPR.T |
| **Topoisomerase DNA II, beta** | TOP2B | S1436S1444S1 | | | **18.6** | **42.2** | K.S*QDFGNLFS*FPSYS*QK.S |
| **Topoisomerase DNA II, beta** | TOP2B | S1519S1521 | | | **7.6** | **26.2** | K.KVVEAVNSDS*DS*EFGIPK.K |

Peak Area: <-10, -5, -3, 0, 3, 5, >10

%CV: 0, 29, 42, 71, 86, >100

**White dots-** Significant change in peptide abundance at **5** %FDR compared to the timepoint with the minimum peak area for a given PSM

| Protein Name | Gene | Phosphosites | CarT 0m | CarT 2m | CarT 5m | RajiB 0m | RajiB 2m | RajiB 5m | Ascor | MOWSE | Sequence |
|---|---|---|---|---|---|---|---|---|---|---|---|
| **Topoisomerase DNA II, beta** | TOP2B | T1570 | | | | | | | **7.2** | **30.0** | K.KT*SFDQDSDVDIFPSDFPTEPPSLPR.T |
| **Topoisomerase DNA II, beta** | TOP2B | S1452S1461 | | | | | | | **4.1** | **12.3** | K.S*EDDSAKFDS*NEEDSASVFSPSFGLK.Q |
| **Topoisomerase DNA II, beta** | TOP2B | Y1365 | | | | | | | **7.6** | **68.0** | K.Y*TFDFSEEEDDDADDDDDDNNDLEELK.V |
| **Topoisomerase DNA II, beta** | TOP2B | T1398S1408T14 | | | | | | | **8.5** | **13.5** | K.ASPIT*NDGEDEFVPS*DGLDKDEYT*FSPGK.S |
| **Topoisomerase DNA II, beta** | TOP2B | S1444 | | | | | | | **14.9** | **40.4** | K.SQDFGNLFS*FPSYSQK.S |
| **Topoisomerase DNA II, beta** | TOP2B | S1436 | | | | | | | **21.5** | **51.3** | K.S*QDFGNLFSFPSYSQK.S |
| **Topoisomerase DNA II, beta** | TOP2B | S1447 | | | | | | | **14.9** | **13.7** | K.SQDFGNLFSFPS*YSQK.S |
| **Topoisomerase DNA II, beta** | TOP2B | S1337S1339 | | | | | | | **61.9** | **48.1** | K.S*ES*DLEETEPVVIPR.D |
| **Topoisomerase DNA II, beta** | TOP2B | S1331S1335S1 | | | | | | | **12.0** | **44.4** | R.NPWS*DDES*KSES*DLEET*EPVVIPR.D |
| **TORC3** | CRTC3 | S62 | | | | | | | **53.0** | **64.6** | R.LTQYHGGS*LPNVSQLR.S |
| **TORC3** | CRTC3 | S329 | | | | | | | **31.8** | **55.3** | R.S*NPSIQATLNK.T |
| **TORC3** | CRTC3 | S391S396 | | | | | | | **22.6** | **38.8** | R.RQPPVS*PLTLS*PGPEAHQGFSR.Q |
| **TORC3** | CRTC3 | S391T394 | | | | | | | **32.0** | **37.9** | R.RQPPVS*PLT*LSPGPEAHQGFSR.Q |
| **TORC3** | CRTC3 | S443 | | | | | | | **14.0** | **44.1** | R.SQLSFLPTEAQAQVS*PPPPYPAPQELTQPLLQQPR.A |
| **Tousled like kinase 1** | TLK1 | S204T213 | | | | | | | **0.1** | **13.2** | R.SPQNSHSHSTPSSS*VRPNSPSPT*ALAFGDHPIVQPK.Q |
| **Tousled like kinase 1** | TLK1 | S180 | | | | | | | **8.2** | **41.6** | K.ISDYFEYQGGNGSS*PVR.G |
| **Tousled like kinase 1** | TLK1 | S209S211 | | | | | | | **13.0** | **19.1** | R.SPQNSHSHSTPSSSVRPNS*PS*PTALAFGDHPIVQPK.Q |
| **Tousled like kinase 1** | TLK1 | S191S195 | | | | | | | | **24.1** | R.S*PQNS*HSHSTPSSSVRPNSPSPTALAFGDHPIVQPK.Q |
| **Tousled like kinase 1** | TLK1 | S179 | | | | | | | **25.6** | **42.6** | K.ISDYFEYQGGNGS*SPVR.G |
| **Tousled like kinase 1** | TLK1 | S199S209 | | | | | | | **-14.8** | **13.5** | R.SPQNSHSHS*TPSSSVRPNS*PSPTALAFGDHPIVQPK.Q |
| **Tousled like kinase 1** | TLK1 | S199S202 | | | | | | | **7.1** | **16.0** | R.SPQNSHSHS*TPS*SSVRPNSPSPTALAFGDHPIVQPK.Q |
| **Tousled like kinase 1** | TLK1 | S204S211 | | | | | | | **10.8** | **14.0** | R.SPQNSHSHSTPSSS*VRPNSPS*PTALAFGDHPIVQPK.Q |
| **Tousled like kinase 2** | TLK2 | S98 | | | | | | | **9.2** | **20.5** | K.ISDYFEFAGGSAPGTS*PGR.S |
| **Tousled like kinase 2** | TLK2 | T97 | | | | | | | **-0.4** | **35.1** | K.ISDYFEFAGGSAPGT*SPGR.S |
| **TRA2A** | TRA2A | S84S86T88 | | | | | | | **12.2** | **26.2** | R.S*RS*YT*PEYR.R |
| **TRA2A** | TRA2A | T202 | | | | | | | **50.5** | **79.1** | R.AHT*PTPGIYMGRPTHSGGGGGGGGGGGGGGGR.R |
| **TRA2A** | TRA2A | Y87T88 | | | | | | | **15.1** | **32.5** | R.SY*T*PEYR.R |
| **TRA2A** | TRA2A | S16S18S20T24 | | | | | | | **28.1** | **26.6** | R.S*QS*KS*PTGT*PAR.V |
| **TRA2A** | TRA2A | S98S100 | | | | | | | **29.0** | **19.5** | R.S*HS*PMSNR.R |
| **TRA2A** | TRA2A | S96S98S100 | | | | | | | **40.2** | **42.1** | R.S*RS*HS*PMSNR.R |
| **TRA2A** | TRA2B | S260S262 | | | | | | | **30.0** | **31.9** | R.RRS*PS*PYYSR.Y |
| **TRA2A** | TRA2B | S260Y264 | | | | | | | **16.6** | **29.4** | R.RRS*PSPY*YSR.Y |
| **TRA2A** | TRA2A | S84S86Y87 | | | | | | | **25.7** | **25.8** | R.S*RS*Y*TPEYR.R |
| **TRA2A** | TRA2A | S86Y87 | | | | | | | **15.8** | **15.7** | R.S*Y*TPEYR.R |
| **TRA2A** | TRA2A | S16S18S20 | | | | | | | **28.1** | **23.4** | R.S*QS*KS*PTGTPAR.V |
| **TRA2A** | TRA2A | S86T88 | | | | | | | **20.2** | **27.6** | R.S*YT*PEYR.R |
| **TRA2A** | TRA2A | S86Y87T88 | | | | | | | **10.9** | **34.1** | R.SRS*Y*T*PEYR.R |
| **TRA2A** | TRA2B | S260 | | | | | | | **44.9** | **29.5** | R.S*PSPYYSR.Y |
| **TRA2A** | TRA2A | S98S103 | | | | | | | **17.5** | **15.4** | R.S*HSPMS*NR.R |
| | TRA2A | T204 | | | | | | | **56.5** | **81.4** | R.AHTPT*PGIYMGRPTHSGGGGGGGGGGGGGGGR.R |
| **TRAF1** | TRAF1 | S66 | | | | | | | **23.9** | **44.4** | R.GEDLQSIS*PGSR.L |
| **TRAF1** | TRAF1 | S172 | | | | | | | **7.3** | **81.0** | R.APCSES*QEELALQHFMK.E |
| **TRAF1** | TRAF1 | S170 | | | | | | | **20.7** | **76.3** | R.APCS*ESQEELALQHFMK.E |
| **TRAF1** | TRAF1 | S64 | | | | | | | **10.9** | **40.5** | R.GEDLQS*ISPGSR.L |
| **TRAF3-interacting JNK-activating modulator** | TRAF3IP3 | S111 | | | | | | | **19.1** | **28.2** | R.RISS*PR.E |
| **TRAF3-interacting JNK-activating modulator** | TRAF3IP3 | S110S111 | | | | | | | **100.0** | **20.2** | R.RIS*S*PR.E |
| **Transcription elongation factor A protein-like 1** | TCEAL1 | S41S42 | | | | | | | **100.0** | **17.2** | R.RS*S*PRR.R |
| **Transcription elongation factor A, 1** | TCEA1 | S100 | | | | | | | **80.1** | **69.8** | K.KKEPAITSQNS*PEAR.E |
| **Transcription elongation factor B, 3** | ELOA | T164S167 | | | | | | | **4.2** | **16.3** | R.MSPT*YSS*DPESSDYGHVQSPPSCTSPHQMYVDHYR.S |
| **Transcription elongation factor B, 3** | ELOA | S125 | | | | | | | **13.6** | **16.2** | R.SYS*PDHR.Q |
| **Transcription elongation factor B, 3** | ELOA | S162S167 | | | | | | | **2.7** | **16.4** | R.MS*PTYSS*DPESSDYGHVQSPPSCTSPHQMYVDHYR.S |
| **Transcription elongation factor B, 3** | ELOA | S162T164 | | | | | | | | **13.0** | R.MS*PT*YSSDPESSDYGHVQSPPSCTSPHQMYVDHYR.S |
| **Transcription factor 12** | TCF12 | S559 | | | | | | | **15.0** | **95.2** | R.TSS*TNEDEDLNPEQK.I |
| **Transcription factor 12** | TCF12 | T560 | | | | | | | **-0.3** | **91.4** | R.TSST*NEDEDLNPEQK.I |
| **Transcription factor 12** | TCF12 | S98 | | | | | | | **13.1** | **51.0** | R.LGAHEGLS*PTPFMNSNLMGK.T |
| **Transcription factor 12** | TCF12 | T100 | | | | | | | **4.2** | **50.8** | R.LGAHEGLSPT*PFMNSNLMGK.T |
| **Transcription factor 12** | TCF12 | S67 | | | | | | | **14.7** | **79.3** | R.GGTTSWGTSGQPS*PSYDSSR.G |
| **Transcription factor 2B related factor** | BRF2 | S385 | | | | | | | **19.3** | **78.8** | R.ICPVPPVSTVTGDENIS*DSEIEQYLR.T |
| **Transcription factor 3** | TCF3 | S354S359 | | | | | | | **33.5** | **76.2** | K.ALASIYSPDHSSNNFSSSPS*TPVGS*PQGLAGTSQWPR.A |
| **Transcription factor 3** | TCF3 | S338Y340 | | | | | | | | **73.5** | K.ALAS*IY*SPDHSSNNFSSSPSTPVGSPQGLAGTSQWPR.A |
| **Transcription factor 3** | TCF3 | S351T355S359 | | | | | | | **12.5** | **78.4** | K.ALASIYSPDHSSNNFSS*SPST*PVGS*PQGLAGTSQWPR.A |
| **Transcription factor 3** | TCF3 | S346T355S359 | | | | | | | **9.3** | **74.4** | K.ALASIYSPDHSS*NNFSSSPST*PVGS*PQGLAGTSQWPR.A |

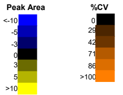


**White dots-** Significant change in peptide abundance at **5** %FDR compared to the timepoint with the minimum peak area for a given PSM

CarT (0m, 2m, 5m) — RajiB (0m, 2m, 5m)

| Protein Name | Gene | Phosphosites | Ascor | MOWSE | Sequence |
|---|---|---|---|---|---|
| Transcription factor 3 | TCF3 | S352T355S359 | 10.4 | 80.4 | K.ALASIYSPDHSSNNFSSS*PST*PVGS*PQGLAGTSQWPR.A |
| Transcription factor 3 | TCF3 | Y340S350S359 | 10.6 | 38.5 | K.ALASIY*SPDHSSNNFS*SSPSTPVGS*PQGLAGTSQWPR.A |
| Transcription factor 3 | TCF3 | S346S359 | 6.2 | 29.6 | K.ALASIYSPDHSS*NNFSSSPSTPVGS*PQGLAGTSQWPR.A |
| Transcription factor 3 | TCF3 | T355S359 | 24.1 | 94.4 | K.ALASIYSPDHSSNNFSSSPST*PVGS*PQGLAGTSQWPR.A |
| Transcription factor 3 | TCF3 | S338Y340S341 | | 61.2 | K.ALAS*IY*S*PDHSSNNFSSSPSTPVGSPQGLAGTSQWPR.A |
| Transcription factor 3 | TCF3 | S352S354S359 | 5.7 | 76.4 | K.ALASIYSPDHSSNNFSSS*PS*TPVGS*PQGLAGTSQWPR.A |
| Transcription factor 3 | TCF3 | S341S350S359 | 9.2 | 47.6 | K.ALASIYS*PDHSSNNFS*SSPSTPVGS*PQGLAGTSQWPR.A |
| Transcription factor 3 | TCF3 | Y340S350S354 | 2.2 | 34.0 | K.ALASIY*SPDHSSNNFS*SSPS*TPVGS*PQGLAGTSQWPR.A |
| Transcription factor 3 | TCF3 | S350S354S359 | 7.9 | 55.7 | K.ALASIYSPDHSSNNFS*SSPS*TPVGS*PQGLAGTSQWPR.A |
| Transcription factor 3 | TCF3 | S346S351S359 | 3.5 | 74.9 | K.ALASIYSPDHSS*NNFSS*SPSTPVGS*PQGLAGTSQWPR.A |
| Transcription factor 3 | TCF3 | S350S351S359 | 5.8 | 49.1 | K.ALASIYSPDHSSNNFS*S*SPSTPVGS*PQGLAGTSQWPR.A |
| Transcription factor 3 | TCF3 | S351S354S359 | 5.3 | 62.8 | K.ALASIYSPDHSSNNFSS*SPS*TPVGS*PQGLAGTSQWPR.A |
| Transcription factor 3 | TCF3 | S345S350S354 | 1.0 | 56.1 | K.ALASIYSPDHS*SNNFS*SSPS*TPVGS*PQGLAGTSQWPR.A |
| Transcription factor 3 | TCF3 | S346S351S354 | 4.2 | 62.0 | K.ALASIYSPDHSS*NNFSS*SPS*TPVGS*PQGLAGTSQWPR.A |
| Transcription factor 3 | TCF3 | Y340S351S354 | 12.9 | 53.0 | K.ALASIY*SPDHSSNNFSS*SPS*TPVGS*PQGLAGTSQWPR.A |
| Transcription factor 3 | TCF3 | S345S351T355 | 5.1 | 70.0 | K.ALASIYSPDHS*SNNFSS*SPST*PVGS*PQGLAGTSQWPR.A |
| Transcription factor 3 | TCF3 | Y340S345S354 | 2.3 | 46.1 | K.ALASIY*SPDHS*SNNFSSSPS*TPVGS*PQGLAGTSQWPR.A |
| Transcription factor 3 | TCF3 | S379 | 46.2 | 61.7 | R.AGAPGALS*PSYDGGLHGLQSK.I |
| Transcription factor 3 | TCF3 | S350S354 | 13.5 | 49.6 | K.ALASIYSPDHSSNNFS*SSPS*TPVGSPQGLAGTSQWPR.A |
| Transcription factor 3 | TCF3 | S351T355 | 4.4 | 84.0 | K.ALASIYSPDHSSNNFSS*SPST*PVGSPQGLAGTSQWPR.A |
| Transcription factor 3 | TCF3 | S345S350S359 | -3.4 | 26.6 | K.ALASIYSPDHS*SNNFS*SSPSTPVGS*PQGLAGTSQWPR.A |
| Transcription factor 3 | TCF3 | S346S352 | 1.9 | 42.7 | K.ALASIYSPDHSS*NNFSSS*PSTPVGSPQGLAGTSQWPR.A |
| Transcription factor 3 | TCF3 | S346S354 | 5.5 | 38.5 | K.ALASIYSPDHSS*NNFSSSPS*TPVGSPQGLAGTSQWPR.A |
| Transcription factor 3 | TCF3 | S350T355S359 | 10.3 | 69.4 | K.ALASIYSPDHSSNNFS*SSPST*PVGS*PQGLAGTSQWPR.A |
| Transcription factor 3 | TCF3 | S350S351S352 | 4.5 | 29.8 | K.ALASIYSPDHSSNNFS*S*S*PSTPVGSPQGLAGTSQWPR.A |
| Transcription factor 3 | TCF3 | Y340T355S359 | 3.0 | 33.4 | K.ALASIY*SPDHSSNNFSSSPST*PVGS*PQGLAGTSQWPR.A |
| Transcription factor 3 | TCF3 | S345S359 | -4.0 | 23.2 | K.ALASIYSPDHS*SNNFSSSPSTPVGS*PQGLAGTSQWPR.A |
| Transcription factor 3 | TCF3 | S351S354 | 7.2 | 56.5 | K.ALASIYSPDHSSNNFSS*SPS*TPVGSPQGLAGTSQWPR.A |
| Transcription factor 3 | TCF3 | S350S359 | 2.5 | 27.1 | K.ALASIYSPDHSSNNFS*SSPSTPVGS*PQGLAGTSQWPR.A |
| Transcription factor 3 | TCF3 | S352S354 | 9.8 | 54.1 | K.ALASIYSPDHSSNNFSSS*PS*TPVGSPQGLAGTSQWPR.A |
| Transcription factor 3 | TCF3 | S127S139 | 6.5 | 39.5 | R.DAGVGGLTQAGFLS*GELALNSPGPLS*PSGMK.G |
| Transcription factor 3 | TCF3 | S134S139 | 10.5 | 60.1 | R.DAGVGGLTQAGFLSGELALNS*PGPLS*PSGMK.G |
| Transcription factor 3 | TCF3 | S346S354S359 | 5.0 | 57.0 | K.ALASIYSPDHSS*NNFSSSPS*TPVGS*PQGLAGTSQWPR.A |
| Transcription factor 3 | TCF3 | S354T355 | 1.2 | 31.5 | K.ALASIYSPDHSSNNFSSSPS*T*PVGSPQGLAGTSQWPR.A |
| Transcription factor 3 | TCF3 | Y340S345S359 | 6.2 | 50.0 | K.ALASIY*SPDHS*SNNFSSSPSTPVGS*PQGLAGTSQWPR.A |
| Transcription factor 3 | TCF3 | S346S350S354 | 3.3 | 25.6 | K.ALASIYSPDHSS*NNFS*SSPS*TPVGSPQGLAGTSQWPR.A |
| Transcription factor AP-2 alpha (activating enhancer binding protein 2 alpha) | TFAP2A | S181 | 15.1 | 21.6 | K.KGPVS*LSK.S |
| Transcription factor Dp-2 | TFDP2 | T40 | 3.5 | 43.9 | K.GNISFVAFPVSNT*NSPTK.I |
| Transcription factor Dp-2 | TFDP2 | S31T40 | 11.8 | 12.3 | R.GFIDQNLSPTKGNIS*FVAFPVSNT*NSPTK.I |
| Transcription factor Dp-2 | TFDP2 | S31T44 | 6.0 | 14.7 | R.GFIDQNLSPTKGNIS*FVAFPVSNTNSPT*K.I |
| Transcription factor E2-alpha | TCF3 | S479 | 13.6 | 90.7 | R.TSS*TDEVLSLEEK.D |
| Transcription factor E3 | TFE3 | S568 | 8.9 | 12.2 | R.RSS*FSMEEES.- |
| Transcription factor E4F1 | E4F1 | S642S643 | 100.0 | 19.6 | R.RS*S*RAPR.Q |
| Transcription factor EB | TFEB | S109S122 | 56.9 | 24.2 | K.FAAHIS*PAQGSPKPPPAAS*PGVR.A |
| Transcription factor EB | TFEB | S467 | 12.1 | 61.3 | R.RSS*FSMEEGDVL.- |
| Transcription factor EB | TFEB | S332 | 31.8 | 97.9 | R.VHGLPTTS*PSGMNMAELAQQVVK.Q |
| Transcription factor EB | TFEB | S133S134S142 | 12.9 | 56.5 | R.AGHVLSS*S*AGNSAPNS*PMAMLHIGS*NPER.E |
| Transcription factor EB | TFEB | S122 | 66.9 | 28.5 | K.FAAHISPAQGSPKPPPAAS*PGVR.A |
| Transcription factor EB | TFEB | S138 | 4.3 | 75.2 | R.AGHVLSSSAGNS*APNSPMAMLHIGSNPER.E |
| Transcription factor EB | TFEB | T331 | 5.8 | 24.8 | R.VHGLPTT*SPSGMNMAELAQQVVK.Q |
| Transcription factor EB | TFEB | S138S142 | 22.4 | 52.2 | R.AGHVLSSSAGNS*APNS*PMAMLHIGSNPER.E |
| Transcription factor EB | TFEB | S114S122 | 26.5 | 21.6 | K.FAAHISPAQGS*PKPPPAAS*PGVR.A |
| Transcription factor EB | TFEB | S133S134 | 10.4 | 86.0 | R.AGHVLSS*S*AGNSAPNSPMAMLHIGSNPER.E |
| Transcription factor EB | TFEB | S133S134S151 | 14.7 | 46.0 | R.AGHVLSS*S*AGNSAPNSPMAMLHIGS*NPER.E |
| Transcription factor EB | TFEB | S134 | | 83.7 | R.AGHVLSSS*AGNSAPNSPMAMLHIGSNPER.E |
| Transcription factor EB | TFEB | S134S138S142 | 6.8 | 16.5 | R.AGHVLSSS*AGNS*APNS*PMAM#LHIGS*NPER.E |
| | TFEB | S134S138S142 | 16.3 | 61.4 | R.AGHVLSSS*AGNS*APNS*PMAMLHIGSNPER.E |
| Transcription factor EB | TFEB | S133S134S138 | 9.9 | 49.7 | R.AGHVLSS*S*AGNS*APNSPMAMLHIGS*NPER.E |
| Transcription factor EB | TFEB | S133 | 15.6 | 78.3 | R.AGHVLSS*SAGNSAPNSPMAMLHIGSNPER.E |
| Transcription factor EB | TFEB | S142 | 3.9 | 16.8 | R.AGHVLSSSAGNSAPNS*PMAMLHIGSNPER.E |
| Transcription factor EB | TFEB | S133S151 | 21.2 | 77.8 | R.AGHVLSS*SAGNSAPNSPMAMLHIGS*NPER.E |
| Transcription factor EB | TFEB | S109S114 | 85.4 | 22.9 | K.FAAHIS*PAQGS*PKPPPAASPGVR.A |

Peak Area: <-10, -5, -3, 0, 3, 5, >10

%CV: 0, 29, 42, 71, 86, >100

**White dots-** Significant change in peptide abundance at **5** %FDR compared to the timepoint with the minimum peak area for a given PSM

| | | | CarT | | | RajiB | | | | | |
|---|---|---|---|---|---|---|---|---|---|---|---|
| **Protein Name** | **Gene** | **Phosphosites** | **0m** | **2m** | **5m** | **0m** | **2m** | **5m** | **Ascor** | **MOWSE** | **Sequence** |
| **Transcription factor EB** | TFEB | S133S138S142 | | | | | | | **6.4** | **38.8** | R.AGHVLSS*SAGNS*APNS*PMAMLHIGS*NPER.E |
| **Transcription factor EB** | TFEB | S138S151 | | | | | | | **16.2** | **24.7** | R.AGHVLSSSAGNS*APNSPMAMLHIGS*NPER.E |
| **Transcription factor EB** | TFEB | S334 | | | | | | | **2.3** | **40.1** | R.VHGLPTTSPS*GM#NMAELAQQVVK.Q |
| **Transcription factor EB** | TFEB | S466S467 | | | | | | | **13.8** | **15.5** | R.RS*S*FSMEEGDVL.- |
| **Transcription factor EB** | TFEB | S134S138 | | | | | | | **11.8** | **74.9** | R.AGHVLSSS*AGNS*APNSPMAMLHIGSNPER.E |
| **Transcription factor EB** | TFEB | S133S138S142 | | | | | | | **14.9** | **58.6** | R.AGHVLSS*SAGNS*APNS*PMAMLHIGSNPER.E |
| **Transcription factor EB** | TFEB | S133S134S142 | | | | | | | **9.1** | **29.2** | R.AGHVLSS*S*AGNSAPNS*PMAMLHIGSNPER.E |
| **Transcription factor ELF 1** | ELF1 | S187 | | | | | | | **21.9** | **16.8** | K.TKPPRPDS*PATTPNISVK.K |
| **Transcription factor ELYS** | AHCTF1 | S1577 | | | | | | | **75.8** | **68.3** | R.NLS*FNELYPSGTLK.L |
| **Transcription factor ELYS** | AHCTF1 | S1980 | | | | | | | **32.1** | **45.6** | R.EVS*PSDVR.E |
| **Transcription factor ELYS** | AHCTF1 | S1195 | | | | | | | **26.9** | **25.6** | R.SLPSSSQLKGS*PQAISR.A |
| **Transcription factor IID** | TAF1 | S328 | | | | | | | **7.6** | **12.3** | K.SLWNYDYAPPPPPEQCLS*DDEITMMAPVESK.F |
| **Transcription factor IIF, alpha subunit** | GTF2F1 | S305S307S308 | | | | | | | **100.0** | **33.4** | K.GVDEQS*DS*S*EES*EEEKPPEEDKEEEEEK.K |
| **Transcription factor IIF, alpha subunit** | GTF2F1 | S385T389 | | | | | | | **31.9** | **76.1** | R.GNS*RPGT*PSAEGGSTSSTLR.A |
| | GTF2F1 | S217S218S221 | | | | | | | **100.0** | **36.4** | R.IHDLEDDLEM#S*S*DAS*DAS*GEEGGRVPK.A |
| **Transcription factor IIF, alpha subunit** | GTF2F1 | S385S391 | | | | | | | **12.8** | **52.0** | R.GNS*RPGTPS*AEGGSTSSTLR.A |
| **Transcription factor like nuclear regulator** | BDP1 | S883 | | | | | | | **100.0** | **11.8** | R.RAIS*PR.E |
| **Transcription factor Sp3** | SP3 | S73 | | | | | | | **176.2** | **105.7** | K.IGPPS*PGDDEEEAAAAAGAPAAAGATGDLASAQLGGAPNR.W |
| **Transcription factor Sp4** | SP4 | S770 | | | | | | | **21.4** | **17.3** | R.IVTVAAISQDS*NPATPNVSTNMEEF.- |
| **Transcription factor Sp4** | SP4 | T774 | | | | | | | **1.1** | **12.0** | R.IVTVAAISQDSNPAT*PNVSTNMEEF.- |
| **Transcription factor YY1** | YY1 | S118 | | | | | | | **100.0** | **82.2** | R.EEVVGGDDS*DGLR.A |
| **Transcription factor YY1** | YY1 | S247 | | | | | | | **67.5** | **87.9** | K.DIDHETVVEEQIIGENS*PPDYSEYMTGK.K |
| **Transcription termination factor, RNA polymerase 1** | TTF1 | S64 | | | | | | | | **18.4** | K.DFQHLIS*SPLKK.S |
| **Transcriptional coactivator CRSP70** | MED26 | S296S314T323 | | | | | | | **14.6** | **10.8** | K.GS*VPSPSPRPQALDATQVPS*PLPLAQPST*PPVR.R |
| **Transcriptional coactivator MMS19** | MMS19 | S1027 | | | | | | | **11.8** | **36.9** | R.GEWFLLGS*PGS.- |
| **Transcriptional intermediary factor 1 beta** | TRIM28 | S473 | | | | | | | **94.5** | **79.0** | R.S*GEGEVSGLMR.K |
| **Transcriptional intermediary factor 1 beta** | TRIM28 | S598 | | | | | | | **7.2** | **95.4** | R.LASPSGS*TSSGLEVVAPEGTSAPGGGPGTLDDSATICR.V |
| **Transcriptional intermediary factor 1 beta** | TRIM28 | T599 | | | | | | | **5.3** | **105.6** | R.LASPSGST*SSGLEVVAPEGTSAPGGGPGTLDDSATICR.V |
| **Transcriptional intermediary factor 1 beta** | TRIM28 | S823 | | | | | | | **16.6** | **35.5** | K.FSAVLVEPPPMSLPGAGLS*SQELSGGPGDGP.- |
| **Transcriptional intermediary factor 1 beta** | TRIM28 | S601 | | | | | | | **6.0** | **46.1** | R.LASPSGSTSS*GLEVVAPEGTSAPGGGPGTLDDSATICR.V |
| **Transcriptional intermediary factor 1 beta** | TRIM28 | S33 | | | | | | | | **31.8** | R.S*TAPSAAASASASAAASSPAGGGAEALELLEHCGVCR.E |
| **Transcriptional intermediary factor 1 beta** | TRIM28 | S816 | | | | | | | **4.9** | **22.1** | K.FSAVLVEPPPMS*LPGAGLSSQELSGGPGDGP.- |
| **Transcriptional intermediary factor 1 beta** | TRIM28 | T541 | | | | | | | **17.6** | **55.1** | R.GAAAAATGQPGTAPAGT*PGAPPLAGMAIVK.E |
| **Transcriptional intermediary factor 1 beta** | TRIM28 | S50 | | | | | | | **18.9** | **71.4** | R.STAPSAAASASASAAASS*PAGGGAEALELLEHCGVCR.E |
| **Transcriptional intermediary factor 1 beta** | TRIM28 | S45 | | | | | | | **3.9** | **24.6** | R.STAPSAAASASAS*AAASSPAGGGAEALELLEHCGVCR.E |
| **Transcriptional intermediary factor 1 beta** | TRIM28 | S49 | | | | | | | **4.6** | **26.3** | R.STAPSAAASASASAAAS*SPAGGGAEALELLEHCGVCR.E |
| **Transcriptional intermediary factor 1 beta** | TRIM28 | S824 | | | | | | | **4.8** | **17.3** | K.FSAVLVEPPPMSLPGAGLSS*QELSGGPGDGP.- |
| **Transcriptional intermediary factor 1 beta** | TRIM28 | S594 | | | | | | | | **76.4** | R.LAS*PSGSTSSGLEVVAPEGTSAPGGGPGTLDDSATICR.V |
| **Transcriptional intermediary factor 1 beta** | TRIM28 | T599S612 | | | | | | | **5.8** | **32.2** | R.LASPSGST*SSGLEVVAPEGTS*APGGGPGTLDDSATICR.V |
| **Transcriptional intermediary factor 1 gamma** | TRIM33 | T1102S1105 | | | | | | | **62.9** | **82.2** | R.TFAPLPEFEQEEDDGEVT*EDS*DEDFIQPR.R |
| **Transcriptional intermediary factor 1 gamma** | TRIM33 | S803S808T815 | | | | | | | **24.1** | **12.6** | R.S*ACMLS*SPESSLT*PPLSTNLHLESELDALASLENHVK.I |
| **Transcriptional intermediary factor 1 gamma** | TRIM33 | S1119 | | | | | | | **100.0** | **12.1** | R.LKS*DERPVHIK.- |
| **Transcriptional intermediary factor 1, alpha** | TRIM24 | S1019S1025 | | | | | | | **4.0** | **59.8** | R.NES*EDNKFS*DDSDDDFVQPR.K |
| **Transcriptional intermediary factor 1, alpha** | TRIM24 | S663S667 | | | | | | | **6.0** | **11.6** | R.TVQSPNS*SVPS*PGLAGPVTMTSVHPPIR.S |
| **Transcriptional intermediary factor 1, alpha** | TRIM24 | S768S771 | | | | | | | **3.8** | **31.1** | R.SILTSLLLNSS*QSS*TSEETVLR.S |
| **Transcriptional intermediary factor 1, alpha** | TRIM24 | S660S667 | | | | | | | **12.8** | **70.6** | R.TVQS*PNSSVPS*PGLAGPVTMTSVHPPIR.S |
| **Transcriptional intermediary factor 1, alpha** | TRIM24 | T657S667 | | | | | | | **4.2** | **14.1** | R.T*VQSPNSSVPS*PGLAGPVTMTSVHPPIR.S |
| **Transcriptional intermediary factor 1, alpha** | TRIM24 | S767S771 | | | | | | | **11.3** | **44.6** | R.SILTSLLLNS*SQSS*TSEETVLR.S |
| **Transcriptional intermediary factor 1, alpha** | TRIM24 | S1019S1028 | | | | | | | **11.7** | **73.3** | R.NES*EDNKFSDDS*DDDFVQPR.K |
| **Transcriptional intermediary factor 1, alpha** | TRIM24 | S768S770 | | | | | | | **-0.4** | **24.4** | R.SILTSLLLNSS*QS*STSEETVLR.S |
| **Transcriptional intermediary factor 1, alpha** | TRIM24 | S1025S1028 | | | | | | | **7.6** | **46.9** | R.NESEDNKFS*DDS*DDDFVQPR.K |
| **Transcriptional regulator protein HCNGP** | SAP30BP | S69S72S77 | | | | | | | **9.6** | **16.1** | R.LGGDEDGYEEEEDENS*RQS*EDDDS*ETEKPEADDPK.D |
| **Transcriptional regulator protein HCNGP** | SAP30BP | Y46 | | | | | | | **1.2** | **20.6** | K.GGLVSDAY*GEDDFSR.L |
| **Transcriptional regulator protein HCNGP** | SAP30BP | S18S22 | | | | | | | **4.8** | **16.5** | K.NVLSSLAVYAEDS*EPES*DGEAGIEAVGSAAEEK.G |
| **Transcriptional regulator protein HCNGP** | SAP30BP | S9S10 | | | | | | | **8.7** | **13.1** | K.NVLS*S*LAVYAEDSEPESDGEAGIEAVGSAAEEK.G |
| **Transcriptional regulator protein HCNGP** | SAP30BP | Y14S22 | | | | | | | **2.3** | **14.9** | K.NVLSSLAVY*AEDSEPES*DGEAGIEAVGSAAEEK.G |
| **Transcriptional repressor p66 alpha** | GATAD2A | S100S113 | | | | | | | **45.1** | **38.1** | R.RPPS*PDVIVLSDNEQPS*SPR.V |
| **Transcriptional repressor p66 alpha** | GATAD2A | S340 | | | | | | | **14.6** | **83.8** | K.GTTATSAQANSTPTSVASVVTSAES*PASR.Q |
| **Transcriptional repressor p66 alpha** | GATAD2A | S100S107S114 | | | | | | | **33.4** | **40.0** | R.RPPS*PDVIVLS*DNEQPSS*PR.V |
| **Transcriptional repressor p66 alpha** | GATAD2A | S100S107S113 | | | | | | | **45.1** | **20.0** | R.RPPS*PDVIVLS*DNEQPS*SPR.V |
| **Transcriptional repressor p66 alpha** | GATAD2A | S100S114 | | | | | | | **42.8** | **16.7** | R.RPPS*PDVIVLSDNEQPSS*PR.V |
| **Transducer of regulated cAMP response element binding protein 2** | CRTC2 | S171 | | | | | | | **-0.3** | **34.9** | R.TSS*DSALHTSVMNPSPQDTYPGPTPPSILPSR.R |

Peak Area: <-10, -5, -3, 0, 3, 5, >10

%CV: 0, 29, 42, 71, 86, >100

**White dots-** Significant change in peptide abundance at **5** %FDR compared to the timepoint with the minimum peak area for a given PSM

| Protein Name | Gene | Phosphosites | CarT (0m 2m 5m) | RajiB (0m 2m 5m) | Ascor | MOWSE | Sequence |
|---|---|---|---|---|---|---|---|
| Transducer of regulated cAMP response element binding protein 2 | CRTC2 | S170 | | | 5.7 | 33.2 | R.TS*SDSALHTSVMNPSPQDTYPGPTPPSILPSR.R |
| Transducer of regulated cAMP response element binding protein 2 | CRTC2 | S173 | | | 0.1 | 14.9 | R.TSSDS*ALHTSVMNPSPQDTYPGPTPPSILPSR.R |
| Transducer of regulated cAMP response element binding protein 2 | CRTC2 | T187 | | | 8.7 | 31.0 | R.TSSDSALHTSVMNPSPQDT*YPGPTPPSILPSR.R |
| Transducer of regulated cAMP response element binding protein 2 | CRTC2 | S178 | | | 6.5 | 35.6 | R.TSSDSALHTS*VM#NPSPQDTYPGPTPPSILPSR.R |
| Transducer of regulated cAMP response element binding protein 2 | CRTC2 | S70S79S90 | | | 27.8 | 38.8 | R.SSHYGGS*LPNVNQIGS*GLAEFQSPLHS*PLDSSR.S |
| Transducer of regulated cAMP response element binding protein 2 | CRTC2 | S456T458 | | | 5.2 | 38.5 | K.QFS*PT*MSPTLSSITQGVPLDTSK.L |
| Transducer of regulated cAMP response element binding protein 2 | CRTC2 | S70S86S90 | | | 25.0 | 31.0 | R.SSHYGGS*LPNVNQIGSGLAEFQS*PLHS*PLDSSR.S |
| Transducer of regulated cAMP response element binding protein 2 | CRTC2 | S456S460 | | | 6.9 | 52.2 | K.QFS*PTMS*PTLSSITQGVPLDTSK.L |
| Transducer of regulated cAMP response element binding protein 2 | CRTC2 | S623 | | | 5.5 | 14.1 | R.HGSGPNIILTGDS*SPGFSK.E |
| Transducer of regulated cAMP response element binding protein 2 | CRTC2 | S613 | | | 26.7 | 23.8 | R.HGS*GPNIILTGDSSPGFSK.E |
| Transducer of regulated cAMP response element binding protein 2 | CRTC2 | S183 | | | 11.8 | 15.0 | R.TSSDSALHTSVMNPS*PQDTYPGPTPPSILPSR.R |
| Transducer of regulated cAMP response element binding protein 2 | CRTC2 | S456 | | | 9.1 | 36.9 | K.QFS*PTMSPTLSSITQGVPLDTSK.L |
| Transducer of regulated cAMP response element binding protein 2 | CRTC2 | S456T462 | | | 5.7 | 25.6 | K.QFS*PTMSPT*LSSITQGVPLDTSK.L |
| Transducer of regulated cAMP response element binding protein 2 | CRTC2 | S456S464 | | | 10.6 | 48.2 | K.QFS*PTMSPTLS*SITQGVPLDTSK.L |
| Transducin like enhancer of split 3 | TLE3 | T285 | | | 27.8 | 46.5 | K.DAPT*SPASVASSSSTPSSK.T |
| Transducin like enhancer of split 3 | TLE3 | S203 | | | 26.0 | 106.2 | R.ESSANNSVS*PSESLR.A |
| Transducin like enhancer of split 3 | TLE3 | S240S245T259 | | | 21.6 | 32.8 | R.YDS*DGDKS*DDLVVDVSNEDPAT*PR.V |
| Transducin like enhancer of split 3 | TLE3 | Y238S240T259 | | | 11.1 | 22.2 | R.Y*DS*DGDKSDDLVVDVSNEDPAT*PR.V |
| Transducin like enhancer of split 3 | TLE3 | S263S267 | | | 100.0 | 15.6 | R.VS*PAHS*PPENGLDK.A |
| Transducin like enhancer of split 3 | TLE3 | S286 | | | 13.9 | 48.7 | K.DAPTS*PASVASSSSTPSSK.T |
| Transducin like enhancer of split 3 | TLE3 | Y238S245T259 | | | 9.3 | 13.1 | R.Y*DSDGDKS*DDLVVDVSNEDPAT*PR.V |
| Transducin-like enhancer protein 3 (Fragment) | TLE3 | S37S38 | | | 100.0 | 11.6 | R.CENELGGPGAS*S*R.F |
| Transgelin 2 | TAGLN2 | S163 | | | 100.0 | 60.5 | R.NFS*DNQLQEGK.N |
| Transketolase | TKT | Y275 | | | 10.9 | 66.5 | K.NMAEQIIQEIY*SQIQSK.K |
| Translation machinery-associated protein 16 | TMA16 | S192S193S195 | | | 3.3 | 20.3 | K.TIITVDQDLGELELNDES*S*DS*DEEMTAVA.- |
| Translocated promoter region | TPR | S2155 | | | 85.5 | 46.5 | R.TDGFAEAIHS*PQVAGVPR.F |
| Translocated promoter region | TPR | T2116 | | | 100.0 | 85.4 | R.GLQLT*PGIGGMQQHFFDDEDR.T |
| Translocated promoter region | TPR | S2136 | | | 15.7 | 53.0 | R.TVPS*TPTLVVPHR.T |
| Translocated promoter region | TPR | S522 | | | 12.1 | 76.9 | R.DEEVS*SADISSSSEVISQHLVSYR.N |
| Translocated promoter region | TPR | S2136S2155 | | | 17.5 | 11.0 | R.TVPS*TPTLVVPHRTDGFAEAIHS*PQVAGVPR.F |
| Translocation associated membrane protein 1 | TRAM1 | S365 | | | 28.7 | 60.2 | K.KGTENGVNGTLTSNVADS*PR.N |
| Transmembrane and coiled coil domains 1 | TMCC1 | S90 | | | 13.9 | 58.9 | K.ALGVISNFQSS*PK.Y |
| Transmembrane and coiled coil domains 1 | TMCC1 | S89 | | | 19.3 | 11.4 | K.ALGVISNFQS*SPK.Y |
| Transmembrane and coiled coil domains 1 | TMCC1 | S58 | | | 100.0 | 29.4 | R.NKFGS*ADNIPNLK.D |
| transmembrane and coiled coil domains 3 | TMCC3 | S46 | | | 4.4 | 25.8 | R.GGS*DTNLNFDVPDGILDFHK.V |
| Transmembrane protein 181 | TMEM181 | S580Y586 | | | 23.5 | 88.6 | K.DNPAFSMLNDS*DDDVIY*GSDYEEMPLQNGQAIR.A |
| Transmembrane protein 181 | TMEM181 | S588 | | | -0.7 | 53.3 | K.DNPAFSMLNDSDDDVIYGS*DYEEMPLQNGQAIR.A |
| Transmembrane protein 181 | TMEM181 | S580 | | | 5.9 | 63.8 | K.DNPAFSMLNDS*DDDVIYGSDYEEMPLQNGQAIR.A |
| Transmembrane protein 181 | TMEM181 | S609S611 | | | 7.5 | 16.2 | K.YKEES*DS*D.- |
| Transmembrane protein 181 | TMEM181 | S580Y590 | | | -1.0 | 41.8 | K.DNPAFSMLNDS*DDDVIYGSDY*EEM#PLQNGQAIR.A |
| Transmembrane protein 181 | TMEM181 | Y586 | | | -0.3 | 74.8 | K.DNPAFSMLNDSDDDVIY*GSDYEEMPLQNGQAIR.A |
| Transmembrane protein 181 | TMEM181 | S580S588 | | | 7.7 | 49.9 | K.DNPAFSMLNDS*DDDVIYGS*DYEEMPLQNGQAIR.A |
| Transmembrane protein 181 | TMEM181 | S575Y586 | | | 8.2 | 43.1 | K.DNPAFS*MLNDSDDDVIY*GSDYEEMPLQNGQAIR.A |
| Transmembrane protein 181 | TMEM181 | Y586S588 | | | | 48.9 | K.DNPAFSMLNDSDDDVIY*GS*DYEEMPLQNGQAIR.A |
| Transmembrane protein 181 | TMEM181 | S588Y590 | | | | 39.0 | K.DNPAFSMLNDSDDDVIYGS*DY*EEMPLQNGQAIR.A |
| | TMEM206 | S9 | | | 1.5 | 59.9 | R.STS*YQELSEELVQVVENSELADEQDKETVR.V |
| Transmembrane protein 35B | TMEM35B | S6 | | | 100.0 | 10.9 | M.ALLLS*VLR.V |
| Transmembrane protein 87A | TMEM87A | S549 | | | 149.2 | 89.0 | K.WVEENVPSSVTDVALPALLDS*DEER.M |
| Transmembrane protein 87A | TMEM87A | S540 | | | 74.9 | 52.0 | K.AQEDDLKWVEENVPSSVTDVALPALLDS*DEER.M |
| Transmembrane protein 87B | TMEM87B | S534 | | | 58.2 | 70.1 | K.WVEENIPSSFTDVALPVLVDS*DEEIMTR.S |
| Transmembrane protein 87B | TMEM87B | S469 | | | 23.2 | 50.0 | R.YAFMPLIDDS*DDEIEEFMVTSENLTEGIK.L |
| Transmembrane protein 9 | TMEM9 | S137 | | | 95.9 | 50.0 | R.S*MAAAAASLGGPR.A |
| TRAP150 | THRAP3 | S243 | | | 52.0 | 70.1 | R.ASAVSELS*PR.E |
| TRAP150 | THRAP3 | S240 | | | 43.9 | 34.5 | R.ASAVS*ELSPR.E |
| TRAP150 | THRAP3 | S939 | | | 29.8 | 113.1 | K.FSGEEGEIEDDES*GTENR.E |
| TRAP150 | THRAP3 | S379 | | | 19.8 | 79.2 | K.GSFS*DTGLGDGK.M |
| TRAP150 | THRAP3 | S928T941 | | | 40.6 | 64.1 | K.FS*GEEGEIEDDESGT*ENR.E |
| TRAP150 | THRAP3 | S682 | | | 36.8 | 24.4 | R.RIDIS*PSTFRK.H |
| TRAP150 | THRAP3 | S928S939 | | | 57.8 | 92.4 | K.WAHDKFS*GEEGEIEDDES*GTENR.E |
| TRAP150 | THRAP3 | S248S253 | | | 81.6 | 59.2 | R.ERS*PALKS*PLQSVVVR.R |
| TRAP150 | THRAP3 | S248 | | | 68.1 | 85.1 | R.ERS*PALKSPLQSVVVR.R |
| TRAP150 | THRAP3 | S575 | | | 65.9 | 42.7 | R.MDS*FDEDLARPSGLLAQER.K |

Peak Area: <-10, -5, -3, 0, 3, 5, >10

%CV: 0, 29, 42, 71, 86, >100

**White dots-** Significant change in peptide abundance at **5** %FDR compared to the timepoint with the minimum peak area for a given PSM

CarT (0m, 2m, 5m) RajiB (0m, 2m, 5m)

| Protein Name | Gene | Phosphosites | Ascor | MOWSE | Sequence |
|---|---|---|---|---|---|
| **TRAP150** | THRAP3 | S534 | -0.3 | 26.9 | K.SS*SPPPR.K |
| **TRAP150** | THRAP3 | S535 | 13.6 | 33.3 | K.SSS*PPPR.K |
| **TRAP150** | THRAP3 | S24S26S28S30 | 100.0 | 13.1 | R.S*RS*FS*KS*R.S |
| **TRAP150** | THRAP3 | Y54S55 | 10.9 | 32.3 | R.SY*S*PAHNR.E |
|  | BCLAF1 | S672 | 100.0 | 23.2 | K.S*PEIHRR.I |
| **TRAP150** | THRAP3 | S119 | 13.9 | 29.8 | R.QAYS*PR.R |
| **TRAP150** | THRAP3 | S320 | 36.3 | 87.3 | K.S*PPSTGSTYGSSQK.E |
| **TRAP150** | THRAP3 | S399S406 | 20.1 | 13.6 | K.TDS*EKPFRGS*QSPK.R |
| **TRAP150** | THRAP3 | S315S320 | 56.7 | 75.4 | K.S*PVGKS*PPSTGSTYGSSQK.E |
| **TRAP150** | THRAP3 | S51S53S55 | 17.7 | 32.4 | R.S*RS*YS*PAHNR.E |
| **TRAP150** | THRAP3 | S26S28S30 | 100.0 | 20.6 | R.S*FS*KS*R.S |
| **TRAP150** | THRAP3 | S406S408 | 23.6 | 19.2 | K.TDSEKPFRGS*QS*PK.R |
| **TRAP150** | ZNF318 | S134S136 | 100.0 | 20.3 | R.RS*PS*PR.R |
| **TRAP150** | THRAP3 | S53Y54S55 | -0.1 | 15.2 | R.SRS*Y*S*PAHNR.E |
| **TRAP150** | THRAP3 | S698 | 100.0 | 36.2 | K.HGLAHDEMKS*PR.E |
| **TRAP150** | THRAP3 | S928 | 5.8 | 29.4 | K.FS*GEEGEIEDDESGTENR.E |
| **TRAP150** | THRAP3 | S253 | 42.4 | 15.2 | K.S*PLQSVVVR.R |
| **TRAP150** | THRAP3 | S51Y54S55 | 13.7 | 13.9 | R.S*RSY*S*PAHNR.E |
| **TRAP150** | THRAP3 | S51S53Y54 | 1.9 | 11.4 | R.S*RS*Y*SPAHNR.E |
| **TRAP150** | THRAP3 | S377 | 7.2 | 72.4 | K.GS*FSDTGLGDGK.M |
| **TRAP150** | THRAP3 | S24S26S28 | 100.0 | 11.5 | R.S*RS*FS*K.S |
| **TRAP150** | THRAP3 | S323 | -20.9 | 87.8 | K.SPPS*TGSTYGSSQK.E |
| **TRAP220** | MED1 | S1207T1215 | 42.4 | 18.6 | K.LAS*PMKPVPGT*PPSSK.A |
| **TRAP220** | MED1 | S1479S1482 | 17.7 | 18.7 | K.SYQNS*PSS*DDGIRPLPEYSTEK.H |
| **Treacle** | TCOF1 | S1301 | 13.9 | 97.9 | K.LGAGEGGEASVS*PEK.T |
| **Treacle** | TCOF1 | S270S272S273 | 65.8 | 46.2 | R.AKKPEEES*ES*S*EEGS*ES*EEEAPAGTR.S |
| **Treacle** | TCOF1 | S1151 | 13.9 | 96.1 | K.LDSS*PSVSSTLAAK.D |
| **Treacle** | TCOF1 | S1073 | 6.5 | 13.5 | K.KQEGPATQVDSAVGTLPATS*PQSTSVQAK.G |
| **Treacle** | TCOF1 | T785S791S794 | 5.2 | 33.3 | K.AVATAAQAQT*GPEEDS*GSS*EEES*DS*EEEAETLAQVKPSGK.T |
| **Treacle** | TCOF1 | T785S791S793 | 9.3 | 29.9 | K.AVATAAQAQT*GPEEDS*GS*SEEES*DS*EEEAETLAQVKPSGK.T |
| **Treacle** | TCOF1 | S1113S1114 | 25.4 | 77.0 | R.TETLVEETAAES*S*EDDVVAPSQSLLSGYMTPGLTPANSQASK.A |
| **Treacle** | TCOF1 | T1102T1104 |  | 53.5 | R.T*ET*LVEETAAESSEDDVVAPSQSLLSGYMTPGLTPANSQASK.A |
| **Treacle** | TCOF1 | S1273 | 100.0 | 74.3 | R.KLS*GDQPAAR.T |
| **Treacle** | TCOF1 | S304 | 24.2 | 55.5 | K.TSQVGAASAPAKES*PR.K |
| **Treacle** | TCOF1 | S829 | 100.0 | 44.8 | R.AALAPAKES*PR.K |
| **Treacle** | TCOF1 | S156 | 11.3 | 42.0 | K.TVANLLSGKS*PR.K |
|  | TCOF1 | S505 | 10.9 | 33.4 | K.SLGNILQAKPTS*SPAKGPPQK.A |
| **Treacle** | TCOF1 | S426 | 72.1 | 71.0 | K.S*PQVKPASTMGMGPLGK.G |
|  | TCOF1 | S506 | 3.5 | 37.3 | K.SLGNILQAKPTSS*PAKGPPQK.A |
| **Treacle** | TCOF1 | S1114S1122 | 3.3 | 51.0 | R.TETLVEETAAESS*EDDVVAPS*QSLLSGYMTPGLTPANSQASK.A |
| **Treacle** | TCOF1 | S1114S1124 | 0.9 | 42.1 | R.TETLVEETAAESS*EDDVVAPSQS*LLSGYMTPGLTPANSQASK.A |
| **Treacle** | TCOF1 | T504 | 79.5 | 29.8 | K.SLGNILQAKPT*SSPAKGPPQK.A |
| **Treacle** | TCOF1 | T1072 | 10.3 | 36.0 | K.KQEGPATQVDSAVGTLPAT*SPQSTSVQAK.G |
| **Treacle** | TCOF1 | T1109S1122S11 | 1.0 | 15.6 | R.TETLVEET*AAESSEDDVVAPS*QS*LLSGYMTPGLTPANSQASK.A |
| **Treacle** | TCOF1 | S1122S1124 | 0.9 | 44.2 | R.TETLVEETAAESSEDDVVAPS*QS*LLSGYMTPGLTPANSQASK.A |
| **Treacle** | TCOF1 | T779T785S791 |  | 54.3 | K.AVAT*AAQAQT*GPEEDS*GS*S*EEESDSEEEAETLAQVKPSGK.T |
| **Trinucleotide repeat containing 6** | TNRC6A | Y1574 | 0.0 | 43.9 | R.LEESPFVPY*DFMNSSTSPASPPGSIGDGWPR.A |
| **Trinucleotide repeat containing 6** | TNRC6A | S1585 | 26.0 | 47.7 | R.LEESPFVPYDFMNSSTSPAS*PPGSIGDGWPR.A |
| **Trinucleotide repeat containing 6** | TNRC6A | Y1574S1585 | 14.6 | 39.6 | R.LEESPFVPY*DFMNSSTSPAS*PPGSIGDGWPR.A |
| **Trinucleotide repeat containing 6** | TNRC6A | Y1574T1581 | 2.1 | 25.3 | R.LEESPFVPY*DFMNSST*SPASPPGSIGDGWPR.A |
| **Trinucleotide repeat containing 6** | TNRC6A | S1579S1585 | 9.7 | 35.6 | R.LEESPFVPYDFMNS*STSPAS*PPGSIGDGWPR.A |
| **Trinucleotide repeat containing 6** | TNRC6A | S1579S1580 | 5.5 | 11.9 | R.LEESPFVPYDFMNS*S*TSPASPPGSIGDGWPR.A |
| **Trinucleotide repeat containing 6** | TNRC6A | T1581S1585 | 9.6 | 44.8 | R.LEESPFVPYDFMNSST*SPAS*PPGSIGDGWPR.A |
| **Trinucleotide repeat containing 6B** | TNRC6B | S1012 | 99.8 | 25.9 | R.MGS*PAPLLPGDLLGGGSDSI.- |
| **Trinucleotide repeat containing 6B** | TNRC6B | Y630T640 | 4.5 | 11.5 | R.GGSPY*NQFDIIPGDT*LGGHTGPAGDSWLPAKSPPTNK.I |
| **Trinucleotide repeat containing 6B** | TNRC6B | Y630S657 | 6.3 | 17.1 | R.GGSPY*NQFDIIPGDTLGGHTGPAGDSWLPAKS*PPTNK.I |
| **Trinucleotide repeat containing 6B** | TNRC6B | Y630 | 9.8 | 15.8 | R.GGSPY*NQFDIIPGDTLGGHTGPAGDSWLPAK.S |
| **Trinucleotide repeat containing 6B** | TNRC6B | T640 | 4.7 | 11.2 | R.GGSPYNQFDIIPGDT*LGGHTGPAGDSWLPAK.S |
| **Trinucleotide repeat containing protein 15** | GIGYF2 | S376 | 31.2 | 21.2 | R.VGVEASEETPQTSSS*SARPGTPSDHQSQEASQFER.K |
| **Trinucleotide repeat containing protein 15** | GIGYF2 | S384 | 0.3 | 30.6 | R.VGVEASEETPQTSSSSARPGTPS*DHQSQEASQFER.K |
| **Trinucleotide repeat containing protein 15** | GIGYF2 | S26S30 | 23.1 | 37.2 | R.ALSSGGSITS*PPLS*PALPK.Y |
| **Trinucleotide repeat containing protein 15** | GIGYF2 | S160 | 20.9 | 32.5 | R.SQS*WEER.G |

Peak Area: <-10, -5, -3, 0, 3, 5, >10

%CV: 0, 29, 42, 71, 86, >100

**White dots-** Significant change in peptide abundance at **5** %FDR compared to the timepoint with the minimum peak area for a given PSM

| Protein Name | Gene | Phosphosites | CarT 0m | CarT 2m | CarT 5m | RajiB 0m | RajiB 2m | RajiB 5m | Ascor | MOWSE | Sequence |
|---|---|---|---|---|---|---|---|---|---|---|---|
| **Trinucleotide repeat containing protein 15** | GIGYF2 | S236 | | | | | | | **100.0** | **49.2** | R.WRPHS*PDGPR.S |
| **Trinucleotide repeat containing protein 15** | GIGYF2 | S26 | | | | | | | **23.9** | **87.6** | R.ALSSGGSITS*PPLSPALPK.Y |
| **Trinucleotide repeat containing protein 15** | GIGYF2 | T25S30 | | | | | | | **22.8** | **35.1** | R.ALSSGGSIT*SPPLS*PALPK.Y |
| **Trinucleotide repeat containing protein 15** | GIGYF2 | T373 | | | | | | | **7.4** | **16.8** | R.VGVEASEETPQT*SSSSARPGTPSDHQSQEASQFER.K |
| **Trinucleotide repeat containing protein 15** | GIGYF2 | S374 | | | | | | | **0.3** | **16.8** | R.VGVEASEETPQTS*SSSARPGTPSDHQSQEASQFER.K |
| **Trinucleotide repeat containing protein 15** | GIGYF2 | T25 | | | | | | | **17.8** | **18.4** | R.ALSSGGSIT*SPPLSPALPK.Y |
| **Trinucleotide repeat-containing gene 6C protein** | TNRC6C | S1000 | | | | | | | **49.0** | **87.2** | R.S*PLLGPVSSGWGEMPNVHSK.T |
| **TRIO** | TRIO | S1724 | | | | | | | **-0.3** | **26.2** | R.SS*MEMEGIFNHK.D |
| **Triosephosphate isomerase 1** | TPI1 | S21 | | | | | | | **108.0** | **119.5** | R.KQS*LGELIGTLNAAK.V |
| **Tripartite motif containing 45** | TRIM45 | S451 | | | | | | | **100.0** | **12.1** | K.KDS*PVR.T |
| **tRNA modification GTPase GTPBP3, mitochondrial** | GTPBP3 | T22S23 | | | | | | | **100.0** | **17.9** | R.RT*S*PRLR.R |
| **tRNA-yW synthesizing protein 3 homolog** | TYW3 | S236 | | | | | | | **14.1** | **15.0** | K.ES*DEELENDDDDDLGINVTIFPEDY.- |
| **Tropomodulin 3** | TMOD3 | S25 | | | | | | | **18.4** | **65.1** | K.YKDLDEDELLGNLS*ETELK.Q |
| **TRPC6** | TRPC6 | S14 | | | | | | | **8.9** | **20.0** | R.RGSS*PR.G |
| **Tryptophanyl tRNA synthetase** | WARS | S467 | | | | | | | **100.0** | **36.6** | R.KLS*FDFQ.- |
| **Tryptophanyl tRNA synthetase** | WARS | S8 | | | | | | | **4.3** | **35.9** | M.PNSEPAS*LLELFNSIATQGELVR.S |
| **TSC22 domain family 4** | TSC22D4 | S279 | | | | | | | **100.0** | **49.6** | K.S*PDPFGAVAAQK.F |
| **TSC22 domain family 4** | TSC22D4 | S165 | | | | | | | **23.2** | **90.8** | R.S*FTGGLGQLVVPSK.A |
| **TSC22 domain family 4** | TSC22D4 | S260S263 | | | | | | | **11.4** | **12.0** | R.MELGAPEEMGQVPPLDS*RPS*SPALYFTHDASLVHK.S |
| **TSC22 domain family 4** | TSC22D4 | T167 | | | | | | | **-0.5** | **82.4** | R.SFT*GGLGQLVVPSK.A |
| **TSC22 domain family 4** | TSC22D4 | T229 | | | | | | | **16.0** | **29.9** | R.T*PPLSR.R |
| **TTK protein kinase** | TTK | S455 | | | | | | | **16.9** | **54.2** | K.TPS*SNTLDDYMSCFR.T |
| **Tuberin** | TSC2 | S1386 | | | | | | | **-0.1** | **87.8** | K.SS*SSPELQTLQDILGDPGDK.A |
| **Tuberin** | TSC2 | S1387S1411 | | | | | | | **22.0** | **21.6** | K.SSS*SPELQTLQDILGDPGDKADVGRLS*PEVK.A |
| **Tuberin** | TSC2 | S1388S1411 | | | | | | | **39.8** | **24.3** | K.SSSS*PELQTLQDILGDPGDKADVGRLS*PEVK.A |
| **Tuberin** | TSC2 | T1422 | | | | | | | **-0.5** | **86.4** | R.SQSGT*LDGESAAWSASGEDSR.G |
| **Tuberin** | TSC2 | S1387 | | | | | | | **9.5** | **40.8** | K.SSS*SPELQTLQDILGDPGDK.A |
| **Tuberous sclerosis 1** | TSC1 | T339T345 | | | | | | | | **38.5** | R.LIT*EPPQAT*LWSPSMVCGMTTPPTSPGNVPPDLSHPYSK.V |
| **Tubulin alpha 1** | TUBA1B | S340 | | | | | | | **104.7** | **72.4** | R.S*IQFVDWCPTGFK.V |
| **Tubulin tyrosine ligase-like family, member 4** | TTLL4 | S121S122 | | | | | | | **13.9** | **16.1** | R.RS*S*YR.Q |
| **Tubulin tyrosine ligase-like family, member 4** | TTLL4 | S122Y123 | | | | | | | **5.0** | **12.1** | R.RSS*Y*R.Q |
| **Tubulin tyrosine ligase-like family, member 4** | TTLL4 | S121Y123 | | | | | | | **11.4** | **13.2** | R.RS*SY*R.Q |
| **Tubulin, alpha 2** | TUBA1C | S48 | | | | | | | **46.9** | **142.5** | K.TIGGGDDS*FNTFFSETGAGK.H |
| **Tubulin, alpha, brain specific** | TUBA1B | S439 | | | | | | | **63.6** | **13.7** | K.DYEEVGVDS*VEGEGEEEGEEY.- |
| **Tubulin, beta** | TUBB | T55 | | | | | | | **12.7** | **30.3** | R.ISVYYNEAT*GGK.Y |
| **Tuftelin interacting protein 11** | TFIP11 | S98 | | | | | | | **100.0** | **66.5** | K.GAAEEAELEDS*DDEEKPVKQDDFPK.D |
| **Tuftelin interacting protein 11** | TFIP11 | S210 | | | | | | | **86.3** | **85.7** | R.TTQSMQDFPVVDS*EEEAEEEFQK.E |
| | TFIP11 | S59 | | | | | | | **40.7** | **26.2** | R.DS*DDERPSFGGK.R |
| **Tuftelin interacting protein 11** | TFIP11 | Y51 | | | | | | | **11.9** | **13.2** | K.EEATY*GVWAERDSDDERPSFGGK.R |
| **Tumor necrosis factor receptor superfamily, member 9** | | Y222 | | | | | | | **100.0** | **26.5** | K.LLY*IFK.Q |
| **Tumor protein D52** | TPD52 | T180 | | | | | | | **7.4** | **47.6** | K.VGGTKPAGGDFGEVLNSAANASATTTEPLPEKT*QESL.- |
| **Tumor protein D52** | TPD52 | S183 | | | | | | | **17.6** | **66.5** | K.VGGTKPAGGDFGEVLNSAANASATTTEPLPEKTQES*L.- |
| **Tumor protein D52 like 2** | TPD52L2 | S21 | | | | | | | **32.1** | **100.2** | K.GLLSDS*MTDVPVDTGVAAR.T |
| **Tumor protein p53 binding protein 1** | TP53BP1 | S1094S1101 | | | | | | | **29.0** | **36.7** | R.QSQQPMKPIS*PVKDPVS*PASQK.M |
| **Tumor protein p53 binding protein 1** | TP53BP1 | S1208S1219 | | | | | | | **7.6** | **13.5** | R.GSGEKPVS*APGDDTESLHS*QGEEEFDMPQPPHGHVLHR.H |
| **Tumor protein p53 binding protein 1** | TP53BP1 | T1214S1219 | | | | | | | **8.6** | **32.9** | R.GSGEKPVSAPGDDT*ESLHS*QGEEEFDMPQPPHGHVLHR.H |
| **Tumor protein p53 binding protein 1** | TP53BP1 | S1216S1219 | | | | | | | **19.2** | **37.7** | R.GSGEKPVSAPGDDTES*LHS*QGEEEFDMPQPPHGHVLHR.H |
| **Tumor protein p53 binding protein 1** | TP53BP1 | S1202S1208 | | | | | | | | **14.6** | R.GS*GEKPVS*APGDDTESLHSQGEEEFDMPQPPHGHVLHR.H |
| **Tumor protein p53 binding protein 1** | TP53BP1 | S380 | | | | | | | **20.1** | **50.4** | R.STPFIVPSS*PTEQEGR.Q |
| **Tumor protein p53 binding protein 1** | TP53BP1 | S552 | | | | | | | **27.3** | **35.8** | R.IDEDGENTQIEDTEPMS*PVLNSK.F |
| **Tumor protein p53 binding protein 1** | TP53BP1 | S500 | | | | | | | **104.3** | **117.8** | K.NS*PEDLGLSLTGDSCK.L |
| **Tumor protein p53 binding protein 1** | TP53BP1 | S294 | | | | | | | **112.6** | **108.0** | K.S*PEPEVLSTQEDLFDQSNK.T |
| **Tumor protein p53 binding protein 1** | TP53BP1 | S366 | | | | | | | **88.2** | **76.0** | R.SLVQDSLSTNSSDLVAPS*PDAFR.S |
| **Tumor protein p53 binding protein 1** | TP53BP1 | S1114 | | | | | | | **9.0** | **87.0** | K.MVIQGPSS*PQGEAMVTDVLEDQK.E |
| **Tumor protein p53 binding protein 1** | TP53BP1 | S1462 | | | | | | | **-0.5** | **96.3** | R.SDS*PEIPFQAAAGPSDGLDASSPGNSFVGLR.V |
| **Tumor protein p53 binding protein 1** | TP53BP1 | S1460 | | | | | | | **7.2** | **52.6** | R.S*DSPEIPFQAAAGPSDGLDASSPGNSFVGLR.V |
| **Tumor protein p53 binding protein 1** | TP53BP1 | S1758S1759 | | | | | | | **20.1** | **48.1** | R.SKLPDGPTGS*S*EEEEEFLEIPPFNK.Q |
| **Tumor protein p53 binding protein 1** | TP53BP1 | S635S639S640 | | | | | | | **100.0** | **87.1** | R.S*EALS*S*VLDQEEAMEIK.E |
| **Tumor protein p53 binding protein 1** | TP53BP1 | S1462S1481 | | | | | | | **5.8** | **56.6** | R.SDS*PEIPFQAAAGPSDGLDASS*PGNSFVGLR.V |
| **Tumor protein p53 binding protein 1** | TP53BP1 | S1426S1430 | | | | | | | **24.4** | **41.4** | R.ETAVPGPLGIEDIS*PNLS*PDDKSFSR.V |
| **Tumor protein p53 binding protein 1** | TP53BP1 | S1462S1480 | | | | | | | **19.5** | **39.2** | R.SDS*PEIPFQAAAGPSDGLDAS*SPGNSFVGLR.V |
| **Tumor protein p53 binding protein 1** | TP53BP1 | S1460S1462 | | | | | | | **-0.5** | **16.3** | R.S*DS*PEIPFQAAAGPSDGLDASSPGNSFVGLR.V |

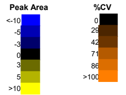


**White dots-** Significant change in peptide abundance at **5** %FDR compared to the timepoint with the minimum peak area for a given PSM

| Protein Name | Gene | Phosphosites | CarT 0m | CarT 2m | CarT 5m | RajiB 0m | RajiB 2m | RajiB 5m | Ascor | MOWSE | Sequence |
|---|---|---|---|---|---|---|---|---|---|---|---|
| **Tumor protein p53 binding protein 1** | TP53BP1 | S525 | | | | | | | 7.8 | 23.3 | K.LMLSTSEYSQS*PK.M |
| **Tumor protein p53 binding protein 1** | TP53BP1 | T1214S1216 | | | | | | | 2.3 | 38.4 | R.GSGEKPVSAPGDDT*ES*LHSQGEEEFDMPQPPHGHVLHR.H |
| | TP53BP1 | S1113 | | | | | | | 8.0 | 41.5 | K.MVIQGPS*SPQGEAM#VTDVLEDQK.E |
| **Tumor protein p53 binding protein 1** | TP53BP1 | T548 | | | | | | | 8.4 | 46.8 | R.IDEDGENTQIEDT*EPMSPVLNSK.F |
| **Tumor protein p53 inducible protein 11** | TP53I11 | S14 | | | | | | | 50.1 | 46.4 | K.KHS*QTDLVSR.L |
| **Tumor protein, translationally controlled 1** | TPT1 | T39 | | | | | | | | 56.0 | R. [illegible] |
| **Tumor rejection antigen 1** | HSP90B1 | S306 | | | | | | | 100.0 | 61.1 | K.EES*DDEAAVEEEEEEKKPK.T |
| **Tumor rejection antigen 1** | HSP90B1 | T468 | | | | | | | 100.0 | 33.9 | R.KT*LDMIKK.I |
| **Tumor suppressing subtransferable candidate 4** | TSSC4 | S320 | | | | | | | -0.6 | 11.9 | K.SS*SPEDPGAEV.- |
| **Tumor suppressing subtransferable candidate 4** | TSSC4 | S132 | | | | | | | 100.0 | 15.1 | R.S*PVEGLGR.A |
| **Tumor suppressing subtransferable candidate 4** | TSSC4 | S321 | | | | | | | 20.1 | 21.6 | K.SSS*PEDPGAEV.- |
| **Twist neighbor** | TWISTNB | S316S328 | | | | | | | 47.2 | 27.0 | R.KHS*EEAEFTPPLKCS*PK.R |
| **Tyrosine protein kinase ABL2** | ABL2 | S631 | | | | | | | 35.4 | 33.7 | R.DKS*PSSLLEDAK.E |
| **Tyrosine-protein kinase PRAG1** | PRAG1 | S696 | | | | | | | 28.9 | 68.8 | K.SAS*FAFEFPK.D |
| **Tyrosine-protein kinase PRAG1** | PRAG1 | S826 | | | | | | | 46.3 | 15.3 | R.AAS*SPDGFFWTQGSPKPGTASPK.L |
| **Tyrosine-protein kinase PRAG1** | PRAG1 | S694 | | | | | | | 8.0 | 40.3 | K.S*ASFAFEFPK.D |
| **Tyrosyl DNA phosphodiesterase 1** | TDP1 | T147S148 | | | | | | | 26.2 | 91.7 | R.LKEEEDEYET*S*GEGQDIWDMLDK.G |
| **Tyrosyl DNA phosphodiesterase 1** | TDP1 | S13S14S15S18 | | | | | | | 10.6 | 25.2 | R.WTIS*S*S*DES*EEEKPKPDKPSTSSLLCAR.Q |
| **Tyrosyl DNA phosphodiesterase 1** | TDP1 | T11S13S14S15 | | | | | | | 6.9 | 16.3 | R.WT*IS*S*S*DESEEEKPKPDKPSTSSLLCAR.Q |
| **Tyrosyl DNA phosphodiesterase 1** | TDP1 | Y145T147 | | | | | | | 4.2 | 68.0 | R.LKEEEDEY*ET*SGEGQDIWDMLDK.G |
| **U2-associated SR140 protein** | U2SURP | S202 | | | | | | | 100.0 | 15.5 | K.KKS*NLELFK.E |
| **U2-associated SR140 protein** | U2SURP | S67 | | | | | | | 28.4 | 44.4 | R.ESLCDS*PHQNLSRPLLENK.L |
| **U2-associated SR140 protein** | U2SURP | S946S948 | | | | | | | 100.0 | 34.5 | R.VKS*PS*PK.S |
| **U2-associated SR140 protein** | U2SURP | S970S974 | | | | | | | 10.2 | 16.7 | R.SS*HKDS*PR.D |
| **U2-associated SR140 protein** | U2SURP | S930T931S934 | | | | | | | 16.3 | 21.3 | R.RHS*T*SPS*PSR.S |
| **U2-associated SR140 protein** | U2SURP | S930S932S934 | | | | | | | 7.3 | 16.6 | R.RHS*TS*PS*PSR.S |
| **U2-associated SR140 protein** | U2SURP | S49 | | | | | | | 25.1 | 11.3 | R.TRPKS*PR.K |
| **U2-associated SR140 protein** | U2SURP | T931S932S934 | | | | | | | 17.1 | 18.7 | R.RHST*S*PS*PSR.S |
| **U2-associated SR140 protein** | U2SURP | S930T931S932 | | | | | | | 12.4 | 22.6 | R.RHS*T*S*PSPSR.S |
| **U2-associated SR140 protein** | U2SURP | S930S934S936 | | | | | | | 10.0 | 14.6 | R.RHS*TSPS*PS*R.S |
| **U2AF65** | U2AF2 | S62 | | | | | | | 42.4 | 27.8 | R.S*KPLTR.G |
| **U2AF65** | U2AF2 | S28S30 | | | | | | | 100.0 | 16.0 | R.KRS*HS*R.S |
| **U2AF65** | U2AF2 | S79 | | | | | | | 100.0 | 16.9 | K.EEHGGLIRS*PR.H |
| **U5 200kD** | SNRNP20 | S225 | | | | | | | 100.0 | 115.0 | R.EEAS*DDDM#EGDEAVVR.C |
| **Ubiquitin activating enzyme 1** | UBA1 | S24 | | | | | | | 2.0 | 26.0 | R.RVSGPDPKPGSNCS*PAQSVLSEVPSVPTNGMAK.N |
| **Ubiquitin activating enzyme 1** | UBA1 | S13S24 | | | | | | | 8.0 | 21.2 | R.RVS*GPDPKPGSNCS*PAQSVLSEVPSVPTNGMAK.N |
| **Ubiquitin activating enzyme 1** | UBA1 | S13S28 | | | | | | | 22.3 | 29.3 | R.RVS*GPDPKPGSNCSPAQS*VLSEVPSVPTNGMAK.N |
| **Ubiquitin activating enzyme 1** | UBA1 | S21 | | | | | | | 49.5 | 23.9 | R.RVSGPDPKPGS*NCSPAQSVLSEVPSVPTNGMAK.N |
| **Ubiquitin activating enzyme 1** | UBA1 | S13 | | | | | | | 4.8 | 13.9 | R.RVS*GPDPKPGSNCSPAQSVLSEVPSVPTNGMAK.N |
| **Ubiquitin activating enzyme 1** | UBA1 | S13S31 | | | | | | | 11.7 | 41.1 | R.RVS*GPDPKPGSNCSPAQSVLS*EVPSVPTNGMAK.N |
| **Ubiquitin activating enzyme 1** | UBA1 | S13S21 | | | | | | | 31.6 | 24.8 | R.RVS*GPDPKPGS*NCSPAQSVLSEVPSVPTNGMAK.N |
| **Ubiquitin associatedand SH3 domain containing protein A** | UBASH3A | S382 | | | | | | | 20.0 | 41.1 | R.S*LSSLQALQATVAR.K |
| **Ubiquitin carboxyl terminal esterase L3** | UCHL3 | S128 | | | | | | | 13.4 | 12.7 | K.FLEESVS*MSPEER.A |
| **Ubiquitin carboxyl terminal esterase L3** | UCHL3 | S130 | | | | | | | 11.3 | 26.9 | K.FLEESVSMS*PEER.A |
| **Ubiquitin carboxyl terminal hydrolase 4** | USP4 | S675S680 | | | | | | | 16.2 | 28.3 | K.EQLS*ETEGS*GEDEPGNDPSETTQKK.I |
| **Ubiquitin carboxyl-terminal hydrolase 24** | USP24 | T42 | | | | | | | | 33.8 | K.NDINEAVALLT*NERPGLDYGGYEPMDSGGGPSPGPGGGPR.G |
| **Ubiquitin carboxyl-terminal hydrolase 8** | USP8 | S718 | | | | | | | 14.3 | 34.1 | R.SYS*SPDITQAIQEEEKR.K |
| **Ubiquitin carboxyl-terminal hydrolase 8** | USP8 | Y717 | | | | | | | -0.5 | 22.8 | R.SY*SSPDITQAIQEEEKR.K |
| **Ubiquitin conjugating enzyme E2, J1** | UBE2J1 | S268 | | | | | | | -1.5 | 48.8 | R.RLSTS*PDVIQGHQPR.D |
| **Ubiquitin conjugating enzyme E2, J1** | UBE2J1 | S251 | | | | | | | 26.0 | 40.1 | K.NTSMS*PR.Q |
| **Ubiquitin conjugating enzyme E2, J1** | UBE2J1 | T267 | | | | | | | -0.4 | 40.8 | R.RLST*SPDVIQGHQPR.D |
| **Ubiquitin conjugating enzyme E2, J1** | UBE2J1 | S266 | | | | | | | 7.2 | 45.0 | R.RLS*TSPDVIQGHQPR.D |
| **Ubiquitin conjugating enzyme E2Q** | UBE2Q1 | Y160T181 | | | | | | | | 15.6 | K. [illegible] |
| **Ubiquitin fusion degradation 1 like** | UFD1 | S299 | | | | | | | 10.4 | 45.7 | R.FVAFSGEGQS*LR.K |
| **Ubiquitin protein ligase E3C** | UBE3C | T347S350 | | | | | | | | 25.9 | R.VLQT*FLS*QLPVSPASASCHDSASDSEEESEEADKPSSPEDGR.L |
| **Ubiquitin protein ligase EDD** | UBR5 | S2486 | | | | | | | 79.1 | 49.8 | R.S*VVDMDLDDTDDGDDNAPLFYQPGKR.G |
| **Ubiquitin protein ligase NEDD4 like** | NEDD4L | S428 | | | | | | | 20.1 | 99.3 | R.SLS*SPTVTLSAPLEGAK.D |
| **Ubiquitin protein ligase NEDD4 like** | NEDD4L | S429 | | | | | | | -0.4 | 79.0 | R.SLSS*PTVTLSAPLEGAK.D |
| **Ubiquitin specifc protease 16** | USP16 | S552 | | | | | | | 10.9 | 89.4 | K.NINMDNDLEVLTSS*PTR.N |
| **Ubiquitin specifc protease 16** | USP16 | S415 | | | | | | | 39.6 | 42.1 | K.TVEDEDQDS*EEEKDNDSYIK.E |
| **Ubiquitin specific peptidase 9, X-linked** | USP9X | S2436 | | | | | | | 17.2 | 51.9 | R.RPYTGNPQYTYNNWS*PPVQSNETSNGYFLER.S |
| **Ubiquitin specific protease 10** | USP10 | S576 | | | | | | | 130.6 | 126.8 | K.NHSVNEEEQEEQGEGS*EDEWEQVGPR.N |

Peak Area: <-10, -5, -3, 0, 3, 5, >10

%CV: 0, 29, 42, 71, 86, >100

**White dots-** Significant change in peptide abundance at **5** %FDR compared to the timepoint with the minimum peak area for a given PSM

| Protein Name | Gene | Phosphosites | CarT 0m | CarT 2m | CarT 5m | RajiB 0m | RajiB 2m | RajiB 5m | Ascor | MOWSE | Sequence |
|---|---|---|---|---|---|---|---|---|---|---|---|
| Ubiquitin specific protease 10 | USP10 | S211 | | | | | | | 24.3 | 71.9 | R.TCNS*PQNSTDSVSDIVPDSPFPGALGSDTR.T |
| Ubiquitin specific protease 10 | USP10 | S80 | | | | | | | 10.0 | 83.3 | R.TPSYSIS*STLNPQAPEFILGCTASK.I |
| Ubiquitin specific protease 10 | USP10 | T74 | | | | | | | 10.9 | 152.0 | R.T*PSYSISSTLNPQAPEFILGCTASK.I |
| Ubiquitin specific protease 10 | USP10 | T82 | | | | | | | 10.3 | 53.7 | R.TPSYSISST*LNPQAPEFILGCTASK.I |
| Ubiquitin specific protease 10 | USP10 | S76 | | | | | | | 17.7 | 68.4 | R.TPS*YSISSTLNPQAPEFILGCTASK.I |
| Ubiquitin specific protease 10 | USP10 | S211S215S220 | | | | | | | 7.5 | 52.6 | R.TCNS*PQNS*TDSVS*DIVPDSPFPGALGSDTR.T |
| Ubiquitin specific protease 10 | USP10 | S81 | | | | | | | 12.2 | 29.7 | R.TPSYSISS*TLNPQAPEFILGCTASK.I |
| Ubiquitin specific protease 10 | USP10 | T216 | | | | | | | 0.2 | 20.3 | R.TCNSPQNST*DSVSDIVPDSPFPGALGSDTR.T |
| Ubiquitin specific protease 10 | USP10 | T208 | | | | | | | 2.5 | 26.1 | R.T*CNSPQNSTDSVSDIVPDSPFPGALGSDTR.T |
| Ubiquitin specific protease 10 | USP10 | S78 | | | | | | | 3.5 | 92.5 | R.TPSYS*ISSTLNPQAPEFILGCTASK.I |
| Ubiquitin specific protease 10 | USP10 | S211S215T216 | | | | | | | 1.7 | 44.8 | R.TCNS*PQNS*T*DSVSDIVPDSPFPGALGSDTR.T |
| Ubiquitin specific protease 10 | USP10 | S211S215S218 | | | | | | | 6.1 | 40.9 | R.TCNS*PQNS*TDS*VSDIVPDSPFPGALGSDTR.T |
| Ubiquitin specific protease 10 | USP10 | S211S218S220 | | | | | | | 3.2 | 55.3 | R.TCNS*PQNSTDS*VS*DIVPDSPFPGALGSDTR.T |
| Ubiquitin specific protease 14 | USP14 | S143 | | | | | | | 16.3 | 85.8 | R.AS*GEMASAQYITAALR.D |
| Ubiquitin specific protease 14 | USP14 | S148 | | | | | | | 18.5 | 67.9 | R.ASGEMAS*AQYITAALR.D |
| Ubiquitin specific protease 15 | USP15 | S965 | | | | | | | 0.7 | 31.4 | K.GASAATGIPLESDEDS*NDNDNDIENENCMHTN.- |
| Ubiquitin specific protease 15 | USP15 | S961 | | | | | | | 25.5 | 42.3 | K.GASAATGIPLES*DEDSNDNDNDIENENCMHTN.- |
| Ubiquitin specific protease 15 | USP15 | S961S965 | | | | | | | 25.3 | 32.9 | K.GASAATGIPLES*DEDS*NDNDNDIENENCMHTN.- |
| Ubiquitin specific protease 15 | USP15 | S229 | | | | | | | 42.4 | 29.0 | K.S*PGASNFSTLPK.I |
| Ubiquitin specific protease 20 | USP20 | S132S134 | | | | | | | 100.0 | 31.9 | K.AVPIAVADEGES*ES*EDDDLKPR.G |
| Ubiquitin specific protease 24 | USP24 | S1887 | | | | | | | 57.8 | 46.8 | R.VSDQNS*PVLPK.K |
| Ubiquitin specific protease 24 | USP24 | S250 | | | | | | | 33.6 | 141.1 | R.TIS*AQDTLAYATALLNEK.E |
| Ubiquitin specific protease 24 | USP24 | T254 | | | | | | | 14.2 | 86.5 | R.TISAQDT*LAYATALLNEK.E |
| Ubiquitin specific protease 34 | USP34 | S483S486S487 | | | | | | | 14.7 | 11.8 | K.AQLS*KQS*S*FAS*LLNTNIPIGNKK.E |
| Ubiquitin specific protease 34 | USP34 | S2488 | | | | | | | 46.9 | 58.7 | K.GPENPQVEVLS*EEEGEEEEEEEDILSLAEEK.Y |
| Ubiquitin specific protease 39 | USP39 | S82 | | | | | | | 100.0 | 28.6 | R.EVDEDS*EPER.E |
| Ubiquitin specific protease 47 | USP47 | T844 | | | | | | | 22.4 | 53.6 | R.ELEQHIQT*SDPENFQSEER.S |
| Ubiquitin specific protease 47 | USP47 | S744 | | | | | | | 22.4 | 18.4 | R.LFVLLPEQS*PVSYSK.R |
| Ubiquitin specific protease 6 | USP6 | S1214 | | | | | | | 100.0 | 31.1 | R.LRLPQIGS*K.N |
| Ubiquitin specific protease 7 | USP7 | S18 | | | | | | | 68.9 | 76.8 | K.AGEQQLS*EPEDMEMEAGDTDDPPR.I |
| Ubiquitin-associated protein 2-like | UBAP2L | S478 | | | | | | | 18.0 | 49.6 | K.STSAPQMS*PGSSDNQSSSPQPAQQK.L |
| Ubiquitin-associated protein 2-like | UBAP2L | S478S488 | | | | | | | 20.8 | 42.4 | K.STSAPQMS*PGSSDNQSSS*PQPAQQK.L |
| Ubiquitin-associated protein 2-like | UBAP2L | S473S488 | | | | | | | 20.8 | 26.0 | K.STS*APQMSPGSSDNQSSS*PQPAQQK.L |
| Ubiquitin-associated protein 2-like | UBAP2L | S473 | | | | | | | 8.7 | 26.7 | K.STS*APQMSPGSSDNQSSSPQPAQQK.L |
| Ubiquitin-associated protein 2-like | UBAP2L | S481 | | | | | | | 7.4 | 33.4 | K.STSAPQMSPGS*SDNQSSSPQPAQQK.L |
| Ubiquitin-associated protein 2-like | UBAP2L | S486S488 | | | | | | | 5.7 | 17.0 | K.STSAPQMSPGSSDNQS*SS*PQPAQQK.L |
| Ubiquitin-conjugating enzyme E2Z | UBE2Z | S231S232 | | | | | | | 4.6 | 35.5 | R.LHNENAEMDSDS*S*SSGTETDLHGSLR.V |
| Ubiquitin-conjugating enzyme E2Z | UBE2Z | S229S231S232 | | | | | | | 19.9 | 77.7 | R.LHNENAEMDS*DS*S*S*SGTETDLHGSLR.V |
| Ubiquitin-conjugating enzyme E2Z | UBE2Z | S229S231S232 | | | | | | | 15.1 | 38.6 | R.LHNENAEMDS*DS*S*S*S*GTETDLHGSLR.V |
| Ubiquitin-conjugating enzyme E2Z | UBE2Z | S229S232 | | | | | | | 6.9 | 65.6 | R.LHNENAEMDS*DSS*SSGTETDLHGSLR.V |
| Ubiquitin-conjugating enzyme E2Z | UBE2Z | S229S232S233 | | | | | | | 11.2 | 54.3 | R.LHNENAEMDS*DSS*S*SGTETDLHGSLR.V |
| Ubiquitin-conjugating enzyme E2Z | UBE2Z | S229S231S232 | | | | | | | 21.9 | 74.0 | R.LHNENAEMDS*DS*S*SS*GTETDLHGSLR.V |
| Ubiquitin-conjugating enzyme E2Z | UBE2Z | S229S232S233 | | | | | | | 13.5 | 20.9 | R.LHNENAEMDS*DSS*S*S*GT*ETDLHGSLR.V |
| Ubiquitin-conjugating enzyme E2Z | UBE2Z | S231S232S233 | | | | | | | 10.6 | 33.6 | R.LHNENAEMDSDS*S*S*S*GTETDLHGSLR.V |
| Ubiquitination factor E4B protein | UBE4B | S79 | | | | | | | 0.3 | 15.6 | R.SQSS*EGVSSLSSSPSNSLETQSQSLSR.S |
| Ubiquitination factor E4B protein | UBE4B | S87 | | | | | | | 6.1 | 48.7 | R.SQSSEGVSSLSS*SPSNSLETQSQSLSR.S |
| Ubiquitously transcribed TPR protein on the X chromosome | KDM6A | S769 | | | | | | | 58.3 | 75.1 | K.S*PGLLSSDNPQLSALLMGK.A |
| UBX domain containing 7 | UBXN7 | S288 | | | | | | | 34.4 | 134.4 | R.SESLIDASEDS*QLEAAIR.A |
| UBX domain containing 7 | UBXN7 | S280S288 | | | | | | | 18.6 | 93.6 | R.SES*LIDASEDS*QLEAAIR.A |
| UBX domain containing 7 | UBXN7 | S350 | | | | | | | 100.0 | 42.5 | K.S*PHKDLGHR.K |
| UBX domain containing 7 | UBXN7 | S285 | | | | | | | 6.0 | 31.7 | R.SESLIDAS*EDSQLEAAIR.A |
| UBX domain containing 7 | UBXN7 | S280 | | | | | | | 13.6 | 29.4 | R.SES*LIDASEDSQLEAAIR.A |
| Unactive progesterone receptor 23KD | PTGES3 | S113 | | | | | | | 75.8 | 100.0 | K.DWEDDS*DEDMSNFDR.F |
| Unactive progesterone receptor 23KD | PTGES3 | S151 | | | | | | | 10.0 | 58.6 | R.FSEMMNNMGGDEDVDLPEVDGADDDSQDS*DDEKMPDLE.- |
| Unactive progesterone receptor 23KD | PTGES3 | S148 | | | | | | | 4.3 | 34.8 | R.FSEMMNNMGGDEDVDLPEVDGADDDS*QDSDDEKMPDLE.- |
| Unactive progesterone receptor 23KD | PTGES3 | S124 | | | | | | | | 19.3 | R.FS*EMMNNMGGDEDVDLPEVDGADDDSQDSDDEKMPDLE.- |
| Unactive progesterone receptor 23KD | PTGES3 | S100 | | | | | | | 17.0 | 58.4 | K.LNWLS*VDFNNWKDWEDDSDEDMSNFDR.F |
| Unactive progesterone receptor 23KD | PTGES3 | S113S118 | | | | | | | 100.0 | 55.0 | K.DWEDDS*DEDMS*NFDR.F |
| Unc 51 like kinase 1 | ULK1 | S556 | | | | | | | 33.7 | 37.8 | R.LHS*APNLSDLHVVRPK.L |
| Unc 51 like kinase 1 | ULK1 | S638 | | | | | | | 16.9 | 101.3 | K.TPS*SQNLLALLAR.Q |
| Unc 51 like kinase 1 | ULK1 | T709S716 | | | | | | | | 42.9 | K.AAFGT*QAPDPGS*TESLQEKPMEIAPSAGFGGSLHPGAR.A |
| Unc 51 like kinase 1 | ULK1 | S716S719 | | | | | | | 30.6 | 43.3 | K.AAFGTQAPDPGS*TES*LQEKPMEIAPSAGFGGSLHPGAR.A |

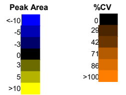


**White dots-** Significant change in peptide abundance at 5 %FDR compared to the timepoint with the minimum peak area for a given PSM

| Protein Name | Gene | Phosphosites | CarT 0m | CarT 2m | CarT 5m | RajiB 0m | RajiB 2m | RajiB 5m | Ascor | MOWSE | Sequence |
|---|---|---|---|---|---|---|---|---|---|---|---|
| Unc 51 like kinase 1 | ULK1 | T660 | | | | | | | 31.4 | 12.8 | R.T*LPDLSEVGPFHGQPLGPGLRPGEDPK.G |
| Unc 93 homolog B1 | UNC93B1 | S547S550 | | | | | | | 27.8 | 25.1 | R.YLEEDNS*DES*DAEGEHGDGAEEEAPPAGPRPGPEPAGLGR.R |
| Unc 93 homolog B1 | UNC93B1 | Y541S547 | | | | | | | | 19.4 | R.Y*LEEDNS*DESDAEGEHGDGAEEEAPPAGPRPGPEPAGLGR.R |
| UNC112 related protein 2 | FERMT3 | S8 | | | | | | | 20.6 | 52.8 | K.TAS*GDYIDSSWELR.V |
| UNC112 related protein 2 | FERMT3 | S484 | | | | | | | 7.6 | 125.8 | R.TGS*GGPGNHPHGPDASAEGLNPYGLVAPR.F |
| UNC112 related protein 2 | FERMT3 | T482 | | | | | | | 5.5 | 44.6 | R.T*GSGGPGNHPHGPDASAEGLNPYGLVAPR.F |
| UNC119 | UNC119 | T12T14S16 | | | | | | | | 12.5 | K. [illegible] |
| | UNC119 | T12T14S16S21 | | | | | | | | 13.0 | K. [illegible] |
| UNC13D | UNC13D | S150 | | | | | | | 12.8 | 61.0 | K.DVSGFSDPYCLLGIEQGVGVPGGS*PGSR.H |
| Uncharacterized membrane protein C19orf24 | C19orf24 | T113S117 | | | | | | | 4.4 | 11.7 | R.YGLLANTEDPT*EM#AS*LDSDEETVFESR.N |
| Uncharacterized membrane protein C19orf24 | C19orf24 | S120 | | | | | | | 24.4 | 84.9 | R.YGLLANTEDPTEMASLDS*DEETVFESR.N |
| Uncharacterized membrane protein C19orf24 | C19orf24 | S117S120 | | | | | | | 31.2 | 119.9 | R.YGLLANTEDPTEMAS*LDS*DEETVFESR.N |
| Uncharacterized membrane protein C19orf24 | C19orf24 | S117 | | | | | | | 12.0 | 97.0 | R.YGLLANTEDPTEMAS*LDSDEETVFESR.N |
| Uncharacterized membrane protein C19orf24 | C19orf24 | T109S120 | | | | | | | 3.9 | 13.7 | R.YGLLANT*EDPTEM#ASLDS*DEETVFESR.N |
| Uncharacterized membrane protein C19orf24 | C19orf24 | T109T113 | | | | | | | 1.7 | 11.7 | R.YGLLANT*EDPT*EM#ASLDSDEETVFESR.N |
| Uncharacterized membrane protein C19orf24 | C19orf24 | T113S120 | | | | | | | 20.0 | 36.8 | R.YGLLANTEDPT*EM#ASLDS*DEETVFESR.N |
| Uncharacterized protein | | S101S102 | | | | | | | 100.0 | 23.4 | R.RS*S*LPRR.G |
| Uncharacterized protein (Fragment) | | S96 | | | | | | | 6.5 | 24.2 | R.RSS*GWPR.T |
| Uncharacterized protein C11orf96 | C11orf96 | S86 | | | | | | | 12.1 | 45.5 | K.SFLQS*LECLR.R |
| Uncharacterized protein C6orf132 | C6orf132 | S1151 | | | | | | | 26.1 | 13.1 | R.S*PYTTTR.Y |
| Unconventional myosin-XVIIIa (Fragment) | MYO18A | S332 | | | | | | | 100.0 | 24.8 | R.AAS*IRR.S |
| UPF0505 protein C16orf62 | C16orf62 | T434 | | | | | | | 100.0 | 17.6 | R.ALLT*EM#MER.C |
| Upstream regulatory element binding protein 1 | HUWE1 | S2362S2365 | | | | | | | 44.1 | 107.4 | R.DGGS*GNS*TIIVSR.S |
| Upstream regulatory element binding protein 1 | HUWE1 | S1907 | | | | | | | 41.9 | 103.2 | R.GSGTAS*DDEFENLR.I |
| Upstream regulatory element binding protein 1 | HUWE1 | S718 | | | | | | | | 66.6 | R. [illegible] |
| Upstream regulatory element binding protein 1 | HUWE1 | S726S727 | | | | | | | -3.7 | 46.5 | R. [illegible] |
| Upstream regulatory element binding protein 1 | HUWE1 | S3919T3924 | | | | | | | 35.3 | 25.7 | K.DEPPPLS*PAPLT*PATPSSLDPFFSR.E |
| Upstream regulatory element binding protein 1 | HUWE1 | S2619 | | | | | | | 40.5 | 102.1 | R.S*DDELLDDFFHDQSTATSQAGTLSSIPTALTR.W |
| Upstream regulatory element binding protein 1 | HUWE1 | S3373 | | | | | | | 22.2 | 83.6 | K.ACS*PCSSQSSSSGICTDFWDLLVK.L |
| Upstream regulatory element binding protein 1 | HUWE1 | S1395 | | | | | | | 100.0 | 51.2 | R.AES*PEEVACR.K |
| Upstream regulatory element binding protein 1 | HUWE1 | T3924 | | | | | | | 0.8 | 29.5 | K.DEPPPLSPAPLT*PATPSSLDPFFSR.E |
| Upstream regulatory element binding protein 1 | HUWE1 | T2635 | | | | | | | 10.9 | 51.1 | R.SDDELLDDFFHDQSTAT*SQAGTLSSIPTALTR.W |
| Upstream regulatory element binding protein 1 | HUWE1 | S3919T3924T39 | | | | | | | 33.6 | 34.1 | K.DEPPPLS*PAPLT*PAT*PSSLDPFFSR.E |
| Upstream regulatory element binding protein 1 | HUWE1 | S740S743 | | | | | | | 6.8 | 66.1 | R. [illegible] |
| Upstream regulatory element binding protein 1 | HUWE1 | S649 | | | | | | | 12.1 | 95.9 | R.RSS*DPLGDTASNLGSAVDELMR.H |
| Upstream regulatory element binding protein 1 | HUWE1 | S3379 | | | | | | | 5.8 | 25.0 | K.ACSPCSSQS*SSSGICTDFWDLLVK.L |
| Upstream regulatory element binding protein 1 | HUWE1 | S3919 | | | | | | | 1.6 | 45.0 | K.DEPPPLS*PAPLTPATPSSLDPFFSR.E |
| Upstream regulatory element binding protein 1 | HUWE1 | S3757 | | | | | | | 41.9 | 112.7 | R.LGSSGLGS*ASSIQAAVR.Q |
| Upstream regulatory element binding protein 1 | HUWE1 | S740T744 | | | | | | | -2.8 | 37.5 | R. [illegible] |
| Upstream regulatory element binding protein 1 | HUWE1 | S3919T3927 | | | | | | | 24.0 | 15.5 | K.DEPPPLS*PAPLTPAT*PSSLDPFFSR.E |
| Upstream regulatory element binding protein 1 | HUWE1 | S3816 | | | | | | | 8.0 | 66.5 | R.REESPMDVDQPS*PSAQDTQSIASDGTPQGEK.E |
| Upstream regulatory element binding protein 1 | HUWE1 | S718S726 | | | | | | | | 86.3 | R. [illegible] |
| Upstream regulatory element binding protein 1 | HUWE1 | S2377 | | | | | | | 3.9 | 66.3 | R.SGEDES*QEDVLMDEAPSNLSQASTLQANR.E |
| Upstream regulatory element binding protein 1 | HUWE1 | S2632 | | | | | | | 2.4 | 22.8 | R.SDDELLDDFFHDQS*TATSQAGTLSSIPTALTR.W |
| Upstream regulatory element binding protein 1 | HUWE1 | S726T749 | | | | | | | 0.1 | 26.4 | R. [illegible] |
| Upstream regulatory element binding protein 1 | HUWE1 | T2633 | | | | | | | 16.0 | 74.4 | R.SDDELLDDFFHDQST*ATSQAGTLSSIPTALTR.W |
| Upstream regulatory element binding protein 1 | HUWE1 | S2636 | | | | | | | -0.1 | 58.7 | R.SDDELLDDFFHDQSTATS*QAGTLSSIPTALTR.W |
| Uracil DNA glycosylase | UNG | S23 | | | | | | | 34.3 | 45.7 | R.HAPS*PEPAVQGTGVAGVPEESGDAAIPAK.K |
| Uracil DNA glycosylase | UNG | T60S63 | | | | | | | 19.6 | 56.7 | K.KAPAGQEEPGT*PPS*SPLSAEQLDR.I |
| Uracil DNA glycosylase | UNG | T60S64 | | | | | | | 33.8 | 60.9 | K.KAPAGQEEPGT*PPSS*PLSAEQLDR.I |
| Uridine monophosphate kinase | UCK2 | S254 | | | | | | | 32.1 | 63.6 | R.QAS*ESSSRPH.- |
| USP6 N terminal like | USP6NL | T276T279 | | | | | | | 100.0 | 11.8 | R.T*PFT*LNLR.I |
| | UTP14A | S29S31 | | | | | | | 51.0 | 102.5 | K.DYLLS*ES*EDEGDNDGER.K |
| | UTP14A | S341S343 | | | | | | | | 37.5 | K.LQVAS*ES*EEEEGGTEDVEELLVPDVVNEVQMNADGPNPWMLR.S |
| UTP14, U3 small nucleolar ribonucleoprotein, homolog A | UTP14A | S405S407 | | | | | | | 38.7 | 24.3 | K.EAATQEDPEQLPELEAHGVS*ES*EGEERPVAEEEILLR.E |
| Utrophin | UTRN | S1866 | | | | | | | -0.3 | 34.4 | R.SS*LLPTDYLVEINK.I |
| Utrophin (Fragment) | UTRN | S211S213 | | | | | | | 10.1 | 19.5 | R.LGYLPVQTVLEGDNLETPS*QS*PQLFHDDTHSR.I |
| Utrophin (Fragment) | UTRN | T200S213 | | | | | | | 9.9 | 17.6 | R.LGYLPVQT*VLEGDNLETPSQS*PQLFHDDTHSR.I |
| Utrophin (Fragment) | UTRN | T209S211 | | | | | | | 8.3 | 24.2 | R.LGYLPVQTVLEGDNLET*PS*QSPQLFHDDTHSR.I |
| Utrophin (Fragment) | UTRN | T209S213 | | | | | | | 10.6 | 31.0 | R.LGYLPVQTVLEGDNLET*PSQS*PQLFHDDTHSR.I |
| Vac14 homolog | VAC14 | S539S540 | | | | | | | 100.0 | 11.3 | K.RFS*S*ER.K |
| Vacuolar protein sorting 13C | VPS13C | S829Y835 | | | | | | | -0.2 | 51.1 | K.GLLGTSLLLDTVES*ESDDEY*FDAEDGEPQTCK.S |

**Peak Area**: <-10, -5, -3, 0, 3, 5, >10

**%CV**: 0, 29, 42, 71, 86, >100

**White dots**- Significant change in peptide abundance at 5 %FDR compared to the timepoint with the minimum peak area for a given PSM

CarT: 0m, 2m, 5m — RajiB: 0m, 2m, 5m

| Protein Name | Gene | Phosphosites | Ascor | MOWSE | Sequence |
|---|---|---|---|---|---|
| Vacuolar protein sorting 13C | VPS13C | T826Y835 | -0.2 | 41.2 | K.GLLGTSLLLDT*VESESDDEY*FDAEDGEPQTCK.S |
| Vacuolar protein sorting 13C | VPS13C | T826S831 | 0.7 | 37.1 | K.GLLGTSLLLDT*VESES*DDEYFDAEDGEPQTCK.S |
| Vacuolar protein sorting 13C | VPS13C | S831Y835 | -2.1 | 27.3 | K.GLLGTSLLLDTVESES*DDEY*FDAEDGEPQTCK.S |
| Vacuolar protein sorting 13C | VPS13C | T826S829 | 13.7 | 43.9 | K.GLLGTSLLLDT*VES*ESDDEYFDAEDGEPQTCK.S |
| Vacuolar protein sorting 13C | VPS13C | S829S831 | 9.4 | 54.7 | K.GLLGTSLLLDTVES*ES*DDEYFDAEDGEPQTCK.S |
| Vacuolar protein sorting 13C | VPS13C | S821T826 | 12.4 | 33.3 | K.GLLGTS*LLLDT*VESESDDEYFDAEDGEPQTCK.S |
| Vacuolar protein sorting 13D | VPS13D | S1724 | 28.1 | 43.8 | R.S*LPSHMEEAPNVFQLYQRPTSASR.K |
| Vacuolar protein sorting 13D | VPS13D | S1034 | 2.3 | 18.1 | K.NLSFDIPTGS*LRDSR.A |
| Vacuolar protein sorting 13D | VPS13D | T2862S2864 | -0.4 | 33.3 | R.ST*AS*LTNLEHQIYAR.A |
| Vacuolar protein sorting 24 | CHMP3 | T185 | | 25.7 | K.VT*DALPEPEPPGAMAASEDEEEEEEALEAMQSR.L |
| Vacuolar protein sorting 24 | CHMP3 | S200 | 45.6 | 33.1 | K.VTDALPEPEPPGAMAAS*EDEEEEEEALEAMQSR.L |
| Valosin containing protein | VCP | S770 | 100.0 | 32.7 | R.GFGS*FR.F |
| Vanilloid receptor like 3 | TRPV3 | S374 | 22.4 | 24.7 | K.RLRSLS*R.K |
| Vanilloid receptor like protein 1 | TRPV2 | S751 | 8.4 | 25.3 | R.TLENPVLASPPKEDEDGAS*EENYVPVQLLQSN.- |
| Vanilloid receptor like protein 1 | TRPV2 | S741 | 3.6 | 20.4 | R.TLENPVLAS*PPKEDEDGASEENYVPVQLLQSN.- |
| Vanilloid receptor like protein 1 | TRPV2 | Y755 | 6.7 | 15.6 | R.TLENPVLASPPKEDEDGASEENY*VPVQLLQSN.- |
| Vasodilator stimulated phosphoprotein | VASP | S323 | -0.2 | 54.7 | K.SS*SSVTTSETQPCTPSSSDYSDLQR.V |
| Vasodilator stimulated phosphoprotein | VASP | T335 | 15.7 | 64.4 | K.SSSSVTTSETQPCT*PSSSDYSDLQR.V |
| VAV1 | VAV1 | Y826 | 100.0 | 24.5 | R.GEIY*GR.V |
| VAV2 | VAV2 | S768S769 | 9.0 | 11.9 | R.AS*S*RSPVFTPR.V |
| VAV2 | VAV2 | S768S771 | 10.0 | 21.1 | R.AS*SRS*PVFTPR.V |
| VAV2 | VAV2 | S768T775 | 6.9 | 14.6 | R.AS*SRSPVFT*PR.V |
| VCIP135 | VCPIP1 | S997S998 | -0.3 | 18.3 | R.SRES*S*PSHGLLK.L |
| VCIP135 | VCPIP1 | S1198 | 3.1 | 40.9 | R.GNS*VEELEEMDSQDAEMTNTTEPMDHS.- |
| VCIP135 | VCPIP1 | S994S997 | 8.0 | 13.5 | R.S*RES*SPSHGLLK.L |
| VDAC1 | VDAC1 | S104 | 26.0 | 85.0 | K.LTFDSSFS*PNTGKK.N |
| Ventricular zone expressed PH domain homolog 1 | VEPH1 | S783 | 100.0 | 15.2 | R.DRS*LPR.A |
| Vesicle associated membrane protein 4 | VAMP4 | S30 | 100.0 | 117.8 | R.NLLEDDS*DEEEDFFLR.G |
| Vesicle docking protein p115 | USO1 | S942 | 62.5 | 69.0 | K.DLGHPVEEEDELES*GDQEDEDDESEDPGKDLDHI.- |
| Vigilin | HDLBP | S944 | 100.0 | 22.1 | K.DCDPGS*PR.R |
| Vimentin | VIM | S55 | 27.3 | 82.0 | R.SLYAS*SPGGVYATR.S |
| Vimentin | VIM | S56 | 29.1 | 83.8 | R.SLYASS*PGGVYATR.S |
| Vimentin | VIM | Y61 | 17.8 | 62.1 | R.SLYASSPGGVY*ATR.S |
| Vimentin | VIM | S39 | 10.8 | 24.6 | R.TYS*LGSALRPSTSR.S |
| Vimentin | VIM | S51 | 10.0 | 28.4 | R.S*LYASSPGGVYATR.S |
| Vimentin | VIM | S83 | 55.6 | 11.6 | R.LLQDS*VDFSLADAINTEFK.N |
| Vinexin beta | SORBS3 | S530 | 8.2 | 42.5 | R.LCDDGPQLPTS*PR.L |
| Vinexin beta | SORBS3 | S606 | 14.7 | 52.8 | R.GPSHPLDLGTSS*PNTSQIHWTPYR.A |
| Vinexin beta | SORBS3 | S605 | 9.2 | 25.8 | R.GPSHPLDLGTS*SPNTSQIHWTPYR.A |
| Vinexin beta | SORBS3 | T529 | | 26.5 | R.LCDDGPQLPT*SPR.L |
| Voltage dependent anion selective channel protein 2 | VDAC2 | S130 | 28.4 | 44.1 | K.LTFDTTFS*PNTGKK.S |
| Vpr binding protein | DCAF1 | S1000 | 4.0 | 53.9 | R.HLPS*PPTLDSIITEYLR.E |
| WAS protein homology region 2 domain containing 1 | WHAMM | S481S484 | 100.0 | 14.0 | R.AS*LRS*R.K |
| WASP interacting protein | WIPF1 | S340 | 44.7 | 82.2 | R.NLS*LSSSTPPLPSPGR.S |
| WAVE1 | WASF1 | S336S339 | 12.0 | 18.0 | R.TPVFVSPTPPPPPPPLPSALS*TSS*LR.A |
| WAVE1 | WASF1 | S336T337 | 3.5 | 11.2 | R.TPVFVSPTPPPPPPPLPSALS*T*SSLR.A |
| WAVE1 | WASF1 | S544S546S550 | 26.1 | 14.3 | R.IAVEYS*DS*EDDS*EFDEVDWLE.- |
| WD repeat and HMG box DNA binding protein 1 | WDHD1 | S868 | 120.5 | 52.7 | R.NQVEEDAEDS*GEADDEEKPEIHKPGQNSFSK.S |
| | WDHD1 | S333 | 55.2 | 56.1 | K.DYNDLFDGDDM#S*NAGDFLNDNAVEIPSFSK.G |
| WD repeat and HMG box DNA binding protein 1 | WDHD1 | S407 | 64.8 | 13.3 | K.EEEEDGQEGS*IHNLPLVTSQRPFYDGPMPTPR.Q |
| WD repeat and HMG box DNA binding protein 1 | WDHD1 | S348 | 17.4 | 20.2 | K.DYNDLFDGDDMSNAGDFLNDNAVEIPS*FSK.G |
| WD repeat domain 22 | DCAF5 | S531S533 | 5.4 | 49.0 | R.LLALSNES*DS*EENVCEVELDTDLFPRPR.S |
| WD repeat domain 3 | WDR3 | S241 | 10.6 | 87.8 | K.GSS*PGIQDTLEAEDGAFETDEAPEDR.I |
| WD repeat domain 42A | DCAF8 | S99 | 100.0 | 81.1 | R.S*EEEEEEEEEEEEQPR.R |
| WD repeat domain 42A | DCAF8 | S129S130 | 69.4 | 34.3 | R.DQDS*S*DDERALEDWVSSETSALPRPR.W |
| WD repeat domain 44 | WDR44 | S403 | -0.3 | 44.2 | K.EYVSNDAAQS*DDEEKLQSQPTDTDGGR.L |
| WD repeat domain 44 | WDR44 | S470S471S472 | 54.0 | 58.8 | R.DEVFHTDQDDPS*S*S*DDEGMPYTRPVK.F |
| WD repeat domain 44 | WDR44 | S161 | 12.1 | 84.5 | K.LTQTS*STEQLNVLETETEVLNK.E |
| WD repeat domain 44 | WDR44 | S162 | 8.0 | 78.7 | K.LTQTSS*TEQLNVLETETEVLNK.E |
| WD repeat domain 50 | UTP18 | T180S186 | 14.5 | 62.9 | R.KT*SSDDES*EEDEDDLLQR.T |
| WD repeat domain 50 | UTP18 | S182S186 | 38.8 | 133.1 | R.KTSS*DDES*EEDEDDLLQR.T |
| WD repeat domain 50 | UTP18 | T180S182S186 | 14.6 | 49.6 | R.KT*SS*DDES*EEDEDDLLQR.T |

Peak Area: <-10, -5, -3, 0, 3, 5, >10

%CV: 0, 29, 42, 71, 86, >100

**White dots**- Significant change in peptide abundance at **5** %FDR compared to the timepoint with the minimum peak area for a given PSM

| Protein Name | Gene | Phosphosites | CarT 0m | CarT 2m | CarT 5m | RajiB 0m | RajiB 2m | RajiB 5m | Ascor | MOWSE | Sequence |
|---|---|---|---|---|---|---|---|---|---|---|---|
| WD repeat domain 50 | UTP18 | S97S100 | | | | | | | 100.0 | 104.8 | R.VQEHEDS*GDS*EVENEAK.G |
| WD repeat domain 50 | UTP18 | S181S182S186 | | | | | | | 23.8 | 53.8 | R.KTS*S*DDES*EEDEDDLLQR.T |
| WD repeat domain 50 | UTP18 | T180S182 | | | | | | | 6.2 | 22.0 | R.KT*SS*DDESEEDEDDLLQR.T |
| WD repeat domain 50 | UTP18 | S181S186 | | | | | | | 15.6 | 68.8 | R.KTS*SDDES*EEDEDDLLQR.T |
| WD repeat domain 50 | UTP18 | T180S181S186 | | | | | | | 2.8 | 47.1 | R.KT*S*SDDES*EEDEDDLLQR.T |
| WD repeat domain 50 | UTP18 | T180S181S182 | | | | | | | 7.8 | 18.2 | R.KT*S*S*DDESEEDEDDLLQR.T |
| WD repeat domain 55 | WDR55 | S14 | | | | | | | 39.2 | 60.9 | R.TCEERPAEDGS*DEEDPDSMEAPTR.I |
| WD repeat domain 70 | WDR70 | S638 | | | | | | | 103.6 | 63.8 | K.TMFAQVES*DDEEAK.N |
| WD repeat domain 75 | WDR75 | S779S782 | | | | | | | 57.6 | 20.0 | K.EIPEDVDMEEEKES*EDS*DEENDFTEK.V |
| WD repeat domain 75 | WDR75 | T798 | | | | | | | 0.3 | 107.9 | K.VQDTSNT*GLGEDIIHQLSK.S |
| WD repeat domain 75 | WDR75 | S796 | | | | | | | 17.7 | 113.8 | K.VQDTS*NTGLGEDIIHQLSK.S |
| WD repeat domain 79 | WRAP53 | T489 | | | | | | | 94.9 | 30.2 | R.VFPEPT*ESGDEGEELGLPLLSTR.H |
| WD repeat domain 79 | WRAP53 | S491 | | | | | | | 2.3 | 38.9 | R.VFPEPTES*GDEGEELGLPLLSTR.H |
| WD repeat domain, phosphoinositide interacting 2 | WIPI2 | S412 | | | | | | | 14.5 | 18.5 | K.GTYVPS*SPTR.L |
| WD repeat domain, phosphoinositide interacting 2 | WIPI2 | T408 | | | | | | | 8.4 | 16.4 | K.GT*YVPSSPTR.L |
| WD repeat protein 13 | WDR13 | S116 | | | | | | | | 36.0 | R.GS*YQLQAQMNR.A |
| WD repeat protein 13 | WDR13 | S113Y117 | | | | | | | 3.2 | 18.1 | R.SVS*RGSY*QLQAQMNR.A |
| WD repeat protein 13 | WDR13 | S113S116 | | | | | | | 6.2 | 26.3 | R.SVS*RGS*YQLQAQMNR.A |
| WD repeat-containing protein 43 | WDR43 | T656S658 | | | | | | | 100.0 | 58.1 | K.DEENGEDRDT*AS*EK.E |
| WEE 1 tyrosine kinase | WEE1 | S165T173 | | | | | | | 100.0 | 15.7 | R.S*PRPDHPGT*PPHK.T |
| WEE 1 tyrosine kinase | WEE1 | S139 | | | | | | | 3.5 | 46.6 | K.SPAAPYFLGSSFS*PVR.C |
| Werner helicase interacting protein 1 | WRNIP1 | S153 | | | | | | | 26.0 | 43.4 | K.RPAAAAAAGSAS*PR.S |
| Williams-Beuren syndrome chromosome region 5 | LAT2 | S106 | | | | | | | 9.2 | 56.1 | R.HGS*EEAYIDPIAMEYYNWGR.F |
| Wilms tumor 1 associated protein | WTAP | S305 | | | | | | | 16.4 | 11.7 | R.EGNTTEDDFPS*SPGNGNK.S |
| Wilms tumor 1 associated protein | WTAP | S306 | | | | | | | 13.9 | 33.6 | R.EGNTTEDDFPSS*PGNGNK.S |
| Wiskott-aldrich syndrome protein | WAS | S483S484 | | | | | | | 100.0 | 50.7 | R.AIHS*S*DEGEDQAGDEDEDDEWDD.- |
| Wiskott-Aldrich syndrome-like | WASL | S484S485 | | | | | | | 100.0 | 30.3 | K.AIHS*S*DEDEDEDDEEDFEDDDEWED.- |
| WNT-4 protein | WNT4 | S251S252 | | | | | | | 100.0 | 12.3 | R.RVGS*S*R.A |
| WNT-4 protein | WNT4 | S252 | | | | | | | 8.9 | 16.4 | R.RVGSS*R.A |
| Wolf-Hirschhorn syndrome candidate 2 | NELFA | T277 | | | | | | | 100.0 | 66.3 | K.T*LDAEVVEKPAK.E |
| Wolf-Hirschhorn syndrome candidate 2 | NELFA | Y377S379 | | | | | | | | 21.8 | R. [illegible] |
| WW domain binding protein 11 | WBP11 | S353S361S364 | | | | | | | 60.2 | 69.2 | R.EVEEFS*EDDDEDDS*DDS*EAEKQSQK.Q |
| WW domain binding protein 11 | WBP11 | S237 | | | | | | | 16.9 | 36.9 | R.DEDMLYS*PELAQR.G |
| WW domain containing adaptor with coiled coil | WAC | S523S535 | | | | | | | 6.8 | 44.1 | R.S*PSPGPNHTSNSS*NASNATVVPQNSSAR.S |
| WW domain containing adaptor with coiled coil | WAC | S64 | | | | | | | 51.2 | 22.5 | R.RSDS*PENK.Y |
| WW domain containing adaptor with coiled coil | WAC | S525S534 | | | | | | | 5.6 | 31.3 | R.SPS*PGPNHTSNS*SNASNATVVPQNSSAR.S |
| WW domain containing adaptor with coiled coil | WAC | T457S470 | | | | | | | 13.7 | 16.2 | R.IST*PQTNTVPIKPLIS*TPPVSSQPK.V |
| WW domain containing adaptor with coiled coil | WAC | S525S538 | | | | | | | 1.7 | 27.7 | R.SPS*PGPNHTSNSSNAS*NATVVPQNSSAR.S |
| WW domain containing adaptor with coiled coil | WAC | T457T471 | | | | | | | 8.9 | 42.1 | R.IST*PQTNTVPIKPLIST*PPVSSQPK.V |
| WW domain containing adaptor with coiled coil | WAC | S534S535 | | | | | | | 7.9 | 31.0 | R.SPSPGPNHTSNS*S*NASNATVVPQNSSAR.S |
| WW domain containing adaptor with coiled coil | WAC | S525S535 | | | | | | | 3.0 | 30.5 | R.SPS*PGPNHTSNSS*NASNATVVPQNSSAR.S |
| WW domain containing adaptor with coiled coil | WAC | S523S534 | | | | | | | 13.1 | 20.0 | R.S*PSPGPNHTSNS*SNASNATVVPQNSSAR.S |
| WW domain containing adaptor with coiled coil | WAC | S523S525 | | | | | | | 4.8 | 27.6 | R.S*PS*PGPNHTSNSSNASNATVVPQNSSAR.S |
| WW domain containing adaptor with coiled coil | WAC | S523S538 | | | | | | | -1.6 | 30.5 | R.S*PSPGPNHTSNSSNAS*NATVVPQNSSAR.S |
| WW domain containing adaptor with coiled coil | WAC | S523T531 | | | | | | | 2.4 | 23.0 | R.S*PSPGPNHT*SNSSNASNATVVPQNSSAR.S |
| WWC family member 3 | WWC3 | S927 | | | | | | | -0.5 | 10.9 | R.SDS*DSSTLPR.K |
| Xeroderma pigmentosum group C complementing protein | XPC | S883S884 | | | | | | | 41.4 | 110.5 | K.SEAAAPHTDAGGGLS*S*DEEEGTSSQAEAAR.I |
| Xeroderma pigmentosum group C complementing protein | XPC | S122S129 | | | | | | | 27.6 | 102.1 | R.GATMNEDS*NEEEEES*ENDWEEVEELSEPVLGDVR.E |
| Xeroderma pigmentosum group C complementing protein | XPC | T117S129 | | | | | | | 3.7 | 49.1 | R.GAT*MNEDSNEEEEES*ENDWEEVEELSEPVLGDVR.E |
| | XPC | T117S122 | | | | | | | 5.4 | 71.1 | R.GAT*M#NEDS*NEEEEESENDWEEVEELSEPVLGDVR.E |
| XPA binding protein 1 | GPN1 | S338 | | | | | | | 53.8 | 118.4 | R.GTLDEEDEEADS*DTDDIDHR.V |
| XPA binding protein 1 | GPN1 | S314 | | | | | | | 23.4 | 56.4 | K.DSLS*PVLHPSDLILTR.G |
| XPA binding protein 1 | GPN1 | S312 | | | | | | | 11.6 | 18.0 | K.DS*LSPVLHPSDLILTR.G |
| XRCC1 | XRCC1 | T198S226 | | | | | | | 11.5 | 68.4 | R.INKT*SPVTASDPAGPSYAAATLQASSAASSAS*PVSR.A |
| XRCC1 | XRCC1 | S518T519T523 | | | | | | | 11.9 | 24.9 | R. [illegible] |
| XRCC1 | XRCC1 | S241 | | | | | | | 32.2 | 39.9 | R.AIGSTSKPQES*PK.G |
| XRCC1 | XRCC1 | S199S226 | | | | | | | 16.7 | 95.3 | R.INKTS*PVTASDPAGPSYAAATLQASSAASSAS*PVSR.A |
| XRCC1 | XRCC1 | S226 | | | | | | | 22.8 | 106.3 | K.TSPVTASDPAGPSYAAATLQASSAASSAS*PVSR.A |
| XRCC1 | XRCC1 | Y515S518T519 | | | | | | | | 19.1 | R. [illegible] |
| | YBX3 | S34 | | | | | | | 11.8 | 86.7 | K. [illegible] |
| Y-box-binding protein 3 | YBX3 | S34S38 | | | | | | | | 29.9 | K. [illegible] |
| YB-1 | YBX1 | S165 | | | | | | | 44.4 | 76.2 | R.NYQQNYQNS*ESGEKNEGSESAPEGQAQQR.R |

Peak Area: <-10, -5, -3, 0, 3, 5, >10

%CV: 0, 29, 42, 71, 86, >100

**White dots-** Significant change in peptide abundance at **5** %FDR compared to the timepoint with the minimum peak area for a given PSM

CarT (0m, 2m, 5m) RajiB (0m, 2m, 5m)

| Protein Name | Gene | Phosphosites | Ascor | MOWSE | Sequence |
|---|---|---|---|---|---|
| | YBX1 | S176 | 22.0 | 56.7 | R.NYQQNYQNSESGEKNEGSES*APEGQAQQR.R |
| YB-1 | YBX1 | Y162S165 | 5.5 | 13.3 | R.NYQQNY*QNS*ESGEKNEGSESAPEGQAQQR.R |
| YB-1 | YBX1 | S314 | 10.8 | 30.1 | K.AADPPAENSS*APEAEQGGAE.- |
| YB-1 | YBX1 | S174 | 32.1 | 83.7 | K.NEGS*ESAPEGQAQQR.R |
| YB-1 | YBX1 | S313 | | 12.5 | K.AADPPAENS*SAPEAEQGGAE.- |
| YB-1 | YBX1 | Y162 | 17.3 | 31.6 | R.NYQQNY*QNSESGEK.N |
| YB-1 | YBX3 | S102 | 30.6 | 42.9 | R.S*VGDGETVEFDVVEGEK.G |
| | YBX1 | S167 | 6.3 | 12.1 | R.NYQQNYQNSES*GEK.N |
| YLP motif containing 1 | YLPM1 | S398 | 22.4 | 64.4 | R.GPASQFYITPSTSLS*PR.Q |
| YLP motif containing 1 | YLPM1 | S1119 | 100.0 | 45.7 | R.RAGS*QER.G |
| YLP motif containing 1 | YLPM1 | S394 | 7.8 | 12.9 | R.GPASQFYITPS*TSLSPR.Q |
| YRDC | YRDC | S37 | 100.0 | 34.5 | R.LFRPPS*PAPAAPGAR.L |
| YTH domain containing 2 | YTHDC2 | S1090 | 25.6 | 40.8 | R.VDGIPNDSS*DSEMEDK.T |
| YTH domain containing 2 | YTHDC2 | S1089 | 22.0 | 49.7 | R.VDGIPNDS*SDSEMEDK.T |
| YTH domain family 1 | YTHDF1 | S291 | | 35.3 | K. APVPQQAPS*PQAAPQPQQVAQPLPAQPPALAQPQYQSPQQPPQT |
| YY1 associated factor 2 | YAF2 | T8 | 1.6 | 30.1 | K.SPT*RPK.R |
| YY1 associated factor 2 | YAF2 | S6 | 22.7 | 27.9 | K.S*PTRPK.R |
| ZAK | MAP3K20 | S648 | 16.9 | 42.6 | K.NFS*SLHLNSR.D |
| ZAP70 | ZAP70 | Y492Y493 | 12.9 | 43.1 | K.ALGADDSY*Y*TAR.S |
| Zinc and ring finger 2 | ZNRF2 | S82 | 106.1 | 77.9 | R.S*LGGAVGSVASGAR.A |
| Zinc and ring finger protein 1 | ZNRF1 | S123 | 100.0 | 67.4 | R.AS*LADALPLHIAPR.W |
| Zinc and ring finger protein 1 | ZNRF1 | S50S53 | 8.1 | 62.8 | R.S*VSS*VAGMGMDPSTAGGVPFGLYTPASR.G |
| Zinc and ring finger protein 1 | ZNRF1 | S120S123 | 14.4 | 31.7 | R.DGMLYLGS*RAS*LADALPLHIAPR.W |
| Zinc and ring finger protein 1 | ZNRF1 | S50S62 | 6.7 | 35.1 | R.S*VSSVAGMGMDPS*TAGGVPFGLYTPASR.G |
| Zinc and ring finger protein 1 | ZNRF1 | S50 | 18.0 | 54.5 | R.S*VSSVAGMGMDPSTAGGVPFGLYTPASR.G |
| Zinc and ring finger protein 1 | ZNRF1 | S52S62 | 0.1 | 34.5 | R.SVS*SVAGMGMDPS*TAGGVPFGLYTPASR.G |
| Zinc and ring finger protein 1 | ZNRF1 | S50S52 | 5.1 | 57.6 | R.S*VS*SVAGMGMDPSTAGGVPFGLYTPASR.G |
| Zinc finger and BTB domain containing 10 | ZBTB10 | T646 | 48.0 | 59.0 | R.NVNANLLAEAGT*SQDGGDAGTSHDFK.Y |
| Zinc finger antiviral protein | ZC3HAV1 | S335 | 82.0 | 101.2 | R.FLENGS*QEDLLHGNPGSTYLASNSTSAPNWK.S |
| Zinc finger antiviral protein | ZC3HAV1 | S312 | 5.2 | 18.6 | R.ARPPSGS*SK.A |
| Zinc finger antiviral protein | ZC3HAV1 | S235 | 12.1 | 18.6 | R.APS*SHRR.N |
| Zinc finger antiviral protein | ZC3HAV1 | S271T273S275 | 46.8 | 15.2 | R.S*CT*PS*PDQISHR.A |
| Zinc finger antiviral protein | ZC3HAV1 | T347S355 | 9.3 | 42.8 | R.FLENGSQEDLLHGNPGST*YLASNSTS*APNWK.S |
| Zinc finger antiviral protein | ZC3HAV1 | S257 | 90.1 | 88.6 | R.FFQGS*QEFLASASASAER.S |
| Zinc finger antiviral protein | ZC3HAV1 | S346 | 4.0 | 56.1 | R.FLENGSQEDLLHGNPGS*TYLASNSTSAPNWK.S |
| Zinc finger antiviral protein | ZC3HAV1 | S335S355 | 16.3 | 44.3 | R.FLENGS*QEDLLHGNPGSTYLASNSTS*APNWK.S |
| Zinc finger antiviral protein | ZC3HAV1 | S346S355 | 13.4 | 38.9 | R.FLENGSQEDLLHGNPGS*TYLASNSTS*APNWK.S |
| Zinc finger antiviral protein | ZC3HAV1 | S387 | -0.4 | 31.3 | R.SS*LGSLQTPEAVTTR.K |
| Zinc finger antiviral protein | ZC3HAV1 | S284 | 116.4 | 62.7 | R.AS*LEDAPVDDLTR.K |
| Zinc finger antiviral protein | ZC3HAV1 | T375 | 22.2 | 28.7 | K.T*VFSPTLPAAR.S |
| Zinc finger antiviral protein | ZC3HAV1 | S378 | 11.4 | 33.8 | K.TVFS*PTLPAAR.S |
| Zinc finger antiviral protein | ZC3HAV1 | S310 | 4.0 | 18.1 | R.ARPPS*GSSK.A |
| Zinc finger CCCH-type antiviral protein 1 | ZC3HAV1 | S572 | 49.7 | 64.1 | R.NLVPTTPGESTAPAQVSTLPQS*PAALSSSNR.A |
| Zinc finger CCCH-type containing 11A | ZC3H11B | S761 | 9.2 | 33.0 | R.RLSSAS*TGKPPLSVEDDFEK.L |
| Zinc finger CCCH-type containing 11A | ZC3H11A | S171 | 27.0 | 51.3 | K.VESSENVPSPTHPPVVINAADDDEDDDDQFS*EEGDETK.T |
| Zinc finger CCCH-type containing 11A | ZC3H11B | S761T762 | 6.1 | 48.2 | R.RLSSAS*T*GKPPLSVEDDFEK.L |
| Zinc finger CCCH-type containing 11A | ZC3H11B | S758 | 3.5 | 38.2 | R.RLS*SASTGKPPLSVEDDFEK.L |
| Zinc finger CCCH-type containing 11A | ZC3H11B | S758S761 | 13.6 | 56.3 | R.RLS*SAS*TGKPPLSVEDDFEK.L |
| Zinc finger CCCH-type containing 13 | ZC3H13 | S198 | 100.0 | 59.0 | K.EVS*PEVVR.S |
| Zinc finger CCCH-type containing 13 | ZC3H13 | S207S209 | 27.0 | 25.0 | R.SKLS*PS*PSLR.K |
| Zinc finger CCCH-type containing 13 | ZC3H13 | S877 | 9.9 | 53.1 | R.SLS*PSHLTEDR.Q |
| Zinc finger CCCH-type containing 13 | ZC3H13 | S372Y374S381 | 2.3 | 32.8 | R.SAS*PY*PSHSLSS*PQR.K |
| Zinc finger CCCH-type containing 13 | ZC3H13 | S1456Y1462S1 | 24.4 | 48.9 | K.LDDAHSLGS*GAGEGY*EPIS*DDELDEILAGDAEK.R |
| Zinc finger CCCH-type containing 13 | ZC3H13 | S387 | 100.0 | 20.4 | R.KQS*PPR.H |
| Zinc finger CCCH-type containing 13 | ZC3H13 | S77 | 18.4 | 28.7 | R.S*PERPTGDLR.E |
| Zinc finger CCCH-type containing 13 | ZC3H13 | S1014S1017 | 31.8 | 71.6 | K.SKGDS*DIS*DEEAAQQSK.K |
| Zinc finger CCCH-type containing 13 | ZC3H13 | T263S265 | 100.0 | 40.3 | R.T*PS*PPPPIPEDIALGKK.Y |
| Zinc finger CCCH-type containing 13 | ZC3H13 | T317S318S325 | 10.2 | 16.3 | R.ST*S*PAGQHHS*PISSR.H |
| Zinc finger CCCH-type containing 13 | ZC3H13 | S370S372S380 | 22.3 | 16.8 | R.S*AS*PYPSHSLS*SPQR.K |
| Zinc finger CCCH-type containing 13 | ZC3H13 | S372Y374S378 | 5.8 | 10.8 | R.SAS*PY*PSHS*LSSPQR.K |
| Zinc finger CCCH-type containing 13 | ZC3H13 | S372Y374S380 | 1.3 | 23.3 | R.SAS*PY*PSHSLS*SPQR.K |
| Zinc finger CCCH-type containing 13 | ZC3H13 | T134S137 | 50.1 | 23.5 | K.ERT*PES*EEENVEWETNR.D |

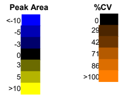


**White dots-** Significant change in peptide abundance at 5 %FDR compared to the timepoint with the minimum peak area for a given PSM

| Protein Name | Gene | Phosphosites | Ascor | MOWSE | Sequence |
|---|---|---|---|---|---|
| Zinc finger CCCH-type containing 13 | ZC3H13 | S1453S1456S1 | 42.4 | 56.5 | K.LDDAHS*LGS*GAGEGYEPIS*DDELDEILAGDAEK.R |
| Zinc finger CCCH-type containing 13 | ZC3H13 | S643 | 6.5 | 18.4 | R.DQRPSS*PIR.H |
| Zinc finger CCCH-type containing 13 | ZC3H13 | S1010S1014S1 | 56.9 | 59.0 | K.S*KGDS*DIS*DEEAAQQSK.K |
| Zinc finger CCCH-type containing 13 | ZC3H13 | S875 | 13.8 | 47.1 | R.S*LSPSHLTEDR.Q |
| Zinc finger CCCH-type containing 13 | ZC3H13 | S370S372S378 | 8.4 | 19.4 | R.S*AS*PYPSHS*LSSPQR.K |
| Zinc finger CCCH-type containing 13 | ZC3H13 | S875S877 | 24.2 | 18.2 | R.S*LS*PSHLTEDR.Q |
| Zinc finger CCCH-type containing 13 | ZC3H13 | S110 | 7.8 | 18.6 | R.NTEESSS*PVRK.E |
| Zinc finger DHHC domain containing 5 | ZDHHC5 | S296S299 | 10.9 | 15.0 | K.S*KGS*LEITESQSADAEPPPPPKPDLSR.Y |
| Zinc finger DHHC domain containing 5 | ZDHHC5 | S423 | 21.2 | 41.6 | K.SFHFDPLSS*GSR.S |
| Zinc finger DHHC domain containing 5 | ZDHHC5 | S429S432 | 11.6 | 64.3 | R.SSS*LKS*AQGTGFELGQLQSIR.S |
| Zinc finger DHHC domain containing 5 | ZDHHC5 | S428S429 | 4.4 | 65.3 | R.SS*S*LKSAQGTGFELGQLQSIR.S |
| Zinc finger DHHC domain containing 5 | ZDHHC5 | S432 | 35.4 | 100.5 | K.S*AQGTGFELGQLQSIR.S |
| Zinc finger DHHC domain containing 5 | ZDHHC5 | S422 | 37.2 | 36.0 | K.SFHFDPLS*SGSR.S |
| Zinc finger DHHC domain containing 5 | ZDHHC5 | S425 | 9.1 | 37.9 | K.SFHFDPLSSGS*R.S |
| Zinc finger DHHC domain containing 5 | ZDHHC5 | S429T436 | 1.9 | 50.0 | R.SSS*LKSAQGT*GFELGQLQSIR.S |
| Zinc finger FYVE domain containing 26 | ZFYVE26 | S1762 | 6.2 | 79.6 | R.S*PSAEFSPAAPPGISSIHSPSLR.E |
| Zinc finger FYVE domain containing 26 | ZFYVE26 | S1764 | 7.6 | 55.0 | R.SPS*AEFSPAAPPGISSIHSPSLR.E |
| Zinc finger protein 106 | ZNF106 | S1370 | 78.7 | 42.1 | R.AAHVPENS*DTEQDVLTVKPVRK.V |
| Zinc finger protein 106 | ZNF106 | S1025S1026 | 20.7 | 53.1 | R.ATGDGS*S*PELPSLER.K |
| Zinc finger protein 183 | RNF113A | S43S45S46S47 | 36.4 | 27.0 | R.KRPACDPEPGES*GS*S*S*DEGCTVVRPEK.K |
| Zinc finger protein 183 | RNF113A | S84S85 | 38.2 | 55.8 | K.AAYGDLS*S*EEEEENEPESLGVVYK.S |
| Zinc finger protein 183 | RNF113A | S253 | 87.6 | 107.2 | R.YGVYEDENYEVGS*DDEEIPFK.C |
| Zinc finger protein 183 | RNF113A | Y80S84 | 25.5 | 13.7 | K.AAY*GDLS*SEEEEENEPESLGVVYK.S |
| Zinc finger protein 206 | ZSCAN10 | S333 | -0.2 | 10.8 | K.SFGRS*SILK.L |
| Zinc finger protein 22 | ZNF22 | S49 | 43.5 | 22.1 | R.S*LDDKPYK.C |
| Zinc finger protein 261 | ZMYM3 | S256T257 | | 28.6 | R.RAEPPKPEVVDS*T*ESIPVSDEDSDAMVDDPNDEDFVPFRPR.R |
| Zinc finger protein 261 | ZMYM3 | S957S960 | 46.5 | 68.9 | K.ASSDLCDLVS*NQS*AEGLLEDCDLFGPAR.D |
| Zinc finger protein 261 | ZMYM3 | S263S267 | 45.8 | 18.5 | R.RAEPPKPEVVDSTESIPVS*DEDS*DAMVDDPNDEDFVPFRPR.R |
| Zinc finger protein 262 | ZMYM4 | S122 | 12.7 | 46.2 | R.VTQHES*DNENEIQIQNK.L |
| Zinc finger protein 262 | ZMYM4 | S1542S1547 | 22.3 | 46.2 | K.SEDS*DVELS*D.- |
| Zinc finger protein 262 | ZMYM4 | S1088T1089 | 8.9 | 32.2 | K.DQGS*T*YSGDLESEAVSTPHSWEEELNHYALK.S |
| Zinc finger protein 262 | ZMYM4 | S1088S1091 | 16.8 | 11.1 | K.DQGS*TYS*GDLESEAVSTPHSWEEELNHYALK.S |
| Zinc finger protein 265 | ZRANB2 | S153 | 112.6 | 109.7 | K.EVEDKES*EGEEEDEDEDLSK.Y |
| Zinc finger protein 265 | ZRANB2 | S305S307 | 53.8 | 36.9 | R.S*RS*PESQVIGENTK.Q |
| | ZRANB2 | S188 | 46.5 | 77.4 | K.YNLDAS*EEEDSNK.K |
| Zinc finger protein 265 | ZRANB2 | S120 | 49.2 | 74.7 | R.EES*DGEYDEFGRK.K |
| Zinc finger protein 265 | ZRANB2 | T303S305S307 | 31.6 | 29.7 | R.T*RS*RS*PESQVIGENTK.Q |
| Zinc finger protein 265 | ZRANB2 | S266S267 | 0.4 | 12.4 | R.SHRGS*S*SPR.K |
| Zinc finger protein 265 | ZRANB2 | S307 | 37.8 | 77.4 | R.S*PESQVIGENTK.Q |
| Zinc finger protein 265 | ZRANB2 | T303S307S310 | 12.3 | 11.5 | R.T*RSRS*PES*QVIGENTK.Q |
| Zinc finger protein 265 | ZRANB2 | S305S307S310 | 10.5 | 11.9 | R.TRS*RS*PES*QVIGENTK.Q |
| Zinc finger protein 272 | ZNF460 | T492 | 9.8 | 19.6 | R.RSPLT*R.H |
| Zinc finger protein 272 | ZNF460 | S489 | 15.8 | 20.1 | R.RS*PLTR.H |
| Zinc finger protein 28 (KOX 24) | ZNF28 | S427 | 100.0 | 14.2 | R.CKS*HLER.H |
| Zinc finger protein 289, ID1 regulated | ARFGAP2 | S319 | 3.5 | 17.2 | R.SSVSHS*VLSEMQVIEQETPVSAK.S |
| Zinc finger protein 289, ID1 regulated | ARFGAP2 | S152 | | 15.3 | K.DS*DEETEHTQPPAWDAPATEPSCTQQPARSTESSCLAQPEHGPNT |
| Zinc finger protein 289, ID1 regulated | ARFGAP2 | S146 | 36.2 | 18.2 | R.HGTDLWIDNMSSAVPNHS*PEKK.D |
| Zinc finger protein 295 | ZBTB21 | S345 | 13.6 | 84.5 | R.SLS*MDSQVPVYSPSIDLK.S |
| Zinc finger protein 316 | ZNF316 | S112 | 100.0 | 14.1 | K.S*PVLQEK.G |
| Zinc finger protein 318 | ZNF318 | S136 | 74.0 | 59.2 | R.S*PGLCSDSLEK.S |
| Zinc finger protein 318 | ZNF318 | S173 | 138.9 | 58.0 | R.LGS*PVDNLEDMDRDDLTDDSVFTR.S |
| Zinc finger protein 318 | ZNF318 | S79S81 | 100.0 | 28.2 | R.RVS*PS*PPR.A |
| Zinc finger protein 318 | ZNF318 | S89S91 | 100.0 | 24.7 | R.RGS*PS*PPR.G |
| Zinc finger protein 318 | ZNF318 | S214 | 3.6 | 45.5 | R.YISQEEGPLS*PFLGQLDEDYR.T |
| Zinc finger protein 318 | ZNF318 | S69S71 | 100.0 | 18.1 | R.RAS*PS*PPR.G |
| Zinc finger protein 318 | ZNF318 | T686S698 | 24.7 | 16.7 | R.EAHHSNT*HSPEVSHPHPPS*PVDPYLLTK.N |
| Zinc finger protein 318 | ZNF318 | S684S698 | 26.3 | 16.9 | R.EAHHS*NTHSPEVSHPHPPS*PVDPYLLTK.N |
| Zinc finger protein 318 | ZNF318 | Y205S214 | 4.1 | 32.0 | R.Y*ISQEEGPLS*PFLGQLDEDYR.T |
| Zinc finger protein 330 | ZNF330 | S291 | 44.7 | 143.7 | R.KDS*DTESSDLFTNLNLGR.T |
| Zinc finger protein 36 | ZKSCAN1 | S208 | 100.0 | 14.7 | R.ALPAAHIPAPPHEGS*PR.D |
| Zinc finger protein 36 C3H type like 2 | ZFP36L2 | S57 | 49.3 | 34.1 | R.RHS*ASNLHALAHPAPSPGSCSPK.F |
| Zinc finger protein 36 C3H type like 2 | ZFP36L2 | S59S75 | 7.3 | 17.0 | R.RHSAS*NLHALAHPAPSPGSCS*PK.F |

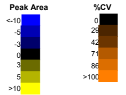


**White dots**- Significant change in peptide abundance at 5 %FDR compared to the timepoint with the minimum peak area for a given PSM

| Protein Name | Gene | Phosphosites | Ascor | MOWSE | Sequence |
|---|---|---|---|---|---|
| Zinc finger protein 36 C3H type like 2 | ZFP36L2 | S59 | 4.6 | 15.7 | R.RHSAS*NLHALAHPAPSPGSCSPK.F |
| Zinc finger protein 363 | RCHY1 | S257 | 100.0 | 25.5 | R.RIS*LDQQ.- |
| Zinc finger protein 40 | HIVEP1 | S536 | 32.1 | 83.5 | K.SSFTPS*SPENVIGDFLLQDR.S |
| Zinc finger protein 40 | HIVEP1 | S537 | 15.2 | 55.3 | K.SSFTPSS*PENVIGDFLLQDR.S |
| Zinc finger protein 40 | HIVEP1 | T569S577 | 6.4 | 15.1 | K.VVVHHVT*VSPLRTDS*PK.A |
| Zinc finger protein 444 | ZNF444 | S104 | 33.0 | 63.2 | R.QPQSGEEAVALLEELWGPAAS*PDGSSATR.V |
| Zinc finger protein 483 | ZNF483 | T404S406 | 17.5 | 18.0 | R.RST*LS*R.R |
| Zinc finger protein 483 | ZNF483 | S403S406 | 13.9 | 12.3 | R.RS*TLS*R.R |
| Zinc finger protein 581 | ZNF581 | S30S36 | | 49.5 | R.S*PEPGPS*SSIGSPQASSPPRPNHYLLIDTQGVPYTVLVDEESQR. |
| Zinc finger protein 581 | ZNF581 | S45T58 | 12.1 | 53.2 | R.SPEPGPSSSIGSPQAS*SPPRPNHYLLIDT*QGVPYTVLVDEESQR. |
| Zinc finger protein 581 | ZNF581 | S45S46 | 7.1 | 24.5 | R.SPEPGPSSSIGSPQAS*S*PPRPNHYLLIDTQGVPYTVLVDEESQR. |
| Zinc finger protein 581 | ZNF581 | Y53T58 | -2.7 | 18.9 | R.SPEPGPSSSIGSPQASSPPRPNHY*LLIDT*QGVPYTVLVDEESQR. |
| Zinc finger protein 581 | ZNF581 | S46T58 | -0.4 | 40.6 | R.SPEPGPSSSIGSPQASS*PPRPNHYLLIDT*QGVPYTVLVDEESQR. |
| Zinc finger protein 592 | ZNF592 | S142S145S146 | 9.9 | 26.4 | K.SEPLPTFNQFS*PIS*S*PEPEDPIKDNGFGIKPK.H |
| Zinc finger protein 592 | ZNF592 | S689 | 4.3 | 42.0 | K.HGLTSGS*ASPPPPALPLYPDPVR.L |
| Zinc finger protein 592 | ZNF592 | T137S145S146 | 10.1 | 21.7 | K.SEPLPT*FNQFSPIS*S*PEPEDPIKDNGFGIKPK.H |
| Zinc finger protein 592 | ZNF592 | S691 | 7.3 | 49.4 | K.HGLTSGSAS*PPPPALPLYPDPVR.L |
| Zinc finger protein 592 | ZNF592 | S132T137S142 | | 21.5 | K.S*EPLPT*FNQFS*PISSPEPEDPIKDNGFGIKPK.H |
| Zinc finger protein 609 | ZNF609 | S842Y845S849 | 22.1 | 58.7 | K.TNS*PAY*SDIS*DAGEDGEGKVDSVK.S |
| Zinc finger protein 609 | ZNF609 | S842S846S849 | 17.8 | 75.8 | K.TNS*PAYS*DIS*DAGEDGEGKVDSVK.S |
| Zinc finger protein 609 | ZNF609 | Y845S846S849 | 8.4 | 12.7 | K.TNSPAY*S*DIS*DAGEDGEGKVDSVK.S |
| Zinc finger protein 638 | ZNF638 | S508S510 | 32.7 | 12.8 | R.S*RS*PMHYMYRPR.S |
| Zinc finger protein 638 | ZNF638 | S552 | 100.0 | 16.1 | R.IRNPFRGS*PK.C |
| Zinc finger protein 646 | ZNF646 | S930 | 54.7 | 123.2 | R.S*PPLQLSEAELLNQLQR.E |
| Zinc finger protein 646 | ZNF646 | S936 | 8.2 | 89.0 | R.SPPLQLS*EAELLNQLQR.E |
| Zinc finger protein 687 | ZNF687 | S251 | 42.3 | 15.2 | K.ATDIPAS*ASPPPVAGVPFFK.Q |
| Zinc finger protein 687 | ZNF687 | S253 | 13.4 | 36.3 | K.ATDIPASAS*PPPVAGVPFFK.Q |
| Zinc finger protein 687 | ZNF687 | S1057 | 100.0 | 25.2 | R.HGLQLGAQS*PGR.G |
| Zinc finger protein 687 | ZNF687 | T191S197 | | 11.9 | R.EGALT*PPPFPS*SFELAQENGPGMQPPVSSPPLGALK.Q |
| Zinc finger protein 687 | ZNF687 | S198S215 | 17.8 | 11.1 | R.EGALTPPPFPSS*FELAQENGPGMQPPVSS*PPLGALK.Q |
| Zinc finger protein 740 | ZNF740 | S44 | 100.0 | 50.4 | R.AGS*PDVLR.C |
| Zinc finger protein 800 | ZNF800 | S334 | 82.1 | 38.8 | R.DSITPDIATKPGQPLFLDS*ISPK.K |
| Zinc finger protein 99 | | S2 | | 13.2 | -.M#S*FWSEK.C |
| Zinc finger protein A20 | TNFAIP3 | S381 | 46.9 | 54.3 | R.EGHAQNPMEPSVPQLS*LMDVK.C |
| Zinc finger protein A20 | TNFAIP3 | S376S381 | 100.0 | 45.0 | R.EGHAQNPMEPS*VPQLS*LMDVK.C |
| Zinc finger protein A20 | TNFAIP3 | T454 | 7.2 | 30.2 | R.GEAYEPLAWNPEEST*GGPHSAPPTAPSPFLFSETTAMK.C |
| Zinc finger protein A20 | TNFAIP3 | S645 | 32.1 | 13.9 | K.VS*PTASR.F |
| Zinc finger protein A20 | TNFAIP3 | S459 | 23.2 | 39.5 | R.GEAYEPLAWNPEESTGGPHS*APPTAPSPFLFSETTAMK.C |
| Zinc finger protein A20 | TNFAIP3 | T463 | 7.1 | 45.2 | R.GEAYEPLAWNPEESTGGPHSAPPT*APSPFLFSETTAMK.C |
| Zinc finger protein A20 | TNFAIP3 | S453 | 4.2 | 45.9 | R.GEAYEPLAWNPEES*TGGPHSAPPTAPSPFLFSETTAMK.C |
| Zinc finger protein A20 | TNFAIP3 | S573S575 | 37.5 | 37.8 | R.S*PS*PHSCHR.A |
| Zinc finger protein A20 | TNFAIP3 | S453S471 | 10.1 | 17.0 | R.GEAYEPLAWNPEES*TGGPHSAPPTAPSPFLFS*ETTAMK.C |
| Zinc finger protein A20 | TNFAIP3 | S459S466 | 12.8 | 14.6 | R.GEAYEPLAWNPEESTGGPHS*APPTAPS*PFLFSETTAMK.C |
| Zinc finger protein A20 | TNFAIP3 | T454S459 | 1.5 | 35.3 | R.GEAYEPLAWNPEEST*GGPHS*APPTAPSPFLFSETTAMK.C |
| Zinc finger protein A20 | TNFAIP3 | S453S459 | 6.4 | 30.0 | R.GEAYEPLAWNPEES*TGGPHS*APPTAPSPFLFSETTAMK.C |
| Zinc finger protein A20 | TNFAIP3 | S466 | | 19.2 | R.GEAYEPLAWNPEESTGGPHSAPPTAPS*PFLFSETTAMK.C |
| Zinc finger protein A20 | TNFAIP3 | S453T454 | 15.3 | 30.1 | R.GEAYEPLAWNPEES*T*GGPHSAPPTAPSPFLFSETTAMK.C |
| Zinc finger protein A20 | TNFAIP3 | S220 | 23.6 | 26.1 | R.SLES*GSNFAPLK.V |
| Zinc finger protein A20 | TNFAIP3 | T454S466 | 13.5 | 21.9 | R.GEAYEPLAWNPEEST*GGPHSAPPTAPS*PFLFSETTAMK.C |
| Zinc finger protein A20 | TNFAIP3 | S453T463 | 0.6 | 37.1 | R.GEAYEPLAWNPEES*TGGPHSAPPT*APSPFLFSETTAMK.C |
| Zinc finger protein AEBP2 | AEBP2 | S206 | 49.2 | 24.9 | R.RGS*LEMSSDGEPLSR.M |
| Zinc finger protein hRit1 alph | BCL11B | S375S381 | 26.8 | 51.2 | R.ELAGNSS*TPPPVS*PGR.G |
| Zinc finger protein hRit1 alph | BCL11B | T376S381 | 14.7 | 57.1 | R.ELAGNSST*PPPVS*PGR.G |
| Zinc finger protein hRit1 alph | BCL11B | S129 | 16.0 | 59.5 | R.KVSEPVEIGIQVTPDEDDHLLS*PTK.G |
| Zinc finger protein hRit1 alph | BCL11B | S496S497 | 21.9 | 43.0 | R.SDDGLSAAS*S*PEPGTSELAGEGLK.A |
| Zinc finger protein hRit1 alph | BCL11B | S398 | 100.0 | 43.7 | R.LLNPFQPS*PK.S |
| Zinc finger protein hRit1 alph | BCL11B | T406T417 | 54.5 | 24.8 | K.SPFLST*PPLPPMPPGGT*PPPQPPAK.S |
| Zinc finger protein hRit1 alph | BCL11B | S493S496 | 6.6 | 25.6 | R.SDDGLS*AAS*SPEPGTSELAGEGLK.A |
| Zinc finger protein hRit1 alph | BCL11B | S493S496S497 | 1.1 | 25.5 | R.SDDGLS*AAS*S*PEPGTSELAGEGLK.A |
| Zinc finger protein hRit1 alph | BCL11B | S401T417 | 56.4 | 19.0 | K.S*PFLSTPPLPPMPPGGT*PPPQPPAK.S |
| Zinc finger protein hRit1 alph | BCL11B | S401 | 30.6 | 19.4 | K.S*PFLSTPPLPPMPPGGTPPPQPPAK.S |
| Zinc finger protein hRit1 alph | BCL11B | S488S493S496 | 25.2 | 30.6 | R.S*DDGLS*AAS*S*PEPGTSELAGEGLK.A |

**Peak Area**: <-10, -5, -3, 0, 3, 5, >10

**%CV**: 0, 29, 42, 71, 86, >100

**White dots-** Significant change in peptide abundance at **5** %FDR compared to the timepoint with the minimum peak area for a given PSM

| Protein Name | Gene | Phosphosites | Ascor | MOWSE | Sequence |
|---|---|---|---|---|---|
| Zinc finger protein hRit1 alph | BCL11B | S401S405T417 | 42.2 | 14.6 | K.S*PFLS*TPPLPPMPPGGT*PPPQPPAK.S |
| Zinc finger protein, subfamily 1A, member 1 | IKZF1 | S409 | 42.6 | 45.1 | R.S*GLIYLTNHIAPHAR.N |
| Zinc finger protein, subfamily 1A, member 1 | IKZF1 | S17T23 | 11.7 | 39.5 | K.ES*PPVSDT*PDEGDEPMPIPEDLSTTSGGQQSSK.S |
| Zinc finger protein, subfamily 1A, member 1 | IKZF1 | S361S364 | 32.1 | 69.0 | R.S*NHS*AQDSAVENLLLLSK.A |
| Zinc finger protein, subfamily 1A, member 1 | IKZF1 | S445 | 34.4 | 57.0 | R.AASENS*QDALR.V |
| Zinc finger protein, subfamily 1A, member 1 | IKZF1 | S364 | 8.7 | 49.9 | R.SNHS*AQDSAVENLLLLSK.A |
| Zinc finger protein, subfamily 1A, member 1 | IKZF1 | S21T23 | 1.0 | 48.9 | K.ESPPVS*DT*PDEGDEPMPIPEDLSTTSGGQQSSK.S |
| Zinc finger protein, subfamily 1A, member 1 | IKZF1 | S361 | 9.2 | 71.3 | R.S*NHSAQDSAVENLLLLSK.A |
| Zinc finger protein, subfamily 1A, member 1 | IKZF1 | S63 | 39.7 | 47.1 | K.VETQS*DEENGR.A |
| Zinc finger protein, subfamily 1A, member 1 | IKZF1 | S17S21T23 | 42.7 | 45.4 | K.ES*PPVS*DT*PDEGDEPMPIPEDLSTTSGGQQSSK.S |
|  | IKZF2 | S56 | 65.7 | 63.4 | K.LEMQS*DEECDRKPLSR.E |
| Zinc finger protein, subfamily 1A, member 2 | IKZF2 | S369T372S375 | 59.6 | 72.2 | R.IERPIS*RET*ADS*HENNMDGPISLIRPK.S |
| Zinc finger protein, subfamily 1A, member 2 | IKZF2 | S433 | 20.3 | 27.5 | R.KQS*PAYMK.E |
| Zinc finger protein, subfamily 1A, member 2 | IKZF2 | S369 | 100.0 | 17.6 | R.IERPIS*R.E |
| Zinc finger protein, subfamily 1A, member 2 | IKZF2 | S40 | 17.0 | 20.5 | R.EHSNMAIDLTSSTPNGQHAS*PSHMTSTNSVK.L |
| Zinc finger protein, subfamily 1A, member 2 | IKZF2 | S31S40 | 20.6 | 35.2 | R.EHSNMAIDLTS*STPNGQHAS*PSHMTSTNSVK.L |
| Zinc finger protein, subfamily 1A, member 2 | IKZF2 | T30S40 | 7.4 | 19.6 | R.EHSNMAIDLT*SSTPNGQHAS*PSHMTSTNSVK.L |
| Zinc finger protein, subfamily 1A, member 2 | IKZF2 | S78S79 | 50.0 | 62.6 | R.GHDEGS*S*LEEPLIESSEVADNR.K |
| Zinc finger protein, subfamily 1A, member 2 | IKZF2 | S32S40 | 7.3 | 39.0 | R.EHSNMAIDLTSS*TPNGQHAS*PSHMTSTNSVK.L |
| Zinc finger protein, subfamily 1A, member 2 | IKZF2 | S23S40 | 23.7 | 32.6 | R.EHS*NMAIDLTSSTPNGQHAS*PSHMTSTNSVK.L |
| Zinc finger protein, subfamily 1A, member 2 | IKZF2 | S23T30 |  | 28.4 | R.EHS*NMAIDLT*SSTPNGQHASPSHMTSTNSVK.L |
| Zinc finger protein, subfamily 1A, member 2 | IKZF2 | S23 |  | 27.9 | R.EHS*NMAIDLTSSTPNGQHASPSHMTSTNSVK.L |
| Zinc finger protein, subfamily 1A, member 3 | IKZF3 | S378S382S386 |  | 12.2 | R.GLS*PNNS*GHDS*T*DT*DS*NHEER.Q |
| Zinc finger, AN1-type domain 2B | ZFAND2B | S183 | 18.6 | 110.8 | R.SPS*WTAPPVIALQNGLSEDEALQR.A |
| Zinc finger, AN1-type domain 2B | ZFAND2B | T185 | 6.0 | 78.8 | R.SPSWT*APPVIALQNGLSEDEALQR.A |
| Zinc finger, CCHC domain containing 6 | ZCCHC6 | T13 | 100.0 | 12.8 | K.RT*KDR.G |
| Zinc finger, CW type with coiled-coil domain 1 | MORC2 | S681 | 50.7 | 32.6 | R.SVAVS*DEEEVEEEAER.R |
| Zinc finger, CW type with coiled-coil domain 1 | MORC2 | S553 | 100.0 | 17.6 | R.S*PPLPAVIR.N |
| Zinc finger, CW type with coiled-coil domain 1 | MORC2 | S715S717 | 29.4 | 62.4 | K.KDSNELS*DS*AGEEDSADLKR.A |
| Zinc finger, FYVE domain containing 19 | ZFYVE19 | S344 | 133.0 | 99.6 | R.LPDS*DDDEDEETAIQR.V |
| Zinc finger, FYVE domain containing 19 | ZFYVE19 | S134 | 67.4 | 30.2 | K.WS*PPQNYK.K |
| Znf HX | ATRX | Y1009S1012S1 | 2.3 | 36.6 | K.MEQQY*ESS*S*DGTEKLPER.E |
| Znf HX | ATRX | T1989S1995 | 5.9 | 39.7 | K.AT*SSSNPS*SPAPDWYKDFVTDADAEVLEHSGK.M |
| Znf HX | ATRX | S1990S1991 |  | 19.3 | K.ATS*S*SNPSSPAPDWYKDFVTDADAEVLEHSGK.M |
| Znf HX | ATRX | T1346S1352 | 20.3 | 58.4 | R.HKLT*VSDGES*GEEKK.T |
| Znf HX | ATRX | S1348S1352 | 13.8 | 61.9 | R.HKLTVS*DGES*GEEKK.T |
| Znf HX | ATRX | S1011S1012S10 | 10.2 | 41.7 | K.MEQQYES*S*S*DGTEKLPER.E |
| Znf HX | ATRX | S729S731 | 61.1 | 71.1 | K.QSETVDQNS*DS*DEMLAILK.E |
| Znf HX | ATRX | T724S731 | 9.3 | 18.1 | K.QSET*VDQNSDS*DEMLAILK.E |
| Znf HX | ATRX | Y1009S1011S10 | 10.1 | 17.6 | K.MEQQY*ES*SS*DGTEKLPER.E |
| Znf HX | ATRX | Y1009S1011S10 | 30.6 | 33.5 | K.MEQQY*ES*S*SDGTEKLPER.E |
| Znf HX | ATRX | T1989Y2002 | 4.3 | 14.2 | K.AT*SSSNPSSPAPDWY*KDFVTDADAEVLEHSGK.M |
| Znf HX | ATRX | S1991S1995 | 10.3 | 74.7 | K.ATSS*SNPS*SPAPDWYKDFVTDADAEVLEHSGK.M |
| Znf HX | ATRX | S1996Y2002 | -2.0 | 25.6 | K.ATSSSNPSS*PAPDWY*KDFVTDADAEVLEHSGK.M |
| ZNF635 | POGZ | S746 | 100.0 | 15.8 | R.KMS*VMGR.Q |
| Zuotin related factor 1 | DNAJC2 | S49 | 12.1 | 55.7 | R.NASAS*FQELEDKK.E |
| Zuotin related factor 1 | DNAJC2 | S47 | 39.3 | 73.0 | R.NAS*ASFQELEDK.K |
| Zuotin related factor 1 | DNAJC2 | S47S49 | 100.0 | 39.3 | R.NAS*AS*FQELEDKK.E |
| Zyxin | ZYX | S308 | 15.7 | 44.6 | K.LGHPEALSAGTGS*PQPPSFTYAQQR.E |
| Zyxin | ZYX | S344 | 54.8 | 40.8 | R.S*PGAPGPLTLK.E |
